# TransDFL: Identification of Disordered Flexible Linkers in Proteins by Transfer Learning

**DOI:** 10.1101/2022.06.03.494673

**Authors:** Yihe Pang, Bin Liu

**Affiliations:** School of Computer Science and Technology, Beijing Institute of Technology, Beijing 100081, China; Advanced Research Institute of Multidisciplinary Science, Beijing Institute of Technology, Beijing 100081, China

**Author notes:** Corresponding author. (Liu B).

**Keywords:** Intrinsically disordered proteins, Disordered flexible linkers, False-positive rate, Computational predictor, Transfer learning

## Abstract

Disordered flexible linkers (DFLs) are the functional disordered regions in proteins, which are the sub-regions of intrinsically disordered regions (IDRs) and play important roles in connecting domains and maintaining inter-domain interactions. Trained with the limited available DFLs, the existing DFL predictors based on the machine learning techniques tend to predict the ordered residues as DFLs leading to a high false-positive rate (FPR) and low prediction accuracy. Previous studies have shown that DFLs are the extremely flexible disordered regions, which are usually predicted as disordered residues with high confidence [P(D) > 0.9] by an IDR predictor. Therefore, transferring an IDR predictor to an accurate DFL predictor is of great significance for understanding the functions of IDRs. In this study, we proposed a new predictor called TransDFL for identifying DFLs by transferring the RFPR-IDP predictor for IDR identification to the DFL prediction. The RFPR-IDP was pre-trained with IDR sequences to learn the general features between IDRs and DFLs, which is helpful to reduce the false positives in the ordered regions. RFPR-IDP was fine-tuned with the DFL sequences to capture the specific features of DFLs so as to be transferred into the TransDFL. Experimental results of two application scenarios (prediction of DFLs only in the IDRs or prediction of DFLs in the entire proteins) showed that TransDFL consistently outperforms the other exiting DFL predictors with higher accuracy. The corresponding web server of TransDFL can be freely accessed from http://bliulab.net/TransDFL/.

## Introduction

Intrinsically disordered regions (IDRs) are protein regions without stable three-dimensional (3D) structures, which are particularly common among the eukaryotic organisms and viral proteomes [1]. Although the IDRs lack well-defined 3D structures, they carry out many critical functions, such as transcriptions, signal transmission, post-translational modifications, multi-protein aggregation, etc [2].

The functions of IDRs derive either from binding to molecular partners (such as DNAs, RNAs, and proteins) or directly from their native disordered states, where the former is called binding functions and the latter is called non-binding functions [3]. According to the DisProt database [4], about 75% of the non-binding functions are disordered flexible linkers (DFLs) [5, 6]. DFLs serve as the linkers in multidomain proteins characterized by extremely structural flexibility, and can be located between inter-and intra-domain, which are different from generic linkers [5, 7-10]. DFLs play essential roles for intramolecular allosteric regulation [11, 12] and phase separation [13]. Identification of DFLs is crucial for comprehensively studying IDR functions. Experimental annotation of DFLs primarily relies on X-ray crystallography, NMR spectroscopy, and circular dichroism. In order to efficiently identify the DFLs, two computational methods have been developed only based on the protein sequences, including DFLpred [5] and APOD [6]. DFLpred identifies the DFLs via combining the logistic regression (LR) and four sequence-based features, including structure domain propensities, putative disordered regions, and two properties of spiral and turn formation. APOD incorporates various sequence profile features into Support Vector Machines (SVMs) to further improve the predictive performance, such as evolutionary conservation, relative solvent accessible area, etc.

These predictors successfully incorporate various sequence profile features for DFL prediction. However, DFLs are continuous regions in proteins, sharing global sequence patterns along the whole protein [5]. The global features of DFLs should be incorporated into the DFL predictors. Furthermore, DFLs are the sub-regions of IDRs, while sequences with unannotated disordered regions are common in nature [14-16]. As a result, DFL predictors tend to predict the ordered residues as DFLs, resulting in high FPR and low prediction accuracy.

According to the recent CAID experiment reports (Necci, et al., 2021), great efforts have been made by researchers for the development of IDR predictors. **Fig. 1** shows DFL prediction results on the DFL dataset TE82 [6] predicted by 6 state-of-the-art IDR predictors, including AUCpred [17], SPINE-D [18], DISOPRED3 [19], SPOT-disorder [20], IDP-Seq2seq [16], and SPOT-disorder2 [15]. We can see that DFLs can be predicted with high disordered probabilities [i.e. extremely disordered state P(D) > 0.9] by different IDR predictors, providing an opportunity to predict the DFLs based on an IDR predictor (see **Fig. 1a**).

**Figure 1.**
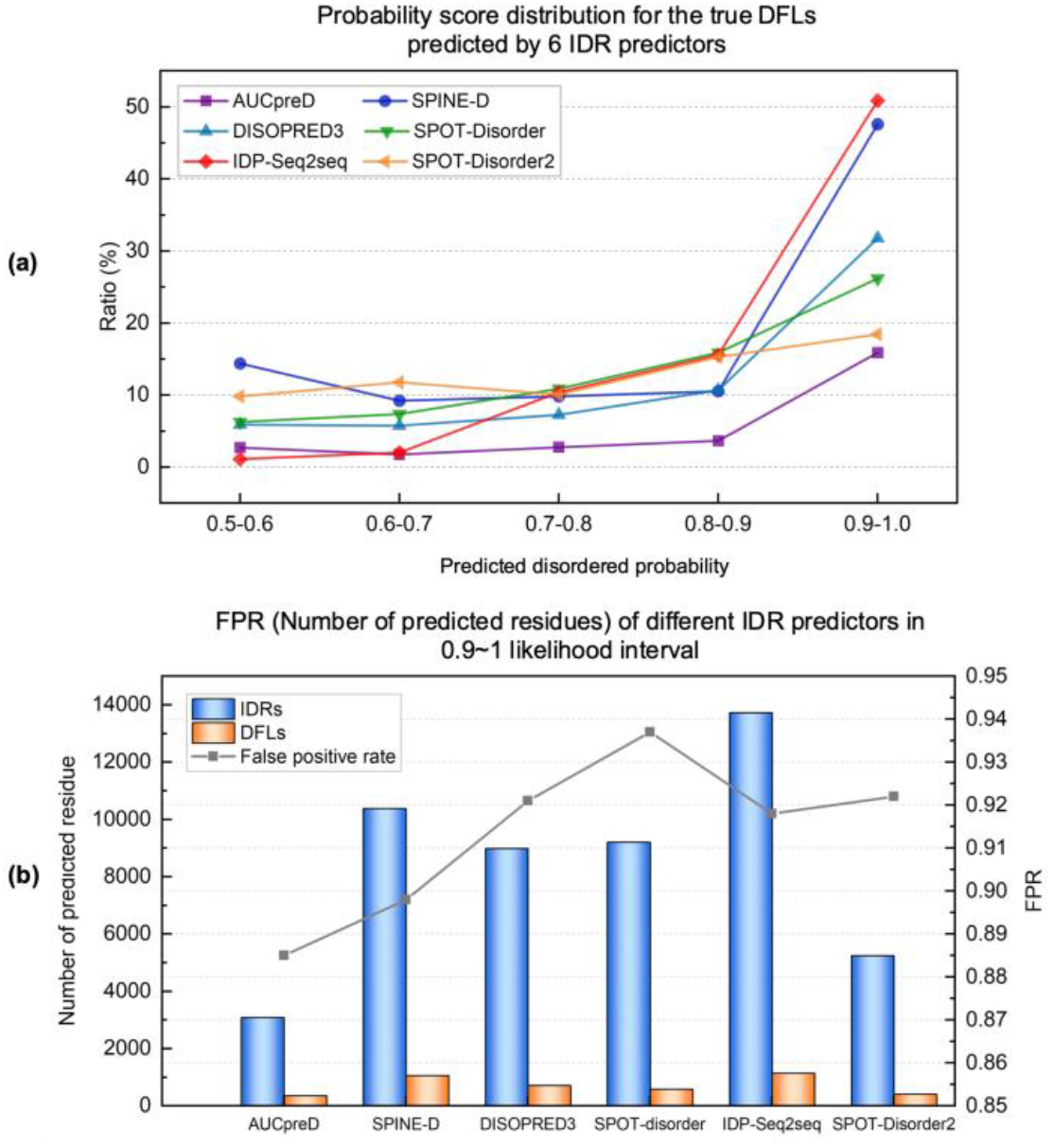
Applying the IDR predictor to DFLs. **A**. The relationships between the true DFLs and their probability scores predicted by IDR predictors. DFLs are preferred to be confidently predicted by IDR predictors with higher disordered probabilities [i.e. extremely disordered state P(D) > 0.9]. **B**. General IDR predictors are insufficient with DFLs because of high FPR, many non-DFLs also show probabilities higher than 0.9, leading to a large number of false positives. Histograms show the number of predicted IDRs and DFLs by different IDR predictors with probabilities between 0.9 and 1.0. Line shows the corresponding FPR of each predictor, which equals to the ratio of non-DFLs in predicted IDRs.

The information of disordered regions and functions are both encoded in their primary sequences, similar as the source language and target language share the same semantic [21] in the field of machine translation. For example, French and Portuguese are both from the Romance language family sharing similar grammatical structures, the pre-trained French translation model can be transferred to Portuguese translation via transfer learning [22, 23] (see **Fig. 2a**). Motivated by the similarities between protein sequences and natural languages, we treated the IDR prediction as the French translation and DFL prediction as the Portuguese translation according to the predictive correlations between IDRs and DFLs as discussed in **Fig. 1** (See **Fig. 2b**). A new DFL predictor was proposed called TransDFL, which was transferred from an IDR predictor by the transfer learning technology. The IDR predictor RFPR-IDP [24] was pre-trained with the IDR data to learn the common features between DFLs and IDRs, then it was transferred to DFL prediction by fine-tuning so as to capture the specific features of DFLs. The proposed TransDFL has the following advantages: (1) The predicted model employed the sequence labelling method by combining Bi-directional Long Short-Term Memory neural network (Bi-LSTM) and Convolutional Neural Network (CNN), which models the protein as a whole and captures the local and long-range interaction features among residues. (2) The disordered features learned from the pre-trained IDR predictor by transfer learning can reduce the incorrectly predicted DFL residues in the ordered regions, leading to a lower FPR.

**Figure 2.**
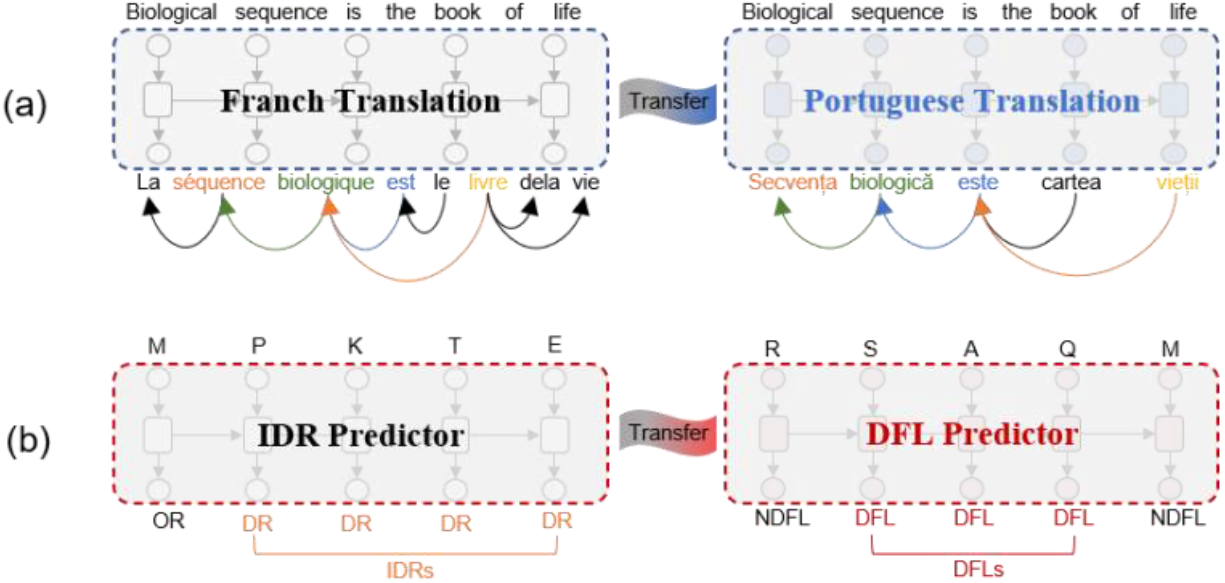
Comparison between the transfer learning frameworks for machine translation and DFL prediction. **A**. The linguistic commonalities learned from the language translation pairs, such as English-French parallel corpus can be adapted to the English-Portuguese translation by the transfer learning. **B**. In DFL prediction, the IDR predictor RFPR-IDP trained with the IDRs datasets was transferred to predict the DFL by transfer learning.

We evaluated the performance of different DFL predictors in two scenarios: prediction of DFLs only in the IDRs (situation-I) and prediction of DFLs in the entire proteins (situation-II). Experimental results showed that TransDFL consistently outperforms existing predictors. Furthermore, the corresponding web server of TransDFL was established, which can be accessed at http://bliulab.net/TransDFL/.

## Methods

### Datasets

In the pre-training phase, the IDR benchmark dataset [16] was used for RFPR-IDP predictor pre-training. To avoid the redundancy between the source and target domains, proteins sharing >25% similarities with any protein in the DFL datasets (TR166, TE82 and TE64) were removed from the IDR benchmark dataset by using Blastclust search tool [25], leading to 2645 training IDR sequences and 1077 validation IDR sequences. The details of IDR benchmark dataset are given in Supplementary Information S1.

In fine-tuning phase, the TR166 [6] DFL benchmark dataset collected by Peng et al [6] was used for model fine-tuning, and any two proteins in the dataset share sequence similarities <25%. We randomly divided the DFL benchmark dataset into five subsets. Four of the subsets with 133 sequences were randomly selected as the training dataset for fine-tuning the model parameters, and the remaining one subset with 33 sequences was employed as the validation dataset for model selection. This way ensures that there is no redundancy between the validation and training datasets.

In this study, TE82 and TE64 independent test sets were used for the performance evaluation of different DFL predictors. The TE82 test set has 82 sequences collected from DisProt database (V8.0) by Peng et al [6], and the sequences identity between the TE82 and the TR166 is less than 25%. We constructed a new independent test set TE64 from the latest released DisProt database (V9.0, September 2021) [4, 26]. Following the previous annotation protocols [5, 6], IDR proteins that have DFL functionally annotated regions in the database were collected as “DFL proteins”. To reduce data redundancies and avoid overestimating the predictive performance, only the sequences sharing <25% similarities with any protein in TE82, TR166, and IDR benchmark dataset were included in TE64. Finally, 64 sequences were collected as the TE64 independent test set. More details were given in Supplementary Information S2.

### The overview of TransDFL predictor

The overall flowchart of TransDFL is shown in **Fig. 3**.

**Figure 3.**
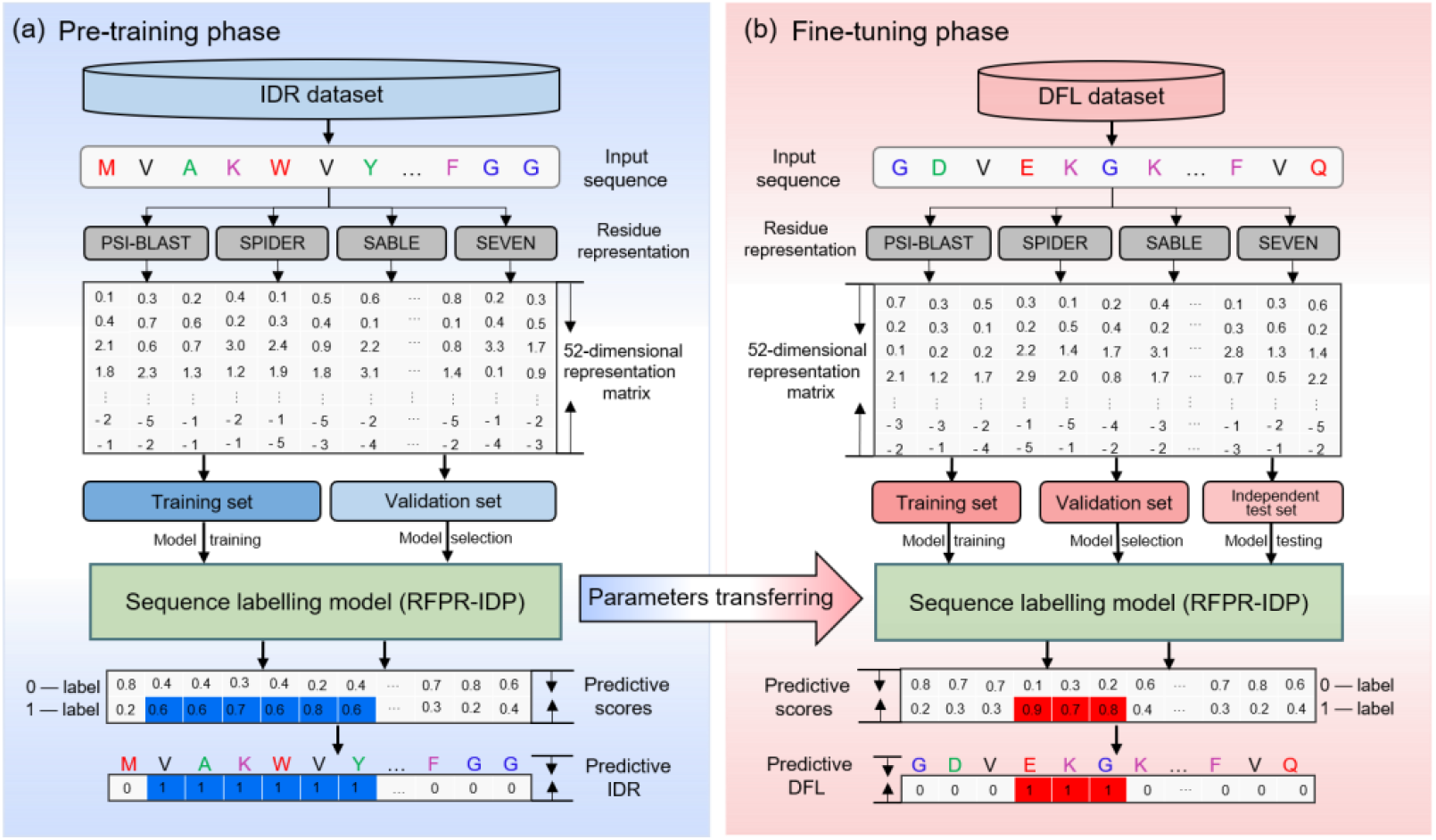
The flowchart of TransDFL predictor. **A**. In the pre-training phase, IDR dataset was used for pre-training the sequence labelling model of the IDR predictor RFPR-IDP. **B**. In the fine-tuning phase, DFL dataset was used to fine-tune the sequence labelling model for DFL prediction through transfer learning.

#### Sequence representation

In this study, the state-of-the-art IDR predictor RFPR-IDP [24] was employed to transfer into the DFL predictor. Two sequential features were used to represent sequence in RFPR-IDP, including seven commonly used physicochemical properties [27] (steric parameter, polarizability, volume, hydrophobicity, isoelectric point, helix probability, and sheet probability), and PSSM features generated by the PSI-BLAST tool [25] searching against the nrdb90 database (downloaded from http://ftp.ebi.ac.uk/pub/databases/nrdb90/). In this study, we also incorporate two additional features into RFPR-IDP so as to more comprehensively represent the DFL sequences, including the secondary structures (SS) features generated by the SPIDER tool [28, 29], and the solvent accessibility (SA) features generated by the SABLE tool [30, 31]. The linear combination of the 4-dimensional SS features, 1-dimensional SA feature, 7-dimensional physicochemical features (Seven), and 40-dimensional PSSM features, leading to a feature vector with 52 dimensions for representing a residue R_*i*_ as:

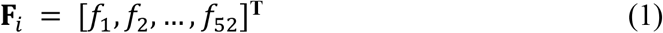

The four features provide complementary information, and their combination leads to the best prediction performance (See **Table S1** in Supplementary Information S3). Following [5, 6], the local sliding window was applied to represent the residues. The target residue R_*i*_ can be represeted as *win***F**_*i*_, which is the combination of the corresponding feature vectors of the R_*i*_ and its *k*-1 neighboring residues.

#### The sequence labelling model transferred to TransDFL

The sequence labelling model is able to incorporate the correlation among adjacent residues, and captures the interaction features of residues along the whole proteins. Two IDR predictors IDP-Seq2seq [16] and RFPR-IDP [24] based on sequence labelling model were used to be transferred to TransDFL. However, due to the insufficient number of DFL training sequences, the IDP-Seq2seq using a more complex network structure is not suitable. Therefore, the RFPR-ID predictor by a combination of Bi-LSTM and CNN is more suitable for transferring to TransDFL. The model architecture is shown in **Fig. 4**. The Bi-LSTM layer with a forward and a backward LSTM layer was adopted to capture the global correlation features. For each residue R_*i*_, the correlation feature vector **H**_*i*_ is calculated by [24]:

**Figure 4.**
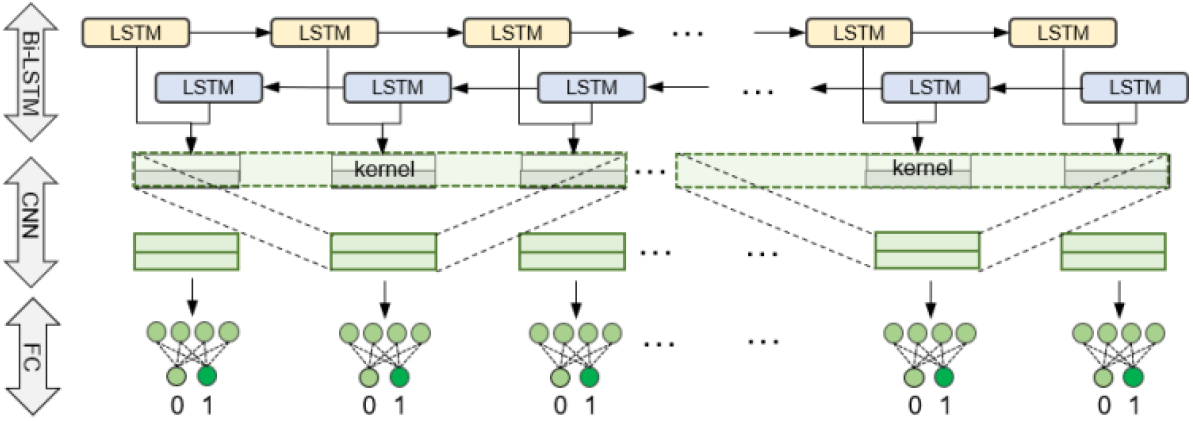
The sequence labelling model architecture transferred to TransDFL. _ENREF_44_ENREF_37

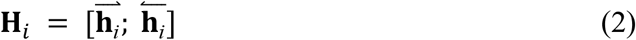

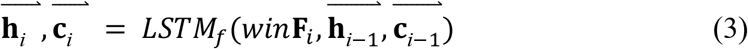

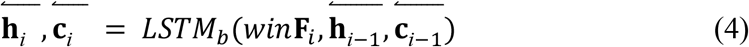

where *win***F**_*i*_ is the feature representation vector of R_*i*_, 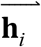 and 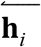 represent the forward and backward output feature vectors of R_*i*_, respectively.

The convolutional layer was used to capture the local correlation features 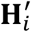 of R_*i*_:

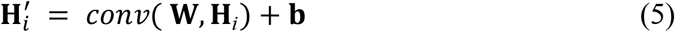

where **W** is the convolutional kernel and **b** is the bias parameter matrix.

Then, a fully-connected layer was used to predict the label of each residue, mapping the output feature vector 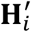 from the CNN layer to a probability score *p*_*i*_ of R_*i*_ being a positive residue, which is calculated by [32]:

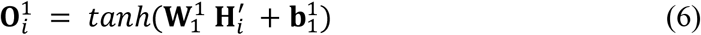

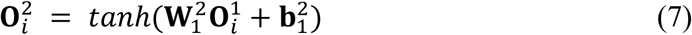

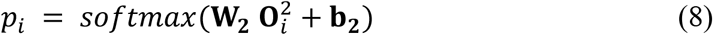

where 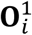 and 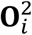 represent the output vector of the first and second fully connected layers, respectively. 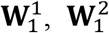, and **W**_2_ are the trainable weight parameter vectors, 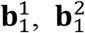, and **b**_2_ are the trainable bias parameter vectors, *tanh* is the hyperbolic tangent activation function [33] and *softmax* is the soft argmax activation function [34].

#### The pre-training phase

In the pre-training phase, the RFPR-IDP predictor was pre-trained with the IDR dataset (see **Fig. 3(a)**). The pre-trained model was optimized based on the binary cross entropy loss function calculated by [35]:

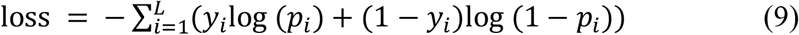

where *L* is the length of sequence, *p*_*i*_ is the predictive probability score of residue R_*i*_ being an IDR residue (cf. **Eq. 8**), *y*_*i*_ is the corresponding real label (0 or 1). All the model parameters were optimized by minimizing the loss function value on the IDRs validation set. The model pre-trained with the source domain IDR dataset learns the common characteristics sharing with IDRs and DFLs, which can be used for the DFL prediction by transfer learning. The hyper-parameters of RFPR-IDP in the pre-training phase were given in Table S1 in Supplementary Information S3.

#### The fine-tuning phase

Because DFLs are the extremely flexible disordered regions predicted by IDR predictors with high probabilities, this pattern can be transferred to identify the DFLs via transfer learning (see **Fig. 3b**). Different from the model directly trained with target dataset with limited number of samples, the model fine-tuned based on the pre-trained parameters avoids over-fitting, and improves the prediction performance in the target domain [36].

The weighted binary cross entropy loss function was employed in the fine-tuning phase:

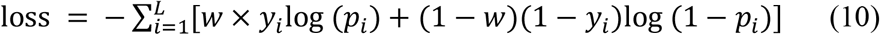

where the *w* is the weight coefficient of DFL residue, optimized according to the best AUC values on the DFL validation dataset (see **Table S2** in Supplementary Information S3). The model was implemented by the Tensorflow framework [37]. Adam algorithm [38] with a learning rate of 0.0008 was used for parameters optimization. All the parameters of the pre-trained RFPR-IDP model were fine-tuned on the validation set of DFL benchmark dataset according to the minimum loss. The hyper-parameters in the fine-tuning phase were given in Table S2 in Supplementary Information S3.

#### Performance evaluation strategy

In this study, the area under receiver operating characteristic curve (AUC) was used to evaluate the overall performance of different methods [39-41]. Besides, following previous studies [6, 42], the Matthews Correlation Coefficient (MCC) [29, 43], Precision (Pre), and Recall (Rec)[44] were used to evaluate the predictive quality of a predictor:

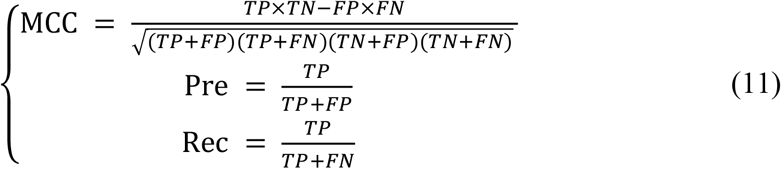

where *TP* is the number of DFL residues correctly predicted as DFLs, *FP* is the number of non-DFL residues incorrectly predicted as DFLs, *TN* is the number of non-DFL residues correctly predicted as non-DFLs, *FN* is the number of DFL residues incorrectly predicted as non-DFLs. Given a threshold *thd*, a residue R_*i*_ is classified as DFL residue, if its predictive probability score *p*_*i*_ ≥ *thd*. Otherwise, it is predicted as a non-DFL residue.

## Results and discussion

### Performance comparison among TransDFL and the other DFL predictors for predicting the DFL residues in IDRs

We compared the performance of TransDFL to the other DFL predictors on two independent test datasets (TE82 and TE64). Following previous studies [5, 6], disordered regions without functional annotations and ordered regions were not evaluated (situation-I). The results of different predictors on the TE82 and TE64 datasets were shown in **Table 1** and **Table 2**, respectively. From these results we can see that TransDFL outperforms DFLpred by 0.198 in terms of AUC on TE82 dataset, and achieves highly comparable performance with APOD. Particularly, the P-R curves (**Fig.S2** in Supplementary Information S3) show that the predictions of TransDFL and APOD are complementary and their differences are significant with a P-value of 0.000 (See P-value in **Table 1** and **Table 2**).

**Table 1.**
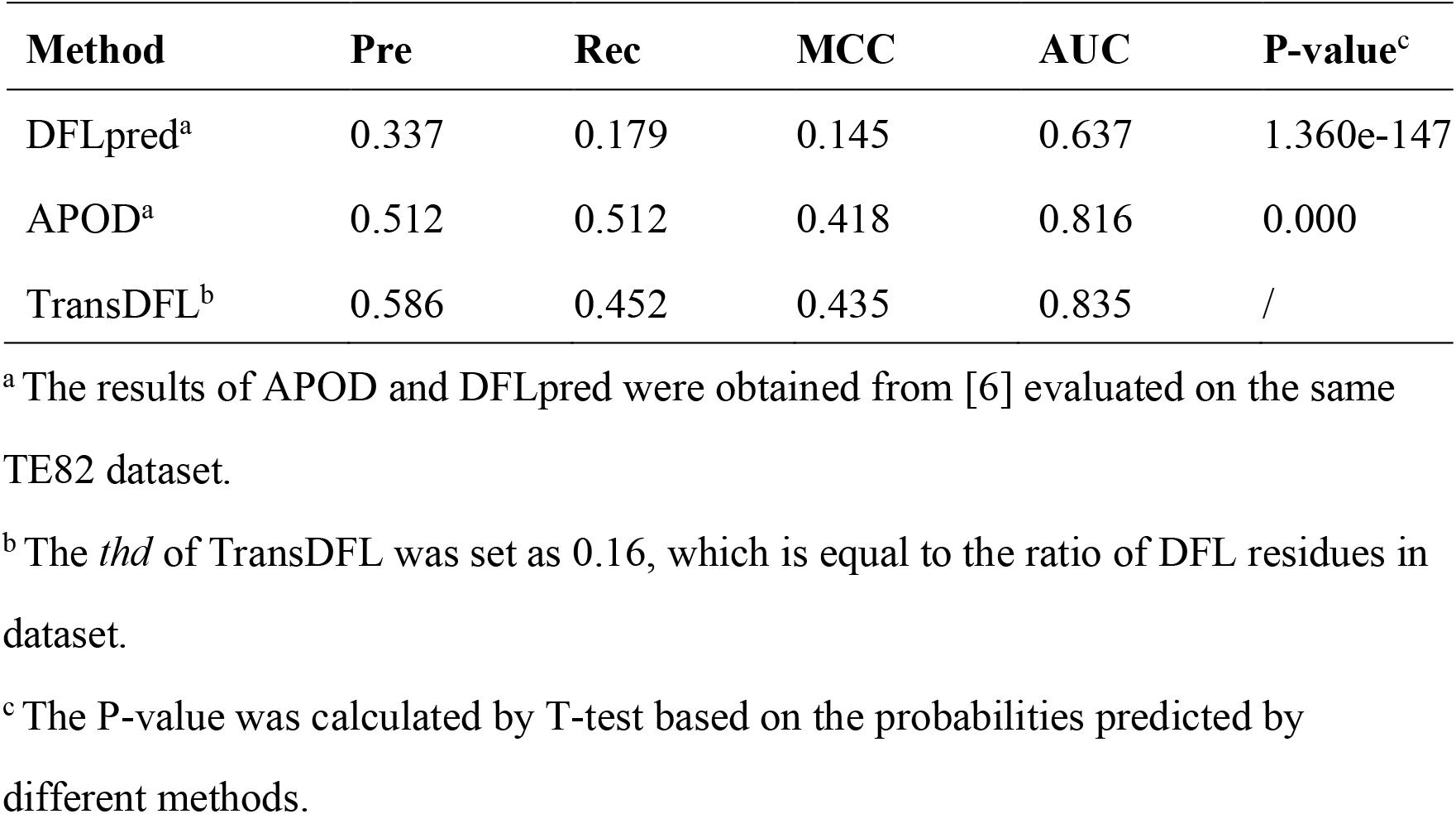
Performance comparison of TransDFL and other predictors on the TE82 independent test dataset evaluated in situation-I.

| Method | Pre | Rec | MCC | AUC | P-value<sup>c</sup> |
| --- | --- | --- | --- | --- | --- |
| DFLpred<sup>a</sup> | 0.337 | 0.179 | 0.145 | 0.637 | 1.360e-147 |
| APOD<sup>a</sup> | 0.512 | 0.512 | 0.418 | 0.816 | 0.000 |
| TransDFL<sup>b</sup> | 0.586 | 0.452 | 0.435 | 0.835 | / |

<sup>a</sup> The results of APOD and DFLpred were obtained from [6] evaluated on the same TE82 dataset.
<sup>b</sup> The *thd* of TransDFL was set as 0.16, which is equal to the ratio of DFL residues in dataset.
<sup>c</sup> The P-value was calculated by T-test based on the probabilities predicted by different methods.

**Table 2.** Performance comparison of TransDFL and other predictors on the TE64 independent test dataset evaluated in situation-I.

| Method | Pre | Rec | MCC | AUC | P-value<sup>c</sup> |
| --- | --- | --- | --- | --- | --- |
| DFLpred<sup>a</sup> | 0.189 | 0.193 | 0.108 | 0.675 | 5.330e-15 |
| APOD<sup>a</sup> | 0.195 | 0.600 | 0.223 | 0.751 | 0.000 |
| TransDFL<sup>b</sup> | 0.207 | 0.722 | 0.273 | 0.784 | / |

<sup>a</sup> The results of APOD and DFLpred were calculated according to the predicted results obtained by running corresponding web servers.
<sup>b</sup> The *thd* of TransDFL was set as same as in Table 1.
<sup>c</sup> The P-value was calculated by T-test based on the probabilities predicted by different methods

### Performance comparison among TransDFL and the other DFL predictors for predicting the DFL residues in the entire proteins

DFLs are flexible linkers in disordered regions. The existing two predictors (APOD and DFLpred) focus on predicting DFLs only in the functionally annotated disordered regions. However, the information of the disordered regions is not always available [16, 45-47]. In order to more comprehensively and fairly evaluate the performance of different methods, they were evaluated for identifying DFLs in the entire protein sequences in TE82 and TE64 datasets (situation-II). Their AUC values were shown in **Fig.5**, and the P-R curves and areas under the P-R curve (AUPR) were shown in **Fig.S3**, from which we can see the followings: 1) Compared with the results in situation-I, the performance of these three predictors decreased. These results indicate that predicting the DFLs in the entire sequence is more challenging. 2) TransDFL obviously outperforms DFLpred and APOD on both the two datasets in terms of AUC value.

**Figure 5.**
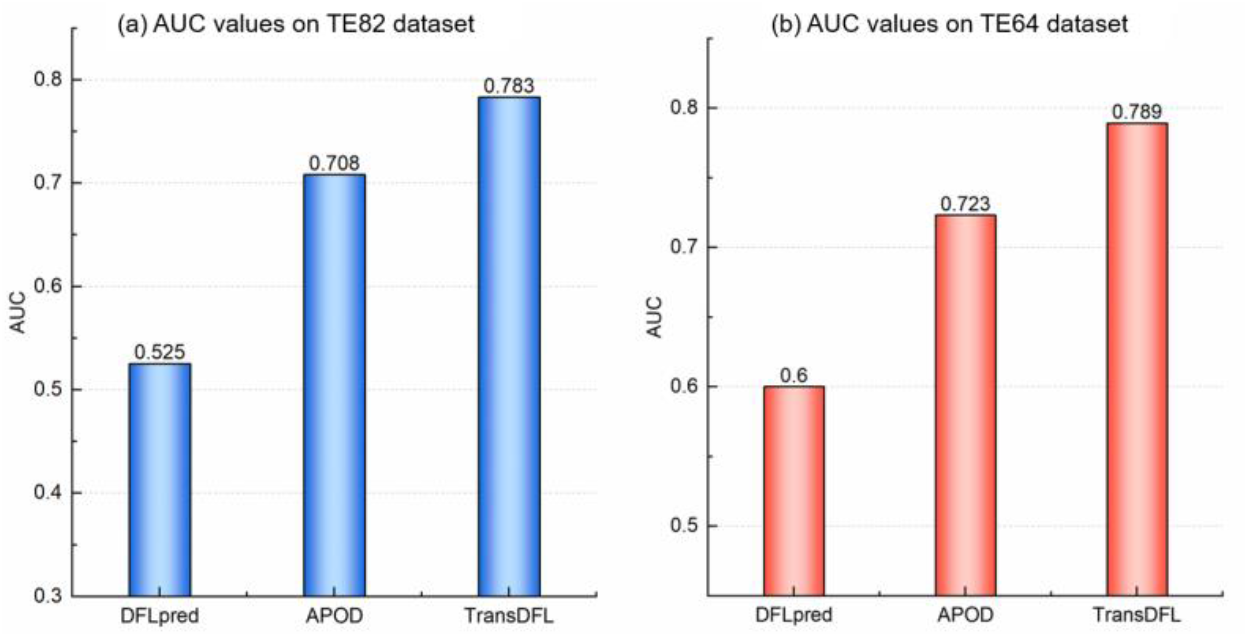
Comparison of different predictors on TE82 (a) and TE64 (b) independent test datasets in situation-II.

### Performance comparison between TransDFL and IDR predictors

In order to investigate the performance of IDR predictors for predicting DFLs, we employed 6 state-of-the-art IDR predictors to identify DFLs. Because the average length of DFLs is 47 in TR166 dataset (see **Fig. S1** in Supplementary Information S3), a target residue is considered as a DFL if its 46 neighbouring residues are disordered residues (the target is in the middle). The results of different methods evaluated in two situations on TE82 independent test dataset are shown in **Table S5 and S6** in Supplementary Information S3. From these results, we can see that the 6 IDR predictor is not effective enough for identifying DFLs compared with the specific DFL predictor TransDFL in both the two evaluation situations.

### Transfer learning obviously reduces the false positives

In order to explore the contribution of transfer learning to the performance improvement of TransDFL, we compared the false-positive rate in all prediction (FPR) and the false-positive rate in ordered region (FPR^OR^) of three different predictors. The FPR^OR^ is calculated as the ratio of the number of predicted DFL residues in ordered regions to all the positive predicted DFL residues, where the ordered residues are annotated according to the DisProt database (version 8.2). For fair evaluation, the FPRs of different predictors were compared under the same numbers of positively predicted residues, and the results were shown in **Fig. 6**. From this figure, we can see that TransDFL achieves the lowest FPR and FPR^OR^. These results are not surprising because the TransDFL employed the transfer learning framework pre-trained with the IDR dataset to capture the common characteristics between IDRs and DFLs. Therefore, compared with the other predictors only trained with DFLs and disordered residues, TransDFL can obviously reduce the incorrectly predicted DFL residues in the ordered regions so as to reduce the overall false positive predictions.

**Figure 6.**
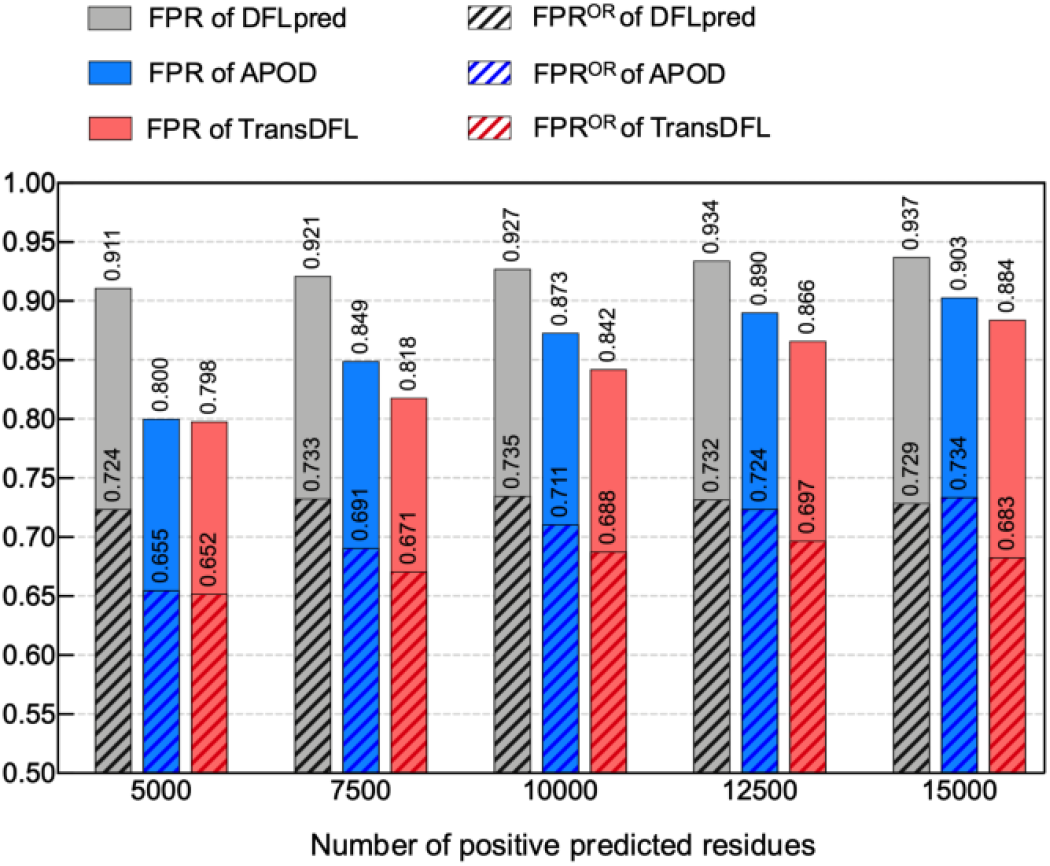
Comparison of false-positive rates of different predictors on TE82 test dataset.

The prediction results of protein (DisProt ID: DP01080, PDB ID: 1OCB) from TE82 independent test dataset obtained by different predictors were visualized and shown in **Fig. 7**, from which we can see that although most of the DFLs can be correctly predicted by the TransDFL, DFLpred, and APOD, the false positives predicted by TransDFL are obviously fewer than those of DFLpred and APOD evaluated in situation-II. The false positives predicted by TransDFL are in the disordered regions near the true DFLs, while most of the false positives predicted by APOD and DFLpred are located in the ordered regions far away from the true DFLs. These results are fully consistent with the observations in **Fig. 6**.

**Figure 7.**
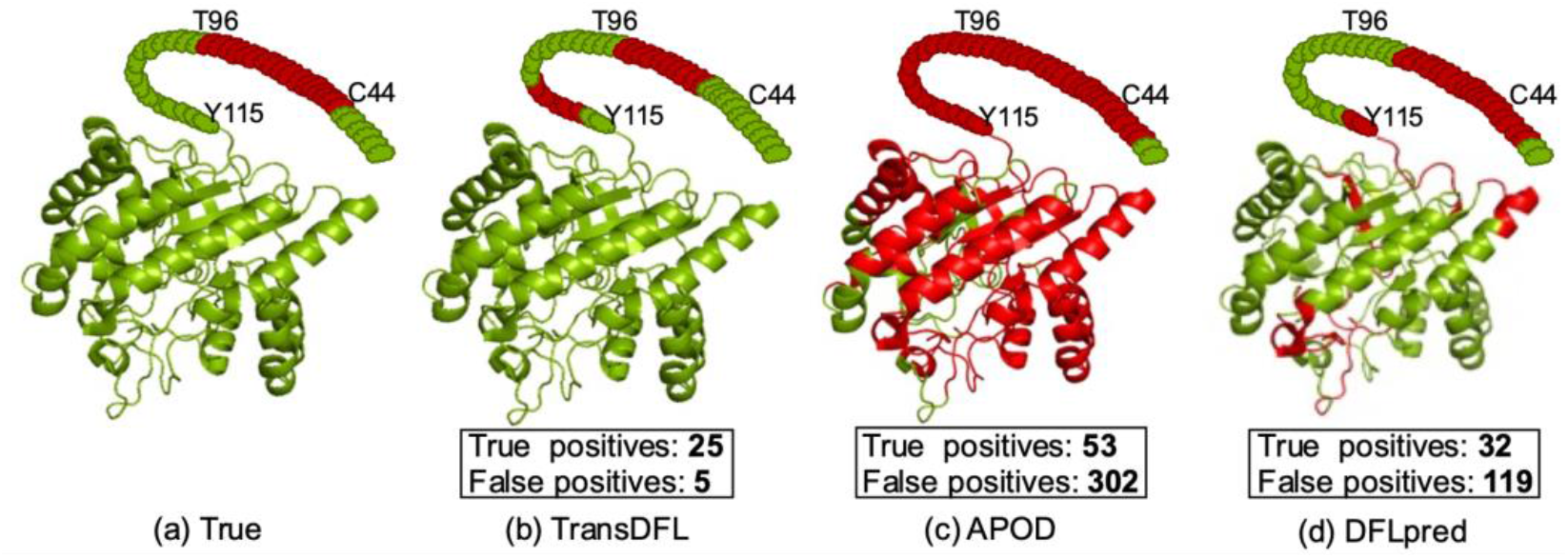
The predictive result visualization of proteins (DisProt ID: DP01080, PDB ID: 1OCB) by using the PyMOL software (https://pymol.org/2/), where the true and predicted DFL residues are shown in red. The true DFLs are shown in (**A**), and the DFLs predicted by TransDFL, APOD and DFLpred are shown in (**B**) (**C**) and (**D**), respectively.

In order to explore the contribution of model pre-trained with disordered proteins, we compare the predictive performance between TransDFL model directly trained with DFLs (TransDFL-DT) and the fine-tuned model based on pre-training with IDRs (TransDFL), and the evaluation results on two independent test datasets were shown in **Table. 3**. From this table, we can see that the model pre-trained with IDRs consistently outperformed the directly trained model (TransDFL-DT) on two datasets in both two situations, indicating that the transfer learning contributes to the predictive performance improvement of TransDFL.

**Table 3.** Performance of TransDFL predictors based on different models.

| Dataset | Model | Situation-I |  |  |  | Situation-II |  |  |  |
| --- | --- | --- | --- | --- | --- | --- | --- | --- | --- |
|  |  | Pre | Rec | MCC | AUC | Pre | Rec | MCC | AUC |
| TE82 | TransDFL-DT<sup>a</sup> | **0.552** | 0.250 | **0.298** | 0.746 | 0.010 | 0.581 | 0.120 | 0.705 |
|  | TransDFL<sup>b</sup> | 0.207 | **0.722** | 0.273 | **0.789** | **0.149** | **0.727** | **0.241** | **0.783** |
| TE64 | TransDFL-DT<sup>a</sup> | 0.254 | 0.481 | 0.166 | 0.697 | 0.080 | 0.680 | 0.113 | 0.642 |
|  | TransDFL<sup>b</sup> | **0.275** | **0.518** | **0.289** | **0.784** | **0.207** | **0.722** | **0.273** | **0.789** |

<sup>a</sup> TransDFL-DT refers to the model directly trained with DFLs training set.
<sup>b</sup> TransDFL refers to the transferred model with pre-training on the IDRs dataset.

### Sequence labelling model facilitates the stable performance on different lengths of DFL regions

In order to investigate the performance of TransDFL for predicting DFL regions with different lengths. We divided the protein sequences in the TE82 independent test dataset according to their DFL lengths (see **Fig. 8**). From this figure, we can see that compared with APOD and DFLpred, TransDFL is insensitive to the lengths of DFL regions, and achieves better and stable performance. There are two reasons: 1) TransDFL employed the sequence labelling model based on deep learning technology, which is able to capture the local and global interactions among the residues and the sequence patterns of the DFLs. In contrast, all the other two classifiers are classification-based methods predicting each residue in a separate manner. 2) Benefitted from the deep neural networks, the sequence labelling model in TransDFL captures the general disordered characteristics of DFLs from the large IDR dataset, which facilitates the DFL prediction.

**Figure 8.**
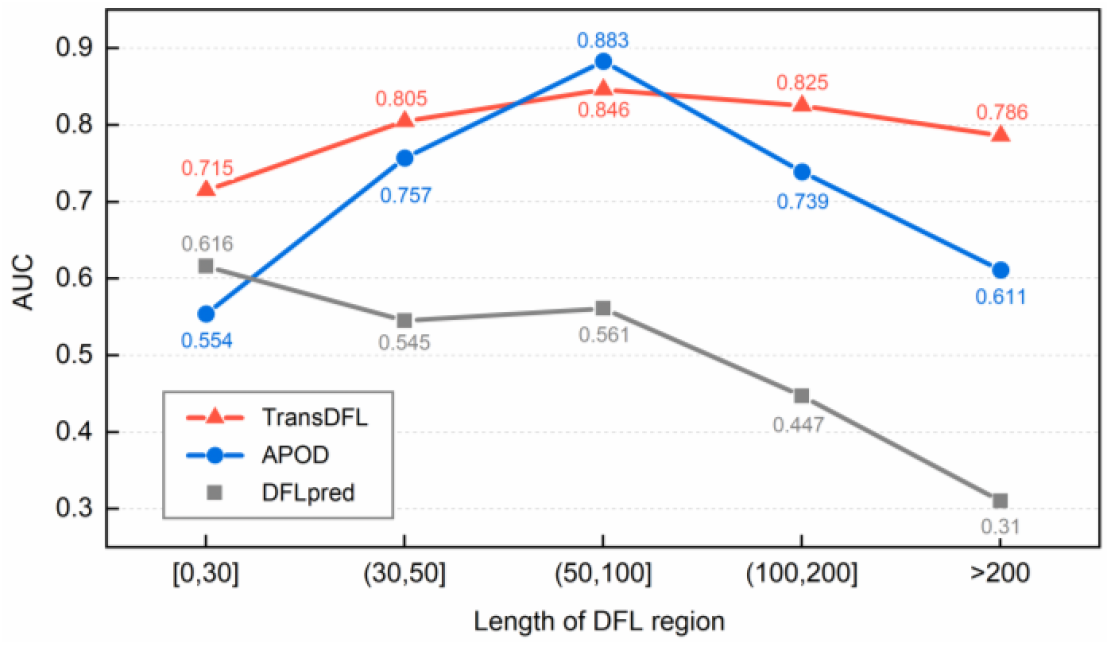
Comparison of TransDFL, APOD and DFLpred for predicting proteins with different lengths of DFL regions.

### Lower FPR leads to better performance in real-world application

According to the latest DisProt database, only 6.3% of the 4438 annotated disordered regions are DFLs [26]. There are even many more disordered proteins without DFLs in MobiDB [48]_ENREF_63_ENREF_63. As a result, the percentage of DFL residues is much lower than 6.3% in nature. Therefore, for real-world applications, it is important for a DFL predictor to deal with the extremely imbalanced problem (the number of non-DFL residues is much higher than the number of DFL residues). In this regard, seven datasets were constructed based on TE82 and TE64 with different ratios of proteins with DFLs and proteins without DFL. The performance of different predictors on the seven datasets is shown in **Fig. 9**, from which we can see that TransDFL consistently outperforms both APOD and DFLpred, especially for the datasets with fewer DFL residues. These results indicate that TransDFL is able to solve the imbalanced problem, and therefore, it is more suitable for real-world applications.

**Figure 9.**
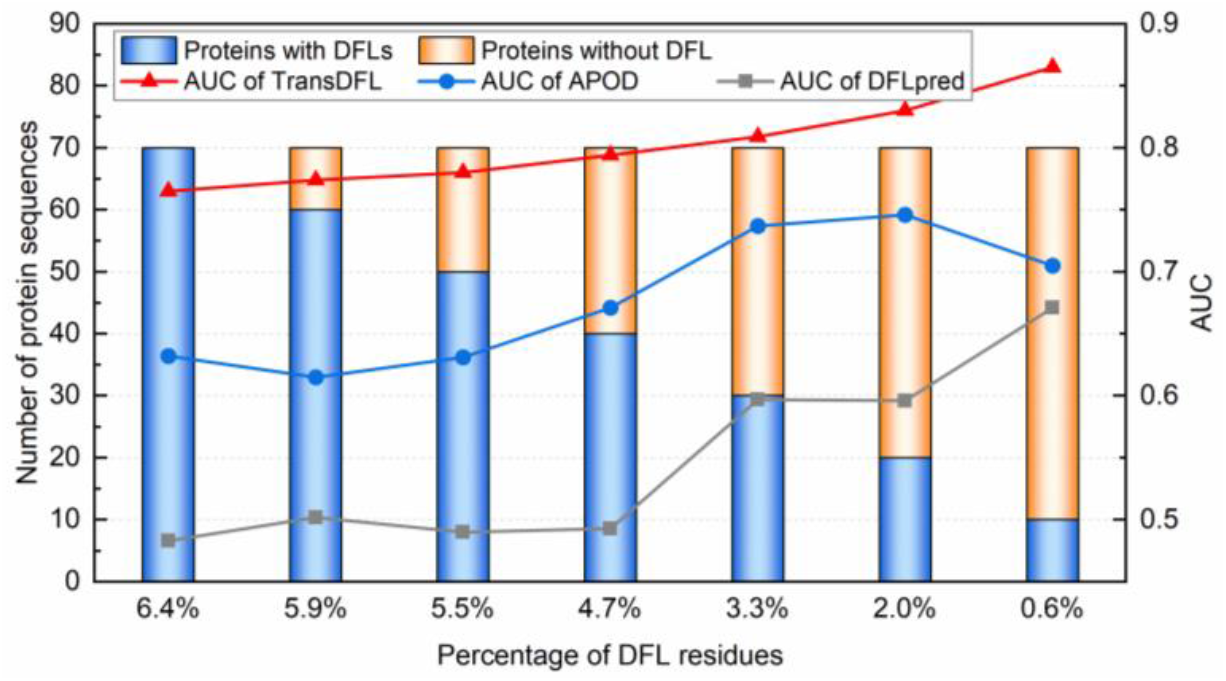
Predictive results of TransDFL, APOD and DFLpred on datasets with different percentages of DFLs.

## Conclusion

Inspired by the similarities between protein sequences and natural language sentences, we applied the transfer learning derived from the machine translation to the DFL identification, and a new predictor TransDFL was proposed. The TransDFL was constructed by transferring the state-of-the-art IDR predictor RFPR-IDP into the current DFL predictor. It has the following advantages: 1) TransDFL employed the sequence labelling model to capture the global sequence patterns of DFLs; 2) Benefitted from transfer learning, TransDFL is the first deep learning predictor for DFL prediction and achieves the state-of-the-art performance with the lowest false positive rate. The web-server of TransDFL was established, which can be freely accessed from http://bliulab.net/TransDFL.

## Data availability

Datasets used in this study is available at http://bliulab.net/TransDFL/benchmark/.

## CRediT author statement

**Yihe Pang**: Methodology, Software, Validation, Formal analysis, Data Curation, Writing - Original Draft, Writing - Review & Editing. **Bin Liu**: Conceptualization, Resources, Writing - Review & Editing, Supervision, Project administration, Funding acquisition. All authors read and approve the final manuscript.

## Acknowledgements

This work was supported by the National Natural Science Foundation of China (No. 61822306, 61861146002), National Key R&D Program of China (No. 2018AAA0100100), and the Beijing Natural Science Foundation (No. JQ19019).

## Supplementary material

**Supplementary Information S1**. The IDR dataset used for model pre-training in the source domain. In the labels, ‘1’ represents that residue is annotated as disordered residue and ‘0’ represents that residue that is annotated as ordered residue

**Supplementary Information S2**. The TE64 independent test set

**Supplementary Information S3**. The length distribution of DFLs on TR166 dataset (Figure S1). The precision-recall curves of TransDFL and the other predictors on TE82 and TE64 datasets in situation-I (Figure S2). The precision-recall curves of TransDFL and the other predictors on TE82 and TE64 datasets in situation-II (Figure S3). The performance of TransDFL based on different feature combinations on DFL validation dataset (Table S1). The performance of TransDFL based on different loss function weight coefficients on DFL validation dataset (Table S2). The hyper-parameters of RFPR-IDP (pre-trained) (Table S3). The hyper-parameters of TransDFL (fine-tuned) (Table S4). The performance of 6 state-of-the-art IDR predictors for predicting DFLs on TE82 dataset (situation-I) (Table S5). The performance of 6 state-of-the-art IDR predictors for predicting DFLs on TE82 dataset (situation-II) (Table S6).

