## Supplementary Information S1 for "TransDFL: Identification of Disordered Flexible Linkers in Proteins by Transfer Learning"

**Supplementary Information S1**. The IDR dataset used for model pre-training in the source domain. In the labels, '1' represents that residue is annotated as disordered residue and '0' represents that residue that is annotated as ordered residue.

The IDRs training dataset.

```
>DM_train26
GLVPRGSHMMEILRGSPALSAFRINKLLARFQAANLQVHNIYAEYVHFADLNAPLNDSEQAQ
LTRLLQYGPALSSHTPAGKLLLVTPRPGTISPWSSKATDIAHNCGLQQVDRLERGVAYYIEA
STLTAEQWRQVAAELHDRMMETVFSSLTDAEKLFIHHQPAPVSSVDLLGEGRQALIDANLRL
GLALAEDEIDYLQEAFTKLGRNPNDIELYMFAQANSEHCRHKIFNADWIIDGKPQPKSLFKM
IKNTFETTPDYVLSAYKDNAAVMEGSAVGRYFADHNTGRYDFHQEPAHILMKVETHNHPTAI
SPWPGAATGSGGEIRDEGATGRGAKPKAGLVGFSVSNLRIPGFEQPWEEDFGKPERIVTALD
IMTEGPLGGAAFNNEFGRPALTGYFRTYEEKVNSHNGEELRGYHKPIMLAGGIGNIRADHVQ
KGEIVVGAKLIVLGGPAMNIGLGGGAASSMASGQSDADLDFASVQRDNPEMERRCQEVIDRC
WQLGDANPILFIHDVGAGGLSNAMPELVSDGGRGGKFELRDILSDEPGMSPLEIWCNESQER
YVLAVAADQLPLFDELCKRERAPYAVIGDATEEQHLSLHDNHFDNQPIDLPLDVLLGKTPKM
TRDVQTLKAKGDALNRADITIADAVKRVLHLPTVAEKTFLVTIGDRTVTGMVARDQMVGPWQ
VPVADCAVTTASLDSYYGEAMSIGERAPVALLDFAASARLAVGEALTNIAATQIGDIKRIKL
SANWMAAAGHPGEDAGLYDAVKAVGEELCPQLGLTIPVGKDSMSMKTRWQEGNEQREMTSPL
SLVISAFARVEDVRHTLTPQLSTEDNALLLIDLGKGHNALGATALAQVYRQLGDKPADVRDV
AQLKGFYDAMQALVAARKLLAWHDRSDGGLLVTLAEMAFAGHCGVQVDIAALGDDHLAALFN
EELGGVIQVRAEDRDAVEALLAQYGLADCVHYLGQALAGDRFVITANDQTVFSESRTTLRVW
WAETTWQMQRLRDNPQCADQEHEAKANDTDPGLNVKLSFDINEDIAAPYIATGARPKVAVLR
EQGVNSHVEMAAAFHRAGFDAIDVHMSDLLGGRIGLGNFHALVACGGFSYGDVLGAGEGWAK
SILFNHRVRDEFETFFHRPQTLALGVCNGCQMMSNLRELIPGSELWPRFVRNHSDRFEARFS
LVEVTQSPSLLLQGMVGSQMPIAVSHGEGRVEVRDDAHLAALESKGLVALRYVDNFGKVTET
YPANPNGSPNGITAVTTENGRVTIMMPHPERVFRTVANSWHPENWGEDSPWMRIFRNARKQL
G
000000000000000000000000000000000000000000000000000000000000
000000000000000000000000000000000000000000000000000000000000
000000000000000000000000000000000000000000000000000000000000
000000000000000000000000000000000000000000000000000000000000
000000000000000000000000000000000000000000000000000000000000
000000000000000000000000000000000000000000000000000000000000
000000000000000000000000000000000000000000000000000000000000
000000000000000000001111111111111111110000000000000000000000
000000000000000000000000000000000000000000000000000000000000
000000000000000000000000000000000000000000000000000000000000
000000000000000000000000000000000000000000000000000000000000
000000000000000000000000000000000000000000000000000000000000
000000000000000000000000000000000000000000000000000000000000
000000000000000000000000000000000000000000000000000000000000
000000000000000000000000000000000000000000000000000000000000
000000000000000000000000000000000000000000000000000000000000
```

00000000000000000000000000000000000000000000000000000000000000
00000000000000000000000000000000000000000000000000000000000000
00000000000000000000000000000000000000000000000000000000000000
00000000000000000000000000000000000000000000000000000000000000
00000000000000000000000000000000000000000000000000000000000000
0
>DM_train27
MANVWGVRLADSLSSPTIETRTRQYTLHDLCSDLDANPGREPWKPLRNQRTNNIVAVQLFRP
LQGLVLDTQLYGFPGAFDDWERFMREKLRVLKYEVLRIYPISNYSNEHVNVFVANALVGAFL
SNQAFYDLLPLLIINDTMIGDLLGTGASLSQFFQSHGDVLEVAAGRKYLQMENYSNDDDDPP
LFAKDLSDYAKAFYSDTYEVLDRFFWTHDSSAGVLVHYDKPTNGHHYLLGTLTQMVSAPPYI
INATDAMLLESCLEQFSANVRARPAQPVTRLDQCYHLRWGAQYVGEDSLTYRLGVLSLLATN
GYQLARPIPRQLTNRWLSSFVSQIMSDGVNETPLWPQERYVQIAYDSPSVVDGATQYGYVRK
NQLRLGMRISALQSLSDTPSPVQWLPQYTIDQAAMDEGDLMVSRLTQLPLRPDYGNIWVGDA
LSYYVDYNRSHRVVLSSELPQLPDTYFDGDEQYGRSLFSLARKIGDRSLVKDTAVLKHAYQA
IDPNTGKEYLRSRQSVAYFGASAGHSGADQPLVIEPWIQGKISGVPPPSSVRQFGYDVARGA
IVDLARPFPSGDYQFVYSDVDQVVDGHDDLSISSGLVESLLSSCMHATAPGGSFVVKINFPT
RPVWHYIEQKILPNITSYMLIKPFVTNNVELFFVAFGVHQHSSLTWTSGVYFFLVDHFYRYE
TLSTISRQLPSFGYVDDGSSVTGIETISIENPGFSNMTQAARIGISGLCANVGNARKSIAIY
ESHGARVLTITSRRSPASARRKSRLRYLPLIDPRSLEVQARTILPADPVLFENVSGASPHVC
LTMMYNFEVSSAVYDGDVVLDLGTGPEAKILELIPATSPVTCVDIRPTAQPSGCWNVRTTFL
ELDYLSDGWITGVRGDIVTCMLSLGAAAAGKSMTFDAAFQQLIKVLSKSTANVVLVQVNCPT
DVVRSIKGYLEIDSTNKRYRFPKFGRDEPYSDMDALEKICRTAWPNCSITWVPLSYDLRWTR
LALLESTTLSSASIRIAELMYKYMPIMRIDIHGLPMEKRGNFIVGQNCSLVIPGFNAQDVFN
CYFNSALAFSTEDVNAAMIPQVSAQFDATKGEWTLDMVFSDAGIYTMQALVGSNANPVSLGS
FVVDSPDVDITDAWPAQLDFTIAGTDVDITVNPYYRLMTFVRIDGQWQIANPDKFQFFSSAS
GTLVMNVKLDIADKYLLYYIRDVQSRDVGFYIQHPLQLLNTITLPTNEDLFLSAPDMREWAV
KESGNTICILNSQGFVLPQDWDVLTDTISWSPSIPTYIVPPGDYTLTPL
00000000000000000000000000000000000000000000000000000000000000
00000000000000000000000000000000000000000000000000000000000000
00000000000000000000000000000000000000000000000000000000000000
00000000000000000000000000000000000000000000000000000000000000
00000000000000000000000000000000000000000000000000000000000000
00000000000000000000000000000000000000000000000000000000000000
00000000000000000000000000000000000000000000000000000000000000
00000000000000000000000000000000000000000000000000000000000000
00000000000000000000000000000000000000000000000000000000000000
00000000000000000000000000000000000000000000000000000000000000
00000000000000000000000000000000000000000000000000000000000000
00000000000000000000000000000000000000000000000000000000000000
00000000000000000000000000000000000000000000000000000000000000
00000000000000000000000000000000000000000000000000000000000000
00000000000000000000000000000000000000000000000000000000000000
00000000000000000000000000000000000000000000000000000000000000

00000000000000000000000000000000000000000000000000000000000000
00000000000000000000000000000000000000000000000000000000000000
00000000000000000000000000000000000000000000000000000000000111
10000000000000000000000000000000000000000000000000000000000000
0000000000000000000000000000000000000000000000000
>DM_train29
MSSMILTQFGPFIESISGITDQSNDVFEDAAKAFSMFTRSDVYKALDEIPFSDDAMLPIPPT
IYTKPSHDSYYYIDALNRVRRKTYQGPDDVYVPNCSIVELLEPHETLTSYGRLSEAIENRAK
DGDSQARIATTYGRIAESQARQIKAPLEKFVLALLVAEAGGSLYDPVLQKYDEIPDLSHNCP
LWCFREICRHISGPLPDRAPYLYLSAGVFWLMSPRMTSAIPPLLSDLVNLAILQQTAGLDPS
LVKLGVQICLHAAASSSYSWFILKTKSIFPQNTLHSMYESLEGGYCPNLEWLEPRSDYKFMY
MGVMPLSAKYARSAPSNDKKARELGEKYGLSSVVGELRKRTKTYVKHDFASVRYIRDAMACT
SGIFLVRTPTETVLQEYTQSPEIKVPIPQKDWTGPIGEIRILKDTTSSIARYLYRTWYLAAA
RMAAQPRTWDPLFQAIMRSQYVTARGGSGAALRESLYAINVSLPDFKGLPVKAATKIFQAAQ
LANLPFSHTSVAILADTSMGLRNQVQRRPRSIMPLNVPQQQVSAPHTLTADYINYHMNLSPT
SGSAVIEKVIPLGVYASSPPNQSINIDISACDASITWDFFLSVIMAAIHEGVASSSIGKPFM
GVPASIVNDESVVGVRAARPISGMQNMIQHLSKLYKRGFSYRVNDSFSPGNDFTHMTTTFPS
GSTATSTEHTANNSTMMETFLTVWGPEHTDDPDVLRLMKSLTIQRNYVCQGDDGLMIIDGTT
AGKVNSETIQNDLELISKYGEEFGWKYDIAYDGTAEYLKLYFIFGCRIPNLSRHPIVGKERA
NSSAEEPWPAILDQIMGVFFNGVHDGLQWQRWIRYSWALCCAFSRQRTMIGESVGYLQYPMW
SFVYWGLPLVKAFGSDPWIFSWYMPTGDLGMYSWISLIRPLMTRWMVANGYVTDRCSTVFGN
ADYRRCFNELKLYQGYYMAQLPRNPKKSGRAASREVREQFTQALSDYLMQNPELKSRVLRGR
SEWEKYGAGIIHNPPSLFDVPHKWYQGAQEAAIATREELAEMDETLMRARRHSYSSFSKLLE
AYLLVKWRMCEAREPSVDLRLPLCAGIDPLNSDPFLKMVSVGPMLQSTRKYFAQTLFMAKTV
SGLDVNAIDSALLRLRTLGADKKALTAQLLMVGLQESEADALAGKIMLQDVNTVQLARVVNL
AVPDTWMSLDFDSMFKHHVKLLPKDGRHLNTDIPPRMGWLRAILRFLGAGMVMTATGVAVDI
YLEDIHGGGRSLGQRFMTWMRQEGRSA
00000000000000000000000000000000000000000000000000000000000000
00000000000000000000000000000000000000000000000000000000000000
00000000000000000000000000000000000000000000000000000000000000
00000000000000000000000000000000000000000000000000000000000000
00000000000000000000000000000000000000000000000000000000000000
00000000000000000000000000000000000000000000000000000000000000
00000000000000000000000000000000000000000000000000000000000000
00000000000000000000000000000000000000000000000000000000000000
00000000000000000000000000000000000000000000000000000000000000
00000000000000000000000000000000000000000000000000000000000000
00000000000000000000000000000000000000000000000000000000000000
00000000000000000000000000000000000000000000000000000000000000
00000000000000000000000000000000000000000000000000000000000000
00000000000000000000000000000000000000000000000000000000000000
00000000000000000000000000000000000000000000000000000000000000
00000000000000000000000000111111110000000000000000000000000000
00000000000000000000000000000000000000000000000000000000000000

00000000000000000000000000000000000000000000000000000000000000
00000000000000000000000000000000000000000000000000000000000000
00000000000000000000000000000000000000000000000000000000000000
000000000000000000000000000
>DM_train31
ESTVTEELKEGIDAVYPSLVGTADSKAEGIKNYFKLSFTLPEEQKSRTVGSEAPLKDVAQAL
SSRARYELFTEKETANPAFNGEVIKRYKELMEHGEGIADILRSRLAKFLNTKDVGKRFAQGT
EANRWVGGKLLNIVEQDGDTFKYNEQLLQTAVLAGLQWRLTATSNTAIKDAKDVAAITGIDQ
ALLPEGLVEQFDTGMTLTEAVSSLAQKIESYWGLSRNPNAPLGYTKGIPTAMAAEILAAFVE
STDVVENIVDMSEIDPDNKKTIGLYTITELDSFDPINSFPTAIEEAVLVNPTEKMFFGDDIP
PVANTQLRNPAVRNTPEQKAALKAEQATEFYVHTPMVQFYETLGKDRILELMGAGTLNKELL
NDNHAKSLEGKNRSVEDSYNQLFSVIEQVRAQSEDISTVPIHYAYNMTRVGRMQMLGKYNPQ
SAKLVREAILPTKATLDLSNQNNEDFSAFQLGLAQALDIKVHTMTREVMSDELTKLLEGNLK
PAIDMMVEFNTTGSLPENAVDVLNTALGDRKSFVALMALMEYSRYLVAEDKSAFVTPLYVEA
DGVTNGPINAMMLMTGGLFTPDWIRNIAKGGLFIGSPNKTMNEHRSTADNNDLYQASTNALM
ESLGKLRSNYASNMPIQSQIDSLLSLMDLFLPDINLGENGALELKRGIAKNPLTITIYGSGA
RGIAGKLVSSVTDAIYERMSDVLKARAKDPNISAAMAMFGKQAASEAHAEELLARFLKDMET
LTSTVPVKRKGVLELQSTGTGAKGKINPKTYTIKGEQLKALQENMLHFFVEPLRNGITQTVG
ESLVYSTEQLQKATQIQSVVLEDMFKQRVQEKLAEKAKDPTWKKGDFLTQKELNDIQASLNN
LAPMIETGSQTFYIAGSENAEVANQVLATNLDDRMRVPMSIYAPAQAGVAGIPFMTIGTGDG
MMMQTLSTMKGAPKNTLKIFDGMNIGLNDITDASRKANEAVYTSWQGNPIKNVYESYAKFMK
NVDFSKLSPEALEAIGKSALEYDQRENATVDDIANAASLIERNLRNIALGVDIRHKVLDKVN
LSIDQMAAVGAPYQNNGKIDLSNMTPEQQADELNKLFREELEARKQKVAKA
11111111110000000000000000000000000000000000000000000000000000
00000000000000000000000000000000000000000000000000000000000000
00000000000000000000000000000000000000000000000000000000000000
00000000000000000000000000000000000000000000000000000000000000
00000000000000000000000000000000000000000000000000000000000000
00000000000000000000000000000000000000000000000000000000000000
00000000000000000000000000000000000000000000000000000000000000
00000000000000000000000000000000000000000000000000000000000000
00000000000000000000000000000000000000000000000000000000000000
00000000000000000000000000000000000000000000000000000000000000
00000000000000000000000000000000000000000000000000000000000000
00000000000000000000000000000000000000000000000000000000000000
00000000000000000000000000000000000000000000000000000000000000
00000000000000000000000000000000000000000000000000000000000000
00000000000000000000000000000000000000000000000000000000000000
00000000000000000000000000000000000000000000000000000000000000
00000000000000000000000000000000000000000000000000000000000000
000000000000000000000000000000000000000000000000011
>DM_train33
MPNFFIDRPIFAWVIAIIIMLAGGLAILKLPVAQYPTIAPPAVTISASYPGADAKTVQDTVT
QVIEQNMNGIDNLMYMSSNSDSTGTVQITLTFESGTDADIAQVQVQNKLQLAMPLLPQEVQQ

QGVSVEKSSSSFLMVVGVINTDGTMTQEDISDYVAANMKDAISRTSGVGDVQLFGSQYAMRI
WMNPNELNKFQLTPVDVITAIKAQNAQVAAGQLGGTPPVKGQQLNASIIAQTRLTSTEEFGK
ILLKVNQDGSRVLLRDVAKIELGGENYDIIAEFNGQPASGLGIKLATGANALDTAAAIRAEL
AKMEPFFPSGLKIVYPYDTTPFVKISIHEVVKTLVEAIILVFLVMYLFLQNFRATLIPTIAV
PVVLLGTFAVLAAFGFSINTLTMFGMVLAIGLLVDDAIVVVENVERVMAEEGLPPKEATRKS
MGQIQGALVGIAMVLSAVFVPMAFFGGSTGAIYRQFSITIVSAMALSVLVALILTPALCATM
LKPIAKGDHGEGKKGFFGWFNRMFEKSTHHYTDSVGGILRSTGRYLVLYLIIVVGMAYLFVR
LPSSFLPDEDQGVFMTMVQLPAGATQERTQKVLNEVTHYYLTKEKNNVESVFAVNGFGFAGR
GQNTGIAFVSLKDWADRPGEENKVEAITMRATRAFSQIKDAMVFAFNLPAIVELGTATGFDF
ELIDQAGLGHEKLTQARNQLLAEAAKHPDMLTSVRPNGLEDTPQFKIDIDQEKAQALGVSIN
DINTTLGAAWGGSYVNDFIDRGRVKKVYVMSEAKYRMLPDDIGDWYVRAADGQMVPFSAFSS
SRWEYGSPRLERYNGLPSMEILGQAAPGKSTGEAMELMEQLASKLPTGVGYDWTGMSYQERL
SGNQAPSLYAISLIVVFLCLAALYESWSIPFSVMLVVPLGVIGALLAATFRGLTNDVYFQVG
LLTTIGLSAKNAILIVEFAKDLMDKEGKGLIEATLDAVRMRLRPILMTSLAFILGVMPLVIS
TGAGSGAQNAVGTGVMGGMVTATVLAIFFVPVFFVVVRRRFSRKNEDIEHSHTVDHHHHHH
11111100000000000000000000000000000000000000000000000000000000
00000000000000000000000000000000000000000000000000000000000000
00000000000000000000000000000000000000000000000000000000000000
00000000000000000000000000000000000000000000000000000000000000
00000000000000000000000000000000000000000000000000000000000000
00000000000000000000000000000000000000000000000000000000000000
00000000000000000000000000000000000000000000000000000000000000
00000000000000000000000000000000000000000000000000000000000000
00111111111111110000000000000000000000000000000000000000000000
00000000000000000000000000000000000000000000000000000000000000
00000000000000000000000000000000000000000000000000000000000000
00000000000000000000000000000000000000000000000000000000000000
00000000000000000000000000000000000000000000000000000000000000
00000000000000000000000000000000000000000000000000000111111111
00000000000000000000000000000000000000000000000000000000000000
00000000000000000000000000000000000000000000000000000000000000
0000000000000000000000000000000000000000000001111111111111111
>DM_train34
DDPIRPPLKVARSPRPGQCQDVVQDVPNVDVQMLELYDRMSFKDIDGGVWKQGWNIKYDPLK
YNAHHKLKVFVVPHSHNDPGWIQTFEEYYQHDTKHILSNALRHLHDNPEMKFIWAEISYFAR
FYHDLGENKKLQMKSIVKNGQLEFVTGGWVMPDEANSHWRNVLLQLTEGQTWLKQFMNVTPT
ASWAIAPFGHSPTMPYILQKSGFKNMLIQRTHYSVKKELAQQRQLEFLWRQIWDNKGDTALF
THMMPFYSYDIPHTCGPDPKVCCQFDFKRMGSFGLSCPWKVPPRTISDQNVAARSDLLVDQW
KKKAELYRTNVLLIPLGDDFRFKQNTEWDVQRVNYERLFEHINSQAHFNVQAQFGTLQEYFD
AVHQAERAGQAEFPTLSGDFFTYADRSDNYWSGYYTSRPYHKRMDRVLMHYVRAAEMLSAWH
SWDGMARIEERLEQARRELSLFQHHDGITGTAKTHVVVDYEQRMQEALKACQMVMQQSVYRL
LTKPSIYSPDFSFSYFTLDDSRWPGSGVEDSRTTIILGEDILPSKHVVMHNTLPHWREQLVD
FYVSSPFVSVTDLANNPVEAQVSPVWSWHHDTLTKTIHPQGSTTKYRIIFKARVPPMGLATY
VLTISDSKPEHTSYASNLLLRKNPTSLPLGQYPEDVKFGDPREISLRVGNGPTLAFSEQGLL

KSIQLTQDSPHVPVHFKFLKYGVRSHGDRSGAYLFLPNGPASPVELGQPVVLVTKGKLESSV
SVGLPSVVHQTIMRGGAPEIRNLVDIGSLDNTEIVMRLETHIDSGDIFYTDLNGLQFIKRRR
LDKLPLQANYYPIPSGMFIEDANTRLTLLTGQPLGGSSLASGELEIMQDRRLASDDERGLGQ
GVLDNKPVLHIYRLVLEKVNNCVRPSKLHPAGYLTSAAHKASQSLLDPLDKFIFAENEWIGA
QGQFGGDHPSAREDLDVSVMRRLTKSSAKTQRVGYVLHRTNLMQCGTPEEHTQKLDVCHLLP
NVARCERTTLTFLQNLEHLDGMVAPEVCPMETAAYVSSHSS
111111111111111111000000000000000000000000000000000000000000
000000000000000000000000000000000000000000000000000000000000
000000000000000000000000000000000000000000000000000000000000
000000000000000000000000000000000000000000000000000000000000
000000000000000000000000000000000000000000000000000000000000
000000000000000000000000000000000000000000000000000000000000
000000000000000000000000000000000000000000000000000000000000
000000000000000000000000000000000000000000000000000000000000
000000000000000000000000000000000000000000000000000000000000
000000000000000000000000000000000000000000000000000000000000
000000000000000000000000000000000000000000000000000000000000
000000000000000000000000000000000000000000000000000000000000
000000000000000000000000000000000000000000000000000000000000
000000000000000000000000000000000000000000000000000000000000
000000000000000000000000000000000000000000000000000000000000
000000000000000000000000000000000000000000000000000000000000
00000000000000000000000000000000000000000
>DM_train35
ALVRDDVDYQIFRDFAENKGRFSVGATNVEVRDKNNHSLGNVLPNGIPMIDFSVVDVDKRIA
TLINPQYVVGVKHVSNGVSELHFGNLNGNMNNGNAKSHRDVSSEENRYFSVEKNEYPTKLNG
KAVTTEDQTQKRREDYYMPRLDKFVTEVAPIEASTASSDAGTYNDQNKYPAFVRLGSGSQFI
YKKGDNYSLILNNHEVGGNNLKLVGDAYTYGIAGTPYKVNHENNGLIGFGNSKEEHSDPKGI
LSQDPLTNYAVLGDSGSPLFVYDREKGKWLFLGSYDFWAGYNKKSWQEWNIYKPEFAKTVLD
KDTAGSLTGSNTQYNWNPTGKTSVISNGSESLNVDLFDSSQDTDSKKNNHGKSVTLRGSGTL
TLNNNIDQGAGGLFFEGDYEVKGTSDSTTWKGAGVSVADGKTVTWKVHNPKSDRLAKIGKGT
LIVEGKGENKGSLKVGDGTVILKQQADANNKVKAFSQVGIVSGRSTVVLNDDKQVDPNSIYF
GFRGGRLDANGNNLTFEHIRNIDDGARLVNHNTSKTSTVTITGESLITDPNTITPYNIDAPD
EDNPYAFRRIKDGGQLYLNLENYTYYALRKGASTRSELPKNSGESNENWLYMGKTSDEAKRN
VMNHINNERMNGFNGYFGEEEGKNNGNLNVTFKGKSEQNRFLLTGGTNLNGDLKVEKGTLFL
SGRPTPHARDIAGISSTKKDQHFAENNEVVVEDDWINRNFKATNINVTNNATLYSGRNVANI
TSNITASDNAKVHIGYKAGDTVCVRSDYTGYVTCTTDKLSDKALNSFNATNVSGNVNLSGNA
NFVLGKANLFGTISGTGNSQVRLTENSHWHLTGDSNVNQLNLDKGHIHLNAQNDANKVTTYN
TLTVNSLSGNGSFYYLTDLSNKQGDKVVVTKSATGNFTLQVADKTGEPTKNELTLFDASNAT
RNNLNVSLVGNTVDLGAWKYKLRNVNGRYDLYNPEVEKRNQTVDTTNITTPHHHHHHVP
000000000000000000000000000000000000000000000000000000000000
000000000000000000000000000000000000000000000000000000000000
000000000000000000000000000000000000000000000000000000000000
000000000000000000000000000000000000000000000000000000000000

000000000000000000000000000000000000000000000000000000000000
000000000000000000000000000000000000000000000000000000000000
000000000000000000000000000000000000000000000000000000000000
000000000000000000000000000000000000000000000000000000000000
000000000000000000000000000000000000000000000000000000000000
000000000000000000000000000000000000000000000000000000000000
000000000000000000000000000000000000000000000000000000000000
000000000000000000000000000000000000000000000000000000000000
000000000000000000000000000000000000000000000000000000000000
000000000000000000000000000000000000000000000000000000000000
000000000000000000000000000000000000000000000000000000000000
000000000011100000000000000000000111111111111111111111111
>DM_train39
MSDLETVAKFLAESVIASTAKTSERNLRQLETQDGFGLTLLHVIASTNLPLSTRLAGALFFK
NFIKRKWVDENGNHLLPANNVELIKKEIVPLMISLPNNLQVQIGEAISSIADSDFPDRWPTL
LSDLASRLSNDDMVTNKGVLTVAHSIFKRWRPLFRSDELFLEIKLVLDVFTAPFLNLLKTVD
EQITANENNKASLNILFDVLLVLIKLYYDFNCQDIPEFFEDNIQVGMGIFHKYLSYSNPLLE
DPDETEHASVLIKVKSSIQELVQLYTTRYEDVFGPMINEFIQITWNLLTSISNQPKYDILVS
KSLSFLTAVTRIPKYFEIFNNESAMNNITEQIILPNVTLREEDVELFEDDPIEYIRRDLEGS
DTDTRRRACTDFLKELKEKNEVLVTNIFLAHMKGFVDQYMSDPSKNWKFKDLYIYLFTALAI
NGNITNAGVSSTNNLLNVVDFFTKEIAPDLTSNNIPHIILRVDAIKYIYTFRNQLTKAQLIE
LMPILATFLQTDEYVVYTYAAITIEKILTIRESNTSPAFIFHKEDISNSTEILLKNLIALIL
KHGSSPEKLAENEFLMRSIFRVLQTSEDSIQPLFPQLLAQFIEIVTIMAKNPSNPRFTHYTF
ESIGAILNYTQRQNLPLLVDSMMPTFLTVFSEDIQEFIPYVFQIIAFVVEQSATIPESIKPL
AQPLLAPNVWELKGNIPAVTRLLKSFIKTDSSIFPDLVPVLGIFQRLIASKAYEVHGFDLLE
HIMLLIDMNRLRPYIKQIAVLLLQRLQNSKTERYVKKLTVFFGLISNKLGSDFLIHFIDEVQ
DGLFQQIWGNFIITTLPTIGNLLDRKIALIGVLNMVINGQFFQSKYPTLISSTMNSIIETAS
SQSIANLKNDYVDLDNLEEISTFGSHFSKLVSISEKPFDPLPEIDVNNGVRLYVAEALNKYN
AISGNTFLNTILPQLTQENQVKLNQLLVGN
000000000000000000000000000000000000000000000000000000000000
000000000000000000000000000000000000000000000000000000000000
000000000000000000000000000000000000000000000000000000000000
000000011000000000000000000000000000000000000000000000000000
111111000000000000000000000000000000000000000000000000000000
000000000000000000000000000000000000000000000000000000000001
111000000000000000000000000000000000011110000000000000000
000000000000000000000000000000000000000000000000000000000000
000000000000000000000000000000000000000000000000000000000000
000000000000000000000000000000000000000000000000000000000000
000000000000000000000000000000000000000000000000000000000000
000000000000000000000000000000000000000000000000000000000000
000000000000000000000000000000000000000000000000000000000000
000000000000000000000000000000000000000000000000000000000000
000000000000111110000000000000000000000000000000000000000000

00000000000000000000000000001
>DM_train40
ATPSMMPQWSYMHISGQDASEYLSPGLVQFARATETYFSLNNKFRNPTVAPTHDVTTDRSQR
LTLRFIPVDREDTAYSYKARFTLAVGDNRVLDMASTYFDIRGVLDRGPTFKPYSGTAYNALA
PKGAPNPCEWDEAATALEINLEEEDDDNEDEVDEQAEQQKTHVFGQAPYSGINITKEGIQIG
VEGQTPKYADKTFQPEPQIGESQWYETEINHAAGRVLKKTTPMKPCYGSYAKPTNENGGQGI
LVKQQNGKLESQVEMQFFSTTEATAGNGDNLTPKVVLYSEDVDIETPDTHISYMPTIKEGNS
RELMGQQSMPNRPNYIAFRDNFIGLMYYNSTGNMGVLAGQASQLNAVVDLQDRNTELSYQLL
LDSIGDRTRYFSMWNQAVDSYDPDVRIIENHGTEDELPNYCFPLGGVINTETLTKVKPKTGQ
ENGWEKDATEFSDKNEIRVGNNFAMEINLNANLWRNFLYSNIALYLPDKLKYSPSNVKISDN
PNTYDYMNKRVVAPGLVDCYINLGARWSLDYMDNVNPFNHHRNAGLRYRSMLLGNGRYVPFH
IQVPQKFFAIKNLLLLPGSYTYEWNFRKDVNMVLQSSLGNDLRVDGASIKFDSICLYATFFP
MAHNTASTLEAMLRNDTNDQSFNDYLSAANMLYPIPANATNVPISIPSRNWAAFRGWAFTRL
KTKETPSLGSGYDPYYTYSGSIPYLDGTFYLNHTFKKVAITFDSSVSWPGNDRLLTPNEFEI
KRSVDGEGYNVAQCNMTKDWFLVQMLANYNIGYQGFYIPESYKDRMYSFFRNFQPMSRQVVD
DTKYKDYQQVGILHQHNNSGFVGYLAPTMREGQAYPANFPYPLIGKTAVDSITQKKFLCDRT
LWRIPFSSNFMSMGALTDLGQNLLYANSAHALDMTFEVDPMDEPTLLYVLFEVFDVVRVHRP
HRGVIETVYLRTPFSAGNATT
111100000000000000000000000000000000000000000000000000000000
000000000000000000000000000000000000000000000000000000000000
000000000001111111111111111111111111111100000000000000000000
111111000000000000000000000000000000000000000000000000000000
001111111000000000000111111110000000000000000000000000000000
000000000000000000000000000000000000000000000000000000000000
000000000000000000000000000000000000000000000000000000001111
110000000000000000000000000000000000000000000000000000000000
000000000000000000000000000000000000000000000000000000000000
000000000000000000000000000000000000000000000000000000000000
000000000000000000000000000000000000000000000000000000000000
000000000000000000000000000000000000000000000000000000000000
000000000000000000000000000000000000000000000000000000000000
000000000000000000000000000000000000000000000000000000000000
000000000000000000000000000000000000000000000000000000000000
000000000000000011111
>DM_train43
MVAKKWVYYFGGGNADGNKNMKELLGGKGANLAEMVNLGIPVPPGFTITTEACKTYQETETI
PQEVADQVRENVSRVEKEMGAKFGDPANPLLFSVRSGAAASMPGMMDTVLNLGLNKVTVDAW
VRRAPRLERFVYDSYRRFITMYADIVMQVGREDFEEALSRMKERRGTKFDTDLTASDLKELC
DGYLELFELKTGCSFPQDPVMQLFAAIKAVFRSWGNPRATIYRRMNNITGLLGTAVNVQAMV
FGNINDRSATGVAFSRSPSTGENFFFGEYLVNAQGEDVVAGIRTPQQINHSLSLRWAKAHGV
GEEERRKRYPSMEEAMPENYRLLCDVRKRLENHYRDMQDLEFTVQDGRLWLLQCRNGKRTIH
AAVRIAIDMVNEGLISREEAVLRIDPYQVDHLMHPNLEPGAEKANKPIGRGLAASPGAAVGQ
VVFDAESAKEWSGRGKKVIMVRLETSPEDLAGMDAACGILTARGGMTSHAAVVARGMGKCCV
SGCGDMVIRGKSFKLNGSVFREGDYITIDGSKGLIYAGKLKLRSPDLKGSFQTILQWCQEMK

RLGVRTNADTPADAAKARSFGAEGVGLCRTEHMFFEGSRINFIREMILADSASGRKAALDKL
LPIQRADFVGILRAMRGLPVTIRLLDPPLHEFVPHDAAAQFELAQKLGMPAEKVRNRVNALH
ELNPMLGHRGCRLGITYPEIYNMQVRAIIEAAIAVSEEGSSVIPEIMVPLVGKKEELSLIRE
EVVKTAEAVITKSGKRVHYTVGTMIEVPRAAVTADSIAQKADFFSFGTNDLTQMGCGFSRDD
AGPFLRHYGNLGIYAQDPFQSIDQEGIGELVRIAVTKGRRVKPMLKMGICGEHGGDPATIGF
CHKVGLDYVSCSPFRVPVAIVAAAHASIKDRRAAMKARKGFAAKL
000000000000000000000000000000000000000000000000000000000000
000000000000000000000000000000000000000011111000000000000000
000000000000000000000000000000000000000000000000000000000000
000000000000000000000000000000000000000000000000000000000000
000000000000000000000000000000000000000000000000000000000000
000000000000000000000000000000000000000000000000000000000000
000000000000000000000000000000000000000000000000000000000000
000000000000000000000000000000000000000000000000000000000000
000000000000000000000000000000000000000000000000000000000000
000000000000000000000000000000000000000000000000000000000000
000000000000000000000000000000000000000000000000000000000000
000000000000000000000000000000000000000000000000000000000000
000000000000000000000000000000000000000000000000000000000000
000000000000000000000000000000000000000000000000000000000000
000000000000000000000000000000000111111111
>DM_train45
MVIASVEDGGDGDTSKDDWLWYKQPASQTDATATAGGNYGNPDNNRWQQTTLPFGNGKIGGT
VWGEVSRERVTFNEETLWTGGPGSSTSYNGGNNETKGQNGATLRALNKQLANGAETVNPGNL
TGGENAAEQGNYLNWGDIYLDYGFNDTTVTEYRRDLNLSKGKADVTFKHDGVTYTREYFASN
PDNVMVARLTASKAGKLNFNVSMPTNTNYSKTGETTTVKGDTLTVKGALGNNGLLYNSQIKV
VLDNGEGTLSEGSDGASLKVSDAKAVTLYIAAATDYKQKYPSYRTGETAAEVNTRVAKVVQD
AANKGYTAVKKAHIDDHSAIYDRVKIDLGQSGHSSDGAVATDALLKAYQRGSATTAQKRELE
TLVYKYGRYLTIGSSRENSQLPSNLQGIWSVTAGDNAHGNTPWGSDFHMNVNLQMNYWPTYS
ANMGELAEPLIEYVEGLVKPGRVTAKVYAGAETTNPETTPIGEGEGYMAHTENTAYGWTAPG
QSFSWGWSPAAVPWILQNVYEAYEYSGDPALLDRVYALLKEESHFYVNYMLHKAGSSSGDRL
TTGVAYSPEQGPLGTDGNTYESSLVWQMLNDAIEAAKAKGDPDGLVGNTTDCSADNWAKNDS
GNFTDANANRSWSCAKSLLKPIEVGDSGQIKEWYFEGALGKKKDGSTISGYQADNQHRHMSH
LLGLFPGDLITIDNSEYMDAAKTSLRYRCFKGNVLQSNTGWAIGQRINSWARTGDGNTTYQL
VELQLKNAMYANLFDYHAPFQIDGNFGNTSGVDEMLLQSNSTFTDTAGKKYVNYTNILPALP
DAWAGGSVSGLVARGNFTVGTTWKNGKATEVRLTSNKGKQAAVKITAGGAQNYEVKNGDTAV
NAKVVTNADGASLLVFDTTAGTTYTITKKAS
111111111000000000000000000000000000000000000000000000000000
000000000000000000000000000000000000000000000000000000000000
000000000000000000000000000000000000000000000000000000000000
000000000000000000000000000000000000000000000000000000000000
000111000000000000000000000000000000000000000000000000000000
000000000000000000000000000000000000000000000000000000000000
000000000000000000000000000000000000000000000000000000000000

000000000000000000000000000000000000000000000000000000000000
000000000000000000000000000000000000000000000000000000000000
000000000000000000000000000000000000000000000000000000000000
000000000000000000000000000000000000000000000000000000000000
000000000000000000000000000000000000000000000000000000000000
000000000000000000000000000000000000000000000000000000000000
000000000000000000000000000000000000000000000000000000011110
000000000000000000000000000011
>DM_train47
MTQQPQAKYRHDYRAPDYQITDIDLTFDLDAQKTVVTAVSQAVRHGASDAPLRLNGEDLKLV
SVHINDEPWTAWKEEEGALVISNLPERFTLKIINEISPAANTALEGLYQSGDALCTQCEAEG
FRHITYYLDRPDVLARFTTKIIADKIKYPFLLSNGNRVAQGELENGRHWVQWQDPFPKPCYL
FALVAGDFDVLRDTFTTRSGREVALELYVDRGNLDRAPWAMTSLKNSMKWDEERFGLEYDLD
IYMIVAVDFFNMGAMENKGLNIFNSKYVLARTDTATDKDYLDIERVIGHEYFHNWTGNRVTC
RDWFQLSLKEGLTVFRDQEFSSDLGSRAVNRINNVRTMRGLQFAEDASPMAHPIRPDMVIEM
NNFYTLTVYEKGAEVIRMIHTLLGEENFQKGMQLYFERHDGSAATCDDFVQAMEDASNVDLS
HFRRWYSQSGTPIVTVKDDYNPETEQYTLTISQRTPATPDQAEKQPLHIPFAIELYDNEGKV
IPLQKGGHPVNSVLNVTQAEQTFVFDNVYFQPVPALLCEFSAPVKLEYKWSDQQLTFLMRHA
RNDFSRWDAAQSLLATYIKLNVARHQQGQPLSLPVHVADAFRAVLLDEKIDPALAAEILTLP
SVNEMAELFDIIDPIAIAEVREALTRTLATELADELLAIYNANYQSEYRVEHEDIAKRTLRN
ACLRFLAFGETHLADVLVSKQFHEANNMTDALAALSAAVAAQLPCRDALMQEYDDKWHQNGL
VMDKWFILQATSPAANVLETVRGLLQHRSFTMSNPNRIRSLIGAFAGSNPAAFHAEDGSGYL
FLVEMLTDLNSRNPQVASRLIEPLIRLKRYDAKRQEKMRAALEQLKGLENLSGDLYEKITKA
LA
111100000000000000000000000000000000000000000000000000000000
000000000000000000000000000000000000000000000000000000000000
000000000000000000000000000000000000000000000000000000000000
000000000000000000000000000000000000000000000000000000000000
000000000000000000000000000000000000000000000000000000000000
000000000000000000000000000000000000000000000000000000000000
000000000000000000000000000000000000000000000000000000000000
000000000000000000000000000000000000000000000000000000000000
000000000000000000000000000000000000000000000000000000000000
000000000000000000000000000000000000000000000000000000000000
000000000000000000000000000000000000000000000000000000000000
000000000000000000000000000000000000000000000000000000000000
000000000000000000000000000000000000000000000000000000000000
000000000000000000000000000000000000000000000000000000000000
00
>DM_train48
SAECPVVNELERINCIPDQPPTKATCDQRGCCWNPQGAVSVPWCYYSKNHSYHVEGNLVNTN
AGFTARLKNLPSSPVFGSNVDNVLLTAEYQTSNRFHFKLTDQTNNRFEVPHEHVQSFSGNAA
ASLTYQVEISRQPFSIKVTRRSNNRVLFDSSIGPLLFADQFLQLSTRLPSTNVYGLGEHVHQ
QYRHDMNWKTWPIFNRDTTPNGNGTNLYGAQTFFLCLEDASGLSFGVFLMNSNAMEVVLQPA

PAITYRTIGGILDFYVFLGNTPEQVVQEYLELIGRPALPSYWALGFHLSRYEYGTLDNMREV
VERNRAAQLPYDVQHADIDYMDERRDFTYDSVDFKGFPEFVNELHNNGQKLVIIVDPAISNN
SSSSKPYGPYDRGSDMKIWVNSSDGVTPLIGEVWPGQTVFPDYTNPNCAVWWTKEFELFHNQ
VEFDGIWIDMNEVSNFVDGSVSGCSTNNLNNPPFTPRILDGYLFCKTLCMDAVQHWGKQYDI
HNLYGYSMAVATAEAAKTVFPNKRSFILTRSTFAGSGKFAAHWLGDNTATWDDLRWSIPGVL
EFNLFGIPMVGPDICGFALDTPEELCRRWMQLGAFYPFSRNHNGQGYKDQDPASFGADSLLL
NSSRHYLNIRYTLLPYLYTLFFRAHSRGDTVARPLLHEFYEDNSTWDVHQQFLWGPGLLITP
VLDEGAEKVMAYVPDAVWYDYETGSQVRWRKQKVEMELPGDKIGLHLRGGYIFPTQQPNTTT
LASRKNPLGLIIALDENKEAKGELFWDDGETKDTVANKVYLLCEFSVTQNRLEVNISQSTYK
DPNNLAFNEIKILGTEEPSNVTVKHNGVPSQTSPTVTYDSNLKVAIITDIDLLLGEAYTVEW
111111000000000000000000000000000000000000000000000000000000
000000000000000000000000000000000000000000000000000000000000
000000000000000000000000000000000000000000000000000000000000
000000000000000000000000000000000000000000000000000000000000
000000000000000000000000000000000000000000000000000000000000
000000000000000000000000000000000000000000000000000000000000
000000000000000000000000000000000000000000000000000000000000
000000000000000000000000000000000000000000000000000000000000
000000000000000000000000000000000000000000000000000000000000
000000000000000000000000000000000000000000000000000000000000
000000000000000000000000000000000000000000000000000000000000
000000000000000000000000000000000000000000000000000000000000
000000000000000000000000000000000000000000000000000000000000
000000000000000000000000000001000000000000000000000000000000
>DM_train50
MAKPLTDQEKRRQISIRGIVGVENVAELKKSFNRHLHFTLVKDRNVATTRDYYFALAHTVRD
HLVGRWIRTQQHYYDKCPKRVYYLSLEFYMGRTLQNTMINLGLQNACDEAIYQLGLDIEELE
EIEEDAGLGNGGLGRLAACFLDSMATLGLAAYGYGIRYEYGIFNQKIRDGWQVEEADDWLRY
GNPWEKSRPEFMLPVHFYGKVEHTNTGTKWIDTQVVLALPYDTPVPGYMNNTVNTMRLWSAR
APNDFNLRDFNVGDYIQAVLDRNLAENISRVLYPNDNFFEGKELRLKQEYFVVAATLQDIIR
RFKASKFGSTRGAGTVFDAFPDQVAIQLNDTHPALAIPELMRIFVDIEKLPWSKAWELTQKT
FAYTNHTVLPEALERWPVDLVEKLLPRHLEIIYEINQKHLDRIVALFPKDVDRLRRMSLIEE
EGSKRINMAHLCIVGSHAVNGVAKIHSDIVKTKVFKDFSELEPDKFQNKTNGITPRRWLLLC
NPGLAELIAEKIGEDYVKDLSQLTKLHSFLGDDVFLRELAKVKQENKLKFSQFLETEYKVKI
NPSSMFDVQVKRIHEYKRQLLNCLHVITMYNRIKKDPKKLFVPRTVIIGGKAAPGYHMAKMI
IKLITSVADVVNNDPMVGSKLKVIFLENYRVSLAEKVIPATDLSEQISTAGTEASGTGNMKF
MLNGALTIGTMDGANVEMAEEAGEENLFIFGMRIDDVAALDKKGYEAKEYYEALPELKLVID
QIDNGFFSPKQPDLFKDIINMLFYHDRFKVFADYEAYVKCQDKVSQLYMNPKAWNTMVLKNI
AASGKFSSDRTIKEYAQNIWNVEPSDLKISLSNESNKVNGN
111111111111111111111110000000000000000000000000000000000000
000000000000000000000000000000000000000000000000000000000000
000000000000000000000000000000000000000000000000000000000000
000000000000000000000000000000000000000000000000000000000000
001111111111000000000000000000000000000000000000000000000000

0000111111111111000000000000000000000000000000000000000000000
000000000000000000000000000000000000000000000000000000000000
000000000000000000000000000000000000000000000000000000000000
000000000000000000000000000000000000000000000000000000000000
000000000000000000000000000000000000000000000000000000000000
000000000000000000000000000000000000000000000000000000000000
000000000000000000000000000000000000000000000000000000000000
000000000000000000000000000000000000000000000000000000000000
0000000000000000000000001111111111111111
>DM_train51
MFSAGHKIKGTVVLMPKNELEVNPDGSAVDNLNAFLGRSVSLQLISATKADAHGKGKVGKDT
FLEGINTSLPTLGAGESAFNIHFEWDGSMGIPGAFYIKNYMQVEFFLKSLTLEAISNQGTIR
FVCNSWVYNTKLYKSVRIFFANHTYVPSETPAPLVSYREEELKSLRGNGTGERKEYDRIYDY
DVYNDLGNPDKSEKLARPVLGGSSTFPYPRRGRTGRGPTVTDPNTEKQGEVFYVPRDENLGH
LKSKDALEIGTKSLSQIVQPAFESAFDLKSTPIEFHSFQDVHDLYEGGIKLPRDVISTIIPL
PVIKELYRTDGQHILKFPQPHVVQVSQSAWMTDEEFAREMIAGVNPCVIRGLEEFPPKSNLD
PAIYGDQSSKITADSLDLDGYTMDEALGSRRLFMLDYHDIFMPYVRQINQLNSAKTYATRTI
LFLREDGTLKPVAIELSLPHSAGDLSAAVSQVVLPAKEGVESTIWLLAKAYVIVNDSCYHQL
MSHWLNTHAAMEPFVIATHRHLSVLHPIYKLLTPHYRNNMNINALARQSLINANGIIETTFL
PSKYSVEMSSAVYKNWVFTDQALPADLIKRGVAIKDPSTPHGVRLLIEDYPYAADGLEIWAA
IKTWVQEYVPLYYARDDDVKNDSELQHWWKEAVEKGHGDLKDKPWWPKLQTLEDLVEVCLII
IWIASALHAAVNFGNYPYGGLIMNRPTASRRLLPEKGTPEYEEMINNHEKAYLRTITSKLPT
LISLSVIEILSTHASDEVYLGQRDNPHWTSDSKALQAFQKFGNKLKEIEEKLVRRNNDPSLQ
GNRLGPVQLPYTLLYPSSEEGLTFRGIPNSISI
111111000000000000001111111111100000000000000000000000000000
000000000000000000000000000000000000000000000000000111111000
000000000000000000000000000000000000000000000000000000000000
000000000000000000000000000000000000000000000000000000000000
000000000000000000000000000000000000000000000000000000000000
000000000000000000000000000000000000000000000000000000000000
000000000000000000000000000000000000000000000000000000000000
000000000000000000000000000000000000000000000000000000000000
000000000000000000000000000000000000000000000000000000000000
000000000000000000000000000000000000000000000000000000000000
000000000000000000000000000000000000000000000000000000000000
000000000000000000000000000000000000000000000000000000000000
000000000000000000000000000000000000000000000000000000000000
000000000000000000000000000000000
>DM_train52
GCPNDRITLPPANAQRTNMTCHFCIVGCGYHVYKWPELEEGGRAPEQNALGLDFRKQLPPLA
STLTPAMTNVVTEHDGARYDIMVVPDKACVVNSGLSSTRGGKMASYMYTPTGDGKERLSAPR
LYAADEWVDTTWDHAMALYAGLIKKTLDSDGPQGVFFSCFDHGGAGGGFENTWGTGKLMFSA
IQTPMVRIHNRPAYNSECHATREMGIGELNNAYEDAQLADVIWSIGNNPYESQTNYFLNHWL
PNLQGATTSKKKERFPNENFPQARIIFVDPRETPSVAIARHVAGNDRVLHLAIEPGTDTALF

NGLFTYVVEQGWIDKPFIEAHTKGFDDAVKTNRLSLDECSNITGVPVDMLKRAAEWSYKPKA
SGQAPRTMHAYEKGIIWGNDNYVIQSALLDLVIATHNVGRRGTGCVRMGGHQEGYTRPPYPG
DKKIYIDQELIKGKGRIMTWWGCNNFQTSNNAQALREAILQRSAIVKQAMQKARGATTEEMV
DVIYEATQNGGLFVTSINLYPTKLAEAAHLMLPAAHPGEMNLTSMNGERRIRLSEKFMDPPG
TAMADCLIAARIANALRDMYQKDGKAEMAAQFEGFDWKTEEDAFNDGFRRAGQPGAPAIDSQ
GGSTGHLVTYDRLRKSGNNGVQLPVVSWDESKGLVGTEMLYTEGKFDTDDGKAHFKPAPWNG
LPATVQQQKDKYRFWLNNGRNNEVWQTAYHDQYNSLMQERYPMAYIEMNPDDCKQLDVTGGD
IVEVYNDFGSTFAMVYPVAEIKRGQTFMLFGYVNGIQGDVTTDWTDRDIIPYYKGTWGDIRK
VGSMSEFKRTVSFKSRRFG
111110000000000000000000000000000000000000000000000000000000
000000000000000000000000000000000000000000000000000000000000
000000000000000000000000000000000000000000000000000000000000
000000000000000000000000000000000000000000000000000000000000
000000000000000000000000000000000000000000000000000000000000
000000000000000000000000000000000000000000000000000000000000
000000000000000000000000000000000000000000000000000000000000
000000000000000000000000000000000000000000000000000000000000
000000000000000000000000000000000000000000000000000000000000
000000000000000000000000000000000000000000000000000000000000
000000000000000000000000000000000000000000000000000000000000
000000000000000000000000000000000000000000000000000000000000
000000000000000000000000000000000000000000000000000000000000
0000000000000000000
>DM_train54
SEHPQPVTTQIEKSVNTALNKNYVFNKADYQYTLTNPSLGKIVGGILYPNATGSTTVKISDK
SGKIIKEVPLSVTASTEDNFTKLLDKWNDVTIGNYVYDTNDSNMQKLNQKLDETNAKNIEAI
KLDSNRTFLWKDLDNLNNSAQLTATYRRLEDLAKQITNPHSTIYKNEKAIRTVKESLAWLHQ
NFYNVNKDIEGSANWWDFEIGVPRSITGTLSLMNNYFTDAEIKTYTDPIEHFVPDAEYFRKT
LVNPFKALGGNLVDMGRVKIIEGLLRKDNTIIEKTSHSLKNLFTTATKAEGFYADGSYIDHT
NVAYTGAYGNVLIDGLTQLLPIIQETDYKISNQELDMVYKWINQSFLPLIVKGELMDMSRGR
SISREAASSHAAAVEVLRGFLRLANMSNEERNLDLKSTIKTIITSNKFYNVFNNLKSYSDIA
NMNKLLNDSTVATKPLKSNLSTFNSMDRLAYYNAKKDFGFALSLHSKRTLNYEGMNDENTRG
WYTGDGMFYIYNSDQSHYSNHFWPTVNPYKMAGTTEKDAKREDTTKEFMSKHSKDAKEKTGQ
VTGTSDFVGSVKLNDHFALAAMDFTNWDRTLTAQKGWVILNDKIVFLGSNIKNTNGIGNVST
TIDQRKDDSKTPYTTYVNGKTIDLKQASSQQFTDTKSVFLESKEPGRNIGYIFFKNSTIDIE
RKEQTGTWNSINRTSKNTSIVSNPFITISQKHDNKGDSYGYMMVPNIDRTSFDKLANSKEVE
LLENSSKQQVIYDKNSQTWAVIKHDNQESLINNQFKMNKAGLYLVQKVGNDYQNVYYQPQTM
TKTDQLAI
000000000000000000000000000000000000000000000000000000000000
000000000000000000000000000000000000000000000000000000000000
000000000000000000000000000000000000000000000000000000000000
000000000000000000000000000000000000000000000000000000000000
000000000000000000000000000000000000000000000000000000000000
000000000000000000000000000000000000000000000000000000000000

000000000000000000000000000000000000000000000000000000000000
000000000000000000000000000000000000000000000000000000000000
000000000000000000000000000000000000000000000111111110000000
000000000000000000000000000000000000000000000000000000000000
000000000000000000000011111100000000000000000000000000000000
000000000000000000000000000000000111100000000000000000000000
000000000000000000000000000000000000000000000000000000000000
00000000
>DM_train56
MKYGYFDNDNREYVITRPDVPAPWTNYLGTEKFCTVISHNAGGYSFYNSPEYNRVTKFRPNA
TFDRPGHYVYLRDDDSGDYWSISWQPVAKSLDEAQYQIRHGLSYSKFQCDYNGIHARKTLFV
PKGEDAEIWDVVIKNTSDQVRTISAFSFVEFSFSHIQSDNQNHQMSLYSAGTAYRPGLIEYD
LYYNTDDFEGFYYLASTFDPDSYDGQRDRFLGLYRDEANPLAVEQGRCSNSAQTCYNHCGSL
HKQFTLQPGEEIRFAYILGIGKGNGERLREHYQDVANIDAAFAAIKAHWDERCAKFQVKSPN
QGLDTMINAWTLYQAETCVVWSRFASFIEVGGRTGLGYRDTAQDAISVPHANPEMTRKRIVD
LLRGQVKAGYGLHLFDPDWFDPEKEDVAPSKSPTVVPTPSDEDKIHGIKDTCSDDHLWLIPT
ICKYVMETGETSFFDQMIPYADGGEASVYEHMKAALDFSAEYVGQTGICKGLRADWNDCLNL
GGGESSMVSFLHFWALQEFIDLAKFLGKDQDVNTYTEMAANVREACETHLWDDEGGWYIRGL
TKNGDKIGTAQQQEGRVHLESNTLAVLSGLASQERGEQAMDAVDEHLFSPYGLHLNAPSFST
PNDDIGFVTRVYQGVKENGAIFSHPNPWAWVAETKLGRGDRAMKFYDALNPYNQNDIIEKRI
AEPYSYVQFIMGRDHQDHGRANHPWLTGTSGWAYFAVTNYILGVQSGFTGLSVDPCIPSDWP
GFEVTRQWRGATYHIQVENPDHVSKGVKSITLNGAPIQGRIPPQAQGSDNQVVVVLG
000000000000000000000000000000000000000000000000000000000000
000000000000000000000000000000000000000000000000000000000000
000000000000000000000000000000000000000000000000000000000000
000000000000000000000000000000000000000000000000000000000000
000000000000000000000000000000000000000000000000000000000000
000000000000000000000000000000000000000000000000000000000000
000000000000000000000011111111111111111111100000000000000000
000000000000000000000000000000000000000000000000000000000000
000000000000000000000000000000000000000000000000000000000000
000000000000000000000000000000000000000000000000000000000000
000000000000000000000000000000000000000000000000000000000000
000000000000000000000000000000000000000000000000000000000000
0000000000000000000000000000000000000000000000000000000
>DM_train58
MRVPFSWLKAYVPELESPEVLEERLAGLGFETDRIERVFPIPRGVVFARVLEAHPIPGTRLK
RLVLDAGRTVEVVSGAENARKGIGVALALPGTELPGLGQKVGERVIQGVRSFGMALSPRELG
VGEYGGGLLEFPEDALPPGTPLSEAWPEEVVLDLEVTPNRPDALGLLGLARDLHALGYALVE
PEAALKAEALPLPFALKVEDPEGAPHFTLGYAFGLRVAPSPLWMQRALFAAGMRPINNVVDV
TNYVMLERAQPMHAFDLRFVGEGIAVRRAREGERLKTLDGVERTLHPEDLVIAGWRGEESFP
LGLAGVMGGAESEVREDTEAIALEVACFDPVSIRKTARRHGLRTEASHRFERGVDPLGQVPA
QRRALSLLQALAGARVAEALLEAGSPKPPEAIPFRPEYANRLLGTSYPEAEQIAILKRLGCR
VEGEGPTYRVTPPSHRLDLRLEEDLVEEVARIQGYETIPLALPAFFPAPDNRGVEAPYRKEQ

RLREVLSGLGFQEVYTYSFMDPEDARRFRLDPPRLLLLNPLAPEKAALRTHLFPGLVRVLKE
NLDLDRPERALLFEVGRVFREREETHLAGLLFGEGVGLPWAKERLSGYFLLKGYLEALFARL
GLAFRVEAQAFPFLHPGVSGRVLVEGEEVGFLGALHPEIAQELELPPVHLFELRLPLPDKPL
AFQDPSRHPAAFRDLAVVVPAPTPYGEVEALVREAAGPYLESLALFDLYQGPPLPEGHKSLA
FHLRFHPKRTLRDEEVEEAVSRVAEALRARGFGLRGLDTP
00000000000000000000000000000000000000000000000000000000000000
00000000000000000000000000000000000000000000000000000000000000
00000000000000000000000000000000000000000000000000000000000000
00000000000000000000000000000000000000000000000000000000000000
00000000000000000000000000000000000000000000000000000000000000
00000000000000000000000000000000000000000000000000000000000000
00000000000000000000000000000000000000000000000000000000000000
00000000000000000000000000000000000000000000000000000000000000
00000000000000000000000000000000000000000000000000000000000000
00000000000000000000000000000000000000000000000000000000000000
00000000000000000000000000000000000000000000000000000000000000
00000000000000000000000000000000000000000000000000000000000000
0000000000000000000000000000001111111111
>DM_train59
MRFKNVKKTALMLAMFGMATSSNAALFDYNATGDTEFDSPAKQGWMQDNTNNGSGVLTNADG
MPAWLVQGIGGRAQWTYSLSTNQHAQASSFGWRMTTEMKVLSGGMITNYYANGTQRVLPIIS
LDSSGNLVVEFEGQTGRTVLATGTAATEYHKFELVFLPGSNPSASFYFDGKLIRDNIQPTAS
KQNMIVWGNGSSNTDGVAAYRDIKFEIQGDVIFRGPDRIPSIVASSVTPGVVTAFAEKRVGG
GDPGALSNTNDIITRTSRDGGITWDTELNLTEQINVSDEFDFSDPRPIYDPSSNTVLVSYAR
WPTDAAQNGDRIKPWMPNGIFYSVYDVASGNWQAPIDVTDQVKERSFQIAGWGGSELYRRNT
SLNSQQDWQSNAKIRIVDGAANQIQVADGSRKYVVTLSIDESGGLVANLNGVSAPIILQSEH
AKVHSFHDYELQYSALNHTTTLFVDGQQITTWAGEVSQENNIQFGNADAQIDGRLHVQKIVL
TQQGHNLVEFDAFYLAQQTPEVEKDLEKLGWTKIKTGNTMSLYGNASVNPGPGHGITLTRQQ
NISGSQNGRLIYPAIVLDRFFLNVMSIYSDDGGSNWQTGSTLPIPFRWKSSSILETLEPSEA
DMVELQNGDLLLTARLDFNQIVNGVNYSPRQQFLSKDGGITWSLLEANNANVFSNISTGTVD
ASITRFEQSDGSHFLLFTNPQGNPAGTNGRQNLGLWFSFDEGVTWKGPIQLVNGASAYSDIY
QLDSENAIVIVETDNSNMRILRMPITLLKQKLTLSQN
11111111111111111111111100000000000000000000000000000000000000
00000000000000000000000000000000000000000000000000000000000000
00000000000000000000000000000000000000000000000000000000000000
00000000000000000000000000000000000000000000000000000000000000
00000000000000000000000000000000000000000000000000000000000000
00000000000000000000000000000000000000000000000000000000000000
00000000000000000000000000000000000000000000000000000000000000
00000000000000000000000000000000000000000000000000000000000000
00000000000000000000000000000000000000000000000000000000000000
00000000000000000000000000000000000000000000000000000000000000
00000000000000000000000000000000000000000000000000000000000000
00000000000000000000000000000000000000000000000000000000000000

0000000000000000000000000000000001111
>DM_train60
AGGHGDVGMHVKEKEKNKDENKRKDEERNKTQEEHLKEIMKHIVKIEVKGEEAVKKEAAEKL
LEKVPSDVLEMYKAIGGKIYIVDGDITKHISLEALSEDKKKIKDIYGKDALLHEHYVYAKEG
YEPVLVIQSSEDYVENTEKALNVYYEIGKILSRDILSKINQPYQKFLDVLNTIKNASDSDGQ
DLLFTNQLKEHPTDFSVEFLEQNSNEVQEVFAKAFAYYIEPQHRDVLQLYAPEAFNYMDKFN
EQEINLSLEELKDQRMLSRYEKWEKIKQHYQHWSDSLSEEGRGLLKKLQIPIEPKKDDIIHS
LSQEEKELLKRIQIDSSDFLSTEEKEFLKKLQIDIRDSLSEEEKELLNRIQVDSSNPLSEKE
KEFLKKLKLDIQPYDINQRLQDTGGLIDSPSINLDVRKQYKRDIQNIDALLHQSIGSTLYNK
IYLYENMNINNLTATLGADLVDSTDNTKINRGIFNEFKKNFKYSISSNYMIVDINERPALDN
ERLKWRIQLSPDTRAGYLENGKLILQRNIGLEIKDVQIIKQSEKEYIRIDAKVVPKSKIDTK
IQEAQLNINQEWNKALGLPKYTKLITFNVHNRYASNIVESAYLILNEWKNNIQSDLIKKVTN
YLVDGNGRFVFTDITLPNIAEQYTHQDEIYEQVHSKGLYVPESRSILLHGPSKGVELRNDSE
GFIHEFGHAVDDYAGYLLDKNQSDLVTNSKKFIDIFKEEGSNLTSYGRTNEAEFFAEAFRLM
HSTDHAERLKVQKNAPKTFQFINDQIKFIINS
11111111111111111111111111000000000000000000000000000000000000
00000000000000000000000000000000000000000000000000000000000000
00000000000000000000000000000000000000000000000000000000000000
00000000000000000000000000000000000000000000000000000000000000
00000000000000000000000000000000000000000000000000000000000000
00000000000000000000000000000000000111111111111111111111000000
00000000000000000000000000000000000000000000000000000000000000
00000000000000000000000000000000000000000000000000000000000000
00000000000000000000000000000000000000000000000000000000000000
00000000000000000000000000000000000000000000000000000000000000
00000000000000000000000000000000000000000000000000000000000000
00000000000000000000000000000000000000000000000000000000000000
00000000000000000000000000000111
>DM_train61
MNQNLLVTKRDGSTERINLDKIHRVLDWAAEGLHNVSISQVELRSHIQFYDGIKTSDIHETI
IKAAADLISRDAPDYQYLAARLAIFHLRKKAYGQFEPPALYDHVVKMVEMGKYDNHLLEDYT
EEEFKQMDTFIDHDRDMTFSYAAVKQLEGKYLVQNRVTGEIYESAQFLYILVAACLFSNYPR
ETRLQYVKRFYDAVSTFKISLPTPIMSGVRTPTRQFSSCVLIECGDSLDSINATSSAIVKYV
SQRAGIGINAGRIRALGSPIRGGEAFHTGCIPFYKHFQTAVKSCSQGGVRGGAATLFYPMWH
LEVESLLVLKNNRGVEGNRVRHMDYGVQINKLMYTRLLKGEDITLFSPSDVPGLYDAFFADQ
EEFERLYTKYEKDDSIRKQRVKAVELFSLMMQERASTGRIYIQNVDHCNTHSPFDPAIAPVR
QSNLCLEIALPTKPLNDVNDENGEIALCTLSAFNLGAINNLDELEELAILAVRALDALLDYQ
DYPIPAAKRGAMGRRTLGIGVINFAYYLAKHGKRYSDGSANNLTHKTFEAIQYYLLKASNEL
AKEQGACPWFNETTYAKGILPIDTYKKDLDTIANEPLHYDWEALRESIKTHGLRNSTLSALM
PSETSSQISNATNGIEPPRGYVSIKASKDGILRQVVPDYEHLHDAYELLWEMPGNDGYLQLV
GIMQKFIDQSISANTNYDPSRFPSGKVPMQQLLKDLLTAYKFGVKTLYYQNTRDGAEDAQDD
LVPSIQDDGCESGACKI
11110000000000000000000000000000000000000000000000000000000000
00000000000000000000000000000000000000000000000000000000000000

00000000000000000000000000000000000000000000000000000000000000
00000000000000000000000000000000000000000000000000000000000000
00000000000000000000000000000000000000000000000000000000000000
00000000000000000000000000000000000000000000000000000000000000
00000000000000000000000000000000000000000000000000000000000000
00000000000000000000000000000000000000000000000000000000000000
00000000000000000000000000000000000000000000000000000000000000
00000000000000000000000000000000000000000000000000000000000000
00000000000000000000000000000000000000000000000000000000000000
00000000000000000000000000000000000000000000000000000001111111
11111111111111111
>DM_train62
ANADNYKNVINRTGAPQYMKDYDYDDHQRFNPFFDLGAWHGHLLPDGPNTMGGFPGVALLTE
EYINFMASNFDRLTVWQDGKKVDFTLEAYSIPGALVQKLTAKDVQVEMTLRFATPRTSLLET
KITSNKPLDLVWDGELLEKLEAKEGKPLSDKTIAGEYPDYQRKISATRDGLKVTFGKVRATW
DLLTSGESEYQVHKSLPVQTEINGNRFTSKAHINGSTTLYTTYSHLLTAQEVSKEQMQIRDI
LARPAFYLTASQQRWEEYLKKGLTNPDATPEQTRVAVKAIETLNGNWRSPGGAVKFNTVTPS
VTGRWFSGNQTWPWDTWKQAFAMAHFNPDIAKENIRAVFSWQIQPGDSVRPQDVGFVPDLIA
WNLSPERGGDGGNWNERNTKPSLAAWSVMEVYNVTQDKTWVAEMYPKLVAYHDWWLRNRDHN
GNGVPEYGATRDKAHNTESGEMLFTVKKGDKEETQSGLNNYARVVEKGQYDSLEIPAQVAAS
WESGRDDAAVFGFIDKEQLDKYVANGGKRSDWTVKFAENRSQDGTLLGYSLLQESVDQASYM
YSDNHYLAEMATILGKPEEAKRYRQLAQQLADYINTCMFDPTTQFYYDVRIEDKPLANGCAG
KPIVERGKGPEGWSPLFNGAATQANADAVVKVMLDPKEFNTFVPLGTAALTNPAFGADIYWR
GRVWVDQFWFGLKGMERYGYRDDALKLADTFFRHAKGLTADGPIQENYNPLTGAQQGAPNFS
WSAAHLYMLYNDFFRKQ
11000000000000000000000000000000000000000000000000000000000000
00000000000000000000000000000000000000000000000000000000000000
00000000000000000000000000000000000000000000000000000000000000
00000000000000000000000000000000000000000000000000000000000000
00000000000000000000000000000000000000000000000000000000000000
00000000000000000000000000000000000000000000000000000000000000
00000000000000000000000000000000000000000000000000000000000000
00000000000000000000000000000000000000000000000000000000000000
00000000000000000000000000000000000000000000000000000000000000
00000000000000000000000000000000000000000000000000000000000000
00000000000000000000000000000000000000000000000000000000000000
00000000000000000000000000000000000000000000000000000000000000
00000000000000001
>DM_train63
MLNQELELSLNMAFARAREHRHEFMTVEHLLLALLSNPSAREALEACSVDLVALRQELEAFI
EQTTPVLPASEEERDTQPTLSFQRVLQRAVFHVQSSGRNEVTGANVLVAIFSEQESQAAYLL
RKHEVSRLDVVNFISHGTRKDEPTQSSDPGSQPNSEEQAGGEERLENFTTNLNQLARVGGID
PLIGREKELERAIQVLCRRRKNNPLLVGESGVGKTAIAEGLAWRIVQGDVPEVMADCTIYSL
DIGSLLAGTKYRGDFEKRFKALLKQLEQDTNSILFIDEIHTIIGAGAASGGQVDAANLIKPL

LSSGKIRVIGSTTYQEFSNIFEKDRALARRFQKIDITEPSIEETVQIINGLKPKYEAHHDVR
YTAKAVRAAVELAVKYINDRHLPDKAIDVIDEAGARARLMPVSKRKKTVNVADIESVVARIA
RIPEKSVSQSDRDTLKNLGDRLKMLVFGQDKAIEALTEAIKMARAGLGHEHKPVGSFLFAGP
TGVGKTEVTVQLSKALGIELLRFDMSEYMERHTVSRLIGAPPGYVGFDQGGLLTDAVIKHPH
AVLLLDEIEKAHPDVFNILLQVMDNGTLTDNNGRKADFRNVVLVMTTNAGVRETERKSIGLI
HQDNSTDAMEEIKKIFTPEFRNRLDNIIWFDHLSTDVIHQVVDKFIVELQVQLDQKGVSLEV
SQEARNWLAEKGYDRAMGARPMARVIQDNLKKPLANELLFGSLVDGGQVTVALDKEKNELTY
GFQSAQKHKAEAAH
00000000000000000000000000000000000000000000000000000000000000
00000000000000000000000000000000000000000000000000000000000000
00000000000000000011111111111111111111111110000000000000000000
00000000000000000000000000000000000000000000000000000000000000
00011100000000000000000000000000000000000000000000000000000000
00000000000000000000000000000000000000000000000000000000000000
00000000000000000000000000000000000000000000000000000000000000
00000000000000000000000000000000000000000000000000000000000000
00000000000000000000000000000000000000000000000000000000000000
00000000000000000000000000000000000000000000000000001111111111
11100000000000000000000000000000000000000000000000000000000000
00000000000000000000000000000000000000000000000000000000000000
00000000000111
>DM_train66
KDWTQYVNPLMGSQSTFELSTGNTYPAIARPWGMNFWTPQTGKMGDGWQYTYTANKIRGFKQ
THQPSPWINDYGQFSIMPIVGQPVFDEEKRASWFAHKGEVATPYYYKVYLAEHDIVTEMTPT
ERAVLFRFTFPENDHSYVVVDAFDKGSYIKIIPEENKIIGYTTRNSGGVPENFKNYFIIEFD
KPFTYKATVENGNLQENVAEQTTDHAGAIIGFKTRKGEQVNARIASSFISFEQAAANMNELG
KDNIEQLAQKGKDAWNQVLGKIEVEGGNLDQYRTFYSCLYRSLLFPRKFYELDANGQPIHYS
PYNGQVLPGYMFTDTGFWDTFRCLFPLLNLMYPSVNKEMQEGLINTYLESGFFPEWASPGHR
GCMVGNNSASILVDAYMKGVKVDDIKTLYEGLIHGTENVHPEVSSTGRLGYEYYNKLGYVPY
DVKINENAARTLEYAYDDWCIYRLAKELKRPKKEISLFAKRAMNYKNLFDKESKLMRGRNED
GTFQSPFSPLKWGDAFTEGNSWHYTWSVFHDPQGLIDLMGGKEMFVTMMDSVFAVPPIFDDS
YYGQVIHEIREMTVMNMGNYAHGNQPIQHMIYLYDYAGQPWKAQYWLRQVMDRMYTPGPDGY
CGDEDNGQTSAWYVFSALGFYPVCPGTDEYVMGTPLFKKATLHFENGNSLVIDAPNNSTENF
YIDSMSFNGADHTKNYLRHEDLFKGGTIKVDMSNRPNLNRGTKEEDMPYSFSKELEHHHHHH
00000000000000000000000000000000000000000000000000000000000000
00000000000000000000000000000000000000000000000000000000000000
00000000000000000000000000000000000000000000000000000000000000
00000000000000000000000000000000000000000000000000000000000000
00000000000000000000000000000000000000000000000000000000000000
00000000000000000000000000000000000000000000000000000000000000
00000000000000000000000000000000000000000000000000000000000000
00000000000000000000000000000000000000000000000000000000000000
00000000000000000000000000000000000000000000000000000000000000
00000000000000000000000000000000000000000000000000000000000000

00000000000000000000000000000000000000000000000000000000000000
00000000000000000000000000000000000000000000000000000011111111
>DM_train67
MTDRVSVGNLRIARVLYDFVNNEALPGTDIDPDSFWAGVDKVVADLTPQNQALLNARDELQA
QIDKWHRRRVIEPIDMDAYRQFLTEIGYLLPEPDDFTITTSGVDAEITTTAGPQLVVPVLNA
RFALNAANARWGSLYDALYGTDVIPETDGAEKGPTYNKVRGDKVIAYARKFLDDSVPLSSGS
FGDATGFTVQDGQLVVALPDKSTGLANPGQFAGYTGAAESPTSVLLINHGLHIEILIDPESQ
VGTTDRAGVKDVILESAITTIMDFEDSVAAVDAADKVLGYRNWLGLNKGDLAAAVDKDGTAF
LRVLNRDRNYTAPGGGQFTLPGRSLMFVRNVGHLMTNDAIVDTDGSEVFEGIMDALFTGLIA
IHGLKASDVNGPLINSRTGSIYIVKPKMHGPAEVAFTCELFSRVEDVLGLPQNTMKIGIMDE
ERRTTVNLKACIKAAADRVVFINTGFLDRTGDEIHTSMEAGPMVRKGTMKSQPWILAYEDHN
VDAGLAAGFSGRAQVGKGMWTMTELMADMVETKIAQPRAGASTAWVPSPTAATLHALHYHQV
DVAAVQQGLAGKRRATIEQLLTIPLAKELAWAPDEIREEVDNNCQSILGYVVRWVDQGVGCS
KVPDIHDVALMEDRATLRISSQLLANWLRHGVITSADVRASLERMAPLVDRQNAGDVAYRPM
APNFDDSIAFLAAQELILSGAQQPNGYTEPILHRRRREFKARAAEKPAPSDRAGDDAAR
00000000000000000000000000000000000000000000000000000000000000
00000000111111000000000000000000000000000000000000000000000000
00000000000000000000000000000000000000000000000000000000000000
00000000000000000000000000000000000000000000000000000000000000
00000000000000000000000000000000000000000000000000000111111111
10000000000000000000000000000000000000000000000000000000000000
00000000111110000000000000000000000000000000000000000000000000
00000000000000000000000000000000000000000000000000000000000000
00000000000000000000000000000000000000000000000000000000000000
00000000000000000000000000000000000000000000000000000000000000
00000000000000000000000000000000000000000000000000000111100000
00000000000000000000000000000000000000000000011111111111111
>DM_train68
MSEEISLSAEFIDRVKASVKPHWGKLGWVTYKRTYARWLPEKGRSENWDETVKRVVEGNINL
DPRLQDSPSLELKQSLTEEAERLYKLIYGLGATPSGRNLWISGTDYQRRTGDSLNNCWFVAI
RPQKYGDSKIVPSYLGKQEKAVSMPFSFLFDELMKGGGVGFSVARSNISQIPRVDFAIDLQL
VVDETSESYDASVKVGAVGKNELVQDADSIYYRLPDTREGWVLANALLIDLHFAQTNPDRKQ
KLILDLSDIRPYGAEIHGFGGTASGPMPLISMLLDVNEVLNNKAGGRLTAVDAADICNLIGK
AVVAGNVRRSAELALGSNDDQDFISMKQDQEKLMHHRWASNNSVAVDSAFSGYQPIAAGIRE
NGEPGIVNLDLSKNYGRIVDGYQAGIDGDVEGTNPCGEISLANGEPCNLFEVFPLIAEEQGW
DLQEVFALAARYAKRVTFSPYDWEISREIIQKNRRIGISMSGIQDWLLTRLGNRVVTGFKDD
FDPETHEAIKVPVYDKRAIKMVDQLYKAVVKADQDYSKTLGCNESIKHTTVKPSGTVAKLAG
ASEGMHFHYGAYLIQRIRFQDSDPLLPALKACGYRTEADIYTENTTCVEFPIKAVGADNPNF
ASAGTVSIAEQFATQAFLQTYWSDNAVSCTITFQDSEGDQVESLLRQYRFITKSTSLLPYFG
GSLQQAPKEPIDKETYEKRSQEITGNVEEVFSQLNSDVKDLELVDQTDCEGGACPIK
11100000000000000000000000000000000000000000000000000000000000
00000000000000000000000000000000000000000000000000000000000000
00000000000000000000000000000000000000000000000000000000000000
00000000000000000000000000000000000000000000000000000000000000

000000000000000000000000000000000000000000000000000000000000
000000111100000000000000000000000000000000000000000000000000
000000000000000000000000000000000000000000000000000000000000
000000000000000000000000000000000000000000000000000000000000
000000000000000000000000000000000000000000000000000000000000
000000000000000000000000000000000000000000000000000000000000
000000000000000000000000000000000000000000000000000000000000
0000000000000000000000000000000000000000111111111111111
>DM_train69
EVKQENRLLNESESSSQGLLGYYFSDLNFQAPMVVTSSTTGDLSIPSSELENIPSENQYFQS
AIWSGFIKVKKSDEYTFATSADNHVTMWVDDQEVINKASNSNKIRLEKGRLYQIKIQYQREN
PTEKGLDFKLYWTDSQNKKEVISSDNLQLPELKQKSSNSRKKRSTSAGPTVPDRDNDGIPDS
LEVEGYTVDVKNKRTFLSPWISNIHEKKGLTKYKSSPEKWSTASDPYSDFEKVTGRIDKNVS
PEARHPLVAAYPIVHVDMENIILSKNEDQSTQNTDSETRTISKNTSTSRTHTSEVHGNAEVH
ASFFDIGGSVSAGFSNSNSSTVAIDHSLSLAGERTWAETMGLNTADTARLNANIRYVNTGTA
PIYNVLPTTSLVLGKNQTLATIKAKENQLSQILAPNNYYPSKNLAPIALNAQDDFSSTPITM
NYNQFLELEKTKQLRLDTDQVYGNIATYNFENGRVRVDTGSNWSEVLPQIQETTARIIFNGK
DLNLVERRIAAVNPSDPLETTKPDMTLKEALKIAFGFNEPNGNLQYQGKDITEFDFNFDQQT
SQNIKNQLAELNATNIYTVLDKIKLNAKMNILIRDKRFHYDRNNIAVGADESVVKEAHREVI
NSSTEGLLLNIDKDIRKILSGYIVEIEDTEGLKEVINDRYDMLNISSLRQDGKTFIDFKKYN
DKLPLYISNPNYKVNVYAVTKENTIINPSENGDTSTNGIKKILIFSKKGYEIG
111111111111100000000000000000000000000000000000000000000000
000000000000000000000000000000000011110000000000000000000000
000000000000000000000000000000000001111111111111000000000000
000000000000000000000000000000000000000000000000000000000000
000000000000000000000000011111111111100000000000000001111111
111111111000000000000000000000111111110000000000000000000000
000000000000000000000000000000000000000000000000000000000000
000000000000000000000000000000000000000000000000000000000000
000000000000001111000000000000000000000000000000000000000000
000000000000000000000000000000000000000000000000000000000000
000000000000000000000000000000000000000000000000000000000000
000000000000000000000000000000000000000000000000000
>DM_train73
MAPSLDSISHSFANGVASAKQAVNGASTNLAVAGSHLPTTQVTQVDIVEKMLAAPTDSTLEL
DGYSLNLGDVVSAARKGRPVRVKDSDEIRSKIDKSVEFLRSQLSMSVYGVTTGFGGSADTRT
EDAISLQKALLEHQLCGVLPSSFDSFRLGRGLENSLPLEVVRGAMTIRVNSLTRGHSAVRLV
VLEALTNFLNHGITPIVPLRGTISASGDLSPLSYIAAAISGHPDSKVHVVHEGKEKILYARE
AMALFNLEPVVLGPKEGLGLVNGTAVSASMATLALHDAHMLSLLSQSLTAMTVEAMVGHAGS
FHPFLHDVTRPHPTQIEVAGNIRKLLEGSRFAVHHEEEVKVKDDEGILRQDRYPLRTSPQWL
GPLVSDLIHAHAVLTIEAGQSTTDNPLIDVENKTSHHGGNFQAAAVANTMEKTRLGLAQIGK
LNFTQLTEMLNAGMNRGLPSCLAAEDPSLSYHCKGLDIAAAAYTSELGHLANPVTTHVQPAE
MANQAVNSLALISARRTTESNDVLSLLLATHLYCVLQAIDLRAIEFEFKKQFGPAIVSLIDQ
HFGSAMTGSNLRDELVEKVNKTLAKRLEQTNSYDLVPRWHDAFSFAAGTVVEVLSSTSLSLA

AVNAWKVAAAESAISLTRQVRETFWSAASTSSPALSYLSPRTQILYAFVREELGVKARRGDV
FLGKQEVTIGSNVSKIYEAIKSGRINNVLLKMLA
111111111111111111111111100000001111100000000000000000000000
000000000000000000000000000000000000000011111111111111111110
000000000000000000000000000000000000000000000000000000000000
000000000000000000000000000000000000000000000000000000000000
000000000000000000000000000000000000000000000000000000000000
000000000000000000000000000000000000001111000000000000000000
000000000000000000000000000000000000000000000000000000000000
000000000000000000000000000000000000000000000000000000000000
000000000000000000000000000000000000000000000000000000000000
000000000000000000000000000000000000000000000000000000000000
000000000000000000000000000000000000000000000000000000000000
0000000000000000000000000000000000
>DM_train74
QNVSLQPPPQQLIVQNKTIDLPAVYQLNGGEEANPHAVKVLKELLSGKQSSKKGMLISIGEK
GDKSVRKYSRQIPDHKEGYYLSVNEKEIVLAGNDERGTYYALQTFAQLLKDGKLPEVEIKDY
PSVRYRGVVEGFYGTPWSHQARLSQLKFYGKNKMNTYIYGPKDDPYHSAPNWRLPYPDKEAA
QLQELVAVANENEVDFVWAIHPGQDIKWNKEDRDLLLAKFEKMYQLGVRSFAVFFDDISGEG
TNPQKQAELLNYIDEKFAQVKPDINQLVMCPTEYNKSWSNPNGNYLTTLGDKLNPSIQIMWT
GDRVISDITRDGISWINERIKRPAYIWWNFPVSDYVRDHLLLGPVYGNDTTIAKEMSGFVTN
PMEHAESSKIAIYSVASYAWNPAKYDTWQTWKDAIRTILPSAAEELECFAMHNSDLGPNGHG
YRREESMDIQPAAERFLKAFKEGKNYDKADFETLQYTFERMKESADILLMNTENKPLIVEIT
PWVHQFKLTAEMGEEVLKMVEGRNESYFLRKYNHVKALQQQMFYIDQTSNQNPYQPGVKTAT
RVIKPLIDRTFATVVKFFNQKFNAHLDATTDYMPHKMISNVEQIKNLPLQVKANRVLISPAN
EVVKWAAGNSVEIELDAIYPGENIQINFGKDAPCTWGRLEISTDAKEWKTVDLKQKESRLSA
GLQKAPVKFVRFTNVSDEEQQVYLRQFVLTIEKK
111100000000000000000000000000000000000011111110000000000
000000000000000000000000000000000000000000000000000000000000
000000000000000000000000000000000000000000000000000000000000
000000000000000000000000000000000000000000000000000000000000
000000000000000000000000000000000000000000000000000000000000
000000000000000000000000000000000000000000000000000000000000
000000000000000000000000000000000000000000000000000000000000
000000000000000000000000000000000000000000000000000000000000
000000000000000000000000000000000000000000000000000000000000
000000000000000000000000000000000001111111100000000000000011
111111111100000000000000000011111111111111111111111111110000
0000000000001111111111111100000001
>DM_train75
GVTVIPRLLGLKDEKKIATTVGEARLSGINYRHPDSALVSYPVAAAAPLGRLPAGNYRIAIV
GGGAGGIAALYELGRLAATLPAGSGIDVQIYEADPDSFLHDRPGIKAIKVRGLKAGRVSAAL
VHNGDPASGDTIYEVGAMRFPEIAGLTWHYASAAFGDAAPIKVFPNPGKVPTEFVFGNRVDR
YVGSDPKDWEDPDSPTLKVLGVVAGGLVGNPQGENVAMYPIANVDPAKIAAILNAATPPADA

LERIQTKYWPEFIAQYDGLTLGAAVREIVTVAFEKGTLPPVDGVLDVDESISYYVELFGRFG
FGTGGFKPLYNISLVEMMRLILWDYSNEYTLPVTENVEFIRNLFLKAQNVGAGKLVVQVRQE
RVANACHSGTASARAQLLSYDSHNAVHSEAYDFVILAVPHDQLTPIVSRSGFEHAASQNLGD
AGLGLETHTYNQVYPPLLLSDSSPAANARIVTAIGQLHMARSSKVFATVKTAALDQPWVPQW
RGEPIKAVVSDSGLAASYVVPSPIVEDGQAPEYSSLLASYTWEDDSTRLRHDFGLYPQNPAT
ETGTADGMYRTMVNRAYRYVKYAGASNAQPWWFYQLLAEARTADRFVFDWTTNKTAGGFKLD
MTGDHHQSNLCFRYHTHALAASLDNRFFIASDSYSHLGGWLEGAFMSALNAVAGLIVRANRG
DVSALSTEARPLVIGLRPVVKVPAAELATSQ
111111111111111000000000000000000000000000000000000000000000
000000000000000000000000000000000000000011110000000000000000
000000000000000000000000000000000000000000000000000000000000
000000000000000000000000000000000000000000000000000000000000
000000000000000000000000000000000000000000000000000000000000
000000000000000000000000000000000000000000000000000000000000
000000000000000000000000000000000000000000000000000000000000
000000000000000000000000000000000000000000000000000000000000
000000000000000000000000011110000000000000000000000000000000
000000000000000000000000000000000000000000000000000000000000
000000000000000000000000000000000000000000000000000000000000
0000000000000000000000000111111
>DM_train77
MKKLFVTCIVFFSILSPALLIAQQTGTAELIMKRVMLDLKKPLRNMDKVAEKNLNTLQPDGS
WKDVPYKDDAMTNWLPNNHLLQLETIIQAYIEKDSHYYGDDKVFDQISKAFKYWYDSDPKSR
NWWHNEIATPQALGEMLILMRYGKKPLDEALVHKLTERMKRGEPEKKTGANKTDIALHYFYR
ALLTSDEALLSFAVKELFYPVQFVHYEEGLQYDYSYLQHGPQLQISSYGAVFITGVLKLANY
VRDTPYALSTEKLAIFSKYYRDSYLKAIRGSYMDFNVEGRGVSRPDILNKKAEKKRLLVAKM
IDLKHTEEWADAIARTDSTVAAGYKIEPYHHQFWNGDYVQHLRPAYSFNVRMVSKRTRRSES
GNKENLLGRYLSDGATNIQLRGPEYYNIMPVWEWDKIPGITSRDYLTDRPLTKLWGEQGSND
FAGGVSDGVYGASAYALDYDSLQAKKAWFFFDKEIVCLGAGINSNAPENITTTLNQSWLNGP
VISTAGKTGRGKITTFKAQGQFWLLHDAIGYYFPEGANLSLSTQSQKGNWFHINNSHSKDEV
SGDVFKLWINHGARPENAQYAYIVLPGINKPEEIKKYNGTAPKVLANTNQLQAVYHQQLDMV
QAIFYTAGKLSVAGIEIETDKPCAVLIKHINGKQVIWAADPLQKEKTAVLSIRDLKTGKTNR
VKIDFPQQEFAGATVELK
111111111111111111111111100000000000000000000000000000000000
000000000000000000000000000000000000000000000000000000000000
000000000000000000000000000000000000000000000000000000000000
000000000000000000000000000000000000000000000000000000000000
000000000000000000000000000000000000000000000000000000000000
000000000000000000000000000000000000000000000000000000000000
000000000000000000000000000000000000000000000000000000000000
000000000000000000000000000000000000000000000000000000000000
000000000000000000000000000000000000000000000000000000000000
000000000000000000000000000000000000000000000000000000000000
000000000000000000000000000000000000000000000000000000000000

00000000000000000
>DM_train79
LMPGGKEFYNFYPEYVVGLQWMGDNYVFIEGDDLVFNKANGKSAQTTRFSAADLNALMPEGC
KFQTTDAFPSFRTLDAGRGLVVLFTQGGLVGFDMLARKVTYLFDTNEETASLDFSPVGDRVA
YVRNHNLYIARGGKLGEGMSRAIAVTIDGTETLVYGQAVHQREFGIEKGTFWSPKGSCLAFY
RMDQSMVKPTPIVDYHPLEAESKPLYYPMAGTPSHHVTVGIYHLATGKTVYLQTGEPKEKFL
TNLSWSPDENILYVAEVNRAQNECKVNAYDAETGRFVRTLFVETDKHYVEPLHPLTFLPGSN
NQFIWQSRRDGWNHLYLYDTTGRLIRQVTKGEWEVTNFAGFDPKGTRLYFESTEASPLERHF
YCIDIKGGKTKDLTPESGMHRTQLSPDGSAIIDIFQSPTVPRKVTVTNIGKGSHTLLEAKNP
DTGYAMPEIRTGTIMAADGQTPLYYKLTMPLHFDPAKKYPVIVYVYGGPHAQLVTKTWRSSV
GGWDIYMAQKGYAVFTVDSRGSANRGAAFEQVIHRRLGQTEMADQMCGVDFLKSQSWVDADR
IGVHGWSYGGFMTTNLMLTHGDVFKVGVAGGPVIDWNRYAIMYGERYFDAPQENPEGYDAAN
LLKRAGDLKGRLMLIHGAIDPVVVWQHSLLFLDACVKARTYPDYYVYPSHEHNVMGPDRVHL
YETITRYFTDHL
11111111111111100000000000000000000000111111100000000000001111
11111110000000000000000000000000000000000000000000000000000000
00000000000000000000000000000000000000000000000000000000000000
00000000000000000000000000000000000000000000000000000000000000
00000000000000000000000000000000000000000000000000000000000000
00000000000000000000000000000000000000000000000000000000000000
00000000000000000000000000000000000000000000000000000000000011
11110000000000000000000000000000000000000000000000000000001111
00000000000000000000000000000000000000000000000000000000000000
00000000000000000000000000000000000000000000000000000000000000
00000000000000000000000000000000000000000000000000000000000000
000000000000
>DM_train83
MAEELVLERCDLELETNGRDHHTADLCREKLVVRRGQPFWLTLHFEGRNYQASVDSLTFSVV
TGPAPSQEAGTKARFPLRDAVEEGDWTATVVDQQDCTLSLQLTTPANAPIGLYRLSLEASTG
YQGSSFVLGHFILLFNAWCPADAVYLDSEEERQEYVLTQQGFIYQGSAKFIKNIPWNFGQFQ
DGILDICLILLDVNPKFLKNAGRDCSRRSSPVYVGRVGSGMVNCNDDQGVLLGRWDNNYGDG
VSPMSWIGSVDILRRWKNHGCQRVKYGQCWVFAAVACTVLRCLGIPTRVVTNYNSAHDQNSN
LLIEYFRNEFGEIQGDKSEMIWNFHCWVESWMTRPDLQPGYEGWQALDPTPQEKSEGTYCCG
PVPVRAIKEGDLSTKYDAPFVFAEVNADVVDWIQQDDGSVHKSINRSLIVGLKISTKSVGRD
EREDITHTYKYPEGSSEEREAFTRANHLNKLAEKEETGMAMRIRVGQSMNMGSDFDVFAHIT
NNTAEEYVCRLLLCARTVSYNGILGPECGTKYLLNLTLEPFSEKSVPLCILYEKYRDCLTES
NLIKVRALLVEPVINSYLLAERDLYLENPEIKIRILGEPKQKRKLVAEVSLQNPLPVALEGC
TFTVEGAGLTEEQKTVEIPDPVEAGEEVKVRMDLVPLHMGLHKLVVNFESDKLKAVKGFRNV
IIGPA
11111111111111000000000000000000000000000001111111111110000000
00000000000000000000000000000000000000000000000000000000000011
11111111000000000000000000000000000000000000000000000000000000
00000000000000000000000000000000000000000000000000000000000000
00000000000000000000000000000000000000000000000000000000000000

00000000000000000000000000000000000000000000000000000000000000
00000000000000000000000000000000000000000000000000000000000000
00000000000000000000000000000000000000000000000000000000000000
00000000000000000000000000000000000000000000000000000000000000
00000000000000000000000000000000000000000000000000000000000000
00000000000000000000000000000000000000000000000000000000000000
00000
>DM_train84
VLSGCSNNVSSIKIDRFNNISAVNGPGEEDTWASAQKQGVGTANNYVSKVWFTLANGAISEV
YYPTIDTADVKEIKFIVTDGKSFVPDETKDAISKVEKFTDKSLGYKLVNTDKKGRYRITKDI
FTDVKRNSLIMKAKFEALEGSIHDYKLYLAYDPHIKNQGSYNEGYVIKANNNEMLMAKRDNV
YTALSSNIGWKGYSIGYYKVNDIMTDLDENKQMTKHYDSARGNIIEGAEIDLTKNSEFEIVL
SFGQSDSEAAKTALETLGEDYNNLKNNYIDEWTKYCNTLNNFNGKANSLYYNSMMILKASED
KTNKGAYIASLSIPWGDGQRDDNTGGYHLVWSRDLYHVANAFIAAGDVDSANRSLDYLAKVV
KDNGMIPQNTWISGKPYWTGIQLDEQADPIILSYRLKRYDLYDSLVKPLADFIIKIGPKTGQ
ERWEEIGGYSPATMAAEVAGLTCAAYIAEQNKDYESAQKYQEKADNWQKLIDNLTYTENGPL
GNGQYYIRIAGLSDPDADFMINIANGGGVYDQKEIVDPSFLELVRLGVKSADDPKILNTLKV
VDSTIKVDTPKGPSWYRYNHDGYGEPSKTELYHGAGKGRLWPLLTGERGMYEIAAGKDATPY
VKAMEKFANEGGIISEQVWEDTGLPTDSASPLNWAHAEYVILFASNIEHKVLDMPDIVYKRY
VA
11111111110000000000000000000000000000000000000000000000000000
00000000000000000000000000000000000000000000000000000000000000
00000000000000000000000000000000000000000000000000000000000000
00000000000000000000000000000000000000000000000000000000000000
00000000000000000000000000000000000000000000000000000000000000
00000000000000000000000000000000000000000000000000000000000000
00000000000000000000000000000000000000000000000000000000000000
00000000000000000000000000000000000000000000000000000000000000
00000000000000000000000000000000000000000000000000000000000000
00000000000000000000000000000000000000000000000000000000000000
00000000000000000000000000000000000000000000000000000000000000
00
>DM_train85
TDAKNNLLYFFDRPNEPCFMQKGEDKVVFEIPDHYYPDKYKSLSNTLSNRFGNEATKRIPIR
NITLPNLEVPMQLPYNDQFSLFVPKHRTMAAKLIDIFMGMRDVEDLQSVCSYCQLRINPYMF
NYCLSVAILHRPDTKGLSIPTFAETFPDKFMDSKVFLRAREVSNVVISGSRMPVNVPINYTA
NTTEPEQRVAYFREDIGINLHHWHWHLVYPFDSADRSIVNKDRRGELFYYMHQQIIGRYNVE
RMCNGLPQVKPFSDFSAPIEEGYFPKLDSQVASRTWPPRFAGSVFRNLDRTVDQVKIDVRKL
FTWRDQFLEAIQKMAIKMPNGRELPLDEVTGIDMLGNLMESSIISPNRGYYGDLHNMGHVFA
AYTHDPDHRHLEQFGVMGDSATAMRDPFFYRWHRFVDDVFNIYKEKLTPYTNERLDFPGVRV
SSVGIEGARPNTLRTLWQQSTVELGRGLDFTPRGSVLARFTHLQHDEFQYVIEVNNTTGGNL
MGTVRIFMAPKVDDNGQPMSFNKQRRLMIELDKFSQALRPGTNTIRRRSVDSSVTIPYERTF
RNQSERPGDPGTAGAAEFDFCGCGWPHHMLIPKGTAQGYPVVLFVMISNWNNDRIEQDLVGS
CNDAASYCGIRDRKYPDKQAMGYPFDRKMANDAATLSDFLRPNMAVRDCSIQFSDTTVERGQ

QG
100000000000000000000000000000000000000000000000000000000000
000000000000000000000000000000000000000000000000000000000000
000000000000000000000000000000000000000000000000000000000000
000000000000000000000000000000000000000000000000000000000000
000000000000000000000000000000000000000000000000000000000000
000000000000000000000000000000000000000000000000000000000000
000000000000000000000000000000000000000000000000000000000000
000000010000000000000000000000000000000000000000000000000000
000000000000000000000000000000000000000000000000000000000001
111111111111111111000000000000000000000000000000000000001100
000000000000000000000000000000000000000000000000000000000111
11
>DM_train86
MLRFVTKNSQDKSSDLFSICSDRGTFVAHNRVRTDFKFDNLVFNRVYGVSQKFTLVGNPTVC
FNEGSSYLEGIAKKYLTLDGGLAIDNVLNELRSTCGIPGNAVASHAYNITSWRWYDNHVALL
MNMLRAYHLQVLTEQGQYSAGDIPMYHDGHVKIKLPVTIDDTAGPTQFAWPSDRSTDSYPDW
AQFSESFPSIDVPYLDVRPLTVTEVNFVLMMMSKWHRRTNLAIDYEAPQLADKFAYRHALTV
QDADEWIEGDRTDDQFRPPSSKVMLSALRKYVNHNRLYNQFYTAAQLLAQIMMKPVPNCAEG
YAWLMHDALVNIPKFGSIRGRYPFLLSGDAALIQATALEDWSAIMAKPELVFTYAMQVSVAL
NTGLYLRRVKKTGFGTTIDDSYEDGAFLQPETFVQAALACCTGQDAPLNGMSDVYVTYPDLL
EFDAVTQVPITVIEPAGYNIVDDHLVVVGVPVACSPYMIFPVAAFDTANPYCGNFVIKAANK
YLRKGAVYDKLEAWKLAWALRVAGYDTHFKVYGDTHGLTKFYADNGDTWTHIPEFVTDGDVM
EVFVTAIERRARHFVELPRLNSPAFFRSVEVSTTIYDTHVQAGAHAVYHASRINLDYVKPVS
TGIQVINAGELKNYWGSVRRTQQGLGVVGLTMPAVMPTGEPTAGAAHEELIEQADNVLVE
000000000000000000000000000000000000000000000000000000000000
000000000000000000000000000000000000000000000000000000000000
000000000000000000000000000000000000000000000000000000000000
000000000000000000000000000000000000000000000000000000000000
000000000000000000000000000000000000000000000000000000000000
000000000000000000000000000000000000000000000000000000000000
000000000000000000000000000000000000000000000000000000000000
000000000000000000000000000000000000000000000000000000000000
000000000000000000000000000000000000000000000000000000000000
000000000000000000000000000000000000000000000000000000000000
0000000000000000000000000000001111111111111111111111111111
>DM_train87
MTAGYEPCWLRYERKDQYSRLRFEEIVAKRTSPIFQAAVEELQKGLRSMMEIEPQVVQEVNE
TANSIWLGTLEDEEFERPLEGTLVHPEGYVIRSDVDDGPFRIYIIGKTDAGVLYGVFHFLRL
LQMGENIAQLSIIEQPKNRLRMINHWDNMDGSIERGYAGRSIFFVDDQFVKQNQRIKDYARL
LASVGINAISINNVNVHKTETKLITDHFLPDVAEVADIFRTYGIKTFLSINYASPIEIGGLP
TADPLDPEVRWWWKETAKRIYQYIPDFGGFVVKADSEFRPGPFTYGRDHAEGANMLAEALAP
FGGLVIWRCFVYNCQQDWRDRTTDRAKAAYDHFKPLDGQFRENVILQIKNGPMDFQVREPVS
PLFGAMPKTNQMMEVQITQEYTGQQKHLCFLIPQWKEVLDFDTYAKGKGSEVKKVIDGSLFD

YRYSGIAGVSNIGSDPNWTGHTLAQANLYGFGRLAWNPDLSAEEIANEWVVQTFGDDSQVVE
TISWMLLSSWRIYENYTSPLGVGWMVNPGHHYGPNVDGYEYSHWGTYHYADRDGIGVDRTVA
TGTGYTAQYFPENAAMYESLDTCPDELLLFFHHVPYTHRLHSGETVIQHIYNTHFEGVEQAK
QLRKRWEQLKGKIDEKRYHDVLERLTIQVEHAKEWRDVINTYFYRKSGIDDQYGRKIYR
111100000000000000000000000000000000000000000000000000000000
000000000000000000000000000000000000000000000000000000000000
000000000000000000000000000000000000000000000000000000000000
000000000000000000000000000000000000000000000000000000000000
000000000000000000000000000000000111110000000000000000000000
000000000000000000000000000000000000000000000000000000000000
000000000000000000000000000000000000000000000000000000000000
000000000000000000000000000000000000000000000000000000000000
000000000000000000000000000000000000000000000000000000000000
000000000000000000000000000000000000000000000000000000000000
000000000000000000000000000000000000000000000000000000000
>DM_train88
SAPGEDEECGRVRDFVAKLANNTHQHVFDDLRGSVSLSWVGDSTGVILVLTTFHVPLVIMTF
GQSKLYRSEDYGKNFKDITDLINNTFIRTEFGMAIGPENSGKVVLTAEVSGGSRGGRIFRSS
DFAKNFVQTDLPFHPLTQMMYSPQNSDYLLALSTENGLWVSKNFGGKWEEIHKAVCLAKWGS
DNTIFFTTYANGSCKADLGALELWRTSDLGKSFKTIGVKIYSFGLGGRFLFASVMADKDTTR
RIHVSTDQGDTWSMAQLPSVGQEQFYSILAANDDMVFMHVDEPGDTGFGTIFTSDDRGIVYS
KSLDRHLYTTTGGETDFTNVTSLRGVYITSVLSEDNSIQTMITFDQGGRWTHLRKPENSECD
ATAKNKNECSLHIHASYSISQKLNVPMAPLSEPNAVGIVIAHGSVGDAISVMVPDVYISDDG
GYSWTKMLEGPHYYTILDSGGIIVAIEHSSRPINVIKFSTDEGQCWQTYTFTRDPIYFTGLA
SEPGARSMNISIWGFTESFLTSQWVSYTIDFKDILERNCEEKDYTIWLAHSTDPEDYEDGCI
LGYKEQFLRLRKSSMCQNGRDYVVTKQPSICLCSLEDFLCDFGYYRPENDSKCVEQPELKGH
DLEFCLYGREEHLTTNGYRKIPGDKCQGGVNPVREVKDLKKKCTSNFLSPEKQNSKSNS
111111111100000000000000000000000000000000000000000000011111
000000000000000000000000000000000000000000000000000000000000
000000000000000000000000000000000000000000000000000000000000
000000000000000000000000000000000000000000000000000000000000
000000000000000000000000000000000000000000000000000000000000
000000000000000000000000000000000000000000000000000000000000
000000000000000000000000000000000000000000000000000000000000
000000000000000000000000000000000000000000000000000000000000
000000000000000000000000000000000000000000000000000000000000
000000000000000000000000000000000000000000000000000000000000
000000000000000000000000000000000000000000000000001111111
>DM_train89
MVKSRKISILLAVAMLVSIMIPTTAFAGPTKAPTKDGTSYKDLFLELYGKIKDPKNGYFSPD
EGIPYHSIETLIVEAPDYGHVTTSEAFSYYVWLEAMYGNLTGNWSGVETAWKVMEDWIIPDS
TEQPGMSSYNPNSPATYADEYEDPSYYPSELKFDTVRVGSDPVHNDLVSAYGPNMYLMHWLM
DVDNWYGFGTGTRATFINTFQRGEQESTWETIPHPSIEEFKYGGPNGFLDLFTKDRSYAKQW
RYTNAPDAEGRAIQAVYWANKWAKEQGKGSAVASVVSKAAKMGDFLRNDMFDKYFMKIGAQD

KTPATGYDSAHYLMAWYTAWGGGIGASWAWKIGCSHAHFGYQNPFQGWVSATQSDFAPKSSN
GKRDWTTSYKRQLEFYQWLQSAEGGIAGGATNSWNGRYEKYPAGTSTFYGMAYVPHPVYADP
GSNQWFGFQAWSMQRVMEYYLETGDSSVKNLIKKWVDWVMSEIKLYDDGTFAIPSDLEWSGQ
PDTWTGTYTGNPNLHVRVTSYGTDLGVAGSLANALATYAAATERWEGKLDTKARDMAAELVN
RAWYNFYCSEGKGVVTEEARADYKRFFEQEVYVPAGWSGTMPNGDKIQPGIKFIDIRTKYRQ
DPYYDIVYQAYLRGEAPVLNYHRFWHEVDLAVAMGVLATYFPDMTYKVPGTPSTKLYG
11111111111111111111111111100000000000000000000000000000000000
00000000000000000000000000000000000000000000000000000000000000
00000000000000000000000000000000000000000000000000000000000000
00000000000000000000000000000000000000000000000000000000000000
00000000000000000000000000000000000000000000000000000000000000
00000000000000000000000000000000000000000000000000000000000000
00000000000000000000000000000000000000000000000000000000000000
00000000000000000000000000000000000000000000000000000000000000
00000000000000000000000000000000000000000000000000000000000000
00000000000000000000000000000000000000000000000000000000000000
0000000000000000000000000000000000000000000000000111111111
>DM_train90
SPCSVVNDLRWDLSAQQIEERTRELIEQTKRVYDQVGTQEFEDVSYESTLKALADVEVTYTV
QRNILDFPQHVSPSKDIRTASTEADKKLSEFDVEMSMREDVYQRIVWLQEKVQKDSLRPEAA
RYLERLIKLGRRNGLHLPRETQENIKRIKKKLSLLCIDFNKNLNEDTTFLPFTLQELGGLPE
DFLNSLEKMEDGKLKVTLKYPHYFPLLKKCHVPETRRKVEEAFNSRCKEENSAILKELVTLR
AQKSRLLGFHTHADYVLEMNMAKTSQTVATFLDELAQKLKPLGEQERAVILELKRAECERRG
LPFDGRIRAWDMRYYMNQVEETRYCVDQNLLKEYFPVQVVTHGLLGIYQELLGLAFHHEEGA
SAWHEDVRLYTARDAASGEVVGKFYLDLYPREGKYGHAACFGLQPGCLRQDGSRQIAIAAMV
ANFTKPTADAPSLLQHDEVRTYFHEFGHVMHQLCSQAEFAMFSGTHVETDFVEAPSQMLENW
VWEQEPLLRMSRHYRTGSAVPRELLEKLIESRQANTGLFNLRQIVLAKVDQALHTQTDADPA
EEYARLCQEILGVPATPGTNMPATFGHLAGGYDAQYYGYLWSEVYSMDMFHTRFKQEGVLNS
KVGMDYRSCILRPGGSEDASAMLRRFLGRDPKQDAFLLSKGLQVGGCEPEPQVC
11111111000000000000000000000000000000000000000000000000000000
00000000000000000000000000000000000000000000000000000000000000
00000000000000000000000000000000000000000000000000000000000000
00000000000000000000000000000000000000000000000000000000000000
00000000000000000000000000000000000000000000000000000000000000
00000000000000000000000000000000000000000000000000000000000000
00000000000000000000000000000000000000000000000000000000000000
00000000000000000000000000000000000000000000000000000000000000
00000000000000000000000000000000000000000000000000000000000000
00000000000000000000000000000000000000000000000000000000000000
000000000000000000000000000000000000000000111111111111
>DM_train91
AGVDEAAIRATEQAGGEWLSHGRTYAEQRFSPLKQIDASNVRSLGLAWYMDLDNTRGLEATP
LFHDGVIYTSMSWSRVIAVDAASGKELWRYDPEVAKVKARTSCCDAVNRGVALWGDKVYVGT
LDGRLIALDAKTGKAIWSQQTTDPAKPYSITGAPRVVKGKVIIGNGGAEYGVRGFVSAYDAD

TGKLAWRFYTVPGDPALPYEHPELREAAKTWQGDQYWKLGGGGTVWDSMAYDPELDLLYVGT
GNGSPWNREVRSPGGGDNLYLSSILAIRPDTGKLAWHYQVTPGDSWDFTATQQITLAELNID
GKPRKVLMQAPKNGFFYVLDRTNGKLISAEKFGKVTWAEKVDLATGRPVEAPGVRYEKEPIV
MWPSPFGAHNWHSMSFNPGTGLVYIPYQEVPGVYRNEGKDFVTRKAFNTAAGFADATDVPAA
VVSGALLAWDPVKQKAAWKVPYPTHWNGGTLSTAGNLVFQGTAAGQMHAYSADKGEALWQFE
AQSGIVAAPMTFELAGRQYVAIMAGWGGVATLTGGESMNLPGMKNRSRLLVFALDGKAQLPP
PAPAPAKVERVPQPVTAAPEQVQAGKQLYGQFCSVCHGMGTISGGLIPDLRQSSDATREHFQ
QIVLQGALKPLGMPSFDDSLKPEEVEQIKLYVMSREYEDYMARHKAAP
00000000000000000000000000000000000000000000000000000000000000
00000000000000000000000000000000000000000000000000000000000000
00000000000000000000000000000000000000000000000000000000000000
00000000000000000000000000000000000000000000000000000000000000
00000000000000000000000000000000000000000000000000000000000000
00000000000000000000000000000000000000000000000000000000000000
00000000000000000000000000000000000000000000000000000000000000
00000000000000000000000000000000000000000000000000000000000000
00000000000000000000000000000000000000000000000000000000000000
00000000000000000000000000000000000000000000000000000000000000
000000000000000000000000000000000000000000001111
>DM_train96
MSLSIPREFSNAIRFLSIDATLKAKSGHPGMPMGMADIATVLWTKFLKHNPNNPHWINRDRF
VLSNGHGSMLLYSLLHLTGYDLSIEDIKNFRQLHSKTPGHPEYGYTPGVETTTGPLGQGVAN
AVGMALGEKLLSDRYNTPDLKVIDHHTYVFLGDGCLMEGVSHEACSLAGTLGLNKLVAFWDD
NNISIDGDTKGWFSDNTPERFRAYGWHVIENVDGHDFVAIEKAINEAHSQQQKPTLICCKTV
IGFGSPEKAGTASVHGSPLSDQERASAAKELNWDYQAFEIPQDVYKYWDAREKGQALEANWQ
GQRNLFKDSPKFDEFERVLSKELPVGLESAINDYIASQLSNPVKVATRKASQMVLEVLCKNM
PEMFGGSADLTGSNNTNWSGSVWLNNTQEGANYLSYGVREFGMAAIMNGLSLYGGIKPYGGT
FLVFSDYSRNAIRMSALMKQPVVHVMSHDSIGLGEDGPTHQPIEHVPSLRLIPNLSVWRPAD
TIETMIAWKEAVKSKDTPSVMVLTRQNLMPVVQTQHQVANIARGGYLVKDNPDAKLTIVATG
SEVELAVKVANEFEKKGIKLNVASIPCVEVFATQAHEYKKTVIKDDIPAVFVEMAQPDMWYK
YMPKAGGEVKGIYSFGESAPAEDLFKRFGFTVENISNIVAKYV
11100000000000000000000000000000000000000000000000000000000000
00000000000000000000000000000000000000000000000000000000000000
00000000000000000000000000000000000000000000000000000000000000
00111111000000000000000000000000000000000000000000000000000000
00000000001111100000000000000000000000000000000000000000000000
00000000000000000000000000000000000000000000000000000000000000
00000000000100000000000000000000000000000000000000000000000000
00000000000000000000000000000000000000000000000000000000000000
00000000000000000000000000000000000000000000000000000000000000
00000000000000000000000000000000000000000000000000000000000000
0000000000000000000000000000000000000000000
>DM_train97
MERINFIFGIHNHQPLGNFGWVFEEAYNRSYRPFMEILEEFPEMKVNVHFSGPLLEWIEENK

PDYLDLLRSLIKRGQLEIVVAGFYEPVLAAIPKEDRLVQIEMLKDYARKLGYDAKGVWLTER
VWQPELVKSLREAGIEYVVVDDYHFMSAGLSKEELFWPYYTEDGGEVITVFPIDEKLRYLIP
FRPVKKTIEYLESLTSDDPSKVAVFHDDGEKFGVWPGTYEWVYEKGWLREFFDAITSNEKIN
LMTYSEYLSKFTPRGLVYLPIASYFEMSEWSLPAKQAKLFVEFVEQLKEEGKFEKYRVFVRG
GIWKNFFFKYPESNFMHKRMLMVSKAVRDNPEARKYILKAQCNDAYWHGVFGGIYLPHLRRT
VWENIIKAQRYLKPENKILDVDFDGRAEIMVENDGFIATIKPHYGGSIFELSSKRKAVNYND
VLPRRWEHYHEVPEATKPEKESEEGIASIHELGKQIPEEIRRELAYDWQLRAILQDHFIKPE
ETLDNYRLVKYHELGDFVNQPYEYEMIENGVKLWREGGVYAEEKIPARVEKKIELTEDGFIA
KYRVLLEKPYKALFGVEINLAVHSVMEKPEEFEAKEFEVNDPYGIGKVRIELDKAAKVWKFP
IKTLSQSEAGWDFIQQGVSYTMLFPIEKELEFTVRFREL
00000000000000000000000000000000000000000000000000000000000000
00000000000000000000000000000000000000000000000000000000000000
00000000000000000000000000000000000000000000000000000000000000
00000000000000000000000000000000000000000000000000000000000000
00000000000000000000000000000000000000000000000000000000000000
00000000000000000000000000000000000000000000000000000000000000
00000000000000000000000000000000000000000000000000000000000000
00000000000111111111111111111111110000000000000000000000000000
00000000000000000000000000000000000000000000000000000000000000
00000000000000000000000000000000000000000000000000000000000000
000000000000000000000000000000000000000
>DM_train98
MSRLLALLALAPLLAGAAETTAPKPPSAFTVEAQRRVEAELPFADRADFERADRGLIRRPER
LLIRNPDGSVAWQLGGYDFLLDGKPRDSINPSLQRQALLNLKYGLFEVAEGIYQVRGFDLAN
ITFIRGDSGWIVVDTLTTPATARAAYELVSRELGERPIRTVIYSHAHADHFGGVRGLVEPQQ
VASGAVQIIAPAGFMEAAIKENVLAGNAMMRRATYQYGTQLPKGPQGQVDMAIGKGLARGPL
SLLAPTRLIEGEGEDLVLDGVPFTFQNTPGTESPAEMNIWLPRQKALLMAENVVGTLHNLYT
LRGAEVRDALGWSKYINQALHRFGRQAEVMFAVHNWPRWGNAEIVEVLEKQRDLYGYLHDQT
LHLANQGVTIGQVHNRLRLPPSLDQEWYDRGYHGSVSHNARAVLNRYLGYYDGNPATLDPLS
PEDSAGRYVEYMGGAERLLEQARASYARGEYRWVVEVVNRLVFAEPDNRAARELQADALEQL
GYQAENAGWRNSYLSAAYELRHGVPRDQPTMKAGSADALAAMDTGLLFDYLGVRLDAGAAEG
KALSINLRLPDIGENYLLELKNSHLNNLRGVQSEDAGQTVSIDRADLNRLLLKEVSAVRLVF
EGKLKSSGNPLLLGQLFGMLGDFDFWFDIVTPAAKSEG
11111111111111111110000000000000000000000000000000000000000000
00000000000000000000000000000000000000000000000000000000000000
00000000000000000000000000000000000000000000000000000000000000
00000000000000000001110000000000000000000000000000000000000000
00000000000000000000000000000000000000000000000000000000000000
00000000000000000000000000000000000000000000000000000000000000
00000000000000000000000000000000000000000000000000000000000000
00000000000000000000000000000000000000000000000000000000000000
00000000000000000000000000001111100000000000000000000000000000
00000000000000000000000000000000000000000000000000000000000000
00000000000000000000000000000000000111

>DM_train100
MNFLRGVMGGQSAGPQHTEAETIQKLCDRVASSTLLDDRRNAVRALKSLSKKYRLEVGIQAM
EHLIHVLQTDRSDSEIIGYALDTLYNIISNDEEEEVEENSTRQSEDLGSQFTEIFIKQQENV
TLLLSLLEEFDFHVRWPGVKLLTSLLKQLGPQVQQIILVSPMGVSRLMDLLADSREVIRNDG
VLLLQALTRSNGAIQKIVAFENAFERLLDIITEEGNSDGGIVVEDCLILLQNLLKNNNSNQN
FFKEGSYIQRMKPWFEVGDENSGWSAQKVTNLHLMLQLVRVLVSPNNPPGATSSCQKAMFQC
GLLQQLCTILMATGVPADILTETINTVSEVIRGCQVNQDYFASVNAPSNPPRPAIVVLLMSM
VNERQPFVLRCAVLYCFQCFLYKNQKGQGEIVSTLLPSTIDATGNTVSAGQLLCGGLFSTDS
LSNWCAAVALAHALQENATQKEQLLRVQLATSIGNPPVSLLQQCTNILSQGSKIQTRVGLLM
LLCTWLSNCPIAVTHFLHNSANVPFLTGQIAENLGEEEQLVQGLCALLLGISIYFNDNSLET
YMKEKLKQLIEKRIGKENFIEKLGFISKHELYSRASQKPQPNFPSPEYMIFDHEFTKLVKEL
EGVITKAIYKSSEEDKKEEEVKKTLEQHDSI
111111111111111100000000000000000000000000000000000000000000
000000000000000000000000000000111111111111100000000000000000
000000000000000000000000000000000000000000000000000000000000
000000000000000000000000000000000000000000000000000000000000
000000000000000000000000000000000000000000000000000000000000
000000000000000000000000000000000000000000000000000000000000
000000000000000000000000000000000000000000000000000000000000
000000000000000000000000000000000000000000000000000000000000
000000000000000000000000000000000000000000000000000000000000
000000000000000000000000000000000000000000000000000000000000
0000000000001111111111111111111
>DM_train101
MGGSHHHHHHGMASLAPGSSRVELFKRQSSKVPFEKDGKVTERVVHSFRLPALVNVDGVMVA
IADARYETSFDNSLIDTVAKYSVDDGETWETQIAIKNSRASSVSRVVDPTVIVKGNKLYVLV
GSYNSSRSYWTSHGDARDWDILLAVGEVTKSTAGGKITASIKWGSPVSLKEFFPAEMEGMHT
NQFLGGAGVAIVASNGNLVYPVQVTNKKKQVFSKIFYSEDEGKTWKFGKGRSAFGCSEPVAL
EWEGKLIINTRVDYRRRLVYESSDMGNTWLEAVGTLSRVWGPSPKSNQPGSQSSFTAVTIEG
MRVMLFTHPLNFKGRWLRDRLNLWLTDNQRIYNVGQVSIGDENSAYSSVLYKDDKLYCLHEI
NSNEVYSLVFARLVGELRIIKSVLQSWKNWDSHLSSICTPADPAASSSERGCGPAVTTVGLV
GFLSHSATKTEWEDAYRCVNASTANAERVPNGLKFAGVGGGALWPVSQQGQNQRYHFANHAF
TLVASVTIHEVPKGASPLLGASLDSSGGKKLLGLSYDKRHQWQPIYGSTPVTPTGSWEMGKR
YHVVLTMANKIGSVYIDGEPLEGSGQTVVPDERTPDISHFYVGGYKRSGMPTDSRVTVNNVL
LYNRQLNAEEIRTLFLSQDLIGTEAHMD
111111111111111100000000000000000000000000000000000000000000
000000000000000000000000000000000000000000000000000000000000
000000000000000000000000000000000000000000000000000000000000
000000000000000000000000000000000000000000000000000000000000
000000000000000000000000000000000000000000000000000000000000
000000000000000000000000000000000000000000000000000000000000
000000000000000000000000000000000000000011111111000000000000
000000000000000000000000000000000000000000000000000000000000
000000000000000000000000000000000000000000000000000000000000

000000000000000000000000000000000000000000000000000000000000
0000000000000000000000000000
>DM_train104
MTHHDCAHCSSDACATEMLNLAEANSIETAWHRYEKQQPQCGFGSAGLCCRICLKGPCRIDP
FGEGPKYGVCGADRDTIVARHLVRMIAAGTAAHSEHGRHIALAMQHISQGELHDYSIRDEAK
LYAIAKTLGVATEGRGLLAIVGDLAAITLGDFQNQDYDKPCAWLAASLTPRRVKRLGDLGLL
PHNIDASVAQTMSRTHVGCDADPTNLILGGLRVAMADLDGSMLATELSDALFGTPQPVVSAA
NLGVMKRGAVNIAVNGHNPMLSDIICDVAADLRDEAIAAGAAEGINIIGICCTGHEVMMRHG
VPLATNYLSQELPILTGALEAMVVDVQCIMPSLPRIAECFHTQIITTDKHNKISGATHVPFD
EHKAVETAKTIIRMAIAAFGRRDPNRVAIPAFKQKSIVGFSAEAVVAALAKVNADDPLKPLV
DNVVNGNIQGIVLFVGCNTTKVQQDSAYVDLAKSLAKRNVLVLATGCAAGAFAKAGLMTSEA
TTQYAGEGLKGVLSAIGTAAGLGGPLPLVMHMGSCVDNSRAVALATALANKLGVDLSDLPLV
ASAPECMSEKALAIGSWAVTIGLPTHVGSVPPVIGSQIVTKLVTETAKDLVGGYFIVDTDPK
SAGDKLYAAIQERRAGLGL
111111111111111111111111111000000000000000000000000000000000
000000000000000000000000000000000000000000000000000000000000
000000000000000000000000000000000000000000000000000000000000
000000000000000000000000000000000000000000000000000000000000
000000000000000000000000000000000000000000000000000000000000
000000000000000000000000000000000000000000000000000000000000
000000000000000000000000000000000000000000000000000000000000
000000000000000000000000000000000000000000000000000000000000
000000000000000000000000000000000000000000000000000000000000
000000000000000000000000000000000000000000000000000000000000
0000000000000000000
>DM_train105
PILEKAPQKMPVKASSWEELDLPKLPVPPLQQTLATYLQCMQHLVPEEQFRKSQAIVKRFGA
PGGLGETLQEKLLERQEKTANWVSEYWLNDMYLNNRLALPVNSSPAVIFARQHFQDTNDQLR
FAACLISGVLSYKTLLDSHSLPTDWAKGQLSGQPLCMKQYYRLFSSYRLPGHTQDTLVAQKS
SIMPEPEHVIVACCNQFFVLDVVINFRRLSEGDLFTQLRKIVKMASNEDERLPPIGLLTSDG
RSEWAKARTVLLKDSTNRDSLDMIERCICLVCLDGPGTGELSDTHRALQLLHGGGCSLNGAN
RWYDKSLQFVVGRDGTCGVVCEHSPFDGIVLVQCTEHLLKHMMTSNKKLVRADSVSELPAPR
RLRWKCSPETQGHLASSAEKLQRIVKNLDFIVYKFDNYGKTFIKKQKYSPDGFIQVALQLAY
YRLYQRLVPTYESASIRRFQEGRVDNIRSATPEALAFVQAMTDHKAAMPASEKLQLLQTAMQ
AQTEYTVMAITGMAIDNHLLALRELARDLCKEPPEMFMDETYLMSNRFVLSTSQVPTTMEMF
CCYGPVVPNGYGACYNPQPEAITFCISSFHSCKETSSVEFAEAVGASLVDMRDLCSSRQPAD
SKPPAPKEKARGPSQAKQS
111111111111111111000000000000000000000000000000000000000000
000000000000000000000000000000000000000000000000000000000000
000000000000000000000000000000000000000000000000000000000000
000000000000000000000000000000000000000000000000000000000000
000000000000000000000000000000000000000000000000000000000000
000000000000000000000000000000000000000000000000000000000000
000000000000000000000000000000000000000000000000000000000000

000000000000000000000000000000000000000000000000000000000000
000000000000000000000000000000000000000000000000000000000000
000000000000000000000000000000000000000000000000000000011111
111111111111111111
>DM_train106
APAADAAQAHDPLSVQTGSDIPASVHMPTDQQRDYIKREVMVPMRDGVKLYTVIVIPKNARN
APILLTRTPYNAKGRANRVPNALTMREVLPQGDDVFVEGGYIRVFQDIRGKYGSQGDYVMTR
PPHGPLNPTKTDETTDAWDTVDWLVHNVPESNGRVGMTGSSYEGFTVVMALLDPHPALKVAA
PESPMVDGWMGDDWFHYGAFRQGAFDYFVSQMTARGGGNDIPRRDADDYTNFLKAGSAGSFA
TQAGLDQYPFWQRMHAHPAYDAFWQGQALDKILAQRKPTVPMLWEQGLWDQEDMWGAIHAWQ
ALKDADVKAPNTLVMGPWRHSGVNYNGSTLGPLEFEGDTAHQYRRDVFRPFFDEYLKPGSAS
VHLPDAIIYNTGDQKWDYYRSWPSVCESNCTGGLTPLYLADGHGLSFTHPAADGADSYVSDP
AHPVPFISRPFAFAQSSRWKPWLVQDQREAESRPDVVTYETEVLDEPVRVSGVPVADLFAAT
SGTDSDWVVKLIDVQPAMTPDDPKMGGYELPVSMDIFRGRYRKDFAKPEALQPDATLHYHFT
LPAVNHVFAKGHRIMVQIQSSWFPLYDRNPQKFVPNIFDAKPADYTVATQSIHHGGKEATSI
LLPVVKQEQKLISEEDL
111111111000000000000111111111000000000000000000000000000000
000000000000000000000000000000000000000000000000000000000000
000000000000000000000000000000000000000000000000000000000000
000000000000000000000000000000000000000000000000000000000000
000000000000000000000000000000000000000000000000000000000000
000000000000000000000000000000000000000000000000000000000000
000000000000000000000000000000000000000000000000000000000000
000000000000000000000000000000000000000000000000000000000000
000000000000000000000000000000000000000000000000000000000000
000000000000000000000000000000000000000000000000000000000000
0000001111111111
>DM_train107
GSHMLDNMDVIGERIKRIKEEHNSTWHYDDENPYKTWAYHGSYEVKATGSASSMINGVVKLL
TKPWDVVPMVTQMAMTDTTPFGQQRVFKEKVDTRTPRPLPGTRKVMEITAEWLWRTLGRNKR
PRLCTREEFTKKVRTNAAMGAVFTEENQWDSAKAAVEDEEFWKLVDRERELHKLGKCGSCVY
NMMGKREKKLGEFGKAKGSRAIWYMWLGARYLEFEALGFLNEDHWFSRENSYSGVEGEGLHK
LGYILRDISKIPGGAMYADDTAGWDTRITEDDLHNEEKIIQQMDPEHRQLANAIFKLTYQNK
VVKVQRPTPTGTVMDIISRKDQRGSGQVGTYGLNTFTNMEAQLVRQMEGEGVLTKADLENPH
LLEKKITQWLETKGVERLKRMAISGDDCVVKPIDDRFANALLALNDMGKVRKDIPQWQPSKG
WHDWQQVPFCSHHFHELIMKDGRKLVVPCRPQDELIGRARISQGAGWSLRETACLGKAYAQM
WSLMYFHRRDLRLASNAICSAVPVHWVPTSRTTWSIHAHHQWMTTEDMLTVWNRVWIEENPW
MEDKTPVTTWENVPYLGKREDQWCGSLIGLTSRATWAQNIPTAIQQVRSLIGNEEFLDYMPS
MKRFRKEEESEGAIW
111111100000000000000000000000000000000000001111100000000000
000000000000000000000000000000000000000000000000000000000000
000000000000000011111111111100000000000000000000000000000001
111111111111111111000000000000000000000000000000000000000000
000000000000000000000000000000000000000000000000000000000000

000000000000000000000000000000000000000000000000000000000000
000000000000000000000000000000000000000000000000000000000000
000000000000000000000000000000000000000000000000000000000000
000000000000000000000000000000000000000000000000000000000000
000000000000000000000000000000000000000000000000000000000011
111111111111111
>DM_train108
MAEQLVEAPAYARTLDRAVEYLLSCQKDEGYWWGPLLSNVTMEAEYVLLCHILDRVDRDRME
KIRRYLLHEQREDGTWALYPGGPPDLDTTIEAYVALKYIGMSRDEEPMQKALRFIQSQGGIE
SSRVFTRMWLALVGEYPWEKVPMVPPEIMFLGKRMPLNIYEFGSWARATVVALSIVMSRQPV
FPLPERARVPELYETDVPPRRRGAKGGGGWIFDALDRALHGYQKLSVHPFRRAAEIRALDWL
LERQAGDGSWGGIQPPWFYALIALKILDMTQHPAFIKGWEGLELYGVELDYGGWMFQASISP
VWDTGLAVLALRAAGLPADHDRLVKAGEWLLDRQITVPGDWAVKRPNLKPGGFAFQFDNVYY
PDVCDTAVVVWALNTLRLPDERRRRDAMTKGFRWIVGMQSSNGGWGAYDVDNTSDLPNHIPF
CDFGEVTDPPSEDVTAHVLECFGSFGYDDAWKVIRRAVEYLKREQKPDGSWFGRWGVNYLYG
TGAVVSALKAVGIDTREPYIQKALDWVEQHQNPDGGWGEDCRSYEDPAYAGKGASTPSQTAW
ALMALIAGGRAESEAARRGVQYLVETQRPDGGWDEPYYTGTGFPGDFYLGYTMYRHVFPTLA
LGRYKQAIERR
111111111000000000000000000000000000000000000000000000000000
000000000000000000000000000000000000000000000000000000000000
000000000000000000000000000000000000000000000000000000000000
000000000000000000000000000000000000000000000000000000000000
000000000000000000000000000000000000000000000000000000000000
000000000000000000000000000000000000000000000000000000000000
000000000000000000000000000000000000000000000000000000000000
000000000000000000000000000000000000000000000000000000000000
000000000000000000000000000000000000000000000000000000000000
000000000000000000000000000000000000000000000000000000000000
00000000111
>DM_train110
GELMPPQLQNGLNLSAKVVQGSLDSLPQAVREFLENNAELCQPDHIHICDGSEEENGRLLGQ
MEEEGILRRLKKYDNCWLALTDPRDVARIESKTVIVTQEQRDTVPIPKTGLSQLGRWMSEED
FEKAFNARFPGCMKGRTMYVIPFSMGPLGSPLSKIGIELTDSPYVVASMRIMTRMGTPVLEA
LGDGEFVKCLHSVGCPLPLQKPLVNNWPCNPELTLIAHLPDRREIISFGSGYGGNSLLGKKC
FALRMASRLAKEEGWLAEHMLVLGITNPEGEKKYLAAAFPSACGKTNLAMMNPSLPGWKVEC
VGDDIAWMKFDAQGHLRAINPENGFFGVAPGTSVKTNPNAIKTIQKNTIFTNVAETSDGGVY
WEGIDEPLASGVTITSWKNKEWSSEDGEPCAHPNSRFCTPASQCPIIDAAWESPEGVPIEGI
IFGGRRPAGVPLVYEALSWQHGVFVGAAMRSEATAAAEHKGKIIMHDPFAMRPFFGYNFGKY
LAHWLSMAQHPAAKLPKIFHVNWFRKDKEGKFLWPGFGENSRVLEWMFNRIDGKASTKLTPI
GYIPKEDALNLKGLGHINMMELFSISKEFWDKEVEDIEKYLVDQVNADLPCEIEREILALKQ
RISQM
111111111111000000000000000000000000000000000000000000000000
000000000000000000000000000000000000000000000000000000000000
000000000000000000000000000000000000000000000000000000000000

000000000000000000000000000000000000000000000000000000000000
000000000000000000000000000000000000000000000000000000000000
000000000000000000000000000000000000000000000000000000000000
000000000000000000000000000000000000000000000000000000000000
000000000000000000000000000000001111111100000000000000000000
000000000000000000000000000000000000000000000000000000000000
000000000000000000000000000000000000000000000000000000000000
00000
>DM_train112
STIRIGGAAVNQTPIDWENNVKNILDAIEEAKNANVEILCLPELCITGYGCEDLFLTDWVAE
TAIEYCFEIAASCTDITVSLGLPMRIAGITYNCVCLVENGIVKGFSAKQFLANEGVHYETRW
FTAWPRNHTTTFLYNDVKYPFGDVLYNVKDARIGFEICEDAWRTDRVGIRHYEKGATLVLNP
SASHFAFGKSAIRYDLVIGGSERFDCTYVYANLLGNEAGRMIYDGEVLIAHKGKLIQRNDRL
SFKNVNLIYADIATDSAETPETVLTQDDLEKEFEFWEATSLGLFDYMRKSRSKGFVLSLSGG
ADSSACAIMVAEMIRKGLKELGLTAFLQKSNMETLFDLPALQHLPFEEQAKKITAVFLTTAY
QSTRNSGDETYTSAKTLAESIGATFYNWSVDEEIEQYKATIENVIERPLTWEKDDITLQNIQ
ARGRAPIIWMLTNVKQALLITTSNRSEGDVGYATMDGDTAGGIAPIAGVDKDFIRSWLRWAE
KNRNQHGLHIVNKLAPTAELRPSEYTQTDERDLMPYDVLARIERKAIKERLSPVQVYTALLT
EGPYTKNEFKYWVKKFFRLWSINQWKRERLAPSFHMDDFNIDPRSWYRFPILSSGFAKELND
LDQ
000000000000000000000000000000000000000000000000000000000000
000000000000000000000000000000000000000000000000000000000000
000000000000000000000000000000000000001111111111000000000000
000000000000000000000000000000000000000000000000000000000000
000000000000001111111111111100000000000000000000000000000000
000000000000000000000000000000000000000000000000000000000000
000000000000000000000000000000000000000000000000000000000000
000000000000000000000000000000000000000000000000000000000000
000000000000111111111111111110000000000000000000000000000000
000000000000000000000000000000000000000000000000000000000000
000
>DM_train115
MYGWWGRILRVNLTTGEVKVQEYPEEVAKKFIGGRGLAAWILWNEARGVEPLSPENKLIFAA
GPFNGLPTPSGGKLVVAAKSPLTGGYGDGNLGTMASVHLRRAGYDALVVEGKAKKPVYIYIE
DDNVSILSAEGLWGKTTFETERELKEIHGKNVGVLTIGPAGENLVKYAVVISQEGRAAGRPG
MGAVMGSKKLKAVVIRGTKEIPVADKEELKKLSQEAYNEILNSPGYPFWKRQGTMAAVEWCN
TNYALPTRNFSDGYFEFARSIDGYTMEGMKVQQRGCPYCNMPCGNVVLDAEGQESELDYENV
ALLGSNLGIGKLNEVSVLNRIADEMGMDTISLGVSIAHVMEAVERGILKEGPTFGDFKGAKQ
LALDIAYRKGELGNLAAEGVKAMAEKLGTHDFAMHVKGLEVSGYNCYIYPAMALAYGTSAIG
AHHKEAWVIAWEIGTAPIEGEKAEKVEYKISYDPIKAQKVVELQRLRGGLFEMLTACRLPWV
EVGLSLDYYPKLLKAITGVTYTWDDLYKAADRVYSLIRAYWVREFNGKWDRKMDYPPKRWFT
EGLKSGPHKGEHLDEKKYDELLSEYYRIRGWDERGIPKKETLKELDLDFVIPELEKVTNLE
000000000000000000000000000000000000000000000000000000000000
000000000000000000000000000000000000000000000000000000000000

000000000000000000000000000000000000000000000000000000000000
000000000000000000000000000000000000000000000000000000000000
000000000000000000000000000000000000000000000000000000000000
000000000000000000000000000000000000000000000000000000000000
000000000000000000000000000000000000000000000000000000000000
000000000000000000111111110000000000000000000000000000000000
000000000000000000000000000000000000000000000000000000000000
00000000000000000000000000000000000000000000000000000000000
>DM_train116
MKPKKRQMEYLTRGLIAVQTEQGVFVSWRFLGTDHETTAFHLYRDGKRITRDPIAESTNFLD
QNGTADSVYQVAAVNKGREEKLSKKARVWQENVLEVPLAKPEGGVTPDGKPYTYSANDASVG
DIDGDGEYEMILKWDPSNSKDNAHDGYTGEVLIDAYKLDGTFLWRINLGRNIRAGAHYTQFM
VYDLDGDGKAEIAMKTADGTTDGKGHIIGDEQADFRNEQGRILSGPEYLTVFKGETGEALTT
VEYEPPRGKLEDWGDGYGNRMDRFLAGTAYLDGERPSLVMARGYYTRTVLVAYDFRNGRLKK
RWVFDSNQPGHEAYAGQGNHSLSVADVDGDGKDEIIYGAMAVDHDGTGLYSTGLGHGDAMHV
GDLDPSRKGLEVFQVHEDATKPYGLSLRDAGTGEILWGVHAGTDVGRGMAAHIDPSYKGSLV
WGIDPPGNDGMSYGLFTSKGEKISDKAPSSANFAIWWDGDLVRELLDHDWDGTIGRPKIEKW
DAENGCLKTIFQPAGVLSNNGTKGNPVLQANLFGDWREEVIWRTEDSSALRIYTTTHLTRHC
FYTLMHDPVYRLGIAWQNTAYNQPPHTSFYLGTGMKKPPKPALYIAGSKAEAPL
110000000000000000000000000000000000000000000000000000000000
000000000000000000000000000000000000000000000000000000000000
000000000000000000000000000000000000000000000000000000000000
000000000000000000000000000000000000000000000000000000000000
000000000000000000000000000000000000000000000000000000000000
000000000000000000000000000000000000000000000000000000000000
000000000000000000000000000000000000000000000000000000000000
000000000000000000000000000000000000000000000000000000000000
000000000000000000000000000000000000000000000000000000000000
0000000000000000000000000000000000000000000000111111
>DM_train117
MRYIAGIDIGNSSTEVALATLDEAGALTITHSALAETTGIKGTLRNVFGIQEALALVARGAG
IAVSDISLIRINEATPVIGDVAMETITETIITESTMIGHNPKTPGGAGLGTGITITPQELLT
RPADAPYILVVSSAFDFADIASVINASLRAGYQITGVILQRDDGVLVSNRLEKPLPIVDEVL
YIDRIPLGMLAAIEVAVPGKVIETLSNPYGIATVFNLSPEETKNIVPMARALIGNRSAVVVK
TPSGDVKARAIPAGNLELLAQGRSVRVDVAAGAEAIMKAVDGCGRLDNVTGESGTNIGGMLE
HVRQTMAELTNKPSSEIFIQDLLAVDTSVPVSVTGGLAGEFSLEQAVGIASMVKSDRLQMAM
IAREIEQKLNIDVQIGGAEAEAAILGALTTPGTTRPLAILDLGAGSTDASIINPKGDIIATH
LAGAGDMVTMIIARELGLEDRYLAEEIKKYPLAKVESLFHLRHEDGSVQFFSTPLPPAVFAR
VCVVKADELVPLPGDLALEKVRAIRRSAKERVFVTNALRALRQVSPTGNIRDIPFVVLVGGS
SLDFEVPQLVTDALAHYRLVAGRGNIRGSEGPRNAVATGLILSWHKEFAHER
000000000000000000000000000000000000000000000000000000000000
000000000000000000000000000000000000000000000000000000000000
000000000000000000000000000000000000000000000000000000000000
000000000000000000000000000000000000000000000000000000000000

00000000000000000000000000000000000000000000000000000000000000
00000000000000000000000000000000000000000000000000000000000000
00000000000000000000000000000000000000000000000000000000000000
00000000000000000000000000000000000000000000000000000000000000
00000000000000000000000000000000000000000000000000000000000000
0000000000000000000000000000000000000000000000111111
>DM_train120
MAKKVNWYVSCSPRSPEKIQPELKVLANFEGSYWKGVKGYKAQEAFAKELAALPQFLGTTYK
KEAAFSTRDRVAPMKTYGFVFVDEEGYLRITEAGKMLANNRRPKDVFLKQLVKWQYPSFQHK
GKEYPEEEWSINPLVFVLSLLKKVGGLSKLDIAMFCLTATNNNQVDEIAEEIMQFRNEREKI
KGQNKKLEFTENYFFKRFEKIYGNVGKIREGKSDSSHKSKIETKMRNARDVADATTRYFRYT
GLFVARGNQLVLNPEKSDLIDEIISSSKVVKNYTRVEEFHEYYGNPSLPQFSFETKEQLLDL
AHRIRDENTRLAEQLVEHFPNVKVEIQVLEDIYNSLNKKVDVETLKDVIYHAKELQLELKKK
KLQADFNDPRQLEEVIDLLEVYHEKKNVIEEKIKARFIANKNTVFEWLTWNGFIILGNALEY
KNNFVIDEELQPVTHAAGNQPDMEIIYEDFIVLGEVTTSKGATQFKMESEPVTRHYLNKKKE
LEKQGVEKELYCLFIAPEINKNTFEEFMKYNIVQNTRIIPLSLKQFNMLLMVQKKLIEKGRR
LSSYDIKNLMVSLYRTTIECERKYTQIKAGLEETLNNWVVDKEVRF
00000000000000000000000000000000000000000000000000000000001111
11100000000000000000000000000000000000000000000000000000000000
00000000000000000000000000000000000000000000000000000000000000
00000000000000000000000001111111111000000000000000000000000000
00000000000000000000000000000000000000000000000000000000000000
00000000000000000000000000000000000000000000000000000000000000
00000000000000000000000000000000000000000000000000000000000000
00000000000000000000000000000000000000000000000000000000000000
00000000000000000000000000000000000000000000000000000000000000
0000000000000000000000000000000000000000000000
>DM_train121
MASEIHMTGPMCLIENTNGRLMANPEALKILSAITQPMVVVAIVGLYRTGKSYLMNKLAGKK
KGFSLGSTVQSHTKGIWMWCVPHPKKPGHILVLLDTEGLGDVEKGDNQNDSWIFALAVLLSS
TFVYNSIGTINQQAMDQLYYVTELTHRIRSKSSPDENENEVEDSADFVSFFPDFVWTLRDFS
LDLEADGQPLTPDEYLTYSLKLKKGTSQKDETFNLPRLCIRKFFPKKKCFVFDRPVHRRKLA
QLEKLQDEELDPEFVQQVADFCSYIFSNSKTKTLSGGIQVNGPRLESLVLTYVNAISSGDLP
CMENAVLALAQIENSAAVQKAIAHYEQQMGQKVQLPTESLQELLDLHRDSEREAIEVFIRSS
FKDVDHLFQKELAAQLEKKRDDFCKQNQEASSDRCSGLLQVIFSPLEEEVKAGIYSKPGGYR
LFVQKLQDLKKKYYEEPRKGIQAEEILQTYLKSKESMTDAILQTDQTLTEKEKEIEVERVKA
ESAQASAKMLHEMQRKNEQMMEQKERSYQEHLKQLTEKMENDRVQLLKEQERTLALKLQEQE
QLLKEGFQKESRIMKNEIQDLQTKMRRRKACTIS
11111000000000000000000000000000000000000000000000000000000000
11111111111000000000000000000000000000000000000000000000000000
00000000000000000000000000000000111111111100000000000000000000
00011110000000000000000000000000000000000000000000000000011111
11111111000000000000000000000000000000000000000000000000000000
00000000000000000000000000000000000000000000000000000000000000

00000000000000000000000000000000000000000000000000000000000000
00000000000000000000000000000000000000000000000000000000000000
00000000000000000000000000000000000000000000000000000000000000
0000000000000000000000000111111111
>DM_train122
MALKTAVHEKIMNDVVLAVKKNAEWKVLIVDQLSMRMVSACCKMHEIMSEGITLVEDINRRR
EPLPLLEAVYLITPTEESVKCLMADFQNPDNPQYRGAHIFFTEACPEELFKELCKSTTARFI
KTLKEINIAFLPYESQIFSLDSPDTFQVYYNPSRAQGGIPNKERCAEQIATLCATLGEYPSV
RYRSDFDENASFAQLVQQKLDAYRADDPTMGEGPQKDRSQLLILDRGFDPISPLLHELTFQA
MAYDLLPIENDVYKYVNTGGNEVPEKEVLLDEKDDLWVEMRHQHIAVVSQNVTKKLKQFADE
KRMGTAADKAGIKDLSQMLKKMPQYQKELSKYSTHLHLAEDCMKQYQQHVDKLCKVEQDLAM
GTDADGEKIRDHMRNIVPILLDQKISAYDKIRIILLYIIHKGGISEENLAKLVQHAHIPAEE
KWIINDMQNLGVPIIQDGGRRKIPQPYHTHNRKERQADHTYQMSRWTPYMKDIMEAAVEDKL
DTRHYPFLNGGGPRPSCQQPVSVRYGHWHKDKGQASYKSGPRLIIFVVGGISYSEMRSAYEV
TQTAKNNWEVILGSTHILTPEGLLRDLRKISNP
00000000000000000000000000000000000000000000000000000000000000
00000000000000000000000000000000000000000000000000000000000000
00000000000000000000000000000000000000000000000000000000000000
00000000000000000000000000000000000000000000000000000000000000
00000000000000001111111100000000000000000000000000000000000000
00111111111110000000000000000000000000000000000000000000000000
00000000000000000000000000000000000000000000000000000000000000
00000000000000000000000000000000000000000000000000000000000000
00000000000011111111111111111111111111100000000000000000000000
000000000000000000000000000000000
>DM_train123
IETGYTPIDISLSLTQFLLSEFVPGAGFVLGLVDIIWGIFGPSQWDAFLVQIEQLINQRIEE
FARNQAISRLEGLSNLYQIYAESFREWEADPTNPALREEMRIQFNDMNSALTTAIPLLAVQN
YQVPLLSVYVQAANLHLSVLRDVSVFGQRWGFDAATINSRYNDLTRLIGNYTDYAVRWYNTG
LERVWGPDSRDWVRYNQFRRELTLTVLDIVALFSNYDSRRYPIRTVSQLTREIYTNPVLENF
DGSFRGMAQRIEQNIRQPHLMDILNSITIYTDVHRGFNYWSGHQITASPVGFSGPEFAFPLF
GNAGNAAPPVLVSLTGLGIFRTLSSPLYRRIILGSGPNNQELFVLDGTEFSFASLTTNLPST
IYRQRGTVDSLDVIPPQDNSVPPRAGFSHRLSHVTMLSQAAGAVYTLRAPTFSWQHRSAEFN
NIIPSSQITQIPLTKSTNLGSGTSVVKGPGFTGGDILRRTSPGQISTLRVNITAPLSQRYRV
RIRYASTTNLQFHTSIDGRPINQGNFSATMSSGSNLQSGSFRTVGFTTPFNFSNGSSVFTLS
AHVFNSGNEVYIDRIEFVPAEVTFEAEYDLER
11110000000000000000000000000000000000000000000000000000000000
00000000000000000000000000000000000000000000000000000000000000
00000000000000000000000000000000000000000000000000000000000000
00000000000000000000000000000000000000000000000000000000000000
00000000000000000000000000000000000000000000000000000000000000
00000000000000000000000000000000000000000000000000000000000000
00000000000000000000000000000000000000000000000000000000000000
00000000000000000000000000000000000000000000000000000000000000

000000000000000000000000000000000000000000000000000000000000
0000000000000000000000111111111
>DM_train124
LVTDEAEASKFVEEYDRTSQVVWNEYAEANWNYNTNITTETSKILLQKNMQIANHTLKYGTQ
ARKFDVNQLQNTTIKRIIKKVQDLERAALPAQELEEYNKILLDMETTYSVATVCHPNGSCLQ
LEPDLTNVMATSRKYEDLLWAWEGWRDKAGRAILQFYPKYVELINQAARLNGYVDAGDSWRS
MYETPSLEQDLERLFQELQPLYLNLHAYVRRALHRHYGAQHINLEGPIPAHLLGNMWAQTWS
NIYDLVVPFPSAPSMDTTEAMLKQGWTPRRMFKEADDFFTSLGLLPVPPEFWNKSMLEKPTD
GREVVCHASAWDFYNGKDFRIKQCTTVNLEDLVVAHHEMGHIQYFMQYKDLPVALREGANPG
FHEAIGDVLALSVSTPKHLHSLNLLSSEGGSDEHDINFLMKMALDKIAFIPFSYLVDQWRWR
VFDGSITKENYNQEWWSLRLKYQGLCPPVPRTQGDFDPGAKFHIPSSVPYIRYFVSFIIQFQ
FHEALCQAAGHTGPLHKCDIYQSKEAGQRLATAMKLGFSRPWPEAMQLITGQPNMSASAMLS
YFKPLLDWLRTENELHGEKLGWPQYNWTPNS
111000000000000000000000000000000000000000000000000000000000
000000000000000000000000000000000000000000000000000000000000
000000000000000000000000000000000000000000000000000000000000
000000000000000000000000000000000000000000000000000000000000
000000000000000000000000000000000000000000000000000000000000
000000000000000000000000000000000000000000000000000000000000
000000000000000000000000011110000000000000000000000000000000
000000000000000000000000000000000000000000000000000000000000
000000000000000000000000000000000000000000000000000000000000
000000000000000000000011111111
>DM_train127
GAMGSPKEHIDLYQQIKWNGWGDTRKFLHQLKPSGTIAMTTPEVSSVPLPSLRGFIKKELTL
PGEEDKPFVLDETPALQIENIHVDPPKQYPEFVRELKAFFLPDQLKDDKLARITHTFGKSLR
DLIRVRIGQVKNAPDLIVLPHSHEEVERLVQLAHKYNVVIIPMGGGSNIVGAIEPVSNERFT
VSIDMRRMNKVLWVDRREMTACIQVGIMGPELEKQLHKQGVSLGHDPDSFEFSTLGGWLATC
SSGHQSDKYGDIEDMAVSFRTVTPTGTLELRNGARSGAGINYKHIILGSEGTLGIITEAVMK
VHAVPQAVEYYGFLFPTFAHAVSALQQIRSSEVIPTMIRVYDPEETQLSFAWKPSKGAVSEF
TSAMVKKYLHYIRSFDFKNVCLSIIGFEGPKKVVDFHRTSVFDILSKNAAFGLGSAPGKTWA
EKRYDLPYIRDFLLDHNMWVDVAETTVSYANLQTLWKDAKQTFVKHFKDQGIPAWICAHISH
TYTNGVCLYFIFASKQNENKDMAQYIEAKKLMTDIIFKYGGSLSHHHGVGYEHVPWMTRYAT
RGWINVYRSLKETIDPKDICNPRKLI
000000000000000000000000000000000000000000000000000000000111
111111100000000000000000000000000000000000000000000000000000
000000000000000000000000000000000000000000000000000000000000
000000000000000000000000000000000000000000000000000000000000
000000000000000000000000000000111111000000000000000000000000
000000000000000000000000000000000000000000000000000001111000
000000000000000000000000000000000000000000000000000000000000
000000000000000000000000000000000000000000000000000000000000
000000000000000001111100000000000000000001111111111111111111
1000000000000000000000011

>DM_train128
PVFSADPGAPAPVNPCCYYPCQHQGICVRFGLDRYQCDCTRTGYSGPNCTIPEIWTWLRTTL
RPSPSFIHFLLTHGRWLWDFVNATFIRDTLMRLVLTVRSNLIPSPPTYNIAHDYISWESFSN
VSYYTRILPSVPRDCPTPMGTKGKKQLPDAEFLSRRFLLRRKFIPDPQGTNLMFAFFAQHFT
HQFFKTSGKMGPGFTKALGHGVDLGHIYGDNLERQYQLRLFKDGKLKYQMLNGEVYPPSVEE
APVLMHYPRGIPPQSQMAVGQEVFGLLPGLMLYATIWLREHNRVCDLLKAEHPTWGDEQLFQ
TARLILIGETIKIVIEEYVQQLSGYFLQLKFDPELLFGAQFQYRNRIAMEFNQLYHWHPLMP
DSFRVGPQDYSYEQFLFNTSMLVDYGVEALVDAFSRQPAGRIGGGRNIDHHILHVAVDVIKE
SRVLRLQPFNEYRKRFGMKPYTSFQELTGEKEMAAELEELYGDIDALEFYPGLLLEKCHPNS
IFGESMIEMGAPFSLKGLLGNPICSPEYWKASTFGGEVGFNLVKTATLKKLVCLNTKTCPYV
SFHVPDPRQEDRPGVERPPTEL
111111111111000000000000000000000000000000000000000000000000
000000000000000000000000000000000000000000000000000000000000
000000000000000000000000000000000000000000000000000000000000
000000000000000000000000000000000000000000000000000000000000
000000000000000000000000000000000000000000000000000000000000
000000000000000000000000000000000000000000000000000000000000
000000000000000000000000000000000000000000000000000000000000
000000000000000000000000000000000000000000000000000000000000
000000000000000000000000000000000000000000000000000000000000
0000011111111111111111
>DM_train129
MVSKIRTFGWVQNPGKFENLKRVVQVFDRNSKVHNEVKNIKIPTLVKESKIQKELVAIMNQH
DLIYTYKELVGTGTSIRSEAPCDAIIQATIADQGNKKGYIDNWSSDGFLRWAHALGFIEYIN
KSDSFVITDVGLAYSKSADGSAIEKEILIEAISSYPPAIRILTLLEDGQHLTKFDLGKNLGF
SGESGFTSLPEGILLDTLANAMPKDKGEIRNNWEGSSDKYARMIGGWLDKLGLVKQGKKEFI
IPTLGKPDNKEFISHAFKITGEGLKVLRRAKGSTKFTRVPKRVYWEMLATNLTDKEYVRTRR
ALILEILIKAGSLKIEQIQDNLKKLGFDEVIETIENDIKGLINTGIFIEIKGRFYQLKDHIL
QFVIPNRLGKPDLVKSELEEKKSELRHKLKYVPHEYIELIEIARNSTQDRILEMKVMEFFMK
VYGYRGKHLGGSRKPDGAIYTVGSPIDYGVIVDTKAYSGGYNLPIGQADEMQRYVEENQTRN
KHINPNEWWKVYPSSVTEFKFLFVSGHFKGNYKAQLTRLNHITNCNGAVLSVEELLIGGEMI
KAGTLTLEEVRRKFNNGEINF
111100000000000000000000000000000000000000000000000000000000
000000000000001111100000000000000000000000000000000000000000
000000000000000000000000000000000000000000000000000000000000
000000000000000000000000000000000000000000000000000000000000
000111110000000000000000000000000000000000000000000000000000
000000000000000000000000000000000000000000000000000000000000
000000011111000000000000000000000000000000000000000000000000
000000000000000000000000000000000000000000000000000000000000
000000000000000000000000000000000000000000000000000000000000
000000000000000000000
>DM_train130
SMSYTWTGALITPCAAEESKLPINALSNSLLRHHNMVYATTSRSAGLRQKKVTFDRLQVLDD

HYRDVLKEMKAKASTVKAKLLSVEEACKLTPPHSAKSKFGYGAKDVRNLSSKAVNHIHSVWK
DLLEDTVTPIDTTIMAKNEVFCVQPEKGGRKPARLIVFPDLGVRVCEKMALYDVVSTLPQVV
MGSSYGFQYSPGQRVEFLVNTWKSKKNPMGFSYDTRCFDSTVTENDIRVEESIYQCCDLAPE
ARQAIKSLTERLYIGGPLTNSKGQNCGYRRCRASGVLTTSCGNTLTCYLKASAACRAAKLQD
CTMLVNGDDLVVICESAGTQEDAASLRVFTEAMTRYSAPPGDPPQPEYDLELITSCSSNVSV
AHDASGKRVYYLTRDPTTPLARAAWETARHTPVNSWLGNIIMYAPTLWARMILMTHFFSILL
AQEQLEKALDCQIYGACYSIEPLDLPQIIERLHGLSAFSLHSYSPGEINRVASCLRKLGVPP
LRVWRHRARSVRARLLSQGGRAATCGKYLFNWAVKTKLKLTPIPAASQLDLSGWFVAGYSGG
DIYHSLSRARPRGSHHHHHH
000000000000000000000000000000000000000000000000000000000000
000000000000000000000000000000000000000000000000000000000000
000000000000000000000000000000000000000000000000000000000000
000000000000000000000000000000000000000000000000000000000000
000000000000000000000000000000000000000000000000000000000000
000000000000000000000000000000000000000000000000000000000000
000000000000000000000000000000000000000000000000000000000000
000000000000000000000000000000000000000000000000000000000000
000000000000000000000000000000000000000000000111000000000011
11111111111111111111
>DM_train131
GSNQSSSTSTKKLKAGNFDVAYQNPDKAIKGGNLKVAYQSDSPMKAQWLSGLSNDATFATMS
GPGGGQDGLFFTDSGFKFIKGGAADVALDKESKTATITLRKDLKWSDGSEVTAKDYEFTYET
IANPAYGSDRWTDSLANIVGLSDYHTGKAKTISGITFPDGENGKVIKVQFKEMKPGMTQSGN
GYFLETVAPYQYLKDVAPKDLASSPKTTTKPLVTGPFKPENVVAGESIKYVPNPYYWGEKPK
LNSITYEVVSTAKSVAALSSSKYDIINGMVSSQYKQVKNLKGYKVLGQQAMYISLMYYNLGH
YDAKNSINVQDRKTPLQDQNVRQAIGYARNVAEVDNKFSNGLSTPANSLIPPIFKQFTSSSV
KGYEKQDLDKANKLLDEDGWKLNKSTGYREKDGKELSLVYAARVGDANAETIAQNYIQQWKK
IGVKVSLYNGKLMEFNSWVDHMTTPPGANDWDITDGSWSLASEPSQQDLFSAAAPYNFGHFN
DSEITKDLNDIDSAKSENPTYRKAAFVKYQEDMNKKAYVIPTNFMLNYTPVNKRVVGMTLDY
GAMNTWSEIGVSSAKLATK
111111111111111111111111111110000000000000000000000000000000
000000000000000000000000000000000000000000000000000000000000
000000000000000000000000000000000000000000000000000000000000
000000000000000000000000000000000000000000000000000000000000
000000000000000000000000000000000000000000000000000000000000
000000000000000000000000000000000000000000000000000000000000
000000000000000000000000000000000000000000000000000000000000
000000000000000000000000000000000000000000000000000000000000
000000000000000000000000000000000000000000000000000000000000
0000000000011111111
>DM_train132
MKHMPRKMYSCAFETTTKVEDCRVWAYGYMNIEDHSEYKIGNSLDEFMAWVLKVQADLYFHN
LKFAGAFIINWLERNGFKWSADGLPNTYNTIISRMGQWYMIDICLGYKGKRKIHTVIYDSLK
KLPFPVKKIAKDFKLTVLKGDIDYHKERPVGYKITPEEYAYIKNDIQIIAEALLIQFKQGLD

RMTAGSDSLKGFKDIITTKKFKKVFPTLSLGLDKEVRYAYRGGFTWLNDRFKEKEIGEGMVF
DVNSLYPAQMYSRLLPYGEPIVFEGKYVWDEDYPLHIQHIRCEFELKEGYIPTIQIKRSRFY
KGNEYLKSSGGEIADLWLSNVDLELMKEHYDLYNVEYISGLKFKATTGLFKDFIDKWTYIKT
TSEGAIKQLAKLMLNSLYGKFASNPDVTGKVPYLKENGALGFRLGEEETKDPVYTPMGVFIT
AWARYTTITAAQACYDRIIYCDTDSIHLTGTEIPDVIKDIVDPKKLGYWAHESTFKRAKYLR
QKTYIQDIYMKEVDGKLVEGSPDDYTDIKFSVKCAGMTDKIKKEVTFENFKVGFSRKMKPKP
VQVPGGVVLVDDTFTIK
111100000000000000000000000000000000000000000000000000000000
000000000000000000000000000000000000000000000000000000000000
000000000000000000000000000000000000000000000000000000000000
000000000000000000000000000000000000000000000000000000000000
000000000000000000000000000000000000000000000000000000111111
111000000000000000000000000000000000000000000000000000000000
000000000000000000000000000000000000000000000000000000000000
000000000000000000000000000000000000000000000000000000000000
000000000000000000000000000000000000000000000000000000000000
00000000000000000
>DM_train133
HGTTANTHCGADFCTWWHDSGEINTQTPVQPGNVRQSHKYSVQVSLAGTNNFHDSFVYESIP
RNGNGRIYAPTDPPNSNTLDSSVDDGISIEPSIGLNMAWSQFEYSHDVDVKILATDGSSLGS
PSDVVIRPVSISYAISQSDDGGIVIRVPADANGRKFSVEFKTDLYTFLSDGNEYVTSGGSVV
GVEPTNALVIFASPFLPSGMIPHMTPDNTQTMTPGPINNGDWGAKSILYFPPGVYWMNQDQS
GNSGKLGSNHIRLNSNTYWVYLAPGAYVKGAIEYFTKQNFYATGHGILSGENYVYQANAGDN
YIAVKSDSTSLRMWWHNNLGGGQTWYCVGPTINAPPFNTMDFNGNSGISSQISDYKQVGAFF
FQTDGPEIYPNSVVHDVFWHVNDDAIKIYYSGASVSRATIWKCHNDPIIQMGWTSRDISGVT
IDTLNVIHTRYIKSETVVPSAIIGASPFYASGMSPDSRKSISMTVSNVVCEGLCPSLFRITP
LQNYKNFVVKNVAFPDGLQTNSIGTGESIIPAASGLTMGLAISAWTIGGQKVTMENFQANSL
GQFNIDGSYWGEWQIS
110000000000000000000000000000000000000000000000000000000000
000000000000000000000000000000000000000000000000000000000000
000000000000000000000000000000000000000000000000000000000000
000000000000000000000000000000000000000000000000000000000000
000000000000000000000000000000000000000000000000000000000000
000000000000000000000000000000000000000000000000000000000000
000000000000000000000000000000000000000000000000000000000000
000000000000000000000000000000000000000000000000000000000000
000000000000000000000000000000000000000000000000000000000000
0000000000000000
>DM_train137
SNISRQAYADMFGPTVGDKVRLADTELWIEVEDDLTTYGEEVKFGGGKVIRDGMGQGQMLAA
DCVDLVLTNALIVDHWGIVKADIGVKDGRIFAIGKAGNPDIQPNVTIPIGAATEVIAAEGKI
VTAGGIDTHIHWICPQQAEEALVSGVTTMVGGGTGPAAGTHATTCTPGPWYISRMLQAADSL
PVNIGLLGKGNVSQPDALREQVAAGVIGLCIHEDWGATPAAIDCALTVADEMDIQVALHSDT
LNESGFVEDTLAAIGGRTIHTFHTEGAGGGHAPDIITACAHPNILPSSTNPTLPYTLNTIDE

HLDMLMVAHHLDPDIAEDVAFAESRIRRETIAAEDVLHDLGAFSLTSSDSQAMGRVGEVILR
TWQVAHRMKVQRGALAEETGDNDNFRVKRYIAKYTINPALTHGIAHEVGSIEVGKLADLVVW
SPAFFGVKPATVIKGGMIAIAPMGDINASIPTPQPVHYRPMFGALGSARHHCRLTFLSQAAA
ANGVAERLNLRSAIAVVKGCRTVQKADMVHNSLQPNITVDAQTYEVRVDGELITSEPADVLP
MAQRYFLF
000000000000000000000000000000000000000000000000000000000000
000000000000000000000000000000000000000000000000000000000000
000000000000000000000000000000000000000000000000000000000000
000000000000000000000000000000000000000000000000000000000000
000000000000000000000000000000000000000000000000000000000111
111111111111111111111111111000000000000000000000000000000000
000000000000000000000000000000000000000000000000000000000000
000000000000000000000000000000000000000000000000000000000000
000000000000000000000000000000000000000000000000000000000000
00000000
>DM_train138
GHMNIESTLNSVASVKDLANEASKYEIILQKGINQVGLKQYTQVVHKLDDMLEDIQSGQANR
EENSEFHGILTHLEQLIKRSEAQLRVYFISILNSIKPFDPQINITKKMPFPYYEDQQLGALS
WILDYFHGNSEGSIIQDILVGERSKLILKCMAFLEPFAKEISTAKNAPYEKGSSGMNSYTEA
LLGFIANEKSLVDDLYSQYTESKPHVLSQILSPLISAYAKLFGANLKIVRSNLENFGFFSFE
LVESINDVKKSLRGKELQNYNLLQDCTQEVRQVTQSLFRDAIDRIIKKANSISTIPSNNGVT
EATVDTMSRLRKFSEYKNGCLGAMDNITRENWLPSNYKEKEYTLQNEALNWEDHNVLLSCFI
SDCIDTLAVNLERKAQIALMPNQEPDVANPNSSKNKHKQRIGFFILMNLTLVEQIVEKSELN
LMLAGEGHSRLERLKKRYISYMVSDWRDLTANLMDSVFIDSSGKKSKDKEQIKEKFRKFNEG
FEDLVSKTKQYKLSDPSLKVTLKSEIISLVMPMYERFYSRYKDSFKNPRKHIKYTPDELTTV
LNQLVR
111111100000000000000000000000000000000000000000000000011110
000000000000000000000000000000000000000000000000000000000000
000000000000000000000000000000000000111111111111000000000000
000000000000000000000000000000000000000000000000111110000000
000000000000000000000000000000000000000000000000000000000000
000000000000000000000000000000000000000000011110000000000000
000000000000000000000000000000000000000000000000000000000000
000000000000000000000000000000000000000000000000000000000000
000000000000000000000000000000000000000000000000000000000000
000000
>DM_train140
MTTAARRPAPTTAGAGWDAGVGALVNPSRRRGGTLRLVSSADVDSLDPARTYYVWVWLLQRL
LNRTLMAYPTDPGPAGLVPAPDLAEGPGEVSDGGRTWTYRLRRGLRYDDGTPITSDDVRHAV
QRVFAQDVLPGGPTYLIPLLDDPERPYPGPYRTDEPLRSVLTPDEHTIVFRLTRPFSDFDHL
MAQPCAAPVPRRSDTGADYGRDPRSSGPYRVARHEPDTLLHLERNPHWDRATDPIRPALPDR
VELTIGLDVDVLDARLIAGEFDINLEGRGLQHAAQRRATADEVLRSHTDNPRTSFLHFVAMQ
PHIPPFDNVHVRRAVQYAADKILLQDARGGPVNGGDLTTALFPPTLPAHQDLDLYPTGPDLR
GDLDAARAELAAAGLPDGFRAVIGTQRGKFRLVADAVVESLARVGIELTVKELDVATYFSLG

AGHPETVREHGLGLLVTDWGADFPTEYGFLAPLVDGRQIKRNGGNWNLPELDDPEVNALIDE
TLHTTDPAARAELWRAVERRVMEHAVLLPLVHDKTLHFRNPWVTNVYVHPAFGLYDIQAMGL
AEED
1111111000000000000000000000000000000000000000000000000000000000
0000000000000000000000000000000000000000000000000000000000000000
0000000000000000000000000000000000000000000000000000000000000000
0000000000000000000000000000000000000000000000000000000000000000
0000000000000000000000000000000000000000000000000000000000000000
0000000000000000000000000000000000000000000000000000000000000000
0000000000000000000000000000000000000000000000000000000000000000
0000000000000000000000000000000000000000000000000000000000000000
0000000000000000000000000000000000000000000000000000000000000000
0011
>DM_train141
STGPVAPLPTPPNFPNDIALFQQAYQNWSKEIMLDATWVCSPKTPQDVVRLANWAHEHDYKI
RPRGAMAGWTPLTVEKGANVEKVILADTMTHLNGITVNTGGPVATVTAGAGASIEAIVTELQ
KHDLGWANLPAPGVLSIGGALAVNAHGAALPAVGQTTLPGHTYGSLSNLVTELTAVVWNGTT
YALETYQRNDPRITPLLTNLGRCFLTSVTMQAGPNFRQRCQSYTDIPWRELFAPKGADGRTF
EKFVAESGGAEAIWYPFTEKPWMKVWTVSPTKPDSSNEVGSLGSAGSLVGKPPQAREVSGPY
NYIFSDNLPEPITDMIGAINAGNPGIAPLFGPAMYEITKLGLAATNANDIWGWSKDVQFYIK
ATTLRLTEGGGAVVTSRANIATVINDFTEWFHERIEFYRAKGEFPLNGPVEIRCCGLDQAAD
VKVPSVGPPTISATRPRPDHPDWDVAIWLNVLGVPGTPGMFEFYREMEQWMRSHYNNDDATF
RPEWSKGWAFGPDPYTDNDIVTNKMRATYIEGVPTTENWDTARARYNQIDPHRVFTNGFMDK
LLP
1111100000000000000000000000000000000000000000000000000000000000
0000000000000000000000000000000000000000000000000000000000000000
0000000000000000000000000000000000000000000000000000000000000000
0000000000000000000000000000000000000000000000000000000000000000
0000000000000000000000000000001111111111111111111110000000000000
0000000000000000000000000000000000000000000000000000000000000000
0000000000000000000000000000000000000000000000000000000000000000
0000000000000000000000000000000000000000000000000000000000000000
0000000000000000000000000000000000000000000000000000000000000000
000
>DM_train142
YDPDQYSIEADKKFKYSVKLSDYPTLQDAASAAVDGLLIDRDYNFYGGETVDFGGKVLTIEC
KAKFIGDGNLIFTKLGKGSRIAGVFMESTTTPWVIKPWTDDNQWLTDAAAVVATLKQSKTDG
YQPTVSDYVKFPGIETLLPPNAKGQNITSTLEIRECIGVEVHRASGLMAGFLFRGCHFCKMV
DANNPSGGKDGIITFENLSGDWGKGNYVIGGRTSYGSVSSAQFLRNNGGFERDGGVIGFTSY
RAGGSGVKTWQGTVGSTTSRNYNLQFRDSVVIYPVWDGFDLGADTDMNPELDRPGDYPITQY
PLHQLPLNHLIDNLLVRGALGVGFGMDGKGMYVSNITVEDCAGSGAYLLTHESVFTNIAIID
TNTKDFQANQIYISGACRVNGLRLIGIRSTDGQSLTIDAPNSTVSGITGMVDPSRINVANLA
EEGLGNIRANSFGYDSAAIKLRIHKLSKTLDSGALYSHINGGAGSGSAYTQLTAISGSTPDA
VSLKVNHKDCRGAEIPFVPDIASDDFIKDSSCFLPYWENNSTSLKALVKKPNGELVRLTLAT

L
111110000000000000000000000000000000000000000000000000000000
000000000000000000000000000000000000000000000000000000000000
000000000000000000000000000000000000000000000000000000000000
000000000000000000000000000000000000000000000000000000000000
000000000000000000000000000000000000000000011111100000000000
000000000000000000000000000000000000000000000000000000000000
000000000000000000000000000111110000000000000000000000000000
000000000000000000000000000000000000000000000000000000000000
000000000000000000000000000000000000000000000000000000000000
0
>DM_train143
MTTANTPVRPKSAIDAVADAYTEKLIELNPSFATTLGLPGHETEYQDYSPAGAAAHAEATRL
ALEALAGLEPSDDVDAVTLDAMRERLGLELEIHQSGWDAADLNNIASPAQDIRAIFDLMPTD
TVEHWEHIAGRAANVPGAIEGYIASLRAAKDDRKVAAARQIRIVIEQTGRYAAEDGFFAKMA
ADASLGDAPLPAEVQDKLDAGTSAARSAYSALGAFLRDELLPVAPEKDAVGRERYSLASRSF
IGAEVDLEETYAWGVQELERLISEQEKVAGQIKPGASIEEAKSILNNDPARQIKGTDALKAW
MQELSDRAVSELADVHFDIPDVMKTLECMIAPTDEGGIYYTGPSDDFSRPGRMWWSVPAGED
TFTTWSETTTVFHEGVPGHHLQVATATYRRELLNNWRRNVCWVSGHGEGWALYAEQLMLELG
YLKDPGDHMGMLDGQRMRAARVVFDIGVHLELPVPERWGTGTWTPEKGFDFLKANLDISEGQ
LQFEFTRYLGWPGQAPSYKVGQRLWEQIRAELESREGFDLKSFHSKALNIGSVGLDVLRRAL
L
111111100000000000000000000000000000000000000000000000000000
000000000000000000000000000000000000000000000000000000000000
000000000000000000000000000000000000000000000000000000000000
000000000000000000000000000000000000000000000000000000000000
000000000000000000000000000000000000000000000000000000000000
000000000000000000000000000000000110000000000000000000000000
000000000000000000000000000000000000000000000000000000000000
000000000000000000000000000000000000000000000000000000000000
000000000000000000000000000000000000000000000000000000000000
0
>DM_train144
MSKVPSDIEIAQAAKMKPVMELARGLGIQEDEVELYGKYKAKISLDVYRRLKDKPDGKLILV
TAITPTPAGEGKTTTSVGLTDALARLGKRVMVCLREPSLGPSFGIKGGAAGGGYAQVVPMED
INLHFTGDIHAVTYAHNLLAAMVDNHLQQGNVLNIDPRTITWRRVIDLNERALRNIVIGLGG
KANGVPRETGFDISVASEVMACLCLASDLMDLKERFSRKVVGYTYDGKPVTAGDLEAQGSMA
LLMKDAIKPNLVQTLENTPAFIHGGPFANIAHGCNSIIATKTALKLADYVVTEAGFGADLGA
EKFYDVKCRYAGFKPDATVIVATVRALKMHGGVPKSDLATENLEALREGFANLEKHIENIGK
FGVPAVVAINAFPTDTEAELNLLYELCAKAGAEVALSWAKGGEGGLELARKVLQTLESRPSN
FHVLYNLDLSIKDKIAKIATEIYGADGVNYTAEADKAIQRYESLGYGNLPVVMAKTQYSFSD
DMTKLGRPRNFTITVREVRLSAGGRLIVPITGAIMTMPGLPKRPAACNIDIDADGVITGLF
111111100000000000000000000000000000000000000000000000000000
000000000000000000000000000000000000000000000000000000000000

000000000000000000000000000000000000000000000000000000000000
000000000000000000000000000000000000000000000000000000000000
000000000000000000000000000000000000000000000000000000000000
000000000000000000000000000000000000000000000000000000000000
000000000000000000000000000000000000000000000000000000000000
000000000000000000000000000000000000000000000000000000000000
00000000000000000000000000000000000000000000000000000000000
>DM_train145
MSVSAFNRRWAAVILEALTRHGVRHICIAPGSRSTLLTLAAAENSAFIHHTHFDERGLGHLA
LGLAKVSKQPVAVIVTSGTAVANLYPALIEAGLTGEKLILLTADRPPELIDCGANQAIRQPG
MFASHPTHSISLPRPTQDIPARWLVSTIDHALGTLHAGGVHINCPFAEPLYGEMDDTGLSWQ
QRLGDWWQDDKPWLREAPRLESEKQRDWFFWRQKRGVVVAGRMSAEEGKKVALWAQTLGWPL
IGDVLSQTGQPLPCADLWLGNAKATSELQQAQIVVQLGSSLTGKRLLQWQASCEPEEYWIVD
DIEGRLDPAHHRGRRLIANIADWLELHPAEKRQPWCVEIPRLAEQAMQAVIARRDAFGEAQL
AHRICDYLPEQGQLFVGNSLVVRLIDALSQLPAGYPVYSNRGASGIDGLLSTAAGVQRASGK
PTLAIVGDLSALYDLNALALLRQVSAPLVLIVVNNNGGQIFSLLPTPQSERERFYLMPQNVH
FEHAAAMFELKYHRPQNWQELETAFADAWRTPTTTVIEMVVNDTDGAQTLQQLLAQVSHL
100000000000000000000000000000000000000000000000000000000000
000000000000000000000000000000000000000000000000000000000000
000000000000000000000000000000000000000000000000110000000000
000000000000000000000000000000000000000000000000000000000000
000000000000000000000000000000000000000000000000000000000000
000000000000000000000000000000000000000000000000000000000000
000000000000000000000000000000000000000000000000000000000000
000000000000000000000000000000000000111111111111111111110000
0000000000000000000000000000000000000000000000000000000000
>DM_train146
MKRSKRFAVLAQRPVNQDGLIGEWPEEGLIAMDSPFDPVSSVKVDNGLIVELDGKRRDQFDM
IDRFIADYAINVERTEQAMRLEAVEIARMLVDIHVSREEIIAITTAITPAKAVEVMAQMNVV
EMMMALQKMRARRTPSNQCHVTNLKDNPVQIAADAAEAGIRGFSEQETTVGIARYAPFNALA
LLVGSQCGRPGVLTQCSVEEATELELGMRGLTSYAETVSVYGTEAVFTDGDDTPWSKAFLAS
AYASRGLKMRYTSGTGSEALMGYSESKSMLYLESRCIFITKGAGVQGLQNGAVSCIGMTGAV
PSGIRAVLAENLIASMLDLEVASANDQTFSHSDIRRTARTLMQMLPGTDFIFSGYSAVPNYD
NMFAGSNFDAEDFDDYNILQRDLMVDGGLRPVTEAETIAIRQKAARAIQAVFRELGLPPIAD
EEVEAATYAHGSNEMPPRNVVEDLSAVEEMMKRNITGLDIVGALSRSGFEDIASNILNMLRQ
RVTGDYLQTSAILDRQFEVVSAVNDINDYQGPGTGYRISAERWAEIKNIPGVVQPDTIE
000000000000000000000000000000000000000000000000000000000000
000000000000000000000000000000000000000000000000000000000000
000000000000000000000000000000000000000000000000000000000000
000000000000000000000000000000000000000000000000000000000000
000000000000000000000000000000000000000000000000000000000000
000000000000000000000000000000000000000000000000000000000000
000000000000000000000000000000000000000000000000000000000000
000000000000000000000000000000000000000000000000000000000000

000000000000000000000000000000000000000000000000000000111
>DM_train147
ANFNVPKLGVFPVAAVFDIDNVPEDSSATGSRWLPSIYQGGNYWGGGPQALHAQVSNFDSSN
RLPYNPRTENNPAGNCAFAFNPFGQYISNISSAQSVHRRIYGIDLNDEPLFSPNAASITNGG
NPTMSQDTGYHNIGPINTAYKAEIFRPVNPLPMSDTAPDPETLEPGQTEPLIKSDGVYSNSG
IASFIFDRPVTEPNPNWPPLPPPVIPIIYPTPALGIGAAAAYGFGYQVTVYRWEEIPVEFIA
DPETCPAQPTTDKVIIRTTDLNPEGSPCAYEAGIILVRQTSNPMNAVAGRLVPYVEDIAVDI
FLTGKFFTLNPPLRITNNYFADDEVKENTVTIGNYTTTLSSAYYAVYKTDGYGGATCFIASG
GAGISALVQLQDNSVLDVLYYSLPLSLGGSKAAIDEWVANNCGLFPMSGGLDKTTLLEIPRR
QLEAINPQDGPGQYDLFILDDSGAYASFSSFIGYPEAAYYVAGAATFMDVENPDEIIFILRN
GAGWYACEIGDALKIADDEFDSVDYFAYRGGVMFIGSARYTEGGDPLPIKYRAIIPGLP
000000000000000000000000000000000000000000000000000000000000
000000000000000000000000000000000000000000000000000000000000
000000000000000000000000000000000000000000000000000000000000
000000000000000000000000000000000000000000000000000000000011
111111111111111111110000000000000000000000000000000000000000
000000000000000000000000000000000000000000000000000000000000
000000000000000000000000000000000000000000000000000000000000
000000000000000000000000000000000000000000000000000000000000
000000000000000000000000000000000000000000000000000000000
>DM_train148
MNTSLFKQERQKYIPKLPNILKKDFNNISLVYGENTEAIQDRQALKEFFKNTYGLPIISFTE
GESSLSFSKALNIGIILSGGPAPGGHNVISGVFDAIKKFNPNSKLFGFKGGPLGLLENDKIE
LTESLINSYRNTGGFDIVSSGRTKIETEEHYNKALFVAKENNLNAIIIIGGDDSNTNAAILA
EYFKKNGENIQVIGVPKTIDADLRNDHIEISFGFDSATKIYSELIGNLCRDAMSTKKYWHFV
KLMGRSASHVALECALKTHPNICIVSEEVLAKKKTLSEIIDEMVSVILKRSLNGDNFGVVIV
PEGLIEFIPEVKSLMLELCDIFDKNEGEFKGLNIEKMKEIFVAKLSDYMKGVYLSLPLFIQF
ELIKSILERDPHGNFNVSRVPTEKLFIEMIQSRLNDMKKRGEYKGSFTPVDHFFGYEGRSAF
PSNFDSDYCYSLGYNAVVLILNGLTGYMSCIKNLNLKPTDWIAGGVPLTMLMNMEERYGEKK
PVIKKALVDLEGRPFKEFVKNRDKWALNNLYLYPGPVQYFGSSEIVDEITETLKLELFK
000000000000000000000000000000000000000000000000000000000000
000000000000000000000000000000000000000000000000000000000000
000000000000000000000000000000000000000000000000000000000000
000000000000000000000000000000000000000000000000000000000000
000000000000000000000000000000000000000000000000000000000000
000000000000000000000011111111111000000000000000000000000000
000000001111111111000000000000000000000000000000000000000000
000000000000000000000000000000000000000000000000000000000000
000000000000000000000000000000000000000000000000000000000
>DM_train149
GSHMPVVHVIDVESGNLQSLTNAIEHLGYEVQLVKSPKDFNISGTSRLILPGVGNYGHFVDN
LFNRGFEKPIREYIESGKPIMGICVGLQALFAGSVESPKSTGLNYIDFKLSRFDDSEKPVPE
IGWNSCIPSENLFFGLDPYKRYYFVHSFAAILNSEKKKNLENDGWKIAKAKYGSEEFIAAVN
KNNIFATQFHPEKSGKAGLNVIENFLKQQSPPIPNYSAEEKELLMNDYSNYGLTRRIIACLD

VRTNDQGDLVVTKGDQYDVREKSDGKGVRNLGKPVQLAQKYYQQGADEVTFLNITSFRDCPL
KDTPMLEVLKQAAKTVFVPLTVGGGIKDIVDVDGTKIPALEVASLYFRSGADKVSIGTDAVY
AAEKYYELGNRGDGTSPIETISKAYGAQAVVISVDPKRVYVNSQADTKNKVFETEYPGPNGE
KYCWYQCTIKGGRESRDLGVWELTRACEALGAGEILLNCIDKDGSNSGYDLELIEHVKDAVK
IPVIASSGAGVPEHFEEAFLKTRADACLGAGMFHRGEFTVNDVKEYLLEHGLKVRMDEE
00000000000000000000000000000000000000000000000000000000000000
00000000000000000000000000000000000000000000000000000000000000
00000000000000000000000000000000000000000000000000000000000000
00000000000000000000000000000000000000000000000000000000000000
00000000000111111111111111111100000000000000000000000001110000
00000000000000000000000000000000000000000000000000000000000000
00000000000000000000000000000000000000000000000000000000000000
00000000000000000000000000000000000000000000000000000000000000
00000000000000000000000000000000000000000000000000000000000
>DM_train151
MGDYDIPTTENLYFQGMTDLNKLVKELNDLGLTDVKEIVYNPSYEQLFEEETKPGLEGFDKG
TLTTLGAVAVDTGIFTGRSPKDKYIVCDETTKDTVWWNSEAAKNDNKPMTQETWKSLRELVA
KQLSGKRLFVVEGYCGASEKHRIGVRMVTEVAWQAHFVKNMFIRPTDEELKNFKADFTVLNG
AKCTNPNWKEQGLNSENFVAFNITEGIQLIGGTWYGGEMKKGMFSMMNYFLPLKGVASMHCS
ANVGKDGDVAIFFGLSGTGKTTLSTDPKRQLIGDDEHGWDESGVFNFEGGCYAKTINLSQEN
EPDIYGAIRRDALLENVVVRADGSVDFDDGSKTENTRVSYPIYHIDNIVRPVSKAGHATKVI
FLTADAFGVLPPVSKLTPEQTEYYFLSGFTAKLAGTERGVTEPTPTFSACFGAAFLSLHPIQ
YADVLVERMKASGAEAYLVNTGWNGTGKRISIKDTRGIIDAILDGSIEKAEMGELPIFNLAI
PKALPGVDPAILDPRDTYADKAQWQVKAEDLANRFVKNFVKYTANPEAAKLVGAGPKA
11111111111111111100000000000000000000000000000000000000000000
00000000000000000000000000000000000000000000000000000000000000
00000000000000000000000000000000000000000000000000000000000000
00000000000000000000000000000000000000000000000000000000000000
00000000000000000000000000000000000000000000000000000000000000
00000000000000000000000000000000000000000000000000000000000000
00000000000000000000000000000001111111111100000000000000000000
00000000000000000000000000000000000000000000000000000000000000
0000000000000000000000000000000000000000000000000000000001
>DM_train152
MSALLLKPHKDLPRRTVLIVVMDGLGIGPEDDYDAVHMASTPFMDAHRRDNRHFRCVRAHGT
AVGLPTDADMGNSEVGHNALGAGRVALQGASLVDDAIKSGEIYTGEGYRYLHGAFSKEGSTL
HLIGLLSDGGVHSRDNQIYSIIEHAVKDGAKRIRVHALYDGRDVPDGSSFRFTDELEAVLAK
VRQNGCDAAIASGGGRMFVTMDRYDADWSIVERGWRAQVLGDARHFHSAKEAITTFREEDPK
VTDQYYPPFIVVDEQDKPLGTIEDGDAVLCVNFRGDRVIEMTRAFEDEDFNKFDRVRVPKVR
YAGMMRYDGDLGIPNNFLVPPPKLTRVSEEYLCGSGLNIFACSETQKFGHVTYFWNGNRSGK
IDEKHETFKEVPSDRVQFNEKPRMQSAAITEAAIEALKSGMYNVVRINFPNGDMVGHTGDLK
ATITGVEAVDESLAKLKDAVDSVNGVYIVTADHGNSDDMAQRDKKGKPMKDGNGNVLPLTSH
TLSPVPVFIGGAGLDPRVAMRTDLPAAGLANVTATFINLLGFEAPEDYEPSLIYVEK
11100000000000000000000000000000000000000000000000000000000000

00000000000000000000000000000000000000000000000000000000000000
00000000000000000000000000000000000000000000000000000000000000
00000000000000000000000000000000000000000000000000000000000000
00000000000000000000000000000000000000000000000000000000000000
00000000000000000000000000000000000000000000000000000000000000
00000000000000000000000000000000000000000000000000000000000000
00000000000000000000000000000000000000000000000000000000000000
00000000000000000000000000000000000000000000000000000001
>DM_train153
MSDMEKPWKEEEKREVLAGHARRQAPQAVDKGPVTGDQRISVTVVLRRQRGDELEAHVERQA
ALAPHARVHLEREAFAASHGASLDDFAEIRKFAEAHGLTLDRAHVAAGTAVLSGPVDAVNQA
FGVELRHFDHPDGSYRSYVGDVRVPASIAPLIEAVLGLDTRPVARPHFRLRRRAEGEFEARS
QSAAPTAYTPLDVAQAYQFPEGLDGQGQCIAIIELGGGYDETSLAQYFASLGVSAPQVVSVS
VDGATNQPTGDPNGPDGEVELDIEVAGALAPGAKIAVYFAPNTDAGFLNAITTAVHDPTHKP
SIVSISWGGPEDSWAPASIAAMNRAFLDAAALGVTVLAAAGDSGSTDGEQDGLYHVDFPAAS
PYVLACGGTRLVASAGRIERETVWNDGPDGGSTGGGVSRIFPLPSWQERANVPPSANPGAGS
GRGVPDVAGNADPATGYEVVIDGETTVIGGTAAVAPLFAALVARINQKLGKPVGYLNPTLYQ
LPPEVFHDITEGNNDIANRARIYQAGPGWDPCTGLGSPIGIRLLQALLPSASQAQP
11111111111000000000000000000000000000000000000000000000000000
00000000000000000000000000000000000000000000000000000000000000
00000000000000000000000000000000000000000000000000000000000000
00000000000000000000000000000000000000000000000000000000000000
00000000000000000000000000000000000000000000000000000000000000
00000000000000000000000000000000000000000000000000000000000000
00000000000000000000000000000000000000000000000000000000000000
00000000000000000000000000000000000000000000000000000000000000
00000000000000000000000000000000000000000000000001111111
>DM_train154
MKRREVLQAGMALGGLAGLGRALAQGGFTLTLVHTNDTHAHLEPVELTLSGEKTPVGGVARR
VALFDRVWARAKNPLFLDAGDVFQGTLYFNQYRGLADRYFMHRLRYRAMALGNHEFDLGPGP
LADFLKGARFKVVSANVDASREPRLKGLFAPYAVVVVGGERVGIIGLTTPDTREISNPGPTV
AFLDPYESAQKAVYELLAKGVNKIVVLSHLGYGEDLKLARRLVGVQVIVGGHSHTLLGSFPH
KELSPAGPYPTVVKNPEGKDVLVVQAWEWGKVVGLLEVTFDAKGELLAYKGEALLMTPEAAP
EDFFAKEALLAYAQPVMALMQQVIAEAKVDLVGERAVVRRRESNLGNLITDGMLWKTRNAGT
QIALQNGGGIRASIPKGPITVGKVYEVLPFGNTLVVMDLKGKEILAALENGVSQWENTAGRF
LQVSGLRYAFDLSRPAGSRVVRVEVKTEKGYVPLDLEATYRVVVNNFIANGGDGFTVLKEAQ
GYRVDTGFSDAESFMDYLKELKVVEAGLEGRIEVLNEPKGERPAYFAYRVPGLVGV
11111111111111111111111111100000000000000000000000000000000000
00000000000000000000000000000000000000000000000000000000000000
00000000000000000000000000000000000000000000000000000000000000
00000000000000000000000000000000000000000000000000000000000000
00000000000000000000000000000000000000000000000000000000000000
00000000000000000000000000000000000000000000000000000000000000
00000000000000000000000000000000000000000000000000000000000000

000000000000000000000000000000000000000000000000000000000000
0000000000000000000000000000000000000011111111111111111
>DM_train155
MTQVAKKILVTCALPYANGSIHLGHMLEHIQADVWVRYQRMRGHEVNFICADDAHGTPIMLK
AQQLGITPEQMIGEMSQEHQTDFAGFNISYDNYHSTHSEENRQLSELIYSRLKENGFIKNRT
ISQLYDPEKGMFLPDRFVKGTCPKCKSPDQYGDNCEVCGATYSPTELIEPKSVVSGATPVMR
DSEHFFFDLPSFSEMLQAWTRSGALQEQVANKMQEWFESGLQQWDISRDAPYFGFEIPNAPG
KYFYVWLDAPIGYMGSFKNLCDKRGDSVSFDEYWKKDSTAELYHFIGKDIVYFHSLFWPAML
EGSNFRKPSNLFVHGYVTVNGAKMSKSRGTFIKASTWLNHFDADSLRYYYTAKLSSRIDDID
LNLEDFVQRVNADIVNKVVNLASRNAGFINKRFDGVLASELADPQLYKTFTDAAEVIGEAWE
SREFGKAVREIMALADLANRYVDEQAPWVVAKQEGRDADLQAICSMGINLFRVLMTYLKPVL
PKLTERAEAFLNTELTWDGIQQPLLGHKVNPFKALYNRIDMRQVEALVEASKEEV
111100000000000000000000000000000000000000000000000000000000
000000000000000000000000000000000000000000000000000000000000
000000000000000000000000000000000000000000000000000000000000
000000000000000000000000000000000000000000000000000000000000
000000000000000000000000000000000000000000000000000000000000
000000000000000000000000000000000000000000000000000000000000
000000000000000000000000000000000000000000000000000000000000
000000000000000000000000000000000000000000000000000000000000
00000000000000000000000000000000000000000000000000000000
>DM_train158
PTVSATPYDYIIVGAGPGGIIAADRLSEAGKKVLLLERGGPSTKQTGGTYVAPWATSSGLTK
FDIPGLFESLFTDSNPFWWCKDITVFAGCLVGGGTSVNGALYWYPNDGDFSSSVGWPSSWTN
HAPYTSKLSSRLPSTDHPSTDGQRYLEQSFNVVSQLLKGQGYNQATINDNPNYKDHVFGYSA
FDFLNGKRAGPVATYLQTALARPNFTFKTNVMVSNVVRNGSQILGVQTNDPTLGPNGFIPVT
PKGRVILSAGAFGTSRILFQSGIGPTDMIQTVQSNPTAAAALPPQNQWINLPVGMNAQDNPS
INLVFTHPSIDAYENWADVWSNPRPADAAQYLANQSGVFAGASPKLNFWRAYSGSDGFTRYA
QGTVRPGAASVNSSLPYNASQIFTITVYLSTGIQSRGRIGIDAALRGTVLTPPWLVNPVDKT
VLLQALHDVVSNIGSIPGLTMITPDVTQTLEEYVDAYDPATMNSNHWVSSTTIGSSPQSAVV
DSNVKVFGTNNLFIVDAGIIPHLPTGNPQGTLMSAAEQAAAKILALAGGP
111110000000000000000000000000000000000000000000000000000000
000000000000000000000000000000000000000000000000000000000000
000000000000000000000000000000000000000000000000000000000000
000000000000000000000000000000000000000000000000000000000000
000000000000000000000000000000000000000000000000000000000000
000000000000000000000000000000000000000000000000000000000000
000000000000000000000000000000000000000000000000000000000000
000000000000000000000000000000000000000000000000000000000000
00000000000000000000000000000000000000000000000000
>DM_train159
MHIDNIENLSDREFDYIVVGGGSAGAAVAARLSEDPAVSVALVEAGPDDRGVPEVLQLDRWM
ELLESGYDWDYPIEPQENGNSFMRHARAKVMGGCSSHNSCIAFWAPREDLDEWEAKYGATGW
NAEAAWPLYKRLETNEDAGPDAPHHGDSGPVHLMNVPPKDPTGVALLDACEQAGIPRAKFNT

GTTVVNGANFFQINRRADGTRSSSSVSYIHPIVEQENFTLLTGLRARQLVFDADRRCTGVDI
VDSAFGHTHRLTARNEVVLSTGAIDTPKLLMLSGIGPAAHLAEHGIEVLVDSPGVGEHLQDH
PEGVVQFEAKQPMVAESTQWWEIGIFTPTEDGLDRPDLMMHYGSVPFDMNTLRHGYPTTENG
FSLTPNVTHARSRGTVRLRSRDFRDKPMVDPRYFTDPEGHDMRVMVAGIRKAREIAAQPAMA
EWTGRELSPGVEAQTDEELQDYIRKTHNTVYHPVGTVRMGAVEDEMSPLDPELRVKGVTGLR
VADASVMPEHVTVNPNITVMMIGERCADLIRSARAGETTTADAELSAALA
00000000000000000000000000000000000000000000000000000000000000
00000000000000000000000000000000000000000000000000000000000000
00000000000000000000000000000000000000000000000000000000000000
00000000000000000000000000000000000000000000000000000000000000
00000000000000000000000000000000000000000000000000000000000000
00000000000000000000000000000000000000000000000000000000000000
00000000000000000000000000000000000000000000000000000000000000
00000000000000000000000000000000000000000000000000000000000000
00000000000000000000000000000001111111111111111111
>DM_train160
ASFPHRNLTWNDINFVHTTDTHGWYSGHINQPLYHANWGDFISFTTHMRRIAHSRNQDLLLI
DSGDRHDGNGLSDITSPNGLKSTPIFIKQDYDLLTIGNHELYLWENSKQEYETVVNHFQDKY
VCSNVDIRLDNGLFVPLGLKYKYFTTPIRGIRVMAFGFLFDFKRFNSGTRVTPMAETIHEPW
FQEALKHEVDLIIIVGHTPISHNWGEFYQVHQYLRQFFPDTIIQYFGGHSHIRDFTVFDSLS
TGLQSGRYCETVGWTSVNLDKADLNLPVRQRFSRSYIDFNTDSFKYHTNLDKEFDTAKGKLV
SKLIRETRKELKLDTLIGYVKTNYYVDYVPIDHPKSIFNLLALKILKTLPKSKHEERITIIN
TGSIRYDLYKGPYTIDSKFIVSPFENIWVNITVPKSVATKVAAKLNDADYISASRLKPPHQY
DLQVQDLSTSPHQAHFEMQEKLPKGYVTHDDFGADGDDTLHRAVVNFPVPNVIQSVEINDEV
DSPVNLVFYSFITPNIIWALKELNFSTEQVPTPYSDIYLGTLLNEFVANN
10000000000000000000000000000000000000000000000000000000000000
00000000000000000000000000000000000000000000000000000000000000
00000000000000000000000000000000000000000000000000000000000000
00000000000000000000000000000000000000000000000000000000000000
00000000000000000000000000000000000000000000000000000000000000
00000000000000000000000000000000000000000000000000000000000000
00000000000000000000000000000000000000000000000110000000000000
00000111111111111100000000000000000000000000000000000000000000
00000000000000000000000000000000000000000000000000
>DM_train161
MAAKEVKFNSDARDRMLKGVNILADAVKVTLGPKGRNVVIDKSFGAPRITKDGVSVAKEIEL
SDKFENMGAQMVREVASRTNDEAGDGTTTATVLAQAIVREGLKAVAAGMNPMDLKRGIDVAT
AKVVEAIKSAARPVNDSSEVAQVGTISANGESFIGQQIAEAMQRVGNEGVITVEENKGMETE
VEVVEGMQFDRGYLSPYFVTNADKMIAELEDAYILLHEKKLSSLQPMVPLLESVIQSQKPLL
IVAEDVEGEALATLVVNKLRGGLKIAAVKAPGFGDRRKAMLQDIAILTGGQVISEDLGMKLE
NVTIDMLGRAKKVSINKDNTTIVDGAGEKAEIEARVSQIRQQIEETTSDYDREKLQERVAKL
AGGVAVIRVGGMTEIEVKERKDRVDDALNATRAAVQEGIVVGGGVALVQGAKVLEGLSGANS
DQDAGIAIIRRALEAPMRQIAENAGVDGAVVAGKVRESSDKAFGFNAQTEEYGDMFKFGVID
PAKVVRTALEDAASVAGLLITTEAMIAEKPEPKAPAGGMPDMGGMGGMM

000000000000000000000000000000000000000000000000000000000000
000000000000000000000000000000000000000000000000000000000000
000000000000000000000000000000000000000000000000000000000000
000000000000000000000000000000000000000000001111111111100000
000000011111111111111111000000000000000000000000111111111111
111000000000000000000000000000000000000000000000000000000000
000000000000000000000000000000000000000000000000000000000000
000000000000000000000000000000000000000000000000000000000000
000000000000000000000000000001111111111111111111
>DM_train163
ASYFIGVDVGTGSARAGVFDLQGRMVGQASREITMFKPKADFVEQSSENIWQAVCNAVRDAV
NQADINPIQVKGLGFDATCSLVVLDKEGNPLTVSPSGRNEQNVIVWMDHRAITQAERINATK
HPVLEFVGGVISPEMQTPKLLWLKQHMPNTWSNVGHLFDLPDFLTWRATKDETRSLCSTVCK
WTYLGHEDRWDPSYFKLVGLADLLDNNAAKIGATVKPMGAPLGHGLSQRAASEMGLIPGTAV
SVSIIDAHAGTIGILGASGVTGENANFDRRIALIGGTSTAHMAMSRSAHFISGIWGPYYSAI
LPEYWLNEGGQSATGALIDHIIQSHPCYPALLEQAKNKGETIYEALNYILRQMAGEPENIAF
LTNDIHMLPYFHGNRSPRANPNLTGIITGLKLSTTPEDMALRYLATIQALALGTRHIIETMN
QNGYNIDTMMASGGGTKNPIFVQEHANATGCAMLLPEESEAMLLGSAMMGTVAAGVFESLPE
AMAAMSRIGKTVTPQTNKIKAYYDRKYRVFHQMYHDHMRYQALMQEG
000000000000000000000000000000000000000000000000000000000000
000000000000000000000000000000000000000000000000000000000000
000000000000000000000000000000000000000000000000000000000000
000000000000000000000000000000000000000000000000000000000000
000000000000000000000000000000000000000000000000000000000000
000000000000000000000000000000000000000000000000000000000000
000000000000000000000000000000000000000000000000000000000000
000000000000000000000000000000000000000000000000000000000000
0000000000000000000000000000000000000000000011
>DM_train164
MASWSHPQFEKGAETAVPNSSSVPGDPSSMRKKNVLLIVVDQWRADFVPHVLRADGKIDFLK
TPNLDRLCREGVTFRNHVTTCVPAGPARASLLTGLYLMNHRAVQNTVPLDQRHLNLGKALRG
VGYDPALIGYTTTVPDPRTTSPNDPRFRVLGDLMDGFHPVGAFEPNMEGYFGWVAQNGFDLP
EHRPDIWLPEGEDAVAGATDRPSRIPKEFSDSTFFTERALTYLKGRDGKPFFLHLGYYRPHP
PFVASAPYHAMYRPEDMPAPIRAANPDIEAAQHPLMKFYVDSIRRGSFFQGAEGSGATLDEA
ELRQMRATYCGLITEVDDCLGRVFSYLDETGQWDDTLIIFTSDHGEQLGDHHLLGKIGYNDP
SFRIPLVIKDAGENARAGAIESGFTESIDVMPTILDWLGGKIPHACDGLSLLPFLSEGRPQD
WRTELHYEYDFRDVYYSEPQSFLGLGMNDCSLCVIQDERYKYVHFAALPPLFFDLRHDPNEF
TNLADDPAYAALVRDYAQKALSWRLKHADRTLTHYRSGPEGLSERSH
111111111111111111111111111110000000000000000000000000000000
000000000000000000000000000000000000000000000000000000000000
000000000000000000000000000000000000000000000000000000000000
000000000000000000000000000000000000000000000000000000000000
000000000000000000000000000000000000000000000000000000000000
000000000000000000000000000000000000000000000000000000000000

000000000000000000000000000000000000000000000000000000000000
000000000000000000000000000000000000000000000000000000000000
00000000000000000000000000000000000000000000000
>DM_train165
MAGQTTVDSRRQPPEEVDVLVVGAGFSGLYALYRLRELGRSVHVIETAGDVGGVWYWNRYPG
ARCDIESIEYCYSFSEEVLQEWNWTERYASQPEILRYINFVADKFDLRSGITFHTTVTAAAF
DEATNTWTVDTNHGDRIRARYLIMASGQLSVPQLPNFPGLKDFAGNLYHTGNWPHEPVDFSG
QRVGVIGTGSSGIQVSPQIAKQAAELFVFQRTPHFAVPARNAPLDPEFLADLKKRYAEFREE
SRNTPGGTHRYQGPKSALEVSDEELVETLERYWQEGGPDILAAYRDILRDRDANERVAEFIR
NKIRNTVRDPEVAERLVPKGYPFGTKRLILEIDYYEMFNRDNVHLVDTLSAPIETITPRGVR
TSEREYELDSLVLATGFDALTGALFKIDIRGVGNVALKEKWAAGPRTYLGLSTAGFPNLFFI
AGPGSPSALSNMLVSIEQHVEWVTDHIAYMFKNGLTRSEAVLEKEDEWVEHVNEIADETLYP
MTASWYTGANVPGKPRVFMLYVGGFHRYRQICDEVAAKGYEGFVLT
111111111000000000000000000000000000000000000000000000000000
000000000000000000000000000000000000000000000000000000000000
000000000000000000000000000000000000000000000000000000000000
000000000000000000000000000000000000000000000000000000000000
000000000000000000000000000000000000000000000000000000000000
000000000000000000000000000000000000000000000000000000000000
000000000000000000000000000000000000000000000000000000000000
000000000000000000000000000000000000000000000000000000000000
0000000000000000000000000000000000000000000000
>DM_train166
ALEEAPWPPPEGAFVGFVLSRKEPMWADLLALAAARGGRVHRAPEPYKALRDLKEARGLLAK
DLSVLALREGLGLPPGDDPMLLAYLLDPSNTTPEGVARRYGGEWTEEAGERAALSERLFANL
WGRLEGEERLLWLYREVERPLSAVLAHMEATGVRLDVAYLRALSLEVAEEIARLEAEVFRLA
GHPFNLNSRDQLERVLFDELGLPAIGKTEKTGKRSTSAAVLEALREAHPIVEKILQYRELTK
LKSTYIDPLPDLIHPRTGRLHTRFNQTATATGRLSSSDPNLQNIPVRTPLGQRIRRAFIAEE
GWLLVALDYSQIELRVLAHLSGDENLIRVFQEGRDIHTETASWMFGVPREAVDPLMRRAAKT
INFGVLYGMSAHRLSQELAIPYEEAQAFIERYFQSFPKVRAWIEKTLEEGRRRGYVETLFGR
RRYVPDLEARVKSVREAAERMAFNMPVQGTAADLMKLAMVKLFPRLEEMGARMLLQVHDELV
LEAPKERAEAVARLAKEVMEGVYPLAVPLEVEVGIGEDWLSAKE
000000000000000000000000000000000000000000000000000000000000
000000000000000000000000000000000000000000000000000000000000
000000000000000000000000000000000000000000000000000000000000
000000000000000000000011111111111111100000000000000000000000
000000000000000000000000000000000000000000000000000000000000
000000000000000000000000000000000000000000000000000000000000
000000000000000000000000000000000000000000000000000000000000
000000000000000000000000000000000000000000000000000000000000
00000000000000000000000000000000000000000000
>DM_train168
MNKPIKNIVIVGGGTAGWMAASYLVRALQQQANITLIESAAIPRIGVGEATIPSLQKVFFDF
LGIPEREWMPQVNGAFKAAIKFVNWRKSPDPSRDDHFYHLFGNVPNCDGVPLTHYWLRKREQ

GFQQPMEYACYPQPGALDGKLAPCLSDGTRQMSHAWHFDAHLVADFLKRWAVERGVNRVVDE
VVDVRLNNRGYISNLLTKEGRTLEADLFIDCSGMRGLLINQALKEPFIDMSDYLLCDSAVAS
AVPNDDARDGVEPYTSSIAMNSGWTWKIPMLGRFGSGYVFSSHFTSRDQATADFLKLWGLSD
NQPLNQIKFRVGRNKRAWVNNCVSIGLSSCFLEPLESTGIYFIYAALYQLVKHFPDTSFDPR
LSDAFNAEIVHMFDDCRDFVQAHYFTTSRDDTPFWLANRHDLRLSDAIKEKVQRYKAGLPLT
TTSFDDSTYYETFDYEFKNFWLNGNYYCIFAGLGMLPDRSLPLLQHRPESIEKAEAMFASIR
REAERLRTSLPTNYDYLRSLRDGDAGLSRGQRGPKLAAQESL
100000000000000000000000000000000000000000000000000000000000
000000000000000000000000000000000000000000000000000000000000
000000000000000000000000000000000000000000000000000000000000
000000000000000000000000000000000000000000000000000000000000
000000000000000000000000000000000000000000000000000000000000
000000000000000000000000000000000000000000000000000000000000
000000000000000000000000000000000000000000000000000000000000
000000000000000000000000000000000000000000000000000000000000
000000000000000000000011111111111111111111
>DM_train169
MSSQKVFGITGPVSTVGATAAENKLNDSLIQELKKEGSFETEQETANRVQVLKILQELAQRF
VYEVSKKKNMSDGMARDAGGKIFTYGSYRLGVHGPGSDIDTLVVVPKHVTREDFFTVFDSLL
RERKELDEIAPVPDAFVPIIKIKFSGISIDLICARLDQPQVPLSLTLSDKNLLRNLDEKDLR
ALNGTRVTDEILELVPKPNVFRIALRAIKLWAQRRAVYANIFGFPGGVAWAMLVARICQLYP
NACSAVILNRFFIILSEWNWPQPVILKPIEDGPLQVRVWNPKIYAQDRSHRMPVITPAYPSM
CATHNITESTKKVILQEFVRGVQITNDIFSNKKSWANLFEKNDFFFRYKFYLEITAYTRGSD
EQHLKWSGLVESKVRLLVMKLEVLAGIKIAHPFTKPFESSYCCPTEDDYEMIQDKYGSHKTE
TALNALKLVTDENKEEESIKDAPKAYLSTMYIGLDFNIENKKEKVDIHIPCTEFVNLCRSFN
EDYGDHKVFNLALRFVKGYDLPDEVFDENEKRPSKKSKRKN
000000000000000000000000000000000000000000000000000000000000
000000000000000000000000000000000000000000000000000000000000
000000000000000000000000000000000000000000000000000000000000
000000000000000000000000000000000000000000000000000000000000
000000000000000000000000000000000000000000000000000000000000
000000000000000000000000000000000000000000000000000000000000
000000000000000000000000000000000000000000000000000001111111
111110000000000000000000000000000000000000000000000000000000
00000000000000000000000000011111111111111
>DM_train170
MPSRVPKSIFYNQVGYLISGDKRFWIQAHEPQPFALRTPEGQAVFAGMTKPVGGNWYVGDFT
ALRVPGTYTLTVGTLEARVVIHRRAYRDVLEAMLRFFDYQLCGVVLPEDEAGPWAHGACHTS
DAKVFGTERALACPGGWHDAGDYGKYTVPAAKAVADLLLAHEYFPAALAHVRPMRSVHRAPH
LPPALEVAREEIAWLLTMQDPATGGVYHKVTTPSFPPLDTRPEDDDAPLVLSPISYAATATF
CAAMAHAALVYRPFDPALSSCCADAARRAYAWLGAHEMQPFHNPDGILTGEYGDAELRDELL
WASCALLRMTGDSAWARVCEPLLDLDLPWELGWADVALYGVMDYLRTPRAAVSDDVRNKVKS
RLLRELDALAAMAESHPFGIPMRDDDFIWGSNMVLLNRAMAFLLAEGVGVLHPAAHTVAQRA
ADYLFGANPLGQCYVTGFGQRPVRHPHHRPSVADDVDHPVPGMVVGGPNRHLQDEIARAQLA

GRPAMEAYIDHQDSYSTNEVAVYWNSPAVFVIAALLEARGR
111111000000000000000000000000000000000000000000000000000000
000000000000000000000000000000000000000000000000000000000000
000000000000000000000000000000000000000000000000000000000000
000000000000000000000000000000000000000000000000000000000000
000000000000000000000000000000000000000000000000000000000000
000000000000000000000000000000000000000000000000000000000000
000000000000000000000000000000000000000000000000000000000000
000000000000000000000000000000000000000000000000000000000000
00000000000000000000000000000000000000000
>DM_train171
GADDVVDSSKSFVMENFSSYHGTKPGYVDSIQKGIQKPKSGTQGNYDDDWKGFYSTDNKYDA
AGYSVDNENPLSGKAGGVVKVTYPGLTKVLALKVDNAETIKKELGLSLTEPLMEQVGTEEFI
KRFGDGASRVVLSLPFAEGSSSVEYINNWEQAKALSVELEINFETRGKRGQDAMYEYMAQAC
AGNRVRRSVGSSLSCINLDWDVIRDKTKTKIESLKEHGPIKNKMSESPNKTVSEEKAKQYLE
EFHQTALEHPELSELKTVTGTNPVFAGANYAAWAVNVAQVIDSETADNLEKTTAALSILPGI
GSVMGIADGAVHHNTEEIVAQSIALSSLMVAQAIPLVGELVDIGFAAYNFVESIINLFQVVH
NSYNRPAYSPGHKTQPFLHDGYAVSWNTVEDSIIRTGFQGESGHDIKITAENTPLPIAGVLL
PTIPGKLDVNKSKTHISVNGRKIRMRCRAIDGDVTFCRPKSPVYVGNGVHANLHVAFHRSSS
EKIHSNEISSDSIGVLGYQKTVDHTKVNSKLSLFFEIKS
000000000000000000000000000000000000111111111000000000000000
000000000000000000000000000000000000000000000000000000000000
000000000000000000000000000000000000000000000000000000000000
011111111111100000000000000000000000000000000000000000000000
000000000000000000000000000000000000000000000000000000000000
000000000000000000000000000000000000000000000000000000000000
000000000000000000000000000000000000000000000000000000000000
000000000000000000000000000000000000000000000000000000000000
000000000000000000000000000000000000000
>DM_train172
EETPVTPQPPDILLGPLFNDVQNAKLFPDQKTFADAVPNSDPLMILADYRMQQNQSGFDLRH
FVNVNFTLPKEGEKYVPPEGQSLREHIDGLWPVLTRSTENTEKWDSLLPLPEPYVVPGGRFR
EVYYWDSYFTMLGLAESGHWDKVADMVANFAHEIDTYGHIPNGNRSYYLSRSQPPFFALMVE
LLAQHEGDAALKQYLPQMQKEYAYWMDGVENLQAGQQEKRVVKLQDGTLLNRYWDDRDTPRP
ESWVEDIATAKSNPNRPATEIYRDLRSAAASGWDFSSRWMDNPQQLNTLRTTSIVPVDLNSL
MFKMEKILARASKAAGDNAMANQYETLANARQKGIEKYLWNDQQGWYADYDLKSHKVRNQLT
AAALFPLYVNAAAKDRANKMATATKTHLLQPGGLNTTSVKSGQQWDAPNGWAPLQWVATEGL
QNYGQKEVAMDISWHFLTNVQHTYDREKKLVEKYDVSTTGTGGGGGEYPLQDGFGWTNGVTL
KMLDLICPKEQPCDNVPATRPTVKSATTQPSTKEAQPTP
111111000000000000000000000000000000000000000000000000000000
000000000011110000000000000000000000000000000000000000000000
000000000000000000000000000000000000000000000000000000000000
000000000000000000000000000000000000000000000000000000000000
000000000000000000000000000000000000000000000000000000000000

000000000000000000000000000000000000000000000000000000000000
000000000000000000000000000000000000000000000000000000000000
000000000000000000000000000000000000000000000000000000000000
000000000000000000000111111111111111111
>DM_train174
MAQLGKLLKEQKYDRQLRLWGDHGQEALESAHVCLINATATGTEILKNLVLPGIGSFTIIDG
NQVSGEDAGNNFFLQRSSIGKNRAEAAMEFLQELNSDVSGSFVEESPENLLDNDPSFFCRFT
VVVATQLPESTSLRLADVLWNSQIPLLICRTYGLVGYMRIIIKEHPVIESHPDNALEDLRLD
KPFPELREHFQSYDLDHMEKKDHSHTPWIVIIAKYLAQWYSETNGRIPKTYKEKEDFRDLIR
QGILKNENGAPEDEENFEEAIKNVNTALNTTQIPSSIEDIFNDDRCINITKQTPSFWILARA
LKEFVAKEGQGNLPVRGTIPDMIADSGKYIKLQNVYREKAKKDAAAVGNHVAKLLQSIGQAP
ESISEKELKLLCSNSAFLRVVRCRSLAEEYGLDTINKDEIISSMDNPDNEIVLYLMLRAVDR
FHKQQGRYPGVSNYQVEEDIGKLKSCLTGFLQEYGLSVMVKDDYVHEFCRYGAAEPHTIAAF
LGGAAAQEVIKIITKQFVIFNNTYIYSGMSQTSATFQL
111111110000000000000000000000000000000000000000000000000000
000000000000000000000000000000000000000000000000000000000000
000000000000000000000000000000000000000000000000000000000000
000000000000000000000000000000000000000000000000000000000000
000000000000000000000000000000000000000000000000000000000000
000000000000000000000000000000000000000000000000000000000000
000000000000000000000000000000000000000000000000000000000000
000000000000000000000000000000000000000000000000000000000000
0000000000000000000000000000000000000
>DM_train175
SHMQSRELKTVSADCKKEAIEKCAQWVVRDCRPFSAVSGSGFIDMIKFFIKVKAEYGEHVNV
EELLPSPITLSRKVTSDAKEKKALIGREIKSAVEKDGASATIDLWTDNYIKRNFLGVTLHYH
ENNELRDLILGLKSLDFERSTAENIYKKLKAIFSQFNVEDLSSIKFVTDRGANVVKSLANNI
RINCSSHLLSNVLENSFEETPELNMPILACKNIVKYFKKANLQHRLRSSLKSECPTRWNSTY
TMLRSILDNWESVIQILSEAGETQRIVHINKSIIQTMVNILDGFERIFKELQTCSSPSLCFV
VPSILKVKEICSPDVGDVADIAKLKVNIIKNVRIIWEENLSIWHYTAFFFYPPALHMQQEKV
AQIKEFCLSKMEDLELINRMSSFNELSATQLNQSDSNSHNSIDLTSHSKDISTTSFFFPQLT
QNNSREPPVCPSDEFEFYRKEIVILSEDFKVMEWWNLNSKKYPKLSKLALSLLSIPASSAAS
ERTFSLAGNIITEKRNRIGQQTVDSLLFLNSFYKNFCK
000000000000000000000000000000000000000000000000000000000000
000000000000000000000000000000000000000000000000000000000000
000000000000000000000000000000000000000000000000000000000000
000000000000000000000000000000000000000000000000000000000000
000000000000000000000000000000000000000000000000000000000000
000000000000000000000000000000000000000000000000000000000000
000000000000000000000000000000011111111111111110000000000000
000000000000000000000000000000000000000000000000000000000000
0000000000000000000000000000000000000
>DM_train177
MSSQVEHPAGGYKKLFETVEELSSPLTAHVTGRIPLWLTGSLLRCGPGLFEVGSEPFYHLFD

GQALLHKFDFKEGHVTYHRRFIRTDAYVRAMTEKRIVITEFGTCAFPDPCKNIFSRFFSYFR
GVEVTDNALVNIYPVGEDYYACTETNFITKVNPETLETIKQVDLCNYVSVNGATAHPHIEND
GTVYNIGNCFGKNFSIAYNIVKIPPLQADKEDPISKSEIVVQFPCSDRFKPSYVHSFGLTPN
YIVFVETPVKINLFKFLSSWSLWGANYMDCFESNETMGVWLHIADKKRKKYINNKYRTSPFN
LFHHINTYEDHEFLIVDLCCWKGFEFVYNYLYLANLRENWEEVKKNARKAPQPEVRRYVLPL
NIDKADTGKNLVTLPNTTATAILCSDETIWLEPEVLFSGPRQAFEFPQINYQKYGGKPYTYA
YGLGLNHFVPDRLCKLNVKTKETWVWQEPDSYPSEPIFVSHPDALEEDDGVVLSVVVSPGAG
QKPAYLLILNAKDLSEVARAEVEINIPVTFHGLFKKS
110000000000000000000000000000000000000000000000000000000000
000000000000000000000000000000000000000000001111111111111111
110000000000000000000000000000000000000000000000000000000000
000000000001110000000000000000000000000000000000000000000000
000000000000111111111111000000000000000000000000000000000000
000000000000000000000000000000000000000000000000000000000000
000000000000000000000000000000000000000000000000000000000000
000000000000000000000000000000000000000000000000000000000000
0000000000000000000000000000000000000
>DM_train179
MKPVKPPRINGRVPVLSAQEAVNYIPDEATLCVLGAGGGILEATTLITALADKYKQTQTPRN
LSIISPTGLGDRADRGISPLAQEGLVKWALCGHWGQSPRISDLAEQNKIIAYNYPQGVLTQT
LRAAAAHQPGIISDIGIGTFVDPRQQGGKLNEVTKEDLIKLVEFDNKEYLYYKAIAPDIAFI
RATTCDSEGYATFEDEVMYLDALVIAQAVHNNGGIVMMQVQKMVKKATLHPKSVRIPGYLVD
IVVVDPDQSQLYGGAPVNRFISGDFTLDDSTKLSLPLNQRKLVARRALFEMRKGAVGNVGVG
IADGIGLVAREEGCADDFILTVETGPIGGITSQGIAFGANVNTRAILDMTSQFDFYHGGGLD
VCYLSFAEVDQHGNVGVHKFNGKIMGTGGFIDISATSKKIIFCGTLTAGSLKTEIADGKLNI
VQEGRVKKFIRELPEITFSGKIALERGLDVRYITERAVFTLKEDGLHLIEIAPGVDLQKDIL
DKMDFTPVISPELKLMDERLFIDAAMGFVLPEAAH
111000000000000000000000000000000000000000000000000000000000
000000000000000000000000000000000000000000000000000000000000
000000000000000000000000000000000000000000000000000000000000
000000000000000000000000000000000000000000000000000000000000
000000000000000000000000000111111000000000000000000000000000
000000000000000000000000000000011111000000000000000000000000
000000000000000000000000000000000000000000000000000000000000
000000000000000000000000000000000000000000000000000000000000
00000000000000000000000000000000011
>DM_train181
MDNGTDSSTSKFVPEYRRTNFKNKGRFSADELRRRRDTQQVELRKAKRDEALAKRRNFIPPT
DGADSDEEDESSVSADQQFYSQLQQELPQMTQQLNSDDMQEQLSATVKFRQILSREHRPPID
VVIQAGVVPRLVEFMRENQPEMLQLEAAWALTNIASGTSAQTKVVVDADAVPLFIQLLYTGS
VEVKEQAIWALGNVAGDSTDYRDYVLQCNAMEPILGLFNSNKPSLIRTATWTLSNLCRGKKP
QPDWSVVSQALPTLAKLIYSMDTETLVDACWAISYLSDGPQEAIQAVIDVRIPKRLVELLSH
ESTLVQTPALRAVGNIVTGNDLQTQVVINAGVLPALRLLLSSPKENIKKEACWTISNITAGN
TEQIQAVIDANLIPPLVKLLEVAEYKTKKEACWAISNASSGGLQRPDIIRYLVSQGCIKPLC

DLLEIADNRIIEVTLDALENILKMGEADKEARGLNINENADFIEKAGGMEKIFNCQQNENDK
IYEKAYKIIETYFGEEEDAVDETMAPQNAGNTFG
11111111111000000001111111111100000000000000000000000000001111
11111111111111111111111100000000000000000000000000000000000000
00000000000000000000000000000000000000000000000000000000000000
00000000000000000000000000000000000000000000000000000000000000
00000000000000000000000000000000000000000000000000000000000000
00000000000000000000000000000000000000000000000000000000000000
00000000000000000000000000000000000000000000000000000000000000
00000000000000000000000000000000000000000000000000000000000000
0000000000000000001111111111111111
>DM_train183
STQTSRVTLVGEMLPAYNEILTPEALSFLKELHENFNERRIELLQKRMKKQQKIDAGEFPKF
LEETKRIREADWTIAKLPKDLEDRRVEITGPVDRKMVINALNSGAHLFMADFEDSNSPTWEN
AIEGQINLRDAVKGTISHKNENGKEYRLNSKTAVLIVRPRGWHLEEKHMQVDGKNMSGSLVD
FGLYFFHNAKALLEKGSGPYFYLPKMESYLEARLWNDVFVFAQKYIGIPNGTIKATVLLETI
HASFEMDEILYELKDHSAGLNCGRWDYIFSFLKAFRNHNEFLLPDRAQVTMTAPFMRAYSLK
VIQTCHRRNAPAIGGMAAQIPIKNNPEANEAAFEKVRADKEREALDGHDGTWVAHPGLVPVA
MEVFNHIMKTPNQIFRKREEIHVTEKDLLEVPVGTITEEGLRMNISVGIQYIASWLSGRGAA
PIYNLMEDAATAEISRAQVWQWIRHEGGKLNDGRNITLELMEELKEEELAKIEREIGKEAFK
KGRFQEATTLFTNLVRNDEFVPFLTLPGYEIL
11111000000001100000000000000000000000000000000000000000000000
00000000000000000000000000000000000000000000000000000000000000
00000000000000000000000000000000000000000000000000000000000000
00000000000000000000000000000000000000000000000000000000000000
00000000000000000000000000000000000000000000000000000000000000
00000000000000111111111111111111100000000000000000000000000000
00000000000000000000000000000000000000000000000000000000000000
00000000000000000000000000000000000000000000000000000000000010
00000000000000000000000000000000
>DM_train184
GCDLERYPLTDLSEETFWNSESNAELALTSLYRGSLTDGVEYNPSDWWSYHGMIMMEHLSDN
AFDRRGENNPFFKISSGNLTADNAFIKRYWETSYKRIGYCNRFLVGIQNSSESEKKTRMIAE
ARFLRATQYFYLASYFKNVPLVENVLTGEEANNVTKTSQADILKWCVTEFTAAAADLPRFSA
IPAGEAGRACKQAALAFLGRTCMLQKDWKSGAKAFHDIMELGDNAINANYQELFYPSTGTSN
KENIFYIQYLENYLGTGLPQHALSAKDGGWSLVNPAADLYESYEFKDGTPFSYDDPRYDPSN
LGKDRDPRLDYTIYYNGAIFMGTEYKMSPDYSAAKKEKLDYTSEASRTGFMMRKYFEESTPI
NDVQSANGLTPVIRYAEVLLGYLECLVEDNQTITQGILDETINAVRGRASVNMPPVTEVTPA
KLREIVRHERRIELAMEGIRYWDIMRWGIAHEVLSQKIWGAPYPGSTQYATTTKEVDPTGNY
RWYVGKRAFRNPTDYTWPIPQSEQNINPNLRD
11111111111111000000000000000000000000000000000000000000000000
00000000000000000000000000000000000000000000000000000000000000
00000000000000000000000000000000000000000000000000000000000000
00000000000000000000000000000000000000000000000000000000000000

00000000000000000000000000000000000000000000000000000000000000
00000000000000000000000000000000000000000000000000000000000000
00000000000000000000000000000000000000000000000000000000000000
00000000000000000000000000000000000000000000000000000000000000
00000000000000000000000000000001
>DM_train185
MADSRDPASDQMQHWKEQRAAQKADVLTTGAGNPVGDKLNVITVGPRGPLLVQDVVFTDEMA
HFDERIPERVVHAKGAGAFGYFEVTHDITKYSKAKVFEHIGKKTPIAVRFSTVAGESGSAD
TVRDPRGFAVKFYTEDGNWDLVGNNTPIFFIRDPILFPSFIHSQKRNPQTHLKDPDMVWDFW
SLRPESLHQVSFLFSDRGIPDGHRHMNGYGSHTFKLVNANGEAVYCKFHYKTDQGIKNLSVE
DAARLSQEDPDYGIRDLFNAIATGKYPSWTFYIQVMTFNQAETFPFNPFDLTKVWPHKDYPL
IPVGKLVLNRNPVNYFAEVEQIAFDPSNMPPGIEASPDKMLQGRLFAYPDTHRHRLGPNYLH
IPVNCPYRARVANYQRDGPMCMQDNQGGAPNYYPNSFGAPEQQPSALEHSIQYSGEVRRFNT
ANDDNVTQVRAFYVNVLNEEQRKRLCENIAGHLKDAQIFIQKKAVKNFTEVHPDYGSHIQAL
LDKYNAEKPKNAIHTFVQSGSHLAAREKANL
11111111111111111111111000000000000000000000000000000000000000
00000000000000000000000000000000000000000000000000000000000000
00000000000000000000000000000000000000000000000000000000000000
00000000000000000000000000000000000000000000000000000000000000
00000000000000000000000000000000000000000000000000000000000000
00000000000000000000000000000000000000000000000000000000000000
00000000000000000000000000000000000000000000000000000000000000
00000000000000000000000000000000000000000000000000000000000000
0000001111111111111111111111111
>DM_train187
SQIAVRTFHDIRAALLARRELALLDVREEDPFAQAHPLFAANLPLSRLELEIHARVPRRDTP
ITVYDDGEGLAPVAAQRLHDLGYSDVALLDGGLSGWRNAGGELFRDVNVPSKAFGELVEAER
HTPSLAAEEVQALLDARAEAVILDARRFDEYQTMSIPGGISVPGAELVLRVAELAPDPRTRV
IVNCAGRTRSIIGTQSLLNAGIPNPVAALRNGTIGWTLAGQQLEHGQTRRFGAISQDTRKAA
AQRARAVADRAGVERLDLAGLAQWQDEHDRTTYLLDVRTPEEYEAGHLPGSRSTPGGQLVQE
TDHVASVRGARLVLVDDDGVRANMSASWLAQMGWQVAVLDGLSEADFSERGAWSAPLPRQPR
ADTIDPTTLADWLGEPGTRVLDFTASANYAKRHIPGAAWVLRSQLKQALERLGTAERYVLTC
GSSLLARFAVAEVQALSGKPVFLLDGGTSAWVAAGLPTEDGESLLASPRIDRYRRPYEGTDN
PREAMQGYLDWEFGLVEQLGRDGTHGFFVI
11000000000000000000000000000000000000000000000000000000000000
00000000000000000000000000000000000000000000000000000000000000
00000000000000000000000000000000000000000000000000000000000000
00000000000000000000000000000000000000000000000000000000000000
00000000000000000000000000000000000000000000000000000000000000
00000000000000000000000000000000000000000000000000000000000000
00000000000000000000000000000000000000000000000000000000000000
00000000000000000000000000000000000000000000000000000000000000
000000000000000000000000000000
>DM_train188

MVAAPPAVGAAMPSLDFDTSVFNKEKVSLAGHEEYIVRGGRNLFPLLPEAFKGIKQIGVIGW
GSQGPAQAQNLRDSLAEAKSDIVVKIGLRKGSKSFDEARAAGFTEESGTLGDIWETVSGSDL
VLLLISDAAQADNYEKIFSHMKPNSILGLSHGFLLGHLQSAGLDFPKNISVIAVCPKGMGPS
VRRLYVQGKEINGAGINSSFAVHQDVDGRATDVALGWSVALGSPFTFATTLEQEYKSDIFGE
RGILLGAVHGIVEALFRRYTEQGMDEEMAYKNTVEGITGIISKTISKKGMLEVYNSLTEEGK
KEFNKAYSASFYPCMDILYECYEDVASGSEIRSVVLAGRRFYEKEGLPAFPMGNIDQTRMWK
VGEKVRSTRPENDLGPLHPFTAGVYVALMMAQIEVLRKKGHSYSEIINESVIESVDSLNPFM
HARGVAFMVDNCSTTARLGSRKWAPRFDYILTQQAFVTVDKDAPINQDLISNFMSDPVHGAI
EVCAELRPTVDISVPANADFVRPELRQSS
111111111111110000000000000000000000000000000000000000000000
000000000000000000000000000000000000000000000000000000000000
000000000000000000000000000000000000000000000000000000000000
000000000000000000000000000000000000000000000000000000000000
000000000000000000000000000000000000000000000000000000000000
000000000000000000000000000000000000000000000000000000000000
000000000000000000000000000000000000000000000000000000000000
000000000000000000000000000000000000000000000000000000000000
00000000111111111111111111111
>DM_train189
AVPAVGEDFPIDYADWLPKRDPNDRRRAGILLHPTSFPGPYGIGDLGPQAFKFLDWLHLAGC
SLWQVLPLVPPGKRGNEDGSPYSGQDANCGNTLLISLEELVDDGLLKMEELPEPLPTDRVNY
STISEIKDPLITKAAKRLLSSEGELKDQLENFRRDPNISSWLEDAAYFAAIDNSVNTISWYD
WPEPLKNRHLAALEEVYQSEKDFIDIFIAQQFLFQRQWKKVRDYARSKGISIMGDMPIYVGY
HSADVWANKKQFLLNRKGFPLIVSGVPPDAFSETGQLWGSPLYDWKAMEKDGFSWWVRRIQR
ATDLFDEFRIDHFRGFAGFWAVPSEEKIAILGRWKVGPGKPLFDAILQAVGKINIIAEDLGV
ITEDVVQLRKSIEAPGMAVLQFAFGSDAENPHLPHNHEQNQVVYTGTHDNDTIRGWWDTLPQ
EEKSNVLKYLSNIEEEEISRGLIEGAVSSVARIAIIPMQDVLGLGSDSRMNIPATQFGNWSW
RIPSSTSFDNLDAEAKKLRDILATYGRL
100000000000000000000000000000000000000000000000000000000000
000000000000000000000000000000000000000000000000000000000000
000000000000000000000000000000000000000000000000000000000000
000000000000000000000000000000000000000000000000000000000000
000000000000000000000000000000000000000000000000000000000000
000000000000000000000000000000000000000000000000000000000000
000000000000000000000000000000000000000000000000000000000000
000000000000000000000000000000000000000000000000000000000000
0000000000000000000000000000
>DM_train191
MASEVPQVVSLDPTSIPIEYNTPIHDIKVQVYDIKGGCNVEEGLTIFLVNNPGKENGPVKIS
SKVNDKQVSEFLKDENMEKFNVKLGTSKHFYMFNDNKNSVAVGYVGCGSVADLSEADMKRVV
LSLVTMLHDNKLSKLTVVFEINVDKNLFRFFLETLFYEYMTDERFKSTDKNVNMEYIKHLGV
YINNADTYKEEVEKARVYYFGTYYASQLIAAPSNYCNPVSLSNAAVELAQKLNLEYKILGVK
ELEELKMGAYLSVGKGSMYPNKFIHLTYKSKGDVKKKIALVGKGITFDSGGYNLKAAPGSMI
DLMKFDMSGCAAVLGCAYCVGTLKPENVEIHFLSAVCENMVSKNSYRPGDIITASNGKTIEV

GNTDAEGRLTLADALVYAEKLGVDYIVDIATLTGAMLYSLGTSYAGVFGNNEELINKILQSS
KTSNEPVWWLPIINEYRATLNSKYADINQISSSVKASSIVASLFLKEFVQNTAWAHIDIAGV
SWNFKARKPKGFGVRLLTEFVLNDAL
110000000000000000000000000000000000000000000000001000000000
000000000000000000000000000000000000000000000000000000000000
000000000000000000000000000000000000000000000011111110000000
000000000000000000000000000000000000000000000000000000000000
000000000000000000000000000000000000000000000000000000000000
000000000000000000000000000000000000000000000000000000000000
000000000000000000000000000000000000000000000000000000000000
000000000000000000000000000000000000000000000000000000000000
00000000000000000000000011
>DM_train193
MPSFASLKSLVVLSLTSLSLAATVALDLHILNANLDPDGTGARSAVTAEGTTIAPLITGNID
DRFQINVIDQLTDANMRRATSIHWHGFFQAGTTEMDGPAFVNQCPIIPNESFVYDFVVPGQA
GTYWYHSHLSTQYCDGLRGAFVVYDPNDPHLSLYDVDDASTVITIADWYHSLSTVLFPNPNK
APPAPDTTLINGLGRNSANPSAGQLAVVSVQSGKRYRFRIVSTSCFPNYAFSIDGHRMTVIE
VDGVSHQPLTVDSLTIFAGQRYSVVVEANQAVGNYWIRANPSNGRNGFTGGINSAIFRYQGA
AVAEPTTSQNSGTALNEANLIPLINPGAPGNPVPGGADINLNLRIGRNATTADFTINGAPFI
PPTVPVLLQILSGVTNPNDLLPGGAVISLPANQVIEISIPGGGNHPFHLHGHNFDVVRTPGS
SVYNYVNPVRRDVVSIGGGGDNVTFRFVTDNPGPWFLHCHIDWHLEAGLAVVFAEDIPNIPI
ANAISPAWDDLCPKYNANNPDSGLA
111111111111111111110000000000000000000000000000000000000000
000000000000000000000000000000000000000000000000000000000000
000000000000000000000000000000000000000000000000000011111110
000000000000000000000000000000000000000000000000000000000000
000000000000000000000000000000000000000000000000000000000000
000000000000000000000000000000000000000000000000000000000000
000000000000000000000000000000000000000000000000000000000000
000000000000000000000000000000000000000000000000000000000000
0000000000000000000111111
>DM_train194
MSASSSSALPPLVPALYRWKSTGSSGRQVQRRCVGAEAIVGLEEKNRRALYDLYIATSLRNI
APASTLLTLQNLKEMFELALLDARFEHPECACTVSWDDEVPAIITYESPESNESARDWARGC
IHVQPTAKSALDLWSEMEEGRAAANDNTPSKSIELFLLSDVSTDSTPIPQDATVEILFHSNH
LFWDGIGCRKFVGDLFRLVGSYIGRSDSREMKKIQWGQEIKNLSPPVVDSLKLDINTLGSEF
DDKCTEYTSALVANYKSRGMKFQPGLALPRCVIHKLSADESIDIVKAVKTRLGPGFTISHLT
QAAIVLALLDHLKPNDLSDDEVFISPTSVDGRRWLREDIASNFYAMCQTAAVVRIENLKSIT
VSHKDEKELQVRALESACRNIKKSYRQWLENPFLQALGLRVHNFEASYLHAKPIPFEGEANP
LFISDGINERFIPHEIKQTATGENVLSVESIDFVVNQSLPYLAIRLDSWRDASTLNIIYNDA
NYTEAEVQKYLQSIVEFMLAFRL
111111110000000000000000000000000000000000000000000000000000
000000000000000000000000000000000000000000000000000000000000
000000000000000000000000010000000000000000000000000000000000

000000000000000000000000000000000000000000000000000000000000
000000000000000000000000000000000000000000000000000000000000
000000000000011100000000000000000000000000000000000000000000
000000000000000000000000000000000000000000000000000000000000
000000000000000000000000000000000000000000000000000000000000
00000000000000000000000
>DM_train195
MEVKREHWATRLGLILAMAGNAVGLGNFLRFPVQAAENGGGAFMIPYIIAFLLVGIPLMWIE
WAMGRYGGAQGHGTTPAIFYLLWRNRFAKILGVFGLWIPLVVAIYYVYIESWTLGFAIKFLV
GLVPEPPPNATDPDSILRPFKEFLYSYIGVPKGDEPILKPSLFAYIVFLITMFINVSILIRG
ISKGIERFAKIAMPTLFILAVFLVIRVFLLETPNGTAADGLNFLWTPDFEKLKDPGVWIAAV
GQIFFTLSLGFGAIITYASYVRKDQDIVLSGLTAATLNEKAEVILGGSISIPAAVAFFGVAN
AVAIAKAGAFNLGFITLPAIFSQTAGGTFLGFLWFFLLFFAGLTSSIAIMQPMIAFLEDELK
LSRKHAVLWTAAIVFFSAHLVMFLNKSLDEMDFWAGTIGVVFFGLTELIIFFWIFGADKAWE
EINRGGIIKVPRIYYYVMRYITPAFLAVLLVVWAREYIPKIMEETHWTVWITRFYIIGLFLF
LTFLVFLAERRRNHESAGTLVPR
111100000000000000000000000000000000000000000000000000000000
000000000000000000000000000000000000000000000000000000000000
000000001100000000000000000000000000000000000000000000000000
000000000000000000000000000000000000000000000000000000000000
000000000000000000000000000000000000000000000000000000000000
000000000000000000000000000000000000000000000000000000000000
000000000000000000000000000000000000000000000000000000000000
000000000000000000000000000000000000000000000000000000000000
00000000000000000001111
>DM_train196
FNYDQPYRGQYHFSPQKNWMNDPNGLLYHNGTYHLFFQYNPGGIEWGNISWGHAISEDLTHW
EEKPVALLARGFGSDVTEMYFSGSAVADVNNTSGFGKDGKTPLVAMYTSYYPVAQTLPSGQT
VQEDQQSQSIAYSLDDGLTWTTYDAANPVIPNPPSPYEAEYQNFRDPFVFWHDESQKWVVVT
SIAELHKLAIYTSDNLKDWKLVSEFGPYNAQGGVWECPGLVKLPLDSGNSTKWVITSGLNPG
GPPGTVGSGTQYFVGEFDGTTFTPDADTVYPGNSTANWMDWGPDFYAAAGYNGLSLNDHVHI
GWMNNWQYGANIPTYPWRSAMAIPRHMALKTIGSKATLVQQPQEAWSSISNKRPIYSRTFKT
LSEGSTNTTTTGETFKVDLSFSAKSKASTFAIALRASANFTEQTLVGYDFAKQQIFLDRTHS
GDVSFDETFASVYHGPLTPDSTGVVKLSIFVDRSSVEVFGGQGETTLTAQIFPSSDAVHARL
ASTGGTTEDVRADIYKIASTWN
000000000000000000000000000000000000000000000000000000000000
000000000000000000000000000000000000000000000000000000000000
000000000000000000000000000000000000000000000000000000000000
000000000000000000000000000000000000000000000000000000000000
000000000000000000000000000000000000000000000000000000000000
000000000000000000000000000000000000000000000000000000000000
000000000000000000000000000000000000000000000000000000000000
000000000000000000000000000000000000000000000000000000000000
0000000000000000000001

>DM_train197
EPAKVVDIRIDTSAERKPISPYIYGSNQELDATVTAKRFGGNRTTGYNWENNFSNAGSDWLH
YSDTYLLEDGGVPKGEWSTPASVVTTFHDKALSKNVPYTLITLQAAGYVSADGNGPVSQEET
APSSRWKEVKFEKGAPFSLTPDTEDDYVYMDEFVNYLVNKYGNASTPTGIKGYSIDNEPALW
SHTHPRIHPDNVTAKELIEKSVALSKAVKKVDPYAEIFGPALYGFAAYETLQSAPDWGTEGE
GYRWFIDYYLDKMKKASDEEGKRLLDVLDVHWYPEARGGGERICFGADPRNIETNKARLQAP
RTLWDPTYIEDSWIGQWKKDFLPILPNLLDSIEKYYPGTKLAITEYDYGGGNHITGGIAQAD
VLGIFGKYGVYLATFWGDASNNYTEAGINLYTNYDGKGGKFGDTSVKCETSDIEVSSAYASI
VGEDDSKLHIILLNKNYDQPTTFNFSIDSSKNYTIGNVWAFDRGSSNITQRTPIVNIKDNTF
TYTVPALTACHIVLEAAEP
110000000000000000000000000000000000000000000000000000000000
000000000000000000000000000000000000000000000000000000000000
000000000000000000000000000000000000000000000000000000000000
000000000000000000000000000000000000000000000000000000000000
000000000000000000000000000000000000000000000000000000000000
000000000000000000000000000000000000000000000000000000000000
000000000000000000000000000000000000000000000000000000000000
000000000000000000000000000000000000000000000000000000000000
0000000000000001111
>DM_train198
AEQVALSRTHVCGILREELFQGDAFHQSDTHIFIIMGASGDLAKKKIYPTIWWLFRDGLLPE
NTFIVGYARSRLTVADIRKQSEPFFKATPEEKLKLEDFFARNSYVAGQYDDAASYQRLNSHM
NALHLGSQANRLFYLALPPTVYEAVTKNIHESCMSQIGWNRIIVEKPFGRDLQSSDRLSNHI
SSLFREDQIYRIDHYLGKEMVQNLMVLRFANRIFGPIWNRDNIACVILTFKEPFGTEGRGGY
FDEFGIIRDVMQNHLLQMLCLVAMEKPASTNSDDVRDEKVKVLKCISEVQANNVVLGQYVGN
PDGEGEATKGYLDDPTVPRGSTTATFAAVVLYVENERWDGVPFILRCGKALNERKAEVRLQF
HDVAGDIFHQQCKRNELVIRVQPNEAVYTKMMTKKPGMFFNPEESELDLTYGNRYKNVKLPD
AYERLILDVFCGSQMHFVRSDELLEAWRIFTPLLHQIELEKPKPIPYIYGSRGPTEADELMK
RVGFQYEGTYKWVNPHKL
111111110000111111111111111000000000000000000000000000000000
000000000000000000000000000000000000000000000000000000000000
000000000000000000000000000000000000000000000000000000000000
000000000000000000000000000000000000000000000000000000000000
000000000000000000000000000000000000000000000000000000000000
000000000000000000000000000000000000000000000000000000000000
000000000000000000000000000000000000000000000000000000000000
000000000000000000000000000000000000000000000000000000000000
000000000000001111
>DM_train199
DPDQFGPDLIEQLAQSGKYSQDNTKGDAMIGVKQPLPKAVLRTQHDKNKEAISILDFGVIDD
GVTDNYQAIQNAIDAVASLPSGGELFIPASNQAVGYIVGSTLLIPGGVNIRGVGKASQLRAK
SGLTGSVLRLSYDSDTIGRYLRNIRVTGNNTCNGIDTNITAEDSVIRQVYGWVFDNVMVNEV
ETAYLMQGLWHSKFIACQAGTCRVGLHFLGQCVSVSVSSCHFSRGNYSADESFGIRIQPQTY
AWSSEAVRSEAIILDSETMCIGFKNAVYVHDCLDLHMEQLDLDYCGSTGVVIENVNGGFSFS

NSWIAADADGTEQFTGIYFRTPTSTQSHKIVSGVHINTANKNTAANNQSIAIEQSAIFVFVS
GCTLTGDEWAVNIVDINECVSFDKCIFNKPLRYLRSGGVSVTDCYLAGITEVQKPEGRYNTY
RGCSGVPSVNGIINVPVAVGATSGSAAIPNPGNLTYRVRSLFGDPASSGDKVSVSGVTINVT
RPSPVGVALPSMVEYLAI
111000000000000000000000000000000000000000000000000000000000000
000000000000000000000000000000000000000000000000000000000000000
000000000000000000000000000000000000000000000000000000000000000
000000000000000000000000000000000000000000000000000000000000000
000000000000000000000000000000000000000000000000000000000000000
000000000000000000000000000000000000000000000000000000000000000
000000000000000000000000000000000000000000000000000000000000000
000000000000000000000000000000000000000000000000000000000000000
000000000000000000
>DM_train200
MPIHDKSPRPQEFAAVDLGSNSFHMVIARVVDGAMQIIGRLKQRVHLADGLGPDNMLSEEAM
TRGLNCLSLFAERLQGFSPASVCIVGTHTLRQALNATDFLKRAEKVIPYPIEIISGNEEARL
IFMGVEHTQPEKGRKLVIDIGGGSTELVIGENFEPILVESRRMGCVSFAQLYFPGGVINKEN
FQRARMAAAQKLETLTWQFRIQGWNVAMGASGTIKAAHEVLMEMGEKDGIITPERLEKLVKE
VLRHRNFASLSLPGLSEERKTVFVPGLAILCGVFDALAIRELRLSDGALREGVLYEMEGRFR
HQDVRSRTASSLANQYHIDSEQARRVLDTTMQMYEQWREQQPKLAHPQLEALLRWAAMLHEV
GLNINHSGLHRHSAYILQNSDLPGFNQEQQLMMATLVRYHRKAIKLDDLPRFTLFKKKQFLP
LIQLLRLGVLLNNQRQATTTPPTLTLITDDSHWTLRFPHDWFSQNALVLLDLEKEQEYWEGV
AGWRLKIEEESTPEIAA
111111111110000000000000000000000000000000000000000000000000000
000000000000000000000000000000000000000000000000000000000000000
000000000000000000000000000000000000000000000000000000000000000
000000000000000000000000000000000000000000000000000000000000000
000000000000000000000000000000000000000000000000000000000000000
000000000000000000000000000000000000000000000000000000000000000
000000000000000000000000000000000000000000000000000000000000000
000000000000000000000000000000000000000000000000000000000000000
00000000000001111
>DM_train201
GSPEFMHTINEEERREFIKHINSVLAGDPDVGSRVPINTETFEFFDQCKDGLILSKLINDSV
PDTIDERVLNKQRNNKPLDNFKCIENNNVVINSAKAMGGISITNIGAGDILEGREHLILGLV
WQIIRRGLLGKIDITLHPELYRLLEEDETLDQFLRLPPEKILLRWFNYHLKAANWPRTVSNF
SKDVSDGENYTVLLNQLAPELCSRAPLQTTDVLQRAEQVLQNAEKLDCRKYLTPTAMVAGNP
KLNLAFVAHLFNTHPGLEPLNEEEKPEIEPFDAEGEREARVFTLWLNSLDVTPSIHDFFNNL
RDGLILLQAYDKITPNTVNWKKVNKAPASGDEMMRFKAVENCNYAVDLGKNQGFSLVGIQGA
DITDGSRTLTLALVWQMMRMNITKTLHSLSRGGKTLSDSDMVAWANSMAAKGGKGSQIRSFR
DPSISTGVFVLDVLHGIKSEYVDYNLVTDGSTEELAIQNARLAISIARKLGAVIFILPEDIV
AVRPRLVLHFIGSLMAV
111111110000000000000000000000000000000000000000000000000000000
000000000000111000000000000000000000000000000000000000000000000

00000000000011111111111111110000000000000000000000000000000000
00000000000000000000000000000000000000000000000000000000000000
00000000000000000001111111111111000000000000000000000000000000
00000000000000000000000000000000000000000000000000000000000000
00000000000000000000000000001111110000000000000000000000000000
00000000000000000000000000000000000000000000000000000000000000
00000000000000000
>DM_train202
GMPEIHQSIAQHYHERTKYDPETIASKSQRLDWAKQPVPFKEYKIGSAIDLKPYLQETPEVF
VNDTNGQWWQRLSRLLFRSYGLTARMPSMGNTVYLRAAPSAGGLYPAEVYVVSRGTPLLSPG
LYNYQCRTHSLIHYWESDVWQSLQEACFWHPALESTQLAIIVTAVFYRSAWRYEDRAYRRIC
LDTGHLLGNIELSAAITDYRPHLIGGFIDEAVNDLLYIDPLQEGAIAVLPLADLLDIQQNIS
PGCTALPSATETNYPQVPDGELLKYFHHHTQISASITGKLNLPTVIQEKSLEDKYNFPFCLK
ISTVSAPIYWGENLSDLEITMHKRRSTRAYNGEELTFDELKALLDFTYQPQNYIDQSLDNSP
DYFDLNLIETFIAVCGVQGLEAGCYYYAPKAQELRQIRFKNFRRELHFLCLGQELGRDAAAV
IFHTSDLKSAIAQYGDRVYRYLHMDAGHLGQRLNLAAIQLNLGVSGIGGFFDDQVNEVLGIP
NDEAVIYITTLGRPR
11111000000000000000000000110000000000000000000000000000000111
11000000000000000000000000000000000000000000000000000000000000
00000000000000000000000000000000000000000000000000000000000000
00000000000000000000000000000000000000000000000000000000000000
00000000000000000000000000000000000000111111111110000000000000
00000000000000000000000000000000000000000000000000000000000000
00000000000000000000000000000000000000000000000000000000000000
00000000000000000000000000000000000000000000000000000000000000
000000000000000
>DM_train204
MSNDKTGKSLEQENSERDVEIRDRNYFRKLSLFDDTVIAGAEMIGTSYDVFGKYCNVGSCMN
SLFDERKINASEDNFKKVTILGKTLKVPYYIDCYSVGDLKYTNASGESIESYQSNISSKSRI
KGNYLFFSASLKVDFDTDSLTDFENAFSRIQYTYDLYILKSSAEALKEFLKESVKTALDKAD
TEEDMNDLFNTWGSHFLSGVVMGGCAQYSSSTNKYTSNLTNSFDVVAAASFAGFIGLSARTG
NSFMEDIKKFRSASNIKTHAIGGDLSRFDPFGGATSADQPSAEEIAAAKKAFEDWKASVPNA
PELVNFADSNPLTGIWELCSDRTQKAKLKKHFETVWAPAESAKRRVHADYIDEIIIGINNTN
TPPEGYIGLKSTKDENLNSKGNICLFMHKAKYDPNIDNKDCITELKFITVRDKSPEGDWVKI
PQDIYISPNQYLYLCYLPAKYSAEKAIKDIQLLCSSCGSSMILPYGYNDVLDERGERANATE
DDNVHYLIYSAGWK
11111110000000000000000000000000000000000000000000000000000000
00000000000000000000000000000000000000000000000000000000000000
00000000000000000000000000000000000000000000000000000000000000
00000000000000000000000000000000000000000000000000000000000000
00000000000000000000000000000000000000000000000000000000000000
00000000000000000000000000000000000000000000000000000000000000
00000000000000000000000000000000000000000000000000000000000000
00000000000000000000000000000000000111100000000000000000000000

00000000000001
>DM_train206
GVSKEEYKDVETAKKEKEQLGELMEPALGYVVKVPVSSFENKKVDISDIEVITNGNLDDVPY
KANSSKYNYPDIKTKDSSLQYVRSGYVIDGEHSGSNEKGYVYYKGNSPAKELPVNQLLTYTG
SWDFTSNANLNNEEGRPNYLNDDYYTKFIGKRVGLVSGDAKPAKHKYTSQFEVDFATKKMTG
KLSDKEKTIYTVNADIRGNRFTGAATASDKNKGKGESYNFFSADSQSLEGGFYGPKAEEMAG
KFVANDKSLFAVFSAKHNGSNVDTVRIIDASKIDLTNFSISELNNFGDASVLIIDGKKIKLA
GSGFTNKHTIEINGKTMVAVACCSNLEYMKFGQLWQQAEGGKPENNSLFLQGERTATDKMPK
GGNYKYIGTWDAQVSKENNWVATADDDRKAGYRTEFDVDFGNKNLSGKLFDKNGVNPVFTVD
PKIDGNGFTGKAKTSDAGFALDSGSSRYENVKFNDVAVSGGFYGPTAAELGGQFHHKSENGS
VGAVFGAKQQVKK
111100000000000000000000000000000000000000000000000000000000
000000000000000000000000000000000000000000000000000000000000
000000000000000000000000000000000000000000000000000000000000
000000000000000000000000000000000000000000000000000000000000
000000000000000000000000000000000000000000000000000000000000
000000000000000000000000000000000000000000000000000000000000
000000000000000000000000000000000000000000000000000000000000
000000000000000000000000000000000000000000000000000000000000
0000000000000
>DM_train207
MFRLGTTPRATTYDPDARIGEVASRFGLPTRVLIEIVRTESFQRSLARVTSGKPVVLDLREL
DSDLASWIATHARLVEPALRELVRTVAPDVEPRVRFRGLPHRFRRVERIRPMDGALISIEGV
VREVRGAERLEHAIVDTGSELVAVRLHGHRLGPGLRVEILGIVRSATLDALEVHKKDPIPEV
HPDPAELEEFRELADKDPLTTFARAIAPLPGAEEVGKMLALQLFSCVGKNSERLHVLLAGYP
VVCSEILHHVLDHLAPRGVYVDLRRTELTDLTAVLKEDRGWALRAGAAVLADGGILAVDHLE
GAPEPHRWALMEAMDKGTVTVDGIALNARCAVLAAINPGEQWPSDPPIARIDLDQDFLSHFD
LIAFLGVDPRPGEPEEQDTEVPSYTLLRRYLLYAIREHPAPELTEEARKRLEHWYETRREEV
EERLGMGLPTLPVTRRQLESVERLAKAHARMRLSDDVEPEDVDIAAELVDWYLETAMQIPGG
DEIRISSLKP
111111111111111110000000000000000000000000000000000000000000
000000000000000000000000000000000000000000000000000000000000
000000000000000000000000000000000000000000000000000000000000
000000000000000000000000000000000000000000000000000000000000
000000000000000000000000000000000000000000000000000000000000
000000000000000000000000000000000000001100000000000000000000
000000011111111111111000000000000000000000000000000000000000
000000000000000000000000000000000000000000000000000000001111
1111111111
>DM_train208
GAASEAEYGKVSKAWTLHADGSQEYRSSMELTLFTHTAMNSTYGESFIVYNPDFQTLKIHSS
YTRQKDGTIVKTPDNAFVEVLPRFAADAPAYNQLKEMVVVHTGLELGATIYLDYSIITKPGY
YPALDINERLQETSPVKECKVSISVPEGTPLACGLYGSPVKAVEESHDGIKEVHWTLRNIPA
SSREAFQPKNREASPHLVASTYPSGKAALATLDKRLKESQGYESKTFAQFLTDKSGNEQEKV

NIIRDHILNNLSTCPIPMAMTGYTVRDIDTVLRSAYGTPLEIAQLLNVMLNAAGIPSEVLAV
YPGHLDTDACGLAAIQTLAVKATVDGKDQYLSASPLTNRGGLDKVVSLSGTSIEIETTPIQI
KESRSVAISADQAKDGFAICVLPAISAGIDSWGMSALNSKRSNLFELPSLIREEVTYTVTPA
EGMKLQTSTQEQVISKPFGKVTRTITPRGNTIEVVRTIELNKQQFTPAEYSDVRSLIHEWTN
PDNRVLLFSL
100000000000000000000000000000000000000000000000000000000000
000000000000000000000000000000000000000000000000000000000000
000000000000000000000000000000000000000000000000000000000000
000000000000000000000000000000000000000000000000000000000000
000000000000000000000000000000000000000000000000000000000000
000000000000000000000000000000000000000000000000000000000000
000000000000000000000000000000000000000000000000000000000000
000000000000000000000000000000000000000000000000000000000000
0000000000
>DM_train212
MATKKATMIIEKDFKIAEIDKRIYGSFIEHLGRAVYGGIYEPGHPQADENGFRQDVIELVKE
LQVPIIRYPGGNFVSGYNWEDGVGPKEQRPRRLDLAWKSVETNEIGLNEFMDWAKMVGAEVN
MAVNLGTRGIDAARNLVEYCNHPSGSYYSDLRIAHGYKEPHKIKTWCLGNAMDGPWQIGHKT
AVEYGRIACEAAKVMKWVDPTIELVVCGSSNRNMPTFAEWEATVLDHTYDHVDYISLHQYYG
NRDNDTANYLALSLEMDDFIRSVVAIADYVKAKKRSKKTIHLSFDEWNVWYHSNEADKLIEP
WTVAPPLLEDIYNFEDALLVGCMLITLMKHADRVKIACLAQLVNVIAPIMTEKNGPAWKQTI
YYPFMHASVYGRGVALHPVISSPKYDSKDFTDVPYLESIAVYNEEKEEVTIFAVNRDMEDAL
LLECDVRSFEDYRVIEHIVLEHDNVKQTNSAQSSPVVPHRNGDAQLSDRKVSATLPKLSWNV
IRLGKR
111100000000000000000000000000000000000000000000000000000000
000000000000000000000000000000000000000000000000000000000000
000000000000000000000000000000000000000000000000000000000000
000000000000000000000000000000000000000000000000000000000000
000000000000000000000000000000000000000000000000000000000000
000000000000000000000000000000000000000000000000000000000000
000000000000000000000000000000000000000000000000000000000000
000000000000000000000000000000000000000000000000000000000000
000001
>DM_train214
METKDLIVIGGGINGAGIAADAAGRGLSVLMLEAQDLACATSSASSKLIHGGLRYLEHYEFR
LVSEALAEREVLLKMAPHIAFPMRFRLPHRPHLRPAWMIRIGLFMYDHLGKRTSLPGSTGLR
FGANSVLKPEIKRGFEYSDCWVDDARLVLANAQMVVRKGGEVLTRTRATSARRENGLWIVEA
EDIDTGKKYSWQARGLVNATGPWVKQFFDDGMHLPSPYGIRLIKGSHIVVPRVHTQKQAYIL
QNEDKRIVFVIPWMDEFSIIGTTDVEYKGDPKAVKIEESEINYLLNVYNTHFKKQLSRDDIV
WTYSGVRPLCDDESDSPQAITRDYTLDIHDENGKAPLLSVFGGKLTTYRKLAEHALEKLTPY
YQGIGPAWTKESVLPGGAIEGDRDDYAARLRRRYPFLTESLARHYARTYGSNSELLLGNAGT
VSDLGEDFGHEFYEAELKYLVDHEWVRRADDALWRRTKQGMWLNADQQSRVSQWLVEYTQQR
LSLAS
110000000000000000000000000000000000000000000000000000000000

000000000000000000000000000000000000000000000000000000000000
000000000000000000000000000000000000000000000000000000000000
000000000000000000000000000000000000000000000000000000000000
000000000000000000000000000000000000000000000000000000000000
000000000000000000000000000000000000000000000000000000000000
000000000000000000000000000000000000000000000000000000000000
000000000000000000000000000000000000000000000000000000000000
00000
>DM_train215
MVYDLIVIGGGSGGMAAARRAARHNAKVALVEKSRLGGTCVNVGCVPKKIMFNAASVHDILE
NSRHYGFDTKFSFNLPLLVERRDKYIQRLNNIYRQNLSKDKVDLYEGTASFLSENRILIKGT
KDNNNKDNGPLNEEILEGRNILIAVGNKPVFPPVKGIENTISSDEFFNIKESKKIGIVGSGY
IAVELINVIKRLGIDSYIFARGNRILRKFDESVINVLENDMKKNNINIVTFADVVEIKKVSD
KNLSIHLSDGRIYEHFDHVIYCVGRSPDTENLKLEKLNVETNNNYIVVDENQRTSVNNIYAV
GDCCMVKKSKEIEDLNLLKLYNEERYLNKKENVTEDIFYNVQLTPVAINAGRLLADRLFLKK
TRKTNYKLIPTVIFSHPPIGTIGLSEEAAIQIYGKENVKIYESKFTNLFFSVYDIEPELKEK
TYLKLVCVGKDELIKGLHIIGLNADEIVQGFAVALKMNATKKDFDETIPIHPTAAEEFLTLQ
PWMK
000000000000000000000000000000000000000000000000000000000000
000000000000000000000000000000000000000000000000001111111111
111111111111111100000000000000000000000000000000000000000000
000000000000000000000000000000000000000000000000000000000000
000000000000000000000000000000000000000000000000000000000000
000000011111111111111111111111111111000000000000000000000000
000000000000000000000000000000000000000000000000000000000000
000000000000000000000000000000000000000000000000000000000000
1111
>DM_train216
TEKKYIVALDQGTTSSRAVVMDHDANIISVSQREFEQIYPKPGWVEHDPMEIWATQSSTLVE
VLAKADISSDQIAAIGITNQRETTIVWEKETGKPIYNAIVWQCRRTAEICEHLKRDGLEDYI
RSNTGLVIDPYFSGTKVKWILDHVEGSRERARRGELLFGTVDTWLIWKMTQGRVHVTDYTNA
SRTMLFNIHTLDWDDKMLEVLDIPREMLPEVRRSSEVYGQTNIDGKGGTRIPISGIAGDQQA
ALFGQLCVKEGMAKNTYGTGCFMLMNTGEKAVKSENGLLTTIACGPTGEVNYALEGAVFMAG
ASIQWLRDEMKLINDAYDSEYFATKVQNTNGVYVVPAFTGLGAPYWDPYARGAIFGLTRGVN
ANHIIRATLESIAYQTRDVLEAMQADSGIRLHALRVDGGAVANNFLMQFQSDILGTRVERPE
VREVTALGAAYLAGLAVGFWQNLDELQEKAVIEREFRPGIETTERNYRYAGWKKAVKRAMAW
EEHD
100000000000000000000000000000000000000000000000000000000000
000000000000000000000000000000000000000000000000000000000000
000000000000000000000000000000000000000000000000000000000000
000000000000000000000000000000000000000011111100000000000000
000000000000000000000000000000000000000000000000000000000000
000000000000011110000000000000000000000000000000000000000000
000000000000000000000000000000000000000000000000000000000001

110000000000000000000000000111000000000000000000000000000000
0000
>DM_train217
AFYIATFDIGTTEVKAALADRDGGLHFQRSIALETYGDGNGPVEQDAGDWYDAVQRIASSWW
QSGVDARRVSAIVLSGQMQNFLPLDQDHEPLHRAVLYSDKRPLKEAEEINARHGADNLWSAL
ENPMTAASILPKLVFWRASFPQAFGRLRHVVLGAKDYVVLRLTGRHATDRTNASTTGLYRPK
DDAWHVELLADYGFSLDLMPRLLEPGEQVGGVSALAARQTGFVSGTPVLCGLGDAGAATLGV
GVLDDEDAYLHLGTTGWLARLTQTDPVGDMPVGTIFRLAGIIAGKTLQVAPVLNAGNILQWA
LTLVGHRPGEDCAEYFHMAAAEVQGVTVPDGLLFVPYLHAERCPVELPAPRGALLGVTGATT
RAQILLAVLEGAALSLRWCAELLGMEKVGLLKVVGGGARSEAWLRMIADNLNVSLLVKPDAH
LHPLRGLAALAAVELEWSHSIQDFLREADLREPASNILHPQPCDEGRRRRKFERFKQCVETL
GRLD
000000000000000000000000000000000000000000000000000000000000
000000000000000000000000000000000000000000000000000000000000
000000000000000000000000000000000000000000000000000000000000
000000000000000000000000000000000000000000000000000000000000
000000000000000000000000000000000000000000000000000000000000
000000000000000000000000000000000000000000000000000000000000
000000000000000000000000000000000000000000000000000000000000
000000000000000000000000000000111000000000000000000000000000
0011
>DM_train218
MASLQLLPSPTPNLEIKYTKIFINNEWQNSESGRVFPVCNPATGEQVCEVQEADKVDIDKAV
QAARLAFSLGSVWRRMDASERGRLLDKLADLVERDRATLATMESLNGGKPFLQAFYIDLQGV
IKTLRYYAGWADKIHGMTIPVDGDYFTFTRHEPIGVCGQIIPWNFPLLMFTWKIAPALCCGN
TVVIKPAEQTPLSALYMGALIKEAGFPPGVVNILPGYGPTAGAAIASHIGIDKIAFTGSTEV
GKLIQEAAGRSNLKRVTLELGGKSPNIIFADADLDYAVEQAHQGVFFNQGQCCTAGSRIFVE
ESIYEEFVKRSVERAKRRIVGSPFDPTTEQGPQIDKKQYNKILELIQSGVAEGAKLECGGKG
LGRKGFFIEPTVFSNVTDDMRIAKEEIFGPVQEILRFKTMDEVIERANNSDFGLVAAVFTND
INKALMVSSAMQAGTVWINCYNALNAQSPFGGFKMSGNGREMGEFGLREYSEVKTVTVKIPQ
KNS
000000000000000000000000000000000000000000000000000000000000
000000000000000000000000000000000000000000000000000000000000
000000000000000000000000000000000000000000000000000000000000
000000000000000000000000000000000000000000000000000000000000
000000000000000000000000000000000000000000000000000000000000
000000000000000000000000000000000000000000000000000000000000
000000000000000000000000000000000000000000000000000000000000
000000000000000000001111111111111111111100000000000000000000
000
>DM_train220
LEPRHAMLLRCRLPGGVITTKQWQAIDKFAGENTIYGSIRLTNRQTFQFHGILKKNVKPVHQ
MLHSVGLDALATANDMNRNVLCTSNPYESQLHAEAYEWAKKISEHLLPRTRAYAEIWLDQEK
VATTDEEPILGQTYLPRKFKTTVVIPPQNDIDLHANDMNFVAIAENGKLVGFNLLVGGGLSI

EHGNKKTYARTASEFGYLPLEHTLAVAEAVVTTQRDWGNRTDRKNAKTKYTLERVGVETFKA
EVERRAGIKFEPIRPYEFTGRGDRIGWVKGIDDNWHLTLFIENGRILDYPARPLKTGLLEIA
KIHKGDFRITANQNLIIAGVPESEKAKIEKIAKESGLMNAVTPQRENSMACVSFPTCPLAMA
EAERFLPSFIDNIDNLMAKHGVSDEHIVMRVTGCPNGCGRAMLAEVGLVGKAPGRYNLHLGG
NRIGTRIPRMYKENITEPEILASLDELIGRWAKEREAGEGFGDFTVRAGIIRPVLDPARDLW
D
11111111000000000000000000000000000000000000000000000111110000
00000000001110000000000000000000000000000000000011111111111111
11111111111111000000000000000000000000000000000000000000000000
00000000000000000000000000000000000000000000000000000000000000
00000000000000000000000000000000000000000000000000000000000000
00000000000000000000000000000000000000000000000000000000000000
00000000000000000000000000000000000000000000000000000000000000
00000000000000000000000000000000000000000000000000000000000000
0
>DM_train221
MAHHHHHHMASPASTNPAHDHFETFVQAQLCQDVLSSFQGLCRALGVESGGGLSQYHKIKAQ
LNYWSAKSLWAKLDKRASQPVYQQGQACTNTKCLVVGAGPCGLRAAVELALLGARVVLVEKR
IKFSRHNVLHLWPFTIHDLRALGAKKFYGRFCTGTLDHISIRQLQLLLLKVALLLGVEIHWG
VKFTGLQPPPRKGSGWRAQLQPNPPAQLASYEFDVLISAAGGKFVPEGFTIREMRGKLAIGI
TANFVNGRTVEETQVPEISGVARIYNQKFFQSLLKATGIDLENIVYYKDETHYFVMTAKKQC
LLRLGVLRQDLSETDQLLGKANVVPEALQRFARAAADFATHGKLGKLEFAQDARGRPDVAAF
DFTSMMRAESSARVQEKHGARLLLGLVGDCLVEPFWPLGTGVARGFLAAFDAAWMVKRWAEG
AGPLEVLAERESLYQLLSQTSPENMHRNVAQYGLDPATRYPNLNLRAVTPNQVQDLYDMMDK
E
11111111111111000000000000000000000000000000000000000000000000
00000000000000000000000000000000000000000000000000000000000000
00000000000000000000000000000000000000000000000000000000000000
00000000000000000000000000000000000000000000000000000000000000
00000000000000000000111100000000000000000000000000000000000000
00000000000000000000000000000000000000000000000000000000000000
00000000000000000000000000000000000000000000000000000000000000
00000000000000000000000000000000000000000000000000000000000000
0
>DM_train222
GKDDDNVETGAFDPSKPVAISDFTPKEGGAYQKLLIYGENFGTDVSKVKVKIGGKDAIVINV
KSTYVYCFVPSGAFSGEIEITVGEGENAVTTTASTTFSYEKKMVVGTLCGYRNNRDDQGWRD
GPFDGPEGVKCCGFSDNGRLAFDPLNKDHLYICYDGHKAIQLIDLKNRMLSSPLNINTIPTN
RIRSIAFNKKIEGYADEAEYMIVAIDYDGKGDESPSVYIIKRNADGTFDDRSDIQLIAAYKQ
CNGATIHPINGELYFNSYEKGQVFRLDLVDYFKTIKNGGSWDPIVKNNPNTFKQLFTIADPS
WEFQIFIHPTGKYAYFGVINNHYFMRSDYDEIKKEFITPYNFVGGYKQSGYRDDVGTEARMN
NPCQGVFVKNPDYTGEEEYDFYFVDRLNFCVRKVTPEGIVSTYAGRGASTSLADGNQWGTDD
GDLREVARFRDVSGLVYDDVKEMFYVHDQVGHTIRTISMEQEENVAGDENIPEDESTVESNE
11111100000000000000000000000000000000000000000000000000000000

000000000000000000000000000000000000000000000000000000000000000
000000000000000000000000000000000000000000000000000000000000000
000000000000000000000000000000000000000000000000000000000000000
000000000000000000000000000000000000000000000000000000000000000
000000000000000000000000000000000000000000000000000000000000000
000000000000000000000000000000000000000000000000000000000000000
000000000000000000000000000000000000000000011111111111111111111
>DM_train223
ATSDSNMLLNYVPVYYVMLPLGVVNVDNVFEDPDGLKEQLLQLRAAGVDGVMVDVWWGIIELK
GPKQYDWRAYRSLLQLVQECGLTLQAIMSFHQCGGNVGDIVNIPIPQWVLDIGESNHDIFYT
NRSGTRNKEYLTVGVDNEPIFHGRTAIEIYSDYMKSFRENMSDFLESGLIIDIEVGLGPAGE
LRYPSYPQSQGWEFPGIGEFQCYDKYLKADFKAAVARAGHPEWELPDDAGKYNDVPESTGFF
KSNGTYVTEKGKFFLTWYSNKLLNHGDQILDEANKAFLGCKVKLAIKVSGIHWWYKVENHAA
ELTAGYYNLNDRDGYRPIARMLSRHHAILNFTCLEMRDSEQPSDAKSGPQELVQQVLSGGWR
EDIRVAGENALPRYDATAYNQIILNARPQGVNNNGPPKLSMFGVTYLRLSDDLLQKSNFNIF
KKFVLKMHADQDYCANPQKYNHAITPLKPSAPKIPIEVLLEATKPTLPFPWLPETDMKVDG
111110000000000000000000000000000000000000000000000000000000000
000000000000000000000000000000000000000000000000000000000000000
000000000000000000000000000000000000000000000000000000000000000
000000000000000000000000000000000000000000000000000000000000000
000000000000000000000000000000000000000000000000000000000000000
000000000000000000000000000000000000000000000000000000000000000
000000000000000000000000000000000000000000000000000000000000000
0000000000000000000000000000000000000000000000000000000000000
>DM_train224
MTSRDGYQWTPETGLTQGVPSLGVISPPTNIEDTDKDGPWDVIVIGGGYCGLTATRDLTVAG
FKTLLLEARDRIGGRSWSSNIDGYPYEMGGTWVHWHQSHVWREITRYKMHNALSPSFNFSRG
VNHFQLRTNPTTSTYMTHEAEDELLRSALHKFTNVDGTNGRTVLPFPHDMFYVPEFRKYDEM
SYSERIDQIRDELSLNERSSLEAFILLCSGGTLENSSFGEFLHWWAMSGYTYQGCMDCLMSY
KFKDGQSAFARRFWEEAAGTGRLGYVFGCPVRSVVNERDAARVTARDGREFVAKRVVCTIPL
NVLSTIQFSPALSTERISAMQAGHVSMCTKVHAEVDNKDMRSWTGIAYPFNKLCYAIGDGTT
PAGNTHLVCFGNSANHIQPDEDVRETLKAVGQLAPGTFGVKRLVFHNWVKDEFAKGAWFFSR
PGMVSECLQGLREKHGGVVFANSDWALGWRSFIDGAIEEGTRAARVVLEELGTKREVKARL
000000000000000000000000000000011111111000000000000000000000000
000000000000000000000000000000000000000000000000000000000000000
000000000000000000000000000000000000000000000000000000000000000
000000000000000000000000000000000000000000000000000000000000000
000000000000000000000000000000000000000000000000000000000000000
000000000000000000000000000000000000000000000000000000000000000
000000000000000000000000000000000000000000000000000000000000000
0000000000000000000000000000000000000000000000000000111111111
>DM_train225
MAQILPIRFQEHLQLQNLGINPANIGFSTLTMESDKFICIREKVGEQAQVVIIDMNDPSNPI
RRPISADSAIMNPASKVIALKAGKTLQIFNIEMKSKMKAHTMTDDVTFWKWISLNTVALVTD

NAVYHWSMEGESQPVKMFDRHSSLAGCQIINYRTDAKQKWLLLTGISAQQNRVVGAMQLYSV
DRKVSQPIEGHAASFAQFKMEGNAEESTLFCFAVRGQAGGKLHIIEVGTPPTGNQPFPKKAV
DVFFPPEAQNDFPVAMQISEKHDVVFLITKYGYIHLYDLETGTCIYMNRISGETIFVTAPHE
ATAGIIGVNRKGQVLSVCVEEENIIPYITNVLQNPDLALRMAVRNNLAGAEELFARKFNALF
AQGNYSEAAKVAANAPKGILRTPDTIRRFQSVPAQPGQTSPLLQYFGILLDQGQLNKYESLE
LCRPVLQQGRKQLLEKWLKEDKLECSEELGDLVKSVDPTLALSVYLRANVPNKVIQCFAE
000000000000000000000000000000000000000000000000000000000000
000000000000000000000000000000000000000000000000000000000000
000000000000000000000000000000000000000000000000000000000000
000000000000000000000000000000000000000000000000000000000000
000000000000000000000000000000000000000000000000000000000000
000000000000000000000000000000000000000000000000000000000000
000000000000000000000000000000000000000000000000000000000000
0000000000000000000000000000000000000000000000000001111111
>DM_train226
MDYTYADDSLTLHTDMYQINMMQTYWELGRADLHAVFECYFREMPFNHGYAIFAGLERLVNY
LENLTFTESDIAYLREVEEYPEDFLTYLANFEFKCTVRSALEGDLVFNNEPLIQIEGPLAQC
QLVETALLNMVNFQTLIATKAARIKSVIGDDPLLEFGTRRAQELDAAIWGTRAAYIGGADAT
SNVRAGKIFGIPVSGTHAHSLVQSYGNDYEAFMAYAKTHRDCVFLVDTYDTLKAGVPSAIRV
AREMGDKINFLGVRIDSGDMAYISKRVREQLDEAGFTEAKIYASNDLDENTILNLKMQKSKI
DVWGVGTKLITAYDQPALGAVFKLVSIEGEDGQMKDTIKLSSNAEKVTTPGKKQVWRITRKS
DKKSEGDYVTLWNEDPRQEEEIYMFHPVHTFINKYVRDFEARPVLQDIFVEGKRVYELPTLD
EIKQYAKENLDSLHEEYKRDLNPQKYPVDLSTDCWNHKMNLLEKVRKDVKHLTETVNKEA
111000000000000000000000000000000000000000000000000000000000
000000000000000000000000000000000000000000000000000000000000
000000000000000000000000000000000000000000000000000000000000
000000000000000000000000000000000000000000000000000000000000
000000000000000000000000000000000000000000000000000000000000
000000000000000000000000000000000000000000000000000000000000
000000000000000000000000000000000000000000000000000000000000
0000000000000000000000000000000000000000000000000111111111
>DM_train227
TGNLVTKNSLTPDVRNGIDFKIADLSLADFGRKELRIAEHEMPGLMSLRREYAEVQPLKGAR
ISGSLHMTVQTAVLIETLTALGAEVRWASCNIFSTQDHAAAAVVVGPHGTPDEPKGVPVFAW
KGETLEEYWWAAEQMLTWPDPDKPANMILDDGGDATMLVLRGMQYEKAGVVPPAEEDDPAEW
KVFLNLLRTRFETDKDKWTKIAESVKGVTEETTTGVLRLYQFAAAGDLAFPAINVNDSVTKS
KFDNKYGTRHSLIDGINRGTDALIGGKKVLICGYGDVGKGCAEAMKGQGARVSVTEIDPINA
LQAMMEGFDVVTVEEAIGDADIVVTATGNKDIIMLEHIKAMKDHAILGNIGHFDNEIDMAGL
ERSGATRVNVKPQVDLWTFGDTGRSIIVLSEGRLLNLGNATGHPSFVMSNSFANQTIAQIEL
WTKNDEYDNEVYRLPKHLDEKVARIHVEALGGHLTKLTKEQAEYLGVDVEGPYKPDHYRY
111111111000000000000000000000000000000000000000000000000000
000000000000000000000000000000000000000000000000000000000000
000000000000000000000000000000000000000000000000000000000000
000000000000000000000000000000000000000000000000000000000000

000000000000000000000000000000000000000000000000000000000000
000000000000000000000000000000000000000000000000000000000000
000000000000000000000000000000000000000000000000000000000000
0000000000000000000000000000000000000000000000000000000000
>DM_train228
MTANGTAEAVQIQFGLINCGNKYLTAEAFGFKVNASASSLKKKQIWTLEQPPDEAGSAAVCL
RSHLGRYLAADKDGNVTCEREVPGPDCRFLIVAHDDGRWSLQSEAHRRYFGGTEDRLSCFAQ
TVSPAEKWSVHIAMHPQVNIYSVTRKRYAHLSARPADEIAVDRDVPWGVDSLITLAFQDQRY
SVQTADHRFLRHDGRLVARPEPATGYTLEFRSGKVAFRDCEGRYLAPSGPSGTLKAGKATKV
GKDELFALEQSCAQVVLQAANERNVSTRQGMDLSANQDEETDQETFQLEIDRDTKKCAFRTH
TGKYWTLTATGGVQSTASSKNASCYFDIEWRDRRITLRASNGKFVTSKKNGQLAASVETAGD
SELFLMKLINRPIIVFRGEHGFIGCRKVTGTLDANRSSYDVFQLEFNDGAYNIKDSTGKYWT
VGSDSAVTSSGDTPVDFFFEFCDYNKVAIKVGGRYLKGDHAGVLKASAETVDPASLWEY
111111100000000000000000000000000000000000000001111111100000
000000000000000000000000000000000000000000000000000000000000
000000000000000000000000000000000000000000000000000000000000
000000000000000000000000000000000000000000000000000000000000
000000000000000000000000000111100000000000000000000000000000
000000000000000000000000000000000000000000000000000000000000
000000000000000000000000000000000000000000000000000000000000
000000000000000000000000000000000000000000000000000000000
>DM_train229
GGTAYWKNPDQFTAFNTGLHALLREKSYNFFLLGEPRADIYGDNPIGGEASQGMERLPFNTI
NKENVGISNYGDMYKIINQINQMIAKTTETTILTEATQNYYLGEAYGMRAYLYFHLLRSWGD
VVLYLDYTEGQNLDLSNITKGVSPATEVMEQIKKDIQASENAFGSDYSFKLGRHFWSAAATQ
MLKGEAYLWSGRQMNGGNSDYTIAKNAFENVKKADVGLVTSSFKDIFSFENKKNKEMIFTIH
NGKDEYEMWGGYYRMRLIPAQDKMVKIYCDENGNSFVGTPDAQLNGLTQLQVRREFYFKGFR
NNDTRWTTSLKAVYKKDAQGVVSYFGPITYKFQGTMLEGGSTRSFLDDFPIYRYADCLLQLA
MAKVLLGEDPTEEINAVRERAYGSKYFNEHKAEIAYPNDNDPEFYTDNKWMKPDNAGALEAI
LKERLREFMFEGKRWYDIRLLGWDYVHQYSSAEQSRLLWPIDAGTLTNNSALKQTPGYE
111110000000000000000000000000000000000000000000000000000000
000000000000000000000000000000000000000000000000000000000000
000000000111111111100000000000000000000000000000000000000000
000000000000000000000000000000000000000000000000000000000000
000000000000000000000000000000000000000000000000000000000000
000000000000000000000000000000000000000000000000000000000000
000000000000000000000000000000000000000000000000000000000000
000000000000000000000000000000000000000000000000000000000
>DM_train231
SVTVQQPQLTLTAAVIGDGAPANGKTAITVEFTVADFEGKPLAGQEVVITTNNGALPNKITE
KTDANGVARIALTNTTDGVTVVTAEVEGQRQSVDTHFVKGTIAADKSTLAAVPTSIIADGLM
ASTITLELKDTYGDPQAGANVAFDTTLGNMGVITDHNDGTYSAPLTSTTLGVATVTVKVDGA
AFSVPSVTVNFTADPIPDAGRSSFTVSTPDILADGTMSSTLSFVPVDKNGHFISGMQGLSFT
QNGVPVSISPITEQPDSYTATVVGNSVGDVTITPQVDTLILSTLQKKISLFPVPTLTGILVN

GQNFATDKGFPKTIFKNATFQLQMDNDVANNTQYEWSSSFTPNVSVNDQGQVTITYQTYSEV
AVTAKSKKFPSYSVSYRFYPNRWIYDGGRSLVSSLEASRQCQGSDMSAVLESSRATNGTRAP
DGTLWGEWGSLTAYSSDWQSGEYWVKKTSTDFETMNMDTGALQPGPAYLAFPLCALSI
111111100000000000000000000000000000000000000000000000000000
000000000000000000000000000000000000000000000000000000000000
000000000000000000000000000000000000000000000000000000000000
000000000000000000000000000000000000000000000000000000000000
000000000000000000000000000000000000000000000000000000000000
000000000000000000000000000000000000000000000000000000000000
000000000000000000000000000000000000000000000000000000000000
0000000000000000000000000000000000000000000000000000000000
>DM_train234
LLAGASSQRSVARMDGDVIIGALFSVHHQPPAEKVPERKCGEIREQYGIQRVEAMFHTLDKI
NADPVLLPNITLGSEIRDSCWHSSVALEQSIEFIRDSLISIRDEKDGINRCLPDGQSLPPGR
TKKPIAGVIGPGSSSVAIQVQNLLQLFDIPQIAYSATSIDLSDKTLYKYFLRVVPSDTLQAR
AMLDIVKRYNWTYVSAVHTEGNYGESGMDAFKELAAQEGLSIAHSDKIYSNAGEKSFDRLLR
KLRERLPKARVVVCFCEGMTVRGLLSAMRRLGVVGEFSLIGSDGWADRDEVIEGYEVEANGG
ITIKLQSPEVRSFDDYFLKLRLDTNTRNPWFPEFWQHRFQCRLPGHLLENPNFKRICTGNES
LEENYVQDSKMGFVINAIYAMAHGLQNMHHALCPGHVGLCDAMKPIDGSKLLDFLIKSSFIG
VSGEEVWFDEKGDAPGRYDIMNLQYTEANRYDYVHVGTWHEGVLNIDDYKIQMNKSG
111111100000000000000000000000000000000000000000000000000000
000000000000000000000000000000000000001111111111111111111111
110000000000000000000000000000000000000000000000000000000000
000000000000000000000000000000000000000000000000000000000000
000000000000000000000000000000000000000000000000000000000000
000000000000000000000000000000000000000000011111100000000000
000000000000000000000000000000000000000000000000000000000000
000000000000000000000000000000000000000000000000111111111
>DM_train235
MKKRILAVAVTSMLLSASVFAQETVVPSRVGDLKFESDFPTQETMKNMLNEMDFQRATQAYL
WGIPASSIMEWLNVSRNDFKFEEGQMGFFNTLKQKQGIITANFTTPYVIGTWNLEKTGPLII
NLPEAKMAGMMLDVHQRVLSDLSLLGPDKGKGGKYLIVPPGEKYKDLNPKGYYVIRPKTNVV
YGGIRILEPDVDRVVKQVVPNITTQPYADGKLGRKIPVAQVPEIDWTHIPKDGLEYWKTIHQ
IIQENPVEERDRFVMAQLKFLGIEKGKPFNPTEEQKKILLEASKVGRAMAQSNDYTKRFTQP
YWKGTNWKDAISVSLDQRSENYDELDERAAWFYEAITVSRGMKSTIPGFGQRYLVTYQDSDG
NWLSGEHTYKLHVPANVPASNFWSTTVYDENNRLMIINDAGSPDISSRKNLKVNSDGSIDVY
YGPKPVKGYENNWVQTNPGEGWFTYFRFYGPTEKMFDKSWTMGDIELVKLEHHHHHH
111111111111111111111000000000000000000000000000000000000000
000000000000000000000000000000000000000000000000000000000000
000000000000000000000000000000000000000000000000000000000000
000000000000000000000000000000000000000000000000000000000000
000000000000000000000000000000000000000000000000000000000000
000000000000000000000000000000000000000000000000000000000000
000000000000000000000000000000000000000000000000000000000000

000000000000000000000000000000000000000000000000111111111
>DM_train236
RLTELREDIDAILEDPALEGAVSGVVVVDTATGEELYSRDGGEQLLPASNMKLFTAAAALEV
LGADHSFGTEVAAESAPGRRGEVQDLYLVGRGDPTLSAEDLDAMAAEVAASGVRTVRGDLYA
DDTWFDSERLVDDWWPEDEPYAYSAQISALTVAHGERFDTGVTEVSVTPAAEGEPADVDLGA
AEGYAELDNRAVTGAAGSANTLVIDRPVGTNTIAVTGSLPADAAPVTALRTVDEPAALAGHL
FEEALESNGVTVKGDVGLGGVPADWQDAEVLADHTSAELSEILVPFMKFSNNGHAEMLVKSI
GQETAGAGTWDAGLVGVEEALSGLGVDTAGLVLNDGSGLSRGNLVTADTVVDLLGQAGSAPW
AQTWSASLPVAGESDPFVGGTLANRMRGTAAEGVVEAKTGTMSGVSALSGYVPGPEGELAFS
IVNNGHSGPAPLAVQDAIAVRLAEYAGHQAPEGARMMRGPVQGSGELECSWVQAC
00000000000000000000000000000000000000000000000000000000000000
00000000000000000000000000000000000000000000000000000000000000
00000000000000000000000000000000000000000000000000000000000000
00000000000000000000000000000000000000000000000000000000000000
00000000000000000000000000000000000000000000000000000000000000
00000000000000000000000000000000000000000000000000000000000000
00000000000000000000000000000000000000000000000000000000000000
0000000000000000000000000000000000011111111111111111111111
>DM_train237
MNSTNSGPPDSGSATGVVPTPDEIASLLQVEHLLDQRWPETRIDPSLTRISALMDLLGSPQR
SYPSIHIAGTNGKTSVARMVDALVTALHRRTGRTTSPHLQSPVERISIDGKPISPAQYVATY
REIEPLVALIDQQSQASAGKGGPAMSKFEVLTAMAFAAFADAPVDVAVVEVGMGGRWDATNV
INAPVAVITPISIDHVDYLGADIAGIAGEKAGIITRAPDGSPDTVAVIGRQVPKVMEVLLAE
SVRADASVAREDSEFAVLRRQIAVGGQVLQLQGLGGVYSDIYLPLHGEHQAHNAVLALASVE
AFFGAGAQRQLDGDAVRAGFAAVTSPGRLERMRSAPTVFIDAAHNPAGASALAQTLAHEFDF
RFLVGVLSVLGDKDVDGILAALEPVFDSVVVTHNGSPRALDVEALALAAGERFGPDRVRTAE
NLRDAIDVATSLVDDAAADPDVAGDAFSRTGIVITGSVVTAGAARTLFGRDPQ
11111111111111111111000000000000000000111000000000000000000000
00000000000000000000000000000000000000000000000000000000000000
00000000000000000111110000000000000000000000000000000000000000
00000000000000000000000000000000000000110000000000000000000000
00000000000000000000000000000000000000000000000000000000000000
00000111100000000000000000000000000000000000000000000000000000
00000000000000000000000000000000000000000000000000000000000000
00000000000000000000000001111000000000000000000000000000
>DM_train238
MNAPESVQAKPRKRYDAGVMKYKMGYWDGDYVPKDTDLLALFRITPQDGVDPVEAAAAVAGE
SSTATWTVVWTDRLTACDMYRAKAYRVDPVPNNPEQFFCYVAYDLSLFEEGSIANLTASIIG
NVFSFKPIKAARLEDMRFPVAYVKTFAGPSTGIIVERERLDKFGRPLLGATTKPKLGLSGRN
YGRVVYEGLKGGLDFMKDDENINSQPFMHWRDRFLFVMDAVNKASAATGEVKGSYLNVTAGT
MEEMYRRAEFAKSLGSVIIMVDLIVGWTCIQSMSNWCRQNDMILHLHRAGHGTYTRQKNHGV
SFRVIAKWLRLAGVDHMHTGTAVGKLEGDPLTVQGYYNVCRDAYTQTDLTRGLFFDQDWASL
RKVMPVASGGIHAGQMHQLIHLFGDDVVLQFGGGTIGHPQGIQAGATANRVALEAMVLARNE
GRDILNEGPEILRDAARWCGPLRAALDTWGDISFNYTPTDTSDFAPTASVA

11111111111111111111100000000000000000000000000000000000000000
00000000000000000000000000000000000000000000000000000000000000
00000000000000000000000000000000000000000000000000000000000000
00000000000000000000000000000000000000000000000000000000000000
00000000000000000000000000000000000000000000000000000000000000
00000000000000000000000000000000000000000000000000000000000000
00000000000000000000000000000000000000000000000000000000000000
000000000000000000000000000000001111111111111111111
>DM_train239
MSQPDATPFDYILSGGTVIDGTNAPGRLADVGVRGDRIAAVGDLSASSARRRIDVAGKVVSP
GFIDSHTHDDNYLLKHRDMTPKISQGVTTVVTGNCGISLAPLAHANPPAPLDLLDEGGSFRF
ARFSDYLEALRAAPPAVNAACMVGHSTLRAAVMPDLRREATADEIQAMQALADDALASGAIG
ISTGAFYPPAAHASTEEIIEVCRPLITHGGVYATHMRDEGEHIVQALEETFRIGRELDVPVV
ISHHKVMGKLNFGRSKETLALIEAAMASQDVSLDAYPYVAGSTMLKQDRVLLAGRTLITWCK
PYPELSGRDLEEIAAERGKSKYDVVPELQPAGAIYFMMDEPDVQRILAFGPTMIGSDGLPHD
ERPHPRLWGTFPRVLGHYSRDLGLFPLETAVWKMTGLTAAKFGLAERGQVQPGYYADLVVFD
PATVADSATFEHPTERAAGIHSVYVNGAAVWEDQSFTGQHAGRVLNRAGA
11111110000000000000000000000000000000000000000000000000000000
00000000000000000000000000000000000000000000000000000000000000
00000000000000000000000000000000000000000000000000000000000000
00000000000000000000000000000000000000000000000000000000000000
00000000000000000000000000000000000000000000000000000000000000
00000000000000000000000000000000000000000000000000000000000000
00000000000000000000000000000000000000000000000000000000000000
00000000000000000000000000000000000000000000000111
>DM_train241
LLSRRRTRRPGEPPLDLGSIPWLGYALDFGKDAASFLTRMKEKHGDIFTILVGGRYVTVLLD
PHSYDAVVWEPRTRLDFHAYAIFLMERIFDVQLPHYSPSDEKARMKLTLLHRELQALTEAMY
TNLHAVLLGDATEAGSGWHEMGLLDFSYSFLLRAGYLTLYGIEALPRTHESQAQDRVHSADV
FHTFRQLDRLLPKLARGSLSVGDKDHMCSVKSRLWKLLSPARLARRAHRSKWLESYLLHLEE
MGVSEEMQARALVLQLWATQGNMGPAAFWLLLFLLKNPEALAAVRGELESILWQAEQPVSQT
TTLPQKVLDSTPVLDSVLSESLRLTAAPFITREVVVDLAMPMADGREFNLRRGDRLLLFPFL
SPQRDPEIYTDPEVFKYNRFLNPDGSEKKDFYKDGKRLKNYNMPWGAGHNHCLGRSYAVNSI
KQFVFLVLVHLDLELINADVEIPEFDLSRYGFGLMQPEHDVPVRYRIRP
11111000000000000000000000000000000000000000000000000000000000
00000000000000000000000000000000000000000000000000000000000000
00000000000000000000000000000000000000000000000000000000000000
00000000000000000000000000000000000000000000000000000000000000
00000000000000000000000000000000000000000000000000000011111111
11000000000000000000000000000000000000000000000000000000000000
00000000000000000000000000000000000000000000000000000000000000
0000000000000000000000000000000000000000000000000
>DM_train242
MMNDGKQQSTFLFHDYETFGTHPALDRPAQFAAIRTDSEFNVIGEPEVFYCKPADDYLPQPG

AVLITGITPQEARAKGENEAAFAARIHSLFTVPKTCILGYNNVRFDDEVTRNIFYRNFYDPY
AWSWQHDNSRWDLLDVMRACYALRPEGINWPENDDGLPSFRLEHLTKANGIEHSNAHDAMAD
VYATIAMAKLVKTRQPRLFDYLFTHRNKHKLMALIDVPQMKPLVHVSGMFGAWRGNTSWVAP
LAWHPENRNAVIMVDLAGDISPLLELDSDTLRERLYTAKTDLGDNAAVPVKLVHINKCPVLA
QANTLRPEDADRLGINRQHCLDNLKILRENPQVREKVVAIFAEAEPFTPSDNVDAQLYNGFF
SDADRAAMKIVLETEPRNLPALDITFVDKRIEKLLFNYRARNFPGTLDYAEQQRWLEHRRQV
FTPEFLQGYADELQMLVQQYADDKEKVALLKALWQYADEIVEHHHHHH
11111110000000000000000000000000000000000000000000000000000000
00000000000000000000000000000000000000000000000000000000000000
00000000000000000000000000000000111000000000000000000000000000
00000000000000000000000000000000000000000000000000000000000000
00000000000000000000000000000000000000000011110000000000000000
00000000000000000000000000000000000000000000111100000000000000
00000000000000000000000000000000000000000000000000000000000000
000000000000000000000000000000000000000000011111
>DM_train243
MSRLVVVSNRIAPPDEHAASAGGLAVGILGALKAAGGLWFGWSGETGNEDQPLKKVKKGNIT
WASFNLSEQDLDEYYNQFSNAVLWPAFHYRLDLVQFQRPAWDGYLRVNALLADKLLPLLQDD
DIIWIHDYHLLPFAHELRKRGVNNRIGFFLHIPFPTPEIFNALPTYDTLLEQLCDYDLLGFQ
TENDRLAFLDCLSNLTRVTTRSAKSHTAWGKAFRTEVYPIGIEPKEIAKQAAGPLPPKLAQL
KAELKNVQNIFSVERLDYSKGLPERFLAYEALLEKYPQHHGKIRYTQIAPTSRGDVQAYQDI
RHQLENEAGRINGKYGQLGWTPLYYLNQHFDRKLLMKIFRYSDVGLVTPLRDGMNLVAKEYV
AAQDPANPGVLVLSQFAGAANELTSALIVNPYDRDEVAAALDRALTMSLAERISRHAEMLDV
IVKNDINHWQECFISDLKQIVPRSAESQQRDKVATFPKLALEHHHHHH
10000000000000111111000000000000000000000000000000000000000000
00000000000000000000000000000000000000000000000000000000000000
00000000000000000000000000000000000000000000000000000000000000
00000000000000000000000000000000000000000000000000000000000000
00000000000000000000000000000000000000000000000000000000000000
00000000000000000000000000000000000000000000000000000000000000
00000000000000000000000000000000000000000000000000000000000000
000000000000000000000001111111111111111111111111
>DM_train244
TKFYTDAVEAVKDIPNGATVLVGGFGLCGIPENLIGALLKTGVKELTAVSNNAGVDNFGLGL
LLQSKQIKRMISSYVGENAEFERQYLAGELEVELTPQGTLAERIRAGGAGVPAFYTSTGYGT
LVQEGGSPIKYNKDGSIAIASKPREVREFNGQHFILEEAIRGDFALVKAWKADQAGNVTFRK
SARNFNLPMCKAAETTVVEVEEIVDIGSFAPEDIHIPKIYVHRLVKGEKYEKRIERLSVRKE
EDVKTRSGKLGDNVRERIIKRAALEFEDGMYANLGIGIPLLASNFISPNMTVHLQSENGILG
LGPYPLQNEVDADLINAGKETVTVLPGASYFSSDESFAMIRGGHVNLTMLGAMQVSKYGDLA
NWMIPGKLVKGMGGAMDLVSSAKTKVVVTMEHSAKGNAHKIMEKCTLPLTGKQCVNRIITEK
AVFDVDRKKGLTLIELWEGLTVDDIKKSTGCDFAVSPKLIPMQQVTT
00000000000000000000000000000000000000000000000000000000000000
00000000000000000000000000000000000000000000000000000000000000
00000000000000000000000000000000000000000000000000000000000000

000000000000000000000000000000000000000000000000000000000001
111111111111000000000000000000000000000000000000000000000000
000000000000000000000000000000000000000000000000001111111111
111111110000000000000000000011111111111111111111111000000000
000000000000000000000000000000000000000000000
>DM_train246
MARTGAEYIEALKTRPPNLWYKGEKVEDPTTHPVFRGIVRTMAALYDLQHDPRYREVLTYEE
EGKRHGMSFLIPKTKEDLKRRGQAYKLWADQNLGMMGRSPDYLNAVVMAYAASADYFGEFAE
NVRNYYRYLRDQDLATTHALTNPQVNRARPPSGQPDPYIPVGVVKQTEKGIVVRGARMTATF
PLADEVLIFPSTLLQAGSEKYALAFALPTSTPGLHFVCREALVGGDSPFDHPLSSRVEEMDC
LVIFDDVLVPWERVFILGNVELCNNAYAATGALNHMAHQVVALKTAKTEAFLGVAALMAEGI
GADVYGHVQEKIAEIIVYLEAMRAFWTRAEEEAKENAYGLLVPDRGALDGARNLYPRLYPRI
REILEQIGASGLITLPSEKDFKGPLGPFLEKFLQGAALEAKERVALFRLAWDMTLSGFGARQ
ELYERFFFGDPVRMYQTLYNVYNKEPYKERIRAFLKESLKVFEEVQA
100000000000000000000000000000000000000000000000000000000000
000000000000000000000000000000000000000000000000000000000000
000000000000000000000000000000000000000000000000000000000000
000000000000000000000000000000000000000000000000000000000000
000000000000000000000000000000000000000000000000000000000000
000000000000000000000000000000000000000000000000000000000000
000000000000000000000000000000000000000000000000000000000000
000000000000000000000000000000000000000001111
>DM_train247
GSMGATKILMDSTHFNEIRSIIRSRSVAWDALARSEELSEIDASTAKALESILVKKNIGDGL
SSSNNAHSGFKVNGKTLIPLIHLLSTSDNEDCKKSVQNLIAELLSSDKYGDDTVKFFQEDPK
QLEQLFDVSLKGDFQTVLISGFNVVSLLVQNGLHNVKLVEKLLKNNNLINILQNIEQMDTCY
VCIRLLQELAVIPEYRDVIWLHEKKFMPTLFKILQRATDSQLATRIVATNSNHLGIQLQYHS
LLLIWLLTFNPVFANELVQKYLSDFLDLLKLVKITIKEKVSRLCISIILQCCSTRVKQHKKV
IKQLLLLGNALPTVQSLSERKYSDEELRQDISNLKEILENEYQELTSFDEYVAELDSKLLCW
SPPHVDNGFWSDNIDEFKKDNYKIFRQLIELLQAKVRNGDVNAKQEKIIIQVALNDITHVVE
LLPESIDVLDKTGGKADIMELLNHSDSRVKYEALKATQAIIGYTFK
111000000000000000000000000000000000000000000000000001111111
111111111100000000000000000000000000000000000000000000000000
000000000000000000000000000000000000000000000000000000000000
000000000000000000000000000000000000001111111111110000000000
000000000000000000000000000000000000000000000000000000000000
000000000000000000000000000000000000000000000000000000000000
000000000000000000000000000000000000000000000000000000000000
00000000000000000000000000000000000000000000
>DM_train249
GASETQQWKTLEDTRSALMGVYGLTRAALADNNTHWICGDLRKGDFTVYKRSDLQAVSDNEL
NKPYDLLKKVSNWRRFYAVINAASVFMEKAPRTVELDRSYSEQNLKYDIAQVRALRAFAYFY
MVRIWGDVPLVTYSYDNGTFPSMPRTDAQTVLSYAKAELLTAIEDLPYQYGTQTNLYYGSYG
AQWQGKLFNKLSAYSVLAHICAWQGNYAEAETYSAFIIDHASEINAKYTSIADLTSETGLFY

SNASVKGSRILGFNFAHNDNEATQSGHLEQLTLAYPLVQKSYPEIYISKDSLFSIFTNFDDL
RFGIIDTIKYSSYYVQNLNEETPVFSKIKIIQDGSAKDNDFGVFGSSIVFTRLEDITLLRAE
ALCALNRSTEAVSYLNMIRTNRGLREVSFKKDFGNNRESLIAEIFEERRRELMGEGWRWYDL
VRRQKLMKDNEAFLRLISSGGIYWPVSEDIITANSQIEQNEFWK
11110000000000000000000000000000000000000000000000000000000000
00000000000000000000000000000000000000000000000000000000000000
00000000000000000000000000000000000000000000000000000000000000
00000000000000000000000000000000000000000000000000000000000000
00000000000000001111111100000000000000000000000000000000000000
00000000000000000000000000000000111111111000000000000000000000
00000000000000000000000000000000000000000000000000000000000000
00000000000000000000000000000000000000000000
>DM_train250
ARTDNFKLSSLANGLKVATSNTPGHFSALGLYIDAGSRFEGRNLKGCTHILDRLAFKSTEHV
EGRAMAETLELLGGNYQCTSSRENLMYQASVFNQDVGKMLQLMSETVRFPKITEQELQEQKL
SAEYEIDEVWMKPELVLPELLHTAAYSGETLGSPLICPRGLIPSISKYYLLDYRNKFYTPEN
TVAAFVGVPHEKALELTGKYLGDWQSTHPPITKKVAQYTGGESCIPPAPVFGNLPELFHIQI
GFEGLPIDHPDIYALATLQTLLGGGGSFSAGGPGKGMYSRLYTHVLNQYYFVENCVAFNHSY
SDSGIFGISLSCIPQAAPQAVEVIAQQMYNTFANKDLRLTEDEVSRAKNQLKSSLLMNLESK
LVELEDMGRQVLMHGRKIPVNEMISKIEDLKPDDISRVAEMIFTGNVNNAGNGKGRATVVMQ
GDRGSFGDVENVLKAYGLGNSSSSKNDSPKKKGWFHHHHHH
00000000000000000000000000000000000000000000000000000000000000
00000000000000000000000000000000000000000000000000000000000000
00000000000000000000000000000000000000000000000000000000000000
00000000000000000000000000000000000000000000000000000000000000
00000000000000000000000000111111111100000000000000000000000000
00000000000000000000000000000000000000000000000000000000000000
00000000000000000000000000000000000000000000000000000000000000
00000000000000000000011111111111111111111
>DM_train251
GGNLEEMNIDPDNATQTHPKLLLTQICMNAFKRGTDGMYATKKVIQADGESADQYYKWTRGS
FGYYDNLRNVQKMGEEAERVNAPVYTALTKFFRAYYFYELTLRFGDIPYSQALKGEKEEIYT
PEYDAQEDVFAGILQELREADEILANDASVIDGDIIYNGNSTQWRKLINSFRLKVLMTLSNH
TTVGNINIASEFKNIATNSPLMNSLADNGQLVYLDQQGNRYPQFNAQWSGYYMDDTFIQRMR
ERRDPRLFIFSAQTNKGKTEGKPIDDFSSYEGGDPAAPYSDAIIKVSEGTISPINDRFRTDP
IVEPTMLMGYAELQQILAEAVVRGWISGNAQTYYEKGIRASFSFYETHAKDYAGYLNENAVA
QYLKEPLVDFTQASGTEEQIERIIMQKYLVTFYQGNWDSFYEQLRTGYPDFRRPAGTEIPKR
WMYPQGEYDNNGTNVETAITRQFGAGNDKINQATWWQKKS
11111111111111100000000000000000000000000000000000000000000000
00000000000000000000000000000000000000000000000000000011111100
00000000000000000000000000000000000000000000000000000000000000
00000000000000000000000000000000000000000000000000000000000000
00000000000000000000000000000000000000000000000111000000000000
00000000000000000000000000000000000000000000000000000000000000

000000000000000000000000000000000000000000000000000000000000
0000000000000000000000000000000000000000
>DM_train252
GALRSLVLIGHGSHHHGESARATQQVAEALRGRGLAGHLPYDEVLEGYWQQEPGLRQVLRTV
AYSDVTVVPVFLSEGYVTETVLPRELGLGHQGPVPTGGVVRVLGGRRVRYTRPLGAHPGMAD
AIAAQARDTLPEGTDPADVTLLLLAARPGNAALETHAQALRERGQFAGVEVVLESREALTPE
SHAASAVPLSEWPSRVEAGQAVLVPFLTHLGKHAAERLQQALAQAAERFPQAPPLHVGGPVG
EHPAVAEVVLALAAEGREDERGGDIDQAHAEAWAALRHLAERGGRLGEVLLTPYGGLFELRH
TLDEGRATLDLQTVVTPEGLRDLTARDEAGRWRPIRTWRTLPRGWRAVLSPADLRLGLELLY
PAVIEESYAHEHRRLHWTPWMSTARRQTGTLARVQRATPDQVDTVAAQVCASCLRTRLWAGH
TLGQTIFSGVPGGLPCAEACTVLLAAVRDEVGREAMGSGD
000000000000000000000000000000000000000000000000000000000000
000000000000000000000000000000000000000000000000000000000000
000000000000000000000000000000000000000000000000000001111111
111110000000000000000000000000000000000000000000000000000000
000000000000000000000000000000000000000000000000000000000000
000000000000000000000000000000000000000000000000000000000000
000000000000000000000000000000000000000000000000000000000000
0000000000000000000000000000000000111111
>DM_train253
SKDVSSTITTVSASPDGTLNLPAAAPLSIASGRLNQTILETGSQFGGVARWGQESHEFGMRR
LAGTALDGAMRDWFTNECESLGCKVKVDKIGNMFAVYPGKNGGKPTATGSHLDTQPEAGKYD
GILGVLAGLEVLRTFKDNNYVPNYDVCVVVWFNAEGARFARSCTGSSVWSHDLSLEEAYGLM
SVGEDKPESVYDSLKNIGYIGDTPASYKENEIDAHFELHIEQGPILEDENKAIGIVTGVQAY
NWQKVTVHGVGAHAGTTPWRLRKDALLMSSKMIVAASEIAQRHNGLFTCGIIDAKPYSVNII
PGEVSFTLDFRHPSDDVLATMLKEAAAEFDRLIKINDGGALSYESETLQVSPAVNFHEVCIE
CVSRSAFAQFKKDQVRQIWSGAGHDSCQTAPHVPTSMIFIPSKDGLSHNYYEYSSPEEIENG
FKVLLQAIINYDNYRVIRGHQFPGDDDDKHHHHHHHHSGD
111111111111111111111111110000000000000000000000000000000000
000000000000000000000000000000000000000000000000000000000000
000000000000000000000000000000000000000000000000000000000000
000000000000000000000000000000000000000000000000000000000000
000000000000000000000000000000000000000000000000000000000000
000000000000000000000000000000000000000000000000000000000000
000000000000000000000000000000000000000000000000000000000000
0000000000000000000000011111111111111111
>DM_train254
MKTDTSTFLAQQIVRLRRRDQIRRLLQRDKTPLAILFMAAVVGTLTGLVGVAFEKAVSWVQN
MRIGALVQVADHAFLLWPLAFILSALLAMVGYFLVRKFAPEAGGSGIPEIEGALEELRPVRW
WRVLPVKFIGGMGTLGAGMVLGREGPTVQIGGNLGRMVLDVFRMRSAEARHTLLATGAAAGL
SAAFNAPLAGILFIIEEMRPQFRYNLISIKAVFTGVIMSSIVFRIFNGEAPIIEVGKLSDAP
VNTLWLYLILGIIFGVVGPVFNSLVLRTQDMFQRFHGGEIKKWVLMGGAIGGLCGILGLIEP
AAAGGGFNLIPIAAAGNFSVGLLLFIFITRVVTTLLCFSSGAPGGIFAPMLALGTLLGTAFG
MAAAVLFPQYHLEAGTFAIAGMGALMAASVRAPLTGIVLVLEMTDNYQLILPMIITCLGATL

LAQFLGGKPLYSTILARTLAKQDAEQAEKNQNAPADENT
11111111111111111111111111111100000000000000000000000000000000
00000000000000000000000000000000000000000000000000000000000000
00000000000000000000000000000000000000000000000000000000000000
00000000000000000000000000000000000000000000000000000000000000
00000000000000000000000000000000000000000000000000000000000000
00000000000000000000000000000000000000000000000000000000000000
00000000000000000000000000000000000000000000000000000000000000
000000000000000000000000001111111111111
>DM_train256
MGSSSSENLYFQGHMATESPATRRVQVAEHPRLLKLKEMFNSKFGSIPKFYVRAPGRVNIIG
EHIDYCGYSVLPMAVEQDVLIAVEPVKTYALQLANTNPLYPDFSTSANNIQIDKTKPLWHNY
FLCGLKGIQEHFGLSNLTGMNCLVDGNIPPSSGLSSSSALVCCAGLVTLTVLGRNLSKVELA
EICAKSERYIGTEGGGMDQSISFLAEEGTAKLIEFSPLRATDVKLPSGAVFVIANSCVEMNK
AATSHFNIRVMECRLAAKLLAKYKSLQWDKVLRLEEVQAKLGISLEEMLLVTEDALHPEPYN
PEEICRCLGISLEELRTQILSPNTQDVLIFKLYQRAKHVYSEAARVLQFKKICEEAPENMVQ
LLGELMNQSHMSCRDMYECSCPELDQLVDICRKFGAQGSRLTGAGWGGCTVSMVPADKLPSF
LANVHKAYYQRSDGSLAPEKQSLFATKPGGGALVLLEA
11111111111111100000000000000000000000000000000000000000000000
00000000000000000000000000000000000000000000000001100000000000
00000000000000000000000000000000000000000000000000000000000000
00000000000000000000000000000000000000000000000000000000000000
00000000000000000000000000000000000000000000000000000000000000
00000000000000000000000000000000000000000000000000000000000000
00000000000000000000000000000000000000000000000000000000000000
00000000001111111110000000000000000000
>DM_train257
ATLDSWLSNEATVARTAILNNIGADGAWVSGADSGIVVASPSTDNPDYFYTWTRDSGLVLKT
LVDLFRNGDTSLLSTIENYISAQAIVQGISNPSGDLSSGAGLGEPKFNVDETAYTGSWGRPQ
RDGPALRATAMIGFGQWLLDNGYTSTATDIVWPLVRNDLSYVAQYWNQTGYDLWEEVNGSSF
FTIAVQHRALVEGSAFATAVGSSCSWCDSQAPEILCYLQSFWTGSFILANFDSSRSGKDANT
LLGSIHTFDPEAACDDSTFQPCSPRALANHKEVVDSFRSIYTLNDGLSDSEAVAVGRYPEDT
YYNGNPWFLCTLAAAEQLYDALYQWDKQGSLEVTDVSLDFFKALYSDAATGTYSSSSSTYSS
IVDAVKTFADGFVSIVETHAASNGSMSEQYDKSDGEQLSARDLTWSYAALLTANNRRNSVVP
ASWGETSASSVPGTCAATSAIGTYSSVTVTSWPSIV
11111000000000000000000000000000000000000000000000000000000000
00000000000000000000000000000000000000000000000000000000000000
00000000000000000000000000000000000000000000000000000000000000
00000000000000000000000000000000000000000000000000000000000000
00000000000000000000000000000000000000000000000000000000000000
00000000000000000000000000000000000000000000000000000000000000
00000000000000000000000000000000000000000000000000000000000000
000000000000000000000000000001111111
>DM_train258

MPEQYRYTLPVKAGEQRLLGELTGAACATLVAEIAERHAGPVVLIAPDMQNALRLHDEISQF
TDQMVMNLADWETLPYDSFSPHQDIISSRLSTLYQLPTMQRGVLIVPVNTLMQRVCPHSFLH
GHALVMKKGQRLSRDALRTQLDSAGYRHVDQVMEHGEYATRGALLDLFPMGSELPYRLDFFD
DEIDSLRVFDVDSQRTLEEVEAINLLPAHEFPTDKAAIELFRSQWRDTFEVKRDPEHIYQQV
SKGTLPAGIEYWQPLFFSEPLPPLFSYFPANTLLVNTGDLETSAERFQADTLARFENRGVDP
MRPLLPPQSLWLRVDELFSELKNWPRVQLKTEHLPTKAANANLGFQKLPDLAVQAQQKAPLD
ALRKFLETFDGPVVFSVESEGRREALGELLARIKIAPQRIMRLDEASDRGRYLMIGAAEHGF
VDTVRNLALICESDLLGERVARRRQDSRRTINPDTL
111100000000000000000000000000000000000000000000000000000000
000000000000000000000000000000000000000000000000000000000000
000000000000000000000000000000000000000000000000000000000000
000000000000000000000000000000000000000000000000000000000000
000000000000000000000000000000000000000000000000000000000000
000000000000000000000000000000000000000000000000000000000000
000000000000000000000000000000000000000000000000000000000000
000000000000000011111111111111111111
>DM_train259
MLSNELRQTLQKGLHDVNSDWTVPAAIINDPEVHDVERERIFGHAWVFLAHESEIPERGDYV
VRYISEDQFIVCRDEGGEIRGHLNACRHRGMQVCRAEMGNTSHFRCPYHGWTYSNTGSLVGV
PAGKDAYGNQLKKSDWNLRPMPNLASYKGLIFGSLDPHADSLEDYLGDLKFYLDIVLDRSDA
GLQVVGAPQRWVIDANWKLGADNFVGDAYHTMMTHRSMVELGLAPPDPQFALYGEHIHTGHG
HGLGIIGPPPGMPLPEFMGLPENIVEELERRLTPEQVEIFRPTAFIHGTVFPNLSIGNFLMG
KDHLSAPTAFLTLRLWHPLGPDKMEVMSFFLVEKDAPDWFKDESYKSYLRTFGISGGFEQDD
AENWRSITRVMGGQFAKTGELNYQMGRGVLEPDPNWTGPGEAYPLDYAEANQRNFLEYWMQL
MLAESPLRDGNSNGSGTADASTPAAAKSKSPAKAEA
000000000000000000000000000000000000000000000000000000000000
000000000000000000000000000000000000000000000000000000000000
000000000000000000000000000000000000000000000000000000000000
000000000000000000000000000000000000000000000000000000000000
000000000000000000000000000000000000000000000000000000000000
000000000000000000000000000000000000000000000000000000000000
000000000000000000000000000000000000000000000000000000000000
000000011111111111111111111111111111
>DM_train260
GSPGIPMVAAHAAHSQSSAEWIACLDKRPLERSSEDVDIIFTRLKGVKAFEKFHPNLLRQIC
LCGYYENLEKGITLFRQGDIGTNWYAVLAGSLDVKVSETSSHQDAVTICTLGIGTAFGESIL
DNTPRHATIVTRESSELLRIEQEDFKALWEKYRQYMAGLLAPPYGVMETGSNNDRIPDKENV
PSEKILRAGKILRIAILSRAPHMIRDRKYHLKTYRQCCVGTELVDWMIQQTSCVHSRTQAVG
MWQVLLEDGVLNHVDQERHFQDKYLFYRFLDDEREDAPLPTEEEKKECDEELQDTMLLLSQM
GPDAHMRMILRKPPGQRTVDDLEIIYDELLHIKALSHLSTTVKRELAGVLIFESHAKGGTVL
FNQGEEGTSWYIILKGSVNVVIYGKGVVCTLHEGDDFGKLALVNDAPRAASIVLREDNCHFL
RVDKEDFNRILRDVEANTVRLKEHDQDVLVLEKVP
111111111111111111000000000000000000000000000000000000000000
000000000000000000000000000000000000000000000000000000000000

000000000000000000000000000000000000000000000001111111111110
000000000000000000000000000000000000000000000000000000000000
000000000000000000000000000000000000000000000000000000000000
000000000000000000000000000000000000000000000000000000000000
000000000000000000000000000000000000000000000000000000000000
0000000000000000011111111111111111
>DM_train261
MRGSHHHHHHSNFAIILAAGKGTRMKSDLPKVLHKVAGISMLEHVFRSVGAIQPEKTVTVVG
HKAELVEEVLAGQTEFVTQSEQLGTGHAVMMTEPILEGLSGHTLVIAGDTPLITGESLKNLI
DFHINHKNVATILTAETDNPFGYGRIVRNDNAEVLRIVEQKDATDFEKQIKEINTGTYVFDN
ERLFEALKNINTNNAQGEYYITDVIGIFRETGEKVGAYTLKDFDESLGVNDRVALATAESVM
RRRINHKHMVNGVSFVNPEATYIDIDVEIAPEVQIEANVILKGQTKIGAETVLTNGTYVVDS
TIGAGAVITNSMIEESSVADGVTVGPYAHIRPNSSLGAQVHIGNFVEVKGSSIGENTKAGHL
TYIGNCEVGSNVNFGAGTITVNYDGKNKYKTVIGDNVFVGSNSTIIAPVELGDNSLVGAGST
ITKDVPADAIAIGRGRQINKDEYATRLPHHPKNQ
111111111100000000000000000000000000000000000000000000000000
000000000000000000000000000000000000000000000000000000000000
000000000000000000000000001111110000000000000000000000000000
000000000000000000000000000000000000000000000000000000000000
000000000000000000000000000000000000000000000000000000000000
000000000000000000000000000000000000000000000000000000000000
000000000000000000000000000000000000000000000000000000000000
0000000000000000000000111111111111
>DM_train264
MEELNIDFDVFKKRIELLYSKYNEFEGSPNSLLFVLGSSNAENPYQKTTILHNWLLSYEFPA
TLIALVPGKVIIITSSAKAKHLQKAIDLFKDPESKITLELWQRNNKEPELNKKLFDDVIALI
NSAGKTVGIPEKDSYQGKFMTEWNPVWEAAVKENEFNVIDISLGLSKVWEVKDVNEQAFLSV
SSKGSDKFMDLLSNEMVRAVDEELKITNAKLSDKIENKIDDVKFLKQLSPDLSALCPPNYKF
NFDLLDWTYSPIIQSGKKFDLRVSARSTNDQLYGNGCILASCGIRYNNYCSNITRTFLIDPS
EEMANNYDFLLTLQKEIVTNILKPGRTPKEVYESVIEYIEKTKPELVPNFTKNIGSLIGLEF
RDSNFILNVKNDYRKIQRGDCFNISFGFNNLKDSQSANNYALQLADTVQIPLDETEPPRFLT
NYTKAKSQISFYFNNEEEDNNKKKSSPATKV
111100000000000000000000000000000000000000000000000000000000
000000000000000000000000000000110000000000000000000000000000
000000000000000000000000000000000000000000000000000000000000
000000000000000000000000000000000000000000000000000000000000
000000000000000000000000000000000000000000000000000000000000
000000000000000000000000000000000000000000000000000000000000
000000000000000000000000000000000000000000000000000000000000
0000000000000111111111111111111
>DM_train265
VSPKTYKDADFYVAPTQQDVNYDLVDDFGANGNDTSDDSNALQRAINAISRKPNGGTLLIPN
GTYHFLGIQMKSNVHIRVESDVIIKPTWNGDGKNHRLFEVGVNNIVRNFSFQGLGNGFLVDF
KDSRDKNLAVFKLGDVRNYKISNFTIDDNKTIFASILVDVTERNGRLHWSRNGIIERIKQNN

ALFGYGLIQTYGADNILFRNLHSEGGIALRMETDNLLMKNYKQGGIRNIFADNIRCSKGLAA
VMFGPHFMKNGDVQVTNVSSVSCGSAVRSDSGFVELFSPTDEVHTRQSWKQAVESKLGRGCA
QTPYARGNGGTRWAARVTQKDACLDKAKLEYGIEPGSFGTVKVFDVTARFGYNADLKQDQLD
YFSTSNPMCKRVCLPTKEQWSKQGQIYIGPSLAAVIDTTPETSKYDYDVKTFNVKRINFPVN
SHKTIDTNTESSRVCNYYGMSECSSSRWER
000000000000000000000000000000000000000000000000000000000000
000000000000000000000000000000000000000000000000000000000000
000000000000000000000000000000000000000000000000000000000000
000000000000000000000000000000000000000000000000000000000000
000000000000000000000000000000000000111111111111111111111000
001111111111110000000000000000000000000000000000000000000000
000000000000000000000000000000000000000000000000000000000000
000000000000000000000000000000
>DM_train266
MCYASMEMSAPIRFGTEGFRGVIAREFTFATLHRLAEAYGRHLLERGGGLVVVGHDTRFLAD
AFARALSGHLAGMGLKVVLLKGPVPTPLLSFAVRHLKAAGGAMLTASHNPPQYLGVKFKDAT
GGPIAQEEAKAIEALVPEEARALEGAYETLDLREAYFEALKAHLDLKALSGFSGVLYHDSMG
GAGAGFLKGFLRHVGLEIPVRPIREEPHPLFHGVNPEPIPKNLGVTLAVLGPETPPSFAVAT
DGDADRVGVVLPGGVFFNPHQVLTTLALYRFRKGHRGRAVKNFAVTWLLDRLGERLGFGVTT
TPVGFKWIKEEFLKGDCFIGGEESGGVGYPEHLPERDGILTSLLLLESVAATGKDLAEQFKE
VEALTGLTHAYDRLDLPLKAPLDLTPFREPRPLAGLTPKGVDTLDGVKWLYEEAWVLFRASG
TEPVVRIYVEAQSPELVRALLEEARKLVEG
111111100000000000000000000000000000000000000000000000000000
000000000000000000000000000000000000000000000000000000000000
000000000000000000000000000000000000000000000000000000000000
000000000000000000000000000000000000000000000000000000000000
000000000000000000000000000000000000000000000000000000000000
000000000000000000000000000000000000000000000000000000000000
000000000000000111111111111110000000000000000000000000000001
111110000000000000000000000000
>DM_train267
GRPANANSGFYVSGTTLYDANGNPFVMRGINHGHAWYKDQATTAIEGIANTGANTVRIVLSD
GGQWTKDDIQTVRNLISLAEDNNLVAVLEVHDATGYDSIASLNRAVDYWIEMRSALIGKEDT
VIINIANEWFGSWDGAAWADGYKQAIPRLRNAGLNNTLMIDAAGWGQFPQSIHDYGREVFNA
DPQRNTMFSIHMYEYAGGNASQVRTNIDRVLNQDLALVIGEFGHRHTNGDVDESTIMSYSEQ
RGVGWLAWSWKGNGPEWEYLDLSNDWAGNNLTAWGNTIVNGPYGLRETSKLSTVFTGGGSDG
RTSPTTLYDFEESTQGWTGSSLSRGPWTVTEWSSKGNHSLKADIQMSSNSQHYLHVIQNRSL
QQNSRIQATVKHANWGSVGNGMTARLYVKTGHGYTWYSGSFVPINGSSGTTLSLDLSNVQNL
SQVREIGVQFQSESNSSGQTSIYIDNVIVE
000001100000000000000000000000000000000000000000000000000000
000000000000000000000000000000000000000000000000000000000000
000000000000000000000000000000000000000000000000000000000000
000000000000000000000000000000000000000000000000000000000000
000000000000000000000000000000000000000000000000000000111111

111000000000000000000000000000000000000000000000000000000000
000000000000011111110000000000000000000000000000000000000000
000000000000000000000000000000
>DM_train268
MGSDKIHHHHHHMFLGEDYLLTNRAAVRLFNEVKDLPIVDPHNHLDAKDIVENKPWNDIWEV
EGATDHYVWELMRRCGVSEEYITGSRSNKEKWLALAKVFPRFVGNPTYEWIHLDLWRRFNIK
KVISEETAEEIWEETKKKLPEMTPQKLLRDMKVEILCTTDDPVSTLEHHRKAKEAVEGVTIL
PTWRPDRAMNVDKEGWREYVEKMGERYGEDTSTLDGFLNALWKSHEHFKEHGCVASDHALLE
PSVYYVDENRARAVHEKAFSGEKLTQDEINDYKAFMMVQFGKMNQETNWVTQLHIGALRDYR
DSLFKTLGPDSGGDISTNFLRIAEGLRYFLNEFDGKLKIVLYVLDPTHLPTISTIARAFPNV
YVGAPWWFNDSPFGMEMHLKYLASVDLLYNLAGMVTDSRKLLSFGSRTEMFRRVLSNVVGEM
VEKGQIPIKEARELVKHVSYDGPKALFFG
111111111111000000000000000000000000000000000000000000000000
000000000000000000000000000000000000000000000000000000000000
000000000000000000000000000000000000000000000000000000000000
000000000000000000000000000000000000000000000000000000000000
000000000000000000000000000000000000000000000000000000000000
000000000000000000000000000000000000000000000000000000000000
000000000000000000000000000000000000000000000000000000000000
00000000000000000000000000000
>DM_train270
MELITILEKTVSPDRLELEAAQKFLERAAVENLPTFLVELSRVLANPGNSQVARVAAGLQIK
NSLTSKDPDIKAQYQQRWLAIDANARREVKNYVLQTLGTETYRPSSASQCVAGIACAEIPVN
QWPELIPQLVANVTNPNSTEHMKESTLEAIGYICQDIDPEQLQDKSNEILTAIIQGMRKEEP
SNNVKLAATNALLNSLEFTKANFDKESERHFIMQVVCEATQCPDTRVRVAALQNLVKIMSLY
YQYMETYMGPALFAITIEAMKSDIDEVALQGIEFWSNVCDEEMDLAIEASEAAEQGRPPEHT
SKFYAKGALQYLVPILTQTLTKQDENDDDDDWNPCKAAGVCLMLLATCCEDDIVPHVLPFIK
EHIKNPDWRYRDAAVMAFGCILEGPEPSQLKPLVIQAMPTLIELMKDPSVVVRDTAAWTVGR
ICELLPEAAINDVYLAPLLQCLIEGLSA
000000000000000000000000000000000000000000000000000000000000
000000000000000000000000000000000000000000000000000000000000
000000000000000000000000000000000000000000000000000000000000
000000000000000000000000000000000000000000000000000000000000
000000000000000000000000000000000000000000000000000000000000
000000000000000000000000000000000000000000000000000000000000
000000000000000000000000000000000000000000000000000000000000
0000011111111111111111111111
>DM_train271
MAIVPFKRDPTFYPSPKMAMKAPPEDLAYVACLYTGTGINRADFIAVVDVNPKSETYSKIVH
KVELPYINDELHHFGWNACSSALCPNGKPNIERRFLIVPGLRSSRIYIIDTKPNPREPKIIK
VIEPEEVKKVSGYSRLHTVHCGPDAIYISALGNEEGEGPGGILMLDHYSFEPLGKWEIDRGD
QYLAYDFWWNLPNEVLVSSEWAVPNTIEDGLKLEHLKDRYGNRIHFWDLRKRKRIHSLTLGE
ENRMALELRPLHDPTKLMGFINMVVSLKDLSSSIWLWFYEDGKWNAEKVIEIPAEPLEGNLP
EILKPFKAVPPLVTDIDISLDDKFLYLSLWGIGEVRQYDISNPFKPVLTGKVKLGGIFHRAD

HPAGHKLTGAPQMLEISRDGRRVYVTNSLYSTWDNQFYPEGLKGWMVKLNANPSGGLEIDKE
FFVDFGEARSHQVRLSGGDASSDSYCYP
1111111000000000000000000000000000000000000000000000000000000000
0000000000000000000000000000000000000000000000000000000000000000
0000000000000000000000000000000000000000000000000000000000000000
0000000000000000000000000000000000000000000000000000000000000000
0000000000000000000000000000000000000000000000000000000000000000
0000000000000000000000000000000000000000000000000000000000000000
0000000000000000000000000000000000000000000000000000000000000000
0000000000000000000000000000
>DM_train274
MSLDIQSLDIQCEELSDARWAELLPLLQQCQVVRLDDCGLTEARCKDISSALRVNPALAELN
LRSNELGDVGVHCVLQGLQTPSCKIQKLSLQNCCLTGAGCGVLSSTLRTLPTLQELHLSDNL
LGDAGLQLLCEGLLDPQCRLEKLQLEYCSLSAASCEPLASVLRAKPDFKELTVSNNDINEAG
VRVLCQGLKDSPCQLEALKLESCGVTSDNCRDLCGIVASKASLRELALGSNKLGDVGMAELC
PGLLHPSSRLRTLWIWECGITAKGCGDLCRVLRAKESLKELSLAGNELGDEGARLLCETLLE
PGCQLESLWVKSCSFTAACCSHFSSVLAQNRFLLELQISNNRLEDAGVRELCQGLGQPGSVL
RVLWLADCDVSDSSCSSLAATLLANHSLRELDLSNNCLGDAGILQLVESVRQPGCLLEQLVL
YDIYWSEEMEDRLQALEKDKPSLRVIS
1000000000000000000000000000000000000000000000000000000000000000
0000000000000000000000000000000000000000000000000000000000000000
0000000000000000000000000000000000000000000000000000000000000000
0000000000000000000000000000000000000000000000000000000000000000
0000000000000000000000000000000000000000000000000000000000000000
0000000000000000000000000000000000000000000000000000000000000000
0000000000000000000000000000000000000000000000000000000000000000
000000000000000000000000000
>DM_train275
MTQPELTPAQRTEVELLARGRADKSRVLRDLKLPETPEAAHALLLRLGVWDEARTPYADRLR
AALNAVELPVPDFDPAEERLDLTHLPTFAIDDEGNQDPDDAVGVEDLGGGLTRLWVHVADVA
ALVAPDSPLDLEARARGATLYLPDRTIGMLPDELVAKAGLGLHEVSPALSICLDLDPDGNAE
AVDVLLTRVKVQRLAYQEAQARLEAGEEPFVTLARLARASRRLREGEGALSIDLPEVRVKAD
ETGASVFPLPKPEMRTVVQECMTLAGWGTAIFADDNEIPLPFATQDYPTREVAGDTLPAMWA
RRKTLARTRFQPSPGPHHGMGLDLYAQATSPMRRYLDLVVHQQLRAFLAGRDPLSSKVMAAH
IAESQMNADATRQAERLSRRHHTLRFIAAQPERVWDAVVVDRRGAQATLLIPDLAFDVQVNT
PAAPGTALQVQFADIDLPQMRVRARSV
1100000000000000000000000000000000000000000000000000000000000000
0000000000000000000000000000000000000000000000000000000000000000
0000000000000000000000000000000000000000000000000000000000000000
0000000000000000000000000000000000000000000000000000000000000000
0000000000000000000000000000000000000000000000000000000000000000
0000000000000000000000000000000000000000000000000000000000000000
0000000000000000000000000000000000000000000000000000000000000000
000000000000000000000000000

>DM_train276
FSYGAAIPQSTQEKQFSQEFRDGYSILKHYGGNGPYSERVSYGIARDPPTSCEVDQVIMVKR
HGERYPSPSAGKDIEEALAKVYSINTTEYKGDLAFLNDWTYYVPNECYYNAETTSGPYAGLL
DAYNHGNDYKARYGHLWNGETVVPFFSSGYGRVIETARKFGEGFFGYNYSTNAALNIISESE
VMGADSLTPTCDTDNDQTTCDNLTYQLPQFKVAAARLNSQNPGMNLTASDVYNLMVMASFEL
NARPFSNWINAFTQDEWVSFGYVEDLNYYYCAGPGDKNMAAVGAVYANASLTLLNQGPKEAG
SLFFNFAHDTNITPILAALGVLIPNEDLPLDRVAFGNPYSIGNIVPMGGHLTIERLSCQATA
LSDEGTYVRLVLNEAVLPFNDCTSGPGYSCPLANYTSILNKNLPDYTTTCNVSASYPQYLSF
WWNYNTTTELNYRSSPIACQEGDAMD
111111111111100000000000000000000000000000000000000000000000
000000000000000000000000000000000000000000000000000000000000
000000000000000000000000000000000000000000000000000000000000
000000000000011110000000000000000000000000000000000000000000
000000000000000000000000000000000000000000000000000000000000
000000000000000000000000000000000000000000000000000000000000
000000000000000000000000000000000000000000000000000000000000
00000000000000000000000000
>DM_train279
LERGRDYEKNKVCKEFSHLGKEDFTSLSLVLYSRKFPSGTFEQVSQLVKEVVSLTEACCAEG
ADPDCYDTRTSALSAKSCESNSPFPVHPGTAECCTKEGLERKLCMAALKHQPQEFPTYVEPT
NDEICEAFRKDPKEYANQFMWEYSTNYGQAPLSLLVSYTKSYLSMVGSCCTSASPTVCFLKE
RLQLKHLSLLTTLSNRVCSQYAAYGEKKSRLSNLIKLAQKVPTADLEDVLPLAEDITNILSK
CCESASEDCMAKELPEHTVKLCDNLSTKNSKFEDCCQEKTAMDVFVCTYFMPAAQLPELPDV
ELPTNKDVCDPGNTKVMDKYTFELSRRTHLPEVFLSKVLEPTLKSLGECCDVEDSTTCFNAK
GPLLKKELSSFIDKGQELCADYSENTFTEYKKKLAERLKAKLPDATPTELAKLVNKRSDFAS
NCCSINSPPLYCDSEIDAELKNIL
111111111111000000000000000000000000000000000000000000000111
111110000000000000000000000000000011110000000000000000000000
000000000000000000000000000000000000000000000000000000000000
000000000000000000000000000000000000000000000000000000000000
000000000000000000000000000000000000000000000000000000000000
000000000000000000000000000000000000000000000000000000000000
000000000000000000000000000000000000000000000000000000000000
000000000000000000000000
>DM_train280
MSLMLSHNPIVEPFALAHATIVTGDKAGTILRNMTIVVGADGRIEQVAPSIETSIPAEYHYL
DGTGKIVMPGLINAHTHLFSQGKPLNPKLATPKGQRMVATFAHSPLGKPYMAATVKHNATTL
LESGVTTIRTLGDVGYEVVTLRDQIDAGQILGPRILASGPLMAIPEGHGAPLIALTSGTPEE
ARTAVAQNLKAGVNAIKIAATGGVTDAQEIGEAGSPQMSVEQMRAICDEAHQYGVIVGAHAQ
SPEGVRRSLLAGVDTIEHGSVLDDELIGMFRHNPNALRGYSALIPTLSAGLPLTLLGQDVTG
ITDIQLENSKNVVGGMVSGARQAHEAGLMIGVGTDTGMTFVPQYATWRELELLVAYAGFSPA
EALHAATAVNASILGVDAETGSLEVGKSADLLVLNANPLDDLRALEHPALVIAAGHPVWRPG
PKRFADIDALLDEAYAEGHHHHHH
111111110000000000000000000000000000000000000000000000000000

000000000000000000001111111111111111111111111110000000000000
000000000000000000000000000000000000000000000000000000000000
000000000000000000000000000011111100000000000000000000000000
000000000000000000000000000000000000000000000000000000000000
000000000000000000000000000000000000000000000000000000000000
000000000000000000000000000000000000000000000000000000000000
000000000000000011111111
>DM_train282
MGSSGHIEGRHMGKNVLLLGSGFVAQPVIDTLAANDDINVTVACRTLANAQALAKPSGSKAI
SLDVTDDSALDKVLADNDVVISLIPYTFHPNVVKSAIRTKTDVVTSSYISPALRELEPEIVK
AGITVMNEIGLDPGIDHLYAVKTIDEVHRAGGKLKSFLSYCGGLPAPEDSDNPLGYKFSWSS
RGVLLALRNSAKYWKDGKIETVSSEDLMATAKPYFIYPGYAFVCYPNRDSTLFKDLYHIPEA
ETVIRGTLRYQGFPEFVKALVDMGMLKDDANEIFSKPIAWNEALKQYLGAKSTSKEDLIASI
DSKATWKDDEDRERILSGFAWLGLFSDAKITPRGNALDTLCARLEELMQYEDNERDMVVLQH
KFGIEWADGTTETRTSTLVDYGKVGGYSSMAATVGYPVAIATKFVLDGTIKGPGLLAPYSPE
INDPIMKELKDKYGIYLKEKTVA
111111111111000000000000000000000000000000000000000000000000
000000000000000000000000000000000000000000000000000000000000
000000000000000000000000000000000000000000000000000000000000
000000000000000000000000000000000000000000000000000000000000
000000000000000000000000000000000000000000000000000000000000
000000000000000000000000000000000000000000000000000000000000
000000000000000000000000000000000000000000000000000000000000
00000000000000000000000
>DM_train283
MSQTTTNPHVAVLAFPFSTHAAPLLAVVRRLAAAAPHAVFSFFSTSQSNASIFHDSMHTMQC
NIKSYDISDGVPEGYVFAGRPQEDIELFTRAAPESFRQGMVMAVAETGRPVSCLVADAFIWF
AADMAAEMGVAWLPFWTAGPNSLSTHVYIDEIREKIGVSGIQGREDELLNFIPGMSKVRFRD
LQEGIVFGNLNSLFSRMLHRMGQVLPKATAVFINSFEELDDSLTNDLKSKLKTYLNIGPFNL
ITPPPVVPNTTGCLQWLKERKPTSVVYISFGTVTTPPPAEVVALSEALEASRVPFIWSLRDK
ARVHLPEGFLEKTRGYGMVVPWAPQAEVLAHEAVGAFVTHCGWNSLWESVAGGVPLICRPFF
GDQRLNGRMVEDVLEIGVRIEGGVFTKSGLMSCFDQILSQEKGKKLRENLRALRETADRAVG
PKGSSTENFITLVDLVSKPKDV
111111000000000000000000000000000000000000000001111111100
000000000000000000000000000000000000000000000000000000000000
000000000000000000000000000000000000000000000000000000000000
000000000000000000000000000000000000000000000000000000000000
001111111110000000000000000000000000000000000000000000000000
000000000000000000000000000000000000000000000000000000000000
000000000000000000000000000000000000000000000000000000000000
0000000000000000000000
>DM_train284
MNYPAEPFRIKSVETVSMIPRDERLKKMQEAGYNTFLLNSKDIYIDLLTDSGTNAMSDKQWA
GMMMGDEAYAGSENFYHLERTVQELFGFKHIVPTHQGRGAENLLSQLAIKPGQYVAGNMYFT

TTRYHQEKNGAVFVDIVRDEAHDAGLNIAFKGDIDLKKLQKLIDEKGAENIAYICLAVTVNL
AGGQPVSMANMRAVRELTEAHGIKVFYDATRCVENAYFIKEQEQGFENKSIAEIVHEMFSYA
DGCTMSGKKDCLVNIGGFLCMNDDEMFSSAKELVVVYEGMPSYGGLAGRDMEAMAIGLREAM
QYEYIEHRVKQVRYLGDKLKAAGVPIVEPVGGHAVFLDARRFCEHLTQDEFPAQSLAASIYV
ETGVRSMERGIISAGRNNVTGEHHRPKLETVRLTIPRRVYTYAHMDVVADGIIKLYQHKEDI
RGLKFIYEPKQLRFFTARFDYI
100000000000000000000000000000000000000000000000000000000000
000000000000000000000000000000000000000000000000000000000000
000000000000000000000000000000000000000000000000000000000000
000000000000000000000000000000000000000000000000000000000000
000000000000000000000000000000000000000000000000000000000000
000000000000000000000000000000000000000000000000000000000000
000000000000000000000000000000000000000000000000000000000000
0000000000000000000000
>DM_train286
CFAKGTNVLMADGSIECIENIEVGNKVMGKDGRPREVIKLPRGRETMYSVVQKSQHRAHKSD
SSREVPELLKFTCNATHELVVRTPRSVRRLSRTIKGVEYFEVITFEMGQKKAPDGRIVELVK
EVSKSYPISEGPERANELVESYRKASNKAYFEWTIEARDLSLLGSHVRKATYQTYAPILYEN
DHFFDYMQKSKFHLTIEGPKVLAYLLGLWIGDGLSDRATFSVDSRDTSLMERVTEYAEKLNL
CAEYKDRKEPQVAKTVNLYSKVVRGNGIRNNLNTENPLWDAIVGLGFLKDGVKNIPSFLSTD
NIGTRETFLAGLIDSDGYVTDEHGIKATIKTIHTSVRDGLVSLARSLGLVVSVNAEPAKVDM
NGTKHKISYAIYMSGGDVLLNVLSKCAGSKKFRPAPAAAFARECRGFYFELQELKEDDYYGI
TLSDDSDHQFLLANQVVVHN
000000000000000000000000000000000000000000000000000000000000
000000000000000000000000000000000000000000000000000000000000
000000000011111111111111110000000000000000000000000000000000
000000001111100000000000000000000000000000000000000000000000
000001111111111111111111111111110000000000000000000000000000
000000000000000000000000000000000000000000000000000000000000
000000000000000000000000000000000000000000000000000000000000
00000000000000000000
>DM_train287
GAPIHDPDFIGGIGKELIVDNASDVTSFYPSAFQEHLNFIPAPTTGSGCTRIPSFDMSATHY
CYTHNVILSGCRDHSHSHQYLALGVLRTTATGRIFFSTLRSISLDDTQNRKSCSVSATPLGC
DMLCSKVTETEEEDYNSAVPTLMAHGRLGFDGQYHEKDLDVTTLFEDWVANYPGVGGGSFID
GRVWFSVYGGLKPNSPSDTVQEGKYVIYKRYNDTCPDEQDYQIRMAKSSYKPGRFGGKRIQQ
AILSIKVSTSLGEDPVLTVPPNTVTLMGAEGRILTVGTSHFLYQRGSSYFSPALLYPMTVSN
KTATLHSPYTFNAFTRPGSIPCQASARCPNSCVTGVYTDPYPLIFYRNHTLRGVFGTMLDSE
QARLNPASAVFDSTSRSRITRVSSSSTKAAYTTSTCFKVVKTNKTYCLSIAEISNTLFGEFR
IVPLLVEILKNDGVREARSG
000000000000000000000000000000000000000000000000000000000000
000000000000000000000000000000000000000000000000000000000000
000000000000000000000000000000000000000000000000000000000000
000000000000000000000000000000000000000000000000000000000000

000000000000000000000000000000000000000000000000000000000000
000000000000000000000000000000000000000000000000000000000000
000000000000000000000000000000000000000000000000000000000000
00000000000011111111
>DM_train289
LEDKDLRSIQEVRNLIESANKAQKELAAMSQQQIDTIVKAIADAGYGAREKLAKMAHEETGF
GIWQDKVIKNVFASKHVYNYIKDMKTIGMLKEDNEKKVMEVAVPLGVVAGLIPSTNPTSTVI
YKTLISIKAGNSIVFSPHPNALKAILETVRIISEAAEKAGCPKGAISCMTVPTIQGTDQLMK
HKDTAVILATGGSAMVKAAYSSGTPAIGVGPGNGPAFIERSANIPRAVKHILDSKTFDNGTI
CASEQSVVVERVNKEAVIAEFRKQGAHFLSDAEAVQLGKFILRPNGSMNPAIVGKSVQHIAN
LAGLTVPADARVLIAEETKVGAKIPYSREKLAPILAFYTAETWQEACELSMDILYHEGAGHT
LIIHSEDKEIIREFALKKPVSRLLVNTPGALGGIGATTNLVPALTLGCGAVGGSSSSDNIGP
ENLFNIRRIATGVLELEDIR
100000000000000000000000000000000000000000000000000000000000
000000000000000000000000000000000000000000000000000000000000
000000000000000000000000000000000000000000000000000000000000
000000000000000000000000000000000000000000000000000000000000
000000000000000000000000000000000000000000000000000000000000
000000000000000000000000000000000000000000000000000000000000
000000000000000000000000000000000000000000000000000000000000
00000000000000000000
>DM_train290
MASLPILTVLEQSQVSPPPDTLGDKSLQLTFFDFFWLRSPPINNLFFYELPITRSQFTETVV
PNIKHSLSITLKHFYPFVGKLVVYPAPTKKPEICYVEGDSVAVTFAECNLDLNELTGNHPRN
CDKFYDLVPILGESTRLSDCIKIPLFSVQVTLFPNQGIAIGITNHHCLGDASTRFCFLKAWT
SIARSGNNDESFLANGTRPLYDRIIKYPMLDEAYLKRAKVESFNEDYVTQSLAGPSDKLRAT
FILTRAVINQLKDRVLAQLPTLEYVSSFTVACAYIWSCIAKSRNDKLQLFGFPIDRRARMKP
PIPTAYFGNCVGGCAAIAKTNLLIGKEGFITAAKLIGENLHKTLTDYKDGVLKDDMESFNDL
VSEGMPTTMTWVSGTPKLRFYDMDFGWGKPKKLETVSIDHNGAISINSCKESNEDLEIGVCI
SATQMEDFVHIFDDGLKAYL
111110000000000000000000000000000000000000000000000000000000
000000000000000000000000000000000000000000000000000000000000
000000000000000000000000000000000000000000000000000000000000
000000000000000000000000000000000000000000000000000000000000
000000000000000000000000000000000000000000000000000000000000
000000000000000000000000000000000000000000000000000011111000
000000000000000000000000000000000000000000000000000000000000
00000000000000001111
>DM_train291
MKTLSPAVITLLWRQDAAEFYFSRLSHLPWAMLLHSGYADHPYSRFDIVVAEPICTLTTFGK
ETVVSESEKRTTTTDDPLQVLQQVLDRADIRPTHNEDLPFQGGALGLFGYDLGRRFESLPEI
AEQDIVLPDMAVGIYDWALIVDHQRHTVSLLSHNDVNARRAWLESQQFSPQEDFTLTSDWQS
NMTREQYGEKFRQVQEYLHSGDCYQVNLAQRFHATYSGDEWQAFLQLNQANRAPFSAFLRLE
QGAILSLSPERFILCDNSEIQTRPIKGTLPRLPDPQEDSKQAVKLANSAKDRAENLMIVDLM

RNDIGRVAVAGSVKVPELFVVEPFPAVHHLVSTITAQLPEQLHASDLLRAAFPGGSITGAPK
VRAMEIIDELEPQRRNAWCGSIGYLSFCGNMDTSITIRTLTAINGQIFCSAGGGIVADSQEE
AEYQETFDKVNRILKQLEK
000000000000000000000000000000000000000000000000000000000000
000000000000000000000000000000000000000000000000000000000000
000000000000000000000000000000000000000000000000000000000000
000000000000000000000000000000000000000000000000000000000000
000000000000000000000001111111111111111111111100000000000000
000000000000000001111111111110000000000000000000000000000000
000000000000000000000000000000000000000000000000000000000000
0000000000000000000
>DM_train293
MEALNQSQAATGAPVITDLKVVPVAGHDSMLLNLSGAHGPLFTRNILILTDSSGHVGVGEVP
GGEGIRKTLEDARHLLINQSIGNYQSLLNKVRNAFADRDVGGRGLQTFDLRIAVHAVTAVES
ALLDLLGQHLQVPVAALLGEGQQRDAVEMLGYLFYVGDRNKTDLGYRSEHEADNEWFRLRNK
EALTPESVVALAEAAYDRYGFKDFKLKGGVLRGEDEIAAVTALSERFPDARITLDPNGAWSL
KEAVALCRDQHHVLAYAEDPCGAENGYSGREVMAEFRRSTGLRTATNMIATDWRQMGHAIQL
QSVDIPLADPHFWTMQGSVRVAQMCNEWGLTWGSHSNNHFDISLAMFTHVAAAAPGNITAID
THWIWQDGQRLTKEPLQIKGGLVEVPKKPGLGVELDWDALMKAHEVYKSMGLGARDDATAMR
YLVSGWEFNNKRPCMVR
111111111110000000000000000000000000000000000000000000000000
000000000000000000000000000000000001111111111110000000000000
000000000000000000000000000000000000000000000000000000000000
000000000000000000000000000000000000000000000000000000000000
000000000000000000000000000000000000000000000000000000000000
000000000000000000000000000000000000000000000000000000000000
000000000000000000000000000000000000000000000000111111111111
11111111111111111
>DM_train294
IIQVSPAGSMDLLSQLEVERLKKTASSDLYQLYRNCSLAVLNSGSHTDNSKELLDKYKNFDI
TVMRRERGIKLELANPPEHAFVDGQIIKGIQEHLFSVLRDIVYVNMHLADSQRLNLTNATHI
TNLVFGILRNAGALIPGATPNLVVCWGGHSINEVEYQYTREVGHELGLRELNICTGCGPGAM
EGPMKGAAVGHAKQRYSEYRYLGLTEPSIIAAEPPNPIVNELVIMPDIEKRLEAFVRMAHGI
IIFPGGPGTAEELLYILGIMMHPENADQPMPIVLTGPKQSEAYFRSLDKFITDTLGEAARKH
YSIAIDNPAEAARIMSNAMPLVRQHRKDKEDAYSFNWSLKIEPEFQLPFEPNHESMANLDLH
LNQRPEVLAANLRRAFSGVVAGNVKAEGIREIERHGPFEMHGDPVLMKKMDQLLNDFVAQNR
MKLPGGSAYEPCYKIVT
000000000000000000000000000000000000000000001100000000000000
000000000000000000000000000000000000000000000011111111100000
000000000000000000000000000000000000000000000000000000000000
000000000000000000000000000000000000000000000000000000000000
000000000000000000000000000000000000000000000000000000000000
000000000000000000000000000000000000000000000000000000000000
000000000000000000000000000000000000000000000000000000000000

0000000000000001
>DM_train295
MKTWPAPTAPTPVRATVTVPGSKSQTNRALVLAALAAAQGRGASTISGALRSRDTELMLDAL
QTLGLRVDGVGSELTVSGRIEPGPGARVDCGLAGTVLRFVPPLAALGSVPVTFDGDQQARGR
PIAPLLDALRELGVAVDGTGLPFRVRGNGSLAGGTVAIDASASSQFVSGLLLSAASFTDGLT
VQHTGSSLPSAPHIAMTAAMLRQAGVDIDDSTPNRWQVRPGPVAARRWDIEPDLTNAVAFLS
AAVVSGGTVRITGWPRVSVQPADHILAILRQLNAVVIHADSSLEVRGPTGYDGFDVDLRAVG
ELTPSVAALAALASPGSVSRLSGIAHLRGHETDRLAALSTEINRLGGTCRETPDGLVITATP
LRPGIWRAYADHRMAMAGAIIGLRVAGVEVDDIAATTKTLPEFPRLWAEMVGPGQGWGYPQP
RSGQRARRATGQGSGG
000000000000000000000000000000000000000000000000000000000000
000000000000000000000000000000000000000000000000000000000000
000000000000000000000000000000000000000000000000000000000000
000000000000000000000000000000000000000000000000000000000000
000000000000000000000000000000000000000000000000000000000000
000000000000000000000000000000000000000000000000000000000000
000000000000000000000000000000000000000000000000001111111111
1111111111111111
>DM_train296
RQGCEARRAHEHLIRLLLEQGKCPEDGWDESTLELFLHELAVMDSNNFLGNCGVGEREGRVA
SALVARRHYRFIHGIGRSGDISAVQPKAAGSSLLNKITNSLVLNVIKLAGVHSVASCFVVPM
ATGMSLTLCFLTLRHKRPKAKYIIWPRIDQKSCFKSMVTAGFEPVVIENVLEGDELRTDLKA
VEAKIQELGPEHILCLHSTTACFAPRVPDRLEELAVICANYDIPHVVNNAYGLQSSKCMHLI
QQGARVGRIDAFVQSLDKNFMVPVGGAIIAGFNEPFIQDISKMYPGRASASPSLDVLITLLS
LGCSGYRKLLKERKEMFVYLSTQLKKLAEAHNERLLQTPHNPISLAMTLKTIDGHHDKAVTQ
LGSMLFTRQVSGARAVPLGNVQTVSGHTFRGFMSHADNYPCAYLNAAAAIGMKMQDVDLFIK
RLDKCLNIVRKEQTRA
111100000000000000000000000000000000000000000000000000000000
000000000000000001111110000000000000000000000000000000000000
000000000000000000000000000000000000000000000000000000000000
000000000000000000000000000000000000000000000000000000000000
000000000000000000000000000000000000000000000000000000000000
000000000000000000000000000000000000000000000000000000000000
000000000000000000000000000000000000000000000000000000000000
0000000000000001
>DM_train297
SKYVDRVIAEVEKKYADEPEFVQTVEEVLSSLGPVVDAHPEYEEVALLERMVIPERVIEFRV
PWEDDNGKVHVNTGYRVQFNGAIGPYLGGLRFAPSVNLSIMKFLGFEQAFKDSLTTLPMGGA
KGGSDFDPNGKSDREVMRFCQAFMTELYRHIGPDIDVPAGDLGVGAREIGYMYGQYRKIVGG
FYNGVLTGKARSFGGSLVRPEATGYGSVYYVEAVMKHENDTLVGKTVALAGFGNVAWGAAKK
LAELGAKAVTLSGPDGYIYDPEGITTEEKINYMLEMRASGRNKVQDYADKFGVQFFPGEKPW
GQKVDIIMPCATQNDVDLEQAKKIVANNVKYYIEVANMPTTNEALRFLMQQPNMVVAPSKAV
NAGGVLVVGFEMSQNSERLSWTAEEVDSKLHQVMTDIHDGSAAAAERYGLGYNLVAGANIVG
FQKIADAMMAQGIAW

000000000000000000000000000000000000000000000000000000000000
000000000000000000000000000000000000000000000000000000000000
000000000000000000000000000000000000000000000000000000000000
000000111111111111000000000000000000000000000000000000000000
000000000000000000000000000000000000000000000000000000000000
000000000000000000000000000000000000000000000000000000000000
000000000001111111111000000000000000000000000000000000000000
000000000000000
>DM_train298
GAADPVVVPANMEPLTIEGNRFVTLCIMIRTTPWEVSRDVKLHPRDEVDWHTLEGVRALREA
FATNNPNGRLTWGFTMNALEDGRKNYREIRDYVVECQKKYGDEVTYFPGYFPAMYLPRERVN
REMSEAIEIISKMVGNGYRPQSIMGGFLSADNLRYLAEKENIHVAHAVIWSQHNIDGGGADG
SPSYPFYPSTEHFCKPAQGKSDFIDCVNLDGWTMDFICARRSGQTGHGIDGYNSRRGVGPIE
TYKGWGLDLGHREVMHTEAIHFDKGLELNGFGWVANIWEAQMVHEFGKDLICDAMKMWVTGT
KERWPDTHFVTFGEFGELWRKQYKSNDDWNYRFVERGSGLGDSYNNLEIKWFMNKEFRLALL
RDWHTKNSPAYVIDFTRYDLQAHEPADPSPEKPAKDWSLINKINQKALRPQDKPVLIDKLEK
EDQDLIRKYYPELLK
111111111111000000000000000000000000000000000000000000000000
000000000000000000000000000000000000000000000000000000000000
000000000000000000000000000000000000000000000000000011000000
000000000000000000000000000000000000000000000000000000000000
000000000000000000000000000000000000000000000000000000000000
000000000000000000000000000000000000000000000000000000000000
000000000000000000000000000000000000000000000000000000000000
000000000000001
>DM_train300
MGSDKIHHHHHHMTFEEFKDRLFALAKKNGVEVQISFLETREFSLRLANGDLDQYTDAGKFN
VEIKVLKDGKTGTFRTQVLENPEKCFEEALSNLQVKDSEEKEYFFEGGKEYREMETYVGRFE
KLSVKEKMDMAKKAHESAAKDERVVMVPTVMYKDMVIKKIITNTLGLDVESQMDGGFLFAMA
IARDANPRSGSWYELARTPEDLNPEEIGKRAAEEAISLIGSKTIPSGKYPVLMRNTALLDLM
EMFIPMISAENVQKNLSPLKGKLGEQVGNPAVSIKDLPYHPKGLSSTPFDDEGVPTTEKFVL
ENGVLKTFLHNLKTARKEGVEPTGNGFVGGIRPVNLMLMPGEKSFEELLKEMDRGVVITEVE
GMHAGANSISGEFSLFAKGYWVENGEIAHGVEDITISGNFLDLLRKIVLVGNDVKVSQHTIA
PSVLVEVLDVAGK
111111111111000000000000000000000000000000000000000000000000
000000000000000000000000000000000011110000000000000000000000
000000000000000000000000000000000000000000000000000000000000
000000000000000000000000000000000000000000000000000000000000
000000000000000000000000000000000000000000000000000000000000
000000000000000000000000000000000000000000000000000000000000
000000000000000000000000000000000000000000000000000000000000
0000000000011
>DM_train301
ASMKETNQKPYKETYGISHITRHDMLQIPEQQKNEKYQVPEFDSSTIKNISSAKGLDVWDSW

PLQNADGTVANYHGYHIVFALAGDPKNADDTSIYMFYQKVGETSIDSWKNAGRVFKDSDKFD
ANDSILKDQTQEWSGSATFTSDGKIRLFYTDFSGKHYGKQTLTTAQVNVSASDSSLNINGVE
DYKSIFDGDGKTYQNVQQFIDEGNYSSGDNHTLRDPHYVEDKGHKYLVFEANTGTEDGYQGE
ESLFNKAYYGKSTSFFRQESQKLLQSDKKRTAELANGALGMIELNDDYTLKKVMKPLIASNT
VTDEIERANVFKMNGKWYLFTDSRGSKMTIDGITSNDIYMLGYVSNSLTGPYKPLNKTGLVL
KMDLDPNDVTFTYSHFAVPQAKGNNVVITSYMTNRGFYADKQSTFAPSFLLNIKGKKTSVVK
DSILEQGQLTVNK
111111100000000000000000000000000000000000000000000000000000
000000000000000000000000000000000000000000000000000000000000
000000000000000000000000000000000000000000000000000000000000
000000000000000000000000000000000000000000000000000000000000
000000000000000000000000000000000000000000000000000000000000
000000000000000000000000000000000000000000000000000000000000
000000000000000000000000000000000000000000000000000000000000
0000000000000
>DM_train302
MKGRLISSDPYRQQFLVERAVSFSHRQRDCSELISVLPRHALQQIDGFGGSFTEGAGVVFNS
MSEKTKAQFLSLYFSAQEHNYTLARMPIQSCDFSLGNYAYVDSSADLQQGRLSFSRDEAHLI
PLISGALRLNPHMKLMASPWSPPAFMKTNNDMNGGGKLRRECYADWADIIINYLLEYRRHGI
NVQALSVQNEPVAVKTWDSCLYSVEEETAFAVQYLRPRLARQGMDEMEIYIWDHDKDGLVDW
AELAFADEANYKGINGLAFHWYTGDHFSQIQYLAQCLPDKKLLFSEGCVPMESDAGSQIRHW
HTYLHDMIGNFKSGCSGFIDWNLLLNSEGGPNHQGNLCEAPIQYDAQNDVLRRNHSWYGIGH
FCRYVRPGARVMLSSSYDNLLEEVGFVNPDGERVLVVYNRDVQERRCRVLDGDKEIALTLPP
SGASTLLWRQESI
000000000000000000000000000000000000000000000000000000000000
000000000000000000000000000000000000000000000000000000000000
000000000000000000000000000000000000000000000000000000000000
000000000000000000000000000000000000000000000000000000000000
000000000000000000000000000000000000000000000000000000000000
000000000000000000000000000000000000000000000000000000000000
000000000000000000000000000000000000000000000000000000000000
0000000000001
>DM_train303
NAWEVNFDGLVGLTHHYAGLSFGNEASTRHRFQVSNPRLAAKQGLLKMKALADAGFPQAVIP
PHERPFIPVLRQLGFSGSDEQVLEKVARQAPHWLSSVSSASPMWVANAATIAPSADTLDGKV
HLTVANLNNKFHRSLEAPVTESLLKAIFNDEEKFSVHSALPQVALLGDEGAANHNRLGGHYG
EPGMQLFVYGREEGNDTRPSRYPARQTREASEAVARLNQVNPQQVIFAQQNPDVIDQGVFHN
DVIAVSNRQVLFCHQQAFARQSQLLANLRARVNGFMAIEVPATQVSVSDTVSTYLFNSQLLS
RDDGSMMLVLPQECREHAGVWGYLNELLAADNPISELKVFDLRESMANGGGPACLRLRVVLT
EEERRAVNPAVMMNDTLFNALNDWVDRYYRDRLTAADLADPQLLREGREALDVLSQLLNLGS
VYPFQREGGGNG
000000000000000000111111111110000000000000000000000000000000
000000000000000000000000000000000000000000000000000000000000
000000000000000000000000000000000000000000000000000000000000

000000000000000000000000000000000000000000000000000000000000
000000000000000000000000000000000000000000000000000000000000
000000000000000000000000000000000000000000000000000000000000
000000000000000000000000000000000000000000000000000000000000
000000111111
>DM_train304
MSDQPIIRRRQVKTGISDARANNAKTQSQYQPYKDAAWGFINHWYPALFTHELEEDQVQGIQ
ICGVPIVLRRVNGKVFALKDQCLHRGVRLSEKPTCFTKSTISCWYHGFTFDLETGKLVTIVA
NPEDKLIGTTGVTTYPVHEVNGMIFVFVREDDFPDEDVPPLAHDLPFRFPERSEQFPHPLWP
SSPSVLDDNAVVHGMHRTGFGNWRIACENGFDNAHILVHKDNTIVHAMDWVLPLGLLPTSDD
CIAVVEDDDGPKGMMQWLFTDKWAPVLENQELGLKVEGLKGRHYRTSVVLPGVLMVENWPEE
HVVQYEWYVPITDDTHEYWEILVRVCPTDEDRKKFQYRYDHMYKPLCLHGFNDSDLYAREAM
QNFYYDGTGWDDEQLVATDISPITWRKLASRWNRGIAKPGRGVAGAVKDTSLIFKQTADGKR
PGYKVEQIKEDH
111111111111111000000000000000000000000000000000000000000000
000000000000000000000000000000000000000000000000000000000000
000000000000000000000000000000000000000000000000000000000000
000000000000000000000000000000000000000000000000000000000000
000000000000000000000000000000000000000000000000000000000000
000000000000000000000000000000000000000000000000000000000000
000000000000000000000000000000000000000000000000000000000000
000000001111
>DM_train305
MREIVHIQAGQCGNQIGAKFWEVISDEHGIDPTGSYHGDSDLQLERINVYYNEAAGNKYVPR
AILVDLEPGTMDSVRSGPFGQIFRPDNFVFGQSGAGNNWAKGHYTEGAELVDSVLDVVRKES
ESCDCLQGFQLTHSLGGGTGSGMGTLLISKIREEYPDRIMNTFSVVPSPKVSDTVVEPYNAT
LSVHQLVENTDETYCIDNEALYDICFRTLKLTTPTYGDLNHLVSATMSGVTTCLRFPGQLNA
DLRKLAVNMVPFPRLHFFMPGFAPLTSRGSQQYRALTVPELTQQMFDAKNMMAACDPRHGRY
LTVAAVFRGRMSMKEVDEQMLNVQNKNSSYFVEWIPNNVKTAVCDIPPRGLKMSATFIGNST
AIQELFKRISEQFTAMFRRKAFLHWYTGEGMDEMEFTEAESNMNDLVSEYQQYQDATADEQG
EFEEEGEEDEA
000000000000000000000000000000111111111111111110000000000000
000000000000000000000000000000000000000000000000000000000000
000000000000000000000000000000000000000000000000000000000000
000000000000000000000000000000000000000000000000000000000000
000000000000000000000000000000000000000000000000000000000000
000000000000000000000000000000000000000000000000000000000000
000000000000000000000000000000000000000000000000000001111111
11111111111
>DM_train306
GKKAVINDSNTPLHLLQPAYQGTYGDLTPEQVKKDIDRVFAYIDKETPARVVDKNTGKVITD
YTAMGDEAQLERGAFRLASYEWGVTYSALIAAAETTGDKRYTDYVQNRFRFLAEVAPHFKRV
YEEKGKTDSQLLQILTPHALDDAGAVCTAMIKLRLKDESLPVDGLIQNYFDFIINKEYRLAD
GTFARNRPQRNTLWLDDMFMGIPAVAQMSRYDKEAKNKYLAEAVKQFLQFADRMFIPEKGLY

RHGWVESSTDHPAFCWARANGWALLTACELLDVLPEDYPQRPKVMDYFRAHVRGVTALQSGE
GFWHQLLDCNDSYLETSATAIYVYCLAHAINKGWIDAIAYGPVAQLGWHAVAGKINEEGQVE
GTCVGTGMAFDPAFYYYRPVNVYAAHGYGPVLWAGAEMIRLLNTQHPQMNDSAVQYYQEKQK
TTAPIFAVDSE
1111100000000000000000000000000000000000000000000000000000000000
0000000000000000000000000000000000000000000000000000000000000000
0000000000000000000000000000000000000000000000000000000000000000
0000000000000000000000000000000000000000000000000000000000000000
0000000000000000000000000000000000000000000000000000000000000000
0000000000000000000000000000000000000000000000000000000000000000
0000000000000000000000000000000000000000000000000000000000000000
00000000001
>DM_train311
MERVNVVGAGLAGSEAAWTLLRLGVPVRLFEMRPKRMTPAHGTDRFAEIVCSNSLGGEGETN
AKGLLQAEMRRAGSLVMEAADLARVPAGGALAVDREEFSGYITERLTGHPLLEVVREEVREI
PPGITVLATGPLTSEALAEALKRRFGDHFLAYYDAASPIVLYESIDLTKCFRAGRYGQSADY
LNCPMTEEEYRRFHQALLEAQRHTPHDWEKLEFFEACVPVEELARRGYQTLLFGPMKPVGLV
DPRTGKEPFAVVQLRQEDKAGRMWSLVGFQTGLKWPEQKRLIQMIPGLENAEIVRYGVMHRN
TYLNAPRLLGETLEFREAEGLYAAGVLAGVEGYLESAATGFLAGLNAARKALGLPPVAPPEE
SMLGGLVRYLATANPEGFQPMYANWGLVPPVEGRMGKKEKRQAMYRRGLEAFSAWLSGLNPP
LPRPEAALV
0000000000000000000000000000000000000000000000000000000000000000
0000000000000000000000000000000000000000000000000000000000000000
0000000000000000000000000000000000000000000000000000000000111110
0000000000000000000000000000111111100000000000000000000000000000
0000000000000000000000000000000000000000000000000000000000000000
0000000000000000000000000000000000000000000000000000000000000000
0000000000000000000000000000000000000000000000000000000000000000
001111111
>DM_train312
MRIDPFKLAHWMNARKYTAAQTADLAGLPLDDLRRLLGDEANEPDPAAATALAEALSVEPSQ
LAADAHRNLTVVHKSAEEMHASRRPIQRDGIHFYNYYTLAAPEGRVAPVVLDILCPSDRLPA
LNNGHLEPAITVNLGPGDINGRWGEEITPQTWRVLHANHGGDRWITGDSYVEPSYCPHSYSL
AGDAPARIVSYTAQSNISPLMTEANNWSTGAFEEALKALSGKVSAGSVLDLFLARRAHTRTS
AAEAAGVPPADLEAALRSPASETGLTVLRTLGRALGFDYRVLLPADDQHDGVGKTWTTIEDS
RRSRRTFGTYEAASMASAAHLPDLVGSFLRVDADGRGADLIDHAENHYVVTEGRLTLEWDGP
DGPASVELEPDGSAWTGPFVRHRWHGTGTVLKFGSGAHLGYQDWLELTNTFEPAATLRRGRR
DLAGWGYDN
0000000000000000000000000000000000000001111111100000000000000000
0000000000000000000000000000000000000000000000000000000000000000
0000000000000000000000000000000000000000000000000000000000000000
0000000000000000000000000000000000000000000000000000000000000000
0000000000000000000000000000000000000000000000000011000000000000
0000000000000000000000000000000000111111000000000000000000000000

00000000000000000000000000000000000000000000000000000000000000
111111111
>DM_train313
MWEANELSSTNTFSHQAEMDWPSANWWQRYQDAQLNHLIEEALQHSPSLCMAMARLKGAQGF
ARQAGAIRSFDLGLAASATESKVSERYQSATPPDGWNDYGTLTLNFQYDFDFWGKNRAAVVA
ATSELAAAEAESVAARLMISTSIANAYAELARLYANQETVHAALQVRNKTVELLEKRYANGL
ETLGSVSQAKAVAASVEAELLGIQESIQLQKNALAALVGQGPDRAASIEEPHITLTSRYGLP
SEAGVGLLGHRADITAARWRAEAAAQQVGIAQAQFYPDVTLSAFIGYQAFGLDHLFDSGNDA
GAIGPAIYLPLFTGGRLEGQLTSAEARYQEAVAQYNGTLVQALHEIADVVTSSQALQARINK
TEQAVQQAEQALHIATNRYQGGLATYLDVLVAEESLLNNQRALVNLQSRAFSLDLALIHALG
GGFETTES
11111111111111111111000000000000000000000000000000000000000000
00000000000000000000000000000000000000000000000000000000000000
00000000000000000000000000000000000000000000000000000000000000
00000000000000000000000000000000000000000000000000000000000111
11110000000000000000000000000000000000000000000000000000000000
00000000000000000000000000000000000000000000000000000000000000
00000000000000000000000000000000000000000000000000000000000000
00001111
>DM_train315
AGTHDYSTALKDSIIFFDANKCGPQAGENNVFDWRGACHTTDGSDVGVDLTGGYHDAGDHVK
FGLPQGYSAAILGWSLYEFKESFDATGNTTKMLQQLKYFTDYFLKSHPNSTTFYYQVGEGNA
DHTYWGAPEEQTGQRPSLYKADPSSPASDILSETSAALTLMYLNYKNIDSAYATKCLNAAKE
LYAMGKANQGVGNGQSFYQATSFGDDLAWAATWLYTATNDSTYITDAEQFITLGNTMNENKM
QDKWTMCWDDMYVPAALRLAQITGKQIYKDAIEFNFNYWKTQVTTTPGGLKWLSNWGVLRYA
AAESMVMLVYCKQNPDQSLLDLAKKQVDYILGDNPANMSYIIGYGSNWCIHPHHRAANGYTY
ANGDNAKPAKHLLTGALVGGPDQNDKFLDDANQYQYTEVALDYNAGLVGVLAGAIKFFGGTI
VNPPVKK
00000000000000000000000000000000000000000000000000000000000000
00000000000000000000000000000000000000000000000000000000000000
00000000000000000000000000000000000000000000000000000000000000
00000000000000000000000000000000000000000000000000000111111000
00000000000000000000000000000000000000000000000000000000000000
00000000000000000000000000000000000000000000000000000000000000
00000000000000000000000000000000000000000000000000000000000111
1111111
>DM_train316
PYFDNISTIAYEGPASKNPLAFKFYNPEEKVGDKTMEEHLRFSVAYWHTFTGDGSDPFGAGN
MIRPWNKYSGMDLAKARVEAAFEFFEKLNIPFFCFHDVDIAPEGETLKETYKNLDIIVDMIE
EYMKTSKTKLLWNTANLFTHPRFVHGAATSCNADVFAYAAAKVKKGLEIAKRLGAENYVFWG
GREGYETLLNTDMKLELDNLARFLHMAVDYAKEIGFDGQFLIEPKPKEPTKHQYDFDVATAL
AFLQTYGLKDYFKFNIEANHATLAGHTFEHELRVARIHGMLGSVDANQGDMLLGWDTDEFPT
DLYSTTLAMYEILKNGGLGRGGLNFDAKVRRGSFEPEDLFYAHIAGMDSFAVGLKVAHRLIE
DRVFDEFIEERYKSYTEGIGREIVEGTADFHKLEAHALQLGEIQNQSGRQERLKTLLNQYLL

EVCAAR
000000000000000000000000000000000000000000000000000000000000
000000000000000000000000000000000000000000000000000000000000
000000000000000000000000000000000000000000000000000000000000
000000000000000000000000000000000000000000000000000000000000
000000000000000000000000000000000000000000000000000000000000
000000000000000000000000000000000000000000000000000000000000
000000000000000000000000000000000000000000000000000000000000
000111
>DM_train317
MLSTVFRRTMATGRHFIAVCQMTSDNDLEKNFQAAKNMIERAGEKKCEMVFLPECFDFIGLN
KNEQIDLAMATDCEYMEKYRELARKHNIWLSLGGLHHKDPSDAAHPWNTHLIIDSDGVTRAE
YNKLHLFDLEIPGKVRLMESEFSKAGTEMIPPVDTPIGRLGLSICYDVRFPELSLWNRKRGA
QLLSFPSAFTLNTGLAHWETLLRARAIENQCYVVAAAQTGAHNPKRQSYGHSMVVDPWGAVV
AQCSERVDMCFAEIDLSYVDTLREMQPVFSHRRSDLYTLHINEKSSETGGLKFARFNIPADH
IFYSTPHSFVFVNLKPVTDGHVLVSPKRVVPRLTDLTDAETADLFIVAKKVQAMLEKHHNVT
STTICVQDGKDAGQTVPHVHIHILPRRAGDFGDNEIYQKLASHDKEPERKPRSNEQMAEEAV
VYRNLM
111111111000000000000000000000000000000000000000000000000000
000000000000000000000000000000000000000000000000000000000000
000000000000000000000000000000000000000000000000000000000000
000000000000000000000000000000000000000000000000000000000000
000000000000000000000000000000000000000000000000000000000000
000000000000000000000000000000000000000000000000000000000000
000000000000000000000000000000111111111111111111110000000000
000000
>DM_train319
MPKLVTWMNNQRVGELTKLANGAHTFKYAPEWLASRYARPLSLSLPLQRGNITSDAVFNFFD
NLLPDSPIVRDRIVKRYHAKSRQPFDLLSEIGRDSVGAVTLIPEDETVTHPIMAWEKLTEAR
LEEVLTAYKADIPLGMIREENDFRISVAGAQEKTALLRIGNDWCIPKGITPTTHIIKLPIGE
IRQPNATLDLSQSVDNEYYCLLLAKELGLNVPDAEIIKAGNVRALAVERFDRRWNAERTVLL
RLPQEDMCQTFGLPSSVKYESDGGPGIARIMAFLMGSSEALKDRYDFMKFQVFQWLIGATQG
HAKNFSVFIQAGGSYRLTPFYDIISAFPVLGGTGIHISDLKLAMGLNASKGKKTAIDKIYPR
HFLATAKVLRFPEVQMHEILSDFARMIPAALDNVKTSLPTDFPENVVTAVESNVLRLHGRLS
REYGSK
100000000000000000000000000000000000000000000000000000000000
000000000000000000000000000000000000000000000001111000000000
000000000011111111111000000000000000000000000000000000000011
111111111000000000000000000000000000000000000000000000000000
000000000000000000000000000000000000000000000000000000000000
000000000000000000000000000000000000000000000000000000000000
000000000000000000000000000000000000000000000000000000000000
000111
>DM_train320

MKPLSSPLQQYWQTVVERLPEPLAEESLSAQAKSVLTFSDFVQDSVIAHPEWLTELESQPPQ
ADEWQHYAAWLQEALCNVSDEAGLMRELRLFRRRIMVRIAWAQTLALVTEESILQQLSYLAE
TLIVAARDWLYDACCREWGTPCNAQGEAQPLLILGMGKLGGGELNFSSDIDLIFAWPEHGCT
QGGRRELDNAQFFTRMGQRLIKVLDQPTQDGFVYRVDMRLRPFGESGPLVLSFAALEDYYQE
QGRDWERYAMVKARIMGDSEGVYANELRAMLRPFVFRRYIDFSVIQSLRNMKGMIAREVRRR
GLTDNIKLGAGGIREIEFIVQVFQLIRGGREPSLQSRSLLPTLSAIAELHLLSENDAEQLRV
AYLFLRRLENLLQSINDEQTQTLPSDELNRARLAWAMDFADWPQLTGALTAHMTNVRRVFNE
LIGDDE
000000000000000000000000000000000000000000000000000000000000
000000000000000000000000000000000000000000000000000000000000
000000000000000000000000000000000000000000000000000000001111
111100000000000000000000000000000000000000000000000000000000
000000000000000000000000000000000000000000000000000000000000
000000000000000000000000000000000000000000000000000000000000
000000000000000000000000000000000000000000000000000000000000
000111
>DM_train322
DIAVVGIGYVGLVSATCFAELGANVRCIDTDRNKIEQLNSGTIPIYEPGLEKMIARNVKAGR
LRFGTEIEQAVPEADIIFIAVGTPAGEDGSADMSYVLDAARSIGRAMSRYILIVTKSTVPVG
SYRLIRKAIQEELDKREVLIDFDIASNPEFLKEGNAIDDFMKPDRVVVGVDSDRARELITSL
YKPMLLNNFRVLFMDIASAEMTKYAANAMLATRISFMNDVANLCERVGADVSMVRLGIGSDS
RIGSKFLYPGCGYGGSCFPKDVKALIRTAEDNGYRMEVLEAVERVNEKQKSILFDKFSTYYK
GNVQGRCVAIWGLSFKPGTDDMREAPSLVLIEKLLEVGCRVRVYDPVAMKEAQKRLGDKVEY
TTDMYDAVRGAEALFHVTEWKEFRMPDWSALSQAMAASLVIDGRNVYELPADSDFTLLNIGN
SAIES
000000000000000000000000000000000000000111100000000000000000
000000000000000000000000110000000000000000000000000000000000
000000000000000000000000000000000000000000000000000000000000
000011111000000000000000000000000000000000000000000000000000
000000000000000000000000000000000000000000000000000000000000
000000000000000000000000000000000000000000000000000000000000
000000000000000000000000000000000000000000000000111000000001
11111
>DM_train323
GRDAPAPAASQPSGCGKHNSPERKVYMDYNATTPLEPEVIQAMTKAMWEAWGNPSSPYSAGR
KAKDIINAARESLAKMIGGKPQDIIFTSGGTESNNLVIHSVVKHFHANQTSKGHTGGHHSPV
KGAKPHFITSSVEHDSIRLPLEHLVEEQVAAVTFVPVSKVSGQTEVDDILAAVRPTTRLVTI
MLANNETGIVMPVPEISQRIKALNQERVAAGLPPILVHTDAAQALGKQRVDVEDLGVDFLTI
VGHKFYGPRIGALYIRGLGEFTPLYPMLFGGGQERNFRPGTENTPMIAGLGKAAELVTQNCE
AYEAHMRDVRDYLEERLEAEFGQKRIHLNSQFPGTQRLPNTCNFSIRGPRLQGHVVLAQCRV
LMASVGAACHSDHGDQPSPVLLSYGVPFDVARNALRLSVGRSTTRAEVDLVVQDLKQAVAQL
EDQA
111111111111111111111110000000000000000000000000000000000000
000000000000000000000000000000000000000000000000011111111111

100000000000000000000000000000000000000000000000000000000000
000000000000000000000000000000000000000000000000000000000000
000000000000000000000000000000000000000000000000000000000000
000000000000000000000000000000000000000000000000000000000000
000000111111111000000000000000000000000000000000000000000000
0011
>DM_train324
MAAGTFTAQNKVRPGVYINFKSEPQAAGTLGERGIVSMPLILSWGEPGKMITIEAGDDVFPK
LGYSIMDAQLRLINEALKRAKTLLLYRLNAGTKAAVTVGNLTVTAKWGGARGNDITLVIQEN
IDDETKFDVSTLVDGAELDKQTVSDIAGLAANDWVIFSGTGALTETAGAPLINGSDGAVTNQ
AYIDYLAAVEIFDFNTIALPSTDDALKATFTAFAKRLRDDEGKKIQVVLENYPAADYEGVIS
VKNGVVLADGTILTAAQATAWVAGATAGARVNESLTYQGYDEAVDVAPRYTNAQIIAALQAG
EFLFTASDNQALVEQDINTLTSFTADKGKQFAKNRVIRVLDGINNDFVRIFSKFYIGKVSNN
ADGRNLLKSECINYMNTLQDIDAIKNFDGQTDLTVQSGNDVDAVYIEAYAWPVDSIEKIYVR
VRIK
111111111111110000000000011110000000000000000000000000000000
000000000000000000000000000000000000000000000000000000000000
000000000000000000000000000000000000000000000000000000000000
000000000000000000000000000000000000000000000000000000000000
000000000000000000000000000000000000000000000000000000000000
000000000000000000000000000000000000000000000000000001111000
000000000000000000000000000000000000011000000000000000000000
0000
>DM_train325
GAMDPSQYASSSSWTSFLKSIASFNGDLSSLSAPPFILSPISLTEFSQYWAEHPELFLEPSF
INDDNYKEHCLIDPEVESPELARMLAVTKWFISTLKSQYCSRNESLGSEKKPLNPFLGELFV
GKWENKEHPEFGETVLLSEQVSHHPPVTAFSIFNDKNKVKLQGYNQIKASFTKSLMLTVKQF
GHTMLDIKDESYLVTPPPLHIEGILVASPFVELEGKSYIQSSTGLLCVIEFSGRGYFSGKKN
SFKARIYKDSKDSKDKEKALYTISGQWSGSSKIIKANKKEESRLFYDAARIPAEHLNVKPLE
EQHPLESRKAWYDVAGAIKLGDFNLIAKTKTELEETQRELRKEEEAKGISWQRRWFKDFDYS
VTPEEGALVPEKDDTFLKLASALNLSTKNAPSGTLVGDKEDRKEDLSSIHWRFQRELWDEEK
EIVL
110000000000000000000000000000000000000000000000000000000000
000000000000000000000000000000000000000000000000000000000000
000000000000000000000000000000000000000000000000000000000000
000000000000000000000000000000000000000000000000000000000000
000000000000000000000000000000000000000000000000000000000000
000000000000000000000000000000000000000000000000000000000000
000000000000000000000000000000000000000000000000000000000000
0000
>DM_train326
STLHGISHIFSYERLSLKRVVWALCFMGSLALLALVCTNRIQYYFLYPHVTKLDEVAATRLT
FPAVTFCNLNEFRFSRVTKNDLYHAGELLALLNNRYEIPDTQTADEKQLEILQDKANFRNFK
PKPFNMLEFYDRAGHDIREMLLSCFFRGEQCSPEDFKVVFTRYGKCYTFNAGQDGKPRLITM

KGGTGNGLEIMLDIQQDEYLPVWGETDETSFEAGIKVQIHSQDEPPLIDQLGFGVAPGFQTF
VSCQEQRLIYLPPPWGDCKATTGDSEFYDTYSITACRIDCETRYLVENCNCRMVHMPGDAPY
CTPEQYKECADPALDFLVEKDNEYCVCEMPCNVTRYGKELSMVKIPSKASAKYLAKKYNKSE
QYIGENILVLDIFFEALNYETIEQKKAYEVAGLLGDIGGQMGLFIGASILTVLELFDYAYEV
IKHR
111111111111111000000000000000000000000000000000000000000000
000000000000000000000000000000000000000000000000000000000000
000000000000000000000000000000000000000000000000000000000000
000000000000000000000000000000000000000000000000000000000000
000000000000000000000000000000000000000000000000000000000000
000000000000000000000000000000000000000000000000000000000000
000000000000000000000000000000000000000000000000000000111111
1111
>DM_train327
MSLPGTGSGTSSGGGNTQGQDVYIIPRPFSNFGKKLSTYTKSHKFMIFGLANNVIGPTGTGT
TAVNRLLTTCLAEIPWQKLPLYMNQSEFDLLPPGSRVVECNVKVIFRTNRIAFETSSTVTKQ
ATLNQISNVQTAIGLNKLGWGINRAFTAFQSDQPMIPTATTAPKYEPVTGDTGYRGMIADYY
GADSTNDTAFGNAGNYPHHQVSSFTFLQNYYCMYQQTNQGTGGWPCLAEHLQQFDSKTVNNQ
CLIDVTYKPKMGLIKSPLNYKIIGQPTVKGTISVGDNLVNMRGAVVTNPPEATQNVAESTHN
LTRNFPADLFNIYSDIEKSQVLHKGPWGHENPQIQPSVHIGIQAVPALTTGALLINSSPLNS
WTDSMGYIDVMSSCTVMEAQPTHFPFSTEANTNPGNTIYRINLTPNSLTSAFNGLYGNGATL
GNV
111111111111111111110000000000000000000000000000000000000000
000000000000000000000000000000000000000000000000000000000000
000000000000000000000000000000000000000000000000000000000000
000000000000000000000000000000000000000000000000000000000000
000000000000000000000000000000000000000000000000000000000000
000000000000000000000000000000000000000000000000000000000000
000000000000000000000000000000000000000000000000000000000000
000
>DM_train328
ADYQGKNVVIIGLGLTGLSCVDFFLARGVTPRVMDTRMTPPGLDKLPEAVERHTGSLNDEWL
MAADLIVASPGIALAHPSLSAAADAGIEIVGDIELFCREAQAPIVAITGSNGKSTVTTLVGE
MAKAAGVNVGVGGNIGLPALMLLDDECELYVLELSSFQLETTSSLQAVAATILNVTEDHMDR
YPFGLQQYRAAKLRIYENAKVCVVNADDALTMPIRGADERCVSFGVNMGDYHLNHQQGETWL
RVKGEKVLNVKEMKLSGQHNYTNALAALALADAAGLPRASSLKALTTFTGLPHRFEVVLEHN
GVRWINDSKATNVGSTEAALNGLHVDGTLHLLLGGDGKSADFSPLARYLNGDNVRLYCFGRD
GAQLAALRPEVAEQTETMEQAMRLLAPRVQPGDMVLLSPACASLDQFKNFEQRGNEFARLAK
ELG
000000000000000000000000000000000000000000000000000000000000
000000000000000000000000000000000000000000000000000000000000
000000000000000000000000000000000000000000000000000000001111
110000000000000000000000000000000000000000000000000000000000
000000000000000000000000000000000000000000000000000000000000

000000000000000000000000000000000000000000000000000000000000
000000000000000000000000000000000000000000000000000000000000
000
>DM_train329
LNSVGQGEFGGAPFKRFLRGTRIVSGGKLKRMTREKAKQVTVAGVPMPRDAEPRHLLVNGAT
GTGKSVLLRELAYTGLLRGDRMVIVDPNGDMLSKFGRDKDIILNPYDQRTKGWSFFNEIRND
YDWQRYALSVVPRGKTDEAEEWASYGRLLLRETAKKLALIGTPSMRELFHWTTIATFDDLRG
FLEGTLAESLFAGSNEASKALTSARFVLSDKLPEHVTMPDGDFSIRSWLEDPNGGNLFITWR
EDMGPALRPLISAWVDVVCTSILSLPEEPKRRLWLFIDELASLEKLASLADALTKGRKAGLR
VVAGLQSTSQLDDVYGVKEAQTLRASFRSLVVLGGSRTDPKTNEDMSLSLGEHEVERDRYSK
NTGKHHSTGRALERVRERVVMPAEIANLPDLTAYVGFAGNRPIAKVPLEIKQFANRQPAFVE
GTI
111111100000000000000000000000000000000000000000000000000000
000000000000000000000000000000000000000000000000000000000000
000000000000000000000000000000000000000000000000000000000000
000000000000000000000000000000000000000000000000000000000000
000000000000000000000000000000000000000000000000000000000000
000000000000000000000000000000000000000000000000000000001111
111111111111000000000000000000000000000000000000000000000000
111
>DM_train330
MASSSSFTYYCPPSSSPVWSEPLYSLRPEHARERLQDDSVETVTSIEQAKVEEKIQEVFSSY
KFNHLVPRLVLQREKHFHYLKRGLRQLTDAYECLDASRPWLCYWILHSLELLDEPIPQIVAT
DVCQFLELCQSPDGGFGGGPGQYPHLAPTYAAVNALCIIGTEEAYNVINREKLLQYLYSLKQ
PDGSFLMHVGGEVDVRSAYCAASVASLTNIITPDLFEGTAEWIARCQNWEGGIGGVPGMEAH
GGYTFCGLAALVILKKERSLNLKSLLQWVTSRQMRFEGGFQGRCNKLVDGCYSFWQAGLLPL
LHRALHAQGDPALSMSHWMFHQQALQEYILMCCQCPAGGLLDKPGKSRDFYHTCYCLSGLSI
AQHFGSGAMLHDVVMGVPENVLQPTHPVYNIGPDKVIQATTHFLQKPVPGFEECEDAVTSDP
ATD
111111111111111111111000000000000000000000000000000000000000
000000000000000000000000000000000000000000000000000000000000
000000000000000000000000000000000000000000000000000000000000
000000000000000000000000000000000000000000000000000000000000
000000000000000000000000000000000000000000000000000000000000
000000000000000000000000000000000000000000000000000000000000
000000000000000000000000000000000000000000000001111111111111
111
>DM_train331
MAGGLSQLVAYGAQDVYLTGNPQITFFKTVYRRYTNFAIESIQQTINGSVGFGNKVSTQISR
NGDLITDIVVEFVLTKGGNGGTTYYPAEELLQDVELEIGGQRIDKHYNDWFRTYDALFRMND
DRYNYRRMTDWVNNELVGAQKRFYVPLIFFFNQTPGLALPLIALQYHEVKLYFTLASQVQGV
NYNGSSAIAGAAQPTMSVWVDYIFLDTQERTRFAQLPHEYLIEQLQFTGSETATPSATTQAS
QNIRLNFNHPTKYLAWNFNNPTNYGQYTALANIPGACSGAGTAAATVTTPDYGNTGTYNEQL
AVLDSAKIQLNGQDRFATRKGSYFNKVQPYQSIGGVTPAGVYLYSFALKPAGRQPSGTCNFS

RIDNATLSLTYKTCSIDATSPAAVLGNTETVTANTATLLTALNIYAKNYNVLRIMSGMGGLA
YAN
11111111111111111111111100000000000000000000000000000000000000
00000000000000000000000000000000000000000000000000000000000000
00000000000000000000000000000000000000000000000000000000000000
00000000000000000000000000000000000000000000000000000000000000
00000000000000000000000000000000000000000000000000000000000000
00000000000000000000000000000000000000000000000000000000000000
00000000000000000000000000000000000000000000000000000000000000
000
>DM_train332
MSLSINSREVLAEKVKNAVNNQPVTDMHTHLFSPNFGEILLWDIDELLTYHYLVAEVMRWTD
VSIEAFWAMSKREQADLIWEELFIKRSPVSEACRGVLTCLQGLGLDPATRDLQVYREYFAKK
TSEEQVDTVLQLANVSDVVMTNDPFDDNERISWLEGKQPDSRFHAALRLDPLLNEYEQTKHR
LRDWGYKVNDEWNEGSIQEVKRFLTDWIERMDPVYMAVSLPPTFSFPEESNRGRIIRDCLLP
VAEKHNIPFAMMIGVKKRVHPALGDAGDFVGKASMDGVEHLLREYPNNKFLVTMLSRENQHE
LVVLARKFSNLMIFGCWWFMNNPEIINEMTRMRMEMLGTSFIPQHSDARVLEQLIYKWHHSK
SIIAEVLIDKYDDILQAGWEVTEEEIKRDVADLFSRNFWRFVGRNDHVTSVKVEQQTEGHHH
HHH
11000000000000000000000000000000000000000000000000000000000000
00000000000000000000000000000000000000000000000000000000000000
00000000000000000000000000000000000000000000000000000000000000
00000000000000000000000000000000000000000000000000000000000000
00000000000000000000000000000000000000000000000000000000000000
00000000000000000000000000000000000000000000000000000000000000
00000000000000000000000000000000000000000000111111111111111111
111
>DM_train333
MLLAQKPFWQRHLAYPHINLDTVAHSLRLTGPLDTTLLLRALHLTVSEIDLFRARFSAQGEL
YWHPFSPPIDYQDLSIHLEAEPLAWRQIEQDLQRSSTLIDAPITSHQVYRLSHSEHLIYTRA
HHIVLDGYGMMLFEQRLSQHYQSLLSGQTPTAAFKPYQSYLEEEAAYLTSHRYWQDKQFWQG
YLREAPDLTLTSATYDPQLSHAVSLSYTLNSQLNHLLLKLANANQIGWPDALVALCALYLES
AEPDAPWLWLPFMNRWGSVAANVPGLMVNSLPLLRLSAQQTSLGNYLKQSGQAIRSLYLHGR
YRIEQIEQDQGLNAEQSYFMSPFINILPFESPHFADCQTELKVLASGSAEGINFTFRGSPQH
ELCLDITADLASYPQSHWQSHCERFPRFFEQLLARFQQVEQDVARLLAEPAALAATTSTRAI
AS
00000000000000000000000000000000000000000000000000000000000000
00000000000000000000000000000000000000000000000000000000000000
00000000000000000000000000000000000000000000000000000000000000
00000000000000000000000000000000000000000000000000000000000000
00000000000000000000000000000000000000000000000000000000000000
00000000000000000000000000000000000000000000000000000000000000
00000000000000000000000000000000000000000000000000001111111111
11

>DM_train334
MSASQKEGKLSTATISVDGKSAEMPVLSGTLGPDVIDIRKLPAQLGVFTFDPGYGETAACNS
KITFIDGDKGVLLHRGYPIAQLAENASYEEVIYLLLNGELPNKAQYDTFTNTLTNHTLLHEQ
IRNFFNGFRRDAHPMAILCGTVGALSAFYPDANDIAIPANRDLAAMRLIAKIPTIAAWAYKY
TQGEAFIYPRNDLNYAENFLSMMFARMSEPYKVNPVLARAMNRILILHADHEQNASTSTVRL
AGSTGANPFACIAAGIAALWGPAHGGANEAVLKMLARIGKKENIPAFIAQVKDKNSGVKLMG
FGHRVYKNFDPRAKIMQQTCHEVLTELGIKDDPLLDLAVELEKIALSDDYFVQRKLYPNVDF
YSGIILKAMGIPTSMFTVLFAVARTTGWVSQWKEMIEEPGQRISRPRQLYIGAPQRDYVPLA
KR
111111111100000000000000000000000000000000000000000000000000
000000000000000000000000000000000000000000000000000000000000
000000000000000000000000000000111111111000000000000000000000
000000000000000000000000000000000000000000000000000000000000
000000000000000000000000000000000000000000000000000000000000
000000000000000000000000000000000000000000000000000000000000
000000000000000000000000000000000000000000000000000000000000
00
>DM_train336
YTPVEEKENGRMIVIVAKKYEGDIKDFVDWKNQRGLRTEVKVAEDIASPVTANAIQQFVKQE
YEKEGNDLTYVLLVGDHKDIPAKITPGIKSDQVYGQIVGNDHYNEVFIGRFSCESKEDLKTQ
IDRTIHYERNITTEDKWLGQALCIASAEGGPSADNGESDIQHENVIANLLTQYGYTKIIKCY
DPGVTPKNIIDAFNGGISLVNYTGHGSETAWGTSHFGTTHVKQLTNSNQLPFIFDVACVNGD
FLFSMPCFAEALMRAQKDGKPTGTVAIIASTIDQYWAPPMRGQDEMNEILCEKHPNNIKRTF
GGVTMNGMFAMVEKYKKDGENMLDTWTVFGDPSLLVRTLVPTEMQVTAPANISASAQTFEVA
CDYNGAIATLSDDGDMVGTAIVKDGKAIIKLNESIADETNLTLTVVGYNKVTVIKDVKVEGT
S
000000000000000000000000000000000000000000000000000000000000
000000000000000000000000000000000000000000000000000000000000
000000000000000000000000000000000000000000000000000000000000
000000000000000000000000000000000000000000000000000000000000
000000000000000000000000000000000000000000000000000000000000
000000000000000000000000000000000000000000000000000000000000
000000000000000000000000000000000000000000000000000000000011
1
>DM_train337
SLMAGLPNSSNALQQWHHLFEAEGTKRSPQAQQHLQQLLRTGLPTRKHENWKYTPLEGLINS
QFVSIAGEISPQQRDALALTLDSVRLVFVDGRYVPALSDATEGSGYEVSINDDRQGLPDAIQ
AEVFLHLTESLAQSVTHIAVKRGQRPAKPLLLMHITQGVAGEEVNTAHYRHHLDLAEGAEAT
VIEHFVSLNDARHFTGARFTINVAANAHLQHIKLAFENPLSHHFAHNDLLLAEDATAFSHSF
LLGGAVLRHNTSTQLNGENSTLRINSLAMPVKNEVCDTRTWLEHNKGFCNSRQLHKTIVSDK
GRAVFNGLINVAQHAIKTDGQMTNNNLLMGKLAEVDTKPQLEIYADDVKCSHGATVGRIDDE
QIFYLRSRGINQQDAQQMIIYAFAAELTEALRDEGLKQQVLARIGQRLPGGAREGGSHHHHH
H
111111111100000000000011100000000000000000000000000000000000

000000000000000000000000000000000000000000000000000000000000
000000000000000000000000000000000000000000000000000000000000
000000000000000000000000000000000000000000000000000000000000
000000000000000000000000000000000000000000000000000000000000
000000000000000000000000000000000000000000000000000000000000
000000000000000000000000000000000000000000000000001111111111
1
>DM_train338
ERTSIAVHALMGLPTGQPANGTKLDSIGLPKVDGMSFTLYRVNEIDLTTQAGWDAASKIKLE
ELYTNGHPTDKVTKVATKKTEGGVAKFDNLTPALYLVVQELNGAEAVVRSQPFLVAAPQTNP
TGDGWLQDVHVYPKHQALSEPVKTAVDPDATQPGFSVGENVKYRVATKIPEIASNTKFEGFT
VADKLPAELGKPDTNKITVTLGGKPINSTDVSVQTYQVGDRTVLSVQLAGATLQSLDQHKDQ
ELVVEFEAPVTKQPENGQLDNQAWVLPSNPTAQWDPEESGDAALRGMPSSRVSSKFGQITIE
KSFDGNTPGADRTATFQLHRCEADGSLVKSDPPISLDGKQEFVTGQDGKAVLSGIHLGTLQL
ESNVMKYTDAWAGKGTEFCLVETATASGYELLPKPVIVKLEANESTNVLVEQKVKIDNKKKN
100000000000000111111111110000000000000000000000000000000000
000000000000000000000000000000000000000000000000000000000000
000000000000000000000000000000000000000000000000000000000000
000000000000000000000000000000000000000000000000000000000000
000000000000000000000000000000000000000000000000000000000000
000000000000000000000000000000000000000000000000000000000000
000000000000000000000000000000000000000000000000000000000011
>DM_train339
MTQDVREPPALKYGIVLDAGSSHTSMFVYKWPADKENDTGIVGQHSSCDVQGGGISSYANDP
SKAGQSLVRCLEQALRDVPRDRHASTPLYLGATAGMRLLNLTSPEATARVLEAVTQTLTQYP
FDFRGARILSGQDEGVFGWVTANYLLENFIKYGWVGRWIRPRKGTLGAMDLGGASTQITFET
TSPSEDPGNEVHLRLYGQHYRVYTHSFLCYGRDQILLRLLASALQIHRFHPCWPKGYSTQVL
LQEVYQSPCTMGQRPRAFNGSAIVSLSGTSNATLCRDLVSRLFNISSCPFSQCSFNGVFQPP
VAGNFIAFSAFYYTVDFLTTVMGLPVGTLKQLEEATEITCNQTWTELQARVPGQKTRLADYC
AVAMFIHQLLSRGYHFDERSFREVVFQKKAADTAVGWALGYMLNLTNLIPADLPGLRKGTHF
111111110000000000000000000000000000000000000000000000000000
000000000000000000000000000000000000000000000000000000000000
000000000000000000000000000000000000000000000000000000000000
000000000000000000000000000000000000000000000000000000000000
000000000000011111100000000000000000000000000000000000000000
000000000000000000000000000000000000000000000000000000000000
000000000000000000000000000000000000000000000000000000000000
>DM_train343
MSNVFYRSSKPYPVAVRGEGVFLYDDAGRRYLDGSSGALVANIGHGRAEVGERMAAQAARLP
FVHGSQFSSDVLEEYAGRLARFVGLPTFRFWAVSGGSEATESAVKLARQYHVERGEPGRFKV
ITRVPSYHGASLGSLAASGMGARRELYTPLMRPEAWPKLPKPDPARNGAEDAEGLRALLERE
GPETVAAFMAEPVVGASDAALAPAPGYYERVRDICDEAGIIFIADEVMSGMGRCGSPLALSR
WSGVTPDIAVLGKGLAAGYAPLAGLLAPQVYETVMGGSGAFMHGFTYAGHPVSVAAGLSVL
DIVEREDLTGAAKERGAQLLAGLQALQARFPQMMQVRGTGLLLGVVLGDLATGQAFETPGIA

SRIGAAALKRGLITYPGSGAEPNGRGDHLLLGPPLSITAAEVDGLLALLAGALEDVLG
11000000000000000000000000000000000000000000000000000000000000
00000000000000000000000000000000000000000000000000000000000000
00000011111111111111111111111110000000000000000000000000000000
00000000000000000000000000000000000000000000000000000000000000
00000000000000000000000000000000000000000000000000000000000000
00000000000000000000000000000000000000000000000011111111111100
0000000000000001111111111100000000000000000000000000000001
>DM_train344
MAEFRIAQDVVARENDRRASALKEDYEALGANLARRGVDIEAVTAKVEKFFVAVPSWGVGTG
GTRFARFPGTGEPRGIFDKLDDCAVIQQLTRATPNVSLHIPWDKADPKELKARGDALGLGFD
AMNSNTFSDAPGQAHSYKYGSLSHTNAATRAQAVEHNLECIEIGKAIGSKALTVWIGDGSNF
PGQSNFTRAFERYLSAMAEIYKGLPDDWKLFSEHKMYEPAFYSTVVQDWGTNYLIAQTLGPK
AQCLVDLGHHAPNTNIEMIVARLIQFGKLGGFHFNDSKYGDDDLDAGAIEPYRLFLVFNELV
DAEARGVKGFHPAHMIDQKHNVTDPIESLINSANEIRRAYAQALLVDRAALSGYQEDNDALM
ATETLKRAYRTDVEPILAEARRRTGGAVDPVATYRASGYRARVAAERPASVAGGGGII
11100000000000000000000000000000000000000000000000000000000000
00000000000000000000000000000000000000000000000000000000000000
00000000000000000000000000000000000000000000000000000000000000
00000000000000000000000000000000000000000000000000000000000000
00000000000000000000000000000000000000000000000000000000000000
00000000000000000000000000000000000000000000000000000000000000
0000000000000000000000000000000000000000000000000111111111
>DM_train345
MTTQTTPAHIAMFSIAAHGHVNPSLEVIRELVARGHRVTYAIPPVFADKVAATGPRPVLYHS
TLPGPDADPEAWGSTLLDNVEPFLNDAIQALPQLADAYADDIPDLVLHDITSYPARVLARRW
GVPAVSLSPNLVAWKGYEEEVAEPMWREPRQTERGRAYYARFEAWLKENGITEHPDTFASHP
PRSLVLIPKALQPHADRVDEDVYTFVGACQGDRAEEGGWQRPAGAEKVVLVSLGSAFTKQPA
FYRECVRAFGNLPGWHLVLQIGRKVTPAELGELPDNVEVHDWVPQLAILRQADLFVTHAGAG
GSQEGLATATPMIAVPQAVDQFGNADMLQGLGVARKLATEEATADLLRETALALVDDPEVAR
RLRRIQAEMAQEGGTRRAADLIEAELPARHERQEPVGDRPNVGDRPAGVRSDRQRSAL
11111110000000000000000000000000000000000000000000000000000000
00000000000000000000000000000000000000000000000000000000000000
00000000000000000000000000000000000000000000000000000000000000
00000000000000000000000000000011111000000000000000000000001000
00000000000000000000001110000000000000000000000000000000000000
00000000000000000000000000000000000000000000000000000000000000
0000000000000000000000000000111111111111111111111111111111
>DM_train346
MNVLVIGRGGREHAIAWKAAQSPLVGKLYVAPGNPGIADVAELVHIDELDIEALVQFAKQQA
IDLTIVGPEAPLASGIVDRFMAEGLRIFGPSQRAALIEGSKAFAKELMKKYGIPTADHAAFT
SYEEAKAYIEQKGAPIVIKADGLAAGKGVTVAQTVEEALAAAKAALVDGQFGTAGSQVVIEE
YLEGEEFSFMAFVNGEKVYPLAIAQDHKRAYDGDEGPNTGGMGAYSPVPQISDEMMDAALEA
ILRPAAKALAAEGRPFLGVLYAGLMATANGPKVIEFNARFGDPEAQVVLPRLKTDLVEAVLA

VMDGKELELEWTDEAVLGVVLAAKGYPGAYERGAEIRGLDRISPDALLFHAGTKREGGAWYT
NGGRVLLLAAKGETLAKAKEKAYEQLAAIDCDGLFYRRDIGRRAIERASAAYTRMKGR
00000000000000000000000000000000000000000000000000000000000000
00000000000000000000000000000000000000000000000000000000000000
00000000000000000000001111000000000000000000000000000000000000
00000000000000000000000000000000000000000000000000000000000000
00000000000000000000000000000000000000000000000000000000000000
00000000000000000000000000000000000000000000000000000000000000
0000000000000000000000000000000000000000000001111111111111
>DM_train347
MGSDKIHHHHHHMFDPESLKKLAIEIVKKSIEAVFPDRAVKETLPKLNLDRVILVAVGKAAW
RMAKAAYEVLGKKIRKGVVVTKYGHSEGPIDDFEIYEAGHPVPDENTIKTTRRVLELVDQLN
ENDTVLFLLSGGGSSLFELPLEGVSLEEIQKLTSALLKSGASIEEINTVRKHLSQVKGGRFA
ERVFPAKVVALVLSDVLGDRLDVIASGPAWPDSSTSEDALKVLEKYGIETSESVKRAILQET
PKHLSNVEIHLIGNVQKVCDEAKSLAKEKGFNAEIITTSLDCEAREAGRFIASIMKEVKFKD
RPLKKPAALIFGGETVVHVKGNGIGGRNQELALSAAIALEGIEGVILCSAGTDGTDGPTDAA
GGIVDGSTAKTLKAMGEDPYQYLKNNDSYNALKKSGALLITGPTGTNVNDLIIGLIV
11111111111111100000000000000000000000000000000000000000000000
00000000000000000000000000000000000000000000000000000000000000
00000000000000000000000000000000000000000000000000000000000000
00000000000000000000000000000000000000000000000000000000000000
00000000000000000000000000000000000000000000000000000000000000
00000000000000000000000000000000000000000000000000000000000000
00000000000000000000000000000000000000000000000000000000000000
000000000000000000000000000000000000000000000000000000000
>DM_train348
MAAVVLENGVLSRKLSDFGQETSYIEDNSNQNGAISLIFSLKEEVGALAKVLRLFEENDINL
THIESRPSRLNKDEYEFFTYLDKRTKPVLGSIIKSLRNDIGATVHELSRDKEKNTVPWFPRT
IQELDRFANQILSYGAELDADHPGFKDPVYRARRKQFADIAYNYRHGQPIPRVEYTEEEKQT
WGTVFRTLKALYKTHACYEHNHIFPLLEKYCGFREDNIPQLEDVSQFLQTCTGFRLRPVAGL
LSSRDFLGGLAFRVFHCTQYIRHGSKPMYTPEPDICHELLGHVPLFSDRSFAQFSQEIGLAS
LGAPDEYIEKLATIYWFTVEFGLCKEGDSIKAYGAGLLSSFGELQYCLSDKPKLLPLELEKT
ACQEYSVTEFQPLYYVAESFSDAKEKVRTFAATIPRPFSVRYDPYTQRVEVLDNTQQ
11111111111111111100000000000000000000000000000000000000000000
00000000000000000000000000000000000000000000000000000000000000
00000000000011111100000000000000000000000000000000000000000000
00000000000000000000000000000000000000000000000000000000000000
00000000000000000000000000000000000000000000000000000000000000
00000000000000000000000000000000000000000000000000000000000000
000000000000000000000000000000000000000000000000000000000
>DM_train349
SQFDILCKTPPKVLVRQFVERFERPSGEKIASCAAELTYLCWMITHNGTAIKRATFMSYNTI
ISNSLSFDIVNKSLQFKYKTQKATILEASLKKLIPAWEFTIIPYNGQKHQSDITDIVSSLQL
QFESSEEADKGNSHSKKMLKALLSEGESIWEITEKILNSFEYTSRFTKTKTLYQFLFLATFI
NCGRFSDIKNVDPKSFKLVQNKYLGVIIQCLVTETKTSVSRHIYFFSARGRIDPLVYLDEFL

RNSEPVLKRVNRTGNSSSNKQEYQLLKDNLVRSYNKALKKNAPYPIFAIKNGPKSHIGRHLM
TSFLSMKGLTELTNVVGNFSDKRASAVARTTYTHQITAIPDHYFALVSRYYAYDPISKEMIA
LKDETNPIEEWQHIEQLKGSAEGSIRYPAWNGIISQEVLDYLSSYINRRIGHHHHHH
00000000000000000000000000000000000000000000000000000000000000
00000000000000000000000000000000000000000000000111000000000000
00001111110000000000000000000000000000000000000000000000000000
00000000000000000000000000000000000000000000000000000000000000
00000000000000000000000000000000000000000000000000000000000000
00000000000000000000001111111111110000000000000000000000000000
000000000000000000000000000000000000000000000000011111111
>DM_train350
MTDFYSLIPSAPKGRFDGIERAHTAEDVKRLRGSVEIKYSLAEMGANRLWKLIHEEDFVNAL
GALSGNQAMQMVRAGLKAIYLSGWQVAADANTASAMYPDQSLYPANAGPELAKRINRTLQRA
DQIETAEGKGLSVDTWFAPIVADAEAGFGDPLDAFEIMKAYIEAGAAGVHFEDQLASEKKCG
HLGGKVLIPTAAHIRNLNAARLAADVMGTPTLIVARTDAEAAKLLTSDIDERDQPFVDYEAG
RTAEGFYQVKNGIEPCIARAIAYAPYCDLIWMETSKPDLAQARRFAEAVHKAHPGKLLAYNC
SPSFNWKKNLDDATIAKFQRELGAMGYKFQFITLAGFHQLNYGMFELARGYKDRQMAAYSEL
QQAEFAAEADGYTATKHQREVGTGYFDAVSLAITGGQSSTTAMKESTETAQFKPAAE
11000000000000000000000000000000000000000000000000000000000000
00000000000000000000001111111111111111111111100000000000000000
00000000000000000000000001110000000000000000000000000000111111
11100000000000000000000000000000000000000000000000000000000000
00000000000000000000000000000000000000000000000000000000000000
00000000000000000000000000000000000000000000000000000000000000
000000000000000111111110000000000001111111111111111111111
>DM_train352
VDFHGYARSGIGWTGSGGEQQCFQATGAQSKYRLGNECETYAELKLGQEVWKEGDKSFYFDT
NVAYSVNQQNDWESTDPAFREANVQGKNLIEWLPGSTIWAGKRFYQRHDVHMIDFYYWDISG
PGAGIENIDLGFGKLSLAATRSTEAGGSYTFSSQNIYDEVKDTANDVFDVRLAGLQTNPDGV
LELGVDYGRANTTDGYKLADGASKDGWMFTAEHTQSMLKGYNKFVVQYATDAMTTQGKGQAR
GSDGSSSFTEELSDGTKINYANKVINNNGNMWRILDHGAISLGDKWDLMYVGMYQNIDWDNN
LGTEWWTVGVRPMYKWTPIMSTLLEVGYDNVKSQQTGDRNNQYKITLAQQWQAGDSIWSRPA
IRIFATYAKWDEKWGYIKDGDNISRYAAATNSGISTNSRGDSDEWTFGAQMEIWW
00000000000000000000000000000000000000000000000000000000000000
00000000000000000000000000000000000000000000000000000000000000
00000000000000000000000000000000000000000000000000000000000000
00000000000000000000000000000000000000000000000000000000000000
00000000001111110000000000000000000000000000000000000000000000
00000000000000000000000000000000000000000000000000000000000000
0000000000000000000000000000000000000000000000000000000
>DM_train353
ELDGKAPSHRNLNVQTWSTAEGAKVLFVEARELPMFDLRLIFAAGSSQDGNAPGVALLTNAM
LNEGVAGKDVGAIAQGFEGLGADFGNGAYKDMAVASLRSLSAVDKREPALKLFAEVVGKPTF
PADSLARIKNQMLAGFEYQKQNPGKLASLELMKRLYGTHPYAHASDGDAKSIPPITLAQLKA

FHAKAYAAGNVVIALVGDLSRSDAEAIAAQVSAALPKGPALAKIEQPAEPKASIGHIEFPSS
QTSLMLAQLGIDRDDPDYAAVSLGNQILGGGGFGTRLMSEVREKRGLTYGVYSGFTPMQARG
PFMINLQTRAEMSEGTLKLVQDVFAEYLKNGPTQKELDDAKRELAGSFPLSTASNADIVGQL
GAMGFYNLPLSYLEDFMRQSQELTVEQVKAAMNKHLNVDKMVIVSAGPTVAQKPL
11111111111000000000000000000000000000000000000000000000000000
00000000000000000000000000000000000000000000000000000000000000
00000000000000000000000000000000000000000000000000000000000000
00000000000000000000000000000000000000000000000000000000000000
00000000000000000000000000000000000000000000000000000000000000
00000000000000000000000000000000000000000000000111110000000000
0000000000000000000000000000000000000000000000000000000
>DM_train354
MMEYKIVENGLTYRIGNGASVPISNTGELIKGLRNYGPYEVPSLKYNQIALIHNNQFSSLIN
QLKSQISSKIDEVWHIHNINISEFIYDSPHFDSIKSQVDNAIDTGVDGIMLVLPEYNTPLYY
KLKSYLINSIPSQFMRYDILSNRNLTFYVDNLLVQFVSKLGGKPWILNVDPEKGSDIIIGTG
ATRIDNVNLFCFAMVFKKDGTMLWNEISPIVTSSEYLTYLKSTIKKVVYGFKKSNPDWDVEK
LTLHVSGKRPKMKDGETKILKETVEELKKQEMVSRDVKYAILHLNETHPFWVMGDPNNRFHP
YEGTKVKLSSKRYLLTLLQPYLKRNGLEMVTPIKPLSVEIVSDNWTSEEYYHNVHEILDEIY
YLSKMNWRGFRSRNLPVTVNYPKLVAGIIANVNRYGGYPINPEGNRSLQTNPWFL
11111111110000000000000000000000000000000000000000000000000000
00000000000000000000000000000000000000000000000000000000000000
00000000000000000001111000000000000000000000000000000000000000
00000000000000000000000000000000000000000000000000000000000000
00000000000000000000000000000000000000000000000000000011111111
00000000000000000001111111111000000000000000000000000000000000
0000000000000000000000000000000000000000111111000000000
>DM_train355
TEAAAQPHALPADAPDIAPERDLLSKFDGLIAERQKLLDSGVTDPFAIVMEQVKSPTEAVIR
GKDTILLGTYNYMGMTFDPDVIAAGKEALEKFGSGTCGSRMLNGTFHDHMEVEQALRDFYGT
TGAIVFSTGYMANLGIISTLAGKGEYVILDADSHASIYDGCQQGNAEIVRFRHNSVEDLDKR
LGRLPKEPAKLVVLEGVYSMLGDIAPLKEMVAVAKKHGAMVLVDEAHSMGFFGPNGRGVYEA
QGLEGQIDFVVGTFSKSVGTVGGFVVSNHPKFEAVRLACRPYIFTASLPPSVVATATTSIRK
LMTAHEKRERLWSNARALHGGLKAMGFRLGTETCDSAIVAVMLEDQEQAAMMWQALLDGGLY
VNMARPPATPAGTFLLRCSICAEHTPAQIQTVLGMFQAAGRAVGVIGLEHHHHHH
11111111111111111111100000000000000000000000000000000000000000
00000000000000000000000000000000000000000000000000000000000000
00000000000000000000000000000000000000000000000000000000000000
00000000000000000000000000000000000000000000000000000000000000
00000000000000000000000000000000000000000000000000000000000000
00000000000000000000000000000000000000000000000000000000000000
0000000000000000000000000000000000000000000000011111111
>DM_train357
GMKVTNYQGATIDPYSKGLGMVPGTSIQLTDAARLEWNLLNEDVSLPAAVLYADRVEHNLKW
MQAFVAEYGVKLAPHGKTTMAPQLFRRQLETGAWGITLATAHQVRAAYHGGVSRVLMANQLV

GRRNMMMVAELLSDPEFEFFCLVDSVEGVEQLGEFFKSVNKQLQVLLELGVPGGRTGVRDAA
QRNAVLEAITRYPDTLKLAGVELYEGVLKEEHEVREFLQSAVAVTRELVEQERFARAPAVLS
GAGSAWYDVVAEEFVKASETGKVEVVLRPGCYLTHDVGIYRKAQTDIFARNPVAKKMGEGLL
PALQLWAYVQSIPEPDRAIIGLGKRDSAFDAGMPEPARHYRPGNEAPRDIAASEGWEIFGLM
DQHAYLRIPAGADLKVGDMIAFDISHPCLTFDKWRQVLVVDPAYRVTEVIETFF
11111111000000000000000000000000000000000000000000000000000000
00000000000000000000000000000000000000000000000000000000000000
00000000000000000000000000000000000000000000000000000000000000
00000000000000000000000000000000000000000000000000000000000000
00000000000000000000000000000000001111111111111111111111000000
00000000000000000000000000000000000000000000000000000000000000
000000000000000000000000000000000000000000000000000000
>DM_train358
MKNTTLYTYRDYFVIRGGKPLTGKVKISGAKNAALPIMFATILTEEPCTITNVPDLLDVRNT
LLLLRELGAELEFLNNTVFINPSINSFITNQEIIRRMRASVLSLGPLLGRFGRAVVGLPGGC
SIGARPIDQHLKFFKEAGADVEVREGYVYVNLKEKRRVHFKFDLVTVTGTENALLYLASVPE
ESILENIALEPEVMDLIEVLKKMGAHVKVEGRSAYVKGSENLKGFTHSVIPDRIEAGTFMVG
AVLTDGEILLENARINHLRAVVEKLKLIGGEVVEENGNLRVFRKESLRACDIETQVYPGFPT
DMQAQFMALLSVAKGKSRIKENIFEHRFHHAQELNRLGANITVRGNTAYVEGVERLYGSEVY
STDLRASASLVLAGLVAQGETVVRDVYHLDRGYEKLEEKLKKLGADIERVSEL
11111111000000000000000000000000000000000000000000000000000000
00000000000000000000000000000000000000000000000000000000000000
00000000000000000000000000000000000000000000000000000000000000
00000000000000000000000000000000000000000000000000000000000000
00000000000000000000000000000000000000000000000000000000000000
00000000000000000000000000000000000000000000000000000000000000
00000000000000000000000000000000000000000000000000001
>DM_train359
GSMSFIPVAEDSDFPIQNLPYGVFSTQSNPKPRIGVAIGDQILDLSVIKHLFTGPALSKHQH
VFDETTLNNFMGLGQAAWKEARASLQNLLSASQARLRDDKELRQRAFTSQASATMHLPATIG
DYTDFYSSRQHATNVGIMFRGKENALLPNWLHLPVGYHGRASSIVVSGTPIRRPMGQMRPDN
SKPPVYGACRLLDMELEMAFFVGPGNRFGEPIPISKAHEHIFGMVLMNDWSARDIQQWEYVP
LGPFLGKSFGTTISPWVVPMDALMPFVVPNPKQDPKPLPYLCHSQPYTFDINLSVSLKGEGM
SQAATICRSNFKHMYWTMLQQLTHHSVNGCNLRPGDLLASGTISGSDPESFGSMLELSWKGT
KAIDVGQGQTRTFLLDGDEVIITGHCQGDGYRVGFGQCAGKVLPALSPAGS
00000000000000000000000000000000000000000000000000000000000000
00000000000000000000000000000000000000000000000000000000000000
00000000000000000000000000000000000000000000000000000000000000
00000000000000000000000000000000000000000000000000000000000000
00000000000000000000000000000000000000000000000000000000000000
00000000000000000000000000000000000000000000000000000000000000
000000000000000000000000000000000000000000000011111
>DM_train360
MSSGVDLGTENLYFQSNAMYMAQPGHIDHIKQINAGRVYKLIDQKGPISRIDLSKESELAPA

SITKITRELIDAHLIHETTVQEAISRGRPAVGLQTNNLGWQFLSMRLGRGYLTIALHELGGE
VLIDTKIDIHEIDQDDVLARLLFEIEEFFQTYAAQLDRVTSIAITLPGLVNSEQGIVLQMPH
YNVKNLALGPEIYKATGLPVFVANDTRAWALAEKLFGHSQDVDNSVLISIHHGLGAGIVLDG
RVLQGRHGNIGELGHIQIDPQGKRCHCGNYGCLETVASSQAIRDQVTARIQAGEPSCLATVE
EISIEDICAAAADGDPLAVDVIQQLGRYLGAAIAIVINLFNPEKILIGGVINQAKSILYPSI
EQCIREQSLPVYHQDLKLVESRFYKQATMPGAALIKQALYDGLLLMKVVEG
111111111111111111111111110000000000000000000000000000000000
000000000000000000000000000000000000000000000000000000000000
000000000000000000000000000000000000000000000000000000000000
000000000000000000000000000000000000000000000000000000000000
000000000000000000000000000000000000000000000000000000000000
000000000000000000000000000000000000000000000000000000000000
000000000000000000000000000000000000000000000000000
>DM_train361
SGDFSLLSSPINREKNGTEIVKFSIHPYKGTVIRLGEEILPFKVLEMDKNIALVEMAIPVYK
DEKEIELKLSSPGFQNSSYRIRKPEELNEKLIALDKEGITHRFISRFKTGFQPKSVRFIDNT
RLAIPLLEDEGMDVLDINSGQTVRLSPPEKYKKKLGFVETISIPEHNELWVSQMQANAVHVF
DLKTLAYKATVDLTGKWSKILLYDPIRDLVYCSNWISEDISVIDRKTKLEIRKTDKIGLPRG
LLLSKDGKELYIAQFSASNQESGGGRLGIYSMDKEKLIDTIGPPGNKRHIVSGNTENKIYVS
DMCCSKIEVYDLKEKKVQKSIPVFDKPNTIALSPDGKYLYVSCRGPNHPTEGYLKKGLVLGK
VYVIDTTTDTVKEFWEAGNQPTGLDVSPDNRYLVISDFLDHQIRVYRRDGF
111111111111111100000000000000000000000000000000000000000000
000000000000000000000000000000000000000000000000000000000000
000000000000000000000000000000000000000000000000000000000000
000000000000000000000000000000000000000000000000000000000000
000000000000000000000000000000000000000000000000000000000000
000000000000000000000000000000000000000000000000000000000000
000000000000000000000000000000000000000000000000000
>DM_train362
MRVLVRDLKAHVGQEVELLGFLHWRRDLGRIQFLLLRDRSGVVQVVTGGLKLPLPESALRVR
GLVVENAKAPGGLEVQAKEVEVLSPALEPTPVEIPKEEWRANPDTLLEYRYVTLRGEKARAP
LKVQAALVRGFRRYLDRQDFTEIFTPKVVRAGAEGGSGLFGVDYFEKRAYLAQSPQLYKQIM
VGVFERVYEVAPVWRMEEHHTSRHLNEYLSLDVEMGFIADEEDLMRLEEALLAEMLEEALNT
AGDEIRLLGATWPSFPQDIPRLTHAEAKRILKEELGYPVGQDLSEEAERLLGEYAKERWGSD
WLFVTRYPRSVRPFYTYPEEDGTTRSFDLLFRGLEITSGGQRIHRYEELLESLKAKGMDPEA
FHGYLEVFKYGMPPHGGFAIGAERLTQKLLGLPNVRYARAFPRDRHRLTP
000000000000000000000000000000000000000000000000000000000000
000000000000000000000000000000011111111111111100000000000000
000000000000000000000000001111111111111111111111111100000000
000000000000011111111111110000000000000000000000000000000000
000000000000000000000000000000000000000000000000000000000000
000000000000000000000000000000000000000000000000000000000000
00000000000000000000000000000000000000000001111111
>DM_train363

GSQSRSIQTFPQPDTSVINGPDRPAGIPDPAGTTVAGGGAVYTVVPHLSMPHWAAQDFAKSL
QSFRLGCANLKNRQGWQDVCAQAFQTPIHSFQAKRFFERYFTPWQVAGNGSLAGTVTGYYEP
VLKGDGRRTERARFPIYGIPDDFISVPLPAGLRGGKNLVRIRQTGKNSGTIDNAGGTHTADL
SRFPITARTTAIKGRFEGSRFLPYHTRNQINGGALDGKAPILGYAEDPVELFFMHIQGSGRL
KTPSGKYIRIGYADKNEHPYVSIGRYMADKGYLKLGQTSMQGIKAYMRQNPQRLAEVLGQNP
SYIFFRELAGSGGDGPVGALGTPLMGEYAGAIDRHYITLGAPLFVATAHPVTRKALNRLIMA
QDTGSAIKGAVRVDYFWGYGDEAGELAGKQKTTGYVWQLLPNGMKPEYRP
111111111111111111100000000000000000000000000000000000000000
000000000000000000000000000000000000000000000000000000000000
000000000000000000000000000111111110000000000001111111000000
000000000000000000000000000000000000000000000000000000000000
000000000000000000000000000000000000000000000000000000000000
000000001111100000000000000000000000000000000000000000000000
00000000000000000000000000000000000000000000000000
>DM_train365
MGLPPLLSLPSNSAPRSLGRVETPPEVVDFMVSLAEAPRGGRVLEPACAHGPFLRAFREAHG
TGYRFVGVEIDPKALDLPPWAEGILADFLLWEPGEAFDLILGNPPYGIVGEASKYPIHVFKA
VKDLYKKAFSTWKGKYNLYGAFLEKAVRLLKPGGVLVFVVPATWLVLEDFALLREFLAREGK
TSVYYLGEVFPQKKVSAVVIRFQKSGKGLSLWDTQESESGFTPILWAEYPHWEGEIIRFETE
ETRKLEISGMPLGDLFHIRFAARSPEFKKHPAVRKEPGPGLVPVLTGRNLKPGWVDYEKNHS
GLWMPKERAKELRDFYATPHLVVAHTKGTRVVAAWDERAYPWREEFHLLPKEGVRLDPSSLV
QWLNSEAMQKHVRTLYRDFVPHLTLRMLERLPVRREYGFHTSPESARNF
111111111111111111100000000000000000000000000000000000000000
000000000000000000000000000000000000000000000001111111111110
000000000000000000000000000000000000000000000000000000000000
000000000000000000000000000000000000000000000000000000000000
000000000000000000000000000000000000000000000000000000000000
000000000000000000000000000000000000000000000000000000000000
0000000000000000000000000000000000000000011111111
>DM_train366
MSRTVMERIEYEMHTPDPKADPDKLHFVQIDEAKCIGCDTCSQYCPTAAIFGEMGEPHSIPH
IEACINCGQCLTHCPENAIYEAQSWVPEVEKKLKDGKVKCIAMPAPAVRYALGDAFGMPVGS
VTTGKMLAALQKLGFAHCWDTEFTADVTIWEEGSEFVERLTKKSDMPLPQFTSCCPGWQKYA
ETYYPELLPHFSTCKSPIGMNGALAKTYGAERMKYDPKQVYTVSIMPCIAKKYEGLRPELKS
SGMRDIDATLTTRELAYMIKKAGIDFAKLPDGKRDSLMGESTGGATIFGVTGGVMEAALRFA
YEAVTGKKPDSWDFKAVRGLDGIKEATVNVGGTDVKVAVVHGAKRFKQVCDDVKAGKSPYHF
IEYMACPGGCVCGGGQPVMPGVLEAMDRTTTRLYAGLKKRLAMASANKA
000000000000000000000000000000000000000000000000000000000000
000000000000000000000000000000000000000000000000000000000000
000000000000000000000000000000000000000000000000000000000000
000000000000000000000000000000000000000000000000000000000000
000000000000000000000000000000000000000000000000000000000000
000000000000000000000000000000000000000000000000000000000000
0000000000000000000000000111111111111111111111111

>DM_train367
MGSDKIHHHHHHMKEKVVLAYSGGLDTSVILKWLCEKGFDVIAYVANVGQKDDFVAIKEKAL
KTGASKVYVEDLRREFVTDYIFTALLGNAMYEGRYLLGTAIARPLIAKRQVEIAEKEGAQYV
AHGATGKGNDQVRFELTYAALNPNLKVISPWKDPEFLAKFKGRTDLINYAMEKGIPIKVSKK
RPYSEDENLMHISHEAGKLEDPAHIPDEDVFTWTVSPKDAPDEETLLEIHFENGIPVKVVNL
KDGTEKTDPLELFEYLNEVGAKNGVGRLDMVENRFIGIKSRGVYETPGATILWIAHRDLEGI
TMDKEVMHLRDMLAPKFAELIYNGFWFSPEMEFLLAAFRKAQENVTGKVTVSIYKGNVMPVA
RYSPYSLYNPELSSMDVEGGFDATDSKGFINIHALRLKVHQLVKKGYQR
111111111111100000000000000000000000000000000000000000000000
000000000000000000000000000000000000000000000000000000000000
000000000000000000000000000000000000000000000000000000011110
000000000000000000000000000000000000000000000000000000000000
000000000000000000000000000000000000000000000000000000000000
000000000000000000000000000000000000000000000000000000000000
0000000001111111111000000000000000000000000011111
>DM_train368
MSRNPSNSDAAHAFWSTQPVPQTEDETEKIVFAGPMDEPKTVADIPEEPYPIASTFEWWTPN
MEAADDIHAIYELLRDNYVEDDDSMFRFNYSEEFLQWALCPPSYIPDWHVAVRRKADKKLLA
FIAGVPVTLRMGTPKYMKVKAQEKGQEEEAAKYDAPRHICEINFLCVHKQLREKRLAPILIK
EVTRRVNRTNVWQAVYTAGVLLPTPYASGQYFHRSLNPEKLVEIRFSGIPAQYQKFQNPMAM
LKRNYQLPNAPKNSGLREMKPSDVPQVRRILMNYLDNFDVGPVFSDAEISHYLLPRDGVVFT
YVVENDKKVTDFFSFYRIPSTVIGNSNYNILNAAYVHYYAATSMPLHQLILDLLIVAHSRGF
DVCNMVEILDNRSFVEQLKFGAGDGHLRYYFYNWAYPKIKPSQVALVML
111111111100000000000000000000000000000000000000000000000000
000000000000000000001110000000000000000000000000000000000000
000000000000000000000000000000000000000000000000000000000000
000000000000000000000000000000000000000000000000000000000000
000000000000000000000000000000000000000000000000000000000000
000000000000000000000001111110000000000000000000000000000000
0000000000000000000000000000000000000000000000000
>DM_train371
MFVFMLLLLLPFTISKAKDFPANPIEKAGYKLDFSDEFNGPTLDREKWTDYYLPHWCKDPES
AKANYRFENGSLVEYITEDQKPWCPEHDGTVRSSAIMSFDKSWIHNFSGTTDNHERNEWRGY
TTKYGYFEIRAKLSNTGGGGHQAWWMVGMQDDTNDWFNSKQTGEIDILETFFSKKDTWRIAA
YGWNDPNFQTSWTISEDKVPSGDPTSEYHIYAMEWTPTALKFYYDNELFKVIYGSPDYEMGT
ILNIYTDAGSGAHNDVWPKEWAIDYMRVWKPVDGYKESESLNNYLIRNRQTGKFLYIEENND
KVSYGDITLKNEKNAKWSKEYRDGYTLLKNNETGEYLNIENQTGYIEHGKVPKTWWSAQWSE
VPVDGYTRFVNRWKPNMSIHTESYEGVLQYGNVPNTYWTSQWQLIPVE
111111111111111111000000000000000000000000000000000000000000
000000000000000000000000000000000000000000000000000000000000
000000000000000000000000000000000000000000000000000000000000
000000000000000000000000000000000000000000000000000000000000
000000000000000000000000000000000000111110000000000000000000
000000000000000000000000000000000000000000000000000000000000

0000000000000000000000000000000000000000000000
>DM_train373
MIDLSKTVFYTSIDIGSRYIKGLVLGKRDQEWEALAFSSVKSRGLDEGEIKDAIAFKESVNT
LLKELEEQLQKSLRSDFVISFSSVSFEREDTVIERDFGEEKRSITLDILSEMQSEALEKLKE
NGKTPLHIFSKRYLLDDERIVFNPLDMKASKIAIEYTSIVVPLKVYEMFYNFLQDTVKSPFQ
LKSSLVSTAEGVLTTPEKDRGVVVVNLGYNFTGLIAYKNGVPIKISYVPVGMKHVIKDVSAV
LDTSFEESERLIITHGNAVYNDLKEEEIQYRGLDGNTIKTTTAKKLSVIIHARLREIMSKSK
KFFREVEAKIVEEGEIGIPGGVVLTGGGAKIPRINELATEVFKSPVRTGCYANSDRPSIINA
DEVANDPSFAAAFGNVFAVSENPYEETPVKSENPLKKIFRLFKELME
111111000000000000000000000000000000000000000000000000000000
000000000000000000000000000000000000000000000000000000000000
000000000000000000000000000000000000000000000000000000000000
000000000000000000000000000000000000000000000000000000000000
000000000000000000000000000000000000000000000000000000000000
000000000011111100000000000000000000000000000000000000000000
00000000000000000011111111111111111111111111111
>DM_train375
MAAKNAFYAQSGGVTAVINASAAGVIEAARKQSGKIGRIYAGRNGIIGALTEDLIDTGQESD
AAISALRYTPSGAFGSCRYKLKSLEQNRREYERLIEVFKAHDIGYFFYNGGGDSADTCLKVS
QLSGTLGYPIQAIHVPKTVDNDLPITDCCPGFGSVAKYIAVSTLEASFDVASMSATSTKVFV
LEVMGRHAGWIAAAGGLASSPEREIPVVILFPEISFDKQKFLAKVDSCVKKFGYCSVVVSEG
VKGDDGKFLSDQGVRDAFGHAQLGGVAPVVASMVKEGLGLKYHWGVADYLQRAARHIASKTD
VEQAYAMGQAAVEFAVQGHNSVMPTIERISARPYQWKVGMAQLSQVANVEKMMPENFITEDG
FGITDLCREYLAPLIEGEDYPPYKDGLPDYVRLKNVAVPKKLSGFTL
100000000000000000000000000000000000000000000000000000000000
000000000000000000001111110000000000000000000000000000000000
000000000000000000000000000000000000000000000000000000000000
000000000000000000000000000000000000000000000000000000000000
000000001111111111111111000000000000000000000000000000000000
000000000000000000000000000000010000000000000000000000000000
00000000000000000000000000000000000000000000001
>DM_train376
MGDKGTRVFKKASPNGKLTVYLGKRDFVDHIDLVEPVDGVVLVDPEYLKERRVYVTLTCAFR
YGREDLDVLGLTFRKDLFVANVQSFPPAPEDKKPLTRLQERLIKKLGEHAYPFTFEIPPNLP
CSVTLQPGPEDTGKACGVDYEVKAFCAENLEEKIHKRNSVRLVIRKVQYAPERPGPQPTAET
TRQFLMSDKPLHLEASLDKEIYYHGEPISVNVHVTNNTNKTVKKIKISVRQYADICLFNTAQ
YKCPVAMEEADDTVAPSSTFCKVYTLTPFLANNREKRGLALDGKLKHEDTNLASSTLLREGA
NREILGIIVSYKVKVKLVVSRGGLLGDLASSDVAVELPFTLMHPKPKEEPPHREVPEHETPV
DTNLIELDTNDDDIVFEDFARQRLKGMKDDKEEEEDGTGSPRLNDR
111110000000000000000000000000000000000000000000000000000000
000000000000000000000000000000000000000000000000000000000000
000000000000000000000000000000000000000000000000000000000000
000000000000000000000000000000000000000000000000000000000000
000000000000000000000000000000000000000000000000000000000000

000000000000000000001111111110000000000000000111111111111111
1111111111000000000000000001111111111111111111
>DM_train377
GSHMTNYKEKLQQYAELLVKVGMNVQPKQPVFIRSSVETLELTHLIVEEAYHCGASDVRVVY
SDPTLKRLKFENESVEHFANHEIKSYDVEARMDYVKRGAANLALISEDPDLMDGIDSQKLQA
FQQQNARAFKGYMESVQKNQFPWVVAAFPSKAWAKRVYPELSVEEAYIKFIDEVFDIVRIDG
NDPVENWRQHIANLSVYAQKLQQKNYHALHYVSEGTDLTVGLAKNHIWEDATSYVNGKEQAF
IANIPTEEVFTAPDRNRVDGYVTNKLPLSYNGTIIDQFKLMFKDGEIIDFSAEKGEAVLKDL
INTDEGSRRLGEVALVPDDSPISNRNTIFYNTLFDENAACHLAIGSAYAFNIQGGTEMTVEE
KIASGLNDSNVHVDFMIGSSDLTIYGIFEDGSKELVFENGNWASTF
111110000000000000000000000000000000000000000000000000000000
000000000000000000000000000000000000000000000000000000000000
000000000000000000000000000000000000000000000000000000000000
000000000000000000000000000000000000000000000000000000000000
000000000000000000000000000000000000000000000000000000000000
000000000000000000000000000000000000000000000000000000000000
0000000000000000000000000000000000000000000000
>DM_train378
MSLSTPLQGIKVLDFTGVQSGPSCTQMLAWFGADVIKIERPGVGDVTRHQLRDIPDIDALYF
TMLNSNKRSIELNTKTAEGKEVMEKLIREADILVENFHPGAIDHMGFTWEHIQEINPRLIFG
SIKGFDECSPYVNVKAYENVAQAAGGAASTTGFWDGPPLVSAAALGDSNTGMHLLIGLLAAL
LHREKTGRGQRVTMSMQDAVLNLCRVKLRDQQRLDKLGYLEEYPQYPNGTFGDAVPRGGNAG
GGGQPGWILKCKGWETDPNAYIYFTIQEQNWENTCKAIGKPEWITDPAYSTAHARQPHIFDI
FAEIEKYTVTIDKHEAVAYLTQFDIPCAPVLSMKEISLDPSLRQSGSVVEVEQPLRGKYLTV
GCPMKFSAFTPDIKAAPLLGEHTAAVLQELGYSDDEIAAMKQNHAI
110000000000000000000000000000000000000000000000000000000000
000000000000000000000000000000000000000111111100000000000000
000000000000000000000000000000000000000000000000000000000000
000000000000000000000000000000000000000000000000000000000000
000000000000000000000000000000000000000000000000000000000000
000000000000000000000000000000000000000000000000000000000000
0000000000000000000000000000000000000000000000
>DM_train379
SESPWIAEDTSQIQTKLLEFAKSPEVFDWMVKIRRKIHENPELGYEELETSKLIRSELELIG
IKYRYPVAITGVIGYIGTGEPPFVALRADMDALPIQEGVEWEHKSKIAGKMHACGHDGHVTM
LLGAAKILHEHRHHLQGTVVLIFQPAEEGLSGAKKMREEGALKNVEAIFGIHLSARIPFGKA
ASRAGSFLAGAGVFEAVITGKGGHAAIPQHTIDPVVAASSIVLSLQQLVSRETDPLDSKVVT
VSKVNGGNAFNVIPDSITIGGTLRAFTGFTQLQQRVKEVITKQAAVHRCNASVNLTPNGREP
MPPTVNNKDLYKQFKKVVRDLLGQEAFVEAAPVMGSEDFSYFAETIPGHFSLLGMQDETNGY
ASSHSPLYRINEDVLPYGAAIHASMAVQYLKEKASKGSVSGFHEEL
111111111111111000000000000000000000000000000000000000000000
000000000000000000000000000000000000000000000000000000000000
000000000000000000000000000000000000000000000000000000000000
000000000000000000000000000000000111111111000000000000000000000000000000

0000011111111000000000000000000000000000000000000000000000000000
00000000000000000000000000000000000000000000000000000000000000
0000000000000000000000000000000000011111111111
>DM_train381
TSPTTTSTGTGTGSSTDQAISPLVEGENAPEVTTAKLGFIALTDAAPLIIAKEKGFYAKYGM
PDVEVLKQASWGTTRDNLVLGSASGGIDGAHILTPMPYLITMGTVTDGKPTPMYILARLNVN
GQGIQLGNNYKDLKVGTDAAPLKEAFAKVTDPKVAMTFPGGTHDMWIRYWLAAGGMEPGKDF
STIVVPPAQMVANVKVNAMESFCVGEPWPLQTVNQGVGYQALTTGQLWKDHPEKAFGMRADW
VDQNPKAAKALLMAVMEAQQWCDQAENKEEMCQILSKREWFKVPFEDIIDRSKGIYNFGNGQ
ETFEDQEIMQKYWVDNASYPYKSHDQWFLTENIRWGYLPASTDTKAIVDKVNREDLWREAAQ
ALEVPADQIPSSPSRGIETFFDGITFDPENPQAYLDSLKIKSIKA
11111111111111111111111111100000000000000000000000000000000000
00000000000000000000000000000000000000000000000000000000000000
00000000000000000000000000000000000000000000000000000000000000
00000000000000000000000000000000000000000000000000000000000000
00000000000000000000000000000000000000000000000000000000000000
00000000000000000000000000000000000000000000000000000000000000
000000000000000000000000000000000000000011111
>DM_train382
MSLSNKLTLDKLDVKGKRVVMRVDFNVPMKNNQITNNQRIKAAVPSIKFCLDNGAKSVVLMS
HLGRPDGVPMPDKYSLEPVAVELKSLLGKDVLFLKDCVGPEVEKACANPAAGSVILLENLRF
HVEEEGKGKDASGNKVKAEPAKIEAFRASLSKLGDVYVNDAFGTAHRAHSSMVGVNLPQKAG
GFLMKKELNYFAKALESPERPFLAILGGAKVADKIQLINNMLDKVNEMIIGGGMAFTFLKVL
NNMEIGTSLFDEEGAKIVKDLMSKAEKNGVKITLPVDFVTADKFDENAKTGQATVASGIPAG
WMGLDCGPESSKKYAEAVTRAKQIVWNGPVGVFEWEAFARGTKALMDEVVKATSRGCITIIG
GGDTATCCAKWNTEDKVSHVSTGGGASLELLEGKVLPGVDALSNI
11000000000000000000000000000000000000000000000000000000000000
00000000000000000000000000000000000000111100000000000000000000
00000000000000000000000000000000000000000000000000000000000000
00000000000000000000000000000000000000000000000000000000000000
00000000000000000000000000000000000000000000000000000000000000
00000000000000000000000000000000000000000000000000000000000000
011111111111100000000000000000000000000000000
>DM_train385
CGSSSHETSYGYATLSYADYWAGELGQSRDVLLAGNAEADRAGDLDAGMFDAVSRATHGHGA
FRQQFQYAVEVLGEKVLSKQETEDSRGRKKWEYETDPSVTKMVRASASFQDLGEDGEIKFEA
VEGAVALADRASSFMVDSEEYKITNVKVHGMKFVPVAVPHELKGIAKEKFHFVEDSRVTENT
NGLKTMLTEDSFSARKVSSMESPHDLVVDTVGTGYHSRFGSDAEASVMLKRADGSELSHREF
IDYVMNFNTVRYDYYGDDASYTNLMASYGTKHSADSWWKTGRVPRISCGINYGFDRFKGSGP
GYYRLTLIANGYRDVVADVRFLPKYEGNIDIGLKGKVLTIGGADAETLMDAAVDVFADGQPK
LVSDQAVSLGQNVLSADFTPGTEYTVEVRFKEFGSVRAKVVAQ
11111100000000000000000000000000001111111110000000000000000000
00000000000000000000000000000000000000000000000000000000000000
00000000000000000000000000000000000000000000000000000000000000

000000000000000000000000000000000000000000000000000000000000
000000000000000000000000000000000000000000000000000000000000
000000000000000000000000000000000000000000000000000000000000
0000000000000000000000000000000000000000000
>DM_train386
MGSDKIHHHHHHPEQQVQQFAQLPVGFKNMGNTCYLNATLQALYRVNDLRDMILNYNPSQGV
SNSGAQDEEIHKQIVIEMKRCFENLQNKSFKSVLPVVLLNTLRKCYPQFAERDSQGGFYKQQ
DAEELFTQLFHSMSIVFGDKFSEDFRIQFKTTIKDTANDNDITVKENESDSKLQCHISGTTN
FMRNGLLEGLNEKIEKRSDLTGANSIYSVEKKISRLPKFLTVQYVRFFWKRSTNKKSKILRK
VVFPFQLDVADMLTPEYAAEKVKVRDELRKVEKEKNEKEREIKRRKFDPSSSENVMTPREQY
ETQVALNESEKDQWLEEYKKHFPPNLEKGENPSCVYNLIGVITHQGANSESGHYQAFIRDEL
DENKWYKFNDDKVSVVEKEKIESLAGGGESDSALILMYKGFGL
111111111111111111000000000000000000000000000000000000000000
000000000000000000000000000010000000000000000000000111100000
000000000000000000000000000000000000000000000000000000000000
000000000000011111111100000000000000000000000000000000000000
000000000000000000000000000000000000111111111111111100000000
000000000000000000000000000000000000000000000000000000000000
0000000000000000000000000000000000000000000
>DM_train387
MSSKPKYVQDQEMIPGVYWVGIVDWMVRIFHGYHTDEGSSYNSYFIDDECPTVIDSVKYPFA
EEWLSRIAACCPLDKIKYVVMNHAEGDHASSLKDHYHKFTNATFVCTKKCQEHLKILYGMEK
ATWLIVDDKYTLKIGKRTLKFIPVPLLHWPDSTFTYCPEDKILFSNDGFGQHYATSRRWADE
CDVSHVMHLFKEYTANILGLFSAQMRKALEVASTVEIKYILSAHGVSWRGDAMGLAIAEYDR
WSKGQHCQKKVTVVLDSMYGTTHRMALALLDGARSTGCETVLLEMTSSDITKVALHTYDSGA
VAFASPTLNNTMMPSVAAALNYVRGLTLIKGKPAFAFGAFGWSNRAVPDIVAELRDGCKADV
YDEKGITFKFNYTEELLEQAYNAGVDLGKRAIAYCEKNAPKQ
111110000000000000000000000000000000000000000000000000000000
000000000000000000000000000000000000000000000000000000000000
000000000000000000000000000000000000000000000000000000000000
000000000000000000000000000000000000000000000000000000000000
000000000000000000000000000000000000000000000000000000000000
000000000000000000000000000000000000000000000000000000000000
000000000000000000000000000000000000000011
>DM_train388
MGQLFSSPKSDENNDLPSSFTGYFKKFNTGRKIISQEILNLIELRMRAGNIQLTNSAISDAL
KEIDSSVLNVAVTGETGSGKSSFINTLRGIGNEEEGAAKTGVVEVTMERHPYKHPNIPNVVF
WDLPGIGSTNFPPDTYLEKMKFYEYDFFIIISATRFKKNDIDIAKAISMMKKEFYFVRTKVD
SDITNEADGEPQTFDKEKVLQDIRLNCVNTFRENGIAEPPIFLLSNKNVCHYDFPVLMDKLI
SDLPIYKRHNFMVSLPNITDSVIEKKRQFLKQRIWLEGFAADLVNIIPSLTFLLDSDLETLK
KSMKFYRTVFGVDETSLQRLARDWEIEVDQVEAMIKSPAVFKPTDEETIQERLSRYIQEFCL
ANGYLLPKNSFLKEIFYLKYYFLDMVTEDAKTLLKEICLRN
111111111111100000000000000000000000000000000000000000000000
000000000000000000000000000000000000000011110000000000000000

000000000000000000000000000000000000000000000000000000000000
000000000000000000000000000000000000000000000000000000000000
000000000000000000000000000000000000000000000000000000000000
000000000000000000000000000000000000000000000000000000000000
00000000000000000000000000000000000000000
>DM_train389
KIQGGSVVEMQGDEMTRIIWELIKEKLILPYVELDLHSYDLGIENRDATNDQVTKDAAEAIK
KYNVGVKCATITPDEKRVEEFKLKQMWKSPNGTIRNILGGTVFREAIICKNIPRLVTGWVKP
IIIGRHAYGDQYRATDFVVPGPGKVEITYTPKDGTQKVTYMVHDFEEGGGVAMGMYNQDKSI
EDFAHSSFQMALSKGWPLYLSTKNTILKKYDGRFKDIFQEIYDKKYKSQFEAQNICYEHRLI
DDMVAQAMKSEGGFIWACKNYDGDVQSDSVAQGYGSLGMMTSVLICPDGKTVEAEAAHGTVT
RHYRMYQKGQETSTNPIASIFAWSRGLAHRAKLDNNTELSFFAKALEDVCIETIEAGFMTKD
LAACIKGLPNVQRSDYLNTFEFMDKLGENLKAKLAQAK
000000000000000000000000000000000000000000000000000000000000
000000000000000000000000000000000000000000000000000000000000
000000000000000000000000000000000000000000000000000000000000
000000000000000000000000000000000000000000000000000000000000
000000000000000000000000000000000000000000000000000000000000
000000000000000000000000000000000000000000000000000000000000
00000000000000000000000000000000000001
>DM_train390
ADGQQGAPLNSPNTYDVTTWRIKAHPEVTAQSDIGAVINDIIADIKQRQTSPDARPGAAIII
PPGDYDLHTQVVVDVSYLTIAGFGHGFFSRSILDNSNPTGWQNLQPGASHIRVLTSPSAPQA
FLVKRAGDPRLSGIVFRDFCLDGVGFTPGKNSYHNGKTGIEVASDNDSFHITGMGFVYLEHA
LIVRGADALRVNDNMIAECGNCVELTGAGQATIVSGNHMGAGPDGVTLLAENHEGLLVTGNN
LFPRGRSLIEFTGCNRCSVTSNRLQGFYPGMLRLLNGCKENLITANHIRRTNEGYPPFIGRG
NGLDDLYGVVHIAGDNNLISDNLFAYNVPPANIAPAGAQPTQILIAGGDANVVALNHVVSDV
ASQHVVLDASTTHSKVLDSGTASQITSYSSDTAIRPTP
111111111110000000000000000000000000000000000000000000000000
000000000000000000000000000000000000000000000000000000000000
000000000000000000000000000000000000000000000000000000000000
000000000000000000000000000000000000000000000000000000000000
000000000000000000000000000000000000000000000000000000000000
000000000000000000000000000000000000000000000000000000000000
00000000000000000000000000000000000000
>DM_train391
MSVFDSKFKGIHVYSEIGELESVLVHEPGREIDYITPARLDELLFSAILESHDARKEHKQFV
AELKANDINVVELIDLVAETYDLASQEAKDKLIEEFLEDSEPVLSEEHKVVVRNFLKAKKTS
RELVEIMMAGITKYDLGIEADHELIVDPMPNLYFTRDPFASVGNGVTIHYMRYKVRQRETLF
SRFVFSNHPKLINTPWYYDPSLKLSIEGGDVFIYNNDTLVVGVSERTDLQTVTLLAKNIVAN
KECEFKRIVAINVPKWTNLMHLDTWLTMLDKDKFLYSPIANDVFKFWDYDLVNGGAEPQPVE
NGLPLEGLLQSIINKKPVLIPIAGEGASQMEIERETHFDGTNYLAIRPGVVIGYSRNEKTNA
ALEAAGIKVLPFHGNQLSLGMGNARCMSMPLSRKDVKW
100000000000000000000000000000000000000000000000000000000000

000000000000000000000000000000000000000000000000000000000000
000000000000000000000000000000000000000000000000000000000000
000000000000000000000000000000000000000000000000000000000000
000000000000000000000000000000000000000000000000000000000000
000000000000000000000000000000000000000000000000000000000000
0000000000000000000000000000000000000
>DM_train392
GMNNIQAIRKKVETQIDDLQNKTDEIAEFNQAKVLDAFQENKVSDFHFHPSTGYGYDDEGRD
TLERVYATVFKTEAALVRPQIISGTHAISTVLFGILRPDDELLYITGQPYDTLEEIVGIRKQ
GQGSLKDFHIGYSSVPLLENGDVDFPRIAKKMTPKTKMIGIQRSRGYADRPSFTIEKIKEMI
VFVKNINPEVIVFVDNCYGEFVEYQEPPEVGADIIAGSLIKNPGGGLAKTGGYIAGKEALVD
LCGYRLTTPGIGREAGASLYSLLEMYQGFFLAPHVTAQAIKGARFTAAMLAEFGVEADPVWD
APRTDLIQSVSFHNKEKMVAFAQAIQAASPVNAHVLPIGAYMPGYEDDVIMAAGTFIQGASL
ELTADGPIREPYQLYVQGGLTYEHIKIAVTRAIQKIV
111100000000000000000000000000000000000000000000000000000000
000000000000000000000000000000000000000000000000000000000000
000000000000000000000000000000000000000000000000000000000000
000000000000000000000000000000000000000000000000000000000000
000000000000000000000000000000000000000000000000000000000000
000000000000011000000000000000000000000000000110000000000000
0000001111110000000000000000000000001
>DM_train393
APQVVDKVAAVVNNGVVLESDVDGLMQSVKLNAAQARQQLPDDATLRHQIMERLIMDQIILQ
MGQKMGVKISDEQLDQAIANIAKQNNMTLDQMRSRLAYDGLNYNTYRNQIRKEMIISEVRNN
EVRRRITILPQEVESLAQQVGNQNDASTELNLSHILIPLPENPTSDQVNEAESQARAIVDQA
RNGADFGKLAIAHSADQQALNGGQMGWGRIQELPGIFAQALSTAKKGDIVGPIRSGVGFHIL
KVNDLRGESKNISVTEVHARHILLKPSPIMTDEQARVKLEQIAADIKSGKTTFAAAAKEFSQ
DPGSANQGGDLGWATPDIFDPAFRDALTRLNKGQMSAPVHSSFGWHLIELLDTRNVDKTDAA
QKDRAYRMLMNRKFSEEAASWMQEQRASAYVKILSN
111100000000000000000000000000000000000000000000000000000000
000000000000000000000000000000000000000000000000000000000000
000000000000000000011111111000000000000000000000000000000000
000000000000000000000000000000000000000000000000000000000000
000001111111110000000000000000000000000000000000000000000000
000000000000000000000000000000000000000000000000000001111111
111000000000000000000000000000000000
>DM_train394
MLFDEKEILAITRWAKLYANQSPDRILLGSNDIPPEYRAAVATQIELWPRLRNKLPQWAGIS
SLYIPSRLSLEQSSGAVTSSYKSRFIREGTKVVDLTGGLGIDFIALMSKASQGIYIERNDET
AVAARHNIPLLLNEGKDVNILTGDFKEYLPLIKTFHPDYIYVDPARRSGADKRVYAIADCEP
DLIPLATELLPFCSSILAKLSPMIDLWDTLQSLLHVQELHVVAAHGEVKELLVRMSLNEATI
PPEKVPIHAINLLLEDTVIPFIFTMEEERSISIPYTDSIDKYVYEPHTALLKAGAFKTVAYR
LGLRKLHPNSHLYTSEAYESAFPGRTFVLEEIIPFSTSVLKQLRKVVPQASISCRNFPLSPI
ELRQRSKMADGGEKTLMGTTMADGKKVLLLLRKAE

0000000000000000000000000000000000000000000000000000000000000000
0000000000000000000000000000000000000000000000000000000000000000
0000000000000000000000000000000000000000000000001111100000000000
0000000000000000000000000000000000000000000000000000000000000000
0000000000000001000000000000000000000000000000000000000000000000
0000000000000000000000000000000000000000000000000000000000000000
000000000000000000000000000000000000
>DM_train395
PGSSVSGTFGCLGGRLTIPGTGVSLLVPNGAIPQGKFYDLYLRINKTESTLPLSEGSQTVLS
PSVTCGPTGLLLCRPVVLTVPHCAEVIAGDWIFQLKTQAHQGHWEEVVTLDEETLNTPCYCQ
LEAKSCHILLDQLGTYVFTGESYSRSAVKRLQLAIFAPALCTSLEYSLRVYCLEDTPAALKE
VLELERTLGGYLVEEPKPLLFKDSYHNLRLSLHDIPHAHWRSKLLAKYQEIPFYHVWNGSQK
ALHCTFTLERHSLASTEFTCKVCVRQVEGEGQIFQLHTTLAETPAGSLDALCSAPGNAATTQ
LGPYAFKIPLSIRQKICNSLDAPNSRGNDWRLLAQKLSMDRYLNYFATKASPTGVILDLWEA
RQQDDGDLNSLASALEEMGKSEMLVAMTTDGDC
0000000000000000000000000000000000000000000000000000000000000000
0000000000000000000000000000000000000000000000000000000000000000
0000000000000000000000000000000000000000000000000000000000000000
0000000000000000000000000000000000000000000000000000000000000000
0000000000000000000000000000000000000000011111111111111111111000
0000000000000000000000000000000000000000000000000000000000000000
000000000000000000000000000000111
>DM_train396
MADLLPEHPEFLWANPEPKKSYDAIIVGGGGHGLATAYFLAKNHGITNVAVLEKGWLAGGNM
ARNTTIIRSNYLWDESAGIYEKSLKLWEQLPEDLEYDFLFSQRGVLNLAHTLGDVRESVRRV
EANKLNGVDAEWLDPSQVKEACPIINTSDDIRYPVMGATWQPRAGIAKHDHVAWAFARKANE
MGVDIIQNCEVTGFIKDGEKVTGVKTTRGTIHAGKVALAGAGHSSVLAEMAGFELPIQSHPL
QALVSELFEPVHPTVVMSNHIHVYVSQAHKGELVMGAGIDSYNGYGQRGAFHVIQEQMAAAV
ELFPIFARAHVLRTWGGIVDTTMDASPIISKTPIQNLYVNCGWGTGGFKGTPGAGFTLAHTI
ANDEPHELNKPFSLERFETGHLIDEHGAAAVAH
1100000000000000000000000000000000000000000000000000000000000000
0000000000000000000000000000000000000000000000000000000000000000
0000000000000000000000000000000000000000000000000000000000000000
0000000000000000000000000000000000000000000000000000000000000000
0000000000000000000000000000000000000000000000000000000000000000
0000000000000000000000000000000000000000000000000000000000000000
000000000000000000000000000000000
>DM_train398
LHLDQTPSRQPIPSEGLQLHLPQVLADAVSRLVLGKFGDLTDNFSSPHARRKVLAGVVMTTG
TDVKDAKVISVSTGTKCINGEYMSDRGLALNDCHAEIISRRSLLRFLYTQLELYLNNKDDQK
RSIFQKSERGGFRLKENVQFHLYISTSPCGDARIFSPHEPILEEPADRHPNRKARGQLRTKI
ESGEGTIPVRSNASIQTWDGVLQGERLLTMSCSDKIARWNVVGIQGSLLSIFVEPIYFSSII
LGSLYHGDHLSRAMYQRISNIEDLPPLYTLNKPLLSGISNAEARQPGKAPNFSVNWTVGDSA
IEVINATTGKDELGRASRLCKHALYCRWMRVHGKVPSHLLRSKITKPNVYHESKLAAKEYQA

AKARLFTAFIKAGLGAWVEKPTEQDQFSLTP
11111110000000111100000000000000000000000000000000000000000000
00000000000000000000000000000000000000000000000000000000000000
00000000000000000000000000000000000000011111111111100000000000
00000000011111111111111111000000000000000000000000000000000000
00000000000000000000000000000000000000000000000000000000000000
00000000000000000000000000000000000000000000000000000000000000
0000000000000000000000000000001
>DM_train399
MSTFNRIHLVVLDSVGIGAAPDANNFSNAGVPDGASDTLGHISKTVGLNVPNMAKIGLGNIP
RDTPLKTVPAENHPTGYVTKLEEVSLGKDTMTGHWEIMGLNITEPFDTFWNGFPEEIISKIE
KFSGRKVIREANKPYSGTAVIDDFGPRQMETGELIIYTSADPVLQIAAHEDVIPLDELYRIC
EYARSITLERPALLGRIIARPYVGKPRNFTRTANRHDYALSPFAPTVLNKLADAGVSTYAVG
KINDIFNGSGITNDMGHNKSNSHGVDTLIKTMGLSAFTKGFSFTNLVDFDALYGHRRNAHGY
RDCLHEFDERLPEIIAAMKVDDLLLITADHGNDPTYAGTDHTREYVPLLAYSPSFTGNGVLP
VGHYADISATIADNFGVDTAMIGESFLDKLI
10000000000000000000000000000000000000000000000000000000000000
00000000000000000000000000000000000000000000000000000000000000
00000000000000000111111111111100000000000000000000000000000000
00000000000000000000011111111110000000000000000000000000000000
00000000000000000000000000000000000000000000000000000000000000
00000000000000000000000000000000000000000000000000000000000000
0000000000000000000000000000000
>DM_train401
EKPGETKEVHPQLTTFRCTKRGGCKPATNFIVLDSLSHPIHRAEGLGPGGCGDWGNPPPKDV
CPDVESCAKNCIMEGIPDYSQYGVTTNGTSLRLQHILPDGRVPSPRVYLLDKTKRRYEMLHL
TGFEFTFDVDATKLPCGMNSALYLSEMHPTGAKSKYNPGGAYYGTGYCDAQCFVTPFINGLG
NIEGKGSCCNSMDIWEANSRASHVAPHTCNKKGLYLCEGEECAFEGVCDKNGCGWNNYRVNV
TDYYGRGEEFKVNTLKPFTVVTQFLANRRGKLEKIHRFYVQDGKVIESFYTNKEGVPYTNMI
DDEFCEATGSRKYMELGATQGMGEALTRGMVLAMSIWWDQGGNMEWLDHGEAGPCAKGEGAP
SNIVQVEPFPEVTYTNLRWGEIGSTYQELQ
00000000000000000000000000000000000000000000000000000000000000
00000000000000000000000000000000000000000000000000000000000000
00000000000000000000000000000000000000000000000000000000000000
00000000000000000000000000000000000000000000000000000000000000
00000000000000000000000000000000000000000000000000000000000000
00000000000000000000000000000000000000000000000000000000000000
000000000000000000000000000111
>DM_train402
GSMRICIFMARGLEGCGVTKFSLEQRDWFIKNGHEVTLVYAKDKSFTRTSSHDHKSFSIPVI
LAKEYDKALKLVNDCDILIINSVPATSVQEATINNYKKLLDNIKPSIRVVVYQHDHSVLSLR
RNLGLEETVRRADVIFSHSDNGDFNKVLMKEWYPETVSLFDDIEEAPTVYNFQPPMDIVKVR
STYWKDVSEINMNINRWIGRTTTWKGFYQMFDFHEKFLKPAGKSTVMEGLERSPAFIAIKEK
GIPYEYYGNREIDKMNLAPNQPAQILDCYINSEMLERMSKSGFGYQLSKLNQKYLQRSLEYT

HLELGACGTIPVFWKSTGENLKFRVDNTPLTSHDSGIIWFDENDMESTFERIKELSSDRALY
DREREKAYEFLYQHQDSSFCFKEQFDIITK
000000000000000000000000000000000000000000000000000000000000
000000000000000000000000000000000000000000000000000000000000
000000000000000000011111111111111111111111111111000000000000
000000000000000000000000000000000000000000000000000000000000
000000000000000000000000000000000000000000000000000000000000
000000000000000000000000000000000000000000000000000000000000
000000000000000000000000000000
>DM_train403
MSEPIDILIAGAGIGGLSCALALHQAGIGKVTLLESSSEIRPLGVGINIQPAAVEALAELGL
GPALAATAIPTHELRYIDQSGATVWSEPRGVEAGNAYPQYSIHRGELQMILLAAVRERLGQQ
AVRTGLGVERIEERDGRVLIGARDGHGKPQALGADVLVGADGIHSAVRAHLHPDQRPLSHGG
ITMWRGVTEFDRFLDGKTMIVANDEHWSRLVAYPISARHAAEGKSLVNWVCMVPSAAVGQLD
NEADWNRDGRLEDVLPFFADWDLGWFDIRDLLTRNQLILQYPMVDRDPLPHWGRGRITLLGD
AAHLMYPMGANGASQAILDGIELAAALARNADVAAALREYEEARRPTANKIILANREREKEE
WAAASRPKTEKSAALEAITGSYRNQVERPR
111000000000000000000000000000000000000000000000000000000000
000000000000000000000000000000000000000000000000000000000000
000000000000000000000000000000000000000000000000000000000000
000000000000000000000000000000000000000000000000000000000000
000000000000000000000000000000000000000000000000000000000000
000000000000000000000000000000000000000000000000000000000000
000000011111111111111111110000
>DM_train404
MQRPSDQTAPGTSSRPILSAKEAQNFDAQHYFASLTPGAAAWNPSPITLPAQPDFVVGPAGT
QGVTHTTIQAAVDAAIIKRTNKRQYIAVMPGEYQGTVYVPAAPGGITLYGTGEKPIDVKIGL
SLDGGMSPADWRHDVNPRGKYMPGKPAWYMYDSCQSKRSDSIGVLCSAVFWSQNNGLQLQNL
TIENTLGDSVDAGNHPAVALRTDGDQVQINNVNILGRQNTFFVTNSGVQNRLETNRQPRTLV
TNSYIEGDVDIVSGRGAVVFDNTEFRVVNSRTQQEAYVFAPATLSNIYYGFLAVNSRFNAFG
DGVAQLGRSLDVDANTNGQVVIRDSAINEGFNTAKPWADAVISNRPFAGNTGSVDDNDEIQR
NLNDTNYNRMWEYNNRGVGSKVVAEAKK
000000000000000000000000000000000000000000000000000000000000
000000000000000000000000000000000000000000000000000000000000
000000000000000000000000000000000000000000000000000000000000
000000000000000000000000000000000000000000000000000000000000
000000000000000000000000000000000000000000000000000000000000
000000000000000000000000000000000000000000000000000000000000
0000000000000000000001111111
>DM_train405
SSLDDKPQFPGASAEFIDKLEFIQPNVISGIPIYRVMDRQGQIINPSEDPHLPKEKVLKLYK
SMTLLNTMDRILYESQRQGRISFYMTNYGEEGTHVGSAAALDNTDLVFGQAREAGVLMYRDY
PLELFMAQCYGNISDLGKGRQMPVHYGCKERHFVTISSPLATQIPQAVGAAYAAKRANANRV
VICYFGEGAASEGDAHAGFNFAATLECPIIFFCRNNGYAISTPTSEQYRGDGIAARGPGYGI

MSIRVDGNDVFAVYNATKEARRRAVAENQPFLIEAMTYRIGHASTSDDSSAFRSVDEVNYWD
KQDHPISRLRHYLLSQGWWDEEQEKAWRKQSRRKVMEAFEQAERKPKPNPNLLFSDVYQEMP
AQLRKQQESLARHLQTYGEHYPLDHFDK
111110000000000000000000000000000000000000000000000000000000
000000000000000000000000000000000000000000000000000000000000
000000000000000000000000000000000000000000000000000000000000
000000000000000000000000000000000000000000000000000000000000
000000000000000000000000000000000000000011001111111111111111
110000000000000000000000000000000000000000000000000000000000
0000000000000000000000000000
>DM_train406
MGVSPAAPASGIRRPCYLVLSSHDFRTPRRANIHFITDQLALRGTTRFFSLRYSRLSRMKGD
MRLPLDDTANTVVSHNGVDCYLWRTTVHPFNTRRSWLRPVEDAMFRWYAAHPPKQLLDWMRE
SDVIVFESGIAVAFIELAKRVNPAAKLVYRASDGLSTINVASYIEREFDRVAPTLDVIALVS
PAMAAEVVSRDNVFHVGHGVDHNLDQLGDPSPYAEGIHAVAVGSMLFDPEFFVVASKAFPQV
TFHVIGSGMGRHPGYGDNVIVYGEMKHAQTIGYIKHARFGIAPYASEQVPVYLADSSMKLLQ
YDFFGLPAVCPNAVVGPYKSRFGYTPGNADSVIAAITQALEAPRVRYRQCLNWSDTTDRVLD
PRAYPETRLYPHPPTAAPQLSSEAALSH
111111111111000000000000000000000000000000000000000000000000
000000000000000000000000000000000000000000000000000000000000
000000000000000000000000000000000000000000000000000000000000
000000000000000000000000000000000000000000000000000000000000
000000000000000000000000000000000000000000000000000000000000
000000000000000000000000000000000000000000000000000000000000
0000000000000111111111111111
>DM_train407
MTALKVGSESWWQSKHGPEWQRLNDEMFEVTFWWRDPQGSEEYSTIKRVWVYITGVTDHHQN
SQPQSMQRIAGTDVWQWTTQLNANWRGSYCFIPTERDDIFSAPSPDRLELREGWRKLLPQAI
ADPLNPQSWKGGLGHAVSALEMPQAPLQPGWDCPQAPEIPAKEIIWKSERLKNSRRVWIFTT
GDVTAEERPLAVLLDGEFWAQSMPVWPVLTSLTHRQQLPPAVYVLIDAIDTTHRAHELPCNA
DFWLAVQQELLPLVKVIAPFSDRADRTVVAGQSFGGLSALYAGLHWPERFGCVLSQSGSYWW
PHRGGQQEGVLLEKLKAGEVSAEGLRIVLEAGIREPMIMRANQALYAQLHPIKESIFWRQVD
GGHDALCWRGGLMQGLIDLWQPLFHDRS
111110000000000000000000000000000000000000000000000000011111
110000000000000000000000000000000000000000001100000000000000
000000000000000000000000000000000000000000000000000000000000
111110000000000000000000000000000000000000000000000000000000
000000000000000000000000000000000000000000000000000000000000
000000000000000000000000000000000000000000000000000000000000
0000000000000000000000000111
>DM_train408
MKTITLYLDPASLPALNQLMDFTQNNEDKTHPRIFGLSRFKIPDNIITQYQNIHFVELKDNR
PTEALFTILDQYPGNIELNIHLNIAHSVQLIRPILAYRFKHLDRVSIQQLNLYDDGSMEYVD
LEKEENKDISAEIKQAEKQLSHYLLTGKIKFDNPTIARYVWQSAFPVKYHFLSTDYFEKAEF

LQPLKEYLAENYQKMDWTAYQQLTPEQQAFYLTLVGFNDEVKQSLEVQQAKFIFTGTTTWEG
NTDVREYYAQQQLNLLNHFTQAEGDLFIGDHYKIYFKGHPRGGEINDYILNNAKNITNIPAN
ISFEVLMMTGLLPDKVGGVASSLYFSLPKEKISHIIFTSNKQVKSKEDALNNPYVKVMRRLG
IIDESQVIFWDSLKQLGGGLEHHHHHH
000000000000000000000000000000000000000000000000000000000000
000000000000000000000000000000000000000000000000000000000000
000000000000000000000000000000000000000000000000000000000000
000000000000000000000000000000000000000000000000000000000000
000000000000000000000000000000000000000000000000000000000000
000000000000000000000000000000000000000000000000000000000000
000000000000000000111111111
>DM_train409
TMAHHHHHHVTNSTDGRADGRLRVVVLGSTGSIGTQALQVIADNPDRFEVVGLAAGGAHLDT
LLRQRAQTGVTNIAVADEHAAQRVGDIPYHGSDAATRLVEQTEADVVLNALVGALGLRPTLA
ALKTGARLALANKESLVAGGSLVLRAARPGQIVPVDSEHSALAQCLRGGTPDEVAKLVLTAS
GGPFRGWSAADLEHVTPEQAGAHPTWSMGPMNTLNSASLVNKGLEVIETHLLFGIPYDRIDV
VVHPQSIIHSMVTFIDGSTIAQASPPDMKLPISLALGWPRRVSGAAAACDFHTASSWEFEPL
DTDVFPAVELARQAGVAGGCMTAVYNAANEEAAAAFLAGRIGFPAIVGIIADVLHAADQWAV
EPATVDDVLDAQRWARERAQRAVSGM
111111111111111111100000000000000000000000000000000000000000
000000000000000000000000000000000000000000000000000000000000
000000000000000000000000000000000000000000000000000000000000
000000000000000000000000000000000000000000000000000000000000
000000000000000000000000000000000000000000000000000000000000
000000000000000000000000000000000000000000000000000000000000
00000000000000000000000000
>DM_train410
MAYDVARVRGLHPSLGDGWVHFDAPAGMLIPDSVATTVSTAFRRSGASTVGAHPSARRSAAV
LDAAREAVADLVNADPGGVVLGADRAVLLSLLAEASSSRAGLGYEVIVSRLDDEANIAPWLR
AAHRYGAKVKWAEVDIETGELPTWQWESLISKSTRLVAVNSASGTLGGVTDLRAMTKLVHDV
GALVVVDHSAAAPYRLLDIRETDADVVTVNAHAWGGPPIGAMVFRDPSVMNSFGSVSTNPYA
TGPARLEIGVHQFGLLAGVVASIEYLAALDESARGSRRERLAVSMQSADAYLNRVFDYLMVS
LRSLPLVMLIGRPEAQIPVVSFAVHKVPADRVVQRLADNGILAIANTGSRVLDVLGVNDVGG
AVTVGLAHYSTMAEVDQLVRALASLG
110000000000000000000000000000000000000000000000000000000000
000000000000000000000000000000000000000000000000000000000000
000000000000000000000000000000000000000000000000000000000000
000000000000000000000000000000000000000000000000000000000000
000000000000000000000000000000000000000000000000000000000000
000000000000000000000000000000000000000000000000000000000000
00000000000000000000000000
>DM_train411
MSKVFIATANAGKAHDADIFSVSACNSFTVSCSGDGYLKVWDNKLLDNENPKDKSYSHFVHK
SGLHHVDVLQAIERDAFELCLVATTSFSGDLLFYRITREDETKKVIFEKLDLLDSDMKKHSF

WALKWGASNDRLLSHRLVATDVKGTTYIWKFHPFADESNSLTLNWSPTLELQGTVESPMTPS
QFATSVDISERGLIATGFNNGTVQISELSTLRPLYNFESQHSMINNSNSIRSVKFSPQGSLL
AIAHDSNSFGCITLYETEFGERIGSLSVPTHSSQASLGEFAHSSWVMSLSFNDSGETLCSAG
WDGKLRFWDVKTKERITTLNMHCDDIEIEEDILAVDEHGDSLAEPGVFDVKFLKKGWRSGMG
ADLNESLCCVCLDRSIRWFREAGGK
11000000000000000000000000000000000000000000000000000000000000
00000000000000000000000000000000000000000000000000000000000000
00000000011110000000000000000000000000000000000000000000000000
00000000000000000000000000000000000000000111000000000000000000
00000000000000000000000000000111111100000000000000000000000000
00000000000000000000000000000000000000000000000000000000000000
0000000000000000000000011
>DM_train412
APARGTLLTSNFLTSYTRDAISAMLASGSQPASGSQPEQAKCNVRVAEFTYATIGVEGEPAT
ASGVLLIPGGERCSGPYPLLGWGHPTEALRAQEQAKEIRDAKGDDPLVTRLASQGYVVVGSD
YLGLGKSNYAYHPYLHSASEASATIDAMRAARSVLQHLKTPLSGKVMLSGYSQGGHTAMATQ
REIEAHLSKEFHLVASAPISGPYALEQTFLDSWSGSNAVGENTFGILLGSYAIVAMQHTYKN
IYLEPGQVFQDPWAAKVEPLFPGKQSLTDMFLNDTLPSIDKVKSYFQPGFYSDFPSNPANPF
RQDLARNNLLEWAPQTPTLLCGSSNDATVPLKNAQTAIASFQQRGSNQVALVDTGTGNASDN
SAFAHMLTKESCIVVVRDQLLDKQR
10000000000000000000000000011111111100000000000000000000000000
00000000000000000000000000000000000000000000000000000000000000
00000000000000000000000000000000000000000000000000000000000000
00000000000000000000000000000000000000000000000000000000000000
00000000000000000000000000000000000000000000000000000000000000
00000000000000000000000000000000000000000000000000000000000000
0000000000000000000000000
>DM_train413
MSLVVFPFKHEHPEVLLHNVRVAAAHPRVHEVLCIGYERDQTYEAVERAAPEISRATGTPVS
VRLQERLGTLRPGKGDGMNTALRYFLEETQWERIHFYDADITSFGPDWITKAEEAADFGYGL
VRHYFPRASTDAMITWMITRTGFALLWPHTELSWIEQPLGGELLMRREVAAMLYEDERVRRR
SDWGIDTLYTFVTVQQGVSIYECYIPEGKAHRLYGGLDDLRTMLVECFAAIQSLQHEVVGQP
AIHRQEHPHRVPVHIAERVGYDVEATLHRLMQHWTPRQVELLELFTTPVREGLRTCQRRPAF
NFMDEMAWAATYHVLLEHFQPGDPDWEELLFKLWTTRVLNYTMTVALRGYDYAQQYLYRMLG
RYRYQAALENGRGHPVPPRAALSTA
10000000000000000000000000000000000000000000000000000000000000
00000000000000000000000000000000000000000000000000000000000000
00000000000000000000000000000000000000000000000000000000000000
00000000000000000000000000000000000000000000000000000000000000
00000000000000000000000000000000000000000000000000000000000000
00000000000000000000000000000000000000000000000000000000000000
0000000001111111111111111
>DM_train414
MGVTKTPLYETLNESSAVALAVKLGLFPSKSTLTCQEIGDGNLNYVFHIYDQEHDRALIIKQ

AVPYAKVVGESWPLTIDRARIESSALIRQGEHVPHLVPRVFYSDTEMAVTVMEDLSHLKIAR
KGLIEGENYPHLSQHIGEFLGKTLFYSSDYALEPKVKKQLVKQFTNPELCDITERLVFTDPF
FDHDTNDFEEELRPFVEKLWNNDSVKIEAAKLKKSFLTSAETLIHGDLHTGSIFASEHETKV
IDPEFAFYGPIGFDVGQFIANLFLNALSRDGADREPLYEHVNQVWETFEETFSEAWQKDSLD
VYANIDGYLTDTLSHIFEEAIGFAGCELIRRTIGLAHVADLDTIVPFDKRIGRKRLALETGT
AFIEKRSEFKTITDVIELFKLLVKE
111111000000000000000000011111110000000000011111000000000000
000111111111000000000000000000000000000000000000000000000000
000000000000000000000000000000000000000000000000000000000000
000000000000000000000000000000000000000000000000000000000000
000000000000000000000000000000000000000000000000000000000000
000000000000000000000000000000000000000000000000000000000000
0000000000000000000000001
>DM_train416
MSPLRKTVPEFLAHLKSLPISKIASNDVLTICVGNESADMDSIASAITYSYCQYIYNEGTYS
EEKKKGSFIVPIIDIPREDLSLRRDVMYVLEKLKIKEEELFFIEDLKSLKQNVSQGTELNSY
LVDNNDTPKNLKNYIDNVVGIIDHHFDLQKHLDAEPRIVKVSGSCSSLVFNYWYEKLQGDRE
VVMNIAPLLMGAILIDTSNMRRKVEESDKLAIERCQAVLSGAVNEVSAQGLEDSSEFYKEIK
SRKNDIKGFSVSDILKKDYKQFNFQGKGHKGLEIGLSSIVKRMSWLFNEHGGEADFVNQCRR
FQAERGLDVLVLLTSWRKAGDSHRELVILGDSNVVRELIERVSDKLQLQLFGGNLDGGVAMF
KQLNVEATRKQVVPYLEEAYSNLEE
111000000000000000000000000000000000000000000000000000000000
000000000000000000000000000000000000000000000000000000000000
000000000000000000000000000000000000000000000000000000000000
000000000000000000000000000000000000000000000000000000000000
000000000000000000000000011111000000000000000000000000000000
000000000000000000000000000000000000000000000000000000000000
0000000000000000000000000
>DM_train417
GMETYDVLVVGGGPGGSTAARYAAKYGLKTLMIEKRPEIGSPVRCGEGLSKGILNEADIKAD
RSFIANEVKGARIYGPSEKRPIILQSEKAGNEVGYVLERDKFDKHLAALAAKAGADVWVKSP
ALGVIKENGKVAGAKIRHNNEIVDVRAKMVIAADGFESEFGRWAGLKSVILARNDIISALQY
RMINVDVDPDYTDFYLGSIAPAGYIWVFPKGEGMANVGIGSSINWIHNRFELKNYLDRFIEN
HPGLKKGQDIQLVTGGVSVSKVKMPITMPGLMLVGDAARLIDPITGGGIANAIVSGMYAAQV
TKEAIESNDYSPQMMQKYEKLIKERFERKHLRNWVAKEKLAMLSDDTLDKLVDIVSEQVLTT
ISVEAILKAIAEKYPEVVKELEDLI
000000000000000000000000000000000000000000000000000000000000
000000000000000000000000011111000000000000000000000000000000
000000000000000000000000000000000000000000000000000000000000
000000000000000000000000000000000000000000000000000000000000
000000000000000000000000000000000000000000000000000000000000
000000000000000000000000000000000000000000000000000000000000
0000000000000001111111111
>DM_train418

TTLLNPYFGEFGGMYVPQILMPALNQLEEAFVRAQKDPEFQAQFADLLKNYAGRPTALTKCQ
NITAGTRTTLYLKREDLLHGGAHKTNQVLGQALLAKRMGKSEIIAETGAGQHGVASALASAL
LGLKCRIYMGAKDVERQSPNVFRMRLMGAEVIPVHSGSATLKDACNEALRDWSGSYETAHYM
LGTAAGPHPYPTIVREFQRMIGEETKAQILDKEGRLPDAVIACVGGGSNAIGMFADFINDTS
VGLIGVEPGGHGIETGEHGAPLKHGRVGIYFGMKAPMMQTADGQIEESYSISAGLDFPSVGP
QHAYLNSIGRADYVSITDDEALEAFKTLCRHEGIIPALESSHALAHALKMMREQPEKEQLLV
VNLSGRGDKDIFTVHDILKARGEI
000000000000000000000000000000000000000000000000000000000000
000000000000000000000000000000000000000000000000000000000000
000000000000111100000000000000011111100000000000000000000000
000000000000000000000000000000000000000000000000000000000000
000000000000000000000000000000000000000000000000000000000000
000000000000000000000000000000000000000000000000000000000000
00000000000000111111111
>DM_train419
PQDLMVTCTAPVNIAVIKYWGKRDEALILPINSSLSVTLHQDQLKTTTTVAISKDFTEDRIW
LNGREEDVGQPRLQACLREIRRLARKRRSTEDGDTLPLSLSYKVHVASVNNFPTAAGLASSA
AGYACLAYTLAQVYGVEGDLSEVARRGSGSACRSLYGGFVEWQMGEQADGKDSIARQIAPEW
HWPQLRILILVVSADKKQTGSTVGMQTSVETSTLLKFRAESVVPERMKEMTRCIQEQDFQGF
AQLTMKDSNQFHATCLDTFPPISYLNDTSRRIIQLVHRFNTHHGQTKVAYTFDAGPNAVIFT
LEDTVAEFVAAVRHSFPPAANGDKFLKGLQVAPVLLSDELKAALVVEPSPGGVQYIIATQVG
PGPQVLDDTHDHLLGQDGLPQRDL
110000000000000000000000000000000000000000000000000000000000
000000000000000000000000000111111111000000000000000111110000
000000000000000000000000000000000000000000000000000000000000
000000000000000000000000000000000000000000000000000000000000
000000000000000000000000000000000000000000000000000000000000
000000000000000000001100000000000000000000011000000000000000
00000000000000000000111
>DM_train420
MFENITAAPADPILGLADLFRADERPGKINLGIGVYKDETGKTPVLTSVKKAEQYLLENETT
KNYLGIDGIPEFGRCTQELLFGKGSALINDKRARTAQTPGGTGALRVAADFLAKNTSVKRVW
VSNPSWPNHKSVFNSAGLEVREYAYYDAENHTLDFDALINSLNEAQAGDVVLFHGCCHNPTG
IDPTLEQWQTLAQLSVEKGWLPLFDFAYQGFARGLEEDAEGLRAFAAMHKELIVASSYSKNF
GLYNERVGACTLVAADSETVDRAFSQMKAAIRANYSNPPAHGASVVATILSNDALRAIWEQE
LTDMRQRIQRMRQLFVNTLQEKGANRDFSFIIKQNGMFSFSGLTKEQVLRLREEFGVYAVAS
GRVNVAGMTPDNMAPLCEAIVAVL
000000000111111111111111110000000000000000000000000000000000
000000000000000000000000000000000000000000000000000000000000
000000000000000000000000000000000000000000000000000000000000
000000000000000000000000000000000000000000000000000000000000
000000000000000000000000000000000000000000000000000000000000
000000000000000000000000000000000000000000000000000000000000
00000000000000000000000

>DM_train421
MSQPQSSQVTKRGLTDPERAAIIAAAVPDHALDTQRKYHYFIQPRWKPLSEYEQLSCYAQPN
PDWIAGGLDWGDWTQKFHGGRPSWGNESTELRTTDWYRHRDPARRWHHPYVKDKSEEARYTQ
RFLAAYSSEGSIRTIDPYWRDEILNKYFGALLYSEYGLFNAHSSVGRDCLSDTIRQTAVFAA
LDKVDNAQMIQMERLFIAKLVPGFDASTDVPKKIWTTDPIYSGARATVQEIWQGVQDWNEIL
WAGHAVYDATFGQFARREFFQRLATVYGDTLTPFFTAQSQTYFQTTRGAIDDLFVYCLANDS
EFGAHNRTFLNAWTEHYLASSVAALKDFVGLYAKVEKVAGATDSAGVSEALQRVFGDWKIDY
ADKIGFRVDVDQKVDAVLAGYKN
111111111100000000000000000000000000000000000000000000000000
000000000000000000000000000000000000000000000000000000000000
000000000000000000000000000000000000000000000000000000000000
000000000000000000000000000000000000000000000000000000000000
000000000000000000000000000000000000000000000000000000000000
000000000000000000000000000000000000000000000000000000000000
00000000000000000000000
>DM_train422
MFAKFPSSASISPNPFTVSIPDEQLDDLKTLVRLSKIAPPTYESLQADGRFGITSEWLTTMR
EKWLSEFDWRPFEARLNSFPQFTTEIEGLTIHFAALFSEREDAVPIALLHGWPGSFVEFYPI
LQLFREEYTPETLPFHLVVPSLPGYTFSSGPPLDKDFGLMDNARVVDQLMKDLGFGSGYIIQ
GGDIGSFVGRLLGVGFDACKAVHLNFCNMSAPPEGPSIESLSAAEKEGIARMEKFMTDGYAY
AMEHSTRPSTIGHVLSSSPIALLAWIGEKYLQWVDKPLPSETILEMVSLYWLTESFPRAIHT
YREWVPTASAPNGATPYQKELYIHKPFGFSFFPKDLVPVPRSWIATTGNLVFFRDHAEGGHF
AALERPRELKTDLTAFVEQVWQK
100000000000000000000000000000000000000000000000000000000000
000000000000000000000000000000000000000000000000000000000000
000000000000000000000000000000000000000000000000000000000000
000000000000000000000000000000001111110000000000000000000000
000000000000000000000000000000000000000000000000000000000000
000001111111100000000000000000000000000000000000000000000000
00000000000000000000011
>DM_train424
AQVQQLTPAQQAALRNQQAMAANLQARQIVLQQSYPVIQQVETQTFDPANRSVFDVTPANVG
IVKGFLVKVTAAITNNHATEAVALTDFGPANLVQRVIYYDPDNQRHTETSGWHLHFVNTAKQ
GAPFLSSMVTDSPIKYGDVMNVIDAPATIAAGATGELTMYYWVPLAYSETDLTGAVLANVPQ
SKQRLKLEFANNNTAFAAVGANPLEAIYQGAGAADCEFEEISYTVYQSYLDQLPVGQNGYIL
PLIDLSTLYNLENSAQAGLTPNVDFVVQYANLYRYLSTIAVFDNGGSFNAGTDINYLSQRTA
NFSDTRKLDPKTWAAQTRRRIATDFPKGVYYCDNRDKPIYTLQYGNVGFVVNPKTVNQNARL
LMGYEYFTSRTELVNAGTISTT
111111111111100000000000000000000000000000000000000000000000
000000000000000000000000000000000000000000000000000000000000
000000000000000000000000000000000000000000000000000000000000
000000000000000000000000000000000000000000000000000000000000
000000000000000000000000000000000000000000000000000000000000
000000000000000000000000000000000000000000000000000000000000

0000000000111111111111
>DM_train425
APLASVPTPAFPFPVPTMVHGPCVAESEPALLTGSKQFGLSRNSHIAIAFDDTKVKNRLTIE
LEVRTEAESGLLFYMARINHADFATVQLRNGFPYFSYDLGSGDTSTMIPTKINDGQWHKIKI
VRVKQEGILYVDDASSQTISPKKADILDVVGILYVGGLPINYTTRRIGPVTYSLDGCVRNLH
MEQAPVDLDQPTSSFHVGTCFANAESGTYFDGTGFAKAVGGFKVGLDLLVEFEFRTTRPTGV
LLGVSSQKMDGMGIEMIDEKLMFHVDNGAGRFTAIYDAEIPGHMCNGQWHKVTAKKIKNRLE
LVVDGNQVDAQSPNSASTSADTNDPVFVGGFPGGLNQFGLTTNIRFRGCIRSLKLTKGTGKP
LEVNFAKALELRGVQPVSCPTT
111111111111111111100000000000000000000000000000000000000000
000000000000000000000000000000000000000000000000000000000000
000000000000000000000000000000000000000000000000000000000000
000000000000000000000000000000000000000000000000000000000000
000000000000000000000000000000000000000000000000000000000000
000000000000000000000000000000000000000000000000000000000000
0000000000000000000000
>DM_train426
MKTQVAIIGAGPSGLLLGQLLHKAGIDNVILERQTPDYVLGKIRAGVLEQGMVDLLREAGVD
RRMARDGLVHEGVEIAFAGQRRRIDLKRLSGGKTVTVYGQTEVTRDLMEAREASGATTVYQA
AEVRLHDLQGERPYVTFERDGERLRLDCDYIAGCDGFHGISRQSIPAERLKVFERVYPFGWL
GLLADTPPVSHELIYANHPRGFALCSQRSATRSRYYVQVPLTEKVEDWSDERFWTELKARLP
AEVAEKLVTGPSLEKSIAPLRSFVVEPMQHGRLFLAGDAAHIVPPTGAKGLNLAASDVSTLY
RLLLKAYREGRGELLERYSAICLRRIWKAERFSWWMTSVLHRFPDTDAFSQRIQQTELEYYL
GSEAGLATIAENYVGLPYEEIE
000000000000000000000000000000000000000000000000000000000000
000000000000000000000000000000000000000000000000000000000000
000000000000000000000000000000000000000000000000000000000000
000000000000000000000000000000000000000000000000000000000000
000000000000000000000000000000000000000000000000000000000000
000000000000000000000000000000000000000000000000000000000000
0000000000000000000111
>DM_train428
GMSSSDRYMQRGVSSQKEDVHKAIKSIDKGIYPRAFCKIIPDILGGDPEYCNIMHADGAGTK
SSLAYVYWKETGDISVWKGIAQDAVIMNIDDLICVGAVDNILLSSTIGRNKNLIPGEVLAAI
INGTEEVLQMLRDNGIGIYSTGGETADVGDLVRTIIVDSTVTCRMKRQDVISNENIKAGNVI
VGFASYGQTSYETEYNGGMGSNGLTSARHDVFNNVLASKYPESFDPKVPENLVYSGEMNLTD
PYLNVPLDAGKLVLSPTRTYAPLMKEIIHQYKGKLDGVVHCSGGGQTKVLHFTDATTHIIKD
NLFDVPPLFQLIQGQSNTPWEEMYKVFNMGHRLEIYTDAAHAEGMIAIAKKFNIEAKIIGRV
EAPVAGKRLTITGPQGTEYTYA
111111111111111111111111111000000000000000000000000000000000
000000000000000000000000000000000000000000000000000000000000
000000000000000000000000000000000000000000000000000000000000
000000000000000000000000000000000000000000000000000000000000
000000000000000000000000000000000000000000000000000000000000

000000000000000000000000000000000000000000000000000000000000
0000000000000000000000
>DM_train429
MRCIGISNRDFVEGVSGGSWVDIVLEHGSCVTTMAKNKPTLDFELIKTEAKQPATLRKYCIE
AKLTNTTTESRCPTQGEPTLNEEQDKRFVCKHSMVDRGWGNGCGLFGKGGIVTCAMFTCKKN
MEGKIVQPENLEYTVVITPHSGEEHAVGNDTGKHGKEVKITPQSSITEAELTGYGTVTMECS
PRTGLDFNEMVLLQMKDKAWLVHRQWFLDLPLPWLPGADTQGSNWIQKETLVTFKNPHAKKQ
DVVVLGSQEGAMHTALTGATEIQMSSGNLLFTGHLKCRLRMDKLQLKGMSYSMCTGKFKVVK
EIAETQHGTIVIRVQYEGDGSPCKIPFEIMDLEKRHVLGRLITVNPIVTEKDSPVNIEAEPP
FGDSYIIIGVEPGQLKLNWFKK
000000000000000000000000000000000000000000000000000000000000
000000000000000000000000000000000000000000000000000000000000
000000000000000000011111111111111000000000000000000000000000
000000000000000000000000000000000000000000000000000000000000
000000000000000000000000000000000000000000000000000000000000
000000000000000000000000000000000000000000000000000000000000
0000000000000000000000
>DM_train430
MKLEIFSWWAGDEGPALEALIRLYKQKYPGVEVINATVTGGAGVNARAVLKTRMLGGDPPDT
FQVHAGMELIGTWVVANRMEDLSALFRQEGWLQAFPKGLIDLISYKGGIWSVPVNIHRSNVM
WYLPAKLKGWGVNPPRTWDKFLATCQTLKQKGLEAPLALGENWTQQHLWESVALAVLGPDDW
NNLWNGKLKFTDPKAVRAWEVFGRVLDCANKDAAGLSWQQAVDRVVQGKAAFNIMGDWAAGY
MTTTLKLKPGTDFAWAPSPGTQGVFMMLSDSFGLPKGAKNRQNAINWLRLVGSKEGQDTSNP
LKGSIAARLDSDPSKYNAYGQSAMRDWRSNRIVGSLVHGAVAPESFMSQFGTVMEIFLQTRN
PQAAANAAQAIADQVGLGRLGQ
000000000000000000000000000000000000000000000000000000000000
000000000000000000000000000000000000000000000000000000000000
000000000000000000000000000000000000000000000000000000000000
000000000000000000000000000000000000000000000000000000000000
000000000000000000000000000000000000000000000000000000000000
000000000000000000000000000000000000000000000000000000000000
0000000000000000000111
>DM_train431
MGSSSSGRENLYFQGMASIVFSTIGNPKGYQKVTYEIDGEKFESNVSVLALRDLLKVDKTVV
ILGISVADVYNCKYADYRSCKECIIQNSKNDLGISESYVVAPNVYQKFKGKPDHYFTYIYYH
SLRILEKEGINEVFIDTTHGINYMGVLAKEAIQLAVSAYAAKSEKEVKVSLYNSDPVGKDVS
DTVKLHEIEAIKISPLSGLKYVVTYQILNKDKNFFNKIFSDSVNAIPRFATALDNGLFIYLSE
KDSSLHLKRLEDDLSKDPLLTPSENEINVVYKDMKYALSHALFYVISRFSGNVDLDTLRHYA
ETYADKVTRAIIENEVDKIEKYQMGSERKLLGEYMKVEGKGFDKRILYAHGGLPYAGTYVYK
EKDKVYVTYGDKIDEIERQIGS
111111111111000000000000000000000000000000000000000000000000
000000000000000000000000000000000000000000000000000000000000
000000000000000000000000000000000000000000000000000000000000
000000000000000000000000000000000000000000000000000000000000

000000000000000000000000000000000000000000000000000000000000
000000000000000000000000000000000111111111000000000000000000
0000000000000000000011
>DM_train432
IESLCMNCYRNGTTRLLLTKIPFFREIIVSSFSCEHCGWNNTEIQSAGRIQDQGVRYTLTVR
SQEDMNREVVKTDSATTRIPELDFEIPAFSQKGALTTVEGLISRAISGLEQDQPTRRAVEGA
IAERIDEFIGKLKDLKQMASPFTLVIDDPSGNSFVENPHAPQKDNALVITYYDRTPQQAEML
GLQAEAPEEKAEEEDLRNEVLQFNTNCPECNAPAQTNMKLVQIPHFKEVIIMATNCENCGHR
TNEVKSGGAVEPLGTRITLHITDPSDMTRDLLKSETCSVEIPELEFELGMAVLGGKFTTLEG
LLKDIRELVTKNPFTLGDSSNPDQSEKLQEFSQKLGQIIEGKMKAHFIMNDPAGNSYLQNVY
APEDDPEMKVERYKRTFDQNEE
000000000000000000000000000000000000000000000000000000000000
000000000000000000000000000000000000000000000000000000000000
000000000000000000000000000000000000000000000000000000000000
001111111111000000000000000000000000000000000000000000000000
000000000000000000000000000000000000000000000000000000000000
000000000000000000000000000000000000000000000000000000000000
0000000000000000000000
>DM_train433
MASMQKLINSVQNYAWGSKTALTELYGIANPQQQPMAELWMGAHPKSSSRITTANGETVSLR
DAIEKNKTAMLGEAVANRFGELPFLFKVLCAAQPLSIQVHPNKRNSEIGFAKENAAGIPMDA
AERNYKDPNHKPELVFALTPFLAMNAFREFSDIVSLLQPVAGAHSAIAHFLQVPNAERLSQL
FASLLNMQGEEKSRALAVLKAALNSQQGEPWQTIRVISEYYPDDSGLFSPLLLNVVKLNPGE
AMFLFAETPHAYLQGVALEVMANSDNVLRAGLTPKYIDIPELVANVKFEPKPAGELLTAPVK
SGAELDFPIPVDDFAFSLHDLALQETSIGQHSAAILFCVEGEAVLRKDEQRLVLKPGESAFI
GADESPVNASGTGRLARVYNKL
000000000000000000000000000000000000000000000000001111100000
000000000000000000000000000000000000000000000000000000000000
000000000000000000000000000000000000000000000000000000000000
000000000000000000000000000000000000000000000000000000000000
000000000000000000000000000000000000000000000000000000000000
000000000000000000000000000000000000000000000000000000000000
0000000000000000000000
>DM_train434
MLLEAPVYKEIFGAVTIHEVQKVIKMDTETEEVPIYTISNIPREKIYDLLGKMAVIVPMKNE
KLHLVDGVLKAIPHKCPIIIVSNSKREGPNRYKLEVDLIRHFYNLTHSKIIMIHQKDPGLAK
AFKEVGYTDILDENGMIRSGKGEGMLVGLLLAKAIGAEYVGFVDADNYIPGAVNEYVKDYAA
GFLMSESEYTMVRLHWRHKPKVTKGTLYFKKWGRVSEITNHYLNLLVSEHTAFETTIMVTGN
AGEHAMTMKLAEILPFSTGYSIEPYEIVYILERFGKWENVEEFKDVFDQGIEIFQIETLNPH
FHEDKGKEHVKEMLLLSLATIYHSKLATDNLRKRILKDLRDHGILGENEEPPKPLVMRPIKE
IPIKEWMDIVEGNSETLLRFEL
000000000000000000000000000011111111000000000000000000000000
000000000000000000000000000000000000000000000000000000000000
000000000000000000000000000000000000000000000000000000000000

00000000000000001111111111111111110000000000000000000000000000
00000000000000000000000000000000000000000000000000000000000000
00000000000000000000000000000000000000011111111000000000000000
0000000000000000000000
>DM_train435
MSNPTKISILGRESIIADFGLWRNYVAKDLISDCSSTTYVLVTDTNIGSIYTPSFEEAFRKR
AAEITPSPRLLIYNRPPGEVSKSRQTKADIEDWMLSQNPPCGRDTVVIALGGGVIGDLTGFV
ASTYMRGVRYVQVPTTLLAMVDSSIGGKTAIDTPLGKNLIGAIWQPTKIYIDLEFLETLPVR
EFINGMAEVIKTAAISSEEEFTALEENAETILKAVRREVTPGEHRFEGTEEILKARILASAR
HKAYVVSADEREGGLRNLLNWGHSIGHAIEAILTPQILHGECVAIGMVKEAELARHLGILKG
VAVSRIVKCLAAYGLPTSLKDARIRKLTAGKHCSVDQLMFNMALDKKNDGPKKKIVLLSAIG
TPYETRASVVANEDIRVVLAP
11100000000000000000000000000000000000000000000000000000000000
00000000000000000000000000000000000000000000000000000000000000
00000000000000000000000000000000000000000000000000000000000000
00000000000000000000000000000000000000000000000000000000000000
00000000000000000000000000000000000000000000000000000000000000
00000000000000000000000000000000000000000011111111111110000000
000000000000000000000
>DM_train436
MAEGKMMSDFLPFSRPAMGAEELAAVKTVLDSGWITTGPKNQELEAAFCRLTGNQYAVAVSS
ATAGMHIALMALGIGEGDEVITPSMTWVSTLNMIVLLGANPVMVDVDRDTLMVTPEHIEAAI
TPQTKAIIPVHYAGAPADLDAIYALGERYGIPVIEDAAHATGTSYKGRHIGARGTAIFSFHA
IKNITCAEGGIVVTDNPQFADKLRSLKFHGLGVDAWDRQSGGRAPQAEVLAPGYKYNLPDLN
AAIALAQLQKLDALNARRAAIAAQYHQAMADLPFQPLSLPSWEHIHAWHLFIIRVDEARCGI
TRDALMASLKTKGIGTGLHFRAAHTQKYYRERFPTLTLPDTEWNSERICSLPLFPDMTESDF
DRVITALHQIAGQGSHHHHHH
11111111000000000000000000000000000000000000000000000000000000
00000000000000000000000000000000000000000000000000000000000000
00000000000000000000000000000000000000000000000000000000000000
00000000000000000000000000000000001111111111100000000000000000
00000000000000000000000000000000000000000000000000000000000000
00000000000000000000000000000000000000000000000000000000000000
000000000001111111111
>DM_train437
GAKSEMAFAGKGEISPRAITMWDFSWLERRWPGAGYEDWDQVLDELSERGYNAIRIDAYPHL
IAENPMKKWLLKEVWNQQDWGSPDMNEVQVQPNLNLFLSKCKERDIKVGLSSWYRLDVDEVC
LKLDTPEKLADCWLTILRSIEEDGLLDTILYVDLCNEWPGDSWAPFFAKTYPNVGWGNWYKE
ESLRWMKTSLEKMRQVYPDMPFLYSFDHGDVKKYEEVDCSFLDLYEHHIWMAQQNGGEFYKL
VGYGYNRFLPDDYKNVVKNAERVYRERPGYWQKLLTDKIELMASVARKNRRPLVTTECWGLV
DYKDWPLLKWDWVKDLCELGTITAARTGMWVGVATSNFCGPQFAGMWRDVEWHKRLTSIIRS
SPLDESLTKNNEVAAKLLKRL
11111111111100000000000000000000000000000000000000000000000000
00000000000000000000000000000000000000000000000000000000000000

00000000000000000000000000000000000000000000000000000000000000
00000000000000000000000000000000000000000000000000000000000000
00000000000000000000000000000000000000000000000000000000000000
00000000000000000000000000000000000000000000000000000000000000
000000000000000000000
>DM_train438
RIGIPRERLTNETRVAATPKTVEQLLKLGFTVAVESGAGQLASFDDKAFVQAGAEIVEGNSV
WQSEIILKVNAPLDDEIALLNPGTTLVSFIWPAQNPELMQKLAERNVTVMAMDSVPRISRAQ
SLDALSSMANIAGYRAIVEAAHEFGRFFTGQITAAGKVPPAKVMVIGAGVAGLAAIGAANSL
GAIVRAFDTRPEVKEQVQSMGAEFLELDFKEEAGSGDGYAKVMSDAFIKAEMELFAAQAKEV
DIIVTTALIPGKPAPKLITREMVDSMKAGSVIVDLAAQNGGNCEYTVPGEIFTTENGVKVIG
YTDLPGRLPTQSSQLYGTNLVNLLKLLCKEKDGNITVDFDDVVIRGVTVIRAGEITWPAPPI
QVSAQPQAAQKAAPEVKTEEK
00000000000000000000000000000000000000000000000000000000000000
00000000000000000000000000000000000000000000000000000000000000
00000000000000000000000000000000000000000000000000000000000000
00000000000000000000000000001111111111111100000000000000000000
00000000000000000000000000000000000000000000000000000000000000
00000000000000000000000000000000000000000000000000000000000000
000011111111111111111
>DM_train439
ASASTPHALLLISIDGLRADMLDRGITPNLSHLAREGVRARWMAPSYPSLTFPNHYTLVTGL
RPDHHGIVHNSMRDPTLGGFWLSKSEAVGDARWWGGEPVWVGVENTGQHAATWSWPGSEAAI
KGVRPSQWRHYQKGVRLDTRVDAVRGWLATDGAQRNRLVTLYFEHVDEAGHDHGPESRQYAD
AVRAVDAAIGRLLAGMQRDGTRARTNIIVVSDHGMAEVAPGHAISVEDIAPPQIATAITDGQ
VIGFEPLPGQQAAAEASVLGAHDHYDCWRKAELPARWQYGSHPRIPSLVCQMHEGWDALFPD
KLAKRAQRGTRGSHGYDPALPSMRAVFLAQGPDLAQGKTLPGFDNVDVYALMSRLLGIPAAP
NDGNPATLLPALRMPPAPDAR
11110000000000000000000000000000000000000000000000000000000000
00000000000000000000000000000000000000000000000000000000000000
00000000000000000000000000000000000000000000000000000000000000
00000000000000000000000000000000000000000000000000000000000000
00000000000000000000000000000000000000000000000000000000000000
00000000000000000000000000000000000000000000000000000000000000
000000000000001111111
>DM_train441
MTAASLDPTAFSLDAASLAARLDAVFDQALRERRLVGAVAIVARHGEILYRRAQGLADREAG
RPMREDTLFRLASVTKPIVALAVLRLVARGELALDAPVTRWLPEFRPRLADGSEPLVTIHHL
LTHTSGLGYWLLEGAGSVYDRLGISDGIDLRDFDLDENLRRLASAPLSFAPGSGWQYSLALD
VLGAVVERATGQPLAAAVDALVAQPLGMRDCGFVSAEPERFAVPYHDGQPEPVRMRDGIEVP
LPEGHGAAVRFAPSRVFEPGAYPSGGAGMYGSADDVLRALEAIRANPGFLPETLADAARRDQ
AGVGAETRGPGWGFGYLSAVLDDPAAAGTPQHAGTLQWGGVYGHSWFVDRALGLSVLLLTNT
AYEGMSGPLTIALRDAVYAR
11111111111111000000000000000000000000000000000000000000000000

000000000000000000000000000000000000000000000000000000000000
000000000000000000000000000000000000000000000000000000000000
000000000000000000000000000000000000000000000000000000000000
000000000000000000000000000000000000000000000000000000000000
000000000000000000000000000000000000000000000000000000000000
00000000000000000000
>DM_train442
MATALYTANDFILISLPQNAQPVTAPGSKTDSWFNETLIGGRAFVSDFKIPEFKIGSLDTLI
VESEELSKVDNQIGASIGKIIEILQGLNETSTNAYRTLPINNMPVPEYLENFQWQTRKFKLD
KSIKDLITLISNESSQLDADVRATYANYNSAKTNLAAAERKKTGDLSVRSLHDIVKPEDFVL
NSEHLTTVLVAVPKSLKSDFEKSYETLSKNVVPASASVIAEDAEYVLFNVHLFKKNVQEFTT
AAREKKFIPREFNYSEELIDQLKKEHDSAASLEQSLRVQLVRLAKTAYVDVFINWFHIKALR
VYVESVLRYGLPPHFNIKIIAVPPKNLSKCKSELIDAFGFLGGNAFMKDKKGKINKQDTSLH
QYASLVDTEYEPFVMYIINL
111100000000000000000000000000000000000000000000000000000000
000000000000000000000000000000000000000000000000000000000000
000000000000000000000000000000000000000000000000000000000000
000000000000000000000000000000000000000000000000000000000000
000000000000000000000000000000000000000000000000000000000000
000000000000000000000000000000000000000000000111111111111111
11111111100000000000
>DM_train443
GAMGSMKPYLFDLKLKDTEKLDWKKGLSSYLKKSYGSSQWRTFYDEKATSELDHLRNNANGE
LAPSSLSEQNLKYYSFLEHLYFRLGSKGSRLKMDFTWYDAEYSSAQKGLKYTQHTLAFEKSC
TLFNIAVIFTQIARENINEDYKNSIANLTKAFSCFEYLSENFLNSPSVDLQSENTRFLANIC
HAEAQELFVLKLLNDQISSKQYTLISKLSRATCNLFQKCHDFMKEIDDDVAIYGEPKWKTTV
TCKLHFYKSLSAYYHGLHLEEENRVGEAIAFLDFSMQQLISSLPFKTWLVEFIDFDGFKETL
EKKQKELIKDNDFIYHESVPAVVQVDSIKALDAIKSPTWEKILEPYMQDVANKYDSLYRGII
PLDVYEKESIYSEEKATLLR
111110000000000000000000000000000000000000000000000000000000
000000000000000000000000000000000000000000000000000000000000
000000000000000000000000000000000000000000000000000000000000
000000000000000000000000000000000000000000000000000000000000
000000000000000000000000000000000000000000000000000000000000
000000000000000000000000000000000000000000000000000000000000
11111111111111111111
>DM_train444
MNARSTGQHPARYPGAAAGEPTLDSWQEPPHNRWAFAHLGEMVPSAAVSRRPVNAPGHALAR
LGAIAAQLPDLEQRLEQTYTDAFLVLRGTEVVAEYYRAGFAPDDRHLLMAVSKSLCGTVVGA
LVDEGRIDPAQPVTEYVPELAGSVYDGPSVLQVLDMQISIDYNEDYVDPASEVQTHDRSAGW
RTRRHGDPADTYEFLTTLRGDGSTGEFQYCSANTDVLAWIVERVTGLRYVEALSTYLWAKLD
ADRDATITVDTTGFGFANGGVSCTARDLARVGRMMLDGGVAPGGRVVSEDWVRRVLAGGSHE
AMTDKGFTNTFPDGSYTRQWWCTGNERGNVSGIGIHGQNLWLDPLTDSVIVKLSSWPDPDTE
HWHRLQNGILLDVSRALDAV

1111000000000000000000000000000000000000000000000011110000000
000000000000000000000000000000000000000000000000000000000000
000000000000000000000000000000000000000000000000000000000000
000000000000000000000000000000000000000000000000000000000000
000000000000000000000000000000000000000000000000000000000000
000000000000000000000000000000000000000000000000000000000000
00000000000000000000
>DM_train445
TSTATDSPLKYYDIGLNLTDPMFHGIYNGKQYHPADYVKLLERAAQRHVKNALVTGSSIAES
QSAIELVSSVKDLSPLKLYHTIGVHPCCVNEFADASQGDKASASIDNPSMDEAYNESLYAKV
ISNPSFAQGKLKELYDLMNQQAKPHDTSFRSIGEIGLDYDRFHYSSKEMQKVFFEEQLKISC
LNDKLSSYPLFLHMRSACDDFVQILERFVVGFTDEKDTFQLQKLGASSSSGFYKFHPDRKLV
VHSFTGSAIDLQKLLNLSPNIFIGVNGCSLRTEENLAVVKQIPTERLLLETDAPWCEIKRTH
ASFQYLAKYQEVRDFEYPAFKSVKKNKLADKLNAEELYMVKGRNEPCNMEQVAIVVSEVKDV
DLATLIDTTWKTTCKIFGE
111111000000000000000000000000000000000000000000000000000000
000000000000000000000000000000011111111111111111100000000000
000000000000000000000000000000000000000000000000000000000000
000000000000000000000000000000000000000000110000000000000000
000000000000000000000000000000000000000000000000000000000000
000000000000000000000000000000000000000000000000000000000000
0000000000000000011
>DM_train446
TSKNQFYKPGMLVPVFSAAGLLKNGRYQFLLQQIETLSLLPTEQYAQLYEALVYRFVEFVQV
LPIRLDEPLCSLMNEGLLRGVNSLNHYIQNHPEATPLERYALFSAGLLLEVAHAVVNQKIFI
TDEEGNFIKQWNPFSGPLIDDVETKHYKIMPLSSYYQRNIPSITPILVRQLLPDEGFLWLTS
DMRVFSDWMQALRDDEGEGGGRFEHVLQLFKHKNIDGLFNTLPALPVNLQDSPATAHADAFL
NWLKEALATNQIKVNTSDAGVHVIPEGVFLEKTGIFKQYIDLHVNVPVNLFTVYQQFGNLFG
LTKLSGIDYRFEQLFSEYPDALKRKSKMGFAGLIGIRRASPTREGVLIADPNLIFTRGEIPS
ATSYLKLSSAQKSQNIPMT
111000000000000000000000000000000000000000000000000000000000
000000000000000000000000000000000000000000000000000000000000
000000000000000000000000000000000000000000000000000000000000
000000000000000111110000000000000000000000000000000000000000
000000000000000000000000000000000000000000000000000000000000
000000000000000111100000000000000111111000000000000000000000
0010000111111111111
>DM_train447
PEFFSVRHLELAGDDPYSNVNCTKILQGDPEEIQKVKLEILTVQFKKRPRWTPHDYINMTRD
CASFIRTRKYIVEPLTKEEVGFPIAYSIVVHHKIEMLDRLLRAIYMPQNFYCIHVDRKAEES
FLAAVQGIASCFDNVFVASQLESVVYASWTRVKADLNCMKDLYRMNANWKYLINLCGMDFPI
KTNLEIVRKLKCSTGENNLETEKMPPNKEERWKKRYAVVDGKLTNTGIVKAPPPLKTPLFSG
SAYFVVTREYVGYVLENENIQKLMEWAQDTYSPDEFLWATIQRIPEVPGSFPSSNKYDLSDM
NAIARFVKWQYFEGDVSNGAPYPPCSGVHVRSVCVFGAGDLSWMLRQHHLFANKFDMDVDPF

AIQCLDEHLRRKALENLEH
111111000001111111000000000000000001111111111000000000000000
000000000000000000000000000000000000000000000000000000000000
000000000000000000000000000000000000000000000000000000000000
000000000000000000000000000000000000000000000000000000000000
000000000000000000000000000000000000000000000000000000000000
000000000000000000000000000000000000000000000000000000000000
0000000000000001111
>DM_train448
YEHSTVMPNVVGFPYKAHIERPGYSPLTLQMQVVETSLEPTLNLEYITCEYKTVVPSPYVKC
CGASECSTKEKPDYQCKVYTGVYPFMWGGAYCFCDSENTQLSEAYVDRSDVCRHDHASAYKA
HTASLKAKVRVMYGNVNQTVDVYVNGDHAVTIGGTQFIFGPLSSAWTPFDNKIVVYKDEVFN
QDFPPYGSGQPGRFGDIQSRTVESNDLYANTALKLARPSPGMVHVPYTQTPSGFKYWLKEKG
TALNTKAPFGCQIKTNPVRAMNCAVGNIPVSMNLPDSAFTRIVEAPTIIDLTCTVATCTHSS
DFGGVLTLTYKTNKNGDCSVHSHSNVATLQEATAKVKTAGKVTLHFSTASASPSFVVSLCSA
RATCSASCEPPKDHIVPY
000000000000000000000000000000000000000000000000000000000000
000000000000000000000000000000000000000000000000000000000000
000000000000000000000000000000000000000000000000000000000000
000000000000000000000000000000000000000000000000000000000000
000000000000000000000000000000000000000000000000000000000001
111100000000000000000000000000000011111100000000000000000000
000000001111111111
>DM_train449
MIDYTAAGFTLLQGAHLYAPEDRGICDVLVANGKIIAVASNIPSDIVPNCTVVDLSGQILCP
GFIDQHVHLIGGGGQAGPTTRTPEVALSRLTEAGVTSVVGLLGTDSISRHPESLLAKTRALN
EEGISAWMLTGAYHVPSRTITGSVEKDVAIIDRVIGVKCAISDHRSAAPDVYHLANMAAESR
VGGLLGGKPGVTVFHMGDSKKALQPIYDLLENCDVPISKLLPTHVNRNVPLFEQALEFARKG
GTIDITSSIDEPVAPAEGIARAVQAGIPLARVTLSSDGNGSQPFFDDEGNLTHIGVAGFETL
LETVQVLVKDYDFSISDALRPLTSSVAGFLNLTGKGEILPGNDADLLVMTPELRIEQVYARG
KLMVKDGKACVKGTFETA
000000000000000000000000000000000000000000000000000000000000
000000000000000000000000000000000000000000000000000000000000
000000000000000000000000000000000000000000000000000000000000
000000000000000000000000000000000000000000000000000000000000
000000000000000000000000000000000000000111111111111110000000
000000000000000000000000000000000000000000000000000000000000
000000000000000001
>DM_train450
MKSKFKLTTAAAMLGLMVLAGGAQAQDKPREVLTGGHSVSAPQENRIYVMDSVFMHLTESRV
HVYDYTNGKFLGMVPTAFNGHVQVSNDGKKIYTMTTYHERITRGKRSDVVEVWDADKLTFEK
EISLPPKRVQGLNYDGLFRQTTDGKFIVLQNASPATSIGIVDVAKGDYVEDVTAAAGCWSVI
PQPNRPRSFMTICGDGGLLTINLGEDGKVASQSRSKQMFSVKDDPIFIAPALDKDKAHFVSY
YGNVYSADFSGDEVKVDGPWSLLNDEDKAKNWVPGGYNLVGLHRASGRMYVFMHPDGKEGTH

KFPAAEIWVMDTKTKQRVARIPGRDALSMTIDQQRNLMLTLDGGNVNVYDISQPEPKLLRTI
EGAAEASLQVQFHPVGGV
111111111111111111111111111000000000000000000000000000000000
000000000000000000000000000000000000000000000000000000000000
000000000000000000000000000000000000000000000000000000000000
000000000000000000000000000000000000000000000000000000000000
000000000000000000000000000000000000000000000000000000000000
000000000000000000000000000000000000000000000000000000000000
000000000000000111
>DM_train451
MTESYDVVVVGGGPVGLATAWQVAERGHRVLVLERHTFFNENGGTSGAERHWRLQYTQEDLF
RLTLETLPLWRALESRCERRLIHEIGSLWFGDTDVVTNEGQISGTAAMMDKLSVRYEWLKAT
DIERRFGFRGLPRDYEGFLQPDGGTIDVRGTLAALFTLAQAAGATLRAGETVTELVPDADGV
SVTTDRGTYRAGKVVLACGPYTNDLLEPLGARLAYSVYEMAIAAYRQATPVTEAPFWFAFQQ
PTPQDTNLFYGFGHNPWAPGEFVRCGPDFEVDPLDHPSAATGVADRRQMDRLSGWLRDHLPT
VDPDPVRTSTCLAVLPTDPERQFFLGTARDLMTHGEKLVVYGAGWAFKFVPLFGRICADLAV
EDSTAYDISRLAPQSAL
110000000000000000000000000000000000000000000000000000000000
000000000000000000000000000000000000000000000000000000000000
000000000000000000000000000000000000000000000000000000000000
000000000000000000000000000000000000000000000000000000000000
000000000000000000000000000000000000000000000000000000000000
000000000000000000000000000000000000000000000000000000000000
00000000000000011
>DM_train452
ASPRANDAPIVLLHGFTGWGREEMLGFKYWGGVRGDIEQWLNDNGYRTYTLAVGPLSSNWDR
ACEAYAQLVGGTVDYGAAHAAKHGHARFGRTYPGLLPELKRGGRVHIIAHSQGGQTARMLVS
LLENGSQEEREYAKEHNVSLSPLFEGGHRFVLSVTTIATPHDGTTLVNMVDFTDRFFDLQKA
VLEAAAVASNAPYTSEIYDFKLDQWGLRREPGESFDHYFERLKRSPVWTSTDTARYDLSVPG
AETLNRWVKASPNTYYLSFSTERTYRGALTGNYYPELGMNAFSAIVCAPFLGSYRNAALGID
SHWLGNDGIVNTISMNGPKRGSNDRIVPYDGTLKKGVWNDMGTYNVDHLEVIGVDPNPSFNI
RAFYLRLAEQLASLRP
111000000000000000000000000000000000000000000000000000000000
000000000000000000000000000000000000000000000000000000000000
000000000000000000000000000000000000000000000000000000000000
000000000000000000000000000000000000000000000000000000000000
000000000000000000000000000000000000000000000000000000000000
000000000000000000000000000000000000000000000000000000000000
0000000000000000
>DM_train453
MNLHEYQAKQLFARYGLPAPVGYACTTPREAEEAASKIGAGPWVVKCQVHAGGRGKAGGVKV
VNSKEDIRAFAENWLGKRLVTYQTDANGQPVNQILVEAATDIAKELYLGAVVDRSSRRVVFM
ASTEGGVEIEKVAEETPHLIHKVALDPLTGPMPYQGRELAFKLGLEGKLVQQFTKIFMGLAT
IFLERDLALIEINPLVITKQGDLICLDGKLGADGNALFRQPDLREMRDQSQEDPREAQAAQW

ELNYVALDGNIGCMVNGAGLAMGTMDIVKLHGGEPANFLDVGGGATKERVTEAFKIILSDDK
VKAVLVNIFGGIVRCDLIADGIIGAVAEVGVNVPVVVRLEGNNAELGAKKLADSGLNIIAAK
GLTDAAQQVVAAVEGK
000000000000000000000000000000000000000000000000000000000000
000000000000000000000000000000000000000000000000000000000000
000000000000000000000000000000000000000000000000000000000000
000000000000000000000000000000000000000000000000000000000000
000000000000000000000000000000000000000000000000000000000000
000000000000000000000000000000000000000000000000000000000000
0000000000000111
>DM_train455
MEKTTEGAFYTREYRNLFKEFGYSEAEIQERVKDTWEQLFGDNPETKIYYEVGDDLGYLLDT
GNLDVRTEGMSYGMMMAVQMDRKDIFDRIWNWTMKNMYMTEGVHAGYFAWSCQPDGTKNSWG
PAPDGEEYFALALFFASHRWGDGDEQPFNYSEQARKLLHTCVHNGEGGPGHPMWNRDNKLIK
FIPEVEFSDPSYHLPHFYELFSLWANEEDRVFWKEAAEASREYLKIACHPETGLAPEYAYYD
GTPNDEKGYGHFFSDSYRVAANIGLDAEWFGGSEWSAEEINKIQAFFADKEPEDYRRYKIDG
EPFEEKSLHPVGLIATNAMGSLASVDGPYAKANVDLFWNTPVRTGNRRYYDNCLYLFAMLAL
SGNFKIWFPEGQEEEH
111110000000000000000000000000000000000000110000000000000000
000000000000000000000000000000000000000000000000000000000000
000000000000000000000000000000000000000000000000000000000000
000000000000000000000000000000000000000000000000000000000000
000000000000000000000000000000000000000000000000000000000000
000000000000000000000000000000000000000000000000000000000000
0000000001111111
>DM_train456
MAGSATVEKRLDFGLLGPLQMTIDGTPVPSGTPKQRAVLAMLVINRNRPVGVDALITALWEE
WPPSGARASIHSYVSNLRKLLGGAGIDPRVVLAAAPPGYRLSIPDNTCDLGRFVAEKTAGVH
AAAAGRFEQASRHLSAALREWRGPVLDDLRDFQFVEPFATALVEDKVLAHTAKAEAEIACGR
ASAVIAELEALTFEHPYREPLWTQLITAYYLSDRQSDALGAYRRVKTTLADDLGIDPGPTLR
ALNERILRQQPLDAKKSAKTTAAGTVTVLDQRTMASGQQAVAYLHDIASGRGYPLQAAATRI
GRLHDNDIVLDSANVSRHHAVIVDTGTNYVINDLRSSNGVHVQHERIRSAVTLNDGDHIRIC
DHEFTFQISAGTHGGT
111111100000000000000000000000000000000000000000000000000000
000000000000000000000000000000000000000000000000000000000000
000000000000000000000000000000000000000000000000000000000000
000000000000000000000000000000000000000000000000000000000000
000000000000000000000000000000000000000000000000000000000000
000000000000000000000000000000000000000000000000000000000000
0000000011111111
>DM_train457
MGKNLQALAQLYKNALLNDVLPFWENHSLDSEGGYFTCLDRQGKVYDTDKFIWLQNRQVWTF
SMLCNQLEKRENWLKIARNGAKFLAQHGRDDEGNWYFALTRGGEPLVQPYNIFSDCFAAMAF
SQYALASGEEWAKDVAMQAYNNVLRRKDNPKGKYTKTYPGTRPMKALAVPMILANLTLEMEW

LLPQETLENVLAATVQEVMGDFLDQEQGLMYENVAPDGSHIDCFEGRLINPGHGIEAMWFIM
DIARRKNDSKTINQAVDVVLNILNFAWDNEYGGLYYFMDAAGHPPQQLEWDQKLWWVHLESL
VALAMGYRLTGRDACWAWYQKMHDYSWQHFADPEYGEWFGYLNRRGEVLLNLKGGKWKGCFH
VPRAMYLCWQQFEALS
11000000000000000000000000000000000000000000000000000000000000
00000000000000000000000000000000000000000000000000000000000000
00000000000000000000000000111111111111110000000000000000000000
00000000000000000000000000000000000000000000000000000000000000
00000000000000000000000000000000000000000000000000000000000000
00000000000000000000000000000000000000000000000000000000000000
0000000000000000
>DM_train458
NSIQIGGLFPRGADQEYSAFRVGMVQFSTSEFRLTPHIDNLEVANSFAVTNAFCSQFSRGVY
AIFGFYDKKSVNTITSFCGTLHVSFITPSFPTDGTHPFVIQMRPDLKGALLSLIEYYQWDKF
AYLYDSDRGLSTLQAVLDSAAEKKWQVTAINVGNINNDKKDEMYRSLFQDLELKKERRVILD
CERDKVNDIVDQVITIGKHVKGYHYIIANLGFTDGDLLKIQFGGANVSGFQIVDYDDSLVSK
FIERWSTLEEKEYPGAHTTTIKYTSALTYDAVQVMTEAFRNLRKQRIEISRRGNAGDCLANP
AVPWGQGVEIERALKQVQVEGLSGNIKFDQNGKRINYTINIMELKTNGPRKIGYWSEVDKMV
VTLTELPSGNDTSGLE
00000000000000000000000000000000000000000000000000000000000000
00000000000000000000000000000000000000000000000000000000000000
00000000000000000000000000000000000000000000000000000000000000
00000000000000000000000000000000000000000000000000000000000000
00000000000000000000000000000000000000000000000001111110000000
00000000000000000000000000000000000000000000000000000000000000
0011111111111111
>DM_train459
QKTQSAPGTLSPDARNEKQPFYGEHQAGILTPQQAAMMLVAFDVLASDKADLERLFRLLTQR
FAFLTQGGAAPETPNPRLPPLDSGILGGYIAPDNLTITLSVGHSLFDERFGLAPQMPKKLQK
MTRFPNDSLDAALCHGDVLLQICANTQDTVIHALRDIIKHTPDLLSVRWKREGFISDHAARS
KGKETPINLLGFKDGTANPDSQNDKLMQKVVWVTADQQEPAWTIGGSYQAVRLIQFRVEFWD
RTPLKEQQTIFGRDKQTGAPLGMQHEHDVPDYASDPEGKVIALDSHIRLANPRTAESESSLM
LRRGYSYSLGVTNSGQLDMGLLFVCYQHDLEKGFLTVQKRLNGEALEEYVKPIGGGYFFALP
GVKDANDYFGSALLRV
11111111111100000000000000000000000000000000000000000000000000
00000000000000000000000000000000000000000000000000000000000000
00000000000000000000000000000000000000000000000000000000000000
00000000000000000000000000000000000000000000000000000000000000
00000000000000000000000000000000000000000000000000011111111000
00000000000000000000000000000000000000000000000000000000000000
0000000000000000
>DM_train460
MEQVVIVDAIRTPMGRSKGGAFRNVRAEDLSAHLMRSLLARNPSLTAATLDDIYWGCVQQTL
EQGFNIARNAALLAEIPHSVPAVTVNRLCGSSMQALHDAARMIMTGDAQVCLVGGVEHMGHV

PMSHGVDFHPGLSRNVAKAAGMMGLTAEMLSRLHGISREMQDQFAARSHARAWAATQSGAFK
TEIIPTGGHDADGVLKQFNYDEVIRPETTVEALSTLRPAFDPVSGTVTAGTSSALSDGAAAM
LVMSESRARELGLKPRARIRSMAVVGCDPSIMGYGPVPASKLALKKAGLSASDIDVFEMNEA
FAAQILPCIKDLGLMEQIDEKINLNGGAIALGHPLGCSGARISTTLINLMERKDAQFGLATM
CIGLGQGIATVFERV
000000000000000000000000000000000000000000000000000000000000
000000000000000000000000000000000000000000000000000000000000
000000000000011111110000000000000000000000000000000000000000
000000000000000000000000000000000000000000000000000000000000
000000000000000000000000000000000000000000000000000000000000
000000000000000000000000000000000000000000000000000000000000
000000000000000
>DM_train461
MSKTIIKNIGKIVSGDIKSPVLQADTIVVEDGLIAAIGGEELMKDAGDATIIDAAGSTVTPG
LLDTHVHVSGGDYAPRQKTMDFISSALHGGVTTMISAGSPHFPGRPKDAAGTKALAITLSKS
YYNARPAGVKVHGGAVILEKGLTEEDFIEMKKEGVWIVGEVGLGTIKNPEDAAPMVEWAHKH
GFKVQMHTGGTSIPGSSTVTADDVIKTKPDVVSHINGGPTAISVQEVDRIMDETDFAMEIVQ
CGNPKIADYVARRAAEKGQLGRVIFGNDAPSGTGLIPLGILRNMCQIASMSDIDPEVAVCMA
TGNSTAVYGLNTGVIAPGKEADLIIMDTPLGSVAEDAMGAIAAGDIPGISVVLIDGEAVVTK
SRNTPPAKRAAKIL
100000000000000000000000000000000000000000000000000000000000
000000000000000000000000000000000000000000000000000000000000
000000000000000000000000000000000000000000000000000000000000
000000000000000000000000000000000000000000000000000000000000
000000000000000000000000000000000000000000000000000000000000
000000000000000000000000000000000000000000000000000000000000
00000000000000
>DM_train462
MKKQNDIPQPIRGDKGATVKIPRNIERDRQNPDMLVPPETDHGTVSNMKFSFSDTHNRLEKG
GYAREVTVRELPISENLASVNMRLKPGAIRELHWHKEAEWAYMIYGSARVTIVDEKGRSFID
DVGEGDLWYFPSGLPHSIQALEEGAEFLLVFDDGSFSENSTFQLTDWLAHTPKEVIAANFGV
TKEEISNLPGKEKYIFENQLPGSLKDDIVEGPNGEVPYPFTYRLLEQEPIESEGGKVYIADS
TNFKVSKTIASALVTVEPGAMRELHWHPNTHEWQYYISGKARMTVFASDGHARTFNYQAGDV
GYVPFAMGHYVENIGDEPLVFLEIFKDDHYADVSLNQWLAMLPETFVQAHLDLGKDFTDVLS
KEKHPVVKKKCSK
111111100000000000000000000000000000000000000000000000000000
000000000000000000000000000000000000000000000000000000000000
000000000000000000000000000000000000000000000000000000000000
000000000000000000000000000000000000000000000000000000000000
000000000000000000000000000000000000000000000000000000000000
000000000000000000000000000000000000000000000000000000000000
0000000111111
>DM_train466
MTPISQTPGLLAEAMVDLGAIEHNVRVLREHAGHAQLMAVVKADGYGHGATRVAQTALGAGA

AELGVATVDEALALRADGITAPVLAWLHPPGIDFGPALLADVQVAVSSLRQLDELLHAVRRT
GRTATVTVKVDTGLNRNGVGPAQFPAMLTALRQAMAEDAVRLRGLMSHMVYADKPDDSINDV
QAQRFTAFLAQAREQGVRFEVAHLSNSSATMARPDLTFDLVRPGIAVYGLSPVPALGDMGLV
PAMTVKCAVALVKSIRAGEGVSYGHTWIAPRDTNLALLPIGYADGVFRSLGGRLEVLINGRR
CPGVGRICMDQFMVDLGPGPLDVAEGDEAILFGPGIRGEPTAQDWADLVGTIHYEVVTSPRG
RITRTYREAENR
111111111100000000000000000000000000000000000000000000000000
000000000000000000000000000000000000000000000000000000000000
000000000000000000000000000000000000000000000000000000000000
000000000000000000000000000000000000000000000000000000000000
000000000000000111111111111111000000000000000000000000000000
000000000000000000000000000000000000000000000000000000000000
000000000111
>DM_train467
MKSKKILIVGAGFSGAVIGRQLAEKGHQVHIIDQRDHIGGNSYDARDSETNVMVHVYGPHIF
HTDNETVWNYVNKHAEMMPYVNRVKATVNGQVFSLPINLHTINQFFSKTCSPDEARALIAEK
GDSTIADPQTFEEEALRFIGKELYEAFFKGYTIKQWGMQPSELPASILKRLPVRFNYDDNYF
NHKFQGMPKCGYTQMIKSILNHENIKVDLQREFIVEERTHYDHVFYSGPLDAFYGYQYGRLG
YRTLDFKKFTYQGDYQGCAVMNYCSVDVPYTRITEHKYFSPWEQHDGSVCYKEYSRACEEND
IPYYPIRQMGEMALLEKYLSLAENETNITFVGRLGTYRYLDMDVTIAEALKTAEVYLNSLTE
NQPMPVFTVSVR
100000000000000000000000000000000000000000000000000000000000
000000000000000000000000000000000000000000000000000000000000
000000000000000000000000000000000000000000000000000000000000
000000000000000000000000000000000000000000000000000000000000
000000000000000000000000000000000000000000000000000000000000
000000000000000000000000000000000000000000000000000000000000
000000000000
>DM_train468
MELQEVLHMNGGEGDTSYAKNSSYNLFLIRVKPVLEQCIQELLRANLPNINKCFKVGDLGCA
SGPNTFSTVRDIVQSIDKVGQEKKNELERPTIQIFLNDLFQNDFNSVFKLLPSFYRNLEKEN
GRKIGSCLIGAMPGSFYSRLFPEESMHFLHSCYCLHWLSQVPSGLVTELGISVNKGCIYSSK
ASRPPIQKAYLDQFTKDFTTFLRIHSEELISRGRMLLTFICKEDEFDHPNSMDLLEMSINDL
VIEGHLEEEKLDSFNVPIYAPSTEEVKRIVEEEGSFEILYLETFNAPYDAGFSIDDDYQGRS
HSPVSCDEHARAAHVASVVRSIYEPILASHFGEAILPDLSHRIAKNAAKVLRSGKGFYDSVI
ISLAKKPEKADM
110000000001111000000000000000000000000000000000000000000000
000000000000000000011111111110000000000000000000000000000000
000000000000000000000000000000000000000000111111000000000000
000000000000000000000000000000000000000000000000000000000000
000000000000000000000000000000000000000000000000000011111111
100000000000000000000000000000000000000000000000000000000000
000000011111
>DM_train469

MRILFVAAGSPATVFALAPLATAARNAGHQVVMAANQDMGPVVTGVGLPAVATTDLPIRHFI
TTDREGRPEAIPSDPVAQARFTGRWFARMAASSLPRMLDFSRAWRPDLIVGGTMSYVAPLLA
LHLGVPHARQTWDAVDADGIHPGADAELRPELSELGLERLPAPDLFIDICPPSLRPANAAPA
RMMRHVATSRQCPLEPWMYTRDTRQRVLVTSGSRVAKESYDRNFDFLRGLAKDLVRWDVELI
VAAPDTVAEALRAEVPQARVGWTPLDVVAPTCDLLVHHAGGVSTLTGLSAGVPQLLIPKGSV
LEAPARRVADYGAAIALLPGEDSTEAIADSCQELQAKDTYARRAQDLSREISGMPLPATVVT
ALEQLAHHHHHH
000000000000000000000000000000000000000000000000000000000000
000000000000000000000000000000000000000000000000000000000000
000000000000000000000000000000000000000000000000000000000000
000000000000000000000000000000001111111000000000000000000000
000000000000000000000000000000000000000000000000000000000000
000000000000000000000000000000000000000000000000000000000000
000000000000
>DM_train471
NDVGAESKQPLLDIALKGLKRTLPQLEQMDGNSLRENFQEMASGNGPLRSLMTNLQNLNKIP
EAKQLNDYVTTLTNIQVGVARFSQWGTCGGEVERWVDKASTHELTQAVKKIHVIAKELKNVT
AELEKIEAGAPMPQTMSGPTLGLARFAVSSIPINQQTQVKLSDGMPVPVNTLTFDGKPVALA
GSYPKNTPDALEAHMKMLLEKECSCLVVLTSEDQMQAKQLPPYFRGSYTFGEVHTNSQKVSS
ASQGEAIDQYNMQLSCGEKRYTIPVLHVKNWPDHQPLPSTDQLEYLADRVKNSNQNGAPGRS
SSDKHLPMIHCLGGVGRTGTMAAALVLKDNPHSNLEQVRADFRDSRNNRMLEDASQFVQLKA
MQAQLLMTTAS
111111111100000000000000000000000001111111111000000000000111
110000000000000000000000011110000000000000000000000000000000
000000111110000000000000000000000000000000000000000000000000
000000000000000000000000000000000000000000000000000000000000
000000000000000000000000000000000000000000000000000000000000
000000000000000000000000000000000000000000000000000000000000
00000001111
>DM_train472
AAASFGQTKIPRGNGPYSVGCTDLMFDHTNKGTFLRLYYPSQDNDRLDTLWIPNKEYFWGLS
KFLGTHWLMGNILRLLFGSMTTPANWNSPLRPGEKYPLVVFSHGLGAFRTLYSAIGIDLASH
GFIVAAVEHRDRSASATYYFKDQSAAEIGDKSWLYLRTLKQEEETHIRNEQVRQRAKECSQA
LSLILDIDHGKPVKNALDLKFDMEQLKDSIDREKIAVIGHSFGGATVIQTLSEDQRFRCGIA
LDAWMFPLGDEVYSRIPQPLFFINSEYFQYPANIIKMKKCYSPDKERKMITIRGSVHQNFAD
FTFATGKIIGHMLKLKGDIDSNVAIDLSNKASLAFLQKHLGLHKDFDQWDCLIEGDDENLIP
GTNINTTNQHI
111111100000000000000000000000000000000000000000000000000000
000000000000000000000000000000000000000000000000000000000000
000000000000000000000000000000000000000000000000000000000000
000000000000000000000000000000000000000000000000000000000000
000000000000000000000000000000000000000000000000000000000000
000000000000000000000000000000000000000000000000000000000000
00000001111

>DM_train473
PQPPADEQPEPRTRRRAYLWCKEFLPGAWRGLREDEFHISVIRGGLSNMLFQCSLPDTTATL
GDEPRKVLLRLYGAILQMRSCNKEGSEQAQKENEFQGAEAMVLESVMFAILAERSLGPKLYG
IFPQGRLEQFIPSRRLDTEELSLPDISAEIAEKMATFHGMKMPFNKEPKWLFGTMEKYLKEV
LRIKFTEESRIKKLHKLLSYNLPLELENLRSLLESTPSPVVFCHNDCQEGNILLLEGRENSE
KQKLMLIDFEYSSYNYRGFDIGNHFCEWMYDYSYEKYPFFRANIRKYPTKKQQLHFISSYLP
AFQNDFENLSTEEKSIIKEEMLLEVNRFALASHFLWGLWSIVQAKISSIEFGYMDYAQARFD
AYFHQKRKLGV
11111111111110000000000000100111111111111111111100000111111111
11111100000111111111111111111111111111110000000000000000000000
00000000000000000000000000000000000000000000000000000000000000
00000000000000000000000000000000000000000000000000000000000000
00000000000000000000000000000000000000000000000000000000000000
00000000000000000000000000000000000000000000000000000000000000
00000000000
>DM_train476
MKALHFGAGNIGRGFIGKLLADAGIQLTFADVNQVVLDALNARHSYQVHVVGETEQVDTVSG
VNAVSSIGDDVVDLIAQVDLVTTAVGPVVLERIAPAIAKGLVKRKEQGNESPLNIIACENMV
RGTTQLKGHVMNALPEDAKAWVEEHVGFVDSAVDRIVPPSASATNDPLEVTVETFSEWIVDK
TQFKGALPNIPGMELTDNLMAFVERKLFTLNTGHAITAYLGKLAGHQTIRDAILDEKIRAVV
KGAMEESGAVLIKRYGFDADKHAAYIQKILGRFENPYLKDDVERVGRQPLRKLSAGDRLIKP
LLGTLEYSLPHKNLIQGIAGAMHFRSEDDPQAQELAALIADKGPQAALAQISGLDANSEVVS
EAVTAYKAMQ
00000000000000000000000000000000000000000000000000000000000000
00000000000000000000000000000000000000000000000000000000000000
00000000000000000000000000000000000000011111000000000000000000
00000000000000000000000000000000000000000000000000000000000000
00000000000000000000000000000000000000000000000000000000000000
00000000000000000000000000000000000000000000000000000000000000
0000000000
>DM_train477
EESRESPAEHGYYMPAEWDSHAQTWIGWPERQDNWRHNALPAQRVFAGVAKAISKFEPVTVC
ASPAQWENARKQLPEDIRVVEMSMNDSWFRDSGPTFIVRKRPVKLSSLNRNIAGIDWNFNAW
GGANDGCYNDWSHDLLVSRKILALERIPRFQHSMILEGGSIHVDGEGTCLVTEECLLNKNRN
PHMSKEQIEEELKKYLGVQSFIWLPRGLYGDEDTNGHIDNMCCFARPGVVLLSWTDDETDPQ
YERSVEALSVLSNSIDARGRKIQVIKLYIPEPLYMTEEESSGITQDGEAIPRLAGTRLAASY
VNFYIANGGIIAPQFGDPIRDKEAIRVLSDTFPHHSVVGIENAREIVLAGGNIHCITQQQPA
EPTSVAENGH
00000000000000000000000000000000000000000000000000000000000000
00000000000000000000000000000000000000000111111100000000000000
00000000000000000000000000000000000000000000000000000000000000
00000000000000000000000000000000000000000000000000000000000000
00000000000000000000000000000000000000000000000000000000000000
00000000000000000000000000000000000000000000000000000000000000

0001111111
>DM_train478
MKSFQLVATAASGLEAIVGKEVARLGYDPKVENGKVYFEGDLSAIARANLWLRVADRVKIVV
GVFKATTFDELFEKTKALPWEDYLPLDAQFPVAGKSVKSTLYSVPDCQAIVKKAIVNRVSEK
YRRSGRLMETGALFKLEVSILKDEVTLTIDTSGAGLHKRGYRLAQGSAPIKETMAAALVLLT
SWHPDRPFYDPVCGSGTIPIEAALIGQNIAPGFNREFVSETWDWMPKQVWADARQEAEDLAN
YDQPLNIIGGDIDARLIEIAKQNAVEAGLGDLITFRQLQVADFQTEDEYGVVVANPPYGERL
EDEEAVRQLYREMGIVYKRMPTWSVYVLTSYELFEEVYGKKATKKRKLYNGYLRTDLYQYWG
PRKPRPKKED
00000000000000000000000000000000000000000000000000000000000000
00000000000000000000000000000000000000000000000000000000000000
00000000000000000000000000000000000000000000000000000000000000
00000000000000000000000000000000000000000000000000000000000000
00000000000000000000000000000000000000000000000000000000000000
00000000000000000000000000000000000000000000000001111000000000
0111111111
>DM_train480
MQRLDALLPTLRERAQETEDLRRIPDDSMKALQETGFFRLLQPEQWGGYQADPVLFYSAVRK
IASACGSTGWVSSIIGVHNWHLALFSQQAQEDVWGNDTDVRISSSYAPMGAGQVVDGGYTVN
GAWAWSSGCDHASWAVLGGPVIKDGRPVDFVSFLIPREDYRIDDVWNVVGLRGTGSNTVVVE
DVFVPTHRVLSFKAMSNLTAPGLERNTAPVYKMPWGTIHPTTISAPIVGMAYGAYDAHVEHQ
GKRVRAAFAGEKAKDDPFAKVRIAEASSDIDAAWRQLSGNVADEYALLVAGEEVPFELRLRA
RRDQVRATGRAISSIDKLFESSGATALANGTPLQRFWRDAHAGRVHAANDPERAYVMYGTGE
FGLPITDTMV
00000000000000000000000000000000000000000000000000000000000000
00000000000000000000000000000000000000000000000000000000000000
00000000000000000000000000000000000000000000000000000000000000
00000000000000000000000000000000000000000000000000000000000000
00001111111111000000000000000000000000000000000000000000000000
00000000000000000000000000000000000000000000000000000000000000
0000000000
>DM_train482
MDINGGGATLPQALYQTSGVLTAGFAQYIGVGSGNGKAAFLNNDYTKFQAGVTNKNVHWAGS
DSKLSATELSTYASAKQPTWGKLIQVPSVGTSVAIPFNKSGSAAVNLSVQELCGVFSGRINT
WDGISGSGRTGPIVVVYRSESSGTTELFTRFLNAKCNAETGNFAVTTTFGTSFSGGLPAGAV
AATGSQGVMTALAAGDGRITYMSPDFAAPTLAGLDDATKVARVGKNVATNTQGVSPAAANVS
AAIGAVPVPAAADRSNPDAWVPVFGPDNTAGVQPYPTSGYPILGFTNLIFSQCYADATQTTQ
VRDFFTKHYGASNNNDAAITANAFVPLPTAWKATVRASFLTASNALSIGNTNVCNGIGRPLL
EAAHHHHHH
00000000000000000000000000000000000000000000000000000000000000
00000000000000000000000000000000000000000000000000000000000000
00000000000000000000000000000000000000000000000000000000000000
00000000000000000000000000000000000000000000000000000000000000
00000000000000000000000000000000000000000000000000000000000000

000000000000000000000000000000000000000000000000000000000000
111111111
>DM_train483
HHHHHHTDPALRAAQQTPLYEQHTLCGARMVDFHGWMMPLHYGSQIDEHHAVRTDAGMFDVS
HMTIVDLRGSRTREFLRYLLANDVAKLTKSGKALYSGMLNASGGVIDDLIVYYFTEDFFRLV
VNSATREKDLSWITQHAEPFGIEITVRDDLSMIAVQGPNAQAKAATLFNDAQRQAVEGMKPF
FGVQAGDLFIATTGYTGEAGYEIALPNEKAADFWRALVEAGVKPCGLGARDTLRLEAGMNLY
GQEMDETISPLAANMGWTIAWEPADRDFIGREALEVQREHGTEKLVGLVMTEKGVLRNELPV
RFTDAQGNQHEGIITSGTFSPTLGYSIALARVPEGIGETAIVQIRNREMPVKVTKPVFVRNG
KAVAGLCGR
111111111111111000000000000000000000000000000000000000000000
000000000000000000000000000000000000000000000000000000000000
000000000000000000000000000000000000000000000000000000000000
000000000000000000000000000000000000000000000000000000000000
000000000000000000000000000000000000000000000000000000000000
000000000000000000000000000000000000000000000000000000000000
000000011
>DM_train484
MSTAVQFRGGTTAQHATFTGAAREITVDTDKNTVVVHDGATAGGFPLARHDLVKTAFIKADK
SAVAFTRTGNATASIKAGTIVEVNGKLVQFTADTAITMPALTAGTDYAIYVCDDGTVRADSN
FSAPTGYTSTTARKVGGFHYAPGSNAAAQAGGNTTAQINEYSLWDIKFRPAALDPRGMTLVA
GAFWADIYLLGVNHLTDGTSKYNVTIADGSASPKKSTKFGGDGSAAYSDGAWYNFAEVMTHH
GKRLPNYNEFQALAFGTTEATSSGGTDVPTTGVNGTGATSAWNIFTSKWGVVQASGCLWTWG
NEFGGVNGASEYTANTGGRGSVYAQPAAALFGGAWNGTSLSGSRAALWYSGPSFSFAFFGAR
GVCDHLILE
111100000000000000000000000000000000000000000000000000000000
000000000000000000000000000000000000000000000000000000000000
000000000000000000000000000000000000000000000000000000000000
000000000000000000000000000000000000000000000000000000000000
000000000000000000000000000000000000000000000000000000000000
000000000000000000000000000000000000000000000000000000000000
000000001
>DM_train488
MIQFSINRTLFIHALNTTKRAISTKNAIPILSSIKIEVTSTGVTLTGSNGQISIENTIPVSN
ENAGLLITSPGAILLEASFFINIISSLPDISINVKEIEQHQVVLTSGKSEITLKGKDVDQYP
RLQEVSTENPLILKTKLLKSIIAETAFAASLQESRPILTGVHIVLSNHKDFKAVATDSHRMS
QRLITLDNTSADFMVVLPSKSLREFSAVFTDDIETVEVFFSPSQILFRSEHISFYTRLLEGN
YPDTDRLLMTEFETEVVFNTQSLRHAMERAFLISNATQNGTVKLEITQNHISAHVNSPEVGK
VNEDLDIVSQSGSDLTISFNPTYLIESLKAIKSETVKIHFLSPVRPFTLTPGDEEESFIQLI
TPVRTN
100000000000000000000000000000000000000000000000000000000011
111000000000000000000000000000000000000000000000000000000000
000000000000000000000000000000000000000000000000000000000000
000000000000000000000000000000000000000000000000000000000000

000000000000000000000000000000000000000000000000000000000000
000000000000000000000000000000000000000000000000000000000000
000001
>DM_train489
MELIEVQEGKAKILIPKAESIYDSPVFYNPRMALNRDIVVVLLNILNPKIVLDALSATGIRG
IRFALETPAEEVWLNDISEDAYELMKRNVMLNFDGELRESKGRAILKGEKTIVINHDDANRL
MAERHRYFHFIDLDPFGSPMEFLDTALRSAKRRGILGVTATDGAPLCGAHPRACLRKYLAVP
LRGELCHEVGTRILVGVIARYAAKYDLGIDVILAYYKDHYFRAFVKLKDGARKGDETLEKLG
YIYFDDKTGKFELEQGFLPTRPNAYGPVWLGPLKDEKIVSKMVKEAESLSLARKKQALKLLK
MIDQELDIPLFYDTHAIGRRLKIETKKVEEIISALREQGYEATRTHFSPTGIKTSAPYEVFI
ETIKRI
110000000000000011111111000000000000000000000000000000000000
000000000000000000000000000000000000000000000000000000000000
000000000000000000000000000000000000000000000000000000000000
000000000000000000000000000000000000000000000000000000000000
000000000000000000000000000000000000000000000000000000000000
000000000000000000000000000000000000000000000000000000000000
000001
>DM_train490
MTKKRVALIFGGNSSEHDVSKRSAQNFYNAIEATGKYEIIVFAIAQNGFFLDTESSKKILAL
EDEQPIVDAFMKTVDASDPLARIHALKSAGDFDIFFPVVHGNLGEDGTLQGLFKLLDKPYVG
APLRGHAVSFDKALTKELLTVNGIRNTKYIVVDPESANNWSWDKIVAELGNIVFVKAANQGS
SVGISRVTNAEEYTEALSDSFQYDYKVLIEEAVNGARELEVGVIGNDQPLVSEIGAHTVPNQ
GSGDGWYDYNNKFVDNSAVHFQIPAQLSPEVTKEVKQMALDAYKVLNLRGEARMDFLLDENN
VPYLGEPNTLPGFTNMSLFKRLWDYSDINNAKLVDMLIDYGFEDFAQNKKLSYSFVSLGEEK
IGKFN
000000000000000000000000000000000000000000000000000000000000
000000000000000000000000000000000000000000000000000000000000
000000000000000000000000000000000000000000000000000000000000
000000000000000000000000000000000000000000000000000000001111
111111111111111111000000000000000000000000000000000000000000
000000000000000000000000000000000000000000000000000000000011
11111
>DM_train491
GLKEHDFNPEEAEETKQVSWKLVTEYAMETKCDDVLLLLGMYLEFQYSFEMCLKCIKKEQPS
HYKYHEKHYANAAIFADSKNQKTICQQAVDTVLAKKRVDSLQLTREQMLTNRFNDLLDRMDI
MFGSTGSADIEEWMAGVAWLHCLLPKMDSVVYDFLKCMVYNIPKKRYWLFKGPIDSGKTTLA
AALLELCGGKALNVNLPLDRLNFELGVAIDQFLVVFEDVKGTGGESRDLPSGQGINNLDNLR
DYLDGSVKVNLEKKHLNKRTQIFPPGIVTMNEYSVPKTLQARFVKQIDFRPKDYLKHCLERS
EFLLEKRIIQSGIALLLMLIWYRPVAEFAQSIQSRIVEWKERLDKEFSLSVYQKMKFNVAMG
IGVLD
111111111111111000000000000000000000000000000000000000000000
000000000000000000000000000000000000000000000000000000000000
000000000000000000000000000000000000000000000000000000000000

000000000000000000000000000000000000000000000000000000000000
000000000000000000000000000000000000000000000000000000000000
000000000000000000000000000000000000000000000000000000000000
00000
>DM_train492
GMKSAPDWIAHPDTPYLLIDEAKLKSNINYLKQRVESLGSHLRPHLKTLRTLEAAGYLLDSK
SAPATVSTLAEAEAYAKAGYTDLLYAVGIAPAKLKRVAALRQQGINLHILLDNITQAQAVVD
YAAEFGQDFSVFIEIDSDDHRGGIKPSDSKLLTIAKTLGEHFTGLMTHAGGSYACNTEQGLK
NFAKQECDAVRIARNNLETAGIHCAITSVGSTPTAHFGEDFSDISEVRAGVYTTFDLVMKNI
GVCDFSHIAMSVVTTVIGHNKEKNWLLTDSGWMALSRDSGTAGQNRDFGYGQVCKIDGSVLD
GLCVNSTSQEHGVIELSDAYQLEDFPVGHQLRIMPNHACATAAMHPVYHVLMSDGSHNTWQR
ITGW
111000000000000000000000000000000000000000000000000000000000
000000000000000000000000000000000000000000000000000000000000
000000000000000000000000000000000000000000000000000000000000
000000000000000000000000000000000000000000000000000000000000
000000000000000000000000000000000000000000000000000000000000
000000000000000000000000000000000000000000000000000000000000
0000
>DM_train493
MREWKIIDSTLREGEQFEKANFSTQDKVEIAKALDEFGIEYIEVTTPVASPQSRKDAEVLAS
LGLKAKVVTHIQCRLDAAKVAVETGVQGIDLLFGTSKYLRAPHGRDIPRIIEEAKEVIAYIR
EAAPHVEVRFSAEDTFRSEEQDLLAVYEAVAPYVDRVGLADTVGVATPRQVYALVREVRRVV
GPRVDIEFHGHNDTGCAIANAYEAIEAGATHVDTTILGIGERNGITPLGGFLARMYTLQPEY
VRRKYKLEMLPELDRMVARMVGVEIPFNNYITGETAFSHKAGMHLKAIYINPEAYEPYPPEV
FGVKRKLIIASRLTGRHAIKARAEELGLHYGEEELHRVTQHIKALADRGQLTLEELDRILRE
WITA
100000000000000000000000000000000000000000000000000000000000
000000000000000000000000000000000000001111110000000000000000
000000000000000000000000000000000000000000000000000000000000
000000000000000000000000000000000000000000000000000000000000
000000000000000000000000000000000000000000000000000000000000
000011111111111111111111100000000000000000000000000000000000
0000
>DM_train494
MTSQTPTRKPRARDLGLPFTGVTGPYNAITDVDGVGVGFQTIIENEPRPGRKRPARSGVTAI
LPHMQSETPVPVYAGVHRFNGNGEMTGTHWIEDGGYFLGPVVITNTHGIGMAHHATVRWMVD
RYASTYQTDDFLWIMPVVAETYDGALNDINGFPVTEADVRKALDNVASGPVQEGNCGGGTGM
ITYGFKGGTGTASRVVEFGGRSFTIGALVQANHGQRDWLTIAGVPVGQHMRDGTPQSQLQER
GSIIVVLATDLPLMPHQLKRLARRASIGIGRNGTPGGNNSGDIFIAFSTANQRPMQHRSAPF
LDVEMVNDEPLDTVYLAAVDSVEEAVVNAMIAAEDMGGTPFDRLLVQAIDHERLRAVLRQYG
RLA
111111110000000000000000000000000000000000000000000000000000
000000000000000000000000000000000000000000000000000000000000

000000000000000000000000000000000000000000000000000000000000
000000000000000000000000000000000000000000000000000000000111
100000000000000000000000000000000000000000000000000000000000
000000000000000000000000000000000000000000000000000000000000
000
>DM_train495
MSLTREAYHRLTPLPHPGGRLFIKPGARGYRDPVHDLLQKTVEPFGERALDLNPGVGWGSLP
LEGRMAVERLETSRAAFRCLTASGLQARLALPWEAAAGAYDLVVLALPAGRGTAYVQASLVA
AARALRMGGRLYLAGDKNKGFERYFKEARALLGYGVVVRREGPYRVALLEKEKEAPPLPSLW
RAFSARILGAEYTFHHLPGVFSAGKVDPASLLLLEALQERLGPEGVRGRQVLDLGAGYGALT
LPLARMGAEVVGVEDDLASVLSLQKGLEANALKAQALHSDVDEALTEEARFDIIVTNPPFHV
GGAVILDVAQAFVNVAAARLRPGGVFFLVSNPFLKYEPLLEEKFGAFQTLKVAEYKVLFAEK
RGR
110000000000000000000000000000000000000000000000000000000000
000000000000000000000000000000000000000000000000000000000000
000000000000000000000000000000000000000000000000000000000000
000000000000000000000000000000000000000000000000000000000000
000000000000000000000000000000000000000000000000000000000000
000000000000000000000000000000000000000000000000000000000000
011
>DM_train497
MENFPTEYFLNTTVRLLEYIRYRDSNYTREERIENLHYAYNKAAHHFAQPRQQQLLKVDPKR
LQASLQTIVGMVVYSWAKVSKECMADLSIHYTYTLVLDDSKDDPYPTMVNYFDDLQAGREQA
HPWWALVNEHFPNVLRHFGPFCSLNLIRSTLDFFEGCWIEQYNFGGFPGSHDYPQFLRRMNG
LGHCVGASLWPKEQFNERSLFLEITSAIAQMENWMVWVNDLMSFYKEFDDERDQISLVKNYV
VSDEISLHEALEKLTQDTLHSSKQMVAVFSDKDPQVMDTIECFMHGYVTWHLCDRRYRLSEI
YEKVKEEKTEDAQKFCKFYEQAANVGAVSPSEWAYPPVAQLANVRSKDVKEVQKPFLSSIEL
VE
111100000000000000000000000000000000000000000000000000000000
000000000000000000000000000000000000000000000000000000000000
000000000000000000000000000000000000000000000000000000000000
000000000000000000000000000000000000000000000000000000000000
000000000000000000000000000000000000000000000000000000000000
000000000000000000000000000000000000000000011111111111111111
11
>DM_train498
MTWKNFGFEIFGEKYGQEELEKRIKDEHTPPPDSPVFGGLKLKLKKEKFKTLFTLGTTLKGF
RRATHTVGTGGIGEITIVNDPKFPEHEFFTAGRTFPARLRHANLKYPDDAGADARSFSIKFA
DSDSDGPLDIVMNTGEANIFWNSPSLEDFVPVEEGDAAEEYVYKNPYYYYNLVEALRRAPDT
FAHLYYYSQVTMPFKAKDGKVRYCRYRALPGDVDIKEEDESGRLTEEEQRKIWIFSRHENEK
RPDDYLRKEYVERLQKGPVNYRLQIQIHEASPDDTATIFHAGILWDKETHPWFDLAKVSIKT
PLSPDVLEKTAFNIANQPASLGLLEAKSPEDYNSIGELRVAVYTWVQHLRKLKIGSLVPAGQ
NA
110000000000000000000000000000000000000000000000000000000000

000000000000000000000000000000000000000000000000000000000000
000000000000000000000000000000000000000000000000000000000000
000000000000000000000000000000000000000000000000000000000000
000000000000000000000000000000000000000000000000000000000000
000000000000000000000000000000000000000000000000000000001111
11
>DM_train499
MGSDKIHHHHHHMTSSPTSTQESSTSWYLLLQQLIDGESLSRSQAAELMQGWLSEAVPPELS
GAILTALNFKGVSADELTGMAEVLQSQSKMGTGENYSQLPITNSPFSIIDTCGTGGDGSSTF
NISTAVAFVAAAYGVPVAKHGNRSASSLTGSADVLEALGVNLGASPEKVQAALQEVGITFLF
APGWHPALKAVATLRRTLRIRTVFNLLGPLVNPLRPTGQVVGLFTPKLLTTVAQALDNLGKQ
KAIVLHGRERLDEAGLGDLTDLAVLSDGELQLTTINPQEVGVTPAPIGALRGGDVQENAEIL
KAVLQGKGTQAQQDAVALNAALALQVAGAVPLLDHAQGVSVAKEILQTGTAWAKLAQLVYFL
GN
111111111111111111111111000000000000000000000000000000000000
000000000000000000000000000001111111111100000000000001110000
000000000000000000000000010000000000000000000000000000000000
001110000000000000000000000000000000000000000000000000000000
000000000000000000000000000000000000000000000000000000000000
000000000000000000000000000000000000000000000000000000000000
00
>DM_train500
MIVAFCLYKYFPFGGLQRDFMRIASTVAARGHHVRVYTQSWEGDCPKAFELIQVPVKSHTNH
GRNAEYYAWVQNHLKEHPADRVVGFNKMPGLDVYFAADVCYAEKVAQEKGFLYRLTSRYRHY
AAFERATFEQGKSTKLMMLTDKQIADFQKHYQTEPERFQILPPGIYPDRKYSEQIPNSREIY
RQKNGIKEQQNLLLQVGSDFGRKGVDRSIEALASLPESLRHNTLLFVVGQDKPRKFEALAEK
LGVRSNVHFFSGRNDVSELMAAADLLLHPAYQEAAGIVLLEAITAGLPVLTTAVCGYAHYIA
DANCGTVIAEPFSQEQLNEVLRKALTQSPLRMAWAENARHYADTQDLYSLPEKAADIITGGL
DG
100000000000000000000000000000000000000000000000000000000000
000000000000000000000000000000000000000000000000000000000000
000000000000000000000000000000000000000000000000000000000000
000000000000000000000000000000000000000000000000000000000000
000000000000000000000000000000000000000000000000000000000000
000000000000000000000000000000000000000000000000000000000001
11
>DM_train501
GHEIKDIREDTMHAEFNALRAQVAINDGNPDEAERLAKLALEELPPGWFYSRIVATSVLGEV
LHCKGELTRSLALMQQTEQMARQHDVWHYALWSLIQQSEILFAQGFLQTAWETQEKAFQLIN
EQHLEQLPMHEFLVRIRAQLLWAWARLDEAEASARSGIEVLSSYQPQQQLQCLAMLIQCSLA
RGDLDNARSQLNRLENLLGNGKYHSDWISNANKVRVIYWQMTGDKAAAANWLRHTAKPEFAN
NHFLQGQWRNIARAQILLGEFEPAEIVLEELNENARSLRLMSDLNRNLLLLNQLYWQAGRKS
DAQRVLLDALKLANRTGFISHFVIEGEAMAQQLRQLIQLNTLPELEQHRAQRILREINQHHG
A

000000000000000000000000000000000000000000000000000000000000
000000000000000000000000000000000000000000000000000000000000
000000000000000000000000000000000000000000000000000000000000
000000000000000000000000000000000000000000000000000000000000
000000000000000000000000000000000000000000000000000000000000
000000000000000000000000000000000000000000000000000000001111
1
>DM_train502
MGSMDQSIAVKSPLTYAEALANTIMNTYTVEELPPANRWHYHQGVFLCGVLRLWEATGEKRY
FEYAKAYADLLIDDNGNLLFRRDELDAIQAGLILFPLYEQTKDERYVKAAKRLRSLYGTLNR
TSEGGFWHKDGYPYQMWLDGLYMGGPFALKYANLKQETELFDQVVLQESLMRKHTKDAKTGL
FYHAWDEAKKMPWANEETGCSPEFWARSIGWYVMSLADMIEELPKKHPNRHVWKNTLQDMIK
SICRYQDKETGLWYQIVDKGDRSDNWLESSGSCLYMYAIAKGINKGYLDRAYETTLLKAYQG
LIQHKTETSEDGAFLVKDICVGTSAGFYDYYVSRERSTNDLHGAGAFILAMTELEPLFRSAG
K
111111111100000000000000000000000000000000000000000000000000
000000000000000000000000000000000000000000000000000000000000
000000000000000000000000000000000000000000000000000000000000
000000000000000000000000000000000000000000000000000000000000
000000000000000000000000000000000000000000000000000000000000
000000000000000000000000000000000000000000000000000000000000
0
>DM_train503
MAAVTLHLRAETKPLEARAALTPTTVKKLIAKGFKIYVEDSPQSTFNINEYRQAGAIIVPAG
SWKTAPRDRIIIGLKEMPETDTFPLVHEHIQFAHCYKDQAGWQNVLMRFIKGHGTLYDLEFL
ENDQGRRVAAFGFYAGFAGAALGVRDWAFKQTHSDDEDLPAVSPYPNEKALVKDVTKDYKEA
LATGARKPTVLIIGALGRCGSGAIDLLHKVGIPDANILKWDIKETSRGGPFDEIPQADIFIN
CIYLSKPIAPFTNMEKLNNPNRRLRTVVDVSADTTNPHNPIPIYTVATVFNKPTVLVPTTAG
PKLSVISIDHLPSLLPREASEFFSHDLLPSLELLPQRKTAPVWVRAKKLFDRHCARVKRSSR
L
100000000000000000000000000000000000000000000000000000000000
000000000000000000000000000000000000000000000000000000000000
000000000000000000000000000000000000000000000000000000000000
000000000000000000000000000000000000000000000000000000000000
000000000000000000000000000000000000000000000000000000000000
000000000000000000000000000000000000000000000000000000000000
0
>DM_train504
MGNSYITKEDNQISATSEQTEDSACLSAMVLTTNLVYPAVLNAAIDLNLFEIIAKATPPGAF
MSPSEIASKLPASTQHSDLPNRLDRMLRLLASYSVLTSTTRTIEDGGAERVYGLSMVGKYLV
PDESRGYLASFTTFLCYPALLQVWMNFKEAVVDEDIDLFKNVHGVTKYEFMGKDKKMNQIFN
KSMVDVCATEMKRMLEIYTGFEGISTLVDVGGGSGRNLELIISKYPLIKGINFDLPQVIENA
PPLSGIEHVGGDMFASVPQGDAMILKAVCHNWSDEKCIEFLSNCHKALSPNGKVIIVEFILP
EEPNTSEESKLVSTLDNLMFITVGGRERTEKQYEKLSKLSGFSKFQVACRAFNSLGVMEFYK

111111111111111111000000000000000000000000000000000000000000000
00000000000000000000000000000000000000000000000000000000000000
00000000000000000000000000000000000111111111111100000000000000
00000000000000000000000000000000000000000000000000000000000000
00000000000000000000000000000000000000000000000000000000000000
00000000000000000000000000000000000000000000000000000000000000
>DM_train505
DPFAGDPPRHPGLRVNSQKPFNAEPPAELLAERFLTPNELFFTRNHLPVPAVEPSSYRLRVD
GPGGRTLSLSLAELRSRFPKHEVTATLQCAGNRRSEMSRVRPVKGLPWDIGAISTARWGGAR
LRDVLLHAGFPEELQGEWHVCFEGLDADPGGAPYGASIPYGRALSPAADVLLAYEMNGTELP
RDHGFPVRVVVPGVVGARSVKWLRRVAVSPDESPSHWQQNDYKGFSPCVDWDTVDYRTAPAI
QELPVQSAVTQPRPGAAVPPGELTVKGYAWSGGGREVVRVDVSLDGGRTWKVARLMGDKAPP
GRAWAWALWELTVPVEAGTELEIVCKAVDSSYNVQPDSVAPIWNLRGVLSTAWHRVRVSVQD
11111111111110000000000000000000000000000000000000000000000000
00000000000000000000000000000000000000000000000000000000000000
00000000000000000000000000000000000000000000000000000000000000
00000000000000000000000000000000000000000000000000000000000000
00000000000000000000000000000000000000000000000000000000000000
00000000000000000000000000000000000000000000000000000000000000
>DM_train507
SPEVTMNISQMITYWGYPNEEYEVVTEDGYILEVNRIPYGKKNSGNTGQRPVVFLQHGLLAS
ATNWISNLPNNSLAFILADAGYDVWLGNSRGNTWARRNLYYSPDSVEFWAFSFDEMAKYDLP
ATIDFIVKKTGQKQLHYVGHSQGTTIGFIAFSTNPSLAKRIKTFYALAPVATVKYTKSLINK
LRFVPQSLFKFIFGDKIFYPHNFFDQFLATEVCSREMLNLLCSNALFIICGFDSKNFNTSRL
DVYLSHNPAGTSVQNMFHWTQAVKSGKFQAYDWGSPVQNRMHYDQSQPPYYNVTAMNVPIAV
WNGGKDLLADPQDVGLLLPKLPNLIYHKEIPFYNHLDFIWAMDAPQEVYNDIVSMISEDKK
00000000000000000000000000000000000000000000011100000000000000
00000000000000000000000000000000000000000000000000000000000000
00000000000000000000000000000000000000000000000000000000000000
00000000000000000000000000000000000000000000000000000000000000
00000000000000000000000000000000000000000000000000000000000000
0000000000000000000000000000000000000000000000000000000000000
>DM_train509
GSSSSGLVPRGSHMSVQNVVLFDTQPLTLMLGGKLSYINVAYQTYGTLNDEKNNAVLICHAL
TGDAEPYFDDGRDGWWQNFMGAGLALDTDRYFFISSNVLGGCKGTTGPSSINPQTGKPYGSQ
FPNIVVQDIVKVQKALLEHLGISHLKAIIGGSFGGMQANQWAIDYPDFMDNIVNLCSSIYFS
AEAIGFNHVMRQAVINDPNFNGGDYYEGTPPDQGLSIARMLGMLTYRTDLQLAKAFGRATKS
DGSFWGDYFQVESYLSYQGKKFLERFDANSYLHLLRALDMYDPSLGYENVKEALSRIKARYT
LVSVTTDQLFKPIDLYKSKQLLEQSGVDLHFYEFPSDYGHDAFLVDYDQFEKRIRDGLAGN
11111111111111000000000000000000000000000000000000000000000000
00000000000000000000000000000000000000000000000000000000000000
00000000000000000000000000000000000000000000000000000000000000
00000000000000000000000000000000000000000000000000000000000000
00000000000000000000000000000000000000000000000000000000000000

00000000000000000000000000000000000000000000000000000000000
>DM_train510
MAHHHHHHMNILQEPIDFLKKEELKNIDLSQMSKKERYKIWKRIPKCELHCHLDLCFSADFF
VSCIRKYNLQPNLSDEEVLDYYLFAKGGKSLGEFVEKAIKVADIFHDYEVIEDLAKHAVFNK
YKEGVVLMEFRYSPTFVAFKYNLDIELIHQAIVKGIKEVVELLDHKIHVALMCIGDTGHEAA
NIKASADFCLKHKADFVGFDHGGHEVDLKEYKEIFDYVRESGVPLSVHAGEDVTLPNLNTLY
SAIQVLKVERIGHGIRVAESQELIDMVKEKNILLEVCPISNVLLKNAKSMDTHPIRQLYDAG
VKVSVNSDDPGMFLTNINDDYEELYTHLNFTLEDFMKMNEWALEKSFMDSNIKDKIKNLYF
111111111111000000000000000000000000000000000000000000000000
000000000000000000000000000000000000000000000000000000000000
000000000000000000000000000000000000000000000000000000000000
000000000000000000000000000000000000000000000000000000000000
000000000000000000000000000000000000000000000000000000000000
00000000000000000000000000000000000000000000000000000000000
>DM_train511
WDGKIDGTGTHAMIVTQGVSILENDLSKNEPESVRKNLEILKENMHELQLGSTYPDYDKNAY
DLYQDHFWDPDIDNNFSKDNSWYLAYSIPDTGESQIRKFSALARYEWQRGNYKQATFYLGEA
MHYFGDIDTPYHPANVTAVDSAGHVKFETFAEERKEQYKINTAGCKTNEAFYTDILKNKDFN
AWSKEYARGFAKTGKSIYYSHASMSHSWDDWDYAAKVTLANSQKGTAGYIYRFLHDVSEGND
PSVGKNVKELVAYISTSGEKDAGTDDYMYFGIKTKDGKTQEWEMDNPGNDFMTGSKDTYTFK
LKDENLKIDDIQNMWIRKRKYTAFPDAYKPENIKIIANGKVVVDKDINEWISGNSTYNIK
000000000000000000000000000000000000000000000000000000000000
000000000000111111111000000000000000000000000000000000000000
000000000000000000000000000000000000000000000000000000000000
000000000000000000000000000000000000000000000000000000000000
000000000000000000000000000000000000000000000000000000000000
0000000000000000000000000000000000000000000000000000000000
>DM_train512
MSTDIDCIVIGAGVVGLAIARALAAGGHEVLVAEAAEGIGTGTSSRNSEVIHAGIYYPADSL
KARLCVRGKHLLYEYCAARGVPHQRLGKLIVATSDAEASQLDSIARRAGANGVDDLQHIDGA
AARRLEPALHCTAALVSPSTGIVDSHALMLAYQGDAESDGAQLVFHTPLIAGRVRPEGGFEL
DFGGAEPMTLSCRVLINAAGLHAPGLARRIEGIPRDSIPPEYLCKGSYFTLAGRAPFSRLIY
PVPQHAGLGVHLTLDLGGQAKFGPDTEWIATEDYTLDPRRADVFYAAVRSYWPALPDGALAP
GYTGIRPKISGPHEPAADFAIAGPASHGVAGLVNLYGIESPGLTASLAIAEETLARLAA
110000000000000000000000000000000000000000000000000000000000
000000000000000000000000000000000000000000000000000000000000
000000000000000000000000000000000000000000000000000000000000
000000000000000000000000000000000000000000000000000000000000
000000000000000000000000000000000000000000000000000000000000
000000000000000000000000000000000000000000000000000000001
>DM_train513
KERRRAVLELLQRPGNARCADCGAPDPDWASYTLGVFICLSCSGIHRNIPQVSKVKSVRLDA
WEEAQVEFMASHGNDAARARFESKVPSFYYRPTPSDCQLLREQWIRAKYERQEFIYPEKQEP
YSAGYREGFLWKRGRDNGQFLSRKFVLTEREGALKYFNRNDAKEPKAVMKIEHLNATFQPAK

IGHPHGLQVTYLKDNSTRNIFIYHEDGKEIVDWFNALRAARFHYLQVAFPGASDADLVPKLS
RNYLKEGYMEKTGPKQTEGFRKRWFTMDDRRLMYFKDPLDAFARGEVFIGSKESGYTVLHGF
PPSTQGHHWPHGITIVTPDRKFLFACETESDQREWVAAFQKAVDRPMLPQEYAVEAHF
000000000000000000000000000000000000000000000000000000000000
000000000000000000000000000000000000000000000000000000000000
000000000000000000000000000000000000000000000000000000000000
000000000000000000000000000000000000000000000000000000000000
000000000000111111000000000000000000000000000000000000000000
00000000000000000000000000000000000000000000011111111111
>DM_train514
GMRVDKHEVRVGELAAGQPLSLPVYRFKGKGAGPSVYIQANVHGAEVQGNAVIYQLMKLLEH
YELLGDISLVPLANPLGINQKSGEFTLGRFDPITGVNWNREYLDHGFNIEVWYQEHSHLDDD
TLITAFRATLVEECARRLNNPWGVTTGHRLAVTLQSMAHRADIVLDLHTGPKSCKHLYCPEY
ERSAAQYFSIPYTLLIPNSFGGAMDEAAFVPWWTLAEVASSHGRELGVRVSALTLELGSQER
IDLDDALEDAEGILAYLSHRGVIAETVLPKPMKRYGCFLKNYRKFHAPKAGMVEYLGKVGVP
MKATDPLVNLLRLDLYGTGEELTVLRLPEDGVPILHFASASVHQGTELYKVMTKVFEL
111000000000000000000000000000000000000000000000000000000000
000000000000000000000000000000000000000000000000000000000000
000000000000000000000000000000000000000000000000000000000000
000000000000000000000000000000000000000000000000000000000000
000000000000000000000000000000000000000000000000000000000000
000000000000000000000000000000000000000000000000000000000000
00000000000000000000000000000000000000000000000000000000
>DM_train517
MPNPPAKEDTWAFGPIGSPFPDNPVKALGQQNMYVALWYKNGRPMHGRAWNNGGVIECSFPY
NKSELTGVKDLGGQIQVLQYKGNHLSLGYWYNWIKYSDRFDKMDKGAEMLRCGDSFPILWSE
RPGGALLGYADNKTEIARFSHDGKVDEVSGSALANMLIIARELKGGPPYCECEECKSEPPKP
IVRVTLNEWADFRCGDPWPTVGTPVRALGRSLDTLPGENPDQYVALWYQSGEPVMGRIWNDG
GKIAACFGWGGHEYRQKIGSIQILYELPEAIRGFDYDWKPFPEAAQFGAKEWIPVHVDHHKG
NISPAVLIVDGKEILGKADIRNERATIGYGGTEKVLVGPAVHSCMVLCRKAKPGCTID
111110000000000000000000000000000000000000000000000000000000
000000000000000000000000000000000000000000000000000000000000
000000000000000000000000000000000000000000000000000000000001
100000000000000000000000000000000000000000000000000000000000
000000000000000000000000000000000000000000001111000000000000
00000000000000000000000000000000000000000000000000000000
>DM_train518
ASHSRKFLDVRSEEELLSCIKKETEAGKLPPNVAAGMEELYQNYRNAVIESGNPKADEIVLS
NMTVALDRILLDVEDPFVFSSHHKAIREPFDYYIFGQNYIRPLIDFGNSFVGNLSLFKDIEE
KLQQGHNVVLISNHQTEADPAIISLLLEKTNPYIAENTIFVAGDRVLADPLCKPFSIGRNLI
CVYSKKHMFDIPELTETKRKANTRSLKEMALLLRGGSQLIWIAPSGGRDRPDPSTGEWYPAP
FDASSVDNMRRLIQHSDVPGHLFPLALLCHDIMPPPSQVEIEIGEKRVIAFNGAGLSVAPEI
SFEEIAATHKNPEEVREAYSKALFDSVAMQYNVLKTAISGKQGLGASTADVSLSQPW
000000000000000000000000000000000000000000000000000000000000
000000000000000000000000000000000000000000000000000000000000

00000000000000000000000000000000000000000000000000000000000000
00000000000000000000000000000000000000000000000000000000000000
00000000000000000000000000000000000011111111110000000000000000
000000000000000000000000000000000000000000000000000000000
>DM_train520
MKMKLAVLPGDGIGPEVMDAAIRVLKTVLDNDGHEAVFENALIGGAAIDEAGTPLPEETLDI
CRRSDAILLGAVGGPKWDHNPASLRPEKGLLGLRKEMGLFANLRPVKAYATLLNASPLKRER
VENVDLVIVRELTGGLYFGRPSERRGPGENEVVDTLAYTREEIERIIEKAFQLAQIRRKKLA
SVDKANVLESSRMWREIAEETAKKYPDVELSHMLVDSTSMQLIANPGQFDVIVTENMFGDIL
SDEASVITGSLGMLPSASLRSDRFGMYEPVHGSAPDIAGQGKANPLGTVLSAALMLRYSFGL
EKEAAAIEKAVDDVLQDGYCTGDLQVANGKVVSTIELTDRLIEKLNNSAARPRIFQ
00000000000000000000000000000000000000000000000000000000000000
00000000000000000000000000000000000000000000000000000000000000
00000000000000000000000000000000000000000000000000000000000000
00000000000000000000000000000000000000000000000000000000000000
00000000000000000000000000000000000000000000000000000000000000
00000000000000000000000000000000000000000000001111111111
>DM_train522
MGSSHHHHHHSSGLVSRRHMNIDMAALHAIEVDRGISVNELLETIKSALLTAYRHTQGHQTD
ARIEIDRKTGVVRVIARETDEAGNLISEWDDTPEGFGRIAATTARQVMLQRFRDAENERTYG
EFSTREGEIVAGVIQRDSRANARGLVVVRIGTETKASEGVIPAAEQVPGESYEHGNRLRCYV
VGVTRGAREPLITLSRTHPNLVRKLFSLEVPEIADGSVEIVAVAREAGHRSKIAVRSNVAGL
NAKGACIGPMGQRVRNVMSELSGEKIDIIDYDDDPARFVANALSPAKVVSVSVIDQTARAAR
VVVPDFQLSLAIGKEGQNARLAARLTGWRIDIRGDAPPPPPGQPEPGVSRGMAHDR
11111111111111000000000000000000000000000000000000000000000000
00000000000000000000000000000000000000000000000000000000111111
11000000000000000000000000000000000000000000000000000000000000
00000000000000000000000000000000000000000000000000000000000000
00000000000000000000000000000000000000000000000000000000000000
00000000000000000000000000000000000000111111111111111111
>DM_train523
MPYRNCWSKIMKKLINDVQDVLDEQLAGLAKAHPSLTLHQDPVYVTRADAPVAGKVALLSGG
GSGHEPMHCGYIGQGMLSGACPGEIFTSPTPDKIFECAMQVDGGEGVLLIIKNYTGDILNFE
TATELLHDSGVKVTTVVIDDDVAVKDSLYTAGRRGVANTVLIEKLVGAAAERGDSLDACAEL
GRKLNNQGHSIGIALGACTVPAAGKPSFTLADNEMEFGVGIHGEPGIDRRPFSSLDQTVDEM
FDTLLVNGSYHRTLRFWDYQQGSWQEEQQTKQPLQSGDRVIALVNNLGATPLSELYGVYNRL
TTRCQQAGLTIERNLIGAYCTSLDMTGFSITLLKVDDETLALWDAPVHTPALNWGK
11111111111111111110000000000000000000000000000000000000000000
00000000000000000000000000000000000000000000000000000000000000
00000000000000000000000000000000000000000000000000000000000000
00000000000000000011111111111000000000000000000000000000000000
00000000000000000000000000000000000000000000000000000000000000
00000000000000000000000000000000000000000000000000000000
>DM_train524

AVPATIPLTITNNSGRAEQIHIYNLGTELSSGRQGWADASGAFHPWPAGGNPPTPAPDASIP
GPAPGRSTTIQIPKFSGRIYFSYGRKMEFRLTTGGLVQPAVQNPTDPNRDILFNWSEYTLND
SGLWINSTQVDMFSAPYTVGVRRGDGTTLSTGKLRPGGYNGVFNALRGQSGGWANLIQTRSD
GTVLRALSPLYGVETGALPASVMDDYINRVWNKYTGTDLIVTPFADRPDVRYTGRVSGGVLR
FTDGSGAVVTTFQKPDASSVFGCHRLLDAPNDQVRGPISRTLCAGFNRTTLLANPHQPDRSA
AGFYQEPVTNHYARIIHAHMADGKAYGFAFDDVGHHESLVHDGDPRGASLTLDPFD
100000000000000000000000000000000000000000000000000000000000
000000000000000000000000000000000000000000000000000000000000
000000000000000000000000000000000000000000000000000000000000
000000000000000000000000000000000000000000000000000000000000
000000000000000000000000000001110000000000000000000000000000
000000000000000000000000000000000000000000000000000000
>DM_train526
KKCKYSFKCVNSLKEDHNQPLFGVQFNWHSKEGDPLVFATVGSNRVTLYECHSQGEIRLLQS
YVDADADENFYTCAWTYDSNTSHPLLAVAGSRGIIRIINPITMQCIKHYVGHGNAINELKFH
PRDPNLLLSVSKDHALRLWNIQTDTLVAIFGGVEGHRDEVLSADYDLLGEKIMSCGMDHSLK
LWRINSKRMMNAIKESYDYNPNKTNRPFISQKIHFPDFSTRDIHRNYVDCVRWLGDLILSKS
CENAIVCWKPGKMEDDIDKIKPSESNVTILGRFDYSQCDIWYMRFSMDFWQKMLALGNQVGK
LYVWDLEVEDPHKAKCTTLTHHKCGAAIRQTSFSRDSSILIAVCDDASIWRWDRLR
111100000000000000000000000000000000000000000000000000000000
000000000000000000000000000000000000000000000000000000000000
000000000000000000000000000000000000000000000000000000000000
000000000000000000000000000000000000000000000000000000000000
000000000000000000000000000000000000000000000000000000000000
000000000000000000000000000000000000000000000000000011
>DM_train528
ESDSEMVPFPQLPMPIENNYRACTIPYRFPSDDPKKATPNEISWINVFANSIPSFKKRAESD
ITVPDAPARAEKFAERYAGILEDLKKDPESHGGPPDGILLCRLREQVLRELGFRDIFKKVKD
EENAKAISLFPQVVSLSDAIEDDGKRLENLVRGIFAGNIFDLGSAQLAEVFSRDGMSFLASC
QNLVPRPWVIDDLENFQAKWINKSWKKAVIFVDNSGADIILGILPFARELLRRGAQVVLAAN
ELPSINDITCTELTEILSQLKDENGQLLGVDTSKLLIANSGNDLPVIDLSRVSQELAYLSSD
ADLVIVEGMGRGIETNLYAQFKCDSLKIGMVKHLEVAEFLGGRLYDCVFKFNEVQS
111100000000000000000000000000000000000000000000000000000000
000000000000000000000000000000000000000000000000000000000000
000000000000000000000000000000000000001111111111111110000000
000000000000000000000000000000000000000000000000000000000000
000000000000000000001100000000000000000000000000000000000000
000000000000000000000000000000000000000000000000000011
>DM_train529
MKPHALIGLLAGMLLSSSLYAADSTKPLPLHIGGRVLVESPANQPVSYTYSWPAVYFETAFK
GQSLTLKFDDDQNIFRLIVDDKAPVVINKPGKVDYPVESLAPGKHRVRLEKLTETQSTSGRF
LGFYTDPSAKPLALPKRKRQIEFIGDSFTVGYGNTSPSRECTDEELFKTTNSQMAFGPLTAK
AFDADYQINASSGFGIVRNYNGTSPDKSLLSLYPYTLNNPDQLYHNKHWKPQVIVIGLGTND
FSTALNDNERWKTREALHADYVANYVKFVKQLHSNNARAQFILMNSDQSNGEIAEQVGKVVA

QLKGGGLHQVEQIVFKGLDYSGCHWHPSANDDQLLANLLITHLQQKKGIWLEHHHH
11111111111111111111111111000000000000000110000000000000000001
00000000000000000000110000000000000011111111100000000000000000
00011111111100000000000000000000000000000000000000000000000000
00000000000000000000000000000000000000000000000000000000000000
00000000000000000000000000000000000000000000000000000000000000
00000000000000000000000000000000000000000000000000011111
>DM_train530
MVVEGVKTDFGPPYFRDLLHPVIAKNYGKWKYHEVVKPGVIKRVAESGDVIYVVRFGTPRLL
SIYTVRELCDIADKYSDGYLRWTSRNNVEFFVTDESKIDDLINEVQERVGFPCGGTWDAVKG
EYGLSNIVHTQGWIHCHTPAIDASGIVKAVMDELYEYFTDHKLPAMCRISLACCANMCGAVH
ASDIAIVGIHRTPPIPNDEAIRKTCEIPSTVAACPTGALKPDMKNKTIKVDVEKCMYCGNCY
TMCPGMPLFDPENDGAAIMVGGKLSEARRMPELSKVVVPWVPNEPPRWPTLVKYVKQILEAW
AANANKHERLIEWVDRIGWERFFELTGLEFTQHLIDDYRITPYFYSEFRASTQFKW
11100000000000000000000000000000000000000000000000000000000000
00000000000000000000000000000000000000000000000000000000000000
00000000000000000000000000000000000000000000000000000000000000
00000000000000000000000000000000000000000000000000000000000000
00000000000000000000000000000000000000000000000000000000000000
00000000000000000000000000000000000000000000000000000000
>DM_train533
NVIDLFSGVGGLSLGAARAGFDVKMAVEIDQHAINTHAINFPRSLHVQEDVSLLNAEIIKGF
FKNDMPIDGIIGGPPCQGFSSIGKGNPDDSRNQLYMHFYRLVSELQPLFFLAENVPGIMQEK
YSGIRNKAFNLVSGDYDILDPIKVKASDYGAPTIRTRYFFIGVKKSLKLDISDEVFMPKMID
PVTVKDALYGLPDIIDANWQSDSESWRTIKKDRKGGFYEKLWGQIPRNVGDTESIAKLKNNI
ISGCTGTLHSKIVQERYASLSFGETDKISRSTRLDPNGFCPTLRAGTARDKGSFQAVRPIHP
YHPRVITPREAARLQGFPDWFRFHVTKWHSFRQIGNSVSPIVAEYILKGLYNLLN
00000000000000000000000000000000000000000000000000000000000000
00000000000000000000111111000000000000000000000000000000000000
00000000000000000000000000000000000000000000000000000000000000
00000000000000000000000000000000000000000000000000000000000000
00000000000000000000000000000000000000000001111111111111000000
0000000000000000000000000000000000000000000000000000000
>DM_train534
MTSRVDPANPGSELDSAIRDTLTYSNSPVPNALLTASESGFLDAAGIELDVLSGQQGTVHFT
YDQPAYTRFGGEIPPLLSEGLRAPGRTRLLGITPLLGRQGFFVRDDSPITAAADLAGRRIGV
SASAIRILRGQLGDYLELDPWRQTLVALGSWEARALLHTLEHGELGVDDVELVPISSPGVDV
PAEQLEESATVKGADLFPDVARGQAAVLASGDVDALYSWLPWAGELQATGARPVVDLGLDER
NAYASVWTVSSGLVRQRPGLVQRLVDAAVDAGLWARDHSDAVTSLHAANLGVSTGAVGQGFG
ADFQQRLVPRLDHDALALLERTQQFLLTNNLLQEPVALDQWAAPEFLNNSLNRHR
11111111111111111110000000000000000000000000000000000000000000
00000000000000000000000000000000000000000000000000000000000000
00000000000000000000000000000000000000000000000000000000000000
00000000000000000000000000000000000000000000000000000000000000

000000000000000000000000000000000000000000000000000000000000
00000000000000000000000000000000000000000000000000011
>DM_train536
MMSGQGKRLMVMAGGTGGHVFPGLAVAHHLMAQGWQVRWLGTADRMEADLVPKHGIEIDFIR
ISGLRGKGIKALIAAPLRIFNAWRQARAIMKAYKPDVVLGMGGYVSGPGGLAAWSLGIPVVL
HEQNGIAGLTNKWLAKIATKVMQAFPGAFPNAEVVGNPVRTDVLALPLPQQRLAGREGPVRV
LVVGGSQGARILNQTMPQVAAKLGDSVTIWHQSGKGSQQSVEQAYAEAGQPQHKVTEFIDDM
AAAYAWADVVVCRSGALTVSEIAAAGLPALFVPFQHKDRQQYWNALPLEKAGAAKIIEQPQL
SVDAVANTLAGWSRETLLTMAERARAASIPDATERVANEVSRVARALEHHHHHH
111111000000000000000000000000000000000000000000000000000000
000000000000000000000000000000000000000000000000000000000000
000000000000000000000000000000000000000000000000000000000000
000000000000000000000000000000000000000000000000000000000000
000000000000000000000000000000000000000000000000000000000000
000000000000000000000000000000000000000000000011111111
>DM_train537
MGFKCGIVGLPNVGKSTLFNALTKAGIEAANYPFCTIEPNTGVVPMPDPRLDALAEIVKPER
ILPTTMEFVDIAGLVAGASKGEGLGNKFLANIRETDAIGHVVRCFENDDIVHVAGKIDPLDD
IDTINTELALADLDSCERAIQRLQKRAKGGDKEAKFELSVMEKILPVLENAGMIRSVGLDKE
ELQAIKSYNFLTLKPTMYIANVNEDGFENNPYLDRVREIAAKEGAVVVPVCAAIESEIAELD
DEEKVEFLQDLGIEEPGLNRVIRAGYALLNLQTYFTAGVKEVRAWTVSVGATAPKAAAVIHT
DFEKGFIRAEVIAYEDFIQFNGENGAKEAGKWRLEGKDYIVQDGDVMHFRFNV
000000000000000000000000111111111111110000000000000000000000
000000000000000000000000000000000000000000001111111111111000
000000000000000000000000000000000000000000000000000000000000
000000000000000000000000000000000000000000000000000000000000
000000000000000000000000000000000000000000000000000000000000
00000000000000000000000000000000000000000000000000000
>DM_train538
ASTLKSGSKEVENLKKPFMPPREVHVQVTHSMPPQKIEIFKSLDNWAEENILVHLKPVEKCW
QPQDFLPDPASDGFDEQVRELRERAKEIPDDYFVVLVGDMITEEALPTYQTMLNTLDGVRDE
TGASPTSWAIWTRAWTAEENRHGDLLNKYLYLSGRVDMRQIEKTIQYLIGSGMDPRTENSPY
LGFIYTSFQERATFISHGNTARQAKEHGDIKLAQICGTIAADEKRHETAYTKIVEKLFEIDP
DGTVLAFADMMRKKISMPAHLMYDGRDDNLFDHFSAVAQRLGVYTAKDYADILEFLVGRWKV
DKLTGLSAEGQKAQDYVCRLPPRIRRLEERAQGRAKEAPTMPFSWIFDRQVKL
111111111111111110000000000000000000000000000000000000000000
000000000000000000000000000000000000000000000000000000000000
000000000000000000000000000000000000000000000000000000000000
000000000000000000000000000000000000000000000000000000000000
000000000000000000000000000000000000000000000000000000000000
00000000000000000000000000111111100000000000000000000
>DM_train539
MFQPLLDAYVESASIEKMASKSPPPLKIAVANWWGDEEIKEFKNSVLYFILSQRYTITLHQN
PNEFSDLVFGNPLGSARKILSYQNAKRVFYTGENESPNFNLFDYAIGFDELDFNDRYLRMPL

YYDRLHHKAESVNDTTAPYKLKDNSLYALKKPSHCFKEKHPNLCAVVNDESDPLKRGFASFV
ASNPNAPIRNAFYDALNSIEPVTGGGSVRNTLGYNVKNKNEFLSQYKFNLCFENTQGYGYVT
EKIIDAYFSHTIPIYWGSPSVAKDFNPKSFVNVHDFKNFDEAIDYIKYLHTHKNAYLDMLYE
NPLNTLDGKAYFYQNLSFKKILAFFKTILENDTIYHDNPFIFCRDLNEPLVTI
00000000000000000000000000000000000000000000000000000000000000
00000000000011111111110000000000000000000000000000000000000000
00000000000000000000000000000000000000000000000000000000000000
00000000000000000000000000000000000000000000000000000000000000
00000000000000000000000000000000000000000000000000000000000000
00000000000000000000000000000000000000111111111111111
>DM_train540
MHSFASLLAYGLVAGATFASASPIEARDSCTFTTAAAAKAGKAKCSTITLNNIEVPAGTTLD
LTGLTSGTKVIFEGTTTFQYEEWAGPLISMSGEHITVTGASGHLINCDGARWWDGKGTSGKK
KPKFFYAHGLDSSSITGLNIKNTPLMAFSVQANDITFTDVTINNADGDTQGGHNTDAFDVGN
SVGVNIIKPWVHNQDDCLAVNSGENIWFTGGTCIGGHGLSIGSVGDRSNNVVKNVTIEHSTV
SNSENAVRIKTISGATGSVSEITYSNIVMSGISDYGVVIQQDYEDGKPTGKPTNGVTIQDVK
LESVTGSVDSGATEIYLLCGSGSCSDWTWDDVKVTGGKKSTACKNFPSVASC
11111111111111111111111111100000000000000000000000000000000000
00000000000000000000000000000000000000000000000000000000000000
00000000000000000000000000000000000000000000000000000000000000
00000000000000000000000000000000000000000000000000000000000000
00000000000000000000000000000000000000000000000000000000000000
0000000000000000000000000000000000000000000000000000
>DM_train541
SVNNSQLVVSVAGTVEGTNQDISLKFFEIDLTSRPAHGGKTEQGLSPKSKPFATDSGAMPHK
LEKADLLKAIQEQLIANVHSNDDYFEVIDFASDATITDRNGKVYFADKDGSVTLPTQPVQEF
LLSGHVRVRPYKEKPIQNQAKSVDVEYTVQFTPLNPDDDFRPGLKDTKLLKTLAIGDTITSQ
ELLAQAQSILNKTHPGYTIYERDSSIVTHDNDIFRTILPMDQEFTYHVKNREQAYEINKKSG
LNEEINNTDLISEKYYVLKKGEKPYDPFDRSHLKLFTIKYVDVNTNELLKSEQLLTASERNL
DFRDLYDPRDKAKLLYNNLDAFGIMDYTLTGKVEDNHDDTNRIITVYMGKRP
00000000000000000000000000000000001111111111111111111111111000
00000000000000000000000000000000000000000000000000000000000000
00000000000000000000000000000000000000011111110000000000000000
00000000000000000000000000000000000000000000000000000011111111
11100000000000000000000000000000000000000000000000000000000000
0000000000000000000000000000000000000000000000000000
>DM_train542
ALTVSAGTKSKLAKPIQDLIKMIFDVESMKKAMVEFEIDLQKMPLGKLSKRQIQSAYSILNE
VQQAVSDGGSESQILDLSNRFYTLIPHDFGMKKPPLLSNLEYIQAKVQMLDNLLDIEVAYSL
LRGGNEDGDKDPIDINYEKLRTDIKVVDKDSEEAKIIKQYVKNTHAATHNAYDLKVVEIFRI
EREGESQRYKPFKQLHNRQLLWHGSRTTNFAGILSQGLRIAPPEAPVTGYMFGKGIYFADMV
SKSANYCHTSQADPIGLILLGEVALGNMYELKNASHITKLPKGKHSVKGLGKTAPDPTATTT
LDGVEVPLGNGISTGINDTCLLYNEYIVYDVAQVNLKYLLKLKFNYKTSLW
11111111000000000000000000000000000000000000000000000000000000

00000000000000000000000000000000000000000000000000000000000000
00000000000000000000000000000000000000000000000000000000000000
00000000000000000000000000000000000000000000000000000000000000
00000000000000000000000000000000000000000000000000000000000000
000000000000000000000000000000000000000000000011111
>DM_train544
VKQVFNFNAGPSALPKPALERAQKELLNFNDTQMSVMELSHRSQSYEEVHEQAQNLLRELLQ
IPNDYQILFLQGGASLQFTMLPMNLLTKGTIGNYVLTGSWSEKALKEAKLLGETHIAASTKA
NSYQSIPDFSEFQLNENDAYLHITSNNTIYGTQYQNFPEINHAPLIADMSSDILSRPLKVNQ
FGMIYAGAQKNLGPSGVTVVIVKKDLLNTKVEQVPTMLQYATHIKSDSLYNTPPTFSIYMLR
NVLDWIKDLGGAEAIAKQNEEKAKIIYDTIDESNGFYVGHAEKGSRSLMNVTFNLRNEELNQ
QFLAKAKEQGFVGLNGHRSVGGCRASIYNAVPIDACIALRELMIQFKENA
00000000000000000000000000000000000000000000000000000000000000
00000000000000000000000000000000000000000000000000000000000000
00000000000000000000000000000000000000000000000000000000000000
00000000000000000000000000011100000000000000000000000000000000
00000000000000000000000000000000000000000000000000000000000000
00000000000000000000000000000000000000000000000000
>DM_train545
PPFPWFGMDIGGTLVKLVYFEPKDITAEEEQEEVENLKSIRKYLTSNTAYGKTGIRDVHLEL
KNLTMCGRKGNLHFIRFPSCAMHRFIQMGSEKNFSSLHTTLCATGGGAFKFEEDFRMIADLQ
LHKLDELDCLIQGLLYVDSVGFNGKPECYYFENPTNPELCQKKPYCLDNPYPMLLVNMGSGV
SILAVYSKDNYKRVTGTSLGGGTFLGLCCLLTGCETFEEALEMAAKGDSTNVDKLVKDIYGG
DYERFGLQGSAVASSFGNMMSKEKRDSISKEDLARATLVTITNNIGSIARMCALNENIDRVV
FVGNFLRINMVSMKLLAYAMDFWSKGQLKALFLEHEGYFGAVGALLELFK
11000000000000000000000011111111111100000000000000000000000000
00000110000000000000000000000111111111111000000000000001111111
00000000000000000000000000000000000000000000000000000000000000
00000000000000000000000000000000000000000000000000000000000000
00000000000000000000000000000000000000000000000000000000000000
00000000000000000000000000000000000000000000000000
>DM_train546
SCDLNATNYIRGCQSKTYDGKIFPGKGGEKQWICKDTIIHGDTNGACIPPRTQNLCVGELWD
KSYGGRSNIKNDTKELLKEKIKNAIHKETELLYEYHDTGTAIISKNDKKGQKGKNDPNGLPK
GFCHAVQRSFIDYKNMILGTSVNIYEHIGKLQEDIKKIIEKGTPQQKDKIGGVGSSTENVNA
WWKGIEREMWDAVRCAITKINKKNNNSIFNGDECGVSPPTGNDEDQSVSWFKEWGEQFCIER
LRYEQNIREACTINGKNEKKCINSKSGQGDKIQGACKRKCEKYKKYISEKKQEWDKQKTKYE
NKYVGKSASDLLKENYPECISANFDFIFNDNIEYKTYYPYGDYSSICSCE
00000000000000000000000000000000000000000000000000000000000000
00000000000000000000000000000000000000000000000000000000000000
00000000000000000000000000000000000000000000001111111110000000
00000000000000000000000000000000000000001111000000000000000000
00000000000011111000000011110000000000000000000000000000000000
00000000000000000000000000000000000000000000000000

>DM_train547
MRGSHHHHHHGSLVPRGSMVAADAPAAGTDITNQLTNVTVGIDSGTTVYPHQAGYVKLNYGF
SVPNSAVKGDTFKITVPKELNLNGVTSTAKVPPIMAGDQVLANGVIDSDGNVIYTFTDYVNT
KDDVKATLTMPAYIDPENVKKTGNVTLATGIGSTTANKTVLVDYEKYGKFYNLSIKGTIDQI
DKTNNTYRQTIYVNPSGDNVIAPVLTGNLKPNTDSNALIDQQNTSIKVYKVDNAADLSESYF
VNPENFEDVTNSVNITFPNPNQYKVEFNTPDDQITTPYIVVVNGHIDPNSKGDLALRSTLYG
YNSNIIWRSMSWDNEVAFNNGSGSGDGIDKPVVPEQPDEPGEIEPIPEK
111111111111111111111111111000000000000000000000000000000000
000000000000000000000000000000000000000000000000000000000000
000000000000000000000000000000000000000000000000000000000000
000000000000000000000000000000000000000000000000000000000000
000000000000000000000000000000000000000000000000000000000000
0000000000000000000000000000000000000000000000000
>DM_train548
MIESKTYAVLGLGNGGHAFAAYLALKGQSVLAWDIDAQRIKEIQDRGAIIAEGPGLAGTAHP
DLLTSDIGLAVKDADVILIVVPAIHHASIAANIASYISEGQLIIILNPGATGGALEFRKILRE
NGAPEVTIGETSSMLFTCRSERPGQVTVNAIKGAMDFACLPAAKAGWALEQIGSVLPQYVAV
ENVLHTSLTNVNAVMHPLPTLLNAARCESGTPFQYYLEGITPSVGSLAEKVDAERIAIAKAF
DLNVPSVCEWYKESYGQSPATIYEAVQGNPAYRGIAGPINLNTRYFFEDVSTGLVPLSELGR
AVNVPTPLIDAVLDLISSLIDTDFRKEGRTLEKLGLSGLTAAGIRSAVE
111000000000000000000000000000000000000000000000000000000000
000000000000000000000000000000000000000000000000000000000000
000000000000000000000000000000000000000000000000000000000000
000000000000000000000000000000000000000000000000000000000000
000000000001111111000000000000000000000000000000000000000000
0000000000000000000000000000000000000000000000000
>DM_train549
MGDVRLSALSTLNYRNLAPGTLNFPEGVTGIYGENGAGKTNLLEAAYLALTGQTDAPRIEQL
IQAGETEAYVRADLQQGGSLSIQEVGLGRGRRQLKVDGVRARTGDLPRGGAVWIRPEDSELV
FGPPSGRRAYLDSLLSRLSARYGEQLSRYERTVSQRNAALRGGEEWAMHVWDDVLLKLGTEI
MLFRRRALTRLDELAREANAQLGSRKTLALTLTESTSPETYAADLRGRRAEELARGSTVTGP
HRDDLLLTLGDFPASDYASRGEGRTVALALRRAELELLREKFGEDPVLLLDDFTAELDPHRR
QYLLDLAASVPQAIVTGTELAPGAALTLRAQAGRFTPVADEEMQAEGTA
100000000000000000000000000000000000000000000000000000000000
000000000000000000000000000000000000000000000000000000000000
000000000000000000000000000000000000000000000000000000000000
000000000000000000000000000000000000000000000000000000000000
000000000000000000000000000000000000000000000000000000000000
0000000000000000000000000000000000000000000000000
>DM_train550
PGDVVIEELFNRIPQANVRTTSEYMQSAADSLVSTSLWNTGQPFRVESELGERPRTLVRGTV
LGQEDPYAYLEATDQETGESFEVHVPYFTERPPSNAIKQMKEEVLRLRLLRGIKNQKQAKVH
LRFIFPFDLVKDPQKKKMIRVRLDERDMWVLSRFFLYPRMQSNLQTFGEVLLSHSSTHKSLV
HHARLQLTLQVIRLLASLHHYGLVHTYLRPVDIVLDQRGGVFLTGFEHLVRDGARVVSSVSR

GFEPPELEARRATISYHRDRRTLMTFSFDAWALGLVIYWIWCADLPITKDAALGGSEWIFRS
CKNIPQPVRALLEGFLRYPKEDRLLPLQAMETPEYEQLRTELSAALPLY
00000000000000000000000100000000000000010000000000000000000000
00000000000000100000000000000010000000000000000000000000000000
00000000000000000000011111100000000000000000000000000000000000
00000000000000000000000000000000000000000000000000000000000000
00000000000000000000000000000000000000000000000111111000000000
0000000000000000000000000000000000000000000000000
>DM_train551
MKIAILGAGCYRTHAAAGITNFMRACEVAKEVGKPEIALTHSSITYGAELLHLVPDVKEVIV
SDPCFAEEPGLVVIDEFDPKEVMEAHLSGNPESIMPKIREVVKAKAKELPKPPKACIHLVHP
EDVGLKVTSDDREAVEGADIVITWLPKGNKQPDIIKKFADAIPEGAIVTHACTIPTTKFAKI
FKDLGREDLNITSYHPGCVPEMKGQVYIAEGYASEEAVNKLYEIGKIARGKAFKMPANLIGP
VCDMCSAVTATVYAGLLAYRDAVTKILGAPADFAQMMADEALTQIHNLMKEKGIANMEEALD
PAALLGTADSMCFGPLAEILPTALKVLEKHKVVEEEGKTKCEIMSQKE
00000000000000000000000000000000000000000000000000000000000000
00000000000000000000000000000000000000000000000000000000000000
00000000000000000000000000000000000000000000000000000000000000
00000000000000000000000000000000000000000000000000000000000000
00000000000000000000000000000000000000000000000000000000000000
000000000000000000000000000000000011111111111111
>DM_train552
MAVKEATSETKKRSGYEIITLTSWLLQQEQKGIIDAELTIVLSSISMACKQIASLVQRANIS
NLTGTQGAVNIQGEDQKKLDVISNEVFSNCLRSSGRTGIIASEEEDVPVAVEESYSGNYIVV
FDPLDGSSNLDAAVSTGSIFGIYSPNDESLPDFGDDSDDNTLGTEEQRCIVNVCQPGSNLLA
AGYCMYSSSVIFVLTIGKGVFVFTLDPLYGEFVLTQENLQIPKSGKIYSFNEGNYKLWDENL
KKYIDDLKEPGPSGKPYSARYIGSLVGDFHRTLLYGGIYGYPRDKKSKNGKLRLLYECAPMS
FIVEQAGGKGSDGHQRVLDIQPTEIHQRVPLYIGSTEEVEKVEKYLA
11111111111111111000000000000000000000000000000000000000000000
00011111111110000000000000000000000000000000000000000000000000
00000000000000000000000000111111111111100000000000000000000000
00000000000000000000000000000000000000000000000000000000000000
00000000000000000000000000000000000000000000000000000000000000
00000000000000000000000000000000000000000000000
>DM_train553
MSAFEKPQIIAHIQKGFNYTVFDCKWVPCSAKFVTMGNFARGTGVIQLYEIQHGDLKLLREI
EKAKPIKCGTFGATSLQQRYLATGDFGGNLHIWNLEAPEMPVYSVKGHKEIINAIDGIGGLG
IGEGAPEIVTGSRDGTVKVWDPRQKDDPVANMEPVQGENKRDCWTVAFGNAYNQEERVVCAG
YDNGDIKLFDLRNMALRWETNIKNGVCSLEFDRKDISMNKLVATSLEGKFHVFDMRTQHPTK
GFASVSEKAHKSTVWQVRHLPQNRELFLTAGGAGGLHLWKYEYPIQRSKKDSEGIEMGVAGS
VSLLQNVTLSTQPISSLDWSPDKRGLCVCSSFDQTVRVLIVTKLNKI
11110000000000000000000000000000000000110000000000000000000000
00000000000000000000000000000000000000000000000000000000000000
00000000000000000000000000000000000000000000000000000000000000

00000000000000000000000000000000000000000000000000000000000000
00000000000000000000000000000000000000000000000110000000000000
00000000000000000000000000000000000000000001111
>DM_train554
GMNRLRTSFQQTTGQISGHGKRNVGVLKTAFAAVADEMASDQYGTGAIIEPFEQKFADVLGM
DDAVFFPSGTMAQQVALRIWSDETDNRTVAYHPLCHLEIHEQDGLKELHPIETILVGAADRL
MTLDEIKALPDIACLLLELPQREIGGVAPAFSELETISRYCRERGIRLHLDGARLFEMLPYY
EKTAAEIAGLFDSIYISFYKGLGGIAGAILAGPAAFCQTARIWKRRYGGDLISLYPYIVSAD
YYYELRKDRMGQYYEQAKQLAEQFNALPGVHTTPEVPVSNMFHLHFDGQAADISPKLEQVQE
ETGLGFVGYLVDKDGYCSTEISVGDAYGELDQQTRDAGFARLRQAFS
11000000000000000000000000000000000000000000000000000000000000
00000000000000000000000000000000000000000000000000000000000000
00000000000000000000000000000000000000000000000000000000000000
00000000000000000000000000000000000000000000000000000000000000
00000000000000000000000000000000000000000000000000000000000000
00000000000000000000000000000000000000000000001
>DM_train555
MLHILCQGTPFEIGYEHGSAAKAVIARSIDFAVDLIRGKTKKTDEELKQVLSQLGRVIEERW
PKYYEEIRGIAKGAERDVSEIVMLNTRTEFAYGLKAARDGCTTAYCQLPNGALQGQNWDFFS
ATKENLIRLTIRQAGLPTIKFITEAGIIGKVGFNSAGVAVNYNALHLQGLRPTGVPSHIALR
IALESTSPSQAYDRIVEQGGMAASAFIMVGNGHEAFGLEFSPTSIRKQVLDANGRMVHTNHC
LLQHGKNEKELDPLPDSWNRHQRMEFLLDGFDGTKQAFAQLWADEDNYPFSICRAYEEGKSR
GATLFNIIYDHARREATVRLGRPTNPDEMFVMRFDEEDERSALNARL
00000000000000000000000000000000000000000000000000000000000000
00000000000000000000000000000000000111110000000000000000000000
00000000000000000000000000000000000000000000000000000000000000
00000000000000000000000000000000000000000000000000000000000000
00000000000000000000000000000000000000000000000000000000000000
00000000000000000000000000000000000000000000001
>DM_train558
MQTLHALLRDIPAPDAEAMARTQQHIDGLLKPPGSLGRLETLAVQLAGMPGLNGTPQVGEKA
VLVMCADHGVWDEGVAVSPKIVTAIQAANMTRGTTGVCVLAAQAGAKVHVIDVGIDAEPIPG
VVNMRVARGCGNIAVGPAMSRLQAEALLLEVSRYTCDLAQRGVTLFGVGELGMANTTPAAAM
VSVFTGSDAKEVVGIGANLPPSRIDNKVDVVRRAIAINQPNPRDGIDVLSKVGGFDLVGMTG
VMLGAARCGLPVLLDGFLSYSAALAACQIAPAVRPYLIPSHFSAEKGARIALAHLSMEPYLH
MAMRLGEGSGAALAMPIVEAACAMFHNMGELAASNIVLPEGNANAT
11100000000000000000000000000000000000000000000000000000000000
00000000000000000000000000000000000000000000000000000000000000
00000000000000000000000000000000000000000000000000000000000000
00000000000000000000000000000000000000000000000000000000000000
00000000000000000000000000000000000000000000000000000000000000
0000000000000000000000000000000001111111111111
>DM_train559
EIKTTTTLHRVVEETTKPLGATLVVETDISRKDVNGLARGHLVDGIPLCTPSFYADIAMQVG

QYSMQRLRAGHPGAGAIDGLVDVSDMVVDKALVPHGKGPQLLRTTLTMEWPPKAAATTRSAK
VKFATYFADGKLDTEHASCTVRFTSDAQLKSLRRSVSEYKTHIRQLHDGHAKGQFMRYNRKT
GYKLMSSMARFNPDYMLLDYLVLNEAENEAASGVDFSLGSSEGTFAAHPAHVDAITQVAGFA
MNANDNVDIEKQVYVNHGWDSFQIYQPLDNSKSYQVYTKMGQAKENDLVHGDVVVLDGEQIV
AFFRGLTLRSVPRGALRVVLQTTVKKADRQLGFKTMPSPPPPTTTM
000000000000000000000000000000000000000000000000000000000000
000000001111111111000000000000000000000000000000000000000000
000000011110000000000000000000000000000000000000000000000000
000000000000000000000000000000000000000000000000000000000000
000000000000000000000000000000000000000001100000000000000000
0000000000000000000000111111111111111111111111
>DM_train560
MKLTLKNLSMAIMMSTIVMGSSAMAADSNEKIVIAHRGASGYLPEHTLPAKAMAYAQGADYL
EQDLVMTKDDNLVVLHDHYLDRVTDVADRFPDRARKDGRYYAIDFTLDEIKSLKFTEGFDIE
NGKKVQTYPGRFPMGKSDFRVHTFEEEIEFVQGLNHSTGKNIGIYPEIKAPWFHHQEGKDIA
AKTLEVLKKYGYTGKDDKVYLQCFDADELKRIKNELEPKMGMELNLVQLIAYTDWNETQQKQ
PDGSWVNYNYDWMFKPGAMKQVAEYADGIGPDYHMLIEETSQPGNIKLTGMVQDAQQNKLVV
HPYTVRSDKLPEYTPDVNQLYDALYNKAGVNGLFTDFPDKAVKFLN
111111111111111111111111111000000000000000000000000000000000
000000000000000000000000000000000000000000000000000000000000
000000000000000000000000000000000000000000000000000000000000
000000000000000000000000000000000000000000000000000000000000
000000000000000000000000000000000000000000000000000000000000
0000000000000000000000000000000000000000000000
>DM_train561
HHHHHHPMSNTNIPTTENLYPEGAMVNKNKEGLNIDGKEVLAGSTNYYELTWDLDQYKGDKS
SKEAIQNGFYYVDDYPEEALDVRPDLVKVADEKGNQVSGVSVQQYDSLEAAPKKVQDLLKKA
NITVKGAFQLFSADNPEEFYKQYVATGTSLVITDPMTVKSEFGKTGGKYENKAYQIDFGNGY
ATEVVVNNVPKITPKKDVTVSLDPTSENLDGQTVQLYQTFNYRLIGGLIPQNHSEELEDYSF
VDDYDQAGDQYTGNYKTFSSLNLTMKDGSVIKAGTDLTSQTTAETDATNGIVTVRFKEDFLQ
KISLDSPFQAETYLQMRRIAIGTFENTYVNTVNKVAYASNTVRTTT
111111111111111111111100000000000000000000000000000000000000
000000000000000000000000000000000000000000000000000000000000
000000000000000000000000000000000000000000000000000000000000
000000000000000000000000000000000000000000000000000000000000
000000000000000000000000000000000000000000000000000000000000
0000000000000000000000000000000000000000000000
>DM_train563
MRICLFLCLCLCMASPALAQVAVFPALSGKTDAQTLVVYSSLDEPLATPMIEGFQKANPDIA
VHYEDMLTGEIYDRIVKETDAGKKTADFAFSSAMDLQVKLSNDGYAQRSDLAMSARWPAWAN
WRNTAYALTFEPAVFVYHKPSFTTEKPPATRAEFVDYLERHAKEVHGRIATYDIERSGVGFL
FMSRDQEQFGDIWSVIKAMGAAGVKVYSTSSAILERVSDGRFVLGYNILGSYAADWASRHPD
VGIVLPKDYTVVMSRIGLVPEAAANPELGRRYLEFFMSKEGQTIMARQLQIPAVSPEVAGEN
TANTMQAIHGAQLRPVPVSPGLMVYLDQVKRSRLIERWNEALRSQ

1111111111111110000000000000000111100000000000000000000000000000
0000000000000000000000000000000000000000000000000000000000000000
0000000000000000000000000000000000000000000000000000000001000000
0000000000000000000000000000000000000000000000000000000000000000
0000000000000000000000000000000000000000000000000000000000000000
000000000000000000000000000000000000000000001
>DM_train564
KELKFVTLVFRHGDRSPIDTFPTDPIKESSWPQGFGQLTQLGMEQHYELGEYIRKRYRKFLN
ESYKHEQVYIRSTDVDRTLMSAMTNLAALFPPEGVSIWNPILLWQPIPVHTVPLSEDQLLYL
PFRNCPRFQELESETLKSEEFQKRLHPYKDFIATLGKLSGLHGQDLFGIWSKVYDPLYCESV
HNFTLPSWATEDTMTKLRELSELSLLSLYGIHKQKEKSRLQGGVLVNEILNHMKRATQIPSY
KKLIMYSAHDTTVSGLQMALDVYNGLLPPYASCHLTELYFEKGEYFVEMYYRNETQHEPYPL
MLPGCSPSCPLERFAELVGPVIPQDWSTECMTTNSHQGTEDSTD
0000000000000000000000000000000000000000000000000000000000000000
0000000000000000000000000000000000000000000000000000000000000000
0000000000000000000000000000000000000000000000000000000000000000
0000000000000000000000000000000000000000000000000000000000000000
0000000000000000000000000000000000000000000000000000000000000000
00000000000000000000000000000000011111111111
>DM_train565
METMKSKANCAQNPNCNIMIFHPTKEEFNDFDKYIAYMESQGAHRAGLAKIIPPKEWKARET
YDNISEILIATPLQQVASGRAGVFTQYHKKKKAMTVGEYRHLANSKKYQTPPHQNFEDLERK
YWKNRIYNSPIYGADISGSLFDENTKQWNLGHLGTIQDLLEKECGVVIEGVNTPYLYFGMWK
TTFAWHTEDMDLYSINYLHLGEPKTWYVVPPEHGQRLERLARELFPGSSRGCGAFLRHKVAL
ISPTVLKENGIPFNRITQEAGEFMVTFPYGYHAGFNHGFNCAEAINFATPRWIDYGKMASQC
SCGEARVTFSMDAFVRILQPERYDLWKRGQDRAVVDHMEPRVPA
1111111111000000000000000000000000000000000000000000000000000000
0000000000000000000000000000000000000000000000000000000000000000
0000000000000000000000000000000000000000000000000000000000000000
0000000000000000000000000000000000000000000000000000000000000000
0000000000000000000000000000000000000000000000000000000000000000
00000000000000000000000000000000111111111111
>DM_train566
MKRVFVFQDFKSQKFWSIDVRGTDVIVNYGKLGTDGQTQVKNFSSAGEAEKAAGKLIAEKTK
KGYVETLEEVAKEMKVEAKKYALSYDEAEEGVNLMDKILKDKKLPSLKQITIGCWGYEGEDC
SDIADGIVENKEKFAHFEGLFWGDIDFEEQEISWIEQVDLSPVLDAMPLLNNLKIKGTNNLS
IGKKPRPNLKSLEIISGGLPDSVVEDILGSDLPNLEKLVLYVGVEDYGFDGDMNVFRPLFSK
DRFPNLKWLGIVDAEEQNVVVEMFLESDILPQLETMDISAGVLTDEGARLLLDHVDKIKHLK
FINMKYNYLSDEMKKELQKSLPMKIDVSDSQEYDDDYSYPMITE
0000000000000000000000000000000000000000000000000000000000000000
0000000000000000000000000000000000000000000000000000000000000000
0000000000000000000000000000000000000000000000000000000000000000
0000000000000000000000000000000000000000000000000000000000000000
0000000000000000000000000000000000000000000000000000000000000000

00000000000000000000000000000000011111100000
>DM_train567
GMYEEWKIVKREAPILGNDQLIENIWKMKREDSPYDIISLHKVNLIGGGNDAVLILPGTWSS
GEQLVTISWNGVHYTIPDYRKSIVLYLARNGFNVYTIDYRTHYVPPFLKDRQLSFTANWGWS
TWISDIKEVVSFIKRDSGQERIYLAGESFGGIAALNYSSLYWKNDIKGLILLDGGPTKHGIR
PKFYTPEVNSIEEMEAKGIYVIPSRGGPNNPIWSYALANPDMPSPDPKYKSISDFLMDSLYV
TGSANPYDYPYSKKEDMFPILASFDPYWPYRLSLERDLKFDYEGILVPTIAFVSERFGIQIF
DSKILPSNSEIILLKGYGHLDVYTGENSEKDVNSVVLKWLSQQR
110000000000000000000000000000000000000000000000000000000000
000000000000000000000000000000000000000000000000000000000000
000000000000000000000000000000000000000000000000000000000000
110000000000000000000000000000000000000000000000000000000000
000000000000000000000000000000000000000000000000000000000000
00000000000000000000000000000000000000000000
>DM_train568
MGGSHHHHHHMASSVPIPGIKDISKLKFFYGFKYLWNPTVYNKIFDKLDLTKTYKHPEELKV
LDLYPGVGIQSAIFYNKYCPRQYSLLEKRSSLYKFLNAKFEGSPLQILKRDPYDWSTYSNLI
DEERIFVPEVQSSDHINDKFLTVANVTGEGSEGLIMQWLSCIGNKNWLYRFGKVKMLLWMPS
TTARKLLARPGMHSRSKCSVVREAFTDTKLIAISDANELKGFDSQCIEEWDPILFSAAEIWP
TKGKPIALVEMDPIDFDFDVDNWDYVTRHLMILKRTPLNTVMDSLGHGGQQYFNSRITDKDL
LKKCPIDLTNDEFIYLTKLFMEWPFKPDILMDFVDMYQTEHSG
111111111111111000000000000000000000000000000000000000000000
000000000000000000000000000000000000000000000000000000000000
000000000000000000000000000000000000000000000000000000000000
000000000000000000000000000000000000000000000000000000000000
000000000000000000000000000000000000000000000000000000000000
0000000000000000000000000001111111111111111
>DM_train572
MLREFSFYDVPPAHVPPVSEPLEIACYSLSRDRELLLDDSKLSYYYPPPLFSDLNTGFPNRF
HPPKSDPDPISIVKDVLMTKGIQMNSSFLTWRGLITKIMCAPLDPRNHWETYLVMDPTSGII
MMEERTRSETSYANQDRMCYWGYKFEAISTLPEIWDACSRDQIEQRDNQDVVPDEQYCSIVK
INIGKSKLILAGEVDCIWDKKPCSAKESDVHSDDGTIEEDASNAENPNLHYVELKTSKKYPL
ENYGMRKKLLKYWAQSFLLGIGRIIIGFRDDNGILIEMKELFTHQIPKMLRPYFKPNDWTPN
RLLVVLEHALEWIKQTVKQHPPSTEFTLSYTGGSKLVLRQII
000000000000000000000000000000000000000000000000000000000000
000000000000000000000000000000000000000000000000000000000000
000000111111000000000000000000000001111111111100000000000000
000000000000000000000011111111111111111111000000000000000000
000000000000000000000000000000000000000000000000000000000000
000000000000000000000000000000000000000000
>DM_train573
MTQIIKIDPLNPEIDKIKIAADVIRNGGTVAFPTETVYGLGANAFDGNACLKIFQAKNRPVD
NPLIVHIADFNQLFEVAKDIPDKVLEIAQIVWPGPLTFVLKKTERVPKEVTAGLDTVAVRMP
AHPIALQLIRESGVPIAAPSANLATRPSPTKAEDVIVDLNGRVDVIIDGGHTFFGVESTIIN

VTVEPPVLLRPGPFTIEELKKLFGEIVIPEFAQGKKEAEIALAPGMKYKHYAPNTRLLLVEN
RNIFKDVVSLLSKKYKVALLIPKELSKEFEGLQQIILGSDENLYEVARNLFDSFRELDKLNV
DLGIMIGFPERGIGFAIMNRARKASGFSIIKAISDVYKYVNI
10000000000000000000000000000000000000000000000000000000000000
00000000000000000000000000000000000000000000000000000000000000
00000000000000000000000000000000000000000000000000000000000000
00001000000000000000000000001111111111111111111000000000000000
00000000000000000000000000000000000000000000000000000000000000
000000000000000000000000000000000000000001
>DM_train574
MDRRRFLVGLLGLGLARGQGLRVEEVVGGLEVPWALAFLPDGGMLIAERPGRIRLFREGRLS
TYAELSVYHRGESGLLGLALHPRFPQEPYVYAYRTVAEGGLRNQVVRLRHLGERGVLDRVVL
DGIPARPHGLHSGGRIAFGPDGMLYVTTGEVYERELAQDLASLGGKILRLTPEGEPAPGNPF
LGRRGARPEVYSLGHRNPQGLAWHPKTGELFSSEHGPSGEQGYGHDEVNLIVPGGNYGWPRV
VGRGNDPRYRDPLYFWPQGFPPGNLAFFRGDLYVAGLRGQALLRLVLEGERGRWRVLRVETA
LSGFGRLREVQVGPDGALYVTTSNRDGRGQVRPGDDRVLRLL
11111111111111111110000000000000000000000000000000000000000000
00000000000000000000000000000000000000000000000000000000000000
00000000000000000000000000000000000000000000000000000000000000
00000000000000000000000000000000000001111110000000000000000000
00000000000000000000000000000000000000000000000000000000000000
000000000000000000000000000000000000000000
>DM_train575
MTYVQDGNKSELRKAKRQLVTVIDLNKCLGCQTCTVACKNIWTKRPGTEHMRWNNVTTYPGK
GYPRDYERKGGGFLRGEPQPGVLPTLIDSGDDFQFNHKEVFYEGKGQTVHFHPTSKSTGKDP
AWGYNWDEDQGGGKWPNPFFFYLARMCNHCTNPACLAACPTGAIYKREDNGIVLVDQERCKG
HRHCVEACPYKAIYFNPVSQTSEKCILCYPRIEKGIANACNRQCPGRVRAFGYLDDTTSHVH
KLVKKWKVALPLHAEYGTGPNIYYVPPMGARGFGEDGEITDKTRIPLDVLEGLFGPEVKRVL
AVLHTERENMRAGRGSELMDLLISKKWSDRFGGFTNDPLTQS
11111111111111100000000000000000000000000000000000000000000000
00000000000000000000000000000000000000000000000000000000000000
00000000000000000000000000000000000000000000000000000000000000
00000000000000000000000000000000000000000000000000000000000000
00000000000000000000000000000000000000000000000000000000000000
000000000000000000000000000000000000000000
>DM_train576
MKIAIINMGNNVINFKTVPSSETIYLFKVISEMGLNVDIISLKNGVYTKSFDEVDVNDYDRL
IVVNSSINFFGGKPNLAILSAQKFMAKYKSKIYYLFTDIRLPFSQSWPNVKNRPWAYLYTEE
ELLIKSPIKVISQGINLDIAKAAHKKVDNVIEFEYFPIEQYKIHMNDFQLSKPTKKTLDVIY
GGSFRSGQRESKMVEFLFDTGLNIEFFGNAREKQFKNPKYPWTKAPVFTGKIPMNMVSEKNS
QAIAALIIGDKNYNDNFITLRVWETMASDAVMLIDEEFDTKHRIINDARFYVNNRAELIDRV
NELKHSDVLRKEMLSIQHDILNKTRAKKAEWQDAFKKAIDL
00000000000000000000000000000000000000000000000000000000000000
00000111111111000000000000000000000000000000001111111111111100

00000000000000000000000000000000000000000000000000000000000000
00000000000000000000000000000000000000000000000000000000000000
00000000000000000000000000000000000000000000000000000000000000
00000000000000000000000000000000000000000
>DM_train577
MGSDKIHHHHHHMIRAGIIGATGYTGLELVRLLKNHPEAKITYLSSRTYAGKKLEEIFPSTL
ENSILSEFDPEKVSKNCDVLFTALPAGASYDLVRELKGVKIIDLGADFRFDDPGVYREWYGK
ELSGYENIKRVYGLPELHREEIKNAQVVGNPGCYPTSVILALAPALKHNLVDPETILVDAKS
GVSGAGRKEKVDYLFSEVNESLRPYNVAKHRHVPEMEQELGKISGKKVNVVFTPHLVPMTRG
ILSTIYVKTDKSLEEIHEAYLEFYKNEPFVHVLPMGIYPSTKWCYGSNHVFIGMQMEERTNT
LILMSAIDNLVKGASGQAVQNMNIMFGLDETKGLEFTPIYP
11111111111100000000000000000000000000000000000000000000000000
00000000000000000000000000000000000000000000000000000000000000
00000000000000000000000000000000000000000000000000000000000000
00001111000000000000000000000000000000000000000000000000000000
00000000000000000000000000000000000000000000000000000000000000
00000000000000000000000000000000000000000
>DM_train578
MTSPSHASDRGGGDGDSVENQSPELRKDPVTNRWVIFSPARAKRPTDFKSKSPQNPNPKPSS
CPFCIGREQECAPELFRVPDHDPNWKLRVIENLYPALSRNLETQSTQPETGTSRTIVGFGFH
DVVIESPVHSIQLSDIDPVGIGDILIAYKKRINQIAQHDSINYIQVFKNQGASAGASMSHSH
SQMMALPVVPPTVSSRLDGTKDYFEETGKCCLCEAKSKHFVIDESSHFVSVAPFAATYPFEI
WIIPKDHSSHFHHLDDVKAVDLGGLLKLMLQKIAKQLNDPPYNYMIHTSPLKVTESQLPYTH
WFLQIVPQLSGVGGFEIGTGCYINPVFPEDVAKVMREVSLT
11111111111111111111000000000000000000011111100000111111111110
00000000000000000000000000000000000000000000011111111000000000
00000000000000000000000000000000000000000000000000000000000000
00000000000000000000000000000000000000000000000000000000000000
00000000000000000000000000000000000000000000000000000000000000
00000000000000000000000000000000000000001
>DM_train580
DTSSLIVEDAPDHVRPYVIRHYSHARAVTVDTQLYRFYVTGPSSGYAFTLMGTNAPHSDALG
VLPHIHQKHYENFYCNKGSFQLWAQSGNETQQTRVLSSGDYGSVPRNVTHTFQIQDPDTEMT
GVIVPGGFEDLFYYLGTNATDTTHTPYIPSSSDSSSTTGPDSSTISTLQSFDVYAELSFTPR
TDTVNGTAPANTVWHTGANALASTAGDPYFIANGWGPKYLNSQYGYQIVAPFVTATQAQDTN
YTLSTISMSTTPSTVTVPTWSFPGACAFQVQEGRVVVQIGDYAATELGSGDVAFIPGGVEFK
YYSEAYFSKVLFVSSGSDGLDQNLVNGGEEWSSVSFPADW
11110000000000000000000000000000000000000000000000000000000000
00000000000000000000000000000000000000000000000000000000000000
00000000000000000000000000000111111111111111100000000000000000
00000000000000000000000000000000000000000000000000000000000000
00000000000000000000000000000000000000000000000000000000000000
0000000000000000000000000000000000000000
>DM_train581

APLVHHHHHHALADNCPLEHNPDQLDSDSDRIGDTCDNNQDIDEDGHQNNLDNCPYVPNANQ
ADHDKDGKGDACDHDDDNDGIPDDKDNCRLVPNPDQKDSDGDGRGDACKDDFDHDSVPDIDD
ICPENVDISETDFRRFQMIPLDPKGTSQNDPNWVVRHQGKELVQTVNSDPGLAVGYDEFNAV
DFSGTFFINTERDDDYAGFVFGYQSSSRFYVVMWKQVTQSYWDTNPTRAQGYSGLSVKVVKS
TTGPGEHLRNALWHTGNTPGQVRTLWHDPRHIGWKDFTAYRWRLSHRPKTGFIRVVMYEGKK
IMADSGPIYDKTYAGGRLGLFVFSQEMVFFSDLKYECRDP
11111111110000000000000000000000000000000000000000000000000000
00000000000000000000000000000000000000000000000000000000000000
00000000000000000000000000000000000111110000000000000000000000
00000000000000000000000000000000000000000000000000000000000000
00000000000000000000000000000000000000000000000000000000000000
0000000000000000000000000000000000000000
>DM_train582
MTYFIDVPTMSDLVHDIGVAPFIGELAAALRDDFKRWQAFDKSARVASHSEVGVIELMPVAD
KSRYAFKYVNGHPANTARNLHTVMAFGVLADVDSGYPVLLSELTIATALRTAATSLMAAQAL
ARPNARKMALIGNGAQSEFQALAFHKHLGIEEIVAYDTDPLATAKLIANLKEYSGLTIRRAS
SVAEAVKGVDIITTVTADKAYATIITPDMLEPGMHLNAVGGDCPGKTELHADVLRNARVFVE
YEPQTRIEGEIQQLPADFPVVDLWRVLRGETEGRQSDSQVTVFDSVGFALEDYTVLRYVLQQ
AEKRGMGTKIDLVPWVEDDPKDLFSHTRGRAGKRRIRRVA
10000000000000000000000000000000000000000000000000000000000000
00000000000000000000000000000000000000000000000000000000000000
00000000000000000000000000000000000000000000000000000000000000
00000000000000000000000000000000000000000000000000000000000000
00000000000000000000000000000000000000000000000000000000000000
0000000000000000000000000000001111111111
>DM_train583
MANVAIIGTEKSGRTSLAANLGKKGTSSDITMYNNDKEGRNMVFVDAHSYPKTLKSLITALN
ISDIAVLCIPPQGLDAHTGECIIALDLLGFKHGIIALTRSDSTHMHAIDELKAKLKVITSGT
VLQDWECISLNTNKSAKNPFEGVDELKARINEVAEKIEAENAELNSLPARIFIDHAFNVTGK
GCVVLGVVKQGISKDKDKTKIFPLDRDIEIRSIQSHDVDIDSAPAGTRVGMRLKNVQAKDIE
RGFIISDKEIVTTDYTLECTVSKFTKKIEPASVLHLFVGLQSEPVRVEKILVDGNEVEEAKP
GSTCVLELSGNKKLAYSKQDRFLLANLDLTQRFAAYGFSK
00000000000000000000000000000000000000000000000000000000000000
00000000000000000000000000000000000000000000000000000000000000
00000000000000000000000000000000000000000000000000000000000011
10000000000000000000000000000000000000000000000000000000000000
00000000000000000000000000000000000000000000000000000000000000
0000000000000000000000000000000000000000
>DM_train584
MAATAAEAVASGSGEPREEAGALGPAWDESQLRSYSFPTRPIPRLSQSDPRAEELIENEEPV
VLTDTNLVYPALKWDLEYLQENIGNGDFSVYSASTHKFLYYDEKKMANFQNFKPRSNREEMK
FHEFVEKLQDIQQRGGEERLYLQQTLNDTVGRKIVMDFLGFNWNWINKQQGKRGWGQLTSNL
LLIGMEGNVTPAHYDEQQNFFAQIKGYKRCILFPPDQFECLYPYPVHHPCDRQSQVDFDNPD
YERFPNFQNVVGYETVVGPGDVLYIPMYWWHHIESLLNGGITITVNFWYKGAPTPKRIEYPL

KAHQKVAIMRNIEKMLGEALGNPQEVGPLLNTMIKGRYN
11111111111111000000000000000000000000000000000000000000000000
00000000000000000000000000000000000000000000000000000000000000
00000000000000000000000000000000000000000000000000000000000000
00000000000000000000000000000000000000000000000000000000000000
00000000000000000000000000000000000000000000000000000001110000
000000000000000000000000000000000000000
>DM_train586
MQNNNEFKIGNRSVGYNHEPLIICEIGINHEGSLKTAFEMVDAAYNAGAEVVKHQTHIVEDE
MSDEAKQVIPGNADVSIYEIMERCALNEEDEIKLKEYVESKGMIFISTLFSRAAALRLQRMD
IPAYKIGSGECNNYPLIKLVASFGKPIILSTGMNSIESIKKSVEIIREAGVPYALLHCTNIY
PTPYEDVRLGGMNDLSEAFPDAIIGLSDHTLDNYACLGAVALGGSILERHFTDRMDRPGPDI
VCSMNPDTFKELKQGAHALKLARGGKKDTIIAGEKPTKDFAFASVVADKDIKKGELLSGDNL
WVKRPGNGDFSVNEYETLFGKVAACNIRKGAQIKKTDIE
10000000000000000000000000000000000000000000000000000000000000
00000000000000000000000000000000000000000000000000000000000000
00000000000000000000000000000000000000000000000000000000000000
00000000000000000000000000000000000000000000000000000000000000
00000000000000000000000000000000000000000000000000000000000000
000000000000000000000000000000000000000
>DM_train588
VNIKTNPFKAVSFVESAIKKALDNAGYLIAEIKYDGVRGNICVDNTANSYWLSRVSKTIPAL
EHLNGFDVRWKRLLNDDRCFYKDGFMLDGELMVKGVDFNTGSGLLRTKWTDTKNQEFHEELF
VEPIRKKDKVPFKLHTGHLHIKLYAILPLHIVESGEDCDVMTLLMQEHVKNMLPLLQEYFPE
IEWQAAESYEVYDMVELQQLYEQKRAEGHEGLIVKDPMCIYKRGKKSGWWKMKPENEADGII
QGLVWGTKGLANEGKVIGFEVLLESGRLVNATNISRALMDEFTETVKEATLSQWGFFSPYGI
GDNDACTINPYDGWACQISYMEETPDGSLRHPSFVMFR
00000000000000000000000000000000000000000000000000000000000000
00000000000000000000000000000000000000000000000000000000001111
11110000000000000000000000000000000000000000000000000000000000
00000000000000000000000000000000000000000000000000000000000000
00000000000000000000000000000000000000000000000000000000011111
11100000000000000000000000000000000000
>DM_train590
MNGTEGPNFYVPFSNKTGVVRSPFEAPQYYLAEPWQFSMLAAYMFLLIMLGFPINFLTLYVT
VQHKKLRTPLNYILLNLAVADLFMVFGGFTTTLYTSLHGYFVFGPTGCNLEGFFATLGGEIA
LWSLVVLAIERYVVVCKPMSNFRFGENHAIMGVAFTWVMALACAAPPLVGWSRYIPEGMQCS
CGIDYYTPHEETNNESFVIYMFVVHFIIPLIVIFFCYGQLVFTVKEAAAQQQESATTQKAEK
EVTRMVIIMVIAFLICWLPYAGVAFYIFTHQGSDFGPIFMTIPAFFAKTSAVYNPVIYIMMN
KQFRNCMVTTLCCGKNPLGDDEASTTVSKTETSQVAPA
00000000000000000000000000000000000000000000000000000000000000
00000000000000000000000000000000000000000000000000000000000000
00000000000000000011111000000000000000000000000000000000000000
00000000000000000000000000000000000000001111111111111111000000

000000000000000000000000000000000000000000000000000000000000
00000000000000001111111111111111111111
>DM_train591
SRLQELAVIPILVIACDRSTVRRCLDKLLHYRPSAELFPIIVSQDCGHEETAQVIASYGSAV
THIRQPDLSNIAVQPDHRKFQGYYKIARHYRWALGQIFHNFNYPAAVVVEDDLEVAPDFFEY
FQATYPLLKADPSLWCVSAWNDNGKEQMVDSSKPELLYRTDFFPGLGWLLLAELWAELEPKW
PKAFWDDWMRRPEQRKGRACVRPEISRTMTFGRKGVSHGQFFDQHLKFIKLNQQFVPFTQLD
LSYLQQEAYDRDFLARVYGAPQLQVEKVRTNDRKELGEVRVQYTGRDSFKAFAKALGVMDDL
KSGVPRAGYRGIVTFLFRGRRVHLAPPQTWDGYDPSWT
111100000000000000000000000000000000000000000000000000000000
000000000000000000000000000000000000000000000000000000000000
000000000000000000000000000000000000000000000000000000000000
000000000000000000000000000000011111111111100000000000000000
000000000000000000000000000000000000000000000000000000000000
00000000000000000000000000000000000000
>DM_train592
MTGREILEKLERREFTREVLKEALSINDRGFNEALFKLADEIRRKYVGDEVHIRAIIEFSNV
CRKNCLYCGLRRDNKNLKRYRMTPEEIVERARLAVQFGAKTIVLQSGEDPYYMPDVISDIVK
EIKKMGVAVTLSLGEWPREYYEKWKEAGADRYLLRHETANPVLHRKLRPDTSFENRLNCLLT
LKELGYETGAGSMVGLPGQTIDDLVDDLLFLKEHDFDMVGIGPFIPHPDTPLANEKKGDFTL
TLKMVALTRILLPDSNIPATTAMGTIVPGGREITLRCGANVIMPNWTPSPYRQLYQLYPGKI
CVFEKDTACIPCVMKMIELLGRKPGRDWGGRKRVFETV
000000000000000000000000000000000000000000000000000000000000
000000000000000000000000000000000000000000000000000000000000
000000000000000000000000000000000000000000000000000000000000
000000000000000000000000000000000000000000000000000000000000
000000000000000000000000000000000000000000000000000000000000
00000000000000000000000000000000000001
>DM_train594
MPKFDVSKSDLERLIGRSFSIEEWEDLVLYAKCELDDVWEENGKVYFKLDSKDTNRPDLWSA
EGVARQIKWALGIEKGLPKYEVKKSNVTVYVDEKLKDIRPYGVYAIVEGLRLDEDSLSQMIQ
LQEKIALTFGRRRREVAIGIFDFDKIKPPIYYKAAEKTEKFAPLGYKEEMTLEEILEKHEKG
REYGHLIKDKQFYPLLIDSEGNVLSMPPIINSEFTGRVTTDTKNVFIDVTGWKLEKVMLALN
VMVTALAERGGKIRSVRVVYKDFEIETPDLTPKEFEVELDYIRKLSGLELNDGEIKELLEKM
MYEVEISRGRAKLKYPAFRDDIMHARDILEDVLIAYGY
100000000000000000000000000000000000000000000000000000000000
000000000000000000000000000000000000000000000000000000000000
000000000000000000000000000000000000000000000000000000000000
000000000000000000000000000000000000000000000000000000000000
000000000000000000000000000000000000000000000000000000000000
00000000000000000000000000000000000000
>DM_train595
MKERLEKLVKFMDENSIDRVFIAKPVNVYYFSGTSPLGGGYIIVDGDEATLYVPELEYEMAK
EESKLPVVKFKKFDEIYEILKNTETLGIEGTLSYSMVENFKEKSNVKEFKKIDDVIKDLRII

KTKEEIEIIEKACEIADKAVMAAIEEITEGKREREVAAKVEYLMKMNGAEKPAFDTIIASGH
RSALPHGVASDKRIERGDLVVIDLGALYNHYNSDITRTIVVGSPNEKQREIYEIVLEAQKRA
VEAAKPGMTAKELDSIAREIIKEYGYGDYFIHSLGHGVGLEIHEWPRISQYDETVLKEGMVI
TIEPGIYIPKLGGVRIEDTVLITENGAKRLTKTERELL
111000000000000000000000000000000000000000000000000000000000
000000000000000000000000000000000000000000000000000000000000
000000000000000000000000000000000000000000000000000000000000
000000000000000000000000000000000000000111111111111111100000
000000000000000000000000001000000000000000000001100000000000
0000000111100000000000000000000000111
>DM_train596
MVKRMIALDGAQGEGGGQILRSALSLSMITGQPFTITSIRAGRAKPGLLRQHLTAVKAATEI
CGATVEGAELGSQRLLFRPGTVRGGDYRFAIGSAGSCTLVLQTVLPALWFADGPSRVEVSGG
TDNPSAPPADFIRRVLEPLLAKIGIHQQTTLLRHGFYPAGGGVVATEVSPVASFNTLQLGER
GNIVQMRGEVLLAGVPRHVAEREIATLAGSFSLHEQNIHNLPRDQGPGNTVSLEVESENITE
RFFVVGEKRVSAEVVAAQLVKEVKRYLASTAAVGEYLADQLVLPMALAGAGEFTVAHPSCHL
LTNIAVVERFLPVRFSLIETDGVTRVSIEGSHHHHHH
111100000000000000000000000000000000000000000000000000000000
000000000000000000000000000000000000000000000000000000000000
000000000000000000000000000000000000000000000000000000000000
000000000000000000000000000000000000000000000000000000000000
000000000000000000000000000000000000000000000000000000000000
000000000000000000000000000111111111
>DM_train597
MRVVVIGAGVIGLSTALCIHERYHSVLQPLDVKVYADRFTPFTTTDVAAGLWQPYTSEPSNP
QEANWNQQTFNYLLSHIGSPNAANMGLTPVSGYNLFREAVPDPYWKDMVLGFRKLTPRELDM
FPDYRYGWFNTSLILEGRKYLQWLTERLTERGVKFFLRKVESFEEVARGGADVIINCTGVWA
GVLQPDPLLQPGRGQIIKVDAPWLKNFIITHDLERGIYNSPYIIPGLQAVTLGGTFQVGNWN
EINNIQDHNTIWEGCCRLEPTLKDAKIVGEYTGFRPVRPQVRLEREQLRFGSSNTEVIHNYG
HGGYGLTIHWGCALEVAKLFGKVLEERNLLTMPPSHL
000000000000000000000000000000000000000000000000000000000000
000000000000000000000000000000000000000000000000000000000000
000000000000000000000000000000000000000000000000000000000000
000000000000000000000000000000000000000000000000000000000000
000000000000000000000000000000000000000000000000000000000000
000000000000000000000000000011111111
>DM_train598
MLMREVTKEERSEFYSKEWSAKKIPKFIVDTLESREFGFDHNGEGPSDRKNQYSDIRDLEDY
IRATSPYAVYSSVAFYENPREMEGWRGAELVFDIDAKDLPLKRCNHEPGTVCPICLEDAKEL
AKDTLIILREELGFENIHVVYSGRGYHIRILDEWALQLDSKSRERILAFISASEIENVEEFR
RFLLEKRGWFVLKHGYPRVFRLRLGYFILRVNVPHLLSIGIRRNIAKKILDHKEEIYEGFVR
KAILASFPEGVGIESMAKLFALSTRFSKAYFDGRVTVDIKRILRLPSTLHSKVGLIATYVGT
KEREVMKFNPFRHAVPKFRKKEVREAYKLWRESLEYE
000000000000000000000000000000000000000000000000000000000000

00000000000000000000000000000000000000000000000000000000000000
00000000000000000000000000000000000000000000000000000000000000
00000000000000000000000000000000000000000000000000000000000000
00000000000000000000000000000000000000000000000000000000000000
0000000000000000000000000000000000111
>DM_train600
PQFMAAIYGGVEGGGTRSEVLLVSEDGKILAEADGLSTNHWLIGTDKCVERINEMVNRAKRK
AGVDPLVPLRSLGLSLSGGDQEDAGRILIEELRDRFPYLSESYLITTDAAGSIATATPDGGV
VLISGTGSNCRLINPDGSESGCGGWGHMMGDEGSAYWIAHQAVKIVFDSIDNLEAAPHDIGY
VKQAMFHYFQVPDRLGILTHLYRDFDKCRFAGFCRKIAEGAQQGDPLSRYIFRKAGEMLGRH
IVAVLPEIDPVLFQGKIGLPILCVGSVWKSWELLKEGFLLALTQGREIQAQNFFSSFTLMKL
RHSSALGGASLGARHIGHLLPMDYSANAIAFYSYTFS
00000000000000000000000000000000000000000000000000000000000000
00000000000000000000000000000000000000000000000000000000000000
00000000000000000000000000000000000000000000000000000000000000
00000000000000000000000000000000000000000000000000000000000000
00000000000000000000000000000000000000000000000011111000000000
0000000000000000000000000000000000000
>DM_train601
MASLSRFRGCLAGALLGDCVGSFYEAHDTVDLTSVLRHVQSLEPDPGTPGSERTEALYYTDD
TAMARALVQSLLAKEAFDEVDMAHRFAQEYKKDPDRGYGAGVVTVFKKLLNPKCRDVFEPAR
AQFNGKGSYGNGGAMRVAGISLAYSSVQDVQKFARLSAQLTHASSLGYNGAILQALAVHLAL
QGESSSEHFLKQLLGHMEDLEGDAQSVLDARELGMEERPYSSRLKKIGELLDQASVTREEVV
SELGNGIAAFESVPTAIYCFLRCMEPDPEIPSAFNSLQRTLIYSISLGGDTDTIATMAGAIA
GAYYGMDQVPESWQQSCEGYEETDILAQSLHRVFQKS
00000000000000000000000000000000000000000000001111111100000000
00000000000000000000000000000000000000000000000000000000000000
00000000000000000000000000000000000000000000000000000000000000
00000000000000000000000000000000000000000000000000000000000000
00000000000000000000000000000000000000000000000000000000000000
0000000000000000000000000000000000000
>DM_train602
MSSIEWNEKTFAKFAYLSDPRTKGELVAYVLTKANLKDNKYENTIVIENLKNNARRFIENAT
MPRISPDGKKIAFMRANEEKKVSEIWVADLETLSSKKILEAKNIRSLEWNEDSRKLLIVGFK
RREDEDFIFEDDVPAWFDDLGFFDGEKTTFWIFDTESEEVIEEFEKPRFSSGIWHRDKIVVN
VPHREIIPQYFKFWDIYIWEDGKEEKMFEKVSFYAVDSDGERILLYGKPEKKYMSEHNKLYI
YDGKEVMGILDEVDRGVGQAKIKDGKVYFTLFEEGSVNLYIWDGEIKPIAKGRHWIMGFDVD
EIVVYLKETATRLRELFTWDGEEKQLTDYNDPIFAKL
11111111111100000000000000000000001110000000000000000000000000
00000000000000000000000000000000000000000000000000000000000000
11111111111111111111111000000000000000000000000000000000000000
00000000000000000000000000000000000000000000000000000000000000
00000000000000000000000000000000000000000000000000000000000000
0000000000000000000000000000000011111

>DM_train604
CSALIKLLPGGHDLLVAHNTWNSYQNMLRIIKKYRLQFREGPQEEYPLVAGNNLVFSSYPGT
IFSGDDFYILGSGLVTLETTIGNKNPALWKYVQPQGCVLEWIRNVVANRLALDGATWADVFK
RFNSGTYNNQWMIVDYKAFLPNGPSPGSRVLTILEQIPGMVVVADKTAELYKTTYWASYNIP
YFETVFNASGLQALVAQYGDWFSYTKNPRAKIFQRDQSLVEDMDAMVRLMRYNDFLHDPLSL
CEACNPKPNAENAISARSDLNPANGSYPFQALHQRAHGGIDVKVTSFTLAKYMSMLAASGPT
WDQCPPFQWSKSPFHSMLHMGQPDLWMFSPIRVPWD
00000000000000000000000000000000000000000000000000000000000000
00000000000000000000000000000000000000000000000000000000000000
00000000000000000000000000000000000000000000000000000000000000
00000000000000000000000000000000000000000000000000000000000000
00000000000000000000000000000000000000000000000000000000000000
000000000000000000000000000000000011
>DM_train605
MTVSPVPSPKGRVLIAGATGFIGQFVATASLDAHRPTYILARPGPRSPSKAKIFKALEDKGA
IIVYGLINEQEAMEKILKEHEIDIVVSTVGGESILDQIALVKAMKAVGTIKRFLPSEFGHDV
NRADPVEPGLNMYREKRRVRQLVEESGIPFTYICCNSIASWPYYNNIHPSEVLPPTDFFQIY
GDGNVKAYFVAGTDIGKFTMKTVDDVRTLNKSVHFRPSCNCLNINELASVWEKKIGRTLPRV
TVTEDDLLAAAGENIIPQSVVAAFTHDIFIKGCQVNFSIDGPEDVEVTTLYPEDSFRTVEEC
FGEYIVKMEEKQPTADSAIANTGPVVGMRQVTATCA
11111111110000000000000000000000000000000000000000000000000000
00000000000000000000000000000000000000000000000000000000000000
00000000000000000000000000000000000000000000001111100000000000
00000000000000000000000000000000000000000000000000000000000000
00000000000000000000000000000000000000000000000000000000000000
000000011111111111111111111111111111
>DM_train606
MSNVYVMALDFGNGFVKGKINDEKFVIPSRIGRKTNENNQLKGFVDNKLDVSEFIINGNNDE
VLLFGNDLDKTTNTGKDTASTNDRYDIKSFKDLVECSIGLLAREVPEEVVNVVIATGMPSNE
IGTDKQAKFEKLLNKSRLIEIDGIAKTINVKGVKIVAQPMGTLLDLNMENGKVFKAFTEGKY
SVLDFGSGTTIIDTYQNMKRVEEESFVINKGTIDFYKRIASHVSKKSEGASITPRMIEKGLE
YKQCKLNQKTVIDFKDEFYKEQDSLIEEVMSNFEITVGNINSIDRIIVTGGGANIHFDSLSH
YYSDVFEKADDSQFSNVRGYEKLGELLKNKVEQESK
10000000000000000000000000000000000000000000000000000000000000
00000000000000000000000000000000000000000000000000000000000000
00000000000000000000000000000000000000000000000000000000000000
00000000000000000000000000000000000000000000111111100000000000
11111111111110000000000000000000000000000000000000000000000000
000000000000000000000000000000000111
>DM_train607
MGSDKIHHHHHHKDLLGREVEIPSNVNRIVAVGPGALRLIAYLKATDMVVGVEDFEKLRPYG
RPYILAYPELKKLPSVGPGGPGKLPDLESLITLQPDVVFITYVDRKTAKDIQEKTGIPVVVL
SYGNLGTFEDEDLFRSIELAGKILGREERAHEVVDFIRKAQEDLVTRSEGVESPTVYVGGIG
YKGAHGIDSTEAKYPPFVVLHARNVVDELGEGHKFIDPEKLLVWNPEYIFIDENGLSLVLDD

YSKHREFYESLSAVKRGKVYGILPYNYYTTNIGTALADAYFIGKVLYPERFTDIDPEEKADE
IYEFLLGKRVYGEMAEQFGGFGKIDLPSGRILRGTW
111111111100000000000000000000000000000000000000000000000000
000000000000000000000000000000000000000000000000000000000000
000000000000000000000000000000000000000000000000000000000000
000000000000000000000000000000000000000000000000000000000000
000000000000000000000000000000000000000000000000000000000000
000000000000000000000000000000000000
>DM_train608
MPSRTELLAEEYLRREGHDPQKFRYVHPDGTPYTDADGLARIARLAVPPAYQDVYVSPDAEN
ELQAFGRDAAGRLQYRYHPDFVQAGALKKWQRLTRFAGALPTLKVATTADLRASGLPPRKVM
ALMTRLLHVARFRVGSDIYARQHKTYGLSTLRQRHVVVDGNTVTFRFKGKHGVSQHKATSDR
TLAANMQKLLDLPGPWLFQTVDAGGGERRRIHSTELNAYLREVIGPFTAKDFRTWGGTLLAA
EYLAQQGTESSERQAKKVLVDCVKFVADDLGNTPAVTRGSYICPVIFDRYLDGKVLDDYEPR
TERQEAELEGLTRSEGALKRMLESERTLRQRGRALK
100000000000000011100000111111111111110000000000000000000000
000000000000000000000000000000000000000000000000000000000000
000000000000001111111111000000000000000000000000000000000000
000000000000000000000000000000000000000000000000000000000000
000000000000000000000000000000000000000000000000000000000000
000000000000000000000000000111111111
>DM_train610
VTDKTSLSYKDAGVDIDAGNALVGRIKGVVKKTRRPEVMGGLGGFGALCALPQKYREPVLVS
GTDGVGTKLRLAMDLKRHDTIGIDLVAMCVNDLVVQGAEPLFFLDYYATGKLDVDTASAVIS
GIAEGCLQSGCSLVGGETAEMPGMYHGEDYDVAGFCVGVVEKSEIIDGSKVSDGDVLIALGS
SGPHSNGYSLVRKILEVSGCDPQTTELDGKPLADHLLAPTRIYVKSVLELIEKVDVHAIAHL
TGGGFWENIPRVLPDNTQAVIDESSWQWPEVFNWLQTAGNVEHHEMYRTFNCGVGMIIALPA
PEVDKALALLNANGENAWKIGIIKASDSEQRVVIE
111111111111111111110000000000000000000000000000000000000000
000000000000000000000000000000000000000000000000000000000000
000000000000000000000000000000000000000000000000000000000000
000000000000000000000000000000000000000000000000000000000000
000000000000000000000000000000000000000000000000000000000000
00000000000000000000000000000000000
>DM_train611
MTFNPSKKLYISDVTLRDGSHAIRHQYTLDDVRAIARALDKAKVDSIEVAHGDGLQGSSFNY
GFGRHTDLEYIEAVAGEISHAQIATLLLPGIGSVHDLKNAYQAGARVVRVATHCTEADVSKQ
HIEYARNLGMDTVGFLMMSHMIPAEKLAEQGKLMESYGATCIYMADSGGAMSMNDIRDRMRA
FKAVLKPETQVGMHAHHNLSLGVANSIVAVEEGCDRVDASLAGMGAGAGNAPLEVFIAVAER
LGWNHGTDLYTLMDAADDIVRPLQDRPVRVDRETLGLGYAGVYSSFLRHAEIAAAKYNLKTL
DILVELGHRRMVGGQEDMIVDVALDLLAAHKENRA
000000000000000000000000000000000000000000000000000000000000
000000000000000000000000000000000000000000000000000000000000
000000000000000000000000000000000000000000000000000000000000

000000000000000000000000000000000000000000000000000000000000
000000000000000000000000000000000000000000000000000000000000
00000000000000000000000000000111111
>DM_train613
MPARALLPRRMGHRTLASTPALWASIPCPRSELRLDLVLPSGQSFRWREQSPAHWSGVLADQ
VWTLTQTEEQLHCTVYRGDKSQASRPTPDELEAVRKYFQLDVTLAQLYHHWGSVDSHFQEVA
QKFQGVRLLRQDPIECLFSFICSSNNNIARITGMVERLCQAFGPRLIQLDDVTYHGFPSLQA
LAGPEVEAHLRKLGLGYRARYVSASARAILEEQGGLAWLQQLRESSYEEAHKALCILPGVGT
KVADCICLMALDKPQAVPVDVHMWHIAQRDYSWHPTTSQAKGPSPQTNKELGNFFRSLWGPY
AGWAQAVLFSADLRQSRHAQEPPAKRRKGSKGPEG
111111111110000000000000000000000000000000000000000000000000
000000000000000000000000000000000000000000000000000000000000
000000000000000000000000000000000000000000000000000000000000
000000000000000000000000000000000000000000000000000000000000
000000000000000000000000000000000000000000000000000000000000
00000000000001111111111111111111111
>DM_train614
MSLRSIFWDDGLKLIDQTKLPEKLEVIECRNVEELADAIKKLAVRGAPALEAAGAYGIALAA
REREFADVDELKEHLKKAADFLASTRPTAVNLFVGIERALNAALKGESVEEVKELALREAEK
LAEEDVERNRKMGEYGAELLEDGDVVLTYCNAGRLATVDWGTALGVVRSAVEQGKEIRVIAC
ETRPLNQGSRLTCWELMEDGIDVTLITDSMVGIVMQKGMVDKVIVGADRIVRDAVFNKIGTY
TVSVVAKHHNIPFYVAAPKATFDWERTAKDVVIEERPREELIFCGKRQIAPLNVKVYNPAFD
PTPLENVTALITEYGVIYPPYEVNVPKVLKFEGGS
100000000000000000000000000000000000000000000000000000000000
000000000000000000000000000000000000000000000000000000000000
000000000000000000000000000000000000000000000000000000000000
000000000000000000000000000000000000000000000000000000000000
000000000000000000000000000000000000000000000000000000000000
00000000000000000000000000000001111
>DM_train617
MIIDIHGHYTTAPKALEDWRNRQIAGIKDPSVMPKVSELKISDDELQASIIENQLKKMQERG
SDLTVFSPRASFMAHHIGDFNVSSTWAAICNELCYRVSQLFPDNFIGAAMLPQSPGVDPKTC
IPELEKCVKEYGFVAINLNPDPSGGHWTSPPLTDRIWYPIYEKMVELEIPAMIHVSTSCNTC
FHTTGAHYLNADTTAFMQCVAGDLFKDFPELKFVIPHGGGAVPYHWGRFRGLAQEMKKPLLE
DHVLNNIFFDTCVYHQPGIDLLNTVIPVDNVLFASEMIGAVRGIDPRTGFYYDDTKRYIEAS
TILTPEEKQQIYEGNARRVYPRLDAALKAKGKLE
000000000000000000000000000000000000000000000000000000000000
000000000001111100000000000000000000000000000000000000000000
000000000000000000000001111000000000000000000000000000111111
111100000000000000000000000000000000000000000000000000000000
000000000000000000000000000000000000000000000000000000000000
0000000000000000000000000000000000
>DM_train618
MFKYVIPLCALTLAAPSFAAQTTLMLSQKSDVNYLGWSTDESKVARQEVYRGTTSNPDLRER

IAVLDAETRTFKDADTNSGLNYWYWVDVVSENQAQVVSNAVTTAPNAGPLRAAKASSECKPG
ATFENRTVDCGGVTIGTSCPNDSDKQKPLIILKNATVKNLRISASGGADGIHCDSGNCTIEN
VIWEDICEDAATNNGKTMTIVGGIAHNAKDGYGGKPDKVLQHNSKNSTTVVKGNFTLTGEHG
KLWRSCGDCSNNGGPRFLTVTSATVNGTIDSIAGVNRNYGDVATISGLKIKNYKEGKPPVCE
EFKGVVKGQGSTEKYGEKWDTTNCKVSRSGVSKL
11111111111111111110000000000000000000000000000000000000000000
00000000000000000000000000000000000000000000011111111111000000
00000000000000000000000000000000000000000000000000000000000000
00000000000000000000000000000000000000000000000000000000000000
00000000000000000000000000000000000000000000000000000000000000
0000000000000000000000000000000000
>DM_train619
MVSFKAGINLGGWISQYQVFSKEHFDTFITEKDIETIAEAGFDHVRLPFDYPIIESDDNVGE
YKEDGLSYIDRCLEWCKKYNLGLVLDMHHAPGYRFQDFKTSTLFEDPNQQKRFVDIWRFLAK
RYINEREHIAFELLNEVVEPDSTRWNKLMLECIKAIREIDSTMWLYIGGNNYNSPDELKNLA
DIDDDYIVYNFHFYNPFFFTHQKAHWSESAMAYNRTVKYPGQYEGIEEFVKNNPKYSFMMEL
NNLKLNKELLRKDLKPAIEFREKKKCKLYCGEFGVIAIADLESRIKWHEDYISLLEEYDIGG
AVWNYKKMDFEIYNEDRKPVSQELVNILARRKT
00000000000000000000000000000000000000000000000000000000000000
00000000000000000000000000000000111111110000000000000000000000
00000000000000000000000000000000000000000000000000000000000000
00000000000000000000000000000000000000000000000000000000000000
00000000000000000000000000000000000000000000000000000000000000
000000000000000000000000000000111
>DM_train620
MSASHADQPTQTVSYPQLIDLLRRIFVVHGTSPEVADVLAENCASAQRDGSHSHGIFRIPGY
LSSLASGWVDGKAVPVVEDVGAAFVRVDACNGFAQPALAAARSLLIDKARSAGVAILAIRGS
HHFAALWPDVEPFAEQGLVALSMVNSMTCVVPHGARQPLFGTNPIAFGAPRAGGEPIVFDLA
TSAIAHGDVQIAAREGRLLPAGMGVDRDGLPTQEPRAILDGGALLPFGGHKGSALSMMVELL
AAGLTGGNFSFEFDWSKHPGAQTPWTGQLLIVIDPDKGAGQHFAQRSEELVRQLHGVGQERL
PGDRRYLERARSMAHGIVIAQADLERLQELAGH
11111111100000000000000000000000000000000000000000000000000000
00000000000000000000000000000000000000000000000000000000000000
00000000000000000000000000000000000000000000000000000000000000
00000000000000000000000000000000000000000000000000000000000000
00000000000000000000000000000000000000000000000000000000000000
000000000000000000000000000000011
>DM_train621
MSLKQTVYIASPESQQIHVWNLNHEGALTLTQVVDVPGQVQPMVVSPDKRYLYVGVRPEFRV
LAYRIAPDDGALTFAAESALPGSLTHISTDHQGQFVFVGSYNAGNVSVTRLEDGLPVGVVDV
VEGLDGCHSANISPDNRTLWVPALKQDRICLFTVSDDGHLVAQDPAEVTTVEGAGPRHMVFH
PNEQYAYCVNELNSSVDVWELKDPHGNIECVQTLDMMPENFSDTRWAADIHITPDGRHLYAC
DRTASLITVFSVSEDGSVLSKEGFQPTETQPRGFNVDHSGKYLIAAGQKSHHISVYEIVGEQ
GLLHEKGRYAVGQGPMWVVVNAHEGGSHHHHHH

10000000000000000000000000000000000000000000000000000000000000
00000000000000000000000000000000000000000000000000000000000000
00000000000000000000000000000000000000000000000000000000000000
00000000000000000000000000000000000000000000000000000000000000
00000000000000000000000000000000000000000000000000000000000000
000000000000000000000000111111111
>DM_train622
GMENLGDRLEVVAELSLAPGNITLTPDGRLFLSLHQFYQPEMQVAELTQDGLIPFPPQSGNA
IITFDTVLGIKSDGNGIVWMLDNGNQSKSVPKLVAWDTLNNQLSRVIYLPPPITLSNSFVND
LAVDLIHNFVYISDPAPDDKAALIRVDLQTGLAARVLQGYPGIAPEDIDLVIDGVPVQIGQP
DGTVIRPHLGVNGIVLDAENEWLYLSPMHSTSMYRIKSADLSNLQLTDAELGSKIERYSEKP
ICDGISIDKDHNIYVGDLAHSAIGVITSADRAYKLLVTDEKLSWTDSFNFGSDGYLYFDCNQ
LHHSAPLNAGENISAPPYYIFRLKPLAAGIVGR
11111100000000000000000000000000000000000000000000000000000000
00000000000000000000000000000000000000000000000000000000000000
00000000000000000000000000000000000000000000000000000000000000
00000000000000000000000000000000000000000000000000000000000000
00000000000000000000000000000000000000000000000000000000000000
000000000000000000000000000000000
>DM_train623
MSGSHHHHHHSSGIEGRGRLQTRQDVLYLYNWTYYTPTSLIKKFEQQYNVQVVYDDYASNED
MFAKLSIGASGYDLVVPSGDFVSIMKRKHLLEKIDLSKIPNVQFIKESVRARIAYDPKMEYS
VPYYLGAAGIAVNKKAVPSYARTWSIFSRKDLAYRMSMMDDMREVMGAALASLGYNVNTKNE
QELAQAAILVTDHWKPNLVKFDSDGYAKSFASGDFVVAHGFAEAFFAETPEAMHEHIDFFIP
QDVASPVYVDSFCIPKGARNRDLAHAFINFFLEPAHYAEFLDTFGFPSTIHREAAAYQKKTP
YYSEHDLERGTLKTDVGAAIEHYNAHWNAVRFR
11111111111111111111111100000000000000000000000000000000000000
00000000000000000000000000000000000000000000000000000000000000
00000000000000000000000000000000000000000000000000000000000000
00000000000000000000000000000000000000000000000000000000000000
00000000000000000000000000000000000000000000000000000000000000
000000000000000000000000000000000
>DM_train624
GMNSDIIVADFWKNNEEILTDFDKDSFCESWTENEMWSIEFKVAQTPKNAHCYSFLDYESSV
YFRGQEFVVKQLSHDAVGKTLSKDIRAPHIYYTCQDGRQDDAITGSFTLEQCLTHIFKTDNR
GFSWEIIDPSNILEKVQQENFGNNNYLTLIDQLLDDYGVVVIPDNRHLVFKPREIYGAKTEN
FIRYKYNTDEASFDIDTLSLKTKIKGYGKVDSNGNNYFSPITYTSPEVEKWGIRWQEPVSDE
RYTVAGNMQRRLKLELQDYPATTGSVILKNDYECEKGDYVLFIYEPLGIDYDVQIVAYKKYP
FTIKAPEITLSNNKKSIVSIMAQLAKVLKGAK
11100000000000000000000000000000000000000000001111100000000000
00000000000000001110000000000000000000000000000000000000000000
00000000000000000000000000000000000000000000000000000000000000
00000000000000000000000000000000000000000000000000000000000000
00000000000000000000000000000111000000000000000000000000000000

00000000000000000000000000000011
>DM_train625
GHMLVMRPAQAADLPQVQRLAADSPVGVTSLPDDAERLRDKILASEASFAAEVSYNGEESYF
FVLEDSASGELVGCSAIVASAGFSEPFYSFRNETFVHASRSLSIHNKIHVLSLCHDLTGNSL
LTSFYVQRDLVQSVYAELNSRGRLLFMASHPERFADAVVVEIVGYSDEQGESPFWNAVGRNF
FDLNYIEAEKLSGLKSRTFLAELMPHYPIYVPLLPDAAQESMGQVHPRAQITFDILMREGFE
TDNYIDIFDGGPTLHARTSGIRSIAQSRVVPVKIGEAPKSGRPYLVTNGQLQDFRAVVLDLD
WAPGKPVALSVEAAEALGVGEGASVRLVAVGS
000000000000000000000000000000000000000000000000000000000000
000000000000000000000000000000000000000000000000000000000000
000000000000000000000000000000000000000000000000000000000000
000000000000000111111111100000000000000000000000000000000000
000000000000000000000000000000000000110000000000000000000000
00000000000000000000000000000000
>DM_train626
MKERRYLPLSQARKSGFQMDWLSEPHPVKPTFIGTQVFEEYDLQKLVDYIDWKPFFDVWQLR
GKYPNRGFPKIFNDKTVGGEARKVYDDAHNMLNTLISQKKLRARGVVGFWPAQSIQDDIHLY
AEAAVPQAAEPIATFYGLRQQAENDSASTEPYYCLSDFIAPLHSGIRDYLGLFAVACFGVEE
LSKAYEDDGDDYSSIMVKALGDRLAEAFAEELHERVRRELWAYCGSEQLDVADLRRLRYKGI
RPAPGYPSQPDHTEKLTMWRLADIEQSTGIRLTESLAMAPASAVSGLYFSNLKSKYFAVGKI
SKDQVEDYALRKNISVAEVEKWLGPILGYDTD
110000000000000000000000000000000000000000000000000000000000
000000000000000111000000000000000000000000000000000000000000
000000000000000000000000111000000000000000000000000000000000
000000000000000000000000000000000000000000000000000000000000
000000000000000000000000000000000000000000000000000000000000
00000000000000000000000000000011
>DM_train627
GLMEPPGPSTPIVQRPRILLVIDDAHTDWSKYFHGKKVNGEIEIRVEQAEFSELNLAAYVTG
GCMVDMQVVRNGTKVVSRSFKPDFILVRQHAYSMALGEDYRSLVIGLQYGGLPAVNSLYSVY
NFCSKPWVFSQLIKIFHSLGPEKFPLVEQTFFPNHKPMVTAPHFPVVVKLGHAHAGMGKIKV
ENQLDFQDITSVVAMAKTYATTEAFIDSKYDIRIQKIGSNYKAYMRTSISGNWKANTGSAML
EQVAMTERYRLWVDSCSEMFGGLDICAVKAVHSKDGRDYIIEVMDSSMPLIGEHVEEDRQLM
ADLVVSKMSQLPMPGGTAPSPLRPWAPQIKSA
111111111111110000000000000000000000000000000000000000000000
000000011111111111000000000000000000000000000000000000000000
000000000000000000000000000000000000000000000000000000000000
000000000000000000000000000000000000000000000000000000000000
000000000000000000000000000000000000000000000000000000000000
00000000000011111111111111111111
>DM_train628
LTCGTNSGFVCKGTQTQYAGGFAPGVGYGGFGGGSCTATKTPVIFIHGNGDNAISFDMPPGN
VSGYGTPARSVYAELKARGYNDCEIFGVTYLSSSEQGSAQYNYHSSTKYAIIKTFIDKVKAY
TGKSQVDIVAHSMGVSMSLATLQYYNNWTSVRKFINLAGGIRGLYSCYYTGYANAAAPTCGS

QNYYNSYTFGFFPEGWYYGVWVSNPWTGSGSTNSMRDMPAKRTAVSFYTLSAGFKDQVGCAT
ASFWAGCDSAAKFASTTSNVKAQINVGAGSNATQADYDWADGMPYNAGGGDTTNGVGHFRTK
TNTGAIIQRMLLTTCTGLDCAAEYTTGPKAAY
00000000000000000000000000000000000000000000000000000000000000
00000000000000000000000000000000000000000000000000000000000000
00000000000000000000000000000000000000000000000000000000000000
00000000000000000000000000000000000000000000000000000000000000
00000000000000000000000000000000000000000000110000000000000000
00000000000000000000000000000000
>DM_train629
PFKESQARGLLQDNMANSYNGGDFEDGLLNLSKEVFPTDKYLYQDGQFLDKKTINAYLNPKY
TKREIDKMSEKDKKDKKANENLGLNPSHEGETNPEKIAEKSPAYLSNILEQDFYGGGDTKGK
NIKGMTIGLAMNSVYYYKKEKDGPTFSKKLDDSEVKKQGKQMASEILSRLRENDDLKDIPIH
FAIYKQSSEDSITPGEFITQATAEKSQTKLNEWHNINEKSALLPSSTAADYDENLNNNFKQF
NDNLQSYFSNFTQAVGKVKFVDKKPQRLVVDLPIDYYGQAETIGITQYVTEQANKYFDKIDN
YEIRIKDGNQPRALISKTKDDKEPQVHIYSN
11111111100000000000000000000000000000000000000000000000000000
00000000000000000000000000000000000000000000000000000001111110
00000000000000000000000000000000000000000000000000000000000000
00000000000000000000000000000010000000000000000000000000000000
00000111111100000000000000000000000000000000000000000000000000
0000000000000000000000000000000
>DM_train630
MPTLTARSVIPPYMLRRIIEHGSLPQRDCALHTLNHVQSLLGNKPLRAPGAKTSTGGEVIRD
IYDAENSTQLPGKQVRNEGQASNHDVAVDEAYDYLGVTYDFFWQAFKRNSLDNQGLPLTGSV
HYGKEYQNAFWNGQQMVFGDGDGEIFNRFTIAIDVVGHALAHGVTESEAGLIYFQQAGALNE
SLSDVFGSLVKQFHLKQTADKADWLIGEGLLAKGINGKGLRSMSAPGTAYDDPLLGKDPQPA
SMKDYIQTKEDNGGVHLNSGIPNRAFYLAATALGGYAWEKAGYIWYDTLCDKALPQDADFAT
FARTTVKHAEQRFDSKVAQKVQQAWHQVGVA
11111000000000000000000000000000000111111111111100000000000000
00000000000000000000000000000000000000000000000000000000000000
00000000000000000000000000000000000000000000000000000000000000
00000000000000000000000000000000000000000000000000000000000000
00000000000000000000000000000000000000000000000000000000000000
0000000000000000000000000000000
>DM_train631
MDRNSLSNSLKTCYKYLNQTSRSFAAVIQALDGEMRNAVCIFYLVLRALDTLEDDMTISVEK
KVPLLHNFHSFLYQPDWRFMESKEKDRQVLEDFPTISLEFRNLAEKYQTVIADICRRMGIGM
AEFLDKHVTSEQEWDKYCHYVAGLVGIGLSRLFSASEFEDPLVGEDTERANSMGLFLQKTNI
IRDYLEDQQGGREFWPQEVWSRYVKKLGDFAKPENIDLAVQCLNELITNALHHIPDVITYLS
RLRNQSVFNFCAIPQVMAIATLAACYNNQQVFKGAVKIRKGQAVTLMMDATNMPAVKAIIYQ
YMEEIYHRIPDSDPSSSKTRQIISTIRTQN
11111110000000000000000000000000000000000000000000000000000000
00000000000000000000000000000000000000000000000000000000000000

00000000000000000000000000000000000000000000000000000000000000
00000000000000000000000000000000000000000000000000000000000000
00000000000000000000000000000000000011111111111110000000000000
000000000000000000000000000000
>DM_train632
GMHPNIATLLSANLGESRTRHLLSLVSVPDGLPSDAEGRATRAEIAQALNMVLFAGILDRVP
TGRAYTDDVAATGGKVVFDHGALRTVKWRDNGALPEGEAAFTRILRPLGYRLNGNYPLDRIS
MTGRSYAHADAPEGIAQFFVSEFHPERFSDAFREAVGRVTGNSADPLTPRAQTLLWQLDRDG
VLTVADGAELIGLLVPCFERQHGVPRLADYETLLRESAEMAWIATEGNAFNHATDRVDDVFG
LSEQQKALGRPMKDKVEVSGSGRVKQTAFRADTVRRQFIGAQGETVERDVPGSFYEFITRDR
FADAPAASPRVDLGFDAGNAQGIFKMTAAV
00000000000000000000000000000000000000000000000000000000000000
00000000000000000000000000000000000000000000000000000000000000
00000000000000000000000000000000000000000000000000000000000000
00000000000000000000000000000000000000000000000000000000000000
00000000000000000000000000000000000000000000000000000000000000
011111111000000000111111111111
>DM_train633
NTATEFDGPYVITPISGQSTAYWICDNRLKTTSIEKLQVNRPEHCGDLPETKLSSEIKQIMP
DTYLGIKKVVALSDVHGQYDVLLTLLKKQKIIDSDGNWAFGEGHMVMTGDIFDRGHQVNEVL
WFMYQLDQQARDAGGMVHLLMGNHEQMVLGGDLRYVHQRYDIATTLINRPYNKLYSADTEIG
QWLRSKNTIIKINDVLYMHGGISSEWISRELTLDKANALYRANVDASKKSLKADDLLNFLFF
GNGPTWYRGYFSETFTEAELDTILQHFNVNHIVVGHTSQERVLGLFHNKVIAVDSSIKVGKS
GELLLLENNRLIRGLYDGTRETLQENSLNQ
11000000000000000000000000000000000000000000000000000000000000
00000000000000000000000000000000000000000000000000000000000000
00000000000000000000000000000000000000000000000000000000000000
00000000000000000000000000000000000000000000000000000000000000
00000000000000000000000000000000000000000000000000000000000000
000000000000000000000000000000
>DM_train634
AEIYNKDGNKVDLYGKAVGLHYFSKGNGENSYGGNGDMTYARLGFKGETQINSDLTGYGQWE
YNFQGNNSEGADAQTGNKTRLAFAGLKYADVGSFDYGRNYGVVYDALGYTDMLPEFGGDTAY
SDDFFVGRVGGVATYRNSNFFGLVDGLNFAVQYLGKNERDTARRSNGDGVGGSISYEYEGFG
IVGAYGAADRTNLQEAQPLGNGKKAEQWATGLKYDANNIYLAANYGETRNATPITNKFTNTS
GFANKTQDVLLVAQYQFDFGLRPSIAYTKSKAKDVEGIGDVDLVNYFEVGATYYFNKNMSTY
VDYIINQIDSDNKLGVGSDDTVAVGIVYQF
00000000000000000000000000100000000000000000000000000000000000
00000000000000000000000000000000000000000000000000000000000000
00000000000000000000000000000000000000000000000000000000000000
00000000000000000000000000000000000000000000000000000000000000
00000000000000000000000000000000000000000000000000000000000000
000000000000000000000000000000
>DM_train635

MRGSPHHHHHHGSAKNVVLDHAGNLDDFVAMVLLASNTEKVRLIGALCTDADCFVENGFNVT
GKIMCLMHNNMNLPLFPIGKSAATAVNPFPKEWRCLAKNMDDMPILNIPENVELWDKIKAEN
EKYEGQQLLADLVMNSEEKVTICVTGPLSNVAWCIDKYGEKFTSKVEECVIMGGAVDVRGNV
FLPSTDGTAEWNIYWDPASAKTVFGCPGLRRIMFSLDSTNTVPVRSPYVQRFGEQTNFLLSI
LVGTMWAMCTHCELLRDGDGYYAWDALTAAYVVDQKVANVDPVPIDVVVDKQPNEGATVRTD
AENYPLTFVARNPEAEFFLDMLLRSARAC
111111111111100000000000000000000000000000000000000000000000
000000000000000000000000000000000000000000000000000000000000
000000000000000000000000000000000000000000000000000000000000
000000000000000000000000000000000000000000000000000000000000
000000001111111111110000000000000000000000000000000000000000
00000000000000000000000000000
>DM_train636
MVSVARELTVGINGFGRIGRLVLRACMEKGVKVVAVNDPFIDPEYMVYMFKYDSTHGRYKGS
VEFRNGQLVVDNHEISVYQCKEPKQIPWRAVGSPYVVESTGVYLSIQAASDHISAGAQRVVI
SAPSPDAPMFVMGVNENDYNPGSMNIVSNASCTTNCLAPLAKVIHERFGIVEGLMTTVHSYT
ATQKTVDGPSRKAWRDGRGAHQNIIPASTGAAKAVTKVIPELKGKLTGMAFRVPTPDVSVVD
LTCRLAQPAPYSAIKEAVKAAAKGPMAGILAYTEDEVVSTDFLGDTHSSIFDAKAGIALNDN
FVKLISWYDNEYGYSHRVVDLLRYMFSRD
111110000000000000000000000000000000000000000000000000000000
000000000000000000000000000000000000000000000000000000000000
000000000000000000000000000000000000000000000000000000000000
000000000000000000000000000000000000000000000000000000000000
000000000000000000000000000000000000000000000000000000000000
00000000000000000000000000000
>DM_train637
PFTYSIEATRNLATTERCIQDIRNAPVRNRSTQFQLAQQNMLAYTFGEVIPGFASAGINGMD
YRDVIGRPVENAVTEGTHFFRDDFRVDSNAKAKVAGDIFEIVSSAVMWNCAARWNSLMVGEG
WRSQPRYSRPTLSPSPRRQVAVLNLPRSFDWVSLLVPESQEVIEEFRAGLRKDGLGLPTSTP
DLAVVVLPEEFQNDEMWREEIAGLTRPNQILLSGAYQRLQGRVQPGEISLAVAFKRSLRSDR
LYQPLYEANVMQLLLEGKLGAPKVEFEVHTLAPEGTNAFVTYEAASLYGLAEGRSAVHRAIR
ELYVPPTAADLARRFFAFLNERMELVNG
000000000000000000000000000000000000000000000000000000000000
000000000000000000000000000000000000000000000000000000000000
000000000000000000000000000000000000000000000000000000000000
000000000000000000000000000000000000000000000000000000000000
000000000000000000000000000000000000000000000000001111000000
0000000000000000000000000000
>DM_train638
MKLATLKDSTRDGKLVVVSKDLTRCSEVGHIARTLQAALDDWAHAGPRLERVAEGIETGAQP
TMRFHEHDAASPLPRAFQWADGSAYVNHVELVRKARNAEMPASFWTDPLIYQGGSDSFLGPR
DPILMADDAWGIDMEGEAAVIVDDVPMGATLDEAKAAIRLVMLVNDVSLRGLIPGELAKGFG
FYQSKPSSAFSPVAVTPEELGEAWDGGKLHLPLHVDLNGEPFGRANAGIDMTFDFPQLIVHA
ARTRPLSAGTIIGSGTVSNKLEGGPGRPVSEGGAGYSCIAELRMIETIEGGAPKTQFLKFGD

VVRIEMKDRTGHSIFGAIEQKVGKYERG
100000000000000000000000000000000000000000000000000000000000
000000000000000000000000000000000000000000000000000000000000
000000000000000000000000000000000000000000000000000000000000
000000000000000000000000000000000000000000000000000000000000
000000000000000000000000000000000000000000000000000000000000
0000000000000000000000000001
>DM_train639
MGSDKIHHHHHHMAFTPFPPRQPTASARLPLTLMTLDDWALATITGADSEKYMQGQVTADVS
QMAEDQHLLAAHCDAKGKMWSNLRLFRDGDGFAWIERRSVREPQLTELKKYAVFSKVTIAPD
DERVLLGVAGFQARAALANLFSELPSKEKQVVKEGATTLLWFEHPAERFLIVTDEATANMLT
DKLRGEAELNNSQQWLALNIEAGFPVIDAANSGQFIPQATNLQALGGISFKKGCYTGQEMVA
RAKFRGANKRALWLLAGSASRLPEAGEDLELKMGENWRRTGTVLAAVKLEDGQVVVQVVMNN
DMEPDSIFRVRDDANTLHIEPLPYSLEE
111111111111110000000000000000000000000000000000000000000000
000000000000000000000000000000000000000000000000000000000000
000000000000000000000000000000000000000000000000000000000000
000000000000000000000000000000000000000000000000000000000000
000000000000000000000000000000000000000000000000000000000000
0000000000000000000000000001
>DM_train640
MGSQSETVCVTGASGFIGSWLVMRLLERGYTVRATVRDPTNVKKVKHLLDLPKAETHLTLWK
ADLADEGSFDEAIKGCTGVFHVATPMDFESKDPENEVIKPTIEGMLGIMKSCAAAKTVRRLV
FTSSAGTVNIQEHQLPVYDESCWSDMEFCRAKKMTAWMYFVSKTLAEQAAWKYAKENNIDFI
TIIPTLVVGPFIMSSMPPSLITALSPITGNEAHYSIIRQGQFVHLDDLCNAHIYLFENPKAE
GRYICSSHDCIILDLAKMLREKYPEYNIPTEFKGVDENLKSVCFSSKKLTDLGFEFKYSLED
MFTGAVDTCRAKGLLPPSHEKPVDGKT
111110000000000000000000000000000000000000000000000000000000
000000000000000000000000000000000000000000000000000000000000
000000000000000000000000000000000000000000000000000000000000
000000000000000000000000000000000000000000000000000000000000
000000000000000000000000000000000000000000000000000000000000
000000000000000000011111111
>DM_train641
MVLVIEIIRKHLPRVLKVPATRILLLVLAVIIYGTAGFHFIEGESWTVSLYWTFVTIATVGY
GDYSPSTPLGMYFTVTLIVLGIGTFAVAVERLLEFLINREQMKLMGLIDVAKSRHVVICGWS
ESTLECLRELRGSEVFVLAEDENVRKKVLRSGANFVHGDPTRVSDLEKANVRGARAVIVDLE
SDSETIHCILGIRKIDESVRIIAEAERYENIEQLRMAGADQVISPFVISGRLMSRSIDDGYE
AMFVQDVLAEESTRRMVEVPIPEGSKLEGVSVLDADIHDVTGVIIIGVGRGDELIIDPPRDY
SFRAGDIILGIGKPEEIERLKNYISA
111111111111111111000000000000000000000000000000000000000000
000000000000000000000000000000000011111111111111111000000000
000000000000000000000000000000000000000000000000000000000000
000000000000000000000000000000000000000000000000000000000000

000000000000000000000000000000000000000000000000000000000000
00000000000000000000000000
>DM_train642
PQDVDFHIPLPGRQSPDHARAEAEQLAWPRSLGLIRSDAAAERHLRGGYADLASRFYPHATG
ADLDLGVDLMSWFFLFDDLFDGPRGENPEDTKQLTDQVAAALDGPLPDTAPPIAHGFADIWR
RTCEGMTPAWCARSARHWRNYFDGYVDEAESRFWNAPCDSAAQYLAMRRHTIGVQPTVDLAE
RAGRFEVPHRVFDSAVMSAMLQIAVDVNLLLLDIASLEKEEARGEQNNMVMILRREHGWSKS
RSVSHMQNEVRARLEQYLLLESCLPKVGEIYQLDTAEREALERYRTDAVRTVIRGSYDWHRS
SGRYDAEFALAAGAQGYLEELGSSAH
111110000000000000000000000000000000000000000000000000000000
000000000000000000000000000000000000000000000000000000000000
000000000000000000000000000000011111110000000000000000000000
000000000000000000000000000000000000000000111100000111111100
000000000000000000000000000000000000000000000000000000000001
11111111111111111111111111
>DM_train643
RENFPGVSWDSLPDELLLGIFSCLCLPELLKVSGVCKRWYRLASDESLWQTLDLTGKNLHPD
VTGRLLSQGVIAFRCPRSFMDQPLAEHFSPFRVQHMDLSNSVIEVSTLHGILSQCSKLQNLS
LEGLRLSDPIVNTLAKNSNLVRLNLSGCSGFSEFALQTLLSSCSRLDELNLSWCFDFTEKHV
QVAVAHVSETITQLNLSGYRKNLQKSDLSTLVRRCPNLVHLDLSDSVMLKNDCFQEFFQLNY
LQHLSLSRCYDIIPETLLELGEIPTLKTLQVFGIVPDGTLQLLKEALPHLQINCSHFTTIAR
PTIGNKKNQEIWGIKCRLTLQKPSCL
111111000000000000000000000000000000000000000000000000000000
000000000000000000000000000000000000000000000000000000000000
000000000000000000000000000000000000000000000000000000000000
000000000000000000000000000000000000000000000000000000000000
000000000000000000000000000000000000000000000000000000000000
00000000000000000000011111
>DM_train644
MPVLNLNDPQAVERYEEFMRQSPYGQVTQDLGWAKVKNNWEPVDVYLEDDQGAIIAAMSMLL
GDTPTDKKFAYASKGPVMDVTDVDLLDRLVDEAVKALDGRAYVLRFDPEVAYSDEFNTTLQD
HGYVTRNRNVADAGMHATIQPRLNMVLDLTKFPDAKTTLDLYPSKTKSKIKRPFRDGVEVHS
GNSATELDEFFKTYTTMAERHGITHRPIEYFQRMQAAFDADTMRIFVAEREGKLLSTGIALK
YGRKIWYMYAGSMDGNTYYAPYAVQSEMIQWALDTNTDLYDLGGIESESTDDSLYVFKHVFV
KDAPREYIGEIDKVLDPEVYAELVKD
100000000000000000000000000000000000000000000000000000000000
000000000000000000000000000000000000000000000000000000000000
000000000000000000000000000000000000000000000000000000000000
000000000000000000000000000000000000000000000000000000000000
000000000000000000000000000000000000000000000000000000000000
00000000000000000000000000
>DM_train645
MALSMTGGQQMGRGSMSRDPLPFFPPLYLGGPEITTENCEREPIHIPGSIQPHGALLTADGH
SGEVLQMSLNAATFLGQEPTVLRGQTLAALLPEQWPALQAALPPGCPDALQYRATLDWPAAG

HLSLTVHRVGELLILEFEPTEAWDSTGPHALRNAMFALESAPNLRALAEVATQTVRELTGFD
RVMLYKFAPDATGEVIAEARREGLHAFLGHRFPASDIPAQARALYTRHLLRLTADTRAAAVP
LDPVLNPQTNAPTPLGGAVLRATSPMHMQYLRNMGVGSSLSVSVVVGGQLWGLIACHHQTPY
VLPPDLRTTLESLGRLLSLQVQVKEA
111111111111111111000000000000000000000000000000000000000000
000000000000000000000000000000000000000000000000000000000000
000000000000000000000000000000000000000000000000000000000000
000000000000000000000000000000000000000000000000000000000000
000000000000000000000000000000000000000000000000000000000000
00000000000000000000000000
>DM_train646
MGADIGDLFEREEVELEYFSGKKIAVDAFNTLYQFISIIRQPDGTPLKDSQGRITSHLSGIL
YRVSNMVEVGIRPVFVFDGEPPEFKKAEIEERKKRRAEAEEMWIAALQAGDKDAKKYAQAAG
RVDEYIVDSAKTLLSYMGIPFVDAPSEGEAQAAYMAAKGDVEYTGSQDYDSLLFGSPRLARN
LAITGKRKLPGKNVYVDVKPEIIILESNLKRLGLTREQLIDIAILVGTDYNEGVKGVGVKKA
LNYIKTYGDIFRALKALKVNIDHVEEIRNFFLNPPVTDDYRIEFREPDFEKAIEFLCEEHDF
SRERVEKALEKLKALKSTQATLERWF
110000000000000000000000000000000000000000000000000000000000
000000000000000000000000000000000000000000000000000000000000
000000000000000000000000000000000000000000000000000000000000
000111111111111100000000000000000000000000000000000000000000
000000000000000000011110000000000000000000000000000000000000
00000000000000111111111111
>DM_train648
SMGECSQKGPVPFSHCLPTEKLQRCEKIGEGVFGEVFQTIADHTPVAIKIIAIEGPDLVNGS
HQKTFEEILPEIIISKELSLLSGEVCNRTEGFIGLNSVHCVQGSYPPLLLKAWDHYNSTKGS
ANDRPDFFKDDQLFIVLEFEFGGIDLEQMRTKLSSLATAKSILHQLTASLAVAEASLRFEHR
DLHWGNVLLKKTSLKKLHYTLNGKSSTIPSCGLQVSIIDYTLSRLERDGIVVFCDVSMDEDL
FTGDGDYQFDIYRLMKKENNNRWGEYHPYSNVLWLHYLTDKMLKQMTFKTKCNTPAMKQIKR
KIQEFHRTMLNFSSATDLLCQHSLFK
111111100000000000000000000000000000000000000000000000000000
000000000000000000000000000000000000000000000000000000000000
000000000000000000000000000000000000000000000000000000000000
000000000000000000000000000000000000000000000000000000000000
000000000000000000000000000000000000000000000000000000000000
00000000000000000000000000
>DM_train649
MAELKLGYKASAEQFAPRELVELAVAAEAHGMDSATVSDHFQPWRHQGGHAPFSLSWMTAVG
ERTNRLLLGTSVLTPTFRYNPAVIAQAFATMGCLYPNRVFLGVGTGEALNEIATGYEGAWPE
FKERFARLRESVGLMRQLWSGDRVDFDGDYYRLKGASIYDVPDGGVPVYIAAGGPAVAKYAG
RAGDGFICTSGKGEELYTEKLMPAVREGAAAADRSVDGIDKMIEIKISYDPDPELAMNNTRF
WAPLSLTAEQKHSIDDPIEMEKAADALPIEQIAKRWIVASDPDEAVEKVGQYVTWGLNHLVF
HAPGHDQRRFLELFQSDLAPRLRRLG
110000000000000000000000000000000000000000000000000000000000

000000000000000000000000000000000000000000000000000000000000
000000000000000000000000000000000000000000000000000000000000
000000000000000000000000000000000000000000000000000000000000
000000000000000000000000000000000000000000000000000000000000
0000000000000000000000011
>DM_train650
MGSDKIHHHHHHMEMRFFVLGAGSWGTVFAQMLHENGEEVILWARRKEIVDLINVSHTSPYV
EESKITVRATNDLEEIKKEDILVIAIPVQYIREHLLRLPVKPSMVLNLSKGIEIKTGKRVSE
IVEEILGCPYAVLSGPSHAEEVAKKLPTAVTLAGENSKELQKRISTEYFRVYTCEDVVGVEI
AGALKNVIAIAAGILDGFGGWDNAKAALETRGIYEIARFGMFFGADQKTFMGLAGIGDLMVT
CNSRYSRNRRFGELIARGFNPLKLLESSNQVVEGAFTVKAVMKIAKENKIDMPISEEVYRVV
YEGKPPLQSMRDLMRRSLKDEFWAS
111111111111000000000000000000000000000000000000000000000000
000000000000000000000000000000000000000000000000000000000000
000000000000000000000000000000000000000000000000000000000000
000000000000000000000000000000000000000000000000000000000000
000000000000000000000000000000000000000000000000000000000000
0000000000000001111111111
>DM_train651
GVLAPVVPGKALEFPQDFGAHNDFRIEWWYVTGWLETPTGKPLGFQITFFRTATEIDRDNPS
HFAPDQLIIAHVALSDPAIGKLQHDQKIARAGFDLAYARTGNTDVKLDDWIFVRETDGRYRT
RIEAEDFTLTFILTPSQPLMLQGENGFSRKGPGAPQASYYYSEPHLQVSGIINRQGEDIPVT
GTAWLDREWSSEYLDPNAAGWDWISANLDDGSALMAFQIRGKDDSKIWAYAALRDASGHTRL
FTPDQVSFHPIRTWRSARTQAVYPVATRVLTGETEWQITPLMDDQELDSRASAGAVYWEGAV
TFTRDGQPAGRGYMELTGYVRPLSM
110000000000000000000000000000000000000000000000000111111110
000000000000000000000000000000000000000000000000000000000000
000000000000000000000000000000000000000000000000000000000000
000000000000000000000000000000000000000000000000000000000000
000000000000000000000000000000000000000000000000000000000000
0000000000000000000011111
>DM_train652
CSSLSIRTTDDKSLFARTMDFTMEPDSKVIIVPRNYGIRLLEKENVVINNSYAFVGMGSTDI
TSPVLYDGVNEKGLMGAMLYYATFATYADEPKKGTRGINPVYVISQVLGNCVTVDDVIEKLT
SYTLLNEANIILGFAPPLHYTFTDASGESIVIEPDKTGITIHRKTIGVMTASPGYEWHQTNL
RAYIGVTPNPPQDIMMGDLDLTPFGQGAGGLGLPGDFTPSARFLRVAYWKKYTEKAKNETEG
VTNLFHILSSVNIPKGVVLTNEGKTDYTIYTSAMCAQSKNYYFKLYDNSRISAVSLMAENLN
SQDLITFEWDRKQDIKQLNQVNVMS
000000000000000000000000000000000000000000000000000000000000
000000000000000000000000000000000000000000000000000000000000
000000000000000000000000000000000000000000000000000000000000
000000000000000000000000000000000000000000000000000000000000
000000000000000000000000000000000000000000000000000000000000
0000000000000000000111111

>DM_train653
MNDSSVIYRAIVTSKFRTEKMLNFYNSIGSGPDKNTIFITFGRSEPWSSNENEVGFAPPYPT
DSVLGVTDMWTHMMGTVKVLPSMLDAVIPRRDWGDTRYPDPYTFRINDIVVCNSAPYNATES
GAGWLVYRCLDVPDTGMCSIASLTDKDECLKLGGKWTPSARSMTPPEGRGDAEGTIEPGDGY
VWEYLFEIPPDVSINRCTNEYIVVPWPEELKEDPTRWGYEDNLTWQQDDFGLIYRVKANTIR
FKAYLDSVYFPEAALPGNKGFRQISIITNPLEAKAHPNDPNVKAEKDYYDPEDLMRHSGEMI
YMENRPPIIMAMDQTEEINILFTF
111111000000000000000000000000000000000000000000000000000000
000000000000000000000000000000000000000000000000000000000000
000000000000000000000000000000000000000000000000000000000000
000000000000000000000000000000000000000000000000000000000000
000000000000000000000000000000000000000000000000000000000000
000000000000000000000000
>DM_train654
GRRRRSVQWCAVSQPEATKCFQWQRNMRRVRGPPVSCIKRDSPIQCIQAIAENRADAVTLDG
GFIYEAGLAPYKLRPVAAEVYGTERQPRTHYYAVAVVKKGGSFQLNELQGLKSCHTGLRRTA
GWNVPIGTLRPFLNWTGPPEPIEAAVARFFSASCVPGADKGQFPNLCRLCAGTGENKCAFSS
QEPYFSYSGAFKCLRDGAGDVAFIEESTVFEDLSDEAERDEYELLCPDNTRKPVDKFKDCHL
ARVPSHAVVARSVNGKEDAIWNLLRQAQEKFGKDKSPKFQLFGSPSGQKDLLFKDSAIGFSR
VPPRIDSGLYLGSGYFTAIQNLRK
111110000000000000000000000000000000000000000000000000000000
000000000000000000000000000000000000000000000000000000000000
000000000000000000000000000000000000000000000000000000000000
000000000000000000000000000000000000000000000000000000000000
000000000000000000000000000000000000000000000000000000000000
000011111111111111111111
>DM_train655
MSEFLTPERTVYDSGVQFLRPKSLDEFIGQENVKKKLSLALEAAKMRGEVLDHVLLAGPPGL
GKTTLAHIIASELQTNIHVTSGPVLVKQGDMAAILTSLERGDVLFIDEIHRLNKAVEELLYS
AIEDFQIDIMIGKGPSAKSIRIDIQPFTLVGATTRSGLLSSPLRSRFGIILELDFYTVKELK
EIIKRAASLMDVEIEDAAAEMIAKRSRGTPRIAIRLTKRVRDMLTVVKADRINTDIVLKTME
VLNIDDEGLDEFDRKILKTIIEIYRGGPVGLNALAASLGVEADTLSEVYEPYLLQAGFLART
PRGRIVTEKAYKHLKYEVPENRLF
111111111111111100000000000000000000000000000000000000000000
000000000000000000000000000000000000000000000000000000000000
000000011111111111111100000000000000000000000000000000000000
000000000000000000000000000000000000000000000000000000000000
000000000000000000000000000000000000000000000000000000000000
000000000000000000011111
>DM_train656
MVYVSNKYLTMSEMKVNAQYILNYLSSNGWTKQAICGMLGNMQSESTINPGLWQNLDEGNTS
LGFGLVQWTPASNYINWANSQGLPYKNMDSELKRIIWEVNNNAQWINLRDMTFKEYIKSTKT
PRELAMIFLASYERPANPNQPERGDQAEYWYKNLSGGGGGGLQLAQFPMDIINISQGENGSF
SHKGTLCIDFVGKTEKYPYYAPCDCTCVWRGDASAYLAWTSDKEVMCADGSVRYITWVNVHE

SPLPFDVGKKLKKGDLMGHTGIGGNVTGDHWHFNVIDGKEYQGWTKKPDSCLAGTELHIYDV
FAVNNVEIINGNGYDWKTSDWQDG
00000000000000000000000000000000000000000000000000000000000000
00000000000000000000000000000000000000000000000000000000000000
00000000000000000000000000000000000111111000000000000000000000
00000000000000000000000000000000000000000000000000000000000000
00000000000000000000000111100000000000000000000000000000000000
000000000000000000000000
>DM_train657
NHKSKKRIREAKRSARPELKDSLDWTRHNYYESFSLSPAAVADNVERADALQLSVEEFVERY
ERPYKPVVLLNAQEGWSAQEKWTLERLKRKYRNQKFKCGEDNDGYSVKMKMKYYIEYMESTR
DDSPLYIFDSSYGEHPKRRKLLEDYKVPKFFTDDLFQYAGEKRRPPYRWFVMGPPRSGTGIH
IDPLGTSAWNALVQGHKRWCLFPTSTPRELIKVTRDEGGNQQDEAITWFNVIYPRTQLPTWP
PEFKPLEILQKPGETVFVPGGWWHVVLNLDTTIAITQNFASSTNFPVVWHKTVRGRPKLSRK
WYRILKQEHPELAVLADSVDLQES
00000000000000000000000000000000000000000000000000000000000000
00000000000000000000000000000000000000000000000000000000000000
00000000000000000000000000000000000000000000000000000000000000
00000000000000000000000000000000000000000000000000000000000000
00000000000000000000000000000000000000000000000000000000000000
000000000000000000000001
>DM_train658
MKVKVGVNGYGTIGKRVAYAVTKQDDMELIGITKTKPDFEAYRAKELGIPVYAASEEFIPRF
EKEGFEVAGTLNDLLEKVDIIVDATPGGIGAKNKPLYEKAGVKAIFQGGEKADVAEVSFVAQ
ANYEAALGKNYVRVVSCNTTGLVRTLSAIREYADYVYAVMIRRAADPNDTKRGPINAIKPTV
EVPSHHGPDVQTVIPINIETMAFVVPTTLMHVHSVMVELKKPLTKDDVIDIFENTTRVLLFE
KEKGFDSTAQIIEFARDLHREWNNLYEIAVWKESINIKGNRLFYIQAVHQESDVIPENIDAI
RAMFELADKWDSIKKTNKSLGILK
10000000000000000000000000000000000000000000000000000000000000
00000000000000000000000000000000000000000000000000000000000000
00000000000000000000000000000000000000000000000000000000000000
00000000000000000000000000000000000000000000000000000000000000
00000000000000000000000000000000000000000000000000000000000000
000000000000000000000001
>DM_train659
ALPDQIDVKVKNLTPEDTIYDRTRQVFYQSNLYKGRIEVYNPKTQSHFNVVIDGASSNGDGE
QQMSGLSLLTHDNSKRLFAVMKNAKSFNFADQSSHGASSFHSFNLPLSENSKPVWSVNFEKV
QDEFEKKAGKRPFGVVQSAQDRDGNSYVAFALGMPAIARVSADGKTVSTFAWESGNGGQRPG
YSGITFDPHSNKLIAFGGPRALTAFDVSKPYAWPEPVKINGDFGTLSGTEKIVTVPVGNESV
LVGARAPYAISFRSWDNWKSANIKKTKRSELQNSGFTAVADYYQGSEQGLYAVSAFFDNGAH
GGRSDYPLYKLDNSIQNFHHHHHH
00000000000000000000000000000000000000000000000000000000000000
00000000000000000000000000000000000000000000000000000000000000
00000000000000000000000000000000000000000000000000000000000000

000000000000000000000000000000000000000000000000000000000000
000000000000000000000000000000000000000000000000000000000000
000000000000000000000100
>DM_train660
MKVRIATYASHSALQILKGAKDEGFETIAFGSSKVKPLYTKYFPVADYFIEEKYPEEELLNL
NAVVVPTGSFVAHLGIELVENMKVPYFGNKRVLRWESDRNLERKWLKKAGIRVPEVYEDPDD
IEKPVIVKPHGAKGGKGYFLAKDPEDFWRKAEKFLGIKRKEDLKNIQIQEYVLGVPVYPHYF
YSKVREELELMSIDRRYESNVDAIGRIPAKDQLEFDMDITYTVIGNIPIVLRESLLMDVIEA
GERVVKAAEELMGGLWGPFCLEGVFTPDLEFVVFEISARIVAGTNIFVNGSPYTWLRYDRPV
STGRRIAMEIREAIENDMLEKVLT
000000000000000000000000000000000000000000000000000000000000
000000000000000000000000000000000000000000000000000000000000
000000000111110000000000000000000000000000000000000000000000
000000000000000000000000000000000000000000000000000000000000
000000000000000000000000000000000000000000000000000000000000
000000000000000000000000
>DM_train661
ETVLADRFERLVELVTMMGDQGELPIAMALANVVPCSQWDELARVLVTLFDSRHLLYQLLWN
MFSKEVELADSMQTLFRGNSLASKIMTFCFKVYGATYLQKLLDPLLRIVITSSDWQHVSFEV
DPTRLEPSESLEENQRNLLQMTEKFFHAIISSSSEFPPQLRSVCHCLYQVVSQRFPQNSIGA
VGSAMFLRFINPAIVSPYEAGILDKKPPPIIERGLKLMSKILQSIANHVLFTKEEHMRPFND
FVKSNFDAARRFFLDIASDCPTSDAVNHSLSFISDGNVLALHRLLWNNQEKIGQYLSSNRDH
KAVGRRPFDKMATLLAYLGPPEH
111111110000000000000000000000000000000000000000000000000000
000000000000000000000000000000000000000000001111111111111111
111111111000000000000000000000000000000000000000000000000000
000000000000000000001111111100000000000000000000000000000000
000000000000000000111111111111111111110000000000000000001111
11111000000000000000000
>DM_train664
MSEPSEKKQKVATSSLEQLKKAGTHVVADSGDFEAISKYEPQDSTTNPSLILAASKLEKYAR
FIDAAVEYGRKHGKTDHEKIENAMDKILVEFGTQILKVVPGRVSTEVDARLSFDKKATVKKA
LHIIKLYKDAGVPKERVLIKIASTWEGIQAARELEVKHGIHCNMTLLFSFTQAVACAEANVT
LISPFVGRIMDFYKALSGKDYTAETDPGVLSVKKIYSYYKRHGYATEVMAASFRNLDELKAL
AGIDNMTLPLNLLEQLYESTDPIENKLNSESAKEEGVEKVSFINDEPHFRYVLNEDQMATEK
LSDGIRKFSADIEALYKLVEEKM
111111111110000000000000000000000000000000000000000000000000
000000000000000000000000000000000000000000000000000000000000
000000000000000000000000000000000000000000000000000000000000
000000000000000111110000000000000000000000000000000000000000
000000000000000000000000000000000000000000000000000000000000
00000000000000000000000
>DM_train665
GMATYNYPEFGAGLWHFANYIDRYAVDGYGPALSTIDQINAAKEVGELSYVDLPYPFTPGVT

LSEVKDALKDAGLKAIGITPEIYLQKWSRGAFTNPDPAARAAAFELMHESAGIVRELGANYV
KVWPGQDGWDYPFQVSHKNLWKLAVDGMRDLAGANPDVKFAIEYKPREPRVKMTWDSAARTL
LGIEDIGLDNVGVLLDFGHALYGGESPADSAQLIIDRGRLFGMDVNDNLRGWDDDLVVGTVH
MTEIFEFFYVLKINNWQGVWQLDQFPFRENHVEAAQLSIRFLKHIYRALDKLDIPALQAAQE
AQNPLQAQRIVQDALLSSITVSE
110000000000000000000000000000000000000000000000000000000000
000000000000000000000000000000000000000000000000000000000000
000000000000000000000000000000000000000000000000000000000000
000000000000000000000000000000000000000000000000000000000000
000000000000000000000000000000000000000000000000000000000000
00000000000000000000001
>DM_train666
MTHSPSFLQHALSSSDTRAEWPLPGGLAARWLAPGCVELNGDARGADSVLLSCGVHGNETAP
IEVVDGMLTDIAAGQLALNCRLLVMFANLDAIRQGVRYGNYDMNRLFNGAHARHPELPESVR
AAELETLAAEFFAGARARKLHYDLHTAIRGSVFEKFAIYPFLHDGRTHKREQLAWLQRCGIE
AVLLHTQPANTFSYFTSQYCEADAFTLELGKARPFGQNDLSRFSGIDGALRGLLSNPQANVP
DLDEDKLPLFRAKYDLVKHSEAFKLNLADSVENFTLLPDGMLIAEDGAVRYQATGGEERILF
PNPAVKPGLRAGIVVEPARLPSR
111000000000000000000000000000000000000000000000000000000000
000000000000000000000000000000000000000000000000000000000000
000000000000000000000000000000000000000000000000000000000000
000000000000000000000000000000000000000000000000000000000000
000000000000000001111111000000000000000000111110000000000000
00011111110000000000011
>DM_train667
SLSQTTNIKEQTDTGLAPPAPAITTNSAAIEEQANSSIRKLYHTLNTMPNTSMADRISQISA
YFKGTKYILGSLGEGPNARYDQFPRYRVDGFDCDTYVNTVLSLALANSLESFQECLKHTRYK
NGKRSYINRNHFTSIDWNNYNQKRGLLKDITFSIRNEKKQPVALYANALINKPQWYNHKTID
TIRLQKQDKNEQEKRLVELKAKGKTLETSLSNVPYIPFTALFSENKPNLHLFSQIPNGAVIE
IIRPNWDLRQQIGTELDISHLGFAIWINNELFFRQASSQYGKVVDVSLIDYLDKARSSPTIK
GINIQVVLPEKPVSNGCQLFDIR
111111111111111111111111100000000000000000000111000000000000
000000000000000000000000000000000000000000000000000000000000
000000000000000000000000000000000000000000000000000000000000
000000000000000000000000000000000000000000000000000000000000
000000000000000000000000000000000000000000000000000000000000
00000000000001110000111
>DM_train668
GLVPRGSHMTGRMLTLDGNPAANWLNNARTKWSASRADVVLSYQQNNGGWPKNLDYNSVGNG
GGGNESGTIANGATITEMVFLAEVYKSGGNTKYRDAVRKAANFLVNSQYSTGALPQFYPLKG
GYSDHATFNDNGMAYALTVLDFAANKRAPFDTDVFSDNDRTRFKTAVTKGTDYILKAQWKQN
GVLTVWCAQHGALDYQPKKARAYELESLSGSESVGVLAFLMTQPQTAEIEQAVRAGVAWFNS
PRTYLEGYTYDSSLAATNPIVPRAGSKMWYRFYDLNTNRGFFSDRDGSKFYDITQMSLERRT
GYSWGGNYGTSIINFAQKVGYL

111111111110000000000000000000000000000000000000000000000000
000000000000000000000000000000000000000000000000000000000000
000000000000000000000000000000000000000000000000000000000000
000000000000000000000000000000000000000000000000000000000000
000000000000000000000000000000000000000000000000000000000000
0000000000000000000000
>DM_train669
MHHHHHHGSTSLYKKAGSETLYIQGMAEFPASLLILNGKSTDNLPLREAIMLLREEGMTIHV
RVTWEKGDAARYVEEARKFGVATVIAGGGDGTINEVSTALIQCEGDDIPALGILPLGTANDF
ATSVGIPEALDKALKLAIAGDAIAIDMAQVNKQTCFINMATGGFGTRITTETPEKLKAALGS
VSYIIHGLMRMDTLQPDRCEIRGENFHWQGDALVIGIGNGRQAGGGQQLCPNALINDGLLQL
RIFTGDEILPALVSTLKSDEDNPNIIEGASSWFDIQAPHDITFNLDGEPLSGQNFHIEILPA
ALRCRLPPDCPLLRSTHHHHHH
111111111111111111111111111110000000000000000000000000000000
000000000000000000000000000000000000000000000000000000000000
000000000000000000000000000000000000000000000011111111111111
000000000000000000000000000000000000000000000000000000000000
000011111111111111111100000000000000000000000000000000000000
0000000000000000111111
>DM_train670
QSSVTLFGIVDTNVAYVNKDAAGDSRYGLGTSGASTSRLGLRGTEDLGGGLKAGFWLEGEIF
GDDGNASGFNFKRRSTVSLSGNFGEVRLGRDLVPTSQKLTSYDLFSATGIGPFMGFRNWAAG
QGADDNGIRANNLISYYTPNFGGFNAGFGYAFDEKQTIGTADSVGRYIGGYVAYDNGPLSAS
LGLAQQKTAVGGLATDRDEITLGASYNFGVAKLSGLLQQTKFKRDIGGDIKTNSYMLGASAP
VGGVGEVKLQYALYDQKAIDSKAHQITLGYVHNLSKRTALYGNLAFLKNKDASTLGLQAKGV
YAGGVQAGESQTGVQVGIRHAF
110000000000000000000000000000000000000000000000000000000000
000000000000000000000000000000000000000000000000000000000000
000000000000000000000000000000000000000000000000000000000000
000000000000000000000000000000000000000000000000000000000000
000000000000000000000000000000000000000000000000000000000000
0000000000000000000000
>DM_train671
GMSEAPHLTFDLDTPGVSTGHLVVPKGADCEALSLPVFSCNRGEGPSLLITGGNHGNELQGP
ILARRLVKWLPEAQRCGRIIIVPEINPLAVQAWTRNTPIDGKNLNRVFPGRSDGSVSERIAD
AISRLLLPVVDTVLDLHSFGPTWDCAPSIISHPIADIDQMTKTVSISKAFKLPVTLLWEHNE
TDGMFDTLVHRQGKTFICTEFGGGVVSAEALTIYEAGVRNGLIALGLVKGKAEYPTFRQQKT
GQTLETTSSDQLKSPSPGIFEPRCSVMDEVEQGDVVGVLHPMGSLSAASIDIRAQSKSTVFA
IRSAMYVQGNEEVAILARPLAR
100000000000000000000000011100000000000000000000000000000000
000000000000000000000000000011111111111111111111111100000000
000000000000000000000000000000011111000000000000000000000111
111000000000000000000000001100000000000000000000000000000000
000000000000000000000000000000000000000000000000000000000000

000000000000000000001
>DM_train672
MGSVSKANVPKIDVSPLFGDDQAAKMRVAQQIDAASRDTGFFYAVNHGINVQRLSQKTKEFH
MSITPEEKWDLAIRAYNKEHQDQVRAGYYLSIPGKKAVESFCYLNPNFTPDHPRIQAKTPTH
EVNVWPDETKHPGFQDFAEQYYWDVFGLSSALLKGYALALGKEENFFARHFKPDDTLASVVL
IRYPYLDPYPEAAIKTAADGTKLSFEWHEDVSLITVLYQSNVQNLQVETAAGYQDIEADDTG
YLINCGSYMAHLTNNYYKAPIHRVKWVNAERQSLPFFVNLGYDSVIDPFDPREPNGKSDREP
LSYGDYLQNGLVSLINKNGQT
00000000000000000000000000000000000000000000000000000000000000
00000000000000000000000000000000000000000000000000000000000000
00000000000000000000000000000000000000000000000000000000000000
00000000000000000000000000000000000000000000000000000000000000
00000000000000000000000000000000000000000000000000000000000000
000000000000011111111
>DM_train673
MLKQPIQAQQLIELLKVHYGIDIHTAQFIQGGADTNAFAYQADSESKSYFIKLKYGYHDEIN
LSIIRLLHDSGIKEIIFPIHTLEAKLFQQLKHFKIIAYPFIHAPNGFTQNLTGKQWKQLGKV
LRQIHETSVPISIQQQLRKEIYSPKWREIVRSFYNQIEFDNSDDKLTAAFKSFFNQNSAAIH
RLVDTSEKLSKKIQPDLDKYVLCHSDIHAGNVLVGNEESIYIIDWDEPMLAPKERDLMFIGG
GVGNVWNKPHEIQYFYEGYGEINVDKTILSYYRHERIVEDIAVYGQDLLSRNQNNQSRLESF
KYFKEMFDPNNVVEIAFATEQ
11000000000000000000000000000000000000000000000000000000000000
00000000000000000000000000000000000000000000000000000000000000
00000000000000000000000000000000000000000000000000000000000000
00000000000000000000000000000000000000000000000000000000000000
00000000000000000000000000000000000000000000000000000000000000
000000000000000000001
>DM_train674
MHNHNNRLVITPGEPAGVGPDLAITLAQQDWPVELVVCADPALLLARASQLNLPLQLREYQA
DQPAIAQQAGSLTILPVKTAVNVVPGKLDVGNSHYVVETLAKACDGAISGEFAALVTGPVQK
SIINDAGIPFIGHTEFFADRSHCQRVVMMLATEELRVALATTHLPLLAVPGAITQASLHEVI
TILDNDLKTKFGITQPQIYVCGLNPHAGEGGHMGHEEIDTIIPALNTLRQQGINLIGPLPAD
TLFQPKYLQHADAVLAMYHDQGLPVLKYQGFGRAVNITLGLPFIRTSVDHGTALELAATGTA
DVGSFITALNLAIKMINNSNE
00000000000000000000000000000000000000000000000000000000000000
00000000000000000000000000000000000000000000000000000000000000
00000000000000000000000000000000000000000000000000000000000000
00000000000000000000000000000000000000000000000000000000000000
00000000000000000000000000000000000000000000000000000000000000
000000000000000000011
>DM_train675
GMQKPHLLITSAGRRAKLVEYFVKEFKTGRVSTADCSPLASALYMADQHYIVPKIDEVEYID
HLLTLCQDEGVTALLTLIDPELGLLAQATERFQAIGVTVIVSPYAACELCFDKYTMYEYCLR
QGIAHARTYATMASFEEALAAGEVQLPVFVKPRNGSASIEVRRVETVEEVEQLFSKNTDLIV

QELLVGQELGVDAYVDLISGKVTSIFIKEKLTMRAGETDKSRSVLRDDVFELVEHVLDGSGL
VGPLDFDLFDVAGTLYLSEINPRFGGGYPHAYECGVNFPAQLYRNLMHEINVPQIGQYLDDI
YMLKHDTVTLISAAELQKIKR
000000000000000000000000000000000000000000000000000000000000
000000000000000000000000000000000000000000000000000000000000
000000000000000000000000000000000011111111111111111111110000
000000000000000000000000000000000000000000000000000000000000
000000000000000000000000000000000000000000000000000000000000
000000000000000000000
>DM_train676
GMELKSAEEKSLYYRNKIKEVIDEVGKVVVGQKYMINRLLIGICTGGHILLEGVPGLAKTLS
VNTLAKTMDLDFHRIQFTPDLLPSDLIGTMIYNQHKGNFEVKKGPVFSNFILADEVNRSPAK
VQSALLECMQEKQVTIGDTTYPLDNPFLVLATQNPVEQEGTYPLPEAQVDRFMMKIHLTYLD
KESELEVMRRVSNMNFNYQVQKIVSKNDVLEIRNEINKVTISESLEKYIIELVFATRFPAEY
GLEAEASYILYGASTRAAINLNRVAKAMAFFNNRDYVLPEDIKEVAYDILNHRIILNYEAEA
EGISTRQIIETILRKVNITKA
000000000000000000000000000000000000000000000000000000000000
000000000000000000000000000000000000000000000000000000000000
000000000000000000000000000000000000000000000000000000000000
000000000000000000000000000000000000000000000000000000000000
000000000000000000000000000000000000000000000000000000000000
000000000000000000001
>DM_train677
MVPPTINTGKNITTERAVLTLNGLQIKLHKVVGESRDDIVAKMKDLAMDDHKFPRLPGPNPV
SIERKDFEKLKQNKYVVSEKTDGIRFMMFFTRVFGFKVCTIIDRAMTVYLLPFKNIPRVLFQ
GSIFDGELCVDIVEKKFAFVLFDAVVVSGVTVSQMDLASRFFAMKRSLKEFKNVPEDPAILR
YKEWIPLEHPTIIKDHLKKANAIYHTDGLIIMSVDEPVIYGRNFNLFKLKPGTHHTIDFIIM
SEDGTIGIFDPNLRKNVPVGKLDGYYNKGSIVECGFADGTWKYIQGRSDKNQANDRLTYEKT
LLNIEENITIDELLDLFKWE
111111111100000000000000000000000000000000000000000000000000
000000000000000000000000000000000000000000000000000000000000
000000000000000000000000000000000000000000000000000000000000
000000000000000000000000000000000000000000000000000000000000
000000000000000000000000000000000000000000000000000000000000
00000000000000000111
>DM_train678
MKTAYIAKQRQISFVKSHFSRQLEERLGLIEVQAPILSRVGDGTQDNLSGAEKAVQVKVKAL
PDAQFEVVHSLAKWKRQTLGQHDFSAGEGLYTHMKALRPDEDRLSPLHSVYVDQWDWERVMG
DGERQFSTLKSTVEAIWAGIKATEAAVSEEFGLAPFLPDQIHFVHSQELLSRYPDLDAKGRE
RAIAKDLGAVFLVGIGGKLSDGHRHDVRAPDYDDWSTPSELGHAGLNGDILVWNPVLEDAFE
LSSMGIRVDADTLKHQLALTGDEDRLELEWHQALLRGEMPQTIGGGIGQSRLTMLLLQLPHI
GQVQAGVWPAAVRESVPSLL
111000000000000000000000000000000000000000000000000000000000
000000000000000000000000000000000000000000000000000000000000

000000000000000000000000000000000000000000000000000000000000
000000000000000000000000000000000000000000000000000000000000
000000000000000000000000000000000000000000000000000000000000
00000000000000000000
>DM_train679
MQPQSVLHSGYFHPLLRAWQTATTTLNASNLIYPIFVTDVPDDIQPITSLPGVARYGVKRLE
EMLRPLVEEGLRCVLIFGVPSRVPKDERGSAADSEESPAIEAIHLLRKTFPNLLVACDVCLC
PYTSHGHCGLLSENGAFRAEESRQRLAEVALAYAKAGCQVVAPSDMMDGRVEAIKEALMAHG
LGNRVSVMSYSAKFASCFYGPFRDAAKSSPAFGDRRCYQLPPGARGLALRAVDRDVREGADM
LMVKPGMPYLDIVREVKDKHPDLPLAVYHVSGEFAMLWHGAQAGAFDLKAAVLEAMTAFRRA
GADIIITYYTPQLLQWLKEE
000000000000000000000000000000000000000000000000000000000000
000000000000000000000000001111100000000000000000000000000000
001111111111111111000000000000000000000000000000000000000000
000000000000000000000000001111111100000000000000000000000000
000000000000000000000000000000000000000000000000000000000000
00000000000000000000
>DM_train680
MLALGIEGTAHTLGIGIVSEDKVLANVFDTLTTEKGGIHPKEAAEHHARLMKPLLRKALSEA
GVSLDDIDVIAFSQGPGLGPALRVVATAARALAVKYRKPIVGVNHCIAHVEITKMFGVKDPV
GLYVSGGNTQVLALEGGRYRVFGETLDIGIGNAIDVFARELGLGFPGGPKVEKLAEKGEKYI
ELPYAVKGMDLSFSGLLTEAIRKYRSGKYRVEDLAYSFQETAFAALVEVTERAVAHTEKDEV
VLVGGVAANNRLREMLRIMTEDRGIKFFVPPYDLCRDNGAMIAYTGLRMYKAGISFRLEETI
VKQKFRTDEVEIVWHHHHHH
000000000000000000000000000000000000000000000000000000000000
000000000000000000000000000000000000000000000000000000000000
000000000000000000000000000000000000000000000000000000000000
000000000000000000000000000000000000000000000000000000000000
000000000000000000000000000000000000000000000000000000000000
00000000000000011111
>DM_train681
MIKISYSPYTLKPVQSLNAATAATAREGVLLKVEWNDGLYGFADLHPWPELGDLSLEEQLSD
LRMGRMTTQIEQSIWLARRDALLRKEKKHVFDGGEKIKNNYLLSHFQDLKPGFLDGLKNEGY
NTVKVKMGRDLQKEADMLTHIAASGMRMRLDFNALGSWQTFEKFMVNLPLTVRPLIEYVEDP
FPFDFHAWGEARKLAKIALDNQYDKVPWGKIASAPFDVIVIKPAKTDVDKAVAQCQKWNLKL
AVTSYMDHPVGVVHAVGVAMELKDKYGDMILESGCLTHRLYQMDSFAAELSTQGPYLLKNKG
TGVGFDKLLEALTWYQLKVR
110000000000001111111111000000000000000000000000000000000000
000000000000000000000000000000000000000000000000000000000000
000000000000000000000000000000000000000000000000000000000000
000000000000000000000000000000000000000000000000000000000000
000000000000000000000000000000000000000000000000000000000000
00000000000000000011
>DM_train682

MTTSDLVAGELAGDGLRDTRPGDTWLADRSWNRPGWTVAELEAAKAGRTISVVLPALDEEDT
IGSVIDSISPLVDGLVDELIVLDSGSTDDTEIRAVAAGARVVSREQALPEVPIRPGKGEALW
RSLAASRGDIVVFVDSDLINPHPMFVPWLVGPLLTGDGVHLVKSFYRRPLNVGDAGGGAGAT
GGGRVTELVARPLLAALRPELGCILQPLGGEYAATRELLTSVPFAPGYGVEIGLLVDTFDRL
GLDAIAQVNLGVREHRNRPLAELGAMSRQVIATLLSRCGIPDSGVGLTQFVADGPEGQSYTQ
HTWPVSLADRPPMQAIRPR
111111111111110000000000000000000000000000000000000000000000
000000000000000000000000000000000000000000000000000000000000
000000000000000000000000000000000000000000000001111111111111
110000000000000000000000000000000000000000000000000000000000
000000000000000000000000000000000000000000000000000000000000
0000000000000000000
>DM_train683
TGLGELKISDTICAQNAVEALPALSVDEPTVTMTFQVNTSPFAGKEGKFVTSRNILERLEKE
LVHNVALRVEQTDDPDKFRVSGRGELHLSILIENMRREGFELAVSRPEVIIKEEDGQLMEPF
ETVTIDVMEEHQGGIMENIGLRKGELKDMAPDGKGRVRMDFIMPSRGLIGFQTEFMTLTSGS
GLLYHTFDHYGPHKGGNIGQRVNGVLIANAAGKALTNALFNLQERGRLFIGHGVEVYEGMVI
GIHSRDNDLTVNALKGKQLTNVRASGTDDAQVLTPPIVMTLEQALEFIDDDELVEVTPESIR
IRKKFLTESDRKRASRSAK
111111111111111111111111100000000000000000000000000000000000
000000000000000000000000000000000000000000000000000000000000
000000000000000000000000000000000000000000000000000000000000
000000000000000000000000000000000000000000000000000000000000
000000000000000111111111111110000000000000000000000000000000
0000000000000001111
>DM_train684
GSLIGLMKDAFQPHHHHHHLSPHPPGTVDKKMVEKCWKLMDKVVRLCQNPKLALKNSPPYI
LDLLPDTYQHLRTILSRYEGKMETLGENEYFRVFMENLMKKTKQTISLFKEGKERMYEENSQ
PRRNLTKLSLIFSHMLAELKGIFPSGLFQGDTFRITKADAAEFWRKAFGEKTIVPWKSFRQA
LHEVHPISSGLEAMALKSTIDLTCNDYISVFEFDIFTRLFQPWSSLLRNWNSLAVTHPGYMA
FLTYDEVKARLQKFIHKPGSYIFRLSCTRLGQWAIGYVTADGNILQTIPHNKPLFQALIDGF
REGFYLFPDGRNQNPDLTG
111111111111111111111111000000000000000000000000000000000000
000000000000000000000000000000000000000000000000000000000000
000000000000000000000000000000000000000000000000000000000000
000000000000000000000000000000000000000000000000000000000000
000000000000000000000000000000000000000000000000000000000000
0000000000000000000
>DM_train685
APPPVPKTPAGPLTLSGQGSFFVGGRDVTSETLSLAPKYDAHGTVTVDQMYVRYQIPQRAKR
YPITLIHGCCLTGMTWETTPDGRMGWDEYFLRKGYSTYVIDQSGRGRSATDISAINAVKLGK
APASSLPDLFAAGHEAAWAIFRFGPRYPDAFKDTQFPVQAQAELWQQMVPDWLGSMPTPNPT
VANLSKLAIKLDGTVLLSHSQSGIYPFQTAAMNPKGITAIVSVEPGECPKPEDVKPLTSIPV
LVVFGDHIEEFPRWAPRLKACHAFIDALNAAGGKGQLMSLPALGVHGNSHMMMQDRNNLQVA

DLILDWIGRNTAKPAHGR
111100000000000000000000000000000000000000000000000000000000
000000000000000000000000000000000000000000000000000000000000
000000000000000000000000000000000000000000000000000000000000
000000000000000000000000000000000000000000000000000000000000
000000000000000000000000000000000000000000000000000000000000
000000000001111111
>DM_train687
MDTWMVLLGLQILLLPLLAHCTAPASTLRFVAVGDWGGVPNAPFHTAREMANAKEIARTVQI
MGADFIMSLGDNFYFTGVHDANDKRFQETFEDVFSDRALRNIPWYVLAGNHDHLGNVSAQIA
YSKISKRWNFPSPYYRLRFKVPRSNITVAIFMLDTVMLCGNSDDFVSQQPEMPRDLGVARTQ
LSWLKKQLAAAKEDYVLVAGHYPIWSIAEHGPTRCLVKNLRPLLAAYGVTAYLCGHDHNLQY
LQDENGVGYVLSGAGNFMDPSVRHQRKVPNGYLRFHYGSEDSLGGFTYVEIGSKEMSITYVE
ASGKSLFKTSLPRRPRP
111111111111111111111111100000000000000000000000000000000000
000000000000000000000000000000000000000000000000000000000000
000000000000000000000000000000000000000000000000000000000000
000000000000000000000000000000000000000000000000000000000000
000000000000000000000000000000000000000000000000000000000000
00000000000000000
>DM_train689
MIEIKDKQLTGLRFIDLFAGLGGFRLALESCGAECVYSNEWDKYAQEVYEMNFGEKPEGDIT
QVNEKTIPDHDILCAGFPCQAFSISGKQKGFEDSRGTLFFDIARIVREKKPKVVFMENVKNF
ASHDNGNTLEVVKNTMNELDYSFHAKVLNALDYGIPQKRERIYMICFRNDLNIQNFQFPKPF
ELNTFVKDLLLPDSEVEHLVIDRKDLVMTNQEIEQTTPKTVRLGIVGKGGQGERIYSTRGIA
ITLSAYGGGIFAKTGGYLVNGKTRKLHPRECARVMGYPDSYKVHPSTSQAYKQFGNSVVINV
LQYIAYNIGSSLNFKPY
000000000000000000000000000000000000000000000000000000000000
000000000000000000000000000000000000000000000000000000000000
000000000000000000000000000000000000000000000000000000000000
000000000000000000000000000000000000000000000000000000000000
000000000000000000000000000000000000000000000000001110000000
00000000000000000
>DM_train690
HLIPAQQRTVEHLDAMQWINGDGYLHNVFVRWFNGDVIRPKTWFWQDVKTRKILGWRCDVSE
NIDSIRLSFMDVVTRYGIPEDFHITIDNTRGAANKWLTGGAPNRYRFKVKEDDPKGLFLLMG
AKMHWTSVVAGKGWGQAKPVERAFGVGGLEEYVDKHPALAGAYTGPNPQAKPDNYGDRAVDA
ELFLKTLAEGVAMFNARTGRETEMCGGKLSFDDVFEREYARTIVRKPTEEQKRMLLLPAEAV
NVSRKGEFTLKVGGSLKGAKNVYYNMALMNAGVKKVVVRFDPQQLHSTVYCYTLDGRFICEA
ECLAPVAFNDAAAGREY
111111111100000000000000000000000000000000000000000000000000
000000000000000000000000000000000000000000000000000000000000
000000000000000000000000000000000000000000001111111100000000
000000000000000000000000000000000000000000000000000000000000

000000000000000000000000000000000000000000000000000000000000
0001111111111111
>DM_train691
MENEKILNPIIIQRADPMIYKHNDGYYYFTASVPEYDRIEVRKAKTIEGLRNAEPVDVWRRH
ESGEMSNLIWAPEIHFINGAWYIYFAAAPDKNIEDDTFNHRMFVIQNENENPFTGNWVEKGR
IKTAWESFSLDATIFEHNEKLYYVWAQQDINIKGHSNIYIAEMENPWTLKTKPVMLTKPELE
WEIKGFWVNEGPAVLKKNGKIFITYSASATDVNYCIGMLTAEENSNLLDKNSWTKSQTPVFK
TSMENHQYGPGHNSFTVSEDGKHDVIVYHARNYTEIKGDPLYDPNRHTRAQIINWREDGTPD
FGVPEVDSLEIETPVKA
111110000000000000000000000000000000000000000000000000000000
000000000000000000000000000000000000000000000000000000000000
000000000000000000000000000000000000000000000000000000000000
000000000000000000000000000000000000000000000000000000000000
000000000000000000000000000000000000000000000000000000000000
00000000011111111
>DM_train692
LDGVLVPESGILVSVGQDVDSVNDYASALGTIPAGVTNYVGIVNLDGLNSDADAGAGRNNIA
ELANAYPTSALVVGVSMNGEVDAVASGRYNANIDTLLNTLAGYDRPVYLRWAYEVDGPWNGH
SPSGIVTSFQYVHDRIIALGHQAKISLVWQVASYCPTPGGQLDQWWPGSEYVDWVGLSYFAP
QDCNWDRVNEAAQFARSKGKPLFLNESTPQRYQVADLTYSADPAKGTNRQSKTSQQLWDEWF
APYFQFMSDNSDIVKGFTYINADWDSQWRWAAPYNEGYWGDSRVQANALIKSNWQQEIAKGQ
YINHSETLFETLGYGST
000000000000000000000000000000000000000000000000000000000000
000000000000000000000000000000000000000000000000000000000000
000000000000000000000000000000000000000000000000000000000000
000000000000000000000000000000000000000000000000000000000000
000000000000000000000000000000000000000000000000000000000000
00000000000000111
>DM_train693
MRGSHHHHHHTDPINKDIYQALQQLIPNEKIKVDEPLKRYTYTKTGGNADFYITPTKNEEVQ
AVVKYAYQNEIPVTYLGNGSNIIIREGGIRGIVISLLSLDHIEVSDDAIIAGSGAAIIDVSR
VARDYALTGLEFACGIPGSIGGAVYMNAGAYGGEVKDCIDYALCVNEQGSLIKLTTKELELD
YRNSIIQKEHLVVLEAAFTLAPGKMTEIQAKMDDLTERRESKQPLEYPSCGSVFQRPPGHFA
GKLIQDSNLQGHRIGGVEVSTKHAGFMVNVDNGTATDYENLIHYVQKTVKEKFGIELNREVR
IIGEHPKESLQPSLIS
111111111111110000000000000000000000000000000000000000000000
000000000000000000000000000000000000000000000000000000000000
000000000000000000000000000000000000000000000000000000000000
000000000000000000000000000000000000000000000000000000000000
000000000000000000000000000000000000000000000000000000000000
0000000111111111
>DM_train694
MSATSTATSTSASQLHLNSTPVTHCLSDIVKKEDWSDFKFAPIRESTVSRAMTSRYFKDLDK
FAVSDVIIVGAGSSGLSAAYVIAKNRPDLKVCIIESSVAPGGGSWLGGQLFSAMVMRKPAHL

FLQELEIPYEDEGDYVVVKHAALFISTVLSKVLQLPNVKLFNATCVEDLVTRPPTEKGEVTV
AGVVTNWTLVTQAHGTQCAMDPNVIELAGYKNDGTRDLSQKHGVILSTTGHDGPFGAFCAKR
IVDIDQNQKLGGMKGLDMNHAEHDVVIHSGAYAGVDNMYFAGMEVAELDGLNRMGPTFGAMA
LSGVHAAEQILKHFAA
111111111111111000000000000000000000000000000000000000000000
000000000000000000000000000000000000000000000000000000000000
000000000000000000000000000000000000000000000000000001111000
000000000000000000000000000000000000000000000000000000000000
000000000000000000000000000000000000000000000000000000000000
0000000000000000
>DM_train695
MGSSSSGRENLYFQGHMFFTDWEGPWILTDFALELCMAVFNNARFFSNLSEYDDYLAYEVRR
EGYEAGYTLKLLTPFLAAAGVKNRDVERIAELSAKFVPDAEKAMATLQERWTPVVISTSYTQ
YLRRTASMIGVRGELHGTEVDFDSIAVPEGLREELLSIIDVIASLSGEELFRKLDELFSRSE
VRKIVESVKAVGAGEKAKIMRGYCESKGIDFPVVVGDSISDYKMFEAARGLGGVAIAFNGNE
YALKHADVVIISPTAMSEAKVIELFMERKERAFEVLSAVSIPETEIYIMENSDFGEVLEKSK
RMRVRLRGLAGELGGS
111111111111100000000000000000000000000000000000000000000000
000000000000000000000000000000000000000000000000000000000000
000000000000000000000000000000000000000000000000000000000000
000000000000000000000000000000000000000000000000000000000000
000000000000000000000000000000000000000000000000000000000000
0000000000000000
>DM_train696
MVVVGLDVGYGDTKVIGVDGKRIIFPSRWAVTETESWGIGGKIPVLSTDGGQTKFIYGKYAS
GNNIRVPQGDGRLASKEAFPLIAAALWESGIHNDGSPVDLVIGSGTPLGTFDLEVKAAKEAL
ENKVLTVTGPEGEVRQFNITRLIMRPQGVGAALYLLNQGIIEQQPGYGVVIDVGSRTTDVLT
INLMDMEPVVELSFSLQIGVGDAISALSRKIAKETGFVVPFDLAQEALSHPVMFRQKQVGGP
EVSGPILEDLANRIIENIRLNLRGEVDRVTSLIPVGGGSNLIGDRFEEIAPGTLVKIKPEDL
QFANALGYRDAAERSM
000000000000000000000000000000000000111100000000000000000000
000000000000000000000000000000001110000000000000000000000000
000000000000000000000000000000000000000000000000000000000000
000000000000000000000000000000000000000000000000000000000000
000000000000000000000000000000000000000000000000000000000000
0000000000000001
>DM_train698
MRVLIVKTSSMGDVLHTLPALTDAQQAIPGIKFDWVVEEGFAQIPSWHAAVERVIPVAIRRW
RKAWFSAPIKAERKAFREALQAKNYDAVIDAQGLVKSAALVTRLAHGVKHGMDWQTAREPLA
SLFYNRKHHIAKQQHAVERTRELFAKSLGYSKPQTQGDYAIAQHFLTNLPTDAGEYAVFLHA
TTRDDKHWPEEHWRELIGLLADSGIRIKLPWGAPHEEERAKRLAEGFAYVEVLPKMSLEGVA
RVLAGAKFVVSVDTGLSHLTAALDRPNITVYGPTDPGLIGGYGKNQMVCRAPGNELSQLTAN
AVKQFIEENAEKAAMI
000000000000000000000000000000000000000000000000000000000000

001111111111000000000000000000000000000000000000000000000000
000000000000000000000000000000000000000000000011111000000000
000000000000000000000000000000000000000000000000000000000000
000000000000000000000000000000000001111111000000000000000000
0000000000000000
>DM_train699
GMRKPKITVIGGGTGSPVILKSLREKDVEIAAIVTVADDGGSSGELRKNMQQLTPPGDLRNV
LVAMSDMPKFYEKVFQYRFSEDAGAFAGHPLGNLIIAGLSEMQGSTYNAMQLLSKFFHTTGK
IYPSSDHPLTLHAVFQDGTEVAGESHIVDHRGIIDNVYVTNALNDDTPLASRRVVQTILESD
MIVLGPGSLFTSILPNIVIKEIGRALLETKAEIAYVCNIMTQRGETEHFTDSDHVEVLHRHL
GRPFIDTVLVNIEKVPQEYMNSNRFDEYLVQVEHDFVGLCKQVSRVISSNFLRLENGGAFHD
GDLIVDELMRIIQVKK
100000000000000000000000000000000000111111111111111111000000
000000000000000000000000000000000000000000000000000000000000
000000000000000000000000000000000000000000000000000000000000
000000000000000000000000000000000000000000000000000000000000
000000000000000000000000000000000000000000000000000000000000
0000000000000011
>DM_train700
MILGIDIGGANTKITELHENGEFKVHHLYFPMWKNNDKLAEVLKTYSNDVSHVALVTTAELA
DSYETKKEGVDNILNAAESAFGSNISVFDSNGNFISLESAKTNNMKVSASNWCGTAKWVSKN
IEENCILVDMGSTTTDIIPIVEGKVVAEKTDLERLMNHELLYVGTLRTPISHLGNTISFKGV
DTNVSSEYFAITADISVVLEKVTTEEYTCDTPDGKGTDKRSSLVRISKVLCSDLDQISEIDA
ENIAKNYYELWKELILENVENVAEKYGSKKVVITGLGENILKDALADFEVISVAERYGKDVS
LATPSFAVAELLKNEL
000000001100000000000000000111111111111111111110000000000111
111000000000000000000000000000000000000000000000000000000000
000000000000000000000000000000000000000000000000000000000000
000000000000000000000000000000000000000000000000000000000000
000000000000000000000000000000000000000000000000000000000000
0000000000000000
>DM_train701
SQLVECVPNFSEGKNQEVIDAISRAVAQTPGCVLLDVDSGPSTNRTVYTFVGRPEDVVEGAL
NAARAAYQLIDMSRHHGEHPRMGALDVCPFIPVRGVTMDECVRCAQAFGQRLAEELGVPVYL
YGEAARTAGRQSLPALRAGEYEALPEKLKQAEWAPDFGPSAFVPSWGATVAGARKFLLAFNI
NLLSTREQAHRIALDLREQGRGKDQPGRLKKVQAIGWYLDEKNLAQVSTNLLDFEVTGLHTV
FEETCREAQELSLPVVGSQLVGLVPLKALLDAAAFYCEKENLFLLQDEHRIRLVVNRLGLDS
LAPFKPKERIIEYLV
000000000000000000000000000000000000000000000000000000000000
000000000000000000000000000000000000000000000000000000000000
000000000000000000000000000000000000000000000000000000000000
000000000000000000011111100000000000000000000000000000000000
000000000000000000000000000000000000000000000000000000000000
000000000000000

>DM_train702
LQTLPPVQLAQKYFQIFSEEKDPLWQDPCEDKRHKDIWSKEKTCDRFPKLLIIGPQKTGTTA
LYLFLGMHPDLSSNYPSSETFEEIQFFNGHNYHKGIDWYMEFFPIPSNTTSDFYFEKSANYF
DSEVAPRRAAALLPKAKVLTILINPADRAYSWYQHQRAHDDPVALKYTFHEVITAGSDASSK
LRALQNRCLVPGWYATHIERWLSAYHANQILVLDGKLLRTEPAKVMDMVQKFLGVTNTIDYH
KTLAFDPKKGFWCQLLEGGKTKCLGKSKGRKYPEMDLDSRAFLKDYYRDHNIELSKLLYKMG
QTLPTWLREDLQNTR
11111111111111111111100000001111111111111110000000000000000000
00000000000000000000000000000000000000000000011111000000000000
00000000000000000000000000000000000000000000000000000000000000
00000000000000000000000000000000000000000000000000000000000000
00000000000000000000000000000000000000000000000000000000000000
000000000000111
>DM_train703
GMKEIRIILMGTGNVGLNVLRIIDASNRRRSAFSIKVVGVSDSRSYASGRNLDISSIISNKE
KTGRISDRAFSGPEDLMGEAADLLVDCTPASRDGVREYSLYRMAFESGMNVVTANKSGLANK
WHDIMDSANQNSKYIRYEATVAGGVPLFSVLDYSILPSKVKRFRGIVSSTINYVIRNMANGR
SLRDVVDDAIKKGIAESNPQDDLNGLDAARKSVILVNHIFGTEYTLNDVEYSGVDERSYNAN
DRLVTEVYVDDRRPVAVSRIISLNKDDFLMSIGMDGLGYQIETDSNGTVNVSDIYDGPYETA
GAVVNDILLLSKVQK
11000000000000000000000000000011000000000000000000000000000000
00000000000000000000000000000000000000000000000000000000000000
00000000000000000000000000000000000000000000000000000000000000
00000000000000000000000000000000000000000000000000000000000000
00000000000000000000000000000000000000000000000000000000000000
000000000000011
>DM_train704
MGSDKIHHHHHHRTKDQYYELRDFALGTSVRIVVSSQKINPRTIAEAILEDMKRITYKFSFT
DERSVVKKINDHPNEWVEVDEETYSLIKAACAFAELTDGAFDPTVGRLLELWGFTGNYENLR
VPSREEIEEALKHTGYKNVLFDDKNMRVMVKNGVKIDLGGIAKGYALDRARQIALSFDENAT
GFVEAGGDVRIIGPKFGKYPWVIGVKDPRGDDVIDYIYLKSGAVATSGDYERYFVVDGVRYH
HILDPSTGYPARGVWSVTIIAEDATTADALSTAGFVMAGKDWRKVVLDFPNMGAHLLIVLEG
GAIERSETFKLFERE
11111111111111110000000000000000000000000000000000000000000000
00000000000000000000000000000000000000000000000000000000000000
00000000000000000000000000000000000000000000000000000000000000
00000000000000000000000000000000000000000000000000000000000000
00000000000000000000000000000000000000000000000000000000000000
000000000000000
>DM_train705
MAMSQQDLPTLFYSGKSNSAVPIISESELQTITAEPWLEISKKGLQLEGLNFDRQGQLFLLD
VFEGNIFKINPETKEIKRPFVSHKANPAAIKIHKDGRLFVCYLGDFKSTGGIFAATENGDNL
QDIIEDLSTAYCIDDMVFDSKGGFYFTDFRGYSTNPLGGVYYVSPDFRTVTPIIQNISVANG
IALSTDEKVLWVTETTANRLHRIALEDDGVTIQPFGATIPYYFTGHEGPDSCCIDSDDNLYV

AMYGQGRVLVFNKRGYPIGQILIPGRDEGHMLRSTHPQFIPGTNQLIICSNDIEMGGGSMLY
TVNGFAKGHQSFQFQ
111110000000000000000000000000000000000000000000000000000000
000000000000000000000000000000000000000000000000000000000000
000000000000000000000000000000000000000000000000000000000000
000000000000000000000000000000000000000000000000000000000000
000000000000000000000000000000000000000000000000000000000000
000000000000000
>DM_train706
AAAADTTVTQFLQSFKENAENGALRKFYEVIMDNGGAVLDDINSLTEVTILAPSNEAWNSSN
INNVLRDRNKMRQILNMHIIKDRLNVDKIRQKNANLIAQVPTVNNNTFLYFNVRGEGSDTVI
TVEGGGVNATVIQADVAQTNGYVHIIDHVLGVPYTTVLGKLESDPMMSDTYKMGKFSHFNDQ
LNNTQRRFTYFVPRDKGWQKTELDYPSAHKKLFMADFSYHSKSILERHLAISDKEYTMKDLV
KFSQESGSVILPTFRDSLSIRVEEEAGRYVIIWNYKKINVYRPDVECTNGIIHVIDYPLLEE
KDVVVAGGSAAAAA
111111111111111111000000000000000000000000000000000000000000
000000000000000000000000000000000000000000000000000000000000
000000000000000000000000000000000000000000000000000000000000
000000000000000000000000000000000000000000000000000000000000
000000000000000000000000000000000000000000000000000000000000
00001111111111
>DM_train708
MTKERFVISRRRTGFGDCLWSLASAWSYAQRTGRTLVIDWRGSCYVEQPFSNAFPAFFEPVE
DIAGVPVICDDRVNQLSFPGPFFPRWWNRPSIDCINRPDEQIFRERDELTELFQAREDSEAN
TIVCDACLMWRCSEEAERLIFRNIKLRSEIRARIDALYEEHFSGHSIIGVHVRHGNGEDIME
HAPYWADSELALHQVCMAIRKAKALSYPKPVKVFLCTDSAQVLDQVSGLFPDVFAVPKRFQA
DRAGPLHSAEMGIEGGASALIDMYLLARCATVIRFPPTSAFTRYARLLVPRIIEFDLSNPGH
LTMIDNPYEHFAAS
100000000000000000000000000000000000000000000000000000000000
000000000000000000000000000000000000000000000000000000000000
000000000000000000000000000000000000000000000000000011111111
111100000000000000000000000000000000000000000000000000001111
111111100000000000000000000000000000000000000000000000110000
00000001111111
>DM_train709
MKTRITELLKIDYPIFQGGMAWVADGDLAGAVSKAGGLGIIGGGNAPKEVVKANIDKIKSLT
DKPFGVNIMLLSPFVEDIVDLVIEEGVKVVTTGAGNPSKYMERFHEAGIIVIPVVPSVALAK
RMEKIGADAVIAEGMEAGGHIGKLTTMTLVRQVATAISIPVIAAGGIADGEGAAAGFMLGAE
AVQVGTRFVVAKESNAHPNYKEKILKARDIDTTISAQHFGHAVRAIKNQLTRDFELAEKDAF
KQEDPDLEIFEQMGAGALAKAVVHGDVDGGSVMAGQIAGLVSKEETAEEILKDLYYGAAKKI
QEEASRWTGVVRND
000000000000000000000000000000000000000000000000000000000000
000000000000000000000000000000000000000000000000000000000000
000000000000000000000000000000000000000000000000000000000000

000000000000000000000000000000000000000000000000000000000000
001110000000000000000000000000000000000000000000000000000000
00000000001111
>DM_train710
EIFSQELTQREANVKKVHENLEELQKKLDHTSFAHKEDRDRLEAQIAQKEQEQKAKLAEYDQ
KVQNEFDARERAEREREAARGDAAAEKQRLASLLKDLEDDASGYNRLRPSKPMLSEEDTNIL
RQLFLSSAVSGSGKFSFQDLKQVLAKYADTIPEGPLKKLFVMVENDTKGRMSYITLVAVAND
LAALVADFRKIDTNSNGTLSRKEFREHFVRLGFDKKSVQDALFRYADEDESDDVGFSEYVHL
GLCLLVLRILYAFADFDKSGQLSKEEVQKVLEDAHIPESARKKFEHQFSVVDVDDSKSLSYQ
EFVMLVLLMFHDD
000000000000000000000000000000000011110000000000000000000000
000000000000000000000000000000000000011111111111000000000000
000000000000000000000000000000000000000000000000000000000000
000000000000000000000000000000000000000000000000000000000000
000000000000000000000000000000000000000000000000000000000000
0000000000000
>DM_train711
FNKIKDSVKQKIDSMGDKGTYGVSASHPLAVEEGMKVLKNGGSAVDAAIVVSYVLGVVELHA
SGIGGGGGMLIISKDKETFIDYRETTPYFTGNQKPHIGVPGFVAGMEYIHDNYGSLPMGELL
QPAINYAEKGFKVDDSLTMRLDLAKPRIYSDKLSIFYPNGEPIETGETLIQTDLARTLKKIQ
KEGAKGFYEGGVARAISKTAKISLEDIKGYKVEVRKPVKGNYMGYDVYTAPPPFSGVTLLQM
LKLAEKKEVYKDVDHTATYMSKMEEISRIAYQDRKKNLGDPNYVNMDPNKMVSDKYISTMKN
ENGDALSEAEHES
111111111111111110000000000000000000000000000000000000000000
000000000000000000000000000011111000000000000000000000000000
000000000000000000000000000000000000000000000000000000000000
000000000000000000000000000000000000000000000000000000000000
000000000000000000000000000000000000001111110000000000000001
1111111111111
>DM_train712
MKNDRTLQAIGRQLKAMGCERFDIGVRDATTGQMMNREWSAAEVLQNTPWLKRMNAQGNDVY
IRPAEQERHGLVLVDDLSEFDLDDMKAEGREPALVVETSPKNYQAWVKVADAAGGELRGQIA
RTLASEYDADPASADSRHYGRLAGFTNRKDKHTTRAGYQPWVLLRESKGKTATAGPALVQQA
GQQIEQAQRQQEKARRLASLELPERQLSRHRRTALDEYRSEMAGLVKRFGDDLSKCDFIAAQ
KLASRGRSAEEIGKAMAEASPALAERKPGHEADYIERTVSKVMGLPSVQLARAELARAPAPR
QRGMDRGGPDFSM
111100000000000000000000000000000000000000000000000000000000
000000000000000000000000000000000000000000000000000000000000
000000000000000000000000000010000111000000000000000000000000
000000000000000000001111111110000000000000000000000000000000
000000000000000000000000001100000000000000000000000000000111
1111111111111
>DM_train713
MADDFAYLNARVRVRRGTLLKESFFQEALDLSFADFLRLLSETVYGGELAGQGLPDVDRAVL

RTQAKLVGDLPRLVTGEAREAVRLLLLRNDLHNLQALLRAKATGRPFEEVLLLPGTLREEVW
RQAYEAQDPAGMAQVLAVPGHPLARALRAVLRETQDLARVEALLAKRFFEDVAKAAKGLDQP
ALRDYLALEVDAENLRTAFKLQGSGLAPDAFFLKGGRFVDRVRFARLMEGDYAVLDELSGTP
FSGLSGVRDLKALERGLRCVLLKEAKKGVQDPLGVGLVLAYVKEREWEAVRLRLLARRAYFG
LPRAQVEEEVVCP
111100000000000000000000000000000000000000000000000000000000
000000000000000000000000000000000000000000000000000000000000
000000000000000000000000000000000000000000000000000011111110
000000000000000000000000000000000000000000000000000000000000
000000000000000000000000000000000000000000000000000000000000
0000000000000
>DM_train714
QLTPTFYDNSCPNVSNIVRDTIVNELRSDPRIAASILRLHFHDCFVNGCDASILLDNTTSFR
TEKDAFGNANSARGFPVIDRMKAAVESACPRTVSCADLLTIAAQQSVTLAGGPSWRVPLGRR
DSLQAFLDLANANLPAPFFTLPQLKDSFRNVGLNRSSDLVALSGGHTFGKNQCRFIMDRLYN
FSNTGLPDPTLNTTYLQTLRGLCPLNGNLSALVDFDLRTPTIFDNKYYVNLEEQKGLIQSDQ
ELFSSPNATDTIPLVRSFANSTQTFFNAFVEAMDRMGNITPLTGTQGQIRLNCRVVNSNSLL
HDMVEVVDFVSSM
000000000000000000000000000000000000000000000000000000000000
000000000000000000000000000000000000000000000000000000000000
000000000000000000000000000000000000000000000000000000000000
000000000000000000000000000000000000000000000000000000000000
000000000000000000000000000000000000000000000000000000001111
1111111111111
>DM_train715
MGSSHHHHHHSSGRENLYFQGHMEGVRWAFSCGTWLPSRAEWLLAVRSIQPEEKERIGQFV
FARDAKAAMAGRLMIRKLVAEKLNIPWNHIRLQRTAKGKPVLAKDSSNPYPNFNFNISHQGD
YAVLAAEPELQVGIDIMKTSFPGRGSIPEFFHIMKRKFTNKEWETIRSFKDEWTQLDMFYRN
WALKESFIKAIGVGLGFELQRLEFDLSPLNLDIGQVYKETRLFLDGEEEKEWAFEESKIDEH
HFVAVALRKPDGSRHQDVPSQDDSKPTQRQFTILNFNDLMSSAVPMTPEDPSFWDCFCFTEE
IPIRNGTKSLESG
111111111111111110000000000000000000000000000000000000000000
000000000000000000000000000000000000000000011000000000000000
000000000000000000000000000000000000000000000000000000000000
000000000000000000000000000000000000000000000000000000000000
000000000111111111111111100000000000000000000000000000000000
0000011111111
>DM_train716
MPLCTLRQMLGEARKHKYGVGAFNVNNMEQIQGIMKAVVQLKSPVILQCSRGALKYSDMIYL
KKLCEAALEKHPDIPICIHLDHGDTLESVKMAIDLGFSSVMIDASHHPFDENVRITKEVVAY
AHARSVSVEAELGTLGGIEEDVQNTVQLTEPQDAKKFVELTGVDALAVAIGTSHGAYKFKSE
SDIRLAIDRVKTISDLTGIPLVMHGSSSVPKDVKDMINKYGGKMPDAVGVPIESIVHAIGEG
VCKINVDSDSRMAMTGAIRKVFVEHPEKFDPRDYLGPGRDAITEMLIPKIKAFGSAGHAGDY
KVVSLEEAKAWYK

10000000000000000000000000000000000000000000000000000000000000
00000000000000000000000000000000000000000000000000000000000000
00000000000001111111111111100000000000000000000000011111111111
11110000000000000000000001110000000000000000000011000000000000
00000000000000000000000000000000000000000000000000000000000000
0000000000000
>DM_train717
MTNSNDIDETAATIDTARALLKSFGFEAQRHNVRSAVTLLALAGLKPGDHWADSTTPRLGVQ
KIMDWSGAYWAKPYATGSREDFRKKTLRQWVDNGFAVLNPDNLNIATNSQLNEYCLSDEAAQ
AIRSYGTDAFESALVDFLSKASDTVRARAEALRAAMISVDLADGDEFLLSPAGQNPLLKKMV
EEFMPRFAPGAKVLYIGDTRGKHTRFEKRIFEETLGLTFDPHGRMPDLVLHDKVRKWLFLME
AVKSKGPFDEERHRTLRELFATPVAGLVFVNCFENREAMRQWLPELAWETEAWVADDPDHLI
HLNGSRFLGPYER
11110000000000000000000000000000000000000000000000000000000000
00000000000000000000000000000000000000000000000000000000000000
00000000000000000000000000000000000000000000000000000000000000
00000000000000000000000000000000000000000000000000000000000000
00000000000000000000000000000000000000000000000000000000000000
0000000000000
>DM_train719
MLRGGVNDCKIKWEFLIGNSIDSSPILAKNGTIYLGSSNKNLYAINTDGSVKWFFKSGEIIE
CRPSIGKDGTIYFGSDKVYAINPDGTEKWRFDTKKAIVSDFTIFEDILYVTSMDGHLYAINT
DGTEKWRFKTKKAIYATPIVSEDGTIYVGSNDNYLYAINPDGTEKWRFKTNDAITSAASIGK
DGTIYFGSDKVYAINPDGTEKWNFYAGYWTVTRPAISEDGTIYVTSLDGHLYAINPDGTEKW
RFKTGKRIESSPVIGNTDTIYFGSYDGHLYAINPDGTEKWNFETGSWIIATPVIDENGTIYF
GTRNGKFYALFN
11111111111000000011110000000000000011100000000000000001110000
00000000000000000000000111111111111111000000000000000000000000
00000000000000000000000000000000000000000000000000000000000000
00000000000000000000000000000000000000000000000000000000000000
00000000000000000000000000000000000000000000000000000000000000
000000000000
>DM_train720
TENKRKSMEKLSVDGVISALAFDQRGALKRMMAQHQTKEPTVEQIEELKSLVSEELTPFASS
ILLDPEYGLPASRVRSEEAGLLLAYEKTGYDATTTSRLPDCLDVWSAKRIKEAGAEAVKFLL
YYDIDGDQDVNEQKKAYIERIGSECRAEDIPFYLEILTYDEKIADNASPEFAKVKAHKVNEA
MKVFSKERFGVDVLKVEVPVNMKFVEGFADGEVLFTKEEAAQAFRDQEASTDLPYIYLSAGV
SAKLFQDTLVFAAESGAKFNGVLCGRATWAGSVKVYIEEGPQAAREWLRTEGFKNIDELNKV
LDKTASPWTEKM
00000000000000000000000000000000000000000000000000000000000000
00000000000000000000000000000000000000000000000000000000000000
00000000000000000000000000000000000000000000000000000000000000
00000000000000000000000000000000000000000000000000000000000000
00000000000000000000000000000000000000000000000000000000000000

000000000111
>DM_train721
SEDPVFTFGLIADVQYADIEDGENYLRTRRRYYRGSADLLRDAVLQWRRERVQCVVQLGDII
DGHNRRRDASDRALDTVMAELDACSVDVHHVWGNHEFYNFSRPSLLSSRLNSAQRTGTDTGS
DLIGDDIYAYEFSPAPNFRFVLLDAYDLSVIGREEESEKHTHSWRILTQHNHNLQDLNLPPV
SVGLEQRFVKFNGGFSEQQLQWLDAVLTLSDHKQERVLIFSHLPVHPCAADPICLAWNHEAV
LSVLRSHQSVLCFIAGHDHDGGRCTDSSGAQHITLEGVIETPPHSHAFATAYLYEDRMVMKG
RGRVEDLTITYS
11000000000000000000000000000000000000000000000000000000000000
00000000000000000000000000000000000000000000000000000001111100
00000000000000000000000000000000000000000000000000000000000000
00000000000000000000000000000000000000000000000000000000000000
00000000000000000000000000000000000000000000000000000000000000
000000000000
>DM_train722
MAVYTDITEDELRNFLTQYDVGSLTSYKGIAEGVENSNFLLHTTKDPLILTLYEKRVEKNDL
PFFLGLMQHLAAKGLSCPLPLPRKDGELLGELSGRPAALISFLEGMWLRKPEAKHCREVGKA
LAAMHLASEGFEIKRPNALSVDGWKVLWDKSEERADEVEKGLREEIRPEIDYLAAHWPKDLP
AGVIHADLFQDNVFFLGDELSGLIDFYFACNDLLAYDVSICLNAWCFEKDGAYNVTKGKALL
EGYQSVRPLSEAELEALPLLSRGSALRFFLTRLYDWLTTPAGALVVKKDPLEYLRKLRFHRT
IANVAEYGLAGE
11110000000000000000000000000001110000000000000000000001110000
00000000000000000000000000000000000000000000000000000000000000
00000000000000000000000000000000000000000000000000000000000000
00000000000000000000000000000000000000000000000000000000000000
00000000000000000000000000000000000000000000000000000000000000
000000000011
>DM_train723
GMTGLSQSQASPMQIQPGNAAFNPWTDAALDTIRDVNQALTLYAEMRVVPAHHDAFLAAIDT
VSAKLRVLPGFLSLALKQMSGDSTMVKNYPETYKGVLATAYLDGVAAGTQPYFYNLFVRFAD
GRAARAAGFEALFETHIHPLLHAMAPRGGDGPELLAYRAVLQSVVAGDRHAIYRGAEEIRSF
LRRPVELPERETVTVENHVMVPEDKHAAWEPQVAILLQVAQDTFEPQDEPSGVGLPGARDNR
YYRKALSTEILRNAHADGGLRAYIMHGVWESVWDHENSHLDPRFLAAAGPVGAAAVVGPVEP
FYLTRRLVVAD
11111111111111111111100000000000011000000000000000000000000000
00000000000000000000000000000000000000000000000000000000000000
00000000000000000000000001111000000000000000000000000000000000
00000000000000000000000000000000000000000000000000000000000000
00000000000000000000000000000000000000000000000000000000000000
00000000000
>DM_train724
MSSGVDLGTENLYFQSNAMYYGFDIGGTKIALGVFDSTRRLQWEKRVPTPHTSYSAFLDAVC
ELVEEADQRFGVKGSVGIGIPGMPETEDGTLYAANVPAASGKPLRADLSARLDRDVRLDNDA
NCFALSEAWDDEFTQYPLVMGLILGTGVGGGLVLNGKPITGQSYITGEFGHMRLPVDALTLM

GFDFPLRRCGCGQMGCIENYLSGRGFAWLYQHYYDQSLQAPEIIALWEQGDEQAHAHVERYL
DLLAVCLGNILTIVDPDLLVIGGGLSNFTAITTQLAERLPRHLLPVARAPRIERARHGDAGG
MRGAAFLHLTD
111111111111111100000000000000000000000000000000000000000000
000000000000000000000000000000000000000000000000000000000000
000000000000000000000000000000000000000000000000000000000000
000000000000000000000000000000000000000000000000000000000000
000000000000000000000000000000000000000000000000000000000000
00000000000
>DM_train725
MLVPRGSMLDFEKPLFEIRNKIESLKESQDKNDVDLQEEIDMLEASLERETKKIYTNLKPWD
RVQIARLQERPTTLDYIPYIFDSFMELHGDRNFRDDPAMIGGIGFLNGRAVTVIGQQRGKDT
KDNIYRNFGMAHPEGYRKALRLMKQAEKFNRPIFTFIDTKGAYPGKAAEERGQSESIATNLI
EMASLKVPVIAIVIGEGGSGGALGIGIANKVLMLENSTYSVISPEGAAALLWKDSNLAKIAA
ETMKITAHDIKQLGIIDDVISEPLGGAHKDIEQQALAIKSAFVAQLDSLESLSRDEIANDRF
EKFRNIGSYIE
111111100000000000000001111111111111111100000000000000000000
000000000000000000000000000000000000000000000000000000000000
000000000000000000000000000000000000000000000000000000000000
000000000000000000000000000000000000000000000000000000000000
000000000000000000000000000000000000000000000000000000000000
00000000000
>DM_train728
MLVFIDDGSTNIKLQWQESDGTIKQHISPNSFKREWAVSFGDKKVFNYTLNGEQYSFDPISP
DAVVTTNIAWQYSDVNVVAVHHALLTSGLPVSEVDIVCTLPLTEYYDRNNQPNTENIERKKA
NFRKKITLNGGDTFTIKDVKVMPESIPAGYEVLQELDELDSLLIIDLGGTTLDISQVMGKLS
GISKIYGDSSLGVSLVTSAVKDALSLARTKGSSYLADDIIIHRKDNNYLKQRINDENKISIV
TEAMNEALRKLEQRVLNTLNEFSGYTHVMVIGGGAELICDAVKKHTQIRDERFFKTNNSQYD
LVNGMYLIGN
000000000000000000000000000000000000000000000000000000000000
000000000000000000000000000000000000000000000000000000000000
000000000000000000000000000000000000000000000000000000000000
000000000000000000000000011111000000000000000000000000000000
000000000000000000000000000000000000000000000000000000000000
0000000000
>DM_train729
MSANDFKPETWTSSANEALRVSIVGENAVQFSPLFTYPIYGDSEKIYGYKDLIIHLAFDSVT
FKPYVNVKYSAKLGDDNIVDVEKKLLSFLPKDDVIVRDEAKWVDCFAEERKTHNLSDVFEKV
SEYSLNGEEFVVYKSSLVDDFARRMHRRVQIFSLLFIEAANYIDETDPSWQIYWLLNKKTKE
LIGFVTTYKYWHYLGAKSFDEDIDKKFRAKISQFLIFPPYQNKGHGSCLYEAIIQSWLEDKS
ITEITVEDPNEAFDDLRDRNDIQRLRKLGYDAVFQKHSDLSDEFLESSRKSLKLEERQFNRL
VEMLLLLNNS
111110000000000000000000000000000000000000000000000000000000
000000000000000000000000000000000000000000000000000000000000

000000000000000000000000000000000000000000000000000000000000
000000000000011111111100000000000000000000000000000000000000
000000000000000000000000000000000000000000000000000000000000
0000000000
>DM_train730
MQLFDLSLEELKKYKPKKTARPDFSDFWKKSLEELRQVEAEPTLESYDYPVKGVKVYRLTYQ
SFGHSKIEGFYAVPDQTGPHPALVRFHGYNASYDGGIHDIVNWALHGYATFGMLVRGQGGSE
DTSVTPGGHALGWMTKGILSKDTYYYRGVYLDAVRALEVIQSFPEVDEHRIGVIGGSQGGAL
AIAAAALSDIPKVVVADYPYLSNFERAVDVALEQPYLEINSYFRRNSDPKVEEKAFETLSYF
DLINLAGWVKQPTLMAIGLIDKITPPSTVFAAYNHLETDKDLKVYRYFGHEFIPAFQTEKLS
FLQKHLLLST
100000000000000000000000000000000000000000000000000000000000
000000000000000000000000000000000000000000000000000000000000
000000000000000000000000000000000000000000000000000000000000
000000000000000000000000000000000000000000000000000000000000
000000000000000000000000000000000000000000000000000000000000
0000000111
>DM_train731
MKKPNGTNGASSSLEPPPSTFQPLCHPLVEEVSKEVDGYFLQHWNFPNEKARKKFVAAGFSR
VTCLYFPKALDDRIHFACRLLTVLFLIDDLLEYMSFEEGSAYNEKLIPISRGDVLPDRSIPV
EYIIYDLWESMRAHDREMADEILEPVFLFMRAQTDRTRARPMGLGGYLEYRERDVGKELLAA
LMRFSMGLKLSPSELQRVREIDANCSKHLSVVNDIYSYEKELYTSKTAHSEGGILCTSVQIL
AQEADVTAEAAKRVLFVMCREWELRHQLLVARLSAEGLETPGLAAYVEGLEYQMSGNELWSQ
TTLRYSVVVD
111111111111000000000000000000000000000000000000000000000000
000000000000000000000000000000000000000000000000000000000000
000000000000000000000000000000000000000000000000000000000000
000000000000000000000000000000000000000000111111111100000000
000000000000000000000000000000000001100000000000000000000000
0000000111
>DM_train733
MTDSPGGVAPASPVEDASDASLGQPEEGAPCQVVLQGAELNGILQAFAPLRTSLLDSLLVMG
DRGILIHNTIFGEQVFLPLEHSQFSRYRWRGPTAAFLSLVDQKRSLLSVFRANQYPDLRRVE
LAITGQAPFRTLVQRIWTTTSDGEAVELASETLMKRELTSFVVLVPQGTPDVQLRLTRPQLT
KVLNATGADSATPTTFELGVNGKFSVFTTSTCVTFAAREEGVSSSTSTQVQILSNALTKAGQ
AAANAKTVYGENTHRTFSVVVDDCSMRAVLRRLQVGGGTLKFFLTTPVPSLCVTATGPNAVS
AVFLLKPQK
111111111111111111111111111100000000000000000000000000000000
000000000000000000000000000000000000000000000000000000000000
000000000000000000000000000000000000000000000000000000000000
000000000000000000000000000000000000000111111111111111111111
111000000000000000000000000000000000000000000000000000000000
000000000
>DM_train734

LEADKCKEREEKIILVSSANEIDVRPCPLNPNEHKGTITWYKDDSKTPVSTEQASRIHQHKE
KLWFVPAKVEDSGHYYCVVRNSSYCLRIKISAKFVENEPNLCYNAQAIFKQKLPVAGDGGLV
CPYMEFFKNENNELPKLQWYKDCKPLLLDNIHFSGVKDRLIVMNVAEKHRGNYTCHASYTYL
GKQYPITRVIEFITLEENKPTRPVIVSPANETMEVDLGSQIQLICNVTGQLSDIAYWKWNGS
VIDEDDPVLGEDYYSVENPANKRRSTLITVLNISEIESRFYKHPFTCFAKNTHGIDAAYIQL
IYPVTNFQK
1110000000000000000000000000000000000000000000000000000000000000
0000000000000000000000000000000000000000000000000000000000000000
0000000000000000000000000000000000000000000000000000000000000000
0000000000000000000000000000000000000000000000000000000000000000
0000000000000000000000000000000000000000000000000000000000000000
000011111
>DM_train735
RHRLFQLNREVDDLEQWIAEREVVAGSHELGQDYEHVTMLQERFREFARDTGNIGQERVDTV
NHLADELINSGHSDAATIAEWKDGLNEAWADLLELIDTRTQILAASYELHKFYHDAKEIFGR
IQDKHKKLPEELGRDQNTVETLQRMHTTFEHDIQALGTQVRQLQEDAARLQAAYAGDKADDI
QKRENEVLEAWKSLLDACESRRVRLVDTGDKFRFFSMVRDLMLWMEDVIRQIEAQEKPRDVS
SVELLMNNHQGIKAEIDARNDSFTTCIELGKSLLARKHYASEEIKEKLLQLTEKRKEMIDKW
EDRWEWLRL
0000000000000000000000000000000000000000000000000000000000000000
0000000000000000000000000000000000000000000000000000000000000000
0000000000000000000000000000000000000000000000000000000000000000
0000000000000000000000000000000000000000000000000000000000000000
0000000000000000000000000000000000000000000000000000000000000000
000000001
>DM_train736
ADTYAATRYPVILVHGLAGTDKFANVVDYWYGIQSDLQSHGAKVYVANLSGFQSDDGPNGRG
EQLLAYVKQVLAATGATKVNLIGHSQGGLTSRYVAAVAPQLVASVTTIGTPHRGSEFADFVQ
DVLKTDPTGLSSTVIAAFVNVFGTLVSSSHNTDQDALAALRTLTTAQTATYNRNFPSAGLGA
PGSCQTGAATETVGGSQHLLYSWGGTAIQPTSTVLGVTGATDTSTGTLDVANVTDPSTLALL
ATGAVMINRASGQNDGLVSRCSSLFGQVISTSYHWNHLDEINQLLGVRGANAEDPVAVIRTH
VNRLKLQGV
0000000000000000000000000000000000000000000000000000000000000000
0000000000000000000000000000000000000000000000000000000000000000
0000000000000000000000000000000000000000000000000000000000000000
0000000000000000000000000000000001111100000000000000000000000000
0000000000000000000000000000000000000000000000000000000000000000
000000000
>DM_train737
SNAMTQSPIFLTPVFKEKIWGGTALRDRFGYSIPSESTGECWAISAHPKGPSTVANGPYKGK
TLIELWEEHREVFGGVEGDRFPLLTKLLDVKEDTSIKVHPDDYYAGENEEGELGKTECWYII
DCKENAEIIYGHTARSKTELVTMINSGDWEGLLRRIKIKPGDFYYVPSGTLHALCKGALVLE
TQQNSDATYRVYDYDRLDSNGSPRELHFAKAVNAATVPHVDGYIDESTESRKGITIKTFVQG
EYFSVYKWDINGEAEMAQDESFLICSVIEGSGLLKYEDKTCPLKKGDHFILPAQMPDFTIKG

TCTLIVSHI
111110000000000000000000000000000000000000000000000000000000
000000000000000000000000000000000000000000000000000000000000
000000000000000000000000000000000000000000000000000000000000
000000000000000000000000000000000000000000000000000000000000
000000000000000000000000000000000000000000000000000000000000
000000000
>DM_train738
AKAPEQTVTEPVGSYARAERPQDFEGFVWRLDNDGKEALPRNFRTSADALRAPEKKFHLDAA
YVPSREGMDALHISGSSAFTPAQLKNVAAKLREKTAGPIYDVDLRQESHGYLDGIPVSWYGE
RDWANLGKSQHEALADERHRLHAALHKTVYIAPLGKHKLPEGGEVRRVQKVQTEQEVAEAAG
MRYFRIAATDHVWPTPENIDRFLAFYRTLPQDAWLHFHCEAGVGRTTAFMVMTDMLKNPSVS
LKDILYRQHEIGGFYYGEFPIKTKDKDSWKTKYYREKIVMIEQFYRYVQENRADGYQTPWSV
WLKSHPAKA
111111000000000000000000000000000000000000000000000000000000
000000000000000000000000000000000000000000000000000000000000
000000000000000000000000000000000000000000000000000000000000
000000000000000000000000000000000000000000000000000000000000
000000000000000000000000000000000000000000000000000000000000
000000000
>DM_train739
GSPGISGGGGGILDSMAEIVADKTVEVVKNAIETADGALDLYNKYLDQVIPWQTFDETIKEL
SRFKQEYSQAASVLVGDIKTLLMDSQDKYFEATQTVYEWCGVATQLLAAYILLFDEYNEKKA
SAQKDILIKVLDDGITKLNEAQKSLLVSSQSFNNASGKLLALDSQLTNDFSEKSSYFQSQVD
KIRKEAYAGAAAGVVVGPFGLIISYSIAAGVVEGKLIPELKNKLKSVQNFFTTLSNTVKQAN
KDIDAAKLKLTTEIAAIGEIKTETETTRFYVDYDDLMLSLLKEAAKKMINTCNEYQKRHGKK
TLFEVPEV
111111111100000000000000000000000000000000000000000000000000
000000000000000000000000000000000000000000000000000000000000
000000000000000000000000000000000000000000000000000000000000
000000000000000000000000000000000000000000000000000000000000
000000000000000000000000000000000000000000000000000000000000
00011111
>DM_train741
VSQIGLLLPLSGDGQILGTTIQSGFNDAKGNSTIPVQVFDTSMNSVQDIIAQAKQAGIKTLV
GPLLKQNLDVILADPAQIQGMDVLALNATPNSRAIPQLCYYGLSPEDEAESAANKMWNDGVR
NPLVAMPQNDLGQRVGNAFNVRWQQLAGTDANIRYYNLPADVTYFVQENNSNTTALYAVASP
TELAEMKGYLTNIVPNLAIYASSRASASATNTNTDFIAQMNGVQFSDIPFFKDTNSPQYQKL
AKSTGGEYQLMRLYAMGADAWLLINQFNELRQVPGYRLSGLTGILSADTNCNVERDMTWYQY
QDGAIVPV
100000000000000000000000000000000000000000000000000000000000
000000000000000000000000000000000000000000000000000000000000
000000000000000000000000000000000000000000000000000000000000
000000000000000000000000000000000000000000000000000000000000

000000000000000000000000000000000000000000000000000000000000
00000000
>DM_train742
MSLEVKVGLAPMAGYTDSAFRTLAFEWGADFAFSEMVSAKGFLMNSQKTEELLPQPHERNVA
VQIFGSEPNELSEAARILSEKYKWIDLNAGCPVRKVVKEGAGGALLKDLRHFRYIVRELRKS
VSGKFSVKTRLGWEKNEVEEIYRILVEEGVDEVFIHTRTVVQSFTGRAEWKALSVLEKRIPT
FVSGDIFTPEDAKRALEESGCDGLLVARGAIGRPWIFKQIKDFLRSGKYSEPSREEILRTFE
RHLELLIKTKGERKAVVEMRKFLAGYTKDLKGARRFREKVMKIEEVQILKEMFYNFIKEVEG
GSHHHHHH
111100000000000000000000000000000000000000000000000000000000
000000000000000000000000000000000000000000000000000000000000
000000000000000000000000000000000000000000000000000000000000
000000000000000000000000000000000000000000000000000000000000
000000000000000000000000000000000000000000000000000000000001
11111111
>DM_train743
MPKPYVAINMAELKNEPKTFEMFASVGPKVCMVTARHPGFVGFQNHIQIGILPFGNRYGGAK
MDMTKESSTVRVLQYTFWKDWKDHEEMHRQNWSYLFRLCYSCASQMIWGPWEPIYEIIYANM
PINTEMTDFTAVVGKKFAEGKPLDIPVISQPYGKRVVAFAEHSVIPGKEKQFEDAIVRTLEM
LKKAPGFLGAMVLKEIGVSGIGSMQFGAKGFHQVLENPGSLEPDPNNVMYSVPEAKNTPQQY
IVHVEWANTDALMFGMGRVLLYPELRQVHDEVLDTLVYGPYIRILNPMMEGTFWREYLNEQA
WRHPQFGG
100000000000000000000000000000000000000000000000000000000000
000000000000000000000000000000000000000000000000000000000000
000000000000000000000000000000000000000000000000000000000000
000000000000000000000000000000000000000000000000000000000000
000000000000000000000000000000000000000000000000000000000011
11111111
>DM_train744
EGGDFVVGMVTDSGDIDDKSFNQQVWEGISRFAQENNAKCKYVTASTDAEYVPSLSAFADEN
MGLVVACGSFLVEAVIETSARFPKQKFLVIDAVVQDRDNVVSAVFGQNEGSFLVGVAAALKA
KEAGKSAVGFIVGMELGMMPLFEAGFEAGVKAVDPDIQVVVEVANTFSDPQKGQALAAKLYD
SGVNVIFQVAGGTGNGVIKEARDRRLNGQDVWVIGVDRDQYMDGVYDGSKSVVLTSMVKRAD
VAAERISKMAYDGSFPGGQSIMFGLEDKAVGIPEENPNLSSAVMEKIRSFEEKIVSKEIVVP
VRSARMMN
110000000000000000000000000000000000000000000000000000000000
000000000000000000000000000000000000000000000000000000000000
000000000000000000000000000000000000000000000000000000000000
000000000000000000000000000000000000000000000000000000000000
000000000000000000000000000000000000000000000000000000000000
00000000
>DM_train745
MEQLRGLYPPLAAYDSGWLDTGDGHRIYWELSGNPNGKPAVFIHGGPGGGISPHHRQLFDPE
RYKVLLFDQRGCGRSRPHASLDNNTTWHLVADIERLREMAGVEQWLVFGGSWGSTLALAYAQ

THPERVSEMVLRGIFTLRKQRLHWYYQDGASRFFPEKWERVLSILSDDERKDVIAAYRQRLT
SADPQVQLEAAKLWSVWEGETVTLLPSRESASFGEDDFALAFARIENHYFTHLGFLESDDQL
LRNVPLIRHIPAVIVHGRYDMACQVQNAWDLAKAWPEAELHIVEGAGHSYDEPGILHQLMIA
TDRFAGK
111000000000000000000000000000000000000000000000000000000000
000000000000000000000000000000000000000000000000000000000000
000000000000000000000000000000000000000000000000000000000000
000000000000000000000000000000000000000000000000000000000000
000000000000000000000000000000000000000000000000000000000000
0000000
>DM_train746
MAELACFCYPHLENDSYKFIPFNNLAIKAMLTAKVDKKDMDKFYDSIIYGIAPPPQFKKRYN
TNDNSRGMNFETIMFTKVAMLICEALNSLKVTQANVSNVLSRVVSIRHLENLVIRKENPQDI
LFHSKDLLLKSTLIAIGQSKEIETTITAEGGEIVFQNAAFTMWKLTYLEHQLMPILDQNFIE
YKVTLNEDKPISDVHVKELVAELRWQYNKFAVITHGKGHYRIVKYSSVANHADRVYATFKSN
VKTGVNNDFNLLDQRIIWQNWYAFTSSMKQGNTLDVCKRLLFQKMKPEKNPFKGLSTDRKMD
EVSQVGV
000000000000000000000000000000000000000000000000000000000000
000000000000000000000000000000000000000000000000000000000000
000000000000000000000000000000000000000000000000000000000000
000000000000000000000000000000000000000000000000000000000000
000000000000000000000000000000000000000000000000000000000000
0001111
>DM_train747
MSNQYGDKNLKIFSLNSNPELAKEIADIVGVQLGKCSVTRFSDGEVQINIEESIRGCDCYII
QSTSDPVNEHIMELLIMVDALKRASAKTINIVIPYYGYARQDRKARSREPITAKLFANLLET
AGATRVIALDLHAPQIQGFFDIPIDHLMGVPILGEYFEGKNLEDIVIVSPDHGGVTRARKLA
DRLKAPIAIIDKRRPRPNVAEVMNIVGNIEGKTAILIDDIIDTAGTITLAANALVENGAKEV
YACCTHPVLSGPAVERINNSTIKELVVTNSIKLPEEKKIERFKQLSVGPLLAEAIIRVHEQQ
SVSYLFS
111111110000000000000000000000000000000000000000000000000000
000000000000000000000000000000000000000000000000000000000000
000000000000000000000000000000000000000000000000000000000000
000000000000011111111100000000000000000000000000000000000000
000000000000000000000000000000000111100000000000000000000000
0000000
>DM_train749
GMPPVWTLPRLYQHFQGAIDLELWTIPYYLTVLYSIKDPTTVPYRLIQAAVYQEMLHAQLVS
NIANAYGYSPTLSAPEYVGTAVPHIDFDLDTPNPTSIFTPYSAELGPLDLTRVNTMCLIEYP
EWRTQREPDLADDVTDYGSIGEFYDALRVGMEQLRGHVRGNQKQMDEFGPFYQNSPPLTVTE
SGDAGFLQALTLVDIIVDQGEGQTQPVETIPTEFQNTADGFQDAWPHFQRFDFIRRMPNWPG
VYTGVTDPPAGSPGAEAQARLIADFAGFLDILNGMFSGGGAPPAFGVQMAKLGGDILSCWKL
GAVPRYS
100000000000000000000000000000000000000000000000000000000000

000000000000000000000000000000000000000000000000000000000000
000000000000000000000000000000000000000000000111111110000000
000000000000000000011111111111111111111111100000000000000000
000000000000000000000000000000000000000000000000000000000000
0000000
>DM_train750
GMNPFQNSFGGHIPQDVAGKQGENVIFIVYNLTDSPDTVDKVKDVCANFSAMIRSMRNRFPD
MQFSCTMGFGADAWTRLFPDKGKPKELSTFSEIKGEKYTAVSTPGDLLFHIRAKQMGLCFEF
ASILDEKLKGAVVSVDETHGFRYMDGKAIIGFVDGTENPAVDENPYHFAVIGEEDADFAGGS
YVFVQKYIHDMVAWNALPVEQQEKVIGRHKFNDVELSDEEKPGNAHNAVTNIGDDLKIVRAN
MPFANTSKGEYGTYFIGYASTFSTTRRMLENMFIGSPAGNTDRLLDFSTAITGTLFFVPSYD
LLGELGE
111111110000000000000000000000000000000000000000000000000000
000000000000000000000000000000000000000000000000000000000000
000000000000000000000000000000000000000000000000000000000000
000000000000000000000000000000000000000000000000000000000000
000000000000000000000000000000000000000000000000000000000000
0000000
>DM_train751
MIKLGIVMDPIANINIKKDSSFAMLLEAQRRGYELHYMEMGDLYLINGEARAHTRTLNVKQN
YEEWFSFVGEQDLPLADLDVILMRKDPPFDTEFIYATYILERAEEKGTLIVNKPQSLRDCNE
KLFTAWFSDLTPETLVTRNKAQLKAFWEKHSDIILKPLDGMGGASIFRVKEGDPNLGVIAET
LTEHGTRYCMAQNYLPAIKDGDKRVLVVDGEPVPYCLARIPQGGETRGNLAAGGRGEPRPLT
ESDWKIARQIGPTLKEKGLIFVGLDIIGDRLTEINVTSPTCIREIEAEFPVSITGMLMDAIE
ARLQQQ
000000000000000000000000000000000000000000000000000000000000
000000000000000000000000000000000000000000000000000000000000
000000000000000000000000000000000000011110000000000000000000
000000000000000000000000000000000000011111111111111110000000
000000000000000000000000000000000000000000000000000000000000
000000
>DM_train752
MPFTIDSARGIFPNTLAADVVPATIARFSQLNAEDQLALIWFAYLEMGKTLTIAAPGAASMQ
LAENALKEIQAMGPLQQTQAMCDLANRADTPLCRTYASWSPNIKLGFWYRLGELMEQGFVAP
IPAGYQLSANANAVLATIQGLESGQQITVLRNAVVDMGFTAGKDGKRIAEPVVPPQDTASRT
KVSIEGVTNATVLNYMDNLNANDFDTLIELFTSDGALQPPFQRPIVGKENVLRFFREECQNL
KLIPERGVTEPAEDGFTQIKVTGKVQTPWFGGNVGMNIAWRFLLNPEGKIFFVAIDLLASPK
ELLNFA
111100000000000000000000000000000000000000000000000000000000
000000000000000000000000000000000000000000000000000000000000
000000000000000000000000000000000000011111111000000000000000
000000000000000000000000000000000000000000000000000000000000
000000000000000000000000000000000000000000000000000000000000
011111

>DM_train753
GSACAEFSFHVPSLEELAGVMQKGLKDNFADVQVSVVDCPDLTKEPFTFPVKGICGKTRIAE
VGGVPYLLPLVNQKKVYDLNKIAKEIKLPGAFILGAGAGPFQTLGFNSEFMPVIQTESEHKP
PVNGSYFAHVNPADGGCLLEKYSEKCHDFQCALLANLFASEGQPGKVIEVKAKRRTGPLNFV
TCMRETLEKHYGNKPIGMGGTFIIQKGKVKSHIMPAEFSSCPLNSDEEVNKWLHFYEMKAPL
VCLPVFVSRDPGFDLRLEHTHFFSRHGEGGHYHYDTTPDIVEYLGYFLPAEFLYRIDQPKET
HSIGRD
111000000000000000000000000000000000000000000000000000000000
000000000000000000000000000000000000000000000000000000000000
000000000000000000000000000000000000000000000000000000000000
000000000000000000000000000000000000000000000000000000000000
000000000000000000000000000000000000000000000000000000000000
000000
>DM_train754
MDSGRLAVIALGGNAIAGPGMDVSVESQTAAVKRASSIIADVLADGWRSVITHGNGPQVGYL
SEAFEALPPERPRQPLYIATAMTQAWIGLLLKHSLEEELRRRGLNVLVPVVISRVLVDVSDP
SFNNPSKPVGPIYGREEAEELSRRYGWVFKRDPRGGFRRVVPSPRPVSIVDRDLIAEASAES
PAVVALGGGGVPVVERPGGVLEPVEAVVDKDLASSLLATQLNADLLVILTDVPGVAVNYGRE
GERWLRRAAASELKKYLREGHFPPGSMGPKVEAAISFVERTGKPAVIGSLEEARQVLSLQAG
TVVMLG
111100000000000000000000000000000000000000000000000000000000
000000000000000000000000000000000000000000000000000000000000
000000000000000000000000000000000000000000000000000000000000
000000000000000000000000000000000000000000000000000000000000
000000000000000000000000000000000000000000000000000000000000
000000
>DM_train755
NAVLTWNNEILPNWETMWCSRKVRDLWWQGIPPSVRGKVWSLAIGNELNITHELFDICLARA
KERWRSLSTGGSEVENEDAGFSAADREASLELIKLDISRTFPNLCIFQQGGPYHDMLHSILG
AYTCYRPDVGYVQGMSFIAAVLILNLDTADAFIAFSNLLNKPCQMAFFRVDHGLMLTYFAAF
EVFFEENLPKLFAHFKKNNLTPDIYLIDWIFTLYSKSLPLDLACRIWDVFCRDGEEFLFRTA
LGILKLFEDILTKMDFIHMAQFLTRLPEDLPAEELFASIATIQMQSRNKKWAQVLTALQKDS
REMEKG
000000000000000110000000000000000000000000000000000000000000
000011111111111111111111100000000000000000000000000000000000
000000000000000000000000000000000000000000000000000000000000
000000000000000000000000000000000000000000000000000000000000
000000000000000000000000000000000000000000000000000000000011
111111
>DM_train756
MKQVKAAFEANKRVYESVLLTFKGVDGYDVYNCSVPFSYKGKTHIYGRVEKRDIWAASHVRL
FEETGKDEFTAVPELSWELEDPYIAKINNEMIFGGTRVRKNGNAILSYYGYFYRGTPDELTY
FTRGPGCMKDIRVLQLQDGRLGVFSRPRVGRKASIGFVILNSIDELGAEVIAKAPPLDILSE
NAWGGVNQAYLLSSGKVGCIGHYSYEDTNEKQQPQSVYVNYAFVLDPQSRAITGAKIIGTKS

CYPPCEPKVPLLADCVFASGIVMRSDGKVDLYSGVGDSHEGRITIDYPFKGHGTIIGDLHFP
MASSL
100000000000000000000000000000000000000000000000000000000000
000000000000000000000000000000000000000111110000000000000000
000000000000000000000000000001110000000000000000000000000011
000000000000000000000000001111111100000000000000000000000000
000000000000000000000000000000000000000000000000000000000000
01111
>DM_train757
GRVYYINSHGTLSRHENTLRFENAEVKKDIPVEDVEEIFVFAELSLNTKLLNFLASKGIPLH
FFNYYGYYTGTFYPRESSVSGHLLIKQVEHYLDAQKRLYLAKSFVIGSILNLEYVYKISADT
YLNKVKETNSIPELMSVEAEFRKLCYKKLEEVTGWELEKRTKRPPQNPLNALISFGNSLTYA
KVLGEIYKTQLNPTVSYLHEPSTKRFSLSLDVAEVFKPIFVDNLIIRLIQENKIDKTHFSTE
LNMTFLNEIGRKVFLKAFNELLETTIFYPKLNRKVSHRTLIKLELYKLIKHLLEEEVYLPLN
YGGLK
000000000000000000000000000000000000000000000000000000000000
000000000000000000000000000000000000000000000000000000000000
000000000000000000000000000000000000001111100000000000000000
000000000000000000000011000000000000000000000000000000000000
000000000000000000000000000000000000000000000000000000000000
00000
>DM_train758
MTRTRSGSLAAGGLNWASLPLKLFAGGNAKFWHPADIDFTRDRADWEKLSDDERDYATRLCT
QFIAGEEAVTEDIQPFMSAMRAEGRLADEMYLTQFAFEEAKHTQVFRMWLDAVGISEDLHRY
LDDLPAYRQIFYAELPECLNALSADPSPAAQVRASVTYNHIVEGMLALTGYYAWHKICVERA
ILPGMQELVRRIGDDERRHMAWGTFTCRRHVAADDANWTVFETRMNELIPLALRLIEEGFAL
YGDQPPFDLSKDDFLQYSTDKGMRRFGTISNARGRPVAEIDVDYSPAQLEDTFADEDRRTLA
AASA
100000000000000000000000000000000000000000000000000000000000
000000000000000000000000000000000000000000000000000000000000
000000000000000000000000000000000000000000000000000000000000
000000000000000000000000000000000000000000000000000000000000
000000000000000000000000000000000000000000111111111111111111
1111
>DM_train759
MARKKNTSDQSRSQAANTPIAGTYEGEYSVIELEADSYTTDGWLISINGVPSSHIVLGQPQA
LEFEYMRWIATGARAFIDAHQDASKLRITHLGGGACTMARYFADVYPQSRNTVVELDAELAR
LSREWFDIPRAPRVKIRVDDARMVAESFTPASRDVIIRDVFAGAITPQNFTTVEFFEHCHRG
LAPGGLYVANCGDHSDLRGAKSELAGMMEVFEHVAVIADPPMLKGRRYGNIILMGSDTEFFS
SNSTEASAITRELLGGGVPAQYKDESWVRKFASGAQARHDGVSTLQMPSDTPQHPAETPEHS
NTQP
111111111111111110000000000000000000000000000000000000000000
000000000000000000000000000000000000000000000000000000000000
000000000000000000000000000000000000000000000000000000000000

000000000000000000000000000000000000000000000000000000000000
000000000000000000000000000000000000000000000001111111111111
1111
>DM_train760
GGNSKSRNDRTETVDKEVIKAPEFDADSAYQYIQVQADFGPRVPNTQAHKECGEYLAGQLEK
FGAKVYNQYADLIAYDGTILKSRNIIGAYKPESKKRILLCAHWDSRPYADNDPDPKNHHTPI
LGVNDGASGVGVLLEIARQIQKEQPALGIDIVFFDSEDYGIPEFYDGKYKQDTWCLGSQYWA
RTPHVQNYNARYGILLDMVGGKDATFYYEGYSARTARSEMKKIWKKAHELGYGKYFVKEDGG
ETVDDHIYVNKLARIPCVDIINYDAGNPQSSFGSFWHTVNDTMENIDRNTLKAVGQTVMDVI
YNEK
111111111111111100000000000000000000000000000000000000000000
000000000000000000000000000000000000000000000001111111111111
111110000000000000000000000000000000001111111111110000000000
000000000000000000000000000000000000000000000000000000000000
000000000000000000000000011111111111111111100000000000000000
0000
>DM_train761
MLDNVLRIATRQSPLALWQAHYVKDKLMASHPGLVVELVPMVTRGDVILDTPLAKVGGKGLF
VKELEVALLENRADIAVHSMKDVPVEFPQGLGLVTICEREDPRDAFVSNNYDSLDALPAGSI
VGTSSLRRQCQLAERRPDLIIRSLRGNVGTRLSKLDNGEYDAIILAVAGLKRLGLESRIRAA
LPPEISLPAVGQGAVGIECRLDDSRTRELLAALNHHETALRVTAERAMNTRLEGGCQVPIGS
YAELIDGEIWLRALVGAPDGSQIIRGERRGAPQDAEQMGISLAEELLNNGAREILAEVYNGD
APA
000000000000000000000000000000000000000001111111111111111000
000000000000000000000000000000000000000000000000000000000000
000000000000000000000000000000000000000000000000000000000000
000000000000000000000000000000000000000000000000000000000000
000000000000000000000000000000000000000000000000000000000000
000
>DM_train762
MERIDSHVSPRYAQIPTFMRLPHDPQPRGYDVVVIGAPYDGGTSYRPGARFGPQAIRSESGL
IHGVGIDRGPGTFDLINCVDAGDINLTPFDMNIAIDTAQSHLSGLLKANAAFLMIGGDHSLT
VAALRAVAEQHGPLAVVHLDAHSDTNPAFYGGRYHHGTPFRHGIDEKLIDPAAMVQIGIRGH
NPKPDSLDYARGHGVRVVTADEFGELGVGGTADLIREKVGQRPVYVSVDIDVVDPAFAPGTG
TPAPGGLLSREVLALLRCVGDLKPVGFDVMEVSPLYDHGGITSILATEIGAELLYQYARAHR
TQL
111111110000000000000000000000000000000000000000000000000000
000000011100000000000000000000000000000000000000000000000000
000000000000000000000000000000000000000000000000000000000000
111111000000000000000000000000000000000000000000000000000000
000000000000000000000000000000000000000000000000000000000001
111
>DM_train763
GSHMSKKFALTAEQRASFEKNGFIGPFDAYSPEEMKETWKRTRLRLLDRSAAAYQDLDAISG

GTNIANYDRHLDDDFLASHICRPEICDRVESILGPNVLCWRTEFFPKYPGDEGTDWHQADTF
ANASGKPQIIWPENEEFGGTITVWTAFTDANIANGCLQFIPGTQNSMNYDETKRMTYEPDAN
NSVVKDGVRRGFFGYDYRQLQIDENWKPDEASAVPMQMKAGQFIIFWSTLMHASYPHSGESQ
EMRMGFASRYVPSFVHVYPDSDHIEEYGGRISLEKYGAVQVIGDETPEYNRLVTHTTRGKKF
EAV
11111100000000000000000000000000000000000000000000000000111111
11000000000000000000000000000000000000000000000000000000000000
00000000000000000000000000000000000000000000000000000000000000
00000000000000000000000000000000000000000000000000000000000000
00000000000000000000000000000000000000000000000000000000000000
000
>DM_train764
MNQKLKVAIIGSGNIGTDLMIKVLRNAKYLEMGAMVGIDAASDGLARAQRMGVTTTYAGVEG
LIKLPEFADIDFVFDATSASAHVQNEALLRQAKPGIRLIDLTPAAIGPYCVPVVNLEEHLGK
LNVNMVTCGGQATIPMVAAVSRVAKVHYAEIVASISSKSAGPGTRANIDEFTETTSKAIEVI
GGAAKGKAIIIMNPAEPPLIMRDTVYVLSAAADQAAVAASVAEMVQAVQAYVPGYRLKQQVQ
FDVIPESAPLNIPGLGRFSGLKTSVFLEVEGAAHYLPAYAGNLDIMTSAALATAERMAQSML
NA
11100000000000000000000000000000000000000000000000000000000000
00000000000000000000000000000000000000000000000000000000000000
00000000000000000000000000000000000000000000000000000000000000
00000000000000000000000000000000000000000000000000000000000000
00000000000000000000000000000000000000000000000000000000000000
00
>DM_train766
MSDLMQKENIVVMLTKKGFLKRLSQNEYKLQGTGGKGLSSFDLNDGDEIVIALCVNTHDYLF
MISNEGKLYLINAYEIKDSSRASKGQNISELINLGDQEEILTIKNSKDLTDDAYLLLTTASG
KIARFESTDFKAVKSRGVIVIKLNDKDFVTSAEIVFKDEKVICLSKKGSAFIFNSRDVRLTN
RGTQGVCGMKLKEGDLFVKVLSVKENPYLLIVSENGYGKRLNMSKISELKRGATGYTSYKKS
DKKAGSVVDAIAVSEDDEILLVSKRSKALRTVAGKVSEQGKDARGIQVLFLDNDSLVSVSKF
IK
11111110000000000000000000000000000000000000000000000000000000
00000000000000000011111110000000000000000000000000000000000000
00000000000011110000000000000000000000000000000000000000000000
00000000000000000000000000000000000000000000000000000000000000
00000000000000000000000000000000000000000000000000000000000000
01
>DM_train767
HMYRNVPIWAQKWKPTIKALQSINVKDLKIDPSFLNIIPDDDLTKSVQDWVYATIYSIAPEL
RSFIELEMKFGVIIDAKGPDRVNPPVSSQCVFTELDAHLTPNIDASLFKELSKYIRGISEVT
ENTGKFSIIESQTRDSVYRVGLSTQRPRFLRMSTDIKTGRVGQFIEKRHVAQLLLYSPKDSY
DVKISLNLELPVPDNDPPEKYKSQSPISERTKDRVSYIHNDSCTRIDITKVENHNQNSKSRQ
SETTHEVELEINTPALLNAFDNITNDSKEYASLIRTFLNNGTIIRRKLSSLSYEIFEGSKKV
M

00000000000000000000000000000000000000000000000000000000000000
00000000000000000000000000000000000000000000000000000000000000
00000000000000000000011111000000000000000000000000000000000000
00000000000000000000000000000000000000000000000000000011111111
00000000000000000000000000000000000000000000000000000111111111
1
>DM_train768
MDTTVPTFSLAELQQGLHQDEFRRCLRDKGLFYLTDCGLTDTELKSAKDIVIDFFEHGSEAE
KRAVTSPVPTMRRGFTGLESESTAQITNTGSYSDYSMCYSMGTADNLFPSGDFERIWTQYFD
RQYTASRAVAREVLRATGTEPDGGVEAFLDCEPLLRFRYFPQVPEHRSAEEQPLRMAPHYDL
SMVTLIQQTPCANGFVSLQAEVGGAFTDLPYRPDAVLVFCGAIATLVTGGQVKAPRHHVAAP
RRDQIAGSSQTSSVFFLRPNADFTFSVPLARECGFDVSLDGETATFQDWIGGNYVNIRRTSK
A
00000000000000000000000000000000000000000000000000000000000000
00000000000000000011111111111111111000000000000000000000000000
00000000000000000000000000000000000000000111111111111000000000
00000000011111000000000000000000000000000000000000000000000001
11111111000000000000000000000000000000000000000000000000000111
1
>DM_train769
GQDYSGFTLTPSAQSPRLLELTFTEQTTKQFLEQVAEWPVQALEYKSFLRFRVAKILDDLCA
NQLQPLLLKTLLNRAEGALLINAVGVDDVKQADEMVKLATAVAHLIGRSNFDAMSGQYYARF
VVKNVDNSDSYLRQPHRVMELHNDGTYVEEITDYVLMMKIDEQNMQGGNSLLLHLDDWEHLD
NYFRHPLARRPMRFAAPPSKNVSKDVFHPVFDVDQQGRPVMRYIDQFVQPKDFEEGVWLSEL
SDAIETSKGILSVPVPVGKFLLINNLFWLHGRDRFTPHPDLRRELMRQRGYFAYASNHYQTH
Q
00000000000000000000000000000000000000000000000000000000000000
00000000000000000000000000000000000000000000000000000000000000
00000111110000000000000000000000000000000000000000000000000000
00000000000000000000000000000000000000000000000000000000000000
00000000000000000000000000000000000000000000000000000000000000
0
>DM_train770
AAASMDAVTAEKFSPASFLDKKETGVLHFVKYHGLGNDFILVDNRDSSEPKITQEQAAKLCD
RNFGVGADGVIFAMPGVNGTDYAMRIFNSDGSEPEMCGNGVRCFARFIAELENLQGKHSFTI
HTGAGLIVPEIQDDGQVKVDMGTPILKAQDVPTKLSGNKGEAVVEAELVVDGVSWNVTCVSM
GNPHCITFGKKGGPNLKVDDLNLPEIGPKFEHHEMFPARTNTEFVEVLSRSHLKMRVWERGA
GATLACGTGACALVVAAVLEGRADRKCTVDLPGGPLEIEWKQEDNHIYMTGPAEAVFYGSAL
L
11111111111111111111111100000000000000000000000000000000000000
00000000000000000000000000000000000000000000000000000000000000
00000000000000000000000000000000000000000000000000000000000000
00000000000000000000000000000000000000000000000000000000000000
00000000000000000000000000000000000000000000000000000000000000

0
>DM_train771
MANPLYHKHIISINDLSRDELELVLRTAASLKKTPQPELLKHKVIASCFFEASTRTRLSFET
SIHRLGASVVGFSDSSNTSLGKKGETLADTMSVISTYVDAIVMRHPQEGASRLAAQFSGNVP
IVNAGDGANQHPTQTLLDLFTIQETQGRLDNINIAMVGDLKYGRTVHSLTQALAKFNGNHFF
FIAPDALAMPAYILQMLEEKEIEYSLHESLEEVVPELDILYMTRVQKERLDPSEYANVKAQF
ILRSSDLTGARDNLKVLHPLPRIDEITTDVDKTPYAYYFQQAGNGIFARQALLALVLNAELA
L
1100000000000000000000000000000000000000000000000000000000000000
0000000000000001111111100000000000000000000000000000000000000000
0000000001100000000000000000000000000000000000000000000000000000
0000000000000000000000000000000000000000000001111111111111111111
0000000000000000000000000000000000000000000000000000000000000011
1
>DM_train772
EANAPGPVPGERQLAHSKMVPIPAGVFTMGTDDPQIKQDGEAPARRVTIDAFYMDAYEVSNT
EFEKFVNSTGYLTEAEKFGDSFVFEGMLSEQVKTNIQQAVAAAPWWLPVKGANWRHPEGPDS
TILHRPDHPVLHVSWNDAVAYCTWAGKRLPTEAEWEYSCRGGLHNRLFPWGNKLQPKGQHYA
NIWQGEFPVTNTGEDGFQGTAPVDAFPPNGYGLYNIVGNAWEWTSDWWTVHHSVEETLNPKG
PPSGKDRVKKGGSYMCHRSYCYRYRCAARSQNTPDSSASNLGFRCAADRLPTMDRGSHHHHH
H
1111111111111000000000000000000000000000000000000000000000000000
0000000000000000000000000000011111111110000000000000000000000000
0000000000000000000000000000000000000000000000000000000000000000
0000000000000000000000000000000000000000000000000000000000000000
0000000000000000000000000000000000000000000000000000011111111111
1
>DM_train773
GHMTSRPSSDQTWQPIDGRVALIAPASAIATDVLEATLRQLEVHGVDYHLGRHVEARYRYLA
GTVEQRLEDLHNAFDMPDITAVWCLRGGYGCGQLLPGLDWGRLQAASPRPLIGFSDISVLLS
AFHRHGLPAIHGPVATGLGLSPLSAPREQQERLASLASVSRLLAGIDHELPVQHLGGHKQRV
EGALIGGNLTALACMAGTLGGLHAPAGSILVLEDVGEPYYRLERSLWQLLESIDARQLGAIC
LGSFTDCPRKEVAHSLERIFGEYAAAIEVPLYHHLPSGHGAQNRAWPYGKTAVLEGNRLRWG
S
1111111100000000000000000000000000000000000000000000000000000000
0000000000000000000000000000000000000000000000000000000000000000
0000000000000000000000000000000000000000000000000000000000000000
0000000000000000000000000000000000000000000000000000000000000000
0000000000000000000000000000000000000000000000000000000000000000
0
>DM_train774
GPNANPIPEHFFAPYIDMSLSVHKPLVEYAKLTGTKYFTLAFILYSSVYNGPAWAGSIPLEK
FVDEVRELREIGGEVIIAFGGAVGPYLCQQASTPEQLAEWYIKVIDTYNATYLDFDIEAGID
ADKLADALLIVQRERPWVKFSFTLPSDPGIGLAGGYGIIETMAKKGVRVDRVNPMTMDYYWT

PSNAENAIKVAENVFRQLKQIYPEKSDEEIWKMIGLTPMIGVNDDKSVFTLEDAQQLVDWAI
QHKIGSLAFWSVDRDHPGPTGEVSPLHRGTNDPDWAFSHVFVKFMEAFGYTFSAQTSEASVP
T
00000000000000000000000000000000000000000000000000000000000000
00000000000000000000000000000000000000000000000000000000000000
00000000000000000000000000000000000000000000000000000000000000
00000000000000000000000000000000000000000000000000000000000000
00000000000000000000000000000000000000000000000000001111111111
1
>DM_train776
MRLKDLGERALLARLAPLGYPPEAPLPPGDDAGGVWAEGRAWLLKTDGFLYREVALKGMGPF
EVGFRGVAATASDLLAKMGRPLGFTLGLFLPEDLEEGFVLELVRGAAEAAKRLGAFLLGGDT
NRGVEVALTVSGYALAEAPLPRKALPGDLLYLAGDRWGRTGAAIRAHYEGRSLEGFPKIREA
AFYPLPRLELLALSGLLRGSLDSSDGLAETLWQLADLGVGVEVEALPLYPDVLAFAGSEEAA
LELVLYGGEEFEAVLVVPQEGAAAVEARAKAKGLPLFRAGRVVAGEGVYLRGAPLPRKGYAH
F
10000000000000000000000000000000000000000000000000000000000000
00000000000000000000000000000000000000000000000000000000000000
00000000000000000000000000000000000000000000000000000000000000
00000000000000000000000000000000000000000000000000000000000000
00000000000000000000000000000000000000000000000000000000011111
1
>DM_train777
MSRKKMGLLVMAYGTPYKEEDIERYYTHIRRGRKPEPEMLQDLKDRYEAIGGISPLAQITEQ
QAHNLEQHLNEIQDEITFKAYIGLKHIEPFIEDAVAEMHKDGITEAVSIVLAPHFSTFSVQS
YNKRAKEEAEKLGGLTITSVESWYDEPKFVTYWVDRVKETYASMPEDERENAMLIVSAHSLP
EKIKEFGDPYPDQLHESAKLIAEGAGVSEYAVGWQSEGNTPDPWLGPDVQDLTRDLFEQKGY
QAFVYVPVGFVADHLEVLYDNDYECKVVTDDIGASYYRPEMPNAKPEFIDALATVVLKKLGR
11110000000000000000000000000000000000000000000000000000000000
00000000000000000000000000000000000000000000000000000000000000
00000000000000000000000000000000000000000000000000000000000000
00000000000000000000000000000000000000000000000000000000000000
00000000000000000000000000000000000000000000000000000000000000
>DM_train778
GMQKQHNVVAGVDMGATHIRFCLRTAEGETLHCEKKRTAEVIAPGLVSGIGEMIDEQLRRFN
ARCHGLVMGFPALVSKDKRTIISTPNLPLTAADLYDLADKLENTLNCPVEFSRDVNLQLSWD
VVENRLTQQLVLAAYLGTGMGFAVWMNGAPWTGAHGVAGELGHIPLGDMTQHCACGNPGCLE
TNCSGMALRRWYEQQPRNYPLRDLFVHAENAPFVQSLLENAARAIATSINLFDPDAVILGGG
VMDMPAFPRETLVAMTQKYLRRPLPHQVVRFIAASSSDFNGAQGAAILAHQRFLPQFCAKAP
11100000000000000000000000000000000000000000000000000000000000
00000000000000000000000000000000000000000000000000000000000000
00000000000000000000000000000000000000001111111111111111111110
00000000000000000000000000000000000000000000000000000000000000
00000000000000000000000000000000000000000000000000000001111111

>DM_train779
MLDANKLQQAVDQAYTQFHSLNGGQNADYIPFLANVPGQLAAVAIVTCDGNVYSAGDSDYRF
ALESISKVCTLALALEDVGPQAVQDKIGADPTGLPFNSVIALELHGGKPLSPLVNAGAIATT
SLINAENVEQRWQRILHIQQQLAGEQVALSDEVNQSEQTTNFHNRAIAWLLYSAGYLYCDAM
EACDVYTRQCSTLLNTIELATLGATLAAGGVNPLTHKRVLQADNVPYILAEMMMEGLYGRSG
DWAYRVGLPGKSGVGGGILAVVPGVMGIAAFSPPLDEDGNSVRGQKMVASVAKQLGYNVFKG
100000000000000000000000000000000000000000000000000000000000
000000000000000000000000000000000000000000000000000000000000
000000000000000000000000000000000000000000000000000000000000
000000000000000000000000000000000000000000000000000000000000
000000000000000000000000000000000000000000000000000000000000
>DM_train780
MAVEVLKEKWNSKVVEVTLGTGDKTVTLGGDSTLPFLTFEGEMPNPPRFALEVFDTPPTDWP
DILVEPFKDVINDPVAWAKKCVEYGADIVALRLVSAHPDGQNRSGAELAEVCKAVADAIDVP
LMIIGCGVEEKDAEIFPVIGEALSGRNCLLSSATKDNYKPIVATCMVHGHSVVASAPLDINL
SKQLNIMIMEMNLAPNRIIMDPLIGALGYGIEYSYSIIERMRLGALTGDKILAMPVVCFIGQ
EAWKAKEAKDPEVAEWGDYALRAIHWETVTTVALIQAGGHLFVMRHPKSLAEVKEHLKRILK
110000000000000000000000000000000000000000000000000000000000
000000000000000000000000000000000000000000000000000000000000
000000000000000000000000000000000000000000000000000000000000
000000000000000000000000000000000000000000000000000000000000
000000000000000000000000000000000000000000000000000000000001
>DM_train781
GPLGSYGDAIPEVKAILEAKNEEELVTFTSRWSAEERKELRTQFQDTTGLEFIAFLKKCIKN
GPYEDVMALGWDCNISARVNVIKKAMKNVNDFRAIHDVVLIATPDERLKLAQAYKEKTGNDL
LQDFVDQIPLTSAASYLCHLAIRENRTPRGSVASDAEVLKHNLIDADEPDHEAVVRLIITST
ADEYKEINHRFEVLTGKSVQEAIETRYADKENARGLCIAHYYNLAPARAVAYAFHSAVETQN
DDMAYEQAARITGLFHDLHKFAWVHYACWGVMRDDILSRFQSKEANKVNFRDACLMFWKLAK
111110000000000000000000000000000000000000000000000000000000
000000000000000000000000000000000000000000000000000000000000
000000000000000000000000000000000000000000000000000000000000
000000000000000000000000000000000000000000000000000000000000
000000000000000000000000000000000000000000000000000000000001
>DM_train782
MRTHDDTWDIKTSVGTTAVMVAAARAAETDRPDALIRDPYAKLLVTNTGAGALWEAMLDPSM
VAKVEAIDAEAAAMVEHMRSYQAVRTNFFDTYFNNAVIDGIRQFVILASGLDSRAYRLDWPT
GTTVYEIDQPKVLAYKSTTLAEHGVTPTADRREVPIDLRQDWPPALRSAGFDPSARTAWLAE
GLLMYLPATAQDGLFTEIGGLSAVGSRIAVETSPLHGDEWREQMQLRFRRVSDALGFEQAVD
VQELIYHDENRAVVADWLNRHGWRATAQSAPDEMRRVGRWGDGVPMADDKDAFAEFVTAHRL
111111111111100000000000000000000000000000000000000000011111
111111100000000000000000000000000000000000000000000000000000
000000000000000000000000000000000000000000000000000000000000
000000000000000000000000000000000000000000000000000011111111
110000000000000000000000000000000000000000000000000000000000

>DM_train783
MARYDLVDRLNTTFRQMEQELAIFAAHLEQHKLLVARVFSLPEVKKEDEHNPLNRIEVKQHL
GNDAQSLALRHFRHLFIQQQSENRSSKAAVRLPGVLCYQVDNLSQAALVSHIQHINKLKTTF
EHIVTVESELPTAARFEWVHRHLPGLITLNAYRTLTVLHDPATLRFGWANKHIIKNLHRDEV
LAQLEKSLKSPRSVAPWTREEWQRKLEREYQDIAALPQNAKLKIKRPVKVQPIARVWYKGDQ
KQVQHACPTPLIALINRDNGAGVPDVGELLNYDADNVQHRYKPQAQPLRLIIPRLHLYVAD
111100000000000000000000000000000000000000000000000000000000
000000000000000000000000000000000000000000000000000000000000
000000000000000000000000000000000000000000000000000000000000
000000000000000000000000000000000000000000000000000000000000
00000000000000000000000000000000000000000000000000000000000
>DM_train784
ASSSYYEQYHSLNEIYSWIEVMTERYPDMVEKIHIGSSYEKYPLYVLKVSKKEQRAKNAMWI
DCGIHAREWISPAFCLWFVGSVTYYYGKEKMHTNLLKHMDFYIMPVVNVDGYDYTWKKDRMW
RKNRSLHEKNACVGTDLNRNFASKHWCGEGASSSSCSEIYCGTYPESEPEVKAVADFLRRNI
KHIKAYISMHSYSQKIVFPYSYSRSRSKDHEELSLVAREAVFAMENIHRNIRYTHGSGSESL
YLAPGGSDDWIYDLGIKYSFTFELRDKGKYGFLLPESYIRPTCSEALVAVAKIASHVVKNV
111000000000000000000000000000000000000000000000011111000000
000000000000000000000000000111110000000000000000000000000000
000000000000000000000000000000000000000000000000000000000000
000000000000000000000000000000000000000000000000000000000000
00000000000000000000000000000000000000000000000000000000000
>DM_train785
NILVIGTGAIGSFYGALLAKTGHCVSVVSRSDYETVKAKGIRIRSATLGDYTFRPAAVVRSA
AELETKPDCTLLCIKVVEGADRVGLLRDAVAPDTGIVLISNGIDIEPEVAAAFPDNEVISGL
AFIGVTRTAPGEIWHQAYGRLMLGNYPGGVSERVKTLAAAFEEAGIDGIATENITTARWQKC
VWNAAFNPLSVLSGGLDTLDILSTQEGFVRAIMQEIRAVAAANGHPLPEDIVEKNVASTYKM
PPYKTSMLVDFEAGQPMETEVILGNAVRAGRRTRVAIPHLESVYALMKLLELRTSKSLWGN
000000000000000000000000000000000000000000000000000000000011
111111100000011111110000000001110000000000000000000011000000
000000000000000000000000000000000000000000000000000000000000
000000000000000000000000000000000000000000000000000000000000
00000000000000000000000000000000000000000000000000000000001
>DM_train786
CLGTIGPVTPLDASDFALDIRMPGVTPKESDTYFCMSMRLPVDEEAFVIDFKPRASMDTVHH
MLLFGCNMPSSTGSYWFCDEGTCTDKANILYAWARNAPPTRLPKGVGFRVGGETGSKYFVLQ
VHYGDISAFRDNHKDCSGVSVHLTRVPQPLIAGMYLMMSVDTVIPPGEKVVNADISCQYKMY
PMHVFAYRVHTHHLGKVVSGYRVRNGQWTLIGRQNPQLPQAFYPVEHPVDVTFGDILAARCV
FTGEGRTEATHIGGTSSDEICNLYIMYYMEAKYALSFMTCTKNVAPDMFRTIPAEANIPIP
000000000000000000000000000000000000000000000000000000000000
000000000000000000000000000000000000000000000000000000000000
000000000000000000000000000000000000000000000000000000000000
000000000000000000000000000000000000000000000000000000000000
00001111111111111100000000000000000000000000000000000000000

>DM_train787
GSRPGPASQLPRFVRVNTLKTCSDDVVDYFKRQGFSYQGRASSLDDLRALKGKHFLLDPLMP
ELLVFPAQTDLHEHPLYRAGHLILQDRASCLPAMLLDPPPGSHVIDACAAPGNKTSHLAALL
KNQGKIFAFDLDAKRLASMATLLARAGVSCCELAEEDFLAVSPSDPRYHEVHYILLDPSCSG
SGMPSRQLEEPGAGTPSPVRLHALAGFQQRALCHALTFPSLQRLVYSTCSLCQEENEDVVRD
ALQQNPGAFRLAPALPAWPHRGLSTFPGAEHCLRASPETTLSSGFFVAVIERVEVPRRARG
111111110000000000000000000000000000000000000000000000000000
000000000000000000000000000000000000000000000000000000000000
000000000000000000000000000000000000000000000000000000000000
111111111111111111000000000000000000000000000000000000000000
00000000000000000000000000000000000000000000000000011111111
>DM_train788
MEKILIFGHQNPDTDTICSAIAYADLKNKLGFNAEPVRLGQVNGETQYALDYFKQESPRLVE
TAANEVNGVILVDHNERQQSIKDIEEVQVLEVIDHHRIANFETAEPLYYRAEPVGCTATILN
KMYKENNVKIEKEIAGLMLSAIISDSLLFKSPTCTDQDVAAAKELAEIAGVDAEEYGLNMLK
AGADLSKKTVEELISLDAKEFTLGSKKVEIAQVNTVDIEDVKKRQAELEAVISKVVAEKNLD
LFLLVITDILENDSLALAIGNEAAKVEKAFNVTLENNTALLKGVVSRKKQVVPVLTDAMAE
100000000000000000000000000000000000000000000000000000000000
000000000000000000000000000000000000000000000000000000000000
000000000000000000000000000000000000000000000000000000000000
000000000000000000000000000000000000000000000000000000000000
00000000000000000000000000000000000000000000000000000000001
>DM_train789
MGSSHHHHHHSSGENLYFQGHMKHGIYYSYWEHEWSAKFGPYIEKVAKLGFDIIEVAAHHINE
YSDAELATIRKSAKDNGIILTAGIGPSKTKNLSSEDAAVRAAGKAFFERTLSNVAKLDIHTI
GGALHSYWPIDYSQPVDKAGDYARGVEGINGIADFANDLGINLCIEVLNRFENHVLNTAAEG
VAFVKDVGKNNVKVMLDTFHMNIEEDSFGDAIRTAGPLLGHFHTGESNRRVPGKGRMPWHEI
GLALRDINYTGAVIMEPFVKTGGTIGSDIKVWRDLSGGADIAKMDEDARNALAFSRFVLGG
111111111111111111110000000000000000000000000000000000000000
000000000000000000000000000000000000000000000000000000000000
000000000000000000000000000000000000000000000000000000000000
000000000000000000000000000000000000000000000000000000000000
00000000000000000000000000000000000000000000000000000000000
>DM_train790
GSSGFKKLVSPSSAVEKCIVSVSYRGNNLNGLWLGDSIYCPRHVLGKFSGDQWGDVLNLANN
HEFEVVTQNGVTLNVVSRRLKGAVLILQTAVANAETPKYKFVKANCGDSFTIACSYGGTVIG
LYPVTMRSNGTIRASFLAGACGSVGFNIEKGVVNFFYMHHLELPNALHTGTDLMGEFYGGYV
DEEVAQRVPPDNLVTNNIVAWLYAAIISVKESSFSQPKWLESTTVSIEDYNRWASDNGFTPF
STSTAITKLSAITGVDVCKLLRTIMVKSAQWGSDPILGQYNFEDELTPESVFNQVGGVRLQ
110000000000000000000000000000000000000000000000000000000000
000000000000000000000000000000000000000000000000000000000000
000000000000000000000000000000000000000000000000000000000000
000000000000000000000000000001111110000000000000000000000000
00000000000000000000000000000000000000000000000000000111111

>DM_train792
MSADYPRDLIGYGNNPPHPHWPGDARIALSFVLNYEEGGERCVLHGDKESEAFLSEMVAAQP
LQGVRHMSMESLYEYGSRAGVWRLLKLFKRRNVPLTVFAVAMAAQRNPEVIRAMVADGHEIC
SHGYRWIDYQYMDEAQEREHMLEAIRILTELTGQRPVGWYTGRTGPNTRRLVMEEGGFLYDS
DTYDDDLPYWDPASTAEKPHLVIPYTLDTNDMRFTQVQGFNNGEQFFQYLKDAFDVLYEEGA
TAPKMLSIGLHCRLIGRPARMAALERFIQYAQSHDKVWFARREDIARHWHREHPFQETEA
11110000000000000000000000000000000000000000000000000000000000
00000000000000000000000000000000000000000000000000000000000000
00000000000000000000000000000000000000000000000000000000000000
00000000000000000000000000000000000000000000000000000000000000
000000000000000000000000000000000000000000000000000000001111
>DM_train793
MNVVIVRYGEIGTKSRQTRSWFEKILMNNIREALVTEEVPYKEIFSRHGRIIVKTNSPKEAA
NVLVRVFGIVSISPAMEVEASLEKINRTALLMFRKKAKEVGKERPKFRVTARRITKEFPLDS
LEIQAKVGEYILNNENCEVDLKNYDIEIGIEIMQGKAYIYTEKIKGWGGLPIGTEGRMIGIL
HDELSALAIFLMMKRGVEVIPVYIGKDDKNLEKVRSLWNLLKRYSYGSKGFLVVAESFDRVL
KLIRDFGVKGVIKGLRPNDLNSEVSEITEDFKMFPVPVYYPLIALPEEYIKSVKERLGL
00000000111110000000000000000000000000000000000000000000000000
00000000000000000000000000000000000000000000000000000000000000
00000000000000000000000000000000000000000000000000000000000000
00000000000000000000000000000000000000000000000000000000000000
00000000000000000111111000000000000000000000000000000000000
>DM_train794
ESTSLYKKAGLKPQVYHVDAFTSQPFRGNSAGVVFPADNLSEAQMQLIARELGHSETAFLLH
SDDSDVRIRYFTPTVEVPICGHATVAAHYVRAKVLGLGNCTIWQTSLAGKHRVTIEKHNDDY
RISLEQGTPGFEPPLEGETRAAIINALHLTEDDILPGLPIQVATTGHSKVMIPLKPEVDIDA
LSPDLNALTAISKKIGCNGFFPFQIRPGKNETDGRMFSPAIGIVEDPVTGNANGPMGAWLVH
HNVLPHDGNVLRVKGHQGRALGRDGMIEVTVTIRDNQPEKVTISGTAVILFHAEWAIEL
11111111110000000000000000000000000000000000000000000000000000
00000000000000000001100000000000000000000000000110000000000000
00000000000000000000000000000000000000000000000000000000000000
00000000000000000000000000000000000000000000000000000000000000
00000000000000000000000000000000000000000000000000000000000
>DM_train795
MAPAEILNGKEISAQIRARLKNQVTQLKEQVPGFTPRLAILQVGNRDDSNLYINVKLKAAEE
IGIKATHIKLPRTTTESEVMKYITSLNEDSTVHGFLVQLPLDSENSINTEEVINAIAPEKDV
DGLTSINAGRLARGDLNDCFIPCTPKGCLELIKETGVPIAGRHAVVVGRSKIVGAPMHDLLL
WNNATVTTCHSKTAHLDEEVNKGDILVVATGQPEMVKGEWIKPGAIVIDCGINYVPDDKKPN
GRKVVGDVAYDEAKERASFITPVPGGVGPMTVAMLMQSTVESAKRFLEKFKPGKWMIQ
00000000000000000000000000000000000000000000000000000000000000
00000000000000000000000000000000000000000000000000000000000000
00000000000000000000000000000000000000000000000000000000000000
00000000000000000000000000000000000000000000000000000011111111
1100000000000000000000000000000000000000000000001111111111

>DM_train796
APADPVPTVFFGDSYTANFGIAPVTNQDSERGWCFQAKENYPAVATRSLADKGITLDVQADV
SCGGALIHHFWEKQELPFGAGELPPQQDALKQDTQLTVGSLGGNTLGFNRILKQCSDELRKP
SLLPGDPVDGDEPAAKCGEFFGTGDGKQWLDDQFERVGAELEELLDRIGYFAPDAKRVLVGY
PRLVPEDTTKCLTAAPGQTQLPFADIPQDALPVLDQIQKRLNDAMKKAAADGGADFVDLYAG
TGANTACDGADRGIGGLLEDSQLELLGTKIPWYAHPNDKGRDIQAKQVADKIEEILNR
111000000000000000000000000000000000000000000000000000000000
000000000000000000000000000000000000000000000000000000000000
000000000000000000000000000000000000000000000000000000000000
000000000000000000000000000000000000000000000000000000000000
0000000000000000000000000000000000000000000000000000000000
>DM_train797
IWELKKDVYVVELDWYPDAPGEMVVLTCDTPEEDGITWTLDQSSEVLGSGKTLTIQVKEFGD
AGQYTCHKGGEVLSHSLLLLHKKEDGIWSTDILKDQKEPKNKTFLRCEAKNYSGRFTCWWLT
TISTDLTFSVKSSRGSSDPQGVTCGAATLSAERVRGDNKEYEYSVECQEDSACPAAEESLPI
EVMVDAVHKLKYENYTSSFFIRDIIKPDPPKNLQLKPLKNSRQVEVSWEYPDTWSTPHSYFS
LTFCVQVQGKSKREKKDRVFTDKTSATVICRKNASISVRAQDRYYSSSWSEWASVPCS
000000000000000000000000000000000000000000000000000000000000
000000111111000000000000000000000000000000000000000000000000
000000000000000000000000000000011111100000000000000000000000
000000000000000000000000000000000000000000000000000000000000
0000000011111110000000000000001110000000000000000000000000
>DM_train798
GMGDSAGLPPIYADKPIELAPAKIITSFPVNTFLENLASAPDGTIFVTNHEVGEIVSITPDG
NQQIHATVEGKVSGLAFTSNGDLVATGWNADSIPVVSLVKSDGTVETLLTLPDAIFLNGITP
LSDTQYLTADSYRGAIWLIDVVQPSGSIWLEHPMLARSNSESVFPAANGLKRFGNFLYVSNT
EKMLLLRIPVDSTDKPGEPEIFVEQTNIDDFAFDVEGNLYGATHIYNSVVRIAPDRSTTIIA
QAEQGVIGSTAVAFGQTEGDCTAIYVVTNGGMFLPPPTGVVPANVVRLEVGKPGYPLG
111100000000000000000000000000000000000000000000000000000000
000000000000000000000000000000000000000000000000000000000000
000000000000000000000000000000000000000000000000000000000000
000000000000000000000000000000000000000000000000000000000000
0000000000000000000000000000000000000000000000000000000000
>DM_train799
MDLEMMLDEDYKEGICLIDFSQIALSTALVNFPDKEKINLSMVRHLILNSIKFNVKKAKTLG
YTKIVLCIDNAKSGYWRRDFAYYYKKNRGKAREESTWDWEGYFESSHKVIDELKAYMPYIVM
DIDKYEADDHIAVLVKKFSLEGHKILIISSDGDFTQLHKYPNVKQWSPMHKKWVKIKSGSAE
IDCMTKILKGDKKDNVASVKVRSDFWFTRVEGERTPSMKTSIVEAIANDREQAKVLLTESEY
NRYKENLVLIDFDYIPDNIASNIVNYYNSYKLPPRGKIYSYFVKAGLSKLTNSINEF
111111111110000000000000000000000000000000000000000000000000
000000000000000000000000011111111100000000000000000000000000
000000000000000000000000000000000000000000000000000000000000
000000000000000000000000000000000000000000000000000000000000
000000000000000000000000000000000000000000000000000000000

>DM_train800
MNNQCKTIAHVLRVNNGQELHVWETPPKENVPFKNNTILIASGFARRMDHFAGLAEYLSTNG
FHVFRYDSLHHVGLSSGSIDEFTMTTGKNSLCTVYHWLQTKGTQNIGLIAASLSARVAYEVI
SDLELSFLITAVGVVNLRDTLEKALGFDYLSLPIDELPNDLDFEGHKLGSEVFVRDCFEHHW
DTLDSTLDKVANTSVPLIAFTANNDDWVKQEEVYDMLAHIRTGHCKLYSLLGSSHDLGENLV
VLRNFYQSVTKAAIAMDGGSLEIDVDFIEPDFEQLTIATVNERRLKAEIENRTPEMA
11100000000000000000000000000000000000000000000000000000000000
00000000000111111111000000000000000000000000000000000000000000
00000000000000000000000000000000000000000000000000000000000000
00000000000000000000000000000000000000000000000000000000000000
000000000000000000000000000000000000000000000000000000000
>DM_train801
GMDISGRDVETRLTLAREFMSGVDELPTVPDIVLRIAGKLNDPDVAIDEVADLLLQDQVLTA
RVVHLANSPLYSAARPISSIRDAVIYLGLDLLREAIFTCAIVDLFKTGKGPLNRSTLWAHSL
GVARIAKLIAERTGFLNPVNVYVAGLLHDVGEVFINFFRGKEFSQVVTLVDEEKITFGQAEE
RLFGTSHCEVGFALAKRWSLNEFICDTILYHHDIEAVPYKQAAIVAMVAFADEYCTLRRLGF
EGHKPVDSVRTLLENHPSWGVIRRSLGGSDFDEKLIVAELDSSIVEIRAAVDELFLL
11111110000000000000000000000000000000000000000000000000000000
00000000000000000000000000000000000000000000000000000000000000
00000000000000000000000000000000000000000000000000000000000000
00000000000000000000000000000000000000000000000000000000000000
000000000000000000000000000000000000000000000000000000000
>DM_train802
MQRLSPGEFKTLISKERKSHFITPFALVYKTFCDLGYDQKNSDYFLNNPSEYIIAMRKNCWK
EFEPFEKEFTTRMLSYLIDEERIKDMSPYDAIRDFTMEYPTHIYDLALSNTQSRRSRAGKEF
ESILELLMMGAGIPVDVQGAIGKSFFQKNQIGKLVDLVMPGVVQYTSNKRNTMLISAKTTLR
ERWQEVPEEVNRTGIREMYLATLDDSFSEETINILYEANVVVVTTVENKNFKYKNNNRVLTF
EDMLQSAMELSRKWNNVSYTDSEKEEIQQSILKQIEKYSDFPYVVNYYRNRLSALFD
11100000000000000000000000000000000000000000000000000000000000
00000000000000000000000000000000000000000000000000000000000000
00000000000000000000011111111100000000000000000000000000000000
00000000000000000000000000000000000000000000000000000000000000
000000000000000000000000000000000000000000000000000000111
>DM_train803
VDYNAPLNPKSELFLDDWHIPKFNRFISFTLDVLIDKYKDIFKDFIKLPSRKFHPQYYYKIQ
QPMSINEIKSRDYEYEDGPSNFLLDVELLTKNCQAYNEYDSLIVKNSMQVVMLIEFEVLKAK
NLKRNYLINSEVKAKLLHYLNKLVDATEKKINQALLGASSPKNLDDKVKLSEPFMELVDKDE
LPEYYEIVHSPMALSIVKQNLEIGQYSKIYDFIIDMLLVFQNAHIFNDPSALIYKDATTLTN
YFNYLIQKEFFPELQDLNERGEINLEFDKFEFENYLAIGGGGPAAAGALAISALDND
10000000000000000000000000000000000000000000000000000000000000
00000000000000000000000000000000000000000000000000000000000000
00000000000000000000000000000000000000000000000000000000000000
00000000000000000000000000000000000000000000000000000000000000
000000000000000000000000000000000000111111111111111111111

>DM_train804
MSSDKNGDILRPLSDSEVDELLDLYKVKFGIRNFHYLLLYNQRKWDRQLSEAQIPRNDLNHI
SLRKQFYTHRRGNFRTWGTYVSLHRDIVQSVSFFSWQPDGAAELWECLEQTQLIEWTQGALL
TNVDLGFCNRVKELAVSRGVTAIQPRQCFGMVLSHEDAFCAKVPDLPSEFEIRRLRAEDAAM
VHDSWPNKGEGSLTYLQALVRFNKSLGICRSDTGELIAWIFQNDFSGLGMLQVLPKAERRGL
GGLLAAAMSREIARGEEITLTAWIVATNWRSEALLKRIGYQKDLVNEWIKLVPNSS
111111000000000000000000000000000000000000000000000000000000
000000000000000000000000000000000000000000000000000000000000
000000000000000000000000000000000000000000000000000000000000
000000000000000000000000000000000000000000000000000000000000
00000000000000000000000000000000000000000000000000000001
>DM_train805
MRLGDAAELCYNLTSSYLQIAAESDSIIAQTQRAINTTKSILINETFPKWSPLNGEISFSYN
GGKDCQVLLLLYLSCLWEYYIVKLSQSQFDGKFHRFPLTKLPTVFIDHDDTFKTLENFIEET
SLRYSLSLYESDRDKCETMAEAFETFLQVFPETKAIVIGIRHTDPFGEHLKPIQKTDANWPD
FYRLQPLLHWNLANIWSFLLYSNEPICELYRYGFTSLGNVEETLPNPHLRKDKNSTPLKLNF
EWEIENRYKHNEVTKAEPIPIADEDLVKIENLHEDYYPGWYLVDDKLERAGRIKKK
000000000000000000000000000000000000000000000000000000000000
000000000000000000000011111111111111111000000000000000000000
000000000000000000000000000000000000000000000000000000000000
000000000000000000000000000000000000000000000000000000000000
00000000000000000000000000000000000000000000000000000000
>DM_train806
GTRPRRKKHDYRIALFGGSQPQSCRYFNPKDYSWTDIRCPFEKRRDAACVFWDNVVYILGGS
QLFPIKRMDCYNVVKDSWYSKLGPPTPRDSLAACAAEGKIYTSGGSEVGNSALYLFECYDTR
TESWHTKPSMLTQRCSHGMVEANGLIYVCGGSLGNNVSGRVLNSCEVYDPATETWTELCPMI
EARKNHGLVFVKDKIFAVGGQNGLGGLDNVEYYDIKLNEWKMVSPMPWKGVTVKCAAVGSIV
YVLAGFQGVGRLGHILEYNTETDKWVANSKVRAFPVTSCLICVVDTCGANEETLET
111111100000000000000000000000000000000000000000000000000000
000000000000000000000000000000000000000000000000000000000000
000000000000000000000000000000000000000000000000000000000000
000000000000000000000000000000000000000000000000000000000000
00000000000000000000000000000000000000000000000111111111
>DM_train807
LKQKILNRESGIITYGITPPKKNNTEEKIKEISQKHIERISGLDIDGLVIYDLQDEKERVSE
ERPFPFIETIDPQIYSENYLKDLKIPKIIYRCVGKYTPDEFRRLTRPVSGQDAFSVFVGAAS
RNQSVLLKLSDAYKIRQDVNPDLLLGGVAIPERHMKNTDEHLRIIDKINKGCKYFITQAVYN
VEAAKDFLSDYYYYSKNNNLKMVPIIFTLTPCGSTKTLEFMKWLGISIPRWLENDLMNCEDI
LNKSVSLSKSIFNELMEFCLEKGIPIGCNIESVSVRKVEIEASIALAKDIKYIMKG
000000000000000000000000000000000000000000000000000011111111
111111000000000000000000000000000000000000000000000000000001
111100000000000000000000000000000000000000000000000000000000
000000000000000000000000000000000000000000000000000000000000
00000000000000000000000000000000000000000000000000000011

>DM_train808
GAMERKANYAVDAYFREALRVSEPKAPKAPRPPKQPIVQDFQFFPPRLFELLDQEIYYFRKT
VGYKVPKNTELGSDATKVQREEQRKIDEAEPLTEEEIQEKENLLSQGFTAWTKRDFNQFIKA
NEKYGRDDIDNIAKDVEGKTPEEVIEYNAVFWERCTELQDIERIMGQIERGEGKIQRRLSIK
KALDQKMSRYRAPFHQLRLQYGNNKGKNYTEIEDRFLVCMLHKLGFDKENVYEELRAAIRAS
PQFRFDWFIKSRTALELQRRCNTLITLIERENIELEEKERAEKKKKAPKGSVSAGS
11111111100000000011111111100000000000000000000000000000000000
00000000111111100000000000000000000000000000000000000000000000
00000000000000000000000000000000000000000000000000000000000000
00000000000000000000000000000000000000000000000000000000000000
00000000000000000000000000000000000000000001111111111111
>DM_train810
TEEPDFTALCQKLKIPDHVRERAWLTWEKVSSVDGVLGGYIQKKKELWGICIFIAAVDLDEM
SFTFTELQKNIEISVHKFFNLLKEIDTSTKVDNAMSRLLKKYDVLFALFSKLERTCELIYLT
QPSSSISTEINSALVLKVSWITFLLAKGEVLQMEDDLVISFQLMLCVLDYFIKLSPPMLLKE
PYKTAVIPINGSPRTPRRGQNRSARIAKQLENDTRIIEVLCKEHECNIDEVKNVYFKNFIPF
MNSLGLVTSNGLPEVENLSKRYEEIYLKNKDLDARLFLDHDKTLQTDSIDSFETQR
00000000000000000000000000000000111111111100000000000000000000
00000000000000000000000000000000000000000000000000000000000000
00000000000000000000000000000001110000000000000000000000000000
00000001111111111111111111111111000000000000000000000000000000
00001111111000000000000000000000000000000000000000000000
>DM_train811
EERPYAYVKISDGCDRGCTFCSIPSFKGSLRSRSIEDITREVEDLLKEGKKEIILVAQDTTS
YGIDLYRKQALPDLLRRLNSLNGEFWIRVMYLHPDHLTEEIISAMLELDKVVKYFDVPVQHG
SDKILKLMGRTKSSEELKKMLSSIRERFPDAVLRTSIIVGFPGETEEDFEELKQFVEEIQFD
KLGAFVYSDEEGTVAFNLKEKVDPEMAKRRQEELLLLQAEISNSRLDRFVGKKLKFLVEGKE
GKFLVGRTWTEAPEVDGVVFVRGKGKIGDFLEVVIKEHDEYDMWGSVILEHHHHHH
10000000000001111111111111110000000000000000000000000000000000
00000000000000000000000000000000000000000000000000000000000000
00000000000000000000000000000000000000000000000000000000000000
00000000011111111111000000000000000000000000000000000000000000
00000000000000000000000000000000000000000000000011111111
>DM_train812
KLFLFEFATCGERIEDSTAVEGLAMFKSAFDGFKNYYEITGFVRPEFSCLFTLPVDSMDSME
KYLEKSDAFLIIAPEDDFLLYTLTKKAEKYCENLGSSSRAIAVTSDKWELYKKLRGEVQVPQ
TSLRPLDCKFIIKPRTACAGEGIGFSDEVPDGHIAQEFIEGINLSVSLAVGEDVKCLSVNEQ
IINNFRYAGAVVPARISDEVKREVVEEAVRAVECVEGLNGYVGVDIVYSDQPYVIEINARLT
TPVVAFSRAYGASVADLLAGGEVKHVRRQMVRKSKSAEKPYVSVGDYTLEIIDLD
00000000000000000000000000000000000000000000000000000000000000
00000000000000000000000000000000000000000000000000000000000000
00000000000000000011110000000000000000000000000000000000000000
00000000000000000000000000000000000000000000000000000000000000
0000000000000000000000000000000000000000000000000000000

>DM_train813
MGPFDPEEMLFIFTRCMEDNLEDGANRLPMLAKWKEWINEPVDSPATQCFGKCVLVRTGLYD
PVAQKFDASVIQEQFKAYPSLGEKSKVEAYANAVKQLPSTNNDCAAVFKAYDPVHKAHKDTS
KNLFHGNKELTKGLYEKLGKDIRQKKQSYFEFCENKYYPAGSDKRQQLCQIRQYTVLDDALF
KEHTDCVMKGIRYITKDNQLDVEEVKRDFKLVNKDTKALEEVLNDCKSKEPSNAKEKSWHYY
KCLVESSVKDDFKEAFDYREVRSQIYAFNLPKNQAYSKPAVQSQVMEIDGKQCPQ
00000000000000000000000000000000000000000000000000000000000000
00000000000000000000000000000000000000000000000000000000000000
00000000000000000000000000000000000000000000000000000000000000
00000000000000000000000000000000000000000000000000000000000000
0000000000000000000000000000000000000000011110000000000
>DM_train814
MKITVLGCGALGQLWLTALCKQGHEVQGWLRVPQPYCSVNLVETDGSIFNESLTANDPDFLA
TSDLLLVTLKAWQVSDAVKSLASTLPVTTPILLIHNGMGTIEELQNIQQPLLMGTTTHAARR
DGNVIIHVANGITHIGPARQQDGDYSYLADILQTVLPDVAWHNNIRAELWRKLAVNCVINPL
TAIWNCPNGELRHHPQEIMQICEEVAAVIEREGHHTSAEDLRDYVMQVIDATAENISSMLQD
IRALRHTEIDYINGFLLRRARAHGIAVPENTRLFEMVKRKESEYERIGTGLPRPW
00000000000000000000000000000000000000000000000000000000000000
00000000000000000000000000000000000000000000000000000000000000
00000000000000000000000000000000000000000000000000000000000000
00000000000000000000000000000000000000000000000000000000000000
0000000000000000000000000000000000000000000011111111111
>DM_train815
GPVDSTDVRRITPKKLRELSDLLRTHLSSAATKQLDMGGVLSDLDTMLVALDKAEREGGVDK
DQLKSFNSLILKTYRVIEDYVKGREGDTKNSSTEVSPYHRSNFMLSIVEPSLQRIQKHLDQT
HSFSDIGSLVRAHKHLETLLEVLVTLSQQGQPVSSETYGFLNRLAEAKITLSQQLNTLQQQQ
ESAKAQLSILINRSGSWADVARQSLQRFDSTRPVVKFGTEQYTAIHRQMMAAHAAITLQEVS
EFTDDMRNFTVDSIPLLIQLGRSSLMDEHLVEQREKLRELTTIAERLNRLEREWM
11111111100000000000000000000000000000000000000000000000011000
00000000000000000000000111111111111111111000000000000000000000
00000000000000000000000000001110000000000000000000000000000000
00000000000000000000000000000000000000000000000000000000000000
0000000000000000000000000000000000000000000000000000001
>DM_train816
MSLTNDERILSWNETPSKPRYTPPPGAIDAHCHVFGPMAQFPFSPKAKYLPRDAGPDMLFAL
RDHLGFARNVIVQASCHGTDNAATLDAIARAQGKARGIAVVDPAIDEAELAALHEGGMRGIR
FNFLKRLVDDAPKDKFLEVAGRLPAGWHVVIYFEADILEELRPFMDAIPVPIVIDHMGRPDV
RQGPDGADMKAFRRLLDSREDIWFKATCPDRLDPAGPPWDDFARSVAPLVADYADRVIWGTD
WPHPNMQDAIPDDGLVVDMIPRIAPTPELQHKMLVTNPMRLYWSEEMEGHHHHHH
11000000000000000000000000000000000000000000000000000000000000
00000000000000000000000000000000000000000000000000000000000000
00000000000000000000000000000000000000000000000000000000000000
00000000000000000000000000000000000000000000000000000000000000
0000000000000000000000000000000000000000000000001111111

>DM_train817
MKGLKERQDRRLGAFLGLAVGDALGAQVEGLPKGTFPEVREMKGGGPHRLPPGFWTDDTSQA
LCLAESLLQRGFDPKDQMDRYLRWYREGYRSATGVCFGLGHATRRALERYAATGDPYAGDEA
GAGNGPLMRLAPLVLAYENHPDLLSLARRAARTTHGAREALEATEVLAWLLREALRGAPKEA
LLALEPFRGADLHPALRRVVEGGFWEAPEEGPGYAPGTLAAALWAFARGRDFEEGMRLAVNL
GGDADTVGAVYGQLAGAYYGLGAIPGRWLRALHLREEMEALALALYRMSMASPRE
11000000000000000000000000000000000000000000000000000000000000
00000000000000000000000000000111111111111000000000000000000000
00000000000000000000000000000000000000000000000000000000000000
00000000000000000000000000000000000000000000000000000000000000
0000000000000000000000000000000000000000000000000000111
>DM_train819
TKIDAYAHILPAKYYQKMLSVEPNIPNMFPFIKIKTLMDLDERLTKWPDQNTKQVISLANIS
PEDFTDSKTSAELCQSANEELSNLVDQHPGKFAGAVAILPMNNIESACKVISSIKDDENLVG
AQIFTRHLGKSIADKEFRPVLAQAAKLHVPLWMHPVFDARKPDNNLVFSWEYELSQAMLQLV
QSDLFQDYPNLKILVHHAGAMVPFFSGRIDHILDEKHAQDFKKFYVDTAILGNTPALQLAID
YYGIDHVLFGTDAPFAVMPSGADQIITQAINDLTISDKDKQKIFHDNYYSLIKE
00000000000000000000000000000000000000000000000000000000000000
00000000000000000000000000000000000000000000000000000000000000
00000000000000000000000000000000000000000000000000000000000000
00000000000000000000000000000000000000000000000000000000000000
000000000000000000000000000000000000000000000000000001
>DM_train820
MSQLLQDYLNWENYILRRVDFPTSYVVEGEVVRIEAMPRLYISGMGGSGVVADLIRDFSLTW
NWEVEVIAVKDYFLKARDGLLIAVSYSGNTIETLYTVEYAKRRRIPAVAITTGGRLAQMGVP
TVIVPKASAPRAALPQLLTAALHVVAKVYGIDVKIPEGLEPPNEALIHKLVEEFQKRPTIIA
AESMRGVAYRVKNEFNENAKIEPSVEILPEAHHNWIEGSERAVVALTSPHIPKEHQERVKAT
VEIVGGSIYAVEMHPKGVLSFLRDVGIASVKLAEIRGVNPLATPRIDALKRRLQ
10000000000000000000000000000000000000000000000000000000000000
00000000000000000000000000000000000000000000000000000000000000
00000000000000000000000000000000000000000000000000000000000000
00000000000000000000000000000000000000000000000000000000000000
000000000000000000000000000000000000000000000000000001
>DM_train821
MAAINTKVKKAVIPVAGLGTRMLPATKAIPKEMLPLVDKPLIQYVVNECIAAGITEIVLVTH
SSKNSIENHFDTSFELEAMLEKRVKRQLLDEVQSICPPHVTIMQVRQGLAKGLGHAVLCAHP
VVGDEPVAVILPDVILDEYESDLSQDNLAEMIRRFDETGHSQIMVEPVADVTAYGVVDCKGV
ELAPGESVPMVGVVEKPKADVAPSNLAIVGRYVLSADIWPLLAKTPPGAGDEIQLTDAIDML
IEKETVEAYHMKGKSHDCGNKLGYMQAFVEYGIRHNTLGTEFKAWLEEEMGIKK
11111000000000000000000000000000000000000000000000000000000000
00000000000000000000111100000000000000000000000011000000000000
00000000000000000000000000000000000000000000000000000000000000
00000000000000011111110000000000000000000000011000000000000000
000000000000000000000000000000000000000000000000001111

>DM_train822
MTATAETSTGTKVVLGQNQYGKAEVRLVKVTRNTARHEIQDLNVTSQLRGDFEAAHTAGDNA
HVVATDTQKNTVYAFARDGFATTEEFLLRLGKHFTEGFDWVTGGRWAAQQFFWDRINDHDHA
FSRNKSEVRTAVLEISGSEQAIVAGIEGLTVLKSTGSEFHGFPRDKYTTLQETTDRILATDV
SARWRYNTVEVDFDAVYASVRGLLLKAFAETHSLALQQTMYEMGRAVIETHPEIDEIKMSLP
NKHHFLVDLQPFGQDNPNEVFYAADRPYGLIEATIQREGSRADHPIWSNIAGFC
111111111100000000000000000000000000000000000000000000000000
000000000000000000000000000000000000000000000000000000000000
000000000000000000000000000000000000000000000000000000000000
000000000000000000000000000000000000000000000000000000000000
000000000000000000000000000000000000000000000000011111
>DM_train823
MRGSHHHHHHTDPHASSVPLEWPLSSQSGSYELRIEVQPKPHHRAHYETEGSRGAVKAPTGG
HPVVQLHGYMENKPLGLQIFIGTADERILKPHAFYQVHRITGKTVTTTSYEKIVGNTKVLEI
PLEPKNNMRATIDCAGILKLRNADIELRKGETDIGRKNTRVRLVFRVHIPESSGRIVSLQTA
SNPIECSQRSAHELPMVERQDTDSCLVYGGQQMILTGQNFTSESKVVFTEKTTDGQQIWEME
ATVDKDKSQPNMLFVEIPEYRNKHIRTPVKVNFYVINGKRKRSQPQHFTYHPV
111111111111111111110000000000000000000000000000000000000000
000000000000000000000000000000000000000000000000000000000000
000000000000000000000000000000000000000000000000000000000000
000000000000000000000000000000000000000000000000000000000000
00000000000000000000000000000000000000000000000000000
>DM_train824
TVNQAKIFAQTTKMLEFAKQLLETDDFSTLREAYYVSKNWGEARFDDQQASNNVIEDLEAAL
GVLREHLGFIPEEDGSSVVGPLKIIEETPEGELVVDCTKLGTGAYNIPNDVTKLNLETDADF
ILAIETSGMFARLNAERFWDKHNCILVSLKGVPARATRRFIKRLHEEHDLPVLVFTDGDPYG
YLNIYRTLKVGSGKAIHLADKLSIPAARLIGVTPQDIIDYDLPTHPLKEQDIKRIKDGLKND
DFVRSFPEWQKALKQMLDMGVRAEQQSLAKYGLKYVVNTYLPEKIKDESTWLP
111000000000000000000000000000000000000000000000000000000000
000000000000000000000000000000000000000000000000000000000000
000000000000000000000000000000000000000000000000000000000000
000000000011111111100000000000000000000000000000000000000000
00000000000000000000000000000000000000000000000000000
>DM_train825
GMSSKAEKDIKWGIAPIGWRNDDIPSIGKDNNLQQLLSDIVVAGFQGTEVGGFFPGPEKLNY
ELKLRNLEIAGQWFSSYIIRDGIEKASEAFEKHCQYLKAINAPVAVVSEQTYTIQRSDTANI
FKDKPYFTDKEWDEVCKGLNHYGEIAAKYGLKVAYHHHMGTGIQTKEETDRLMANTDPKLVG
LLYDTGHIAVSDGDYMALLNAHIDRVVHVHFKDVRRSKEEECRAKGLTFQGSFLNGMFTVPG
DGDLDFKPVYDKLIANNYKGWIVVEAEQDPSKANPLEMAQIAHRYIKQHLIEN
100000000000000000000000000000000000000000000000000000000000
000000000000000000000000000000000000000000000000000000000000
000000000000000000000000000000000000000000000000000000000000
000000000000000000000000000000000000000000000000000000000000
00000000000000000000000000000000000000000000000000000

>DM_train826
MKPADLTNADRIALELGHAGRNAIPYLDDDRNADRPFTLNTYRPYGYTPDRPVVVVQHGVLR
NGADYRDFWIPAADRHKLLIVAPTFSDEIWPGVESYNNGRAFTAAGNPRHVDGWTYALVARV
LANIRAAEIADCEQVYLFGHSAGGQFVHRLMSSQPHAPFHAVTAANPGWYTLPTFEHRFPEG
LDGVGLTEDHLARLLAYPMTILAGDQDIATDDPNLPSEPAALRQGPHRYARARHYYEAGQRA
AAQRGLPFGWQLQVVPGIGHDGQAMSQVCASLWFDGRMPDAAELARLAGSQSA
111100000000000000000000000011100000000000000000000000000000
000000000000000000000000000000000000000000000000000000000000
000000000000000000000000000000000000000000000000000000000000
000000000000000000000000000000110000000000000000000000000000
00000000000000000000000000000000000000000000011111
>DM_train827
ATGDFPNKPLDIIVTFPPGGGTDMLARLIGNYLTESLGQTAVVENRPGASGNVGARLVADRA
PDGYSLLMVNSSFAVNPGVFRNLPFDPKKDFAAVINVAYVPSVFVVPAGSKYKTLGELMAAA
KQTNTQVTYGSCGNGTPQHLAGELLNVSAKTHMVHVPYKGCGPALNDVLGSQIGLAVVTASS
AIPFIKAGKLQALAVTSKERSALLPEVPTVAEQGVAGYELNQWHGLLVPGATPMAVRQKLYD
GIAKVMQRDDVQKKLADLGYSTASDGPEVFQKMVETDIDRFSALTKQIGLKVD
111100000000000001111110000000000000000000000000000000000000
000000000000000000000000000000000000000000000000000000000000
000000000000000000000000000000000000000000000000000000000000
000000000000000000000000000000000000000000000000000000000000
00000000000000000000000000000000000000000000000000
>DM_train828
QGLVPQGQTQVLQGGNKVPVVNIADPNSGGVSHNKFQQFNVANPGVVFNNGLTDGVSRIGGA
LTKNPNLTRQASAILAEVTDTSPSRLAGTLEVYGKGADLIIANPNGISVNGLSTLNASNLTL
TTGRPSVNGGRIGLDVQQGTVTIERGGVNATGLGYFDVVARLVKLQGAVSSKQGKPLADIAV
VAGANRYDHATRRATPIAAGARGAAAGAYAIDGTAAGAMYGKHITLVSSDSGLGVRQLGSLS
SPSAITVSSQGEIALGDATVQRGPLSLKGAGVVSAGKLASGGGAVNVAGGGAV
000000000000000000000000000000000000000000000000000000000000
000000000000000000000000000000000000000000000000000000000000
000000000000000000000000000000000000000000000000000000000000
000000000000000000000000000000000000000000000000000000000000
00000000000000000000000000000000000000000000001111
>DM_train829
MPIIIDKDLPARKVLQEENIFVMTKERAETQDIRALKIAILNLMPTKQETEAQLLRLIGNTP
LQLDVHLLHMESHLSRNVAQEHLTSFYKTFRDIENEKFDGLIITGAPVETLSFEEVDYWEEL
KRIMEYSKTNVTSTLHICWGAQAGLYHHYGVQKYPLKEKMFGVFEHEVREQHVKLLQGFDEL
FFAVHSRHTEVRESDIREVKELTLLANSEEAGVHLVIGQEGRQVFALGHSEYSCDTLKQEYE
RDRDKGLNIDVPKNYFKHDNPNEKPLVRWRSHGNLLFSNWLNYYVYQETPYVL
111111111111111100000000000000000000000000000000000000000000
000000000000111111111111000000000000000000000000000000000000
000000000000000000000000000000000000000000000000000000000000
000000000000000000000000000000000000000000000000000000000000
00000000000000000000000000000000000000000000011111

>DM_train831
MTDDKKIRAAIVGYGNIGRYALQALREAPDFEIAGIVRRNPAEVPFELQPFRVVSDIEQLES
VDVALVCSPSREVERTALEILKKGICTADSFDIHDGILALRRSLGDAAGKSGAAAVIASGWD
PGSDSVVRTLMQAIVPKGITYTNFGPGMSMGHTVAVKAIDGVKAALSMTIPLGTGVHRRMVY
VELLPGHNLEEVSAAIKADEYFVHDETHVIQVDEVDALIDMGHGVRMVRKGVSGSTQNQRMS
FDMEINNPALTGQVLVCAARAAMRQQPGAYTLQEIPVIDLLPGDREQWIGKLC
11110000000000000000000000000000000000011111100000000000000000
00000000000000000000000000000000000000000000000000000000000000
00000000000000000000000000000000000000000000000000000000000000
00000000000000000000000000000000000000000000000000000000000000
00000000000000000000000000000000000000000000000000000
>DM_train832
MILLEVNNRIIEETLALKFENAAAGNKPEAVEVTFADFDGVLYHISNPNGDKTKVMVSISLK
FYKELQAHGADELLKRVYGSYLVNPESGYNVSLLYDLENLPASKDSIVHQAGMLKRNCFASV
FEKYFQFQEEGKEGENRAVIHYRDDETMYVESKKDRVTVVFSTVFKDDDDVVIGKVFMQEFK
EGRRASHTAPQVLFSHREPPLELKDTDAAVGDNIGYITFVLFPRHTNASARDNTINLIHTFR
DYLHYHIKCSKAYIHTRMRAKTSDFLKVLNRARPDAEKKEMKTITGKTFSSR
00000000000000000000000000000000000000000000000000000000000000
00000000000000000000000000000000000000000000000000000000000000
00000000000000000000000000000000000000000000000000000000000000
00000000000000000000000000000000000000000000000000000000000000
0000000000000000000000000000000000001111111111111111
>DM_train833
MAIPAFHPGELNVYSAPGDVADVSRALRLTGRRVMLVPTMGALHEGHLALVRAAKRVPGSVV
VVSIFVNPMQFGAGGDLDAYPRTPDDDLAQLRAEGVEIAFTPTTAAMYPDGLRTTVQPGPLA
AELEGGPRPTHFAGVLTVVLKLLQIVRPDRVFFGEKDYQQLVLIRQLVADFNLDVAVVGVPT
VREADGLAMSSRNRYLDPAQRAAAVALSAALTAAAHAATAGAQAALDAARAVLDAAPGVAVD
YLELRDIGLGPMPLNGSGRLLVAARLGTTRLLDNIAIEIGTFAGTDRPDGYR
10000000000000000000000000000000000000000000000000000000000000
00000000001111111111000000000000000000000000000000000000000000
00000000000000000000000000000000000000000000000000000000000000
00000000000000000000000000000000000000000000000000000000000000
0000000000000000000000000000000000000000111111111111
>DM_train834
MEDLYPSRQNNQPKILKRKDPVIYTDRSKDNQAPITKEQLDSYEKNGFLQIKNFFSEDEVID
MQKAIFELQDSIKDVASDKVIREPESNDIRSIFHVHQDDNYFQDVANDKRILDIVRHLLGSD
VYVHQSRINYKPGFKGKEFDWHSDFETWHVEDGMPRMRAISVSIALSDNYSFNGPLMLIPGS
HNYFVSCVGETPDNNYKESLKKQKLGVPDEESLRELTRIGGGISVPTGKAGSVTLFESNTMH
GSTSNITPYPRNNLFMVYNSVKNRLVEPFSGGEKRPEYIAVREKQPVYSAVN
00000000000000000000000000000000000000000000000000000000000000
00000000000000000000000000000000000000000000000000000000000000
00000000000000000000000000000000000000000000000000000000000000
00000000111111111111111110000000000000000000000000000000000000
0000000000000000000000000000000000000000000000001111

>DM_train835
MDLVEEILRLKEERNAIILAHNYQLPEVQDIADFIGDSLELARRATRVDADVIVFAGVDFMA
ETAKILNPDKVVLIPSREATCAMANMLKVEHILEAKRKYPNAPVVLYVNSTAEAKAYADVTV
TSANAVEVVKKLDSDVVIFGPDKNLAHYVAKMTGKKIIPVPSKGHCYVHQKFTLDDVERAKK
LHPNAKLMIHPECIPEVQEKADIIASTGGMIKRACEWDEWVVFTEREMVYRLRKLYPQKKFY
PAREDAFCIGMKAITLKNIYESLKDMKYKVEVPEEIARKARKAIERMLEMSK
00000000000000000000000000000000000000000000000000000000000000
00000000000000001111111111110000000000000000000000000000000000
00000000000000000000000000000000000000001111111111100000000000
00000000000000000000000000000000000000000000000000000000000000
0000000011110000000000000000000000000000000000000011
>DM_train836
MKKSALEKLLSLIENLTNQEFKQATNSLISFIYKLNRNEVIELVRSIGILPEAIKPSSTQEK
LFSKAGDIVLAKAFQLLNLNSKPLEQRGNAGDVIALSKEFNYGLVADAKSFRLSRTAKNQKD
FKVKALSEWREDKDYAVLTAPFFQYPTTKSQIFKQSLDENVLLFSWEHLAILLQLDLEETNI
FSFEQLWNFPKKQSKKTSVSDAENNFMRDFNKYFMDLFKIDKDTLNQLLQKEINFIEERSLI
EKEYWKKQINIIKNFTREEAIEALLKDINMSSKIETIDSFIKGIKSNDRLYL
11000000000000000000000000000000000000000000000000000000000000
00000000000000000000000000000000000000000000000000000000000000
00000000000000000000000000000000000000000000000000000000000000
00000000000000000000000000000000000000000000000000000000000000
0000000000000000000000000000000000000000000000000000
>DM_train837
FVSSHRYVETMLVADQSMAEFHGSGLKHYLLTLFSVAARLYKHPSIRNSVSLVVVKILVIHD
EQKGPEVTSNAALTLRNFCNWQKQHNPPSDRDAEHYDTAILFTRQDLCGSQTCDTLGMADVG
TVCDPSRSCSVIEDDGLQAAFTTAHELGHVFNMPHDDAKQCASLNGVNQDSHMMASMLSNLD
HSQPWSPCSAYMITSFLDNGHGECLMDKPQNPIQLPGDLPGTSYDANRQCQFTFGEDSKHCP
DAASTCSTLWCTGTSGGVLVCQTKHFPWADGTSCGEGKWCINGKCVNKLVPR
11100000000000000000000000000000000000000000000000000000000000
00000000000000000000000000000000000000000000000000000000000000
00000000000000000000000000000000000000000000011111000000011100
00000000000000000000000000000000000000000000000000000000000000
1111000000000111100000000000000000000000000000000001
>DM_train838
MMPEGKIIKALSGFYYVLDESEDSDKVIQCRGRGIFRKNKITPLVGDYVVYQAENDKEGYLM
EIKERTNELIRPPICNVDQAVLVFSAVQPSFSTALLDRFLVLVEANDIQPIICITKMDLIED
QDTEDTIQAYAEDYRNIGYDVYLTSSKDQDSLADIIPHFQDKTTVFAGQSGVGKSSLLNAIS
PELGLRTNEISEHLGRGKHTTRHVELIHTSGGLVADTPGFSSLEFTDIEEEELGYTFPDIRE
KSSSCKFRGCLHLKEPKCAVKQAVEDGELKQYRYDHYVEFMTEIKDRKPRY
10000000000000000000000000000000000000000000000000000000000000
00000000000000000000000000000000000000000000000000000000000000
00000000000000000000000000000000000000000000000000000000000000
01111111111111111111000000000000000000000000000000000000000000
000000000000000000000000000000000000000000000000000

>DM_train839
GLDVAISQNGFFRLVDSNGSVFYSRNGQFKLDENRNLVNMQGMQLTGYPATGTPPTIQQGAN
PAPITIPNTLMAAKSTTTASMQINLNSTDPVPSKTPFSVSDADSYNKKGTVTVYDSQGNAHD
MNVYFVKTKDNEWAVYTHDSSDPAATAPTTASTTLKFNENGILESGGTVNITTGTINGATAA
TFSLSFLNSMQQNTGANNIVATNQNGYKPGDLVSYQINNDGTVVGNYSNEQEQVLGQIVLAN
FANNEGLASQGDNVWAATQASGVALLGTAGSGNFGKLTNGALEASNVDLSK
00000000000000000000000000000000000000000000000000000000000000
00000000000000000000000000000000000000000000000000000000000000
00000000000000000000000000000000000000000000000000000000000000
00000000000000000000000000000000000000000000000000000000000000
000000000000000000000000000000000000000000000111111
>DM_train840
MIKQRTLKNIIRATGVGLHSGEKVYLTLKPAPVDTGIVFSRTDLDPVVEIPARAENVGETTM
STTLVKGDVKVDTVEHLLSAMAGLGIDNAYVELSASEVPIMDGSAGPFVFLIQSAGLQEQEA
AKKFIRIKREVSVEEGDKRAVFVPFDGFKVSFEIDFDHPVFRGRTQQASVDFSSTSFVKEVS
RARTFGFMRDIEYLRSQNLALGGSVENAIVVDENRVLNEDGLRYEDEFVKHKILDAIGDLYL
LGNSLIGEFRGFKSGHALNNQLLRTLIADKDAWEVVTFEDARTAPISYMRP
00000000000000000000000000000000000000000000000000000000000000
00000000000000000000000000000000000000000000000000000000000000
00000000000000000000000000000000000000000000000000000000000000
00000000000000000000000000000000000000000000000000000000000000
000000000000000000000000000000000000000011111111111
>DM_train841
MEFKLQELNLTNQDTGPYGITVSDKGKVWITQHKANMISCINLDGKITEYPLPTPDAKVMCL
TISSDGEVWFTENAANKIGRITKKGIIKEYTLPNPDSAPYGITEGPNGDIWFTEMNGNRIGR
ITDDGKIREYELPNKGSYPSFITLGSDNALWFTENQNNAIGRITESGDITEFKIPTPASGPV
GITKGNDDALWFVEIIGNKIGRITTSGEITEFKIPTPNARPHAITAGAGIDLWFTEWGANKI
GRLTSNNIIEEYPIQIKSAEPHGICFDGETIWFAMECDKIGKLTLIKDNME
10000000000000000000000000000000000000000000000000000000000000
00000000000000000000000000000000000000000000000000000000000000
00000000000000000000000000000000000000000000000000000000000000
00000000000000000000000000000000000000000000000000000000000000
000000000000000000000000000000000000000000000011111
>DM_train842
CVIFPVEIDVSQTIIRDCQVDKQTRELVYINKIMNTQLTKPVLMMFNISGPIRSVTRKNNNL
RDRIKSKPDEQFDQLEKDYSDQMDGFHDSIKYFKDEHYSVSCQNGSVLKSKFAKILKSHDYT
DKKSIEAYEKYCLPKLVDERNDYYVAVCVLKPGFENGSNQVLSFEYNPIGNKVIVPFAHEIN
DTGLYEYDVVAYVDSVQFDGEQFEEFVQSLILPSSFKNSEKVLYYNEASKNKSMIYKALEFT
TESSWGKSEKYNWKIFCNGFIYDKKSKVLYVKLHNVTSALNKNVILNTIK
11110000000000000000000000000000000000000000000000000000000000
00000000000000000001111111100000000000000000000000000000000000
00000000000000000000000000000000000000000000000000000000000000
00000000000000000000000000000000000000000000000011100000000000
01111111100000000000000000000000000000000000000000

>DM_train843
MLIDPIELYRYPEKWIKDRDAEKKVRSGLYILTEDGYLRRGITTGTTASAAAVAAIASLKEK
VEKVKVSTPAGVDVEVEVEAEKGFARVRKFSGDHEFDVTNGIIFEAEVCETSGIFFGRGVGV
KAGEKAVSRSAKLQILENFIKASREFNFSGGVRISVPDGEEVAKKTGNEKVGIKGGISILGT
TGFVEPWCKKLVETKLKIAMQYHRIAITTGRKAWLYARKKFPEYQPFVFGVHIDEALKHPGE
KIIVGFPGLLKIWAGSRDRIEERAREEGVRVVVIEDDMDSWVWDVQGTDH
00000000000000000000000000000000000000000000000000000000000000
00000000000000000000000000000000000000000000000000000000000000
00000000000000000000000000000000000000000000000000000000000000
00000000000000000000000000000000000000000000000000000000000000
00000000000000000000000000000000001111111111111111
>DM_train844
SDESFEFDVSVIGLGAMGTIMAQVLLKQGKRVAIWNRSPGKAAALVAAGAHLCESVKAALSA
SPATIFVLLDNHATHEVLGMPGVARALAHRTIVDYTTNAQDEGLALQGLVNQAGGHYVKGMI
VAYPRNVGHRESHSIHTGDREAFEQHRALLEGLAGHTVFLPWDEALAFATVLHAHAFAAMVT
FFEAVGAGDRFGLPVSKTARLLLETSRFFVADALEEAVRRLETQDFKGDQARLDVHADAFAH
IAQSLHAQGVWTPVFDAVCQVVQRAAAMGYGDQDIAATTKSFAREQEEGH
00000000000000000000000000000000000000000000000000000000000000
00000000000000000000000000000000000000000000000000000000000000
00000000000000000000000000000000000000000000000000000000000000
00000000000000000000000000000000000000000000000000000000000000
00000000000000000000000000000000000000000001111111
>DM_train845
MTDTQYIGRFAPSPSGELHFGSLIAALGSYLQARARQGRWLVRIEDIDPPREVPGAAETILR
QLEHYGLHWDGDVLWQSQRHDAYREALAWLHEQGLSYYCTCTRARIQSIGGIYDGHCRVLHH
GPDNAAVRIRQQHPVTQFTDQLRGIIHADEKLAREDFIIHRRDGLFAYNLAVVVDDHFQGVT
EIVRGADLIEPTVRQISLYQLFGWKVPDYIHLPLALNPQGAKLSKQNHAPALPKGDPRPVLI
AALQFLGQQAEAHWQDFSVEQILQSAVKNWRLTAVPESAIVNSTFSNASC
11100000000000000000000000000000000000000000000000000000000000
00000000000000000000000000000000000000000000000000000000000000
00000000000000000000000000000000000000000000000000000000000000
00000000000000000000000000000000000001111111111110000000000000
00000000000000000000000000000000000000000111111111
>DM_train846
MGSDKIHHHHHHMITFIGHVSKDVNVVDGKREIAYGGGVVMGAITSSLLGVKTKVITKCTRE
DVSKFSFLRDNGVEVVFLKSPRTTSIENRYGSDPDTRESFLISAADPFTESDLAFIEGEAVH
INPLWYGEFPEDLIPVLRRKVMFLSADAQGFVRVPENEKLVYRDWEMKEKYLKYLDLFKVDS
REAETLTGTNDLRESCRIIRSFGAKIILATHASGVIVFDGNFYEASFRSWSLEGRTGRGDTC
TAAFLVGFVFKKMSIEKATKFAAAVTSVKMRHPGPLRREDLEAISGDQYF
11111000000000000000000000000000000000000000000000000000000000
00000000000000000000000000000011111000000000000000000000000000
00000000000000000000000000000000000000000000000000000000000000
00000000000000000000000000000000000000000000000000000000000000
00000000000000000000000000000000000000000000011111

>DM_train847
GAMVVSPAGADRRIPTWASRVVSGLARDRPVVVTKEDLTQRLTEAGCGRDPDSAIRELRRIG
WLVQLPVKGTWAFIPPGEAAISDPYLPLRSWLARDQNAGFMLAGASAAWHLGYLDRQPDGRI
PIWLPPAKRLPDGLASYVSVVRIPWNAADTALLAPRPALLVRRRLDLVAWATGLPALGPEAL
LVQIATRPASFGPWADLVPHLDDLVADCSDERLERLLSGRPTSAWQRASYLLDSGGEPARGQ
ALLAKRHTEVMPVTRFTTAHSRDRGESVWAPEYQLVDELVVPLLRVIGKA
100000000000000000000000000000000000000000000000000000000000
000000000000000000000000000000000000000000000000000000000000
000000000000000000000000000000000000000000000000000000000000
000000000000000000000000000000000000000000000000000000000000
00000000000000000000011100000000000000000000000001
>DM_train848
MSERIVPSGDVELWSDDFGDPADPALLLVMGGNLSALGWPDEFARRLADGGLHVIRYDHRDT
GRSTTRDFAAHPYGFGELAADAVAVLDGWGVDRAHVVGLSMGATITQVIALDHHDRLSSLTM
LLGGGLDIDFDANIERVMRGEPTLDGLPGPQQPFLDALALMNQPAEGRAAEVAKRVSKWRIL
SGTGVPFDDAEYARWEERAIDHAGGVLAEPYAHYSLTLPPPSRAAELREVTVPTLVIQAEHD
PIAPAPHGKHLAGLIPTARLAEIPGMGHALPSSVHGPLAEVILAHTRSAA
100000000000000000000000000000000000000000000000000000000000
000000000000000000000000000000000000000000000000000000000000
000000000000000000000000000000000000000000000000000000000000
000000000000000000000000000000000000000000000000000000000000
00000000000000000000000000000000000000000000000000
>DM_train849
MEEKQILCVGLVVLDVISLVDKYPKEDSEIRCLSQRWQRGGNASNSCTVLSLLGAPCAFMGS
MAPGHVADFVLDDLRRYSVDLRYTVFQTTGSVPIATVIINEASGSRTILYYDRSLPDVSATD
FEKVDLTQFKWIHIEGRNASEQVKMLQRIDAHNTRQPPEQKIRVSVEVEKPREELFQLFGYG
DVVFVSKDVAKHLGFQSAEEALRGLYGRVRKGAVLVCAWAEEGADALGPDGKLLHSDAFPPP
RVVDTLGAGDTFNASVIFSLSQGRSVQEALRFGCQVAGKKCGLQGFDGIV
110000000000000000000000000000000000000000000000000000000000
000000000000000000000000000000000000000000000000000000000000
000000000000000000000000000000000000000000000000000000000000
000000000000000000000000000000000000000000000000000000000000
00000000000000000000000000000000000000000000000000
>DM_train850
MPAPEAPTSTLPPERPLTNLQQQIQQLVSRQPNLTAGLYFFNLDSGASLNVGGDQVFPAAST
IKFPILVAFFKAVDEGRVTLQERLTMRPDLIAPEAGTLQYQKPNSQYAALEVAELMITISDN
TATNMIIDRLGGAAELNQQFQEWGLENTVINNPEPDMKGTNTTSPRDLATLMLKIGQGEILS
PRSRDRLLDIMRRTVTNTLLPAGLGKGATIAHKTGDIGIVVGDAGMVDMPNGQRYVAAMMVK
RPYNDPRGSELIRQVSRMVYQAFEKLSPPEQKLISEEDLNSAVDHHHHHH
111111111100000000000000000000000000000000000000000000000000
000000000000000000000000000000000000000000000000000000000000
000000000000000000000000000000000000000000000000000000000000
000000000000000000000000000000000000000000000000000000000000
00000000000000000000000000111111111111111111111111

>DM_train851
GASTVEKELLRSRRLENSIIEQKGTMRCYAYVMEQNLPENLLFDYENGVITQGLSEHVYKFN
RVIPHLKVSEDKFFTQEYSVYHDMCLNQKKNFNLISLSTTPHGSLRESLIKFLAEKDTIYQK
QYVITLQFVFLSDDEFSQDMLLDYSHNDKDSIKLKFEKHSISLDSKLVIIENGLEDLPLNFS
CDEHPNLPHSGMGIIKVQFFPRDSKSDGNNDPVPVDFYFIELNNLKSIEQFDKSIFKKESCE
TPIALVLKKLISDTKSFFLLNLNDSKNVNKLLTISEEVQTQLCKRKKKLT
10000000000000000000000000000000000000000000000000000000000000
00000000000000000000000000000000000000000000000000000000000000
00000000000000000000000011111000000000000000000000000000000000
00111111100000000000000011111110000000000000000000000000001000
00000000000000000000000000000000000000000001111111
>DM_train852
ELSENFKKLMKYPYRPCTCTRCIEEQRVSAWFDERFNRSMQPLLTAKNAHLEEDTYKWWLRL
QREKQPNNLNDTIRELFQVVPGNVDPLLEKRLVSCRRCAVVGNSGNLKESYYGPQIDSHDFV
LRMNKAPTEGFEADVGSKTTHHFVYPESFRELAQEVSMILVPFKTTDLEWVISATTTGRISH
TYVPVPAKIKVKKEKILIYHPAFIKYVFDRWLQGHGRYPSTGILSVIFSLHICDEVDLYGFG
ADSKGNWHHYWENNPSAGAFRKTGVHDGDFESNVTTILASINKIRIFKGR
11111111111111100000000000000000000000000000000000000000000000
00000000000000000000000000000000000000000000000000000000000000
00000000000000000000000000000000000000000000000000000000000000
00000000000000000000000000000000000000000000000000000000000000
00000000000011111111111000000000000000000000000000
>DM_train853
AITKPLLAATLENIEDVQFPCLATPKIAGIRSVKQTQMLSRTFKPIRNSVMNRLLTELLPEG
SDGEISIEGATFQDTTSAVMTGHKMYNAKFSYYWFDYVTDDPLKKYIDRVEDMKNYITVHPH
ILEHAQVKIIPLIPVEINNITELLQYERDVLSKGFEGVMIRKPDGKYKFGRSTLKEGILLKM
KQFKDAEATIISMTALFKNTNTKTKDNFGYSKRSTHKSGKVEEDVMGSIEVDYDGVVFSIGT
GFDADQRRDFWQNKESYIGKMVKFKYFEMGSKDCPRFPVFIGIRHEEDR
00000000000000000000000000000000000000000000000000000000000000
00000000000000000000000111100000000000000000000000000000000000
00000000000000000000000000000000000000000000000000000000000000
00000000000000000011111111111111111110000000000000000000000000
0000000000000000000000000000011111000000000011111
>DM_train854
LEVKPEINFVNIGERCNVAGSRKFLRLVNEKKYDEALSIARQQVEDGALVIDVNMDDGLLDA
RTEMTTFLNLIMSEPEIARVPVMIDSSKWEVIEAGLKCLQGKSIVNSISLKEGEEVFLEHAR
IIKQYGAATVVMAFDEKGQADTAARKIEVCERAYRLLVDKVGFNPHDIIFDPNVLAVATGIE
EHNNYAVDFIEATGWIRKNLPGAHVSGGVSNLSFSFRGNNYIREAMHAVFLYHAIQQGMDMG
IVNPGTSVLYSDIPADTLEKIEDVVLNRRPDAAERLIELAEALKETMGG
11111000000000000000000000000000000000000000000000000000000000
00000000000000000000000000000000000000000000000000000000000000
00000000000000000000000000000000000000000000000000000000000000
00000000000000000000000000000000000000000000000000000000000000
0000010000000000000000000000000000000000000011111

>DM_train855
MPIKETDKEVVLTHPADETTSVHILKYGATVYSWKLKSEEQLWLSTAAKLDGSKPVRGGIPL
VFPVFGKNSTDEHLSKLPQHGLARNSTWEFLGQTKENPPTVQFGLKPEIANPELTKLWPMDY
LLILTVELGSDYLKTAIEVENTSSSKELKFNWLFHTYFRIEDIEGTMVSNLAGMKLYDQLLK
ESYVDKHPVVTFNQETDVIYQNVSAERAIQIVDKGVQIHTLKRYNLPDTVVWNPWIEKSQGM
ADFEPKTGYQQMICIEPGHVHDFISLAPGKKWNAYQLLCKEELKYQAIQ
100000000000000000000000000000000000000000000000000000000000
000000000000000000000000000000000000000000000000000000000000
000000000000000000000000000000000000000000000000000000000000
000000000000000000000000000000000000000000000000000000000000
0000000000000000000000000000000000000000001111111
>DM_train856
SRVENSNGQSQPAATSKTVKDNAEIYYDDDDSDRFYFHVWGGEDIHVGLYKEPVDQDEIREA
SLRTDEWLASELAMTGVLQRQAKGLDLGAGYGGAARFLVRKFGVSIDCLNIAPVQNKRNEEY
NNQAGLADNITVKYGSFLEIPCEDNSYDFIWSQDAFLHSPDKLKVFQECARVLKPRGVMAIT
DPMKEDGIDKSSIQPILDRIKLHDMGSLGLYRSLAKECGLVTLRTFSRPDSLVHHYSKVKAE
LIKRSSEIASFCSPEFQANMKRGLEHWIEGGRAGKLTWGGMLFRKSDKI
111111111111111111111111111110000000000000000000000000000000
000000000000000000000000000000000000000000000000000000000000
000000000000000000000000000000000000000000000000000000000000
000000000000000000000000000000000000000000000000000000000000
0000000111000000000000000000000000000000000000000
>DM_train857
MEGGMEKGTFQIKTGFAEMFKGGVIMDVTTPEQAVIAEEAGAVAVMALERVPADIRAQGGVA
RMSDPKIIKEIMAAVSIPVMAKVRIGHFVEAMILEAIGVDFIDESEVLTPADEEHHIDKWKF
KVPFVCGARNLGEALRRIAEGAAMIRTKGEAGTGNVVEAVRHARTMWKEIRYVQSLREDELM
AYAKEIGAPFELVKWVHDHGRLPVVNFAAGGIATPADAALMMHLGMDGVFVGSGIFKSGDPR
KRARAIVRAVAHYNDPEVLAEVSEDLGEPMVGINLDQLKEEERLAKRGW
111111110000000000000000000000000000000000000000000000000000
000000000000000000000000000000000000000000000000000000000000
000000000000000000000000000000000000000000000000000000000000
000000000000000000000000000000000000000000000000000000000000
0000000000000000000000000000001111111111111111111
>DM_train858
EPVQVFTDDLGRKVTVPAHPKRIVSLHDLDITIPLIELGVPPVASHGRTRPDGSHFIRSGAL
LTGVDFDNSSIAFIGTADIDIEAIVAAKPDLIITEPTRNTPIERLEKIAPTVSIDHLKGGAP
EIYRKLAELTGTQSQLAILERRYQAQINALKATLDSQKITVSVIQANQGKINVMHSYHSLGR
VLRDAGFRFPPLIESIPEGGRMDVSAERLPELDADFVFATWRGDTGGKPQDELATMEKVMPG
WCQFLTACRSGRYVLISREEAISNSFASLGLMAAQIQSQIAGRPLPEAK
110000000000000000000000000000000000000000000000000000000000
000000000000000000000000000000000000000000000000000000000000
000000000000000000000000000000000000000000000000000000000000
000000000000000000000000000000000000000000000000000000000000
0000000000000000000000000000000000000000000000111

>DM_train859
MRERIREHLSSEESVGLLFSGCTGINDKTLGARGGEVIMVTSGSGMGKSTFVRQQALQWGTA
MGKKVGLAMLEESVEETAEDLIGLHNRVRLRQSDSLKREIIENGKFDQWFDELFGNDTFHLY
DSFAEAETDRLLAKLAYMRSGLGCDVIILDHISIVVSASGESDERKMIDNLMTKLKGFAKST
GVVLVVICHLKNPDKGKAHEEGRPVSITDLRGSGALRQLSDTIIALERNQQGDMPNLVLVRI
LKCRFTGDTGIAGYMEYNKETGWLEPSSYSGEEESHSESTDWSNDTDF
00000000000000000000000000000000000000000000000000000000000000
00000000000000000000000000000000000000000000000000000000000000
00000000000000000000000000000000001111111111000000000000000000
00000000000111111111111111111111000000000000000001111110000000
000000000000000000000000000001111111111111111111
>DM_train860
MSFFHASQRDALNQSLAEVQGQINVSFEFFPPRTSEMEQTLWNSIDRLSSLKPKFVSVTYGA
NSGERDRTHSIIKGIKDRTGLEAAPHLTCIDATPDELRTIARDYWNNGIRHIVALRGDLPPG
SGKPEMYASDLVTLLKEVADFDISVAAYPEVHPEAKSAQADLLNLKRKVDAGANRAITQFFF
DVESYLRFRDRCVSAGIDVEIIPGILPVSNFKQAKKLADMTNVRIPAWMAQMFDGLDDDAET
RKLVGANIAMDMVKILSREGVKDFHFYTLNRAEMSYAICHTLGVRPGL
11111111111111111111100000000000000000000000000000000000000000
00000000000000000000000000000000000000000000000000000000001111
11110000000000000000000000000000000000000000000000000000000000
00000000000000000000000000000000000000000000000000000000000000
000000000000000000000000000000000000000000000011
>DM_train863
MWRIELKHAVNWELKMKFFVLPELPTPDVVESGVWRRAIVLDGRAVAVMAYPESERTIVVEG
NFENREWEAVRRKLVEYLGLQNPEELYRFMDGDEKLRMLKNRFYGFGRAGLMSMSVFEGIAK
AIIQQQISFVVAEKLAAKIVGRFGDEVEWNGLKFYGFPTQEAILKAGVEGLRECGLSRRKAE
LIVEIAKEENLEELKEWGEEEAYEYLTSFKGIGRWTAELVLSIALGKNVFPADDLGVRRAVS
RLYFNGEIQSAEKVREIARERFGRFARDILFYLFLYDRFFSKKTELV
00000000000000000000000000000000000000000000000000000000000000
00000000000000000000000000000000000000000000000000000000000000
00000000000000000000000000000000000000000000000000000000000000
00000000000000000000000000000000000000000000000000000000000000
00000000000000000000000000000000000000000111100
>DM_train865
SKLGYIPNQAARTLITNKTLTIGLIQKSSAPEIRQNPFNSDVLNGINQACNVRGYSTRMTVS
ENSGDLYHEVKTMIQSKSVDGFILLYSLKDDPIEHLLNEFKVPYLIVGKSLNYENIIHIDND
NIDAAYQLTQYLYHLGHRHILFLQESGHYAVTEDRSVGFKQYCDDVKISNDCVVIKSMNDLR
DFIKQYCIDASHMPSVIITSDVMLNMQLLNVLYEYQLRIPEDIQTATFNTSFLTENATPSQT
SVNINPDVLGFTAGNTIIDVLRNETISFREKLISTQIVERVSTTKI
11111111111111111111000000000000000000000000000000000000000000
00000000000000000000000000000000000000000000000000000000000000
00000000000000000000000000000000000000000000000000000000000000
00011111111100000000000000000000000000000000000000000000000000
0000000000000000000000111111000000000000000000

>DM_train866
MEARNMTPFTYFSLPMQKLFLRNQAAVRNKPYAKYFRSEMRVPLSAVRKIQQGPMALEDTLT
PSIEDINRLLEPDFVSEESGYALLPGPMAYVQSRKFFPGCTAQMFKWWFIWHPAESERYTLW
FPYAHVSNPCVHHQRLRDESLSFEERLYGNTFCASEYVGDRLMHLHIDFQQPASLGLNTDLY
REAKIDGSVSALMSLADHPEVPVSLMVHLFKEVPDGMYLTSRYWVGAHPSMARFPGAEKAAS
LLKENGFGEAELETLAYEFAVHDMCEFNHLASFLPDLYREFGTPAA
111000000000000000000000000000000000000000000000000000000000
000000000000000000000000000000000000000000000000000000000000
000000000000000000000000000000000000000000000000000000000000
000000000000000000000000000000000000000000000000000000000000
0000000000000000000000000000000000000000000111
>DM_train867
MGAPPLGPTIIPPVRTLWLRDRALDLDRVRLLGVLNLTPDSFSDGGRYLDPERALERAREMV
AEGADILDLGAESTRPGAAPVPVEEEKRRLLPVLEAVLSLGVPVSVDTRKPEVAEEALKLGA
HLLNDVTGLRDERMVALAARHGVAAVVMHMPVPDPATMMAHARYRDVVAEVKAFLEAQARRA
LSAGVPQVVLDPGFGFGKLLEHNLALLRRLDEIVALGHPVLVGLSRKRTIGELSGVEDPAQR
VHGSVAAHLFAVMKGVRLLRVHDVRAHREALGVWEALYGGDRPSRA
111111111111110000000000000000000000011111111111000000000000
000000000000011111111000000000000000000000000000000000000000
000000000000000000000000000000011111100000000000000000000000
000000000000000000000000000000000000000000000000000000000000
0000000000000000000000000000000000000011111111
>DM_train868
MVLRDKMILLDENTKAIVQGITGRQGSFHTKKMLECGTKIVGGVTPGKGGQNVHGVPVFDTV
KEAVKETDANASVIFVPAPFAKDAVFEAIDAGIELIVVITEHIPVHDTMEFVNYAEDVGVKI
IGPNTPGIASPKVGKLGIIPMEVLKEGSVGMVSRSGTLTYEIAHQIKKAGFGVSTCVGIGGD
PIVGLRYKEVLDLFEKDDETEAIVMIGEIGGGAEEEAAKFIEKMKKPVIGYIAGQSAPEGKR
MGHAGAIVEKGKGTAESKMKALEEAGAYVAKNISDIPKLLAGILGK
111111000000000000000000000000000000000000000000000000000000
000000000000000000000000000000000000000000000000000000000000
000000000000000000000000000000000000000000000000000000000000
000000000000000000000000000000000000000000000000000001111111
1111111111110000000000000000000000000000000001
>DM_train869
PVYEHLLPVNGAWRQDVTNWLSEDVPSFDFGGYVVGSDLKEANLYCKQDGMLCGVPFAQEVF
NQCELQVEWLFKEGSFLEPSKNDSGKIVVAKITGPAKNILLAERTALNILSRSSGIATASHK
IISLARSTGYKGTIAGTRKTTPGLRRLEKYSMLVGGCDTHRYDLSSMVMLKDNHIWATGSIT
NAVKNARAVCGFAVKIEVECLSEDEATEAIEAGADVIMLDNFKGDGLKMCAQSLKNKWNGKK
HFLLECSGGLNLDNLEEYLCDDIDIYSTSSIHQGTPVIDFSLKLAH
000000000000000000000000000000000000000000000000000000000000
000000000000000000000000000000000000000000000000000000000000
000000000000000000000000000000000000000000000000000000000000
000000000000000000000000000000000000000111111111111111111111
0000000000001111110000000000000000000000000000

>DM_train870
MDQECIENYAKVNGIYIYYKLCKAPEEKAKLMTMHGGPGMSHDYLLSLRDMTKEGITVLFYD
QFGCGRSEEPDQSKFTIDYGVEEAEALRSKLFGNEKVFLMGSSYGGALALAYAVKYQDHLKG
LIVSGGLSSVPLTVKEMNRLIDELPAKYRDAIKKYGSSGSYENPEYQEAVNYFYHQHLLRSE
DWPPEVLKSLEYAERRNVYRIMNGPNEFTITGTIKDWDITDKISAIKIPTLITVGEYDEVTP
NVARVIHEKIAGSELHVFRDCSHLTMWEDREGYNKLLSDFILKHL
111000000000000000000000000000000000000000000000000000000000
000000000000000000000000000000000000000000000000000000000000
000000000000000000000000000000000000000000000000000000000000
000000000000000000000000000000000000000000000000000000000000
000000000000000000000000000000000000000000000
>DM_train871
MTTNLTNSNCVEEYKENGKTKIRIKPFNALIELYHHQTPTGSIKENLDKLENYVKDVVKAKG
LAIPTSGAFSNTRGTWFEVMIAIQSWNYRVKRELNDYLIIKMPNVKTFDFRKIFDNETREKL
HQLEKSLLTHKQQVRLITSNPDLLIIRQKDLIKSEYNLPINKLTHENIDVALTLFKDIEGKC
KWDSLVAGVGLKTSLRPDRRLQLVHEGNILKSLFAHLKMRYWNPKAEFKYYGASSEPVSKAD
DDALQTAATHTIVNVNSTPERAVDDIFSLTSFEDIDKMLDQIIKK
111000000000000000000000000000000000000000000000000000000000
000000000000000000000000000000000000000000000000000000000000
000000000000000000000000000000000000000000000000000000000000
000000000000000000000000000000000000000000000000000000000000
000000000000000000000000000000000000000000000
>DM_train872
TNLLPNIKEDKMLTLRREIKSPSKSALDSFTLIMQTYNRTDLLLRLLNHYQAVPSLHKVIVV
WNNVGEKGPEELWNSLGPHPIPVIFKPQTANKMRNRLQVFPEVETNAVLMVDDDTLISAQDL
VFAFSIWQQFPDQIIGFVPRKHVSTSSGIYSYGGFELQTPGPGNGDQYSMVLIGASFFNSKY
LELFQKQPAAVHALIDETQNCDDIAMNFLVTRHTGKPSGIFVKPINMVNLEKETNGYSGMWH
RAEHFLQRSYCINKLVNIYDGMPLKYSNIMISQFGFPYANHKSKM
111111111111111111111111000000000000000000000000000000000000
000000000000000000000000000000000000000000000000000000000000
000000000000000000000000000000000000000000000000000000000000
000000000000000000000000000000000000000000000000111111111111
110000000000000000000000000000000000000000111
>DM_train873
TAKNLHPLLGEKLNLARIENQHHFQSYLTAESPAYLSQHQVFNKVLFPATGYLEIAAAVGKN
LLTTGEQVVVSDVTIVRGLVIPETDIKTVQTVISTLENNSYKLEIFSTSEGDNQQANQWTLH
AEGKIFLDSTTNTKAKIDLEQYQRECSQVIDIQQHYQQFKSRGIDYGNSFQGIKQLWKGQGK
ALGKIALPEEIAGQATDYQLHPALLDAALQILGHAIGNTETDDKAYLPVGIDKLKQYRQTIT
QVWAIVEIPENTLKGSIKLVDNQGSLLAEIEGLRVTATTADALLK
111100000000000000000000000000000000000000000000000000000000
000000000000000000000000000000000000000000000000011111000000
000000001111100000000000000000000000000000000000000000000000
000000000000000000000000000000000000000000000000000000000000
000000000000000000000000000000000000000000000

>DM_train874
MLSHMVLTRQDIGRAASYYEDGADDYYAKDGDASEWQGKGAEELGLSGEVDSKRFRELLAGN
IGEGHRIMRSATRQDSKERIGLDLTFSAPKSVSLQALVAGDAEIIKAHDRAVARTLEQAEAR
AQARQKIQGKTRIETTGNLVIGKFRHETSRERDPQLHTHAVILNMTKRSDGQWRALKNDEIV
KATRYLGAVYNAELAHELQKLGYQLRYGKDGNFDLAHIDRQQIEGFSKRTEQIAEWYAARGL
DPNSVSLEQKQAAKVLSRAKKTSVDREALRAEWQATAKELGIDFS
00000000000000000001111111111100000000000000000000000000000000
00000000000000000000000000000000000000000000000000000000000000
00000000000000000000000000000000000000000000000000000000000000
00000000000000000000000000000000000000000000000000000000000000
000000000000000000000000000000000000000000000
>DM_train875
MGSSSSGRENLYFQGQHLLKNPGILDKIIYAAKIKSSDIVLEIGCGTGNLTVKLLPLAKKVI
TIDIDSRMISEVKKRCLYEGYNNLEVYEGDAIKTVFPKFDVCTANIPYKISSPLIFKLISHR
PLFKCAVLMFQKEFAERMLANVGDSNYSRLTINVKLFCKVTKVCNVNRSSFNPPPKVDSVIV
KLIPKESSFLTNFDEWDNLLRICFSRKRKTLHAIFKRNAVLNMLEHNYKNWCTLNKQVPVNF
PFKKYCLDVLEHLDMCEKRSINLDENDFLKLLLEFNKKGIHFFNI
11111111111111100000000000000000000000000000000000000000000000
00000000000000000000000000011110000000000000000000000000000000
00000000000000000000000000000000000000000000000000000000000000
00000000000000000000000000000000000000000000000000000000000000
000000000000000000000000000000000000000000011
>DM_train876
EHEHNYHPNAVAAGLRAGRTRSIGLVIPDLENTSYTRIANYLERQARQRGYQLLIACSEDQP
DNEMRCIEHLLQRQVDAIIVSTSLPPEHPFYQRWANDPFPIVALDRALDREHFTSVVGADQD
DAEMLAEELRKFPAETVLYLGALPELSVSFLREQGFRTAWKDDPREVHFLYANSYEREAAAQ
LFEKWLETHPMPQALFTTSFALLQGVMDVTLRRDGKLPSDLAIATFGDNELLDFLQCPVLAV
AQRHRDVAERVLEIVLASLDEPRKPKPGLTRIKRNLYRRGVLSRS
11111111111111111000000000000000000000000000000000000000000000
00000000000000000000000000000000000000000000000000000000000000
00000000000000000000000000000000000000000000000000000000000000
00000000000000000000000000000000000000000000000000000000000000
000000000000000000001110000000000000000000001
>DM_train877
ISDPVELLKRAEKKGVPSSGFMKLFSGSDSYKFEEAADLCVQAATIYRLRKELNLAGDSFLK
AADYQKKAGNEDEAGNTYVEAYKCFKSGGNSVNAVDSLENAIQIFTHRGQFRRGANFKFELG
EILENDLHDYAKAIDCYELAGEWYAQDQSVALSNKCFIKCADLKALDGQYIEASDIYSKLIK
SSMGNRLSQWSLKDYFLKKGLCQLAATDAVAAARTLQEGQSEDPNFADSRESNFLKSLIDAV
NEGDSEQLSEHCKEFDNFMRLDKWKITILNKIKESIQQQEDDLL
00000000000000000000000000000000000000000000000000000000000000
00000000000000000000000000000000000000000000000000000000000000
00000000000000000000000000000000000000000000000000000000000000
00000000000000000000000000000000000000011111111100000000000000
00000000000000000000000000000000000000000000

>DM_train879
MASSMRSLFSDHGKYVESFRRFLNHSTEHQCMQEFMDKKLPGIIGRIGDTKSEIKILSIGGG
AGEIDLQILSKVQAQYPGVCINNEVVEPSAEQIAKYKELVAKISNLENVKFAWHKETSSEYQ
SRMLEKKELQKWDFIHMIQMLYYVKDIPATLKFFHSLLGTNAKMLIIVVSGSSGWDKLWKKY
GSRFPQDDLCQYITSDDLTQMLDNLGLKYECYDLLSTMDISDCFIDGNENGDLLWDFLTETC
NFNATAPPDLRAELGKDLQEPEFSAKKEGKVLFNNTLSFIVIEA
111111111111000000000000000000000000000000000000000000000000
000000000000000000000000000000000000000000000000000000000000
000000000000000000000000000000000000000000000000000000000000
000000000000000000000000000000000000000000000000000000000000
00000000000000000000000000000000000000000000
>DM_train880
HRPADVSALGVRGAEHPLLLAAVDVPGHGGAVFTGRLSTDEQPWLAEHVVGGRTLVPGSVLV
DLALAAGEDVGLPVLEELVLQRPLVLAGAGALLRMSVGAPDESGRRTIDVHAAEDVADLADA
QWSQHATGTLAQGVAAGPRDTEQWPPEDAVRIPLDDHYDGLAEQGYEYGPSFQALRAAWRKD
DSVYAEVSIAADEEGYAFHPVLLDAVAQTLSLGALGEPGGGKLPFAWNTVTLHASGATSVRV
VATPAGADAMALRVTDPAGHLVATVDSLVVRSTGEKWEQPEPRG
111100000000000000000000000000000000000000000000000000000000
000000000000000000000000000000000000000000000000000000000000
000000000000000000000000000000000000000000000000000000000000
000000000000000000000000000000000000011110000000000000000000
00000000000000000000000000000001111111111111
>DM_train881
TGTDFHVGIVGLGSMGMGAARSCLRAGLSTWGADLNPQACANLLAEGACGAAASAREFAGVV
DALVILVVNAAQVRQVLFGEDGVAHLMKPGSAVMVSSTISSADAQEIAAALTALNLNMLDAP
VSGGAVKAAQGEMTVMASGSEAAFTRLKPVLDAVASNVYRISDTPGAGSTVKIIHQLLAGVH
IAAAAEAMALAARAGIPLDVMYDVVTHAAGNSWMFENRMQHVVDGDYTPRSAVDIFVKDLGL
VADTAKALRFPLPLASTALNMFTSASNAGYGKEDDSAVIKIFSG
111000000000000000000000000000000000000000000000000000000000
000000000000000000110000000000000000000000000000000000000000
000000000000000000000000000000000000000000000000000000000000
000000000000000000000000000000000000000000000000000000000000
00000000000000000000000000000000000000000011
>DM_train882
MGSSSSGRENLYFQGHMKVIMTTKVDKASMNIMNKLIENFGFKETEYVFEGNPVYKRGDVLI
LTTNDEMIYYDYLDREIENQLGFKPEIIAFASRHSSKQKLPALTTHVTGNWGKAMYGGKDES
FAVAIPSAMKLSLLKMSELNDLGWTVCYEATHHGPTELEVPSFFIEIGSSEEEWINDRAGEI
IAETIIYVLDNYEKGRSKFKVALGIGGGHYAPKQTKRALEGDLAFGHILPKYAQPVSRDVMI
KALNRFGEKVEAIYVDWKGSRGETRQLAKSLAQELGLEFIKDGS
111111111111000000000000000000000000000000000000000000000000
000000000000000000000000000000000000000000000000000000000000
000000000000000000000000000000000000000000000000000000000000
000000000000000001000000000000000000000000000000000000000000
00000000000000000000000000000000000000000001

>DM_train884
MSKSWGKFIEEEEAEMASRRNLMIVDGTNLGFRFKHNNSKKPFASSYVSTIQSLAKSYSART
TIVLGDKGKSVFRLEHLPEYAGNRDEKYAQRTEEEKALDEQFFEYLKDAFELCKTTFPTFTI
RGVEADDMAAYIVKLIGHLYDHVWLISTDGDWDTLLTDKVSRFSFTTRREYHLRDMYEHHNV
DDVEQFISLKAIMGDLGDNIRGVEGIGAKRGYNIIREFGNVLDIIDQLPLPGKQKYIQNLNA
SEELLFRNLILVDLPTYCVDAIAAVGQDVLDKFTKDILEIAEQ
111111111111111111000000000000000011111110000000000000000000
000000000000000000000111111111111111111110000000000000000000
000000000000000000000000000000000000000000000000000000000000
000000000000000000000000000000000000000000000000000000000000
0000000000000000000000000000000000000000000
>DM_train885
MDYLVKALAYDGKVRAYAARTTDMVNEGQRRHGTWPTASAALGRTMTASLMLGAMLKGDDKL
TVKIEGGGPIGAIVADANAKGEVRAYVSNPQVHFDLNAAGKLDVRRAVGTNGTLSVVKDLGL
REFFTGQVEIVSGELGDDFTYYLVSSEQVPSSVGVGVLVNPDNTILAAGGFIIQLMPGTDDE
TITKIEQRLSQVEPISKLIQKGLTPEEILEEVLGEKPEILETMPVRFHCPCSKERFETAILG
LGKKEIQDMIEEDGQAEAVCHFCNEKYLFTKEELEGLRDQTTR
000000000000000000000000000000000000000000000000000000000000
000000000000000000000000000000000000000000000000000000000000
000000000000000000000000000000000000000000000000000000000000
000000000000000000000000000000000000000000000000000000000000
0000000000000000000000000000000000000011111
>DM_train888
MACLSPSQLQKFQQDGFLVLEGFLSAEECVAMQQRIGEIVAEMDVPLHCRTEFSTQEEEQLR
AQGSTDYFLSSGDKIRFFFEKGVFDEKGNFLVPPEKSINKIGHALHAHDPVFKSITHSFKVQ
TLARSLGLQMPVVVQSMYIFKQPHFGGEVSPHQDASFLYTEPLGRVLGVWIAVEDATLENGC
LWFIPGSHTSGVSRRMVRAPVGSAPGTSFLGSEPARDNSLFVPTPVQRGALVLIHGEVVHKS
KQNLSDRSRQAYTFHLMEASGTTWSPENWLQPTAELPFPQLYT
100000000000000000000000000000000000000000000000000000000000
000000000000000000000000000000000000000000000000000000000000
000000000000000000000000000000000000000000000000000000000000
000000000000000000001111000000000000000000000000000000000000
0000000000000000000000000000000000000000000
>DM_train890
AEPESCGTVRFSDVGWTDITATTATATTILEALGYETDVKVLSVPVTYTSLKNKDIDVFLGN
WMPTMEADIAPYREDKSVETVRENLAGAKYTLATNAKGAELGIKDFKDIAAHKDELDGKIYG
IEPGNDGNRLIIDMVEKGTFDLKGFEVVESSEQGMLAQVARAEKSGDPIVFLGWEPHPMNAN
FKLTYLSGGDDVFGPNYGGATVHTNVRAGYTTECPNVDKLLQNLSFSLQMENEIMGKILNDG
EDPEKAAAAWLKDNPQSIEPWLSGVATKDGGDGLAAVKAALGL
111000000000000000000000000000000000000000000000000000000000
000000000000000000000000000000000000000000000000000000000000
000000000000000000000000000000000000000000000000000000000000
000000000000000000000000000000000000000000000000000000000000
0000000000000000000000000000000000000000000

>DM_train891
QDLARIEQFLDALWLEKNLAENTLNAYRRDLSMMVEWLHHRGLTLATAQSDDLQALLAERLE
GGYKATSSARLLSAVRRLFQYLYREKFREDDPSAHLASPKLPQRLPKDLSEAQVERLLQAPL
IDQPLELRDKAMLEVLYATGLRVSELVGLTMSDISLRQGVVRVIGKGNKERLVPLGEEAVYW
LETYLEHGRPWLLNGVSIDVLFPSQRAQQMTRQTFWHRIKHYAVLAGIDSEKLSPHVLRHAF
ATHLLNHGADLRVVQMLLGHSDLSTTQIYTHVATERLRQLHQ
000000000000000000000000000000000000000000000000000000000001
111111000000000000000000000000000111111111000000000000000000
000000000000000000000000000000000000000000000000000000000000
000000000000000000000000000000000000000000000000000000000000
0000000000000000000000000000000000000000
>DM_train892
MRSHILGKIELDQTRLAPDLAYLAAVPTVEEEYDEFSNGFWKHVPLWNASGDSEDRLYRDLK
DAAAQPTAHVEHVPYLKEIVTTVFDGTHLQMARSRNLKNAIVIPHRDFVELDREVDRYFRTF
MVLEDSPLAFHSNEDTVIHMRPGEIWFLDAATVHSAVNFSEISRQSLCVDFAFDGPFDEKEI
FADATLYAPGSTPDLPERRPFTAEHRRRILSLGQVIERENFRDILFLLSKVHYKYDVHPSET
YDWLIEISKQAGDEKMVVKAEQIRDFAVEARALSERFSLTSW
000000000000000000000000000001111000000000000000111111111111
111110000000000000000000000000000000000000000000111111100000
000000000000000000000000000000000000000000000000001111111111
111111000000000000000000000000000000000000000000000000000000
0000000000000000000000000000000000000000
>DM_train893
ATALAGSNGVCIAYYITDGRNPTFKLKDIPDKVDMVILFGLKYWSLQDTTKLPGGTGMMGSF
KSYKDLDTQIRSLQSRGIKVLQNIDDDVSWQSSKPGGFASAAAYGDAIKSIVIDKWKLDGIS
LDIEHSGAKPNPIPTFPGYAATGYNGWYSGSMAATPAFLNVISELTKYFGTTAPNNKQLQIA
SGIDVYAWNKIMENFRNNFNYIQLQSYGANVSRTQLMMNYATGTNKIPASKMVFGAYAEGGT
NQANDVEVAKWTPTQGAKGGMMIYTYNSNVSYANAVRDAVKN
111111100000000000000000000000000000000000000000000000000000
000000000000000000000000000000000000000000000000000000000000
000000000000000000000000000000000000000000000000000000000000
000000000000000000000000000000000000000000000000000000000000
0000000000000000000000000000000000000000
>DM_train895
MDRKTRLSEPPTLALRLKPYKTAIQQLRSVIRALKENTTVTFLPTPSLILQTVRSHCVSKIT
FNSSCLYITDKSFQPKTINNSTPLLGNFMYLTSSKDLTKFYVQDISDLSAKISMCAPDFNME
FSSACVHGQDIVRESENSAVHVDLDFGVVADLLKWIGPHTRVKRNVKKAPCPTGTVQILVHA
GPPAIKFILTNGSELEFTSNNRVSFHGVKNMRINVQLKNFYQTLLNCAVTKLPCTLRIVTEH
DTLLYVASRNGLFAVENFLTEEPFQRGDPFDKNYVGNSGKSR
111111111000000000000000000000000000000000000000000000000000
000000000000000000000000000000000000000000000000000000000000
000000000000000000000000000000000000011111111111000000000000
000000000000000000000000000000000000000000000000000000000000
0000000000000000000001111111111111111111

>DM_train896
GMEQATRTIYSEYAAYPETQGIIAVEKRQPRDSLTDQFDVLLLVITRDPSVEWTVKHYRLNT
LRVSLHLVHEQVLSRWLILNANRRAVHWVSEGTIIFERNDYLTDLKKQLRNFPETERCLQMS
LSFAKLLRRFQDGRNLFSRGNYYDAYTHVHHALHHLARLSVLEKGAHPEVVVWEQARLDDPD
VYKLYEQLLLSEETLEQRIHLALIGLEHLLQSKVLSGGKYLFEVMRERDRPWTMHELMEESR
LTELKVDLGSLVDFFIRKGLIRISYQRTKGLGVELVTYEPVV
00000000000000000000000000000000000000000000000000000000000000
00000000000000000000000000000000000000000000000000000000000000
00000000000000000000000000000000000000000000000000000000000000
00000000000000000000000000000000000000000000000000000000000000
000000000000000000000000000000000000000001
>DM_train897
AYSLNNWNKPIKNSVTTKQWMSALPDTTNLAALSIPGTHDTMSYNGDITWTLTKPLAQTQTM
SLYQQLEAGIRYIDIRAKDNLNIYHGPIFLNASLSGVLETITQFLKKNPKETIIMRLKDEQN
SNDSFDYRIQPLINIYKDYFYTTPRTDTSNKIPTLKDVRGKILLLSENHTKKPLVINSRKFG
MQFGAPNQVIQDDYNGPSVKTKFKEIVQTAYQASKADNKLFLNHISATSLTFTPRQYAAALN
NKVEQFVLNLTSEKVRGLGILIMDFPEKQTIKNIIKNNKFN
11111111111111000000000000000000000000000000000000000000000000
00000000000000000000000000000000000000000000000000000000000000
00000000000000000000000000000000000000000000000000000000000000
00000000000000000000000000000000000000000000000000000000000000
00000000000000000000000000000000000000000
>DM_train898
MANGYTYEDYQDTAKWLLSHTEQRPQVAVICGSGLGGLVNKLTQAQTFDYSEIPNFPESTVP
GHAGRLVFGILNGRACVMMQGRFHMYEGYPFWKVTFPVRVFRLLGVETLVVTNAAGGLNPNF
EVGDIMLIRDHINLPGFSGENPLRGPNEERFGVRFPAMSDAYDRDMRQKAHSTWKQMGEQRE
LQEGTYVMLGGPNFETVAECRLLRNLGADAVGMSTVPEVIVARHCGLRVFGFSLITNKVIMD
TESQGKANHEEVLEAGKQAAQKLEQFVSLLMASIPVSGHTG
00000000000000000000000000000000000000000000000000000000000000
00000000000000000000000000000000000000000000000000000000000000
00000000000000000000000000000000000000000000000000000000000000
00000000000000000000000000000000000000000000000000000000000111
11111111111111111100000000000000000011111
>DM_train899
SFLKHVQDCNTHDLSNFVRFVIEGRRVGWVRKALAQRLKAHGRVFDVTRDAVLLSASLRTPQ
SRTRAVADVVDRLADEGVVPAPRGELYRVNQSWGEPTLMLLDRAVVPTFGVRAYGVHLNGYV
GAGADLHLWIGRRSPDKSVAPGKLDNMVAGGQPADLSLRQNLIKECAEEADLPEALARQAIP
VGAITYCMESPAGIKPDTLFLYDLALPEDFRPHNTDGEMADFMLWPAAKVVEAVRTTEAFKF
NVNLTVIDFAIRHGLIDPDNEPDYQEILAGLRGRPRERAAS
00000000000000000000000000000000000000000000000000000000000000
00000000000000000000000000000000000000000000000000000000000000
00000000000000000000000000000000000000000000000000000000000000
00000000000000000000000000000000000000000000000000000000000000
00000000000000000000000000000000011111111

>DM_train900
NAMPIEIITDSGADLPQSYIREHRIAFLPLVVHWNGQDYKDGITIEPKQVYDAMRQGHTVKT
AQPSPLAMKELFLPYAKENRPCLYIAFSSKLSGTYQTAMAVRSELLDEYPEFRLTIIDSKCA
SLGQGLAVMKAVELAKQNTPYNLLCETIESYCRHMEHIFTVDNLDYLARGGRISKTAAAFGG
LLNIKPLLHVEDGALIPLEKWRGRKKVLKRMVELMGERGDDLQKQTIGISHADDEETALELK
QMIEETHGCTRFFLSDIGSAIGAHAGPGTIALFFLNKYIEI
110000000000000000000000000000000000000000000000000000000000
000000000000000000000000000000000000000000000000000000000000
000000000000000000000000000000000000000000000000000011111111
110000000000000000000000000000000000000000000000000000000000
00000000000000000000000000000000000000000
>DM_train903
MPKERVDVLAYKQGLFETREQAKRGVMAGLVVNVINGERYDKPGEKIDDGTELKLKGEKLRY
VSRGGLKLEKALAVFNLSVEDMITIDIGASTGGFTDVMLQNGAKLVYAVDVGTNQLVWKLRQ
DDRVRSMEQYNFRYAEPVDFTEGLPSFASIDVSFISLNLILPALAKILVDGGQVVALVKPQF
EAGREQIGKNGIVRESSIHEKVLETVTAFAVDYGFSVKGLDFSPIQGGHGNIEFLAHLEKTD
SPQNDVPTSIKEVVAQAHKEFKKNEEERSFGVNPKDCPRK
000000000000000000000000000000000000000000000000000000000000
000000000000000000000000000000000000000000000000000000000000
000000000000000000000000000000000000000000000000000000000000
000000001000000000000000000000000000000000000000000000000000
0000000000000000000000000001111111111111
>DM_train904
VIQNSLNDKIVTISCKADTNLFFYQVAGNVSLFQQTRNYLERWRLIYDSNKAAYKIKSMDIH
NTNLVLTWNAPTHNISTQQDSNADNQYWLLLKDIGNNSFIIASYKNPNLVLYADTVARNLKL
STLNNSNYIKFIIEDYIISDLNNFTCKISPILDLNKVVQQVDVTNLNVNLYTWDYGRNQKWT
IRYNEEKAAYQFFNTILSNGVLTWIFSNGNTVRVSSSNDQNNDAQYWLINPVSDTDETYTIT
NLRDTTKALDLYGGQTANGTAIQVFNYHGDDNQKWNIRNP
111100000000000000000000000000000000000000000000000000000000
000000000000000000000000000000000000000000000000000000000000
000000000000000000000000000000000000000000000000000000000000
000000000000000000000000000000000000000000000000000000000000
0000000000000000000000000000000000000000
>DM_train905
MSVQLTPHFGNVQAHYDLSDDFFRLFLDPTQTYSCAYFERDDMTLQEAQIAKIDLALGKLNL
EPGMTLLDIGCGWGATMRRAIEKYDVNVVGLTLSENQAGHVQKMFDQMDTPRSRRVLLEGWE
KFDEPVDRIVSIGAFEHFGHQRYHHFFEVTHRTLPADGKMLLHTIVRPTFKEGREKGLTLTH
ELVHFTKFILAEIFPGGWLPSIPTVHEYAEKVGFRVTAVQSLQLHYARTLDMWATALEANKD
QAIAIQSQTVYDRYMKYLTGCAKLFRQGYTDVDQFTLEK
111111111111111000000000000000000000000000000000000000000000
000000000000000000000000000000000000000000000000000000000000
000000000000000000000000000000000000000000000000001111111111
100000000000000000000000000000000000000000000000000000000000
000000000000000000000000000000000000000

>DM_train906
MSKQLLALNIDGALLRSNGKIHQATKDAIEYVKKKGIYVTLVTNRHFRSAQKIAKSLKLDAK
LITHSGAYIAEKIDAPFFEKRISDDHTFNIVQVLESYQCNIRLLHEKYSIGNKKKVNSNLLG
KALIHPSDPIFYPVQFVESLSDLLMDEPVSAPVIEVYTEHDIQHDITETITKAFPAVDVIRV
NDEKLNIVPKGVSKEAGLALVASELGLSMDDVVAIGHQYDDLPMIELAGLGVAMGNAVPEIK
RKADWVTRSNDEQGVAYMMKEYFRMQQRKGFLDKFHMKR
110000000000000000000000000000000000000000000000000000000000
000000000000000000000000000000000000000000000000000000000000
000000000000000000000000000000000000000000000000000000000000
000000000000000000000000000000000000000000000000000000000000
000000000000000000000000000001111111111
>DM_train907
GSCSSKPANTAKGEKGSDKTGKDLELKLSFQEGIAPGESLNEKLDFMEKLGVVGFEPGGGL
AGRVNEIKQALNGRNIKVSAICAGFKGFILSTDPAIRKECMDTMKEIIAAAGELGSTGVIIV
PAFNGQVPALPHTMETRDFLCEQFNEMGTFAAQHGTSVIFEPLNRKECFYLRQVADAASLCR
DINNPGVRCMGDFWHMTWEETSDMGAFISGGEYLQHVHVASRKRRSMPGEDGDADNYINGFK
GLKMIGYNNYVSFECGCQGDRNVVVPAAVKLLREQWEQA
111111111111111111111100000000000000000000000000000000000000
000000000000000000000000000000000000000000000000000000000000
000000000000000000000000000000000000000000000000000000000000
000000000000000000000000000000000000000000000000000000000000
000000000000000000000000000000000000000
>DM_train908
MADFEDRVSDEEKVRIAAKFITHAPPGEFNEVFNDVRLLLNNDNLLREGAAHAFAQYNMDQF
TPVKIEGYDDQVLITEHGDLGNGRFLDPRNKISFKFDHLRKEASDPQPEDTESALKQWRDAC
DSALRAYVKDHYPNGFCTVYGKSIDGQQTIIACIESHQFQPKNFWNGRWRSEWKFTITPPTA
QVAAVLKIQVHYYEDGNVQLVSHKDIQDSVQVSSDVQTAKEFIKIIENAENEYQTAISENYQ
TMSDTTFKALRRQLPVTRTKIDWNKILSYKIGKEMQNA
111100000000000000000000000000000000000000000000000000000000
000000000000000000000000000000000000000000000000000000000000
000000000000000000000000000000000000000000000000000000000000
000000000000000000000000000000000000000000000000000000000000
00000000000000000000000000000111111111
>DM_train909
GMTDAPADAPLDADARRAVKPVICYPNDSLPRPDLALYRAARASARKTGEVLVPPREGRCFE
VKAGQFFRISSVEGPQVGDLNLHNLHDLTERFFSGKTRALHGTHVTTGERLWSNLPYLRPMA
TIIEDTLGWYGIDQYGGSVHDVIGTRCDPYTGNLLAGGHYHHCCHSNLTRALADHTGLPLHE
AEMLVHDVLNVFMCTGFTRDTGQYFMKASPVRPGDYLEFFAEIDLLGNLSACPGGDCSSEHS
SDTASCHPLLVEIFAPAEGMLGDWPSPSVNGYDRSHGR
111111100000000000000000000000000000000000000000000000000000
000000000000000000000000000000000000000000000000000000000000
000000000000000000000000000000000000000000000000000000000000
000000000000000000000000000000000000000000000000000000000011
11100000000000000000000000000000000000

>DM_train910
GTHMKSQIKRQRMIYHCKFGEFGVMEGQFTEPSGVAVNAQNDIIVADTNNHRIQIFDKEGRF
KFQFGECGKRDSQLLYPNRVAVVRNSGDIIVTERSPTHQIQIYNQYGQFVRKFGATILQHPR
GVTVDNKGRIIVVECKVMRVIIFDQNGNVLHKFGCSKHLEFPNGVVVNDKQEIFISDNRAHC
VKVFNYEGQYLRQIGGEGITNYPIGVGINSNGEILIADNHNNFNLTIFTQDGQLISALESKV
KHAQCFDVALMDDGSVVLASKDYRLYIYRYVQLAPVGM
111111000000000000000000000000000000000000000000000000000000
000000000000000000000000000000000000000000000000000000000000
000000000000000000000000000000000000000000000000000000000000
000000000000000000000000000000000000000000000000000000000000
00000000000000000000000000000000000000
>DM_train912
MGSDKIHHHHHHTHSVETPYGSVTFTVYGTPKPKRPAIFTYHDVGLNYKSCFQPLFRFGDMQ
EIIQNFVRVHVDAPGMEEGAPVFPLGYQYPSLDQLADMIPCILQYLNFSTIIGVGVGAGAYI
LSRYALNHPDTVEGLVLINIDPNAKGWMDWAAHKLTGLTSSIPDMILGHLFSQEELSGNSEL
IQKYRGIIQHAPNLENIELYWNSYNNRRDLNFERGGETTLKCPVMLVVGDQAPHEDAVVECN
SKLDPTQTSFLKMADSGGQPQLTQPGKLTEAFKYFLQG
111111100000000000000000000000000000000000000000000000000000
000000000000000000000000000000000000000000000000000000000000
000000000000000000000000000000000000000000000000000000000000
000000000000000000000000000000000000000000000000000000000000
00000000000000000000000000000000000000
>DM_train913
MGYAHPEVLVSTDWVQEHLEDPKVRVLEVDEDILLYDTGHIPGAQKIDWQRDFWDPVVRDFI
SEEEFAKLMERLGISNDTTVVLYGDKNNWWAAYAFWFFKYNGHKDVRLMNGGRQKWVEEGRP
LTTEVPSYPPGRYEVPYRDESIRAYRDDVLEHIIKVKEGKGALVDVRSPQEYRGELTHMPDY
PQEGALRAGHIPGAKNIPWAKAVNPDGTFKSAEELRALYEPLGITKDKDIVVYCRIAERSSH
SWFVLKYLLGYPHVKNYDGSWTEWGNLVGVPIAKGEE
100000000000000000000000000000000000000000000000000000000000
000000000000000000000000000000000000000000000000000000000000
000000000000000000000000000000000000000000000000000000111111
110000000000000000000000000000000000000000000000000000000000
0000000000000000000000000000000000000
>DM_train914
MRPMTHVPVLYQEALDLLAVRPGGVYVDATLGGAGHARGILERGGRVIGLDQDPEAVARAKG
LHLPGLTVVQGNFRHLKRHLAALGVERVDGILADLGVSSFHLDDPSRGFSYQKEGPLDMRMG
LEGPTAKEVVNRLPLEALARLLRELGEEPQAYRIARAIVAAREKAPIETTTQLAEIVRKAVG
FRRAGHPARKTFQALRIYVNDELNALKEFLEQAAEVLAPGGRLVVIAFHSLEDRVVKRFLRE
SGLKVLTKKPLVPSEKEAAQNPRARSAKLRAAEKEAP
111100000000000000000000000000000000000000000000000000000000
000000000000000000000000000000000000000000000000000000000000
000000000000000000000000000000000000000000000000000000000000
000000000000000000000000000000000000000000000000000000000000
0000000000000000000000000000000000001

>DM_train915
MFKDFFNRTKKKKYLTVQDSKNNDVPAGIMTKCPKCKKIMYTKELAENLNVCFNCDHHIALT
AYKRIEAISDEGSFTEFDKGMTSANPLDFPSYLEKIEKDQQKTGLKEAVVTGTAQLDGMKFG
VAVMDSRFRMGSMGSVIGEKICRIIDYCTENRLPFILFSASGGARMQEGIISLMQMGKTSVS
LKRHSDAGLLYISYLTHPTTGGVSASFASVGDINLSEPKALIGFAGRRVIEQTINEKLPDDF
QTAEFLLEHGQLDKVVHRNDMRQTLSEILKIHQEVTK
111111111111111111111111111000000000000000000000000000000000
000000000000000000000000000000000000000000000000000000000000
000000000000000000000000000000000000000000000000000000000000
000000000000000000000000000000000000000000000000000000000000
0000000000000000000000000000000000011
>DM_train916
MGSSSSGLVPRGSHMASAPALTKSQTDRLEVLLNPKDEISLNSGKPFRELESELLSRRKKDL
QQIYAEERENYLGKLEREITRFFVDRGFLEIKSPILIPLEYIERMGIDNDTELSKQIFRVDK
NFCLRPMLAPNLYNYLRKLDRALPDPIKIFEIGPCYRKESDGKEHLEEFTMLNFCQMGSGCT
RENLESIITDFLNHLGIDFKIVGDSCMVYGDTLDVMHGDLELSSAVVGPIPLDREWGIDKPW
IGAGFGLERLLKVKHDFKNIKRAARSESYYNGISTNL
111111111111111110000000000000000000111111000000000000000000
000000000000000000000000000000000000000000000001110000000000
000000000000000000000000000000000000000000000000000000000000
000000000000000000000000000000000000000000000000000000000000
0000000000000000000000000000000000000
>DM_train917
MNSLLFVYGTLRKHEKNHHLLAQSACINEQARTKGSLFAAKEGPTVVFNDEDEGYIYGEVYE
ADELCIHKLDQFFQGYHKQTVFVETDVGIKIALIYFMNKDGCAGFTKISSGDWKEHQMISKS
KNPIYYFAYGSCMDNARFQKAGVDHYFQDPVGRAVLKGYTTRFTLKREDGSRADMLEDGGTT
EGVLYRIPYSALSYLYKREGVESLTYRPAFVDVEAGGRHYKDCLTFLVLQKEAEIAPPQHYQ
IEIERGAELYLSPEFTEKLKRHMNSLPKGLEHHHHHH
000000000000000000000000000000000000000110000000000000000000
000000000000000000000000000000000000111111000000000000000000
000000000000000000000000000000000000000000000000000000000000
000000000000000000000000000000000000000000000000000000000000
0000000000000000000000000000011111111
>DM_train918
MTNVTGDYTDCTPLLGDRAALDSFYEEHGYLFLRNVLDRDLVKTVAEQMREGLVALGAADPH
ATLEELTIDSFESVDEVAMHDYVKYDAFWNNPSTIKVFEQVFGEPVFVFLSTTIRYYPSQAG
SEEPSFHYLTPFHQDGFYIGPNQDFRTFWIPLIRTTRESGGVALADGSHRRGKRDHVLNESF
RRFGHPVRGIPPTEVSEDEHLLHSPMEPGDILLFHAHMCHKSIPNLSKDPRLMRMSMDTRVQ
PAKSHRGFNAMTPWTESAKDASKGIMAKITGTPTDVE
100000000000000000000000000000000000000000000000000000000000
000000000000000000000000000000000000000000000000000000000000
000000000000000000000000000000000000000000000000000000000000
000000000000000000000000000000000000000000000000000000000000
0000000000000000011111111111111111111

>DM_train919
MNYNPEEQFRCTIIRGKAKNMLDNLLPAYANIIDDICPCDKASFVKDFNNRLIEILGEETTK
KTLDNHRTEIAGKLFGMFYEDDEVIFPSGRTNKYIEDSDQPAFFKDICFKFQFPNGMDKLDK
VIEKVGAKIQIRQFPYILQVLLTADNNNIQLSKDDIAYYVLNSLQVLQGKIKPIEVIEKIIE
DRSNDITKKVRHPGKETSYSMQHIREQLNYLELANLIRIDGNLVKLNYREAENINYIAQFWG
NKPEFNAYKYDFTSEDDKKSFFKDWQQYYSNVNSHKS
00000000000000000000000000000000000000000000000000000000000000
00000000000000000000000000000000000000000000000000000000000000
00000000000000000000000000000000000000000000000000000000000000
00000000000000000000000000000000000000000000000000000000000000
0000000000000000000000000000000011111
>DM_train920
GAMDTRKNVSGAERKIWSLIRDCSGKLEGVTETSVLEVLLIVSRVLGIRKEDLFLKDLGVSP
TEEKRILELVEKRASGYPLHYILGEKEFMGLSFLVEEGVFVPRPETEELVELALELIRKYGI
KTVADIGTGSGAIGVSVAKFSDAIVFATDVSSKAVEIARKNAERHGVSDRFFVRKGEFLEPF
KEKFASIEMILSNPPYVKSSAHLPKDVLFEPPEALFGGEDGLDFYREFFGRYDTSGKIVLME
IGEDQVEELKKIVSDTVFLKDSAGKYRFLLLNRRSS
11111111111110000000000000000000000000000000000000000000000000
00000000000000000000000000000000000000000000000000000000000000
00000000000000000000000000000000000000000000000000000000000000
00000000000000000000011111100000000000000000000000000000000000
000000000000000000000000000000000000
>DM_train921
MRGSHHHHHHGSMDAQSAAKCLTAVRRHSPLVHSITNNVVTNFTANGLLALGASPVMAYAKE
EVADMAKIAGALVLNIGTLSKESVEAMIIAGKSANEHGVPVILDPVGAGATPFRTESARDII
REVRLAAIRGNAAEIAHTVGVTDWLIKGVDAGEGGGDIIRLAQQAAQKLNTVIAITGEVDVI
ADTSHVYTLHNGHKLLTKVTGAGSLLTSVVGAFCAVEENPLFAAIAAISSYGVAAQLAAQQT
ADKGPGSFQIELLNKLSTVTEQDVQEWATIERVTVS
11111111111100000000000000000000000000000000000000000000000000
00000000000000000000000000000000000000000000000000000000000000
00000000000000000000001111111111111000000000000000000000000000
00000000000000000000000000000000000000000000000000000000000000
000000000000000000000000000000000000
>DM_train922
MAGVDPFTGNSAYRSAASKTMNIYFPKKEAVTFDQANPTQILGKLKELNGTAPEEKKLTDDL
ILLEKILSLICNSSSEKPTVQQLQILWKAINCPEDIVFPALDILRLSIKHPSVNENFCNEKE
GAQFSSHLINLLNPKGKPANQLLALRTFCNCFVGQAGQKLMMSQRESLMSHAIELKSGSNKN
IHIALATLALNYSVCFHKDHNIEGKAQCLSLISTILEVVQDLEATFRLLVALGTLISDDSNA
VQLAKSLGVDSQIKKYSSVSEPAKVSECCRFILNLL
11111111111111111111000000000000000000000000000000000000000000
00000000000000000000000000000000000000000000000000000000000000
00000000000000000000000000000000000000000000000000000000010000
00000000000000000001100000000000000000000000000000000000000000
000000000000000000000000000000000000

>DM_train923
MTGSLNRHSLLNGVKKMRIILCDTNEVVTNLWQESIPHAYIQNDKYLCIHHGHLQSLMDSMR
KGDAIHHGHSYAIVSPGNSYGYLGGGFDKALYNYFGGKPFETWFRNQLGGRYHTVGSATVVD
LQRCLEEKTIECRDGIRYIIHVPTVVAPSAPIFNPQNPLKTGFEPVFNAMWNALMHSPKDID
GLIIPGLCTGYAGVPPIISCKSMAFALRLYMAGDHISKELKNVLIMYYLQYPFEPFFPESCK
IECQKLGIDIEMLKSFNVEKDAIELLIPRRILTLDL
11111111111111100000000000000000000001111111000000000000000000
00001111000000000000000000000000000000000000000000000000000000
00000011110000000000000000000000000000000000000000000000000000
00000000000000000000000000000000000000000000000000000000000000
000000000000000000000000000000011111
>DM_train924
TAQIKGLTQAQRNANDGISLAQTAEGALGEISNNLQRIRELAVQASNGTNTQTDRDALQAEV
TQLQSEIQRVAEQTSFNGQKLLDGSFNGVQFQIGANAGETIGVSKIMNAQTASLGGSLTRTT
STIDATDLTKYDTAMAAGDLTINGVDVGKIDAASTAQERAAQLTEAINRVSSQTNVGASYDK
TTGQVTLTSNAAIAVAGAANDATVAGWANNATTGTATTTTGINSLTVSSFTNAQQTITQIDN
ALKDINTARADLGAVQNRFTSTVANLQSMTENLSSA
11111110000000000000000000000000000000000000000000000000000000
00000000000000000000000000000000000000000000000000000000000000
00000000000000000000000000000000000000000000000000000000000000
00000000000000000000000000000000000000000000000000000000000000
000000000000000000000000111111111111
>DM_train926
AVDRDTAKNSSPIAGNIEYTISTPGSNYAVGDKITVKYVSDDIETEGKITEVDADGKIKKIN
IPTAKIIAKAKEVGEYPTLGSNWTAEISSSSSGLAAVITLGKIITDSGILLAEIENAEAAMT
AVDFQANLKKYGIPGVVALYPGELGDKIEIEIVSKADYAKGASALLPIYPGGGTRASTAKAV
FGYGPQTDSQYAIIVRRNDAIVQSVVLSTKRGGKDIYDSNIYIDDFFAKGGSEYIFATAQNW
PEGFSGILTLSGGLSSNAEVTAGDLMEAWDFFADR
11110000000000000000000000000000000000000000000000000000000000
00000000000000000000000000000000000000000000000000000000000000
00000000000000000000000000000000000000000000000000000000000000
00000000000000000000000000000000000000000000000000000000000000
00000000000000000000000000000011111
>DM_train927
GTEDTVIKVSVLRGPSVIAFADWLENPPIIDNKKVQVKVVDSPDLAQALLIKQETDIAVLPM
INAANLYNKGIKIKLAGCPIWGTLYLVEKTPLKEPALYVFGNGTTPDILTRYYLGRQRLDYP
LNYAFNTAGEITQGILAGKVNRAVLGEPFLSIALRKDSSLRITADLNHLTDNDTLGFAQTAV
VYTPTMEKYRIAFEDALRASCQKAVRYPKETIHSLEEHGIFAQGALTPKSIERCKIYYLSAI
EAKDAVMGFLRLIEQYEPKAVGGRLPDAGFIPEKQ
11100000000000000000000000000000000000000000000000000000000000
00000000000000000000000000000000000000000000000000000000000000
00000000000000000000000000000000000000000000000000000000000000
00000000000000000000000000000000000000000000000000000000000000
00000000000000000000000000000000000

>DM_train928
MTGTLLSFGHGYTARVLSRALAPQGWRIIGTSRNPDQMEAIRASGAEPLLWPGEEPSLDGVT
HLLISTAPDSGGDPVLAALGDQIAARAAQFRWVGYLSTTAVYGDHDGAWVDETTPLTPTAAR
GRWRVMAEQQWQAVPNLPLHVFRLAGIYGPGRGPFSKLGKGGIRRIIKPGQVFSRIHVEDIA
QVLAASMARPDPGAVYNVCDDEPVPPQDVIAYAAELQGLPLPPAVDFDKADLTPMARSFYSE
NKRVRNDRIKEELGVRLKYPNYRVGLEALQADAET
10000000000000000000000000000000000000000000000000000000000000
00000000000000000000000000000000000000000000000000000000000000
00000000000000000000000000000000000111110000000000000000000000
00000000000000000000000000000000000000000000000000000000000000
00000000000000000000000000000000000
>DM_train929
TITDDQIEVTDATELVQSSSTGKICNNPHRILDGIDCTLIDALLGDPHCDVFQDETWDLFVE
RSKAFSNCYPYDVPDYASLRSLVASSGTLEFITEGFTWTGVTQNGGSNACKRGPGSGFFSRL
NWLTKSGSTYPVLDVTMPNNDNFDKLYIWGIHHPSTNQEQTSLYVQASGRVTVSTRRSQQTI
IPNIGSRPWVRGLSSRISIYWTIVKPGDVLVINSNGNLIAPRGYFKMRTGKSSIMRSDAPID
TCISECITPDGSIPNDKPFQNVNKITYGACPKYV
11111111111111100000000000000000000000000000000000000000000000
00000000000000000000000000000000000000000000000000000000000000
00000000000000000000000000000000000000000000000000000000000000
00000000000000000000000000000000000000000000000000000000000000
0000000000000000000000000000000000
>DM_train930
MSGADRSPNAGAAPDSAPGQAAVASAYQRFEPRAYLRNNYAPPRGDLCNPNGVGPWKLRCLA
QTFATGEVSGRTLIDIGSGPTVYQLLSACSHFEDITMTDFLEVNRQELGRWLQEEPGAFNWS
MYSQHACLIEGKGECWQDKERQLRARVKRVLPIDVHQPQPLGAGSPAPLPADALVSAFCLEA
VSPDLASFQRALDHITTLLRPGGHLLLIGALEESWYLAGEARLTVVPVSEEEVREALVRSGY
KVRDLRTYIMPAHLQTGVDDVKGVFFAWAQKVGL
11111111111111111111100000000000000000000000000000000000000000
00000000000000000000000000000000000000000000000000000000000000
00000000000000000000000000000000000000000000000000000000000000
00000000000000000000000000000000000000000000000000000000000000
0000000000000000000000000000000000
>DM_train931
GAKESEQKVTIDSAKHEKHTKDKEENNSANTVFFDKINDLLVASVKEFEGTVGISYLDLETG
EQRSVNGQHEFYTASTIKVPLTMLVADTVASGQKKWTDLIPYNAEEDYEEGTGIIAYNIQPE
YPLKTLQEYAITYSDNIAKNMLYDTLGGDAKAKREMYQRYLHKTPSIEEPQFSSEDALVILQ
KLYTEKATKPDYQAIYDSMKQSVFHERMETPTTQGKVAHKIGSYDEFIHDMGILETPHPFAL
AIFTKGPDNAKSAAFIASVTDKLWQLQVSEYPNQ
11111111111111111111111111100000000000000000000000000000000000
00000000000000000000000000000000000000000000000000000000000000
00000000000000000000000000000000000000000000000000000000000000
00000000000000000000000000000000000000000000000000000000000000
0000000000000000000000000000000000

>DM_train932
MDIAQDWNGIPVPANPGNGMTWQLQDNVSDSFNYTSSEGNRPTAFTSKWKPSYINGWTGPGS
TIFNAPQAWTNGSQLAIQAQPAGNGKSYNGIITSKNKIQYPVYMEIKAKIMDQVLANAFWTL
TDDETQEIDIMEGYGSDRGGTWFAQRMHLSHHTFIRNPFTDYQPMGDATWYYNGGTPWRSAY
HRYGCYWKDPFTLEYYIDGVKVRTVTRAEIDPNNHLGGTGLNQATNIIIDCENQTDWRPAAT
QEELADDSKNIFWVDWIRVYKPVAVSGGGNNSLE
111000000000000000000000000000000000000000000000000000000000
000000000000000000000000000000000000000000000000000000000000
000000000000000000000000000000000000000000000000000000000000
000000000000000000000000000000000000000000000000000000000000
0000000000000000000000000111111111
>DM_train933
GSAMDVRQSIHSAHAKTLDTQGLRNEFLVEKVFVADEYTMVYSHIDRIIVGGIMPITKTVSV
GGEVGKQLGVSYFLERRELGVINIGGAGTITVDGQCYEIGHRDALYVGKGAKEVVFASIDTG
TPAKFYYNCAPAHTTYPTKKVTPDEVSPVTLGDNLTSNRRTINKYFVPDVLETCQLSMGLTE
LAPGNLWNTMPCHTHERRMEVYFYFNMDDDACVFHMMGQPQETRHIVMHNEQAVISPSWSIH
SGVGTKAYTFIWGMVGENQVFDDMDHVAVKEICA
000000000000000000000000000000000000000000000000000000000000
000000000000000000000000000000000000000000000000000000000000
000000000000000000000000000000000000000000000000000000000000
000000000000000000000000000000000000000000000000000000000000
0000000000000000000000000000000001
>DM_train934
MTVRKNQATLTADEKRRFVAAVLELKRSGRYDEFVRTHNEFIMSDTDSGERTGHRSPSFLPW
HRRFLLDFEQALQSVDSSVTLPYWDWSADRTVRASLWAPDFLGGTGRSTDGRVMDGPFAAST
GNWPINVRVDSRTYLRRSLGGSVAELPTRAEVESVLAISAYDLPPYNSASEGFRNHLEGWRG
VNLHNRVHVWVGGQMATGVSPNDPVFWLHHAYVDKLWAEWQRRHPDSAYVPTGGTPDVVDLN
ETMKPWNTVRPADLLDHTAYYTFDALEHHHHHH
100000000000000000000000000000000000000000000000000000000000
000000000000000000000000000000000000000000000000000000000000
000000000000000000000000000000000000000000000000000000000000
000000000000000000000000000000000000000000000000000000000000
000000000000000000000000001111111
>DM_train936
MSNQLERGPVRTPHADVLLASVGERGVLCDFYDEGAADTYRDLIQDADGTSEAREFATRTGP
VSGPVLELAAGMGRLTFPFLDLGWEVTALELSTSVLAAFRKRLAEAPADVRDRCTLVQGDMS
AFALDKRFGTVVISSGSINELDEADRRGLYASVREHLEPGGKFLLSLAMSEAAESEPLERKQ
ELPGRSGRRYVLHVRHLPAEEIQEITIHPADETTDPFVVCTHRRRLLAPDQVVRELVRSGFD
VIAQTPFASGGAGRKDMVLVEAVMPGATADAR
111111111110000000111111111111111111111111111000000000000001
000000000000000000000000000000000000000000000000000000000000
000000000000000000000000000000000000000000000000000000000000
000111100000000000000000001111111111110000000000000000000000
00000000000000000000000001111111

>DM_train937
MTTTESPTAAGSGSAATDKFKAERATADTSPERLAAIAKDALGALNDVILKHGVTYPEYRVF
KQWLIDVGEGGEWPLFLDVFIEHSVEEVLARSRKGTMGSIEGPYYIENSPELPSKCTLPMRE
EDEKITPLVFSGQVTDLDGNGLAGAKVELWHADNDGYYSQFAPHLPEWNLRGTIIADEEGRY
EITTIQPAPYQIPTDGPTGQFIEAQNGHPWRPAHLHLIVSAPGKESVTTQLYFKGGEWIDSD
VASATKPELILDPKTGDDGKNYVTYNFVLDPA
111111111111111111111110000000000000000000000000000000000000
000000000000000000000000000000000000000000000000000000000000
000000000000000000000000000000000000000000000000000000000000
000000000000000000000000000000000000000000000000000000000000
00000000000000000000000000000000
>DM_train938
MEDDPVWRFDDRDVILYNIALGATTKQLKYVYENDSDFQVIPTFGHLITFNSGKSQNSFAKL
LRNFNPMLLLHGEHYLKVHSWPPPTEGEIKTTFEPIATTPKGTNVVIVHGSKSVDNKSGELI
YSNEATYFIRNCQADNKVYADRPAFATNQFLAPKRAPDYQVDVPVSEDLAALYRLSGDRNPL
HIDPNFAKGAKFPKPILHGMCTYGLSAKALIDKFGMFNEIKARFTGIVFPGETLRVLAWKES
DDTIVFQTHVVDRGTIAINNAAIKLVGDKAKI
111100000000000000000000000000000000000000000000001111000000
000000000000000000000000000000000000000000000000000000000000
000000000000000000000000000000000000000000000000000000000000
000000000000000000000000000000000000000000000000000000000000
00000000000000000000000000011111
>DM_train939
GPMNPGPLELQLGDIFIAVKTTWAFHRSRLDLLLDTWVSRIRQQTFIFTDSPDERLQERLGP
HLVVTQCSAEHSHPALSCKMAAEFDAFLVSGLRWFCHVDDDNYVNPKALLQLLKTFPQDRDV
YVGKPSLNRPIHASELQSKQRTKLVRFWFATGGAGFCINRQLALKMVPWASGSHFVDTSALI
RLPDDCTVGYIIECKLGGRLQPSPLFHSHLETLQLLGAAQLPEQVTLSYGVFEGKLNVIKLP
GPFSHEEDPSRFRSLHCLLYPDTPWCPLLAAP
111111110000000000000000000000000000000000000000000000000000
000000011111110000000000000000000000000000000000000000000000
000000011111111111111111110000000000000000000000000000000000
000000000000000000000000000000000000000000000000000000000000
00000000000000000000000000000111
>DM_train940
MDIKINDITLGNNSPFVLFGGINVLESLDSTLQTCAHYVEVTRKLGIPYIFKASFDKANRSS
IHSYRGVGLEEGLKIFEKVKAEFGIPVITDVHEPHQCQPVAEVCDVIQLPAFLARQTDLVVA
MAKTGNVVNIKKPQFLSPSQMKNIVEKFHEAGNGKLILCERGSSFGYDNLVVDMLGFGVMKQ
TCGNLPVIFDVTHSLQTRDAGSAASGGRRAQALDLALAGMATRLAGLFLESHPDPKLAKCDG
PSALPLHLLEDFLIRIKALDDLIKSQPILTIE
000000000000000000000000000000000000000000000000000000000000
000000000000000000000000000000000000000000000000000000000000
000000000000000000000000000000000000000000000000000000000000
000000000000000011111111110000000000000000000000001111111111
11111000000000000000000000000111

>DM_train942
MATHKPINILEAFAAAPPPLDYVLPNMVAGTVGALVSPGGAGKSMLALQLAAQIAGGPDLLE
VGELPTGPVIYLPAEDPPTAIHHRLHALGAHLSAEERQAVADGLLIQPLIGSLPNIMAPEWF
DGLKRAAEGRRLMVLDTLRRFHIEEENASGPMAQVIGRMEAIAADTGCSIVFLHHASKGAAM
MGAGDQQQASRGSSVLVDNIRWQSYLSSMTSAEAEEWGVDDDQRRFFVRFGVSKANYGAPFA
DRWFRRHDGGVLKPAVLERQRKSKGVPRGEA
000000000000000000000000000000000000000000000000000000000000
000000000000000000000000000000000000000000000000000000000000
000000000000000000000000000000000000000000000000000000111111
111111111111110000000000000000000000000000000000000000000000
0000000000000001111111111111111
>DM_train944
MKKNRAFLKWAGGKYPLLDDIKRHLPKGECLVEPFVGAGSVFLNTDFSRYILADINSDLISL
YNIVKMRTDEYVQAARELFVPETNCAEVYYQFREEFNKSQDPFRRAVLFLYLNRYGYNGLCR
YNLRGEFNVPFGRYKKPYFPEAELYHFAEKAQNAFFYCESYADSMARADDSSVVYCDPPYAP
LSATANFTAYHTNSFTLEQQAHLAEIAEGLVERHIPVLISNHDTMLTREWYQRAKLHVVKVR
RSISSNGGTRKKVDELLALYKPGVVSPAKK
110000000000000000000000000000000000000000000000000000000000
000000000000000000000000000000000000000000000000000000000000
000000000000000000000000000000000000000000000000000000000000
011111111111000000000000000000000000000000000000000000000011
111111111110000000000011111111
>DM_train945
MSDQQLDCALDLMRRLPPQQIEKNLSDLIDLVPSLCEDLLSSVDQPLKIARDKVVGKDYLLC
DYNRDGDSYRSPWSNKYDPPLEDGAMPSARLRKLEVEANNAFDQYRDLYFEGGVSSVYLWDL
DHGFAGVILIKKAGDGSKKIKGCWDSIHVVEVQEKSSGRTAHYKLTSTVMLWLQTNKTGSGT
MNLGGSLTRQMEKDETVSDSSPHIANIGRLVEDMENKIRSTLNEIYFGKTKDIVNGLRSIDA
IPDNQKYKQLQRELSQVLTQRQIYIQPDN
000000000000000000000000000000000000000000000000000000000000
000000000000000000000000000000000000000000000000000000000000
000000000000000000000000000000000000000000000000000000000000
000000000000000000000000000000000000000000000000000000000000
00000000000000000000000111111
>DM_train946
GMGGAALPDNGWPADRIDTESAHSARIYDYIIGGKDYYPADKEAGDAMSREWPALPVHMRAN
RDWMNRAVAHLAKEAGIRQFLDIGTGIPTSPNLHEIAQSVAPESRVVYVDNDPIVLTLSQGL
LASTPEGRTAYVEADMLDPASILDAPELRDTLDLTRPVALTVIAIVHFVLDEDDAVGIVRRL
LEPLPSGSYLAMSIGTAEFAPQEVGRVAREYAARNMPMRLRTHAEAEEFFEGLELVEPGIVQ
VHKWHPDAATADGIRDEDIAMYGAVARKP
111111111111111000000000000000000000000000000000000000000000
000000000000000000000000000000000000000000000000000000000000
000000000000000000000000000000000000000000000000000000000000
000000000000000000000000000000000000000000000000000000000000
00000000000000000000000000000

>DM_train948
GMQVEQRTLNTAAHPFQITAYWLDQISDFETAVDYPIMIICPGGGFTYHSGREEAPIATRMM
AAGMHTVVLNYQLIVGDQSVYPWALQQLGATIDWITTQASAHHVDCQRIILAGFSAGGHVVA
TYNGVATQPELRTRYHLDHYQGQHAAIILGYPVIDLTAGFPTTSAARNQITTDARLWAAQRL
VTPASKPAFVWQTATDESVPPINSLKYVQAMLQHQVATAYHLFGSGIHGLALANHVTQKPGK
DKYLNDQAAIWPQLALRWLQEQGLLAGNY
000000000000000000000000000000000000000000000000000000000000
000000000000000000000000000000000000000000000000000000000000
000000000000000000000000000000000000000000000000000000000000
000000000000000000000000000000000000000000001111111111111111
00000000000000000000000000111
>DM_train949
SNKKQSNRLTEQHKLSQGVIGIFGDYAKAHDLAVGEVSKLVKKALSNEYPQLSFRYRDSIKK
TEINEALKKIDPDLGGTLFVSNSSIKPDGGIVEVKDDYGEWRVVLVAEAKHQGKDIINIRNG
LLVGKRGDQDLMAAGNAIERSHKNISEIANFMLSESHFPYVLFLEGSNFLTENISITRPDGR
VVNLEYNSGILNRLDRLTAANYGMPINSNLCINKFVNHKDKSIMLQAASIYTQGDGREWDSK
IMFEIMFDISTTSLRVLGRDLFEQLTSK
111111111111111000000000000000000000000000000000000000000000
000000000000000000000000000000000000000000000000000000001111
111100000000000000000000000000000000000000000000000000000000
000000000000000000000000000000000000000000000000000000000000
0000000000000000000000000000
>DM_train951
GSAMAKANSSFSEVQIARRIKEGRGQGHGKDYIPWLTVQEVPSSGRSHRIYSHKTGRVHHLL
SDLELAVFLSLEWESSVLDIREQFPLLPSDTRQIAIDSGIKHPVIRGVDQVMSTDFLVDCKD
GPFEQFAIQVKPAAALQDERTLEKLELERRYWQQKQIPWFIFTDKEINPVVKENIEWLYSVK
TEEVSAELLAQLSPLAHILQEKGDENIINVCKQVDIAYDLELGKTLSEIRALTANGFIKFNI
YKSFRANKCADLCISQVVNMEELRYVAN
111111111000000000000000000000000000000000000000000000000000
000000000000000000000000000000000000000000000000000000000000
000000000000000000000000000000000000000000000000000000000000
000000000000000000000000000000000000000000000000000000000000
0000000000000000000000011111
>DM_train952
MSWVQLAGHTGSFKAAGTSGLILKRCSEPERYCLARLMADALRGCVPAFHGVVERDGESYLQ
LQDLLDGFDGPCVLDCKMGVRTYLEEELTKARERPKLRKDMYKKMLAVDPEAPTEEEHAQRA
VTKPRYMQWREGISSSTTLGFRIEGIKKADGSCSTDFKTTRSREQVLRVFEEFVQGDEEVLR
RYLNRLQQIRDTLEVSEFFRRHEVIGSSLLFVHDHCHRAGVWLIDFGKTTPLPDGQILDHRR
PWEEGNREDGYLLGLDNLIGILASLAER
000000011110000000000000000000000000000000000000000000000000
000000000000000000000000000000000000000000000000000000000000
000000000000000000000000000000000000000000000000000000000000
000000000000000000000000000000000000000000000000000000000000
0000000000000000000000000000

>DM_train953
SLRCMQCKTNGDCRVEECALGQDLCRTTIVRLWEEGEELELVEKSCTHSEKTNRTLSYRTGL
KITSLTEVVCGLDLCNQGNSGRAVTYSRSRYLECISCGSSDMSCERGRHQSLQCRSPEEQCL
DVVTHWIQEGEEGRPKDDRHLRGCGYLPGCPGSNGFHNNDTFHFLKCCNTTKCNEGPILELE
NLPQNGRQCYSCKGNSTHGCSSEETFLIDCRGPMNQCLVATGTHEPKNQSYMVRGCATASMC
QHAHLGDAFSMNHIDVSCCTKSGCNHPD
000000000000000000000000000000000111000000000000000000000000
000000000000000000011111111111000000000000000000000000000000
000000111111111100000000000000000000000000000000000000000000
000000000000000000000000000000000000000000000000000000000000
0111000000000000000000000000
>DM_train954
MTDTTNSIKHVISPLARQTLQDRDLTRPVAGKRPIRLLPWLQVVKIGGRVMDRGADAILPLV
EELRKLLPEHRLLILTGAGVRARHVFSVGLDLGLPVGSLAPLAASEAGQNGHILAAMLASEG
VSYVEHPTVADQLAIHLSATRAVVGSAFPPYHHHEFPGSRIPPHRADTGAFLLADAFGAAGL
TIVENVDGIYTADPNGPDRGQARFLPETSATDLAKSEGPLPVDRALLDVMATARHIERVQVV
NGLVPGRLTAALRGEHVGTLIRTGVRPA
111111111111111111111111111110000000000000000000000000000000
000000000000000000000000000000000000000000000000000000000000
000000000000000000000000000000000000000000000000000000000000
000000000000000000000000000000000000000000000000000000000000
0000000000000000000000000000
>DM_train955
MAHHHHHHGICSSAPTLGEIWKRKLNQLDAKEFMAYRRRFVVEVDRNEAREALAKGKTNTGH
AVSRGTAKLAWIDERGGVELKGTVVDLGCGRGSWSYYAASQPNVREVKAYTLGTSGHEKPRL
VETFGWNLITFKSKVDVTKMEPFQADTVLCDIGESNPTAAVEASRTLTVLNVISRWLEYNQG
CGFCVKVLNPYSCDVLEALMKMQARFGGGLIRVPLSRNSTHEMYFVSGIKNNIMGNVTAVSR
QLLKRMEEQGGERVVPDYKFSTGTRSNL
111111111111000000000000000000000000000001111111111111111111
100000000000000000000000000000000000000000000000000000000000
000000000000000000000000000000000000000000000000000000000000
000000000000000000000000000000000000000000000000000000000000
0000000111111111111111111111
>DM_train956
MTDTYLHETLVFDNKLSYIDNQRDTDGPAILLLPGWCHDHRVYKYLIQELDADFRVIVPNWR
GHGLSPSEVPDFGYQEQVKDALEILDQLGVETFLPVSHSHGGWVLVELLEQAGPERAPRGII
MDWLMWAPKPDFAKSLTLLKDPERWREGTHGLFDVWLDGHDEKRVRHHLLEEMADYGYDCWG
RSGRVIEDAYGRNGSPMQMMANLTKTRPIRHIFSQPTEPEYEKINSDFAEQHPWFSYAKLGG
PTHFPAIDVPDRAAVHIREFATAIRQGQ
100000000000000000000000000000000000000000000000000000000000
000000000000000000000000000000000000000000000000000000000000
000000000000000000000000000000000000000000000000000000000000
000000000000000000000000000000000000000000000000000000000000
0000000000000000000000000011

>DM_train957
MSYYHHHHHHLESTSLYKAGLTRKFFVGGNWKMNGDYASVDGIVTFLNASADNSSVDVVVAP
PAPYLAYAKSKLKAGVLVAAQNCYKVPKGAFTGEISPAMIKDLGLEWVILGHSERRHVFGES
DALIAEKTVHALEAGIKVVFCIGEKLEEREAGHTKDVNFRQLQAIVDKGVSWENIVIAYEPV
WAIGTGKTASGEQAQEVHEWIRAFLKEKVSPAVADATRIIYGGSVTADNAAELGKKPDIDGF
LVGGASLKPDFVKIINARSTALSCTCW
111111111111111111111100000000000000000000000000000000000000
000000000000000000000000000000000000000000000000000000000000
000000000000000000000000000000000000000000000000000000000000
000000000000000000000000000000000000000000000000000000000000
000000000000000000011111111
>DM_train958
MERLAPAKVNLGLSVRFRREDGYHELHTLFAPFSLADRLVVEPVSSGLHFQGPYGRENLAYR
AASLYLEAAGQPGGVRILLEKRIPEGAGLGGGSSDAAQVLLALQALYPAEVDLFALARTLGA
DVPFFLLGRGAEARGVGERLKPLALPPVPAVVFFPGLRVPTPLVYRAVRPEDFGPDLPVEAI
LEALARGEEPPYWNSLEGPAFRLFPELKEVRGRMRALGLRGVLMSGSGSAFFGLAEGPDHAR
RAAEALRAWGRAWAGTLGGGDAGSGPA
000000000000000000000000000000000000000000000000000000000000
000000000000000000000000000000000000000000000000000000000000
000000000000000000000000000000000000000000000000000000000000
000000000000000000000000000000000000000000000000000000000000
000000000000000000001111111
>DM_train959
GSQSSERVVIGSKPFNEQYILANMIAILLEENGYKAEVKEGLGGTLVNYEALKRNDIQLYVE
YTGTAYNVILRKQPPELWDQQYIFDEVKKGLLEADGVVVAAKLGFRDDYALAVRADWAEENG
VEKISDLAEFADQLVFGSDPEFASRPDGLPQIKKVYGFEFKEVKQMEPTLMYEAIKNKQVDV
IPAYTTDSRVDLFNLKILEDDKGALPPYDAIIIVNGNTAKDEKLISVLKLLEDRIDTDTMRA
LNYQYDVEKKDAREIAMSFLKEQGLVK
111110000000000000000000000000000000000000000000000000000000
000000000000000000000000000000000000000000000000000000000000
000000000000000000000000000000000000000000000000000000000000
000000000000000000000000000000000000000000000000000000000000
000000000000000000000000000
>DM_train960
MTEIKYKVITKDAFALPYTIIKAKNQPTKGVIVYIHGGGLMFGKANDLSPQYIDILTEHYDL
IQLSYRLLPEVSLDCIIEDVYASFDAIQSQYSNCPIFTFGRSSGAYLSLLIARDRDIDGVID
FYGYSRINTEPFKTTNSYYAKIAQSINETMIAQLTSPTPVVQDQIAQRFLIYVYARGTGKWI
NMINIADYTDSKYNIAPDELKTLPPVFIAHCNGDYDVPVEESEHIMNHVPHSTFERVNKNEH
DFDRRPNDEAITIYRKVVDFLNAITMV
110000000000000000000000000000000000000000000000000000000000
000000000000000000000000000000000000000000000000000000000000
000000000000000000000000000000000000000000000000000000000000
000000000000000000000000000000000000000000000000000000000000
000000000000000000000000001

>DM_train961
MGSSSSGLVPRGSMANYTEKFAAWSVICLTDHTFLDENGTEDDIRELCNESVKTCPFAAAVC
VYPKFVKFINEKIKQEINPFKPKIACVINFPYGTDSMEKVLNDTEKALDDGADEIDLVINYK
KIIENTDEGLKEATKLTQSVKKLLTNKILKVIIEVGELKTEDLIIKTTLAVLNGNADFIKTS
TGKVQINATPSSVEYIIKAIKEYIKNNPEKNNKIGLKVSGGISDLNTASHYILLARRFLSSL
ACHPDNFRIGSSSLVIKLRKVISQCPL
11111111111111100000000000000000000000000000000000000000000000
00000000000000000000000000000000000000000000000000000000000000
00000000000000000000000000000000000000000000000000000000000000
00000000000000000000000000000000000000000000000000000000000111
111111000000000000000001111
>DM_train962
GMEKRLDYITDLMALGPTANARTIQRRQTAHRLAIAEAWQVKPGEKILEIGCGQGDLSAVLA
DQVGSSGHVTGIDIASPDYGAPLTLGQAWNHLLAGPLGDRLTVHFNTNLSDDLGPIADQHFD
RVVLAHSLWYFASANALALLFKNMAAVCDHVDVAEWSMQPTALDQIGHLQAAMIQGLLYAIA
PSDVANIRTLITPDTLAQIAHDNTWTYTAGTIVEDPTLDDAHWEIATTNALLTELKLSTDLR
DRVKPLLEAMSHNGTASLATFTGRITF
10000000000000000000000000000000000000000000000000000000000000
00000000000000000000000000000000000000000000000000000000000000
00000000000000000000000000000000000000000000000000000000000000
00000000000000000000000000000000000000000000000000000000000000
000000000000000000000000000
>DM_train964
QTIKDFLAVAMKKWTAPFEPFQLIDNIYYVGTDGIAVYVIKTSQGLILMDTAMPQSTGMIKD
NIAKLGFKVADIKLILNTHAHLDHTGGFAEIKKETGAQLVAGERDKPLLEGGYYPGDEKNED
LAFPAVKVDRAVKEGDRVTLGDTTLTAHATPGHSPGCTSWEMTVKDGKEDREVLFFCSGTVA
LNRLVGQPTYAGIVDDYRATFAKAKAMKIDVLLGPHPEVYGMQAKRAEMKDGAPNPFIKPGE
LVTYATSLSEDFDKQLAKQTAALEKK
11111111000000000000000000000000000000000000000000000000000000
00000000000000000000000000000000000000000000000000000000000000
00000000000000000000000000000000000000000000000000000000000000
00000000000000000000000000000000000000000000000000000000000000
00000000000000000000000011
>DM_train965
MAEPTAKAAPYENLMVPSPSMGRDIPVAFLAGGPHAVYLLDAFNAGPDVSNWVTAGNAMNTL
AGKGISVVAPAGGAYSMYTNWEQDGSKQWDTFLSAELPDWLAANRGLAPGGHAAVGAAQGGY
GAMALAAFHPDRFGFAGSMSGFLYPSNTTTNGAIAAGMQQFGGVDTNGMWGAPQLGRWKWHD
PWVHASLLAQNNTRVWVWSPTNPGASDPAAMIGQAAEAMGNSRMFYNQYRSVGGHNGHFDFP
ASGDNGWGSWAPQLGAMSGDIVGAIR
11111110000000000000000000000000000000000000000000000000000000
00000000000000000000000000000000000000000000000000000000000000
00000000000000000000000000000000000000000000000000000000000000
00000000000000000000000000000000000000000000000000000000000000
00000000000000000000000000

>DM_train966
GQGKLISVKTDVLDLTINTRGGDVEQALLPAYPKELNSTQPFQLLETSPQFIYQAQSGLTGR
DGPDNPANGPRPLYNVEKDAYVLAEGQNELQVPMTYTDAAGNTFTKTFVLKRGDYAVNVNYN
VQNAGEKPLEISSFGQLKQSITLPPHLDTGSSNFALHTFRGAAYSTPDEKYEKYKFDTIADN
ENLNISSKGGWVAMLQQYFATAWIPHNDGTNNFYTANLGNGIAAIGYKSQPVLVQPGQTGAM
NSTLWVGPEIQDKMAAVAPHLDLTVD
111000000000000000000000000000000000000000000000000000000000
000000000000000000000000000000000000000000000000000000000000
000000000000000000000001111111111110000000000000000000000000
000000000000000000000000000000000000000000000000000000000000
00000000000000000000000000
>DM_train967
GAHMGDFLASLEQDIKPAFGTNQEDYASYIMNGIIKWGDPVTRVLDDGELLVQQTKNSDRTP
LVSVLLEGPPHSGKTALAAKIAEESNFPFIKICSPDKMIGFSETAKCQAMKKIFDDAYKSQL
SCVVVDDIERLLDYVPIGPRFSNLVLQALLVLLKKAPPQGRKLLIIGTTSRKDVLQEMEMLN
AFSTTIHVPNIATGEQLLEALELLGNFKDKERTTIAQQVKGKKVWIGIKKLLMLIEMSLQMD
PEYRVRKFLALLREEGASPLDFDLE
111111111111111000000000000000000000000000000000000000000000
000000000000000000000000000000000000000000000000000000000000
000000000000000000000000000000000000000000000000000000000000
000000000000000000000000000000000000000000000000000000000000
0000000000000011111111111
>DM_train968
MHDANIRVAIAGAGGRMGRQLIQAALALEGVQLGAALEREGSSLLGSDAGELAGAGKTGVTV
QSSLDAVKDDFDVFIDFTRPEGTLNHLAFCRQHGKGMVIGTTGFDEAGKQAIRDAAADIAIV
FAANFSVGVNVMLKLLEKAAKVMGDYTDIEIIEAHHRHKVDAPSGTALAMGEAIAHALDKDL
KDCAVYSREGHTGERVPGTIGFATVRAGDIVGEHTAMFADIGERLEITHKASSRMTFANGAV
RSALWLSGKESGLFDMRDVLDLNNL
111100000000000000000000000000000000000000000000000000000000
000000000000000000000000000000000000000000000000000000000000
000000000000000000000000000000000000000000000000000000000000
000000000000000000000000000000000000000000000000000000000000
0000000000000000000000000
>DM_train969
MPHDPGATRLLAISPHLDDAVLSFGAGLAQAAQDGANVLVYTVFAGAAQPPYSPAAQRMHTI
WGLAPDDDAVLYRRKEDIAALDHLRVAHRHGRFLDSIYRKLPDGRWLTAHVEGRQKLAVNDH
SPDSDHDLVGEVADDIRSIIDEFDPTLVVTCAAIGEHPDHEATRDAALFATHEKNVPVRLWE
DLPYAVFKSGAVELPQGFRLGSADVSSVKPEMRSQKFQAVERYSSQMVLLNGSENNLFDRLD
EHARQNAPHGGYGETTWPVVRSDDS
111111100000000000000000000000000000000000000000000000000000
000000000000000000000000000000000000000000000000000000000000
000000000000000000000000000000000000000000000000000000000000
000000000000000000000000000000000000000000000000000000000000
0000000000000000000000000

>DM_train970
MALKRMGIVSDYEKIRTFAVAIVGVGGVGSVTAEMLTRCGIGKLLLFDYDKVELANMNRLFF
QPHQAGLSKVQAAEHTLRNINPDVLFEVHNYNITTVENFQHFMDRISNGGLEEGKPVDLVLS
CVDNFEARMTINTACNELGQTWMESGVSENAVSGHIQLIIPGESACFACAPPLVVAANIDEK
TLKREGVCAASLPTTMGVVAGILVQNVLKFLLNFGTVSFYLGYNAMQDFFPTMSMKPNPQCD
DRNCRKQQEEYKKKVAALPKQEVIQ
111111111111100000000000000000000000000000000000111111111111
111111000000000000000000000000000000000000000000000000000000
000000000000000000000000000000000000000000000000000000011111
111111000000000000000000000000000000000000000000000000000000
0000000000000000111111111
>DM_train971
MGSDKIHHHHHHMSNTVRAISPDITLFNKTLTFQEISQNTREAVIYIHGGAWNDPENTPNDF
NQLANTIKSMDTESTVCQYSIEYRLSPEITNPRNLYDAVSNITRLVKEKGLTNINMVGHSVG
ATFIWQILAALKDPQEKMSEAQLQMLGLLQIVKRVFLLDGIYSLKELLIEYPEYDCFTRLAF
PDGIQMYEEEPSRVMPYVKKALSRFSIDMHLVHSYSDELLTLRQTNCLISCLQDYQLSFKLY
LDDLGLHNDVYKNGKVAKYIFDNIC
111111111111111111111000000000000000000000000000000000000000
000000000000000000000000000000000000000000000000000000000000
000000000000000000000000000000000000000000000000000000000000
000000000000000000000000000000000000000000000000000000000000
0000000000000000000000000
>DM_train972
MSSGVDLGTENLYFQSNAMRLRHLSDPDSLPALDKSFAIERPALGLAPDAPPVRILLLYGSL
RARSFSRLAVEEAARLLQFFGAETRIFDPSDLPLPDQVQSDDHPAVKELRALSEWSEGQVWC
SPERHGQITSVMKAQIDHLPLEMAGIRPTQGRTLAVMQVSGGSQSFNAVNTLRLLGRWMRMF
TIPNQSSIAKAFQEFDAAGRMKPSPYYDRIADVMEELVRFTALVRPHREALTDRYSERKAAG
HVIDEATDLSSIAIAPQPLPESETS
111111111111111100000000000000000000000000000000000000000000
000000000000000000000000000000000000000000000000000000000000
000000000000000000000000000000000000000000000000000000000000
000000000000000000000000000000000000000000000000000000000000
0001111111111111111111111
>DM_train973
TTVNETTPIAPAKVLPDAQTVTSVRHWTDTLFSFRVTRPQTLRFRSGEFVMIGLLDDNGKPI
MRAYSIASPAWDEELEFYSIKVPDGPLTSRLQHIKVGEQIILRPKPVGTLVIDALLPGKRLW
FLATGTGIAPFASLMREPEAYEKFDEVIMMHACRTVAELEYGRQLVEALQEDPLIGELVEGK
LKYYPTTTREEFHHMGRITDNLASGKVFEDLGIAPMNPETDRAMVCGSLAFNVDVMKVLESY
GLREGANSEPREFVVEKAFVGEGI
111111111111111000000000000000000000000000000000000000000000
000000000000000000000000000000000000000000000000000000000000
000000000000000000000000000000000000000000000000000000000000
000000000000000000000000000000000000000000000000000000000000
000000000000000000000000

>DM_train974
MEALRLGFSPCPNDTFIFYALVHGRVESPVPLEPVLEDVETLNRWALEGRLPLTKLSYAAYA
QVRDRYVALRSGGALGRGVGPLVVARGPLQALEGLRVAVPGRHTTAYFLLSLYAQGFVPVEV
RYDRILPMVAQGEVEAGLIIHESRFTYPRYGLVQVVDLGAWWEERTGLPLPLGAILARRDLG
EGLIRALDEAVRRSVAYALAHPEEALDYMRAHAQELSDEVIWAHVHTYVNAFSLDVGEEGER
AVARLFAEAEARGLAAPSPRPLFV
11000000000000000000000000000000000000000000000000000000000000
00000000000000000000000000000000000000000000000000000000000000
00000000000000000000000000000000000000000000000000000000000000
00000000000000000000000000000000000000000000000000000000000000
000000000000000000000000
>DM_train975
MVRNLVIDITKKPTQNIPPTNEIIEEAITELNVDELLDRLFEKDESGEVITPSRIAKMLEEK
AFEIYKEYEKQVREAYLSAGYSREKLEQSFQQARFSRGGKAFEIIFTKLLNKFGIRYEHDRV
IKIYDYITEGEKPAFIIPSVRTFLNDPSSAILITVKRKVRERWREAVGEAQILRNKFGDEIN
FWFVGFDEEFTIYSAIAMLDNGIDRVYVIDGRYDSLIEEIKRISDPNFNEDKYIQKIRRFSD
IFDDIIQFLNKHGNKKRGKQLTLV
11000000000000000000000000000000000000000000000000000000000000
00000000000000000000000000000000000000000000000000000000000000
00000000000000000000000000000000000000000000000000000000000000
00000000000000000000000000000000000000000000000000000000000000
000000000000111111111111
>DM_train976
APAPVKQGPTSVAYVEVNNNSMLNVGKYTLADGGGNAFDVAVIFAANINYDTGTKTAYLHFN
ENVQRVLDNAVTQIRPLQQQGIKVLLSVLGNHQGAGFANFPSQQAASAFAKQLSDAVAKYGL
DGVDFDDEYAEYGNNGTAQPNDSSFVHLVTALRANMPDKIISLYNIGPAASRLSYGGVDVSD
KFDYAWNPYYGTWQVPGIALPKAQLSPAAVEIGRTSRSTVADLARRTVDEGYGVYLTYNLDG
GDRTADVSAFTRELYGSEAVRTP
11111000000000000000000000000000000000000000000000000000000000
00000000000000000000000000000000000000000000000000000000000000
00000000000000000000000000000000000000000000000000000000000000
00000000000000000000000000000000000000000000000000000000000000
00000000000000000000000
>DM_train978
METLTVHAPSPSTNLPSYGNGAFSLSAPHVPGAGPLLVQVVYSFFQSPNMCLQALTQLEDYI
KKHGASNPLTLQIISTNIGYFCNADRNLVLHPGISVYDAYHFAKPAPSQYDYRSMNMKQMSG
NVTTPIVALAHYLWGNGAERSVNIANIGLKISPMKINQIKDIIKSGVVGTFPVSTKFTHATG
DYNVITGAYLGNITLKTEGTLTISANGSWTYNGVVRSYDDKYDFNASTHRGIIGESLTRLGA
MFSGKEYQILLPGEIHIKESGKR
10000000000000000000000000000000000000000000000000000000000000
00000000000000000000000000000000000000000000000000000000000000
00000000000000000000000000000000000000000000000000000000000000
00000000000000000000000000000000000000000000000000000000000000
00000000000000000000000

>DM_train979
MPLSSQPAILIIGGAEDKVHGREILQTFWSRSGGNDAIIGIIPSASREPLLIGERYQTIFSD
MGVKELKVLDIRDRAQGDDSGYRLFVEQCTGIFMTGGDQLRLCGLLADTPLMDRIRQRVHNG
EISLAGTSAGAAVMGHHMIAGGSSGEWPNRALVDMAVGLGIVPEIVVDQHFHNRNRMARLLS
AISTHPELLGLGIDEDTCAMFERDGSVKVIGQGTVSFVDARDMSYTNAALVGANAPLSLHNL
RLNILVHGEVYHQVKQRAFPRVT
111000000000000000000000000000000000000000000000000000000000
000000000000001111110000000000000000000000000000000000000000
000000000000000000000000000000000000000000000000000000000000
000000000000000000000000000000000000000000000000000000000000
00000000000000000000011
>DM_train980
MSDWTYSEGALDEAHWPQHYPACGGQRQSPINLQRTKVRYNPSLKGLNMTGYETQAGEFPMV
NNGHTVQISLPSTMRMTVADGTVYIAQQMHFHWGGASSEISGSEHTVDGIRHVIEIHIVHYN
SKYKSYDIAQDAPDGLAVLAAFVEVKNYPENTYYSNFISHLANIKYPGQRTTLTGLDVQDML
PRNLQHYYTYHGSLTTPPCTENVHWFVLADFVKLSRTQVWKLENSLLDHRNKTIHNDYRRTQ
PLNHRVVESNFPNQEYTLGSEFQ
111111111111000000000000000000000000000000000000000000000000
000000000000000000000000000000000011110000000000000000000000
000000000000000000000000000000000000000000000000000000000000
000000000000000000000000000000000000000000000000000000000000
00000000000011111111111
>DM_train981
WGALGHATVAYVAQHYVSPEAASWAQGILGSSSSSYLASIASWADEYRLTSAGKWSASLHFI
DAEDNPPTNCNVDYERDCGSSGCSISAIANYTQRVSDSSLSSENHAEALRFLVHFIGDMTQP
LHDEAYAVGGNKINVTFDGYHDNLHSDWDTYMPQKLIGGHALSDAESWAKTLVQNIESGNYT
AQAIGWIKGDNISEPITTATRWASDANALVCTVVMPHGAAALQTGDLYPTYYDSVIDTIELQ
IAKGGYRLANWINEIHGSEIAK
000000000000000000000000000000000000000000000000000000000000
000000000000000000000000000000000000000000000000000000000000
000000000000000000000000000000000000000000000000000000000000
000000000000000000000000000000000000000000000000000000000000
0000000000000000111111
>DM_train982
MSIYTLGIDVGSTASKCIILKDGKEIVAKSLVAVGTGTSGPARSISEVLENAHMKKEDMAFT
LATGYGRNSLEGIADKQMSELSCHAMGASFIWPNVHTVIDIGGQDVKVIHVENGTMTNFQMN
DKCAAGTGRFLDVMANILEVKVSDLAELGAKSTKRVAISSTCTVFAESEVISQLSKGTDKID
IIAGIHRSVASRVIGLANRVGIVKDVVMTGGVAQNYGVRGALEEGLGVEIKTSPLAQYNGAL
GAALYAYKKAAKSAWSHPQFEK
000000000000000000000000000000000000000000000000000000000000
000000000000000000000000000000000000000000000000000000000000
000000000000000000000000000000000000000000000000000000000000
000000000000000000000000000000000000000000000000000000000000
0000000000001111111111

>DM_train983
GASCSGGDKSKAPVVSTADIENAAEVIKYYNTSLGVLKDMVKEKDVNAVLDYMEQKGKTPAL
SAIVPPAVVSKDSAIVLNPGNCFNEETRRNLKQNYTGLFQARTEFYANFDTYLSYLKKKDVT
NAKKLLDVNYQLSTQMSEYKQNIFDILSPFTEQAELVLLVDNPLKAQIMSVRKMSSTMQSIL
NLYARKHRMDGPRIDLKVAELTKQLDAAKKLPVVNGHEGEMKSYQAFLSQVETFIKQVKKVR
EKGEYSDADYDMLTSAFETSII
111111111111110000000000000000000000000000000000000000000000
000000000000000000000000000000000000000000000000000000000000
000000000000000000000000000000000000000000000000000000000000
000000000000000000000000000000000000000000000000000000000000
0000000000000000000000
>DM_train984
GMTNVGRPIETHAITVGQGSDARSIAALVRAPAQDERPTCIWLGGYRSDMTGTKALEMDDLA
ASLGVGAIRFDYSGHGASGGAFRDGTISRWLEEALAVLDHFKPEKAILVGSSMGGWIALRLI
QELKARHDNPTQVSGMVLIAPAPDFTSDLIEPLLGDRERAELAENGYFEEVSEYSPEPNIFT
RALMEDGRANRVMAGMIDTGCPVHILQGMADPDVPYQHALKLVEHLPADDVVLTLVRDGDHR
LSRPQDIDRMRNAIRAMIEPRP
111111000000000000000000000000000000000000000000000000000000
000000000000000000000000000000000000000000000000000000000000
000000000000000000000000000000000000000000000000000000000000
000000000000000000000000000000000000000000000000000000000000
0000000000000000000111
>DM_train985
GMKSQWECFLQNLGVWEGSFSNFSPEGTLLNDTSSRLCLEGLNNNQTVRLTLSRSGKDDVIR
EFRSVGGGLLFFENGSFSEGLIQLGPFSEFGGELAFVHENRRLRLVQLFDRNGHLNGLTLIR
EHLAGTPVAERPLLQINDLLGEWRGQAVTIYRDLRPPDIYSTTLKIQLDDAGRLMQSTSFGE
RTITSTATIKGSIVLFDQDPEKQVQVLLLPDGASATSPLKVQLRQPLFLEAGWLIQSDLRQR
MIRSYNDKGEWVSLTLVTEERV
100000000000000000000000000000000000000000000000000000000000
000000000000000000000000000000000000000000000000000000000000
000000000000000000000000000000001000000000000000000000000000
000000000000000000000000000000000000000000000000000000000000
0000000000000000000000
>DM_train986
GMANLLSTCTSESGNIQHISPQNAGWEYVGFDVWQLKAGESITLPSDERERCLVLVAGLASV
KAADSFFYRIGQRMSPFERIPAYSVYLPHHTEAKVTAETDLELAVCSAPGFGELPVRLISPQ
EVGVEHRGKGRNQRLVHNILPDSQLADSLLVVEVYTNAGATSSWPAHKHDTAVEGQETYLEE
TYYHRFNPPQGFCLQRVYTDDRSLDECMAVYNRDVVKVPKGYHPVATIAGYDNYYLNVMAGP
LRKWRFTWEENHAWINSPDYPR
110000000000000000000000000000000000000000000000000000000000
000000000000000000000000000000000000000000000000000000000000
000000000000000000000000000000000000000000000000000000000000
000000000000000000000000000000000000000000000000000000000000
0000000000000000011111

>DM_train987
AMSNNKIKRKDASPEQEAIESFTSLTKCDPKVSRKYLQRNHWNINYALNDYYDKEIGTFTDE
VSTVAHPPVYPKELTQVFEHYINNNLFDIDSLVKFIEELGYNLEDLATLCLAHLLGYKKLEE
PLKREDFLSTWFMQGCSTISDMQECIKTLDVKLHEDLQYFTQIYNYAFNLILDPNRKDIDTD
EGIQYWKLFFQPEYPVRMEPDLLEAWFRFLRDEGKTTISKDTWRMLLLFFKRYPTIQKIISD
YDETAAWPFIIDEFYECLQDQQ
111111111111000000000000000000000000000000000000000000011111
111000000000000000000000000000000000000000000000000000000000
000000000000000000000000000000000000000000000000000000000000
000000000000000000000000000000000000000000000000000000000000
0000000000000000000000
>DM_train988
SDGGKVVAATTAQIQEFTKYAGIAATAYCRSVVPGNKWDCVQCQKWVPDGKIITTFTSLLSD
TNGYVLRSDKQKTIYLVFRGTNSFRSAITDIVFNFSDYKPVKGAKVHAGFLSSYEQVVNDYF
PVVQEQLTAHPTYKVIVTGHSLGGAQALLAGMDLYQREPRLSPKNLSIFTVGGPRVGNPTFA
YYVESTGIPFQRTVHKRDIVPHVPPQSFGFLHPGVESWIKSGTSNVQICTSEIETKDCSNSI
VPFTSILDHLSYFDINEGSCL
111100000000000000000000000000000000000000000000000000000000
000000000000000000000000000000000000000000000000000000000000
000000000000000000000000000000000000000000000000000000000000
000000000000000000000000000000000000000000000000000000000000
000000000000000000000
>DM_train989
ATISRRGILVIRHGERVDQVFGKSWLQQCTTADGKYYRPDLNFPRSLPRRSNGIKDFENDPP
LSSCGIFQARLAGEALLDSGVRVTAVFASPALRCVQTAKHILEELKLEKKLKIRVEPGIFEW
MKWEASKATLTFLTLEELKEANFNVDLDYRPALPRCSLMPAESYDQYVERCAVSMGQIINTC
PQDMGITLIVSHSSALDSCTRPLLGLPPRECGDFAQLVRKIPSLGMCFCEENREDGKWDLVN
PPVKTLTHGANSVFNWRNWIS
111100000000000000000000000000000000000000000000000000000000
000000000000000000000000000000000000000000000000000000000000
000000000000000000000000000000000000000000000000000000000000
000000000000000000000000000000000000000000000000000000000000
000000000000000000011
>DM_train990
GMQGKKFISPGAWFSMNYPSDWNEFEDGEGSFLFYNPDVWTGNFRISAFKGNASYGKDAIRQ
ELKENDSASLVKIGTWDCAYSKEMFQEEGTYYTSHLWITGTGNIAFECSFTVPKGGSAKEAE
EVIATLEARKEGEKYPAELIPVRLSEIYQINEGYEWVVSTVKQELKKDFQGVEEDLEKIQQV
IDSGKISPKKKDEWLAIGITVCAILTNEVEGMEWKTLIDGNREVPVLEYQGRTIDPMKIAWS
KVKAGQPCNIAEAYQSAIDHH
100000000000000000000000000000000000000000000000000000000000
000000000000000000000000000000000000000000000000000000000000
000000000000000000000000000000000000000000000000000000000000
000000000000000000000000000000000000000000000000000000000000
000000000000000000000

>DM_train991
GSHMEEKRLSAKKGLPPGTLVYTGKYREDFEIEVMNYSIEEFREFKTTDVESVLPFRDSSTP
TWINITGIHRTDVVQRVGEFFGTHPLVLEDILNVHQRPKVEFFENYVFIVLKMFTYDKNLHE
LESEQVSLILTKNCVLMFQEKIGDVFDPVRERIRYNRGIIRKKRADYLLYSLIDALVDDYFV
LLEKIDDEIDVLEEEVLERPEKETVQRTHQLKRNLVELRKTIWPLREVLSSLYRDVPPLIEK
ETVPYFRDVYDHTIQIADTVE
111111111111111000000000000000000000000000000000000000000000
000000000000000000000000000000000000000000000000000000001100
000000000000000000000000000000000000000000000000000000000000
000000000000000011111110000000000000000000000000000000000001
111111111111111111111
>DM_train992
MIVVKVGGAEGINYEAVAKDAASLWKEGVKLLLVHGGSAETNKVAEALGHPPRFLTHPGGQV
SRLTDRKTLEIFEMVYCGLVNKRLVELLQKEGANAIGLSGLDGRLFVGRRKTAVKYVENGKV
KVHRGDYTGTVEEVNKALLDLLLQAGYLPVLTPPALSYENEAINTDGDQIAALLATLYGAEA
LVYLSNVPGLLARYPDEASLVREIPVERIEDPEYLALAQGRMKRKVMGAVEAVKGGVKRVVF
ADGRVENPIRRALSGEGTVVR
000000000000000000000000000000000000000000000000000011111110
000000000000000000000000000000000000000000000000011111111111
111110000000000000000000000000000000000000000000000000000000
000000000000111111000000000000000000000000000000000000000000
000000000000000000000
>DM_train994
NGHLQHHPPMPPHPGHYWPVHNELAFQPPISNHPAPEYWCSIAYFEMDVQVGETFKVPSSCP
IVTVDGYVDPSGGDRFCLGQLSNVHRTEAIERARLHIGKGVQLECKGEGDVWVRCLSDHAVF
VQSYYLDREAGRAPGDAVHKIYPSAYIKVFDLRQCHRQMQQQAATAQAAAAAQAAAVAGNIP
GPGSVGGIAPAISLSAAAGIGVDDLRRLCILRMSFVKGWGPDYPRQSIKETPCWIEIHLHRA
LQLLDEVLHTMPIADPQPLD
000000000000111111111111110000000000000000000000000000000000
000000000000000000000000000000000000000000000000000000000000
000000000000000000000000000000000000000000000000001111111111
111111111111111100000000000000000000000000000000000000000000
00000000000001111111
>DM_train995
MNRVDLSLFIPDSLTAETGDLKIKTYKVVLIARAASIFGVKRIVIYHDDADGEARFIRDILT
YMDTPQYLRRKVFPIMRELKHVGILPPLRTPHHPTGKPVTGEYRQGLTVKRVKKGTLVDIGA
DKLALCREKLTVNRIMSFRVVRLGKEILIEPDEPEDRYWGYEVLDTRRNLAESLKTVGADVV
VATSRNASPITSILDEVKTRMRGAREAAILFGGPYKGLPEIDADIWVNTLPGQCTETVRTEE
AVLATLSVFNMLTQIDEKDE
000000000000000000000000000000000000000000000000000000000000
000000000000000000000000000000000000000000000000000000000000
000000000000000000000000000000000000000000000000000000000000
000000000000000000000000000000000000000000000000000000000000
00000000000000111111

>DM_train996
MERYESLFAQLKERKEGAFVPFVTLGDLGIEQSLKIIDTLIEAGADALELGIPFSDPLADGP
TIQNATLRAFAAGVTPAQCFEMLAIIREKHPTIPIGLLMYANLVFNKGIDEFYARCEKVGVD
SVLVADVPVEESAPFRQAALRHNVAPIFICPPNADDDLLRQIASYGRGFTYLLSRAGVTGAE
NRAALPLNHLVAKLKEYNAAPPLQGFGISAPDQVKAAIDAGAAGAISGSAIVKIIEQHINEP
EKMLAALKVFVQPMKAATRS
000000000000000000000000000000000000000000000000000011111111
111111111111100000000000000000000000000000000000000000000000
000000000000000000000000000000000000000000000000000000111111
110000000000000000000000000000000000000000000000000000000000
00000000000000000001
>DM_train997
MIQITVIQIDNYGPWTVTPNPRRESDLQALQSRLYADLNLMFGAHKGLVFYTRFDNLIAITN
GIDLITHKRIQESIRNRYPFTVSMVIASAETPYEAQKLATETLQEYGSAQDENRKEVLDVAN
ELVVDGYVQIAHIDINNITGTLTDIVSAYDTYLNVNKVKLALMEELLKYNALLFFIGGDNFM
APSNGMSEEDFLDIFNRINKKYKIELKAGIGIGRTAEDASNLADIGLEKIRGKLVDKNVCTL
KQDDFLESKMGMGKIYHPQF
000000000000000000000000000000000000000000000000000000000000
000000000000000000000000000000000000000000000000000000000000
000000000000000000000000000000000000000000000000000000000000
000000000000000000000000000000000000000000000000000000000000
00111111111111111111
>DM_train999
MKDRTQELRTAKDSDDDDDVTVTVDRDRFMDEFFEQVEEIRGFIDKIAENVEEVKRKHSAIL
ASPNPDEKTKEELEELMSDIKKTANKVRSKLKSIEQSIEQEEGLNRSSADLRIRKTQHSTLS
RKFVEVMSEYNATQSDYRERCKGRIQRQLEITGRTTTSEELEDMLESGNPAIFASGIIMDSS
ISKQALSEIETRHSEIIKLENSIRELHDMFMDMAMLVESQGEMIDRIEYNVEHAVDYVERAV
SDTKKAVKYQSKARRKKIM
111111111111111111111111100000000000000000000000000000000000
000000000000000000000000000000000000000000000000000000000000
000000000000000000000000000000000000000000000000000000000000
000000000000000000000000000000000000000000000000000000000000
1111111111111111111
>DM_train1001
SEDNGVDFNKNTALDKNYLLNNKIPSNNVEEAGSKNYVHYIIQLQGDDISYEATCNLFSKNP
KNSIIIQRNMNESAKSYFLSDDGESILELNKYRIPERLKNKEKVKVTFIGHGKDEFNTSEFA
RLSVDSLSNEISSFLDTIKLDISPKNVEVNLLGCNMFSYDFNVEETYPGKLLLSIMDKITST
LPDVNKNSITIGANQYEVRINSEGRKELLAHSGKWINKEEAIMSDLSSKEYIFFDSIDNKLK
AKSKNIPGLASISEDIKTL
111110000000000000000000000111111100000000000000000000000000
000000000000000000000000000000000000000000000000011111000000
000000000000000000000000000000000000000000000000000000000000
000000000000000000000000000000000000000000000000000000000000
0000000000000111111

>DM_train1002
GMHTDVNALFAALWQDYIKMTPSAAKIHQLLGHGAPIINDHIALRTFNIAKVNLSVLAKHFT
SIGYVDSGDYKFEQKKLIAKHFEHPDPKQPKVFISELLVEEFSPEVQKSIHGLIDQVDIAAT
TADNFIYSGRHWDVDKATYQALLAESEYAAWVAALGYRANHFTVSINDLPEFERIEDVNQAL
KQAGFVLNSSGGEVKGSPEVLLEQSSTMADKVVVNFTDGDVEIPSCFYEFARRYPMANGQLY
TGFVAASADKIFESTNAMM
000000000000000000000000000000000000000000000000000000000000
000000000000000000000000000000000000000000000000000000000000
000000000000000000000000000000000000000000000000000000000000
000000000000000000000000000000000000000000000000000000000000
0000001111111111111
>DM_train1003
MKEGKHFVLVHGACHGGWSWYKLKPLLEAAGHKVTALDLAASGTDLRKIEELRTLYDYTLPL
MELMESLSADEKVILVGHSLGGMNLGLAMEKYPQKIYAAVFLAAFMPDSVHNSSFVLEQYNE
RTPAENWLDTQFLPYGSPEEPLTSMFFGPKFLAHKLYQLCSPEDLALASSLVRPSSLFMEDL
SKAKYFTDERFGSVKRVYIVCTEDKGIPEEFQRWQIDNIGVTEAIEIKGADHMAMLCEPQKL
CASLLEIAHKYNMAGDPLE
110000000000000000000000000000000000000000000000000000000000
000000000000000000000000000000000000000000000000000000000000
000000000000000000000000000000000000000000000000000000000000
000000000000000000000000000000000000000000000000000000000000
0000000000111111111
>DM_train1004
MAPHGDGLSDIEEPEVDAQSEILRPISSVVFVIAMQAEALPLVNKFGLSETTDSPLGKGLPW
VLYHGVHKDLRINVVCPGRDAALGIDSVGTVPASLITFASIQALKPDIIINAGTCGGFKVKG
ANIGDVFLVSDVVFHDRRIPIPMFDLYGVGLRQAFSTPNLLKELNLKIGRLSTGDSLDMSTQ
DETLIIANDATLKDMEGAAVAYVADLLKIPVVFLKAVTDLVDGDKPTAEEFLQNLTVVTAAL
EGTATKVINFINGRNLSDL
111111111111111111111110000000000000000000000000000000000000
000000000000000000000000000000000000000000000000000000000000
000000000000000000000000000000000000000000000000000000000000
000000000000000000000000000000000000000000000000000000000000
0000000000000000000
>DM_train1005
MKGFQFKVIQEKLQGRQFINSDKIKPDHPQTIIKKTLLKEYQSKNFSCQEERDLFLEFTEKI
VQNFHNINFNYLLKKFCKLPENYQSLKSQVKQIVQSENKANQQSCENLFNSLYDTEISYKQI
TNFLRQIIQNCVPNQLLGKKNFKVFLEKLYEFVQMKRFENQKVLDYICFMDVFDVEWFVDLK
NQKFTQKRKYISDKRKILGDLIVFIINKIVIPVLRYNFYITEKHKEGSQIFYYRKPIWKLVS
KLTIVKLEEENLEKVEEKL
111111000000011111111111000000000000000000000000000000000000
000000000000000000000000000000000000000000000000000000000000
000000000000000000000000000000000000000000000000000000000000
000000000000000000000000000000000000000000000000000000000000
0000000000000000000

>DM_train1006
MVDSHVHTPLCGHAEGHPEAYLEEARAKGLKGVVFTDHSPMPPWYDPESRMRLEALPFYLLA
LERVRERAQDLYVGIGLEADFHPGTEGFLAQLLRRYPFDYVIGSVHYLGAWPLDHPDHQEEY
AWRDLKEVFRAYFQEVEKAARSGLFHAIGHLDLPKKFGHRLPEEALLELAEPALRAVAEAGL
FLDVNTAGLRRPAKEVYPAPALLRRARELGIGLVLGSDAHRPEEVGFAFPEVQALLAGLGFR
EAYYFVEGSPVAYPLSRAS
000000000000000000000000000000000000000000000000000000000000
000000000000000000000000000000000000000000000000000000000000
000000000000000000000000000000000000000000000000000000000000
000000000000000000000000000000000000000000000000000000000000
0000000000000001111
>DM_train1007
MRLIPLTTAEQVGKWAARHIVNRINAFKPTADRPFVLGLPTGGTPMTTYKALVEMHKAGQVS
FKHVVTFNMDEYVGLPKEHPESYYSFMHRNFFDHVDIPAENINLLNGNAPDIDAECRQYEEK
IRSYGKIHLFMGGVGNDGHIAFNEPASSLASRTRIKTLTHDTRVANSRFADNDVNQVPKYAL
TVGVGTLLDAEEVMILVLGSQKALALQAAVEGCVNHMWTISCLQLHPKAIMVCDEPSTMELK
VKTLRYFNELEAENIKGL
000000000000000000000000000000000000000000000000000000000000
000000000000000000000000000000000000000000000000000000000000
000000000000000000000000000000000000111111111111111111100000
000000000000000000000000000000000000000000000000000000000000
000000000000000000
>DM_train1008
MTPDFLAIKVGGSLFSRKDEPGSLDDDAVTRFARNFARLAETYRGRMVLISGGGAFGHGAIR
DHDSTHAFSLAGLTEATFEVKKRWAEKLRGIGVDAFPLQLAAMCTLRNGIPQLRSEVLRDVL
DHGALPVLAGDALFDEHGKLWAFSSDRVPEVLLPMVEGRLRVVTLTDVDGIVTDGAGGDTIL
PEVDARSPEQAYAALWGSSEWDATGAMHTKLDALVTCARRGAECFIMRGDPGSDLEFLTAPF
SSWPAHVRSTRITTTASA
000000000000000000000000000000000000000000000000000000111111
111111000000000000000000000000000000000000000000000000000000
000000000000000000000000000000000000000000000000000011100000
000000000000000000001111000000000000000000000000000000000000
000000000000001111
>DM_train1009
MHDLRISLVQGSTRWHDPAGNRDYYGALLEPLAGQSDLVILPETFTSGFSNEAIDKAEDMDG
PTVAWIRTQAARLGAAITGSVQLRTEHGVFNRLLWATPDGALQYYDKRHLFRFGNEHLRYAA
GRERLCVEWKGWRINPQVCYDLRFPVFCRNRFDVERPGQLDFDLQLFVANWPSARAYAWKTL
LRARAIENLCFVAAVNRVGVDGNQLHYAGDSAVIDFLGQPQVEIREQEQVVTTTISAAALAE
HRARFPAMLDGDSFVLGE
000000000000000000000000000000000000000000000000000000000000
000000000000000000000000000000000000000000000000000000000000
000000000000000000000000000000000000000000000000000000000000
000000000000000000000000000000000000000000000000000000000000
000000000000000001

>DM_train1010
SNANFDTNMFKLENYVKEKYSLESLEIIPNEFDDTPTILSERISQVAAGVLRNLIDDNMKIG
FSWGKSLSNLVDLIHSKSVRNVHFYPLAGGPSHIHAKYHVNTLIYEMSRKFHGECTFMNATI
VQENKLLADGILQSRYFENLKNSWKDLDIAVVGIGDFSNKGKHQWLDMLTEDDFKELTKVKT
VGEICCRFFDSKGKEVYENLQERTIAISLEDLKNIPQSLAVAYGDTKVSSILSVLRANLVNH
LITDKNTILKVLEEDGDL
11100000000000000000000000000000000000000000000000000000000000
00000000000000000000000000000000000000000000000000000000000000
00000000000000000000000000000000000000000000000000000000000000
00000000000000000000000000000000000000000000000000000000000000
000000000000000001
>DM_train1011
RPSSRVYIVEVLNEFPHDPYAFTQGLVYAENDTLFESTGLYGRSSVRQVALQTGKVENIHKM
DDSYFGEGLTLLNEKLYQVVWLKNIGFIYDRRTLSNIKNFTHQMKDGWGLATDGKILYGSDG
TSILYEIDPHTFKLIKKHNVKYNGHRVIRLNELEYINGEVWANIWQTDCIARISAKDGTLLG
WILLPNLRKKLIDEGFRDIDVLNGIAWDQENKRIFVTGKLWPKLFEIKLHLVRHRIPDGYIE
RHCLNLRDNTLSLKTDID
00000000000000000000000000000000000000000000000000000000000000
00000000000000000000000000000000000000000000000000000000000000
00000000000000000000000000000000000000000000000000000000000000
00000000000000000000000000000000000000000000000000000000000000
000000111111111111
>DM_train1012
GMSADIVFDRGSPSDIDEIKTFTSNTWKVGYYTDLYSKLADTGTMDDYVDKVIERWVNDGSV
YVLRVSGRPVATIHMEKLPDGSVMLGGLRVHPEYRGSRLGMSIMQETIQFLRGKTERLRSAV
YSWNEPSLRLVHRLGFHQVEEYPIYTFQGGSTAVPALKPVNERYAGRWRCFFIDWKYMCSDD
PDIIHEEYSNNLIVDGSTFVYFDIYEGGIDLFVNDSDDASSFIEKYRSMNGRITFYVRKALA
NGLPYVPASSLTVWEYRY
11111000000000000000000000000000000000000000000000000000000000
00000000000000000000000000000000000000000000000000000000000000
00000000000000000000000000000001111000000000000000000000000000
00000000000000000000000000000000000000000000000000000000000000
000000000000000000
>DM_train1013
MNITPFPTLSPATIDAINVIGQWLAQDDFSGEVPYQADCVILAGNAVMPTIDAACKIARDQQ
IPLLISGGIGHSTTFLYSAIAQHPHYNTIRTTGRAEATILADIAHQFWHIPHEKIWIEDQST
NCGENARFSIALLNQAVERVHTAIVVQDPTMQRRTMATFRRMTGDNPDAPRWLSYPGFVPQL
GNNADSVIFINQLQGLWPVERYLSLLTGELPRLRDDSDGYGPRGRDFIVHVDFPAEVIHAWQ
TLKHDAVLIEAMESRSLR
11110000000000000000000000000000000000000000000000000000000000
00000000000000000000000000000000000000000000000000000000000000
00000000000000000000000000000000000000000000000000000000000000
00000000000000000000000000000000000000000000000000000000000000
000000000000000111

>DM_train1014
GSHMQAEILLTLKLQQKLFADPRRISLLKHIALSGSISQGAKDAGISYKSAWDAINEMNQLS
EHILVERATGGKGGGGAVLTRYGQRLIQLYDLLAQIQQKAFDVLSDDDALPLNSLLAAISRF
SLQTSARNQWFGTITARDHDDVQQHVDVLLADGKTRLKVAITAQSGARLGLDEGKEVLILLK
APWVGITQDEAVAQNADNQLPGIISHIERGAEQCEVLMALPDGQTLCATVPVNEATSLQQGQ
NVTAYFNADSVIIATLC
111000000000000000000000000000000000000000000000000000000000
000000111111111100000000000000000000000000000000000000000000
000000000000000011111110000000000000000000000000000000000000
000000000000000000000000000000000000000000000000000000000000
00000000000000000
>DM_train1015
MRVLALSAVFLVASIIGMPAVAKEWQENKSWNAHFTEHKSQGVVVLWNENKQQGFTNNLKRA
NQAFLPASTFKIPNSLIALDLGVVKDEHQVFKWDGQTRDIATWNRDHNLITAMKYSVVPVYQ
EFARQIGEARMSKMLHAFDYGNEDISGNVDSFWLDGGIRISATEQISFLRKLYHNKLHVSER
SQRIVKQAMLTEANGDYIIRAKTGYSTRIEPKIGWWVGWVELDDNVWFFAMNMDMPTSDGLG
LRQAITKEVLKQEKIIP
111111111111111111111111000000000000000000000001100000000000
000000000000000000000000000000000000000000000000000000000000
000000000000000000000000000000000000000000000000000000000000
000000000000000000000000000000000000000000000000000000000000
00000000000000000
>DM_train1016
MNDIVASTQLPNTIKTITNDLRKLGLKKGMTVIVHSSLSSIGWISGGAVAVVEALMEVITEE
GTIIMPTQSSDLSDPKHWSRPPVPEEWWQIIRDNVPAFEPHITPTRAMGKVVECFRTYPNVV
RSNHPLGSFAAWGRHAEEITVNQSLSMSLGEESPLRKIYDLDGYILLIGVGYDSNTSVHLSE
VRSGACELIKVGAPIIENGERVWKEFVDMDYDSDKFVEIGVEFEQKGTVTMGKIGNAKCRLM
KQRDIVDFGTEWFRKKN
000000000000000000000000000000000000000000000000000000000000
000000000000000000000000000000000000000000000000000000000000
000000000000000000000000000000000000000000000000000000000000
000000000000000000000000000000000000000000000000000000000000
00000000000000001
>DM_train1017
TQSSKLIAVIVANIDDYFSTELFKGISSILESRGYIGVLFDANADIEREKTLLRAIGSRGFD
GLILQSFSNPQTVQEILHQQMPVVSVDREMDACPWPQVVTDNFEAAKAATTAFRQQGYQHVV
VLTSELELSRTRQERYRGILAAAQDVDVLEVSESSYNHSEVHQRLTQLITQNDQKTVAFALK
ERWLLEFFPNLIISGLIDNQTVTATGFADTDFIRRMEPKLTLITQNPFLMGASSAEIMLRQL
AGEKVAPEKMVIPAKLQ
111100000000000000000000000000000000000000000000000000000000
000000000000000000000000000000000000000000000000000100000000
000000000000000000000000000000000000000000000000111100000000
000000000000000000000000000000000011000000000000000000000000
00000000000000000

>DM_train1018
MAEYKDNLLGEANSFLEVLEQVSHLAPLDKPVLIIGERGTGKELIASRLHYLSSRWQGPFIS
LNCAALNENLLDSELFGHEAGAFTGAQKRHPGRFERADGGTLFLDELATAPMMVQEKLLRVI
EYGELERVGGSQPLQVNVRLVCATNADLPAMVNEGTFRADLLDALAFDVVQLPPLRERESDI
MLMAEYFAIQMCREIKLPLFPGFTERARETLLNYRWPGNIRELKNVVERSVYRHGTSDYPLD
DIIIDPFKRRPPEDAIA
111111111000000000000000000000000000000000000000000000000000
000000000000000000011111111100000000000000000000000000000000
000000000011000000000000000000000000000000000000000000000000
000000000000000000000000000000000000000000000000000000000000
00000000011111111
>DM_train1021
MKILVAVKQTAALEEDFEIREDGMDVDEDFMMYDLNEWDDFSLEEAMKIKESSDTDVEVVVV
SVGPDRVDESLRKCLAKGADRAVRVWDDAAEGSDAIVVGRILTEVIKKEAPDMVFAGVQSSD
QAYASTGISVASYLNWPHAAVVADLQYKPGDNKAVIRRELEGGMLQEVEINCPAVLTIQLGI
NKPRYASLRGIKQAATKPIEEVSLADIGLSANDVGAAQSMSRVRRMYIPEKGRATMIEGTIS
EQAAKIIQIINEFKGA
000000000000000000000000000000000000000000000000000000000000
000000000000000000000000000000000000000000000000000000000000
000000000000000000000000000000000000000000000000000000000000
000000000000000000000000000000000000000000000001111111111111
1111111111111111
>DM_train1022
GHMSLHPVSISLSSYGADLVRSRGQASFLPLLAMAGAQRVELREELFAGPPDTEALTAAIQL
QGLECVFSSPLELWREDGQLNPELEPTLRRAEACGAGWLKVSLGLLPEQPDLAALGRRLARH
GLQLLVENDQTPQGGRIEVLERFFRLAERQQLDLAMTFDIGNWRWQEQAADEAALRLGRYVG
YVHCKAVIRNRDGKLVAVPPSAADLQYWQRLLQHFPEGVARAIEYPLQGDDLLSLSRRHIAA
LARLGQPQEERQHGGS
111100000000000000000000000000000000000000000000000000000000
000000000000000000000000000000000000000000000000000000000000
000000000000000000000000000000000000000000000000000000000000
000000000000000000000000000000000000000000000000000000000000
0000001111111111
>DM_train1023
SNAMNHLPEKMRPYRDLLEKSAKEYVKLNVRKGKTGRYDSKIAGDPYFPKHETYPTDENGQP
MKLLAQINFSHIPQLDGYPSSGILQFYISVHDDVYGLNFDDRCEQKNFRVIYFENIVENDDE
LVSDFSFIGTGECDFPILSEAAVEPVKSSEWVLPTDFQFEQYTGMETMEFFGQFGEDEEDIY
NELAENGFGHKIGGYASFTQHDPREYAYKEHTIMLLQIDSDDDIDSMWGDVGIANFFITPED
LRKKDFSNVLYNWDCS
111000000000000000000000000000000000000000000000000000000000
000000000000000000000000000000000000000000000000000000000000
000000000000000000000000000000000000000000000000000000000000
000000000000000000000000000000000000000000000000000000000000
0000000000000000

>DM_train1024
SLLSNQLHFAKPTARTPLVVLVHGLLGSGADWQPVLSHLARTQCAALTLDLPGHGTNPERHC
DNFAEAVEMIEQTVQAHVTSEVPVILVGYSLGGRLIMHGLAQGAFSRLNLRGAIIEGGHFGL
QENEEKAARWQHDQQWAQRFSQQPIEHVLSDWYQQAVFSSLNHEQRQTLIAQRSANLGSSVA
HMLLATSLAKQPYLLPALQALKLPIHYVCGEQDSKFQQLAESSGLSYSQVAQAGHNVHHEQP
QAFAKIVQAMIHSIID
000000000000000000000000000000000000000000000000000000001111
111000000000000000000000000000000000000000000000000000000000
000000000000000000000000000000000000000000000000000000000000
000000000000000000000000000000000000000000000000000000000000
0000000000000000
>DM_train1025
RPSWSPTQASGQTVLPFTGIDFRLSPSGVAVDSAGNVYVTSEGMYGRVVKLATGSTGTTVLP
FNGLYQPQGLAVDGAGTVYVTDFNNRVVTLAAGSNNQTVLPFDGLNYPEGLAVDTQGAVYVA
DRGNNRVVKLAAGSKTQTVLPFTGLNDPDGVAVDNSGNVYVTDTDNNRVVKLEAESNNQVVL
PFTDITAPWGIAVDEAGTVYVTEHNTNQVVKLLAGSTTSTVLPFTGLNTPLAVAVDSDRTVY
VADRGNDRVVKLTSLE
111111111110000000000000000000000000000000000000001111100000
000000000000000000000000000000000000000000000000000000000000
000000000000000000000000000000000000000000000000000000000000
000000000000000000000000000000000000000000000000000000000000
0000000000000000
>DM_train1027
MVRTITVASGKGGTGKTTITANLGVALAQLGHDVTIVDADITMANLELILGMEGLPVTLQNV
LAGEARIDEAIYVGPGGVKVVPAGVSLEGLRKANPEKLEDVLTQIMESTDILLLDAPAGLER
SAVIAIAAAQELLLVVNPEISSITDGLKTKIVAERLGTKVLGVVVNRITTLGIEMAKNEIEA
ILEAKVIGLIPEDPEVRRAAAYGKPVVLRSPNSPAARAIVELANYIAGGAKKKVPAEVKEKK
KEGALAKMLRIFRRR
000000000000000000000000000000000000000000000000000000000000
000000000000000000000000000000000000000000000000000000000000
000000000000000000000000000000000000000000000000000000000000
000000000000000000000000000000000000000000000111111111111111
111111111111111
>DM_train1028
MANLTEKFLRIFARRGKSIILAYDHGIEHGPADFMDNPDSADPEYILRLARDAGFDGVVFQR
GIAEKYYDGSVPLILKLNGKTTLYNGEPVSVANCSVEEAVSLGASAVGYTIYPGSGFEWKMF
EELARIKRDAVKFDLPLVVESFPRGGKVVNETAPEIVAYAARIALELGADAMKIKYTGDPKT
FSWAVKVAGKVPVLMSGGPKTKTEEDFLKQVEGVLEAGALGIAVGRNVWQRRDALKFARALA
ELVYGGKKLAEPLNV
110000000000000000000000000000000000000000000000000000000000
000000000000000000000000000000000000000000000000000000000000
000000000000000000000000000000000000000000000000000000000000
000000000000000000000000000000000000000000000000000000000000
000011111111111

>DM_train1029
MWELRSASFWRAIFAEFFATLFYVFFGLGASLRWAPGPLHVLQVALAFGLALATLVQAVGHI
SGAHVNPAVTFAFLVGSQMSLLRAICYVVAQLLGAVAGAAVLYSVTPPAVRGNLALNTLHPG
VSVGQATIVEIFLTLQFVLCIFATYDERRNGRLGSVALAVGFSLTLGHLFGMYYTGAGMNPA
RSFAPAILTRNFTNHWVYWVGPVIGAGLGSLLYDFLLFPRLKSVSERLSILKGTRPSESNGQ
PEVTGEPVELKTQAL
111100000000000000000000000000000000000000000000000000000000
000000000000000000000000000000000000000000000000000000000000
000000000000000000000000000000000000000000000000000000000000
000000000000000000000000000000000000000000000000000111111111
111111111111111
>DM_train1030
GMSNKKLIINADDFGYTPAVTQGIIEAHKRGVVTSTTALPTSPYFLEAMESARISAPTLAIG
VHLTLTLNQAKPILPREMVPSLVDEAGYFWHQSIFEEKVNLEEVYNEWDAQIISFMKSGRRP
DHIDSHHNVHGKNKKLLGVALALARKYQLPLRNASRSIETKDYLELYQDVRTPDEMLYQFYD
KAISTETILQLLDMVVCSEGEVFEINCHPAFIDTILQNQSGYCMPRIREVEILTSQEVKEAI
EERGILLANYESLAM
110000000000000000000000000000000000000000000000000000000000
000000000000000000000000000000000000000000000000000000000000
000000000000000000000000000000000000000000000000000000000000
000000000000000000000000000000000000000000000000000000000000
000000000000000
>DM_train1031
GAMDPAVMKIEYYSQVLDMEWGVNVLYPDANRVEEPECEDIPVLYLLHGMSGNHNSWLKRTN
VERLLRGTNLIVVMPNTSNGWYTDTQYGFDYYTALAEELPQVLKRFFPNMTSKREKTFIAGL
SMGGYGCFKLALTTNRFSHAASFSGALSFQNFSPESQNLGSPAYWRGVFGEIRDWTTSPYSL
ESLAKKSDKKTKLWAWCGEQDFLYEANNLAVKNLKKLGFDVTYSHSAGTHEWYYWEKQLEVF
LTTLPIDFKLEERLT
000000000000000000000000000011111111000000000000000000000000
000000000000000000000000000000000000000000000000000000000000
000000000000000000000000000000111111100000000000000000000000
000000000000000000000000000000000000000000000000000000000000
000000000000001
>DM_train1032
EQASVTNDFSENKQGCIQHPWQGKKVGYIGDSITDPNCYGDNIKKYWDFLKEWLGITPFVYG
ISGRQWDDVPRQAEKLKKEHGGEVDAILVFMGTNDYNSSVPIGEWFTEQEEQVLSAHGEMKK
MVTRKKRTPVMTQDTYRGRINIGITQLKKLFPDKQIVLLTPLHRSLANFGDKNVQPDESYQN
GCGEYIDAYVQAIKEAGNIWGIPVIDFNAVTGMNPMVEEQLIYFYDAGYDRLHPDTKGQERM
ARTLMYQLLALPVAF
111111111111111000000000000000000000111100000000000000000000
000000000000000000000000000000000000000000000000000000000000
000000000000000000000000000000000000000000000000000000000000
000000000000000000000000000000000000000000000000000000000000
000000000000000

>DM_train1033
MSWYPGHIEKAKRQIKDLLRLVNTVVEVRDARAPFATSAYGVDFSRKETIILLNKVDIADEK
TTKKWVEFFKKQGKRVITTHKGEPRKVLLKKLSFDRLARVLIVGVPNTGKSTIINKLKGKRA
SSVGAQPGITKGIQWFSLENGVKILDTPGILYKNIFSEDLAAKLLLVGSLPVERIEDQRIFE
RAFEIFARSIGIESSFSEFFEDFARKRGLLKKGGVPDIERALMLFFTEVAQGKAGRVSFERP
EDITPVQQEQTRGV
111111110000000000000000000000000000000000000000000000000000
000000000000000000000000000000000000000000000000000000000000
111111111100000000000000000000000000000000000000000000000000
000000000000000000000000000000000000000000000000000000000000
00011111111111
>DM_train1034
MSFTVIIPARFASSRLPGKPLADIKGKPMIQHVFEKALQSGASRVIIATDNENVADVAKSFG
AEVCMTSVNHNSGTERLAEVVEKLAIPDNEIIVNIQGDEPLIPPVIVRQVADNLAKFNVNMA
SLAVKIHDAEELFNPNAVKVLTDKDGYVLYFSRSVIPYDRDQFMNLQDVQKVQLSDAYLRHI
GIYAYRAGFIKQYVQWAPTQLENLEKLEQLRVLYNGERIHVELAKEVPAVGVDTAEDLEKVR
AILAANGSHHHHHH
100000000000000000000000000000000000000000000000000000000000
000000000000000000000000000000000000000000000000000000000000
000000000000000000000000000000000000000000011111000000000000
000000000000000000000000000000000000000000000000000000000000
00000011111111
>DM_train1035
MSSGVDLGTENLYFQSNAMKMTVVGFWGGFPEAGEATSGYLFEHDGFRLLVDCGSGVLAQLQ
KYITPSDIDAVVLSHYHHDHVADIGVLQYARLITSATKGQLPELPIYGHTFDENGFHSLTHE
PHTKGIPYNPEETLQIGPFSISFLKTVHPVTCFAMRITAGNDIVVYSADSSYIPEFIPFTKD
ADLFICECNMYAHQEAAKAGHMNSTEVASIAKDANVKELLLTHLPHTGNPADLVTEAKQIFS
GHITLAHSGYVWNS
111111111111111110000000000000000000000000000000000000000000
000000000000000000000000000000000000000000000000000000000000
000000000000000000000000000000000000000000000000000000000000
000000000000000000000000000000000000000000000000000000000000
00000000000000
>DM_train1036
MEGRLLLLETPGNTRMSLAYDEAIYRSFQYGDKPILRFYRHDRSVIIGYFQVAEEEVDLDYM
KKNGIMLARRYTGGGAVYHDLGDLNFSVVRSSDDMDITSMFRTMNEAVVNSLRILGLDARPG
ELNDVSIPVNKKTDIMAGEKKIMGAAGAMRKGAKLWHAAMLVHTDLDMLSAVLKVPDEKFRD
KIAKSTRERVANVTDFVDVSIDEVRNALIRGFSETLHIDFREDTITEKEESLARELFDKKYS
TEEWNMGLLRKEVV
000000000000000000000000000000000000000000000000000000000000
000000000000000000000000000000000000000000000000000000000000
000000000000000000000000000000000000000000000000000011111111
111111100000000000000000000000000000000000000000000000000000
00000000111111

>DM_train1037
MAVTAQAARRKERVLCLFDVDGTLTPARQKIDPEVAAFLQKLRSRVQIGVVGGSDYCKIAEQ
LGDGDEVIEKFDYVFAENGTVQYKHGRLLSKQTIQNHLGEELLQDLINFCLSYMALLRLPKK
RGTFIEFRNGMLNISPIGRSCTLEERIEFSELDKKEKIREKFVEALKTEFAGKGLRFSRGGM
ISFDVFPEGWDKRYCLDSLDQDSFDTIHFFGNETSPGGNDFEIFADPRTVGHSVVSPQDTVQ
RCREIFFPETAHEA
111111111110000000000000000000000000000000000000000000000000
000000000000000000000000000000000000000000000000000000000000
000000000000000000000000000000000000000000000000000000000000
000000000000000000000000000000000000000000000000000000000000
00000000001111
>DM_train1038
MIFVLDVGNTNAVLGVFEEGELRQHWRMETDRHKTEDEYGMLVKQLLEHEGLSFEDVKGIIV
SSVVPPIMFALERMCEKYFKIKPLVVGPGIKTGLNIKYENPREVGADRIVNAVAGIHLYGSP
LIIVDFGTATTYCYINEEKHYMGGVITPGIMISAEALYSRAAKLPRIEITKPSSVVGKNTVS
AMQSGILYGYVGQVEGIVKRMKEEAKQEPKVIATGGLAKLISEESNVIDVVDPFLTLKGLYM
LYERNANLQHEKGE
000000000000000000000000000000000000000000000000000000000000
000000000000000000000000000000000000000000000000000000000000
000000000000000000000000000000000000000111111100000000000000
000000000000000000000000000000000000000000000000000000000000
00000011111111
>DM_train1039
GMATLGANASLYSEQHRITYYECDRTGRATLTTLIDIAVLASEDQSDALGLTTEMVQSHGVG
WVVTQYAIDITRMPRQDEVVTIAVRGSAYNPYFAYREFWIRDADGQQLAYITSIWVMMSQTT
RRIVKILPELVAPYQSEVVKRIPRLPRPISFEATDTTITKPYHVRFFDIDPNRHVNNAHYFD
WLVDTLPATFLLQHDLVHVDVRYENEVKYGQTVTAHANILPSEVADQVTTSHLIEVDDEKCC
EVTIQWRTLPEPIQ
111000000000000000000000000000000000000000000000000000000000
000000000000000000000000000000000000000000000000000000000000
000000000000000000000000000000000000000000000000000000000000
000000000000000000000000000000000000000000000000000000000000
00000000000111
>DM_train1041
SPTVEACGYSDRIIQITRGDSTITSQDVANAIVAYGVWPHYLSSKDASAIDKPSQPDTSSNR
FYTLRSVTWSSSSKGWWWKLPDALKDMGIFGENMFYHYLGRSGYTIHVQCNASKFHQGTLIV
ALIPEHQIASALHGNVNVGYNYTHPGETGREVKAETRLNPDLQPTEEYWLNFDGTLLGNITI
FPHQFINLRSNNSATIIAPYVNAVPMDSMRSHNNWSLVIIPICPLETSSAINTIPITISISP
MCAEFSGARAKRQ
111111111110000000000000000000000000000000000000000000000000
000000000000000000000000000000000000000000000000000000000000
000000000000000000000000000000000000000000000000000000000000
000000000000000000000000000000000000000000000000000000000000
0000000000000

>DM_train1042
NLSDFKVATWNLQGSSAVNESKWNINVRQLLSGEQGADILMVQEAGSLPSSAVRTSRVIQHG
GTPIEEYTWNLGTRSRPNMVYIYYSRLDVGANRVNLAIVSRRQADEAFIVHSDSSVLQSRPA
VGIRIGTDVFFTVHALATGGSDAVSLIRNIFTTFNSSSSPPERRVYSWMVVGDFNRAPANLE
VALRQEPAVSENTIIIAPTEPTHRSGNILDYAILHDAHLPRREQARERIGASLMLNQLRSQI
TSDHFPVSFVRDR
000000000000000000000000000000000000000000000000000000000000
000000000000000000000000000000000000000000000000000000000000
000000000000000000000000000000000001110000000000000000000000
000000000000000000000000000000000000000000000000000000000000
0000000000000
>DM_train1043
EPVKYGELVVLGYNGALPNGDRGRRKSRFALYKRPKANGVKPSTVHMISTPQASKAISCKGQ
HSISYTLSRNQTVVVEYTHDKDTDMFQVGRSTESPIDFVVTDTISGSQNTDEAQITQSTISR
FACRIVCDRNEPYTARIFAAGFDSSKNIFLGEKAAKWKNPDGHMDGLTTNGVLVMHPRGGFT
EESQPGVWREISVCGDVYTLRETRSAQQRGKMVESETNVLQDGSLIDLCGATLLWRTADGLF
HTPTQKHIEALRQ
000000000000011111111110000000000000000000000000000000000000
000000000000000000000000000000000000000001111111111000000000
000000000000000000000000000000000000000000000000000000001111
100000000000000000000000000000000000000000000000000000001111
1111111111111
>DM_train1044
QQGKFDTKGIESTDEAKMSFLVTLNNVEVCSENISTLKKTLESDCTKLFSQGIGGEQAQAKF
DSCLSDLAAVSNKFRDLLQEGLTELNSTAIKPQVQPWINSFFSVSHNIEEEEFNDYEANDPW
VQQFILNLEQQMAEFKASLSPVIYDSLTGLMTSLVAVELEKVVLKSTFNRLGGLQFDKELRS
LIAYLTTVTTWTIRDKFARLSQMATILNLERVTEILDYWGPNSGPLTWRLTPAEVRQVLALR
IDFRSEDIKRLRL
111111111111000000000000000000000000000000000000000000000000
000000000000000000000000000000000000000000000000000000000000
000000000000000000000000000000000000000000000000000000000000
000000000000000000000000000000000000000000000000000000000000
0000000000000
>DM_train1045
IAPTVWKLADRSKIYMADLESALHYILRIEVGRFPVLEGQRLVALKKFVAVLAKYFPGRPLV
QNFLHSVNEWLKRQKRNKIPYSFFKTALDDRKEGAVLAKKVNWIGCQGSEPHFRGFPCSLWV
LFHFLTVQAARQNVDHSQEAAKAKEVLPAIRGYVHYFFGCRDCASHFEQMAAASMHRVGSPN
AAVLWLWSSHNRVNARLAGAPSEDPQFPKVQWPPRELCSACHNERLDVPVWDVEATLNFLKA
HFSPSNIILDFPA
111111100000000000000000000000000000000000000000000000000000
000000000000000000000000000000000000000000000000000000000000
000000000000111111111000000000000000000000000000000000000000
000000000000000000000000000000000000000000000000000000000000
0000000001111

>DM_train1047
MVVLSVPAEVTVILLDIEGTTTPIAFVKDILFPYIEENVKEYLQTHWEEEECQQDVSLLRKQ
AEEDAHLDGAVPIPAASGNGVDDLQQMIQAVVDNVCWQMSLDRKTTALKQLQGHMWRAAFTA
GRMKAEFFADVVPAVRKWREAGMKVYIYSSGSVEAQKLLFGHSTEGDILELVDGHFDTKIGH
KVESESYRKIADSIGCSTNNILFLTDVTREASAAEEADVHVAVVVRPGNAGLTDDEKTYYSL
ITSFSELYLPSST
11100000000000000000000000000000000000000000000000000000000000
00000000000000000000000000000000000000000010000000000000000000
00000000000000000000000000000000000000000000000000000000000000
00000000000000000000000000000000000000000000000000000000000000
0000000001111
>DM_train1048
DKLPIEETLEDSPQTRSLLGVFEEDATAISNYMNQLYQAMHRIYDAQNELSAATHLTSKLLK
EYEKQRFPLGGDDEVMSSTLQQFSKVIDELSSCHAVLSTQLADAMMFPITQFKERDLKEILT
LKEVFQIASNDHDAAINRYSRLSKKRENDKVKYEVTEDVYTSRKKQHQTMMHYFCALNTLQY
KKKIALLEPLLGYMQAQISFFKMGSENLNEQLEEFLANIGTSVQNVRREMDSDIETMQQTIE
DLEVASDPLYVPD
11111111000000000000000000000000000000000000000000000000000000
00000000111110000000000000000000000000000000000000000000000000
00000000000000000000000000000000000000000000000000000000000000
00000000000000000000000000000000000000000000000000000000000000
0000000111111
>DM_train1049
MGAGEKGIVEKEGYQLDTRRQAQAAYPRIKVLVIHYTADDFDSSLATLTDKQVSSHYLVPAV
PPRYNGKPRIWQLVPEQELAWHAGISAWRGATRLNDTSIGIELENRGWQKSAGVKYFAPFEP
AQIQALIPLAKDIIARYHIKPENVVAHADIAPQRKDDPGPLFPWQQLAQQGIGAWPDAQRVN
FYLAGRAPHTPVDTASLLELLARYGYDVKPDMTPREQRRVIMAFQMHFRPTLYNGEADAETQ
AIAEALLEKYGQD
11110000000000000000000000000000000000000000000000000000000000
00000000000000000000000000000000000000000000000000000000000000
00000000000000000000000000000000000000000000000000000000000000
00000000000000000000000000000000000000000000000000000000000000
0000000000000
>DM_train1050
MHNNENVSGISAYLLGLIWGDGGLYKLKYKGNRSEYRVVITQKSENLIKQFIAPRMQFLIDE
LNVKSKIQIVKGDTRYELRVSSKKLYYYFANMLERIRLFNGNRFLAYLAGIVDGDGSIIAQI
KPNQSYKFKHQLSLTFQVTQKTERRWFLDKLVDEIGVGYVRDRGSVSDYILSEIKPLHNFLT
QLQPFLNFKQKQANLVLKIIEQLPSAKESPDKFLEVCTWVDQIAALNDSKTRKTTSETVRAV
LDSLSEKKKSSP
11111100000000000000000000011110000000000000000000000000000000
00000000000000000000000000000000000000000000000000000000000000
00000000000000000000000000000000000000000000000000000000000000
00000000000000000000000000000000000000000000000000000000000000
001111111111

>DM_train1051
MVGVTAFGNFDLASLAIYSFWIFLAGLIYYLQTENMREGYPLENEDGTPAANQGPFPLPKPK
TFILPHGRGTLTVPGPESEDRPIALARTAVSEGFPHAPTGDPMKDGVGPASWVARRDLPELD
GHGHNKIKPMKAAAGFHVSAGKNPIGLPVRGCDLEIAGKVVDIWVDIPEQMARFLEVELKDG
STRLLPMQMVKVQSNRVHVNALSSDLFAGIPTIKSPTEVTLLEEDKICGYVAGGLMYAAPKR
KSVVAAMLAEYA
111111111100000000000000000000000000000000000000000000000000
000000000000000000000000000000000000000000000000000000000000
000000000000000000000000000000000000000000000000000000000000
000000000000000000000000000000000000000000000000000000000011
111111111111
>DM_train1052
MGSDKIHHHHHHMYVTSQIHAKGSDLAKLMQIAALKGNEEVLDVATGGGHVANAFAPFVKKV
VAFDLTEDILKVARAFIEGNGHQQVEYVQGDAEQMPFTDERFHIVTCRIAAHHFPNPASFVS
EAYRVLKKGGQLLLVDNSAPENDAFDVFYNYVEKERDYSHHRAWKKSDWLKMLEEAGFELEE
LHCFHKTFIFEDWCDRMNVTTEKKQELSDFIKSKPTEYYQKFKIVVEDGRVYSFRGESILMK
ARKPTVKNKGSS
111111111111111111111100000000000000000000000000000000000000
000001111111111111111110000000000000000000000000000000000000
000000000000000000000000000000000000000000000000000000000000
000000000000000000000000000000000000000000000000000000000000
000001111111
>DM_train1053
TKFIGCIDLHNGEVKQIVGGTLTSKKEDVPKTNFVSQHPSSYYAKLYKDRDVQGCHVIKLGP
NNDDAAREALQESPQFLQVGGGINDTNCLEWLKWASKVIVTSWLFTKEGHFQLKRLERLTEL
CGKDRIVVDLSCRKTQDGRWIVAMNKWQTLTDLELNADTFRELRKYTNEFLIHAADVEGLCG
GIDELLVSKLFEWTKDYDDLKIVYAGGAKSVDDLKLVDELSHGKVDLTFGSSLDIFGGNLVK
FEDCCRWNEKQG
000000000000000011111111111111111111000000000000000000000000
000000000000000000000000000000000000000000000000000000000000
000000000000000000000000000000000000000000000000000011111110
000000000000000000000000000000000000000000000000000000000000
000000000000
>DM_train1054
MASVPENQAPKAIFTFSPEDPVTDENVVFNASNSIDEDGTIAYYVWDFGDGYEGTSTTPTIT
YKYKNPGTYKVKLIVTDNQGASSSFTATIKVTSATGDNSKFNFEDGTLGGFTTSGTNATGVV
VNTTEKAFKGERGLKWTVTSEGEGTAELKLDGGTIVVPGTTMTFRIWIPSGAPIAAIQPYIM
PHTPDWSEVLWNSTWKGYTMVKTDDWNEITLTLPEDVDPTWPQQMGIQVQTIDEGEFTIYVD
AIDWLEHHHHHH
111100000000000000000000000000000000000000000000000000000000
000000000000000000000000000000000000000000000000000000000000
000000000000000000000000000000000000000000000000000000000000
000000000000000000000000000000000000000000000000000000000000
000000111111

>DM_train1055
MGIMPRKVLAIKNFKERFDLLPKLKGVIAEKQPDILVVVGNILKNEALEKEYERAHLARREP
NRKVIHENEHYIIETLDKFFREIGELGVKTFVVPGKNDAPLKIFLRAAYEAETAYPNIRVLH
EGFAGWRGEFEVIGFGGLLTEHEFEEDFVLKYPRWYVEYILKFVNELKPRRLVTIFYTPPIG
EFVDRTPEDPKHHGSAVVNTIIKSLNPEVAIVGHVGKGHELVGNTIVVNPGEFEEGRYAFLD
LTQHKIKLEQFS
111000000000000000000000000000000000000000000000000000000000
000000000000000000000000000000000000000000000000000000000000
000000000000000000000000000000000000000000000000000000000000
000000000000000000000000000000000000000000000000000000000000
000000000000
>DM_train1056
MYTKNFSNSRMEVKGNNGCSAPIIRKPFKHIVLTVPSSDLDNFNTVFYVQPQYINQALHLAN
AFQGAIDPLNLNFNFEKALQIANGIPNSAIVKTLNQSVIQQTVEISVMVEQLKKIIQEVLGL
VINSTSFWNSVEATIKGTFTNLDTQIDEAWIFWHSLSAHNTSYYYNILFSIQNEDTGAVMAV
LPLAFEVSVDVEKQKVLFFTIKDSARYEVKMKALTLVQALHSSNAPIVDIFNVNNYNLYHSN
HKIIQNLNLSN
111111111111111111000000000000000000000000000000000000000000
000000000000000000000000000000000000000000000000000000000000
000000000000000000000000000000000000000000000000000000000000
000000000000000000000000000000000000000000000000000000000111
11111111111
>DM_train1057
MHRPEAEKWLRRFERAPDARARLVCLPHAGGSASFFFPLAKALAPAVEVLAVQYPGRQDRRH
EPPVDSIGGLTNRLLEVLRPFGDRPLALFGHSMGAIIGYELALRMPEAGLPAPVHLFASGRR
APSRYRDDDVRGASDERLVAELRKLGGSDAAMLADPELLAMVLPAIRSDYRAVETYRHEPGR
RVDCPVTVFTGDHDPRVSVGEARAWEEHTTGPADLRVLPGGHFFLVDQAAPMIATMTEKLAG
PALTGSTGGNS
100000000000000000000000000000000000000000000000000000000000
000000000000000000000000000000000000000000000000000000000000
000000000000000000000000000000000000000000000000000000000000
000000000000000000000000000000000000000000000000000000000001
11111111111
>DM_train1058
MSNENKTIHDFELNLICDFFSNMERQGPGSPEVTLKALSFIDNLTEKSLIADIGCGTGGQTM
VLAGHVTGQVTGLDFLSGFIDIFNRNARQSGLQNRVTGIVGSMDDLPFRNEELDLIWSEGAI
YNIGFERGLNEWRKYLKKGGYLAVSECSWFTDERPAEINDFWMDAYPEIDTIPNQVAKIHKA
GYLPVATFILPENCWTDHYFTPKVAAQKIFLTKYAGNKIAEEFSMLQSIEEELYHKYKEYYG
YTFFIAKKIRL
111111111110000000000000000000000000000000000000000000000000
000000000000000000000000000000000000000000000000000000000000
000000000000000000000000000000000000000000000000000000000000
000000000000000000000000000000000000000000000000000000000000
00000000000

>DM_train1059
MSMLTIGGKSFQSRLLLGTGKYPSFDIQKEAVAVSESDILTFAVRRMNIFEASQPNFLEQLD
LSKYTLLPNTAGASTAEEAVRIARLAKASGLCDMIKVEVIGCSRSLLPDPVETLKASEQLLE
EGFIVLPYTSDDVVLARKLEELGVHAIMPGASPIGSGQGILNPLNLSFIIEQAKVPVIVDAG
IGSPKDAAYAMELGADGVLLNTAVSGADDPVKMARAMKLAVEAGRLSYEAGRIPLKQYGTAS
SPGEGLPVLEH
100000000000000000000000000000000000000000000001111111 00000
000000000000000000000000000000000000000000000000000000000000
000000000000000000000000000000000000000000000000000000000000
000000000000000000000000000000000000000000000000000000000000
00001111111
>DM_train1060
MKKVIIAGNGPSLKEIDYSRLPNDFDVFRCNQFYFEDKYYLGKKCKAVFYNPSLFFEQYYTL
KHLIQNQEYETELIMCSNYNQAHLENENFVKTFYDYFPDAHLGYDFFKQLKDFNAYFKFHEI
YFNQRITSGVYMCAVAIALGYKEIYLSGIDFYQNGSSYAFDTKQKNLLKLAPNFKNDNSHYI
GHSKNTDIKALEFLEKTYKIKLYCLCPNSLLANFIELAPNLNSNFIIQEKNNYTKDILIPSS
EAYGKFSKNIN
000000000000000000000000000000000000000000000000000000000000
000000000000000000000000000000000000000000000000000000000000
000000000000000000000000000001111111111110000000000011111111
100000000000000000000000000000000000000000000000000000000000
00000000001
>DM_train1061
MKTITETLNLAPKGKNFLTADWPAPANVKTLITTRNGGVSQGAYQSLNLGTHVGDNPEAVRR
NREIVQQQVGLPVAYLNQIHSTVVVNAAEALGGTPDADASVDDTGKVACAVMTADCLPVLFC
DRAGTAVAAAHAGWRGLAGGVLQNTIAAMKVPPVEMMAYLGPAISADAFEVGQDVFDAFCTP
MPEAATAFEGIGSGKFLADLYALARLILKREGVGGVYGGTHCTVLERDTFFSYRRDGATGRM
ASLIWLDGNAV
111111111111110000000000000000000000000000000000000000000000
000000000000000000000000000000000000000000000000000000000000
000000000000000000000000000000000000000000000000000000000000
000000000000000000000000000000000000000000000000000000000000
00000000111
>DM_train1062
RFQYQALLNEHQSQLDRFSSHIVATLDKYAHIPHLISKDKELVDALLSAQNSAQIDITNRYL
EQVNEVIQAADTYLIDRFGNTIASSNWNLDRSFIGRNFAWRPYFYLSIAGQKSQYFALGSTS
GQRGYYYAYPVIYAAEILGVIVVKMDLSAIEQGWQNKSSYFVATDDHQVVFMSSQPAWLFHS
VADLSPAQLNDIRQSQQYLDSPIPSLGWQGDLQAEQSEWRKPEKHWLQDDYIVSSRPLPELA
LTIRVLSPKIE
000000000000000000000000000000000000000000000000000000000000
000000000000000000000000000000000000000000000000000000000000
000000000000000000000000000000000000000000000000000000000000
000000000000000000000000000000000000000000000000000000000000
00000000001

>DM_train1063
LRYNLWFGVYDGKEIKLSENFEESFLKAENPSPLPFNVSEVGAKALGKDYYRILRKTALAVS
EKMVEKELRREDRYVVALVKALEEIDESINMLNEKLEDIRAVKESEITEKFEKKIRELRELR
RDVEREIEEVMEKIAPNMTELVGAKVAAKLLERAGSMERLVRLPASKIQVIGAEKSLYKAFA
RMKKGKKAKIPKHGIIFLHPFIRTLPKAKRGKMARFLAAKLAIAAKIDYFRGEIDESLYESI
RRRYEELRRK
000000000000000000000000000000000000000000000000000000000000
000000000000000000000000000000000000000000000000000000000000
000000000000000000000000000000000000000000000000000011111111
111111111111000000000000000000000000000000000000000000000000
0000000000
>DM_train1064
MISGILASPGIAFGKALLLKEDEIVIDRKKISADQVDQEVERFLSGRAKASAQLETIKTKAG
ETFGEEKEAIFEGHIMLLEDEELEQEIIALIKDKHMTADAAAHEVIEGQASALEELDDEYLK
ERAADVRDIGKRLLRNILGLKIIDLSAIQDEVILVAADLTPSETAQLNLKKVLGFITDAGGR
TSHTSIMARSLELPAIVGTGSVTSQVKNDDYLILDAVNNQVYVNPTNEVIDKMRAVQEQVAS
EKAELAKLKD
000000000000000000000000000000000000000000000000000000000000
000000000000000000000000000000000000000000000000000000000000
000000000000000000000000000000000000000000000000000000000000
000000000000000000000000000000000000000000000000000000000000
0111111111
>DM_train1065
MAEKAQFYYPGETDVAENRRKYMNPNYELKKLREIPDEDIVRLMGHREPGEEYPSVHPPLEE
MEEPECPIRELVEPTEGAKAGDRIRYIQFTDSVYFAPIHPYIRARMYMWRYRGVDTGSLSGR
QIIEVRERDLEKIAKELLETEIFDPARSGVRGATVHGHALRLDENGLMLHALRRYRLNEETG
EVEYVKDQVGIELDEPIPVGAPADEDDLKERTTIYRIDGTPYREDEELLQVVQRIHELRTLA
GYRPEEAEGK
111111000000000000000000000000000000000000000000000000000000
000000000000000000000000000000000000000000000000000000000000
000000000000000000000000000000000000000000000000000000000000
000000000000000000000000000000000000000000000000000000000000
0000001111
>DM_train1066
MATVKLSFLQHICKLTGLSRSGRKDELLRRIVDSPIYPTSRVLGIDLGIKNFSYCFASQNED
SKVIIHNWSVENLTEKNGLDIQWTEDFQPSSMADLSIQLFNTLHEKFNPHVILMERQRYRSG
IATIPEWTLRVNMLESMLYALHYAEKRNSIEQKIQYPFLLSLSPKSTYSYWASVLNTKASFS
KKKSRVQMVKELIDGQKILFENEEALYKWNNGSRVEFKKDDMADSALIASGWMRWQAQLKHY
RNFCKQFLKQ
111100000000000000000111100000000000000000000000000000000000
000000000000000000000000000000000000000000000000000000000111
111110000000000000000000000000000000000000000000000000111111
111000000000000000000000000000001110000000000000000000000000
0000000000

>DM_train1067
MPVFHTRTIESILEPVAQQISHLVIMHEEGEVDGKAIPDLTAPVAAVQAAVSNLVRVGKETV
QTTEDQILKRDMPPAFIKVENACTKLVQAAQMLQSDPYSVPARDYLIDGSRGILSGTSDLLL
TFDEAEVRKIIRVCKGILEYLTVAEVVETMEDLVTYTKNLGPGMTKMAKMIDERQQELTHQE
HRVMLVNSMNTVKELLPVLISAMKIFVTTKNSKNQGIEEALKNRNFTVEKMSAEINEIIRVL
QLTSWDEDAW
000000000000000000000000000111111000000000000000000000000000
000000000000000000000000000000000000000000000000000000000000
000000000000000000000000000000000000000000000000000000000000
000000000000000000000000000000000000000000000000000000000000
0000000000
>DM_train1068
MGSDKIHHHHHHMNDFKNQWLRKRTFAIPASRLTGRLTTLKSDVPAADSLFWKLWNGSLDTA
VQVLQTDYFKGIAAGTLDPNAYGSLMVQDGYYCFRGRDDYATAATCAQDETLREFFKAKAKS
YDEYNETYHQTWHLREASGLIPGTDIKDYADYEAYVAGSLASPYMCVVMLPCEYLWPWIANF
LDGYTPTNSLYRFWIEWNGGTPNGAYQMGNMLEQYRDKIDEDKAVEIFNTAMNYELKVFTSS
TILTTIENGK
111111111111111111110000000000000000000000000000000000000000
000000000000000000000000000000000000000000000000000000000000
000000000000000000000000000000000000000000000000000000000000
000000000000000000000000000000000000000000000000000000000000
0000111111
>DM_train1069
SGTTEKTVAKEEKVKLTDQQLMADLWYQTAGEMKALYYQGYNTGQLKLDAALAKGTEKKPAI
VLDLDETVLDNSPHQAMSVKTGKGYPYKWDDWINKAEAEALPGSIDFLKYTESKGVDIYYIS
NRKTNQLDATIKNLERVGAPQATKEHILLQDPKEKGKEKRRELVSQTHDIVLFFGDNLSDFT
GFDGKSVKDRNQAVTDSKAQFGEKFIIFPNPMYGDWEGALYDYNFKKSDAEKDKIRHDNLKS
FDAKHHHHHH
111111111111100000000000000000000000000000000000000000000000
000000000000000000000000000000000000000000000000000000000000
000000000000000000000000000000001000000000000000000000000000
000000000000000000000000000000000000000000000000000000000000
0011111111
>DM_train1070
GSVTSANLDHTKPCWYWDKKDLAHTPSQLEGLDPATEARYRREGARFIFDVGTRLGLHYDTL
ATGIIYFHRFYMFHSFKQFPRYVTGACCLFLAGKVEETPKKCKDIIKTARSLLNDVQFGQFG
DDPKEEVMVLERILLQTIKFDLQVEHPYQFLLKYAKQLKGDKNKIQKLVQMAWTFVNDSLCT
TLSLQWEPEIIAVAVMYLAGRLCKFEIQEWTSKPMYRRWWEQFVQDVPVDVLEDICHQILDL
YSQGKQQMPH
111100000000000000000000000000000000000000000000000000000000
000000000000000000000000000000000000000000000000000000000000
000000000000000000000000000000000000000000000000000000000000
000000000000000000000000000000000000000000000000000000000000
0000000000

>DM_train1071
MVEDSRVRDALKGGDQKALPASLVPQAPPVLTSKDKITKRMIVVLAMASLETHKISSNGPGG
DKYVLLNCDDHQGLLKKMGRDISEARPDITHQCLLTLLDSPINKAGKLQVYIQTSRGILIEV
NPTVRIPRTFKRFSGLMVQLLHKLSIRSVNSEEKLLKVIKNPITDHLPTKCRKVTLSFDAPV
IRVQDYIEKLDDDESICVFVGAMARGKDNFADEYVDEKVGLSNYPLSASVACSKFCHGAEDA
WNILHHHHHH
111111111111111111111110000000000000000000000000000001111111
110000000000000000000000000000000000000000000000000000000000
000000000000000000000000000000000000000000000000000000000000
000000000000000000000000000000000000000000000000000000000000
0001111111
>DM_train1072
MSTGITYDEDRKTQLIAQYESVREVVNAEAKNVHVNENASKILLLVVSKLKPASDIQILYDH
GVREFGENYVQELIEKAKLLPDDIKWHFIGGLQTNKCKDLAKVPNLYSVETIDSLKKAKKLN
ESRAKFQPDCNPILCNVQINTSHEDQKSGLNNEAEIFEVIDFFLSEECKYIKLNGLMTIGSW
NVSHEDSKENRDFATLVEWKKKIDAKFGTSLKLSMGMSADFREAIRQGTAEVRIGTDIFGAR
PPKNEARII
111000000000000000000000000000001111100000000000000000000000
000000000000000000000000000000000000000000000000000000000000
000000000000000000000000000000000000000000000000000000000000
111111110000000000000000000000000000000000000000000000000011
111111111
>DM_train1073
MATANNPPSGLLDPTLCEDRIFYNILEIEPRFLTSDSVFGTFQQSLTSHMRKLLGTWMFSVC
QEYNLEPNVVALALNLLDRLLLIKQVSKEHFQKTGSACLLVASKLRSLTPISTSSLCYAAAD
SFSRQELIDQEKELLEKLAWRTEAVLATDVTSFLLLKLVGGSQHLDFWHHEVNTLITKALVD
PLTGSLPASIISAAGCALLVPANVIPQDTHSGGVVPQLASILGCDVSVLQAAVEQILTSVSD
FDLRILDSY
111111111111111000000000000000000000000000000000000000000000
000000000000000000000000000000000000000000000000000000000000
000000000000000000000000000000000000000000000000000000000000
000000000000000000000000001111100000000000000000000000000000
000001111
>DM_train1074
AKQNVSSLDEKNSVSVDLPGEMKVLVSKEKNKDGKYDLIATVDKLELKGTSDKNNGSGVLEG
VKADKCKVKLTISDDLGQTTLEVFKEDGKTLVSKKVTSKDKSSTEEKFNEKGEVSEKIITRA
DGTRLEYTGIKSDGSGKAKEVLKGYVLEGTLTAEKTTLVVKEGTVTLSKNISKSGEVSVELN
DTDSSAATKKTAAWNSGTSTLTITVNSKKTKDLVFTKENTITVQQYDSNGTKLEGSAVEITK
LDEIKNALK
111111000000000000000000000000000000000000000000000000000000
000000000000000000000000000000000000000000000000000000000000
000000000000000000000000000000000000000000000000000000000000
000000000000000000000000000000000000000000000000000000000000
000000000

>DM_train1075
MCELLGMSANVPTDIVFSFTGLMQRGGGTGPHRDGWGIAFYEGRGVRLFQDPLASVDSEVAR
LVQRFPIKSETVIGHIRQANVGKVGLSNTHPFIRELGGRYWTFAHNGQLADFQPKPGFYRPV
GETDSEAAFCDLLNRVRRAFPEPVPVEVLLPVLISACDEYRKKGVFNALISDGDWLFTFCSS
KLAYITRRAPFGPARLKDADLTVDFHAETTPDDVVTVIATEPLTDNENWTLQQSGEWVLWWG
GEVLAKGRV
100000000000000000000000000000000000000000000000000000000000
000000000000000000000000000000000000000000000000000000000000
000000000000000000000000000000000000000000000000000000000000
000000000000000000000000000000000000000000000000000000000000
000000111
>DM_train1076
GMAGAAHRLMEARMRRYDNKFARISDIDINQPESWRGRIFLTFDIDWAADFVLQDTIDLIEG
AGVCATWFATHSTPLLENIRRNPLFELGVHPNFNPLLAGAHAEGVQEILDRTLELAPGCVSV
RSHSLVQATSILNMFGERRLRYDCNILVPWDAGIVLQPWRHWTGDMVRVPYLWEDDVACLYD
WEFDSTFDYWYQPDGINVLDFHPIHVYMNTESLRRYEDSREVHRNPVDLIRWRNTSAGSRTF
LQSLLARNI
111111111111111000000000000000000000000000000000000000000000
000000000000000000000000000000000000000000000000000000000000
000000000000000000000000000000000000000000000000000000000000
000000000000000000000000000000000000000000000000000000000000
000000000
>DM_train1077
PDPMKNTCKLLVVADHRFYRYMGRGEESTTTNYLIELIDRVDDIYRNTAWDNAGFKGYGIQI
EQIRILKSPQEVKPGEKHYNMAKSYPNEEKDAWDVKMLLEQFSFDIAEEASKVCLAHLFTYQ
DFDMGTLGLAYGGSPRANSHGGVCPKAYYSPVGKKNIYLNSGLTSTKNYGKTILTKEADLVT
THELGHNFGAEHDPDGLAECAPNEDQGGKYVMYPIAVSGDHENNKMFSQCSKQSIYKTIESK
AQECFQERS
000000000000000000000000000000000000000000000000000000000000
000000000000000000000000000000000000000000000000000000000000
000000000000000111100000000000000000000000000000000000000000
000000000000000000000000000000000000000000000000000000000000
000000000
>DM_train1078
MTVRSRFFTEAEGKAVGVENAAAKGDVLLVCEHASATIPQKYGTLGLSADVLSSHAAWDPGA
LAVARLLSEKFHATLVYQRFSRLVYDCNRPPESPSAMPVKSEIYDIPGNFDLDEAERFARTS
ALYVPFHDRVSEIIAERQAAGRKVVVVTIHSFTPVYHGRFREVEIGILHDNDSRLADAMLAG
AEGASLTVRRNDPYGPEDGVTHTLRLHALPDGLLNVMIEIRNDLIANEGEQAAIAGFLHELM
GKALSSIEE
111111000000000000000000000000000000000000000000000000000000
000000000000000000000000000000000000000000000000000000000000
000000000000000000000000000000000000000000000000000000000000
000000000000000000000000000000000000000000000000000000000000
000000011

>DM_train1079
MPVEITVKELLEAGVHFGHERKRWNPKFARYIYAERNGIHIIDLQKTMEELERTFRFIEDLA
MRGGTILFVGTKKQAQDIVRMEAERAGMPYVNQRWLGGMLTNFKTISQRVHRLEELEALFAS
PEIEERPKKEQVRLKHELERLQKYLSGFRLLKRLPDAIFVVDPTKEAIAVREARKLFIPVIA
LADTDSDPDLVDYIIPGNDDAIRSIQLILSRAVDLIIQARGGVVEPSPSYALVQEAEATETP
EGESEVEA
11111100000000000000000000000000000000000000000000000000000000
00000000000000000000000000000000000000000000000000000000000000
00000000000000000000000000000000000000000000000000000000000000
00000000000000000000000000000000000000000000000000000011111111
11111111
>DM_train1080
MDFTKPETVLNLQNIRDELVRMEDSIIFKFIERSHFATCPSVYEANHPGLEIPNFKGSFLDW
ALSNLEIAHSRIRRFESPDETPFFPDKIQKSFLPSINYPQILAPYAPEVNYNDKIKKVYIEK
IIPLISKRDGDDKNNFGSVATRDIECLQSLSRRIHFGKFVAEAKFQSDIPLYTKLIKSKDVE
GIMKNITNSAVEEKILERLTKKAEVYGVDPTNESGERRITPEYLVKIYKEIVIPITKEVEVE
YLLRRLEE
00000000000000000000000000000000000000000000000000000000000000
00000000000000000000000000000000000000000000000000000000000000
00000000000000000000000000000000000000000000000000000000000000
00000000000000000000000000001111111100000000000000000000000000
00000000
>DM_train1081
MTMDEQQSQAVAPVYVGGFLARYDQSPDEAELLLPRDVVEHWLHAQGQGQPSLSVALPLNIN
HDDTAVVGHVAAMQSVRDGLFCLGCVTSPRFLEIVRRASEKSELVSRGPVSPLQPDKVVEFL
SGSYAGLSLSSRRCDDAEAPTSLSGSETTPFKHVALCSVGRRRGTLAVYGRDPEWVTQRFPD
LTAADRDGLRAQWQRCGSTAVDASGDPFRSDSYGLLGNSVDALYIRERLPKLRYDKQLVGVT
ERESYVKA
11111111110000000000000000001111000000000111111111111100000000
00000000000000000000000000000000000000000000000000000000000000
00000000001111111111111111111000000000000000000000000000000000
00000000000001111111111100000000000000000000000000000000000000
00000000
>DM_train1082
RPQPGDGERRYREASARKKIRLDRKYIVSCKQTEVPLSVPWDPSNQVYLSYNNVSSLKMLVA
KDNWVLSSEISQVRLYTLEDDKFLSFHMEMVVHVDAAQAFLLLSDLRQRPEWDKHYRSVELV
QQVDEDDAIYHVTSPALGGHTKPQDFVILASRRKPCDNGDPYVIALRSVTLPTHRETPEYRR
GETLCSGFCLWREGDQLTKVSYYNQATPGVLNYVTTNVAGLSSEFYTTFKACEQFLLDNRND
LAPSLQTL
11111111111110000000000000001111111100000000000000000000000000
00000000000000000000000000000000000000000000000000000000000000
00000000000000000000000000000000000000000000000000000000000000
00000000000000000000000000000000000000000000000000000000000000
00000011

>DM_train1084
GSCEKENIGIEVTPVNAKFIITPVVIDATTGTDVTQSAEISFSKGNGTYEGTPELASESINI
NAKYKGMTGSASVTIPALKAGQFGAKEVTIILSENFFAQEESSNSQIETTKHSGFKNNTSDY
WYYITVTYTKKEGSEVIKNDYEGDDSEIKNIIDAYNKGVREDKVTLNDVQVLAHSRFSVFVD
YMKTTSVYQIIEKSPKRDGNPVASFTVDSYNTIVSPKNEQIPGHGHAPSHGHGHGHGDDSNA
GGGIIIAD
11111111111100000000000000000000000000000000000000000000000000
00000000000000000000000000000000000000000000000000000000000000
00000000000000000000000000000000000000000000000000000000000000
00000000000000011000000000000000000000000000000000000011111111
11111111
>DM_train1085
MAAESQLKRVIETLRRLGIEEVLKLERRDPQYRAVCNVVKRHGETVGSRLAMLNALISYRLT
GKGEEHWEYFGKYFSQLEVIDLCRDFLKYIETSPFLKIGVEARKKRALKACDYVPNLEDLGL
TLRQLSHIVGARREQKTLVFTIKILNYAYMCSRGVNRVLPFDIPIPVDYRVARLTWCAGLID
FPPEEALRRYEAVQKIWDAVARETGIPPLHLDTLLWLAGRAVLYGENLHGVPKEVIALFQWR
GGCRPPSE
10000000000000000000000000000000000000000000000000000000000000
00000000000000000000000000000000000000000000000000000000000000
00000000000000000000000000000000000000000000000000000000000000
00000000000000000000000000000000000000000000000000000000000000
00000011
>DM_train1086
SEPVTIVLSQMGWVRSAKGHDIDAPGLNYKAGDSFKAAVKGKSNQPVVFVDSTGRSYAIDPI
TLPSARGQGEPLTGKLTLPPGATVDHMLMESDDQKLLMASDAGYGFVCTFNDLVARNRAGKA
LITLPENAHVMPPVVIEDASDMLLAITQAGRMLMFPVSDLPQLSKGKGNKIINIPSAEAARG
EDGLAQLYVLPPQSTLTIHVGKRKIKLRPEELQKVTGERGRRGTLMRGLQRIDRVEIDSPRR
ASSGDSEE
00000000000000000000000000000000000000000000000000000000000000
00000000000000000000000000000000000000000000000000000000000000
00000000000000000000000000000000000000000000000000000000000000
00000000000100000000000000000000000000000000000000000000000011
11111111
>DM_train1087
GSHPLGDTSDASKLSSDYSLPDLINTRKVPNNWQTGEQASLEEGRIVLTSNQNSKGSLWLKQ
GFDLKDSFTMEWTFRSVGYSGQTDGGISFWFVQDSNIPRDKQLYNGPVNYDGLQLLVDNNGP
LGPTLRGQLNDGQKPVDKTKIYDQSFASCLMGYQDSSVPSTIRVTYDLEDDNLLKVQVDNKV
CFQTRKVRFPSGSYRIGVTAQNGAVNNNAESFEIFKMQFFNGVIEDSLIPNVNAMGQPKLIT
KYIDQQTG
11111111100000000000000000000000000000000000000000000000000000
00000000000000000000000000000000000000000000000000000000000000
00000000000000000000000000000000000000000000000000000000000000
00000000000000000000000000000000000000000000001111111111110000
00011111

>DM_train1088
MTNESILESYSGVTPERKKSRMPAKLDWWQSATGLFLGLFMIGHMFFVSTILLGDNVMLWVT
KKFELDFIFEGGKPIVVSFLAAFVFAVFIAHAFLAMRKFPINYRQYLTFKTHKDLMRHGDTT
LWWIQAMTGFAMFFLGSVHLYIMMTQPQTIGPVSSSFRMVSEWMWPLYLVLLFAVELHGSVG
LYRLAVKWGWFDGETPDKTRANLKKLKTLMSAFLIVLGLLTFGAYVKKGLEQTDPNIDYKYF
DYKRTHEE
000000000000000000000000000000000000000000000000000000000000
000000000000000000000000000000000000000000000000000000000000
000000000000000000000000000000000000000000000000000000000000
000000000000000000000000000000000000000000000000000000000000
00000001
>DM_train1089
MHHHHHHGTSLYKKAGSENLYFQGITTRTARQCGQADYGWLQARYTFSFGHYFDPKLLGYAS
LRVLNQEVLAPGAAFQPRTYPKVDILNVILDGEAEYRDSEGNHVQASAGEALLLSTQPGVSY
SEHNLSKDKPLTRMQLWLDACPQRENPLIQKLALNMGKQQLIASPEGAMGSLQLRQQVWLHH
IVLDKGESANFQLHGPRAYLQSIHGKFHALTHHEEKAALTCGDGAFIRDEANITLVADSPLR
ALLIDLPV
111111111111111111111111111110000000000000000000000000000000
000000000000000000000000000000000000000000000000000000000000
000000000000000000000000000000000000000000000000000000000000
000000000000000000000000000000000000000000000000000000000000
00000000
>DM_train1091
PKVSVIMTSYNKSDYVAKSISSILSQTFSDFELFIMDDNSNEETLNVIRPFLNDNRVRFYQS
DISGVKERTEKTRYAALINQAIEMAEGEYITYATDDNIYMPDRLLKMVRELDTHPEKAVIYS
ASKTYHLNENRDIVKETVRPAAQVTWNAPCAIDHCSVMHRYSVLEKVKEKFGSYWDESPAFY
RIGDARFFWRVNHFYPFYPLDEELDLNYITDQSIHFQLFELEKNEFVRNLPPQRNCRELRES
LKKLGMG
000000000000000000000000000000000000000000000000000000000000
000000000000000000000000000000000000000000000000000000000000
000000001110000000000000000000000000000000000000000000000000
000000000000000000000000000000111111111111110000000000000000
0000000
>DM_train1092
MIMPQHDQLHRYLFENFAVRGELVTVSETLQQILENHDYPQPVKNVLAELLVATSLLTATLK
FDGDITVQLQGDGPMNLAVINGNNNQQMRGVARVQGEIPENADLKTLVGNGYVVITITPSEG
ERYQGVVGLEGDTLAACLEDYFMRSEQLPTRLFIRTGDVDGKPAAGGMLLQVMPAQNAQQDD
FDHLATLTETIKTEELLTLPANEVLWRLYHEEEVTVYDPQDVEFKCTCSRERCADALKTLPD
EEVDSIL
111110000000000000000000000000000000000000000000000000000000
000000000000000000000000000000000000000000000000000000000000
000000000000000000000000000000000000000000000000000000000000
000000000000000000000000000000000000000000000011111111111111
1111111

>DM_train1093
ATPWQKITQPVPGSAQSIGSFSNGCIVGADTLPIQSEHYQVMRTDQRRYFGHPDLVMFIQRL
SSQVSNLGMGTVLIGDMGMPAGGRFNGGHASHQTGLDVDIFLQLPKTRWTSAQLLRPQALDL
VSRDGKHVVSTLWKPEIFSLIKLAAQDKDVTRIFVNPAIKQQLCLDAGTDRDWLRKVRPWFQ
HRAHMHVRLRCPADSLECEDQPLPPSGDGCGAELQSWFEPPKPGTTKPEKKTPPPLPPSCQA
LLDEHVI
111000000000000000000000000000000000000000000000000000000000
000000000000000000000000000000000000000000000000000000000000
000000000000000000000000000000000000000000000000000000000000
000000000000000000000000000000000000011111111111111000000000
0000011
>DM_train1094
MRPVYFLSDFGLEDPYVAVVKAVLAERAPGPAVVDLAHALPPQDLRRAAYALFEALPYLPEG
AVVLAVVDPGVGTARRAVAALGRWTYVGPDNGLFTLAWLLDPPRRAFLLEPPRPRPKAALPG
WAPGEATFHGRDVFAPAAAHLALGLPPEGLGPEVPVETLARLPLALTEGPEGEVLTFDRFGN
AITTLLRAPVGGFVEVGGRRVPVRRTFGEVPEGAPVAYLGSAGLLEVAVNRGSAREALGLKE
GMPVRLL
000000000000000000000000000000000000000000000000000000000000
000000011111100000000000000000000000000000000000111111111111
111111111000000000000000000000000000000000000000000000000000
000000000000000000000000001111000000000000000000000000000000
0000000
>DM_train1095
MPAEIPEFKKLGLPDKVLELCHRKMGLILVTGPTGSGKSTTIASMIDYINQTKSYHIITIED
PIEYVFKHKKSIVNQREVGEDTKSFADALRAALREDPDVIFVGEMRDLETVETALRAAETGH
LVFGTLHTNTAIDTIHRIVDIFPLNQQEQVRIVLSFILQGIISQRLLPKIGGGRVLAYGLLI
PNTAIRNLIRENKLQQVYSLMQSGQAETGMQTMNQTLYKLYKQGLITLEDAMEASPDPKELE
RMIRGGR
111110000000000000000000000000000000000000000000000000000000
000000000000000000000000000000000000000000000000000000000000
000000000000000000000000000000000000000000000000000000000000
000000000000000000000000000000000000000000000000000000000000
0000111
>DM_train1096
MGSSSSGLVPRGSHMTTFHDLPLEERLTLARLGTSHYSRQLSLVDNAEFGEHSLLEGWTRSH
LIAHVAYNAIALCNLMHWANTGEETPMYVSPEARNEEIAYGSTLNPDALRNLHEHSVARLDV
AWRETSEDAWSHEVLTAQGRTVPASETLWMRSREVWIHAVDLGAVATFGDIPEVILRTLAAE
ITQKWTSQGAGEGLVLLDEPSSTRYPAAPGQDEVVVSGSLAGIVRYAAGRGSDGVTSSTGEV
PEPPRWL
111111111111110000000000000000000000000000000000000000000000
000000000000000000000000000000000000000000000000000000000000
000000000000000000000000000000000000000000000000000000000000
000000000000000000000000000000000000000000000000000000000000
0000001

>DM_train1097
GMDKYLSANSLEGVIDNEFSMPAPRWLNTYPAGPYRFINREFFIIAYETDPDLLQAILPPDM
ELLEPVVKFEFIRMPDSTGFGDYTESGQVVPVRYKGEEGGFTISMFLDCHAPIAGGREIWGF
PKKLAKPKLFVEEDTLIGILKYGSIDIAIATMGYKHRPLDAEKVLESVKKPVFLLKNIPNVD
GTPLVNQLTKTYLTDITVKGAWTGPGSLELHPHALAPISNLYIKKIVSVSHFITDLTLPYGK
VVADYLA
111111000000000000000000000000000000000000000000000000000000
000000000000000000000000000000000000000000000000000000000000
000000000000000000000000000000000000000000000000000000000000
000000000000000000000000000000000000000000000000000000000000
0000000
>DM_train1098
MLDVAPPVITPRGTKIEPSAGAPFEAVRVARDVLHTSRTAALATLDPVSGYPYTTATNIGIE
PDGTPFFFAAGLTLHARNMETDARISVTLAPFGKGDALTLPRLTLVGRADRIGPDEVPLAIA
RYIARYPKAKLYLSLPDTRLYRLRTEGVQINGGPARNASNITPADLRTDLSGAEELMAAAES
EATRLNAIKGEASRLAVLAGAKTGRWKITSIDPDGIDLASASDLARLWFAERVETLKQFEKA
LAQLLK
111100000000111111110000000000000000000000000000000000000000
000000000000000000000000000000000000000000000000000000000000
000000000000000000000000000000001111111000000000000000000000
000000000000000000000000000000000000000000000000000000000000
000011
>DM_train1099
KDQIPIILIHGSGGNASSLDKMADQLMNEYRSSNEALTMTVNSEGKIKFEGKLTKDAKRPII
KFGFEQNQATPDDWSKWLKIAMEDLKSRYGFTQMDGVGHSNGGLALTYYAEDYAGDKTVPTL
RKLVAIGSPFNDLDPNDNGMDLSFKKLPNSTPQMDYFIKNQTEVSPDLEVLAIAGELSEDNP
TDGIVPTISSLATRLFMPGSAKAYIEDIQVGEDAVHQTLHETPKSIEKTYWFLEKFKTDETV
IQLDYK
100000000000000000000000000000000000000000000000000000000000
000000000000000000000000000000000000000000000000000000000000
000000000000000000000000000000000000000000000000000000000000
000000000000000000000000000000000000000000000000000000000000
000000
>DM_train1100
MSKPWLFTVHGTGQPDPLGPGLPADTARDVLDIYRWQPIGNYPAAAFPMWPSVEKGVAELIL
QIELKLDADPYADFAMAGYSQGAIVVGQVLKHHILPPTGRLHRFLHRLKKVIFWGNPMRQKG
FAHSDEWIHPVAAPDTLGILEDRLENLEQYGFEVRDYAHDGDMYASIKEDDLHEYEVAIGRI
VMKASGFIGGRDSVVAQLIELGQRPITEGIALAGAIIDALTFFARSRMGDKWPHLYNRYPAV
EFLRQI
100000000000000000000000000000000000000000000000000000000000
000000000000000000000000000000000000000000000000000000000000
000000000000000000000000000000000000000000000000000000000000
000000000000000000000000000000000000000000000000000000000000
000001

>DM_train1102
MWVYRLKGTLEALDPILPGLFDGGARGLWEREGEVWAFFPAPVDLPYEGVWEEVGDEDWLEA
WRRDLKPALAPPFVVLAPWHTWEGAEIPLVIEPGMAFGTGHHETTRLALKALARHLRPGDKV
LDLGTGSGVLAIAAEKLGGKALGVDIDPMVLPQAEANAKRNGVRPRFLEGSLEAALPFGPFD
LLVANLYAELHAALAPRYREALVPGGRALLTGILKDRAPLVREAMAGAGFRPLEEAAEGEWV
LLAYGR
00000000000000000000000000000000000000000000000000000000000000
00000000000000000000000000000000011111000000000000000000000000
00000000000000000000000000000000000000000000000000000000000000
00000000000000000000000000000000000000000000000000000000000000
000000
>DM_train1103
GAMSSKNEWRKSAIANTLLYLRLKNIYVSADDFVEEQNVYVLPKNLLKKFIEISDVKIQVAA
FIYGMSAKDHPKVKEIKTVVLVPQLGHVGSVQISNIPDIGDLPDTEGLELLGWIHTQTEELK
FMAASEVATHSKLFADKKRDCIDISIFSTPGSVSLSAYNLTDEGYQWGEENKDIMNVLSEGF
EPTFSTHAQLLLSDRITGNFIIPSGNVWNYTFMGTAFNQEGDYNFKYGIPLEFYNEMHRPVH
FLQFSE
11110000000000000000000000000000000000000000000000000000000000
00000000000000000000000000000000000000000000000000000000000000
00000000000000000000000000000000000000000000000000000000000000
00000000000000000000000000000000000000000000000000000000000000
000001
>DM_train1104
SFQEQTTKSRDVNSFQIPLRDGVRELLPEDASRNRASIKSPVDIWIGGENMTALNGIVDGGR
KFEAGQEFQINTFGSVNYWVSDEEIRVFKEYSARAKYAQNEGRTALEANNVPFFDIDVPPEL
DGVPFSLKARVRHKSKGVDGLGDYTSISVKPAFYITEGDETTDTLIKYTSYGSTGSHSGYDF
DDNTLDVMVTLSAGVHRVFPVETELDYDAVQEVQHDWYDESFTTFIEVYSDDPLLTVKGYAQ
ILMERT
11100000000000000000000000000000000000000000000000000000000000
00000000000000000000000000000000000000000000000000000000000000
00000000000000000000000000000000000000000000000000000000000000
00000000000000000000000000000000000000000000000000000000000000
000000
>DM_train1107
MAHTGQTAKSQGDTATMELTAKPRTPKQKLDESMIAAVAYNKENNVSFALDRKAFDRAFRQQ
STTGLFDITVEGGETFPALVKAVQMDKRKRAPIHVDFYMVTYGEPVEVSVPVHTTGRSQGEV
QGGLVDIVVHNLQIVAPGPRRIPQELVVDVTKMNIGDHITAGDIKLPEGCTLAADPELTVVS
VLPPRLTAEELEAEVQAAQVAGLVAAGELSEEAAEAVLEGDASLEEVKAEASEDNAGTDSED
NSDAQ
11111111111111110000000000000000000000000000000000000000000000
00000000000000000000000000000000000000000000000000000000000000
00000000000000000000000000000000000000000000000000000000000000
00000000000000000000000000000000000000000000000000001111111111
11111

>DM_train1108
MGSDKIHHHHHHMTVLIIGMGNIGKKLVELGNFEKIYAYDRISKDIPGVVRLDEFQVPSDVS
TVVECASPEAVKEYSLQILKNPVNYIIISTSAFADEVFRERFFSELKNSPARVFFPSGAIGG
LDVLSSIKDFVKNVRIETIKPPKSLGLDLKGKTVVFEGSVEEASKLFPRNINVASTIGLIVG
FEKVKVTIVADPAMDHNIHIVRISSAIGNYEFKIENIPSPENPKTSMLTVYSILRTLRNLES
KIIFG
111111111100000000000000000000000000000000000000000000000000
000000000000000000000000000000000000000000000000000000000000
000000000000000000000000000000000000000000000000000000000000
000000000000000000000000000000000000111111110000000000000000
00000
>DM_train1109
MAMQVPPLTELSPSISVHLLLGNPSGATPTKLTPDNYLMVKNQYALSYNNSKGTANWVAWQL
NSSWLGNAERQDNFRPDKTLPAGWVRVTPSMYSGSGYARGHIAPSADRTKTTEDNAATFLMT
NMMPQTPDNNRNTWGNLEDYCRELVSQGKELYIVAGPNGSLGKPLKGKVTVPKSTWKIVVVL
DSPGSGLEGITANTRVIAVNIPNDPELNNDWRAYKVSVDELESLTGYDFLSNVSPNIQTSIE
SKVDN
111111111111110000000000000000000000000000000000000000000000
000000000000000000000000000000000000000000000000000000000000
000000000000000000000000000000000000000000000000000000000000
000000000000000000000000000000000000000000000000000000000000
00000
>DM_train1110
MNNQGSIISVSLGPGDPGLITVKALSQLREADVIYYPGTVSASGAVTSVALDILKEFDLDPS
KLRGMLVPMSRSRGAAEASYAANYASMAEEVQAGRRVAVVSVGDGGFYSTASAIIERARRDG
LDCSMTPGIPAFIAAGSAAGMPLALQSDSVLVLAQIDEIGELERALVTHSTVVVMKLSTVRD
ELVSFLERYAKPFLYAEKVGMAGEFITMEVDALRSRAIPYFSLLVCSPHCRQSTLSPFASKL
AAALE
111100000000000000000000000000000000000000000000000000000000
000000000111111111000000000000000000000000000000000000000000
000000000000000000000000000000000000000000000000000000000000
000000000000000000000000000000000000000000000000000000111111
11111
>DM_train1111
MSKVVPIASLTPYQSKWTICARVTNKSQIRTWSNSRGEGKLFSLELVDESGEIRATAFNEQV
DKFFPLIEVNKVYYFSKGTLKIANKQFTAVKNDYEMTFNNETSVMPCEDDHHLPTVQFDFTG
IDDLENKSKDSLVDIIGICKSYEDATKITVRSNNREVAKRNIYLMDTSGKVVTATLWGEDAD
KFDGSRQPVLAIKGARVSDFGGRSLSVLSSSTIIANPDIPEAYKLRGWFDAEGQALDGVSIS
DLKS
000000000000000000000000000000000000000000000000000000000000
000000000000000000000000000000000000000000000111111110000000
000000000000000000000000000000000000000000000000000000000000
000000000000000000000000000000000000000000000000000000000011
1111

>DM_train1112
SNAMTTPIRVVVWNEFRHEKKDEQVRAIYPEGMHTVIASYLAEAGFDAATAVLDEPEHGLTD
EVLDRCDVLVWWGHIAHDEVKDEVVERVHRRVLEGMGLIVLHSGHFSKIFKKLMGTTCNLKW
READEKERLWVVAPGHPIVEGIGPYIELEQEEMYGEFFDIPEPDETIFISWFEGGEVFRSGC
TFTRGKGKIFYFRPGHETYPTYHHPDVLKVIANAVRWAAPVNRGEIVFGNVKPLEPIKAKQG
GVTQ
111110000000000000000000000000000000000000000000000000000000
000000000000000000000000000000000000000000000000000000000000
000000000000000000000000000000000000000000000000000000000000
000000000000000000000000000000000000000000000000000000000111
1111
>DM_train1113
PGMAGDSEQTLQNHQQPNGGEPFLIGVSGGTASGKSSVCAKIVQLLGQNEVDYRQKQVVILS
QDSFYRVLTSEQKAKALKGQFNFDHPDAFDNELILKTLKEITEGKTVQIPVYDFVSHSRKEE
TVTVYPADVVLFEGILAFYSQEVRDLFQMKLFVDTDADTRLSRRVLRDISERGRDLEQILSQ
YITFVKPAFEEFCLPTKKYADVIIPRGADNLVAINLIVQHIQDILNGGPSKRQTNGCLNGYT
PSRK
111111111111111111100000000000000000000000000000000000000000
000000000000000000000000000000000000000000000000000000000000
000000000000000000000000000000000000000000000000000000000000
000000000000000000000000000000000000000000000111111111111111
1111
>DM_train1114
LHWPTSLPSGDAFSSVGTHRFVQKVEEMVQNHMTYSLQDVGGDANWQLVVEEGEMKVYRREV
EENGIVLDPLKATHAVKGVTGHEVCNYFWNVDVRNDWETTIENFHVVETLADNAIIIYQTHK
RVWPASQRDVLYLSVIRKIPALTENDPETWIVCNFSVDHDSAPLNNRCVRAKINVAMICQTL
VSPPEGNQEISRDNILCKITYVANVNPGGWAPASVLRAVAKREYPKFLKRFTSYVQEKTAGK
PILF
111111111111111110000000000000000000000000000000000000000000
000000000000000000000000000000000000000000000000000000000000
000000000000000000000000000000000000000000000000000000000000
001110000000000000000000000000000000000000000000000000000000
0000
>DM_train1115
MGSDKIHHHHHHMSLDMKEHPDAEVQKNQVLTLEDWKEKWVTRHISFHQEQGHQLLKKHLDT
FLKGQSGLRVFFPLCGKAIEMKWFADRGHTVVGVEISEIGIREFFAEQNLSYTEEPLAEIAG
AKVFKSSSGSISLYCCSIFDLPRANIGKFDRIWDRGALVAINPGDHDRYADIILSLLRKEFQ
YLVAVLSYDPTKHAGPPFYVPSAELKRLFGTKCSMQCLEEVDALEERHKAWGLDYLFEKLYL
LTEK
111111111111111111111000000000000000000000000000000000000000
000000000000000000000000000000000000000000000000000000000000
000000000000000000000000000000000000000000000000000000000000
000000000000000000000000000000000000000000000000000000000000
0000

>DM_train1116
GPLGSPASVPLRTEEEFKKFISDKDASIVGFFDDSFSEAHSEFLKAASNLRDNYRFAHTNVE
SLVNEYDDNGEGIILFRPSHLTNKFEDKTVAYTEQKMTSGKIKKFIQENIFGICPHMTEDNK
DLIQGKDLLIAYYDVDYEKNAKGSNYWRNRVMMVAKKFLDAGHKLNFAVASRKTFSHELSDF
GLESTAGEIPVVAIRTAKGEKFVMQEEFSRDGKALERFLQDYFDGNLKRYLKSEPIPESNDG
AAAS
11111100000000000000000000000000000000000000000000000000000000
00000000000000000000000000000000000000000000000000000000000000
00000000000000000000000000000000000000000000000000000000000000
00000000000000000000000000000000000000000000000000111111111111
1111
>DM_train1117
MAETAVINHKKRKNSPRIVQSNDLTEAAYSLSRDQKRMLYLFVDQIRKSDGTLQEHDGICEI
HVAKYAEIFGLTSAEASKDIRQALKSFAGKEVVFYRPEEDAGDEKGYESFPWFIKPAHSPSR
GLYSVHINPYLIPFFIGLQNRFTQFRLSETKEITNPYAMRLYESLCQYRKPDGSGIVSLKID
WIIERYQLPQSYQRMPDFRRRFLQVCVNEINSRTPMRLSYIEKKKGRQTTHIVFSFRDITSM
TTG
11111111111111000000000000000000000000000000000001111110000000
00000000000000000000000000000000000111111111111000000000000000
00000000000000000000000000000000000000000000000000000000000000
00000000000000000000000000000000000000000000000000000000000011
111
>DM_train1118
MGSLVREWVGFQQFPAATQEKLIEFFGKLKQKDMNSMTVLVLGKGGVGKSSTVNSLIGEQVV
RVSPFQAEGLRPVMVSRTMGGFTINIIDTPGLVEAGYVNHQALELIKGFLVNRTIDVLLYVD
RLDVYAVDELDKQVVIAITQTFGKEIWCKTLLVLTHAQFSPPDELSYETFSSKRSDSLLKTI
RAGSKMRKQEFEDSAIAVVYAENSGRCSKNDKDEKALPNGEAWIPNLVKAITDVATNQRKAI
HVD
11111000000000000000000000000000000000000000000000000000000000
00000001000000000000000000000000000000000000000000000000000000
00000000000000000000000000000000000000000000000000000000000000
00000000000000000000000000000000000000000000000000000000000000
001
>DM_train1119
MNINDALTSILASKKYRALCPDTVRRILTEEWGRHKSPKQTVEAARTRLHGICGAYVTPESL
KAAAAALSAGDVKKALSLHASTKERLAELDTLYDFIFSAETPRRVLDIACGLNPLALYERGI
ASVWGCDIHQGLGDVITPFAREKDWDFTFALQDVLCAPPAEAGDLALIFKLLPLLEREQAGS
AMALLQSLNTPRMAVSFPTRSLGGRGKGMEANYAAWFEGGLPAEFEIEDKKTIGTELIYLIK
KNG
00000000000000000000000000000000000000000000000000000000000000
00000000000000000000000000000000000000000000000000000000000000
00000000000000000000000000000000000000000000000000000000000000
00000000000000000000111111111110000000000000000000000000000000
001

>DM_train1121
GHMNMSQSQEIVLQPRSIVVPGELLAEGEFQIPWSPYILKINSKYYSTVVGLFDVKDTQFEV
IPLEGSFYYPKINDIVIGLVEDVEIYGWVVDIKAPYKAYLPASNLLGRSINVGEDLRRYLDV
GDYVIARIENFDRSIDPVLSVKGKDLGRVSNGIVIDIMPVKVPRVIGKNKSMYETLTSKSGC
SIFVANNGRIWATCPSRFSEEILIEAIRKIENESHIKGLTDRIKQFIEEKLGERNASSGETK
TNS
111111100000000000000000000000000000000000000000000000000000
000000000000000000000000000000000000000000000000000000000000
000000000000000000000000000000000000000000000000000000000011
100000000000111110000000000000000000111111111111111111111111
111
>DM_train1122
GMTPIAQRDGQAIQLVGFDGDDTLWKSEDYYRTAEADFEAILSGYLDLGDSRMQQHLLAVER
RNLKIFGYGAKGMTLSMIETAIELTEARIEARDIQRIVEIGRATLQHPVEVIAGVREAVAAI
AADYAVVLITKGDLFHQEQKIEQSGLSDLFPRIEVVSEKDPQTYARVLSEFDLPAERFVMIG
NSLRSDVEPVLAIGGWGIYTPYAVTWAHEQDHGVAADEPRLREVPDPSGWPAAVRALDAQAG
RQQ
111111111000000000000000000000000000000000000000000000000001
111110000000000000000000000000000000000000000000000000000000
000000000000000000000000000000000000000000000000000000000000
000000000000000000000011111100000000000000000000000000000000
001
>DM_train1123
MPTFLLVNDDGYFSPGINALREALKSLGRVVVVAPDRNLSGVGHSLTFTEPLKMRKIDTDFY
TVIDGTPADCVHLGYRVILEEKKPDLVLSGINEGPNLGEDITYSGTVSGAMEGRILGIPSIA
FSAFGRENIMFEEIAKVCVDIVKKVLNEGIPEDTYLNVNIPNLRYEEIKGIKVTRQGKRAYK
ERVFKYIDPYGKPFYWIAAEEFGWHAEEGTDYWAVLNGYVSVTPLHLDLTNYKVMKSIKYLE
DSP
100000000000000000000000000000000000000000000000000000000000
000000000000000000000000000000000000000000000000000000000000
000000000000000000000000000000000000000000000000000000000000
000000000000000000000000000000000000000000000000000000000000
011
>DM_train1124
SGAMYIDCDGIKLNAYLDMPKNNPEKCPLCIIIHGFTGHSEERHIVAVQETLNEIGVATLRA
DMYGHGKSDGKFEDHTLFKWLTNILAVVDYAKKLDFVTDIYMAGHSQGGLSVMLAAAMERDI
IKALIPLSPAAMIPEIARTGELLGLKFDPENIPDELDAWDGRKLKGNYVRVAQTIRVEDFVD
KYTKPVLIVHGDQDEAVPYEASVAFSKQYKNCKLVTIPGDTHCYDHHLELVTEAVKEFMLEQ
IAK
000000000000000000000000000000000000000000000000000000000000
000011111111100000000000000000000000000000000000000000000000
000000000000000000000000000000000000000000000000000000000000
000000000000000000000000000000000000000000000000000000000000
000

>DM_train1126
MTFGEQPAYLRVAGDLRKKIVDGSLPPHTRLPSQARIREEYGVSDTVALEARKVLMAEGLVE
GRSGSGTYVRERPVPRRVARSGYRPDSGATPFRQEQADGAVRGTWESHSEQAEASGAIAERL
DIRPGERVMCTKYVFRDAGEVMMLSTSWEPLAVTGRTPVMLPEEGPVGGMGVVERMAAIDVI
VDNVTEEVGARPGLAEELLTLGGVPGHVVLVIQRTYFASGRPVETADVVVPADRYRVAYHLP
VK
111111110000000000000000000000000000000000000000000000000000
111111000000000000000000110000000000000000000000000000000000
000000000000000000000000000000000000000000000000000000000000
000000000000000000000000000000000000000000000000000000000000
00
>DM_train1127
SLPMQHVHTSPVRDYRNRCARREGETVFQVVVEETDLRVTALAELATPMAAYVGELRAQLKV
WMEFQPAFRHSLVPVEVPEGAPEVVRRMAHGARLVGVGPFAAVAGTIAQMVAERFVDVSPEL
IVENGGDLYLYSERDRVVGILPDPASGDMVGILVRAGTAPVSLCGSSARIGHSLSLGDGDLA
VVRARDASLADAAATAFGNMLRRADDVAAVTERAAQLASIGIEGVYAQCGGRIGIWGDMELA
VA
111111111111111111100000000000000000000000000000000000000000
000000000000000000000000000000000000000000000000000000000000
000000000000000000000000000000000000000000000000000000000000
000000000000000000000000000000000000000000000000000000000000
00
>DM_train1128
GQLPVVGADGLRPMEQYARPWSGARGTRVAIVVGGLGLSQTGSQKAIRDLPPEVTLGFAASG
NSLQRWMQDARREGHEILLQIPLEPFGYPGTNPGPDTLLAGDPAKVNIDRLHRSMAKITNYT
GVMNYLGGRFLAEQSALEPVMRDIGKRGLLFLDDGSSAQSLSGGIAKAISAPQGFADVLLDG
EVTEASILRKLDDLERIARRNGQAIGVASAFDESIAAISKWSREAGGRGIEIVGVSALVSGQ
AG
000000000000000000000000000000000000000000000000000000000000
000000000000000000000000000000000000000000000000000000000000
000000000000000000000000000000000000000000000000000000000000
000000000000000000000000000000000000000000000000000000000111
11
>DM_train1129
MTDILIDDTATEAVRTLIRAFPLVPVSQPPEQGSYLLAEHDTVSLRLVGEKSNVIVDFTSGA
AQYRRTKGGGELIAKAVNHTAHPTVWDATAGLGRDSFVLASLGLTVTAFEQHPAVACLLSDG
IRRALLNPETQDTAARINLHFGNAAEQMPALVKTQGKPDIVYLDPMYPERRKSAAVKKEMAY
FHRLVGEAQDEVVLLHTARQTAKKRVVVKRPRLGEHLAGQAPAYQYTGKSTRFDVYLPYGAD
KG
100000000000000000000000000000000000000000000000000000000111
111111111100000000000000000000000000000000000000000000000000
000000000000000000000000000000000000000000001111111111111000
000000000000000000000000000000000000000000000000000000000000
00

>DM_train1130
GSPGILAPLAPGSEDNFARFVCKNNGVLFENQLLQIGLKSEFRQNLGRMFIFYGNKTSTQFL
NFTPTLICADDLQTNLNLQTKPVDPTVDGGAQVQQVINIECISDFTEAPVLNIQFRYGGTFQ
NVSVKLPITLNKFFQPTEMASQDFFQRWKQLSNPQQEVQNIFKAKHPMDTEITKAKIIGFGS
ALLEEVDPNPANFVGAGIIHTKTTQIGCLLRLEPNLQAQMYRLTLRTSKDTVSQRLCELLSE
QF
11110000000000000000000000000000000000000000000000000000000000
00000000000000000000000000000000000000000000000000000000000000
00000000000000000000000000000000000000000000000000000000000000
00000000000000000000000000000000000000000000000000000000000000
00
>DM_train1132
MTTNDTYEATYAIPMHCENCVNDIKACLKNVPGINSLNFDIEQQIMSVESSVAPSTIINTLR
NCGKDAIIRGAGKPNSSAVAILETFQKYTIDQKKDTAVRGLARIVQVGENKTLFDITVNGVP
EAGNYHASIHEKGDVSKGVESTGKVWHKFDEPIECFNESDLGKNLYSGKTFLSAPLPTWQLI
GRSFVISKSLNHPENEPSSVKDYSFLGVIARSAGVWENNKQVCACTGKTVWEERKDALANNI
K
00000000000000000000000000000000000000000000000000000000000000
00000000000000000000000000000000000000000000000000000000000000
00000000000000000000000000000000000000000000000000000000000000
00000000000000000000000000000000000000000000000000000000000111
1
>DM_train1133
MKSMSEYLNLLKEAIQNVVDGGWHETKRKGNTGIGKTFEDLLEKEEDNLDAPDFHDIEIKTH
ETAAKSLLTLFTKSPTNPRGANTMLRNRYGKKDEYGNNILHQTVSGNRKTNSNSYNYDFKID
IDWESQVVRLEVFDKQDIMIDNSVYWSFDSLQNQLDKKLKYIAVISAESKIENEKKYYKYNS
ANLFTDLTVQSLCRGIENGDIKVDIRIGAYHSGKKKGKTHDHGTAFRINMEKLLEYGEVKVI
V
11111100000000000000000000011111000000000000000000000000000000
00000000000000000000000000000000000000000000000000000000000000
00000000000000000000000000000000000000000000000000000000000000
00000000000000000000000000000111111111111100000000000000000000
0
>DM_train1134
SWEIEELDREIGKIKKHSLILIHEEDASSRGKDILFYILSRKLKSDNLVGMFSISYPLQLII
RILSRFGVDVIKYLENHRLAIVDTFGSFHGIKATMPGVWYLEGMLSSETLPIKYAKAVEDHK
KVWMDLNLFEGRELYGFAISMSGYLEVFTPEETLRYLETSAEVRYGHPAYKKYPRGTNFWLW
EGVKDKRVLLSVYRRADYVLKTRSSLGENGIKRELLVIKTPKPIEELVRFEYEFKGNEPKLR
R
00000000000000000000000000000000000000000000000000000000000000
00000000000000000000000000000001110000000000000000000000000000
00000000000000000000000000000000000000000000000000000000000000
00000000000000000000000000000000000000000011110000000000000000
0

>DM_train1135
MTAAEHGLHAPAYAWSHNGPFETFDHASIRRGYQVYREVCAACHSLDRVAWRTLVGVSHTNE
EVRNMAEEFEYDDEPDEQGNPKKRPGKLSDYIPGPYPNEQAARAANQGALPPDLSLIVKARH
GGCDYIFSLLTGYPDEPPAGVALPPGSNYNPYFPGGSIAMARVLFDDMVEYEDGTPATTSQM
AKDVTTFLNWCAEPEHDERKRLGLKTVIILSSLYLLSIWVKKFKWAGIKTRKFVFNPPKPRK
000000000000000000000000000000000000000000000000000000000000
000000000000000000000000000000000000000000000000000000000000
000000000000000000000000000000000000000000000000000000000000
000000000000000000000000000000000000000000000000000000000111
>DM_train1136
VRSFIYEPFQIPSGSMMPTLLIGDFILVEKFAYGIKDPIYQKTLIETGHPKRGDIVVFKYPE
DPKLDYIKRAVGLPGDKVTYDPVSKELTIQPGCSSGQACENALPVTYSNVEPSDFVQTFSRR
NGGEATSGFFEVPKNETKENGIRLSERKETLGDVTHRILTVPIAQDQVGMYYQQPGQQLATW
IVPPGQYFMMGDNRDNSADSRYWGFVPEANLVGRATAIWMSFDKQEGEWPTGLRLSRIGGIH
111000000000000000000000000011111111111111110000000000000000
000000000000000000000000000000011111000000000000000000000001
111000000000000000000000000000000000000000000000000000000000
000000000000000000000000000000000000000001111111110000000000
>DM_train1137
GSHMASMTGGQQMGRGSEFKKYLETNGIKPKQFHKKELIFNQWDPQEYCIFLYDGITKLTSI
SENGTIMNLQYYKGAFVIMSGFIDTETSVGYYNLEVISEQATAYVIKINELKELLSKNLTHF
FYVFQTLQKQVSYSLAKFNDFSINGKLGSICGQLLILTYVYGKETPDGIKITLDNLTMQELG
YSSGIAHSSAVSRIISKLKQEKVIVYKNSCFYVQNLDYLKRYAPKLDEWFYLACPATWGKLN
111111111111111000000000000000000000000000000000000000000000
000000000000000000000000000000000000000000000000000000000000
000000000000000000000000000000000000000000000000000000000001
111111110000000000000000000000000000000000000000000000000000
>DM_train1138
MSEAAQTLDGWYCLHDFRTIDWSAWKTLPNEEREAAISEFLALVDQWETTESEKQGSHAVYT
IVGQKADILFMILRPTLDELHEIETALNKTKLADYLLPAYSYVSVVELSNYLASGSEDPYQI
PEVRRRLYPILPKTNYICFYPMDKRRQGNDNWYMLSMEQRRELMRAHGMTGRKYAGKVTQII
TGSVGLDDFEWGVTLFSDDALQFKKLVYEMRFDEVSARFGEFGSFFVGTRLPMENVSSFFHV
111110000000000000000000000000000000000000000000000000000000
000000000000000000000000000000000000000000000000000000000000
000000000000000000000000000000000000000000000000000000000000
000000000000000000000000000000000000000000000000000000000000
>DM_train1139
MAVTDLSLTNSSLMPTLNPMIQQLALAIAASWQSLPLKPYQLPEDLGYVEGRLEGEKLVIEN
RCYQTPQFRKMHLELAKVGKGLDILHCVMFPEPLYGLPLFGCDIVAGPGGVSAAIADLSPTQ
SDRQLPAAYQKSLAELGQPEFEQQRELPPWGEIFSEYCLFIRPSNVTEEERFVQRVVDFLQI
HCHQSIVAEPLSEAQTLEHRQGQIHYCQQQQKNDKTRRVLEKAFGEAWAERYMSQVLFDVIQ
111110000000000000000000000000000000000000000000000000000000
000000000000000000000000000000000000000000000000000000000000
000000000000000000000000000000000000000000000000000000000000

000000000000000000000000000000000000000000000000000000000000
>DM_train1140
MSDDPIAASQAVVLSQAQAQNSQYALARTFATQKVSLEESVLSQVTTAIQTAQEKIVYAGNG
TLSDDDRASLATDLQGIRDQLMNLANSTDGNGRYIFAGYKTEAAPFDQATGGYHGGEKSVTQ
QVDSARTMVIGHTGAQIFNSITSNAVPEPDGSDSEKNLFVMLDTAIAALKTPVEGNNVEKEK
AAAAIDKTNRGLKNSLNNVLTVRAELGTQLSELSTLDSLGSDRALGQKLQMSNLVDVDWNSV
111111111110000000000000000000000000000000000000000000000000
000000000000000000000000000000000000000000000000000000000000
000000000000000000000000000000000000000000000000000000000000
000000000000000000000000000000000000000111111111111111111111
>DM_train1141
GMSEENKIGTYQFVAEPFHVDFNGRLTMGVLGNHLLNCAGFHASDRGFGIATLNEDNYTWVL
SRLAIELDEMPYQYEKFSVQTWVENVYRLFTDRNFAVIDKDGKKIGYARSVWAMINLNTRKP
ADLLALHGGSIVDYICDEPCPIEKPSRIKVTSNQPVATLTAKYSDIDINGHVNSIRYIEHIL
DLFPIELYQTKRIRRFEMAYVAESYFGDELSFFCDEVSENEFHVEVKKNGSEVVCRSKVIFE
100000000000000000000000000000000000000000000000000000000000
000000000000000000000000000000000000000000000000000000000000
011110000000000000000000000000000000000000000000000000000000
000000000000000000000000000000000000000000000000000000000000
>DM_train1142
MGSDKIHHHHHHMEDGMNTFDLYYWPVPFRGQLIRGILAHCGCSWDEHDVDAIEGLMDCGAE
KQPVAFMGPPVLIDRERNFAISQMPAIAIYLGERLDILPATVEGRTLSAKIVNDANDVLDEL
TLNGGREMWTPEKWQEFVPRLQKWIRIFADTGARNGLSAASGFMLGTEKIGVADIVTAILWT
TVADRFPAIKGIIEDTSPIIWGLSRRVVATAPLAALNSKSFEEYGNAYCGGEIEKSLRKVAS
111111111111110000000000000000000000000000000000000000000000
000000000000000000000000000000000000000000000000000000000000
000000000000000000000000000000000000000000000000000000000000
000000000000000000000000000000000000000000000000000000000000
>DM_train1143
GEAQPASGRFDQVGGAFGWKPHKLDPKECAQVAYDGYWYKGFGCGFGAFYSIVGLMGEKYGA
PYNQFPFAMLEANKGGISDWGTICGALYGAAATFSLFWGRKEVHPMVNELFRWYEVTKLPIF
NPGDAAQGVKGDLPMSASDSVLCHISVSKWCYENKIEATSKQRSERCGRLTADAAFKAAEII
NTKIDQGKDFKSTFPMQASVSSCGECHMTKGNDANWAKGIMDCTPCHSGTAATQNKFVNHP
111111100000000000000000000000000000000000000000000000000000
000000000000000000000000000000000000000000000000000000000000
000000000000000000000000000000000000000000000000000000000000
00000000000000000000000000000000000000000000000000000000000
>DM_train1144
MRSRRVDVMDVMNRLILAMDLMNRDDALRVTGEVREYIDTVKIGYPLVLSEGMDIIAEFRKR
FGCRIIAAFKVADIPETNEKICRATFKAGADAIIVHGFPGADSVRACLNVAEEMGREVFLLT
EMSHPGAEMFIQGAADEIARMGVDLGVKNYVGPSTRPERLSRLREIIGQDSFLISPGVGAQG
GDPGETLRFADAIIVGRSIYLADNPAAAAAGIIESIKDLLNPEDPAANKARKEAELAAATA
111111111111100000000000000000000000000000000000000000000000
000000000000000000000000000000000000000000000000000000000000

00000000000000000000000000000000000000000000000000000000000000
0000000000000000000000000000000000001111111111111111111111111
>DM_train1145
MFKAIVSAATLRDALDSVSVLVDECKIRLNEESLSIRAVDPANVGMVDLTLDAAAFESYEAH
GGVIGVNLSRLEEVAGMAGAGDLIHLTLDEETRKLNIRIDGLSYTLALIDPDSIRQEPDIPD
LDLAANIVLEGTHLDRGIKAADMVSDHIRLRVDGAEETFHIEAEGDTDDVDLSLPPADLISI
EAGAADSLFSLDYLKDMNKAIPTDAEVTVELGEEFPVKLHYQIAEGMGTITYMLAPRIQSD
10000000000000000000000000011100000000000000000000000000000111
11000000000000000011110000011111100000000000001110000111100000
00000000000000000000000000000000000000000000000000000000000000
0000000000000000000000000000000000000000000000000000000001111
>DM_train1146
MAHHHHHHLPALKLALEYIVPCMNKHGICVVDDFLGKETGQQIGDEVRALHDTGKFTDGQLV
SQKSDSSKDIRGDKITWIEGKEPGCETIGLLMSSMDDLIRHCNGKLGSYKINGRTKAMVACY
PGNGTGYVRHVDNPNGDGRCVTCIYYLNKDWDAKVSGGILRIFPEGKAQFADIEPKFDRLLF
FWSDRRNPHEVQPAYATRYAITVWYFDADERARAKVKYLTGEKGVRVELNKPSDSVGKDVF
11111111000000000000000000000000000000000000000000000000000000
00000000000000000000000000000000000000000000000000000000000000
00000000000000000000000000000000000000000000000000000000000000
0000000000000000000000000000000000000001110000000111111111111
>DM_train1147
SNATTHKFEHPLNEKTRIYLRVESLLRQAHLASGFADNHQYQLFFRALFDMVEIFEQIQLKS
ELAKDLEKQRLSYRHWLNVEGVDQEALNSLLNEIDVVHSQLMGAERFGQALKEDRFLSSIRQ
RFNLPGGSCCFDLPALHYWLHLPIERKKHDANQWQKSLKPLSDALTLWLKLARETGHFKAQI
ARAGFFQSDADEANILRLHIPMKYGVYPMISGHKNRFAIKFMAFENGQACSQDVEFELAVC
11000000000000000000000000000000000000000000000000000000000000
00000000000000000000000000000000000000000000000000000000000000
00001111000000000000000000000000000000000000000000000000000000
0000000000000000000000000000000000000000000000000000000000000
>DM_train1148
GAMGMIDIHSHIVFDVDDGPKSREESKALLAESYRQGVRTIVSTSHRRKGMFETPEEKIAEN
FLQVREIAKEVASDLVIAYGAEIYYTPDVLDKLEKKRIPTLNDSRYALIEFSMNTPYRDIHS
ALSKILMLGITPVIAHIERYDALENNEKRVRELIDMGCYTQVNSSHVLKPKLFGERYKFMKK
RAQYFLEQDLVHVIASDMHNLDGRPPHMAEAYDLVTQKYGEAKAQELFIDNPRKIVMDQLI
11100000000000000000000000000000000000000000000000000000000000
00000000000000000000000000000000000000000000000000000000000000
00000000000000000000000000000000000000000000000000000000000000
0000000000000000000000000000000000000000000000000000000000000
>DM_train1149
MISDSMTVEEIRLHLGLALKEKDFVVDKTGVKTIEIIGASFVADEPFIFGALNDEYIQRELE
WYKSKSLFVKDIPGETPKIWQQVASSKGEINSNYGWAIWSEDNYAQYDMCLAELGQNPDSRR
GIMIYTRPSMQFDYNKDGMSDFMCTNTVQYLIRDKKINAVVNMRSNDVVFGFRNDYAWQKYV
LDKLVSDLNAGDSTRQYKAGSIIWNVGSLHVYSRHFYLVDHWWKTGETHISKKDYVGKYA
00000000000000000000000000000000000000000000000000000000000000

000000000000000000000000000000000000000000000000000000000000
000000000000000000000000000000000000000000000000000000000000
0000000000000000000000000000000000000000000000000000011111
>DM_train1151
MEATIYDLDGNTDGEVDLPDVFETPVRSDLIGKAVRAAQANRKQDYGSDEYAGLRTPAESFG
SGRGQAHVPKLDGRARRVPQAVKGRSAHPPKTEKDRSLDLNDKERQLAVRSALAATADADLV
ADRGHEFDRDEVPVVVSDDFEDLVKTQEVVSLLEALDVHADIDRADETKIKAGQGSARGRKY
RRPASILFVTSDEPSTAARNLAGADVATASEVNTEDLAPGGAPGRLTVFTESALAEVAER
000000011100000000000000000000000000000000000000000000000000
000000011111111100000000000000000000000000000000000000001111
111111111111100000000000000000000000000000111111111111111111
1111110000000000000000000000000000000111100000000000000000
>DM_train1153
EQLLPDLLISPHMLPLTDLEIKFQYRGRPPRALTISNPHGCRLFYSQLEATQEQVELFGPIS
LEQVRFPSPEDIPSDKQRFYTNQLLDVLDRGLILQLQGQDLYAIRLCQCKVFWSGPCASAHD
SCPNPIQREVKTKLFSLEHFLNELILFQKGQTNTPPPFEIFFCFGEEWPDRKPREKKLITVQ
VVPVAARLLLEMFSGELSWSADDIRLQISNPDLKDRMVEQFKELHHIWQSQQRLQPVAQA
000000000000000000000000000000000000000000000000000000000000
000000000000000000000000000000000000000000000000000000000000
000000000000000000000000000000000000000000000000000000000000
0000000000000000000000000000000000000000000000001111111111
>DM_train1154
MKLARFLAKGRVHQGVYREGLLLDEAGEAHRPEDVTWLLPFTPGKILGVALNYADHAEELGL
SRPEEPALFWKPNTSLLPHKGVVLYPKGARFVHYEVELAVVVGRPMKRVRAKDALDYVLGYT
IANDLVARDYVTNTFRPPIRAKGRDTFLPLGPFLVVEEVEDPQDLWLRAYVNGELRQEGHTS
RMLYSVAELLEFISEFMTLEPYDVLLTGTPKGISQVRPGDVMRLEIEGLGALENPIEEEP
000000000000000000000000000000000000000000000000000001111100
000000000000000000000000000000000000000000000000000000000000
000000000001111000000000000000000000000000000000000000000000
0000000000000000000000000000000000000000000000000000000000
>DM_train1155
MGSDKIHHHHHHMVRLSEAEVQNPKLGEYQCLADLNLFDSPECTRLATQSASGRHLWVTSNH
QNLAVEVYLCEDDYPGWLSLSDFDSLQPATVPYQAATFSESEIKKLLAEVIAFTQKAMQQSN
YYLWGGTVGPNYDCSGLMQAAFASVGIWLPRDAYQQEGFTQPITIAELVAGDLVFFGTSQKA
THVGLYLADGYYIHSSGKDQGRDGIGIDILSEQGDAVSLSYYQQLRGAGRVFKSYEPQRR
111111111111111111111111000000000000000000000000000000000000
000000000000000000000000000000000000000000000000000000000000
000000000000000000000000000000000000000000000000000000000000
0000000000000000000000000000000000000000000000000000000000
>DM_train1156
VGPEAYGEASHSHSPASGRYIQQMLDQRCQEIAAELCQSGLRKMCVPSSRIVARNAVGITHQ
NTLQWRCFDTASLLESNQENNGVNCVDDCGHTIPCPGGVHRQNSNHATRHEILSKLVEEGVQ
RFCSPYQASANKYCNDKFPGTIARRSKGFGNNVEVAWRCYEKASLLYSVYAECASNCGTTWY
CPGGRRGTSTELDKRHYTEEEGIRQAIGSVDSPCSEVEVCLPKDENPPLCLDESGQISRT

11111111111000000000000000000000000000000000000000000000000000
00000000000000000000000000000000000000000000000000000000000000
00000000000000000000000000000000000000000000000000000000000000
000000000000000000000000000000000000000000000000000000000000
>DM_train1157
KHYSIQPANLEFNAEGTPVSRDFDDVYFSNDNGLEETRYVFLGGNQLEARFPEHPHPLFVVA
ESGFGTGLNFLTLWQAFDQFREAHPQAQLQRLHFISFEKFPLTRADLALAHQHWPELAPWAE
QLQAQWPMPLPGCHRLLLDEGRVTLDLWFGDINELISQLDDSLNQKVDAWFLDGFAPAKNPD
MWTQNLFNAMARLARPGGTLATFTSAGFVRRGLQEAGFTMQKRKGFGRKREMLCGVMEQT
00000000000000000000000000000000000000000000000000000000000000
00000000000000000000000000000000000000000000000000000000000000
00000000000000000001100000000000000000000000000000000000000000
000000000000000000000000000000000000000000000000000000000011
>DM_train1158
MYPVDLHMHTVASTHAYSTLSDYIAQAKQKGIKLFAITDHGPDMEDAPHHWHFINMRIWPRV
VDGVGILRGIEANIKNVDGEIDCSGKMFDSLDLIIAGFHEPVFAPHDKATNTQAMIATIASG
NVHIISHPGNPKYEIDVKAVAEAAAKHQVALEINNSSFLHSRKGSEDNCREVAAAVRDAGGW
VALGSDSHTAFTMGEFEECLKILDAVDFPPERILNVSPRRLLNFLESRGMAPIAEFADL
00000000000000000000000000000000000000000000000000000000000000
00000000000000000000000000000000000000000000000000000000000000
00000000000000000000000000000000000001111111111000000000000000
00000000000000000000000000000000000000000000000000000000000
>DM_train1159
MDKNELVQKAKLAEQAERYDDMAACMKSVTEQGAELSNEERNLLSVAYKNVVGARRSSWRVV
SSIEQKTEGAEKKQQMAREYREKIETELRDICNDVLSLLEKFLIPNASQAESKVFYLKMKGD
YYRYLAEVAAGDDKKGIVDQSQQAYQEAFEISKKEMQPTHPIRLGLALNFSVFYYEILNSPE
KACSLAKTAFDEAIAELDTLSEESYKDSTLIMQLLRDNLTLWTSDTQGDEAEAGEGGEN
00000000000000000000000000000000000000000000000000000000000000
00000011110000000000000000000000000000000000000000000000000000
00000000000000000000000000000000000011111000000000000001111111
10000000000000000111111111111000000000000011111111111111111
>DM_train1160
EMGTADLGPSSVPTPTNVTIESYNMNPIVYWEYQIMPQVPVFTVEVKNYGVKNSEWIDACIN
ISHHYCNISDHVGDPSNSLWVRVKARVGQKESAYAKSEEFAVCRDGKIGPPKLDIRKEEKQI
MIDIFHPSVFVNGDEQEVDYDPETTCYIRVYNVYVRMNGSEIQYKILTQKEDDCDEIQCQLA
IPVSSLNSQYCVSAEGVLHVWGVTTEKSKEVCITIFNSSIKGSLWIPVVAALLLFLVLS
11111111111000000000000000000000000000000000000000000000000000
00000000000000000000000000000000011000000000000000000000000000
00000000111111111111100000000000000000000000000000000000000000
00000000000000000000000000000000000111111111111111111111111
>DM_train1162
TESDMRATLVWSDEFDGPAGSAPDPANWNHETGDHGWGNNELQNYTDSRANSALDGNGNLVI
TARQEADGGYTSARLTTQNKVQPQYGRVEASIQIPRGQGIWPAFWMLGADFPNTPWPDSGEI
DIMENIGREPHLVHGSLHGPGYFGGEPLTGSYMHPQGWSFADTFHTFAVDWRPGSITWSVDG

VAYQTYTSADTRGNPWVFDQPFFMILNVAVGGDWPGYPDGSTQFPQEMRVDYVRVYELG
11111100000000000000000000000000000000000000000000000000000000
00000000000000000000000000000000000000000000000000000000000000
00000000000000000000000000000000000000000000000000000000000000
00000000000000000000000000000000000000000000000000000000011
>DM_train1163
AVKRRPRFPVNSNSNGGNELCPKIRIGQDDLPGFDLISQFQVDKAASRRAIQRVVGSATLQV
AYKLGNNVDFRIPTRNLYPSGLPEEYSFLTTFRMTGSTLKKNWNIWQIQDSSGKEQVGIKIN
GQTQSVVFSYKGLDGSLQTAAFSNLSSLFDSQWHKIMIGVERSSATLFVDCNRIESLPIKPR
GPIDIDGFAVLGKLADNPQVSVPFELQWMLIHCDPLRPRRETCHELPARITPSQTTDER
11111111111111111100000011111000000000000000000000000000000000
00000000000000000000000000000000000000000000000000000000000000
00000000000000000000000000000000000000000000000000000000000000
00000000000000000000000000000000000000000000000111111111111
>DM_train1164
MPDYSYRPTIGRTYVYDNKYYKNLDAVIKNAKRKKHFAEHEIEEATLDPLDNYLVAEDPFLG
PGKNQKLTLFKEIRNVKPDTMKLVVGWKGKEFYRETWTRFMEDSFPIVNDQEVMDVFLVVNM
RPTRPNRCYKFLAQHALRCDPDYVPHDVIRIVEPSWVGSNNEYRISLAKKGGGCPIMNLHSE
YTNSFEQFIDRVIWENFYKPIVYIGTDSAEEEEILLEVSLVFKVKEFAPDAPLFTGPAY
11000000000000000000000000000001111111111111111100000000000000
00000000000000000000000000000000000000000000000000000000000000
00000000000000000000000000000000000000000000000001111111111111
00000000000000000000000000000000000000000000000000000000000
>DM_train1165
MADDRGDITPIGDCRVCSFRMSLLLTGRCTPGDACVAVESGRQIDRFFRNNPHLAVQYLADP
FWERRAIAVRYSPVEALTPLIRDSDEVVRRAVAYRLPREQLSALMFDEDREVRITVADRLPL
EQLEQMAADRDYLVRAYVVQRIPPGRLFRFMRDEDRQVRKLVAKRLPEESLGLMTQDPEPEV
RRIVASRLRGDDLLELLHDPDWTVRLAAVEHASLEALRELDEPDPEVRLAIAGRLGIA
11111111000000000000000000000000000000000000000000000000000000
00000000000000000000000000000000000000000000000000000000000000
00000000000000000000000000000000000000000000000000000000000000
0000000000000000000000000000000000000000000000000000000111
>DM_train1166
SLRSDLINALYDENQKYDVCGIISAEGKIYPLGSDTKVLSTIFELFSRPIINKIAEKHGYIV
EEPKQQNHYPDFTLYKPSEPNKKIAIDIKTTYTNKENEKIKFTLGGYTSFIRNNTKNIVYPF
DQYIAHWIIGYVYTRVATRKSSLKTYNINELNEIPKPYKGVKVFLQDKWVIAGDLAGSGNTT
NIGSIHAHYKDFVEGKGIFDSEDEFLDYWRNYERTSQLRNDKYNNISEYRNWIYRGRK
00000000111111110000000000000000111111100000000000000000000000
00000000000000000000000000000000000000000000000000000000000000
00000000000000001111111000000000000000000000000000000000011111
0000000000000000000000000000001111111111111111111111111111
>DM_train1167
MDLTNGGVSPAATSAPLDWTTFRRVFLIDDAWRPLMEPELANPLTAHLLAEYNRRCQTEEVL
PPREDVFSWTRYCTPDEVRVVIIGQDPYHHPGQAHGLAFSVRANVPPPPSLRNVLAAVKNCY

PEARMSGHGCLEKWARDGVLLLNTTLTVKRGAAASHSRIGWDRFVGGVIRRLAARRPGLVFM
LWGTHAQNAIRPDPRVHCVLKFSHPSPLSKVPFGTCQHFLVANRYLETRSISPIDWSV
11111111111111111000000000000000000000000000000000000000000000
00000000000000000000000000000000000000000000000000000000000000
00000000000000000000000000000000000000000000000000000000000000
0000000000000000000000000000000000000000000000000000000000
>DM_train1168
TNVISTLDLNLLTKGGGSWNVDGVNMKKSAVTTFDGKRVVKAVYDKNSGTSANPGVGGFSFS
AVPDGLNKNAITFAWEVFYPKGFDFARGGKHGGTFIGHGAASGYQHSKTGASNRIMWQEKGG
VIDYIYPPSDLKQKIPGLDPEGHGIGFFQDDFKNALKYDVWNRIEIGTKMNTFKNGIPQLDG
ESYVIVNGKKEVLKRINWSRSPDLLISRFDWNTFFGGPLPSPKNQVAYFTNFQMKKYE
10000000000000000000000000000000000000000000000000000000000000
00000000000000000000000000000000000000000000000000000000000000
00000000000000000000000000000000000000000000000000000000000000
0000000000000000000000000000000000000000000000000000000001
>DM_train1169
QRSYERDFPHHDRICIVKTHGTRQSQEGDKPEELDFSQVSGGVAPAIQEEIPGVELATRTTL
YGTSKMILEDNKTYETKTLLAEPAFLDMFGVELIAGVRDSALRDNMTCLISESLARKMGGDV
LGKRLRPAESKSDRAITIGGVFEDLPHNSSIQADMLLPITWMPAESLNNWIGNDRYIAYVRL
RPGVSPESLDEALLEMQKRHQDMEVFRKAGVELHYSLTPFNRLRLEDPTLVNMLRIQQ
11111100000000000000011111111111100000000000000000000000000000
00000000000000000000000000000000000000000000000000000000000000
00000000000000000000000000000000000000000000000000000000000000
0000000000000000000000011111100000000000000111000000000000
>DM_train1170
MRSRTANRSGIVIRRRVTPAGDIIVTLLTPQGKLKAIARGGVKGPLSSSLNLFHHVGVQVYQ
GPHNDLASVKQAVLEGALPTLAEPERYAFAHLMAEFADALFQEGEFSEQAFDLFAASLRGVA
HQPDPEWVALVMSYKLLGLAGVIPQTARCARCGAPDPEHPDPLGGQLLCSKCAALPPYPPAV
LDFLRHAVRRTVRASFEQPVPSADRPALWRALEKFVTVQVGGVHSWRQLVPSGVPVLS
00000000000000000000000000000000000000001111000000000000000000
00000000000000000000000000000000000000000010000000000000000000
00000000000000000000000000000000000000000000000000000000000000
0000000000000000000000000000000000000000000000000001111111
>DM_train1171
MKIDLNADLGEGCASDAELLTLVSSANIACGFHAGDAQIMQACVREAIKNGVAIGAHPSFPD
RENFGRSAMQLPPETVYAQTLYQIGALATIARAQGGVMRHVKPHGMLYNQAAKEAQLADAIA
RAVYACDPALILVGLAGSELIRAGKQYGLTTREEVFADRGYQADGSLVPRSQSGALIENEEQ
ALAQTLEMVQHGRVKSITGEWATVAAQTVCLHGDGEHALAFARRLRSAFAEKGIVVAA
00000000000000000000000000000000000000000000000000000000000001
11111100000000000000000000000000000000000000000000000000000000
00000000000000000000000000000000000000000000000000000011111000
0000000000000000000000000000000000010000000000000111100000
>DM_train1172
MARSQLSAAGEQHVIQLNQQGGKNLFCFPPISGFGIYFKDLALQLNHKAAVYGFHFIEEDSR

IEQYVSRITEIQPEGPYVLLGYSAGGNLAFEVVQAMEQKGLEVSDFIIVDAYKKDQSITADT
ENDDSAAYLPEAVRETVMQKKRCYQEYWAQLINEGRIKSNIHFIEAGIQTETSGAMVLQKWQ
DAAEEGYAEYTGYGAHKDMLEGEFAEKNANIILNILDKINSDQKVLPNKHGSHHHHHH
111111000000000000000000000000000000000000000000000000000000
000000000000000000000000000000000000000000000000000000000001
111111000000000000000000000000000000000000000000000000000000
0000000000000000000000000000000000000001111111111111111111
>DM_train1173
GMPTLEMPVAAVLDSTVGSSEALPDFTSDRYKDAYSRINAIVIEGEQEAHDNYIAIGTLLPD
HVEELKRLAKMEMRHKKGFTACGKNLGVEADMDFAREFFAPLRDNFQTALGQGKTPTCLLIQ
ALLIEAFAISAYHTYIPVSDPFARKITEGVVKDEYTHLNYGEAWLKANLESCREELLEANRE
NLPLIRRMLDQVAGDAAVLQMDKEDLIEDFLIAYQESLTEIGFNTREITRMAAAALVS
111111111111111111110000000000000000000000000000000000000000
000000000000000000000000000000000000000000000000000000000000
000000000000000000000000000000000000000000000000000000000000
0000000000000000000000000000000000000000000000000000000011
>DM_train1174
SMLPEETILDKLPKDFQERITSSKWKDRVEALEEFWDSVLSQTKKLKSTSQNYSNLLGIYGH
IIQKDANIQAVALAAQSVELICDKLKTPGFSKDYVSLVFTPLLDRTKEKKPSVIEAIRKALL
TICKYYDPLASSGRNEDMLKDILEHMKHKTPQIRMECTQLFNASMKEEKDGYSTLQRYLKDE
VVPIVIQIVNDTQPAIRTIGFESFAILIKIFGMNTFVKTLEHLDNLKRKKIEETVKTL
000000000000000000000000000000000000000000000000000000000000
000000000000000000000000000000000000000000000000000000000000
000000000000000000000000000000000000000000000000000000000000
0000000000000000000000000000000000000000000000000000000001
>DM_train1175
GMEFNHLTKQLNQLLAQDYVAFSITENPVVQMLSQASFAQIAYVMQQYSIFPKELVGFTELA
RRKALGAGWNGVAQELQENIDEEMGSTTGGISHYTLLADGLEEGLGVAVKNTMPSVATSKLL
RTVLSLFDRQVDYVLGATYAIEATSIPELTLIVKLVEWLHEGAIPKDLQYFFSKHLDEWEIE
HEAGLRTSVAAYIQPEEFGEFAAGFRAMIDAMQVWWQELAQEAISSEVVLSTAIAQHH
000000000000000000000000000000000000000000000000000000000000
000000000000000000000000000000000000000000000000000000000000
000000000000000000000000000000000000000000000000000000011111
1100000000000000000000000000000000000000000000000001111111
>DM_train1176
ATSMTDIKGFNMISIEQEKELGNKFAVEIEKQQQPVNDPEVQRYVDKVGKRLLSGARAVEFD
YVFKVVKDDSVNAFAIPGGRVYVHTGLLKAADNETELAGVLAHEINHAVARHGTRQMTQEYG
YSLVLSLVLGDNPNMLAQLAGQLFGKAGMMSYSREYENQADFLGVETMYKAGYNPNGLTSFF
QKLNAMDGGTQSNVARFFSTHPLTSERIQRVQAEIAKLPPQRYLTDETEFKKIKGRLK
111111100000000000000000000000000000000000000000000000000000
000000000000000000000000000000000000000000000000000000000000
000000000011100000000000000000000000000000000000000000000000
0000011111111111111000000000000000000000000000000000000000
>DM_train1178

FADRMVIKAQSPAGFAEEYIIESIWNNRFPPGTILPAERELSELIGVTRTTLREVLQRLARD
GWLTIQHGKPTKVNNFWETSGLNILETLARLDHESVPQLIDNLLSVRTNISTIFIRTAFRQH
PDKAQEVLATANEVADHADAFAELDYNIFRGLAFASGNPIYGLILNGMKGLYTRIGRHYFAN
PEARSLALGFYHKLSALCSEGAHDQVYETVRRYGHESGEIWHRMQKNLPGDLAIQGR
11111111000000000000000000000000000000000000000000000000000000
00000000000000000000000000000000000000000000000000000000000000
00000000000000000000000000000000000000000000000000000000000000
000000000000000000000000000000000000000000000111111111111
>DM_train1179
GMTSSFTDYCKFFNRILSEVQETQEQAIIKGAHLVSEAVMNGGRFYVFGSGHSHMIAEEIYN
RAGGLALVTAILPPELMLHERPNKSTYLERIEGLSKSYLKLHQVTNKDVIMIISNSGRNTVP
VEMAIESRNIGAKVIAMTSMKHSQKVTSRHKSGKKLYEYADVVLDNGAPVGDAGFQIANSEI
YSGATSDSIGCFLAQALIVETLHLLVQQGFEPPVFKSSNVDGADLYNDKIFNEYVKW
10000000000000000000000000000000000000000000000000000000000000
00000000000000000000000000000000000000000000000000000000000000
00000000000000000000000000000000000000000000000000000000000000
000000000000000000000000000000000000000000000000000000000
>DM_train1180
GHMHAPLVSKDLDYISTANHDQPPRHLGSRFSAEGEFLPEPGNTVVCHLVEGSQTESAIVST
RQRFLDMPEASQLAFTPVSSLHMTVFQGVIESRRALPYWPQTLPLDTPIDAVTDYYRDRLST
FPTLPAFNMRVTGLRPVGMVMKGATAEDDSIVALWRDTFADFFGYRHPDHDTYEFHITLSYI
VSWFEPECLPRWQAMLDEELEKLRVAAPVIQMRPPAFCEFKDMNHFKELVVFDKRGS
11111110000000000000000000000000000000000000000000000000000000
00000000000000000000000000000000000000000000000000000000000000
00000000000000000000000000000000000000000000000000000000000000
000000000000000000000000000000000000000000000000000001111
>DM_train1181
MARYYVLDLSEDFRRELRETLAEMVNPVEVHVFLSKSGCETCEDTLRLMKLFEEESPTRNGG
KLLKLNVYYRESDSDKFSEFKVERVPTVAFLGGEVRWTGIPAGEEIRALVEVIMRLSEDESG
LEDATKEALKSLKGRVHIETIITPSCPYCPYAVLLAHMFAYEAWKQGNPVILSEAVEAYENP
DIADKYGVMSVPSIAINGYLVFVGVPYEEDFLDYVKSAAEGRLTVKGPIRAGEAEEL
11000000000000000000000000000000000000000000000000000000000000
00000000000000000000000000000000000000000000000000000000000000
00000000000000000000000000000000000000000000000000000000000000
000000000000000000000000000000000000000000000111111111111
>DM_train1182
MIRLIFLDIDKTLIPGYEPDPAKPIIEELKDMGFEIIFNSSKTRAEQEYYRKELEVETPFIS
ENGSAIFIPKGYFPFDVKGKEVGNYIVIELGIRVEKIREELKKLENIYGLKYYGNSTKEEIE
KFTGMPPELVPLAMEREYSETIFEWSRDGWEEVLVEGGFKVTMGSRFYTVHGNSDKGKAAKI
LLDFYKRLGQIESYAVGDSYNDFPMFEVVDKVFIVGSLKHKKAQNVSSIIDVLEVIK
00000000000000000000000000000000000000000000000000000000000000
00000000000000111111000000000000000000000000000000000000000000
00000000000000000000000000000000000000000000000000000000000000
000000000000000000000000000000000000000000000000000000000

>DM_train1183
MSADHDHDHDHDHDHKPAPMVEEVSDFEILEMAVRELAIEKGLFSAEDHRVWKDYVHTLGPL
PAARLVAKAWLDPEYKKLCIEDGVEASKAVGVNWVTSPPTQFGTPSDYCNLRVLADSPTLKH
VVVCTLCSCYPRPILGQSPEWYRSPNYRRRLVRWPRQVLAEFGLQLPSEVQIRVADSNQKTR
YIVMPVRPEGTDGWTEDQLAEIVTRDCLIGVAVPKPGITVNAKRPVLKANRPVHHDH
11111111111111111111111000000000000000000000000000000000000000
00000000000000000000000000000000000000000000000000000000000000
00000000000000000000000000000000000000000000000000000000000000
000000000000000000000000000000000000000000000000000001111
>DM_train1185
AELLLGVNIDHIATLRNARGTAYPDPVQAAFIAEQAGADGITVHLREDRRHITDRDVRILRQ
TLDTRMNLEMAVTEEMLAIAVETKPHFCCLVPEKRQEVTTEGGLDVAGQRDKMRDACKRLAD
AGIQVSLFIDADEEQIKAAAEVGAPFIEIHTGCYADAKTDAEQAQELARIAKAATFAASLGL
KVNAGHGLTYHNVKAIAAIPEMHELNIGHAIIGRAVMTGLKDAVAEMKRLMLEARG
00000000000000000000000000000000000000000000000000000000000000
00000000000000000000000000000000011111110000000000000000000000
00000000000000000000000000000000000000000000000000000000000000
00000000000000000000000000000000000000000000000000000000
>DM_train1186
MGSDKIHHHHHHMIVQRRNHRFLEVLSGKERVKIIAEFKKASPSAGDINADASLEDFIRMYD
ELADAISILTEKHYFKGDPAFVRAARNLTCRPILAKDFYIDTVQVKLASSVGADAILIIARI
LTAEQIKEIYEAAEELGMDSLVEVHSREDLEKVFSVIRPKIIGINTRDLDTFEIKKNVLWEL
LPLVPDDTVVVAESGIKDPRELKDLRGKVNAVLVGTSIMKAENPRRFLEEMRAWSE
11111100011100000000000000000000000000000000000000000000000000
00000000000000000000000000000000000000000000000000000000000000
00000000000000000000000000000000000000000000000000000000000000
00000000000000000000000000000000000000000000000000000000
>DM_train1187
MGVEKVPKYDIPVKKVEYVFIELDKMKPHEQLVQRELEDFIESVTGSGIFWKPMLLAKIPGT
DEYLIVDGHHRWAGLQKLGAKRAPSVILDYFDEGVKVYTWYPAFKGDVNKVIERLKAEGLEV
IEDEKAEEKAEKGEIAFALIGEKSFAIPGGLEEQKKVSKVLDEMDQAKEIELVYYGLKEDAK
ADMEKGEIDYVFIRKAPTKEEVMELVKRGEVFSPKTTRHVLPFIPDKIDVKLEDLF
11111111110000000000000000000000000000000000000000000000000000
00000000000000000000000000000000000000000000000000000000000000
00000000000000000000000000000000000000000000000000000000000000
00000000000000000000000000000000000000000000000000000000
>DM_train1188
GSYFSKQIIWQEVDRTKQQTSALIHNIFDSHFAAIQIHHDSNSKSEVIRDFYTDRDTDVLNF
FFLSIDQSDPSHTPEFRFLTDHKGIIWDDGNAHFYGVNDLILDSLANRVSFSNNWYYINVMT
SIGSRHMLVRRVPILDPSTGEVLGFSFNAVVLDNNFALMEKLKSESNVDNVVLVANSVPLAN
SLIGDEPYNVADVLQRKSSDKRLDKLLVIETPIVVNAVTTELCLLTVQDNQSVVTL
11111111111111000000000000000000000000000000000000000000000000
00000000000000000000000000000000000000000000000000000000000000
00000000000000000000000000000000000000000000000000000000000000

00000000000000111111111111100000000000000000000001111111
>DM_train1189
MAETEALSKLREDFRMQNKSVFILGASGETGRVLLKEILEQGLFSKVTLIGRRKLTFDEEAY
KNVNQEVVDFEKLDDYASAFQGHDVGFCCLGTTRGKAGAEGFVRVDRDYVLKSAELAKAGGC
KHFNLLSSKGADKSSNFLYLQVKGEVEAKVEELKFDRYSVFRPGVLLCDRQESRPGEWLVRK
FFGSLPDSWASGHSVPVVTVVRAMLNNVVRPRDKQMELLENKAIHDLGKAHGSLKP
111100000000000000000000000000000000000000000000000000000000
000000000000000000000000000000000000000000000000000000000000
000000000000000000000000000000000000000000000000000000000000
00000000000000000000000000000000000000000000000000111111
>DM_train1190
AIPKVPPGPNITATYGDKWLDAKSTWYGKPTGAGPKDNGGACGYKDVDKPPFSGMTGCGNTP
IFKSGRGCGSCFEIKCTKPEACSGEPVVVHITDDNEEPIAPYHFDLSGHAFGAMAKKGDEQK
LRSAGELELQFRRVKCKYPEGTKVTFHVEKGSNPNYLALLVKYVNGDGDVVAVDIKEKGKDK
WIELKESWGAIWRIDTPDKLTGPFTVRYTTEGGTKTEAEDVIPEGWKADTSYESK
000000000000000000000000000111111111100000000000000000000000
000000000000000000000000000000000000000000000000000000000000
000000000000000000000000000000000000000000000000000000000000
0000000000000000000000000000000000000000000000000000000
>DM_train1191
MKLIFLSGVKRSGKDTTADFIMSNYSAVKYQLAGPIKDALAYAWGVFAANTDYPCLTRKEFE
GIDYDRETNLNLTKLEVITIMEQAFCYLNGKSPIKGVFVFDDEGKESVNFVAFNKITDVINN
IEDQWSVRRLMQALGTDLIVNNFDRMYWVKLFALDYLDKFNSGYDYYIVPDTRQDHEMDAAR
AMGATVIHVVRPGQKSNDTHITEAGLPIRDGDLVITNDGSLEELFSKIKNTLKVL
000000000000000000000000000000000000000000000000000000000000
000000000000000000000000000000000000000000000000000000000000
000000000000000000000000000000000000000000000000000000000000
0000000000000111111111100000000000000000000000000000000
>DM_train1192
EEACELPECQEDAGNKVCSLQCNNHACGWDGGDCSLNFNDPWKNCTQSLQCWKYFSDGHCDS
QCNSAGCLFDGFDCQRAEGQCNPLYDQYCKDHFSDGHCDQGCNSAECEWDGLDCAEHVPERL
AAGTLVVVVLMPPEQLRNSSFHFLRELSRVLHTNVVFKRDAHGQQMIFPYYGMDVRGSIVYL
EIDNRQCVQASSQCFQSATDVAAFLGALASLGSLNIPYKIEAVQSETVEPPPPAQ
110000000000000000000000000000000000000000000000000000000000
000000000000000000000000000000000000000000000000000000000000
000000000000000000000000000000000000000000000000000000000000
0000000000000000000000000000000000000000000000001111111
>DM_train1193
GAGTQTAQTQALDSDGIPTGGEWITMFDGKTLNGWRGYCRQDVPLGWVVEDGSITYKGSDNK
ADTGFGDLIYDKKFKNFVFEIEWKIDKAGNSGIFYTAQEIEGTPIYYSSPEYQLLDNENMPD
AWEGCDGNRQAGAVYDMIMPDPQPVKPYGNWNKTRIVVYNQRVIHYMNDVKILEFQFGTPVW
RALVDHSKFSKFSTSPEKCPEAYDLMLQCGKQPGYIGMQDHGYGVCFRNIRIKEL
111111111100000000000000000000000000000000000000000000001111
111100000000000000000000000000000000000000000000000000000000

00000000000000000000000000000000000000000000000000000000000000
0000000000000000000000000000000000000000000000000000000
>DM_train1194
EELPAPVKAIEKQGITIIKTFDAPGGMKGYLGKYQDMGVTIYLTPDGKHAISGYMYNEKGEN
LSNTLIEKEIYAPAGREMWQRMEQSHWLLDGKKDAPVIVYVFADPFCPYCKQFWQQARPWVD
SGKVQLRTLLVGVIKPESPATAAAILASKDPAKTWQQYEASGGKLKLNVPANVSTEQMKVLS
DNEKLMDDLGANVTPAIYYMSKENTLQQAVGLPDQKTLNIIMGNKLQHHHHHHHH
10000000000000000000000000000000000000000000000000000000000000
00000000000000000000000000000000000000000000000000000000000000
00000000000000000000000000000000000000000000000000000000000000
0000000000000000000000000000000000000000000011111111111
>DM_train1195
MWDTSKDYRLLVAEKSVELFIRTIEGAKFRGQWDKKRSIQLAKEMIPDIQALRYSYIDPEEL
VDTPQMKDLKEKAKGIIEALGGEDWHHKFLSQASREDREKVEEQVARIKFFLNTILNLDRRL
KLGKINDPVIAVDIVVGEVMSVGKHPSADRLLVTNVNIGERAVTVVTNDLTVKEGNRVAVAL
LPPRNFFGIVSEGMFLGAGEGVLKNVKGEIGGLPKGIPLEALNETRNAVEAFLKG
00000000000000000000000000000000000000000000000000000000000000
00000000000000000000000000000000000000000000000000000000000000
00000000000000000000000000000000000000000000000000000000000000
0000000000000000000000000000000000000000000000000000001
>DM_train1196
MPINRPNLNLNIPPLNIVAAYDGAEIPSTNKHLKNNFNSLHNQMRKMPVSHFKEALDVPDYS
GMRQSGFFAMSQGFQLNNHGYDVFIHARRESPQSQGKFAGDKFHISVLRDMVPQAFQALSGL
LFSEDSPVDKWAVTDMEKVVQQARVSLGAQFTLYIKPDQENSQYSASFLHKTRQFIECLESR
LSENGVISGQCPESDVHPENWKYLSYRNELRSGRDGGEMQRQALREEPFYRLMTE
11111111111111111111111100000000000000000000000000000000000000
00000000000000000000000000000000000000000000000000000000000000
00000000000000000000000000000000000000000000000000000000000000
0000000000000000000000000000000000000000000000000000000
>DM_train1198
MGRAFEYRRAAKEKRWDKMSKVFPKLAKAITLAAKDGGSEPDTNAKLRTAILNAKAQNMPKD
NIDAAIKRASSKEGNLSEITYEGKANFGVLIIMECMTDNPTRTIANLKSYFNKTQGASIVPN
GSLEFMFNRKSVFECLKNEVENLKLSLEDLEFALIDYGLEELEEVEDKIIIRGDYNSFKLLN
EGFESLKLPILKASLQRIATTPIELNDEQMELTEKLLDRIEDDDDVVALYTNIE
11111111111111111111000000000000000000000000000000000000000000
00000000000000000000000000000000000000000000000000000000000000
00000000000000000000000000000000000000000000000000000000000000
000000000000000000000000000000000000000000000000000000
>DM_train1199
MQQASTPTIGMIVPPAAGLVPADGARLYPDLPFIASGLGLGSVTPEGYDAVIESVVDHARRL
QKQGAAVVSLMCTSLSFYRGAAFNAALTVAMREATGLPCTTMSTAVLNGLRALGVRRVALAT
AYIDDVNERLAAFLAEESLVPTGCRSLGITGVEAMARVDTATLVDLCVRAFEAAPDSDGILL
SSGGLLTLDAIPEVERRLGVPVVSSSPAGFWDAVRLAGGGAKARPGYGRLFDES
11111000000000000000000000000000000000000000000000000000000000

00000000000000000000000000000000000000000000000000000000000000
00000000000000000000000000000000000000000000000000000000000000
000000000000000000000000000000000000000000000000000000
>DM_train1200
MSLYDVTEWKHVFKLDPNKDLPDEQLEILCESGTDAVIIGGSDGVTEDNVLRMMSKVRRFLV
PCVLEVSAIEAIVPGFDLYFIPSVLNSKNADWIVGMHQKAMKEYGELMSMEEIVAEGYCIAN
PDCKAAALTEADADLNMDDIVAYARVSELLQLPIFYLEYSGVLGDIEAVKKTKAVLETSTLF
YGGGIKDAETAKQYAEHADVIVVGNAVYEDFDRALKTVAAVKGEEGGSHHHHHH
11000000000000000000000000000000000000000111111000000000000000
00000000000000000000000000000000000000000000000000000000000000
00000000000000000000000000000000000000000000000000000000000000
000000000000000000000000000000000000000000111111111111
>DM_train1201
GMKYKAAIFDMDGTILDTSADLTSALNYAFEQTGHRHDFTVEDIKNFFGSGVVVAVTRALAY
EAGSSRESLVAFGTKDEQIPEAVTQTEVNRVLEVFKPYYADHCQIKTGPFPGILDLMKNLRQ
KGVKLAVVSNKPNEAVQVLVEELFPGSFDFALGEKSGIRRKPAPDMTSECVKVLGVPRDKCV
YIGDSEIDIQTARNSEMDEIAVNWGFRSVPFLQKHGATVIVDTAEKLEEAILGE
10000000000000000000000000000000000000000000000000000000000000
00000000000000000000000000000000000000000000000000000000000000
00000000000000000000000000000000000000000000000000000000000000
000000000000000000000000000000000000000000000000000000
>DM_train1204
MRLVQLSRHSIAFPSPEGALREPNGLLALGGDLSPARLLMAYQRGIFPWFSPGDPILWWSPD
PRAVLWPESLHISRSMKRFHKRSPYRVTMNYAFGQVIEGCASDREEGTWITRGVVEAYHRLH
ELGHAHSIEVWREDELVGGMYGVAQGTLFCGESMFSRMENASKTALLVFCEEFIGHGGKLID
CQVLNDHTASLGACEIPRRDYLNYLNQMRLGRLPNNFWVPRCLFSPQEGLCGR
00000000000000000000000000000000000000000000000000000000000000
00000000000000000000000000000000000000000001110000000000000000
00000000000000000000000000000000000000000000000000000000000000
00000000000000000000000000000000000000000000001111111
>DM_train1205
MDIRPRLRIGQINLGGAEDATRELPSIARDLGLDIVLVQEQYSMVGFLAQCGAHPKAGVYIR
NRVLPCAVLHHLSSTHITVVHIGGWDLYMVSAYFQYSDPIDPYLHRLGNILDRLRGARVVIC
ADTNAHSPLWHSLPRHYVGRGQEVADRRAKMEDFIGARRLVVHNADGHLPTFSTANGESYVD
VTLSTRGVRVSEWRVTNESSSDHRLIVFGVGGGTTGERDEDEEARSDLRPGEP
11110000000000000000000000000000000000000000000000000000000000
00000000000000000000000000000000000000000000000000000000000000
00000000000000000000000000000000000000000000000000000000000000
00000000000000000000000000000000111111111111111111111
>DM_train1206
GLPVMNTPGSNQYLTADNFQSPCALPEFDVTPPIDIPGEVKNMMELAEIDTMIPFDLSATKK
NTMEMYRVRLSDKPHTDDPILCLSLSPASDPRLSHTMLGEILNYYTHWAGSLKFTFLFCGSM
MATGKLLVSYAPPGADPPKKRKEAMLGTHVIWDIGLQSSCTMVVPWISNTTYRQTIDDSFTE
GGYISVFYQTRIVVPLSTPREMDILGFVSACNDFSVRLLRDTTHIEQKALAQ

000000000000000000000000000000000000000000000000000000000000
000000000000000000000000000000000000000000000000000000000000
000000000000000000000000000000000000000000000000000000000000
00000000000000000000000000000000000000000000000111
>DM_train1207
GVNPRGLGPLQIWQTDFTLEPRMAPRSWLAVTVDTASSAIVVTQHGRVTSVAVQHHWATAIA
VLGRPKAIKTDNGSCFTSKSTREWLARWGIAHTTGIPGNSQGQAMVERANRLLKDRIRVLAE
GDGFMKRIPTSKQGELLAKAMYALNHKERGENTKTPIQKHWRPTVLTEGPPVKIRIETGEWE
KGWNVLVWGRGYAAVKNRDTDKVIWVPSRKVKPDITQKDEVTKKDEASPLFA
111111000000000000000000000000000000000000000000000000000000
000000000000000000000000000000000000000000000000000000000000
000000000000000000000000000011111000000000000000000000000000
00000000000000000000000000000000011111111111111111
>DM_train1209
MSLATTHLDVCAVVPAAGFGRRMQTECPKQYLSIGNQTILEHSVHALLAHPRVKRVVIAISP
GDSRFAQLPLANHPQITVVDGGDERADSVLAGLKAAGDAQWVLVHDAARPCLHQDDLARLLA
LSETSRTGGILAAPVRDTMKRAEPGKNAIAHTVDRNGLWHALTPQFFPRELLHDCLTRALNE
GATITDEASALEYCGFHPQLVEGRADNIKVTRPEDLALAEFYLTRTIHQENT
111111100000000011111111110000000000000000000000000000000001
110000000000000000000000000000000000000000000000000000000000
000000000000000000000000000000000000000000000000000000000000
00000000000000000000000000000110000000000111111111
>DM_train1210
MKEEAVRVIEEVLKQGRTAMVEYEAKQVLKAYGLPVPEEKLAKTLDEALEYAKEIGYPVVLK
LMSPQILHKSDAKVVMLNIKNEEELKKKWEEIHENAKKYRPDAEILGVLVAPMLKPGREVII
GVTEDPQFGHAIMFGLGGIFVEILKDVTFRLVPITEKDARKMIQEIKAYPILAGARGEEPAD
IDAIVDMLLKVSKLVDDLKDYIKEMDLNPVFVYNKGEGAVIVDSRIILKPKD
000000000000000000000000000000000000000000000000000000000000
000000000000000000000000000000000000000000000000000000000000
000000000000000000000000000000000000000000000000000001100000
00000000000000000000000000000000000000000000000001
>DM_train1211
MASMTGGQQMGRGSEFSDKMSSFLHIGDICSLYAEGSTNGFISTLGLVDDRCVVQPEAGDLN
NPPKKFRDCLFKLCPMNRYSAQKQFWKAAKPGANSTTDAVLLNKLHHAADLEKKQNETENRK
LLGTVIQYGNVIQLLHLKSNKYLTVNKRLPALLEKNAMRVTLDEAGNEGSWFYIQPFYKLRS
IGDSVVIGDKVVLNPVNAGQPLHASSHQLVDNPGCNEVNSVNCNTSWKIVLF
111111111111111111110000000000000000000000000000000000000000
000000000000000000000000000111110000000000000000000000000000
000000000000000000000000000000000000000000000000000000000000
00000000000000000000000000000000000000000000000000
>DM_train1212
PEFMVTTEPALADLQEQLYNGNEKSQLAAMSTLSTAGTEGYHLLQEFLKDSATFSPPPAPWI
RGQAYRLLFHSPEASVQAFLQQHYPQGVIPLRSDRGVDYQELAKLLVAEKFEAADRLTTQKL
CELAGPLAQKRRWLYFTEVEQLPIPDLQTIDQLWLAFSLGRFGYSVQRQLWLGCGQNWDRLW

EKIGWRQGKRWPRYPNEFIWDLSAPRGHLPLTNQLRGVQVLNALLNHPAWTA
00000011000000000000000000000000000000000000000000000000000000
00000000000000000000000000000000000000000000000000000000000000
00000000000000000000000000000000000000000000000000000000000000
0000000000000000000000000000000000000000000000000000
>DM_train1213
MKIWSKEEVVNKLHEIKNKGYLSVPTDMFRTDDGVVGQILERQFGVQENNITLGDLGEFELK
GMRNRKAKSNLTLFHKKPVAGQTVIQIFNRFGYVKPSSRNPEVMKKKLFTTIKGGRLNNLGL
TLNAKHASEINLYYQDEYLSTWDLNLSKIEKLVLVFAETIGRANSPEEQFHFTKAYMLTEIN
DITSLINDGVLVMDLCIDQDLSKSKGPHDRGPHLRIPISKLDKLYRNIERLL
00000000000000000000000000000000000000000000000000000000000000
00000000000000000000000000000000000011111000000000000000000000
00000000000000000000000000000000000000000000000000000000000000
0000000000000000000000000000000000000000000000000000
>DM_train1214
MVHGKLKEAAEVTGSVSLEALEEVQVGENIEVGVGIDELVNAEAFAYDFTLNYDENAFEYVE
AISDDGVFVNAKKIEDGKVRVLVSSLTGEPLPAKEVLAKVVLRAEAKTEGSNLSVTNSSVGD
GEGLVHEIAGTEKTVNIIEGTSPEIVVNPVRDFKASEINKKNVTVTWTEPETTEGLEGYILY
KDGKKVAEIGKDETSYTFKKLNRHTIYNFKIAAKYSNGEVSSKESLTLRTAR
11111111100000000000000000000000000000000000000000000000000000
00000000000000000000000000000000000000000000000000000000000000
00000000000000000000000000000000000000000000000000000000000000
0000000000000000000000000000000000000000000000000001
>DM_train1215
SNAMTNVLYQHGTLGTLMAGLLEGTATINELLEHGNLGIATLTGSDGEVIFLDGKAYHANEH
KEFIELKGDEKVPYASITNFKASKTFPLQQLSQDDVFAQIKNEMLSENLFSAVKIYGTFKHM
HVRMMPAQQPPYTRLIDSARRQPEEKRQDIRGAIVGFFTPELFHGVGSAGFHIHFADDERAY
GGHVLDFEVDDVVVEIQNFETFQQHFPVNNETFVKAKIDYKDVAEEIREAE
11111000000000000000000000000000000000000000000000000000000000
00000000000000000000000000000000000000000000000000000000000000
00000000000000000000000000000000000000000000000000000000000000
000000000000000000000000000000000000000000000000000
>DM_train1216
MRLRVDVIPGEHLAYPDVVLVVDVIRATTTAAAFLEAGAEALYWTPSLESALAFKDEDVVLA
GETGGLKPPRFDLGNSPREALSAQVAGRVVVMSTTNGTKAAHAAARTAKHVLLASLYNAHAA
ARLARELATEEVAILCAGKEGRAGLDDLYTAGVLAEYLGFLGEVEPEDGARVALAVKRAYPD
PLEALSLSAAALALKQVGLEADVPFCAQVAKSAAVPLRGRVGEALIFKRA
00000000000000000000000000000000000000000000000000000000000000
00000000000000000000000000000000000000000000000000000000000000
00000000000000000000000000000000000000000000000000000000000000
000000000000000000000000000000000000000001111000000
>DM_train1217
MGTKLKKSNNDITIFSENEYNEIVEMLRDYSNGDNLEFEVSFKNINYPNFMRITEHYINITP
ENKIESNNYLDISLIFPDKNVYRVSLFNQEQIGEFITKFSKASSNDISRYIVSLDPSDDIEI

VYKNRGSGKLIGIDNWAITIKSTEEIPLVAGKSKISKPKITGSERIMYRYKTRYSFTINKNS
RIDITDVKSSPIIWKLMTVPSNYELELELINKIDINTLESELLNVFMIIQD
111111111111000000000000000000000000000000000000000000000000
000000000000000000000000000000000000000000000000000000000000
000000000000000000000000000000100000000000000000000000000000
000000000000000000000000000000000000000000000000000
>DM_train1218
TYNISVVGLSGTEKEKGQCGIGKSCLCNRFVRPSADEFHLDHTSVLSTSDFGGRVVNNDHFL
YWGEVSRSLEDCVECKMHIVEQTEFIDDQTFQPHRSTALQPYIKRAAATKLASAEKLMYFCT
DQLGLEQDFEQKQMPDGKLLVDGFLLGIDVSRGMNRNFDDQLKFVSNLYNQLAKTKKPIVVV
LTKCDEGVERYIRDAHTFALSKKNLQVVETSARSNVNVDLAFSTLVQLIDK
000000000000000000000000000000000000000000000000000000000000
000000111111111000000000000000000000000000000000000000000000
000000000000000000000000000000011110000000000000000000000000
000000000000000000000000000000000000000000000000000
>DM_train1219
NNILFGLSHEGSHPQTLHAAQSLELSSFRFTMQSDCNLVLFDSDVRVWASNTAGATGCRAVL
QSDGLLVILTAQNTIRWSSGTKGSIGNYVLVLQPDRTVTIYGPGLWDSGTSNKGSVVVANNG
NSILYSTQGNDNHPQTLHATQSLQLSPYRLSMETDCNLVLFDRDDRVWSTNTAGKGTGCRAV
LQPNGRMDVLTNQNIAVWTSGNSRSAGRYVFVLQPDRNLAIYGGALWTTG
000000000000000000000000000000000000000000000000000000000000
000000000000000000000000000000000000000000000000111111111110
000000011110000000000000000000000000000000000000011100000000
00000000000000000000000000000000000000000000000000
>DM_train1220
MTDGEAVSAAAGPGAEGPPGPVGHWAPGSHILWRYRENGGPHVHIARPVTVVRDDADLLAVW
LAPGTECVKPVLADGTPVHLEPLATRYTKPRTVQRDQWFGTGVLKLARPGEAWSVWLFWDPG
WRFKNWYVNLERPLTRWEGGVDSEDHFLDISVHPDRTWHWRDEDEFAQALRDGLMDPASAGR
VRRAGRSAVAEIRAWGSPFADGWEHWRPDPAWPVPSLPGDWDRTPAHVSS
111111111111111111111000000000000000000000000000000000000000
000000000000000000000000000000000000000000000000000000000000
000000000000000000000000000000000000000000000000000000000000
00000000000000000000000000000000000000000000001111
>DM_train1221
EVKKKKNKKKNKKVNGLPPEIAAVPELAKYWAQRYRLFSRFDDGIKLDREGWFSVTPEKIAE
HIAGRVSQSFKCDVVVDAFCGVGGNTIQFALTGMRVIAIDIDPVKIALARNNAEVYGIADKI
EFICGDFLLLASFLKADVVFLSPPWGGPDYATAETFDIRTMMSPDGFEIFRLSKKITNNIVY
FLPRNADIDQVASLAGPGGQVEIEQNFLNNKLKTITAYFGDLIRRPASET
111111111111111111111100000000000000000000000000000000000000
000000000000000000000000000000000000000000000000000000000000
000000000000000000000000000000000000000000000000000000000000
00000000000000000000000000000000000000000000111111
>DM_train1222
MLTEGISIQSYDGHTFGALVGSPAKAPAPVIVIAQDIFGVNAFMRETVSWLVDQGYAAVCPD

LYARQAPGTALDPQDERQREQAYKLWQAFDMEAGVGDLEAAIRYARHQPYSNGKVGLVGYSL
GGALAFLVASKGYVDRAVGYYGVGLEKQLNKVPEVKHPALFHMGGQDHFVPAPSRQLITEGF
GANPLLQVHWYEEAGHSFARTGSSGYVASAAALANERTLDFLVPLQSRKP
000000000000000000000000000000000000000000000000000000000000
000000000000000000000000000000000000000000000000000000000000
000000000000000000000000000000000000000000000000000000000000
00000000000000000000000000000000000000000000000111
>DM_train1223
GSAPSATPIFAPGENCSPAWGAAPAAYDAADTHLRILGKPVMERWETPYMHALAAAASSKGG
RVLEVGFGMAIAASKVQEAPIDEHWIIECNDGVFQRLRDWAPRQTHKVIPLKGLWEDVAPTL
PDGHFDGILYDTYPLSEETWHTHQFNFIKNHAFRLLKPGGVLTYCNLTSWGELMKSKYSDIT
IMFEETQVPALLEAGFRRENIRTEVMALVPPADCRYYAFPQMITPLVTKG
111111111111111110000000000000000000000000000000000000000000
000000000000000000000000000000000000000000000000000000000000
000000000000000000000000000000000000000000000000000000000000
00000000000000000000000000000000000000000000000000
>DM_train1224
GMTTRKPFIICDFDGTITMNDNIINIMKTFAPPEWMALKDGVLSKTLSIKEGVGRMFGLLPS
SLKEEITSFVLEDAKIREGFREFVAFINEHEIPFYVISGGMDFFVYPLLEGIVEKDRIYCNH
ASFDNDYIHIDWPHSCKGTCSNQCGCCKPSVIHELSEPNQYIIMIGDSVTDVEAAKLSDLCF
ARDYLLNECREQNLNHLPYQDFYEIRKEIENVKEVQEWLQNKNAGESSLK
111000000000000000000000000000000000000000000000000000000000
000000000000000000000000000000000000000000000000000000000000
000000000000000000000000000000000000000000000000000000000000
00000000000000000000000000000000000000000111111111
>DM_train1225
MSPVLTTPKRHAPPPEQLQNVTDFNYTPSHQKPFLQPQAGTTVTTHQDIKQIVEMTLGSEGV
LNQAVKLPRGEDENEWLAVHCVDFYNQINMLYGSITEFCSPQTCPRMIATNEYEYLWAFQKG
QPPVSVSAPKYVECLMRWCQDQFDDESLFPSKVTGTFPEGFIQRVIQPILRRLFRVYAHIYC
HHFNEILELNLQTVLNTSFRHFCLFAQEFELLRPADFGPLLELVMELRDR
110000011111111111100000000000000111111111111000000000000000
111000000000000000000000000000000000000000000000000000000000
000000000000000000000000000000000000000000000000000000000000
00000000000000000000000000000000000000000000000001
>DM_train1226
MSEWKARRFWASVGIHKEEGGWAVLLDERPLRTPGKQPLRLPTEALALAIAEEWQAVQEVID
PNAMPLTRSANSAIEKVAPQFDAVAAMLGDYGGTDLLSYRADAPEALVRAQAEGWDPLIDWA
ATELRAPLRITHGVIPVPQDPVVLLKLRAEVASLDPFGLTALHDLVTLPGSLILGLAVIRGR
IDAPTAHALSRIDEEFQAERWGRDEEAEAQAASRLAAMRDSERFWHLTRG
000000000000000000000000000000000000000000000000000000000000
000000000000000000000000000000000000000000000000000000000000
000000000000000000000000000000000000000000000000000000000000
00000000000000000000000000000000000000000000000001
>DM_train1227

MHLYSSDFPLMMDEKELYEKWMRTVEMLKAEGIIRSKEVERAFLKYPRYLSVEDKYKKYAHI
DEPLPIPAGQTVSAPHMVAIMLEIANLKPGMNILEVGTGSGWNAALISEIVKTDVYTIERIP
ELVEFAKRNLERAGVKNVHVILGDGSKGFPPKAPYDVIIVTAGAPKIPEPLIEQLKIGGKLI
IPVGSYHLWQELLEVRKTKDGIKIKNHGGVAFVPLIGEYGWKEHHHHHH
11111111111110000000000000000000000000000000000000000000000000
00000000000000000000000000000000000000000000000000000000000000
00000000000000000000000000000000000000000000000000000000000000
0000000000000000000000000000000000000000001111111
>DM_train1228
GSPEFATMSVSEIFVELQGFLAAEQDIREEIRKVVQSLEQTAREILTLLQGVHQGTGFQDIP
KRCLKAREHFSTVKTHLTSLKTKFPAEQYYRFHEHWRFVLQRLVFLAAFVVYLETETLVTRE
AVTEILGIEPDREKGFHLDVEDYLSGVLILASELSRLSVNSVTAGDYSRPLHISTFINELDS
GFRLLNLKNDSLRKRYDGLKYDVKKVEEVVYDLSIRGFNKETAAACGEK
11111110000000000000000000000000000000000000001111111000
00000000000000000000000000000000000000000000000000000000000000
00000000000000000000000000000000000000000000000000000000000000
0000000000000000000000000000000000000011111111111
>DM_train1229
MNRLLNEKVEEFKKGVLKAGWFIEKMFRNSISSLVERNESLAREVIADEEVVDQMEVEIQEK
AMEVLGLFSPIGKPLLTVTAGIRVAELIENIADKCHDIAKNVLELMEEPPLKPLEDIPAMAN
QTSEMLKFALRMFADVNVEKSFEVCRMDSKVDDLYEKVREELLLYMMESPKYVKRALLLLEI
AGNIEIIADYATNIVEVSVYMVQGEAYKCYHDELLLFKKSGGVLFESSD
10000000000000000000000000000000000000000000000000000000000000
00000000000000000000000000000000000000000000000000000000000000
00000000000000000000000000000000000000000000000000000000000000
0000000000000000000000000000000000000000111111111
>DM_train1230
MKRYRTLLFDVDDTILDFQAAEALALRLLFEDQNIPLTNDMKAQYKTINQGLWRAFEEGKMT
RDEVVNTRFSALLKEYGYEADGALLEQKYRRFLEEGHQLIDGAFDLISNLQQQFDLYIVTNG
VSHTQYKRLRDSGLFPFFKDIFVSEDTGFQKPMKEYFNYVFERIPQFSAEHTLIIGDSLTAD
IKGGQLAGLDTCWMNPDMKPNVPEIIPTYEIRKLEELYHILNIENTVSC
00000000000000000000000000000000000000000000000000000000000000
00000000000000000000000000000000000000000000000000000000000000
00000000000000000000000000000000000000000000000000000000000000
0000000000000000000000000000000000000000000000111
>DM_train1231
MGLMKFQLFIQPKLDVLQGNIVEYEILLRDDSAVPRFPLSELEAVLADEELYLAFSEWFSEA
FLDVLKKYPNDRFAINIAPQQLFYIETLHWLDKLKSESHRITVEMTEDIFDVPGHKRHLNAN
DKNAFILNKIKVIHGLGYHIAIDDVSCGLNSLERVMSYLPYIIEIKFSLIHFKNIPLEDLLL
FIKAWANFAQKNKLDFVVEGIETKETMTLLESHGVSIFQGYLVNKPFPV
11110000000000000000000000000000000000000000000000000000000000
00000000000000000000000000000000000000000000000000000000000000
00000000000000000000000000000000000000000000000000000000000000
0000000000000000000000000000000000000000000000000

>DM_train1232
MGSSSSGLVPRGSTSEVIEDEKQFYSKAKTYWKQIPPTVDGMLGGYGHISSIDINSSRKFLQ
RFLREGPNKTGTSCALDCGAGIGRITKRLLLPLFREVDMVDITEDFLVQAKTYLGEEGKRVR
NYFCCGLQDFTPEPDSYDVIWIQWVIGHLTDQHLAEFLRRCKGSLRPNGIIVIKDNMAQEGV
ILDDVDSSVCRDLDVVRRIICSAGLSLLAEERQENLPDEIYHVYSFALR
111111111100000000000000000000000000000000000000000000000000
000011110000000000000000000000000000000000000000000000000000
000000000000000000000000000000000000000000000000000000000000
0000000000000000000000000000000000000000000000000
>DM_train1233
MVSRLSTGILDFDKLIQGGIPQGFFIALTGEPGTGKTIFSLHFIAKGLRDGDPCIYVTTEES
RDSIIRQAKQFNWDFEEYIEKKLIIIDALMKEKEDQWSLVNLTPEELVNKVIEAKQKLGYGK
ARLVIDSVSALFLDKPAMARKISYYLKRVLNKWNFTIYATSQYAITTSQAFGFGVEHVADGI
IRFRRMIRNGELHRYILIEKMRQTDHDKHVWEIDIVNGKGIVLKGRLEE
000000000000000000000000000000000000000000000000000000000000
000000000000000000000000000000111100000000000000000000000000
000000000000000000000000000000000000000000111111111100000000
0000000000000000000000000000000000000000000000000
>DM_train1234
GMSETITFPPLMTGEAAGPGQDPFDLACQKAELGVDAGLVVYELGTDVLRAALVLAPEVPLA
KAMAMLPVCGVGFQNALGALAPPEVAVHLDWNGALRINGARCGRLRIAASTDDPDTQPDWLV
VGLDLPLWPEGDGGETPDETALYAEGCADVAAPRLLESWARHCLHWINRWDEGELETIHGEW
RGLAHGMGEARTEAGRSGTFLGVDEDFGMLLRDETTTHLIPLTTVLVQD
111100000000000000000000000000000000000000000000000000000000
000000000000000000000000000000000000000000000000000000000000
000000000000000000000000000000000000000000000000000000000000
0000000000000000000000000000000000000000000000001
>DM_train1235
NAKRDRIFVRVYNVMKRINCFINKNIKKSSTDSNYQLAVFMLMETMFFIRFGKMKYLKENET
VGLLTLKNKHIEISPDEIVIKFVGKDKVSHEFVVHKSNRLYKPLLKLTDDSSPEEFLFNKLS
ERKVYECIKQFGIRIKDLRTYGVNYTFLYNFWTNVKSISPLPSPKKLIALTIKQTAEVVGHT
PSISKRAYMATTILEMVKDKNFLDVVSKTTFDEFLSIVVDHVKSSTDG
000000000000000000000000000000000000000000000011111111100000
000000000000000000000000000000000000000000000000000000000000
000000000000000000000000000000000000000000000000000000000000
000000000000000000000000000000000000000000001111
>DM_train1236
SETNTIFKLEGVSVLSPLRKKLDLVFYLSNVDGSPVITLLKGNDRELSIYQLNKNIKMASFL
PVPEKPNLIYLFMTYTSCEDNKFSEPVVMTLNKENTLNQFKKLGLLDSNVTDFEKCVEYIRK
QAILTGFKISNPFVNSTLVDTDAEKINSFHLQCHRGTKEGTLYFLPDHIIFGFKKPILLLDA
SDIESITYSSITRLTFNASLVTKDGEKYEFSMIDQTEYAKIDDYVKRK
110000000000000000000000000000000000000000000000000000000000
000000000000000000000000000000000000000000000000000000000000
000000000000001111111110000000000000000000000000000000000000

000000000000000000000000000000000000000000000000
>DM_train1237
LNEEQLSKRLEKVASYITKNERIADIGSDHAYLPCFAVKNQTASFAIAGEVVDGPFQSAQKQ
VRSSGLTEQIDVRKGNGLAVIEKKDAIDTIVIAGMGGTLIRTILEEGAAKLAGVTKLILQPN
IAAWQLREWSEQNNWLITSEAILREDNKVYEIMVLAPSEKPVTWTKQEIFFGPCLLKEQSAI
FKSKWRHEANTWQNIIQTISNNQPVSTENQAKIRELEHKIALVEDVLK
111100000000000000000000000000000000000000000000000000000000
000000000000000000000000000000000000000000000000000000000000
000000000000000000000000000000000000000000000000000000000000
000000000000000000000000000000000000000000000000
>DM_train1238
DIKVTPGTSELVEQILALLSRYLSSYIHVLNKFISHLRRVATLRFERTTLIKFVKKLRFYND
SVLSYNASEFINEGKNELDPEADSFDKVILPIASMFVKSVETFDLLNYYLTQSLQKEILSKT
LNEDLTLTAESILAIDDTYNHFVKFSQWMIESLRIGSNLLDLEVVQFAIKSADEDGTNIGET
DNIFLQEILPVNSEEEFQTLSAAWHSILDGKLSALDEEFDVVATKWHD
111111100000000000000000000000000000000000000000000000000000
000000000111111111111110000000000000000000000000000000000000
000000000000000000000000000000000000000000000000111111111111
111111111111000000000000000000000000000000000011
>DM_train1239
GMSRTLVLFDIDGTLLKVESMNRRVLADALIEVYGTEGSTGSHDFSGKMDGAIIYEVLSNVG
LERAEIADKFDKAKETYIALFRERARREDITLLEGVRELLDALSSRSDVLLGLLTGNFEASG
RHKLKLPGIDHYFPFGAFADDALDRNELPHIALERARRMTGANYSPSQIVIIGDTEHDIRCA
RELDARSIAVATGNFTMEELARHKPGTLFKNFAETDEVLASILTPKHS
110000000000000000000000000000000000000011100000000000000000
000000000000000000000000000000000000000000000000000000000000
000000000000000000000000000000000000000000000000000000000000
000000000000000000000000000000000000000000001111
>DM_train1240
MASVTEQFNDIISLYSTKLEHTSLRQDSPEYQGLLLSTIKKLLNLKTAIFDRLALFSTNETI
DDVSTASIKFLAVDYYLGLLISRRQSNDSDVAQRQSMKLIYLKKSVESFINFLTLLQDYKLL
DPLVGEKLGNFKDRYNPQLSELYAQPKNNKDLSGAQLKRKEKIELFQRNKEISTKLHCLELE
LKNNDEDHDHDELLRELYLMRLHHFSLDTINNIEQNLFECEMLSNFLK
100000000000000000000110000000000000000000000000000000000000
000000000000000000000000000000000000000000000000000000000000
000000000111111111111111110000000000000000000000000000000000
000000000000000000000000000000000000000000000000
>DM_train1241
MSGDENPASKPTPVQDVQGDGRWMSLHHRFVADSKDKEPEVVFIGDSAVQLMHQCEIWRELF
SPLHALNFGIGGDSTQHVLWRLENGELEHIRPKIVVVWVGTNNHGHTAEQVTGGIKAIVQLV
NERQPQARVVVLGLLPRGQHPNPLREKNRRVNELVRAALAGHPRAHFLDADPGFVHSDGTIS
HHDMYDYLHLSRLGYTPVCRALHSLLLRLLTQDQGQGGAPLPEPSPP
111100000000000000000000000000000000000000000000000000000000
000000000000000000000000000000000000000000000000000000000000

00000000000000000000000000000000000000000000000000000000000000
00000000000000000000000000000011111111111111111
>DM_train1242
ARSEKRVPMTRLRKRVAERLLEAKNSTAMLTTFNEVNMKPIMDLRKQYGEAFEKRHGIRLGF
MSFYVKAVVEALKRYPEVNASIDGDDVVYHNYFDVSMAVSTPRGLVTPVLRDVDTLGMADIE
KKIKELAVKGRDGKLTVEDLTGGNFTITNGGVFGSLMSTPIINPPQSAILGMHAIKDRPMAV
NGQVEILPMMYLALSYDHRLIDGRESVGFLVTIKELLEDPTRLLLDV
11100011111100000000000000000000000000000000000000000000000000
00000000000000000000001111000000000000000000000000000000000000
00000000000000000000000000000000000000000000000000000000000000
00000000000000000000000000000000000000000000000
>DM_train1243
SVPFEYTPIAQSVLDECEHLDTASLSDALDSLGIDGGLPGIASQVPGTRCVGIAFTVQYQPV
DASEGFRGAANYIDQVPSGSVIVSSNSGRHDCTVWGDIMTHFALANGIKGTVIDGVARDIDT
VINCNYPLFSRGRFMQSAKNRTQLKAVQVPLVIDGITIQPGDLMVCDGSGCVVVPQQLAAEV
VLRARAVEQTERRIIEAISSGSTLEQARMTYRYDQPWLSEAEHGGTQ
11111000000000000000000000000000000000000000000000000000000000
11111111110000000000000000000000000000000000000000000000000000
00000000000000000000000000000000000000000000000000000000000000
00000000000000000000000000000111111111111111111
>DM_train1244
KGFSSEDKGEWKLKLDASGNGQAVIRFLPAKTDDALPFAILVNHGFKKNGKWYIETCSSTHG
DYDSCPVCQYISKNDLYNTNKTEYSQLKRKTSYWANILVVKDPQAPDNEGKVFKYRFGKKIW
DKINAMIAVDTEMGETPVDVTCPWEGANFVLKVKQVSGFSNYDESKFLNQSAIPNIDDESFQ
KELFEQMVDLSEMTSKDKFKSFEELNTKFNQVLGTAALGGAAAAAAS
11111111000000000000000000000000000000000000000000000000000000
00000000000000000000000000000000000000000000000000000000000000
00000000000000000000000000000000000000000000000000000000000000
00000000000000000000000000000000001111111111111
>DM_train1245
RAQANEDIVVSKTSLWKEMQYNRDITIRFKFASSIVKSGGLGYKWSEISFKPANYQYTYTRD
GEEVTAHTTCSVNGMNDFNFNGGSLPTDFVISRYEVIKENSYVYVDYWDDSQAFRNMVYVRS
LAANLNSVICTGGDYSFALPVGQWPVMTGGAVSLHSAGVTLSTQFTDFVSLNSLRFRFRLTV
EEPSFSITRTRVSRLYGLPAANPNNGKEYYEVAGRFSLISLVPSNDD
11111111111100000000000000000000000000000000000000000000000000
00000000000000000000000000000000000000000000000000000000000000
00000000000000000000000000000000000000000000000000000000000000
00000000000000000000000000000000000000000000111
>DM_train1246
DNSNGSTIPSSITSGSIFDLEGDNPNPLVDDSTLVFVPLEAQHITPNGNGWRHEYKVKESLR
VAMTQTYEVFEATVKVEMSDGGKTIISQHHASDTGTISKVYVSDTDESGFNDSVANNGIFDV
YVRLRNTSGNEEKFALGTMTSGETFNLRVVNNYGDVEVTAFGNSFGIPVEDDSQSYFKFGNY
LQSQDPYTLDKCGEAGNSNSFKNCFEDLGITESKVTMTNVTYTRETN
11111000000000000000000000000000000000000000000000000000000000

00000000000000000000000000000000000000000000000000000000000000
00000000000000000000000000000000000000000000000000000000000000
00000000000000000000000000000000000000000000000
>DM_train1247
TTVYLAGDSTMAKNGGGSGTNGWGEYLASYLSATVVNDAVAGRSARSYTREGRFENIADVVT
AGDYVIVEFGHNDGGSLSTDNGRTDCSGTGAEVCYSVYDGVNETILTFPAYLENAAKLFTAK
GAKVILSSQTPNNPWETGTFVNSPTRFVEYAELAAEVAGVEYVDHWSYVDSIYETLGNATVN
SYFPINHTHTSPAGAEVVAEAFLKAVVCTGTSLKSVLTTTSFEGTCL
00000000000000000000000000000000000000000000000000000000000000
00000000000000001100000000000000000000000000000000000000000000
00000000000000000000000000000000000000000000000000000000000000
00000000000000000000000000000000000000000000000
>DM_train1248
MGSDKIHHHHHHMKIDILDKGFVELVDVMGNDLSAVRAARVSFDMGLKDEERDRHLIEYLMK
HGHETPFEHIVFTFHVKAPIFVARQWFRHRIASYNELSGRYSKLSYEFYIPSPERLEGYKTT
IPPERVTEKISEIVDKAYRTYLELIESGVPREVARIVLPLNLYTRFFWTVNARSLMNFLNLR
ADSHAQWEIQQYALAIARIFKEKCPWTFEAFLKYAYKGDILKEVQV
11111111110000000000000000000000000000000000001110000000000000
00000000000000000000000000000000000000111110000000000000000000
00000000000000000000000000000000000000000000000000000000000000
0000000000000000000000000000000000000000011111
>DM_train1249
GSHMSQPRPLLSPPETEEQLLAQAQQLSGYTLGELAALVGLVTPENLKRDKGWIGVLLEIWL
GASAGSKPEQDFAALGVELKTIPVDSLGRPLETTFVCVAPLTGNSGVTWETSHVRHKLKRVL
WIPVEGERSIPLAQRRVGSPLLWSPNEEEDRQLREDWEELMDMIVLGQVERITARHGEYLQI
RPKAANAKALTEAIGARGERILTLPRGFYLKKNFTSALLARHFLIQ
11111000000000000000000000000000000000000000000000000000000000
00000111111000000000000000000000000000000000000000000000000000
00000000000000000000000000000000000000000000000000000000000000
0000111000000000000000000000000000000000000000
>DM_train1250
MSVEGLGKDFCGAIIPDNFFPIEKLRNYTQMGLIRDFAKGSAVIMPGEEITSMIFLVEGKIK
LDIIFEDGSEKLLYYAGGNSLIGKLYPTGNNIYATAMEPTRTCWFSEKSLRTVFRTDEDMIF
EIFKNYLTKVAYYARQVAEMNTYNPTIRILRLFYELCSSQGKRVGDTYEITMPLSQKSIGEI
TGVHHVTVSRVLASLKRENILDKKKNKIIVYNLGELKHLSEQTSYY
11111111000000000000000000000000000000000000000000000000000000
00000000000000000000000000000000000000000000000000000000000000
00000000000000000000000000000000000000000000000000000000000000
0000000000000000000000011110000000000000011111
>DM_train1251
MSLITMKKIGVILSGCGVYDGSEIHEAVLTLLAISRSGAQAVCFAPDKQQVDVINHLTGEAM
TETRNVLIEAARITRGEIRPLAQADAAELDALIVPGGFGAAKNLSNFASLGSECTVDRELKA
LAQAMHQAGKPLGFMCIAPAMLPKIFDFPLRLTIGTDIDTAEVLEEMGAEHVPCPVDDIVVD
EDNKIVTTPAYMLAQNIAEAASGIDKLVSRVLVLAEEGGSHHHHHH

11111000000000000000000000000000000000000000000000000000000000
00000000000000000000000000000000000000000000000000000000000000
00000000000000000000000000000000000000000000000000000000000000
0000000000000000000000000000000000011111111111
>DM_train1252
MAAYQVLIVGAGFSGAETAFWLAQKGVRVGLLTQSLDAVMMPFLPPKPPFPPGSLLERAYDP
KDERVWAFHARAKYLLEGLRPLHLFQATATGLLLEGNRVVGVRTWEGPPARGEKVVLAVGSF
LGARLFLGGVVEEAGRLSEASYPDLLEDLSRLGFRFVEREGEVPETPSTPGYRVRYLAFHPE
EWEEKTFRLKRLEGLYAVGLCVREGDYARMSEEGKRLAEHLLHELG
10000000000000000000000000000000000000000000000000000000000000
00000000000000000000000000000000000000000000000000000000000000
00000000000000000000000000000000000000000000111110000000000000
0000000000000000000000000000000000000000000001
>DM_train1253
GSHMAKTLKDLDGWQVIITDDQGRVIDDNNRRRSRKRGGENVFLKRISDGLSFGKGESVIFN
DNVTETYSVYLIHEIRLNTLNNVVEIWVFSYLRWFELKPKLYYEQFRPDLIKEDHPLEFYKD
KFFNEVNKSELYLTAELSEIWLKDFIAVGQILPESQWNDSSIDKIEDRDFLVRYACEPTAEK
FVPIDIFQIIRRVKEMEPKQSDEYLKRVSVPVSGQKTNRQVMHKMG
11111111110000000000000000111111111111100000000000000000000000
00000000000000000111110000000000000000000000000000000000000000
00000000000000000000000000000000000001111111000000000000000000
0000000000000000000000000000000011111111111111
>DM_train1254
MWKTASDKGLGHLLRDVWSLLNEEERELLDKEIQPFPCKKASTVFSEGDIPNNLFYLYEGKI
KILREGVYGRFHISRIVKPGQFFGMRPYFAEETCSSTAIAVENSKVLAIPVEAIEALLKGNT
SFCRYFLKALAKELGYAERRTVTLTQKHVRGRLAETLLILKENFGFENDGATLSIYLSREEL
ATLSNMTVSNAIRTLSTFVSERMLALDGKRIKIIDCDRLQKTARSG
11111111100000000000000000000000000000000000000000000000000000
00001111100000000000000000000000000000000000000000000000000000
00000000000000000000000000000000000000000000000000000000000000
0000000000000000000000000000000000000000000000
>DM_train1255
EIAVQNPLVSERLELSVLYKEYAEDDNIYQQKIKDLHKKYSYIRKTRPDGNCFYRAFGFSHL
EALLDDSKELQRFKAVSAKSKEDLVSQGFTEFTIEDFHNTFMDLIEQVEKQTSVADLLASFN
DQSTSDYLVVYLRLLTSGYLQRESKFFEHFIEGGRTVKEFCQQEVEPMCKESDHIHIIALAQ
ALSVSIQVEYMDRGEGGTTNPHIFPEGSEPKVYLLYRPGHYDILYK
11111000000000000000000000000000000000000000000000000000000000
00000000000000000000000000000000000000000000000000000000000000
00000000000000000000000000000000000000000000000000000000000000
0000000000001000000000000000000000000000000000
>DM_train1256
MPPKRAALIQNLRDSYTETSSFAVIEEWAAGTLQEIEGIAKAAAEAHGVIRNSTYGRAQAEK
SPEQLLGVLQRYQDLCHNVYCQAETIRTVIAIRIPEHKEEDNLGVAVQHAVLKIIDELEIKT
LGSGEKSGSGGAPTPIGMYALREYLSARSTVEDKLLGSVDAESGKTKGGSQSPSLLLELRQI

DADFMLKVELATTHLSTMVRAVINAYLLNWKKLIQPRTGTDHMVS
11111000000000000000000000000000000000000000000000000000000000
00000000000000000000000000000000000000000000000000000000000000
00000000000000000000000111111111111111111111111111000000000000
000000000000000000000000000000000000000000000
>DM_train1257
MSLIFLDYDGTLVPIIMNPEESYADAGLLSLISDLKERFDTYIVTGRSPEEISRFLPLDINM
ICYHGACSKINGQIVYNNGSDRFLGVFDRIYEDTRSWVSDFPGLRIYRKNLAVLYHLGLMGA
DMKPKLRSRIEEIARIFGVETYYGKMIIELRVPGVNKGSAIRSVRGERPAIIAGDDATDEAA
FEANDDALTIKVGEGETHAKFHVADYIEMRKILKFIEMLGVQKKQ
10000000000000000000000000000000000000000000000000000000000000
00000000000000000000000000000000000000000000000000000000000000
00000000000000000000000000000000000000000000000000000000000000
000000000000000000000000000000000000000000001
>DM_train1258
MEGQRWLPLEANPEVTNQFLKQLGLHPNWQFVDVYGMDPELLSMVPRPVCAVLLLFPITEKY
EVFRTEEEEKIKSQGQDVTSSVYFMKQTISNACGTIGLIHAIANNKDKMHFESGSTLKKFLE
ESVSMSPEERARYLENYDAIRVTHETSAHEGQTEAPSIDEKVDLHFIALVHVDGHLYELDGR
KPFPINHGETSDETLLEDAIEVCKKFMERDPDELRFNAIALSAA
11110000000000000000000000000000000000000000000000000000000000
00000000000000000000000000000000000000000000000000000000000000
00000000000000000000001111111111111111111000000000000000000000
00000000000000000000000000000000000000000000
>DM_train1260
DLGAPQNPNAKAAGSRKIHFNWLPPSGKPMGYRVKYWIQGDSESEAHLLDSKVPSVELTNLY
PYCDYEMKVCAYGAQGEGPYSSLVSCRTHQEVPSEPGRLAFNVVSSTVTQLSWAEPAETNGE
ITAYEVCYGLVNDDNRPIGPMKKVLVDNPKNRMLLIENLRESQPYRYTVKARNGAGWGPERE
AIINLATQPKRPMSIPIIPDIPIVDAQSGEDYDSFLMYSDDVLR
00000000000000000000000000000000000000000000000000000000000000
00000000000000000000000000000000000000000000000000000000000000
00000000000000000000000000000000000000000000000000000000000000
00000000000000000000000000001111111111111111
>DM_train1261
QSCVWYGECGIAYGDKRYNCEYSGPPKPLPKDGYDLVQELCPGFFFGQVSLCCDVRQLQTLK
DNLQLPLQFLSRCPSCFYNLLNLFCELTCSPRQSQFLQVTATEDYVDPVTNQTKTNVKELQY
YVGQSFANAMYNACRDVEAPSSNDKALGLLCGKDADACQATNWIEYMFNKDNGQAPFTITPV
FSDFPVHGMEPMNNATKGCDESVDEVTAPCSCQDCSIVCGPKPQ
00000000000000000000000000000000000000000000000000000000000000
00000000000000000000000000000000000000000000000000000000000000
00000000000000000000000000000000000000000000000000000000000000
00000000000000000000000000000000000000011111
>DM_train1262
GMADAPIDIFQSILSRKSIRAFTDQPVTQETIREILKLAARAPSGTNLQPWQVIVLTGKILQ
KVGQELSQLVLSGIKGEREYHYYPRQWREPYLSRRRKVGLDLYKSLGIQKGDQEKMLHQKAK

NFLFYGAPVGLLFTIDHDMEMGSWLDLGMFMQTIMLAARGFGLDTCAQAAFADYHKQIRSLL
SVPSDRHIICGMALGYRDMNAPENNFETEREPIDNFVHFIKSYP
11110000000000000000000000000000000000000000000000000000000000
00000000000000000000000000000000000000000000000000000000000000
00000000000000000000000000000000000000000000000000000000000000
00000000000000000000000000000000000000000000
>DM_train1263
MSKNIVVAPSILSADFSRLGEEIKAVDEAGADWIHVDVMDGRFVPNITIGPLIVDAIRPLTK
KTLDVHLMIVEPEKYVEDFAKAGADIISVHVEHNASPHLHRTLCQIRELGKKAGAVLNPSTP
LDFLEYVLPVCDLILIMSVNPGFGGQSFIPEVLPKIRALRQMCDERGLDPWIEVDGGLKPNN
TWQVLEAGANAIVAGSAVFNAPNYAEAIAGVRNSKRPEPQLATV
11000000000000000000000000000000000000000000000000000000000000
00000000000000000000000000000000000110000000000000000000000000
00000000000000000000111111000000000000000000000000000000000000
00000000000000000000000000000000000000111111
>DM_train1264
TKLYYEDQYIKEFKGEIIEVKEIDGKFHVLLDQTAFFPGGGGQMGDLGLIDGIKVLDVYEEE
GKVYHVLEKEPKKLKNLQCELDWERRFDGMQQHLGQHLLSGCFYDLFGANTCGFHLGKEIST
VDIVGFLDEKTIREAEKEANRLIFENLEVKSYAPSKKELKKVKTRRALPKTDEEIRIVEIVG
LDLNACCGVHPRNTRDLQVIKIRRWEKHKNATRIEYVAGNRAVS
00000000000000000000000000000000000000000000000000000000000000
00000000000000000000000000000000000000000000000000000000000000
00000000000000000000000000000000000000000000000000110000000000
00000000000000000000000000000000000000000001
>DM_train1266
LLHKRVVLASASPRRQEILSNAGLRFEVVPSKFKEKLDKASFATPYGYAMETAKQKALEVAN
RLYQKDLRAPDVVIGADTIVTVGGLILEKPVDKQDAYRMLSRLSGREHSVFTGVAIVHCSSK
DHQLDTRVSEFYEETKVKFSELSEELLWEYVHSGEPMDKAGGYGIQALGGMLVESVHGDFLN
VVGFPLNHFCKQLVKLYYPPRPEDLRRSVKHDSIPAADTFEDLS
00000000000000000000000000000000000000000000000000000000000000
00000000000000000000000000000000000000000000000000000000000000
00000000000000000000000000000000000000000000001110000000000000
00000000000000000011111111111111111111111111
>DM_train1267
MTTGVRRRMGVEERRQQLIGVALDLFSRRSPDEVSIDEIASAAGISRPLVYHYFPGKLSLYE
AALQRASDDLADRFVEPRQGPLGARLLRVMGRYFDFVDEHGPGFSALMRGGPAVGSTTTNAL
VDSVRQAAYVQILSHLDVTEPPARLELVVRSWISLAESTALLWLDGRRIPRAELETQLVHDF
AALMAVSAAYDEEMGALVRRVLADEPEDGPFGDLVDRLLALSAR
11111111111110000000000000000000000000000000000000000000000000
00000000000000000000000000000000000000000000000001111110000000
00000000000000000000000000000000000000000000000000000000000000
00000000000000000000000000010000000000000011
>DM_train1268
TVSTINSTDALAMVEHSSELTLSITTPVGTKFVCRTPFIGTHTDKFLLVEMPKISADDLQYF

FQEGFWMNIRAISPRGEGALIHFRSQLMHILQEPVPMAFLSIPNTMQVSQLRKEPRFELNLA
GKVLFDEHRGDCELRDLSRSGCRFITPPLGKTYQVGDLVALEIFSDLRGTKTFPPLTGKICN
LQRSLHHARYGLEFNEEGRNNAKNLLAQLKFNGTKLTLNAEKKA
10000000000000000000000000000000000000000000000000000000000000
00000000000000000000000000000000000000000000000000000000000000
00000000000000000000000000000000000000000000000000000000000000
00000000000000000000000000000000000000011111
>DM_train1269
GHHHHHHHGESLFKGPRDYNPISSTICHLTNESDGHTTSLYGIGFGPFIITNKHLFRRNNGT
LLVQSLHGVFKVKNTTTLQQHLIDGRDMIIIRMPKDFPPFPQKLKFREPQREERICLVTTNF
QTKSMSSMVSDTSCTFPSSDGIFWKHWIQTKDGQCGSPLVSTRDGFIVGIHSASNFTNTNNY
FTSVPKNFMELLTNQEAQQWVSGWRLNADSVLWGGHKVFMDKP
11111111110000000000000000000000000000000000000000000000000000
00000000000000000000000000000000000000000000000000000000000000
00000000000000000000000000000000000000000000000000000000000000
0000000000000000000000000000000000000000000
>DM_train1270
KERAERDISMVVSGLTPKEVMIVQKFAEKYRLALTDVITEETTHVIIKTDAEFVCERTLKYF
LGIAGGKWIVSYSWVIKSIQERKLLSVHEFEVKGDVVTGSNHQGPRRSRESQEKLFEGLQIY
CCEPFTNMPKDELERMLQLCGASVVKELPLLTRDTGAHPIVLVQPSAWTEDNDCPDIGQLCK
GRLVMWDWVLDSISVYRCRDLDAYLVQNITCGRDGSEPQDSND
00000000000000000000000000000000000000000000000000000000000000
00000000000000000000000000000000000000000000000000000000000000
00000000000000000000000000000000000000000000111111111111111111
1000000000000000000000000001111111111111111
>DM_train1271
MELLLLSNSTLPGKAWLEHALPLIANQLNGRRSAVFIPFAGVTQTWDEYTDKTAEVLAPLGV
NVTGIHRVADPLAAIEKAEIIIVGGGNTFQLLKESRERGLLAPMADRVKRGALYIGWSAGAN
LACPTIRTTNDMPIVDPNGFDALDLFPLQINPHFTNALPEGHKGETREQRIRELLVVAPELT
VIGLPEGNWIQVSNGQAVLGGPNTTWVFKAGEEAVALEAGHRF
00000000000000000000000000000000000000000000000000000000000000
00000000000000000000000000000000000000000000000000000000000000
00000000000000000000000000000000000011111111000000000000000000
0000000000000000000000000000000000000000000
>DM_train1272
MDKIKQLFANNYSWAQRMKEENSTYFKELADHQTPHYLWIACSDSRVPAEKLTNLEPGELFV
HRNVANQVIHTDFNCLSVVQYAVDVLKIEHIIICGHTNCGGIHAAMADKDLGLINNWLLHIR
DIWFKHGHLLGKLSPEKRADMLTKINVAEQVYNLGRTSIVKSAWERGQKLSLHGWVYDVNDG
FLVDQGVMATSRETLEISYRNAIARLSILDEENILKKDHLENT
00000000000000000000011111111110000000000000000000000000000000
00000000000000000000000000000000000000000000000000000000000000
00000000000000000000000000000000000000000000000000000000000000
0000000000000000000000000000011111111111111
>DM_train1273

SVDHGFLVTRHSQTTDDPQCPPGTKILYHGYSLLYVQGNERAHGQDLGTAGSCLRKFSTMPF
LFCNINNVCNFASRNDYSYWLSTPEPMPMSMAPITGENIRPFISRCAVCEAPAMVMAVHSQT
IQIPQCPTGWSSLWIGYSFVMHTSAGAEGSGQALASPGSCLEEFRSAPFIECHGRGTCNYYA
NAYSFWLATIERSEMFKKPTPSTLKAGELRTHVSRCQVCMRRT
11110000000000000000000000000000000000000000000000000000000000
00000000000000000000000000000000000000000000000000000000000000
00000000000000000000000000000000000000000000000000000000000000
0000000000000000000000000000000000000000011
>DM_train1274
MTTVRIPAGWPATEEEARAVQDELRGRVILDEPGPPPGTGRVTGVDVAYDDERDVVVAAAVV
LDAATLDVVAEATAVGEVSFPYVPGLLAFREIPTVLAALDALPCPPGLIVCDGYGVAHPRRF
GLASHLGVLTGLPTIGVAKNPFTFSYEDPGAPRGSAAPLLAGADEVGRALRTQSGVKPVFVS
VGHRVDLDHACAHTLALTPKYRIPETTRRADSLCRRALKEATA
10000000000000000000000000000000000000000000000000000000000000
00000000000000000000000000000000000000000000000000000000000000
00000000000000000000000000000000000000000000000000000000000000
0000000000000000000000000000000000000000000
>DM_train1275
MNGIHDVGGMDGFGKVMYVKEEEDIYFTHDWERLALGLVAGCMAQGLGMKAFDEFRIGIELM
RPVDYLTSSYYGHWIATVAYNLVDTGVLDEKELDERTEVFSKKPDTKIPRREDPALVKLVEK
ALNDGLSPLREISASPRFKVGERIKTKNIHPTGHTRFPRYARDKYGVIDEVYGAHVFPDDAA
HRKGENPQYLYRVRFEAEELWGYKQKDSVYIDLWESYMEPVSH
00000000000000000000000000000000000000000000000000000000000000
00000000000000000000000000000000000000000000000000000000000000
00000000000000000000000000000000000000000000000000000000000000
0000000000000000000000000000000000000000011
>DM_train1276
SHMTPDIILQRTGIDVRAVEQGDDAWHKLRLGVITASEVHNVIAKPRSGKKWPDMKMSYFHT
LLAEVCTGVAPEVNAKALAWGKQYENDARTLFEFTSGVNVTESPIIYRDESMRTACSPDGLC
SDGNGLELKCPFTSRDFMKFRLGGFEAIKSAYMAQVQYSMWVTRKNAWYFANYDPRMKREGL
HYVVIERDEKYMASFDEIVPEFIEKMDEALAEIGFVFGEQWR
11100000000000000000000000000000000000000000000000000000000000
00000000000000000000000000000000000000000000000000000000000000
00000000000000000000000000000000000000000000000000000000000000
000000000000000000000000000000000000000000
>DM_train1277
AESLPDLKIEKLDEGVYVHTSFEEVNGWGVVPKHGLVVLVNAEAYLIDTPFTAKDTEKLVTW
FVERGYKIKGSISSHFHSDSTGGIEWLNSRSIPTYASELTNELLKKDGKVQATNSFSGVNYW
LVKNKIEVFYPGPGHTPDNVVVWLPERKILFGGCFIKPYGLGNLGDANIEAWPKSAKLLKSK
YGKAKLVVPSHSEVGDASLLKLTLEQAVKGLNESKKPSKPSN
11100000000000000000000000000000000000000000000000000000000000
00000000000000000000000000000000000000000000000000000000000000
00000000000000000000000000000000000000000000000000000000000000
000000000000000000000000000000000111111111

>DM_train1278
GMSEDHPLDRPVWNSLGGPQSELDVASGNLRRLDPAYGPFAAAAPGAEAGLASLLQGDADEI
WLVEPEPVAPPPGTRVIRVAPLLQMIADGPVPSFDDPGIVALGETDVPEMTALALATEPGPW
ASGTWRYGQFYGVRIDGRLAAMAGERMRPAPNLAEVSGVCTWPEYRGRGLAARLIRKVIAGM
AARGEVPYLHSYASNASAIRLYESLGFRARRAMTATLLGKST
111100000000000000000000000000000000000000000000000000000000
000000000000000000000000000000000000000000000000000000000100
000000000000000000000000000000000000000000000000000000000000
000000000000000000000000000000000000000000
>DM_train1279
RTEQQAASLEQTAASMEQLTATVKQNAENARQASHLALSASETAQRGGKVVDNVVQTMRDIS
TSSQKIADIISVIDGIAFQTNILALNAAVEAARAGEQGRGFAVVAGEVRNLAQRSAQAAREI
KSLIEDSVGKVDVGSTLVESAGETMAEIVSAVTRVTDIMGEIASASDEQSRGIDQVGLAVAE
MDRVTQQNAALVEQSAAAAAALEEQASRLTEAVAVFRIQQQ
111111000000000000000000000000000000000000000000000000000000
000000000000000000000000000000000000000000000000000000000000
000000000000000000000000000000000000000000000000000000000000
00000000000000000000000000000000000000000
>DM_train1280
REFPPPNPDPKFESKAALLAARGPEELLCFTERLEDLVCFWEEAASAGVGPGQYSFSYQLED
EPWKLCRLHQAPTARGAVRFWCSLPTADTSSFVPLELRVTAASGAPRYHRVIHINEVVLLDA
PVGLVARLADESGHVVLRWLPPPETPMTSHIRYEVDVSAGQGAGSVQRVEILEGRTECVLSN
LRGRTRYTFAVRARMAEPSFGGFWSEWSEPVSLLTPSDLDP
111111110000000000000000000000000000000000000000000000000000
000000000000000000000000000000000000000000000000000000000000
000000000000000000000000000000000000000000000000000000000000
00000000000000000000000000000000000111111
>DM_train1281
MGSDKIHHHHHHMKNSFFSKFTPKEPKFFPLLKQLSDVLSASSVLLVESMEHDLPTERADYY
KQIKDMEREGDRLTHLIFDELSTTFITPFDREDIHDLASCMDDVIDGINSSAKRIVIYNPRP
ISESGKELSRLIHEEAINIGKAMDELETFRKNPKPLRDYCTQLHDIENQADDVYELFITKLF
EEEKDCIELIKIKEIMHELEKTTDAAEHVGKILKNLIVKYS
111111111111111111000000000000000000000000000000000000000000
000000000000000000000000000000000000000000000000000000000000
000000000000000000000000000000000000000000000000000000000000
00000000000000000000000000000000000000000
>DM_train1282
MERPLESYKKEAAHAAIAYVQDGMVVGLGTGSTARYAVLELARRLREGELKGVVGVPTSRAT
EELAKREGIPLVDLPPEGVDLAIDGADEIAPGLALIKGMGGALLREKIVERVAKEFIVIADH
TKKVPVLGRGPVPVEIVPFGYRATLKAIADLGGEPELRMDGDEFYFTDGGHLIADCRFGPIG
DPLGLHRALLEIPGVVETGLFVGMATRALVAGPFGVEELLP
110000000000000000000000000000000000000000000000000000000000
000000000000000000000000000000000000000000000000000000000000
000000000000000000000000000000000000000000000000000000000000

00000000000000000000000000000000000000000
>DM_train1283
MSLIEERLKHYLEKQIPARDQYIEQMEREAHEQQVPIMDLLGMESLLHLLKMAAPARILEIG
TAIGYSAIRMAQALPEATIVSIERDERRYEEAHKHVKALGLESRIELLFGDALQLGEKLELY
PLFDVLFIDAAKGQYRRFFDMYSPMVRPGGLILSDNVLFRGLVAETDIEHKRHKQLATKIDT
YNQWLLEHPQYDTRIFPVGDGIAISIKRETKGDTDDEKAEG
11110000000000000000000000000000000000000000000000000000000000
00000000000000000000000000000000000000000000000000000000000000
00000000000000000000000000000000000000011111111111111111111111
11000000000000000000000000001111111111111
>DM_train1284
GEMDERLLQKYRAQVFEWGGCFDKMFEALKSLIYLSEFENSEFDDEERHLLTLCIKHKISDY
RTMTSQVLQEQTKQLNNDELVKICSEYVFSLRKDIKAFLQSFEDCVDRLVEKSFFSKFFKLK
VKSDISRYKLEFGLCSLEDSKKIHQDAFTLLCEHPDKIEQLPLGFIQNLAYILSEKYGEKKQ
VFNMLNSLGKILELQIKEQENMDRKAQITVYLQGIKDYIEK
11000000000000000000000000000000000000000000000000000000000000
00000000000000000000000000000000000000000000000000000000000000
00000000000000000000000000000000000000000000000000000000000000
00000000000000000000000000000000000011111
>DM_train1285
SLTGEESDTLRKIVLEECLPNQQQNQNPSPCAEVKPNAGYVVLKDLNGPLQYLLMPTYRING
TESPLLTDPSTPNFFWLAWQARDFMSKKYGQPVPDRAVSLAINSRTGRTQNHFHIHISCIRP
DVREQLDNNLANISSRWLPLPGGLRGHEYLARRVTESELVQRSPFMMLAEEVPEAREHMGSY
GLAMVRQSDNSFVLLATQRNLLTLNRASAEEIQDHQCEILR
11111100000000000000000000000000000000000000000000000000000000
00000000000000000000000000000000000000000000000000000000000000
00000000000000000000000000000000000000000000000000000000000000
00000000000000000000000000000000000000001
>DM_train1287
MRDWVIPPISCPENEKGEFPKNLVQIKSNRDKETKVFYSITGQGADKPPVGVFIIERETGWL
KVTQPLDREAIAKYILYSHAVSSNGEAVEDPMEIVITVTDQNDNRPEFTQEVFEGSVAEGAV
PGTSVMKVSATDADDDVNTYNAAIAYTIVSQDPELPHKNMFTVNRDTGVISVLTSGLDRESY
PTYTLVVQAADLQGEGLSTTAKAVITVKDINDNAPVFNPS
11110000000000000000000000000000000000000000000000000000000000
00000000000000000000000000000000000000000000000000000000000000
00000000000000000000000000000000000000000000000000000000000000
0000000000000000000000000000011111111111
>DM_train1288
HGIRMTRISREMMKELLSVYFIMGSNNTKADPVTVVQKALKGGATLYQFREKGGDALTGEAR
IKFAEKAQAACREAGVPFIVNDDVELALNLKADGIHIGQEDANAKEVRAAIGDMILGVAAHT
MSEVKQAEEDGADYVGLGPIYPTETKKDTRAVQGVSLIEAVRRQGISIPIVGIGGITIDNAA
PVIQAGADGVSMISAISQAEDPESAARKFREEIQTYKTGR
00000000000000000000000000000000000000000000000000000000000000
00000000000000000000000000000000000000000000000000000000000000

000000000000000000000011111110000000000000000000000000000000000
000000000000000000000000000000000000000000
>DM_train1289
LTGPLMLDTAPNAFDDQYEGCVNKMEEKAPLLLQEDFNMNAKLKVAWEEAKKRWNNIKPSRS
YPKGFNDFHGTALVAYTGSIAVDFNRAVREFKENPGQFHYKAFHYYLTRALQLLSNGDCHSV
YRGTKTRFHYTGAGSVRFGQFTSSSLSKKVAQSQEFFSDHGTLFIIKTCLGVYIKEFSFRPD
QEEVLIPGYEVYQKVRTQGYNEIFLDSPKRKKSNYNCLYS
111000000000000000000000000000000000000000000000000000000000000
000000000000000000000000000000000000000000000000000000000000000
000000000000000000000000000000000000000000000000000000000000000
000000000000000000000000000000000000000000
>DM_train1290
WVPHELTENQKNRRFEVSSSLILRNHNEPFLDRIVTCDEKWILYDNRRRSAQWLDQEEAPKH
FPKPILHPKKVMVTIWWSAAGLIHYSFLNPGETITSEKYAQEIDEMNQKLQRLQLALVNRKG
PILLHDNARPHVAQPTLQKLNELGYEVLPHPPYSPDLLPTNYHVFKHLNNFLQGKRFHNQQD
AENAFQEFVESQSTDFYATGINQLISRWQKCVDCNGSYFD
111000000000000000000000000000000000000001111111111111111111111
111111110000000000000000000000111111100000000000000000000011111000
000000000000000000000000000000000000000000000000000000000000000
000000000000000000000000000000000000000001
>DM_train1291
GMAEDQTVLAIDIGGSHVKIGLSTDGEERKVESGKTMTGPEMVAAVTAMAKDMTYDVIAMGY
PGPVVHNKPLREPVNLGEGWVGYDYEGAFGRPVRIVNDALMQAIGSYNGGRMLFLGLGTGLG
AAMIVENVAQPMEIAHLPYRKGKTYEHYVSEAYREKKGNAKWQKRVQDVVERLSAALEPDEV
VIGGGNVERLENLPPKCRRGDNAMAFEGGFRLWKNADLIV
111100000000000000000000000000000000000000000000000000000000000
000000000000000000000000000000000000000000000000000000000000000
000000000000000000000000000000000000000000000000000000000000000
000000000000000000000000000000000000000000
>DM_train1292
MGPLPRTVELFYDVLSPYSWLGFEILCRYQNIWNINLQLRPSLITGIMKDSGNKPPGLLPRK
GLYMANDLKLLRHHLQIPIHFPKDFLSVMLEKGSLSAMRFLTAVNLEHPEMLEKASRELWMR
VWSRNEDITEPQSILAAAEKAGMSAEQAQGLLEKIATPKVKNQLKETTEAACRYGAFGLPIT
VAHVDGQTHMLFGSDRMELLAHLLGEKWMGPIPPAVNARL
111000000000000000000000000000000000000000000000000000000000000
000000000000000000000000000000000000000000000000000000000000000
000000000000000000000000000000000000000000000000000000000000000
000000000000000000000000000000000011111
>DM_train1293
MLSVAGHRFASLATTPVLRMGLARCFSSVTGPFQCPPLPYVKNALEPHMSAETLTYHHDKHH
QTYVDTLNSIAAENSTIASKTLEQIIKTETGKPFNQAAQVYNHTFFFNNLAPNGGGEPTGKI
AELITRDFGSFEKFKEDFSAAAVGHFGSGWVWLIADDGKLKIVQGHDAGNPIRESKTPLMNI
DVWEHAYYIDYRNARAQYVKNYWNLVNWDFVNDNVAKAGI
111111111111111111111111111110000000000000000000000000000000000

000000000000000000000000000000000000000000000000000000000000
000000000000000000000000000000000000000000000000000000000000
0000000000000000000000000000000000000000
>DM_train1294
MSSGAGSDATGAGGVHAAGSGDRAVAAAVERAKATAARNIPAFDDLPVPADTANLREGADLN
NALLALLPLVGVWRGEGEGRGPDGDYRFGQQIVVSHDGGDYLNWESRSWRLTATGDYQEPGL
REAGFWRFVADPYDPSESQAIELLLAHSAGYVELFYGRPRTQSSWELVTDALARSRSGVLVG
GAKRLYGIVEGGDLAYVEERVDADGGLVPHLSARLSRFVG
111111111111111111111111100000000000000000000000000000000000
000000000000000000000000000000000000000000000000000000000000
000000000111111111000000000000000000000000000000000000000000
0000000000000000000000000000000000000000
>DM_train1295
KSAVTAQSILEKADEIRFPQDSFQVNVAIRTAAPDHAEDLYRYQVLSKGNENSIVMITEPAS
ERGQAILMKGRDLWVFMPSVSQPIRLSLSQRLTGQVANGDIARANFTGDYHPQLLRNESIDD
EDYYVLELTGIDRSVTYQKVLLWVNQSNFRPYKAEFYSVSGRLLKTSRYENFDNILGEMRPT
RIIMEDALKSGEVSVLDYSDMKLRDLPDKIFTKDYLKRLE
110000000000000000000000000000000000000000000000000000000000
000000000000000000000000000000000000000000000000000000000000
000000000000000000000000000000000000000000000000000000000000
0000000000000000000000000000000000001111
>DM_train1296
MISHSELRKLFYSADAVCFDVDSTVIREEGIDELAKICGVEDAVSEMTRRAMGGAVPFKAAL
TERLALIQPSREQVQRLIAEQPPHLTPGIRELVSRLQERNVQVFLISGGFRSIVEHVASKLN
IPATNVFANRLKFYFNGEYAGFDETQPTAESGGKGKVIKLLKEKFHFKKIIMIGDGATDMEA
CPPADAFIGFGGNVIRQQVKDNAKWYITDFVELLGELEE
111100000000000000000000000000000000000001111111111100000 0
000000000000000000000000000000000000000000000000000000000000
000000000000000000000000000000000000000000000000000000000000
000000000000000000000000000000000001111
>DM_train1298
SKRYFVTGTDTEVGKTVASCALLQAAKAAGYRTAGYKPVASGSEKTPEGLRNSDALALQRNS
SLQLDYATVNPYTFAEPTSPHIISAQEGRPIESLVMSAGLRALEQQADWVLVEGAGGWFTPL
SDTFTFADWVTQEQLPVILVVGVKLGCINHAMLTAQVIQHAGLTLAGWVANDVTPPGKRHAE
YMTTLTRMIPAPLLGEIPWLAENPENAATGKYINLALL
000000000000000000000000000000000000000000000000000000000000
000000000000000000000000000000000000000000000000000000000000
000000000000000000000000000000000000000000000000000000000000
00000000000000000000011111000000000000
>DM_train1299
MADPESPWSQIGRKIKLEGLSDVASISTKLQNTLIQYHSIEEDEWRVAKKAKDVTVWRKPSE
EFNGYLYKAQGVMDDVVNNVIDHIRPGPWRLDWDRLMTSLDVLEHFEENCCVMRYTTAGQLL
NIISPREFVDFSYTVGYEEGLLSCGVSVEWSETRPEFVRGYNHPCGWFCVPLKDSPSQSLLT
GYIQTDLRGMIPQSAVDTAMASTLANFYSDLRKGLRKA

1111111111111111111111000000000000000000000000000000000000000
000000000000000000000000000000000000000000000000000000000000
000000000000000000000000000000000000000000000000000000000000
00000000000000000000000000000000000000
>DM_train1300
GSRTVDLELELQIELLRETKRKYESVLQLGRALTAHLYSLLQTQHALGDAFADLSQKSPELQ
EEFGYNAETQKLLCKNGETLLGAVNFFVSSINTLVTKTMEDTLMTVKQYEAARLEYDAYRTD
LEELSLGPRDAGTRGRLESAQATFQAHRDKYEKLRGDVAIKLKFLEENKIKVMHKQLLLFHN
AVSAYFAGNQKQLEQTLQQFNIKLRPPGAEKPSWLEEQ
111110000000000000000000000000000000000000000000000000000000
000000000000000000000000000000000000000000000000000000000000
000000001111111111111111111000000000000000000000000000000000
0000000000000000111111111111111111111111
>DM_train1301
GDNDAIYYPVGDVDIERGGPALEVGEEDVLVARSFNEEDYVLDTIAQYPNDPTLGKLTFMID
LKNQQKDQNVADFNGVGKSKLTMSLGYKDGNYPSESQVPIYTSQDVTAKYAVKLRLKGELLV
SGDEWMIDYVYAQLASLFQPYPPANFPEVFMCKGGMKLGTFDSFRRTCTFDITYDRSDLSFS
QLYFNLFINLAGQKRENRVRLRIDKESYFELYEQSEEM
111000000000000000000000000000000000000000000000000000000000
000000000000000000000000000000000000000000000000000000000000
000000000000000000000000000000000000000000000000000000000000
00000000000000000000000000000000000111
>DM_train1303
SNAMSKRVLMVVTNHTTITDDHKTGLWLEEFAVPYLVFQEKGYDVKVASIQGGEVPLDPRSI
NEKDPSWAEAEAALKHTARLSKDDAHGFDAIFLPGGHGTMFDFPDNETLQYVLQQFAEDGRI
IAAVCHGPSGLVNATYKDGTPIVKGKTVTSFTDEEEREVGLDVHMPFLLESTLRLRGANFVR
GGKWTDFSVRDGNLITGQNPQSSRSTAEKVVAALEERE
111000000000000000000000000000000000000000000000000000000000
000000000000000000000000000000000000000000000000000000000000
000000000000000000000000000000000000000000000000000000000000
00000000000000000000000000000000000000
>DM_train1304
AIAENSSKEDYKIQSFDLETQKLLKTALKDPGSVDLEKVSSVIVDQSLKDQVFSREAGRICY
TIVQAEAKQTNGSVFRRNLLNRLQQEFKAREETRKRSTQEWVCLVSFICNIFDYLKVNNMPM
VALVHPVYDCLFRLAQSDALKNEEEVDCLVLQLHRIGDQLEKMNVQLMDELFNLLRDGFLLQ
EDLSSMGRLLLLEILEFRAGGWKLSDTAQKYYYSEVTD
111111111000000000000000000000000000000000000000000000000000
000000000000000000000000000000000000000000000000000000000000
000000000000000000000000000000000000000000000000000000000000
00000000000000000000000000000000011111
>DM_train1305
SWNKKNEEKEDFGIIVILWKQVTVKEDGKVPLEPFLTAAKEVLRVVDAFGSGFRIVKNDIAG
NIKKLYRANQTVHAETLQELIIAENSPDGLATVALLWLKRAFQFIASFLRRLVVTDKSLEQC
VTEAYNCTLRPCHSAVIQKVFWGGVKLAPSRERFYRKLHPDLNIAKAKIEEFLIELHDPLCC

IVQFFFQRELEDQCWGDEVYQRKDSSEWLKVSCEQDSV
111111111100000000000000000000000000000000000000000000000000
000000000000000000000000000000000000000000000000000000000000
000000000000000000000000000000000000000000000000000000000000
00000000000000000000000000000011111111
>DM_train1306
QPLGGESICSARAPAKYSITFTGKWSQTAFPKQYPLFRPPAQWSSLLGAAHSSDYSMWRKNQ
YVSNGLRDFAERGEAWALMKEIEAAGEALQSVHEVFSAPAVPSGTGQTSAELEVQRRHSLVS
FVVRIVPSPDWFVGVDSLDLCDGDRWREQAALDLYPYDAGTDSGFTFSSPNFATIPQDTVTE
ITSSSPSHPANSFYYPRLKALPPIARVTLLRLRQSPR
111111000000000000000000000000000000000000000000000000000000
000000000000000000000000000000000000000000000000000000000000
000000000000000000000000000000000000000000000000000000000000
0000000000000000000000000000000011111
>DM_train1307
APSCPQNVNISGGTFTLSHGWAPGSLLTYSCPQGLYPSPASRLCKSSGQWQTPGATRSLSKA
VCKPVRCPAPVSFENGIYTPRLGSYPVGGNVSFECEDGFILRGSPVRQCRPNGMWDGETAVC
DNGAGHCPNPGISLGAVRTGFRFGHGDKVRYRCSSNLVLTGSSERECQGNGVWSGTEPICRQ
PYSYDFPEDVAPALGTSFSHMLGATNPTQKTKESLGR
110000000000000000000000000000000000000000000000000111111100
000000000000000000000000000000000000000000000000000000000000
000000000000000000000000000000000000000000000000000000000000
0000000000011111111111111111111111111
>DM_train1308
GMNDLKRLPYEPVKGLLPRPAVGTSERVITLPEPDRTSGMPLMGTLWLRKSTREFDQQPLPL
KQLSELLWAAAGVNRSLGGGRTAPSPYGETVIDVYVALPAGLYRYDPVHHCLELKRAADLRS
MTGYQDFVGMAPLDLVFVANHGRMQEMPPKLRETFSAAAAGAMAENAYLYCASAGLGAVVRG
WLNRRQLAEHMSLNEDEEPILSQTIGRAASHVTTTQA
111111000000000000000000000000000000000000000000000000000000
000000000000000000000000000000000000000000000000000000000000
000000000000000000000000000000000000000000000000000000000000
0000000000000000000000000000000111111
>DM_train1309
SNAMYKEGACLYRNPLRSKSDVKDWRMEGGGQISFDDHSLHLSHVQDEAHFVFWCPETFPDG
IIVTWDFSPIEQPGLCMLFFAAAGIRGEDLFDPSLRKRTGTYPEYHSGDINALHLSYFRRKY
AEERAFRTCNLRKSRGFHLAAMGADPLPSPDDADSPYRMKLIKDKGYVHFSINGLPILEWMD
DGSTYGPVLTKGKIGFRQMAPMKAVYRDFAVHQAVRR
111000000000000000000000000000000000000000000000000000000000
000000000000000000000000000000000000000000000000000000000000
000000000000000000000000000000000000000000000000000000000000
0000000000000000000000000000000000000
>DM_train1310
GEQQKRKIVLDPSGSMNIYLVLDGSDSIGASNFTGAKKSLVNLIEKVASYGVKPRYGLVTYA
TYPKIWVKVSEADSSNADWVTKQLNEINYEDHKLKSGTNTKKALQAVYSMMSWPDDVPPEGW

NRTRHVIILMTDGLHNMGGDPITVIDEIRDLLYIGKDRKNPREDYLDVYVFGVGPLVNQVNI
NALASKKDNEQHVFKVKDMENLEDVFYQMIDESQSLS
11111111111111000000000000000000000000000000000000000000000000
00000000000000000000000000000000000000000000000000000000000000
00000000000000000000000000000000000000000000000000000000000000
0000000000000000001111111111111111100
>DM_train1311
GMDRTLTHQPQEYAVSVSVGEVKLKGNLVIPNGATGIVLFAHGSGSSRYSPRNRYVAEVLQQ
AGLATLLIDLLTQEEEEIDLRTRHLRFDIGLLASRLVGATDWLTHNPDTQHLKVGYFGASTG
GGAALVAAAERPETVQAVVSRGGRPDLAPSALPHVKAPTLLIVGGYDLPVIAMNEDALEQLQ
TSKRLVIIPRASHLFEEPGALTAVAQLASEWFMHYLR
11111110000000000000000000000000000000000000000000000000000000
00000000000000000000000000000000000000000000000000000000000000
00000000000000000000000000000000000000000000000000000000000000
0000000000000000000000000000000000000
>DM_train1312
MAASITAITVENLEYPAVVTSPVTGKSYFLGGAGERGLTIEGNFIKFTAIGVYLEDIAVASL
AAKWKGKSSEELLETLDFYRDIISGPFEKLIRGSKIRELSGPEFSRKVMENCVAHLKSVGTY
GDAEAEAMQKFAEAFKPVNFPPGASVFYRQSPDGILGLSFSPDTSIPEKEAALIENKAVSSA
VLETMIGEHAVSPDLKRCLAARLPALLNEGAFKIGN
11100000000000000000000000000000001111111100000000000000000000
00000000000000000000000000000000000000000000000000000000000000
00000000000000000000000000000000000000000000000000000000000000
000000000000000000000000000000011111
>DM_train1313
GELFSNQIIWFVDDTNVYRVTIHKTFEGNLTTKPINGAIFIFNPRTGQLFLKIIHTSVWAGQ
KRLGQLAKWKTAEEVAALIRSLPVEEQPKQIIVTRKGMLDPLEVHLLDFPNIVIKGSELQLP
FQACLKVEKFGDLILKATEPQMVLFNLYDDWLKTISSYTAFSRLILILRALHVNNDRAKVIL
KPDKTTITEPHHIWPTLTDEEWIKVEVQLKDLILAD
11111000000000000000011111111111100000000000000000000000000000
00000000000000000000000000000000000000000000000000000000000000
00000000000000000000000000000000000000000000000000000000000000
000000000000000000000000000000000011
>DM_train1314
GMPIDTPTDASPADGSASDTVFFGIMSGLELGTFVPGQRLVETDLVAHFGVGRNSVREALQR
LAAEGIVDLQRHRGAVIRRLSLQETLDVLDVAERMTGLLARAATRGSGNQPQVQALRASVQA
LVAAEKAQDGETFSNARRHFYRTLLEMGDNRELRRLFPTIHMPIVHAQHRLASLRQMRLDDY
RRIATAVLAGEPDAAEAAGAAHVKNVRGAILDRQPA
11111111111111100000000000000000000000000000000000000000000000
00000000000000000000000000000000000000000000000000000000000000
00000000000000000000000000000000000000000000000000000000000000
000000000000000000000000000000000111
>DM_train1315
MEIPPTHYAASRAASVAENCINYQQGTPHKLFLVQTVQQASKEDIPGRGHKYHLKFSVEEII

QKQVTVNCTAEVLYPQMGQGSAPEVNFTFEGEIGKNPDEEDNTFYQSLMSLKRPLEAQDIPD
NFGNVSPQMKPVQHLAWVACGYVMWQNSTEDTWYKMLKIQTVKQVQRNDDFIELDYTILLHD
IASQEIIPWQMQVLWHPQYGTKVKHNSRLPKEGQAE
0000000000000000000000000000000000000000000000000000000000000000
0000000000000000000000000000000000000000000000000000000000000000
0000000000000000000000000000000000000000000000000000000000000000
000000000000000000000000000000011111
>DM_train1316
METQDYAFQPGLTVGELLKSSQKDWQAAINHRFVKELFAGTIENKVLKDYLIQDYHFFDAFL
SMLGACVAHADKLESKLRFAKQLGFLEADEDGYFQKAFKELKVAENDYLEVTLHPVTKAFQD
LMYSAVASSDYAHLLVMLVIAEGLYLDWGSKDLALPEVYIHSEWINLHRGPFFAEWVQFLVD
ELNRVGKNREDLTELQQRWNQAVALELAFFDIGYDV
1100000000000000000000000000000000000000000000000000000000000000
0000000000000000000000000000000000000000000000000000000000000000
0000000000000000000000000000000000000000000000000000000000000000
000000010110000000000000000000000011
>DM_train1320
SVLVLDDRIVDAATKDLYVNGFQEEIQYQNPTPENLQHMFHQGIEILDSARMINVTHLALWK
PSSFKLGNPVDFALDDNYDTFWQSDGGQPHQLDIMFSKRMDICVMAIFFSMIADESYAPSLV
KVYAGHSPSDARFYKMLEVRNVNGWVALRFLDNREDDQLLKCQFIRLLFPVNHENGKDTHLR
GIRLYVPSNEPHQDTHEWAQTLPETNNVFQDAILR
0000000000000000000000011111100000000000000000000000000000000000
0000000000000000000000000000000000000000000000000000000000000000
0000000000000000000000000000000111111110000000000000000000000000
00000000111111111111111111111111111
>DM_train1321
GAMEKFKTLLYDIPIECMEVSEEIISYAKLQLGKKLNDSIYVSLTDHINFAIQRNQKGLDIK
NALLWETKRLYKDEFAIGKEALVMVKNKTGVSLPEDEAGFIALHIVNAELNEEMPNIINITK
VMQEILSIVKYHFKIEFNEESLHYYRFVTHLKFFAQRLFNGTHMESQDDFLLDTVKEKYHRA
YECTKKIQTYIEREYEHKLTSDELLYLTIHIERVV
0000000000000000000000000000000000000000000000000000000000000000
0000000000000000000000000000000000000000000000000000000000000000
0000000000000000001100000000000000000000000000111111111111110000
00000000000000000000000000000000000
>DM_train1322
SLNCFENNYYNLRHPKIEDLRDLIALETLCWSENLQVDNEEIYRRIFKIPQGQFILELEDKI
VGAIYSQRIDNPQLLDNKTCTQVPLLHTESGVVVQLLAVNILPELQNQGLGDRLLEFMLQYC
AQISGVEKVVAVTLCRNYPDYSPMPMAEYIHQKNESGLLVDPLLRFHQIHGAKIEKLLPGYR
PKDWENQTCGVLVSYDIQHRQRFDGATVEKNRQTK
1111111000000000000000000000000000000000000000000000000000000000
0000000000000000000000000000000000000000000000000000000000000000
0000000000000000000000000000000000000000000000000000000000000000
00000000000000000000111111111111111
>DM_train1323

SLKNEELYHQSYDCVCVMFASIPDFKEFYTESDVNKEGLECLRLLNEIIADFDDLLSKPKFS
GVEKIKTIGSTYMAATGLSAIPSQEHAQEPERQYMHIGTMVEFAYALVGKLDAINKHSFNDF
KLRVGINHGPVIAGVIGAQKPQYDIWGNTVNVASRMDSTGVLDKIQVTEETSLILQTLGYTC
TCRGIINVKGKGDLKTYFVNTEMSRSLSQSNLAS
111100000000000000000000000000000000000000000000000000000000
000000000000000000111111111111100000000000000000000000000000
000000000000000000000000000000000000000000000000000000000000
0011111111111110000011111111111111
>DM_train1324
GVLPVEKCNLEDSACMTSAFQQALPTFVAGLPDHGVEVMDVLDLDDFAFDLSGLQFTLKEGK
LKGLKGAVIDNVKWDLKKKNIEVDFHLDATVKGHYTAGGRILILPITGDGQMKLKLKNIHIH
LVVSYEMEKDAEGVDHVIFKKYTVTFDVKDNAQFGLTNLFNGNKELSDTMLTFLNQNWKQVS
EEFGKPVMEAAAKKIFKNIKHFLAKVPIAEIANV
111100000000000000000000000000000000000000000000000000000000
000000000000000000000000000000000000000000000000000000000000
000000000000000000000000000000000000000000000000000000000000
0000000000000000000000000000000000
>DM_train1325
GMDETPIFKIKKLTIAENDRSEYIRYAEKNMHDSIPAEEGTLLIGSGHDDAHGEDNYEIEVF
RNKGAEDLHIAGSHADDFVETVNKIATKQKVIDLHPEVITTKAQRALNSYADNFVMRLIKVE
VKDADAEKFSHAVKKEMTTSMASEPGMEIMMSGTNIDNPNEWYFIEVYANDEAYDIHVKTPH
YKEYIEETDGMVKSRDVKTLVRDTLATQGAIVLD
111000000000000000000000000000000000000000000000000000000000
000000000000000000000000000000000000000000011111100000000000
000000000000000000000000000000000000000000000000000000000000
0000000000000000000000000000000000
>DM_train1328
MGSSHHHHHHMDRYEIKGVDVASYQGDIDWRELEKQNMKFAFIKATEGSAFVDKYFSKNWTN
ANKTSMRVGAYHFFSFDSKGETQAEQFIRNVPKYKQALPPVIDVEFYANKKDNPPKREDVTK
ELSVMIEMLEKHYGKKVILYATQEAYDLYIKDAYPQCDIWIRSVLTKPSLSDERKWTFWQYT
NRGKLSGYNGKEKYIDLNVFYGNEEEFENYGMKD
100000000000000000000000000000000000000000000000000000000000
000000000000000000000000000000000000000000000000000000000000
000000000000000000000000000000000000000000000000000000000000
0000000000000000000000000000000011
>DM_train1329
MDREALLQAVKEARELAKPRNFTQSFEFIATLKEIDMRKPENRIKTEVVLPHGRGKEAKIAV
IGTGDLAKQAEELGLTVIRKEEIEELGKNKRKLRKIAKAHDFFIAQADLMPLIGRYMGVILG
PRGKMPKPVPANANIKPLVERLKKTVVINTRDKPYFQVLVGNEKMTDEQIVDNIEAVLNVVA
KKYEKGLYHIKDAYVKLTMGPAVKVKKEKAKKK
000000000000000000000000000000000000000000000000000000000000
000000000000000000000000000000000000000000000000000000000000
000000000000000000000000000000000000000000000000000000000000
000000000000000000000000001111111

>DM_train1330
MRGSHHHHHHPEYDYIRDGNAIYERSFAIIRAEADLSRFSEEEADLAVRMVHACGSVEATRQ
FVFSPDFVSSARAALKAGAPILCDAEMVAHGVTRARLPAGNEVICTLRDPRTPALAAEIGNT
RSAAALKLWSERLAGSVVAIGNAPTALFFLLEMLRDGAPKPAAILGMPVGFVGAAESKDALA
ENSYGVPFAIVRGRLGGSAMTAAALNSLARPGL
11111111110000000000000000000000000000000000000000000000000000
00000000000000000000000000000000000000000000000000000000000000
00000000000000000000000000000000000000000000000000000000000000
000000000000000000000000000000000
>DM_train1331
KGHLTRLGLEFFDQPAVPLARAFLGQVLVRRLPNGTELRGRIVETEAYLGPEDEAAHSRGGR
QTPRNRGMFMKPGTLYVYIIYGMYFCMNISSQGDGACVLLRALEPLEGLETMRQLRSTLRKG
TASRVLKDRELCSGPSKLCQALAINKSFDQRDLAQDEAVWLERGPLEPSEPAVVAAARVGVG
HAGEWARKPLRFYVRGSPWVSVVDRVAEQDTQA
00000000000000000000000000000000000000000000000000000000000000
00000000000000000000000000000000000000000000000000000000001111
11110000000000000000000000000000000000000111111000000000000000
000000000000000000000000000001111
>DM_train1333
ETMSGPLHIGLIPTVGPYLLPHIIPMLHQTFPKLEMYLHEAQTHQLLAQLDSGKLDAVILAL
VKESEAFIEVPLFDEPMLLAIYEDHPWANREAVPMADLAGEKLLMLEDGHSLRDQAMGFCFE
AGADEDTHFRATSLETLRNMVAAGSGITLLPALAVPPERKRDGVVYLPAIKPEPRRTIGLVY
RPGSPLRSRYEQLAEAIRARMDGHFDKVLKQAV
00000000000000000000000000000000000000000000000000000000000000
00000000000000000000000000000000000000000000000000000000000001
11111000000000000000000000000000000000000000000000000000000000
000000000000000000000000001111111
>DM_train1334
ASHHLRMHFKTLPAGESLGSLGLWVWGDVDQPSKDWPNGAITMTKAKKDDYGYYLDVPLAAK
HRQQVSYLINNKAGENLSKDQHISLLTPKMNEVWIDENYHAHAYRPLKEGYLRINYHNQSGH
YDNLAVWTFKDVKTPTTDWPNGLDLSHKGHYGAYVDVPLKEGANEIGFLILDKSKTGDAIKV
QPKDYLFKELDNHTQVFVKDTDPKVYNNPYYID
10000000000000000000000000000000000000000000000000000000000000
00000000000000000000000000000000000000000000000000000000000000
00000000000000000000000000000000000000000000000000000000000000
000000000000000000000000000000000
>DM_train1335
GSHMGIKQKSYQPSIIIAGPQNSGKTSLLTLLTTDSVRPTVVSQEPLSAADYDGSGVTLVDF
PGHVKLRYKLSDYLKTRAKFVKGLIFMVDSTVDPKKLTTTAEFLVDILSITESSCENGIDIL
IACNKSELFTARPPSKIKDALESEIQKVIERRKKSLNEVERKINEEDYAENTLDVLQSTDGF
KFANLEASVVAFEGSINKRKISQWREWIDEKL
11111111100000000000000000000000000000000000000000000000000000
00000000000000000000000000000000000000000000000000000000000000
00000000000000000000000000000000000000111111111111110000111100

00000000000000000000000000000000
>DM_train1337
HTPVRTVMNTIQQLMMILNSASDQPSENLISYFNNCTVNPKESILKRVKDIGYIFKEKFAKA
VGQGCVEIGSQRYKLGVRLYYRVMESMLKSEEERLSIQNFSKLLNDNIFHMSLLACALEVVM
ATYSRSTSQNLDSGTDLSFPWILNVLNLKAFDFYKVIESFIKAEGNLTREMIKHLERCEHRI
MESLAWLSDSPLFDLIKQSKDREGPTDHLESA
11111111000000000000000000000000000000000000000000000000000000
00000000000000000000000000000001111111000000000000000000000000
00001111111111111100000000000000000000000000000000000000000000
00000000000000000000111111111111
>DM_train1338
MKEVAKPLQERFFIPHEIKVMGRWSTEDVEVKDPSLKPYINLEPRLLPHTHGRHAKKHFGKA
NVHIVERLINKVMRSGGSHYKVAGHFMRREHRSLNSKKVRAYEVVKEAFKIIEKRTGKNPIQ
VLVWAIENAAPREDTTSVMFGGIRYHVAVDISPLRRLDVALRNIALGASAKCYRTKMSFAEA
LAEEIILAANKDPKSYAYSKKLEIERIAESSR
11111111111111111000000000000000000000000000011111111111111111
10000000000000000011111111111111111000000000000000000000000000
00000000000000000000000000000000000000000000000000000000000000
00000000000000000000000000000000
>DM_train1339
QNPADSPHIGKVFFSTNQGDFVCSANIVASANQSTVATAGHCLHDGNGGQFARNFVFAPAYD
YGESEHGVWAAEELVTSAEWANRGDFEHDYAFAVLETKGGTTVQQQVGTASPIAFNQPRGQY
YSAYGYPAAAPFNGQELHSCHGTATNDPMGSSTQGIPCNMTGGSSGGPWFLGNGTGGAQNST
NSYGYTFLPNVMFGPYFGSGAQQNYNYASTTN
11000000000000000000000000000000000000000000000000000000000000
00000000000000000000000000000000000000000000000000000000000000
00000000000000000000000000000000000000000000000000000000000000
00000000000000000000000000000000
>DM_train1340
EKVKKIVLIGASGFVGSALLNEALNRGFEVTAVVRHPEKIKIENEHLKVKKADVSSLDEVCE
VCKGADAVISAFNPGWNNPDIYDETIKVYLTIIDGVKKAGVNRFLMVGGAGSLFIAPGLRLM
DSGEVPENILPGVKALGEFYLNFLMKEKEIDWVFFSPAADMRPGVRTGRYRLGKDDMIVDIV
GNSHISVEDYAAAMIDELEHPKHHQERFTIGY
10000000000000000000000000000000000000000000000000000000000000
00000000000000111111000000000000000000000000000000000000000000
00000000000000000000000000000000000000000000000000000000000000
00000000000000000000000000000000
>DM_train1341
MGSDKIHHHHHHMNIFEAIENRHSVRDFLERKMPERVKDDIENLLVKFITKKLDWKINLSSF
PSYIYAKAEKHFDELVEYGFQGEQIVLFLTAQGFGTCWMARSPHPDVPYIIVFGYPRTRNFT
RKRRPITSFLENDLEELPPEIVKIVEMTILAPSALNRQPWKIKYTGGELCISSERPVDLGIA
LSHAYLTAREIFKREPVIQKRGEDTYCLILNP
10000000000000000000000000000000000000000000000000000000000000
00000000000000000000000000000000000000000000000000000000000000

0000000000000000000000000000000000000000000000000000000000000000
000000000000000000000000000000000
>DM_train1342
SLRGGHAGTTYIFSKGGGQITYKWPPNDRPSTRADRLAIGFSTVQKEAVLVRVDSSSGLGDY
LELHIHQGKIGVKFNVGTDDIAIEESNAIINDGKYHVVRFTRSGGNATLQVDSWPVIERYPA
GNNDNERLAIARQRIPYRLGRVVDEWLLDKGRQLTIFNSQATIIGGKEQGQPFQGQLSGLY
YNGLKVLNMAAENDANIAIVGNVRLVGEVPSS
1110000000000000000000000000000000000000000000000000000000000000
0000000000000000000000000000000000000000000000000000000000000011
1111111111111110000000000000000000000000000000000000000000000000
000000000000000000000000000000011
>DM_train1343
MISLKRVAALLCVAGLGMSAANAQTCLTSSQTGTNNGFYYSFWKDSPGTVNFCLQSGGRYTS
NWSGINNWVGGKGWQTGSRRNITYSGSFNSPGNGYLALYGWTTNPLVEYYVVDSWGSWRPPG
SDGTFLGTVNSDGGTYDIYRAQRVNAPSIIGNATFYQYWSVRQSKRVGGTITTGNHFDAWAS
VGLNLGTHNYQIMATEGYQSSGSSDITVSEGG
1111111111111111111111000000000000000000000000000000000000000000
0000000000000000000000000000000000000000000000000000000000000000
0000000000000000000000000000000000000000000000000000000000000000
000000000000000000000000000000001
>DM_train1344
STLEIAGLVRKNLVQFGVGEKNGSVRWVMNALGVKDDWLLVPSHAYKFEKDYEMMEFYFNRG
GTYYSISAGNVVIQSLDVGAQDVVLMKVPTIPKFRDITQHFIKKGDVPRALNRLATLVTTVN
GTPMLISEGPLKMEEKATYVHKKNDGTTVDLTVDQAWRGKGEGLPGMCGGALVSSNQSIQNA
ILGIHVAGGNSILVAKLVTQEMFQNIDKKIE
0000000000000000000000000000000000000000000000000000000000000000
0000000000000000000000000000000000000000000000000000000000000000
0000000000000000000000000000000000000000000000000000000000000000
0000000000000000000000000001111
>DM_train1345
SRLDKSKVINSALELLNEVGIEGLTTRKLAQKLGVEQPTLYWHVKNKRALLDALAVEILARH
HDYSLPAAGESWQSFLRNNAMSFRRALLRYRDGAKVHLGTRPDEKQYDTVETQLRFMTENGF
SLRDGLYAISAVSHFTLGAVLEQQEHTAALTDRPAAPDENLPPLLREALQIMDSDDGEQAFL
HGLESLIRGFEVQLTALLQIVGGDKLIIPFC
0000000000000000000000000000000000000000000000000000000000000000
0000000000000000000000000000000000000000000000000000000000000000
0000000000000000000000000001111111111111100000000000000000000000
0000000000000000000001111111111
>DM_train1346
YVEFAQDFDFFYFVQQWPASYCDTRRSCCYPTTGKPDEDFSIHGLWPNYENGKWPQNCDRES
SLDESEISDLISTMEKNWPSLACPSSDGVRFWSHEWLKHGTCSALGERAYFQAALDFRKKSN
LLENLKNAEITPRNGEHYTLESIKKAIEEGVGHSPYIECNVDTQGNHQIYQVYLCVDKTATD
FIDCPIFPHGRGCGSKIEFPPFSSESDHDEF
1111000000000000000000000000000000000000000000000000000000000000

000000000000000000000000000000000000000000000000000000000000
000000000000000000000000000000000000000000000000000000000000
0000000000000000000000111111111
>DM_train1348
MEQPYVYAYPQGSGPSGAPTPQAGGVVVDPKYCAPYPIDMAIVRKMMSLTDGNFVITDVNGN
LLFKVKEPVFGLHDKRVLLDGSGTPVVTLREKMVSMHDRWQVFRGGSTDQRDLLYTVKRSSM
LQLKTKLDVFLGHNKDEKRCDFRVKGSWLERSCVVYAGESDAIVAQMHRKHTVQSVFLGKDN
FSVTVYPNVDYAFIASLVVILDDVNREDRAA
111111111111111111111110000000000000000000011111000000000000
000000000000000000000000000001111110000000000000000000000111
111100000000000010000000000000000000000000000000111111110000
0000000000000000000000000011111
>DM_train1349
GEINGPESVLVNTYQWIIRNWETVKIQWSQNPTMLYNKMEFEPFQSLVPKAIRGQYSGFVRT
LFQQMRDVLGTFDTTQIIKLLPFAAAPPKQSRMQFSSLTVNVRGSGMRILVRGNSPVFNYNK
TTKRLTILGKDAGTLIEDPDESTSGVESAVLRGFLILGKEDRRYGPALSINELSNLAKGEKA
NVLIGQGDVVLVMKRKRDSSILTDSQTATKR
100000000000000000000000000000000000000000000000000000000000
000000000000000000000000000000000000000000000000000000000000
000000000000000001111111100000000000000000000000000000000000
0000000000000000000111111111111
>DM_train1350
MPYSDAKAMLDEVAKIRELGVQRVTRIENLENAKKLWDNANSMLEKGNISGYLKAANELHKF
MKEKNLKEDDLRPELSDKTISPKGYAILQSLWGAASDYSRAAATLTESTVEPGLVSAVNKMS
AFFMDCKLSPNERATPDPDFKVGKSKILVGIMQFIKDVADPTSKIWMHNTKALMNHKIAAIQ
KLERSNNVNDETLESVLSSKGENLSEYLSYK
111111111111111110000000000000000000000000000000000000000000
000000000000000000000000000000000000000000000000000000000000
000000000000000000000000000000000000000000000000000000000000
0000000000000000000000000000000
>DM_train1351
RSIIGGKHGDRDVRILYQVGDSEEDLPVCAPNAVCSKIDLYETPWIERQCRCPDGRTCPSSL
GVEDGHTIADKTRHYKMCQPVHKLPVCKHFRDYTWTLTTAAELNVTEQIVHCRCPRNSVTYL
TKREPIGNDSPGYRYLFACSPLTRLRCQRKQPCKLFTVRKRQEFLDEVNINSLCQCPKGHRC
PSHHTQSGVIAGESFLEDNIQTYSGYCMAND
111111111000000000000000000000000000000000000000000000000000
000000000000000000000000000000000000000000000000000000000000
000000000000000000000000000000000000000000000000000000000000
0000000000000000000000000000001
>DM_train1353
EVTIEAVPPQVAEDNNVLLLVHNLPLALGAFAWYKGNTTAIDKEIARFVPNSNMNFTGQAYS
GREIIYSNGSLLFQMITMKDMGVYTLDMTDENYRRTQATVRFHVHQPVTQPFLQVTNTTVKE
LDSVTLTCLSNDIGANIQWLFNSQSLQLTERMTLSQNNSILRIDPIKREDAGEYQCEISNPV
SVRRSNSIKLDIIFDPSRLVPRGSHHHHHH

000000000000000000000000000000000000000000000000000000000000
000000000000000000000000000000000000000000000000000000000000
000000000000000000000000000000000000000000000000000000000000
000000000000000001111111111111
>DM_train1354
VVKQDLSDNQYIQKKPNEITSNELLLPLPNDKLLGDPKAPILMIEYASLTCYHCSLFHRNVF
PKIKEKYIDTGKMLYIFRHFPLDYRGLKAAMLSHCYEKQEDYFNFNKAVFNSIDSWNYYNLS
DLTLLQRIAALSNLKQDAFNQCINDKKIMDKIVNDKSLAINKLGITAVPIFFIKLNDDKSYI
EHNKVKHGGYKELKYFTNVIDKLYGKAIVK
111111111111111111100000000000000000000000000000000000000000
000000000000000000000000000000000000000000000000000000000000
000000000000000000000000000000000000000000000000000000000000
000000000000000000000000000000
>DM_train1355
GHMAMEKPVLWYIADPMCSWCWGFAPVIENIRQEYSAFLTVKIMPGGLRPGTNTPLLPEKRA
QILHHWHSVHITTGQPFTFENALPEGFIYDTEPACRGVVSVSLIEPEKVFPFFAAIQRAFYV
GQEDVAQLAILKKLAVDLGIPESRFTPVFQSDEAKQRTLAGFQRVAQWGISGFPALVVESGT
DRYLITTGYRPIEALRQLLDTWLQQHGVGS
111110000000000000000000000000000000000000001111000000000000
000000000000000000000000000000000000000000000000000000000000
000000000000000000000000000000000000000000000000000000000000
000000000000000000000000000111
>DM_train1356
VSVGTHPTFENSPSGTVLTSPPDGSAVDRATDAARRVVDALLRTDRGNANLERVAEELNSIA
GHLEEHAPAVAERLIDMWNGEGVTRHDPVTGPENALAPPVVLEGLSDGSVRGTVTLTIPYQG
PPGHVHGGVSALLLDHVLGVANAWGGKAGMTAQLSTRYHRPTPLFEPLTLTGKLMSVDGRKI
TTAGDIRTADGQVCVSVEGLFVDKTVPRPR
111111111111110000000010000000000000000000000000000000000000
000000000000000000000000000000000000000000000000000000000000
000000000000000000000000000000000000000000000000000000000000
000000000000000000000011111111
>DM_train1357
MAKRILCFGDSLTWGWVPVEDGAPTERFAPDVRWTGVLAQQLGADFEVIEEGLSARTTNIDD
PTDPRLNGASYLPSCLATHLPLDLVIIMLGTNDTKAYFRRTPLDIALGMSVLVTQVLTSAGG
VGTTYPAPKVLVVSPPPLAPMPHPWFQLIFEGGEQKTTELARVYSALASFMKVPFFDAGSVI
STDGVDGIHFTEANNRDLGVALAEQVRSLL
100000000000000000000000000000000000000000000000000000000000
000000000000000000000000000000000000000000000000000000000000
000000000000000000000000000000000000000000000000000000000000
000000000000000000000000000000
>DM_train1358
MDQENERNISRLWRAFRTVKEMVKDRGYFITQEEVELPLEDFKAKYCDSMGRPQRKMMSFQA
NPTEESISKFPDMGSLWVEFCDEPSVGVKTMKTFVIHIQEKNFQTGIFVYQNNITPSAMKLV
PSIPPATIETFNEAALVVNITHHELVPKHIRLSSDEKRELLKRYRLKESQLPRIQRADPVAL

YLGLKRGEVVKIIRKSETSGRYASYRICM
111100000000000000000000000000000000000000000000000000000000
000000000000000000000000000000000000000000000000000000000000
000000000000000000000000000000000000000000000000000000000000
00000000000000000000000000000
>DM_train1359
MSSSEEVSWISWFCGLRGNEFFCEVDEDYIQDKFNLTGLNEQVPHYRQALDMILDLEPDEEL
EDNPNQSDLIEQAAEMLYGLIHARYILTNRGIAQMLEKYQQGDFGYCPRVYCENQPMLPIGL
SDIPGEAMVKLYCPKCMDVYTPKSSRHHHTDGAYFGTGFPHMLFMVHPEYRPKRPANQFVPR
LYGFKIHPMAYQLQLQAASNFKSPVKTIR
111110000000000000000000000000000000000000000000000000000000
000000000000000000000000000000000000000000000000000000000000
000000000000000000000000000000000000000000000000000000000000
00000000000000000001111111111
>DM_train1360
MKTVLLTGFDPFGGESINPAWEVAKSLHEKTIGEYKIISKQVPTVFHKSISVLKEYIEELAP
EFIICIGQAGGRPDITIERVAINIDDARIADNEGNQPVDVPVVEEGPAAYWSTLPMKAIVKK
LQEEGIPASVSQTAGTFVCNHLFYGLMHELEKHDTKMKGGFIHIPFLPEQASNYPGQPSMSL
STIRKGIELAVEVTTTVEVDIVEVGGTTH
000000000000000000000000000000000000000000000000000000000000
000000000000000000000000000000000000000000000000000000000000
000000000000000000000000000000000000000000000000000000000000
00000000000000000011111111111
>DM_train1361
PQLEDFPALIKQASLDALFKCGKDAEALKEVFTNSNNVAGKKAIMEFAGLFRSALNATSDSP
EAKTLLMKVGAEYTAQIIKDGLKEKSAFGPWLPETKKAEAKLENLEKQLLDIIKNNTGGELS
KLSTNLVMQEVMPYIASCIEHNFGCTLDPLTRSNLTHLVDKAAAKAVEALDMCHQKLTQEQG
TSVGREARHLEMQTLIPLLLRNVFAQIPA
000000000000000000000000000000000000000000000000000000000000
000000000000000000000000000000000000000000000000000000000000
000000000000000000000000000000000000000000000000000000111111
11111111000000000000000000000
>DM_train1362
GHMVPQTGRAARATRESGRQTRDALFTAATELFLEHGEGVPITQICAAAGAHPNQVTYYYGS
KERLFVEVACAAVLRAGKRAEDDAATAETVGDYTEKLVGSLLGPGAPSVELFTSAMLMTGRR
SELRDLITDTLRTLHSSGEVALIRTLMRTGWQLRAGIDVESKAFWSAIFGLVIQKTATGESF
GYSLEEAVAVIFANLQIPETVRNTSIDGS
111111111111111111110000000000000000000000000000000000000000
000000000000000000000000000000000000000000000000000000000000
000000000000000000000000000000000000000000000000000000011111
11000000000000000000000000111
>DM_train1363
GSGYREVEYWDQRYQGAADSAPYDWFGDFSSFRALLEPELRPEDRILVLGCGNSALSYELFL
GGFPNVTSVDYSSVVVAAMQACYAHVPQLRWETMDVRKLDFPSASFDVVLEKGTLDALLAGE

RDPWTVSSEGVHTVDQVLSEVSRVLVPGGRFISMTSAAPHFRTRHYAQAYYGWSLRHATYGS
GFHFHLYLMHKGGKLSVAQLALGAQILSP
000000000000000000000000000000000000000000000000000000000000
000000000000000000000000000000000000000000000000000000000000
000000000000000000000000000000000000000000000000000000000000
00000000000000000000000000011
>DM_train1364
MEIKGESMQKSKFRRICVFCGSSQGKKSSYQDAAVDLGNELVSRNIDLVYGGGSIGLMGLVS
QAVHDGGRHVIGIIPKTLMPRELTGETVGEVRAVADMHQRKAEMAKHSDAFIALPGGYGTLE
ELLEVITWAQLGIHDKPVGLLNVDGYYNSLLSFIDKAVEEGFISPTAREIIVSAPTAKELVK
KLEEYAPCHERVATKLCWEMERIGYSSEE
111111111000000000000000000000000000000000000000000000000000
000000000000000011111111000000000000000000000000000000000000
000000000000000000000000000000000000000000000000000000000000
00001111111111111111111111111
>DM_train1365
IQHATASLIQTNTDQELLVVDVTPSFASLWLVPNINDFHQRHPNIRVKILTGDGAVKNIHGE
SDLHVRCLPLSTHYEYSQLLCEETLLLIGNTNLPKLSDNQAISHYPFIPQTTRPQLWEQFKQ
ENDLECPITYHSVGFEHFYLACEAVRMEKGLALLPDFMAQFSILRGDIQHIGNLKLHSGYGY
YVVIPNFRLTSRKVALFHDWLKDKLTHHT
111111111111111000000000000000000000000000000000000000111110
000000000000000000000000000000000111111100000000000000000000
001111100000000000000000000000000000000000000000000000000000
00000000000000000000000000111
>DM_train1366
MNKDQSKIPQATAKRLPLYYRFLKNLHASGKQRVSSAELSDAVKVDSATIRRDFSYFGALGK
KGYGYNVDYLLSFFRKTLDQDEMTDVILIGVGNLGTAFLHYNFTKNNNTKISMAFDINESKI
GTEVGGVPVYNLDDLEQHVKDESVAILTVPAVAAQSITDRLVALGIKGILNFTPARLNVPEH
IRIHHIDLAVELQSLVYFLKHYSVLEEIE
111111111100000000000000000000000000000000000000000000000011
111000000000000000001110000000000000000011111000000000000000
000000000000000000000000000000000000000000000000000000000000
00000000000000000000000111111
>DM_train1367
MAQGLIEVERKFLPGPGTEERLQELGGTLEYRVTFRDTYYDTPELSLMQADHWLRRREDSGW
ELKCPGAAGVLGPHTEYKELTAEPTIVAQLCKVLRADGLGAGDVAAVLGPLGLQEVASFVTK
RSAWKLVLLGADEEEPQLRVDLDTADFGYAVGEVEALVHEEAEVPTALEKIHRLSSMLGVPA
QETAPAKLIVYLQRFRPQDYQRLLEVNSS
110000000000000000000000000000000000000000000000000000000000
000000000000000000000000000000000000000000000000000000000000
000000000000000000000000000000000000000000000000000000000000
00000000000000000000001111111
>DM_train1368
MGSDKIHHHHHHMPKVEIAPSEIKIPDNVLKAKLGFGGAEEIPEEFRKTVNRAYEELLDAAK

PVVLWRDFEVDGSLSFDDMRLTGELATKHLSGSKIITVFLATLGKKVDEKIEEYFRKGEDLL
AFFIDGIASEMVEYALRKVDAELRMKRSNLEGSFRISPGYGDLPLSLNKKIAEIFKEEVDVN
VIEDSYVLVPRKTITAFVGWREKNEKQT
111111111100000000000000000000000000000000000000000000000000
000000000000000000000000000000000000000000000000000000000000
000000000000000000000000000000000000000000000000000000000000
0000000000000000000001111111
>DM_train1369
HGTELPSPPSVWFEAEFFHHILHWTPIPQQSESTCYEVALLRYGIESWNSISQCSQTLSYDL
TAVTLDLYHSNGYRARVRAVDGSRHSQWTVTNTRFSVDEVTLTVGSVNLEIHNGFILGKIQL
PRPKMAPAQDTYESIFSHFREYEIAIRKVPGQFTFTHKKVKHEQFSLLTSGEVGEFCVQVKP
SVASRSNKGMWSKEECISLTRQYFTVTN
000000000000000000000000000000000000000000000000000000000000
000000000000000000000000000000000000000000000000000000000000
000000000000000000000000000000000000000000000000000000000000
0000000000000000000011111111
>DM_train1370
MLIAIEGVDGAGKRTLVEKLSGAFRAAGRSVATLAFPRYGQSVAADIAAEALHGEHGDLASS
VYAMATLFALDRAGAVHTIQGLCRGYDVVILDRYVASNAAYSAARLHENAAGKAAAWVQRIE
FARLGLPKPDWQVLLAVSAELAGERSRGRAQRDPGRARDNYERDAELQQRTGAVYAELAAQG
WGGRWLVVGADVDPGRLAATLAPPDVPS
000000000000000000000000000000000000000000000000000000000000
000000000000000000000000000000000000000000000000000000000000
000000000000000000000000011111111111111110000000000000000000
0000000000000000000000011111
>DM_train1371
SNAMAVQLLENWLLKEQEKIQTKYRHLNHISVVEPNILFIGDSIVEYYPLQELFGTSKTIVN
RGIRGYQTGLLLENLDAHLYGGAVDKIFLLIGTNDIGKDVPVNEALNNLEAIIQSVARDYPL
TEIKLLSILPVNEREEYQQAVYIRSNEKIQNWNQAYQELASAYMQVEFVPVFDCLTDQAGQL
KKEYTTDGLHLSIAGYQALSKSLKDYLY
111000000000000000000000000000000000000000000000000000000000
000000000000000000000000000000000000000000000000000000000000
000000000000000000000000000000000000000000000000000000000000
0000000000000000000000000000
>DM_train1372
GMSIVGNVENLINGVGELWNKYVKHEFILKMRDGSLPLDIFRYYLIQDGKYVEDMLRALLIA
SSKGPIDKVTKILNLVFSSRDKGLETHGKLYSKLDISRDVIVKTGYNLINYAYTRHLYYYAN
LDWNKFLVAWTPCMFGYSIVGDYVIDSPNEVYKTWASFYASTEYKKRIEAILYALDEVSITE
DLLNIFINSVRFEIGFWDASLRKDPTVY
111110000000000000000000000000000000000000000000000000000000
000000000000000000011111000000000000000000000000000000000000
000000000000000000000000000000000000000000000000000000000000
0000000000000000000000000000
>DM_train1373

GSHMKITYPLPPNLPEQLPLLTNCQLEDEAILENHLYQQIDLPNQEVRNLVFRDAVFDHLSL
ANGQFASFDCSNVRFEACDFSNVEWLSGSFHRVTFLRCNLTGTNFADSYLKDCLFEDCKADY
ASFRFANFNLVHFNQTRLVESEFFEVTWKKLLLEACDLTESNWLNTSLKGLDFSQNTFERLT
FSPNYLSGLKVTPEQAIYLASALGLVIT
11100000000111100000000000000000000000000000000000000000000000
00000000000000000000000000000000000000000000000000000000000000
00000000000000000000000000000000000000000000000000000000000000
0000000000000000000000000000
>DM_train1374
NSNMGSDSGTMGSDSDEENVAARASPEPELQLRPYQMEVAQPALEGKNIIICLPTGSGKTRV
AVYIAKDHLDKKKKASEPGKVIVLVNKVLLVEQLFRKEFQPFLKKWYRVIGLSGDTQLKISF
PEVVKSCDIIISTAQILENSLLNLENGEDAGVQLSDFSLIIIDECHHTNKEAVYNNIMRHYL
MQKLKNNRLKKENKPVIPLPQILGLTAS
11111111111111100000000000000000000000000000000000000000000000
00000000000000000000000000000000000000000000000000000011100000
00000000000000000000000111111100000000000000011111110000000000
0000000000000000000000000000
>DM_train1377
MEVEKEFITDEAKELLSKDKLIQQAYNEVKTSICSPIWPATSKTFTINNTEKNCNGVVPIKE
LCYTLLEDTYNWYREKPLDILKLEKKKGGPIDVYKEFIENSELKRVGMEFETGNISSAHRSM
NKLLLGLKHGEIDLAIILMPIKQLAYYLTDRVTNFEELEPYFELTEGQPFIFIGFNAEAYNS
NVPLIPKGSDGMSKRSIKKWKDKVENK
00000000000000000000000000000000000000000000000000000000000000
00000000000000000000000000000000000000000000000000000000000000
00000000000000000000000000000000000000000000000000000000000000
000000000000111111111111111
>DM_train1378
ADGKSTRYWDCCKPSCGWAKKAPVNQPVFSCNANFQRLTDFDAKSGCEPGGVAYSCADQTPW
AVNDDFAFGFAATSIAGSNEAGWCCACYELTFTSGPVAGKKMVVQSTSTGGDLGSNHFDLNI
PGGGVGIFDGCTPQFGGLPGQRYGGISSRNECDRFPDALKPGCYWRFDWFKNADNPSFSFRQ
VQCPAELVARTGCRRNDDGNFPAVQIP
00000000000000000000000000000000000000000000000000000000000000
00000000000000000000000000000000000000000000000001111100000000
00000000000000000000000000000000000000000000000000000000000000
000000000000000000000000111
>DM_train1379
MFTGIVQGTAKLVSIDEKPNFRTHVVELPDHMLDGLETGASVAHNGCCLTVTEINGNHVSFD
LMKETLRITNLGDLKVGDWVNVERAAKFSDEIGGHLMSGHIMTTAEVAKILTSENNRQIWFK
VQDSQLMKYILYKGFIGIDGISLTVGEVTPTRFCVHLIPETLERTTLGKKKLGARVNIEIDP
QTQAVVDTVERVLAARENAMNQPGTEA
00000000000000000000000000000000000000000000000000000000000000
00000000000000000000000000001111110000000000000011111111110000
00000000000000000000000000000000000000000000000000000000000000
000000000000000011111111111

>DM_train1380
MERKISRIHLVSEPSITHFLQVSWEKTLESGFVITLTDGHSAWTGTVSESEISQEADDMAME
KGKYVGELRKALLSGAGPADVYTFNFSKESAYFFFEKNLKDVSFRLGSFNLEKVENPAEVIR
ELIAYALDTIAENQAKNEHLQKENERLLRDWNDVQGRFEKAVSAKEALETDLYKRFILVLNE
KKTKIRSLHNKLLNAAQEREKDIKQEG
00000000000000000000000000000000000000000000000000000000000000
00000000000000111111000000000000000000000000000000000000000000
00000000000000000000000000000000000000000000000000000000000000
000000000000000011111111111
>DM_train1381
VSSQKTSSLPKYTPKVNSSINNYIRKKNMKAPRIEEDYTSYFPKYGYRNGVGRPEGIVVHDT
ANDNSTIDGEIAFMKRNYTNAFVHAFVDGNRIIETAPTDYLSWGAGPYGNQRFINVEIVHTH
DYDSFARSMNNYADYAATQLQYYNLKPDSAENDGRGTVWTHAAISNFLGGTDHADPHQYLRS
HNYSYAELYDLIYEKYLIKTKQVAPWG
11111100000000000000000000000000000000000000000000000000000000
00000000000000000000000000000000000000000000000000000000000000
00000000000000000000000000000000000000000000000000000000000000
000000000000000000000000000
>DM_train1382
MGSDKIHHHHHHMTPVLSLDMEDPIRFIDENGSFEVVKVGHNLAIHGKKIFDELAKRNLKII
LDLKFCDIPSTVERSIKSWDHPAIIGFTVHSCAGYESVERALSATDKHVFVVVKLTSMEGSL
EDYMDRIEKLNKLGCDFVLPGPWAKALREKIKGKILVPGIRMEVKADDQKDVVTLEEMKGIA
NFAVLGREIYLSENPREKIKRIKEMRL
11111111110000000000000000000000000000000000000000000000000000
00000000000000000000000000000000000000000000000000000000000000
00000000000000000000000000000000000000000111111111000000000000
000000000000000000000000111
>DM_train1384
GSECSTQLDIVIVLDGSNSIYPWDSVTAFLNDLLKRMDIGPKQTQVGIVQYGENVTHEFNLN
KYSSTEEVLVAAKKIVQRGGRQTMTALGTDTARKEAFTEARGARRGVKKVMVIVTDGESHDN
HRLKKVIQDCEDENIQRFSIAILGSYNRGNLSTEKFVEEIKSIASEPTEKHFFNVSDELALV
TIVKTLGERIFALEATAVDSSGRIVTD
11111100000000000000000000000000000000000000000000000000000000
00000000000000000000000000000000000000000000000000000000000000
00000000000000000000000000000000000000000000000000000000000000
000000000000111111111111111
>DM_train1385
KLNMLKPDSDLCLKFAMLCTLNDKCDRLRKAYGEACSGPHCQRHVCLRQLLTFFEKAAEPHA
QGLLLCPCAPNDRGCGERRRNTIAPNCALPPVAPNCLELRRLCFSDPLCRSRLVDFQTHCHP
MDILGTCATEQSRCLRAYLGLIGTAMTPNFVSNVNTSVALSCTCRGSGNLQEECEMLEGFFS
HNPCLTEAIAAKMRFHSQLFSQDWPHP
11111111000000000000000000000000000000000000000000000000000000
00000000000000000000000000000000000000000000000000000000000000
00000000000000000000000000000000000000000000000000000000000000

000000000000000000000111111
>DM_train1386
SLIGFLSDRQGFEKQYHDIDLSSFGIREGISEIIASTGFEHPNAAPIGIVMKGERPFVRLFK
GSHTWENVLKEKCLASNVVYDPILFVRSTFSDLVPSEFEYVDAGEFKFPVLKEAIAWVVFEC
INLRNTDQSLVADLVPLNAGFNERNIKELPVPNRGFNAVLEATVHATRYQLTGEEKYLELIR
HYESLASKCGGDAEKKAMKLIYEALGL
111111111111111111100000000000000000000000000000000000000000
000000000000000000000000000001100000000000000000000000000000
000000110000000000000000000000000000000000000000011111100000
000000000000000000000000011
>DM_train1387
ASMTTPCITILSGHFPKETIYARKTKELVEEYCSIHGYNFYYEESEPLETEEHALHFRRSWI
IQQAAEKFPSTEWFLWLDSDVYVNPKNKNKPITSFIDLSDPNILYHTFHEAPWGSYPINTGV
KFVHKDALEIEKIVWSLRNEAPWNTFPYEQKTVYEYVFPRIPGRYIVHDPYTLNCIVKAYPE
HVKDALFVHMCGTSRAERDEHMEMVAT
111110000000000000000000000000000000000000000000000000000000
000000000000000000000000000000000000000000000000000000000000
000000000000000000000000000000000000000000000000000000000000
000000000000000000000000011
>DM_train1388
GMSSDVQKGDNLETPGARDLLLQTASNIMREGDVVDISLSELSLRSGLNSALVKYYFGNKAG
LLKALLDRDMENIVKSVDALLAKDDMSPEAKLRRHISKCIDTYYDYPYLNRLLMRLVRDSDE
AEAKRIADQYLLPLHRAYNRFIGEGVKAGVFRPINPQLFYFTVTGAADRFFSARLVLKHCFD
QDTLTEQLRDSYREHTVDFIMAGILAH
111111111111111000000000000000000000000000000000000000000000
000000000000000000000000000000000000000000000000000000000000
000000000000000000000000000000000000000000000000000000000000
000000000000000000000000001
>DM_train1390
KGSVYTAQSDVQVPENESIKLTCTYSGFSSPRVEWKFVQGSTTALVCYNSQITAPYADRVTF
SSSGITFSSVTRKDNGEYTCMVSEEGGQNYGEVSIHLTVLVPPSKPTISVPSSVTIGNRAVL
TCSEHDGSPPSEYSWFKDGISMLTADAKKTRAFMNSSFTIDPKSGDLIFDPVTAFDSGEYYC
QAQNGYGTAMRSEAAHMDAVELNVGG
000000000000000000000000000000000000000000000000000000000000
000000000000000000000000000000000000000000000000000000000000
000000000000000000000000111110000000000000000000000000000000
00000000000000000000000000
>DM_train1391
MKPPSSIQTSEFDSSDEEPIEDEQTPIHISWLSLSRVNCSQFLGLCALPGCKFKDVRRNVQK
DTEELKSCGIQDIFVFCTRGELSKYRVPNLLDLYQQCGIITHHHPIADGGTPDIASCCEIME
ELTTCLKNYRKTLIHSYGGLGRSCLVAACLLLYLSDTISPEQAIDSLRDLRGSGAIQTIKQY
NYLHEFRDKLAAHLSSRDSQSRSVSR
111111111111111111111111000000000000000000000000000000000000
000000000000000000000000000000000000000000000000000000000000

000000000000000000000000000000000000000000000000000000000000
00000000000000111111111111
>DM_train1392
MSGTEEAILGGRDSHPAAGGGSVLCFGQCQYTAEEYQAIQKALRQRLGPEYISSRMAGGGQK
VCYIEGHRVINLANEMFGYNGWAHSITQQNVDFVDLNNGKFYVGVCAFVRVQLKDGSYHEDV
GYGVSEGLKSKALSLEKARKEAVTDGLKRALRSFGNALGNCILDKDYLRSLNKLPRQLPLEV
DLTKAKRQDLEPSVEEARYNSCRPNM
111111111111111111111111000000000000000000000000000000000000
000000000000000000000000000000000000000000000000000000000000
000000000000000000000000000000000000000000000000000000000000
00000000000000000000001111
>DM_train1393
MVGGGGVKFIEMDIRDKEAYELAKEWFDEVVVSIKFNEEVDKEKLREARKEYGKVAILLSNP
KPSLVRDTVQKFKSYLIYVESNDLRVIRYSIEKGVDAIISPWVNRKDPGIDHVLAKLMVKKN
VALGFSLRPLLYSNPYERANLLRFMMKAWKLVEKYKVRRFLTSSAQEKWDVRYPRDLISLGV
VIGMEIPQAKASISMYPEIILKRLKY
111111000000000000000000000000000000000000000000000000000000
000000000000000000000000000000000000000000000000000000000000
000000000000000000000000000000000000000000000000000000000000
00000000000000000000001111
>DM_train1394
SMQELQGLDYCKPTRLDLLLDMPPVSYDVQLLHSWNNNDRSLNVFVKEDDKLIFHRHPVAQS
TDAIRGKVGYTRGLHVWQITWAMRQRGTHAVVGVATADAPLHSVGYTTLVGNNHESWGWDLG
RNRLYHDGKNQPSKTYPAFLEPDETFIVPDSFLVALDMDDGTLSFIVDGQYMGVAFRGLKGK
KLYPVVSAVWGHCEIRMRYLNGLDPE
111111110000000000000000000000000000000000000000000000000000
000000000000000000000000000000000000000000000000000000000000
000000000000000000000000000000000000000000000000000000000000
00000000000000000000000011
>DM_train1395
GMASSGTEHDAMRARLVDVAQAIVEERGGAGLTLSELAARAGISQANLSRYFETREDLMEAI
ADYWFHPMVEIMEDVLASDLPPRRKMYEFFARRFVVMRRKWEADPVKLQTYIEVGNDYFEQV
RSYIDLADHYLGEIIGEAMSDGAFSGLEVDETISLVNQMCAPYCALNTMTTFMERLSEDKLA
RIVDAVFDGLSAQDRGARSLTGLRAA
111110000000000000000000000000000000000000000000000000000000
000000000000000000000000000000000000000000000000000000000000
000000000000000000000000000000000000000000000000000000000000
00000000000000000001111111
>DM_train1396
ESNQQIADFDKEKATLDEADIDERMKLAQAFNDSLNNVVSGDPWSEEMKKKGRAEYARMLEI
HERMGHVEIPVIDVDLPVYAGTAEEVLQQGAGHLEGTSLPIGGNSTHAVITAHTGLPTAKMF
TDLTKLKVGDKFYVHNIKEVMAYQVDQVKVIEPTNFDDLLIVPGHDYVTLLTCTPYMINTHR
LLVRGHRIPYVAEVEEEFIAANKLSH
111000000000000000000000000000000000000000000000000000000000

000000000000000000000000000000000000000000000000000000000000
000000000000000000000000000000000000000000000000000000000000
0000000000001111111111111
>DM_train1397
MMTTLTRQDLNFGQVVADVLCEFLEVAVHLILYVREVYPVGIFQKRKKYNVPVQMSCHPELN
QYIQDTLHCVKPLLEKNDVEKVVVVILDKEHRPVEKFVFEITQPPLLSISSDSLLSHVEQLL
AAFILKISVCDAVLDHNPPGCTFTVLVHTREAATRNMEKIQVIKDFPWILADEQDVHMHDPR
LIPLKTMTSDILKMQLYVEERAHKGS
111111111110000000000000000000000000000000000000000000000000
000000000000000000000000000000000000000000011111100000000000
000000000000000000000000000000111111111111100000000000000000
0000000000000000000000000
>DM_train1398
MKGILGTKIGMTQIWKNDRAIPVTVVLAGPCPIVQRKTAQTDGYEAVQIGYAPKAERKVNKP
MQGHFAKAGVAPTRILREFRGFAPDGDSVNVDIFAEGEKIDATGTSKGKGTQGVMKRWNFAG
GPASHGSKKWHRRPGSIGQRKTPGRVYKGKRMAGHMGMERVTVQNLEVVEIRAGENLILVKG
AIPGANGGLVVLRSAAKASAAKGGK
000000000000000000000000000000000000000000000000000000000000
000000000000000000000000000000000000000000000000000000000000
000000000000000000000000000000000000000000000000000000000000
000000000000000000111111
>DM_train1399
NPSPHQVYNVTWTITNLVTGTKANATSMLGTLTDAFPTMYFDLCDIIGNTWNPSDQEPFPGY
GCDQPMRRWQQRNTPFYVCPGHANRKQCGGPQDGFCAVWGCETTGETYWRPTSSWDYITVKK
GVTQGIYQCSGGGWCGPCYDKAVHSSTTGASEGGRCNPLILQFTQKGRQTSWDGPKSWGLRL
YRSGYDPIALFSVSRQVMTITPPQA
111000000000000000000000000000000000000000000000000000000000
000000000000000000000000000000000000000000000000000000000000
000000000000000000000000000000000000000000000000000000000000
000000000000000000000111
>DM_train1400
MGACGSKGSTSDKGLASDKDGKKAKDRKEAWERIRQAIPREKTAEAKQRRIELFKKFDKNET
GKLCYDEVYSGCLEVLKLDEFTSRVRDITKRAFDKSRTLGSKLENKGSEDFVEFLEFRLMLC
YIYDFFELTVMFDEIDASGNMLVDEEEFKRAVPKLEAWGAKVEDPAALFKELDKNGTGSVTF
DEFAAWASAVKLDADGDPDNVPESA
111111111111111000000000000000000000000000000000000000000000
000000000000000000000000000000000000000000000000000000000000
000000000000000000000000000000000000000000000000000000000000
000000000000000000000111
>DM_train1401
PEDPSDLLQHVKFQSSNFENILTWDSGPEGTPDTVYSIEYKTYGERDWVAKKGCQRITRKSC
NLTVETGNLTELYYARVTAVSAGGRSATKMTDRFSSLQHTTLKPPDVTCISKVRSIQMIVHP
TPTPIRAGDGHRLTLEDIFHDLFYHLELQVNRTYQMHLGGKQREYEFFGLTPDTEFLGTIMI
LVPTWAKESAPYMCRVKTLPDRTWT

11000000000000000000000000000000000000000000000000000000000000
00000000000000000000000000000000000000000000000000000000000000
00000001111000000000000000000011110000000000000000000000000000
0000000000000000000000000
>DM_train1402
ALADGKILHNQNVNSWGPITVTPTTDGGETRFDGQIIVQMENDPVVAKAAANLAGKHAESSV
VVQLDSDGNYRVVYGDPSKLDGKLRWQLVGHGRDHSETNNTRLSGYSADELAVKLAKFQQSF
NQAENINNKPDHISIVGSSLVSDDKQKGFGHQFINAMDANGLRVDVSVRSSELAVDEAGRKH
TKDANGDWVQKAENNKVSLSWDAQG
00000000000000000000000000000000000000000000000000000000000000
00000000000000000000000000000000000000000000000000000000000000
00000000000000000000000000000000000000000000000000000000000000
0000000000000000000000111
>DM_train1403
GAMQGEVSKAASADSTTEGTPADGFTVLSTKSLFLGQKLQVVQADIASIDSDAVVHPTNTDF
YIGGEVGNTLEKKGGKEFVEAVLELRKKNGPLEVAGAAVSAGHGLPAKFVIHCNSPVWGADK
CEELLEKTVKNCLALADDKKLKSIAFPSIGSGRNGFPKQTAAQLILKAISSYFVSTMSSSIK
TVYFVLFDSESIGIYVQEMAKLDAN
11111111111111111111111100000000000000000000000000000000000000
00000000000000000000000000000000000000000000000000000000000000
00000000000000000000000000000000000000000000000000000000000000
0000000000000000000000111
>DM_train1404
APISFSHAKNEAVKIYRDHPVSFYCGCEIRWQGKKGIPDLESCGYQVRKNENRASRIEWEHV
VPAWQFGHQLQCWQQGGRKNCTRTSPEFNQMEADLHNLTPAIGEVNGNRSNFSFSQWNGIDG
VTYGQCEMQVNFKERTAMPPERARGAIARTYLYMSEQYGLRLSKAQNQLMQAWNNQYPVSEW
ECVRDQKIEKVQGNSNRFVREQCPN
11110000000000000000000000000000000000000000000000000000000000
00000000000000000000000000000000000000000000000000000000000000
00000000000000000000000000000000000000000000000000000000000000
0000000000000000000000000
>DM_train1405
MFKKLLHSLVAGLTFVAAVAAVPAHAQEADAQATVKTAVDDVLATIKGDPDLRGGNLQKVFQ
LVDQKIVPRADFKRTTQIAMGRFWSQATPEQQQQIQDGFKSLLIRTYAGALANVRNQTVAYK
PFRAAADDTDVVVRSTVNNNGEPVALDYRVEKSPNGWKVYDINISGLWLSETYKNQFADVIS
KRGGVGGLVQFLDERNAQLAKGPAK
11111111111111111111111111110000000000000000000000000000000000
00000000000000000000000000000000000000000000000000000000000000
00000000000000000000000000000000000000000000000000000000000000
0000000000000000000001111
>DM_train1406
MIMSEMITRQQVTSGETIHVRTDPTACIGSHPNCRMFIDSLTIAGEKLDKNIVAIDGGEDVT
KADSATAAASVIRMSITPGSINPTISITLGVLIKSNVRTKIEEKVSSILQASATDMKIKLGN
SNKKQEYKTDEAWGIMIDLSNLELYPISAKAFSISIEPTELMGVSKDGMRYHIISIDGLTTS

QGSLPVCCAASTDKGVAKIGYIAAA
11000000000000000000000000000000000000000000000000000000000000
00000000000000000000000000000000000000000000000000000000000000
00000000000000000000000000000000000000000000000000000000000000
0000000000000000000000011
>DM_train1408
QSLQVKPLQVEPPEPVVAVALGASRQLTCRLACADRGASVQWRGLDTSLGAVQSDTGRSVLT
VRNASLSAAGTRVCVGSCGGRTFQHTVQLLVYAFPNQLTVSPAALVPGDPEVACTAHKVTPV
DPNALSFSLLVGGQELEGAQALGPEVQEEEEPQGDEDVLFRVTERWRLPPLGTPVPPALYC
QATMRLPGLELSHRQAIPVLIEGR
11110000000000000000000000000000000000000000000000000000000000
00000000000000000000000000000000000000000000000000000000000000
00000000000000000000000000000000000000000000000000000000000000
000000000000000000000000
>DM_train1409
MVDFKMTKEGLVLLIKDYQNLEEVLNAISARITQMGGFFAKGDRISLMIENHNKHSQDIPRI
VSHLRNLGLEVSQILVGSTVEGKENDLKVQSRTTVESTGKVIKRNIRSGQTVVHSGDVIVFG
NVNKGAEILAGGSVVVFGKAQGNIRAGLNEGGQAVVAALDLQTSLIQIAGFITHSKGEENVP
SIAHVKGNRIVIEPFDKVSFERSE
00000000000000000000000000000000000000000000000000000000000000
00000000000000001111111111111100000000000000000000000000000000
00000000000000000000000000000000000000000000000000000000000000
000000000000000000001111
>DM_train1410
IGATEQSYVASAVTPLRLNSSTKVLDMLIDSSTLEINSSGQLTVRSTSPNLRYPIADVSGGI
GMSPNYRFRQSMWIGIVSYSGSGLNWRVQVNSDIFIVDDYIHICLPAFDGFSIADGGDLSLN
FVTGLLPPLLTGDTEPAFHNDVVTYGAQTVAIGLSSGGAPQYMSKNLWVEQWQDGVLRLRVE
GGGSITHSNSKWPAMTVSYPRSFT
11111000000000000000000000000000000000000000000000000000000000
00000000000000000000000000000000000000000000000000000000000000
00000000000000000000000000000000000000000000000000000000000000
000000000000000000000000
>DM_train1411
MTNLNYQQTHFVMSAPDIRHLPSDTGIEVAFAGRSNAGKSSALNTLTNQKSLARTSKTPGRT
QLINLFEVADGKRLVDLPGYGYAEVPEEMKRKWQRALGEYLEKRQSLQGLVVLMDIRHPLKD
LDQQMIEWAVDSNIAVLVLLTKADKLASGARKAQLNMVREAVLAFNGDVQVETFSSLKKQGV
DKLRQKLDTWFSEMQPVEETQDGE
11111111111000000000000000000000000000000000000011111111111111
00000000000000000000111111111100000000000000000000000000000000
00000000000000000000000000000000000000000000000000000000000000
000000000000011111111111
>DM_train1412
TVPITSAKAAELPPGNYRLVVFELENFQGRRAEFSGECSNLADRGFDRVRSIIVSAGPWVAF
EQSNFRGEMFILEKGEYPRWNTWSSSYRSDRLMSFRPIKMDAQEHKISLFEGANFKGNTIEI

QGDDAPSLWVYGFSDRVGSVKVSSGTWVGYQYPGYRGYQYLLEPGDFRHWNEWGAFQPQMQS
LRRLRDKQWHLEGSFPVLATEPPK
111111111111000000000000000000000000000000000000000000000000
000000000000000000000000000000000000000000000000000000000000
000000000000000000000000000000000000000000000000000000000000
000000000111111111111111
>DM_train1414
GMEGSLSRGVEIMKLRELVARSRSIRRFDEHVAVNDATLRDLVELVCYTPSAANRQLLRFLP
VTGADMSDKVFPCLKWAGYLEDWPGPEPGERPAAALVMLCRNEDLPGAACDSGIAAQTIMLG
AAEKELGGCIVAAIDRERLMASLGIPDAWTVLLVIALGKPAETVVIDQIKPGDDIRYWRDKH
GIHHVPKRQVDELLVTAEQLRERG
111111111111100000000000000000000000000000000000000000000000
000000000000000000000000000000000000000000000000000000000000
000000000000000000000000000000000000000000000000000000000000
000000000000000000000011
>DM_train1415
SISPAQIAEALQGRGWDAEIVTDASMAGQLVDVRPEGILKCVDGRGSDNTRMGGPKMPGGIY
AIAHNRGVTSIEGLKQITKEVASKGHLPSVHGDHSSDMLGCGFFKLWVTGRFDDMGYPRPQF
DADQGANAVKDAGGIIEMHHGSHTEKVVYINLLANKTLEPNENDQRFIVDGWAADKFGLDVP
KFLIAAAATVEMLGGPKNAKIVVP
100000000000000000000000000000000000000000000000000000000000
000000000000000000000000000000000000000000000000000000000000
000000000000000000000000000000000000000000000000000000000000
000000000000000000000000
>DM_train1416
MENGKRDRQDMEVNTTPRKPRVLLAASGSVAAIKFGNLCHCFTEWAEVRAVVTKSSLHFLDK
LSLPQEVTLYTDEDEWSSWNKIGDPVLHIELRRWADVLVIAPLSANTLGKIAGGLCDNLLTC
IIRAWDYTKPLFVAPAMNTLMWNNPFTERHLLSLDELGITLIPPIKKRLASGDYGNGAMAEP
SLIYSTVRLFWESQAHQQTGGTS
111111111111111111000000000000000000000000000000000000000000
000000000000000000000000000000000000000000000000000000000000
000000000000000000000000000000000000000000001111111110000000
00000000000000111111111
>DM_train1418
MHTDGDKAFVDFLSDEIKEERKIQKHKTLPKMSGGWELELNGTEAKLVRKVAGEKITVTFNI
NNSIPPTFDGEEEPSQGQKVEEQEPELTSTPNFVVEVIKNDDGKKALVLDCHYPEDEVGQED
EAESDIFSIREVSFQSTGESEWKDTNYTLNTDSLDWALYDHLMDFLADRGVDNTFADELVEL
STALEHQEYITFLEDLKSFVKSQ
000000000000000000000000000000000000000000000000000000000000
000111111111111111111111111000000000000000000000000111111111
111100000000000000000000000000000000000000000000000000000000
00000000000000000000000
>DM_train1419
GSGAPPTDLQKMVMGNTKPVELNLDGKTVAICCATGVFGTAYLVPRHLFAEKYDKIMLDGRA

MTDSDYRVFEFEIKVKGQDMLSDAALMVLHRGNKVRDITKHFRDTARMKKGTPVVGVVNNAD
VGRLIFSGEALTYKDIVVSMDGDTMPGLFAYKAATRAGYAGGAVLAKDGADTFIVGTHSAGG
NGVGYCSCVSRSMLQKMKAHVEH
111111100000000000000000000000000000000000000000000000000000
000000000000111111110000000000000000000000000000000000000000
000000000000000001111111000000000000000000000000000000000000
00000000000000000000111
>DM_train1420
GMAVECIKDKVTRYPERASYSDEDLVAMLDRNFTCTVSFIDGGIPYAIPMMLASEGKTIYLH
GSMKSRIYGILKTGQLIAISLLEINGIVLAKEIKNNSINYVSALIFGRPYEIDDTEKKIEVF
RLLTEKLVKGRWDNSIKPSYEDLNGVFVFAVKPETFSMKARTGPPHDTSTDDIWSGVLPIQH
TISEAGENAPEYVKSLYGKRIFI
111111111111111111100000000000000000000000000000000000000000
000000000000000000000000000000000000000000000000000000000000
000000000000000000000000000000000000000000000000000000000000
00000000000000000000000
>DM_train1422
DISEPPLHDFYCSRLLDLVFLLDGSSRLSEAEFEVLKAFVVDMMEQLRISQKWVRVAVVEYH
DGSHAYIGLKDRKRPSELRRIASQVKYAGSQVASTSEVLKYTLFQIFSKIDRPEASRIALLL
MASQEPQRMSRNFVRYVQGLKKKKVIVIPVGIGPHANLKQIRLIEKQAPENKAFVLSSVDEL
EQQRDEIVSYLCDLAPEAPPPT
111111100000000000000000000000000000000000000000000000000000
000000000000000000000000000000000000000000000000000000000000
000000000000000000000000000000000000000000000000000000000000
0000000000000000000000
>DM_train1423
MTKPYVRLDKNDAAVLLVDHQAGLLSLVRDIEPDKFKNNVLALGDLAKYFNLPTILTTSAET
GPNGPLVPELKAQFPDAPYIARPGNINAWDNEDFVKAVKATGKKQLIIAGVVTEVCVAFPAL
SAIEEGFDVFVVTDASGTFNEITRHSAWDRMSQAGAQLMTWFGVACELHRDWRNDIAGLATL
FSNHIPDYRNLMTSYDTLTKQK
000000000000000000000000000000000000000000000000000000000000
000000000000000000000000000000000000000000000000000000000000
000000000000000000000000000000000000000000000000000000000000
0000000000000000000111
>DM_train1424
VIHVTKEVKEVATLSCGHNVSVEELAQTRIYWQKEKKMVLTMMSGDMNIWPEYKNRTIFDIT
NNLSIVILALRPSDEGTYECVVLKYEKDAFKREHLAEVTLSVKADFPTPSISDFEIPTSNIR
RIICSTSGGFPEPHLSWLENGEELNAINTTVSQDPETELYAVSSKLDFNMTTNHSFMCLIKY
GHLRVNQTFNWNTTKQEHFPDN
000000000000000000000000000000000000000000000000000000000000
000000000000000000000000000000000000000000000000000000000000
000000000000000000000000000000000000000000000000000000000000
0000000000000111111111
>DM_train1426

GMVEQRNRGRPAVPDAVVRESIVGAAQRLLSSGGAAAMTMEGVASEAGIAKKTLYRFASGRA
DLIGLLVESWIAPIFPGFEADPQDAAAALERIVYDIAQAVLSREAVSLFRMLASDADLRNRF
LPAYNANGIERSRRELARWLDQQASAGRLPLPIPAERVADLLLSAVIAEPLRQITLGLREPL
PAWDIAPRVADAVRLIAPGRER
111111111111000000000000000000000000000000000000000000000000
000000000000000011000000000000000000000000000000000000000000
000000000000000000000000000000000000000000000000000000000000
0000000000000000001111
>DM_train1427
LKNEETEKKLNLNQQRLGTVVAVLKSVNAKKVIDLGCGEGNLLSLLLKDKSFEQITGVDVSY
SVLERAKDRLKIDRLPEMQRKRISLFQSSLVYRDKRFSGYDAATVIEVIEHLDENRLQAFEK
VLFEFTRPQTVIVSTPNKEYNFHYGNLFEGNLRHRDHRFEWTRKEFQTWAVKVAEKYGYSVR
FLQIGEIDDEFGSPTQMGVFTL
111111111100000000000000000000000000000000000000000000000000
000000000000000000000000000000000000000000000000000000000000
000000000000000000000000111110000000000000000000000000000000
0000000000000000000000
>DM_train1428
MSLEKEYEYLVPPDDYLAAGVHIGTQIKTGDMKKFIFKVRQDGLYVLDIRKLDERIRVAAKF
LSRYEPSKILLVAARQYAHKPVQMFSKVVGSDYIVGRFIPGTLTNPMLSEYREPEVVFVNDP
AIDKQAVSEATAVGIPVVALCDSNNSSADVDLVIPTNNKGRRALAIVYWLLAREIAKIRGQD
FTYSIEDFEAELEGGSHHHHHH
111110000000000000000000000000000000000000000000000000000000
000000000000000000000000000000000000000000000000000000000000
000000000000000000000000000000000000000000000000000000000000
0000000000001111111111
>DM_train1429
MGSDKIHHHHHHMKKLTVYLATTNPHKVEEIKMIAPEWMEILPSPEKIEVVEDGETFLENSV
KKAVVYGKKLKHPVMADDSGLVIYSLGGFPGVMSARFMEEHSYKEKMRTILKMLEGKDRRAA
FVCSATFFDPVENTLISVEDRVEGRIANEIRGTGGFGYDPFFIPDGYDKTFGEIPHLKEKIS
HRSKAFRKLFSVLEKILESENR
111111111111110000000000000000000000000000000000000000000000
000000000000000000000000000000000000000000000000000000000000
000000000000000000000000000000000000000000000000000000000000
0000000000000000011111
>DM_train1430
MGSDKIHHHHHHMNVINETANVLKLETGSVTSAKGFSAVGIHTGVKRKRKDLGAIVCEVPAS
SAAVYTLNKVQAAPLKVTQESIAVEGKLQAMIVNSGIANACTGKRGLDDAYTMRAVGAETFH
IPEHYVAVTSTGVIGEFLPMDVITNGIRQLKPEATIEGAHAFNEAILTTDTVEKHTCYQTIV
NGKTVTVGGVAKGSGMIHPNMA
111111111111111110000000000000000000000000000000000000000000
000000000000000000000000000000000000000000000000000000000000
000000000000000000000000000000000000000000000000000000000000
0000000000000000000000

>DM_train1431
KSVLEDNLPFLEFPGSIVYSYEASDCSFLSEDISMRLSDGDVVGFDMEWPPIYKPGKRSRVA
VIQLCVSESKCYLFHISSMSVFPQGLKMLLENKSIKKAGVGIEGDQWKLLRDFDVKLESFVE
LTDVANEKLKCAETWSLNGLVKHVLGKQLLKDKSIRCSNWSNFPLTEDQKLYAATDAYAGLI
IYQKLGNLGDTVQVFALNKAEE
11111100000000000000000000000000000000000000000000011100000000
00000000000000000000000000000000000000000000000000000000000000
00000000000000000000000000000000000000000000000000000000000000
0000000001111111111111
>DM_train1432
GMSESPTAGRASLAPLREFITGLSALLDEQPGEARILREGGALLARLVARDDWLPDAFAQPH
PEYYQQMLLHCDSAERFSIVSFVWGPGQRTPIHDHTVWGLIGMLRGAEYSQPFVLDGSGRPV
LHGEPTRLEPGHVEAVSPTVGDIHRVHNAYDDRVSISIHVYGANIGGVRRSVYTEAGERKPF
ISGYSNPYLPNPWDRSKDSAAS
11111111111000000000000000000000000000000000000000000000000000
00000000000000000000000000000000000000000000000000000000000000
00000000000000000000000000000000000000000000000000000000000000
0000000000000000011111
>DM_train1433
MTERLNEQNNESYRYLRSVGNQWQFNVEDLHPKMLSRLYKRFDTFDLDSDGKMEMDEVLYWP
DRMRQLVNATDEQVEKMRDAVRVFFLHKGVEPVNGLLREDWVEANRVFAEAERERERRGEPS
LIALLSNSYYDVLDDDGDGTVDVDELKTMMKAFDVPQEAAYTFFEKADTDKSGKLERTELVH
LFRKFWMEPYDPQWDGVYAYKY
11000000000000000000000000111111111000000000000000000000000000
00000000000000000000000000000000000000000000000000000000000000
00000000000000000000000000000000000000000000000000000000000000
0000000111111111111111
>DM_train1434
LERTIEERVNILFDFVKKKKEEGVIDSSDKEIVAEAERLDVKAMGPLVLTEVLFNEKIREQI
KKYRRHFLRFCHNNKKAQRYLLHGLECVVAMHQAQLISKIPHILKEMYDADLLEEEVIISWS
EKASKKYVSKELAKEIRVKAEPFIKWLKEAEEESSGGEEEDEDENIEVVYSKLESVPKVETV
KSDNKDDDIDIDAILEHHHHHH
00000000000000000000000000000000000000000000000000000000000000
00000000000000000000000000000000000000000000000000000000000000
00000000000000000000000000000000000000000000000000000001111111
1111111111111111111111
>DM_train1435
MTEKKIDFKKEEKKFYAPKRKPERIFVPEMNFLMVDGKGDPDGEEYQKAVQSLYAIAYTIKM
SKMGETRLDGYSDFVVPPLEGFWWSEGKFDLKDRDAWLWTSILRQPDFVTEEVLEWAKEVAR
KKKPDVDTSRVKLVRFEEGECVQMMHVGPFSEEVHTVAEMHQFMETEGLRNDTGAIRKHHEI
YLSDPRKANPEKMKTILRLPVS
11110000000000000000000000000000000000000000000000000000000000
00000000000000000000000000000000000000000000000000000000000000
00000000000000000000000000000000000000000000000000000000000000

000000000000000000000
>DM_train1436
MLNKLSLLLKDAGISLTDHQKNQLIAYVNMLHKWNKAYNLTSVRDPNEMLVRHILDSIVVAP
YLQGERFIDVGTGPGLPGIPLSIVRPEAHFTLLDSLGKRVRFLRQVQHELKLENIEPVQSRV
EEFPSEPPFDGVISRAFASLNDMVSWCHHLPGEQGRFYALKGQMPEDEIALLPEEYQVESVV
KLQVPALDGERHLVVIKANKI
000000000000000000000000000000000111111111111000000000000000
000000000000000000000000000000000000000000000000000000000000
000000000000000000000000000000000000000000000000000000000000
000000000000000000000
>DM_train1437
FKVLQEPTCVSDYMSISTCEWKMNGPTNCSTELRLLYQLVFLLSEAHTCIPENNGGAGCVCH
LLMDDVVSADNYTLDLWAGQQLLWKGSFKPSEHVKPRAPGNLTVHTNVSDTLLLTWSNPYPP
DNYLYNHLTYAVNIWSENDPADFRIYNVTYLEPSLRIAASTLKSGISYRARVRAWAQAYNTT
WSEWSPSTKWHNSYREPFEQH
000000000000000000000000000000000000000000000000000000000000
000000000000000000000000000000000000000000011110000000000000
000000000000000000000000000000000000011111000000000000000000
000000000001111111111
>DM_train1438
MGSDKIHHHHHHKGKIQEADNAAKLSAIKFVLKDPLTGDYLVDEKEIEEIVKKTGIETVVLK
EYKEGVVLGPLYEFVTKDGRNAYVLSGYAPGFGGNVTVVACFIKTEDGFMLNSVRVIDYSQE
TPGLGAKIGEESIQRRFFPVPPEGLKNGLRVDKDAGLPKGSPEELKKQGIVKVSDVMTGATI
TPRAVVTALNLMYRYLEEVSK
111111111111111111100000000000000000000000000000000000000000
000000000000000000000000000000001000000000000000000000000001
111111111100000000000000000000000000000000000000000000111110
000000000000000000000
>DM_train1439
GDLIVSMRILGKKRTKTWHKGTLIAIQTVGPGKKYKVKFDNKGKSLLSGNHIAYDYHPPADK
LYVGSRVVAKYKDGNQVWLYAGIVAETPNVKNKLRFLIFFDDGYASYVTQSELYPICRPLKK
TWEDIEDISCRDFIEEYVTAYPNRPMVLLKSGQLIKTEWEGTWWKSRVEEVDGSLVRILFLD
DKRCEWIYRGSTRLEPMFSMK
000000000000000000000000000000000000000000000000000000000000
000000000000000000000000000000000000000000000000000000000000
000000000000000000000000000000000000000000000000000000000000
000000000000000111111
>DM_train1440
GQEHALENYWPVMGVEFSEDMLDFPALFGREAPVTLEIGFGMGASLVAMAKDRPEQDFLGIE
VHSPGVGACLASAHEEGLSNLRVMCHDAVEVLHKMIPDNSLRMVQLFFPDPWHKARHNKRRI
VQVPFAELVKSKLQLGGVFHMATDWEPYAEHMLEVMSSIDGYKNLSESNDYVPRPASRPVTK
FEQRGHRLGHGVWDLMFERVK
111000000000000000000000000000000000000000000000000000000000
000000000000000000000000000000000000000000000000000000000000

00000000000000000000000000000000000000000000000000000000000000
000000000000000000000
>DM_train1441
MTKKSTKSEAASKTKKSGVPETGAQGVRAGGADHADAAHLGTVNNALVNHHYLEEKEFQTVA
ETLQRNLATTISLYLKFKKYHWDIRGRFFRDLHLAYDEFIAEIFPSIDEQAERLVALGGSPL
AAPADLARYSTVQVPQETVRDARTQVADLVQDLSRVGKGYRDDSQACDEANDPVTADMYNGY
AATIDKIRWMLQAIMDDERLD
11111111111111111111111111111000000000000000000000000000000000
00000000000000000000000000000000000000000000000000000000000000
00000000000000000000000000000000000000000000000000000000000000
000000000000000000000
>DM_train1442
MTDHVREADDANIDDLLGDLGGTARAERAKLVEWLLEQGITPDEIRATNPPLLLATRHLVGD
DGTYVSAREISENYGVDLELLQRVQRAVGLARVDDPDAVVHMRADGEAAARAQRFVELGLNP
DQVVLVVRVLAEGLSHAAEAMRYTALEAIMRPGATELDIAKGSQALVSQIVPLLGPMIQDML
FMQLRHMMETEAVNAGERAAG
11111111111100000000000000000000000000000000000000000000000000
00000000000000000000000000000000000000000000000000000000000000
00000000000000000000000000000000000000000000000000000000000000
000000011111111111111
>DM_train1443
MAPAKSPRAWQRMLSGRRLDLLDPSPLDVEIADIAHGLARVARWNGQTRGDHAFTVAQHCLI
VETIFCRMCPGATPDEMQMALLHDAPEYVIGDMISPFKSVVGGGYKTVEKRLEAAVHLRFGL
PPHASRELKDRIKKADTVAAFFEATELAGFSTAEAQKFFGLPRGITRDMFDIIPLPSTEAQR
LFIARFEAIETLRVTRTGGAV
11111000000000000000000000000000000000000000000000000000000000
00000000000000000000000000000000000000000000000000000000000000
00000000000000000000000000000000000000000000000000000000000000
000000000000000111111
>DM_train1444
GYFEGMLIKQTDYFRIYRVINSLLISQNADPASASMYFSTFGAFILQQHYKVKAVPKGGLAA
YNLGGTVLLFADHREDGYVTGAGENFHCWVEADGWAIDFMAPAFSEGTDALAVPAKMFQRPL
SAMAASINDLGQSGDFFYRSEPEATARRFADWHKQAMIGDMASVAANWFRKSPKQMAASLSV
TDRDGKARTVPLTGEMLTGAW
00000000000000000000000000000000000000000000000000000000000000
00000000000000011000000000000000000000000000000000000000000000
00000000000000000000000000000000000000000000000000000000000000
000000000000000000000
>DM_train1445
QTDMSRKAFVFPKESDTSYVSLKAPLTKPLKAFTVCLHFYTELSSTRGYSIFSYATKRQDNE
ILIFWSKDIGYSFTVGGSEILFEVPEVTVAPVHICTSWESASGIVEFWVDGKPRVRKSLKKG
YTVGAEASIILGQEQDSFGGNFEGSQSLVGDIGNVNMWDFVLSPDEINTIYLGGPFSPNVLN
WRALKYEVQGEVFTKPQLWP
00000000000000000000000000000000000000000000000000000000000000

000000000000000000000000000000000000000000000000000000000000
000000000000000111111111000000000000000000000000000000000000
00000000000000000000
>DM_train1446
MQHYLHIRPAPSDNLPLVDLIEHPDPIFDPKEKDLNETLLRSLLGGHYDPGFMATSPPEDRP
GGGGGAAGGAEDLAELDQLLRQRPSGAMPSEIKGLEFSEGLAQGKKQRLSKKLRRKLQMWLW
SQTFCPVLYAWNDLGSRFWPRYVKVGSCFSKRSCSVPEGMVCKPSKSVHLTVLRWRCQRRGG
QRCGWIPIQYPIISECKCSC
000000000000000000000000000000000000000000000000000000000000
111111100000000000000000000000000000000000000000000000000000
000000000000000000000000000000000000000000000000000000000000
00000000000000000000
>DM_train1447
MAKKIFTSALGTAEPYAYIAKPDYGNEERGFGNPRGVYKVDLTIPNKDPRCQRMVDEIVKCH
EEAYAAAVEEYEANPPAVARGKKPLKPYEGDMPFFDNGDGTTTFKFKCYASFQDKKTKETKH
INLVVVDSKGKKMEDVPIIGGGSKLKVKYSLVPYKWNTAVGASVKLQLESVMLVELATFGGG
EDDWADEVEENGYVASGSAK
000000000000000000000000001110000000000000000000000000000000
000000000000000011111110000000000000000000000000000000000000
000000000000000000000000000000000000000000000000000000001111
11000000000111111111
>DM_train1448
HMKISERQKDLLKEIGNIGAGNAATAISYMINKKVEISVPNVEIVPISKVIFIAKDPEEIVV
GVKMPVTGDIEGSVLLIMGTTVVKKILEILTGRAPDNLLNLDEFSASALREIGNIMCGTYVS
ALADFLGFKIDTLPPQLVIDMISAIFAEASIEELEDNSEDQIVFVETLLKVEEEEEPLTSYM
MMIPKPGYLVKIFERMGIQE
000000000000000000000000000000000000000000000000000000000000
000000000000000000000000000000000000000000000000000000000000
000000000000000000000000000000000000000000000000000000000000
00000000000000000001
>DM_train1449
DEIIDETDNLYVSQGPGVDISGDVNLTDFDKIGWPNVEAVQSYQREFNAVSNIFDTIYPIGT
IYENAVNPNNPVTYMGFGSWKLFGQGKVLVGWNEDISDPNFALNNNDLDSGGNPSHTAGGTG
GSTSVTLENTNLPATETDEEVLIVDENGSVIVGGCQYDPDESGPIYTKYREAKASTNSTHTP
PTSITNIQPYITVYRWIRIA
111111111000000000000000000000000000000000000000000000000000
000000000000000000000000000000000000000000000000000000000000
000000000000000000000000000000001111111111110000000000000000
00000000000000000000
>DM_train1450
GSPEFSLDVRQEELGAVVDKEMAATSAAIEDAVRRIEDMMNQARHASSGVKLEVNERILNSC
TDLMKAIRLLVTTSTSLQKEIVESGRGAATQQEFYAKNSRWTEGLISASKAVGWGATQLVEA
ADKVVLHTGKYEELIVCSHEIAASTAQLVAASKVKANKHSPHLSRLQECSRTVNERAANVVA
STKSGQEQIEDRDTMDFSGL

11111110000000000000000000000000000000000000000000000000000000
00000000000000000000000000000000000000000000000000000000000000
00000000000000000000000000000000000000000000000000000000000000
00000000000111111111
>DM_train1451
GSHMQVTPRRNVLQKRPVIVKVLSTTKPFEYETPEMEKKIMFHATVATQTQFFHVKVLNTSL
KEKFNGKKIIIISDYLEYDSLLEVNEESTVSEAGPNQTFEVPNKIINRAKETLKIDILHKQA
SGNIVYGVFMLHKKTVNQKTTIYEIQDDRGKMDVVGTGQCHNIPCEEGDKLQLFCFRLRKKN
QMSKLISEMHSFIQIKKKTN
11111111111100000000000000000000000000000000000000000000000000
00000000000000000000000000000001111111000000000000000000000000
00000000000000000000000000000000000000000000000000000000000000
00000000000000000111
>DM_train1452
GSPNSRVDELGFNEAERQKILDSNSSLMRNANEVRDKFIQNYATSLKDSNDPQDFLRRVQEL
RINMQKNFISFDAYYNYLNNLVLASYNRCKQEKTFAESTIKNELTLGEFVAEISDNFNNFTC
DEVARISDLVASYLPREYLPPFIDGNMMGVAFQILGIDDFGKKLNEIVQDIGTKYIILSKNK
TYLTSLERAKLITQLKLNLE
11111100000000000000000000000000000000000000000000000000000000
00000000000000000000000000000000000000000000000000000000000000
00000000000000000000000000000000000000000000000000000000000000
11111111111111111111
>DM_train1453
GTLIRATTLSRLSFIDVVNDNGFYQFEKPKEGRKYVATLDCSEGRGQDYHALQIIDITEFPY
KQVAVYHSNTTSHFILPDIVFKYLMMYNECPVYIELNSTGVSIAKSLAMDLEYDNIICDSFI
DLGMKQSKRSKAMGCSALKDLIEKDKLIINHKGTIQELRTFSEKGVSWAAEEGFHDDLVMSL
VIFGWLTTQEKFAEYAGKDE
00000000000000000000000000000000000000000000000000000000000000
00000000000000000000000000000000000000000000000000000000000000
00000000000000000000000000000000000000000000011111111000000000
00000000000000000000
>DM_train1454
MVRVKICGITNLEDALFSVESGADAVGFVFYPKSKRYISPEDARRISVELPPFVFRVGVFVN
EEPEKILDVASYVQLNAVQLHGEEPIELCRKIAERILVIKAVGVSNERDMERALNYREFPIL
LDTKTPEYGGSGKTFDWSLILPYRDRFRYLVLSGGLNPENVRSAIDVVRPFAVDVSSGVEAF
PGKKDHDSIKMFIKNAKGL
00000000000000000000000000000000000000000000000000000000000000
00000000000000000000000000000000000000000000000000000000000000
00011111111111000000000000000000000000000000000000000000000000
0000000000000000000
>DM_train1455
MRGSHHHHHHGSMYLDSQLAERRSMHGVLVDIYGLGVLITGDSGVGKSETALELVQRGHRLI
ADDRVDVYQQDEQTIVGAAPPILSHLLEIRGLGIIDVMNLFGAGAVREDTTISLIVHLENWT
PDKTFDRLGSGEQTQLIFDVPVPKITVPVKVGRNLAIIIEVAAMNFRAKSMGYDATKTFEKN

LNHLIEHNEETDQNSSGDK
111111111111111111110000000000000000000000000000000000000000
000000000000000000000000000000000000000000000000000000000000
011111111100000000000000000000000000000000000000000000000000
0000000001111111111
>DM_train1456
TMLCVLTTDAAAEPAALERALRRAAAATFDRLDIDGSCSTNDTVLLLSSGASEIPPAQADLD
EAVLRVCDDLCAQLQADAEGVTKRVTVTVTGAATEDDALVAARQIARDSLVKTALFGSDPNW
GRVLAAVGMAPITLDPDRISVSFNGAAVCVHGVGAPGAREVDLSDADIDITVDLGVGDGQAR
IRTTDLSHAYVEENSAYSS
000000000000000000000000000000000000000000000000000000000000
000000000000000000000000000000000000000000000000000000000000
000000000000000000000000000000000011110000000000000000000000
0000000000000000111
>DM_train1457
MHPKDQSVTLLSIIRDYLKLSDKEREAARILANGLVKVDGKTVREKKFAVGFMDVIEINGES
YRVVYNDQGALVLMKETKERASMKLLKVRSKVIAPGNRIQLGTHDGRTFITDDKSIKVGDVL
AVSVPDMKISEIIKMQPGNKAYITAGSHVNQTGTISKIEAKEGSSANLVHFQEGFSTIKDHV
FMIGSSKFSFVLSPEEVIP
111000000000000000000000000000000000000000000000000000000000
000000000000000000000000000000000000000000000000000000000000
000000000000000000000000000000000000000000110000000000000000
0000000000011111111
>DM_train1458
MEEVEEFKYEPKSVKEIFIEMKDTVELMVDLAYASLLFGDKEIAEEVLELEERIDLLNYQLM
MHSVLAARNVKEAEQVITILQIANAIEDISNAAGDLAKMVLEGVELHPVIKETILEGEEIIG
KIQVYPESVIVGKTLGELDLATNTGVWIIAVRRGKRWIFGPNENFKIRAGDVLIGRGTRTSI
DHLKEIARGAIRVIGNERA
111111110000000000000000000000000000000000000000000000000000
000000000000000000000000000000000000000000000000000000000000
000000000000000000000000000000000000000000000000000000000000
0000000000000001111
>DM_train1459
APEGYRKLLDVQIFKDSPVVGWSGSGMGELETIGDTLPVDTTVTYNGLPTLRLNVQTTVQSG
WWISLLTLRGWNTHDLSQYVENGYLEFDIKGKEGGEDFVIGFRDKVYERVYGLEIDVTTVIS
NYVTVTTDWQHVKIPLRDLMKINNGFDPSSVTCLVFSKRYADPFTVWFSDIKITSEDNEKSA
PAIKVNQLGFIPEAEKYAL
111111100000000000000000000000000000000000000000000000000000
000000000000000000000000000000000000000000000000000000000000
000000000000000000000000000000000000000000000000000000111111
1111111111111111111
>DM_train1460
ASNAMHEWGLSEELKIQTKQMIEIAEKELSIMRNAIDKEDECILCKMEDIHHMLANVQTLAA
TYYIQAYLSPYTESSSFITTAIQHLSARKHGALIVVERNETLEALIQTGTTLNAHLTAPLLE

SIFYPGNPLHDGAVLVKNNHIVSAANILPLTKSTEVDPELGTRHRAAIGLSEKSDALILVVS
EETGRTSFALNGILYTISL
1000000000000000000000000000000000000000000000000000000000000000
0000000000000000000000000000000000000000000000000000000000000000
0000000000000000000000000000000000000000000000000000000000000000
0000000000000000000
>DM_train1461
MSPKVTKEHKDKRQAEILEAAKTVFKRKGFELTTMKDVVEESGFSRGGVYLYFSSTEEMFRR
IIETGLDEGLRKLDKSAEHQSVWASISSYLDELTEGLRDVADTLAPVQFEYLVTAWRNEERR
QYLEKRYDLFVERFSRLLQKGIDQGEFQPVQPLATIAKFFLNMNDGIIQNALYFDEEKADVS
GLAESAKLYLKTVLQADEK
1111100000000000000000000000000000000000000000000000000000000000
0000000000000000000000000000000000000000000000000000000000000000
0000000000000000000000000000000000000000000000000000000000000000
0000000000000000000
>DM_train1462
AQIKSLFATRLYHAPLSEHGPALDPAEFAASCYSIAEDDDAGQEWCEREGYPGYTSYASLTD
LPWRFPIFADLVKSLDAHVAAFAEDLEFELDGKALRLEDIWINILPEGGVHGSHIHPHSVIS
GTTYVAMPEGTSALKLEDPRLPFMMAAPTRRKGAREELRTFRSVAPKVGDVLLWESWLRHEV
PMNMAEEDRISVSFNYAWG
0000000000000000000000000000000000000000000000000000000000000000
0000000000000000000000000000000000000000000000000000000000000000
0000000000000000000000000000000000000000000000000000000000000000
0000000000000000001
>DM_train1464
MATDTLTHATAHAHEHGHHDAGGTKIFGFWIYLMSDCILFSILFATYAVLVNGTAGGPTGKD
IFELPFVLVETFLLLFSSITYGMAAIAMYKNNKSQVISWLALTWLFGAGFIGMEIYEFHHLI
VNGMGPDRSGFLSAFFALVGTHGLHVTSGLIWMAVLMVQIARRGLTSTNRTRIMCLSLFWHF
LDVVWICVFTVVYLMGAM
1111111111111111110000000000000000000000000000000000000000000000
0000000000000000000000000000000000000000000000000000000000000000
0000000000000000000000000000000000000000000000000000000000000000
000000000000000000
>DM_train1465
MILSDKDIIDYVTSKRIIIKPFNKDFVGPCSYDVTLGDEFIIYDDEVYDLSKELNYKRIKIK
NSILVCPLNYNLTEEKINYFKEKYNVDYVVEGGVLGTTNEYIELPNDISAQYQGRSSLGRVF
LTSHQTAGWIDAGFKGKITLEIVAFDKPVILYKNQRIGQLIFSKLLSPADVGYSERKTSKYA
YQKSVMPSLIHLDNHKKD
0000000000000000000000000000000000000000000000000000001111111111
111111111111111111
>DM_train1466
GAFRTQKPSLNTVNVVGSSMGSGGVFTIDGKIKCVTAAHVLTGNSARVSGVGFNQMLDFDVK

GDFAIADCPNWQGVAPKAQFCEDGWTGRAYWLTSSGVEPGVIGNGFAFCFTACGDSGSPVIT
EAGELVGVHTGSNKQGGGIVTRPSGQFCNVKPIKLSELSEFFAGPKVPLGDVKIGSHIIKDT
CEVPSDLCALLAAKPELE
110000000000000000000000000000000000000000000000000000000000
000000000000000000000000000000000000000000000000000000000000
000000000001111100000000000000000000000000000000000000000000
000000000000011111
>DM_train1467
GSHMADSYYDYICIIDFEATCEEGNPPEFVHEIIEFPVVLLNTHTLEIEDTFQQYVRPEINT
QLSDFCISLTGITQDQVDRADTFPQVLKKVIDWMKLKELGTKYKYSLLTDGSWDMSKFLNIQ
CQLSRLKYPPFAKKWINIRKSYGNFYKVPRSQTKLTIMLEKLGMDYDGRPHCGLDDSKNIAR
IAVRMLQDGCELRINEKM
111100000000000000000000000000000000000000000000000000000000
000000000000000000000000000000000000000000000000000000000000
000000000000000000000000000000000000000000000000000000000000
000000000000000000
>DM_train1468
MRGSGSGSMGMTLETAFMLPVQDAQHSFRRLLKAMSEPGVIVALHQLKRGWQPLNIATTSVL
LTLADNDTPVWLSTPLNNDIVNQSLRFHTNAPLVSQPEQATFAVTDEAISSEQLNALSTGTA
VAPEAGATLILQVASLSGGRMLRLTGAGIAEERMIAPQLPECILHELTERPHPFPLGIDLIL
TCGERLLAIPRTTHVEVC
111111111111111111111000000000000000000000000000000000000000
000000000000000000000000000000000000000000000000000000001111
111110000000000000000000001111000000000000000000000000000000
000000000000000000
>DM_train1469
GMEKLNFGIPEWAFEFHGHKCPYMPMGYRAGSYALKIAGLEKEKDHRTYLLSEMSPEDMNGC
FNDGAQAATGCTYGKGLFSLLGYGKLALILYRPGRKAIRVHVRNSFMDELSTRASDFFRYRK
QGYEPSEIPAGAIDPVLEWISSLEDEEIFEYREIDGFTFEPVKKNGAKVRCDVCGEYTYEAD
AKLLNGKPVCKPDYYGKK
100000000000000000000000000000000000000000000000000000000000
000000000000000000000000000000000000000000000000000000000000
000000000000000000000000000000000000000000000000000000000000
000000000000000011
>DM_train1470
AEHHHHHHLVPRGSIAIDDSAAVQRLTGLLNKAQTLTARFSQLTLDGSGTRLQETAGQLSLK
RPGLFRWHTDAPNEQLLISNGEKVWLYDPDLEQVTIQKLDQRLTQTPALLLSGDISKISESF
AITYKEGGNVVDFVLKPKTKDTLFDTLRLSFRSGKVNDMQMIDGVGQRTNILFFDVKMNEAL
DAKQFTFDVPPGVDVIQE
111111111111111111100000000000000000000000000000000000000000
000000000000000000001000000000000000000000000000000000000000
000000000000000000111100000000000000000000000000000000000000
000000000000000000
>DM_train1471

SEHKAKYENVTKDIFDLRDYYSGASKELKNVTGYRYSKGGKHYLIFDKHQKFTRIQIFGKDI
ERLKTRKNPGLDIFVVKEAENRNGTVFSYGGVTKKNQGAYYDYLNAPKFVIKKEVDAGVYTH
VKRHYIYKEEVSLKELDFKLRQYLIQNFDLYKKFPKDSKIKVIMKDGGYYTFELNKKLQPHR
MSDVIDGRNIEKMEANIR
111111111111000000000000000000000000000000000000000000000000
000000000000000000000000000000000000000000000000000000000000
000000000000000000000000000000000000000000000000000000000000
000000000000000000
>DM_train1472
EGNGPAAVHYQPASPPRDACVYSSCYCEENVWKLCEYIKNHDQYPLEECYAVFISNERKMIP
IWKQQARPGDGPVIWDYHVVLLHVSSGGQSFIYDLDTVLPFPCLFDTYVEDAIKSDDDIHPQ
FRRKFRVICADSYLKNFASDRSHMKDSSGNWREPPPPYPCIETGDSKMNLNDFISMDPKVGW
GAVYTLSEFTHRFGSKNC
111111000000000000000000000000000000000000000000000000000000
000000000000000000000000000000000000000000000000000000000000
000000000000000000000000000000000000000000000000000000000000
000000000000000111
>DM_train1474
LKAGPLLSSEKLIAIGASTGGTEAIRHVLQPLPLSSPAVIITQHMPPGFTRSFAERLNKLCQ
ISVKEAEDGERVLPGHAYIAPGDKHMELARSGANYQIKIHDGPPVNRHRPSVDVLFHSVAKH
AGRNAVGVILTGMGNDGAAGMLAMYQAGAWTIAQNEASCVVFGMPREAINMGGVSEVVDLSQ
VSQQMLAKISAGQAIRI
111110000000000000000000000000000000000000000000000000000000
000000000000000000000000000000000000000000000000000000000000
000000000000000000000000000000000000000000000000000000000000
00000000000000000
>DM_train1475
MEPDMTQAFDDDTLRFYRGNATAYAERQPRSATLTKFLGELPAGAKILELGCGAGYQAEAML
AAGFDVDATDGSPELAAEASRRLGRPVRTMLFHQLDAIDAYDAVWAHACLLHVPRDELADVL
KLIWRALKPGGLFYASYKSGEGEGRDKLARYYNYPSEEWLRARYAEAGTWASVAVESSEGKG
FDQELAQFLHVSVRKPE
111111000000000000000000000000000000000000000000000000000000
000000000000000000000000000000000000000000000000000000000000
000000000000000000000000000000000000000000000000000000000000
00000000000000000
>DM_train1476
MRTLFFIPSMGSVRLPLIDFLVKNDIEYVILSRRNHVAVQREIALDMFLEMKDYDTLAFLDE
DVVPIEIDFQKVEAKFNEGYDVVCGYYYLKTLRGYSVYRKDWEKEIFDGEVNGCGLGFTFIK
REFLEKIKRPAFLAFKPIESPHWIGEDVYFFSTHKPRTYALSSLKAYHFIDERLALSPDRKL
ILQNDHVARIKHHHHHH
000000000000000000000000000000000000000000000000000000000000
000000000000000000000000000000000000000000000000000000000000
000000000000001111111110000000000000000000000000000000000000
00000000000000111

>DM_train1477
MRERRYSSTSPERLAAALFDVAAESGLEGASVREVAKRAGVSIGAVQHHFSTKDEMFAFALR
TLVDKLLARLSEVERGGDPARALFAAMSQLLPLDEARSREAHVMAAFAVRAATSPSLAEIRR
KTLFTIRTGLSAVLIGIGTPEAETRAALLLATVDGLALDAIGSPALYPPEYLEHALDIQIGM
ILQGADVVPSSSIELAS
111111111000000000000000000111111111111111111111110000000000
000000000000000000000000000000000000000000000000000000000000
000000000000000000000000000000000000000000000000000000000000
00000011111111111
>DM_train1478
MLSINPNEQTEKDNYKLLTGSIIPRPVAFVTSVTKEGVLNGAPYSYFNIVAANPPLISVSVQ
RKAGERKDTSRNAIEKGEFVVHISDESYVAAINETAANLPPNESEIELAKLTPIESEVISVP
GVKEANIRMECVLERAIPLGGTEDSPACDLLIGRVVRFHVAEHLYEKGRIHAEGLKPISRLA
GHNYAKLGEQFELVRPI
000000000000000000000000000000000000000000000000000000000000
000000000000000000000000000000000000001111000000000000000000
000000000000000000000000000000000000000000000000000000000000
00000000000001111
>DM_train1479
FKIETTPESRYLAQIGDSVSLTCSTTGCESPFFSWRTQIDSPLNGKVTNEGTTSTLTMNPVS
FGNEHSYLCTATCESRKLEKGIQVEIYSFPKDPEIHLSGPLEAGKPITVKCSVADVYPFDRL
EIDLLKGDHLMKSQEFLEDADRKSLETKSLEVTFTPVIEDIGKVLVCRAKLHIDEMDSVPTV
RQAVKELQVYISPKNT
000000000000000000000000000000000000000000000000000000000000
000000000000000000000000000000000000000000000000000000000000
000000000000000000000000000000000000000000000000000000000000
0000000000000111
>DM_train1481
GMDDQSGDQMRPELTMPPAEAEALRMAYEEAEVILEYGSGGSTVVAAELPGKHVTSVESDRA
WARMMKAWLAANPPAEGTEVNIVWTDIGPTGDWGHPVSDAKWRSYPDYPLAVWRTEGFRHPD
VVLVDGRFRVGCALATAFSITRPVTLLFDDYSQRRWQHQVEEFLGAPLMIGRLAAFQVEPQP
IPPGSLMQLIRTMTSP
111111110000000000000000000000000000000000000000000000000000
000000000000000000000000000000000000000000000000000000000000
000000000000000000000000000000000000000000000000000000000000
0000000000000000
>DM_train1482
QKEQHSQLNQTKIAYEQRLLNDLEDMDDPLDLFLDYMIWISTSYIEVDSESGQEVLRSTMER
CLIYIQDMETYRNDPRFLKIWIWYINLFLSNNFHESENTFKYMFNKGIGTKLSLFYEEFSKL
LENAQFFLEAKVLLELGAENNCRPYNRLLRSLSNYEDRLREMNIVENQNSVPDSRERLKGRL
IYRTAPFFIRKFLTSS
110000000000000000000000000000000000000000000000000000000000
000000000000000000000000000000000000000000000000000000000000
000000000000000000000000000000000000000000000000111100000000

0000000000000001
>DM_train1483
LPTLGPGWQRQNPDPPVSRTRSLLLDAASGQLRLEDGFHPDAVAWANLTNAIRETGWAYLDL
STNGRYNDSLQAYAAGVVEASVSEELIYMHWMNTVVNYCGPFEYEVGYCEKLKNFLEANLEW
MQREMELNPDSPYWHQVRLTLLQLKGLEDSYEGRLTFPTGRFTIKPLGFLLLQISGDLEDLE
PALNKTNTKPSLGSGS
1111111111111111000000000000000000000000000000000000000000000000
0000000000000000000000000000000000000000000000000000000000000000
0000000000000000000000000000000000000000000000000000000000000000
0000001111110000
>DM_train1484
ISTLHHQTMPSDLTQDEFTQLSQSIAEFHTYQLGNGRCSSLLAQRIHAPPETVWSVVRRFDR
PQIYKHFIKSCNVSEDFEMRVGCTRDVNVISGLPANTSRERLDLLDDDRRVTGFSITGGEHR
LRNYKSVTTVHRFEKEEEEERIWTVVLESYVVDVPEGNSEEDTRLFADTVIRLNLQKLASIT
EAMNRNNNNNNSSQVR
1111111111100000000000000000000000000000000000000000000000000000
0000000000000000000000000000000000000000000000000000000000000000
0000000000000000000000000000000000000000000000000000000000000000
0000111111111111
>DM_train1485
MGIHLFWDSRVPVGLSRPVSEELSAVLEMPVSRIDDGIFPLEGFDPVRNQYDAVKVLLKLDM
FRRRMPQIFKPADMDLEFYNKFNHLHEKILLVTPGDLYEPLADFVFGLAYPKLGVAIVSPHR
LQNEFYGKYADDSALIDRIVKEGAHEIGHLFGLGHCDNPGCIMYCPRNLDELDRKRKYFCGK
CRVQLNGDTLEDDLFS
1000000000000000000000000000000000000000000000000000000000000000
0000000000000000000000000000000000000000000000000000000000000000
0000000000000000000000000000000000000000000000000000000000000000
0000001111111111
>DM_train1486
KLIGMLDSPYVRRVAISLKSLGLPFEHHSLSVFSTFEQFKAINPVVKAPTLVCEGGEVLMDS
SLIIDYLETLAGPQRSLMPTALPQRLRELRLVGLALAACEKSVQIVYERNLRPAEKQHGPWL
ERVGGQLQAAYGELEQELQKQPLPRDGSLGQAGISLAVAWSFSQMMVADQFNPGQFPAVRGF
AEYAEQLPVFLATPAT
0000000000000000000000000000000000000000000000000000000000000000
0000000000000000000000000000000000000000000000000000000000000000
0000000000000000000000000000000000000000000000000000000000000000
0000000000000001
>DM_train1488
MKLFYKPGACSLASHITLRESGKDFTLVSVDLMKKRLENGDDYFAVNPKGQVPALLLDDGTL
LTEGVAIMQYLADSVPDRQLLAPVNSISRYKTIEWLNYIATELHKGFTPLFRPDTPEEYKPT
VRAQLEKKLQYVNEALKDEHWICGQRFTIADAYLFTVLRWAYAVKLNLEGLEHIAAFMQRMA
ERPEVQDALSAEGLK
0000000000000000000000000000000011111111110000000000000000000000
0000000000000000000000000000000000000000000000000000000000000000

000000000000000011100000000000000000000000000000000000000000
000000000000000
>DM_train1489
AATSRTERIWPGGVIPYVIGGNFTGSQRAMFKQAMRHWEKHTCVTFIERSDEESYIVFTYRP
CGCCSYVGRRGNGPQAISIGKNCDKFGIVVHELGHVIGFWHEHTRPDRDNHVTIIRENIQPG
QEYNFLKMEPGEVNSLGERYDFDSIMHYARNTFSRGMFLDTILPSRDDNGIRPAIGQRTRLS
KGDIAQARKLYRCPA
000000000000000000000000000000000000000000000000000000000001
111000000000000000110000000000000000000000000000000000000000
000000000000000000000000000000000000000000000000000000000000
000000000000000
>DM_train1490
GADNNKWKPLFGKNLENANYNPEVWSETDGVLGAVKDESIWTKDEYENFELDLDFKTDVGTN
SGVVVYCTDTKDWIPNSVEIQIADDHCEKWGNGKPYEKCGAIYGHLGAVQDKVVKKPGEWNH
MRIKCAGQHIMVILNGKKVTEMDMSKWTSGTKNPDGSDIPSWLPKPFAELPTKGFIGLQGKH
GDSLIWFRNIKIRSL
111100000000000000000000000000000000000000000000000000000000
000000000000000000000000000000000000000000000000000000000000
000000000000000000000000000000000000000000000000000000000000
000000000000000
>DM_train1491
MGAAGDAAIGRESRELMSAANVGRTISRIAHQIIEKTALDDPVGPDAPRVVLLGIPTRGVTL
ANRLAGNITEYSGIHVGHGALDITLYRDDLMIKPPRPLASTSIPAGGIDDALVILVDDVLYS
GRSVRSALDALRDVGRPRAVQLAVLVDRGHRELPLRADYVGKNVPTSRSESVHVRLREHDGR
DGVVISRGSHHHHHH
111111111110000000000000000000000000000000000000000000000000
000000000000000000000000000011111111000000000000000000000000
000000000000000000000000000000000000000000000000000000000000
000000011111111
>DM_train1492
MSANVQEAANAAIEPASFVKVPMPEPPSSLQQLINDWQLIKHREGGYFKETDRSPYTMEVEK
PVNGGSGNTEMVTRNQSTLIYYLLTPDSPIGKFHKNINRIIHILQRGKGQYVLVYPDGQVKS
FKVGFDYKNGEVSQWVVPGGVFKASFLLPNEEFDNGFLISEVVVPGFDFEDHTFLKGEDELK
HLVGPEKAAELAFLA
111111110000000000000000000000000000000000000000000000000000
111111111000000000000000000000000000000000000000000000000000
000000000000000000000000000000000000000000000000000000000000
000000000000000
>DM_train1493
MRTFTWRKGSLSKVNTCYVLAGGKSKRFGEDKLLYEIKGKKVIERVYETAKSVFKEVYIVAK
DREKFSFLNAPVVLDEFEESASIIGLYTALKHAKEENVFVLSGDLPLMKKETVLYVLENFKE
PVSVAKTEKLHTLVGVYSKKLLEKIEERIKKGDYRIWALLKDVGYNEVEIPEELRYTLLNMN
TKEDLKRILAIENHY
111000000000000000000000000000000000000000000000000000000000

000000000000000000000000000000000000000000000000000000000000
000000000000000000000000000000000000000000000000000000000000
001111111111111
>DM_train1494
MGSSSSGRENLYFQGHMKNDTAALAADIVDFWKKAGPDKWFDKDAAFDNHFHDRFRDAHFAA
ARRELDGWLEGAESSLALMLLLDQFPRNCFRGTAHMYATDPLARFFADEAIRRGHDQAVSED
LRVFFYLPFSHAEDIAAQQRACDLNQPLGGLYLHHAEEHRDIVERFGRFPHRNGILLRETTP
EERQYLEEGGFSGGS
111111111111111111100000000000000000000000000000000000000000
000000000000000000000000000000000000000000000000000000000000
000000000000000000000000000000000000000000000000000000000000
000000000111111
>DM_train1495
MELRDRIDFLCKTILAIKTAGRLVLGIDGLSRSGKTTLANQLSQTLREQGISVCVFHMDDHI
VERAKRYHTGNEEWFEYYYLQWDVEWLTHQLFRQLKASHQLTLPFYDHETDTHSKRTVYLSD
SDMIMIEGVFLQRKEWRPFFDFVVYLDCPREIRFARENDQVKQNIQKFINRYWKAEDYYLET
EEPIKRADVVFDMTS
000000000000000000000000000000000000000000000000000000000000
000000000000000000000000000000000000000000000000000000000000
000000000000000000000000000001111111111111100000000000000000
000000000000111
>DM_train1496
VSDAAERNLAFALSELDRITAQLKLPRHVEEEAARLYREAVRKGLIRGRSIESVMAACVYAA
CRLLKVPRTLDEIADIARVDKKEIGRSYRFIARNLNLTPKKLFVKPTDYVNKFADELGLSEK
VRRRAIEILDEAYKRGLTSGKSPAGLVAAALYIASLLEGEKRTQREVAEVARVTEVTVRNRY
KELVEKLKIKVPIA
111111100000000000000000000000000000000000000000000000000000
000000000000000000000000000000000000000000000000000000000000
000000000000000000000000000000000000000000000000000000000000
00000000000000
>DM_train1497
MHTKGPTPQQHDGSALRIGIVHARWNETIIEPLLAGTKAKLLACGVKESNIVVQSVPGSWEL
PIAVQRLYSASQLQTPSSGPSLSAGDLLGSSTTDLTALPTTTASSTGPFDALIAIGVLIKGE
TMHFEYIADSVSHGLMRVQLDTGVPVIFGVLTVLTDDQAKARAGVIEGSHNHGEDWGLAAVE
MGVRRRDWAAGKTE
111100000000000000000000000000000000000000000000000000000000
000000000000001111111111111111111111111111000000000000000000
000000000000000000000000000000000000000000000000000000000000
00000000000000
>DM_train1498
GMTIKALFWDIGGVLLTNGWDREQRADVAQRFGLDTDDFTERHRLAAPELELGRMTLAEYLE
QVVFYQPRDFTPEDFRAVMEEQSQPRPEVLALARDLGQRYRMYSLNNEGRDLNEYRIRTFGL
GEFLLAFFTSSALGVMKPNPAMYRLGLTLAQVRPEEAVMVDDRLQNVQAARAVGMHAVQCVD
AAQLREELAALGVR

100000000000000000000000000000000000000000000000000000000000
000000000000000000000000000000000000000000000000000000000000
000000000000000000000000000000000000000000000000000000000000
00000000000000
>DM_train1499
MSPSTEAAARTPSEARARLLGTATRIFYAEGIHSVGIDRITAEAQVTRATLYRHFSGKDDLI
LAYLDQADRGIRAQVTAARGSSPAADGQVRAVARSIVDGIRSPGFRGCAFLNAVAEYPDPAH
PVHRAVLAHRQWFLDTVTELLAQVGDGDGVAAGRHLVMLRDGAMAAGCLFDPELVSETFLHG
VEGVLRDVSEKTSA
111111111111000000000000000000000000000000000000000000000000
000000000000000000000000000000000000000000000000000000000000
000000000000000000000000000000000000000000000000000000000000
00000000001111
>DM_train1500
GMIPFPESRLAAQMSFVVEIDKLKTILRQTLLTDSSRRENDAEHSWHIATMAFLLAEYADEA
VQIGRVARMLLIHDIVEIDAGDTFIHDEAGNEDKEERERKAAARLFGLLPPDQAAEYSALWQ
EYEARETADARFADALDRLQPLLHNFETEGGTWKPHGVTRAKVDKLLPRIEAGSKRLGAYAR
ALVDEAVRRGYLAP
111111000000000000000000000000000000000000000000000000000000
000000000000000000000000000111110000000000000000000000000000
000000000000000000000000000000000000000000000000000000000000
00000000000000
>DM_train1501
MGHHHHHHGGHMSLLRFLEVVSEHIKNLRNHIDLETVGEMIKLIDSARSIFVIGAGRSGYIA
KAFAMRLMHLGYTVYVVGETVTPRITDQDVLVGISGSGETTSVVNISKKAKDIGSKLVAVTG
KRDSSLAKMADVVMVVKGKMKQERDEILSQLAPLGTMFELTAMIFLDALVAEIMMQKHLTEK
DLEARHAVLEEGGS
111111111100000000000000000000000000000000000000000000000000
000000000000000000000000000000000000000000000000000000000000
000000000000000000000000000000000000000000000000000000000000
00000000000011
>DM_train1502
MASMTGGQQMGRGSIVAASTIPGSAATLNTSITKNIQNGNAYIDLYDVKLGKIDPLQLIVLE
QGFTAKYVFRQGTKYYGDVSQLQSTGRASLTYNIFGEDGLPHVKTDGQIDIVSVALTIYDST
TLRDKIEEVRTNANDPKWTEESRTEVLTGLDTIKTDIDNNPKTQTDIDSKIVEVNELEKLLV
LKLAAALEHHHHHH
111111111111111111000000000000000000000000000000000000000000
000000000000000000000000000000000000000000000000000000000000
000000000000000000000000000000000000000000000000000000000000
00000000000011
>DM_train1503
GSMSSPYILVLYYSRHGATAEMARQIARGVEQGGFEARVRTVPAVSTECEAVAPDIPAEGAL
YATLEDLKNCAGLALGSPTRFGNMASPLKYFLDGTSSLWLTGSLVGKPAAVFTSTASLHGGQ
ETTQLSMLLPLLHHGMLVLGIPYSEPALLETRGGGTPYGASHFAGADGKRSLDEHELTLCRA

LGKRLAETAGKLGS
111100000000000000000000000000000000000000000111111111111100
000000000000000000000000000000000000000000000000000000000000
000000000000000000000011111111111100000000000000000000000000
00000000000000
>DM_train1504
RRLETQKEAEANRGYGKPSLGGPFHLEDMYGNEFTEKNLLGKFSIIYFGFSNCPDICPDELD
KLGLWLNTLSSKYGITLQPLFITCDPARDSPAVLKEYLSDFHPSILGLTGTFDEVKNACKKY
RVYFSTPPNVKPGQDYLVDHSIFFYLMDPEGQFVDALGRNYDEKTGVDKIVEHVKSYVPAEQ
RAKQKEAWYSFLFK
111111111111111000000000000000000000000000000000000000000000
000000000000000000000000000000000000000000000000000000000000
001111111111111100000000000000000000000000000000000000000011
11111111111111
>DM_train1505
MALKYPSGKEYRGNKPNAARRPAADYANRGMTLEDDLNATNEYYRERGIAVIHKKPTPVQIV
RVDYPKRSAAVITEAYFRQASTTDYNGVYRGKYIDFEAKETKNKTAFPLKNFHAHQIRHMEQ
VVAHGGICFAILRFSLLNETYLLDASHLIAWWNKQEAGGRKSIPKQEIERHGHSIPLGYQPR
IDYISVVDNVYFTR
111111111111111111111111111110000000000000000000000000000111
111111111111110000000000000000000000000000000000000000000000
000000000000000000000000000000000000000000000000000000000000
00000000000000
>DM_train1506
MVGQVIRKRGAYPEYTVEDVLAVIFLLKEPLGRKQISERLELGEGSVRTLLRKLSHLDIIRS
KQRGHFLTLKGKEIRDKLLSMFSEPIGVSVDGYPGIAIVVKNPPEFKSIELRDEAIKFDAKG
AMILTVKDNEIVFPEDFRPLKEMYPEVAKKIVDYEDGDAVIITWAETPAKALKSAIHVAYIL
KKEEITPEILEVVK
111111111111100000000000000000000000000000000000000000000000
011000000000000000000000000000000000000000000000000000000000
000000000000000000000000000000000000000000000000000000000000
00000000000001
>DM_train1507
GMTKKTGFFKRLKALTLPQKQLFATALCQRMLPNYQLFSEVCEFGDPAVLSTALELLWQSLY
DPKLKFNIDVHLQRLEDNTPEPADFEAYGVYPAMDAVVAISTLLGAIQGKIEEDIVNISKLS
SSTVANYIEAISDVDLVDEALDDFVFAHEVMEEEKELQNSLLEIIEENPKITAELVKGLRKD
IIETGVSNIGISVA
111100000000000000000000000000000000000000000000000000000000
000000000000000000000000000000000000000000000000000000000000
000000000000000000000000000000000000000000000000000000000000
00000000000000
>DM_train1508
DYGRADNENRVGQVTGLAWTEVGGDLLTIETACVPGKGKLTYTGSLGEVMQESIQAALTVVR
ARAEKLGINPDFYEKRDIHVHVPEGATPKDGPAAGIAMCTALVSCLTGNPVRADVAMTGEIT

LRGQVLPIGGLKEKLLAAHRGGIKTVLIPFENKRDLEEIPDNVIADLDIHPVKRIEEVLTLA
LQNEPSGMQVVTAK
11111111100000000000000000000000000000000000000000000000000000
00000000000000000000000000000000000000000000000000000000000000
00000000000000000000000000000000000000000000000000000000000000
00000111111111
>DM_train1509
SLGLAQSAQEILDRVEKNLSTPWQATVQGRIQGPGGEEELLARVYALPQARLFRVEFLKPGS
LEGNFTVITEKEVWNYLYLTNQLVISPREKAKIQGLGFAPQGLGDLKALSEQVDLRLEGEVR
LPEGMAWKLVGRSKENQGFAAMELYILKADPRPLRFVFLDEKGKVLADLKVVEFKRTNLTEA
QLKRYPKDAQVVRR
11111000000000000000000000000000111110000000000000000000000000
00000000000000000000000000011111111111111111111111100000000000
00000000000001110000000000000000000000000000000000000000000000
00000000000000
>DM_train1510
SKVLVLKSSILAGYSQSNQLSDYFVEQWREKHSADEITVRDLAANPIPVLDGELVGALRPSD
APLTPRQQEALALSDELIAELKAHDVIVIAAPMYNFNISTQLKNYFDLVARAGVTFRYTENG
PEGLVTGKKAIVITSRGGIHKDGPTDLVTPYLSTFLGFIGITDVKFVFAEGIAYGPEMAAKA
QSDAKAAIDSIVSA
00000000000000000000000000000000000000000000000000000000001111
11000000000000000000000000000000000000000000000000000000000000
00000000000000000000000000000000000000000000000000000000000000
00000000000000
>DM_train1511
MEHICGTSRIAGFRFSLYPMTDDFISVIKSALAATDTSKVWTKTDHISTVLRGSIDHVFDAA
KAIYLHAANSEQHIVMNGTFSIGCPGDTQGDTYLSKGDKRVNEDAVRGLKAEAPCQFALYPM
NEPDYMGLIMEAVDIAKAQGTFVQGVHYASELDGDAHDVFSTLEAVFRMAEQQTNHITMTVN
LSANSPSRKNRKQG
11111111000000000000000000000000000000000000000000000000000000
00000000000000000000000000000000011110000000000000000000000000
00000000000000000000000000000000000000000000000000000000000000
00000011111111
>DM_train1512
VAPGIVIPPKALFTQQGGAYGRCPNGTRALTVAELRGNAELQTYLRQITPGWSIYGLYDGTY
LGQAYGGIIKDAPPGAGFIYRETFCITTIYKTGQPAADHYYSKVTATRLLASTNSRLCAVFV
RDGQSVIGACASPYEGRYRDMYDALRRLLYMIYMSGLAVRVHVSKEEQYYDYEDATFQTYAL
TGISLCNPAASIC
11100000000000000000000000000000000000000000000000000000000000
00000000000000000000000000000000000000000000000000000000000000
00000000000000000000000000000000000000000000000000000000000000
0000000000000
>DM_train1514
MKSERILVVKTEDFLKEFGEFEGFMRVNFEDFLNFLDQYGFFRERDEAEYDETTKQVIPYVV

IMDGDRVLITKRTTKQSEKRLHNLYSLGIGGHVREGDGATPREAFLKGLEREVNEEVDVSLR
ELEFLGLINSSTTEVSRVHLGALFLGRGKFFSVKEKDLFEWELIKLEELEKFSGVMEGWSKI
SAAVLLNLFLTQN
111000000000000000000000000000000000000000000000000000000000
000000000000111111111110000000000000000000000000000000000000
000000000000000000000000000000000000000000000000000000000000
0000000001111
>DM_train1515
AHMGIQRPTSTTTDKKEIKAYLKQVDKIKDDEEPIKTVGKKIAELDEKKKKLTEDVNSKDTA
VRGKAVKDLIKNADDRLKEFEKEEDAIKKSEQDFKKAKSHVDNIDNDVKRKEVKQLDDVLKE
KYKLHSDYAKAYKKAVNSEKTLFKYLNQNDATQQGVNEKSKAIEQNYKKLKEVSDKYTKVLN
KVQKEKQDVDQFK
000000000000000000000000000000000000000000000000000000000000
000000000000000000000000000000000000011110000000000000000000
000000000000000000000000000000000000000000000000000000000000
0000000000111
>DM_train1516
MTAPHDVPDLTGSGEYLVPDVLQPGLTLVLVGTAPSGISARARAYYANPENKFWRTLHAVGL
TPRQLVPQEYATLPQYGLGLTDVAKRHSGVDAALPGEAWRPDELRRKVEHYRPRIVAFTSKR
GASETLGVPTGKLPYGPQPQPLDWPAETELWVLPSTSPLGHNHFRLEPWQALGDRVRELRGA
AEAGNPSPETPVL
111111000000000000000000000000000000000000000000000000000000
000000000000000000000000000000000000000000000000000000000000
000000000000000000000000000000000000000000000000000000000000
0001111111111
>DM_train1517
MRHDFVFSGVIMLELNAKTTALVVIDLQEGILPFAGGPHTADEVVNRAGKLAAKFRASGQPV
FLVRVGWSADYAEALKQPVDAPSPAKVLPENWWQHPAALGTTDSDIEIIKRQWGAFYGTDLE
LQLRRRGIDTIVLCGISTNIGVESTARNAWELGFNLVIAEDACSAASAEQHNNSINHIYPRI
ARVRSVEEILNAL
111111111111110000000000000000000000000000000000000000000000
000000000000000000000000000000000000000000000000000000000000
000000000000000000000000000000000000000000000000000000000000
0000000000000
>DM_train1518
GMTVRDENYFTEKYGLTRTHSDVLAAAKVVAPGRTLDLGCGNGRNSLYLAANGYDVTAWDKN
PASMANLERIKAAEGLDNLQTDLVDLNTLTFDGEYDFILSTVVMMFLEAQTIPGLIANMQRC
TKPGGYNLIVAAMDTPDFPCTVGFPFAFKEGELRRYYEGWDMLKYNEDVGELHRTDENGNRI
KLRFATMLARKTA
000000000000000000000000000000000000000000000000000000000000
000000000000000000000000000000000000000000000000000000000000
000000000000001111111110000000000000000000000000111111111111
0000000000000
>DM_train1519

MSWENPGPPKRADARRNYDRIIEAAAAEVARHGADASLEEIARRAGVGSATLHRHFPSRWGL
LQAVFQERVAQLCDEARSLAAEHPPATALTRWLTSLAVFGAVTRGAARSLLPATGTNTGAAL
DSRCEQLLTEAGADLLARAQEDGTVRDDVTALELLSLANAVSLAAEHTPDAAHHATRLMGIA
LGGLGAPGPRPQG
111111111110000000000000000000000000000000000000000000000000
000000000000000000000000000000000000000000000000111111111000
000000000000000000000000000000000000000000000000000000000000
0000001111111
>DM_train1521
MDAPDISYEHQETSVPNRSGIIPTLQNVVATVNLSCKLDLKNIALRARNAEYNPKRFAAVIM
RIREPKTTALIFASGKMVITGAKSEKSSRMAAQRYAKIIHKLGFNATFDDFKIQNIVSSCDI
KFSIRLEGLAYAHSNYCSYEPELFPGLIYRMVKPKIVLLIFVSGKIVLTGAKVRDDIYQAFN
NIYPVLIQHRKA
111111111111111111000000000000000000000000000000000000000000
000000000000000000000000000000000000000000000000000000000000
000000000000000000000000000000000000000000000000000000000000
000000000001
>DM_train1522
MGSSGSSNILIINGAKKFAHSNGQLNDTLTEVADGTLRDLGHDVRIVRADSDYDVKAEVQNF
LWADVVIWQMPGWWMGAPWTVKKYIDDVFTEGHGTLYASDGRTRKDPSKKYGSGGLVQGKKY
MLSLTWNAPMEAFTEKDQFFHGVGVDGVYLPFHKANQFLGMEPLPTFIANDVIKMPDVPRYT
EEYRKHLVEIFG
111110000000000011111111100000000000000000000000000000000000
000000000000000000000000000000000000001111111110000000000000
000000000000000000000000000000000000000000000000000000000000
000000000000
>DM_train1524
MFADRLFNAMERNEPAPGMVLVAAPSMESEDFARSVILIIEHSEYATFGVNLASRSDVAVFN
VIPEWVPCVTKPQALYIGGPLNQQSVVGVGVTAQGVDAARVDNLTRLANRLVMVNLGADPEE
IKPLVSGMRLFAGHAEWAPGQLAQEIENGDWFVAPALPSDVTAPGSVDVWGDVMRRQPMPLP
LYSTFPVNVGEN
000000000000000000000000000000000000000000000000000000000000
000000000000000000000000000000000000000000000000000000000000
000000000000000000000000000000000000000000000000000000000000
000000011111
>DM_train1525
MTSKGPEEEHPSVTLFRQYLRIRTVQPKPDYGAAVAFFEETARQLGLGCQKVEVAPGYVVTV
LTWPGTNPTLSSILLNSHTDVVPVFKEHWSHDPFEAFKDSEGYIYARGAQDMKCVSIQYLEA
VRRLKVEGHRFPRTIHMTFVPDEEVGGHQGMELFVQRPEFHALRAGFALDEGIANPTDAFTV
FYSERSPWWVRV
111111000000000000000000000000000000000000000000000000000000
000000000000000000000000000000000000000000000000000000000000
000000000000000000000000000000000000000000000000000000000000
000000000001

>DM_train1526
MHALYITHPQVKIDPAVPVPEWGLSERGAERAREASRLPWAKALRRIVSSAETKAIETAHML
AETSGAAIEIIEAMHENDRSATGFLPPPEFEKAADWFFAHPEESFQGWERAIDAQARIVEAV
KAVLDRHDARQPIAFVGHGGVGTLLKCHIEGRGISRSKDQPAGGGNLFRFSIAEFSLAAASP
TCDWTAMETWQG
000000000000000000000000000000000000000000000000000000000000
000000000000000000000000000000000000000000000000000000000000
000000000000000000000000000000000000110000000000000000100011
000000000000
>DM_train1527
MDTLAPESTRQNLRSQRLNLLTNEPHQRLESLVKSKEPFASRDNFARFVAAQYLFQHDLEPL
YRNEALARLFPGLASRARDDAARADLADLGHPVPEGDQSVREADLSLAEALGWLFVSEGSKL
GAAFLFKKAAALELDENFGARHLAEPEGGRAQGWKSFVAILDGIELNEEEERLAAKGASDAF
NRFGDLLERTFA
111111111110000000000000000000000000000000000000000000000000
000000000000000000000000000000000000000000000000000000000000
000000000000000000000000000000000000000000000000000000000000
000000000000
>DM_train1528
GMAGKVEARKAALREKLIDLAEAQIEAEGLASLRARELARQADCAVGAIYTHFQDLNALTLE
VNGRTFARLGAAVGAVVADGQDDHPNERLIAMSHAYLAFAREHPKLWRALFDVEMRSDGPVP
QWYGHAMAQLFSYITTPLAKIFPESDDAELDLMTRTLFSSVHGIVLLGLENRISGVPGEQLK
TMIRLLLEQVGR
111111111000000000000000000000000000000000000000000000000000
000000000000001111111000000000000000000000000000000000000000
000000000000000000000000000000000000000000000000000000000000
000000000000
>DM_train1529
MATFYEVIVRVPFDVEEHLPGISDSFVDWVTGQIWELPPESDLNLTLVEQPQLTVADRIRRV
FLYEWNKFSKQESKFFVQFEKGSEYFHLHTLVETSGISSMVLGRYVSQIRAQLVKVVFQGIE
PQINDWVAITKVKKGGANKVVDSGYIPAYLLPKVQPELQWAWTNLDEYKLAALNLEERKRLV
AQFLAESSQRS
000000000000000000000000000000000000000000000000000000000000
000000000000000000000000000000000000000000000000000000000000
000000000000000000000000000000000000000000000000000000000000
00000001111
>DM_train1530
RNLPVATPDPGMFPCLHHSQNLLRAVSNMLQKARQTLEFYPCTSEEIDHEDITKDKTSTVEA
CLPLELTKNESCLNSRETSFITNGSCLASRKTSFMMALCLSSIYEDLKMYQVEFKTMNAKLL
MDPKRQIFLDQNMLAVIDELMQALNFNSETVPQKSSLEEPDFYKTKIKLCILLHAFRIRAVT
IDRVMSYLNAS
111111111111111111100000000000000000000011111111111111100000
000000000000111110000000111111100000000000000000000000000000
000000000000000000000000111111111111111110000000000000000000

00000000000
>DM_train1531
GSHGTTFNGIMQSYVPRIVPDEPDNDTDVESCINRLREDDTDLKEVNINNMKRVSKERIRSL
IEAACNSKHIEKFSLANTAISDSEARGLIELIETSPSLRVLNVESNFLTPELLARLLRSTLV
TQSIVEFKADNQRQSVLGNQVEMDMMMAIEENESLLRVGISFASMEARHRVSEALERNYERV
RLRRLGKDPNV
11111111111111111111111111000000000000000000000000000000000000
00000000000000000000000000000000000000000000000000000000000000
00000000000000000000000000000000000000000000000000000000000000
00000001111
>DM_train1532
SNAMEALKRKIEEEGVVLSDQVLKVDSFLNHQIDPLLMQRIGDEFASRFAKDGITKIVTIES
SGIAPAVMTGLKLGVPVVFARKHKSLTLTDNLLTASVYSFTKQTESQIAVSGTHLSDQDHVL
IIDDFLANGQAAHGLVSIVKQAGASIAGIGIVIEKSFQPGRDELVKLGYRVESLARIQSLEE
GKVSFVQEVHS
00000000000000000000000000000000000000000000000000000000000000
00000000000000000000000000000000000000001110000000000000000000
00000000000000000000000000000000000000000000000000000000000000
00000011111
>DM_train1533
GHMNDALHIGLPPFLVQANNEPRVLAAPEARMGYVLELVRANIAADGGPFAAAVFERDSGLL
IAAGTNRVVPGRCSAAHAEILALSLAQAKLDTHDLSADGLPACELVTSAEPCVMCFGAVIWS
GVRSLVCAARSDDVEAIGFDEGPRPENWMGGLEARGITVTTGLLRDAACALLREYNACNGVI
YNARCGVHKGS
00000000000000000000000000000000000000000000000000000000000000
00000000000000000000000000000000000000000000000000000000000000
00000000000000000000000000000000000000000000000000000000001111
11111111111
>DM_train1534
MSLEKPKVERSEGKEKLEDYAYVLDFMPYGHPDDKRPIHRREPLAQVVGERNFTLLEVSIRK
GKQPLVMDRVYIGKGERDVVYKIKRRLRYEDLTPAAKTELPYVIEHIIKQDEKKYVDFFNKA
DSITTRMHQLELLPGVGKKMMWAIIEERKKRPFESFEDIAQRVKGIQRPEKLIVSRIIYEIK
NPQTKYKLFTA
11111111111111111000000000000000000000000000000000000000000000
00000000000000000000000000000000000000000000000000000000000011
00000000000000000000000000000000000000000000000000000000000000
00000000000
>DM_train1536
MKLIVGMTGATGAPLGVALLQALREMPNVETHLVMSKWAKTTIELETPYSARDVAALADFSH
NPADQAATISSGSFRTDGMIVIPCSMKTLAGIRAGYADGLVGRAADVVLKEGRKLVLVPREM
PLSTIHLENMLALSRMGVAMVPPMPAFYNHPETVDDIVHHVVARVLDQFGLEHPYARRWQGL
PQARNFSQENE
00000000000000000000000000000000000000000000000000000000000000
00000000000000000000000000000000000000000000000000000000000000

000000000000000000000000000000000000000000000000001111111111
11111111111
>DM_train1538
MPLPTFPALLFGLSGCLVDFGAQAATSDTPDDEHAQLTPGAQNALKALRDQGMPCAWIDELP
EALSTPLAAPVNDWMIAAPRPTAGWPQPDACWMALMALNVSQLEGCVLISGDPRLLQSGLNA
GLWTIGLASCGPLCGLSPSQWQALNNAEREQRRAQATLKLYSLGVHSVIDHLGELESCLADI
ALRRSKGEKP
111100000000000000000000000000000000000000000000000000000000
000000000000000000000000000000000000000000000000000000000000
000000000000000000000000000000000000000000000000000000000000
0000000000
>DM_train1539
GMTYKDYTGLDRTELLSKVRHMMSDKRFNHVLGVERAAIELAERYGYDKEKAGLAALLHDYA
KELSDDEFLRLIDKYQPDPDLKKWGNNIWHGLVGIYKIQEDLAIKDQDILAAIAKHTVGSAQ
MSTLDKIVYVADYIEHNRDFPGVEEARELAKVDLNKAVAYETARTVAFLASKAQPIYPKTIE
TYNAYIPYLD
111000000000000000000000000000000000000000000000000000000000
000000000000000000000000000000000000000000000000000000000000
000000000000000000000000000000000000000000000000000000000000
0000000000
>DM_train1540
GMTTTQPEAKDRISSLPVPDATQVPEGVRKLWAKAEANIGFVPNVFRAQAVNGEQFLAWWNY
FNLLLNKEGYLTNAERELVAVVVSGVNRCLYCAVSHGAALREFLGDPQKADAVAVNWRHADL
TEREQALAAYAEKLTRHPAEVTAADLEPLRAVGLDDHQIMELVQVIGMFNLTNRVSSALGFV
PNPEYYRQAR
111111111100000000000000000000000000000000000000000000000000
000000000000000000000000000000000000000000000000000000000000
000000000000000000000000000000000000000000000000000000000000
0000000000
>DM_train1541
MSLSLKIALISQNENLLNLFPKLALEKNFIPITKTASLTRASKIAFGLQDEVDAIISRGATS
DYIKKSVSIPSISIKVTRFDTMRAVYNAKRFGNELALIAYKHSIVDKHEIEAMLGVKIKEFL
FSSEDEITTLISKVKTENIKIVVSGKTVTDEAIKQGLYGETINSGEESLRRAIEEALNLIEV
RNEGHHHHHH
100000000000000000000000000000000000000000000000000000000000
000000000000000000000000000000000000000000000000000000000000
000000000000000000000000000000000000000000000000000000000000
0011111111
>DM_train1542
MAEICLITGTPGSGKTLKMVSMMANDEMFKPDENGIRRKVFTNIKGLKIPHTYIETDAKKLP
KSTDEQLSAHDMYEWIKKPENIGSIVIVDEAQDVWPARSAGSKIPENVQWLNTHRHQGIDIF
VLTQGPKLLDQNLRTLVRKHYHIASNKMGMRTLLEWKICADDPVKMASSAFSSIYTLDKKVY
DLYESAEVHT
000000000000000000000000000000000000000000000000000000000000

000000000000000000000000000000000000000000000000000000000000
000000000000000000000000000000000000000000000000000000000000
0000111111
>DM_train1543
DILECDYFDTVDISAAQKLQNGSYLFEGLLVPAILTGEYDFRILPDDSKQKVARHIRGCVCK
LKPCVRFCCPHDHIMDNGVCYDNMSDEELAELDPFLNVTLDDGSVSRRHFKNELIVQWDLPM
PCDGMFYLDNREEQDKYTLFENGTFFRHFDRVTLRKREYCLQHLTFADGNATSIRIAPHNCL
IVPSITGQT
000000000000000000000000000000000000000000000000000000000000
000000000000000000000000000000000000000000000000000000000000
000000000000000000000000000000000000000000000000000000000000
001111111
>DM_train1544
MRGSHHHHHHLVPRGSKTRGQIQVILGPMFSGKSTELMRRVRRFQIAQYKCLVIKYAKDTRY
SSSFCTHDRNTMEALPACLLRDVAQEALGVAVIGIDEGQFFPDIVEFCEAMANAGKTVIVAA
LDGTFQRKPFGAILNLVPLAESVVKLTAVCMECFREAAYTKRLGTEKEVEVIGGADKYHSVC
RLCYFKKAS
111111111111111111000000000000000000000000000000000000000001
111111111111100000000000000000000000000000000000000000000000
000000000000000000000000000000000000000000000000000000000000
000000111
>DM_train1545
SNAMWKRMTAKAEGLYIADTKSFVTKQMDKLDFDYGGIPGDLHFGLTKKAGAREPMFSRGTE
IFNRRQISIVSIEECNEIALKMGVPRILPEWLGANVAVSGMPDLTSLKEGSRIIFPSGAALL
CEGENDPCIQPGEVIQSYYPDQPKLASAFVRHALGIRGIVCIVERPGAVYTGDEIEVHSYQR
KVKRKAERV
111000000000000000000000000000000000000000000000000000000000
000000000000000000000000000000000000000000000000000000000000
000000000000000000000000000000000000000000000000000000000001
111111111
>DM_train1546
EAASSSDADGKEVGSSGEGNRATGGKWRRPSLAQQRARRAQLPPAFDVVHWNDEDISRGHLL
RVLHRDTFVVLDYHRQARMLTEEGNKAERVVSVMLPAVYTARFLAVLEGRSEKVEVHSRYTN
ATFTPNPAAPYTFTLKCTSTRPAQQKQQVAGEEGDETFEWTVEFDVAESLMLQRFLTQALHY
NTGFARTSV
111111111111111111111111100000000000000000000000000000000000
000000000000000000000000000000000000000000000000000000000000
000000000000000000000011111111111000000000000000000000000000
000000111
>DM_train1547
GMSLAPWRGAIAHALHRNRSLVYARYLQLATVQPNGRPANRTLVFRGFLEDTNQLRFITDTR
SAKADQIQQQPWAEICWYFPNTREQFRMAGDLTLISSDDSHQDLQPARIAMWQELSDAARLQ
FGWPYPGKPRIKESGAFEPSPPDPIEPVPNFCLLLLDPVQVDHLELRGEPQNRWLYHRNDQQ
EWSSEAINP

000000000000000000000000000000000000000000000000000000000000
000000000000000000000000000000000000000000000000000000000000
000000000011110000000000000000000000000000000000000000000000
000000000
>DM_train1548
MSAFFLNMEKSIITQKIIAKAFKDLMQSNAYHQISVSDIMQTAKIRRQTFYNYFQNQEELLS
WIFENDFAELINDNSDYYGWQNELLLLLRYLDENQIFYQKIFVIDKNFEHFFLIQWENLLDK
VIFDQEKKSDYHWSDLEKSFICRYNAAAICAITRESIIRGNSLEKLYSQIVNLLLAQIKIFE
SLEHHHHHH
111111111100000000000000000000000000000000000000000000000000
000000000000000000000000000000000000000000000000000000000000
000000000000000000000000000000000000000000000000000000000000
001111111
>DM_train1549
SETQSFVVSVAGSDRVGIVHDFSWALKNISANVESSRMACLGGDFAMIVLVSLNAKDGKLIQ
SALESALPGFQISTRRASSVAERHVSPDTREYELYVEGPDSEGIVEAVTAVLAKKGANIVEL
ETETLPAPFAGFTLFRMGSRVAFPFPLYQEVVTALSRVEEEFGVDIDLEEVVEGEDSEEDDD
SPNSPVGRH
100000000000000000000000000000000000000000000000000000000000
000000000000000000111110000000000000000000000000000000000000
000000000000000000000000000000000000000000000000001111111111
111111111
>DM_train1550
MSVEKIPGYTYGETENRAPFNLEDLKLLKEAVMFTAEDEEYIQKAGEVLEDQVEEILDTWYG
FVGSHPHLLYYFTSPDGTPNEKYLAAVRKRFSRWILDTSNRSYDQAWLDYQYEIGLRHHRTK
KNQTDNVESVPNIGYRYLVAFIYPITATMKPFLARKGHTPEEVEKMYQAWFKATTLQVALWS
YPYVKYGDF
111110000000000000000000000000000000000000000000000000000000
000000000000000000000000000000000000000000000000000000000000
000000000000000000000000000000000000000000000000000000000000
000000000
>DM_train1551
MARTEKIYIYGASGHGLVCEDVAKNMGYKECIFLDDFKGMKFESTLPKYDFFIAIGNNEIRK
KIYQKISENGFKIVNLIHKSALISPSAIVEENAGILIMPYVVINAKAKIEKGVILNTSSVIE
HECVIGEFSHVSVGAKCAGNVKIGKNCFLGINSCVLPNLSLADDSILGGGATLVKNQDEKGV
FVGVPAKRM
110000000001100000000000000000000000000000000000000000000000
000000000000000000000000000000000000000000000000000000000000
000000000000000000000000000000000000000000000000000000000000
000000000
>DM_train1552
MTAPHDPLDDIQADPWALWLSGRTRTALELPHYRRAAVLVALTREADPRVLLTVRSSELPTH
KGQIAFPGGSLDAGETPTQAALREAQEEVALDPAAVTLLGELDDVFTPVGFHVTPVLGRIAP
EALDTLRVTPEVAQIITPTLAELRAVPLVRERRTLPDGTEVPLYRYPWRGLDIWGMTARVLH

DLLEQGPG
1111000000000000000000011111111110000000000000000000000000111111
0000000000000000000000000000000000000000000000000000000000000000
0000000000000000000000000000000000000000000000000000000000000000
00001111
>DM_train1553
STNDNIKDLLDWYSSGSDTFTNSEVLDNSLGSMRIKNTDGSISLIIFPSPYYSPAFTKGEKV
DLNTKRIKKSQHTSEGTWIHFQISGVTNTEKLPTPIELPLKVKVHGKDSPLKYWPKFDKKQL
AISTLDFKIRHQLTQTHGLYRSSDKTGGYWKITMNDGSTYQSDLSKKFEYNTEKPPINIDEI
KTIEAEIN
1111000000000000000000000000000000000000000000000000000000000000
0000000000000000000000000000000000000000000000000000000000000000
0000000000000000000000000000000000000000000000000000000000000000
00000000
>DM_train1554
CSCSPVHPQQAFCNADVVIRAKAVSEKEVDSGNDIYGNPIKRIQYEIKQIKMFKGPEKDIEF
IYTAPSSAVCGVSLDVGGKKEYLIAGKAEGDGKMHITLCDFIVPWDTLSTTQKKSLNHRYQM
GCECKITRCPMIPCYISSPDECLWMDWVTEKNINGHQAKFFACIKRSDGSCAWYRGAAPPKQ
EFLDIEDP
0000000000000000000000000000000000000000000000000000000000000000
0000000000000000000000000000000000000000000000000000000000000000
0000000000000000000000000000000000000000000000000000000000001111
11111111
>DM_train1555
HMPGGSKKIGIFGGTFDPPHNGHLLMANEVLYQAGLDEIWFMPNQIPPHKQNEDYTDSFHRV
EMLKLAIQSNPSFKLELVEMEREGPSYTFDTVSLLKQRYPNDQLFFIIGADMIEYLPKWYKL
DELLNLIQFIGVKRPGFHVETPYPLLFADVPEFEVSSTMIRERFKSKKPTDYLIPDKVKKYV
EENGLYES
1111110000000000000000000000000000000000000000001111111111000000
0000000000000000000000000000000000000000000000000000000000000000
0000000000001111110000000000000000000000000000000000000000000000
00011111
>DM_train1556
GSHMSEQLTDQVLVERVQKGDQKAFNLLVVRYQHKVASLVSRYVPSGDVPDVVQEAFIKAYR
ALDSFRGDSAFYTWLYRIAVNTAKNYLVAQGRRPPSSDVDAIEAENFESGGALKEISNPENL
MLSEELRQIVFRTIESLPEDLRMAITLRELDGLSYEEIAAIMDCPVGTVRSRIFRAREAIDN
KVQPLIRR
0000000000000000000000000000000000000000000000000000000000000000
0000000000000000000000000000000001111111111111111111111111111100
0000000000000000000000000000000000000000000000000000000000000000
00000000
>DM_train1557
LTTISSILSLKREKPDNLAIILQIDFTKLKEEDSLIVVYNSLKALTIKFARLQFCFVDRNNY
VLDYGSVLHKIDSLDSISNLKSKSSSTQFSPIWLKNTLYPENIHEHLGIVAVSNSNMEAKKS

ILFQDYRCFTSFGRRRPNELKIKVGYLNVDYSDKIDELVEASSWTFVLETLCYSFGLSFDEH
DDDDEEDN
00000000000000000000000000000000000000000000000000000000000000
00000000000000000000000000000000000000000000000000000000000000
00000000000001111000000000000000100000000000000000000000000000
00111111
>DM_train1558
MAHHHHHHMTIRVMLQAMDQGHLLVNNVDKYVRAGRGVMVYIAFLSDRDSAPITDEALRHAV
GVLLHTKIFTHFSPEKMINQPQSLEECPEMDILIVPQASLGGKVKGRSVQFHQLVAKDVGAA
LYDRFCHFVRVARGVDESRVDANGAPRSEGDAPKAEGWIKYNSRVISGTFGNRQGLRFESEG
PFTHMFDI
11111111000000000000000000000000000000000000000000000000000000
00000000000000000000000000000000000000000000000000000000000000
00000000000000000000000000000000000000000000000000000000000000
00000000
>DM_train1559
MNHLISGFENFEKKINYRFKNKAYLLQAFTHASYHYNTITDCYQRLEFLGDAILDYLITKHL
YEDPRQHSPGVLTDLRSALVNNTIFASLAVKYDYHKYFKAVSPELFHVIDDFVQFQLEKNEM
QGMDSELRRSEEDEEKEEDIEVPKAMGDIFESLAGAIYMDSGMSLETVWQVYYPMMRPLIEK
FSANVPRS
10000000000000000000000000000000000000000000000000000000000000
00000000000000000000000000000000000000000000000000000000000111
11111111111111111000000000000000000000000000000000000000000000
00001111
>DM_train1560
GSMKNPQSKADDILQCARTLIIRGGYNSFSYADISQVVGIRNASIHHHFPSKSDLVCKLVSQ
YRQEAEAGIAELEKNISDPLEQLRAYIGYWEGCIADATHPFCVCALLASEIPVLPETVVLEV
RAHFRSLSDWLTAVLERGIAQGRLVLTGTARANAEIFMATVHGAMLSARAHGDAATFGAITR
PMLERITA
11110000000000000000000000000000000000000000000000000000000000
00000000000000000000000000000000000000000000000000000000000000
00000000000000000000000000000000000000000000000000000000000000
00000000
>DM_train1561
SIEFSTAITRQIVLDTETTGMNQIGAHYEGHKIIEIGAVEVVNRRLTGNNFHVYLKPDRLVD
PEAFGVHGIADEFLLDKPTFAEVADEFMDYIRGAELVIHNAAFDIGFMDYEFSLLKRDIPKT
NTFCKVTDSLAVARKMFPGKRNSLDALCARYEIDNSKRTLHGALLDAQILAEVYLAMTGGQT
SMATRESG
11111110000000000000000000000000000000000000000000000000000000
00000000000000000000000000000000000000000000000000000000000000
00000000000000000000000000000000000000000000000000000000000111
11111111
>DM_train1563
MEERVLGMPAVESNMFPLGKQAPPFALTNVIDGNVVRLEDVKSDAATVIMFICNHCPFVKHV

QHELVRLANDYMPKGVSFVAINSNDAEQYPEDSPENMKKVAEELGYPFPYLYDETQEVAKAY
DAACTPDFYIFDRDLKCVYRGQLDDSRPNNGIPVTGESIRAALDALLEGRPVPEKQKPSIGC
SIKWKPSA
111110000000000000000000000000000000000000000000000000000000
000000000000000000000000000000000000000000000000000000000000
000000000000000000000000000000000000000000000000000000000000
00000000
>DM_train1564
GPLGSPEFIILQTYRAIADYEKTSGSEMALSTGDVVEVVEKSESGWWFCQMKAKRGWIPASF
LEPLDSPDETEDPEPNYAGEPYVAIKAYTAVEGDEVSLLEGEAVEVIHKLLDGWWVIRKDDV
TGYFPSMYLQKSGQDVSQAQRQIKRGAPPRRSSIRNAHSIHQRSRKRLSQDAYRRNSVRFLQ
QRRRQAR
111111111000000000000000000000000000000000000000000000000000
000000000000000000000000000000000000000000000000000000000000
000000000000000000000000000000000000000000000000000000000001
1111111
>DM_train1565
SQDASDGLQRLHMLQISYFRDPYHVWYQGNASLGGHLTHVLEGPDTNTTIIQLQPLQEPESW
ARTQSGLQSYLLQFHGLVRLVHQERTLAFPLTIRCFLGCELPPEGSRAHVFFEVAVNGSSFV
SFRPERALWQADTQVTSGVVTFTLQQLNAYNRTRYELREFLEDTCVQYVQKHISAENTKGSQ
TSRSYTS
111111100000000000000000000000000000000000000000000000000000
000000000000000000000000000000000000000000000000000000000000
000000000000000000000000000000000000000000000000000111111111
1111111
>DM_train1566
MKDDPKVHLEAKELWDQFHKRGTEMVITKSGRRMFPPFKVRCSGLDKKAKYILLMDIIAADD
CRYKFHNSRWMVAGKADPEMPKRMYIHPDSPATGEQWMSKVVTFHKLKLTNNISDKHGFTIL
NSMHKYQPRFHIVRANDILKLPYSTFRTYLFPETEFIAVTAYQNDKITQLKIDNNPFAKGFR
DTGNGRR
111000000000000000000000000000000000000000000000000000000000
000000000000000000000000000000000000000000000000000000000000
000000000000000000000000000000000000000000000000000000000000
0111111
>DM_train1567
GMQTPALIIVTGHPATGKTTLSQALATGLRLPLLSKDAFKEVMFDGLGWSDREWSRRVGATA
IMMLYHTAATILQSGQSLIMESNFRVDLDTERMQNLHTIAPFTPIQIRCVASGDVLVERILS
RIAQGARHPGHCDDRSPADLELVRSRGDIPPLPLGGPLLTVDTTFPEQIDMNAIVQWVRQHL
QSGTAFL
000000000000000000000000000000000000000000000000000000000000
000000000000000000000000000000000000000000000000000000000000
000000111111110000000000000000000000000000000000000000000000
0000111
>DM_train1568

TTHFTVADRWGNVVSYTTTIEQLFGTGIMVPDYGVILNNELTDFDAIPGGANEVQPNKRPLS
SMTPTILFKDDKPVLTVGSPGGATIISSVLQTILYHIEYGMELKAAVEEPRIYTNSMSSYRY
EDGVPKDVLSKLNGMGHKFGTSPVDIGNVQSISIDHENGTFKGVADSSRNGAAIGINLKRKL
EHHHHHH
00000000000000000000000000000000000000000000000000000000000000
00000000000000000000000000000000000000000000000000000000000000
00000011111111110000000000000000000000000000000000000000000001
1111111
>DM_train1569
GMSLIEIRKRTLIVETTYHENGPAPAQPLKLAASCAVIRNPYAGRYEPDLMPFMAELRSLGT
LLATELVDTLGKDNIEVYSKAAIVGVDGEMEHGAVWHEAGGWAMRSVLGEPKAMVPAVKAVA
TAGYRMMVPVHYIHASYVRSHFNSIEIGIQDAPRPREILFALVMGTGARVHARLGGLTKEAV
SVHDGQR
11000000000000000000000000000000000000000000000000000000000000
00000000000000000000000000000000000000000000000000000000000000
00000000000000000000000000000000000000000000000000000000000000
0000000
>DM_train1570
MKDVSKNQEENISDTALTNELIHLLGHSRHDWMNKLQLIKGNLSLQKYDRVFEMIEEMVIDA
KHESKLSNLKTPHLAFDFLTFNWKTHYMTLEYEVLGEIKDLSAYDQKLAKLMRKLFHLFDQA
VSRESENHLTVSLQTDHPDRQLILYLDFHGAFADPSAFDDIRQNGYEDVDIMRFEITSHECL
IEIGLD
11111111111100000000000000000000000000000000000000000000000000
00000000000000000000000000000000000000000000000000000000000000
00000000000000000000000000000000000000111111111100000000000000
000000
>DM_train1571
MLEKVYVALIHYPIKGKDGSIISTAVTNLDVHDIARTARTYNLKGYYIVTNLRAQQDMVSKM
LKFWREGFGSRYNPSRAESLKLVKLKSYLEDVLEDIESVEGERPLIFFTSAKKRENDISFEE
GRRIIIETEKPVLILLGTGWGLPDEILEISDYVLEPIRAQSDFNHLSVRAAAAIIIDRLIGE
NYARRD
00000000000000000000000000000000000000000000000000000000000000
00000000000000000000000000000000000000000000000000000000000000
00000000000000000000000000000000000000000000000000000000000000
001111
>DM_train1572
GRGTKQVYIIHGYRASSTNHWFPWLKKRLLADGVQADILNMPNPLQPRLEDWLDTLSLYQHT
LHENTYLVAHSLGCPAILRFLEHLQLRAALGGIILVSGFAKSLPTLQMLDEFTQGSFDHQKI
IESAKHRAVIASKDDQIVPFSFSKDLAQQIDAALYEVQHGGHFLEDEGFTSLPIVYDVLTSY
FSKETR
11100000000000000000000000000000000000000000000000000000000000
00000000000000000000000000000000000000000000000000000000000000
00000000000000000000000000000000000000000000000000000000000000
000111

>DM_train1573
MVYPIRLYGDPVLRRKARPVEDFSGIKRLAEDMLETMFEAKGVGLAAPQIGLSQRLFVAVEY
ADEPEGEEERPLRELVRRVYVVANPVITYREGLVEGTEGCLSLPGLYSEEVPRAERIRVEYQ
DEEGRGRVLELEGYMARVFQHEIDHLDGILFFERLPKPKREAFLEANRAELVRFQKEARALL
KELSQG
100000000000000000000000000000000000000000000000000000000000
111111111111111000000000000000000000000000000000000000000000
000000000000000000000000000000000000000000000000000000000111
111111
>DM_train1574
GSPEFGTRDRMLVLVLGDLHIPHRCNSLPAKFKKLLVPGKIQHILCTGNLCTKESYDYLKTL
AGDVHIVRGDFDENLNYPEQKVVTVGQFKIGLIHGHQVIPWGDMASLALLQRQFDVDILISG
HTHKFEAFEHENKFYINPGSATGAYNALETNIIPSFVLMDIQASTVVTYVYQLIGDDVKVER
IEYKKS
111111111100000000000000000000000000000000000000000000000000
000000000000000000000000000000000000000000000000000000000000
000000000000000000000000000000000000000000000000000000000000
000000
>DM_train1575
MYQDLIRNELNEAAETLANFLKDDANIHAIQRAAVLLADSFKAGGKVLSCGNGGSHCDAMHF
AEELTGRYRENRPGYPAIAISDVSHISCVGNDFGFNDIFSRYVEAVGREGDVLLGISTSGNS
ANVIKAIAAAREKGMKVITLTGKDGGKMAGTADIEIRVPHFGYADRIQEIHIKVIHILIQLI
EKEMVK
000000000000000000000000000000000000000000000000000000000000
000000000000000000001111111111111110000000000000000000000000
000000000000000000000000000000000000000000000000000000000000
000000
>DM_train1576
MGSDKIHHHHHHMVMEYELRTPLVKDQILKLKVGDVVYITGEIFTARDEAHARALEWMEEGK
ELPFSFDKGVVYHCGPLVKKNDEWRVVSAGPTTSARMNPFTPKILEKVECMGIIGKGGMSEE
VVEAMRGKAAYFAFTGGAGALAAMSIKKVKGVVWEDLGMPEAVWLLEVERFGPCIVAIDAHG
NSLYRR
111111110000000000000000000000000000000000000000000000000000
000000000000000000000000000000000000000000000000000000000000
000000000000000000000000000000000000000000000000000000000000
000000
>DM_train1577
MAPLGGAPRLVLLFSGKRKSGKDFVTEALQSRLGADVCAVLRLSGPLKEQYAQEHGLNFQRL
LDTSTYKEAFRKDMIRWGEEKRQADPGFFCRKIVEGISQPIWLVSDTRRVSDIQWFREAYGA
VTQTVRVVALEQSRQQRGWVFTPGVDDAESECGLDNFGDFDWVIENHGVEQRLEEQLENLIE
FIRSRL
000000000000000000000000000000000000000000000000000000000011
111110000000000000000000000000000000000000000000000000000000
000000000000000000000000000000000000000000000000000000000000

000000
>DM_train1578
KVEQAVETEPEPELRQQTEWQSGQRWELALGRFWDYLRWVQTLSEQVQEELLSSQVTQELRA
LMDETMKELKAYKSELEEQLTPVAEETRARLSKELQAAQARLGADMEDVCGRLVQYRGEVQA
MLGQSTEELRVRLASHLRKLRKRLLRDADALQKCLAVYQAGAREGAERGLSAIRERLGPLVE
QGRVR
111111111111111111111000000000000000000000000000000000000000
000000000000000000111111111100000000000000000000000000000000
000000000000000000000000000000000000001111111111111111111111
11111
>DM_train1579
MILRTSEETISTVQEKQQNISPLVRERGPQRVAAHITGTRGRSNTLSSPNSKNEKALGRKIN
SWESSRSGHSFLSNLHLRNGELVIHEKGFYYIYSQTYFRFQEEIKENTKNDKQMVQYIYKYT
SYPAPILLMKSARNSCWSKDAEYGLYSIYQGGIFELKENDRIFVSVTNEHLIDMDHEASFFG
AFLVG
111111111111111111111111111100000000000111111111100000000000
000000000000000000000000000000000000000000000000000000000000
000000000000000000000000000000000000000000000000000000000000
00000
>DM_train1581
FPTIPLSRLADNAWLRADRLNQLAFDTYQEFEEAYIPKEQIHSFWWNPQTSLCPSESIPTPS
NKEETQQKSNLELLRISLLLIQSWLEPVQFLRSVFANSLVYGASDSNVYDLLKDLEEGIQTL
MGRLEDGSPRTGQIFKQTYSKFDTNSHNDDALLKNYGLLYCFNKDMSKVSTYLRTVQCRSVE
GSCGF
000000000000000000000000000000000000000000000000000000000000
000000000000000000000000000000000000000000000000000000000000
000011111111111111111111111111000000000000000000000000000011
10000
>DM_train1582
GMTARNILSYSYEEYVEKITAFHGYPAPGVLIGGFMVDLAVKNLPEGILYDAICETRTCLPD
AVQLLTPCTFGNGWLTVLPMGLFAVSLYDKFTGEGVRVFLDVEKMGPWQEIRNWFLKLKTKK
EQDSERLFKEIREAGPDILELRNVKLKPGFLEKKHKGKIVLCPQCREAYPAQDGELCLSCQG
GSPYL
100000000000000000000000000000000000000000000000000000000000
000000000000000000000000000000000000000000000000000000000111
111000000000000000000000000000000000000000000000000000000000
00000
>DM_train1584
MVQAQTRDQIVAAADELFYRQGFAQTSFVDISAAVGISRGNFYYHFKTKDEILAEVIRLRLA
RTAQMLADWQGTGDSPRARIASFIDLMIMNRAKITRYGCPVGSLCTELSKLDHAAQGQANGL
FTLFRDWLQRQFAEAGCTTEAPALAMHLLARSQGAATLAQSFHDEGFLRSEVADMHRWLDNT
LPMTT
110000000000000000000000000000000000000000000000000000000000
000000000000000000000000000000000000000000000000000000000000

00000000000000000000000000000000000000000000000000000000000000
00001
>DM_train1585
GATDSATAAPAAAATTQVQKEAADVLQVAVQGANAMRDIQFARLALFHGQPDSAKKLTDDAA
ALLAADDASWAKFVKTDAKAKMIADRYVIINASIALSEDYVATPEKESAIQSANEKLAKGDQ
KGAIDTLRLAGIGVIENQYLMPLNQTRKAVAQSQELLKAGKYYEANLVLKGAEEGIVVDSEM
LVAGN
11111111111110000000000000000000000000000000000000000000000000
00000000000000000000000000000000000000000000000000000000000000
00000000000000000000000000000000000000000000000000000000000000
00111
>DM_train1586
MSRQANRGTESKKMSSELFTLTYGALVTQLCKDYENDEDVNKQLDKMGFNIGVRLIEDFLAR
SNVGRCHDFRETADVIAKVAFKMYLGITPSITNWSPAGDEFSLILENNPLVDFVELPDNHSS
LIYSNLLCGVLRGALEMVQMAVEAKFVQDTLKGDGVTEIRMRFIRRIEDNLPAGEEAAAWSH
PQFEK
11111111111100000000000000000000000000000000000000000000000000
00000000000000000000000000000000000000000000000000000000000000
00000000000000000000000000000000000000000000000111111111111111
11111
>DM_train1587
MKVGIIMGSVRAKRVCPEIAAYVKRTIENSEELIDQKLKIQVVDLQQIALPLYEDDDELIPA
QIKSVDEYADSKTRSWSRIVNALDIIVFVTPQYNWGYPAALKNAIDRLYHEWHGKPALVVSY
GGHGGSKCNDQLQEVLHGLKMNVIGGVAVKIPVGTIPLPEDIVPQLSVHNEEILQLLASCIE
TTRNK
10000000000000000000000000000011111111000000000000000000000000
00000000000000000000000000000000000000000000000000000000000000
00000000000000000000000000000000000000000000000000000000000000
11111
>DM_train1588
DEPNQPSTPEAVTKTVTIDASKYETWQYFSFSKGEVVNVTDYKNDLNWDMALHRYDVRLNCG
ESGKGKGGAVFSGKTEMDQATTVPTDGYTVDVLGRITVKYEMGPDGHQMEYEEQGFSEVITG
KKNAQGFASGGWLEFSHGPAGPTYKLSKRVFFVRGADGNIAKVQFTDYQDAELKKGVITFTY
TYPVK
11111111100000000000000000000000000000000000000000000000000000
00000000000000000000000000000000000000000000000000000000000000
00000000000000000000000000000000000000000000000000000000000000
00000
>DM_train1589
MGVSLSKGGNVSLTKEAPGLTAVIVGLGWDIRTTTGTDFDLDASALLLNSGGKVASDAHFIF
FNNLKSPDGSVEHTGDNITGEGEGDDEQIKINLATVPADIEKIVFPVSIYDAENRQQSFGQV
RNAFIRVVNQAGEAEIARYDLSEDASTETAMVFGELYRHGAEWKFRAIGQGYASGLRGIAQD
FGVNV
11111111111100001100000000000000000000000000000000000000000000

000000000000000000000000000000000000000000000000000000000000
000000000000000000000000000000000000000000000000000000000000
00000
>DM_train1591
MSLTLADKVVYEEEIQKSRFIAKAAPVASEEEALAFLAENREPEATHNGHAYKIGLLYRFSD
DGEPSGTAGRPILHAIEAQGLDRVAVLVVRYFGGVKLGAGGLVRAYGGVAAEALRRAPKVPL
VERVGLAFLVPFAEVGRVYALLEARALKAEETYTPEGVRFALLLPKPEREGFLRALLDATRG
QVALE
100000000000000000000000000000000000000000000000000000000000
000000000000000000000000000000000000000000000000000000000000
000000000000000000000000000000000000000000000000000000000000
00000
>DM_train1592
GAPEVSHVRGVTVRMETPEAILFSPGETFSTNVSIHAIAHDDQTYSMDVVWLRFDVPTSCAE
MRIYESCLYHPQLPECLSPADAPCAASTWTSRLAVRSYAGCSRTNPPPRCSAEAHMEPVPGL
AWQAASVNLEFRDASPQHSGLYLCVVYVNDHIHAWGHITISTAAQYRNAVVEQPLDIEGRGH
HHHHH
111111100000000000000000000000000000000000000000000000000000
000000000000000000000000000000000000000000000000000000000000
000000000000000000000000000000000000000000000000000000000001
11111
>DM_train1593
GMQTTEIDLRLTEVSQQLTMVLVPGLRDSDDEHWQSHWERRFPHWQRIRQREWYQADLDRWV
LAIRRELSVCTQPVILIGHSFGALAACHVVQQGQEGIAGVMLVAPAEPMRFEIDDRIQASPL
SVPTLTFASHNDPLMSFTRAQYWAQAWDSELVDVGEAGHINAEAGFGPWEYGLKRLAEFSEI
LIPNR
111100000000000000000000000000000000000000000000000000000000
000000000000000000000000000000000000000000000000000000000000
000000000000000000000000000000000000000000000000000000000000
00011
>DM_train1594
MTSRGDSREKILHTASRLSQLQGYHATGLNQIVKESGAPKGSLYHFFPNGKEELAIEAVTYT
GKIVEHLIQQSMDESSDPVEAIQLFIKKTASQFDNTESIKGIPVGLLASETALISEPLRTVC
MKVFKSWEAVFARKLMENGFAEEEANQLGTLINSMIEGGIMLSLTNKDKTPLLLIAEQIPVL
VRKKG
111100000000000000000000000000000000000000000000000000000000
000000000000000000000000000000000000000000000000000000000000
000000000000000000000000000000000000000000000000000000000000
00111
>DM_train1595
MENREPPLLPARWSSAYVSYWSPMLPDDQLTSGYCWFDYERDICRIDGLFNPWSERDTGYRL
WMSEVGNAASGRTWKQKVAYGRERTALGEQLCERPLDDETGPFAELFLPRDVLRRLGARHIG
RRVVLGREADGWRYQRPGKGPSTLYLDAASGTPLRMVTGDEASRASLRDFPNVSEAEIPDAV
FAAKR

111100000000000000000000000000000000000000000000000000000000
000000000000000000000000000000000000000000000000000000000000
000000000000000000000000000000000000000000000000000000000000
00011
>DM_train1597
ADTLLILGDSLSAGYRMSASAAWPALLNDKWQSKTSVVNASISGDTSQQGLARLPALLKQHQ
PRWVLVELGGNDGLRGFQPQQTEQTLRQILQDVKAANAEPLLMQIRLPANYGRRYNEAFSAI
YPKLAKEFDVPLLPFFMEEVYLKPQWMQDDGIHPNRDAQPFIADWMAKQLQPLVNHDSLEHH
HHHH
000000000000000000000000000000000000000000000000000000000000
000000000000000000000000000000000000000000000000000000000000
000000000000000000000000000000000000000000000000000001111111
1111
>DM_train1598
MKYKEPFGVKLDFETGIIENAKKSVRRLSDMKGYFIDEEAWKKMVEEGDPVVYEVYAIEQEE
KEGDLNFATTVLYPGKVGNEFFMTKGHYHSKIDRAEVYFALKGKGGMLLQTPEGEARFIEME
PGTIVYVPPYWAHRTINTGDKPFIFLALYPADAGHDYGTIAEKGFSKIVVEENGKVVVKDNP
KWRM
110000000000000000000000000000000000000000000000000000000000
000000000000000000000000000000000000000000000000000000000000
000000000000000000000000000000000000000000000000000000000001
1111
>DM_train1599
GMPTFNPELHAQTLNSERAYFVQPDADPAFTPHIGALVEMLTYARLTTLQAVEGLPEDQLWA
TAPGFANSIGTLLAHIAAVERVYHVLSFQGRDVTPEDDGAAYWGLTMGKEGTAPARLPTLDE
LRAELADARAETLRVFAAKDDAWLAEPLGPGWANQHWAWFHVMEDEVNHRGQLRLLRQVLAP
EEGG
110000000000000000000000000000000000000000000000000000000000
000000000000000000000000000000000000000000000000000000000000
000000000000000000000000000000000000000000000000000000000001
1111
>DM_train1600
GMTSPSIESGARERTRRAILDAAMLVLADHPTAALGDIAAAAGVGRSTVHRYYPERTDLLRA
LARHVHDLSNAAIERADPTSGPVDAALRRVVESQLDLGPIVLFVYYEPSILADPELAAYFDI
GDEAIVEVLNRASTERPEYPPGWARRVFWALMQAGYEAAKDGMPRHQIVDAIMTSLTSGIIT
LPRT
111111111111000000000000000000000000000000000000000000000000
000000000000000000000000000000000000000000000000000000000000
000000000000011111000000000000000000000000000000000000000000
0011
>DM_train1601
MTNATIAPTTTAAPVTKSVDAPTADENTPLFSPSLISPDVLAVLPADYTIRPLCRSDYKRGY
LDVLRVLTTVGDINEEQWNSRYEWIRARSDEYYLLVVCDGEGRIVGTGSLVVERKFIHSLGM
VGHIEDIAVEKGQQGKKLGLRIIQALDYVAEKVGCYKTILDCSEANEGFYIKCGFKRAGLEM

AHYY
111111111111111111111111000000000000000000000000000000000000
000000000000000000000000000000000000000000000000000000000000
000000000000000000000000000000000000000000000000000000000000
0000
>DM_train1602
GDRFYDLISALHKSVRGSAPDAALYWYARILTAGGDPLYVARRLLAIASEDVGNADPRAMQV
ALAAWDCFTRVGAYEGERAIAQAIIYLSVAPKSNAVYTAFNTAKQQAKDLPDYDVPPHLRNA
PTNLMKELGYGAEYRYAHDEPNAYAAGENYFPPELKDTQYYFPTNRGMEIQIKEKLERLREQ
DKST
000000000000000000000000000000000000000000000000000000000000
000000000000000000000000000000000000000000000000000000000000
000011111111111111111111000000000000000000000000000000000011
1111
>DM_train1603
MATEGMILTNHDHQIRVGVLTVSDSCFRNLAEDRSGINLKDLVQDPSLLGGTISAYKIVPDE
IEEIKETLIDWCDEKELNLILTTGGTGFAPRDVTPEATKEVIEREAPGMALAMLMGSLNVTP
LGMLSRPVCGIRGKTLIINLPGSKKGSQECFQFILPALPHAIDLLRDAIVKVKEVHDRSHHH
HHH
111111111111000000000000000000000000000000000000000000000000
000000000000000000000000000000000000000000000000000000000000
000000000000000000000000000000000000000000000000000000011111
111
>DM_train1604
GSHMVPISFVFNRFPRMVRDLAKKMNKEVNFIMRGEDTELDRTFVEEIGEPLLHLLRNAIDH
GIEPKEERIAKGKPPIGTLILSARHEGNNVVIEVEDDGRGIDKEKIIRKAIEKGLIDESKAA
TLSDQEILNFLFVPGFSTKEKVSEVSGRGVGMDVVKNVVESLNGSISIESEKDKGTKVTIRL
PLT
000000000000000000000000000000000000000000000000000000000000
000000000000000000000000000000000000000000000000000000000000
000000000000000001111111111110000000000000000000000000000000
000
>DM_train1605
MGSSSSGRENLYFQGHMNHSIPTLRTERLTLRPLAMADFPAYRDFMASPRSTGVGGPYDLPS
TWGVFCHDLANWHFFGHGALMIDLGETGECIGQIGINHGPLFPEKELGWLLYEGHEGRGYAA
EAAVALRDWAFETLNLPTLVSYVSPQNRKSAAVAERIGGTLDPLAPRSDPEDLVYRYHQVKT
AGS
111111000000011111100000000000000000000000000000000000000000
000000000000000000000000000000000000000000000000000000000000
000000000000000000000000000000000000000000000000000000000111
111
>DM_train1606
LERKEILFIPSENEKISKQFHLRYDIVRDRYIRVSDNNTNISGWENGVWKMESIFRKVEKDW
NMVYLARKEGSSFAYISWKFECGSAGLKVDTVSIRTSSQSFESGSVRWKLRSETAQVNLLGD

KNLRSYNDFSGATEVTLEAELSRGDGDVAWQHTQLFRQSLNDSGENGLEIIITFNDLLEHHH
HHH
100000000000000000000000000000000000000000000000000000000000000
000000000000000000000000000000000000000000000000000000000000000
000000000000000000000000000000000000000000000000000000000011111
111
>DM_train1608
MAFSVNYDSSFGGYSIHDYLGQWASTFGDVNHTNGNVTDANSGGFYGGSLSGSQYAISSTAN
QVTAFVAGGNLTYTLFNEPAHTLYGQLDSLSFGDGLSGGDTSPYSIQVPDVSFGGLNLSSLQ
AQGHDGVVHQVVYGLMSGDTGALETALNGILDDYGLSVNSTFDQVAAATAVGVQHADSPELL
AA
000000000000000000000000000000000000000000000000000000000000000
000000000000000000000000000000000000000000000000000000000000000
000000000000000000000000000000000000000000000000001111111111111
11
>DM_train1609
MANSGCKDVTGPDEESFLYFAYGSNLLTERIHLRNPSAAFFCVARLQDFKLDFGNSQGKTSQ
TWHGGIATIFQSPGDEVWGVVWKMNKSNLNSLDEQQGVKSGMYVVIEVKVATQEGKEITCRS
YLMTNYESAPPSPQYKKIICMGAKENGLPLEYQEKLKAIEPNDYTGKVSEEIEDIIKKGETQ
TL
111111111111100000000000000000000000000000000000000000000000000
000000000000000000000000000000000000000000000000000000000000000
000000000000000000000000000000000000000000000000000000000001111
11
>DM_train1610
GMNLDSYERTGLRVSLDLVNIATPGSRRGTPHTGGCVIEDLHDLLKDDPASVAQLGDDHVEG
FVELARLLHTAIDALSNGQVATAATALNHLLRKHPATPELAQDPDGTWRLHHHPLDAELVPM
WTAICAEGLAREIGHQNVRRFGICNAHRCDRVYFDTSRNGTRQYCSLACQNRVKAAAFRERR
AT
100000000000000000000000000000000000000000000000000000000000000
000000000000000000000000000000000000000000000000000000000000000
000000000000000000000000000000000000000000000000000000000000001
11
>DM_train1611
MLGTSFNNFGISLSHKRYFSGKVDEIIRCTMGKRIVKISSTKINTSILSSVSEQIGENITDW
KNDEKKVYVSRVVNQCIDKFCAEHSRKIGDNLRKQIFKQVEKDYRISLDINAAQSSINHLVS
GSSYFKKKMDELCEGMNRSVKNDTTSNVANLISDQFFEKNVQYIDLKKLRGNMSDYITNLES
PF
111111100000000000000000000000000000000000000000000000000000000
000000000000000000000000000000000000000000000000000000000000000
000000000000000000000000000000000000000000000000000000000000000
00
>DM_train1612
MGSDKIHHHHHHMFKGIIAALWDMDSIGEIEPDVVFLLKSDILNLKFHLKILKDRGKTVFVD

MDFVNGLGEGEEAILFVKKAGADGIITIKPKNYVVAKKNGIPAVLRFFALDSKAVERGIEQI
ETLGVDVVEVLPGAVAPKVARKIPGRTVIAAGLVETEEEAREILKHVSAISTSSRILWKMKF
LP
111111111111000000000000000000000000000000000000000000000000
000000000000000000000000000000000000000000000000000000000000
000000000000000000000000000000000000000000000000000000000011
11
>DM_train1613
MTAPSCAFPVQFRQPSVSGLSQITKSLYISNGVAANNKLMLSSNQITMVINVSVEVVNTLYE
DIQYMQVPVADSPNSRLCDFFDPIADHIHSVEMKQGRTLLHCAAGVSRSAALCLAYLMKYHA
MSLLDAHTWTKSCRPIIRPNSGFWEQLIHYEFQLFGKNTVHMVSSPVGMIPDIYEKEVRLMI
PL
111111111111111110000000000000000000000000000000000000000000
000000000000000000000000000000000000000000000000000000000000
000000000000000000000000000000000000000000000000000001111111
11
>DM_train1614
MADDDDKAPPTLWSRVTKFGSGWGFWVSPTVFITTTHVVPTGVKEFFGEPLSSIAIHQAGEF
TQFRFSKKMRPDLTGMVLEEGCPEGTVCSVLIKRDSGELLPLAVRMGAIASMRIQGRLVHGQ
SGMLLTGANAKGMDLGTIPGDCGAPYVHKRGNDWVVCGVHAAATKSGNTVVCAVQAGEGETA
LE
111111100000000000000000000000000000000000000000000000000000
000000000000000000000000000000000000000000000000000000000000
000001111111111100000000000000000000000000000000000000000000
00
>DM_train1615
GMSRDRLYTWAGLWRSPSSSWEALRLEDDQAESQLRAPDERSGLPYQLDYRLRWDADWHLRE
AVFHVESETGVRKLHLLADGRGHWQDGDGEALPAFDGCLDIDIWPSPFTNTFPIRRLGLADG
QRAEIRALYIEAPALEPRSMRQAYTRLDASHYLYENLEGSAFKAVLLVDEQGLVIDYPGLFQ
RL
111100000000000000000000000000000000000000000000000000000000
000000000000000000000000000000000000000000000000000000000000
000000000000000000000000000000000000000000000000000000000000
00
>DM_train1616
KTFIIGISGVTNSGKTTLAKNLQKHLPNCSVISQDDFFKPESEIETDKNGFLQYDVLEALNM
EKMMSAISCWMESARHSVVSTDQESAEEIPILIIEGFLLFNYKPLDTIWNRSYFLTIPYEEC
KRRRSTRVYQPPDSPGYFDGHVWPMYLKYRQEMQDITWEVVYLDGTKSEEDLFLQVYEDLIQ
EL
000000000000000000000000000000000000000000000000000000000000
000000000000000000000111110000000000000000000000000000000000
000000000000000000000000000000000000000000000000000000000000
00
>DM_train1617

MSKAGASLATCYGPVSADVMAKAENIRLLILDVDGVLSDGLIYMGNNGEELKAFNVRDGYGI
RCALTSDIEVAIITGRKAKLVEDRCATLGITHLYQGQSNKLIAFSDLLEKLAIAPENVAYVG
DDLIDWPVMEKVGLSVAVADAHPLLIPRADYVTRIAGGRGAVREVCDLLLLAQGKLDEAKGQ
SI
111111000000000000000000000000000000000000000000000000000000
000000000000000000000000000000000000000000000000000000000000
000000000000000000000000000000000000000000000000000000111111
11
>DM_train1618
AHHHHHHMVHITLDRNTANSWLIISKDRRQVRMGDTHQNVSDNKERFSNYPMVLGAQRFSSG
KMYWEVDVTQKEAWDLGVCRDSVQRKGQFSLSPENGFWTIWLWQDSYEAGTSPQTTLHIQVP
PCQIGIFVDYEAGVVSFYNITDHGSLIYTFSECVFAGPLRPFFNVGFNYSGGNAAPLKLCPL
KM
100000000000000000000000000000000000000000000000000000000000
000000000000000000000000000000000000000000000000000000000000
000000000000000000000000000000000000000000000000000000000000
11
>DM_train1619
AAIGSCSKEYRVLLGQLQKQTDLMQDTSRLLDPYIRIQGLDVPKLREHCRERPGAFPSEETL
RGLGRRGFLQTLNATLGCVLHRLADLEQRLPKAQDLERSGLNIEDLEKLQMARPNILGLRNN
IYCMAQLLDNSDTAEPTKAGRGASQPPTPTPASDAFQRKLEGCRFLHGYHRFMHSVGRVFSK
W
111000000000000000000000000000000000000000000000000000000000
000000000000000000000000000000000000000000000000000000000000
000000000011111111111111111111000000000000000000000000000000
0
>DM_train1620
MAFTEHSPLTPHRRDLCSRSIWLARKIRSDLTALTESYVKHQGLNKNINLDSADGMPVASTD
QWSELTEAERLQENLQAYRTFHVLLARLLEDQQVHFTPTEGDFHQAIHTLLLQVAAFAYQIE
ELMILLEYKIPRNEADGMPINVGDGGLFEKKLWGLKVLQELSQWTVRSIHDLRFISSHQTGI
P
111111111110000000000000000000000000000000011111111111111111
111000000000000000000000000000001111100000000000000000000000
000000000000011111111111111100000000000000000000000000111111
1
>DM_train1621
MPIASYAQELKLALHQYPNFPSEGILFEDFLPIFRNPGLFQKLIDAFKLHLEEAFPEVKIDY
IVGLESRGFLFGPTLALALGVGFVPVRKAGKLPGECFKATYEKEYGSDLFEIQKNAIPAGSN
VIIVDDIIATGGSAAAAGELVEQLEANLLEYNFVMELDFLKGRSKLNAPVFTLLNAQKEALK
K
111100000000000000000000000000000000000000000000000000000000
000000000000000000000000000000000000000001111000000000000000
000000000000000000000000000000000000000000000000000011111111
1

>DM_train1622
MYKILEIADVVKVPPEEFGKDLKETVKKILMEKYEGRLDKDVGFVLSIVDVKDIGEGKVVHG
DGSAYHPVVFETLVYIPEMYELIEGEVVDVVEFGSFVRLGPLDGLIHVSQIMDDYVSYDPKR
EAIIGKETGKVLEIGDYVRARIVAISLKAERKRGSKIALTMRQPYLGKLEWIEEEKAKKQNQ
E
000000000000000000000000000000000000000000000000000000000000
000000000000000000000000000000000000000000000000000000000000
000000000000000000000000000111111100000000000000000000011111
1
>DM_train1624
GSDEDFVTCYSVLKFINANDGSRLHSHDVKYGSGSGQQSVTAVKNSDDINSHWQIFPALNAK
CNRGDAIKCGDKIRLKHLTTGTFLHSHHFTAPLSKQHQEVSAFGSEAESDTGDDWTVICNGD
EWLESEQFKLRHAVTGSYLSLSGQQFGRPIHGQREVVGTDSITGGSAWKVAEGIYIKHQQKD
L
000000000000000000000000000000000000000000000000000000000000
000000000000000000000000000000000000000000000000000000000000
000000000000000000000000000000000000000000000000000011111111
1
>DM_train1625
VGYGDITQVETSGASSKTSRQDKLEYDGVRASHTMAQTDAGRMEKYKSFINNVAKKHVVDPA
VIAAIISRESRAGNVIFNTTPPGWGDNYNGFGLMQVDKRYHEPRGAWNSEEHIDQATGILVN
FIQLIQKKFPSWSTEQQLKGAIAAYNTGDGRVESYESVDSRTTGKDYSNDVVARAQWYKKNG
F
000000000000000000000000000000000000000000000000000000000000
000000000000001111111111111000000000000000000000000000000000
000000000000000000000000000000000000000000000000000000000000
0
>DM_train1626
MSDSKEVPSLPFLRHLLEELDSHEDSLLLFLCHDAAPGCTTVTQALCSLSQQRKLTLAALVE
MLYVLQRMDLLKSRFGLSKEGAEQLLGTSFLTRYRKLMVCVGEELDSSELRALRLFACNLNP
SLSTALSESSRFVELVLALENVGLVSPSSVSVLADMLRTLRRLDLCQQLVEYEQQEQARYRY
C
100000000000000000000000000000000000000000000000000000000000
000000000000000000000000000000000000000000000000000000000000
000000000000000000000000000000000000000000000000000000000000
0
>DM_train1627
GMSQTTVVNGVNVDQLMATIEQIKAKPEIAQFKFRATNQWMGGTHNQATIKDFYGACAEDDT
RKPMVFDLDEPPVLLGENRGANPVEYLLVALSGCLTTSLVAHAAARGIALRGVKSRYEGDID
LRGFLGLSEEVPVGYREIRVFFSIDADLTDGQKEELIRMAQKYSPVYNTVAKPVPVAVLLDR
G
111110000000000000000000000000000000000000000000000000000000
000000000000000000000000000000000000000000000000000000000000
000000000000000000000000000000000000000000000000000000000000

0
>DM_train1628
INAGTDLDNPKDLEVSDPTETTLSLRWRRPVAKFDRYRLTYVSPSGKKNEMEIPVDSTSFIL
RGLDAGTEYTISLVAEKGRHKSKPTTIKGSTVVGSPKGISFSDITENSATVSWTPPRSRVDS
YRVSYVPITGGTPNVVTVDGSKTRTKLVKLVPGVDYNVNIISVKGFEESEPISGILKTALDS
11111110000000000000000000000000000000000000000000000000000000
00000000000000000000000000000000000000000000000000000000000000
00000000000000000000000000000000000000000000000000000000001111
>DM_train1629
MAPVPPGEDSKDVAAPHRQPLTSSERIDKQIRYILDGISALRKETCNKSNMCESSKEALAEN
NLNLPKMAEKDGCFQSGFNEETCLVKIITGLLEFEVYLEYLQNRFESSEEQARAVQMSTKVL
IQFLQKKAKNLDAITTPDPTTNASLLTKLQAQNQWLQDMTTHLILRSFKEFLQSSLRALRQM
11111111111111111111000000000000000000000000000000000011111111
00000000000000000000000000000000000000000000000000000000000000
00000000000000000000000000000000000000000000000000000000000000
>DM_train1631
EKEMIERDMREYRGFSRAVRAVFEEKERFPGLVDVVSNLIEVDEKYSLAVSVLLGGTAQNIV
VRNVDTAKAIVEFLKQNEAGRVTILPLDLIDGSFNRISGLENERGFVGYAVDLVKFPSDLEV
LGGFLFGNSVVVETLDDAIRMKKKYRLNTRIATLDGELISGRGAITGGREERSSNVFERRIK
11111111111111110000000000000000000000000000000000000000000000
00000000000000000000000000000000000000000000000000000000000000
00000000000000000000000000000000000000000000000011111111111111
>DM_train1632
ASVTTPKGPGKSPDSPQWRQHQQDVRNLNQYQTRGAFAYISDQQKVYARFFWQQTGQDRYRL
LLTNPLGSTELELNAQPGNVQLVDNKGQRYTADDAEEMIGKLTGMPIPLNSLRQWILGLPGD
ATDYKLDDQYRLSEITYSQNGKNWKVVYGGYDTKTQPAMPANMELTDGGQRIKLKMDNWIVK
11111111100000000000000000000000000000000000000000000000000000
00000000000000000000000000000000000000000000000000000000000000
00000000000000000000000000000000000000000000000000000000000000
>DM_train1633
KQGIVKSDPKLPDDTTFPLPPPRPKNVIFEDEEKSKMLARLLKSSHPEDLRAANKLIKEMVQ
EDQKRMEKISKRVNAIEEVNNNVKLLTEMVMSHSQGGAAAGSSEDLMKELYQRCERMRPTLF
RLASDTEDNDEALAEILQANDNLTQVINLYKQLVRGEEVNGDATAGSIPGSTSALLDLSGLD
11111111111111111111111111111100000000000000000000000000000000
00000000000000000000000000000000000000000000000000000000000000
00000000000000000000000000000000000000111111111111111111111111
>DM_train1634
MRKDETSNTSPDISVAQPASALRYHLRPPRRNDGAAIHQLVSECPPLDLNSLYAYLLLCEHH
AHTCVVAESPGGRIDGFVSAYLLPTRPDVLFVWQVAVHSRARGHRLGRAMLGHILERQECRH
VRHLETTVGPDNQASRRTFAGLAGERGAHVSEQPFFDRQAFGGADHDDEMLLRIGPFTHPPH
11111111111111111111110000000000000000000000000000000000000000
00000000000000000000000000000000000000000000000000000000000000
00000000111100000000000000000000000111111111111000000000000111
>DM_train1635

KMDMEDADMTLWTEAEFEEKCTYIVNDHPWDSGADGGTSVQAEASLPRNLLFKYATNSEEVI
GVMSKEYIPKGTRFGPLIGEIYTNDTVPKNANRKYFWRIYSRGELHHFIDGFNEEKSNWMRY
VNPAHSPREQNLAACQNGMNIYFYTIKPIPANQELLVWYCRDFAERLHYPYPGELTMMNLTQ
111111000000000000000000000011111111110000000000000000000100000
000000000000000000000000000011100000000000000000000000000000000
000000000000000000000000000000000000000000000000000000000000000
>DM_train1636
GSHMQQWVDCEFTGRDFRDEDLSRLHTERAMFSECDFSGVNLAESQHRGSAFRNCTFERTTL
WHSTFAQCSMLGSVFVACRLRPLTLDDVDFTLAVLGGNDLRGLNLTGCRLRETSLVDTDLRK
CVLRGADLSGARTTGARLDDADLRGATVDPVLWRTASLVGARVDVDQAVAFAAAHGLCLAGG
111000000000000000000000000000000000000000000000000000000000000
000000000000000000000000000000000000000000000000000000000000000
000000000000000000000000000000000000000000000000000000000000011
>DM_train1637
SLWQPQTGDIIFQISRSSQSKAIQLATHSDYSHTGMLVMRNKKPYVFEAVGPVKYTPLKQWI
AHGEKGKYVVRRVEGGLSVEQQQKLAQTAKRYLGKPYDFSFSWSDDRQYCSEVVWKVYQNAL
GMRVGEQQKLKEFDLSNPLVQAKLKERYGKNIPLEETVVSPQAVFDAPQLTTVAKEWPLFSW
110000000000000000000000000000000000000000000000000000000000000
000000000000000000000000000000000000000000000000000000000000000
000000000000000000000000000000000000000000000000000000000001111
>DM_train1638
GVSIRSMKNFYDWIKEFVRDQGEFIAQQSGWLELERSSYAKLIAQTISHVLNGGSLLVSADS
SRHWFLNYILSNLNPKDLKERPLLSVIDFNASSFYPKNDANLSLATIEMTYQNPMFWHVGKI
ENEGLKTILLSKIPSFLWLFEELKEDCLLLKEHDSLLDYKLLQLFKLFENALFSVLYNKVTL
111111100000000000000000000000000000000000000000000000000000000
000000000000000000000000000000000000111000000000000000000000000
000000000000000000000000000000000000000000000000000000000000000
>DM_train1639
KKVVLGKKGDTVELTCTASQKKSIQFHWKNSNQIKILGNQGSFLTKGPSKLNDRADSRRSLW
DQGNFPLIIKNLKIEDSDTYICEVEDQKEEVQLLVFGLTANSDTHLLQGQSLTLTLESPPGS
SPSVQCRSPRGKNIQGGKTLSVSQLELQDSGTWTCTVLQNQKKVEFKIDIVVLAFQKASNT
000000000000000000000000000000000000000000000000000000000000000
000000000000000000000000000000000000000000000000000000000000000
00000000000000000000000000000000000000000000000000000000001111
>DM_train1640
MKFCSQCGGEVILRIPEGDTLPRYICPKCHTIHYQNPKVIVGCIPEWENKVLLCKRAIAPYR
GKWTLPAGFMENNETLVQGAARETLEEANARVEIRELYAVYSLPHISQVYMLFRAKLLDLDF
FPGIESLEVRLFGEQEIPWNDIAFRVIHDPLKRYMEERHHGQPAFHLGIINKPQAGSNSNE
000000000000000000000000000000000000000000000000000000000000000
000000000000000000000000000000000000000000000000000000000000000
00000000000000000000000000000000000000000000000000000111111111
>DM_train1641
RKQPIGPEDVLGLQRITGDYLCSPEENIYKIDFVRFKIRDMDSGTVLFEIKKPPVSERLPIN
RRDLDPNAGRFVVRYQFTPAFLRLRQVGATVEFTVGDKPVNNFRMIERHYFRNQLLKSFDHF

GFCIPSSKNTCEHIYDFPPLSEELISEMIRHPYETQSDSFYFVDDRLVMHNKADYSYSGTP
11100000000000000000000000000000000000000000000000000111111111
11111111000000000000000000000000000000000000000000000000000000
0000000000000000000000000000000000000000000000000000000000001
>DM_train1642
MQKVHVQYIDGETDQMLRQDDLDGYTDETIPYSTAEGIKKFEGDGYELFKDNFPAGEKFDND
DTNDQFYTVIFKHHRENVDPNHSSADGTKGTKTLTETVHYKYANGTKAAEDQTAQVTFTRNG
VLDDVTGIVAWGKWNEASQSYKALTSPTIAGYAPSEAVVKRSSNSDAEQGPTLTVIYTADA
00000000000000000000000000000000000000000000000000000000000000
00000000000000000000000000000000000000000000000000000000000000
0000000000000000000000000000000000000000000000000000000000001
>DM_train1643
MKGKTAAGGGAICAIAVMITIVMGNGNVRTNQAGLELIGNAEGCRRDPYMCPAGVWTDGIGN
THGVTPGVRKTDQQIAADWEKNILIAERCINQHFRGKDMPDNAFSAMTSAAFNMGCNSLRTY
YSKARGMRVETSIHKWAQKGEWVNMCNHLPDFVNSNGVPLRGLKIRREKERQLCLTGLVNE
11111110000000000000000000000000000000000000000000000000000000
00000000000000000000000000000000000000000000000000000000000000
0000000000000000000000000000000000000000000000000000000000000
>DM_train1645
ESSVNPVVLDFEDGTVMSFGEAWGDSLKCIKKVSVSQDLQRPGNKYALRLDVEFNPNNGWDQ
GDLGTWIGGVVEGQFDFTGYKSVEFEMFIPYDEFSKSQGGFAYKVVINDGWKELGSEFNITA
NAGKKVKINGKDYTVIHKAFAIPEDFRTKKRAQLVFQFAGQNSNYKGPIYLDNVRIRPEDA
11000000000000000000000000000000000000000000000000000000000000
00000000000000000000000000000000000000000000000000000000000000
0000000000000000000000000000000000000000000000000000000000000
>DM_train1646
AGVLWDVPSPPPVGKAELEDGAYRIKQKGILGYSQIGAGVYKEGTFHTMWHVTRGAVLMHKG
KRIEPSWADVKKDLISYGGGWKLEGEWKEGEEVQVLALEPGKNPRAVQTKPGLFKTNTGTIG
AVSLDFSPGTSGSPIVDKKGKVVGLYGNGVVTRSGAYVSAIANTEKSIEDNPEIEDDIFRK
11111111111111111000000000000000000000000000000000000000000000
00000000000000000000000000000000000000000000000000000000000000
0000000000000000000000000000000000000000000011111111111111111
>DM_train1648
AVPIAQKSEPHSLSSEALMRRAVSLVTDSTSTFLSQTTYALIEAITEYTKAVYTLTSLYRQY
TSLLGKMNSEEEDEVWQVIIGARAEMTSKHQEYLKLETTWMTAVGLSEMAAEAAYQTGADQA
SITARNHIQLVKLQVEEVHQLSRKAETKLAEAQIEELKQKTQEEGEERAESEQEAYLRED
11111111111000000000000000000000000000000000000000000000000000
00000000000000000000000000000000000000000000000000000000000000
000000000000000000000000000000000000000000000000000000000000
>DM_train1649
MPKTLIILAHPNISQSTVHKHWSDAVRQHTDRFTVHELYAVYPQGKIDVAAEQKLIETHDSL
VWQFPIYWFNCPPLLKQWLDEVLTYGWAYGSKGKALKGRKIALAVSLGAPAADYRADGAVGC
SVAEVLRPFELTAKYCNADYRPPFTFHTIDSNAGYSEAARQEVERSARDYLAWLDALQQT
10000000000000000000000000000000000000000000000000000000000000

000000000000000000000000000000000000000000000000000000000000
0000000000000000000000000001111111000000000000000000000011
>DM_train1650
VNSDSHPLFVRSLAKNMTWQLADTSTQKVLASGASATSGDKQSLLMQSVNLSYQEDGRGFNW
RAQAALSLSYLEPTPLDSKFSTGYLELKMRIDKAPEQGANLQVMCSESNCLRDIDFSSFSQL
MADKSWHTLAIPLHCDDSDQAEQPITDALRITSQNLSLAIADVALTIKPSDDSISLTCAK
111100000000000000000000000000000000000000000000000000000000
000000000000000000000000000000000000000000000000000000000000
0000000000000001111111000000000000000000000000000000000000
>DM_train1651
MKYQLRFGGEGGQGVITAGEILAEAAIKEGRQAFKASTYTSQVRGGPTKVDIIIDDKEILFP
YAVEGEVDFMLSTADKGYKGFRGGVKEGGIIVVEPNLVHPESEDYKKWQIFEIPIITIAKDE
VGNVATQSVVALAIAAYMSKCIDLDVLKETMLHMVPAKTRDANAKAFDLGVKYATQAKPH
100000000011100000000000000000000000000000000000000000000000
000000000000000000000000000000000000000000000000000000000000
0000000000000000000000000000000000000000000000000000000000
>DM_train1652
MTKAELRRRARAAWRRLDLKALSRAVGAALLPWLRERGFRHILLYHPLPHELNLLPLMEAYP
ARYYLPKVAGKGLTVHPFGPLAPGPFGLLEPTTPPEDPRVLDLVVVPGLAFDREGYRLGHGQ
GFYDRFLKEVRAATVGVVPQALLFPALPRDPWDVPVDHLATEAGVEAVKRPAPGPGGLLD
000000000000000000000000000000000000000000000000000000000000
000000000000000000000011111110000000000000000000000000000000
0000000000000000000000000000000000000000000000000111111111
>DM_train1653
GPGSMEQPKGINILITGTPGTGKTSMAEMIAAELDGFQHLEVGKLVKENHFYTEYDTELDTH
IIEEKDEDRLLDFMEPIMVSRGNHVVDYHSSELFPERWFHMVVVLHTSTEVLFERLTKRQYS
EAKRAENMEAEIQCICEEEARDAYEDDIVLVRENDTLEQMAATVEEIRERVEVLKVERGL
111110000000000000000000000000000000000000000000000111110
000000000000000000000000000000000000000000000000000000000000
0000000000000000000000000000000000000000000000000000011111
>DM_train1654
GSMLISHSDMNQQLKSAGIGFNATELHGFLSGLLCGGLKDQSWLPLLYQFSNDNHAYPTGLV
QPVTELYEQISQTLSDVEGFTFELGLTEDENVFTQADSLSDWANQFLLGIGLAQPELAKEKG
EIGEAVDDLQDICQLGYDEDDNEEELAEALEEIIEYVRTIAMLFYSHFNEGEIESKPVLH
111000000000000000000000000000000000000000000000000000000000
000000000000000000000000000000000000000000000000000000000000
0000000000000000000000000000000000000000000000011111111111
>DM_train1655
NSRVSDLDLELLFQDRAARLGKSISRLIVVASLIDKPTNLGGLCRTCEVFGASVLVVGSLQC
ISDKQFQHLSVSAEQWLPLVEVKPPQLIDYLQQKKTEGYTIIGVEQTAKSLDLTQYCFPEKS
LLLLGNEREGIPANLIQQLDVCVEIPQQGIIRSLNVHVSGALLIWEYTRQQLLSHGDTKP
111111111111111111111111100000000000000000000000000000000000
000000000000000000000000000000000000000000000000000000000000
0000000000000000000000000000000000000000000000000000011111

>DM_train1656
GMLFSIQTCPCQINPALNAVSTPLLYQDCCQPYHDGLYNQEVEDSDAIRADTAEHLMRTRYS
AFVLVKPEYIVKTTLPAQQDLLDIKAIENWAKETDWAGLEVVAHTPKLSKRHAQVEFKAYFK
TPDGLQAHHELSTFVKIKNKANSDASWYFLDPTVSMSVTQKQPCICGSGEKFKRCCGMYI
10000000000000000000000000000000000000001111110000000000000000
00000000000000000000000000000000000000000000000000000000000000
000000000000000000000000000000000000000000000000000000000000
>DM_train1657
SNAMKIAVINGGTRSGGNTDVLAEKAVQGFDAEHIYLQKYPIQPIEDLRHAQGGFRPVQDDY
DSIIERILQCHILIFATPIYWFGMSGTLKLFIDRWSQTLRDPRFPDFKQQMSVKQAYVIAVG
GDNPKIKGLPLIQQFEHIFHFMGMSFKGYVLGEGNRPGDILRDHQALSAASRLLKRSDAI
11110000000000000000000000000000000001111111111111111111111100
00000000000000000000000000000000000000000000000000000000000000
000000000000000000000000000000000000000000000000000000000001
>DM_train1658
GPGFDYAVAMAKRNIVTATTSKGEFTMLGVHDNVAILPTHASPGESIVIDGKEVEILDAKAL
EDQAGTNLEITIITLKRNEKFRDIRPHIPTQITETNDGVLIVNTSKYPNMYVPVGAVTEQGY
LNLGGRQTARTLMYNFPTRAGQCGGVITCTGKVIGMHVGGNGSHGFAAALKRSYFTQSQ
00000000000000000000000000000000000000000000000000000000000000
00000000000000000000000000000000000000000000000000000000000000
00000000000000000000000000000000000000000000000000000000111
>DM_train1660
VGSADAPAPYRVSSDCMVCHGMTGRDTLYPIVPRLAGQHKSYMEAQLKAYKDHSRADQNGEI
YMWPVAQALDSAKITALADYFNAQKPPMQSSGIKHAGAKEGKAIFNQGVTNEQIPACMECHG
SDGQGAGPFPRLAGQRYGYIIQQLTYFHNGTRVNTLMNQIAKNITVAQMKDVAAYLSSL
11111111111100000000000000000000000000000000000000000000000000
00000000000000000000000000000000000000000000000000000000000000
00000000000000000000000000000000000000000000000000000000000
>DM_train1661
GMDYNQTVLSHLQKFWKHHDIKGFTWTLGRIVEELPDFQVFQVIPNHEDEPWVYVSSGIGQF
LGQEFFIISPFETPEHIETLAMLASASMHYPDQFQLGKTVNIGRPWVEQSSFRHFLISLPYP
YGQELEYMDNVRFFWLLPITQTERLFLNTHSVEELETKFDEAGIDYLDINRASTVWQAG
00000000000000000000000000000000000000000000000000000000000000
00000000000000000000000000000000000000000000000000000000000000
00000000000000000000000000000000000000000000000000000000001
>DM_train1662
SNAMTRYALLVRGINVGGKNKVVMAELRQELTNLGLEKVESYINSGNIFFTSIDSKAQLVEK
LETFFAVHYPFIQSFSLLSLEDFEAELENLPAWWSRDLARKDFLFYTEGLDVDQVIATVESL
ELKDEVLYFGKLGIFWGKFSEESYSKTAYHKYLLKVPFYRHITIRNAKTFDKIGQMLKK
11000000000000000100000000000000000000000000000000000000000000
00000000000000000000000000000000000000000000000000000000000000
00000000000000000000000000000000000000000000000000000000000
>DM_train1663
MQRCDWVSQDPLYIAYHDNEWGVPETDSRKLFEMICLEGQQAGLSWITVLKKRENYRACFHQ

FDPIRIAAMQEEDVERLLQNTGIIRHRGKIQAIISNARAWLAMEQNGESFADFVWSFVDGQP
QITQAASLDKIPTSTPASDALAKALKKRGFKFVGTTICYSFMQACGLVNDHITGCFCHP
10000000000000000000000000000000000000000000000000000000000000
00000000000000000000000000000000000000000000000000000000000000
00000000000000000000000000000000000000000000000000000000000
>DM_train1664
TPMRSPIIWINGPFGVGKTHTAHTLHERLPGSFVFEPEEMGQALRKLTPGFSGDPQEHPMWI
PLMLDALQYASREAAGPLIVPVSISDTARHRRLMSGLKDRGLSVHHFTLIAPLNVVLERLRR
DGQPQVNVGTVEDRLNELRGEQFQTHIDTAGLGTQQVAEQIAAQVGLTLAPPPQGALHW
11110000000001111100000000000000000000000000000000000000000000
00000000000000000000000000000000000000000000000000000000000000
01111000000000000000000000000000000000000000000000000001111111
>DM_train1666
MSLTDVMFKSQIANQLKNLRKSRGLSLDATAQLTGVSKAMLGQIERGESSPTIATLWKIASG
LEASFSAFFANDPQLLSSERSFPDDLNMKIHTLFPYAADTGLEIFEITLLDHHQQMSSPHAL
GVIEYIHVLEGIMKVFFDEQWHELQQGEHIRFFSDQPHGYAAVTEKAVFQNIVAYPRR
11100000000000000000000000000000000000000000000000000000000000
00000000000000000000000000000000000000000000000000000000000000
0000000000000000000000000000000000000000000000000000000001
>DM_train1667
MALTATVRRAELQISDMDRGYYANHSLTLAQHPSETDERLMVRLLAFALFADDRLEFGRGLS
NDDEPDLWRRDYTGDPDLWIDLGQPDESRVRKACNRSREAVVIGYGGQATETWWKKHANAMG
RYRNLRVIELDSQATEALGALIQRGMRFDVIIQDGEVQMLADHGSVTLTPMVRQAPAE
11100000000000000000000000000000000000000000000000000000000000
00000000000000000000000000000000000000000000000000000000000000
0000000000000000000000000000000000000000000000000000000011
>DM_train1668
QKPQRPRRPASPISTIQPKANFDAQQFAGTWLLVAVGSAARFLQEQGHRAEATTLHVAPQGT
AMAVSTFRKLDGICWQVRQLWGDTGVLGRFLLQARGARGAVHVVVAETDYQSFAVLYLERAG
QLSVKLYARSLPVSDSVLSGFEQRVQEAHLTEDQIFYFPKYGFCEAADQFHVLDEVRR
11111111100000000000000000000000000000000111111000000000000000
00000000000000000000000000000000000000000000000000000000000000
0000000000000000000000000000000000000000000000000000000011
>DM_train1669
AGLKLMGAPVKLTVSQGQPVKLNCSVEGMEEPDIQWVKDGAVVQNLDQLYIPVSEQHWIGFL
SLKSVERSDAGRYWCQVEDGGETEISQPVWLTVEGVPFFTVEPKDLAVPPNAPFQLSCEAVG
PPEPVTIVWWRGTTKIGGPAPSPSVLNVTGVTQSTMFSCEAHNLKGLASSRTATVHLQ
11111100000000000000000000000000000000000000000000000000000000
00000000000000000000000000000000000000000000000000000000000000
0000000000000000000000000000000000000000000000000000000000
>DM_train1670
ELPSLCMLNNSFYYMKGGANIFLIRVSDVSVLMKEYDVSVYEPEDLGNCLNKSDSSWAIHWF
SIALGHDWLMDPPMLCRNKTKKEGSNIQFNISKADESRVYGKKIRNGMRHLFRGFYDPCEEG
KVCYVTINQCGDPSSFEYCGTNYLSKCQFDHVNTLHFLVRSKTHLNFGTGTKHHHHHH

000000000000000000000000000000000000000000000000000000000000
000000000000000000000000000000000000000000000000000000000000
0000000000000000000000000000011111111111111111111111111111
>DM_train1671
GPAAGQAYDAGNLDVASSPVKPTLSITKKTLTAAEAPNAKVTMELSVEGAADKYAATGLHIQ
FDPKLKLIPDEDGALATAGRAARLLELKKAEADTDNSFFTATGSSTNNGKDGVLWSFVLQVP
ADAQPGDKYDVQVAYQSRTTNEDLFTNVKKDEEGLLMQAWTFTQGIEQGYIQVESTTS
000000000000000000000000000000000000000000000000000000000000
000000000000000000000000000000000000000000000000000000000000
0000000000000000000000000000000000000000000000000000000111
>DM_train1672
GSCRRAFDLYFVLDKSGSVANNWIEIYNFVQQLAERFVSPEMRLSFIVFSSQATIILPLTGD
RGKISKGLEDLKRVSPVGETYIHEGLKLANEQIQKAGGLKTSSIIIALTDGKLDGLVPSYAE
KEAKISRSLGASVYCVGVLDFEQAQLERIADSKEQVFPVKGGFQALKGIINSILAQSC
100000000000000000000000000000000000000000000000000000000000
000000000000000000000000000000000000000000000000000000000000
0000000000000000000000000000000000000000000000000000000000
>DM_train1673
MHTQVHTARLVHTADLDSETRQDIRQMVTGAFAGDFTETDWEHTLGGMHALIWHHGAIIAHA
AVIQRRLIYRGNALRCGYVEGVAVRADWRGQRLVSALLDAVEQVMRGAYQLGALSSSARARR
LYASRGWLPWHGPTSVLAPTGPVRTPDDDGTVFVLPIDISLDTSAELMCDWRAGDVW
111110000000000000000000000000000000000000000000000000000000
000000000000000000000000000000000000000000000000000000000000
000000000000000000000000000000000000000000000000000000000
>DM_train1674
GGSNTNWKTLYEVKSENLGQGDKPDYFSSVATVVYLRKENCMYQACPTQDCNKKVIDQQNGL
YRCEKCDTEFPNFKYRMILSVNIADFQENQWVTCFQESAEAILGQNAAYLGELKDKNEQAFE
EVFQNANFRSFIFRVRVKVETYNDESRIKATVMDVKPVDYREYGRRLVMSIRRSALM
111000000000000000000000000000000000000000000000000000000000
000000000000000000000000000000000000000000000000000000000000
000000000000000000001111110000000000000000000000000000000
>DM_train1675
MSKIICLTAGHSNTDPGAVNGSDREADLAQDMRNIVASILRNDYGLTVKTDGTGKGNMPLRD
AVKLIRGSDVAIEFHTNAAANKTATGIEALSTPKNKRWCQVLGKAVAKKTGWKLRGEDGFKP
DNAGQHSRLAYAQAGGIVFEPFFISNDTDLALFKTTKWGICRAIADAIAMELGAAKV
100000000000000000000000000000000000000000000000000000000000
000000000000000000000000000000000000000000000000000000000000
000000000000000000000000000000000000000000000000000000000
>DM_train1676
MGSDKIHHHHHHMTENSRKTATELFEFLDGLGISHTTKQHEPVFTVAESQSLRDLIPGGHTK
NLFVKDKKDQYFVLTVEENAVVDLKSVHKTIGAASRVSFGRPEKMLEYLGVVPGSVTVFGAI
NDTARQVTFVLDSDLLENELVNGHPLSNDQTTTIASKDLIRFLEATGHAPLVLKVSE
111111111111111100000000000000000000000000000000000000000000
000000000000000000000000000000000000000000000000000000000000

0000000000000000000000000000000000000000000000000000000
>DM_train1677
NQKCSGNPRRYNGKSCASTTNYHDSHKGACGCGPASGDAQFGWNAGSFVAAASQMYFDSGNK
GWCGQHCGQCIKLTTTGGYVPGQGGPVREGLSKTFMITNLCPNIYPNQDWCNQGSQYGGHNK
YGYELHLDLENGRSQVTGMGWNNPETTWEVVNCDSEHNHDHRTPSNSMYGQCQCAHQ
000000000000000000000000000000000000000000000000000000000000
000000000000000000000000000000000000000000000000000000000000
0000000000000000000000000000000000000000000000000000001
>DM_train1678
MAGSREVVAMDCEMVGLGPHRESGLARCSLVNVHGAVLYDKFIRPEGEITDYRTRVSGVTPQ
HMVGATPFAVARLEILQLLKGKLVVGHDLKHDFQALKEDMSGYTIYDTSTDRLLWREAKLDH
CRRVSLRVLSERLLHKSIQNSLLGHSSVEDARATMELYQISQRIRARRGLPRLAVSD
111110000000000000000000000000000000000000000000000000000000
000000000000000000000000000000000000000000000000000000000011
1110000000000000000000000000000000000000000000000000111
>DM_train1679
RIAISNSNRTRSVPSLTTIWSISPTPNCSIYETQDANLFLCLTKNGAHVLGTITIKGLKGAL
REMHDNALSLKLPFDNQGNLLNCALESSTWRYQETNAVASNALTFMPNSTVYPRNKTAHPGN
MLIQISPNITFSVVYNEINSGYAFTFKWSAEPGKPFHPPTAVFCYITEQGSHHHHHH
111111111111110000000000000000000000000000000000000000000000
000000000000000000000000000000000000000000000000000000001111
1111111100000000000000000000000000000000000000001111111
>DM_train1680
MELEIRPLFLVPDTNGFIDHLASLARLLESRKYILVVPLIVINELDGLAKGQETDHRAGGYA
RVVQEKARKSIEFLEQRFESRDSCLRALTSRGNELESIAFRSEDITGQLGNNDDLILSCCLH
YCKDKAKDFMPASKEEPIRLLREVVLLTDDRNLRVKALTRNVPVRDIPAFLTWAQVG
000000000000000000000000000000000000000000000000011111100000
000000000000000000000000000000000000000000111111000000000000
0000000000011111000000000000000000000000000000000000001
>DM_train1681
HQYLLPEAKAQDSDKICVVINLDETLVHSSFKPVNNADFIIPVEIDGVVHQVYVLKRPHVDE
FLQRMGELFECVLFTASLAKYADPVADLLDKWGAFRARLFRESCVFHRGNYVKDLSRLGRDL
RRVLILDNSPASYVFHPDNAVPVASWFDNMSDTELHDLLPFFEQLSRVDDVYSVLRQ
100000000000000000000000000000000000000000000000000000000000
000000000000000000000000000000000000000000000000000000000000
0000000000000000000000000000000000000000000000000000000
>DM_train1682
SLKVLAITGYKPFELGIFKQDDKALYYIKKAIKNRLIAFLDEGLEWILISGQLGVELWAAEA
AYDLQEEYPDLKVAVITPFYEQEKNWKEPNKEQYEAVLAQADYEASLTHRPYESPLQFKQKN
QFFIDKSDGLLLLYDPEKEGSPKYMLGTAEKRREQDGYPIYFITMDDLRVTVEEDSY
000000000000000000000000000000000000000000000000000000000000
000000000000000000000000000000000000000000000000000000000000
0000000000000000000000000000000000000000000000000000111
>DM_train1683

MKKWRFLGIDDSFDDRKCCVVGCVTCGGYVEGFLYTEIDIDGLDATDKLISMVRRSKFREQI
KCIFLPGITLGGFNLVDIQRVYRETKIPVVVVMRRKPDMEEFDSAMRNLENYELRRKIVEVA
GEIHRIGDIYIQTAGLTPSEAEKLVKASLIKGNMPEPVRISHLVASAIIHGESRGKA
11000000000000000000000000000000000000000000000000000000000000
00000000000000000000000000000000000000000000000000000000000000
000000000000000000000000000000000000000000000000000111111
>DM_train1684
CAVSSFRVKENFDKARFSGTWYAMAKKDPEGLFLQDNIVAEFSVDETGQMSATAKGRVRLLN
NWDVCADMVGTFTDTEDPAKFKMKYWGVASFLQKGNDDHWIVDTDYDTYAVQYSCRLLNLDG
TCADSYSFVFSRDPNGLPPEAQKIVAQRQEELCLAAQYRLIVHNGYCDGRSERNLL
00000000000000000000000000000000000000000000000000000000000000
00000000000000000000000000000000000000000000000000000000000000
00000000000000000000000000000000000000000000000111111111
>DM_train1685
MRGSHHHHHHGSITSGNKSTNVTVHKSEAGTSSVFYYKTGDMLPEDTTHVRWFLNINNEKSY
VSKDITIKDQIQGGQQLDLSTLNINVTGTHSNYYSGQSAITDFEKAFPGSKITVDNTKNTID
VTIPQGYGSYNSFSINYKTKITNEQQKEFVNNSQAWYQEHGKEEVNGKSFNHTVHN
11111111111111111111111111111100000000000000000000000000000000
00000000000000000000000000000000000000000000000000000000000000
00000000000000000000000000000000000000000000000000000000
>DM_train1686
APLAYERPVLHLVALNTPVAGDIRADFQCFQQARAAGLLSTFRAFLSSHLQDLSTVVRKAER
FGLPIVNLKGQVLFNNWDSIFSGDGGQFNTHIPIYSFDGRDVMTDPSWPQKVVWHGSNPHGV
RLVDKYCEAWRTTDMAVTGFASPLSTGKILDQKAYSCANRLIVLCIENSFMTDTRK
11111100000000000000000000000000000000000000000000000000000000
00000000000000000000000000000000000000000000000000000000000000
00000000000000000000000000000000000000000000000000111111
>DM_train1687
MSVKMKKCSREDLQTLQQLSIETFNDTFKEQNSPENMKAYLESAFNTEQLEKELSNMSSQFF
FIYFDHEIAGYVKVNIDDAQSEEMGAESLEIERIYIKNSFQKHGLGKHLLNKAIEIALERNK
KNIWLGVWEKNENAIAFYKKMGFVQTGAHSFYMGDEEQTDLIMAKTLILEHHHHHH
10000000000000000000000000111111110000000000000000000000000000
00000000000000000000000000000000000000000000000000000000000000
00000000000000000000000000000000000000000000000000000111
>DM_train1688
GSHMENMGTRVIEPLIMGRVVGDVLDFFTPTTKMNVSYNKKQVSNGHELFPSSVSSKPRVEI
HGGDLRSFFTLVMIDPDVPGPSDPFLKEHLHWIVTNIPGTTDATFGKEVVSYELPRPSIGIH
RFVFVLFRQKQRRVIFPNIPSRDHFNTRKFAVEYDLGLPVAAVFFNAQRETAARKR
11111111100000000000000000000000000000000000000000000000000000
00000000000000000000000000000000000000000000000000000000000000
00000000000000000000000000000000000000000000000000111111
>DM_train1689
VMLTRQQKELIVKEMSEIFKKTSLILFADFLGFTVADLTELRSRLREKYGDGARFRVVKNTL
LNLALKNAEYEGYEEFLKGPTAVLYVTEGDPVEAVKIIYNFYKDKKADLSRLKGGFLEGKKF

TAEEVENIAKLPSKEELYAMLVGRVKAPITGLVFALSGILRNLVYVLNAIKEKKSE
000000000000000000000000000000000000000000000000000000000000
000000000000000000000000000000000000000000000000000000000000
000000000000000000000000000000000000000000000000000011
>DM_train1690
MGSSSSGLVPRGSGAVSGSVQASDRLMKELRDIYRSQSYKTGIYSVELINDSLYDWHVKLQK
VDPDSPLHSDLQILKEKEGIEYILLNFSFKDNFPFDPPFVRVVLPVLSGGYVLGGGALCMEL
LTKQGWSSAYSIESVIMQINATLVKGKARVQFGANKNQYNLARAQQSYNSIVQIHE
111111111111111100000000000000000000000000000000000000000000
000000000000000000000000000000000000000000000000000000000000
000000000000000000000000000000001111111111111111111111
>DM_train1691
GSSGSSGMANERMNLMNMAKLSIKGLIESALNLGRTLDSDYAPLQQFFVVMEHCLKHGLKAK
KTFLGQNKSFWGPLELVEKLVPEAAEITASVKDLPGLKTPVGRGRAWLRLALMQKKLSEYMK
ALINKKELLSEFYEVNALMMEEEGAIIAGLLVGLNVIDANFCMKGEDLDSQVGVID
111111100000000000000000000000000000000000000000000000000001
111111000000000000000000000000000000000000000000000000000000
000000000000000000000000000000000000000000000000001111
>DM_train1692
MEQRRLASTEWVDIVNEENEVIAQASREQMRAQCLRHRATYIVVHDGMGKILVQRRTETKDF
LPGMLDATAGGVVQADEQLLESARREAEEELGIAGVPFAEHGQFYFEDKNCRVWGALFSCVS
HGPFALQEDEVSEVCWLTPEEITARCDEFTPDSLKALALWMKRNAKNEAVETETAE
111111100000000000000000000000000000000000000000000000000000
000000000000000000000000000000000000000000000000000000000000
000000000000000000000000000000000000000000111111111111
>DM_train1693
GAPRSHEVPLLVTLEELYLGKRKKIKVTRKRFIEHKVRNEENIVEVEIKPGWKDGTKLTYSG
EGDQESPGTSPGDLVLIIQTKTHPRFTRDDCHLIMKVTIPLVRALTGFTCPVTTLDNRNLQI
PIKEIVNPKTRKIVPNEGMPIKNQPGQKGDLILEFDICFPKSLTPEQKKLIKEALD
111111000000000000000000000011111111110000000000000000000000
000000000000000000000000000000000000000000000000000000100000
000000000000001000000000000000000000000000000000000001
>DM_train1694
GMFADNTLTPKEAVRLCALGTIASQPMRYSELAGSVRHFTSRIMGPSLELMGISIELLRYEG
LVEAVDDGQGMEDDAMLAISAAGRRELHSLLTARLRPGSDLSKLVVALKMRFLGLMEAEERA
HQIDLLIEGVDSELARLADLRGSDGEGGSALAAWLDHDMALLESRLAWLEDFRARL
111111000000000000000000000000000000000000000000000000000000
000001111111100000000000000000000000000000000000000000000000
000000000000000000000011000000000000000000000000000000
>DM_train1695
EVYALEESRDVYLSDLDWLNATHGDDTKSKIVQKNHPFTPGNNNQSTKISLKMEDGSISEFE
KGLGTIAGSPSTITYDISGAGVTKFFSYLGIDRSANPINEQYAKVDKIEVVVDGKVIYSTIN
QFPNGLTYETPAIKVDLNIPENAKRLQLKSYAGEKTWGDEVVYADAKFTAKGDFVN
111111100000000000000000000000000000000000000000000000000000

00000000000000000000000000000000000000000000000000000000000000
00000000000000000000000000000000000000000000000000000011
>DM_train1697
AAQGKEVVLLDFAAAGGELGWLTHPYGKGWDLMQNIMNDMPIYMYSVCNVMSGDQDNWLRTN
WVYRGEAERIFIELKFTVRDCNSFPGGASSCKETFNLYYAESDLDYGTNFQKRLFTKIDTIA
PDEITVSSDFEARHVKLNVEERSVGPLTRKGFYLAFQDIGACVALLSVRVYYKKCP
11110000000000011100000000000000000000000000000000000000000000
00000000000000000000000000000000000000000000000000000000000000
00111111111110000000000000000000000000000000000000000011
>DM_train1698
RSRLTADEYLKIYQAAESSPCWLRLAMELAVVTGQRVGDLCEMKWSDIVDGYLYVEQSKTGV
KIAIPTALHIDALGISMKETLDKCKEILGGETIIASTRREPLSSGTVSRYFMRARKASGLSF
EGDPPTFHELRSLSARLYEKQISDKFAQHLLGHKSDTMASQFRDDRGREWDKIEI
00000000000000000000000000000000000000000000000000000000000000
00000000000000000000000000000000000000000000000000000000000000
0000000000000000000000000000000000111111111111110000000
>DM_train1699
MAAGQNGHEEWVGSAYLFVESSLDKVVLSDAYAHPQQKVAVYRALQAALAESGGSPDVLQML
KIHRSDPQLIVQLRFCGRQPCGRFLRAYREGALRAALQRSLAAALAQHSVPLQLELRAGAER
LDALLADEERCLSCILAQQPDRLRDEELAELEDALRNLKCGSGARGGDGEVASAP
11111100000000000000000000000000000000000000000000000000000000
00000000000000000000000000000000000000000000000000000000000000
0000000000000000000000000000000000000000011111111111111
>DM_train1701
SADVAGAVIDGASLSFDILKTVLEALGNVKRKIAVGVDNESGKTWTALNTYFRSGTSDIVLP
HKVPHGKALLYNGQKDRGPVATGAVGVLAYLMSDGNTLAVLFSVPYDYNWYSNWWNVRIYKG
KRRADQRMYEELYYNLSPFRGDNGWHTRNLGYGLKSRGFMNSSGHAILEIHVSKA
11100000000000000000000000000000000000000000000000000000000000
00000000000000000000000000000000000000000000000000000000000000
0000000000000000000000000000000000000000000000000000000
>DM_train1702
MASNEARYVLAEEVDFSSPEEVKNWWNSGTWQAEFGSPDIEWNGEVGNGALQLNVKLPGKSD
WEEVRVARKFERLSECEILEYDIYIPNVEGLKGRLRPYAVLNPGWVKIGLDMNNANVESAEI
ITFGGKEYRRFHVRIEFDRTAGVKELHIGVVGDHLRYDGPIFIDNVRLYKRTGGM
11111000000000000000000000000000000000000000000000000000000000
00000000000000000000000000000000000000000000000000000000000000
0000000000000000000000000000000000000000000000000000111
>DM_train1703
MSEHFVGKYEVELKFRVMDLTTLHEQLVAQKATAFTLNNHEKDIYLDANGQDLAKQQISMVL
REMNPSGIRLWIVKGPGAERCEASNIEDVSKVQSMLATLGYHPAFTIEKQRSIYFVGKFHIT
VDHLTGLGDFAEIAIMTDDATELDKLKAECRDFANTFGLQVDQQEPRSYRQLLGF
11100000000000000000000000000000000000000000000000000000000000
00000000000000000000000000000000000000000000000000000000000000
0000000000000000000000000000000000000000000000000000000

>DM_train1704
MAEEYSWDSYLNDRLLATNQVSGAGLASEEDGVVYACVAQGEESDPNFDKWSLFYKEDYDIE
VEDENGTKTTKTINEGQTILVVFNEGYAPDGVWLGGTKYQFINIERDLEFEGYNFDVATCAK
LKGGLHLVKVPGGNILVVLYDEEKEQDRGNSKIAALTFAKELAESSQLQHHHHHH
11110000000000000000000000000000000000000000000000000000000000
00000000000000000000000000000000000000000000000000000000000000
0000000000000000000000000000000000000000000001111111111
>DM_train1705
ASWSHPQFEKGAMALTEAWLIEKANRKLNAGGMYKITSDKTRNVIKKMAKEGIYLCVAQGYR
STAEQNALYAQGRTKPGAIVTNAKGGQSNHNYGVAVDLCLYTNDGKDVIWESTTSRWKKVVA
AMKAEGFKWGGDWKSFKDYPHFELCDAVSGEKIPAATQNTNTNSNRYEGKVIDSA
11111111111000000000000000000000000000000000000000000000000000
00000000000000000000000000000000000000000000000000000000000000
0000000000000000000000000000000000001111111111111111111
>DM_train1706
HHHHHHLAYSLDATASFLNFVSSKKTHVLETHRFDVLSGGISTAGEAQLVIDLNSVNTGIDV
RNGRMRDYLFETATYSVATVTVPVDLAAVAGLAVGEDMLVDVSATLDLHGVPGVIDTQLNVQ
RLSATRIMVQNQSPLLIKAADYSLEAGIETLRNLASLNVISTTVPVDFVLFYEAP
11111000000000000000000000000000000000000000000000000000000000
00000000000000000000000000000000000000000000000000000000000000
0000000000000000000000000000000000000000000000000000000
>DM_train1707
MVEIIPVSTTLELRAADESHVPALHQLVLKNKAWLQQSLDWPQYVTSQEETRKHVQGNILLH
QRGYAKMYLIFCQNEMAGVLSFNAIEPINKAAYIGYWLDESFQGQGIMSQSLQALMTHYARR
GDIRRFVIKCRVDNQASNAVARRNHFTLEGCMKQAEYLNGDYHDVNMYARIIDAD
11000000000000000000000000000001111111111111111111000000000000
00000000000000000000000000000000000000000000000000000000000000
0000000000000000000000000000000000000000000000000000000
>DM_train1708
MISYEFQTHLPKGKDSSLNASSENKELYVQATHFNNTILLQIRLNGEMDSTYEVSSKGLNPI
LDINVPLAGNLGNTGGDYDDEEEEFVRDHLSDYQVVTKLGDSADPKVPVVCVQIAELYRRVI
LPEVSGTMAQDNMQFSLLISMSSKIWRATKEQSADDNDFGKLVFVLKCIKDMYAK
00000000000011111111110000000000000000000000000000000000000111
11111111111111111000000000000000000000000000000000000000000000
0001111111111000000000000011111100000000000000000000001
>DM_train1709
GGGVTFCGGEPLLHPEFLIDILKRCGQQGIHRAVDTTLLARKETVDEVMRNCELLLIDLKSM
DSTVHQTFCDVPNELILKNIRRVAEADFPYYIRIPLIEGVNADEKNIKLSAEFLASLPRHPE
IINLLPYHDIGKGKHAKLGSIYNPKGYKMQTPSEEVQQQCIQILTDYGLKATIGG
00000000000000000000000000000000000000000000000000000000000000
00000000000000000000000000000000000000000000000000000000000000
0000000001111111111111111110000000000000000000000000000
>DM_train1710
MEDSCVLRGVMINKDVTHPRMRRYIKNPRIVLLDSSLEYKKGESQTDIEITREEDFTRILQM

EEEYIHQLCEDIIQLKPDVVITEKGISDLAQHYLMRANVTAIRRVRKTDNNRIARACGARIV
SRPEELREDDVGTGAGLLEIKKIGDEYFTFITDCKDPKACTILLRGASHHHHHH
111111100000000000000000000000000000011111111111111111111111
110000000000000000000000000000000000000000000000000000000000
0000000000000000000000000000000000000000001111111111
>DM_train1711
MPAYHSSLMDPDTKLIGNMALLPIRSQFKGPAPRETKDTDIVDEAIYYFKANVFFKNYEIKN
EADRTLIYITLYISECLKKLQKCNSKSQGEKEMYTLGITNFPIPGEPGFPLNAIYAKPANKQ
EDEVMRAYLQQLRQETGLRLCEKVFDPQNDKPSKWWTCFVKRQFMNKSLSGPGQ
000000000000000000000000000000000000000000000000000000000000
000000000000000000000000000000000000000000000000000000000000
0000000000000000000000000000000000000000000000000111
>DM_train1713
GMARKVILFIAMSIDNYIADDQGAVDWLEKNVHGTESDDSYEKMYSKIDTVIMGRTTYEQVT
QKLSPEKYVYADRQTYIVTSHLGEDTDKIKYWKQSPVELVKRIQKEKGKDVWIVGGAKIIDP
LVQANLIDTYILTTVPIFLGSGIRLFDRLEEQVPVRLIDVYQKNELVYSIYQRG
110000000000000000000000000000000000000000000000000000000000
001111000000000000000000000000000000000000000000000000000000
0000000000000000000000000000000000000000000000000000
>DM_train1714
ADFTETFESSTHGEAPAEWTTIDADGDGQGWLCLSSGQLDWLTAHGGSNVVSSFSWNGMALN
PDNYLISKDVTGATKVKYYYAVNDGFPGDHYAVMISKTGTNAGDFTVVFEETPNGINKGGAR
FGLSTEANGAKPQSVWIERTVDLPAGTKYVAFRHYNCSDLNYILLDDIQFTMGG
000000000000000000000000000000000000000000000000000000000000
000000000000000000000000000000000000000000000000000000000000
0000001110000000000000000000000000000000000000000011
>DM_train1715
SKPRILLMGLRRSGKSSIQKVVFHKMSPNETLFLESTNKIYKDDISNSSFVNFQIWDFPGQM
DFFDPTFDYEMIFRGTGALIYVIDAQDDYMEALTRLHITVSKAYKVNPDMNFEVFIHKVDGL
SDDHKIETQRDIHQRANDDLADAGLEKLHLSFYLTSIYDHSIFEAFSKVVQKLI
100000000000000000000000000000000000000000000000000000000000
000000000000000000000000000000000000000000000000000000000000
0000000000000000000000000000000000000000000000000000
>DM_train1716
MASAVGEKMLDDFEGVLNWGSYSGEGAKVSTKIVSGKTGNGMEVSYTGTTDGYWGTVYSLPD
GDWSKWLKISFDIKSVDGSANEIRFMIAEKSINGVGDGEHWVYSITPDSSWKTIEIPFSSFR
RRLDYQPPGQDMSGTLDLDNIDSIHFMYANNKSGKFVVDNIKLIGALEHHHHHH
110000000000000000000000000000000000000000000000000000000000
000000000000000011100000000000000000000000000000000000000000
0000000000000000000000000000000000000000000000000111
>DM_train1718
MKWNLDPSHTSIDFKVRHMGIASVRGSLKVLSGSVETDEAGRPIQVEAVIDAASIATGEPQR
DGHLRSADFLHAEQYPEIRFVSTQIEPLGGNRYRIQGNLTIRDITKPVTLEAEVSAPIKDPW
GMQRVAASASGQINRKDWNLTWNQVLELGALLVGEEVKFNLEVEAVAPAPVAAQ

00000000000000000000000000000000000000000000000000000000000000
00000000000000000000000000000000000000000000000000000000000000
000000000000000000000000000000000000000000000000000011
>DM_train1719
EAEASAVTVDTICKNGQLVQMSNHFKCMCNEGLVHLSENTCEEKNECKKETLGKACGEFGQC
IENPDPAQVNMYKCGCIEGYTLKEDTCVLDVCQYKNCGESGECIVEYLSEIQSAGCSCAIGK
VPNPEDEKKCTKTGETACQLKCNTDNEVCKNVEGVYKCQCMEGFTFDKEKNVCL
11111000000000000000000000000000000000000000000000000000000000
00000000000000000000000000000000000000000000000000000000000000
000000000000000000000000000000000000000000000000000000
>DM_train1720
PLGSQYFLKVLIPSYAAGSIIGKGGQTIVQLQKETGATIKLSKSKDFYPGTTERVCLIQGTI
EALNAVHGFIAEKIREMPQNVAKTEPVSILQPQTTVNPDRANQVKIIVPNSTAGLIIGKGGA
TVKAIMEQSGAWVQLSQKPDGINLQNRVVTVSGEPEQNRKAVELIIQKIQEDPQ
11000000000000000000000000000000000000000000000000000000000000
00000000000000000000111111111111111110000000000000000000000000
000000000000000000011110000000000000000000000000000000
>DM_train1721
MTSVNLSYPIGEYKPRESISKEQKDKWIQVLEEVPAKLKQAVEVMTDSQLDTPYRDGGWTVR
QVVHHLADSHMNSYIRFKLSLTEETPAIRPYDEKAWSELKDSKTADPSGSLALLQELHGRWT
ALLRTLTDQQFKRGFYHPDTKEIITLENALGLYVWHSHHHIAHITELSRRMGWS
11111111100000000000000000000000000000000000000000000000000000
00000000000000000000000000000000000000000000000000000000000000
000000000000000000000000000000000000000000000000000000
>DM_train1722
APLVHHHHHHALANPAICRYPLGMSGGQIPDEDITASSQWSESTAAKYGRLDSEEGDGAWCP
EIPVEPDDLKEFLQIDLHTLHFITLVGTQGRHAGGHGIEFAPMYKINYSRDGTRWISWRNRH
GKQVLDGNSNPYDIFLKDLEPPIVARFVRFIPVTDHSMNVCMRVELYGCVWLDG
11111111111110000000000000000000000000000000000000000000000000
00000000000000000000000000000000000000000000000000000000000000
000000000000000000000000000000000000000000000000000001
>DM_train1723
MLAKVLKKRGAVLRGDFVLSSGRRSSVYIDMRRLLGDESSYSVALDLLLEVGGQDLARSSAV
IGVATGGLPWAAMLALRLSKPLGYVRPERKGHGTLSQVEGDPPKGRVVVVDDVATTGTSIAK
SIEVLRSNGYTVGTALVLVDRGEGAGELLARMGVRLVSVATLKTILEKLGWGGE
00000000000000000000000000000000000000000000000000000000000000
00000000000000000000000000111111111000000000000000000000000000
000000000000000000000000000000000000000000000000000111
>DM_train1725
AVSFIGSTENDVGPSLGSYSRTHAMDNLPFVYDTRNKIGYQNANVWHISKGFCVGLDGKVDL
PVVGSLDGQSIYGLTEEVGLLIWMGDTKYSRGTAMSGNSWENVFSGWCVGANTASTQGLSVR
VTPVILKRNSSARYSVQKTSIGSIRMRPYNGSSAGSVQTTVNFSLNPFTLNDT
00000000000000000000011111100000000000000000000000000000000000
00000000000000000000000000000000000000000000000000000000000000

00000000000000000000000000000000000000000000000000000
>DM_train1726
MSIEKLKAALPEYAKDIKLNLSSITRSSVLDQEQLWGTLLASAAATRNPQVLADIGAEATDH
LSAAARHAALGAAAIMGMNNVFYRGRGFLEGRYDDLRPGLRMNIIANPGIPKANFELWSFAV
SAINGCSQCLVAHEHTLRTVGVDREAIFEALKAAAIVSGVAQALATIEALSPS
00000000000000000000000000000000000000000000000000000000000000
00000000000000000000000000000000000000000000000000000000000000
00000000000000000000000000000000000000000000000111111
>DM_train1727
GKSRQRSPANCPTIKLKRQWGGKPSLGLHYQVRPIRYVVIHHTVTGECSGLLKCAEILQNMQ
AYHQNELDFNDISYNFLIGNDGIVYEGTGWGLRGAHTYGYNAIGTGIAFIGNFVDKLPSDAA
LQAAKDLLACGVQQGELSEDYALIAGSQVISTQSPGLTLYNEIQEWPHWLSNP
11111111110000000000000000000000000000000000000000000000000000
00000000000000000000000000000000000000000000000000000000000000
00000000000000000000000000000000000000000000000000000
>DM_train1728
MLDAFSRVVTNADSKAAYVGGADLQALKKFISEGNKRLDSVNSIVSNASCIVSDAVSGMICE
NPSLISPSGNCYTNRRMAACLRDGEIILRYVSYALLSGDASVLEDRCLNGLKETYSSLGVPA
NSNARAVSIMKACAVAFVNNTASQKKLSTPQGDCSGLASEVGGYFDKVTAAIS
11000000001000000000000000000000000000000000000000000000000000
00000000000000000000000000000000000000000000000000000000000000
00000000000000000000000000000000000000000000000000000
>DM_train1729
IIAMEKFAEYALSFGTEILTVEYGPLPIGKIVSEEINCSVYSVDPEGRVYTQAIAQWHDRGE
QEVLEYELEDGSVIRATSDHRFLTTDYQLLAIEEIFARQLDLLTLENIKQTEEALDNHRLPF
PLLDAGTIKMVKVIGRRSLGVQRIFDIGLPQDHNFLLANGAIAAACFNISTGP
11000000000000000000000000000000000000000000000000000000000000
00000000000000000000000000000000000000000000001111111110000000
00000000000000000000000000000000000000000000000011111
>DM_train1730
MGHHHHHHSHPVETSKFKFKCYGDFWLSVNRIYPEKRIELQLEVFKKLQDEKLYIVGWFSKG
DHAERYARKIMKIAPDNVKFLGSVSEEELIDLYSRCKGLLCTAKDEDFGLTPIEAMASGKPV
IAVNEGGFKETVINEKTGYLVNADVNEIIDAMKKVSKNPDKFKKDCFRRAKEF
11111111111000000000000000000000000000000000000000000000000000
00000000000000000000000000000000000000000000000000000000000000
00000000000000000000000000000000000000000000000000000
>DM_train1731
ISSASGESQMSLDEIEDVYHTRPGYRPEEYRWGQGGAKIIDYHIQSAGVDFPPSLTGNQQTD
FLMKVVFEYDFDCVVPGILIKTLDGLFLYGTNSFLASEGRENISVSRGDVRVFKFSLPVDLN
SGDYLLSFGISAGNPQTDMTPLDRRYDSIILHVTKSMDFWGVIDLKSSFTSYQ
11111111000000000000000000000000000000000000000000000000000000
00000000000000000000000000000000000000000000000000000000000000
00000000000000000000000000000000000000000000000000011
>DM_train1732

GMPQTLKPDTPAGTPAAGAVAFRRQVLPQDALLVRRVVESTGFFTPEEADVAQELVDEHLMH
GAACGYHFVFATEDDDMAGYACYGPTPATEGTYDLYWIAVAPHRQHSGLGRALLAEVVHDVR
LTGGRLLFAETSGIRKYAPTRRFYERAGFSAEAVLKAFYRAGDDKIIYRLEVA
111111111111111111000000000000000000000000000000000000000000
110000000000000000000000000000000000000000000000000000000000
00000000000000000000000000000000000000000000000000000
>DM_train1733
MRTSKKEMILRTAIDYIGEYSLETLSYDSLAEATGLSKSGLIYHFPSRHALLLGMHELLADD
WDKELRDITRDPEDPLERLRAVVVTLAENVSRPELLLLIDAPSHPDFLNAWRTVNHQWIPDT
DDLENDAHKRAVYLVQLAADGLFVHDYIHDDVLSKSKRQAMLETILELIPSQT
110000000000000000000000000000000000000000000000000000000000
000000000000000000000000000000000000000000000000000000000000
00000000000000000000000000000000000000000000000000111
>DM_train1734
MVKLTAELIEQAAQYTNAVRDRELDLRGYKIPVIENLGATLDQFDAIDFSDNEIRKLDGFPL
LRRLKTLLVNNNRICRIGEGLDQALPDLTELILTNNSLVELGDLDPLASLKSLTYLCILRNP
VTNKKHYRLYVIYKVPQVRVLDFQKVKLKERQEAEKMFKGKRGAQLAKDIAR
000000000000000000000000000000000000000000000000000000000000
000000000000000000000000000000000000000000000000000000000000
0000000000000000000000000000000000000001111111111111
>DM_train1735
ATDTLANEDCFRHESLVPNLDYERFRGSWIIAAGTSEALTQYKCWIDRFSYDDALVSKYTDS
QGKNRTTIRGRTKFEGNKFTIDYNDKGKAFSAPYSVLATDYENYAIVEGCPAAANGHVIYVQ
LRMTLRRFHPKLGDKEMLQHYTLDQVNQNKKAIEEDLKHFNLKYEDLHSTCH
111111000000000000000000000000000000000000000000000000000000
000000000000000000000000000000000000000000000000000000000000
0000000000011111100000000000000000000000000000000000
>DM_train1736
SHQVGVAGAVFDESTRKILVVQDRNKLKNMWKFPGGLSEPEEDIGDTAVREVFEETGIKSEF
RSVLSIRQQHTNPGAFGKSDMYIICRLKPYSFTINFCQEECLRCEWMDLNDLAKTENTTPIT
SRVARLLLYGYREGFDKIDLTVEELPAVYTGLFYKLYHKELPENYKTMKGID
110000000000000000000000000000000000000000000000000000000000
000000000011111111100000000000000000000000000000000000000000
0000000000000000000000000000000000000000000000001111
>DM_train1737
MSSHTATKYDVFLSFRGHDTRHNFISFLYKELVRRSIRTFKDDKELENGQRFSPELKSPIEV
SRFAVVVVSENYAASSWCLDELVTIMDFEKKGSITVMPIFYGVEPNHVRWQTGVLAEQFKKH
ASREDPEKVLKWRQALTNFAQLSGDCSGDDDSKLVDKIANEISNKKTIYATI
111111000000000000000000000000000000111111111111110000
000000000000000000000000000000000000000000000000000000000000
0000000000000000000000000000000000000000000000011111
>DM_train1738
HHHHHHGCMSGRKPIIGVMGPGKADTAENQLVMANELGKQIATHGWILLTGGRSLGVMHEAM
KGAKEAGGTTIGVLPGPDTSEISDAVDIPIVTGLGSARDNINALSSNVLVAVGMGPGTAAEV

ALALKAKKPVVLLGTQPEAEKFFTSLDAGLVHVAADVAGAIAAVKQLLAKLN
11111111111000000000000000000000000000000000000000000000000000
00000000000000011111100000000000000000000000000000000000000000
0000000000000000000000000000000000000000000000000011
>DM_train1739
MAPEAQVSVQPNFQQDKFLGRWFSAGLASNSSWLREKKAALSMCKSVVAPATDGGLNLTSTF
LRKNQCETRTMLLQPAGSLGSYSYRSPHWGSTYSVSVVETDYDQYALLYSQGSKGPGEDFRM
ATLYSRTQTPRAELKEKFTAFCKAQGFTEDTIVFLPQTDKCMTEQAHHHHHH
11111100000000000000000000000000011111100000000000000000000000
00000000000000000000000000000000000000000000000000000000000000
0000000000000000000000000000000000000111111111111111
>DM_train1740
SATTSSKNGKPLVVVYGDYKCPYCKELDEKVMPKLRKNYIDNHKVEYQFVNLAFLGKDSIVG
SRASHAVLMYAPKSFLDFQKQLFAAQQDENKEWLTKELLDKHIKQLHLDKETENKIIKDYKT
KDSKSWKAAEKDKKIAKDNHIKTTPTAFINGEKVEDPYDYESYEKLLKDKIK
11111111000000000000000000000000000000000000000000000000000000
00000000000000000000000000000000000000000000000000000000000000
0000000000000000000000000000000000000000000000000111
>DM_train1741
LRPPLPPATPQEVQFVQHMLQHHAQALDLAAPMLERSQQRTVRSLALDIQLSQREQMRQMEA
MLGRWGQPPGEPISPEHARMMGMASEAEVAGLSTLPVEQAERQFLRLMIRHHQGAVAMTLPM
LDAAARPEVERLARQIVVTQRGEIRTMEGVLGRLDGEVPAAPMRPVEHGHGH
11111000000000000000000000000000000000000000000000000000000000
00000000000000000000000000000000000000000000000000000000000000
0011000000000000000000000000000000111111111111111111
>DM_train1742
AITGIFFGSDTGNTENIAKMIQKQLGKDVADVHDIAKSSKEDLEAYDILLLGIPTWYYGEAQ
CDWDDFFPTLEEIDFNGKLVALFGCGDQEDYAEYFCDALGTIRDIIEPRGATIVGHWPTAGY
HFEASKGLADDDHFVGLAIDEDRQPELTAERVEKWVKQISEELHLDEILNA
00000000000000000000000000000000000000000000000000000000000000
00000000000000000000000000000000000000000000000000000000000000
000000000000000000000000000000000000000000000111111
>DM_train1743
IFGIDDLIIGVLFVAIVETGIGGYLLGSRKESGGGVTKESAEKGFEKIGNDIQILKSSINIA
IEKLNDRISHDEQAIRDLTLEIENARSEALLGELGIIRALLVGNISIGLQESLWELASEITN
RAGDLAVEVSPGCWIIDNNICDQSCQNFIFKFNETAPVPTIPPLDTKIDLQ
11100000000000000000000000000000000000000000000000000000000000
00000000000000000000000000000000000000000000000000000000000000
000000000000000000000000000000000000000000111111111
>DM_train1744
MSTTNHDHHIYVLMGVSGSGKSAVASEVAHQLHAAFLDGDFLHPRRNIEKMASGEPLNDDDR
KPWLQALNDAAFAMQRTNKVSLIVCSALKKHYRDLLREGNPNLSFIYLKGDFDVIESRLKAR
KGHFFKTQMLVTQFETLQEPGADETDVLVVDIDQPLEGVVASTIEVIKKGK
00000000000000000000000000000000000000000000000000000000000000

000000000000000000000000000000000000000000000000000000000111
1111110000000000000000000000000000000000000000000
>DM_train1745
MAKKVAVLAVNPVNGCGLFQYLEAFFENGISYKVFAVSDTKEIKTNSGMVLIVDDVIANLKG
HEDEFDALVFSCGDAVPVFQQYANQPYNVDLMEVIKTFGEKGKMMIGHCAGAMMFDFTGITK
GKKVAVHPLAKPAIQNGIATDEKSEIDGNFFTAQDENTIWTMLPKVIEALK
110000000000000000000000000000000000000000000000000000000000
000000000000000000000000000000000000000000000000000000000000
0000000000000000000000000000000000000000000000000
>DM_train1746
METYPITVGGVTRHVPLIEPLPGRRIPLVEFLGDPEFTRAAAEALRPLVPKEAEILFTTETS
PIPLTHVLAEALGLPYVVARRRRRPYMEDPIIQEVQTLTLGVGEVLWLDRRFAEKLLNQRVV
LVSDVVASGETMRAMEKMVLRAGGHVVARLAVFRQGTPGLAVDTVAELPVL
100000000000000000000000000000111000000000000000000000000000
000000000111100000000000000000000000000000001111111100000000000000110000
0000000011000001000000000110000000000000000000000
>DM_train1747
MSLAKNIVFIGFMGSGKSTLARALAKDLDLVFLDSDFLIEQKFNQKVSEIFEQKRENFFREQ
EQKMADFFSSCEKACIATGGGFVNVSNLEKAGFCIYLKADFEYLKKRLDKDEISKRPLFYDE
IKAKKLYNERLSKYEQKANFILNIENKNIDELLSEIKKVIKEGGSHHHHHH
100000000000000000000000000000000000000000000000000000000000
000000000000000000000000000000000000000000000001111111111000
0000000000000000000000000000000000000000011111111
>DM_train1748
AHHHHHHGSMLKEVHELLNRIWGDIFELREELKEELKGFTVEEVSEVFNAYLYIDGKWEEMK
YPHPAFAVKPGGEVGATPQGFYFVFAFPKEELSKEFIEDVIRAFEKLFIYGAENFLEDFYNF
EHPISGDEVWDRIVNSDEEMINFEVDLGFDKEEVKREIKRFIELARRYNLL
111111111000000000000000000000000000000000000000000000000000
000000000000000000000000000000000000000000000000000000000000
0000000000000000000000000000000000000000000000000
>DM_train1749
PLGSPNSSIVSLLGIKVLNNPAKFTDPYEFEITFECLESLKHDLEWKLTYVGSSRSLDHDQE
LDSILVGPVPVGVNKFVFSADPPSAELIPASELVSVTVILLSCSYDGREFVRVGYYVNNEYD
EEELRENPPAKVQVDHIVRNILAEKPRVTRFNIVWDNENEGDLYPPEQPGV
111111000000000000000000000000000000000000000000000000000000
000000000000000000000000000000000000000000000000000000000000
0000000000000000000000000000000000000000000011111
>DM_train1751
MAFPDTDVSPPRGGPSSPAKSPLLRSFKVKTYIPKNDEQNWVVVDASGVPLGRLATLIASRI
RGKHRPDFTPNMIQGDFVVVINAAQVALTGKKLDDKVYTRYTGYQGGLKTETAREALSKHPE
RVIEHAVFGMLPKGRQGRAMHTRLKVYAGETHPHSAQKPQVLKTQPLEVK
111111111111111111111111111000000000000000000000000000000000
000000000000000000000000000000000000000000000000000000000000
000000000000000000000000000000000000000000000111

>DM_train1752
AGKTMIPLLAFAAWSGTGKTTLLKKLIPALCARGIRPGLIKHTHHDMDVDKPGKDSYELRKA
GAAQTIVASQQRWALMTETPDEEELDLQFLASRMDTSKLDLILVEGFKHEEIAKIVLFRDGA
GHRPEELVIDRHVIAVASDVPLNLDVALLDINDVEGLADFVVEWMQKQNG
111100000000000000000000000000000000000001111111111100000
000000000000000000000000000000000000000000000000000000000000
00000000000000000000000000000000000000000000000000
>DM_train1753
MKGPNLTEISKKITDSNAVLLAVKEVEALLSSIDEIAAKAIGKKIHQNNGLDTENNHNGSLL
AGAYAISTLIKQKLDGLKNEGLKEKIDAAKKCSETFTNKLKEKHTDLGKEGVTDADAKEAIL
KTNGTKTKGAEELGKLFESVEVLSKAAKEMLANSVKELTSPVVAESPKKP
111000000000000000000000000000000000000000000000000000000000
000000000000000000000000000000000000000000000000000000000000
00000000000000000000000000000000000000000111111111
>DM_train1754
GMDIMLNTADISAAAILLPVEGAQLSELRQIPAEGGPVLHMLRLDSPQFSQFGEIYFSEVLP
RRVKAWKRHSLMTQLFAVPVGCIHVVLYDGREKSPTSGRLAQVTLGRPDNYRLLRIPPQVWY
GFAATGDTPALVANCTDIPHRQGESERAPQDAPFIPFSWAGADLSGTPVM
111111111111110000000000000000000000000000000000000000000000
000000000000000000000000000000000000000000000000000000000000
00000000000000000000000000000000000000000000000111
>DM_train1755
DDIKVFQNEVLGTLQRGKEENISCDNLVLEINSLKYAYNISLKEVMQVLSHVVLEFPLQQMD
SPLDSSRYCALLLPLLKAWSPVFRNYIKRAADHLEALAAIEDFFLEHEALGISMAKVLMAFY
QLEILAGETILSWFSQRDTTDKGQQLRKNQQLQRFIQWLKEAEEESSEDD
000000000000000000000000000000000000000000000000000000000000
000000000000000000000000000000000000000000000000000000000000
00000000000000000011000000000000000000000000111111
>DM_train1756
MAHHHHHHMSDEDHDFSHQGGGDNASKTYPLAAGALKKGGYVCINGRPCKVIDLSVSKTGKH
GHAKVSIVATDIFTGNRLEDQAPSTHNVEVPFVKTYTYSVLDIQANEDPSLPAHLSLMDDEG
ESREDLDMPPDPALATQIKEQFDSGKDVLVVVVSAMGTEQVLQTKNAAEK
111111111111111111111111110000000000000000000000000000000111
100000000000000000000000000000000000000000000000000000000000
00000000000000000000000000000000000000000000000001
>DM_train1757
GSQQIKDPLNYEVEPFTFQNQDGKNVSLESLKGEVWLADFIFTNCETICPPMTAHMTDLQKK
LKAENIDVRIISFSVDPENDKPKQLKKFAANYPLSFDNWDFLTGYSQSEIEEFALKSFKAIV
KKPEGEDQVIHQSSFYLVGPDGKVLKDYNGVENTPYDDIISDVKSASTLK
110000000000000000000000000000000000000000000000000000000000
000000000000000000000000000000000000000000000000000000000000
00000100000000000000000000000000000000000000000000
>DM_train1758
AKALEQPFDVANIPGPKMATLLEKGKPVANMIKKAKRPLLIVGPDMTDEMFERVKKFVEKDI

TVVATGSAITRFIDAGLGEKVNYAVLHELTQFLLDPDWKGFDGQGNYDLVLMLGSIYYHGSQ
MLAAIKNFAPHIRALAIDRYYHPNADMSFGNLWKKEEDYLKLLDEILAEL
11111111111111111000000000000000000000000000000000000000000000
00000000000000000000000000000000000000000000000000000000000000
00000000000000000000000000000000000000000000000000
>DM_train1759
EFMIIYRDLISHDEMFSDIYKIREIADGLCLEVEGKMVSRTEGNIDDSLIGGNASAEGPEGE
GTESTVITGVDIVMNHHLQETSFTKEAYKKYIKDYMKSIKGKLEEQRPERVKPFMTGAAEQI
KHILANFKNYQFFIGENMNPDGMVALLDYREDGVTPYMIFFKDGLEMEKC
00000000000000000000000000000000000000001111111111111111111111
11111111000000000000000000000000000000000000000000000000000000
00000000000000000000000000000000000000000000000000
>DM_train1760
MDINVVNALAYEDFVKLFGNVVEKCPLISAAIWSYRPFKDLADIEARISEFIHSLPDSGKEG
ILRCHPDLAGRDLQSGTLTPESQEEQSQAGMTTLDSAEIVHMYRLNSEYKERFGFPFVICAR
LNNKADIVRQLSERLKNRRTAELECAIEEVKKICSLRLHSIVLSDIQTKL
10000000000000000000000000000000000000000000000000000000000000
00000000000000000000000000000000000000000000000000000000000000
00000000000000000000000000000000000000000111111111
>DM_train1762
MRIQEAIAQDKTISVIIDPSQIGSTEGKPLLSMKCNLYIHEILSRWKASLEAYHPELFLDTK
KALFPLLLQLRRNQLAPDLLISLATVLYHLQQPKEINLAVQSYMKLSIGNVAWPIGVTSVGI
HARSAHSKIQGGRNAANIMIDERTRLWITSIKRLITFEEWYTSNHDSLA
00000000000000000000000000000000000000000000000000000000000000
00000000000000000000000000000000000000000000000000000000011111
1111111111111110000000000000000000000000000001111
>DM_train1763
MALVDGFLELERSSGKLEWSAILQKMASDLGFSKILFGLLPKDSQDYENAFIVGNYPAAWRE
HYDRAGYARVDPTVSHCTQSVLPIFWEPSIYQTRKQHEFFEEASAAGLVYGLTMPLHGARGE
LGALSLSVEAENRAEANRFMESVLPTLWMLKDYALQSGAGLAFEHPVSK
11111100000000000000000000000000000000000000000000000000000000
00000000000000000000000000000000000000000000000000000000000000
0000000000000000000000000000000000000000000001111
>DM_train1764
MCIIKPMDDVVKFIHEVGSLKLTPRSGWLKLGIRLPESVAEHNFRAAIIAFILALKSGESVE
KACKAATAALFHDLHEARTMDLHKIARRYVSCDEEGAREEQLSWMESKPDFSDVEVYVSDAD
KLELAFQGVEYSQQVSYAIRFAENVELKTDAAKEIYRVLMERKNPVWWR
11111100000000000000000000000000000000000000000000000000000000
00000000000000000000000000000000000000000000000000000000000000
0000000000000000000000000000000000000000000000000
>DM_train1765
GHMGFFSRDIQTMEDLLLHGLRDIYYAEQQITKALPKMIEQATNRDLSQGLTSHLEETQKQI
ERLDQVFKKLGQKPSGVNCPAIDGLIKEADETAGEIADKTVLDAAIVANAQAVEHYEIARYG
TLIAWAEELGHDDIVRFLTTNLNEEKAANTKLNTVALRKGVNRKAASGS

11110000000000000000000000000000000000000000000000000000000000
00000000000000000000000000000000000000000000000000000000000000
0000000000000000000000000000000000000111111111111
>DM_train1766
MHHHHHHSSGRENLYFQGMSDSGSQLGSMGSLTMKSQLQITVISAKLKENKKNWFGPSPYVE
VTVDGQSKKTEKCNNTNSPKWKQPLTVIVTPVSKLHFRVWSHQTLKSDVLLGTAALDIYETL
KSNNMKLEEVVVTLQLGGDKEPTETIGDLSICLDGLQLESEVVTNGETT
11111111111111111111111111111100000000000000000000011000000000
00000000000000000000000000000000000000000000000000000000000000
0000000000000000000000000000000000000001111111111
>DM_train1767
SNAMRVIVTEYQPAWVEQFEEEAQALKQILKENCLKVEHIGSTSVPNLAAKPIIDFLVIVEE
IEKVDLLQWEFERIGYEYMGEFGLSGRRYLRKGPIKRTHHVHIYQFDNTQEILRHLAFRNYL
RENPAIATTYGTLKKQLAQAHPDSIDKYMDGKDAFIKKIEKEALKKYWE
11111100000000000000000000000000000000000000000000000000000000
00000000000000000000000000000000000000000000000000000000000000
0000000000000000000000000000011000000000000000000
>DM_train1768
SSKKYKLSEIHHLHPEYVRVSEHFKASMKNFKIEKIKKIENSELLDKFTWKKSQMKEEGKLL
FYATSRAYVESICSNNFDSFLHETHENKYGKGIYFAKDAIYSHKNCPYDAKNVVMFVAQVLV
GKFIEGNITYTSPPPQFDSCVDTRSNPSVFVIFQKDQVYPQYVIEYTED
11000000000000000000000000000000000000000000000000000000000000
00000000000000000000000000000000000000000000000000000000000000
0000000000000000000000000000000000000000000000000
>DM_train1769
MGSQVQKSDEITFSDYLGLMTCVYEWADSYDSKDWDRLRKVIAPTLRIDYRSFLDKLWEAMP
AEEFVGMVSSKQVLGDPTLRTQHFIGGTRWEKVSEDEVIGYHQLRVPHQRYKDTTMKEVTMK
GHAHSANLHWYKKIDGVWKFAGLKPDIRWGEFDFDRIAEDGRETFGDK
11111111000000000000000000000000000000000000000000000000000000
00000000000000000000000000000000000000000000000000000000000000
000000000000000000000000000000011111111111111111
>DM_train1770
GMNLQTDQTTTTADESAIRAFHRQMIDAWNRGSGEGFAAPFSETADFITFEGTHLKGRKEIA
AFHQQAFDTVVKGTRLEGEVDFVRFVNSQLALMLVVIRVILPGQTETSASRDSLPLYVVTKG
DEGWQIEGLLNTRKLTLERQFFLDDFDSLSAEAQRQVTDLVASLKQSH
11111111100000000000000000000000000000000000000000000000000000
00000000000000000000000000000000000000000000000000000000000000
000000000000000000000000000000000000000000000001
>DM_train1771
GMSEALSDSGLPPVQRVVTGHDAHGRAVFKSEDVTPTRMIPSGDASFLLVWTTATVPADNND
ETDGRQREAGLTLDGGSVIRVVDMLPGKESPMHRTNSIDYGIVLEGEIELELDDGAKRTVRQ
GGIIVQRGTNHLWRNTTDKPCRIAFILIEAPAYLHNGQPLPEDKPDHK
11100000000000000000000000000000000000000000000000000000000000
00000000000000000000000000000000000000000000000000000000000000

000000000000000000000000000000000000000000111111
>DM_train1772
MNLDNVTGGDNVPKEINVIIEISQNSCPVKYEFDKEKNLFCVDRFLPTSMYYPCNYGFIPHT
CAGDGDPVDVLVASRFPVMSGAVIRARPVGVLVMHDESGEDVKILAVPTHKVDQYYNNIKDY
SDFPVSFLNSISHFFTFYKKLEEDKFVSVEGWKDVTVAEKLILSALIK
11100000000000000000000000000000000000000000000000000000000000
00000000000000000000000000000000000011100000000000000000000000
000000000000000000111111110000000000000000000011
>DM_train1773
MNPHPVEPRDQIAELLVESPLFSFNCAHFIAFKGFRETLHGHNYNVSLRLRGNIQGDGYVID
FSILKEKVRKVCKQLDHHFILPMYSDVLNIQEVNDNFKITCEDNSEYSFPKRDCVQIPIKHS
STEEIGLYILNQLIEEIDLPFLKTRSVNYMEVTVSESPSQKATVHRNI
11111111100000000000000000000000000000000000000000000000000000
00000000000000000000000000000000000000000000000000000000000000
000000000000000000000000000000000000000000000000
>DM_train1774
EKEKKNVFMRTFEAISRNFSEIFAKLSPGGSARLILENPEDPFSGGLEIEAKPAGKDVKRIE
AMSGGEKALTALAFVFAIQKFKPAPFYLFDEIDAHLDDANVKRVADLIKESSKESQFIVITL
RDVMMANADKIIGVSMRDGVSKVVSLSLEKAMKILEEIRKKQGWEHGN
11111100000000000000000000000000000000000000000000000000000000
00000000000000000000000000000000000000000000000000000000000000
000000000000000011000000000000000000000000001111
>DM_train1775
LPPREDAARVARFVTHVSDWGALATISTLEAVRGRPFADVLSLSDGPPGAGSGVPYFYLSPL
QLSVSNLQENPYATLTMTLAQTNFCKKHGFDPQSPLCVHIMLSGTVTKVNETEMDIAKHSLF
IRHPEMKTWPSSHNWFFAKLNITNIWVLDYFGGPKIVTPEEYYNVTVQ
00000000000000000000000000000000000000000000000000000000000000
00000000000000000000000000000000000000000000000000000000000000
000000000000000000000000000000000000000000000011
>DM_train1776
QPQSGPQTGVVFRNTVTGEALDVSQGKEGGRDTPAVKKFLETGENLYIDDKSCLRNGESLFA
TSCSGCHGHLAEGKLGPGLNDNYWTYPSNTTDVGLFATIFGGANGMMGPHNENLTPDEMLQT
IAWIRHLYTGPKQDAVWLNDEQKKAYTPYKQGEVIPKDAKGQCKPLDE
11111111111111111111111000000000000000000000000000000000000000
00000000000000000000000000000000000000000000000000000000000000
000000000000000000000000000000000000000000000000
>DM_train1777
SLEKQIESYYQEIAQLIIDMIPEEWAEVRFYAQEDHDGWKIFFFHYLSASSDEWTKDIDIRD
VIKVPQDEFMEKYNELSFCISDFRKDYAEAFGEPWMSFQMTFYASGKFNIDFYYDKNPFDTF
LTRLAWQYEHFGTIPEDSFYKETLNEYLEEKAQGKRYPFLEPLKEEEG
00000000000000000000000000000000000000000000000000000000000000
00000000000000000000000000000000000000000000000000000000000000
000000000000000100000000000000000000000000011111
>DM_train1778

SNAMNPFETKSVTFAEPIEMLYACHGKVRRFCGQVAMLSDYIAENGCNQIVLQTIRQIAQYF
NVAAPLHHEDEEENFFPLLLQYAPQAQESVDELLRQHIGLHDNWAAVSAEFAKLEADNAYVP
DEEAFKRFVAGYDVHLAIEEPLFDMGNTFIPKEKLTEIGEIMAARRRK
11111111111111000000000000000000000000000000000000000000000000
00000000000000000000000000000000000000000000000000000000000000
000000000000000000000000000000000000000000000000
>DM_train1779
SKLQATKTLAADVIMRSPVSWKQELTLDAGRSKGASENMLAIANGGLIGSVSKVEENSTIVN
LLTNTENADKISVKIQHGSTTIYGIIIGYDKENDVLKISQLNSNSDISAGDKVTTGGLGNFN
VADIPVGEVVATTHSTDYLTREVTVKLSADTHNVDVIELVGNSKLVPR
00000000000000000000000000000000000000000000000000000000000000
00000000000000000000000000000000000000000000000000000000000000
000000000000000000000000000000000000000000011111
>DM_train1780
GMDAGRGAEHDEAVTSDDSASVEVPRRATPADAATVAQMLHDFNTEFGAPTPGTDELASRLS
HLLAGEDVVVLLAGEPPTGLAVLSFRPNVWYPGPVAILDELYVRPGRRGHRLGSALLAASCG
LVRSRGGALLEINVDGEDTDARRFYEARGFTNTEPNGTEPMLYYYREL
11111111111111110000000000000000000000000000000000000000000000
00000000000000000000000000000000000000000000000000000000000000
000000000000000000000000000000000000000000000000
>DM_train1781
GSHMNQCPEHSQLTTLGVDGKEFPEVHLGQWYFIAGAAPTKEELATFDPVDNIVFNMAAGSA
PMQLHLRATIRMKDGLCVPRKWIYHLTEGSTDLRTEGRPDMKTELFSSSCPGGIMLNETGQG
YQRFLLYNRSPHPPEKCVEEFKSLTSCLDSKAFLLTPRNQEACELSNN
10000000000000000000000000000000000000000000000000000000000000
00000000000000000000000000000000000000000000000000000000000000
000000000000000000000000000000000000000000000001
>DM_train1782
APKDNTWYTGAKLGWSQYHDTGLINNNGPTHENKLGAGAFGGYQVNPYVGFEMGYDWLGRMP
YKGSVENGAYKAQGVQLTAKLGYPITDDLDIYTRLGGMVWRADTYSNVYGKNHDTGVSPVFA
GGVEYAITPEIATRLEYQWTNNIGDAHTIGTRPDNGMLSLGVSYRFG
00000000000000000111111111111100000000000000000000000000000000
01111111000000000000000000000000000000000000000000000000000000
00000000000000000000011111111111111000000000000
>DM_train1783
SNSGEPAPVLSSPTPTDLSTFLSFPSPEKLLRLGPKVSVLIVQQTDTSDPEKVVSAFLKVAS
VFRDDASVKTAVLDAIDALMKKAFSCSSFNSNTFLTRLLIHMGLLKSEDKIKAIPSLHGPLM
VLNHVVRQDYFPKALAPLLLAFVTKPNGALETCSFARHNLLQTLYNI
11111111111111110000000000000000000000000000000000000000000000
00000000000000000000000000000000000000000000000000000000000000
00000000000000000000000000000000000000000000000
>DM_train1784
GAMGSMAEAEGESLESWLNKATNPSNRQEDWEYIIGFCDQINKELEGPQIAVRLLAHKIQSP
QEWEALQALTVLEACMKNCGRRFHNEVGKFRFLNELIKVVSPKYLGDRVSEKVKTKVIELLY

SWTMALPEEAKIKDAYHMLKRQGIVQSDPPIPVDRTLIPSPPPRPKN
111111111111000000000000000000000000000000000000000000000000
000000000000000000000000000000000000000000000000000000000000
000000000000000000000000000000000001111111111
>DM_train1785
AALGPSSQNVTEYVVRVPKNTTKKYNIMAFNAADKVNFATWNQARLERDLSNKKIYQEEEMP
ESGAGSEFNRKLREEARRKKYGIVLKEFRPEDQPWLLRVNGKSGRKFKGIKKGGVTENTSYY
IFTQCPDGAFEAFPVHNWYNFTPLARHRTLTAEEAEEEWERRNKVLN
111110000000000000000000000000000000000000000000000000000001
111111111000000000000000000000000000000000000000000000000000
000000000000000000000000001111111111111111111
>DM_train1786
SYRSTQQITDFTKEILVNGEAVTAFDRQGDLPNVVVTPNFEAGVDQVVDQLAMNDSERDTTA
IIGKSLAECEALTKALKARGEQVTLIQTENQRLAPGVIVVPSFLAKGLEFDAVIVWNANQEN
YQREDERQLLYTICSRAMHELTLVAVGSLSPLLARVNHALYTLNEAK
000000000000000001111111100000000000000000000000000000000000
000000000000000000000000000001000000000000000000000000000000
000000000000000000000000000000000000000000011
>DM_train1787
NQPDWADEAANGAHQDAWKSLKARVENVYYMVKATYKNDPVWGNDFTCVGVMANDVNEDEKS
IQAEFLFMNNADTNMQFATEKVTAVKMYGYNRENAFRYETEDGQVFTDVIAYSDDNCDVIYV
PGTDGNEEGYELWTTDYDNIPANCLNKFNEYAVGRETRDVFTSACLE
100000000000000000000000000000000000000000000000000000000000
000000000000000000000000000000000000000000000000000000000000
000000000000000000000000000000000000000000000
>DM_train1788
SSQTSNFLAEQYERDRKAIINCCFSRPDHKTGEPPNNYITHVRIIEDSKFPSSRPPPDSKLE
NKKKRLLILSAKPNNAKLIQIHKARENSDGSFQIGRTWQLTELVRVEKDLEISEGFILTMSK
KYYWETNSAKERTVFIKSLITLYIQTFEGHVPELVNWDLSLFYLDER
111110000000000000000000011111111000000000000000000000000000
000000000000000000000000000000000000000000000000000000000000
000000000000000000000000000000000000000000000
>DM_train1789
MKEQLYTGLTEKEANQMQALLLSNDVNVSKEMDKSGNMTLSVAAADFVRAITILNNNGFPKK
KFADIEVIFPPSQLVASPSQENAKINYLKEQDIERLLSKIPGVIDCSVSLNVNNNESQPSSA
AVLVISSPEVNLAPSVIQIKNLVKNSVDDLKLENISVVIKSSSGQDG
100000000000000000000000000000000000000000000000000000000000
000000000011111100000000000000000000000000000000001111110000
000000000000000000000000000000000000000011111
>DM_train1790
GSMSLFPGDIEELARRIITDFTPLGLMVSTAESCTGGLIAGALTEIAGSSAVVDRGFVTYTN
DAKRDMLGVGTETLTTFGAVSRQTALQMAHGALYRSRANFAVAVTGIAGPGGGSAEKPVGLV
HLATKARNGNVLHHEMRYGDIGRTEIRLATVRTALEMLIALNQAGSV
110000000000000000000000000000000000000000000000000000000000

0000000000000000000000000000000000000000000000000000000000000000
000000000000000000000000000000000000000000001111
>DM_train1791
MSLKTLDGKITRPTSDKVRGAIFNMIGPYFNGGRVLDLFAGSGGLAIEAVSRGMSAAVLVEK
NRKAQAIIQDNIIMTKAENRFTLLKMEAERAIDCLTGRFDLVFLDPPYAKETIVATIEALAA
KNLLSEQVMVVCETDKTVLLPKEIATLGIWKEKIYGISKVTVYVNEG
1111111111111111000000000000000000000000000000000000000000000000
0000000000000000000000000000000000000000000000000000000000000000
000000000000000000000000000000000000000000000000
>DM_train1792
GMGILSGNPQDEPLHYGEVFSTWTYLSTNNGLINGYRSFINHTGDEDLKNLIDEAIQAMQDE
NHQLEELLRSNGVGLPPAPPDRPAARLDDIPVGARFNDPEISATISMDVAKGLVTCSQIIGQ
SIREDVALMFSQFHMAKVQFGGKMLKLNKNKGWLIPPPLHSDRPIKE
1111111111000000000000000000000000000000000000000000000000000000
0000000000000000000000000000000000000000000000000000000000000000
000000000000000000000000000000000000000000011111
>DM_train1793
GMKQPNYYQDVKQFHQTFHHPGADQPTAIPLDRGVKRATWTAEEAVVEFLHQSSQNETEFLA
AIETFKAGLDQAVKKSLKETYPVTEVERLVGQGDALTDALYFIMGSFVEAGLEPGPLFEIVQ
QANMAKLGPDGQPIFRESDQKVMKPDGWLPPEPQLEAEVVRQMKEKA
1100000000000000000000000000000000000000000000000000000000000000
0000000000000000000000000000000000000000000000000000000000000000
000000000000000000000000000000000000000000000000
>DM_train1794
MIIKDILRENQDFRFRDLSDLKHSPKLCIITCMDSRLIDLLERALGIGRGDAKVIKNAGNIV
DDGVIRSAAVAIYALGDNEIIIVGHTDCGMARLDEDLIVSRMRELGVEEEVIENFSIDVLNP
VGDEEENVIEGVKRLKSSPLIPESIGVHGLIIDINTGRLKPLYLDE
0000000000111111111111110000000000000000000000000000000000000000
0000000000000000000000000000000000000000000000000000000000000000
00000000000000000000000000000000000000000000000
>DM_train1795
MEEPDANGYYYHDTFEGSVGQWTARGPAEVLLSGRTAYKGSESLLVRNRTAAWNGAQRALNP
RTFVPGNTYCFSVVASFIEGASSTTFCMKLQYVDGSGTQRYDTIDMKTVGPNQWVHLYNPQY
RIPSDATDMYVYVATADDTINFYIDEAIGAVAGTVIEGLEHHHHHH
1110000000000000000000000000000000000000000000000000000000000000
0000000000000000000000000000000000000000000000000000000000000000
00000000000000000000000000000000000001111111111
>DM_train1796
MAKLGIHSNDTRDAWVNKIAQLNTLEKAAEMLKQFRMDHTTPFRNSYELDNDYLWIEAKLEE
KVAVLKARAFNEVDFRHKTAFGEDAKSVLDGTVAKMNAAKDKWEAEKIHIGFRQAYKPPIMP
VNYFLDGERQLGTRLMELRNLNYYDTPLEELRKQRGVRVVHLQSPH
1110000000000000000000000000000000000000000000000000000000000000
0000000000000000000000000000000000000000000000000000000000000000
000000000000000000000000000000000000000000011111

>DM_train1797
GVPFFSCQRGYKGVWRGDGIMQTTCPCGAQITGHVKNGSMRIVGPRTCSNTWHGTFPINAYT
TGPCTPSPAPNYSRALWRVAAEEYVEVTRVGDFHYVTGMTTDNVKCPCQVPAPEFFTEVDGV
RLHRYAPACKPLLREEVTFLVGLNQYLVGSQLPCEPEPDVAVLTSM
00000000000000000000000000000000000000000000000000000000000000
00000000000000000000000000000000000000000000000000000000000000
0000000000000000000000000000000000001111111111
>DM_train1798
MNADEWMTTFRENIAQTWQQPEHYDLYIPAITWHARFAYDKEKTDRYNERPWGGGFGLSRWD
EKGNWHGLYAMAFKDSWNKWEPIAGYGWESTWRPLADENFHLGLGFTAGVTARDNWNYIPLP
VLLPLASVGYGPVTFQMTYIPGTYNNGNVYFAWMRFQFLEHHHHHH
11111110000000000000000000000000000111111111100000000000000000
00000000000000000000000000000000000000000000000000000000000000
0000000000000000000000000000000000000000111111
>DM_train1799
GMLKILATQFNGKLQTLTKQEDELFDVVRLLAQALVGQGKVYLDAYGEFEGLYPMLSDGPDQ
MKRVTKIKDHKTLHAVDRVLIFTPDTERSDLLASLARYDAWHTPYSIITLGDVTETLERSIA
PLALKFDKGLLPAEDGSRHGLPSLALGAFLLTHILTQLQEMTEEWE
00000000000000000000000000000000000000000000000000000000000000
00000000000000000000000000000000000000000000000000000000000000
0000000000000000000000000000000000000000000001
>DM_train1800
MSLKLISNDLRDGDKLPHRHVFNGMGYDGDNISPHLAWDDVPAGTKSFVVTCYDPDAPTGSG
WWHWVVVNLPADTRVLPQGFGSGLVAMPDGVLQTRTDFGKTGYDGAAPPKGETHRYIFTVHA
LDIERIDVDEGASGAMVGFNVHFHSLASASITAMFSEGGSHHHHHH
10000000000000000000000000000000000000000000000000000000000000
00000000000000000000000000000000000000000000000000000000000000
0000000000000000000000000000000000000111111111
>DM_train1801
MSTTPIDAELDLMLKRELAVPVNLVWRGLTEPELLKKWFVPKPWSISDCRVDLRPGGEFYTV
MQDPEGNKFPNSGCFLEVTDEKRLIWTSALVKNYRPAVPATTSDKECAHIVMTAVIELQPTS
SGTRYTACAMHNTPGQRKLHEEMGFHEGWGTTITQLEELLKQEKAY
11111000000000000000000000000000000000000000000000000000000000
00000000000000000000000000000000000000011111111111000000000000
0000000000000000000000000000000000000000000000
>DM_train1802
GAMGPVDEQWIEILRIQALCARYCLTINTQDGEGWAGCFTEDGAFEFDGWVIRGRPALREYA
DAHARVVRGRHLTTDLLYEVDGDVATGRSASVVTLATAAGYKILGSGEYQDRLIKQDGQWRI
AYRRLRNDRLVSDPSVAVNVADADVAAVVGHLLAARRLGTQMSDT
11111111000000000000000000000000000000000000000000000000000000
00000000000000000000000000000000000000000000000000000000000000
000000000000000000000000000000000000000000111
>DM_train1803
MAARTDNSIVVNAPFELVWDVTNDIEAWPELFSEYAEAEILRQDGDGFDFRLKTRPDANGRV

WEWVSHRVPDKGSRTVRAHRVETGPFAYMNLHWTYRAVAGGTEMRWVQEFDMKPGAPFDNAH
MTAHLNTTTRANMERIKKIIEDRHREGQRTPASVLPTELHAQQLLL
000000000000000000000000000000000000000000000000000000000000
000000000000000000000000000000000000000000000000000000000000
0000000000000000000000000000000111111111111111
>DM_train1804
MDPSQTPDANASISVSYKCGVKDGTKNTIRATINIKNTGTTPVNLSDIKVRYWFTSDGNEQN
NFVCDYAAFGTDKVKGIVKKIENSVPGADTYCEISFTEDAGRLAPGGSTGTIPFRIEGAAEY
DQTDDYSYNSEMSDDFGDNTKITAYIKDKLKYGVEAAALEHHHHHH
111110000000000000000000000000000000000000000000000000000000
000000000000000000000000000000000000000000000000000000000000
0000000000000000000000000000000000000011111111
>DM_train1806
AAVILESIFLKRSQQKKKTSPLNFKKCLFLLTVHKLSYYEYDFERGRRGSKKGSIDVEKITC
VETVVPEKNPPPERQIPRRGEESSEMEQISIIERFPYPFQVVYDEGPLYVFSPTEELRKRWI
HQLKNVIRYNSDLVQKYHPCFWIDGQYLCCSQTAKNAMGCQILEN
000000000000000000000000000000000000000000000000000000000000
000000000000000011111111100000000000000000000000000000000000
000000000000000000000000000000000000000000000
>DM_train1807
RNRIKKVTERLPLPPNARRVEDPEMNMSREVVEDPGVPGTQDVTFAVAEVNGVETGRLPVAN
VVVTPAHEAVVRVGTKPGTEVPPVIDGSIWDAIAGCEAGGNWAINTGNGYYGGVQFDQGTWE
ANGGLRYAPRADLATREEQIAVAEVTRLRQGWGAWPVCAARAGAR
000000000000000000000000000000000000000000000000000000000000
000000000000000000000000011000000000000000000000000000000000
000000000000000000000000000000000000000000000
>DM_train1808
SVLQAQMAAQQLPVIGGIAIPELGINLPIFKGLGNTELIYGAGTMKEEQVMGGENNYSLASH
HIFGITGSSQMLFSPLERAQNGMSIYLTDKEKIYEYIIKDVFTVAPERVDVIDDTAGLKEVT
LVTCTDIEATERIIVKGELKTEYDFDKAPADVLKAFNHSYNQVST
111111110000000000000000000000000000000000000000000000000000
000000000000000000000000000000000000000000000000000000000000
000000000000000000000000000000000000000000000
>DM_train1809
MGFEDGFYTILHLAEGQHPNSKIPGGMYASSKDGKDVPVTAEPLGPQSKIRWWIARDPQAGD
DMYTITEFRIDNSIPGQWSRSPVETEVPVYLYDRIKAEETGYTCAWRIQPADHGADGVYHIV
GNVRIGSTDWADLREEYGEPQVYMKPVPVIPNVYIPRWFILGYEE
100000000000000000000000000000000000000000000000000000000000
000000000000000000000000000000000000000000000000000000000000
000000000000000000000000000000000000000000000
>DM_train1810
KNASVITVGNEILKGRTVNTNAAFIGNFLTYHGYQVRRGFVVMDDLDEIGWAFRVALEVSDL
VVSSGGLGPTFDDMTVEGFAKCIGQDLRIDEDALAMIKKKYGQADLTPQRLKMAKIPPSCRP
IENPVGTAPGLICAVGGKKVIILPGVPKEMEALLKAMEKDIIIPD

000000000000000000000000000000000000000000000000000000000000
000000000000000000000000000000000000000001110000000000000000
00000000000000000000000000000000000000000011
>DM_train1811
MARYINITLEKRGVTCKALLLDDVAPRTSKAVWDALPQSSQVFHGKYARNEIYNLVPAFAPK
EPGAENTTVTPIPGDVCYFTFTSNDLKTPSHGYEADSGTDEVQTIVDLAVFYGRNNLLLNGD
TGWVPGNVFATIVEGLDEMAAACQDIWMGGARDETLTFSRAEDTA
000000000000000000000000000000000000000000000000000000000000
000000000000000000000000001111111111111000000000000000000000
00000000000000000000000000000000000000000111
>DM_train1812
MGSDKIHHHHHHMIQSYPVERSRTIQTRLVLPPDTNHLGTIFGGKVLAYIDEIAALTAMKHA
NSAVVTASIDSVDFKSSATVGDALELEGFVTHTGRTSMEVYVRVHSNNLLTGERTLTTESFL
TMVAVDESGKPKPVPQVEPQTEEEKRLYETAPARKENRKKRAALR
111111111111110000000000000000000000000000000000000000000000
000000000000000000000000000000000000000000000000000000000000
00000000000000000000000000000000000000001111
>DM_train1813
AKITKVQVGEALVGDGNEVAHIDLIIGPRGSPAETAFCNGLVNNKHGFTSLLAVIAPNLPCK
PNTLMFNKVTINDARQAVQMFGPAQHGVAMAVQDAVAEGIIPADEADDLYVLVGVFIHWEAA
DDAKIQKYNYEATKLSIQRAVNGEPKASVVTEQRKSATHPFAANA
000000000000000000000000000000000000000000000000000000000000
000000000000000000000000000000000000000000000000000000000000
00000000000000000000000000000000000000000001
>DM_train1814
MGSDKIHHHHHHMIRPEYLRVLRKIYDRLKNEKVNWVVTGSLSFALQGVPVEVHDIDIQTDE
EGAYEIERIFSEFVSKKVRFSSTEKICSHFGELIIDGIKVEIMGDIRKRLEDGTWEDPVDLN
KYKRFVETHGMKIPVLSLEYEYQAYLKLGRVEKAETLRKWLNERK
111111111111000000000000000000000000000000000000000000000000
000000000000000000000000000000000000000000000000000000000000
00000000000000000000000000000000000000000000
>DM_train1815
GPAQLVGRQTLATPAMGDIQIGMEDKKGQLEVEVIRARSLTQKPGSKSTPAPYVKVYLLENG
ACIAKKKTRIARKTLDPLYQQSLVFDESPQGKVLQVIVWGDYGRMDHKCFMGVAQILLEELD
LSSMVIGWYKLFPPSSLVDPTLAPLTRRASQSSLESSSGPPCIRS
000000000111110000000000000000000000000001110000000000000000
000000000000000000000000000000000000000000000000000000000000
00000000000000000000000111111111111111111111
>DM_train1816
ASVRIREAKEGDCGDILRLIRELAEFEKLSDQVKISEEALRADGFGDNPFYHCLVAEILPAP
GKLLGPCVVGYGIYYFIYSTWKGRTIYLEDIYVMPEYRGQGIGSKIIKKVAEVALDKGCSQF
RLAVLDWNQRAMDLYKALGAQDLTEAEGWHFFCFQGEATRKLAGK
100000000000000000000000000011110000000000000000000000000111
111100000000000000000000000000000000000000000000000000000000

00000000000000000000000000000000000000000001
>DM_train1817
GLFDKLKSLVSDDKKDTGTIEIIAPLSGEIVNIEDVPDVVFAEKIVGDGIAIKPTGNKMVAP
VDGTIGKIFETNHAFSIESDSGVELFVHFGIDTVELKGEGFKRIAEEGQRVKVGDTVIEFDL
PLLEEKAKSTLTPVVISNMDEIKELIKLSGSVTVGETPVIRIKK
11111111111111111000000000000000000000000000000000000000000000
00000000000000000000000000000000000000000000000000000000000000
00000000000000000000000000000000000000000000
>DM_train1818
MKVIEFHVVGNSLNQKPNKKILMWLVGLQNVFSHQLPRMPKEYITRLVFDPKHKTLALIKDG
RVIGGICFRMFPSQGFTEIVFCAVTSNEQVKGYGTHLMNHLKEYHIKHDILNFLTYADEYAI
GYFKKQGFSKEIKIPKTKYVGYIKDYEGATLMGCELNPRIPYTE
00000000000000000000000000000000000000000000000000000000000000
00000000000000000000000000000000000000000000000000000000000000
00000000000000000000000000000000000000111111
>DM_train1819
MLRSSNDVTQQGSRPKTKLGGSSMGIIRTCRLGPDQVKSMRAALDLFGREFGDVATYSQHQP
DSDYLGNLLRSKTFIALAAFDQEAVVGALAAYVLPKFEQPRSEIYIYDLAVSGEHRRQGIAT
ALINLLKHEANALGAYVIYVQADYGDDPAVALYTKLGIREEVMH
11111111111111111111111100000000000000000000000000000000000000
00000000000000000000000000000000000000000000000000000000000000
00000000000000000000000000000000000011111111
>DM_train1820
LLKLEKELLKQIKLDPSPQTIYKWIKDNISPKLHVDKGFVNILMTSFLQYISSEVNPPSDET
DSSSAPSKEQLEQEKQLLLSFKPVMQKFLHDHVDLQVSALYALQVHCYNSNFPKGMLLRFFV
HFYDMEIIEEEAFLAWKEDITQEFPGKGKALFQVNQWLTWLETA
00000000000000000000000000000000000000000000000000000000011111
11110000000000000000000000000000000000000000000000000000000000
00000000000000000000000000000000000000000011
>DM_train1822
MFIKAERLLIRKFEFKDWEAVHEYTSDSDVMKYIPEGVFTEEDTRNFVNKNMGENAKNFPVI
LIGENILVGHIVFHKYFGEHTYEIGWVFNPKYFNKGYASEAAQATLKYGFKEMKLHRIIATC
QPENTPSYRVMEKIGMRREGYFKKCIPHGNEWWDEYYYAILEEE
00000000000000000000000000000000000000000000000000000110000000
00000000000000000000000000000000000000000000000000000000000000
00000000000000000000000000000000000000000000
>DM_train1823
MIDEPTGLYNRLRLQEDVSLRLQRDGALTVIAADLLPLALLNTIIRTLGYPFSNDLMLEARD
RIRAELPDFTLYKISPTRFGLLLPRQQQEETESVCLRLLRAFESPVVCRGIPIKANVGLGVL
PLADDTLDGDQDWLRLVVSAADDARDRGVGWARYNPPLDQAQQR
11111111000000000000000000000000000000000000000000000000000000
00000000000000000000000000000000000000000000000000000000000000
00000000000000000000000000000000000000000001
>DM_train1824

MVRYPLEPVLSIKKDRVDRAEKVVKEKRRLLELEQEKLRERESERDKVKNHYMQKIRQLREQ
LDDGTTSDAILKMKAYIKVVAIQLSEEEEKVNKQKENVLAASKELERAEVELTKRRKEEEKT
RLHKEEWMKEALKEEARQEEKEQDEMGQLLHQLHKQKQRESGEN
00000000000000000000000000000000000000000000000000000000000000
00000000000000000000000000000000000000000000000000000000000000
00000000000000000000000000000000000000111111
>DM_train1825
MSLVEKQSALESKARSWLIERGVEIDDIAELVLFLQQKYHPGLELDICRQNVEHVLRKREVQ
NAVLTGIQLDVMAEKGELVQPLQNIISADEGLYGVDEILALSIVNVYGSIGFTNYGYIDKVK
PGILAKLNEHDGIAVHTFLDDIVGAIAAAAASRLAHSYHDDIVN
11111000000000000000000000000000000000000000000000000000000000
00000000000000000000000000000000000000000000000000000000000000
00000000000000000000000000000000000000001111
>DM_train1826
MALLWQLARNAEGDDPTNLESALIGKPVPKFRLESLDNPGQFYQADVLTQGKPVLLNVWATW
CPTCRAEHQYLNQLSAQGIRVVGMNYKDDRQKAISWLKELGNPYALSLFDGDGMLGLDLGVY
GAPETFLIDGNGIIRYRHAGDLNPRVWEEEIKPLWEKYSKEAAQ
11111111111111111000000000000000000000000000000000000000000000
00000000000000000000000000000000000000000000000000000000000000
00000000000000000000000000000000000000000001
>DM_train1827
MSLAEVGSKSVLFVCLGNICRSPIAEAVFRKLVTDEKVSDNWAIDSSAVSDWNVGRPPDPRA
VSCLRNHGISTAHKARQITKEDFATFDYILCMDESNLRDLNRKSNQVKNCKAKIELLGSYDP
QKQLIIEDPYYGNDSDFEVVYQQCLRCCKAFLEKTYEGHHHHHH
11111110000000000000000000000000000000000000000000000000000000
00000000000000000000000000000000000000000000000000000000000000
00000000000000000000000000000000000011111111
>DM_train1828
DIEQKQAEIIDQLVKRASTCKSEALGPLIIEATSHPSLFAFSEILALPNVAQLEGTTDSVYL
DLLRLFAHGTWGDYKCNATRLPHLSPDQILKLKQLTVLTLAESNKVLPYDTLMVELDVSNVR
ELEDFLINECMYAGIVRGKLDQLKRCFEVPFAAGRDLRPGQLGN
11000000000000000000000000000000000000000000000000000000000000
00000000000000000000000000000000000000000000000000000000000000
00000000000000000000000000000000000000011111
>DM_train1829
MSNMQLDTLRRIVQEINSSVSLHDSLDIMVNQVADAMKVDVCSIYLLDERNQRYLLMASKGL
NPESVGHVSLQLSEGLVGLVGQREEIVNLENASKHERFAYLPETGEEIYNSFLGVPVMYRRK
VMGVLVVQNKQPQDFSEAAESFLVTLCAQLSGVIAHAHAVGNID
00000000000000000000000000000000000000000000000000000000000000
00000000000000000000000000000000000000111111111000000000000000
00000000000000000000000000000000000000000001
>DM_train1830
MELTLYNGEKKTFYSRPNNHDNAWLNAILQLFRYVEEPFFDWVYSSPENLTLEAIKQLEDLT
GLELHEGGPPALVIWNIKHLLHTGIGTASRPSEVCVVDGTDMSLADFHAGIFLKGQEHAVFA

CVTSNGWYAIDDEDFYPWTPDPSDVLVFVPYDQEPLNGEWKAK
00000000000000000000000000000000000000000000000000000000000000
00000000000000000000000000000000000000000000000000000000000000
0000000000000000000000000000000011111111111
>DM_train1831
KPGAMYRNSSKPAYTRREYISGIPGKKIAQFDMGNNGAGPTYPAQVELVVEKPVQIRHNALE
AARVAANRYVQNSGAAANYKFRIRKFPFHVIRENKAAAAAAAAAAADGMRAPFGKPVGTAAR
VHGANHIFIAWVNPDPNVEEAWRRAKMKVTPTINIDSSPAGNA
00000000000000000000000000000000000000000000000000000000000000
00000000000000000000000000000000000111111111110000000000000000
0000000000000000000000000000000000000000000
>DM_train1833
MASYAELFDIGEDFAAFVGHGLATEQGAVARFRQKLESNGLPSALTERLQRIERRYRLLVAG
EMWCPDCQINLAALDFAQRLQPNIELAIISKGRAEDDLRQRLALERIAIPLVLVLDEEFNLL
GRFVERPQAVLDGGPQALAAYKAGDYLEHAIGDVLAIIEGAAR
00000000000000000000000000000000000000000000000000000000000000
00000000000000000000000000000000000000000000000000000000000000
0000000000000000000000000000000000000000001
>DM_train1834
SNAMTFNQEQDYWAGYKANERALIIQTWSGFGRYAPDHLYPPHILPLDTDNETLGTTVLQAL
ANSRTFVYDSPEDQDFFDTEKIRQRYEDWVAKLCGNLGYKTRRALFKNMMSVDIWLHNGCLK
ISPSRHVKLEAWDAIDADDVILSLDNSPEEIGAGLKLALSRCR
11100000000000000000000000000000000000000000000000000000000000
00000000000000000000000000000000000000000000000000000000000000
0000000000000000000000000000000000000000000
>DM_train1835
MMFLRQEDFATVVRSTPLVSLDFIVENSRGEFLLGKRTNRPAQGYWFVPGGRVQKDETLEAA
FERLTMAELGLRLPITAGQFYGVWQHFYDDNFSGTDFTTHYVVLGFRFRVAEEELLLPDEQH
DDYRWLTPDALLASENVHANSRVYFNNDPRAIIGLNKKEVKNV
00000000000000000000000000000000000000000000000000000000000000
00000000000000000000000000000000000000000000000000000000000000
0000000000000000000000000111111111111111111
>DM_train1836
GSHGMADEEKLPPGWEKAMSRSSGRVYYFNHITNASQWERPSGNSSSGGKNGQGEPARVRCS
HLLVKHSQSRRPSSWRQEKITRTKEEALELINGYIQKIKSGEEDFESLASQFSDCSSAKARG
DLGAFSRGQMQKPFEDASFALRTGEMSGPVFTDSGIHIILRTE
11111111110000000000000000000000000000000011111111111100000000
00000000000000000000000000000000000000000000000000000000000000
0000000000000000000000000000000000000000000
>DM_train1837
MGMIGYFAEIDSEKINQLLESTEKPLMDNIHDTLSGLRRLDIDKRWDFLHFGLTGTSAFDPA
KNDPLSRAVLGEHSLEDGIDGFLGLTWNQELAATIDRLESLDRNELRKQFSIKRLNEMEIYP
GVTFSEELEGQLFASIMLDMEKLISAYRRMLRQGNHALTVIVG
11000000000000000000111111111000000000000000000000000000000000

000000000000000111111000000000000000000000000000000000000000000
0000000000000000000000000000000000000000001
>DM_train1838
MKGLTEKLMILSHQSIKNLLGKVILNYSEENVRENGYDLRICGDKYYELVQGAELPEKKATL
REIEFKERAILSANHTYLFESCEEFNMPADLAVLITLKSTLARNGFLAPPTVIDAGYKGKVN
VAITAVYNSSLKKGMATHHLIFLKLDKPTERLYNGKYQGGILI
111111111000000000000000000000000000000000000000000000000000000
000000000000000000000000000000000000000000000000000000000000000
0000000000000000000000000000000000000000000
>DM_train1839
MADEATRRVVSEIPVLKTNAGPRDRELWVQRLKEEYQSLIRYVENNKNADNDWFRLESNKEG
TRWFGKCWYIHDLLKYEFDIEFDIPITYPTTAPEIAVPELDGKTAKMYRGGKICLTDHFKPL
WARNVPKFGLAHLMALGLGPWLAVEIPDLIQKGVIQHKEKCNQ
000000000000000000000000000000000000000000000000000000000000000
000000000000000000000000000000000000000000000000000000000000000
0000000000000000000000000000000000000000001
>DM_train1842
GAMDPREVILCKDQDGKIGLRLKSIDNGIFVQLVQANSPASLVGLRFGDQVLQINGENCAGW
SSDKAHKVLKQAFGEKITMTIRDRPFERTITMHKDSTGHVGFIFKNGKITSIVKDSSAARNG
LLTEHNICEINGQNVIGLKDSQIADILSTSGTVVTITIMPAF
111000000000000000000000000000000000000000000000000000000000000
000000000000000000000000000000000000000000000000000000000000000
000000000000000000000000000000000000000000
>DM_train1843
FSDETLDKVPKSEGYCSRILRAQGTRREGYTEFSLRVEGDPDFYKPGTSYRVTLSAAPPSYF
RGFTLIALRENREGDKEEDHAGTFQIIDEEETQFMSNCPVAVTESTPRRRTRIQVFWIAPPA
GTGCVILKASIVQKRIIYFQDEGSLTKKLCEQDSTFDGVTDK
111111111111100000000000000000000000000000000000000000000000000
000000000011111100000000000000000000000000000000000000000000000
000000000000000000000000000000000111111111
>DM_train1844
MATASHNIDDILQLKDDTGVITVTADNYPLLSRGVPGYFNILYITMRGTNSNGMSCQLCHDF
EKTYHAVADVIRSQAPQSLNLFFTVDVNEVPQLVKDLKLQNVPHLVVYPPAESNKQSQFEWK
TSPFYQYSLVPENAENTLQFGDFLAKILNISITVPQAFNVQE
111111100000000000000000000000000000000000000000000000000000000
000000000000000000000000000000000000000000000000000000000000000
000000000000000000000000000000000000000111
>DM_train1845
MAMSRKRVRAEDIHYWLLKSEPHKFSIDDLAKQKTSPWDGVRNYAARNNMRAMSVGDKVLFY
HSNTKEPGVAGLAEVVRLAYDDFTALDKTSEYFDPKATKEKNPWKMVDVKFVARWDTVLTLH
ELKSRRELQKMALFTQRRLSVQPVSASEYAYILRMNEEQQRK
111111110000000000000000000000000000000000000000000000000000000
000000000000000000000000000000000000000000000000000000000000000
000000000000000000000000000000000000000001

>DM_train1846
MSKNIGLNAIEMSYLRQSLSLSAAQVGQLTNHSEAEVLAWENAETQAPELAQKKLLDIDDII
EMQVLNTTDGIEALFKKEPKRHLAFVVYPTQAIYTQYNPEFLSSLPLTELYNTAAWRIKKEC
KLVLEVDVSLINLNVEAYKAYREQNGLSESRESRAKWAATQL
111110000000000000000000000000000000000000000000000000000000
000000000000000000000000000000000000000000000000000000000000
000000000000000000000000000000000000000000
>DM_train1847
MSGELRVRRASSWELDLILKEAEKYGELLHEFFCVVEGKYRDVYAVNEEVWKIIEDINMRPY
SLGTFVGTIRVDENLVEKFYPNLEFFSLIKLEKNYVILGPKASFLFTTGKDAPKEAVREIKW
QGSKRVVVLNDLGDIIGIGLINPKSDRRFIKNLKDVGEFLRR
111100000000000000000000000000000000000000000000000000000000
000000000000000000000000000000000000000000000000000000000000
000000000000000000000000000000000001111111
>DM_train1848
MAQEDVTAVATNGAGPVETHLVFTEFKQMLLVEAQKVGDAVTFYKSAFGAIESGHSLYPKRK
LDQELPHVLSSELNLAGSSFVVCDVSSLPGFSTAKSEGSGVTFLLGTKDAEAAVAKAVDAGA
VKVEVTEAEVELGFKGKVTDPFGVTWIFAEKKTVITDENKEV
111111111111111111000000000000000000000000000000000111111111
111111000000000000000000000000000000000000000000000000000000
000000000000000000000000000001111111111111
>DM_train1849
TPEKEPLKPGDILVYAQGGGEPKPIRLEELKPGDPFVLAYPMDPKTKVVKSGEAKNTLLVAR
FDPEELAPEVAQHAAEGVVAYSAVCTHLGCIVSQWVADEEAALCPCHGGVYDLRHGAQVIAG
PPPRPVPQLPVRVEDGVLVAAGEFLGPVGVQASAGAYTWRV
000000000000000000000000000000000000000000000000000000000000
000000000000000000000000000000000000000000000000000000000000
00000000000000000000000000000000111111111
>DM_train1850
CDLPQTHSLGSRRTLMLLAQMRRISLFSCLKDRHDFGFPQEEFGNQFQKAETIPVLHEMIQQ
IFNLFSTKDSSAAWDETLLDKFYTELYQQLNDLEACVIQGVGVTETPLMNEDSILAVRKYFQ
RITLYLKEKKYSPCAWEVVRAEIMRSFSLSTNLQESLRSKE
111111110000000000000000000000000000000001111111100000000000
000000000000000000000000000000111111111111111110000000000000
00000000000000000000000000000000111111111
>DM_train1852
MNAEINPLHAYFKLPNTVSLVAGSSEGETPLNAFDGALLNAGIGNVALIRISSIMPPEAEIV
PLPKLPMGALVPTAYGYIISDVPGETISAAISVAIPKDKSLCGLIMEYEGKCSKKEAEKTVR
EMAKIGFEMRGWELDRIESIAVEHTVEKLGCAFAAAALWYK
111111111111100000000000000000000000000000000000000000000000
000000000000000000000000000000000000000000000000000000000000
00000000000000000000000000000000000000000
>DM_train1853
MPIGHSREVLVRLRDILALLADGCKTTSLIQQRLGLSHGRAKALIYVLEKEGRVTRVAFGNV

ALVCLSMDQYRQLVDGMIREVERLVTTNKLKFISPPRLHDLIIKDPQARKFFSSIIPIAHRT
AIILSFLNHLLKMIYGEPYVKTDETVYLTANRKVHIEPPAK
111100000000000000000000000000000000000000000000000000000000
000000000000000000000000000000000000000000000000000000000000
00000000000000000000000000000000011111111
>DM_train1854
FELPSVPFPAPGSDEILFVVRDTTFNTNAPVNVEVSDFWTNRNVKRKPYKDVYGQSVFTTSG
TKWLTSYMTVNINDKDYTMAAVSGYKHGHSAVFVKSDQVQLQHSYDSVASFVGEDEDSIPSK
MYLDETPEYFVNVEAYESGSGNILVMCISNKESFFECKHQQ
111111111110000000000000000000000000000000000000000000000000
000000000000000000000000000000000000000000000000000000000000
00000000000000000000000000000000000000000
>DM_train1855
MSLLTEVETYVLSIIPSGPLKAEIAQRLEDVFAGKNTDLEVLMEWLKTRPILSPLTKGILGF
VFTLTVPSERGLQRRRFVQNALNGNGDPNNMDKAVKLYRKLKREITFHGAKEISLSYSAGAL
ASCMGLIYNRMGAVTTEVAFGLVCATCEQIADSQHRSHRQ
000000000000000000000000000000000000000000000000000000000000
000000000000000000000000000000000000000000000000000000000000
0000000000000000000000000000000000111111
>DM_train1856
MRRFLALLLVLTLWLGFTPLASADVAGLVPCKDSPAFQKRAAAAVNTTADPASGQKRFERYS
QALCGEDGLPHLVVDGRLSRAGDFLIPSVLFLYIAGWIGWVGRAYLIAVRNSGEANEKEIII
DVPLAIKCMLTGFAWPLAALKELASGELTAKDNEITVSPR
111111111111111111111110000000000000000000000000000000000000
000000000000000000000000000000000000000000000000000000000000
0000000000000000000000000000000000000000
>DM_train1857
LGSMESTQQMAVSIINSSFEAAVVAATSALENMGIEYDYQDIYSRVKNKFDFVMDDSGVKNN
PIGKAITIDQALNNKFGSAIRNRNWLADTSRPAKLDEDVNKLRMMLSSKGIDQKMRVLNACF
SVKRIPGKSSSIIKCTKLMRDKLERGEVEVDDSFVDEKME
000000000000000000000000000000000000000000000000000000000000
000000000000011111111111111000000000000000000000000000000000
0000000000000000000000000000011111111111
>DM_train1858
GPGRGFGKRRHPKKLTPLAYKQFIPNVAEKTLGASGRYEGKITRNSERFKELTPNYNPDIIF
KDEENTGADRLMTQRCKDKLNALAISVMNQWPGVKLRVTEGWDEDGHHSEESLHYEGRAVDI
TTSDRDRSKYGMLARLAVEAGFDWVYYESKAHIHCSVKAE
111111111111111000000000000000000000000000000000000000000000
000000000000000000000000000000000000000000000000000000000000
0000000000000000000000000000000000000001
>DM_train1859
GMKTIDLADIQAFLYRESRLLDDKAWDAWLDCYRADAVFWMPSWDDADALVTDPQREISLIY
YPNRQGLEDRVFRIKTERSSATVPDTRTSHNIANVERESADGDVHTVRFNWHTLSYRYKTVS
SYFGMSRYAIDFSGDAPKIVSKYVVLKNDYINQLIDIYHI

10000000000000000000000000000000000000000000001111111111100000
00000000000000000000000000000000000000000000000000000000000000
0000000000000000000000000000001110000000
>DM_train1860
MELYLIRHGIAEAQKTGIKDEERELTQEGKQKTEKVAYRLVKLGRQFDLIVTSPLIRARQTA
EILLASGLSCQLEESNHLAPNGNIFNWLDYWLKPKNFPENAQIAIVGHEPCLSNWTEILLWG
EAKDSLVLKKAGMIGLKLPEIGSPVGRSQMFWLTPPRYLL
00000000000000111100000000000000000000000000000000000000000000
00000000000000000000000000000000000000000000000000000000000000
0000000000000000000000000000000000000000
>DM_train1861
MGSDKIHHHHHHMRKAWVKTLALDRVSNTPVVILGIEGTNRVLPIWIGACEGHALALAMEKM
EFPRPLTHDLLLSVLESLEARVDKVIIHSLKDNTFYATLVIRDLTYTDEEDEEAALIDIDSR
PSDAIILAVKTGAPIFVSDNLVEKHSIELEVNERDLINSR
11111111111000000000000000000000000000000000000000000000000000
00000000000000000000000000000000000000000000001111100000000000
0000000000000000000000000000011111111111
>DM_train1862
MARRILSVLLENESGALSRVIGLFSQRGYNIESLTVAPTDDPTLSRMTIQTVGDEKVLEQIE
KQLHKLVDVLRVSELGQGAHVEREIMLVKIQASGYGRDEVKRNTEIFRGQIIDVTPSLYTVQ
LAGTSGKLDAFLASIRDVAKIVEVARSGVVGLSRGDKIMR
10000000000000000000000000000000000000000000000000000000000000
00000000000000000000000000000000000000000000000000000000000000
0000000000000000000000000000000000011111
>DM_train1863
GSHMSPKIVDAGTATVPHTMSTMKTADNRPASVYLKKDKPTLIKFWASWCPLCLSELGQAEK
WAQDAKFSSANLITVASPGFLHEKKDGEFQKWYAGLNYPKLPVVTDNGGTIAQNLNISVYPS
WALIGKDGDVQRIVKGSINEAQALALIRNPNADLGSLKHS
11111111111110000000000000000000000000000000000000000000000000
00000000000000000000000000000000000000000000000000000000000000
0000000000000000000000000000000000000000
>DM_train1864
GILAEAHGTRPDLTDQPLPDADHTWYTDGSSLLQEGQRKAGAAVTTETEVIWAKALPAGTSA
QRAELIALTQALKMAEGKKLNVYTDSRYAFATAHLTSEGKEIKNKDEILALLKALFLPKRLS
IIHCPGHQKGHSAEARGNRMADQAARKAAITETPDTSTLL
11111100000000000000000000000000000000000000000000000000000000
00000000000000000000000000000000000000000000000000000000000000
0000111111000000000000000000000000000111
>DM_train1865
GEFETLEQLKESLKKEGKEIYDVEMKESMREQLLEKLPEIVEIEISDRTLEILVNEAINRLK
REGRYEQIVSSYESEEKFREELKERILDDIKRDRVIEVLAQEKGISVNDEELEKEAEELAPF
WGISPDRAKSLVKARQDLREELRWAILKRKVLDLLLQEVE
10000000000000000000000000000000000000000000000000000000000000
00000000000000000000000000000000000000000000000000000000000000

0000000000000000000000000000000000000000
>DM_train1866
PRILGRNIGSHHVRVEHFMNHSITTLAKDTPLEEVVKVVTSTDVTEYPLVESTESQILVGIV
QRAQLVQALQAEPPSRAPGHQQCLQDILARGCPTEPVTLTLFSETTLHQAQNLFKLLNLQSL
FVTSRGRAVGCVSWVEMKKAISNLTNPPAPKEFLEVLFQG
111111111000000000000000000000000000000000000000000000000000
000000000001111111000000000000000000000000000000000000000000
0000000000000000000000000000000000000111
>DM_train1867
MFHILRLESTVDLSEPLKDNGIIVFQSDKLDLEPSPNLGPTGIDNTNVNLINAKGDVLLHIG
IRRRENAFVFNSIPYGESRGPEERIPLEGTFGDRRDPSITIFDHPDRYQIMIDYKTVYYYKK
RLEGRCEKVSYKINEGQTPPFSDVLGVTVLYFANVMPRAN
000000000000000000000000000000000000000000000000000000000000
000000000000000000000000000000000000000000000000000000000000
0000000000000000000000000000000000111111
>DM_train1868
LPLDERAFEKTLTPIIQEYFEHGDTNEVAEMLRDLNLGEMKSGVPVLAVSLALEGKASHREM
TSKLLSDLCGTVMSTTDVEKSFDKLLKDLPELALDTPRAPQLVGQFIARAVGDGILCNTYID
SYKGTVDCVQARAALDKATVLLSMSKGGKRKDSVWGSGGG
000000000000000000000000000000000000000000000000000000000000
000000000000000000000000000000000000000000000000000000000000
0000000000000000000000001111111111111111
>DM_train1869
MGGSHHHHHHGMASSSDGHNGLGKGFGDHIHWRTLEDGKKEAAASGLPLMVIIHKSWCGACK
ALKPKFAESTEISELSHNFVMVNLEDEEEPKDEDFSPDGGYIPRILFLDPSGKVHPEIINEN
GNPSYKYFYVSAEQVVQGMKEAQERLTGDAFRKKHLEDEL
111111111111111111111000000000000000000000000000000000000000
000000000000000000000000000000000000000000000000000000000000
0000000000000000000000000000000011111111
>DM_train1870
MTVNLEEKRPEIVGPEKVQSPYPIRFEGKVVHGFGRGSKELGIPTANISEDAIQELLRYRDS
GVYFGYAMVQKRVFPMVMSVGWNPYYKNKLRSAEVHLIERQGEDFYEEIMRVIVLGYIRPEL
NYAGLDKLIEDIHTDIRVALNSMDRPSYSSYKKDPFFKV
111111111000000000000000000000000000000000000000000000000000
000000000000000000001111111111100000000000000000000000000000
000000000000000000000000000000000000000
>DM_train1871
GSREEPSDLEELEQFAKTFKQRRIKLGFTQGDVGLAMGKLYGNDFSQTTISRFEALNLSFKN
MCKLKPLLEKWLNDAENLSSDSSLSSPSALNSPGIEGLSRRRKKRTSIETNIRVALEKSFLE
NQKPTSEEITMIADQLNMEKEVIRVWFCNRRQKEKRINP
111111100000000000000000000000000000000000000000000000000000
000000000000000011111111111111111111111100000000000000000000
000000000000000000000000000000000000000
>DM_train1872

MTNNDTTLQLSSVLNRECTRSRVHCQSKKRALEIISELAAKQLSLPPQVVFEAILTREKMGS
TGIGNGIAIPHGKLEEDTLRAVGVFVQLETPIAFDAIDNQPVDLLFALLVPADQTKTHLHTL
SLVAKRLADKTICRRLRAAQSDEELYQIITDTEGTPDEA
111111100000000000000000000000000000000000000000000000000000
000000000000000000000000000000000000000000000000000000000000
000000000000000000000000000000000111111
>DM_train1873
GMSVAAEVAQVAQSAIDDFNAAYGLCLDDDRLEQWPTLFVDDCLYQVIARENVDNGLPAAVM
YCDSKGMLADRVVALRKANVFPEHFNRHLIGRAVITGVEGDQVSAEASYVVFQTRNDGETRI
YNAGKYVDRFDLSGGTVRLKSRTCIYDTLRIATLLATPI
111111000000000000000000000000000000000000000000000000000000
000000000000000000011110000000000000000000000000000000000000
000000000000000000000000000000000000000
>DM_train1874
GMLDNAIPQGFEDAVELRRKNRETVVKYMNTKGQDRLRRHELFVEDGCGGLWTTDTGSPIVI
RGKDKLAEHAVWSLKCFPDWEWYNIKVFETDDPNHFWVECDGHGKILFPGYPEGYYENHFLH
SFELDDGKIKRNREFMNVFQQLRALSIPVPQIKREGIPT
111111111000000000000000000000000000000000000000000000000000
000000000000000000000000000000000000000000000000000000000000
000000000000000000000000000000000000000
>DM_train1875
QSRERLVKWLQDAYAMEKEAETMMAAMASRIEHYPELKRRIEQHVEETQQQSAGVQRCLELL
NGSIPTAKGMLSSVLASMHAAGNSMMTDEVTKGVGISYAFEHLEIASYRALVVAARSAGEQE
VAQICEDILQQEIEMAEWLIEHQEAIVVAFLEREQLEGV
000000000000000000000000000000000000000000000000000000000000
000000011111111111111111000000000000000000000000000000000000
000000000000000000000000000000000000111
>DM_train1876
GHMNSKNTICLWYDSAALEAATFYAETFPDSAVLAVHRAPGDYPSGKEGDVLTVEFRVMGIP
CLGLNGGPAFRHSEAFSFQVATDDQAETDRLWNAIVDNGGEESACGWCRDKWGISWQITPRV
LSEAIASPDRAAARRAFEAMMTMGRIDIATIEKAFKGGS
111100000000000000000000000000000000001111111110000000000000
000000000000000000000000000000000000000000000000000000000000
000000000000000000000000000000000000111
>DM_train1877
MSLRVRIAVIGKLDGFIKEGIKHYEKFLRRFCKPEVLEIKRVHRGSIEEIVRKETEDLTNRI
LPGSFVMVMDKRGEEVSSEEFADFLKDLEMKGKDITILIGGPYGLNEEIFAKAHRVFSLSKM
TFTHGMTVLIVLEQIFRAFKIIHGENYHYEGGSHHHHHH
110000000000000000000000000000000000000000000000000000000000
000000000000000000000000000000000000000000000000000000000000
000000000000000000000000011111111111111
>DM_train1878
GMTLPPYHTPLPAETLRALSIPAPWTFGLADRVRFGELDAIGHVNHTAYLRWYESFRLPFLK
ARHVTDYGPTSPRLVLKQVHCTYLAEMGMGEDYVITGRVSNFRTTSFTMEFACWRLGDAVEC

TSEGSAVVVLLNRDGSGRYPIPEAGRASFVTEDGVLAAG
111000000000000000000000000000000000000000000000000000000000
000000000000000000000000000000000000000000000000000000000000
000000000000000000000000000000000000001
>DM_train1879
GMAFMEKIFPDILEAIRNEEIIKESKKIPMPYFGLFALVIFDKVKELGSETSLYEIGEEFGK
MLSPKNIEELKKIFKLMNFGDLEIDENKILLKNPPYKIKLSNPPYQWVSKEEPIHDFIAGIL
AGCLEEIFYYYFVVNEVECVSQGKDKCVFEVKEVDELNK
000000000000000000000000000000000000000000011000000000000000
000000000000000000000000000000000000000000000000000000000000
000000000000000000000000000000000001111
>DM_train1880
ATPAAQASDDRYEVTQQRNPDAACLDCHKPDTEGMHGKHASVINPNNKLPVTCTNCHGQPSP
QHREGVKDVMRFNEPMYKVGEQNSVCMSCHLPEQLQKAFWPHDVHVTKVACASCHSLHPQQD
TMQTLSDKGRIKICVDCHSDQRTNPNFNPASVPLLKEQP
111111111111111111000000000000000000000000000000000000000000
000000000000000000000000000000000000000000000000000000000000
000000000000000000000000000000000000111
>DM_train1881
KHHHHHHIKPGALCVIDTPEGKGTGFFSGNDIVTAAHVVGNNTFVNVCYEGLMYEAKVRYMP
EKDIAFITCPGDLHPTARLKLSKNPDYSCVTVMAYVNEDLVVSTAAAMVYGNTLSYAVRTQD
GMSGAPVCDKYCRVLAVHQTNTGYTGGAVIIDPTDFHPV
000000000000000000000000000000000000000000000000000000000000
000000000000000000000000000000000000000000000000000000000000
000000000000000000000000000000000000001
>DM_train1882
MSVMYKKILYPTDFSETAEIALKHVKAFKTLKAEEVILLHVIDEREIKKRDIFSLLLGVAGL
NKSVEEFENELKNKLTEEAKNKMENIKKELEDVGFKVKDIIVVGIPHEEIVKIAEDEGVDII
IMGSHGKTNLKEILLGSVTENVIKKSNKPVLVVKRKNS
000000000000000000000000000000000000000000000001111111111111
111000000000000000000000000000000000000000000000000000000000
00000000000000000000000000000000000000
>DM_train1883
MPETDFEEKMILIRRTARMQAGGRRFRFGALVVVGDRQGRVGLGFGKAPEVPLAVQKAGYYA
RRNMVEVPLQNGTIPHEIEVEFGASKIVLKPAAPGTGVIAGAVPRAILELAGVTDILTKELG
SRNPINIAYATMEALRQLRTKADVERLRKGEAHAQAQG
111100000000000000000000000000000000000000000000000000000000
000000000000000000000000000000000000000000000000000000000000
00000000000000000000000000000011111111
>DM_train1884
LSSPADSYNEGVKLQPQEISPPPTANLDRSNDKVYENVTGLVKAVIEMSSKIQPAPPEEYVP
MVKEVGLALRTLLATVDETIPLLPASTHREIEMAQKLLNSDLGELINKMKLAQQYVMTSLQQ
EYKKQMLTAAHALAVDAKNLLDVIDQARLKMLGQTRPH
111111111111111111111111000000000000000000000000000000000000

0000000000000000000000000000000000000000000000000000000000000000
00000000000000000000000000000000000000
>DM_train1885
MSEQAPAPKRGRRFKEQTPVQRALGLLVHREHSKKELNRKLQARGIEPEAAQAAVERLAGEG
WQDDVRFAASVVRNRASSGYGPLHIRAELGTHGLDSDAVSAAMATFEGDWTENALDLIRRRF
GEDGPVDLAQRRKAADLLARRGFDGNSIRLATRFDLED
1111111111111111000000000000000000000000000000000000000000000000
0000000000000000000000000000000000000000000000000000000000000000
00000000000000000000000000000000001111
>DM_train1886
LIEGSERLSGLTDVDEVIKDLSRLLRKLVKTRWIAVYFFDRERRDFAPARSTGLPASFLPVF
REMPLAPDKIPLLKSMLRKRQHLMLTDPGSSDLLTPKLRKLLRNLCVLAVPMVVRTQVIGAV
FMARTRDNPPFSDAETAIIRDLVSHAALVVSHMQLFDE
1111111100000000000000000000000000000000011100000000000000000000
0000000000000000000000000000000000000000000000000000000000000000
00000000000000000000000000000000000000
>DM_train1887
MDFRIGQGYDVHQLVPGRPLIIGGVTIPYERGLLGHSDADVLLHAITDALFGAAALGDIGRH
FSDTDPRFKGADSRALLRECASRVAQAGFAIRNVDSTIIAQAPKLAPHIDAMRANIAADLDL
PLDRVNVKAKTNEKLGYLGRGEGIEAQAAALVVREAAA
0000000000000001111000000000111100011000000000000000000000000000
0001111000000000000000000000000000000000000000000000000000000000
00000000000000000000000000000000001111
>DM_train1888
MPKSEIRKLLQEIKKQVDNPGNSSTTEIKKMASEAGIDEQTAEEIYHLLTEFYQAVEEHGGI
EKYMHSNISWLKIELELLSACYQIAILEDMKVLDISEMLSLNDLRIFPKTPSQLQNTYYKLK
KELIQVEDIPKNKPGRKRKTQKNTKKEKTNIFGKVVPA
1110000000000000000000000000000000000000000000000000000000000000
0000000000000000000000000000000000000000000000000000000000000000
00000000000000000001111111111111111111
>DM_train1889
EDYSVVNRFESHGGGWGYSAHSVEAIRFSADTDILLGGLGLFGGRGEYTAKIKLFELGPDGG
DHETDGDLLAETDVLAYDCAAREKYAMMFDEPVLLQAGWWYVAWARVSGPSSDCGSHGQASI
TTDDGVIFQFKSSKKSNNGTDVNAGQIPQLLYRLPTSD
0000000000000000000000000000000000000000000000000000000000000000
0000000000000000000000000000000000000000000000000000000000000000
00000000000000000000000000000000000111
>DM_train1890
PRAAGHSEEVLEREASFSLTTIDGEVISLNNVGGDVVILWFMAAWCPSCVYMADLLDRLTEK
YREISVIAIDFWTAEALKALGLNKPGYPPPDTPEMFRKFIANYGDPSWIMVMDDGSLVEKFN
VRSIDYIVIMDKSSNVLYAGTTPSLGELESVIKSVQGG
0000000000000000000000000000000000000000000000000000000000000000
0000000000000000000000000000000000000000000000000000000000000000
00000000000000000000000000000000000111

>DM_train1891
GMGGSNMKLDFSVAVHSILYLDAHRDSKVASRELAQSLHLNPVMIRNILSVLHKHGYLTGTV
GKNGGYQLDLALADMNLGDLYDLTIPPTISYARFITGPSKTDEQADQSPIAANISETLTDLF
TVADRQYRAYYHQFTMADLQADLNHHGTFLQHEQDSES
11111100000000000000000000000000000000000000000000000000000000
00000000000000000000000000000000000000001111000000000000000000
00000000000000000000000000000000000001
>DM_train1892
GMDTRKLLLTAQEISRMKGEHKVHFLNPGAVRVNKSLGDAVGLRHMGIHLIQIEPGKESTEY
HLHHYEEEAVYVLSGKGTLTMENDQYPIAPGDFVGFPCHAAAHSISNDGTETLVCLVIGQRL
DQDVVDYPNQHKRLYRNNGEWNLVDMADIRVLREPTQK
11000000000000000000000000000000000000000000000000000000000000
00000000000000000000000000000000000000000000000000000000000000
00000000000000000000000000000000001111
>DM_train1893
TDVNECLDPTTCISGNCVNTPGSYICDCPPDFELNPTRVGCVDTRSGNCYLDIRPRGDNGDT
ACSNEIGVGVSKASCCCSLGKAWGTPCEMCPAVNTSEYKILCPGGEGFRPNPITVILEDIDE
CQELPGLCQGGKCINTFGSFQCRCPTGYYLNEDTRVCD
00000000000000000000000000000000000000000000000000000111111111
10000000000000000000000000000000000000000000000000000000000000
00000000000000000000000000000000000000
>DM_train1894
ASDEEIQDVSGTWYLKAMTVDREFPEMNLESVTPMTLTTLEGGNLEAKVTMLISGRCQEVKA
VLEKTDEPGKYTADGGKHVAYIIRSHVKDHYIFYSEGELHGKPVRGVKLVGRDPKNNLEALE
DFEKAAGARGLSTESILIPRQSETCSPGSAWSHPQFEK
11111110000000000000111111100000000000000000000000011000000000
00000000000000000000000000000000000000110000000000000000000000
00000000000000000000001111111111111111
>DM_train1895
MSFGGKSMSEHSAIVTWKRKDSEAFTDNQYSRAHTWEFDGGSKILASASPHVVPVPLSVEAN
VDPEEAFVAALSSCHMLVFLSIAAKQRYLVESYTDNAVGILGKNSKGKTSVTKVVLRPQVVF
SGTSKPTLQQLEKMHHLAHENCFIANSVETEVVTEIIA
11111110000000000000000000000000000000000000000000000000000000
00000000000000000000000000000000000000000000000000000000000000
00000000000000000000000000000000000001
>DM_train1896
VKMAETCPIFYDVFFAVANGNELLLDLSLTKVNATEPERTAMKKIQDCYVENGLISRVLDGL
VMTTISSSKDCMGEAVQNTVEDLKLNTLGREICPAVKRDVDLFLTGTPDEYVEQVAQYKALP
VVLENARILKNCVDAKMTEEDKENALSLLDKIYTSPLC
11110000000000000000000000000000000000000000000000000000000000
00000000000111111111111111111110000000000000000000000000000000
00000000000000000000000000000000000000
>DM_train1897
MTSSEPAHGATPKRSPSEGSADNAALCDALAVEHATIYGYGIVSALSPPGVNFLVADALKQH

RHRRDDVIVMLSARGVTAPIAAAGYQLPMQVSSAADAARLAVRMENDGATAWRAVVEHAETA
DDRVFASTALTESAVMATRWNRVLGAWPITAAFPGGDE
11111111111111110000000000000000000000000000000000000000000000
00000000000000000000000000000000000000000000000000000000000000
00000000000000000000000011111111111111
>DM_train1898
GPMPQLTLDKTDIKILQVLQENGRLTNVELSERVALSPSPCLRRLKQLEDAGIVRQYAALLS
PESVNLGLQAFIRVSIRKAKDAREDFAASVRKWPEVLSCFALTGETDYLLQAFFTDMNAFSH
FVLDTLLSHHGVQDAQSSFVLKEIKHTTSLPLNHLLKE
11111100000000000000000000000000000000000000000000000000000000
00000000000000000000000000000000000000000000000000000000000000
00000000000000000000000000000000000011
>DM_train1899
MTDVIKPDYLEYDDLLDRDEMFTILEEYFMYRGLLGLRIKYGRLFNEIKKFDNDAEEQFGTI
EELKQKLRLNSEEGADNFIDYIKVQKQDIVKLTVYDCISMIGLCACVVDVWRNEKLFSRWKY
CLRAIKLFINDHMLDKIKSILQNRLVYVEMSKHHHHHH
11110000000000000000000000000000000000000000000000000000000000
00000000000000000000000000000000000000000000000000000000000000
00000000000000000000000000000011111111
>DM_train1900
MAQLTHINAAGEAHMVDVSAKAETVREARAEAFVTMRSETLAMIIDGRHHKGDVFATARIAG
IQAAKRTWDLIPLCHPLMLSKVEVNLQAEPEHNRVRIETLCRLTGKTGVEMEALTAASVAAL
TIYAMCKAVQKDMVIGPVRLLAKSGGKSGDFKVEADD
11111111111111000000000000000000000000000000000000000000000000
00000000000000000000000000000000000000000000000000000000000000
0000000000000000000000000000000011111
>DM_train1901
GSSSTLTKELIKDAAEKCCTRNRQECCIEIMKFGTPIRCGYDRDPKLPGYVYKCLQNVLFAK
EPKKKINLDDSVCCSVFGNDQEDSGRRCENRCKNLMTSPSIDAATRLDSIKSCSLLDNVLYK
CFEKCRSLRKDGIKIEVLQFEEYCEATFIQKRTFRGV
11111000000000000000000000000000000000000000000000000000000000
00000000000000000000000000000000000000000000000000000000000000
0000000000000000000000001111111111111
>DM_train1902
SVSVDLNVDPSLQIDIPDALSERDKVKFTVHTKTTLPTFQSPEFSVTRQHEDFVWLHDTLTE
TTDYAGLIIPPAPTKPDFDGPREKMQKLGEGEGSMTKEEFAKMKQELEAEYLAVFKKTVSSH
EVFLQRLSSHPVLSKDRNFHVFLEYDQDLSVRRKNTK
00000000000000000000000000000000000000000000000000000000000000
00000000000000000000000000000000000000000000000000000000000000
0000000000000000000000000000000111111
>DM_train1903
MAGTSTNTQTTGTSQTIEVGLWGGPGGNAWDDGSYTGIREINLSHGDAIGAFSVIYDLNGQP
FTGPTHPGNEPSFKTVKITLDFPNEFLVSVSGYTGVLARLATGKDVIRSLTFKTNKKTYGPY
GKEEGTPFSLPIENGLIVGFKGRSGFVVDAIGFHLSL

111111100000000000000000000000000000000000000000000000000000
000000000000000000000000000000000000000000000000000000000000
0000000000000000000000000000000000000
>DM_train1904
DLYFQGGSGMQCEEKLEVFENGFKDEKFNVEVKFYGNDARKVLLAMIYELYLPEYGREYVYP
FECAKEFWNIYLEGEEIQDEEFQLKPIKFTSEQVIKKLQEEIKKIKPPLEIKIEEAKIYKTK
EGYLAVGNYFILDPRGRLFIFNKPSIANKILKYIWKW
000000000000000000000000000000000000000000000000000000000000
000000000000000000011100000000000000000000000000000000000000
0000000000000000000000000000000000000
>DM_train1905
SLDDHTRDPTVKAPDGNPSGWRTDGQWEHETLRRAVVHGVRLYNSGEFHESHDCFEDEWYNY
GRGNTESKFLHGMVQVAAGAYKHFDFEDDDGMRSLFRTSLQYFRGVPNDYYGVDLLDVRTTV
TNALSDPSALHGWQIRLDGEYPTCRPEDIEFAESLEH
111111111111111111111111110000000000000000000000000000000000
000000000000000000000000000000000000000000000000000000000000
0000000000000000000011111111111111111
>DM_train1907
MSSLQSTRERVLGTPRRRYKVLVAKMGLDGHDRGAKVVARALRDAGFEVVYTGLRQTPEQVA
MAAVQEDVDVIGVSILNGAHLHLMKRLMAKLRELGADDIPVVLGGTIPIPDLEPLRSLGIRE
IFLPGTSLGEIIEKVRKLAEEKRMREEAEASEQVGQA
111111111111111000000000000000000000000000000000000000000000
000000000000000000000000000000000000000000000000000000000000
0000000000000000000000000000001111111
>DM_train1908
MEKTDEYLLARFKGDGVKYKAKLIGIDDVPDARGDKMSQDSMMKLKGMAAAGRSQGQHKQRI
WVNISLSGIKIIDEKTGVIEHEHPVNKISFIARDVTDNRAFGYVCGGEGQHQFFAIKTGQQA
EPLVVDLKDLFQVIYNVKKKEEDKKKVEEANKAEEN
000000000000000000000000000000000000000000000000000000000000
000000000000000000000000000000000000000000000000000000000000
000000000000000000000001111111111111
>DM_train1909
MPSTDFADIPAAKKMKIDTCVLRFAKLTENALEPVRGSAKAAGVDLRSAYDVVVPARGKAIV
KTDLQVQVPEGSYGRVAPRSGLAVKNFIDVGAGVVDEDYRGNLGVVLFNHSDVDFEVKHGDR
IAQFICERIFYPQLVMVDKLEDTERGEAGFGSTGVK
111111111111111111000000000000000000000000000000000000000000
000000000000000000011111110000000000000000000000000000000000
000000000000000000001111111111111111
>DM_train1910
APSYRVKRMDIAKNDEECVVNAANPRGLPGDGVCKAVYKKWPESFKNSATPVGTAKTVMCGT
YPVIHAVGPNFSNYSESEGDRELAAAYREVAKEVTRLGVNSVAIPLLSTGVYSGGKDRLTQS
LNHLFTAMDSTDADVVIYCRDKEWEKKISEAIQMRT
100000000000000000000000000000000000000000000000000000000000
000000000000010010000000000000000000000000000000000000000000

000000000001100000000000000011111111
>DM_train1911
MQLSHRPAETGDLETVAGFPQDRDELFYCYPKAIWPFSVAQLAAAIAERRGSTVAVHDGQVL
GFANFYQWQHGDFCALGNMMVAPAARGLGVARYLIGVMENLAREQYKARLMKISCFNANAAG
LLLYTQLGYQPRAIAERHDPDGRRVALIQMDKPLEP
000000000000000000000000000000000000000000000000000000000000
000000000000000000000000000000000000000000000000000000000000
000000000000000000000000000000000001
>DM_train1912
GMASSFLEEVDRLITLSGITFHASGTGTPELIKIYQDALGNEFPETYKLFLEKYGTLTFNGV
SFYGISKRGLSAASIPDVKFATEQARTFGDINKEMIMIKNSGYGSIFSIDTSIIGSEGEPVI
VETNLSFKDNTEKKVVANSFGEFLLEEIELSLTDLG
000000000000000000000000000000000000000000000000000000000000
000000000000000000000000000000000000000000000000000000000000
000000000000000000000000000000000001
>DM_train1913
MSSNAQVRPPLPPFTRESAIEKIRLAEDGWNSRDPERVSLAYTLDTQWRNRAEFAHNREEAK
AFLTRKWAKELDYRLIKELWAFTDNRIAVRYAYEWHDDSGNWFRSYGNENWEFDEQGLMARR
FACINDMPIKAQERKFHWPLGRRPDDHPGLSELGLE
111000000000000000000000000000000000000000000000000000000000
000000000000000000000000000000000000000000000000000000000000
000000000000000000000000000000001111
>DM_train1914
LPGTRVTESIHKVAERVVTLVSMELPDNVRLIRDYDPSLPELAHDPDQIEQVLLNIVRNALQ
ALGPEGGEIILRTRTAFQLTLHGERYRLAARIDVEDNGPGIPPHLQDTLFYPMVSGREGGTG
LGLSIARNLIDQHSGKIEFTSWPGHTEFSVYLPIRK
111100000000000000000000000000000000000000000000000000000000
000000000000000000000000000000000000000111111111111111111110
000000000000000000000000000000000000
>DM_train1916
GMQGIAFQDAYPDDLSHCYGCGRNNEQGHQLKSYWRGEQTIAHFMPKPFHTAIPGFVYGGLI
ASLIDCHGTGSASAAAQRALEQAGEQLDEPPRFVTAALNIDYLAPTPMGVELELVGEIKEVK
PRKVVVEIALSADGKLCARGHMVAVKMPETMAATSA
111111000001111000001111111111000000000000000000000000000000
000000000000000001111111111110000000000000000000000000000001
100000000000111000000000000011111111
>DM_train1917
EPTMYGEILSPNYPQAYPSEVEKSWDIEVPEGYGIHLYFTHLDIELSENCAYDSVQIISGDT
EEGRLCGQRSSNNPHSPIVEEFQVPYNKLQVIFKSDFSNEERFTGFAAYYVATDINECTDFV
DVPCSHFCNNFIGGYFCSCPPEYFLHDDMKNCGVN
000000000000000000000000000000000000000000000000000000000111
000000000000000000000000000000000000000000000000000000000000
00000000000000000000000000000000000
>DM_train1918

GDSAPLNEAMAVLQHHDAVSGTSRQHVANDYARQLSEGWRPCEVLMSNALAHLSGLKEDFAF
CRKLNISICPLTQTAERFQVIVYNPLGRKVDWMVRLPVSKHVYLVKDPGGKIVPSDVVTIPS
SDSQELLFSALVPAVGFSIYSVSQMPNQRPQKSWS
000000000000000000000000000000000000000000000000000000000000
000000000000000000000000000000000000000000000000000000000000
00000000000000000000000000111111111
>DM_train1919
MPPRKELVGNKWFIENYENETESLVIDANKDESIFIGKCSQVLVQIKGKVNAISLSETESCS
VVLDSSISGMDVIKSNKFGIQVNHSLPQISIDKSDGGNIYLSKESLNTEIYTSCSTAINVNL
PIGEDDDYVEFPISEQMKHSFADGKFKSAVFEHAG
000000000000000000000000000000000000000000000000000000000000
000000000000000000000000000000000000000000000000000000000000
00000000000000000000000000000000111
>DM_train1920
SVDNHGLRRCLISTDMHHIEESFQEIKRAIQAKDTFPNVTILSTLETLQIIKPLDVCCVTKN
LLAFYVDRVFKDHQEPNPKILRKISSIANSFLYMQKTLRQCQEQRQCHCRQEATNATRVIHD
NYDQLEVHAAAIKSLGELDVFLAWINKNHEVMSSA
111000000000000000000000000000000000000000000000000000000000
000000000000000000000000000000000000000001110000000000000000
00000000000000000000000000000000000
>DM_train1921
RLGEVFVLDEEEIRIIQTEAEGIGPENVLNASYSKLKKFIESADGFTNNSYGATFSFQPYTS
IDEMTYRHIFTPVLTNKQISSITPDMEITTIPKGRYACIAYNFSPEHYFLNLQKLIKYIADR
QLTVVSDVYELIIPIHYSPKKQEEYRVEMKIRILE
000000000000000000000000000000001111111111111100000000000000
000000000000001110000000000000000000000000000000000000000000
00000000000000011111110000000000000
>DM_train1923
MGRGKVKPNRKSTGDNSNVVTMIRAGSYPKVNPTPTWVRAIPFEVSVQSGIAFKVPVGSLFS
ANFRTDSFTSVTVMSVRAWTQLTPPVNEYSFVRLKPLFKTGDSTEEFEGRASNINTRASVGY
RIPTNLRQNTVAADNVCEVRSNCRQVALVISCCFN
111111111111000000000000000000000000000000000000000000000000
000000000000000000000000000000000000000000000000000000000000
00000000000000000000000000000000000
>DM_train1924
MMESVNTSFLSPSLVTIRDFDNGQFAVLRIGRTGFPADKGDIDLCLDKMKGVRDAQQSIGDD
TEFGFKGPHIRIRCVDIDDKHTYNAMVYVDLIVGTGASEVERETAEELAKEKLRAALQVDIA
DEHSCVTQFEMKLREELLSSDSFHPDKDEYYKDFL
111000000000000000000000000000000000000000000000000000000000
000000000000000000000000000000000000000000000000000000000000
00000000000000000000000000000000000
>DM_train1925
QASGNTDQAALESAQARLAALSILVGAVGATGPGVMITIDDPGPGVAPEVMIDVINELRAAG
AEAIQINDAHRSVRVGVDTWVVGVPGSLTVDTKVLSPPYSILAIGDPPTLAAMNIPGGAQD

GVKRVGGRMVVQQADRVDVTALRQPKQHQYAQPVK
111111111111111000000000000000000000000000000000000000000000
000000000000000000000000000000000000000000000000000000000000
00000000000000000000000000000000001
>DM_train1926
GCGVEEKTEVQLLKEMPKPKAMTIDPSLSQKEATEMVHAAQRFYAFWDTGKEELIPQTVTEN
FFDHTLPKGRPQGTEGLKFAAQNFRKIVPNIHCEIEDLLVVGDKVTARLSFTGTHNDKKIDF
FAIDILHVKDGKITEDWHLEDNLTLKQQLGLIAEE
111111110000000000000000000000000000000000000000000000000000
000000000000000000000000000000000000000000000000000000000000
00000000000000000000000000000000011
>DM_train1927
MDEIKIEKLKKLDKKALNELIDVYMSGYEGLEEYGGEGRDYARNYIKWCWKKASDGFFVAKV
GDKIVGFIVCDKDWFSKYEGRIVGAIHEFVVDKKFQGKGIGRKLLITCLDFLGKYNDTIELW
VGEKNYGAMNLYEKFGFKKVGKSGIWVRMIKRQNL
000000000000000000000000000000000000000000000000000000000000
000000000000000000000000000000000000000000000000000000000000
00000000000000000000000000000000011
>DM_train1928
NAMQDIHFRRHYVRHLPKEVSQNDIIKALASPLINDGMVVSDFADHVITREQNFPTGLPVEP
VGVAIPHTDHKYVRQNAISVGILAEPVNFEDMGGEPDPVPVRVVFMLALGESNKQLNVLGWI
MDVIQDEDFMQQLLVMNDDEIYQSIYTRISERGEV
000000000000000000000000000000000000000000000000000000000000
000000000000000000000000000000000000000000000000000000000000
00000000000000000000000000000001111
>DM_train1929
SSDVNELEIGDPIPDLSLLNEDNDSISLKKITENNRVVVFFVYPRASTPGSTRQASGFRDNY
QELKEYAAVFGLSADSVTSQKKFQSKQNLPYHLLSDPKREFIGLLGAKKTPLSGSIRSHFIF
VDGKLKFKRVKISPEVSVNDAKKEVLEVAEKFKEE
110000000000000000000000000000000000000000000000000000000000
000000000000000000000000000000000000000000000000000000000000
00000000000000000000000000000000001
>DM_train1930
MGSDKIHHHHHHMRTLNKDEHNYIKQIANIHETLLSQVESNYKCTKLSIALRYEMICSRLEH
TNDKIYIYENEGQLIAFIWGHFSNEKSMVNIELLYVEPQFRKLGIATQLKIALEKWAKTMNA
KRISNTIHKNNLPMISLNKDLGYQVSHVKMYKDID
111111111100000000000000000000000000000000000000000000000000
000000000000000000000000000000000000000000000000000000000000
00000011111111111111111111111111111
>DM_train1931
MTFQASGTTGITTLTVTEECRVQVGNVTATLARSKLKDDTAIGVIGVTALGCNGLQAALQAD
PDNYDATNLYMTSRNHDKLNVKLKATDGSSWTYGNGVFYKTEGGNWGGHVGISVDGNQTDKP
TGEYTLNLTGGYWTNDNKQTFQASGTTGITTLTLE
100000000000000000000000000000000000000000000000000000000000

0000000000000000000000000000000000000000000000000000000000000000
0000000000000011111111111111111111
>DM_train1932
GSAMASNAVLYKSNHNVVYSCKYHIVWCPKYRRKVLVGAVEMRLKEIIQEVAKELRVEIIEM
QTDKDHIHILADIDPSFGVMKFIKTAKGRSSRILRQEFNHLKTKLPTLWTNSCFISTVGGAP
LNVVKQYIENQQNSNRPKQKEKWKSYVDNLQTKAL
1111111110000000000000000000000000000000000000000000000000000000
0000000000000000000000000000000000000000000000000000000000000000
000000000011111111111111111111111111
>DM_train1933
MFDQLAVFTPQGQVLYQYNCLGKKFSEIQINSFISQLITSPVTRKESVANANTDGFDFNLLT
INSEHKNSPSFNALFYLNKQPELYFVVTFAEQTLELNQETQQTLALVLKLWNSLHLSESILK
NRQGQNEKNKHNYVDILQGIEDDLKKFEQYFRIK
0000000000000000000000000000000000000000000000000000000000000000
0011111110000000000000000000000000000000000000000000000000000000
00000000000000000000000000000000111
>DM_train1934
TDENLKIAIASSPAPRYITDMSIEEISREWYMLMPRQKITGGLMVKMDQAIMDKRITLKANF
SVLFDQLETLVSLRAFTDDGAIVAEISPIPSMPGHSTEDVKNAIGILIGGLEWNDNSIRASE
NIQRFAWGIRDENGGPPLPPKQKRYMARRVESEV
1111111111111111000000000000000000000000000000000000000000000000
0000000000000000000000000000000000000000000000000000000000000000
00000011111111111111111111111111111
>DM_train1935
PPTIQEIKQKIDSYNSREKHCLGMKLSEDGTYTGFIKVHLKLRRPVTVPAGIRPQSIYDAIK
EVNPAATTDKRTSFYLPLDAIKQMHISSTTTVSEVIQGLLDKFMVVDNPQKFALFKRIHKDG
QVLFQKLSIADYPLYLRLLAGPDTDVLSFVLKEN
0000000000000000000000000000000000000000000000000001111111111111
1111111111110000000000000000000000000000000000000000000000000000
00000000000000000000000000000000000
>DM_train1936
ERLRVAAYCRVSTDSEDQLNSYKSQVQYYTDMIKKNKEWVLADIYADEAITGTQVTKREDFQ
RMINDCMNGEIDMVFTKSISRFARNTLDTLKYVRMLKERNIAVYFEDEKINTLTMDGELLLV
VLSSVAQQEVENISANVKKGLKMKMKRGELVGFQ
0000000000000000000000000000000000000000000000000000000000000000
0000000000000000000000000000000000000000000000000000000000000000
00000000000011111111111111111111111
>DM_train1937
GSSMALLQKTRIINSMLQAAAGKPVNFKEMAETLRDVIDSNIFVVSRRGKLLGYSINQQIEN
DRMKKMLEDRQFPEEYTKNLFNVPETSSNLDINSEYTAFPVENRDLFQAGLTTIVPIIGGGE
RLGTLILSRLQDQFNDDDLILAEYGATVVGMEIL
1110000000000000000000000000000000000000000000000000000000000000
0000000000000000000000000000000000000000000000000000000000000000
00000000000000000000000000000000000

>DM_train1938
GSHMNTAEEDMEDDTSWRSEATFQFTVERFSRLSESVLSPPCFVRNLPWKIMVMPRFYPDRP
HQKSVGFFLQCNAESDSTSWSCHAQAVLKIINYRDDEKSFSRRISHLFFHKENDWGFSNFMA
WSEVTDPEKGFIDDDKVTFEVFVQADAPHGVAWD
111111111111110000000000000000000000000000000000000000000000
000000000000000000000000000000000000000000000000000000000000
0000000000000000000000000000000000
>DM_train1939
SNAMEILYIKGDATAPIGSGVKVITHICNDIGGWGKGFVLALSKKWKMPEEAYRQWYKSQEE
FTLGAVQFVNVENKLYVANMIGQHGIYKDSKGLPPIRYDAVRQCLKEVALFTIAHKASVHMP
RIGCGLAGGKWELMEQIIKEELITKEIAVTVYDL
100000000000000000000000000000000000000000000000000000000000
000000000000000000000000000000000000000000000000000000000000
0000000000000000000000000000000000
>DM_train1940
GMSDDLTDAQTAAIPEGFSQLNWSRGFGRQIGPLFEHREGPGQARLAFRVEEHHTNGLGNCH
GGMLMSFADMAWGRIISLQKSYSWVTVRLMCDFLSGAKLGDWVEGEGELISEEDMLFTVRGR
IWAGERTLITGTGVFKALSARKPRPGELAYKEEA
111111111111000000000000000000000000000000000000000000000000
000000000000000000000000000000000000000000000000000000000000
0000000000000000000000000000001111
>DM_train1941
SGAISGDSLISLASTGKRVSIKDLLDEKDFEIWAINEQTMKLESAKVSRVFMTGKKLVYILK
TRLGRTIKATANHRFLTIDGWKRLDELSLKEHIALPRKLESSSLQLSPEIEKLSQSDISWDS
IVSITETGVEEVFDLTVPGPHNFVANDIIVHASI
000000000000000000000000000000000000000000000000000000000000
000000000000000000000000000000000000111111111111111110000000
0000000000000000000000000000000000
>DM_train1942
MGVFTFQDEYTSTIAPAKLYKALVTDADIIIPKAVETIQSVEIVEGNGGPGTIKKLTFIEGG
ESKYVLHKIEAIDEANLGYNYSIVGGVGLPDTIEKISFETKLVEGANGGSIGKVTIKIETKG
DAQPNEEEGKAAKARGDAFFKAIESYLSAHPDYN
100000000000000000000000000000000000000000000000000000000000
000000000000000000000000000000000000000000000000000000000000
0000000000000000000000000000000000
>DM_train1943
VDLGSKSSNSTCRLNVTELASIHPGETWTLHGMCISICYYENVTEDEIIGVAFTWQHNESVV
DLWLYQNDTVIRNFSDITTNILQDGLKMRTVPVTKLYTSRMVTNLTVGRYDCLRCENGTTKI
IERLYVRLGSLYPRPPGSGLAKHPSVSADEELSA
000000000000000000000000000000000000000000000000000000000000
000000000000000000000000000000000000000000000000000000000000
0000000001111111111111111111111111
>DM_train1944
MAPKRSRRSNRRAGSRAAATSLVYDTCYVTLTERATTSFQRQSFPTLKGMGDRAFQVVAFTI

QGVSAAPLMYNARLYNPGDTDSVHATGVQLMGTVPRTVRLTPRVGQNNWFFGNTEEAETILA
IDGLVSTKGANAPSNTVIVTGCFRLAPSELQSS
11111111111111110000000000000000000000000000000000000000000000
00000000000000000000000000000000000000000000000000000000000000
000000000000000000000000000000000
>DM_train1945
VMKLESDKTFPIMLEGKINGYACVVGGKLFRPMHVEGKIDNDVLAALKTKKASKYDLEYADV
PQNMRADTFKYTHEKPQGYYSWHHGAVQYENGRFTVPKGVGAKGDSGRPILDNQGRVVAIVL
GGVNEGSRTALSVVMWNEKGVTVKYTPENCEQW
11111000000000000000000000000000000000000000000000000000000000
00000000000000000000000000000000000000000000000000000000000000
000000000000000000000000000000000
>DM_train1946
MSHPDLNKLLELWPHIQEYQDLALKHGINDIFQDNGGKLLQVLLITGLTVLPGREGNDAVDN
AGQEYELKSINIDLTKGFSTHHHMNPVIIAKYRQVPWIFAIYRGIAIEAIYRLEPKDLEFYY
DKWERKWYSDGHKDINNPKIPVKYVMEHGTKIY
00000000000000000000000000000000000000000000000000001110000000
00000000000000000000000000000000000000000000000000000000000000
000000000000000000000000000000000
>DM_train1948
SDVAFRGNLLDRPCHVSGDSLNKHVVFKTRASRDFWYPPGRSPTESFVIRLENCHATAVGKI
VTLTFKGTEEAALPGHLKVTGVNAGRLGIALLDTDGSSLLKPGTSHNKGQGEKVTGNSLELP
FGAYVVATPEALRTKSVVPGDYEATATFELTYR
11111111000000000000000000000000000000000000000000000000000000
00000000000000000000000000000000000000000000000000000000000000
000000000000000000000000000000000
>DM_train1949
MGAMHVLTLDLAPVTPKDAPLLHRVFHLSPSYFALIGMELPTLEDVVRDLQTLEVDPRRRAF
LLFLGQEPVGYLDAKLGYPEAEDATLSLLLIREDHQGRGLGRQALERFAAGLDGVRRLYAVV
YGHNPKAKAFFQAQGFRYVKDGGPTLTWYVRPL
11100000000000000000000000000000000000000000000000000000000000
00000000000000000000000000000000000000000000000000000000000000
000000000000000000000000000000000
>DM_train1950
SSPNILLTRIDNRLVHGQVGVTWTSTIGANLLVVVDDVVANDDIQQKLMGITAETYGFGIRF
FTIEKTINVIGKAAPHQKIFLICRTPQTVRKLVEGGIDLKDVNVGNMHFSEGKKQISSKVYV
DDQDLTDLRFIKQRGVNVFIQDVPGDQKEQIPD
11100000000000000000000000000000000000000000000000000000000000
00000000000000000000000000000000000000000000000000000000000000
000000000000000000000000000000001
>DM_train1951
YFGKLESKLSVIRNLNDQVLFIDQGNRPLFEDMTDSDSRDNAPRTIFIISMYKDSQPRGMAV
TISVKSEKISTLSSENKIISFKEMNPPDNIKDTKSDIIFFQRSVPGHDNKMQFESSSYEGYF
LASEKERDLFKLILKKEDELGDRSIMFTVQNED

000000000000000000000000000000011111111000000000000000000000
000000000000000000000000000000000000000000000000000000000000
000001110000000000000000000000001
>DM_train1952
GNIESIENLQGIRALQQQAPQLLSSGLPNEQQFSLLKQAGVDVVINLMPDSSKDAHPDEGKL
VTQAGMDYVYIPVDWQNPKVEDVEAFFAAMDQHKGKDVLVHCLANYRASAFAYLYQLKQGQN
PNMAQTMTPWNDELAIYPKWQALLTEVSAKYGH
111100000000000000000000000000000000000000000000000000000000
000000000000000000000000000000000000000000000000000000000000
000000000001100000000000000000000
>DM_train1953
MYSIEVRTHSALHVVKGAVVKVLGSEAKWTYSTYVKGNKGVLIVKFDRKPSDEEIREIERLA
NEKVKENAPIKIYELPREEAEKMFGEDMYDLFPVPEDVRILKVVVIEDWNVNACNKEHTKTT
GEIGPIKIRKVRFRKSKGLLEIHFELLELENPS
000000000000000000000000000000000000000000000000000000000000
000000000000000000000000000000000000000000000000000000000000
000000000000000000000000000111111
>DM_train1954
MALFSAQSPYINPIIPFTGPIQGGLQEGLQVTLQGTTKSFAQRFVVNFQNSFNGNDIAFHFN
PRFEEGGYVVCNTKQNGQWGPEERKMQMPFQKGMPFELCFLVQRSEFKVMVNKKFFVQYQHR
VPYHLVDTIAVSGCLKLSFITFQTQNFRPAHQA
000000000000000000000000000000000000000000000000000000000000
000000000000000000000000000000000000000000000000000000000000
000000000000000000000000011111111
>DM_train1955
GMPALITYRTTVQEDWVDYNGHLRDAFYLLIFSYATDALMDRIGLDADSRGQSGNSLFTLEA
HINYLHEVKLGTEVWVQTQILGFDRKRLHVYHSLHRAGFDEVLAASEQMLLHVDLAGPQSAP
FGHTTVCRLNHLVEQQEGAQAPQYMGRTIKLPA
000000000000000000000000000000000000000000000000000000000000
000000000000000000000000000000000000000000000000000001110000
000000000000000000000000000000000
>DM_train1956
MSSSIREEVHRHLGTVALMQPALHQQTHAPAPTEITHTLFRAYTRVPHDVGGEADVPIEYHE
KEEEIWELNTFATCECLAWRGVWTAEERRRKQNCDVGQTVYLGMPYYGRWLLTAARILVDKQ
FVTLTELHNKIVEMRERVASGQGLGEYLPPKAK
111000000000000000000000000000000000000000000000000000000000
000000000000000000000000000000000000000000000000000000000000
000000000000000000000000000000111
>DM_train1957
MKLHDLKPTPGSRKDRKRVGRGPGGTDKTAGRGHKGQKSRSGAGKGAFFEGGRSRLIARLPK
RGFNNVGTTYEVVKLSQLQDLEDTTFDRDTLEAYRLVRRKNRPVKLLASGEISRAVTVHVDA
ASAAAIKAVEAAGGRVVLPEVQTQQDDAQKAE
111000000000000000000000000000000000000000000000000000000000
000000000000000000000000000000000000000000000000000000000000

00000000000000000000111111111111
>DM_train1958
MSDRKPVRGRHQARKRAVALLFEAEVRGISAAEVVDTRAALAEAKPDIARLHPYTAAVARGV
SEHAAHIDDLITAHLRGWTLDRLPAVDRAILRVSVWELLHAADVPEPVVVDEAVQLAKELST
DDSPGFVNGVLGQVMLVTPQLRAAAQAVRGGA
111111110000000000000000000000000000000000000000000000000000
000000000000000000000000000000000000000000000000000000000000
00000000000000011111111111111111
>DM_train1959
MLKQVEIFTDGSALGNPGPGGYGAILRYRGREKTFSAGYTRTTNNRMELKAAIEGLKALKEP
AEVDLYTDSHYLKKAFTEGWLEGWRKRGWRTAEGKPVKNRDLWEALLLAMAPHRVRFHFVKG
HAGHPENERADELARAAAMNPTLEDTGYQVEV
000000000000000000000000000000000000000000000000000000000000
000000000000000000111111111111111110000000000000000000000000
00000000000000000000000000001111
>DM_train1960
MIVTSGVLVENGKVLLVKHKRLGVYIYPGGHVEHNETPIEAVKREFEEETGIVVEPIGFTYG
IIDENAVERPMPLVILEEVVKYPEETHIHFDLIYLVKRVGGDLKNGEWIDVREIDRIETFPN
VRKVVSLALSTLYRLGKISKLAAALEHHHHHH
000000000000000000000000000000000000000000000000000000000000
000000000000000000000000000000000000000000000000000000000000
00000000000000000000000111111111
>DM_train1961
MSSRSELLLEKFAEKIGIGSISFNENRLCSFAIDEIYYISLSDANDEYMMIYGVCGKFPTDN
SNFALEILNANLWFAENGGPYLCYEAGAQSLLLALRFPLDDATPEKLENEIEVVVKSMENLY
LVLHNQGITLENEHMKIEEISSSDNKHYYAGR
000000000000000000000000000000000000000000000000000000000000
000000000000000000000000000000000000000000000000000000000000
00000000001111100000001111111111
>DM_train1962
MWGKKDAGTELTNYQTLATNTIGMMKGVDGYAFTSGAKMTDTLIQAGAAKGMTVSGDPASGS
ATLWNSWGGQIVVAPDTAGGTGFNNGFTITTNKVPQSACVSISTGMSRSGGTSGIKINGNNH
TDAKVTAEIASSECTADNGRTGTNTLVFNYNG
111111000000000000000000000000000000000000000000000000000000
000000000000000000000000000000000000000000000000000000000000
00000000000000000000000000000000
>DM_train1963
GMPENTAMAALLPEESLFFRELVKQWRAQDSYGTWEKKSDMELLAPYVLDKEQRRAIPIIGD
PDPEILWRVELFYNAVGLATERASGVMVSPMMKMSHEGFGRMVLIAGRLIVVNKQLRDVHRF
GFPSMEKLAEEGDKLVAGALEMIEKFPEVARF
111111111110000000000000000000000000000000000000000000000000
000000000000000000000000000000000000000000000000000000000000
00000000000000000000000000000000
>DM_train1964

GGGVMHYTDKAALPADGEAREVAALFDTWNAALATGNPHKVADLYAPDGVLLPTVSNEVRAS
REQIENYFEMFLTKKPKGVINYRTVRLLDDDSAVDAGVYTFTLTDKNGKKSDVQARYTFVYE
KRDGKWLIINHHSSAMPEVDTATATAAVTKAK
111000000000000000000000000000000000000000000000000000000000
000000000000000000000000000000000000000000000000000000000000
00000000000000000001111111111111
>DM_train1965
HMLLTTSRKPSQRTRSFSQRLSRIMGWRYINRGKMSLRDVLIEARGPVAVVSERHGNPARIT
FLDERGGERGYILFNPSFEMKKPELADKAVRVSSCPPGSEGLCNLMGLEVDESSSRDAWSIR
TDEEYAWVMELMDARGTPAGFKLLIRDFRVGE
100000000000000000000000000000000000000000000000000000000000
000000000000000000000001110000000000000000000000000110000000
00000000000000000000000000000001
>DM_train1966
VTDVLLCVGNSMMGDDGAGPLLAEKCAAAPKGNWVVIDGGSAPENDIVAIRELRPTRLLIVD
ATDMGLNPGEIRIIDPDDIAEMFMMTTHNMPLNYLIDQLKEDIGEVIFLGIQPDIVGFYYPM
TQPIKDAVETVYQRLEGWEGNGGFAQLAVEEE
000000000000000000000000000000000000000000000000000000000000
000000000000000000000000000000000000000000000000000000000000
00000000000000000000000000000111
>DM_train1968
SITKPQNKSAKLVDGELLVKASVESFGLEDEKYWFLVCCELSNGKTRQLKRYYQDFYDLQVQ
LLDAFPAEAGKLRDAGGQWSKRIMPYIPGPVPYVTNSITKKRKEDLNIYVADLVNLPDYISR
SEMVHSLFVVLNNGFDREFERDENGSHHHHHH
111111111100000000000000000000000000000000000000000000000000
000000000000000000000000000000000000000000000000000000000000
00000000000000000000011111111111
>DM_train1969
MPLLSASIVSAPVVTSETYVDIPGLYLDVAKAGIRDGKLQVILNVPTPYATGNNFPGIYFAI
ATNQGVVADGCFTYSSKVPESTGRMPFTLVATIDVGSGVTFVKGQWKSVRGSAMHIDSYASL
SAIWGTAAPSSQGSGNQGAETGGTGAGNIGGG
100000000000000000000000000000000000000000000000000000000000
000000000000000000000000000000000000000000000000000000000000
00000000111111111111111111111111
>DM_train1970
MLEDLIGKAYLESAEDRRRGDRSEEVEAIRKYIRSARRTVVPNWNAEKVDAINDVLRSFNLR
EAEHLQFNTNWADLTRMPAVTKALMALDISGADLVIARGRLGVPGSGSLLVIMDSRGRLLSA
AMSPPHVIHSMEVREAVRSEMTHALERIGFKR
000000000000000000000000000000000000000000000000000000000000
000000000000000000000000000000000000000000000000000000000000
00000000000000000000000000000001
>DM_train1971
ALSPPRRKSFKKWTPPRSPFNLVQEILFHDPWKLLIATIFLNRTSGKMAIPVLWEFLEKYPS
AEVARAADWRDVSELLKPLGLYDLRAKTIIKFSDEYLTKQWRYPIELHGIGKYGNDSYRIFC

VNEWKQVHPEDHKLNKYHDWLWENHEKLSLS
111111111110000000000000000000000000000000000000000000000000
000000000000000000000000000000000000000000000000000000000000
000000000000000000000000000000
>DM_train1972
GTGQAADLKSTQAAIDQINGKLNRVIEKTNEKFHQIEKEFSEVEGRIQDLEKYVEDTKIDLW
SYNAELLVALENQHTIDLTDSEMNKLFEKTRRQLRENAEEMGNGSFKIYHKCDNACIESIRN
GTYDHDVYRDEALNNRFQIKGVELKSGYKDW
111000000000000000000000000000000000000000000000000000000000
000000000000000000000000000000000111111110000000000000000000
000000000000000000001111111111
>DM_train1974
GTLDSTWKEATLPQVKAMLEKDDGKVSGDTVTYSGKTVHVVAAAVLPGFPFPSFEVHDKKNP
TLEIPAGATVDVTFINTNKGFGHSFDITKKGPPYAVMPVIDPIVAGTGFSPVPKDGKFGYTD
FTWHPTAGTYYYVCQIPGHAATGMFGKIVVK
111110000000000000000000000000000000000000000000000000000000
000000000000000000000000000000000000000000000000000000000000
000000000000000000000000000000
>DM_train1975
MIISSLTNPNFKVGLPKVIAEVCDYLNTLDLNALENGRHDINDQIYMNVMEPETAEPSSKKA
ELHHEYLDVQVLIRGTENIEVGATYPNLSKYEDYNEADDYQLCADIDDKFTVTMKPKMFAVF
YPYEPHKPCCVVNGKTEKIKKLVVKVPVKLI
000000000000000000000000000000000000000000000000001111111100
000000000000000000000000000000000000000000000000000000000000
000000000001111110000000000000
>DM_train1976
MKAPELQIQQWFNSATDLTLADLRGKVIVIEAFQMLCPGCVMHGIPLAQKVRAAFPEDKVAV
LGLHTVFEHHEAMTPISLKAFLHEYRIKFPVGVDQPGDGAMPRTMAAYQMRGTPSLLLIDKA
GDLRAHHFGDVSELLLGAEIATLLGEAAPSV
000000000000000000000000000000000000000000000000000000000000
000000000000000000000000000000000000000000000000000000000000
000000000000000000000000000111
>DM_train1977
MAMRKIYIAGPAVFNPDMGASYYNKVRELLKKENVMPLIPTDNEATEALDIRQKNIQMIKDC
DAVIADLSPFRGHEPDCGTAFEVGCAAALNKMVLTFTSDRRNMREKYGSGVDKDNLRVEGFG
LPFNLMLYDGVEVFDSFESAFKYFLANFPSK
110000000000000000000000000000000000000000000000000000000000
000000000000000000000000000000000000000000000000000000000000
000000000000000000000000000001
>DM_train1978
MGSDKIHHHHHHMSKKQKSKYIVIFGCGRLGSLIANLASSSGHSVVVVDKNEYAFHRLNSEF
SGFTVVGDAAEFETLKECGMEKADMVFAFTNDDSTNFFISMNARYMFNVENVIARVYDPEKI
KIFEENGIKTICPAVLMIEKVKEFIIGSEED
111111111111111000000000000000000000000000000000000000000000

000000000000000000000000000000000000000000000000000000000000
0000000000000000000000000000111
>DM_train1979
MDIREIEQERASFAFKVVSDIKDKYSQNKKVQGKYSSYAEKAPTIILNNGLGATLAFFLSKL
EKPIDDVDYKSINPESFGNAENIAYAFLYKHLSTWLAEGNGKDSAFSGLTNGEDPLKYIMEK
TAIDVAISTEEALSILNWIKKFAKAMLEEEL
100000000000000000000000000000000000000000000000000000000000
000000000000000000000000000000000000000000000000000000000000
0000000000000000000000000000011
>DM_train1980
MSLIPKFRAWDTYEKEMLENVTPLFDDSNSMIAIITDFQIKGSPGTSEIEIGSYDTTFNWDE
FPYVIMQSTGLKDKNGVEIFEGDILVYDAPKKYAHRRSMHEIAYADGRFFWEFLDLVFCQSN
ILYRDGYLVIGNIHENPELLEGNEGHHHHHH
100000000000000000000000000000000000000000000000000000000000
000000000000000000000000000000000000000000000000000000000000
0000000000000000000011111111111
>DM_train1981
GSRATVFKLGLFKSLFLCSFHDITRLFKNDKTTNQQWVLAVFGLAEVFFEASFELLKKQCSF
LQMQKRSHEGGTCAVYLICFNTAKSRETVRNLMANMLNVREECLMLQPPKIRGLSAALFWFK
SSLSPATLKHGALPEWIRAQTTLNESLQTE
111000000000000000000000000000000000000000000000000000000000
000000000000000000000000000000000000000000000000000000000000
000000000000000000000000111111
>DM_train1982
PVVIKTEGPAWTPLEPKLITRLADTVRTKGLRSPITMAEVEALMSSPLLPHDVTNLMRVILG
PAPYALWMDAWGVQLQTVIAAATRDPRHPANGQGRGERTNLNRLKGLADGMVGNPQGQAALL
RPGELVAITASALQAFREVARLAEPAGPWA
000001110000000000000000000000000000000000000000000000000000
000000000000000000000000000000001110000000000000000000000000
000000000000000000000001111111
>DM_train1983
EELIEIREARMDDLDTIAKFNYNLAKETEGKELDMDVLTKGVKALLLDERKGKYHVYTVFDK
VVAQIMYTYEWSDWRNGNFLWIQSVYVDKEYRRKGIFNYLFNYIKNICDKDENIVGMRLYVE
KENINAKATYESLNMYECDYNMYEYEVIHS
100000000000000000000000000000000000000000000000000000000000
000000000000000000000000000000000000000000000000000000000000
000000000000000000000000000001
>DM_train1984
GMFASLVIPVSAQANSGEMPQEQQLAVKYMDALTEHDYKTLITFYNRDSIFFDKTANRKYTG
GRFIIDFLERAHQGVLEYDFNIEHMYNAGSLVVMIGNYHFKGPGEQFGKPGKIIDVAIPAVT
SLKLDMLNRRVTEHVDLIDYQTMSDQLAMQ
111111111111111110000000000000000000000000000000000000000000
000000000000000000000000000000000000000000000000000000000000
000000000000000000000000000000

>DM_train1985
MKKILLLNGPNLNMLGKREPHIYGSQTLSDIEQHLQQSAQAQGYELDYFQANGEESLINRIH
QAFQNTDFIIINPGAFTHTSVAIRDALLAVSIPFIEVHLSNVHAREPFRHHSYLSDVAKGVI
CGLGAKGYDYALDFAISELQKIQLGEMMNG
000000000000000011111111000000000000000000000000000000000000000
000000000000000000000000000000000000000000000000000000000000000
0000000000000000000000011111111
>DM_train1986
MGSDKIHHHHHHMSDSLVVCEVDPELKETLRKFRFRKETNNAAIIMKVDKDRQMVVLEDELQ
NISPEELKLELPERQPRFVVYSYKYVHDDGRVSYPLCFIFSSPVGCKPEQQMMYAGSKNRLV
QTAELTKVFEIRTTDDLTETWLKEKLAFFR
111111111111111110000000000000000000000000000000000000000000000
000000000000000000000000000000000000000000000000000000000000000
000000000000000000000000000000
>DM_train1987
MGSSSSGLVPRGSSFMLQHLNSGWAVDQAIVNEDERLVCIRFGHDYDPDCMKMDELLYKVAD
DIKNFCVIYLVDITEVPDFNTMYELYDPVSVMFFYRNKHMMIDLGTGNNNKINWPMNNKQEF
IDIVETIFRGARKGRGLVISPKDYSTKYKY
111111111111111000000000000000000000000000000000000000000000000
000000000000000000000000000000000000000000000000000000000000000
000000000000000000000000111111
>DM_train1988
MAGDPNSMTVSHHNASTARFYALRLLPGQEVFSQLHAFVQQNQLRAAWIAGCTGSLTDVALR
YAGQEATTSLTGTFEVISLNGTLELTGEHLHLAVSDPYGVMLGGHMMPGCTVRTTLELVIGE
LPALTFSRQPCAISGYDELHISSRLEHHHH
111111111110000000000000000000000000000000000000000000000000000
000000000000000000000000000000000000000000000000000000000000000
000000000000000000000000011111
>DM_train1989
GMTRTMSFHKNCELCTTAGGEILWQDALCRVVHVENQDYPGFCRVILNRHVKEMSDLRPAER
DHLMLVVFAVEEAVREVMRPDKINLASLGNMTPHVHWHVIPRFKRDRHFPNSVWGETKRESL
PQALDQGSTTALKKAISVRLDQGEPVFMGM
111111000000000000000000000000000000000000000000000000000000000
000000000000000000000000000000000000000000000000000000000000000
000000000000000000000111111111
>DM_train1990
SNAMTQQDFRTKVDNTVFGVRATALIVQNHKLLVTKDKGKYYTIGGAIQVNESTEDAVVREV
KEELGVKAQAGQLAFVVENRFEVDGVSYHNIEFHYLVDLLEDAPLTMQEDEKRQPCEWIDLD
KLQNIQLVPVFLKTALPDWEGQLRHIHLEE
111000000000000000000000000000000000000000000000000000000000000
000000000000000000000000000000000000000000000000000000000000000
000000000000000000000000000001
>DM_train1991
GHMPLPTELARHLTEEKIAFVQRSGLRAEVLEPGYVRLRMPGAGNENHIGSMYAGALFTLAE

LPGGALFLTSFDSARFYPIVKEMTLRFRRPAKGDIRVEARLDAERIRQLETEAGERGKAEYS
LELQLTDEQGEVVAESAALYQLRSHARPGS
000000000000000000000000000000000000000000000000000000000000
000000000000000000000000000000000000000000000000000000000000
000000000000000000000000011111
>DM_train1992
MGKSASKQFHNEVLKAHNEYRQKHGVPPLKLCKNLNREAQQYSEALASTRILKHSPESSRGQ
CGENLAWASYDQTGKEVADRWYSEIKNYNFQQPGFTSGTGHFTAMVWKNTKKMGVGKASASD
GSSFVVARYFPAGNVVNEGFFEENVLPPKK
111000000000000000000000000000000000000000000000000000000000
000000000000000000000000000000000000000000000000000000000000
000000000000000000000000000011
>DM_train1993
APVRSLNCTLRDSQQKSLVMSGPYELKALHLQGQDMEQQVVWSMSFVQGEESNDKIPVALGL
KEKNLYLSCVLKDDKPTLQLESVDPKNYPKKKMEKRFVFNKIEINNKLEFESAQFPNFYIST
SQAENMPVFLGGTKGGQDITDFTMQFVSS
111000000000000000000000000000000000000000000000000000000000
000000000000000000000000000000000000000000000000000000000000
00000000000111100000000000000
>DM_train1994
MNVRATYTVIFKNASGLPNGYDNWGWGCTLSYYGGAMIINPQEGKYGAVSLKRNSGSFRGGS
LRFDMKNEGKVKILVENSEADEKFEVETISPSDEYVTYILDVDFDLPFDRIDFQDAPGNGDR
IWIKNLVHSTGSADDFVDPINLEHHHHHH
000000000000000000000000000000000000000000000000000000000000
000000000000000000000000000000000000000000000000000000000000
00000000000000000011111111111
>DM_train1995
SNAMAKNKLLRMDNVSIVVESLDNAISFFEEIGLNLEGRANVEGEWAGRVTGLGSQCVEIAM
MVTPDGHSRIELSRFLTPPTIADHRTAPVNALGYLRVMFTVEDIDEMVSRLTKHGAELVGEV
VQYENSYRLCYIRGVEGILIGLAEELGNK
111111000000000000000000000000000000000000000000000000000000
000000000000000000000000000000000000000000000000000000000000
00000000000000000000000000011
>DM_train1996
MTDDSAVESKQKKSKIRKGHWIPVVAGFLRKDGKILVGQRPENNSLAGQWEFPGGKIENGET
PEEALARELNEELGIEAEVGELKLACTHSYGDVGILILFYEILYWKGEPRAKHHMMLEWIHP
EELKHRNIPEANRKILHKIYKALGLEWRK
111111111111111100000000000000000000000111111000000000000000
000000000000000000000000000000000000000000000000000000000000
00000000000000000000000000011
>DM_train1997
LEDRKNERLPSDLARRATAIIEMPDGVLVTASRGGRYNLPGGKANRGELRSQALIREIREET
GLRINSMLYLFDHITPFNAHKVYLCIAQGQPKPQNEIERIALVSSPDTDMDLFVEGRAILRR
YARLRNEETAKGEALRALLGLARYIAKVD

11111111000000000000000000000000111000000000000000000000000000
00000000000000000000000000000000000000000000000000000000000000
00000000000000000000000000000
>DM_train1998
MENQKMQEPLVYRRILLTVDEDDNTSSERAFRYATTLAHDYDVPLGICSVLESEDINIFDSL
TPSKIQAKRKHVEDVVAEYVQLAEQRGVNQVEPLVYEGGDVDDVILEQVIPEFKPDLLVTGA
DTEFPHSKIAGAIGPRLARKAPISVIVVR
10000000000000000000000000000000000000000000000000000111111111
11000000000000000000000000000000000000000000000000000000000000
00000000000000000000000000000
>DM_train1999
LSSTQSVNSVLAPGNLRKVHHIALNVQDMQASRYFYGTILGLHELTDDEVPATLTELVASGK
VANFITPDGTILDLFGEPELSPPDPNPEKTFTRAYHLAFDIDPQLFDRAVTVIGENKIAIAH
GPVTRPTGRGVYFYDPDGFMIEIRCDPEA
11111000000000000000000000000000000000000000000000000000000000
00000000000000000000000000000000000000000000000000000000000000
00000100000000000000000000001
>DM_train2000
GHMTEIERKFLVATFPDGELHAVPLRQGYLTTPTDSIELRLRQQGTEYFMTLKSEGGLSRQE
YEIQIDVTQFEMLWPATEGRRVEKTRYSGKLPDGQLFELDVFAGHLSPLMLVEVEFLSEDAA
QAFIPPPWFGEEVTEDKRYKNKALALSIP
11100000000000000000000000000000000000000000000000000011111100
00000000000000000000000000000000000000000000000000000000000000
00000000000000000000000000000
>DM_train2001
AGHMSLYRLIYSSQGIPNLQPQDLKDILESSQRNNPANGITGLLCYSKPAFLQVLEGECEQV
NETYHRIVQDERHHSPQIIECMPIRRRNFEVWSMQAITVNDLSTEQVKTLVLKYSGFTTLRP
SAMDPEQCLNFLLDIAKIYELSDNFFLDL
11110000000000000000000000000000000000000000000000000000000000
00000000000000000000000000000000000000000000000000000000000000
00000000000000000001111111111
>DM_train2002
GMIYSKVENFINENKQNAIFTEGASHENIGRIEENLQCDLPNSYKWFLEKYGAGGLFGVLVL
GYNFDHASVVNRTNEYKEHYGLTDGLVVIEDVDYFAYCLDTNKMKDGECPVVEWDRVIGYQD
TVADSFIEFFYNKIQEAKDDWDEDEDWDD
00000000000000000000000000000000000000000000000000000000000000
00000000000000000000000000000000000000000000000000000000000000
00000000000000000000000000001
>DM_train2003
MEKLKEFRGIKEHLGVFREAVKDAERIGFAGVPGVCTPFAQLFAYAVRDKDNIFIPNTDFSK
ARKLEVTEYGVELGEISPGNVDVLVLLGGLSMPGIGSDIEDVKKLVEDALEEGGELMGLCYM
DMFARAGWYELLDFDCVINADIDGYVLRG
00000000000000000000000000000000000000000000000000000000000000
00000000000000000000000000000000011000000000000000000000000000

0000000000000000000000000000
>DM_train2004
GMQQPTASVVSYVAEYHKATETTMGRYKKVIEITGHDEVAAKLLEGLIDAGTRYFSKVVEME
HRMASARFRLDGEELRELTETLDRSRRLAHESLISSLHVFNRYIVKEYGEELKEAGIEGGIF
PKPEANRDRIAIADWAGELLTGIYENRHR
1111111111111111111000000000000000000000000000000000000000000000
0000000000000000000000000000000000000000000000000000000000000000
0000000000000000000000000000
>DM_train2005
GAMDPDLEATLRAIVHSATSLVDARYGAMEVHDRQHRVLHFVYEGIDEETVRRIGHLPKGLG
VIGLLIEDPKPLRLDDVSAHPASIGFPPYHPPMRTFLGVPVRVRDESFGTLYLTDKTNGQPF
SDDDEVLVQALAAAAGIAVANARLYQQAK
1111000000000000000000000000000000000000000000000000000000000000
0000000000000000000000000000000000000000000000000000000000000000
0000000000000000000000000111
>DM_train2006
SEDNITVRFVTENDKEGWQRLWKSYQDFYEVSFPDDLDDFNFGRFLDPNIKMWAAVAVESSS
EKIIGMINFFNHMTTWDFKDKIYINDLYVDENSRVKGAGGKLIQFVYDEADKLGTPSVYWCT
DESNHRAQLLYVKVGYKAPKILYKRKGY
1110000000000000000000000000000000000000000000000000000000000000
0000000000000000000000000000000000000000000000000000000000000000
000000000000000000000000000
>DM_train2007
EKTCDLVGEKGKESEKELALLKRLTPLFQKSFESTVGQSPDMYSYVFRVCREAGQHSSGAGL
VQIQKSNGKETVVGRFNETQIFQGSNWIMLIYKGGDEYDNHCGREQRRAVVMISCNRHTLAD
NFNPVSEERGKVQDCFYLFEMDSSLACS
0000000000000000000000000000000000111111000000000000000000000000
0000000000000000000000000000000000000000000000000000000000000000
000000000000000000000000000
>DM_train2008
GSMSAKDERAREILRGFKLNWMNLRDAETGKILWQGTEDLSVPGVEHEARVPKKILKCKAVS
RELNFSSTEQMEKFRLEQKVYFKGQCLEEWFFEFGFVIPNSTNTWQSLIEAAPESQMMPASV
LTGNVIIETKFFDDDLLVSTSRVRLFYV
1111100000000000000000000000000000000000000000000000000000000000
0000000000000000000000000000000000000000000000000001111111111111
111111000000000000000000000
>DM_train2009
MELTTRTLPARKHIALVAHDHCKQMLMSWVERHQPLLEQHVLYATGTTGNLISRATGMNVNA
MLSGPMGGDQQVGALISEGKIDVLIFFWDPLNAVPHDPDVKALLRLATVWNIPVATNVATAD
FIIQSPHFNDAVDILIPDYQRYLADRLK
0000000000000000000000000000000000000000000000000000000000000000
0000000000000000000000000000000000000000000000000000000000000000
000000000000000000011111111
>DM_train2010

MLEKVEEWIKARGLTWRLLIMQKPTRTVAEAAALLGVSESEIVKTLIVLDNAGGVYAVVIPG
DKRLNINSMKELAGKPVRLARANEVVELTGYPVGGVPPVALPPNIVLVVDRILLSRKKVYGG
GGRENALLEFSPRELVEATGAVVADVSE
11000000000000000000000000000000000000000000000000000000000000
00000000000000000000000000000000000000000000000000000000000000
0000000000000000000000000000
>DM_train2011
MKPKSGAVRRLEKNTMNFVLSDVADAEAEKAIRDPLVAYNLARFGESDKRDLNITIRNDDNS
VTGGLVGHTARGWLYVQLLFVPEAMRGQGIAPKLLAMAEEEARKRGCMGAYIDTMNPDALRT
YERYGFTKIGSLGPLSSGQSITWLEKRF
11111111111111100000000000000000000000000000000000000000000000
00000000000000000000000000000000000000000000000000000000000000
0000000000000000000000000000
>DM_train2012
GSHMSSSPAKDPPIQRLRGAVTRCEDGQLFISSYKNEYQTMEVQNNSVVIKCDGLYIIYLKG
SFFQEVKIDLHFREDHNPISIPMLNDGRRIVFTVVASLAFKDKVYLTVNAPDTLCEHLQIND
GELIVVQLTPGYCAPEGSYHSTVNQVPL
11111111111000000000000000000000000000000000000000000000000000
00000000000000000000000000000000000000000000000000000000000000
0000000000000001111111111111
>DM_train2013
GMNPENWLLLRRVVRFGDTDAAGVMHFHQLFRWCHESWEESLESYGLNPADIFPGSRKSEVT
PEVALPIIHCQADFRRPIHTGDALAMELRPERLNPNSFQVHFEFRCEEQIAAHALIRHLAIN
AQTRHCALPEGIDRWLEASGVGKIGSI
10000000000000000000000000000000000000000000000000000000000000
00000000000000000000000000000000000000000000000000000000000000
000000000000000000001111111
>DM_train2014
MNPAHLLVLSAVCVSLLGASSIPPQPLHLIQFGNMIQCTVPGFLSWIKYADYGCYCGAGGSG
TPVDKLDRCCQVHDNCYTQAQKLPACSSIMDSPYVKIYSYDCSERTVTCKADNDECAAFICN
CDRVAAHCFAASPYNNNNYNIDTTTRC
11111111111111111111111111100000000000000000000000000000000000
00000000000000000000000000000000000000000000000000000000000000
000000000000000000000000000
>DM_train2015
MRGSHHHHHHGSENLYFQSGLFDAPGRSQFVPADQAFAFDFQQNQHDLNLTWQIKDGYYLYR
KQIRITPEHAKIADVQLPQGVWHEDEFYGKSEIYRDRLTLPVTINQASAGATLTVTYQGCAD
AGFCYPPETKTVPLSEVVANNAAPQPV
11111111111111111111111111110000000000000000000000000000000000
00000000000000000000000000000000000000000000000000000000000000
000000000000000000001111111
>DM_train2016
MSKNTVSSARFRKVDVDEYDENKFVDEDDGGDGQAGPDEGEVDSCLRQGNMTAALQAALKNP
PINTKSQAVKDRAGSIVLKVLISFKANDIEKAVQSLDKNGVDLLMKYIYKGFESPSDNSSAV

LLQWHEKALAAGGVGSIVRVLTARKTV
1111111100000000000000000000001111000000000000000000000000000000
0000000000000000000000000000000000000000000000000000000000000000
000000000000000000000000000
>DM_train2018
QGGAAAPISSHARLDKSNFQQPYITNRTFMLAKEASLADNNTDVRLIGEKLFHGVSMSERCY
LMKQVLNFTLEEVLFPQSDRFQPYMQEVVPFLARLSNRLSTCHIEGDDLHIQRNVQKLKDTV
KKLGESGEIKAIGELDLLFMSLRNACI
1111111111100000000000000000000000000000000000000000000000000000
0000000000000000000000000000000000000000001111111110000000000000
000000000000000000000000000
>DM_train2019
GMAKKTLILYYSWSGETKKMAEKINSEIKDSELKEVKVSEGTFDADMYKTSDIALDQIQGNK
DFPEIQLDNIDYNNYDLILIGSPVWSGYPATPIKTLLDQMKNYRGEVASFFTSAGTNHKAYV
SHFNEWADGLNVIGVARDDSEVDKWSK
1100000000000000000000000000000000000000000000000000000000000000
0000000000000000000000000000000000000000000000000000000000000000
000000000000000000000000000
>DM_train2020
MRTFESVADLAAAAGEKVGQSDWVTITQEEVNLFADATGDHQWIHVDPERAAAGPFGTTIAH
GFMTLALLPRLQHQMYTVKGVKLAINYGLNKVRFPAPVPVGSRVRATSSLVGVEDLGNGTVQ
ATVSTTVEVEGSAKPACVAESIVRYVA
1000000000000000000000000000000000000000000000000000000000000000
0000000000000000000000000000000000000000000000000000000000000000
000000000000000000000000001
>DM_train2021
MDCMRKLDRVDMQLVKILSENSRLTYRELADILNTTRQRIARRIDKLKKLGIIRKFTIIPDI
DKLGYMYAIVLIKSKVPSDADKVISEISDIEYVKSVEKGVGRYNIIVRLLLPKDIKDAENLI
SEFLQRIKNAENVEVILISEVRKFEII
1111100000000000000000000000000000000000000000000000000000000000
0000000000000000000000000000000000000000000000000000000000000000
000000000000000000000000000
>DM_train2022
MGSSSSGLVPRGSMSSGTMKFNGYLRVRIGEAVGLQPTRWSLRHSLFKKGHQLLDPYLTVSV
DQVRVGQTSTKQKTNKPTYNEEFCANVTDGGHLELAVFHETPLGYDHFVANCTLQFQELLRT
TGASDTFEGWVDLEPEGKVFVVITLTG
1111000000111111100000000000000000000000000000000000000000000000
0000000000000000000000000000000000000000000000000000000000000000
000000000000000000000000001
>DM_train2023
MHRTSNGSHATGGNLPDVASHYPVAYEQTLDGTVGFVIDEMTPERATASVEVTDTLRQRWGL
VHGGAYCALAEMLATEATVAVVHEKGMMAVGQSNHTSFFRPVKEGHVRAEAVRIHAGSTTWF
WDVSLRDDAGRLCAVSSMSIAVRPRRD
1111111111100000000000000000000000000000000000000000000000000000

0000000000000000000000000000000000000000000000000000000000000000
000000000000000000000000001
>DM_train2024
MAITVKGLTNKLTRTQRRIAVVEFIFSLLFFLPKEAEVIQADFLEYDTKERQLNEWQKLIVK
AFSENIFSFQKKIEEQQLKNQLEIQTKYNKISGKKIDLLTTAVVLCALSEQKAHNTDKPLLI
SEALLIMDHYSQGAEKKQTHALLDKLL
1111111111100000000000000000000000000000000000000000000000000000
0000000000000000000000000011111111100000000000000000000000000000
000000000000000000000000000
>DM_train2025
SNAMVHDSALPFDALPMPPQGREGFEECPYLDSQWVADTNGQRMTGQGVDTRFDTPACVFWS
YPEAPQATVMVRHMPSEEEAIRVVDWAAPIDTTEPAEEPDGWSGGRAGHEEGAVYAVQKGPV
AVVVWSNQQQSLKAELMAKEAIARLGL
1111000000000000000011100000000000000000000000000000000000000000
0000000000000000000000000000000000000000000000000000000000000000
000000000000000000000000000
>DM_train2026
GMQGAGRLMQPRPDSAFVHDVRVTWGDCDPAKIAYTGHLPRFALEAIDAWWSEYHGPGGWYH
LELDTNVGTPFVRLEMDFKSPVTPRHILKCHTWPTRLGTKSITFRVDGVQDGVTCFVGAFTC
VFTIADQFKSQPAPDHLRALIEPHIPA
1111111100000000000000000000000000000000000000000000000000000000
0000000000000000000000000000000000000000000000000000000000000000
000000000000000000000000000
>DM_train2027
MSLDRTTQQPFGNGYLSVEQANLILNHLPLEITFVNKDDIFQYYNDSVPAAEMVFKRTPSQV
GRNVELCHPPKVLDKVKKVFELLRNGQRDKVNMWFQSERLGKFVYVTYAAVRDQAGDFQGVL
EYVQDIKPFFELDSEFNRDEGHHHHHH
1110000000000000000000000000000000000000000000000000000000000000
0000000000000000000000000000000000000000000000000000000000000000
000000000000011111111111111
>DM_train2029
MRNEMHLQFSARSENESFARVTVAAFVAQLDPTMDELTEIKTVVSEAVTNAIIHGYNNDPNG
IVSISVIIEDGVVHLTVRDEGVGIPDIEEARQPLFTTKPELERSGMGFTIMENFMDEVIVES
EVNKGTTVYLKKHIVKSKRYLEGSSF
0000000000000000000000000000000000000000000000000000000000000000
0000000000000000000000000000000000000000000000000000000000000000
00000000000000000111111111
>DM_train2031
GMTAVFRNTVLVRFKHCDAAGIVFYPRYFEMLNDFIEDWFAQALDWPFDAMHGAGQAGVPTA
DLHCRFVAPSRLGETLTRELRVVKLGQSSFTVQVRFMGPDSGLRLEVTQRLVCVDTDKIAPR
PLPDPVRQAMATYVDETLAATGSPGL
0000000000000000000000000000000000000000000000000000000000000000
0000000000000000000000000000000000000000000000000000000000000000
00000000000000000001111111

>DM_train2032
MPMTLDLQPGQRLARGVARHLRAHGFVSVEEFVPARGLRVDVMGLGPKGEIWVIECKSSRAD
FQADAKWQGYLEWCDRYFWAVDMEFPAELLPAESGLLIADAYDAEIVRMAPEQKLAPARRKV
LIQKFATHAARRLQALRDPEGHGIFE
111000000000000000000000000000000000000000000000000000000000
000000000000000000000000000000000000000000000000000000000000
00000000000000000000000000
>DM_train2033
EADSGDGDSSEYQDGKEFGIGDLVWGKIKGFSWWPAMVVSWKATSKRQAMSGMRWVQWFGDG
KFSEVSADKLVALGLFSQHFNLATFNKLVSYRKAMYHALEKARVRAGKTFPSSPGDSLEDQL
KPMLEWAHGGFKPTGIEGLKPNNTQP
111111111100000000000000000000000000000000000000000000000000
000000000000000000000000000000000000000000000000000000000000
00000000000000000000011111
>DM_train2034
TGKTGTQLLADKLKKLQVKDFQSIPVVIHENVSVYDAICTMFLEDVGTLFVVDRDAVLVGVL
SRKDLLRASIGQQELTSVPVHIIMTRMPNITVCRREDYVMDIAKHLIEKQIDALPVIKDTDK
GFEVIGRVTKTNMTKILVSLSENEIL
111110000000000000000000000000000000000000000000000000000000
000000000000000000000000000000000000000000000000000000000000
00000000000000000000000000
>DM_train2035
GMDVTIERVNDFDGYNWLPLLAKSSQEGFQLVERMLRNRREESFQEDGEAMFVALSTTNQVL
ACGGYMKQSGQARTGRIRHVYVLPEARSHGIGTALLEKIMSEAFLTYDRLVLYSEQADPFYQ
GLGFQLVSGEKITHTLDKTAFADSNR
100000000000000000000000000000000000000000000000000000000000
000000000000000000000000000000000000000000000000000000000000
00000000000000000000000000
>DM_train2036
MGSDKIHHHHHHMQARWIGNMMFHVRTDSNHDVLMDTKEEVGGKDAAPRPLELVLTGLMGCT
GMDVVSILRKMKVIDQMKDFRIEIEYERTEEHPRIFTKVHLKYIFKFDGEPPKDKVEKAVQL
SQEKYCSVSAILKCSSKVTYEIVYEN
111111111111000000000000000000000000000000000000000000000000
000000000000000000000000000000000000000000000000000000000000
00000000000000000000000001
>DM_train2037
MSSTKRQKNRVLVKLKKRKVRKDKIEKWAELALSALGLNNVELSVYITDDQEIRELNKTYRK
KDKPTDVLSFPMGEEFGGYKILGDVVISQDTAERQARELGHSLEEEVKRLIVHGIVHLLGYD
HEKGGEEEKKFRELENYVLSKLSKAL
111111100000000000000000000000000000000000000000000000000000
000000000000000000000000000000000000000000000000000000000000
00000000000000000000000011
>DM_train2038
ADVSAAVGATGQSGMTYRLGLSWDWDKSWWQTSTGRLTGYWDAGYTYWEGGDEGAGKHSLSF

APVFVYEFAGDSIKPFIEAGIGVAAFSGTRVGDQNLGSSLNFEDRIGAGLKFANGQSVGVRA
IHYSNAGLKQPNDGIESYSLFYKIPI
000000000000000000000000000000000000000000000001111000000000
000000000000000000000000000000000000000000000000000000000000
00000000000000000000000000
>DM_train2039
MTNRLSWQDYFMANAELISKRSTCNRAYVGAVLVKNNRIIATGYNGGVADTDNCDDVGHEME
DGHCIRTVHAEMNALIQCAKEGISANNTEIYVTHFPCINCTKALLQAGVKKITYNTAYRIHP
FAIELMTQKEVEYVQHDVPRVKLGEK
111000000000000000000000000000000000000000000000000000000000
000000000000000000000000000000000000000000000000000000000000
00000000000000000000000001
>DM_train2040
MKDKVYKRPVSILVVIYAQDTKRVLMLQRRDDPDFWQSVTGSVEEGETAPQAAMREVKEEVT
IDVVAEQLTLIDCQRTVEFEIFSHLRHRYAPGVTRNTESWFCLALPHERQIVFTEHLAYKWL
DAPAAAALTKSWSNRQAIEQFVINAA
111110000000000000000000000000000000000000000000000000000000
000000000000000000000000000000000000000000000000000000000000
00000000000000000000000111
>DM_train2041
GQHTLKQFAADSALTTTTPLCSEVPLFDINALGDWTYLGTSLPAKFAKLFASILHCIDDEYF
LITPVEKVRVQVEDAPLLIVDFERAQPHSLLNVSTSIGTLHHNVDIKQMKLTDDSVYLPLER
GLWGKLGRACYYNFVNEFNLSDLNEQ
111111111111111111110000000000000000000000000000000000000000
000000000000000000000000000000000000000000000000000000000000
00000000000000000000000011
>DM_train2042
MDIDPYKEFGATVELLSFLPSDFFPSVRDLLDTASALYREALESPEHCSPHHTALRQAILCW
GELMTLATWVGNNLEDPASRDLVVNYVNTNMGLKIRQLLWFHISCLTFGRETVLEYLVSFGV
WIRTPPAYRPPNAPILSTLPETTVV
000000000000000000000000000000000000000000000000000000000000
000000000000000000000000000000000000000000000000000000000000
0000000000000000001111111
>DM_train2043
MTSKIEQPRWASKDSAAGAASTPDEKIVLEFMDALTSNDAAKLIEYFAEDTMYQNMPLPPAY
GRDAVEQTLAGLFTVMSIDAVETFHIGSSNGLVYTERVDVLRALPTGKSYNLSILGVFQLTE
GKITGWRDYFDLREFEEAVDLPLRG
111100000000000000000000000000000000000000000000000000000000
000000000000000000000000000000000000000000000000000000000000
0000000000000000000000000
>DM_train2044
ATSTKKLHKEPATLIKAIDGDTVKLMYKGQPMTFRLLLVDTPETKHPKKGVEKYGPEASAFT
KKMVENAKKIEVEFDKGQRTDKYGRGLAYIYADGKMVNEALVRQGLAKVAYVYKGNNTHEQL
LRKSEAQAKKEKLNIWSEDNADSGQ

11111000000000000000000000000000000000000000111111000000000000
00000000000000000000000000000000000000000000000000000000000000
0000000000000000011111111
>DM_train2045
MKLERVTVKNFRSHSDTVVEFKEGINLIIGQNGSGKSSLLDAILVGLYWPLRIKDIKKDEFT
KVGARDTYIDLIFEKDGTKYRITRRFLKGYSSGEIHAMKRLVGNEWKHVTEPSSKAISAFME
KLIPYNIFLNAIYIRQGQIDAILES
00000000000000000000000000000000000000000000000000000000000000
00000000000000000000000000001111000000000000000000000000000000
0000000000000000000000000
>DM_train2046
QFLFSMSTGPFICTVKDNQVFVANLPWTMLEGDDIQVGKEFAARVEDCTNVKHDMAPTCTKP
PPFCGPQDMKMFNFVGCSVLGNKLFIDQKYVRDLTAKDHAEVQTFREKIAAFEEQQENQPPS
SGMPHGAVPAGGLSPPPPPSFCTVQ
00000000000000000000000000000000000000000000000000000000000000
00000000000000000000000000000000000000000000000000000001111111
1111111111111000000000000
>DM_train2047
ASPIGEGTFDDGPEGWVAYGTDGPLDTSTGALCVAVPAGSAQYGVGVVLNGVAIEEGTTYTL
RYTATASTDVTVRALVGQNGAPYGTVLDTSPALTSEPRQVTETFTASATYPATPAADDPEGQ
IAFQLGGFSADAWTLCLDDVALDSE
11111110000000000000000000000000000000000000000000000000000000
00000000000000000000000000000000000000000000000000000000000000
0000000000000000000000000
>DM_train2048
ETVSTQELYSIPEDQEPEDQEPEEPTPAPWARLWALQDGFANLECVNDNYWFGRDKSCEYCF
DEPLLKRTDKYRTYSKKHFRIFREVGPKNSYIAYIEDHSGNGTFVNTELVGKGKRRPLNNNS
EIALSLSRNKVFVFFDLTVDDQSVY
11111111111111111111111111110000000000000000000000000000000000
00000000000000000000000000000000000000000000000000000000000000
0000000000000000000011111
>DM_train2050
MNISEINGFEVTGFVVRTTNADEMNPMTAKIGNLWEKFYLNAAPKLTDKSKVYGLYTNYESD
FTGAFDVIACSDTLSPQLLSESVKTKVSSGKYVTFSATGEMPQVVIDLWNEVWNYFASEHCP
HKRAYTTDFEYYKSANTVEISIAVR
00000000000000000000000000000000000000000000000000000000000000
00000000000000000000000000000000000000000000000000000000011100
0000000000000000000000000
>DM_train2051
GAQNEKNMYSYKKIGNKYIVSINNHTEIVKALNAFCKEKGILSGSINGIGAIGELTLRFFNP
KTKAYDDKTFREQMEISNLTGNISSMNEQVYLHLHITVGRSDYSALAGHLLSAIQNGAGEFV
VEDYSERISRTYNPDLGLNIYDFER
11111100000000000000000000000000000000000000000000000000000000
01111000000000000000000000000000000000000000000000000000000000

0000000000000000000000000
>DM_train2052
MATKLDYEDAVFYFVDDDKICSRDSIIDLIDEYITWRNHVIVFNKDITSCGRLYKELMKFDD
VAIRYYGIDKINEIVEAMSEGDHYINFTKVHDQESLFATIGICAKITEHWGYKKISESRFQS
LGNITDLMTDDNINILILFLEKKLN
11111000000000000000000000000000000000000000000000000000000000
00000000000000000001100000000000000000000000000000000000000000
0000000000000000000000000
>DM_train2053
MGIVTGPIIDHSKENDVMNEAEQNAETVRRGYAAFNSGDMKTLTELFDENASWHTPGRSRIA
GDHKGREAIFAQFGRYGGETGGTFKAVLLHVLKSDDGRVIGIHRNTAERGGKRLDVGCCIVF
EFKNGRVIDGREHFYDLYAWDEFWR
11111111111111111100000000000000000000000000000000000000000000
00000000000000000000000000000000000000000000000000000000000000
0000000000000000000000000
>DM_train2054
MPLPPDITFDSLALIKMHSQNMKRILEVTLAKFTVNLSIVTVYRYLTARAYLKKNIEAEFEI
LKDIYNIVPLLDDIAIKAAQIEANLIKKEITLDMEDIITATTAIYTNSLLVTDDPKRYEPIR
RFGLDTMPLDKFIKEVELMVEKELI
10000000000000000000000000000000000000000000000000000000000000
00000000000000000000000000000000000000000000000000000000000000
0000000000000000000001111
>DM_train2055
EARPIVVGPPPPLSGGLPGTENSDQARDGTLPYTKDRFYLQPLPPTEAAQRAKVSASEILNV
KQFIDRKAWPSLQNDLRLRASYLRYDLKTVISAKPKDEKKSLQELTSKLFSSIDNLDHAAKI
KSPTEAEKYYGQTVSNINEVLAKLG
00000000000001111111111111111111100000000000000000000000000000
00000000000000000000000000000000000000000000000000000000000000
0000000000000000000000000
>DM_train2056
MEQRSAETRIVEALLERRRLKDTDLVRARQLQAESGMGLLALLGRLGLVSERDHAETCAEVL
GLPLVDARQLGDTPPEMLPEVQGLSLRFLKQFHLCPVGERDGRLDLWIADPYDDYAIDAVRL
ATGLPLLLHVGLRSEIDDLIERWYG
00000000000000000000000000000011100000000000000000000000000000
00000000000000000110000000000000000000000000000000000000000000
0000000000000000000000000
>DM_train2057
GMEIHFEKVTSDNRKAVENLQVFAEQQAFIESMAENLKESDQFPEWESAGIYDGNQLIGYAM
YGRWQDGRVWLDRFLIDQRFQGQGYGKAACRLLMLKLIEKYQTNKLYLSVYDTNSSAIRLYQ
QLGFVFNGELDTNGERVMEWTHQNK
00000000000000000000000000000000000000000000000000000000000000
00000000000000000000000000000000000000000000000000000000000000
0000000000000000000000011
>DM_train2058

GMTDWQQALDRHVGVGVRTTRDLIRLIQPEDWDKRPISGKRSVYEVAVHLAVLLEADLRIAT
GATADEMAQFYAVPVLPEQLVDRLDQSWQYYQDRLMADFSTETTYWGVTDSTTGWLLEAAVH
LYHHRSQLLDYLNLLGYDIKLDLFE
000000000000000000000000000000000000000000000000000000000000
000000000000000000000000000000000000000000000000000000000000
0000000000000000000000001
>DM_train2059
GMLVRTLSALECTKVLTANRVGRLACAKDGQPYVVPLYYAYSDAHLYAFSMPGKKIEWMRAN
PRVSVQVDEHGQGRGWKSVVVDGRYEELPDLIGHKLQRDHAWSVLSKHTDWWEPGALKPVTP
PTADSAPHVFFRILIEQVSGREASE
000000000000000000000000000000000000000000000000000000000000
000000000000000000000000000000000000000000000000000111111111
1111100000000000000000000
>DM_train2060
STGKKPKHHVLSHDIIPASKPIAEKLQIQPESPVVELKRILYNDDQPLTFEVTHYPLDLFPG
IDTFIADGVSMHDILKQQYKVVPTHNTKLLNVVYAQQEESKYLDCDIGDALFEIDKTAFTSN
DQPIYCSLFLMHTNRVTFTINSPYT
111111100000000000000000000000000000000000000000000000000000
000000000000000000000000000000000000000000000000000000000000
0000000000000000000001111
>DM_train2061
TTSGALFPSLVPGSRGSSTKYLVEFRAGKMSLKGTTVTPDKRKGLVYIQQTDDSLIHFCWKD
RTSGTVEDDLIIFPDDCEFKRVPQCPSGRVYVLKFKAGSKRLFFWMQEPKTDQDEEHCRKVN
ECLNNPPMPGSLGASGSSGHELSAL
111111111111111111110000000000000000000000000000000000000000
000000000000000000000000000000000000000000000000000000000000
0000011111111111111111111
>DM_train2062
DPATVFRLVAAEALTLTGADGTLVAVPADPDASAAEELVIVEVAGAVPAEVEASAIPVQDNA
IGQAFRDRAPRRLDVLDGPGLGGPALVLPLRATDTVAGVLVAVQGSGARPFTAEQLEMMTGF
ADQAAVAWQLASSQRRMSELDILAD
000000000000000000000000000011110000000000000000000000000000
000000000000000000000000000000000000000000000000000000000000
0000000000000000001111111
>DM_train2063
MIPMEELKVKKITNGTVIDHIDAGKALMVFKVLNVPKETSVMIAINVPSKKKGKKDILKIEG
IELKKEDVDKISLISPDVTINIIRNGKVVEKLKPQIPDEIEGTLKCTNPNCITNKEKVRGKF
KIESKNPLKIRCYYCEKFLNEVIFE
111111100000000000000000000000000000000000000000000000000000
000000000000000000000000000000000000000000000000000000000000
0000000000000000000000001
>DM_train2064
TDLSAQKRLAADVLDVGKNRVWFNPERQGDIADAITREDVRELVDEGAIQAKDKKGNSRGRA
RERQKKRAKGHQKGAGSRKGKAGARQNSKEDWESRIRAQRTKLRELRDEGTLSSSQYRDLYD

KAGGGEFDSVADLERYIDANHGDA
0000000000000000000000000000000000000000000000000000000000000000
0000000000000000000000000000000000000000000000000000000000000000
000000000000000000011111
>DM_train2065
KYKALAREAALSKIGELASEIFAEFTEGKYSEVVVRAEENKVRLFVVWEGKERPLTFLSGGE
RIALGLAFRLAMSLYLAGEISLLILDEPTPYLDEERRRKLITIMERYLKKIPQVILVSHDEE
LKDAADHVIRISLENGSSKVEVVS
1111110000000000000000000000000000000000000000000000000000000000
0000000000000000000000000000000000000000000000000000000000000000
000000000000000000000000
>DM_train2066
MANLFKLGAENIFLGRKAATKEEAIRFAGEQLVKGGYVEPEYVQAMLDREKLTPTYLGESIA
VPHGTVEAKDRVLKTGVVFCQYPEGVRFGEEEDDIARLVIGIAARNNEHIQVITSLTNALDD
ESVIERLAHTTSVDEVLELLAGRK
1111000000000000000000000000000000000000000000000000000000000000
0000000000000000000000000000000000000000000000000000000000000000
000000000000000000000111
>DM_train2067
MSNEDLFICIDHVAYACPDADEASKYYQETFGWHELHREENPEQGVVEIMMAPAAKLTEHMT
QVQVMAPLNDESTVAKWLAKHNGRAGLHHMAWRVDDIDAVSATLRERGVQLLYDEPKLGTGG
NRINFMHPKSGKGVLIELTQYPKN
1110000000000000000000000000000000000000000000000000000000000000
0000000000000000000000000000000000000000000000000000000000000000
000000000000000000000000
>DM_train2068
MKVALGGTFEPLHEGHKKLIDVAIKLGGRDITIGVTSDRMARARIRSVLPFAIRAENVKRYV
MRKYGFEPEIVKITNPYGKTLDVDFEYLVVSPETYEMALKINQKREELGKRKITIVKVDWMM
AEDGKPISSTRIKRGEIDRYGGII
1000000000000000000000000000000000000000000000000000000000000000
0000000000000000000000000000000000000000000000000000000000000000
111111111111111111111111
>DM_train2069
DVTPLSLGIETLGGIMTKLITRNTTIPTKKSQVFSTAADGQTQVQIKVFQGEREMATSNKLL
GQFSLVGIPPAPRGVPQVEVTFDIDANGIVNVSARDRGTGKEQQIVIQSSGGLSKDQIENMI
KEAEKNAAEDAKRKELVEVINQAE
0000000000000000000000000000000000000000000000000000000000000000
0000000000000000000000000000000000000000000000001111111111000
000000000000000000000000
>DM_train2070
SPAPSRPFSVLRANDVLWLSLTAAEYDQSTYGSSTGPVYYVSDSVTLVNVATGAQAVARSLDW
TKVTLDGRPLSTIQQHSKTFFVLPLRGKLSFWEAGTTKAGYPYNYNTTASDQLLVENAAGHR
VAISTYTTSLGAGPVSISAVAVLA
1111000000000000000000000000000000000000000000000000000000000000

000000000000000000000000000000000000000000000000000000000000
000000000000000000000000
>DM_train2071
MNYSHPISLKTLVQEDDIGVNAPIIHQSVIARLTAGLYPLYQSKKIPFEPLPETMLTEGYSS
PVPDVLLYDHQTEEAKVIIEVCQNSGLKHDTSKIVKLIEDNAYGILEGFVFNYKTQQWLRYR
LGDGGVATNSSFSEVLQVDLNTFV
111000000000000000000000000000000000000000000000000000000000
000000000000000000000000000000000000000000000000000000000000
000000000000000000000000
>DM_train2072
SMLPVLTLQHFQHMHIKVGDRAELRRAFTQTDVATFSELTGDVNPLHLNEDFAKHTKFGNTI
VHGVLINGLISALLGTKMPGPGCVFLSQEISFPAPLYIGEVVLASAEVKKLKRFIAIIAVSC
SVIESKKTVMEGWVKVMVPEASKS
111111111111100000000000000000000000000000001111111111110000
000000000000000000000000000000000000000000000000000000000000
000000000000000000000001
>DM_train2073
GHMTLYRLHEADLEIPDAWQDQSINIFKLPASGPAREASFVISRDASQGDAPFADYVARQLE
NAEKQLPGFKLHKRWDINIHGHAAVLLDYQWQREGRDLMLRQVFIERRPAVLITTLTTTPAD
LPHHEPAWKQAMQTLVPRPTPSGS
000000000000000000000000000000000000000000000000000000000000
000000000000000000000000000000000000000000000000000000000000
000000000000000000001111
>DM_train2074
RASVGSPGIPAQDLPFVIKAGYLEKRRKDHSFLGFEWQKRWCALSKTVFYYYGSDKDKQQKG
EFAIDGYDVRMNNTLRKDGKKDCCFEICAPDKRIYQFTAASPKDAEEWVQQLKFILQDLGSD
VIPEDDEERGELYDDVDHPAAVSS
000000000000000000000000000000000000000000000000000000000000
000000000000000000000000000000000000000000000000000000011111
111111111111111111111111
>DM_train2075
MKTPTFGKREETLTYQTRYAAYIIVSKPENNTMVLVQAPNGAYFLPGGEIEGTETKEEAIHR
EVLEELGISVEIGCYLGEADEYFYSNHRQTAYYNPGYFYVANTWRQLSEPLERTNTLHWVAP
EEAVRLLKRGSHRWAVEKWLAAAS
100000000000000000000000000000000000000000000000000000000000
000000000000000000000000000000000000000000000000010000000000
000000000000000000000000
>DM_train2076
VSYAGDITPLQAWEMLSDNPRAVLVDVRCEAEWRFVGVPDLSSLGREVVYVEWATSDGTHND
NFLAELRDRIPADADQHERPVIFLCRSGNRSIGAAEVATEAGITPAYNVLDGFEGHLDAEGH
RGATGWRAVGLPWRQGRSHHHHHH
100000000000000000000000000000000000000000000000000000000000
000000000001111110000000000000000000000000000000000000000000
000000000000000011111111

>DM_train2077
GMATSTDRTQEIQKLHELIKNIDYGMFTTVDDDGSLHSYPMSKSGDINSEATLWFFTYAGSH
KVTEIEHHEQVNVSFSSPEQQRYVSISGTSQLVKDRNKMRELWKPELQTWFPKGLDEPDIAL
LKVNINQVNYWDSTSSFKPQTISF
111110000000000000000000000000000000000000000111000000000000
000000000000000000000000000000000000000000000000000000000000
000000000000000000000000
>DM_train2078
GFQSNAMKQETPITLHVKSVIEDDGNQEVIEFRTTGFYYVKQNKVYLSYYEEHDLGKVKTIV
KVSEGEVLVMRSGAVKMNQRFVTGASTIAKYKMSFGELELKTSTKSIQSDLDEEKGRISIAY
DMHVGDEQEHLHNMTITYEGGTHA
000000000000000000000000000000000000000000000000000000000000
000000000000000000000000000000000000000000000000000000000000
000001111000000000000011
>DM_train2079
SNAARGHRHTCKVMVLKEEAAGSERALALDMREGQRVFHSLIVHFENDIPVQIEDRFVNAQV
APDYLKQDFTLQTPYAYLSQVAPLTEGEHVVEAILAEADECKLLQIDAGEPCLLIRRRTWSG
RQPVTAARLIHPGSRHRLEGRFTK
111111000000000000000000000000000000000000000000000000000000
000000000000000000000000000000000000000000000000000000000000
000000000000000000000000
>DM_train2080
GPLGSGGGTIAMLNEISSDTLEQLYSLAFNQYQSGKYEDAHKVFQALCVLDHYDSRFFLGLG
ACRQAMGQYDLAIHSYSYGAVMDIKEPRFPFHAAECLLQKGELAEAESGLFLAQELIANKPE
FKELSTRVSSMLEAIKLKKEMKHE
111110000000000000000000000000000000000000000000000000000000
000000000000000000000000000000000000000000000000000000000000
000000000000000111111111
>DM_train2081
GMSAGQSSLSFAHGDETHPIEQKAFIGVDSAAGNVVKQFHAALQMGNEAIVRQSLAANVQIY
EGGKVERSLTEYANHHMLADMAYLKGLTITPKEHQITITGDIAISTSISHAQGEYKGKSIDS
MTMETLVLIKQADGRWKITHVHWS
111111111111111111111111111000000000000000000000000000000000
000000000000000000000000000000000000000000000000000000000000
000000000000000000000000
>DM_train2082
SFDKTTFRLLRGESNLFYTLAKWENNKLHVELPENIDKQSPTMTLIYDENGQLLWAQRDVPW
LMKMIQPDWLKSNGFHEIEADVNDTSLLLSGDHSIQQQLQEVRQNNNNAQMTHSVAVNVYPA
TSRMPKLTIVVVDTIPVELKSSYM
000000000000000000000000000000000000000000000000000000000000
000000000000000000000000000000000000000000000000000000000000
000000000000000000000011
>DM_train2083
AHGSVVKSEDYALPSYVDRRDYPLPDVAHVKNLSASQKALKEKEKASWSSLSIDEKVELYRL

KFKESFAEMNRSTNEWKTVVGAAMFFIGFTALLLIWEKHYVYGPIPHTFEEEWVAKQTKRML
DMKVAPIQGFSAKWDYDKNEWKK
11100000000000000000000000000000000000000000000000000000000000
00000000000000000000000000000000000000000000000000000000000000
00000000000000000000000
>DM_train2085
GPHMTTIEDEDTKVRKQEIIKVTEQLIEAISNGDFESYTKMCDPGMTAFEPEALGNLVEGLD
FHRFYFENLWSRNSKPVHTTILNPHIHLMGDESACIAYIRITQYLDAGGIPRTAQSEETRVW
HRRDGKWQIVHFHRSGAPSVLPH
11111111100000000000000000000000000000000000000000000000000000
00000000000000000000000000000000000000000000000000000000000000
00000000000000000001111
>DM_train2086
ADLTSHHHHHHVEPARITLTYKEGAPITIMDNGNIDTELLVGTLTLGGYKTGTTSTSVNFTD
AAGDPMYLTFTSQDGNNHQFTTKVIGKDSRDFDISPKVNGENLVGDDVVLATGSQDFFVRSI
GSKGGKLAAGKYTDAVTVTVSNQ
11111111111110000000000000000000000000000000000000000000000000
00000000000000000000000000000000000000000011110000000000000000
00000000000000000000000
>DM_train2087
MRQDAEVIMKLTDADNAADGIFFPALEQNMMGAVLINENDEVMFFNPAAEKLWGYKREEVIG
NNIDMLIPRDLRPAHPEYIRHNREGGKARVEGMSRELQLEKKDGSKIWTRFALSKVSAEGKV
YYLALVRDASVEMAQKEQTRQLI
11111111111111111110000000000000000000000000000000000000000000
00000000000000000000000011111111100000000000000000000000000000
00000000111111111111111
>DM_train2088
MTPGKLYTLRYEVEGGGFIEIATVRPETVFADQAIAVHPEDERYRHLLGKRARIPLTEVWIP
ILADPAVEKDFGTGALKVTPAHDPLDYEIGERHGLKPVSVINLEGRMEGERVPEALRGLDRF
EARRKAVELFREAGHLVKEEDYT
11100000000000000000000000000000000000000000000000000000000000
00000000000000000000000000000000000000000000000000000000000000
00000000000000000000001
>DM_train2089
TLHKERRIGRLSVLLLLNEAEESTQVEELERDGWKVCLGKVGSMDAHKVIAAIETASKKSGV
IQSEGYRESHALYHATMEALHGVTRGEMLLGSLLRTVGLRFAVLRGNPYESEAEGDWIAVSL
YGTIGAPIKGLEHETFGVGINHI
00000000000000000011110000000000000000000000000000000000000000
00000000000000000000000000000000000000000000000000000000000000
00000000000000000000000
>DM_train2090
AFMSQMMQMYQQVGPAQFSAMIGQFAPYFASIAPQFVELRPGYAEVTFPKRREVLNHIGTVH
AIALCNAAELAAGTMTDASIPAGHRWIPRGMTVEYLAKATGDVRAVADGSQIDWQATGNLVV
PVVAYVDDKPVFRAEITMYVSQA

1100000000000000000000000000000000000000000000000000000000000
0000000000000000000000000000000000000000000000000000000000000
00000000000000000000000
>DM_train2091
MSMPRYYEYRHVVGFEETNLVGNVYYVNYLRWQGRCREMFLYEHAPEILDELRADLKLFTLK
AECEFFAELAPFDRLAVRMRLVELTQTQMELGFDYLRLGGDDLLVARGRQRIACMRGPNGRT
EPVRVPAGLVRAFAPFRSATVGQ
1111000000000000000000000000000000000000000000000000000000000
0000000000000000000000000000000000000000000000000000000000000
00000000000000000011111
>DM_train2092
MRSLLILVLCFLPLAALGKVFGRCELAAAMKRHGLDNYRGYSLGNWVCAAKFESNFNTQATN
RNTDGSTDYGILQINSRWWCNDGRTPGSRNLCNIPCSALLSSDITASVNCAKKIVSDGNGMN
AWVAWRNRCKGTDVQAWIRGCRL
1111111111111111110000000000000000000000000000000000000000000
0000000000000000000000000000000000000000000000000000000000000
00000000000000000000000
>DM_train2093
MSNTLFDDIFQVSEVDPGRYNKVCRIEAASTTQDQCKLTLDINVELFPVAAQDSLTVTIASS
LNLEDTPANDSSATRSWRPPQAGDRSLADDYDYVMYGTAYKFEEVSKDLIAVYYSFGGLLMR
LEGNYRNLNNLKQENAYLLIRR
0000000000000000000000000000000000000000000000000000000000000
0111111111111000000000000000000000000000000000000000000000000
0000000000000000000000
>DM_train2095
ICYDAFVSYSERDAYWVENLMVQELENFNPPFKLCLHKRDFIPGKWIIDNIIDSIEKSHKTV
FVLSENFVKSEWSKYELDFSHFRLFDENNDAAILILLEPIEKKAIPQRFCKLRKIMNTKTYL
EWPMDEAQREGFWVNLRAAIKS
0000000000000000000000000000000000000000000000000000000000000
0000000000000000000111111111110000000000000000000000000000000
0000000000000000000000
>DM_train2096
VPIQKVQDDTKTLIKTIVTRINDISHTQSVSSKQKVTGLDFIPGLHPILTLSKMDQTLAVYQ
QILTSMPSRNVIQISNDLENLRDLLHVLAFSKSCHLPEASGLETLDSLGGVLEASGYSTEVV
ALSRLQGSLQDMLWQLDLSPGC
0000000000000000000000001111111111111100000000000000000000000
0000000000000000000000000000000000000000000000000000000000000
0000000000000000000000
>DM_train2097
MLDQQTINIIKATVPVLKEHGVTITTTFYKNLFAKHPEVRPLFDMGRQESLEQPKALAMTVL
AAAQNIENLPAILPAVKKIAVKHCQAGVAAAHYPIVGQELLGAIKEVLGDAATDDILDAWGK
AYGVIADVFIQVEADLYAQAVE
0000000000000000000000000000000000000000000111111111000000000
0000000000000000000000000000000000000000000000000000000000000

0000000000000000000000
>DM_train2099
MAESQALSDDIGFLLSRVGGMVLGAVNKALVPTGLRVRSYSVLVLACEQAEGVNQRGVAATM
GLDPSQIVGLVDELEERGLVVRTLDPSDRRNKLIAATEEGRRLRDDAKARVDAAHGRYFEGI
PDTVVNQMRDTLQSIAFPTFVE
000000000000000000000000000000000000000000000000000000000000
000000000000000000000111111111110000000000000000000000000000
0000000000000000001111
>DM_train2100
GETVVRDAVTIGKPAEQLYAVWRDLPGLPLLMTHLRSVEVLDDKRSRWTVEAPAPLGTVSWE
AELTADEPGKRIAWRSLPGARIENSGEVLFRPAPGARGTEVVVRLTYRPPGGSAGAVIARMF
NQEPSQQLRDDLMRFKREQELG
110000000000000000000000000000000000000000000000000000000000
000000000000000000000000000000000000000000000011111111111111
1100000000000000000000
>DM_train2101
GGEGTDAIQALIQAYFTAWNTNAPERFAEIFWPDGSWVNVVGMHWRGRDQIVFAHTAFLKTI
FKDCKQELVTIEARTIAPGSALAVVTLIQDAYVTPDGRQMPRAHDRLTLLAVEREGVWRFIH
GHNTIVNPDAANNDPVLRMKPA
100000000000000000000000000000000000000000000000000000000000
000000000000000000000000000000000000000000000000000000000000
0000000000000000000111
>DM_train2102
STAADLLRQGAACSVLYLTSVETESLTGPQAVARASSAALSCSPRPTPAVVHFKVSAQGITL
TDNQRKLFFRRHYPVNSITFSSTDPQDRRWTNPDGTTSKIFGFVAKKPGSPWENVCHLFAEL
DPDQPAGAIVTFITKVMLGQRK
000000000000000000000000000000000000000000111000000000000000
000000000000000000000000000000000000000000000000000000000000
0000000000000000000111
>DM_train2103
GQDRSPIETQNVETVLRLFDEGWGAQDGWRDVWRETMTPGFRSIFHSNQAVEGIEQAIAFNA
VLFEGFPRLEVVVENVTVEGDNVVVQARLTGAQDGPFLGVPPSGQMVDVPDVTLFTLADGQV
IEMRYFTDLLAVMTAISAPPEN
000000000000000000000000000000000000000000000000000000000000
000000000000000000000000000000000000000000000000000000000000
0000000000000000000011
>DM_train2104
MSSERTFLPNGNYKIKSLFSDSLYLTYSSGSLSFLNTSSLDNQKWKLEYISSSNGFRFSNVA
EPNKYLAYNDYGFIYLSSSSNNSLWNPIKIAINSYIICTLSIVNVTDYAWTIYDNNNNITDQ
PILNLPNFDINNSNQILKLEKL
111000000000000000000000000000000000000000000000000000000000
000000000000000000000000000000000000000000000000000000000000
0000000000000000000000
>DM_train2105

MVKRTSHKQASCPVARPLDVIGDGWSMLIVRDAFEGLTRFGEFQKSLGLAKNILAARLRNLV
EHGVMVAVPAESGSHQEYRLTDKGRALFPLLVAIRQWGEDYFFAPDESHVRLVERDSGQPVP
RLQVRAGDGSPLAAEDTRVSRD
111110000000000000000000000000000000000000000000000000000000
000000000000000000000000000000000000000000000000000000000000
0000000000000000000001
>DM_train2106
GHMAQTDKHYFGTLLAQTSRAWRAELDRRLSHLGLSQARWLVLLHLARHRDSPTQRELAQSV
GVEGPTLARLLDGLESQGLVRRLAVAEDRRAKHIVLTPKADVLIADIEAIAASVRNDVLTGI
DESEQALCQQVLLRILANLENR
111111111000000000000000000000000000000000000000000000000000
000000000000000000000000000000000000000000000000000000000000
0000000000000000000000
>DM_train2107
MAQERPSCAVEPEHVQRLLLSSREAKKSAYCPYSRFPVGAALLTGDGRIFSGCNIENACYPL
GVCAERTAIQKAISEGYKDFRAIAISSDLQEEFISPCGACRQVMREFGTDWAVYMTKPDGTF
VVRTVQELLPASFGPEDLQKIQ
111111111100000000000000000000000000000000000000000000000000
000000000000000000000000000000000000000000000000000000000000
0000000000000000000011
>DM_train2108
MIEVKPINAEDTYELRHRILRPNQPIEACMFESDLLRGAFHLGGYYGGKLISIASFHQAEHS
ELQGQKQYQLRGMATLEGYREQKAGSSLIKHAEEILRKRGADLLWCNARTSASGYYKKLGFS
EQGEVFDTPPVGPHILMYKRIT
100000000000000000000000000000000000000000000000000000000000
000000000000000000000000000000000000000000000000000000000000
0000000000000000000000
>DM_train2109
SNAIYGYVEKATLIDQNLTLSAKLDTGAKSASLHAVNITEIEKKGIPYLRFTVPTKTGDYSF
EGEYVGKVKIKVRSSETNPGLLRTTPIKRPVVLLNIKLGDKVRTIKVNLTNRKRFLYPLLLG
RDAIIDFNGAVDPALTFTTKSK
000000000000000000000000000000000000000000000000000000000000
000000001111111111111111100000000000000000000000000000000000
0000000000000000000011
>DM_train2110
MESVADIQQLEEKVDESDVRIYFNEKSSGGKISIDNASYNARKLGLAPSSIDEKKIKELYGD
NLTYEQYLEYLSICVHDKDNVEELIKMFAHFDNNCTGYLTKSQMKNILTTWGDALTDQEAID
ALNAFSSEDNIDYKLFCEDILQ
110000000000000000000000000000000000000000000000000000000000
000000000000000000000000000000000000000000000000000000000000
0000000000000000000000
>DM_train2111
MSHDVKELIEGLNEDLAGEYSAIIMYNHNAATVSGIYRQVLKPFFESEISDEQGHALYLAEK
IKTLGGTPTTIPLRVKQAEDVREMLEYARQSEYETIKRYEKRKEQAANLNMTELVVKLEDMI

ADETNHMEELDRLLNDKAMVLN
000000000000000000000000000000000000000000000000000000000000
000000000000000000000000000000000000000000000000000000000000
0000000000000001111111
>DM_train2113
AREGIIGHYIHHNQRVGVLVELNCETDFVARNELFQNLAKDLAMHIAMMNPRYVSAEEIPAE
ELEKERQIYIQAALNEGKPQQIAEKIAEGRLKKYLEEVVLLEQPFVKDDKVKVKELIQQAIA
KIGENIVVRRFCRFELGAMMC
000000000000000000000000000000000000000000000000000000000000
000000000000000000000000000000000000000000000000000000000000
000000000000000000111
>DM_train2114
MMGIMSDPITILDSSDSLSRLSSESVGRLVVHRKDDLDIFPVNFVLDYSAEQPRVYFRTAEG
TKLFSVNLNSDVLFEVDRFDDAEGWSVVLKGNAYVVRDTEEARHADTLGLKPWLPTLKYNFV
RIDVREVSGRAFVFGEEPERY
111111000000000000000000000000000000000000000000000000000011
000000000000000000001100000000000000000000000000000000000000
000000000000011111111
>DM_train2115
MRNKNMSTIYQTSATASAGRNGVVSTEDKLLELNLSYPKEMGGSGTATNPEQLFAVGYAACF
SNAILHVAREAKVALKEAPVTATVGIGPNGQGGFALSVALAAHIALEDEQARQLVTVAHQVC
PYSNAVRGNIDVQVSVNGLAL
111110000000000000000000000000000000000000000000000000000000
000000000000000000000000000000000000000000000000000000000000
000000000000000000000
>DM_train2116
MSNPQPEKVRVVVGDDHPLFREGVVRALSLSGSVNVVGEADDGAAALELIKAHLPDVALLDY
RMPGMDGAQVAAAVRSYELPTRVLLISAHDEPAIVYQALQQGAAGFLLKDSTRTEIVKAVLD
CAKGRDVVAPSLVGGLAGEIR
111111111110000000000000000000000000000011111111111100000000
000111111111111111111000000001111111111000000000000000000000
111111111111111111111
>DM_train2117
KPQIPKDKSKVAGYIEIPDADIKEPVYPGPATPEQLNRGVSFAEENESLDDQNISIAGHTFI
DRPNYQFTNLKAAKKGSMVYFKVGNETRKYKMTSIRDVKPTDVGVLDEQKGKDKQLTLITAD
DYNEKTGVWEKRKIFVATEVK
111000000000000000000000000000000000000000000000000000000000
000000000000000000000000000000000000000001111111111000000000
000000000000000000000
>DM_train2118
MDELLNRLRQTWHSTIPVSEFMQIAPLSFTDGELSVSAPLAPNINLHHTMFAGSIYTIMTLT
GWGMVWLQQQLLNVDGDIVLADAHIRYLAPVTSAPEVKVRWPDTNLSPLQRGRKAKVKLEVQ
LFCDGKLCAQFDGLYVSVPKM
000000000000000000000000000000000000000000000000000000000000

000000000000000000000000000000000000000011111100000000000000
000000000000000000011
>DM_train2119
GSMKQPFERILREICFMVKVEGRKVLRDFGITPAQFDILQKIYFEGPKRPGELSVLLGVAKS
TVTGLVKRLEADGYLTRTPDPADRRAYFLVITRKGEEVIEKVIERRENFIEKITSDLGKEKS
SKILDYLKELKGVMERNFSKQ
111000000000000000000000000000000000000000000000000000000000
000000000000000000011110000000000000000000000000000000000000
000000000000000000001
>DM_train2120
ERVPEGSIRVAIASNNGEQLDGHFGSCLRFLVYQVSAKDASLVDIRSTLDVALAEDKNAWRV
EQIQDCQVLYVVSIGGPAAAKVVRAGIHPLKKPKGCAAQEAIAELQTVMAGSPPPWLAKLVG
VSAEERVRFSVSDDEDEAARA
000000000000000000000000000000000000000000000000000000000000
000000000000000000000000000000000000000000000000000000000001
111111111111111111111
>DM_train2121
MAAHELKEALETLKETGVRITPQRHAILEYLVNSMAHPTADDIYKALEGKFPNMSVATVYNN
LRVFRESGLVKELTYGDASSRFDFVTSDHYHAICENCGKIVDFHYPGLDEVEQLAAHVTGFK
VSHHRLEIYGVCQECSKKENH
111000000000000000000000000000000000000000000000000000000000
000000000000000000000000000000000000000000000000000000000000
000000000000000000001
>DM_train2122
MSMAERKALCLEMVAAWNRWDLSGIIKHWSPDIVHYSEDNEVSSADMVKLMEGGLKAFPDLQ
LEVKSIMAEEDRVALRITVTATHQGEFMGVQPTGQRVSWHLVEELRFVDGKVVEHWDVINMR
PLLVRLGKLPDVPKVVLEASA
100000000000000000000000000000000000000000000000000000000000
000000000000000000000000000000000000000000000000000000000000
000000000001111111111
>DM_train2123
GSHHPFTAQIVAVTASGYDSEKGHVPANIADGDVKTRWAASGESWVQLELDKEQSIENILIV
PFKPTERKLKFSIFYSNDGKNWQPLAEGLETSSADKNGEKLTFTPVTAKYIKLDTFGTDVNN
WSAINEIAINSAAALPSRAIK
111111000000000000000000000000000000000000000000000000000000
000000000000000000000000000000000000000000000000000000000000
000000000000000000000
>DM_train2124
ATSANTDRIATAELGIAENKKDAQIAKAQANENKDGIAKNQADIQLHDKKITNLGILHSMVA
RAVGNNTQGVATNKADIAKNQADIANNIKNIYELAQQQDQHSSDIKTLAKVSAANTDRIAKN
KAEADASFETLTKNQKLCSQA
111100000000000000000000000000000000000000000000000000000000
000000000000000000000000000000000000000000000000000000000000
000000000000000011111

>DM_train2125
MDVQRVGKAGLHRVDSKKQQTAAGVSFSEVMGKQRDEKAYERLQALMSKIDDQGKLLSETRT
IEELRKYKELVKEFVGDAVELGLRLEERRGFNRRGRTKIYKIVKEVDRKLLDLTDAVLAKEK
KGLDILNMVGEIKGLLINIYA
111111111111111111111100000000000000000000000000000000000000
000000000000000000000000000111000000000000000000000000000000
000000000000000000000
>DM_train2126
MRGSHHHHHHGSMSNINLVQLVRDSLFTIGCPPSIITDLDSHSAITISLDSMPAINIALVNE
QVMLWANFDAPSDVKLQSSAYNILNLMLMNFSYSINELVELHRSDEYLQLRVVIKDDYVHDG
IVFAEILHEFYQRMEILNGVL
111111111111000000000000000000000000000000000000000000000000
000000000000000000000000000000000000000000000000000000000000
000000000000000000000
>DM_train2127
RPRGPRAVFILPVTAQGEAVLIRQFRYPLRATITEIVAGGVEKGEDLGAAAARELLEEVGGA
ASEWVPLPGFYPQPSISGVVFYPLLALGVTLGAAQLEDTETIERVVLPLAEVYRMLEAGEIQ
DGPSSLTLWQARGELTRRGLL
111100000000000000000000000000000000000000000000000000000000
000000000000000000000000000000111111111100000000000000000000
000000000000000000000
>DM_train2128
GSHMPLPIPSLLIAGIGCRRGCSAEHLRALLERTLGEHGRSLAELDALASIDGKRDEPGLRQ
LATLLERPVHFLAPAVLHDYEPRLLSPSAVALRETGCSSVAEAAALALAERLGGGRADLLGA
KRSDDRASIALARLLTERELP
000000000000000000000000000000000000000000000000000000000000
000000000000000000000000000000000000000000000000000000000000
000000000000000000111
>DM_train2129
MSATGLEVFDRTLHKTHAWLKAIMEELGTEDRHKAYLALRAVLHALRDRLTVEEVAQLAAQL
PMLVRGLYYEGWDPTGKPLKERHKEAFLAHVAEELKTPSGPAVDPEAATRAVFKVLSREISQ
GELEDVLGLLPKELRALWPQG
100000000000000000000000000000000000000000000000000000000000
000000000000000000000000000000000000000000000000000000000000
000000000000000000000
>DM_train2130
GMDISLLKQVVQSTNKIALSTAVNNEADVKIVNFVWYEAQPDTLYFSSVKTSPALKVYDQNP
DIAFITIPNDGTAGNPYLRAQHVKLQRSTKTMTDLLPQYLETVPNYQQVWDAIGSTLVVFEL
KLTDLFVDAGVGGEKQTLTFN
100000000000000000000000000000000000000000000000000000000000
000000000000000000000000000000000000000000000000000000000000
000000000000000000001
>DM_train2131
STARMQGAGKALHELLLSAQRQGCLTAGVYESAKVLNVDPDNVTFCVLAADEEDEGDIALQI

HFTLIQAFCCENDIDIVRVGDVQRLAAIVGSDEEGGAPGDLHCILISNPNEDTWKDPALEKL
SLFCEESRSFNDWVPSITLPE
111000000000000000000000000000000000000000000000000000000000
000000000000000000000000000001111111100000000001111100000000
000000000000000000000
>DM_train2132
GMEVIESKWYKKDGASSASIDDVEKLLNTTLPKQYKSFLLWSNGGEGKLGDNYIYIWAIEDV
IAYNHDYGIQKYLQKEYWAFGMDGDIGYILHLSDNSIYRVDLGDLDITSIKYIAPSFDDFLG
KAIYLNFNKLQNVANNNLTT
000000000000000000000000000000000000000000000000000000000000
000000000000000000000000000000000000000000000000000000000000
00000000011111111111
>DM_train2133
MYFVDRSKIEKTLGFFEHQLALFDSQTDWQSEIGELALQRIGHLLIECILDTGNDMIDGFIM
RDPGSYDDIMDILVDEKVVTEKEGDELKKLIAYRKTLVQQYLLADSGELYRLIKAHQTALQD
FPKRIRSYLETELGPVSAFK
000000000000000000000000000000000000000000000000000000000000
000000000000000000000000000000000000000000000000000000000000
00000000000000000001
>DM_train2134
AGSLEIRTKVGEICISKVWLTDEQINKLFDRFKGDYQVVNAECADKVIFATIIAIKAVKEGR
SIAKTVPGEILVRLSGNRQIKEAIKKVGAKEGENYIVTFGENASALLQKILSTLEIKELELE
RCDLEYAKKAFEDIAIIEAL
100000000000000000000000000000000000000000000000000000000000
000000000000000000000000000000000000000000000000000000000000
00000000000000011111
>DM_train2135
GMDTNGVLYAANMTNALAKEIPESKWDIQLIPELGTLRKLFIHIVRVRDVYRDGLKTGSIKF
PGRLASDEHRLLDELERSMEELVFEFKQTTFNSIKMGENYLSIMELLGTVIQHEGIHQGQYY
VALKQSGINLPKQWVQDWHM
000000000000000000000000000000000000000000000000000000000000
000000000000000000000000000000000000000000000000000000000000
00000000000000000011
>DM_train2136
MTAVRRIRAAALPDLPDASWSNALLVGEELVMSGMTAHPATRQAAERGAALDAHAQALVVLG
KVKALLEAAGGHVGNLYKLNVYVTRIADKDAIGRARQEFFAGQGTFPASTLVEVSGLVFPEL
LVEIDAWARLDIDLANCDEA
000000000000000000000000000000000000000000000000000000000000
000000000000000000000000000000000000000000000000000000000000
00000000000000000011
>DM_train2137
RETHFLKIPIGDLNIITQDNMKSCQMTTPVIDVIQMLTQGRVSSVPIIDENGYLINVYEAYD
VLGLIKGGIYNDLSLSVGEALMRRSDDFEGVYTCTKNDKLSTIMDNIRKARVHRFFVVDDVG
RLVGVLTLSDILKYILLGSN

110000000000000000000000000000000000000000000000000000000000
000000001111000000000000011111100000000000000000000000000000
00000000000000000011
>DM_train2138
SNAMEIRVFRQEDFEEVITLWERCDLLRPWNDPEMDIERKMNHDVSLFLVAEVNGEVVGTVM
GGYDGHRGSAYYLGVHPEFRGRGIANALLNRLEKKLIARGCPKIQINVPEDNDMVLGMYERL
GYEHADVLSLGKRLIEDEEY
111000000000000000000000000000000000000000000000000000000000
000000000000000000000000000000000000000000000000000000000000
00000000000000000111
>DM_train2139
ASRFHEQFIVREDLMGLAIGTHGANIQQARKVPGVTAIDLDEDTCTFHIYGEDQDAVKKARS
FLEFAEDVIQVPRNLVGKVIGKNGKLIQEIVDKSGVVRVRIEAENEKNVPQEEGMVPFVFVG
TKDSIANATVLLDYHLNYLK
111000000000000000000000000000000000000000000000000000000000
000000000000000000000000000000000000000000000000000000000000
00000000000000000000
>DM_train2140
GMELTALDKLEIMELAARFEMSLDKEDVENYLATFASDGALQGFWGIAKGKEELRQGFYAML
DTFARGKRHCSSNAIIQGNYDEATMESYLTVVNREDLNRAGSAFVKDQVRKINGKWYLILRQ
IEVDPSLPLLQQSQQAGANA
100000000000000000000000000000000000000000000000000000000000
000000000000000000000000000000000000000000000000000000000000
00000000001111111111
>DM_train2141
MLYLTQRLEIPAAATASVTLPIDVRVKSRVKVTLNDGRDAGLLLPRGLLLRGGDVLSNEEGT
EFVQVIAADEEVSVVRCDDPFMLAKACYHLGNRHVPLQIMPGELRYHADHVLDDMLRQFGLT
VTFGQLPFEPEAGAYASES
000000000000000000000000000000000000000000000000000000000000
000000000000000000000000000000000000000000000000000000000000
0000000000000111111
>DM_train2142
MQEGVKVVVDSRELRSEVVKRLKLLGVKLEVKTLDVGDYIISEDVAIERKSANDLIQSIIDG
GLFDQVKRLKEAYSRPIMIVEGSLYGIRNVHPNAIRGAIAAVTVDFGVPIIFSSTPEETAQY
IFLIAKREQEEREKPVRIR
111100000000000000000000000000000000000000000000000000000000
000000000000000000000000000000000000000000000000000000000000
0000000000011111111
>DM_train2143
MSGDRDDPRYPYPKDDAELRRRLTPMQYEVTQHAATEPPFTGEYTDTEDAGIYHCVVCGTAL
FESGAKYHSGCGWPSYFKPIDGEVIDEKMDYTHGMTRVEVRCNQCGAHLGHVFEDGPRDKTG
LRYCINSAALNFEAKPERK
111111111000000000000000000000000000000000000000000000000000
000000000000000000000000000000000000000000000000000000000000

0000000000000000111
>DM_train2144
MTEHSLWRFSRALHRALNDRQTEELATIIDDNIDWAIYGPIDMFPFFGARQGKAAVLEVCRQ
IADSVRIYRYHRESVMLGIDSAASMVRYSLTAAGTNRPISVRMALFTQFQNGRLTNLRMVLD
TFDLVEQALGRPIHLPKIA
000000000000000000000000000000000000000000000000000000000000
000000000000000000000000000000000000000000000000000000000000
0000000000011111111
>DM_train2145
MRKIDLCLSSEGSEVILATSSDEKHPPENIIDGNPETFWTTTGMFPQEFIICFHKHVRIERL
VIQSYFVQTLKIEKSTSKEPVDFEQWIEKDLVHTEGQLQNEEIVAHGSATYLRFIIVSAFDH
FASVHSVSAEGTVVSNLSS
111000000000000000000000000000000000000000000000000000000000
000000000000000000000000000000000000000000000000000000000000
0000000000000001111
>DM_train2146
MGSDKIHHHHHHMKLSDFIKTEDFKKEKHVPVIEAPEKVKKDEKVQIVVTVGKEIPHPNTTE
HHIRWIKVFFQPDGDPYVYEVGRYEFNAHGESVQGPNIGAVYTEPTVTTVVKLNRSGTIIAL
SYCNIHGLWESSQKITVEE
111111111110000000000000000000000000000000000000000000000000
000000000000000000000000000000000000000000000000000000000000
0000000000000000000
>DM_train2147
MTIHKKGQAHWESDIKRGKGTVSTESGVLNQQPYGFNTRFEGEKGTNPEELIGAAHAACFSM
ALSLMLGEAGFTPTSIDTTADVSLDKVDAGFAITKIALKSEVAVPGIDASTFDGIIQKAKAG
CPVSQVLKAEITLDYQLKS
100000000000000000000000000000000000000000000000000000000000
000000000000000000000000000000000000000000000000000000000000
0000000000000000000
>DM_train2148
GMQDERWNHPLYTTTAINDEELEGHAYIPGGLKVQTSSPMNDHPGTNPEQLLGLSLSTCLEA
TLEAVEKEHGLPHTGAVRVKVAFIGARAEYQFLVHAQVMVKGVDFDTAKAFTNEIENRCPVS
KLLKNSGNYTIETVTDFKD
110000000000000000000000000000000000000000000000000000000000
000000000000000000000000000000000000000000000000000000000000
0000000000000000000
>DM_train2150
MMLLPKRTKFRKQFRGRMTGDAKGGDYVAFGDYGLIAMEPAWIKSNQIEACRIVMSRHFRRG
GKIYIRIFPDKPVTKKPAETRMGKGKGAVEYWVSVVKPGRVMFEVAGVTEEQAKEAFRLAGH
KLPIQTKMVKREVYDEAQ
111111100000000000000000000000000000000000000000000000000000
000000000000000000000000000000000000000000000000000000000000
000000011111111111
>DM_train2151

MSHQTGIQASEDVKEIFARARNGKYRLLKISIENEQLVVGSCSPPSDSWEQDYDSFVLPLLE
DKQPCYVLFRLDSQNAQGYEWIFIAWSPDHSHVRQKMLYAATRATLKKEFGGGHIKDEVFGT
VKEDVSLHGYKKYLLSQS
111111000000000000000000000000000000000000000000000000000000
000000000000000000000000000000000000000000000000000000000000
000000000000000111
>DM_train2152
GPSDMYVHVGNLIYRNLHLFNSEMHESILVSYSSDLIIYRTNTVGDDYIPSCDCTQATYYCK
HKNRYFPITVTSHDWYEIQESEYYPKHIQYNLLIGEGPCEPGDCGGKLLCKHGVIGIVTAGG
DNHVAFIDLRHFHCAEEQ
000000000000000000000000000000000000000000000000000000000000
000000000000000000000000000000000000000000000000000000000000
000000000000000111
>DM_train2153
GSHMGQGGSNPKFENIAEGLRALLARSHVERTTDEGTWVAGVFVYGGSKTSLYNLRRGTALA
IPQCRLTPLSRLPFGMAPGPGPQPGPLRESIVCYFMVFLQTHIFAEVLKDAIKDLVMTKPAP
TCNIRVTVCSFDDGVDLP
111111111100000000000000000000000000000000000000000000000000
000000000000000000000000000000000000000000000000000000000000
000000000000000000
>DM_train2154
AESVQPLEKIAPYPQAEKGMKRQVIQLTPQEDESTLKVELLIGQTLEVDCNLHRLGGKLENK
TLEGAAAAYYVFDKVSSPVSTRMACPDGKKEKKFVTAYLGDAGMLRYNSKLPIVVYTPDNVD
VKYRVWKAEEKIDNAVVR
111111111110000000000000000000000000000000000000000000000000
011111100000000000000000000000000000000000000000000000000000
000000000000000000
>DM_train2155
MIRVMATGVFDILHLGHIHYLKESKKLGDELVVVVARDSTARNNGKIPIFDENSRLALISEL
KVVDRAILGHEGDMMKTVIEVKPDIITLGYDQKFDEAELQSKINKLGITVKIVRISKYDGQL
NSSSSVRKKIMELIGERY
000000000000000000000000000000000000000000000000000000000000
000000000000000000000000000000000000000000000000000000000111
111111111111111111
>DM_train2156
SDRPAGRMPLTVHRNVGRWLSEILHASIRDTGVSSRIEFVRRTLHGWVREEYSETELPNAVY
RNLYFPVLDAQPAHAGSGKIETISECDRLKNLVRNVTDTLVENYPQGLESEALLIALDGVKL
ELARIRKDIEMYGDPRKR
111000000000000000000000000000000000000000000000000000000000
000000011111111110000000000000000000000000000000000000000000
000000000000000111
>DM_train2157
MGSDKIHHHHHHPPEAYSLDTAIFVLETRDYRLSDVKEIDSYGDVEMKGKVAVFETEYGPVF
LYVYKGEEAKKIWKKLNGRAGFVSIRSVLDLPNMGKFSTVSNGKKIVAWWRKNWLFIVEGKN

GVEEFVKHVYRVYEEMKQ
111100000000000000000000000000000000000000000000000000000000
000000000000000000011100000000000000000000000000000000000000
000000000000000000
>DM_train2158
ATEATNATNNQSTQVSQATSQPINFQVQKDGSSEKSHMDDYMQHPGKVIKQNNKYYFQTVLN
NASFWKEYKFYNANNQELATTVVNDNKKADTRTINVAVEPGYKSLTTKVHIVVPQINYNHRY
TTHLEFEKAIPTLADAAK
111111111111111100000000000000000000000000000000000000000000
000000000000000000000000000000000000000000000000000000000000
000000000000001111
>DM_train2159
DNGCSVAAESTNFTVDLMENAAKQFNNIGATTPVVPFRILLSPCGNAVSAVKVGFTGVADSH
NANLLALENTVSAASGLGIQLLNEQQNQIPLNAPSSALSWTTLTPGKPNTLNFYARLMATQV
PVTAGHINATATFTLEYQ
000000000000011111111111111000000000000000000000000000000000
000000000000000000000000000000000000000000000000000000000000
000000000000000000
>DM_train2160
EGICRNRVTNNVKDVTKLVANLPKDYMITLKYVPGMDVLPSHCWISEMVVQLSDSLTDLLDK
FSNISEGLSNYSIIDKLVNIVDDLVECVKENSSKDLKKSFKSPEPRLFTPEEFFRIFNRSID
AFKDFVVASETSDCVVS
111111110000000000000000000000000000000000000000000000000000
000000000000000000000000000000000000000000000000000000000000
00000000111111111
>DM_train2161
FDDVVVSRQEQSYVQRGMVNFLDEEMHKLVKRFRDMRWNLGPGFVFLLKKVNRERMMRYCMD
YARYSKKILQLKHLPVNKKTLTKMGRFVGYRNYGVIRELYADVFRDVQGFRGPKMTAAMRKY
SSKDPGTFPCKNEKRRG
000000000000000000000000000000000000000000000000000000000000
000000000000000000000000000000000000000000000000000000000000
00000000000001111
>DM_train2162
MALEPIDYTTHSREIDAEYLKIVRGSDPDTTWLIISPNAKKEYEPESTGSSFHDFLQLFDET
KVQYGLARVSPPGSDVEKIIIIGWCPDSAPLKTRASFAANFAAVANNLFKGYHVQVTARDED
DLDENELLMKISNAAGA
111100000000000000000000000000000000000000000000000000000000
000000000000000000000000000000000000000000000000000000000000
00000000000011111
>DM_train2163
MPRVVPDQRSKFENEEFFRKLSRECEIKYTGFRDRPHEERQTRFQNACRDGRSEIAFVATGT
NLSLQFFPASWQGEQRQTPSREYVDLEREAGKVYLKAPMILNGVCVIWKGWIDLHRLDGMGC
LEFDEERAQQEDALAQQ
000000000000000000000000000000000000000000000000000000000000

000000001111111111000000000000000000000000000000000000000000
00000000000000000
>DM_train2165
EPRFAGYAQKVRDSFARQPVMATLGARIDTLLPGRVELCMPYDRALTQQHGFLHAGIVSTVL
DSACGYAAFSLMEEEAAVLTVEFKVNFLNPAEGERFAFRAEVVKPGRTLTVATATAYAFRDG
EERAIATMTATLMALIG
111110000000000000000000000000000000000000000000000000000000
000000000000000000000000000000000000000000000000000000000000
00000000000000000
>DM_train2166
FNVASLKYKHHESIQMIIDRIDHLVLTVSDISTTIRFYEEVLGFSAVTFKQNRKALIFGAQK
INLHQQEMEFEPKASRPTPGSADLCFITSTPINDVVSEILQAGISIVEGPVERTGATGEIMS
IYIRDPDGNLIEISQYV
111111111111111100000000000000000000000000000000000000000000
000001111100000000000000000000000000000000000000000000000000
00000000000000001
>DM_train2167
MTENGRYEYIVIGSEAAGVGLVRELTAAGKKVLAVDKSKEKIELLEDEGFDAVIADPTDESF
YRSLDLEGVSAVLITGSDDEFNLKILKALRSVSDVYAIVRVSSPKKKEEFEEAGANLVVLVA
DAVKQAFMDKIKKMETL
111110000000000000000000000000000000000000000000000000000000
000000000000000000000000000000000000000000000000000000000000
00000000000000111
>DM_train2168
MRPIPEGYEAVFETVVTPEMTVRFEELGPVHPVYATYWMVKHMELAGRKIILPFLEEGEEGI
GSYVEARHLASALPGMRVRVVARHEKTEGNRVYARVEAYNELGDLIGVGRTEQVILPKAKVE
ALFRRLKERWEAERSPS
000000000000000000000000000000000000000000000000000000000000
000000000000000000000000000000000000000000000000000000000000
00000000000001111
>DM_train2169
MKFLTTNFLKCSVKACDTSNDNFPLQYDGSKCQLVQDESIEFNPEFLLNIVDRVDWPAVLTV
AAELGNNALPPTKPSFPSSIQELTDDDMAILNDLHTLLLQTSIAEGEMKCRNCGHIYYIKNG
IPNLLLPPHLVHHHHHH
000000000000000000000000000000000000000000000000000000000000
000000000000000000000000000000000000000000000000000000000000
00000000000011111
>DM_train2170
GMEMTKVTGKFDVKLTPENAYATGVGGVNLGRMALDKTFYGELEARSQGEMLSAMTAVKGSA
GYVAIEQVVGKLCGRQGSFVLQHFGIMTDGQNRLHLEVVPHSGAGELTGLYGTMAISIENGQ
HFYEFSFCFEPASEVEG
110000000000000000000000000000000000000000000000000000000000
000000000000000000000000000011000000000000000000000000000000
00000000000111111

>DM_train2171
MAHHHHHHMGSRISKEAAPVTFKNGKPTVKGTKTYPMFSNILYRIADTEARRWAFYNDSKEL
IIHVAVLFDYDSQIVPLGDTTAFRIDDPDEGNEDDFGKYLCEVDVRPLETQMFVEGSVTGWR
VDTLEARTAEDERGYRL
11111111111111111110000000000000000000000000000000000000000000
00000000000000000000000001111111111100000000000000000000000000
00000000000000001
>DM_train2172
GHMENKVINFKKIIDSRGSLVAIEENKNIPFSIKRVYYIFDTKGEEPRGFHAHKKLEQVLVC
LNGSCRVILDDGNIIQEITLDSPAVGLYVGPAVWHEMHDFSSDCVMMVLASDYYDETDYIRQ
YDNFKKYIAKINLEKEG
11100000000000000000000000000000000000000000000000000000000000
00000000000000000000000000000000000000000000000000000000000000
00000000000001111
>DM_train2173
GMDQESNATSQDILKQHAAHYESDMGGLPEALVQLAEYAPETFDAYSRMRTTMLKSEADGAK
LPLKYKHLILVVLDAIRDEPIGIVNHTRAAMNAGLSVDELIEGILLGIIVYGMPAWGKTGRK
AVTFAVEFEKELAGKRT
11111110000000000000000000000000000000000000000000000000000000
00000000000000000000000000000000000000000000000000000000000000
00000000000000011
>DM_train2174
NLYFQGMRFVNPIPFVRDINRSKSFYRDRLGLKILEDFGSFVLFETGFAIHEGRSLEETIWR
TSSDAQEAYGRRNMLLYFEHADVDAAFQDIAPHVELIHPLERQAWGQRVFRFYDPDGHAIEV
GESLSQSGENLYFQGGS
00000000000000000000000000000000000000000000000000000000000000
00011000000000000000000000000000000000000000000000000000000000
00001111111111111
>DM_train2175
NIEQQLYDVVKQLIEQRYPNDWGGAAAIRVEDGTIYTSVAPDVINASTELCMETGAILEAHK
FQKKVTHSICLARENEHSELKVLSPCGVCQERLFYWGPEVQCAITNAKQDIIFKPLKELQPY
HWTEAYHDEMVKEWSTR
00000000000000000000000000000000000000000000000000000000000000
00000000000000000000000000000000000000000000000000000000000000
00000000000000001
>DM_train2176
PKQVGNAQHLYDNYDLVPAMIAEVNPRDMVVMALVNTNVDPTLPPRWALATRNITAIPGIEG
DTRKVGTRIPAVAVTGQRSVGNQDSWDQISPMPIAWATPDSSVIARAESTIPSEQWTTLSKN
LNKLDQVRETKFDLLEL
11111111111111000000000000000000000000000000000000000000000000
00000000000000000000000000000000000000000000000000000000000000
00000000000000000
>DM_train2177
MAETKIVVGPQPFSVGEEYPWLAERDEDGAVVTFTGKVRNHNLGDSVNALTLEHYPGMTEKA

LAEIVDEARNRWPLGRVTVIHRIGELWPGDEIVFVGVTSAHRSSAFEAGQFIMDYLKTRAPF
WKREATPEGDRWVEAR
000000000000000000000000000000000000011111111000000000000000
000000000000000000000000000000000000000000000000000000000000
0001111111111111
>DM_train2178
SGDRSMVEVLADHPGELVRTDSPNFLSSVLPTHWRSNKTLPIAFKVVALGDVPDGTLVTVMA
GNDENYSAELRNATAAMKNQVARFNDLRFVGRSGRGKSFTLTITVFTNPPQVATYHRAIKIT
VDGPREPRRHRQKLDD
111111111111110000000000000000000000000000000000000000000000
000000000000000000000000000000000000000000000000000000000000
0000111111111111
>DM_train2179
MASVDFKTYVDQACRAAEEFVNVYYTTMDKRRRLLSRLYMGTATLVWNGNAVSGQESLSEFF
EMLPSSEFQISVVDCQPVHDEATPSQTTVLVVICGSVKFEGNKQRDFNQNFILTAQASPSNT
VWKIASDCFRFQDWAS
111000000000000000000000000000000000000000000000000000000000
000000000000000000000000000000000000000000000000000000000000
0000000000000000
>DM_train2180
GMRNPSEDHMMRTPYQIVADHYAASDRHDPAAMMADIAPAIEWTEMAGFPCAGTYRSADEIV
RNVFRRLGEEWDGYTFKLDALHDAGDTVIGVGRYSGTYRRTGKSFECRVAHVWRVDAGKIVH
FEQFTDTLLVAQAMQP
111111111100000000000000000000000000000000000000000000000000
000000000000000000000000000000000000000000000000000000000000
0000000000000000
>DM_train2181
TDQRWLIDKSALVRLTDSPDMEIWSNRIERGLVHITGVTRLEVGFSAECGEIARREFREPPL
SAMPVEYLTPRIEDRALEVQTLLADRGHHRGPSIPDLLIAATAELSGLTVLHVDKDFDAIAA
LTGQKTERLTHRPPSA
000000000000000000000000000000000000000000000000000000000000
000000000000000000000000000000000000000000000000000000000000
0000000000001111
>DM_train2182
MATEPKAGRPSDYMPEVADDICSLLSSGESLLKVCKRPGMPDKSTVFRWLAKHEDFRDKYAK
ATEARADSIFEEIFEIADNAIPDAAEVAKARLRVDTRKWALARMNPRKYGDKVTNELVGKDG
GAIQIETSPMSTLFGK
111111111000000000000000000000000000000000000000000000000000
000000000000000000000000000000000000000000000000000000000000
0000000011111111
>DM_train2183
ASRTMVSSGSEFESAVGYSRAVRIGPLVVVAGTTGSGDDIAAQTRDALRRIEIALGQAGATL
ADVVRTRIYVTDISRWREVGEVHAQAFGKIRPVTSMVEVTALIAPGLLVEIEADAYVGSAVA
DRNSGAGPKDPSPAGG

1000000000000000000000000000000000000000000000000000000000000000
0000000000000000000000000000000000000000000000000000000000011111
1111111111111111
>DM_train2184
KVILYARVSSNTQKDDLANQVKYLEEQVKEYDLVITDIGSGLNMKRKGFLKLLRMILNNEVS
RVITAYPDRLVRFGFEILEEVCKAHNCEIVVLNQEDKTPEEELVEDLATILVSFSGKLHGMR
SQKYEKVKKCAEELKN
0000000000001100000000000000000000000000000000000000000000000000
0000000000000000000000000000000000000000000000000000000000000000
0000000000000000
>DM_train2185
VYGSEVESKIIEFTIVGADEIIAEKLGISVGDFVYKIIRLRIIHSIPTIMEHTWMPISVIPG
VEVSVLEESIYSHIQNKLGLQVGTSVVRVKGIRPDDKEKQFMNLTNQDFLMRVEQVAYLTDG
RTFEYSYADHLPETFE
0000000000000000000000000000000000000000000000000000000000000000
0011111111111111100000000000000000000000000000000000000000000000
0000000000000001
>DM_train2187
MGSDKIHHHHHHMAYVKERIYESMFIIAPNVPEEERENLVERVKKIIEERVKGKIDKVERMG
MRKFAYEIKKFNEGDYTVIYFRCDGQNLQELENFYRVTPEIIRWQTFRRFDLEKKERKAQRE
KAAAEATESSEGGSED
1111111111111111100000000000000000000000000000000000000000000000
0000000000000000000000000000000000000000000000000000000000000001
1111111111111111
>DM_train2189
KRKEAVIIMNVAAHHGSELNGELLLNSIQQAGFIFGDMNIYHRHLSPDGSGPALFSLANMVK
PGTFDPEMKDFTTPGVTIFMQVPSYGDELQNFKLMLQSAQHIADEVGGVVLDDQRRMMTPQK
LREYQDIIREVKDANA
1000000000000000000000000000000000000000000000000000000000000000
0000000000000000000000000000000000000000000000000000000000000000
0000000000000011
>DM_train2190
STVVMADVSQYHVNHLVTFCLGEEDGVHTVEDASRKLAVMDSQGRVWAQEMLLRVSPSQVTL
LDPVSKEELESYPLDAIVRCDAVMPRGRSRSLLLLVCQEPERAQPDVHFFQGLLLGAELIRE
DIQGALQNYRSGRGER
1111000000000000000000000000000000000000000000000000000000000000
0000000000000000000000000000000000000000000000000000000000000000
0000000000111111
>DM_train2191
MATTAKEEMARFWEKNTKSSRPLSPHISIYKWSLPMAMSITHRGTGVALSLGVSLFSLAALL
LPEQFPHYVAVVKSLSLSPALIYSAKFALVFPLSYHTWNGIRHLVWDMGKGFKLSQVEQSGV
VVLILTLLSSAAIASE
1000000000000000000000000000000000000000000000000000000000000000
0000000000000000000000000000000000000000000000000000000000000000

0000000000000000
>DM_train2192
GMSAKDVFHQVVKIALEKDGWQITNDPLTISVGGVNLSIDLGAEKLIAAEREGEKIAVEVKS
FLERSSAISEFHTALGQFINYRGALRRRQPERVLYLAVPLTTYKTFFQLDFPKEMIAENQVK
MLIYDVEQEVIFQWIN
111110000000000000000000000000000000111111000000000000000000
000000000000000000000000000000000000000000000000000000000000
0000000000000000
>DM_train2193
PVFENNNQRYYESLPFKQLKELKIACSQYGPTAPFTIAMIENLGTQALPPNDWKQTARACLS
GGDYLLWKSEFFEQCARIADVNRQQGIQTSYEMLIGEGPYQATDTQLNFLPGAYAQISNAAR
QAWKRLPSLEHHHHHH
000000000000000000000000000000000000000000000000000000000000
000000000000000000000000000000000000000000000000000000000000
0000000111111111
>DM_train2194
GAMSVASLPECVKNFFPTEQLEFSSSITADEKPVLHEVFQKHSCFSQCGEMIDEVSKKHPEL
GKRLATVLEGNKKRLDGLSPAAVEYAKKLIHMVTTTLCSLTVGKPIDDADAKRLHQEFQSLS
SEDQAALRKNNPDIKF
000000000000000000000000000000000000000000111100000000000000
000000000000000000000000000000000000000000000000000000000000
0000000000000000
>DM_train2195
MTEKKERIDVHAYLAEFDDIPGTRVFTAQRARKGYNLNQFAMSLMKAENRERFKADESAYLD
EWNLTPAAKAAVLARDYNAMIDEGGNVYFLSKLFSTDGKSFQFAAGSMTGMTQEEYAQMMID
GGRSPAGVRSIKGGY
111111100000000000000000000000000000000000000000000000000000
000000000000000000000000000000000000000000000000000000000000
000000000000000
>DM_train2196
MANAASGMAVHDDCKLRFLELKAKRTHRFIVYKIEEKQKQVVVEKVGQPIQTYEEFAACLPA
DECRYAIYDFDFVTAENCQKSKIFFIAWCPDIAKVRSKMIYASSKDRFKRELDGIQVELQAT
DPTEMDLDVFRSRAN
111100000000000000000000000000000000000000000000000000000000
000000000000000000000000000000000000000000000000000000000000
000011111111111
>DM_train2197
MRITQGTFSFLPELTDEQITKQLEYCLNQGWAVGLEYTDDPHPRNTYWEMFGLPMFDLRDAA
GILMEINNARNTFPNHYIRVTAFDSTHTVESVVMSFIVNRPADEPGFRLVRQEEPGRTLRYS
IESYAVQAGPEGSRY
000000000000000000000000000000000000000000000000000000000000
000000000000000000000000000000000000000000000000000000000000
000001111111111
>DM_train2198

MRGSGQFRVIGPGHPIRALVGDEAELPCRISPGKNATGMEVGWYRSPFSRVVHLYRNGKDQD
AEQAPEYRGRTELLKESIGEGKVALRIQNVRFSDEGGYTCFFRDHSYQEEAAVELKVEDPFY
WINPGRSRSHHHHHH
111100000000000000000000000000000000000000000000000000000000
000000000000000000000000000000000000000000000000000000000011
111111111111111
>DM_train2199
GMKAADFTYVTVHGDNSRMSRLKAQYTMLFFYDPDCSNCRKFEKLFAEIPAFVEMVENGTLR
VLAIYPDENREEWATKAVYMPQGWIVGWNKAGDIRTRQLYDIRATPTIYLLDGRKRVILKDT
SMEQLIDYLATQAGK
000000000000000000000000000000000000000000000000000000000000
000000000000000000000000000000000000000000000000000000000000
000000000011111
>DM_train2200
MGSVVPHYSWFSGITQFQKGKEFQFAEGQGVPIANGIPASEQKGYWYRHNRRSFKTPDGQQK
QLLPRWYFYYLGTGPHAGASYGDSIEGVFWVANSQADTNTRSDIVERDPSSHEAIPTRFAPG
TVLPQGFYVEGSGRS
111110000000000000000000000000000000000000000000000000000000
000000000000000000000000000000000000000000000000000000000000
000000000000111
>DM_train2201
KVVKFSYMWTINNFSFCREEMGEVIKSSTFSSGANDKLKWCLRVNPKGLDEESKDYLSLYLL
LVSCPKSEVRAKFKFSILNAKGEETKAMESQRAYRFVQGKDWGFKKFIRRDFLLDEANGLLP
DDKLTLFCEVSVVQD
000000000000000000000000000000000000000000000000000000000000
000000000000000000000000000000000000000000000000000000000000
000000000000001
>DM_train2202
GMQTINATEIRNNFSYYIDTVVRDKPIAVKRNRDVLLFFSEQIIKDLLQDLKIHAELSKEDG
IIIGTIDGFDLVVSGESEQEVIQKLAEDLLEYAQDYMNDFKLFYNAPNRKTHYPYILKVLLS
SNIDEVKGYIYAEMV
100000000000000000000000000000000000000000000000000000000000
000000000000000000000000000000000000000000000000000000000000
000000000000000
>DM_train2203
MGKRFSSFQAAQIRIARPTGQLDEIIRFYEEGLCLKRIGEFSQHNGYDGVMFGLPHADYHLE
FTQYEGGSTAPVPHPDSLLVFYVPNAVELAAITSKLKHMGYQEVESENPYWSNGGVTIEDPD
GWRIVFMNSKGISGK
110000000000000000000000000000000000000000000000000000000000
000000000000000000000000000000000000000000000000000000000000
000000000000000
>DM_train2204
GGGGSSGGSTGGGSGSGPVTIEIGSKGEELAFDKTELTVSAGQTVTIRFKNNSAVQQHNWIL
VKGGEAEAANIANAGLSAGPAANYLPADKSNIIAESPLANGNETVEVTFTAPAAGTYLYICT

VPGHYPLMQGKLVVN
111111111111111000000000000000000000000000000000000000000000
000000000000000000000000000000000000000000000000000000000000
000000000000000
>DM_train2205
DDDGEHGGRRAALVVTDPLTRTECSACHMAYPAALLPARSWTALMADLPNHFGEDASLDEAS
RGQIESYLVANAADSSGAGRALRGLVQTDTPLRISELPWFKRKHADEVSPRMLEKARSMSNC
AACHTGAERGLFDDD
111111111110000000000000000000000000000000000000000000000000
000000000000000000000000000000000000000000000000000000000000
000000000000111
>DM_train2206
GMPQTITRGIKAMLDEANSSIETLTTADAIALHKSGASDVVIVDIRDPREIERDGKIPGSFS
CTRGMLEFWIDPQSPYAKPIFQEDKKFVFYCAGGLRSALAAKTAQDMGLKPVAHIEGGFGAW
RDAGGPIEAWAPKKK
000000000000000000000000000000000000000000000000000000000000
000000000000000000000000000000000000000000000000000000000000
000000001111111
>DM_train2207
GAMGDSIKQLLMAGQINKAFHQALLANDLGLVEFTLRHTDSNQAFAPEGCRLEQKVLLSLIQ
QISADMTNHNELKQRYLNEALLAINMADPITREHAPKVLTELYRNCQQFIKNSPKNSQFSNV
RLLMKAIITYRDQLK
000000000000000000000000000000000000001111111111000000000000
000000000000000000000000000000000000000000000000000000000000
000000000000001
>DM_train2208
MGFNVKTPLLATDVIIRLWDGENFKGIVLIERKYPPVGLALPGGFVEVGERVEEAAAREMRE
ETGLEVRLHKLMGVYSDPERDPRAHVVSVVWIGDAQGEPKAGSDAKKVKVYRLEEIPLDKLV
FDHKKIILDFLKGNY
111100000000000000000000000000000000000000000000000000000000
000000000000000000000000000000000000000000000000000000000000
000000000000000
>DM_train2209
MKPKLILMSHGRMAEETLASTQMIVGELADAAIVSMTAEDGLSGTQAKLAAILKEAGNVPTL
VLADLKGGTPCNVAMMAMGTYPQLRVVAGLNLAMAIEAAVSPVENVDELAAYLTQIGQSAVT
TIDLPELTDEEEFEE
000000000000000000000000000000000000000000000000000000000000
000000000000000000000000000000000000000000000000000000000000
000001111111111
>DM_train2210
QQVAVITLGIGDLEASARFYGEGFGWAPVFRNPEIIFYQMNGFVLATWLVQNLQEDVGVAVT
SRPGSMALAHNVRAETEVAPLMERLVAAGGQLLRPADAPPHGGLRGYVADPDGHIWEIAFNP
VWPIGADGSVTFAAK
000000000000000000000000000000000000000000000000000000000000

000000000000000000000000000000000000000000000000000000000000
000000000000001
>DM_train2211
VGAFRSGLEDKVSKQLESKGIKFEYEEWKVPYVIPASNHTYTPDFLLPNGIFVETKGLWESD
DRKKHLLIREQHPELDIRIVFSSSRTKLYKGSPTSYGEFCEKHGIKFADKLIPAEWIKEPKK
EVPFDRLKRKGGKK
111110000000000000000000000000000000000000000000000000000000
000000000000000000000000000000000000000000000000000000000000
00000000001111
>DM_train2212
MVVKAAGLVIYRKLAGKIEFLLLQASYPPHHWTPPKGHVDPGEDEWQAAIRETKEEANITKE
QLTIHEDCHETLFYEAKGKPKSVKYWLAKLNNPDDVQLSHEHQNWKWCELEDAIKIADYAEM
GSLLRKFSAFLAGF
111000000000000000000000000000000000000000000000000000000000
000000000000000111000000000000000000000000000000000000000000
00000000000000
>DM_train2213
MQSTVHIVGDNTGWSVPSSPNFYSQWAAGKTFRVGDSLQFNFPANAHNVHEMETKQSFDACN
FVNSDNDVERTSPVIERLDELGMHYFVCTVGTHCSNGQKLSINVVAANATVSMPPPSSSPPS
SVMPPPVMPPPSPS
000000000000000000000000000000000000000000000000000000000000
000000000000000000000000000000000000000000000011111111111111
11111111111111
>DM_train2214
MMTMFEEVEVEAYVYPTEDIRKVKKAMLNLIPGLQFEAFDKGEYVILVGRTKDKRALQRLYE
LFRGQQILDTARMMLEEGYFGEEIIIKVHKQVAYVGKVNFNEDSPLGPITITIRTKEPQKLM
KWLAPRTKDGVPIE
111100000000000000000000000000000000000000000000000000000000
000000000000000000000000000000000000000000000000000000000000
00000000000000
>DM_train2215
ENQRQQFPVEHVQLLCINCMVAVGHGSDLRKVEGTHHVNVNPNFSNYYNVSRDPVVINKVFK
DWKPGGVISCRNCGEVWGLQMIYKSVKLPVLKVRSMLLETPQGRIQAKKWSRVPFSVPDFDF
LQHCAENLSDLSLD
111110000000000000000000000000000000000000000000000000000000
000000000000000000000000000000000000000000000000000000000000
00000000111111
>DM_train2216
MKKLLFAIDDTEACERAAQYILDMFGKDADCTLTLIHVKPEFMLYGEAVLAAYDEIEMKEEE
KAKLLTQKFSTFFTEKGINPFVVIKEGEPVEMVLEEAKDYNLLIIGSSENSFLNKIFASHQD
DFIQKAPIPVLIVK
000000000000000000000000000000000000001110000000000000000000
000000000000000000000000000000000000000000000000000000000000
00000000000000

>DM_train2217
AHHHHHHGSNSKEFRKIALVGASKNPAKYGNIILKDLLSKGFEVLPVNPNYDEIEGLKCYRS
VRELPKDVDVIVFVVPPKVGLQVAKEAVEAGFKKLWFQPGAESEEIRRFLEKAGVEYSFGRC
IMVETSNKKIFLEV
111111111111100000000000000000000000000000000000000000000000
000000000000000000000000000000000000000000000000000000000000
00000111111111
>DM_train2218
MSARISLFAVVVEDMAKSLEFYRKLGVEIPAEADSAPHTEAVLDGGIRLAWDTVETVRSYDP
EWQAPTGGHRFAIAFEFPDTASVDKKYAELVDAGYEGHLKPWNAVWGQRYAIVKDPDGNVVD
LFAPLPLEHHHHHH
100000000000000000000000000000000000000000000000000000000000
000000000000000000000000000000000000000000000000000000000000
00000111111111
>DM_train2219
MAKLTRLGDLERAVMDHLWSRTEPQTVRQVHEALSARRDLAYTTVMAVLQRLAKKNLVLQIR
DDRAHRYAPVHGRDELVAGLMVDALAQAEDSGSRQAALVHFVERVGADEADALRRALAELEA
GHGNRPPAGAATET
110000000000000000000000000000000000000000000000000000000000
111000000000000000000000000000000000000000000000000000000000
11111111111111
>DM_train2220
MSSGVDLGTENLYFQSMDLALIPDVDIDSDGVFKYVLIRVHSAPRSGAPAAESKEIVRGYKW
AEYHADIYDKVSGDMQKQGCDCECLGGGRISHQSQDKKIHVYGYSMAYGPAQHAISTEKIKA
KYPDYEVTWANDGY
111111111110000000000000000000000000000001111111100000000000
000000000000000000000000000000000000000000000000000000000000
00000000000111
>DM_train2221
MARYLVVAHRTAKSPELAAKLKELLAQDPEARFVLLVPAVPPPGWVYEENEVRRRAEEEAAA
AKRALEAQGIPVEEAKAGDISPLLAIEEELLAHPGAYQGIVLSTLPPGLSRWLRLDVHTQAE
RFGLPVIHVIAQAA
110000000000000000000000111110000000000000000001000000000000
000000000000000000000000000000000000000000000000000000000000
00000000000011
>DM_train2222
GMATVRLLDDAEISTLPEVKAVFDDIRATRGSDFVNNIWRGLANDPALLKRTWEQVKTVMVG
EGALDPLTREMIYLAVSTANSCSYCAHSHTAAARAKGMTPAQHAEVLAIIGLAAQTNALVTA
MQIPVDEAFLVDGK
000000000000000000000000000000000000000000000000000000000000
000000000000000000000000000000000000000000000000000000000000
00000000000111
>DM_train2223
MSVLDELYREILLDHYQSPRNFGVLPQATKQAGGMNPSCGDQVEVMVLLEGDTIADIRFQGQ

GCAISTASASLMTEAVKGKKVAEALELSRKFQAMVVEGAPPDPTLGDLLALQGVAKLPARVK
CATLAWHALEEALR
11110000000000000000000000000000000000000000000000000000000000
00000000000000000000000000000000000000000000000000000000000000
00000000000000
>DM_train2224
GHMNTTLFRWPVRVYYEDTDAGGVVYHASYVAFYERARTEMLRHHHFSQQALMAERVAFVVR
KMTVEYYAPARLDDMLEIQTEITSMRGTSLVFTQRIVNAENTLLNEAEVLVVCVDPLKMKPR
ALPKSIVAEFKQGS
11111100000000000000000000000000000000000000000000000000000000
00000000000000000000000000000000000000000000000000000000000000
00000000001111
>DM_train2225
EEQPRLFELLGQPGYKATWHAMFKGESDVPKWVSDASGPSSPSTSLSLEGQPYVLANSCKPH
DCGNNRLLVAFRGDKSAAYGLQVSLPDEPAEVMQTPSKYATYRWYGEPSRQVRELLMKQLES
DPNWKLEHHHHHH
00000000000000000000000000000000000000000000000000000000000000
00000000000000000000000000000000000000000000000000000000000000
0000011111111
>DM_train2226
VDICNHALLVGYGRVGSLLGEKLLASDIPLVVIETSRTRVDELRERGVRAVLGNAANEEIMQ
LAHLECAKWLILTIPNGYEAGEIVASARAKNPDIEIIARAHYDDEVAYITERGANQVVMGER
EIARTMLELLETP
00000000000000000000000000000000000000000000000000000000000000
00000000000000000000000000000000000000000000000000000000000000
0000000000011
>DM_train2227
ALYLGNKRMNDMYYTIGQVAKMQHLTISQIRYYDKQGLFPFLQRNEKGDRIFNEEALKYLEM
ILCLKNTGMPIQKIKQFIDWSMEGDSTILHRLKLMKQQEANVLQLIQDTEKNLKKIQQKIAK
YEDEISSANATTK
11111111111000000000000000000000000000000000000000000000000000
00000000000000000000000000000000000000000000000000000000000000
0111111111111
>DM_train2228
MQNLLKNLAASLGRKPFVADKQGVYRLTIDKHLVMLAPHGSELVLRTPIDAPMLREGNNVNV
TLLRSLMQQALAWAKRYPQTLVLDDCGQLVLEARLRLQELDTHGLQEVINKQLALLEHLIPQ
LTPFSVASRVGWN
10000000000000000000000000000000000000000000000000000000000000
00000000000000000000000000000000000000000000000000000000000000
0001111111111
>DM_train2229
VAMVNTTTRLSDDALAFLSERHLAMLTTLRADNSPHVVAVGFTFDPKTHIARVITTGGSQKA
VNADRSGLAVLSQVDGARWLSLEGRAAVNSDIDAVRDAELRYAQRYRTPRPNPRRVVIEVQI
ERVLGSADLLDRA

111111111100000000000000000000000000000000000000000000000000
000000000000000000000000000000000000000000000000000000000000
0000000000011
>DM_train2230
RRWTFVRYHYINKAYEVTMKIQIISGFDRQLTAWLRVHGRRLTNNQKKTLFFVNRRYMQTHW
QNYMLWVKRKIKALGRPAAVGDYTRLGAEIGRRVDMVFFYNFLSGRKMIPPYSAYMAKLNAL
RPADVPVKNHGK
111111111100000000000000000000000000000000000000000000000000
000000000000000000000000000000000000000000000000000000000000
000000001111
>DM_train2231
MKGGIVHSDGSYPKDKFEKINGTWYYFDSSGYMLADRWRKHTDGNWYWFDNSGEMATGWKKI
ADKWYYFNEEGAMKTGWVKYKDTWYYLDAKEGAMVSNAFIQSADGTGWYYLKPDGTLADRPE
FTVEPDGLITVK
111111111111100000000000000000000000000000000000000000000000
000000000000000000000000000000000000000000000000000000000000
000000000000
>DM_train2232
SSSFDKGKYKKGDDASYFEPTGPYLMVNVTGVDGKGNELLSPHYVEFPIKPGTTLTKEKIEY
YVEWALDATAYKEFRVVELDPSAKIEVTYYDKNKKKEETKSFPITEKGFVVPDLSEHIKNPG
FNLITKVVIEKK
111111111111111000000000000000000000000000000000000000000000
000000000000000000000000000000000000000000000000000000000000
000000000000
>DM_train2233
MEAVKTFNSELYSLNDYKPPISKAKMTQITKAAIKAIKFYKHVVQSVEKFIQKCKPEYKVPG
LYVIDSIVRQSRHQFGQEKDVFAPRFSNNIISTFQNLYRCPGDDKSKIVRVLNLWQKNNVFK
SEIIQPLLDMAA
100000000000000000000000000000000000000000000000000000000000
000000000000000000000000000000000000000000000000000000000000
000000000000
>DM_train2234
NVGVFVLMHGDSTASSMLKTAQELLGTSIGTAMNMPLTMEVQTMYEQLRNQVITQKESLNNG
ILLLTDMGSLNSFGNMLFEETGIRTKAITMTSTMIVLEAIRMASVGRSLEDIYQNIQLSFES
VVREQFRSSLQK
000000000000000000000000000000000000000000000000000000000000
000000000000000000000000000000000000000000000000000000000000
000000001111
>DM_train2235
MTKLEQIQQWTAQHHASMTYLSNPKTIEYLTGFGSDPIERVLALVVFPDQDPFIFAPALEVE
VIKETGWQFPVIGYLDHENPWAMIADQVKQRHVNPEHVAIEKGQLQVARMEALAAQFSAPSF
DLDITSFIEHMR
100000000000000000000000000000000000000000000000000000000000
000000000000000000000000000000000000000000000000000000000000

000000000001
>DM_train2236
TVPAQLQFSAKTLDGHDFHGESLLGKPAVLWFWAPWCPTCQGEAPVVGQVAASHPEVTFVGV
AGLDQVPAMQEFVNKYPVKTFTQLADTDGSVWANFGVTQQPAYAFVDPHGNVDVVRGRMSQD
ELTRRVTALTSR
00000000000000000000000000000000000000000000000000000000000000
00000000000000000000000000000000000000000000000000000000000000
000000000011
>DM_train2237
ADLQVRDIMVPRSQMISIKATQTPREFLPAVIDAAHSRYPVIGESHDDVLGVLLAKDLLPLI
LKADGDSDDVKKLLRPATFVPESKRLNVLLREFRANHNHMAIVIDEYGGVAGLVTIEDVLEQ
IVGDIEDEHDVE
11000000000000000000000000000000000000000000000000000000000000
00000000000000000000000000000000000000000000000000000000000000
011111111111
>DM_train2238
MRDPFMEALGLKVLHLAPGEAVVAGEVRADHLNLHGTAHGGFLYALADSAFALASNTRGPAV
ALSCRMDYFRPLGAGARVEARAVEVNLSRRTATYRVEVVSEGKLVALFTGTVFRLGGDGDDV
PAGTGNLAPREA
00000000000000000000000000000000000000000000000000000000000000
00000000000000000000000000000000000000000000000000000001111111
111111111111
>DM_train2239
MGSDKIHHHHHHLKMKKYTKTHEWVSIEDKVATVGITNHAQEQLGDVVYVDLPEVGREVKKG
EVVASIESVKAAADVYAPLSGKIVEVNEKLDTEPELINKDPEGEGWLFKMEISDEGELEDLL
DEQAYQEFCAQE
11111110000000000000000000000000000000000000000000000000000000
00000000000000000000000000000000000000000000000000000000000000
000000000001
>DM_train2240
MGSDKIHHHHHHMRVELLFESGKCVIDLNEEYEVVKLLKEKIPFESVVNTWGEEIYFSTPVN
VQKMENPREVVEIGDVGYWPPGKALCLFFGKTPMSDDKIQPASAVNVIGKIVEGLEDLKKIK
DGEKVAVRFASS
11111111100000000000000000000000000000000000000000000000000000
00000000000000000000000000000000000000000000000000000000000000
000000000000
>DM_train2241
GDADVLVKWSEDLANLPSIDTQHKRLVDYINDLYRAARRRDMDKAREVFDALKNYAVEHFGY
EERLFADYAYPEATRHKEIHRRFVETVLKWEKQLAAGDPEVVMTTLRGLVDWLVNHIMKEDK
KYEAYLRERGVS
11100000000000000000000000000000000000000000000000000000000000
00000000000000000000000000000000000000000000000000000000000000
000000000000
>DM_train2242

SNASAMAQKEEGHEAAAAEAKKETDLAHVDQVETFVGKEKYYVVKGTDKKGTALYVWVPADK
KAKILSKEAKEGISEDKAAKIIKDEGLVSKQKEVHLAREGNVLLWEVTYLDKEGQYSLSYVD
FTTGKILKNITP
111111110000000000000000000000000000000000000000000000000000
000000000000000000000000000000000000000000000000000000000000
000000000000
>DM_train2243
GSHMTSPFKLPDESPSWTEWRLHNDETNSNQDNPLGFKESWGFGKVVFKRYLRYDRTEASLH
RVLGSWTGDSVNYAASRFFGFDQIGCTYSIRFRGVSITVSGGSRTLQHLCEMAIRSKQEMLQ
MAPIEVESNVSR
110000000000000000000000001110000000000000000000000000000000
000000000000000000000000000000000000000000000000000000000000
001111111111
>DM_train2244
MVKLMIIEGEVVSGLGEGRYFLSLPPYKEIFKKILGFEPYEGTLNLKLDREFDINKFKYIET
EDFEFNGKRFFGVKVLPIKILIGNKKIDGAIVVPKKTYHSSEIIEIIAPMKLREQFNLKDGD
VIKILIKGDKDE
100000000000000000000000000000000000000000000000000000000000
000000000000000000000000000000000000000000000000000000000000
000000001111
>DM_train2245
HTAFLKAMLSTGFKIPQKGILIGIQQSFRPRFLGVAEQLHNEGFKLFATEATSDWLNANNVP
ATPVAWPSQEGQNPSLSSIRKLIRDGSIDLVINLPNNNTKFVHDNYVIRRTAVDSGIPLLTN
FQVTKLFAEAVQ
000000000000000000000000000000000000000000000000000000000000
000000000011111000000000000000000000000000000000000000000000
000000000000
>DM_train2247
SNAMGLFDEVVGAFLKGDAGKYQAILSWVEEQGGIQVLLEKLQSGGLGAILSTWLSNQQRNQ
SVSGEQLESALGTNAVSDLGQKLGVDTSTASSLLAEQLPKIIDALSPQGEVSAQANNDLLSA
GMELLKGKLFR
111100000000000000000000000000000000000000000000000000000000
000000000000000000000000000000000000000000000000011000000000
00000000001
>DM_train2248
GSSQNEASEDTGFNPKAFPLASPDLNNKIINLVQQACNYKQLRKGANEATKALNRGIAEIVL
LAADAEPLEILLHLPLVCEDKNTPYVFVRSKVALGRACGVSRPVIAAAITSKDGSSLSSQIT
ELKDQIEQILV
111111111111111110000000000000000000000000000000000000000000
000000000000000000000000000000000000000000000000000000000000
00000000111
>DM_train2249
GMENIEYVFEDVVRIYDTDAQGIAHYAAYYRFFTNTIEKFIKEKVGIPYPIVNENLWFVIAE
SHAIYHRPVKLGDKLTVLLNPKILSNKTIKFEFKVLKDGELTTEGYVIQIAINPKIWKSTEM

PKEIMDKLSIK
11111100000000000000000000000000000000000000000000000000000000
00000000000000000000000000000000000000000000000000000000000000
00000111111
>DM_train2250
MQISSDYIPDSKFYKVEAIVRPWRIQQVSSALLKIGIRGVTVSDVRGFGAQGGSTERHGGSE
FSEDKFVAKVKMEIVVKKDQVESVINTIIEGARTGEIGDGKIFVLPVSDVIRVRTGERGEKA
EKMTGDMLSPS
11100000000000000000000000000000000000000000001111111111111111
11100000000000000000000000000000000000000000000000000000000000
00111111111
>DM_train2251
AGGGGSSSGADHISLNPDLANEDEVNSCDYWRHCAVDGFLCSCCGGTTTTCPPGSTPSPISW
IGTCHNPHDGKDYLISYHDCCGKTACGRCQCNTQTRERPGYEFFLHNDVNWCMANENSTFHC
TTSVLVGLAKN
11111111111111111111110000000000000000000000000000000000000000
00000000000000000000000000000000000000000000000000000000000000
00000000011
>DM_train2254
EAPFTLKVNTLPLNFDKAEHHRKFQIHINVSYIGERPNSNMVIVDVKMVSGFIPVKPSVKKL
QDQSNIQRTEVNTNHVLIYIEKLTNQTMGFSFAVEQDIPVKNLKPAPVKVYDYYETDEFAIE
EYSAPFSSDS
00000000000000000000000000000000000000000000000000000000000000
00000000000000000000000000000000000000000000000000000000000000
0000001111
>DM_train2255
MARIPMASAGTSCVPGWAIPHNPLPSCRWYVTSRTCGIGPRLPWPELKRRCCRELADIPAYC
RCTALSILMDGAIPPGPDAQLEGRLEDLPGCPREVQRGFAATLVTEAECNLATISGVAECPW
ILGGGTMPSK
11111111111000000000000000000000000000000000000000000000000000
00000000000000000000000000000000000000000000000000000000000000
0001111111
>DM_train2256
STTPAAGVLDTSVFIATESGRQLDEALIPDRVATTVVTLAELRVGVLAAATTDIRAQRLATL
ESVADMETLPVDDDAARMWARLRIHLAESGRRVRINDLWIAAVAASRALPVITQDDDFAALD
GAASVEIIRV
11100000000000000111110000000000000000000000000000000000000000
00000000000000000000000000000000000000000000000000000000000000
0000000000
>DM_train2257
MIYMPIVVAVDKKSDRAERVLRFAAEEARLRGVPVYVVHSLPGGGRTKDEDIIEAKETLSWA
VSIIRKEGAEGEEHLLVRGKEPPDDIVDFADEVDAIAIVIGIRKRSPTGKLIFGSVARDVIL
KANKPVICIK
10000000000000000000000000000000000000000000000000000000000000

000000000000000000000000000000000000000011111000000000000000
0000000000
>DM_train2258
GMSNALATLASPADDIAFYEERLRAAMLTGDLKGLETLLADDLAFVDHTGCVKTKQTHLEPY
RAGLLKLSRLDLSDAVVRAAGEDGRVVVVRAVTAGVYDGEAFTETLRFTRIWRRTQGPAGWK
LVAGHCSVIL
111111111111100000000000000000000000000000000000000000000000
000000000000000000000000000000000000000000000000000000000000
0000000000
>DM_train2260
GMTKAREFLGTGWKFPVAAGADGAMVLSSAEEDIAESIRIILGTARGERVMRPDFGCGIHDR
VFSVINTTTLGLIENEVKEALILWEPRIELLSVTASPREAAEGRLLIDIEYRVRSTNTRFNL
VYPFYLKESA
111111111111111111111100000000000000000000000000000000000000
000000000000000000000000000000000000000000000000000000000000
0000000000
>DM_train2261
GMKLQTTIQHEPKDGSGFDRGLREFFEYRDTGVNEATGGMFGAHVIRAIPGKEAKPTWHTHT
VGFQLFYVLRGWVEFEYEDIGAVMLEAGGSAFQPPGVRHRELRHSDDLEVLEIVSPAGFATS
VVDLEEARKP
000000000000000000001100000000000000000000000000111110000000
000000000000000000000000000000000000000000000000000000000000
0000011111
>DM_train2262
AGGRSWCLRRVGMSAGWLLLEDGCEVTVGRGFGVTYQLVSKICPLMISRNHCVLKQNPEGQW
TIMDNKSLNGVWLNRARLEPLRVYSIHQGDYIQLGVPLENKENAEYEYEVTEEDWETIYPCL
SPKNDQMIEK
000000000000000000000000000000000000000000000000000000000000
000000000000000000000000000000000000000000000000000000000000
0000111111
>DM_train2263
TLRVVPELYCFDINVSQSFFVDVLGFEVKYERPDEEFVYLTLDGVDVMLEGIAGKSRKWLSG
DLEFPLGSGVNFQWDVIDIEPLYQRVNESAADSIYLALESKSYQCGDSIATQKQFMVQTPDG
YLFRFCQDIH
000000000000000000000000000000000000000000000000001111111111
100000000000000000000000000000000000000011111000000000000000
0000000011
>DM_train2264
NAIMDLQTRNTRGLSTLVVRDIGELMMAGDMAVIERYVADVRGKGAVLDLRIYDAAGRPAGK
KQDAPDGEVQAALTSGATAEKRHKVDGRHVLSFIVPLANEVRCQSCHEQGARFNGAMLLTTS
LEEGYAGARN
110000000000000000000000000000000000000000000000000000000011
111000000000000000000000111000000000000000000000000000000000
0000000111

>DM_train2265
MKEKPAVEVRLDKWLWAARFYKTRALAREMIEGGKVHYNGQRSKPSKIVELNATLTLRQGND
ERTVIVKAITEQRRPASEAALLYEETAESVEKREKMALARKLNALTMPHPDRRPDKKERRDL
LRFKHGDSE
11110000000000000000000000000000000000000000000000000000000000
00000000000000000000000000000000000000000000000111111111111111
111111111
>DM_train2266
RTTLAYPATAVSVAKNLAANEPSSFTYPDTSSPCVAVKLGAPVPGGVGPDDDIVAYSVLCTH
MGCPTSYDSSSKTFSCPCHFTEFDAEKAGQMICGEATADLPRVLLRYDAASDALTAVGVDGL
IYGRQANVI
11111000000000000000000000000000000000000000000000000000000000
00000000000000000000000000000000000000000000000000000000000000
000000000
>DM_train2267
HMGARASVLSGGELDKWEKIRLRPGGKKQYKLKHIVWASRELERFAVNPGLLETSEGCRQIL
GQLQPSLQTGSEELRSLYNTIAVLYCVHQRIDVKDTKEALDKIEEEQNKSKKKAQQAAADTG
NNSQVSQNY
11111110000000000000000000000000000000000000000000000000000000
00000000000000000000000000000000000000000000000011111111111111
111111111
>DM_train2268
MAAASVTPPGSLELLQPGFSKTLLGTKLEAKYLCSACRNVLRRPFQAQCGHRYCSFCLASIL
SSGPQNCAACVHEGIYEEGISILESSSAFPDNAARREVESLPAVCPSDGCTWKGTLKEYESC
HEGRCPLML
11111111111111000000000000000000000000000000000000000000000000
00000000000000000000000000000000000000000000000000000000000000
000000000
>DM_train2269
DEHLCANLMQLLQESLAQARLGSRRPARLLMPSQLVSQVGKELLRLAYSEPCGLRGALLDVC
VEQGKSCHSVGQLALDPSLVPTFQLTLVLRLDSRLWPKIQGLFSSANSPSQSLTLSTGFRVI
KKKLYSSEQ
00000000000000000000000000000000000000000000000000000000000000
00000000000000000000000000000000000010000000111000000000000000
000001111
>DM_train2271
GSPAAAPPTLPPYFMKGSIIQLANGELKKVEDLKTEDFIQSAEISNDLKIDSSTVERIEDSH
SPGVAVIQFAVGEHRAQVSVEVLVEYPFFVFGQGWSSCCPERTSQLFDLPCSKLSVGDVCIS
LTLKNLKNG
00000000000000000000000000000000000000000000000000000000000000
00000000000000000000000000000000000000000000000000000000000000
000011111
>DM_train2272
AHHHHHHGSEIDKGLAKFGDSLINFLYSLALTEFLGKPTGDRVPNASLAIALELTGLSKNLR

RVDKHAKGDYAEALIAKAWLMGLISEREAVEIIKKNLYPEVLDFSKKKEAIGRALAPLLVII
SERLYSSQV
111111110000000000000000000000000000000000000000000000000000
000000000000000000000000000000000000000000000000000000000000
000000000
>DM_train2273
GAMGQTTFTELMQQLFLKLGLNHQVNENDVYTFEVDGHIQVLIACYHQQWVQLFSELGADLP
TNDNLFGEHWPAHVQGRLDGKSILWSQQSLVGLDIDEMQAWLERFIDDIEQRKEPQNTKFQP
NSTSPILFI
111100000000000000000000000000000000000000000000000000000000
000000000000000000000000000000000000000000000000000000001111
111000000
>DM_train2274
GPLDLSRDECKRILRKLELEAYAGVISALRAQGDLTKEKKDLLGELSKVLSISTERHRAEVR
RAVNDERLTTIAHNMSGPNSSSEWSIEGRRLVPLMPRLVPQTAFTVTANAVANAAIQHNASL
PVPAETGSK
000000000000000000000000000000000000000000000000000000000000
000000000000000000000000000000000000000000000000000000111111
111111111
>DM_train2275
GQSLELGTMQPSFTSVTGKGGVKVIDGSSVKFGRFDGAEPHCVGLTDLVTEQDGSSMAAGFM
QWDNAFFPWTLNYDEIDMVLEGELHVRHEGETMIAKAGDVMFIPKGSSIEFGTPTSVRFLYV
AWPANWQSV
111111000000000000000000000000000000000000000000000000000000
000000000000000000000000000000000000000000000000000000000000
000001111
>DM_train2276
AKLETVTLGNIGKDGKQTLVLNPRGVNPTNGVASLSQAGAVPALEKRVTVSVSQPSRNRKNY
KVQVKIQNPTACTANGSCDPSVTRQAYADVTFSFTQYSTDEERAFVRTELAALLASPLLIDA
IDQLNPAY
000000000000000000000000000000000000000000000000000001111100
000000000000011110000000000000000000000000000000000000000000
00000000
>DM_train2277
MSSILPFTPPIVKRLLGWKKGEQNGQEEKWCEKAVKSLVKKLKKTGQLDELEKAITTQNVNT
KCITIPRSLDGRLQVSHRKGLPHVIYCRLWRWPDLHSHHELRAMELCEFAFNMKKDEVCVNP
YHYQRVET
111111111000000000000000000000000000000000000000000000000000
000000000000000000000000000000000000000000000000000000000000
00000000
>DM_train2278
MYSFEQAITQLFQQLSLSIPDTIEPVIGVKVGEFACHITEHPVGQILMFTLPSLDNNNEKET
LLSHNIFSQDILKPILSWDEVGGHPVLWNRQPLNNLDNNSLYTQLEMLVQGAERLQTSSLIS
PPRSFSPG

000000000000000000000000000000000000000000000000000000000000
000000000000000000000000000000000000000000000000000000111111
10000111
>DM_train2279
MLPAVPPQQWALSAGNGSSEVEVVPFQEVWGRSYCRALERLVDVVSEYPSEVEHMFSPSCVS
LLRCTGCCGDENLHCVPVETANVTMQLLKIRSGDRPSYVELTFSQHVRCECRPLREKMKPER
CGDAVPRR
111111111111111110000000000000000000000000000000000000000000
000000000000000000000000000000000000000000000000000001111111
11111111
>DM_train2280
MFSITVRDHIMIAHSFRGDVFGPAQRLHGATFLVDATFRREQLDEDNIVVDIGLATQELGAV
VGALNYRNLDNEPDFAGVNTSTEFLAKVIADRLAERVHKGALGEGARGLAGLTVTLHESHVA
WASYERAL
100000000000000000000000000000000000000000000000000000000000
000000000000000000000000000000000000000000000000000000000000
00000000
>DM_train2281
HMVTLYTSPSCTSCRKARAWLEEHEIPFVERNIFSEPLSIDEIKQILRMTEDGTDEIISTRS
KVFQKLNVNVESMPLQDLYRLINEHPGLLRRPIIIDEKRLQVGYNEDEIRRFLPRKVRSFQL
REAQRLAN
000000000000000000000000000000000000000000000000000000000000
000000000000000000000000000000000000000000000000000000011111
11111111
>DM_train2282
GSSSSGLVPRGSLTHRKFGGSGGSPFSGLSSIAVRSGSYLDAIIIDGVHHGGSGGNLSPTFT
FGSGEYISNMTIRSGDYIDNISFETNMGRRFGPYGGSGGSANTLSNVKVIQINGSAGDYLDS
LDIYYEQY
111111000000000000000000000000000000000000000000000000000000
000000000000000000000000000000000000000000000000000000000000
00000000
>DM_train2283
VASPAIVEAKPVKTIKDKVRAMRELLLSDEYAEQKRAVNRFMLLLSTLYSLDAQAFAEATES
LHGRTRVYFAADEQTLLKNGNQTKPKHVPGTPYWVITNTNTGRKCSMIEHIMQSMQFPAELI
EKVCGTI
111111111111100000000000000000000000000000000000000000000000
000000000000000000000000000000000000000000000000000000000000
0000000
>DM_train2284
SNRQVQLMARQQRLKAIEDRLEKFYIPLIKAFSSYVYTAQTEDEIETIITCRRYLAGNNLLR
VLPMHFKFKADKIAGSANWTFYAKEDFEQWKEALDVLWEEFLEVLKEYYTLSGTEISLPEKP
DWLIGYK
111111111111000000000000000000000000000000000000000000000000
000000000000000000000000000000000000000000000000000000000000

0000000
>DM_train2285
MAKYEPKKGDYAGGAVKILDMFENGQLGYPEVTLKLAGEEANARRAGDERTKEAIHAIVKMI
SDAMKPYRNKGSGFQSQPIPGEVIAQVTSNPEYQQAKAFLASPATQVRNIEREEVLSKGAKK
LAQAMAS
111111000000000000000000000000000000000000000000000000000000
000000001111111000000000000000000000000000000000001111111111
1111111
>DM_train2286
MSVIKIEKALKLDKVIFVDVRTEGEYEEDHILNAINMPLFKNNEHNEVGTIYKMQGKHEAIQ
KGFDYVSYKLKDIYLQAAELALNYDNIVIYCARGGMRSGSIVNLLSSLGVNVYQLEGGYKAY
RNFVLEY
100000000000000000000000000000000000000000000000000000000000
000000000000000000000000000000000000000000000000000000000000
0000001
>DM_train2287
WQFTPNTGGNTITKVEAENMKIGGTYAGKISAPFDGVALYANADYVSYSQYFANSTHNISVR
GASSNAGTAKVDLVIGGVTVGSFNFTGKTPTVQTLSNITHATGDQEIKLALTSDDGTWDAYV
DFIEFSL
111111111110000000000000000000000000000000000000000000000000
000000000000000000000000000000000000000000000000000000000000
0000000
>DM_train2288
MIPVEQRTHKLTSRILVGKPILIKEGYAEVELETIDEMKVDEKGLVHGGFTFGLADYAAMLA
VNEPTVVLGKAEVRFTKPVKVGDKLVAKAKIIEDLGKKKIVEVKVYREEEVVLEGKFYCYVL
EKHVLDN
000000000000000000000000000000000000000000000000000000000000
000000000000000000000000000000000000000000000000000000000000
0000001
>DM_train2289
MRRAGYLHLYGLNLVFDRVGKGPPVLLVAEEASRWPEALPEGYAFYLLDLPGYGRTEGPRMA
PEELAHFVAGFAVMMNLGAPWVLLRGLGLALGPHLEALGLRALPAEGVEVAEVLSSKLSYGN
IDLGGNL
100000000000000000000000000000000000000000000000000000000000
000000000000000000000000000000000000000000000000000000000001
1111111
>DM_train2291
QVVAAIRHITTGTYIARIREEYQQTEVKPELQPMKEALARMTDRAEALIAFVTEQKDQELLD
FQARRLVEMTAHAVFGHLLMLAANDDDSFRQSAEVYLRYGQAEQEKIDSYVRAFRPEELTFY
SRCSRPE
110000000000000000000000000000000000000000000000000000000000
000000000000000000000000000000000000000000000000000000111111
1111111
>DM_train2292

GSHMVPGQKQHYVQPTAANSGEISYADFEKVDIRVGTIVEAVPFPEARKPAIKVKIDFGPEI
GIKKSSAQITVHYTPESLVGRQVLGVVNFPPRQIGPFRSEVLTLGFADANGDIVLAAVERPV
PNGEKMC
11111111111111111110000000000000000000000000111110000000000000
00000000000000000000000000000000000000000000000000000000000000
0000000
>DM_train2293
MKTVKHLLQEKGHTVVAIGPDDSVFNAMQKMAADNIGALLVMKDEKLVGILTERDFSRKSYL
LDKPVKDTQVKEIMTRQVAYVDLNNTNEDCMALITEMRVRHLPVLDDGKVIGLLSIGDLVKD
AISQHQF
00000000000000000000000000000000000000000000000000000000000000
01000000000000000000000000000000000000000000000000000000000000
0001111
>DM_train2294
MDDSEDTSWDFGPQAFKLLSAVDILGEKFGIGLPILFLRGSNSQRLADQYRRHSLFGTGKDQ
TESWWKAFSRQLITEGFLVEVSRYNKFMKICALTKKGRNWLHKANTESQSLILQANEELCPK
KFLLPSS
11111110000000000000000000000000000000000000000000000000000000
00000000000000000000000000000000000000000000000000000011111111
1111111
>DM_train2295
MSKDEVKREAKDTDGNPEIKGERRRLHSEIQSGSLANNIKKSTVIVKNATHIAICLYYKLGE
TPLPLVIETGKDAKALQIIKLAELYDIPVIEDIPLARSLYKNIHKGQYITEDFFEPVAQLIR
IAIDLDY
11111111111111111111111111111000000000000000000000000000000000
00000000000000000000000000000000000000000000000000000000000000
0000000
>DM_train2296
MSNQIYSARYSGVDVYEFIHSTGSIMKRKKDDWVNATHILKAANFAKAKRTRILEKEVLKET
HEKVQGGFGKYQGTWVPLNIAKQLAEKFSVYDQLKPLFDFTQTDGSASPPPAPKHHHASKVD
HHHHHH
00000000000000000000000000000000000000000000000000000000000000
00000000000000000000000000000000000000001111111111111111111111
111111
>DM_train2297
ALITQQDLAPQQRAAPQQKRSSPSEGLCPPGHHISEDGRDCISCKYGQDYSTHWNDLLFCLR
CTRCDSGEVELSPCTTTRNTVCQCEEGTFREEDSPEMCRKCRTGCPRGMVKVGDCTPWSDIE
CVHKES
11111111111111111111000000000000000000000000000000000000000000
00000000000000000000000000000000000000001111111111111000000001
111111
>DM_train2298
GSASAKELACQEITVPLCKGIGYEYTYMPNQFNHDTQDEAGLEVHQFWPLVEIQCSPDLKFF
LCSMYTPICLEDYKKPLPPCRSVCERAKAGCAPLMRQYGFAWPDRMRCDRLPEQGNPDTLCM

DYERTD
111111000000000000000000000000000000000000000000000000000000
000000000000000000000000000000000000000000000000000000000000
000000
>DM_train2299
GMATEKYHEILKKYFLSFETGDFSQVQFSCNLEFLSPISGNTLKGTEEVIPFLKGVTTRVAE
VNIMSTTVEYPRASGVWQMRTTKGTLYTLHNFFRLDEEGIVYVWPMFDPKAVMENPDALIQW
LTGKDY
110000000000000000000000000000000000000000000000000000000000
000000000000000000000000000000000000000000000000000000000000
000000
>DM_train2300
MAKTDRSQPSVLARIAAFRTGLSAEASAADYLERQGYRILARRFKTRCGEIDLVAQRDALVA
FVEVKARGNVDDAAYAVTPRQQSRIVAAAEAWLSRHPEHAMSELRFDAILIAPNTAPRHLPG
AFDATP
111111111111111111111000000000000000000000000000000000000000
000000011111100000000000000000000000000000000000000000000000
000000
>DM_train2302
DEPDLKDLFITVDEPESHVTTIETFITYRIITKTSRGEFDSSEFEVRRRYQDFLWLKGKLEE
AHPTLIIPPLPEKFIVKGMVERFNDDFIETRRKALHKFLNRIADHPTLTFNEDFKIFLTAQA
WELSSH
000000000000000000000000000000000000000000000000000000000000
000000000000001111000000000000000000000000000000000000000000
001111
>DM_train2303
MRKSLFAVMVLALVAAFALPVIAAEAPADGLKMENTKMPVIFNHSSHSSYQCADCHHPVDGK
ENLAKCATAGCHDVFDKKDKSVHSYYKIIHDRKATTVATCMSCHLEAAGSDKDLKKELTGCK
KSKCHP
111111111111111111111100000000000000000000000000000000000000
000000000000000000000000000000000000000000000000000000000000
000000
>DM_train2304
MELFTEAWAQAYCRKLNESEAYRKAASTWEGSLALAVRPDPKAGFPKGVAVVLDLWHGACRG
AKAVEGEAEADFVIEADLATWQEVLEGRLEPLSALMRGLLELKKGTIAALAPYAQAAQELVK
VAREVA
100000000000000000000000000000000000000000000000000000000000
000000000000000000000000000000000000000000000000000000000000
000000
>DM_train2305
MMRCPECSTEGWRVLPLTVGAHVKEGLWSKIKGDFYFCSLESCEVVYFNEQTVFRKGELKTR
VGVKEREEPKPVCYCNRVTEKMLLEAAEKFGKEKAVEITGAGKGKWCVVTNPSGRCCHWHLE
RLGFPV
100000000000000000000000000000000000000000000000000000000000

000000000000000000000000000000000000000000000000000000000000
000000
>DM_train2306
SNAMFHYTVDVSTGMNETIERLEESLKQEGFGVLWQFSVTEKLQEKGLDFSTPMVILEVCNP
QEAARVLNENLLVGYFLPCKLVVYQENGTTKIGMPKPTMLVGMMNDPALKEIAADIEKRLAA
CLDRCR
110000000000000000000000000000000000000000000000000000000000
000000000000000000000000000000000000000000000000000000000000
000000
>DM_train2307
MVQIIFDSKTGNVQRFVNKTGFQQIRKVDEMDHVDTPFVLVTYTTNFGQVPASTQSFLEKYA
HLLLGVAASGNKVWGDNFAKSADTISRQYQVPILHKFELSGTSKDVELFTQEVERVVTKSSA
KMDPVK
000000000000000000000000000000000000000000000000000000000000
000000000000000000000000000000000000000000000000000000000000
001111
>DM_train2308
KNMADLSYVNIFTRDIVAMSAFYQQVFGFQEIESIRSPIFRGLDTGKSCIGFNAHEAYELMQ
LAQFSETSGIKFLLNFDVDTKEAVDKLVPVAIAAGATLIKAPYETYYHWYQAVLLDPERNVF
RINNVL
111000000000000000000000000000000000000000000000000000000000
000000000000000000000000000000000000000000000000000000000000
000000
>DM_train2309
MGLNIGDLLGSTPAKDQGCSRTCESQFCTIAPLLRYGKYCGILYSGCPGERPCDALDACCMV
HDHCVDTHNDDYLNTMCNENLLSCIDRVSGATFPGNKCNVGQTASVIRGVIETAVFAGKILH
KRDDGQ
111000000111110000000000000000000000000000000000000000000000
000000000000000000000000000000000000000000000000000000000000
000111
>DM_train2310
MPDRTTDDFGPFTEQIRGTIDGMGTAEFDALPVGAIQVDGSGVIHRYNRTESRLSGRIPERV
IGRNFFTEVAPCTNIPAFSGRFMDGVTSGTLDARFDFVFDFQMAPVRVQIRMQNAGVPDRYW
IFVRK
111111111111111111100111000000000000000000000000000000000000
000000000000000000000000000000000000000000000000000000000000
00000
>DM_train2311
MAKKPSKKKVKRQVASGRAYIHASYNNTIVTITDPDGNPITWSSGGVIGYKGSRKGTPYAAQ
LAALDAAKKAMAYGMQSVDVIVRGTGAGREQAIRALQASGLQVKSIVDDTPVPHNGCRPKKK
FRKAS
111111111000000000000000000000000000000000000000000000000000
000000000000000000000000000000000000000000000000000000000000
00000

>DM_train2312
ATLHSFVLVDNGGTGNVTVVPVSNANGVAEWLSNNSRSQAYRVTASYRASGADKRKYTIKLE
VPKIVTQVVNGVELPVSAWKAYASIDLTIPIFAATDDVTVISKSLTGLFKVGNPIAEAISSQ
SGFYA
000000000000000000000000000000000000000000000000000000000000
000001111111000000000000000000000000000000000000000000000000
00000
>DM_train2313
APQSITELCSEYHNTQIYTINDKILSYTESMAGKREMVIITFKSGATFQVEVPGSQHIDSQK
KAIERMKDTLRITYLTETKIDKLCVWNNKTPNSIAAISMEKLAGFGAVGAGATAEETRMLHA
FDTLA
000000000000000000000000000000000000000000000000000000000000
000000000000000000000000000000000000000000111111111111111111
11111
>DM_train2314
MDIGINSDPNSSTQSYKDAMGPLVRECMGSVSATEDDFKTVLNRNPLESRTAQCLLACALDK
VGLISPEGAIYTGDDLMPVMNRLYGFNDFKTVMKAKAVNDCANQVNGAYPDRCDLIKNFTDC
VRNSY
111111111111000000000000000000000000000000000000000000000000
000000000000000000000000000000000000000000000000000000000000
00000
>DM_train2315
MNSPTSRPLRRDPLHNALVTTNVLVVLDQLEVAAEPTRRRLVQLLTSGEQTVNNLAAHFPAS
RSAISQHLRVLTEAGLVTPRKDGRFRYYRLDPQGLAQLRALFDSFWIDELDRLVADATEEAA
SKGDS
111111111111111111111111100000000000000000000000000000000000
000000000000000000000000000000000000000000000000000000000111
11111
>DM_train2316
MVAVPEGYESLLERPLYGHLATVRPDGTPQVNAMWFAWDGEVLRFTHTTKRQKYRNIKANPA
VAMSVIDPDNPYRYLEVRGLVEDIVPDPTGAFYLKLNDRYDGPLTEPPADKADRVIIVVRPT
AFSKQ
111000000000000000000000000000000000000000000000000000000000
000000000000000000000000000000000000000000000000000000000000
00000
>DM_train2317
GMKQDSIITLHPDDRSEQTAEIMRRFNDVFQLHDPAALPELIAEECVIENTVPAPDGARHAG
RQACVQLWSAIATQPGTRFDLEETFVAGDRATIRWRYWMADGNSVRGVNLMRVQDGRIVEAM
GYVKG
111111100000000000000000000000000000000000000000000000000000
000000000000000000000000000000000000000000000000000000000000
00000
>DM_train2318
EDSPLDALDLVWAKCRGYPSYPALIIDPKMPREGMFHHGVPIPVPPLEVLKLGEQMTQEARE

HLYLVLFFDNKRTWQWLPRTKLVPLGVNQDLDKEKMLEGRKSNIRKSVQIAYHRALQHRSKV
QGEQS
000000000000000000000000000000000000000000000000000000000000000
000000000000000000000000000000000000000000000000000000000000000
00011
>DM_train2319
MEYKSSPKRPYLLRAYYDWLVDNSFTPYLVVDATYLGVNPVEYVKDGQIVLNLSASATGNL
QLTNDFIQFNARFKGVSRELYIPMGAALAIYARENGDGVMFEPEEIYDELNIEPDTEQPTGF
YEAVD
111000000000000000000000000000000000000000000000000000000000000
000000000000000000000000000000000000000000000000000001111111111
11111
>DM_train2320
GSRRASVGSHEVEKSGLLNMTKIAQGGRKLRKNWGPSWVVLTGNSLVFYREPPPTAPSSGWG
PAGSRPESSVDLRGAALAHGRHLSSRRNVLHIRTIPGHEFLLQSDHETELRAWHRALRTVIE
RLVRW
111111110000000000000000000000000000000000000000000001111111111
111100000000000000000000000000000000000000000000000000000000000
00000
>DM_train2321
MTKVDFWPTLKDAYEPLYPQQLEILRQQVVSEGGPTATIQSRFNYAWGLIKSTDVNDERLGV
KILTDIYKEAESRRRECLYYLTIGCYKLGEYSMAKRYVDTLFEHERNNKQVGALKSMVEDKI
QKETL
111100000000000000000000000000000000000000000000000000000000000
000000000000000000000000000000000000000011111111111111111111111
11111
>DM_train2322
EEITDGVNNMNLATDSQKKNRIQVSNTKKPLFFYVNLAKRYMQQYNDVELSALGMAIATVVT
VTEILKNNGFAVEKKIMTSIVDIKDDARGRPVQKAKIEITLVKSEKFDELMAAANEEKEDAE
TQVQN
111111111111111110000000000000000000000000000000000000000000000
000000000000000000000000111111000000000000000000000000000011111111
11111
>DM_train2323
MDFITDMSKNQRLELQNRLAQYETSLMVMSHNGDVPVITGFNVMRVTTMLDALKVELPAVAV
LGDDAQDLAYVFGARPLAVGVNIIRVVDVPGQQPSALVDAELGALHEVSMVRVLNDIADEQL
VKANM
111111111111110000000000000000000000000000000000000000001100000
000000000000000000000000000000000000000000000000000000000000000
00000
>DM_train2324
KPQAPELRIFPKKMDAELGQKVDLVCEVLGSVSQGCSWLFQNSSSKLPQPTFVVYMASSHNK
ITWDEKLNSSKLFSAMRDTNNKYVLTLNKFSKENEGYYFCSVISNSVMYFSSVVPVLQKVSS
ALVP

000000000000000000000000000000000000000000000000000000000000000
000000000000000000000000000000000000000000000000000000000001111
1111
>DM_train2325
RKKPHIVISMPQDFRPVSSIIDVDILPETHRRVRLCKYGTEKPLGFYIRDGSSVRVTPHGLE
KVPGIFISRLVPGGLAQSTGLLAVNDEVLEVNGIEVSGKSLDQVTDMMIANSRNLIITVRPA
NQRN
111110000000000000000000000000000000000000000000000000000000000
000000000000000000000000000000000000000000000000000000000000000
0000
>DM_train2326
MACAAARSPADQDRFICIYPAYLNNKKTIAEGRRIPISKAVENPTATEIQDVCSAVGLNVFL
EKNKMYSREWNRDVQYRGRVRVQLKQEDGSLCLVQFPSRKSVMLYAAEMIPKLKTRTQLEHH
HHHH
111100000000000000000000000000000000000000000000000000000000000
000000000000000000000000000000000000000000000000000000000111111
1111
>DM_train2327
GMENELEDKILAILEQHQVGVLTSVQGDFPHARYMTFLHDGLTLYTPSGKELPKTEEVRRNP
HVCVLIGYDSPGSAFLEINGLASLEEDESIKERIWENISKDWFQGEDSPSFVVIKIVPEQIR
ILNS
100000000000000000000000000000000000000000000011110000000000
000000000000000000000000000000000000000000001111110000000000000
0001
>DM_train2328
GAGDVVTRDVNKLPVAAREMIGKHFSQTKVAYIKIEKDLFQTTSYDVKLADGIELEFNSKGE
WLEIDCKNKSVPSTFIPQAISKYMKANYNGHKTVKIERNRKGYELTLENGLEVDFDQFGGFL
KLSD
110000000000000000000000000000000000000000000000000000000000000
000000000000000000000000000000000000000000000000000000000000000
0000
>DM_train2329
MTCAYRREIHHAHVAIRDWLAGDSRADALDALMARFAEDFSMVTPHGVVLDKTALGELFRSK
GGTRPGLRIEIDGESLLASGVDGATLAYREIQSDAAGRSERLSTVVLHRDDEGRLYWRHLQE
TFCG
110000000000000000000000000000000000000000000000000000000000000
000000000000000000000000000000000000000000000000000000000000000
0000
>DM_train2330
GSHMAKHEQILVLDPPSDLKFKGPFTDVVTTNLKLQNPSDRKVCFKVKTTAPRRYCVRPNSG
VIDPGSIVTVSVMLQPFDYDPNEKSKHKFMVQTIFAPPNISDMEAVWKEAKPDELMDSKLRC
VFEM
110000000000000000000000000000000000000000000000000000000000000
000000000000000000000000000000000000000000000000000000000000000

0000
>DM_train2331
MSHAIQIREAHFPGRAPIDAYGNGGFRFADMSHRGSIICIPSGIYGIDMTGPVPTQEDISRV
LEESDQIEVLLIGTGVELLRLPEELRVLLWEKRISSDTMSTGAAVRTFNVLLAEDRAVAALL
FAVE
11111111110000000000000000000000000000000000000000000000000000
00000000000000000000000000000000000000000000000000000000000000
0000
>DM_train2332
ADSQTSSNRAGEFSIPPNTDFRAIFFANAAEQQHIKLFIGDSQEPAAYHKLTTRDGPREATL
NSGNGKIRFEVSVNGKPSATDARLAPINGKKSDGSPFTVNFGIVVSEDGHDSDYNDGIVVLQ
WPIG
11110000000000000000000000000000000000000000000000000000000000
00000000000000000000000000000001100000000000000000000000000000
0000
>DM_train2333
GMTNTTYVQEYHAIVEVLSKYNEGGKKADSTIMRPAFSSQATIFGVDVDNKLTGGPIQGLFD
VIDNVFHPSPEAKAAIARIDIVGTAASARIDTDDISGFRFTDFFNLLKVEGKWTVVSKIYHT
HPSA
11100000000000000000000000000000000000000000000000000000000000
00000000000000000000000000000000000000000000000000000000000000
0011
>DM_train2334
APARSPSPSTQPWEHVNAIQEARRLLNLSRDTAAEMNETVEVISEMFDLQEPTCLQTRLELY
KQGLRGSLTKLKGPLTMMASHYKQHCPPTPETSCATQIITFESFKENLKDFLLVIPFDCWEP
VQE
11100000000000000000000000000000000000000000000000000000000000
00000000000000000000000000000000000000000000000000000000000001
111
>DM_train2335
GSHMGSLDETTYERLAEETLDSLAEFFEDLADKPYTFEDYDVSFGSGVLTVKLGGDLGTYVI
NKQTPNKQIWLSSPSSGPKRYDWTGKNWVYSHDGVSLHELLAAELTKALKTKLDLSSLAYSG
KDA
11111100000000000000000000000000000000000000000000000000000000
00000000000000000000000000000000000000000000000000000000000000
000
>DM_train2336
GENIVCRVICTTGQIPIRDLSADISQVLKEKRSIKKVWTFGRNPACDYHLGNISRLSNKHFQ
ILLGEDGNLLLNDISTNGTWLNGQKVEKNSNQLLSQGDEITVGVGVESDILSLVIFINDKFK
QCL
11100000000000000000000000000000000000000000000000000000000000
00000000000000000000000000000000000000000000000000000000000000
000
>DM_train2337

MGDKPIWEQIGSSFIQHYYQLFDNDRTQLGAIYIDASCLTWEGQQFQGKAAIVEKLSSLPFQ
KIQASITAQDHQPTPDSCIISMVVGQLKADEDPIMGFHQMFLLKNINDAWVCTNDMFRLALH
NFG
111000000000000000000000000000000000000000000000000000000000
000000000000000000000000000000000000000000000000000000000001
111
>DM_train2338
KVLTIKSCNIHSGIGIRPHAQIELEYQGKIHKEISEGDGGYDAFMNALTKITNRLGISIPKL
IDYEVRIPPGGKTDALVETRITWNKSLDLEEDQTFKTMGVHPDQTVAAVHATEKMLNQILQP
WQI
000000000000000000000000000000000000000000000000000000000000
000000000000000000000000111111100000000000000000000000000001
111
>DM_train2339
QEQRMSHHYATIEVSQQLRQLLGDQLVILLRETPDGQALERSQNDFRRVLEQGRANTVDSAE
QAALDGVRDAYLQLQAHTPALLEAPMADNDGFSEAFNGLRLRLQDLQQLALAGISEAETSAR
HRA
110000000000000000000000000000000000000000000000000000000000
000000000000000000000000000000000000000000000000000000011111
111
>DM_train2340
GAMGSGDTQLFNRAVSMVEKNKDIRSLLQCDDGITGKERLKAYGELITNDKWTRNRPIVSTK
KLDKEGRTHHYMRFHVESKKKIALVHLEAKESKQNYQPDFINMYVDVPGEKRYYLIKPKLHP
VSN
111100000000000000000000000000000000000000000000000000000000
000000000000000000000000000000000000000000000000000000000000
000
>DM_train2341
GEEEKKRTARTDYWLQPEIIVKIITKKLGEKYHKKKAIVKEVIDKYTAVVKMIDSGDKLKLD
QTHLETVIPAPGKRILVLNGGYRGNEGTLESINEKTFSATIVIETGPLKGRRVEGIQYEDIS
KLA
111111000000000000000000000000000000000000000000000000000000
000000000000000000000000000000000000000000000000000000000000
000
>DM_train2342
SNAMNLKREQEFVSQYHFDARNFEWENENGAPETKVDVNFQLLQHDQENQVTSLIVILSFMI
VFDKFVISGTISQVNHIDGRIVNEPSELNQEEVETLARPCLNMLNRLTYEVTEIALDLPGIN
LEF
110000000000000000000000000000000000000000000000000000000000
000000000000000000000000000000000000000000000000000000000000
000
>DM_train2343
MSDTLPGTTPPDDNHDRPWWGLPCTVTPCFGARLVQEGNRLHYLADRAGIRGRFSDVDAYHL
DQAFPLLMKQLELMLTGGELNPRHQHTVTLYAKGLTCEADTLGSCGYVYLAVYPTPAAPATT

VLE
111000000000000000000000000000000000000000000000000000000000
000000000000000000000000000000000000000000000000000000011111
111
>DM_train2344
GMIKKEGPGWRIIFDSSRDNFSTLIGGETWAIELDKSEWKILVEVVMELCDQYKLVKEQLMG
DEDITLELERRPWLAILNGDQYGWNLRLILSASGLFNRGAEVYWPRHVTNNVVNAMRSMWDS
DYL
000000000000000000000000000000000000000000000000000000000000
000000000000000000000000000000011100000000000000000000000001
111
>DM_train2345
ELLFNKTKSVEFTFGNDTVVIPCFVTNMEAQNTTEVYVKWKFKGRDIYTFDGALNKSTVPTD
FSSAKIEVSQLLKGDASLKMDKSDAVSHTGNYTCEVTELTREGETIIELKYRVVSWSTRHHH
HHH
000000000000000000000000000000000000000000000000000000000000
000000000000000000000000000000000000000000000000000011111111
111
>DM_train2346
MEHVPAELTRVSEGYTRFYRSPTASVILSGLVKVKWDNEQMTMPLFKWIGGEQAEELHFCVH
IAHSSGPKLNRARSLGTVNSNMDQHWAQAQRNSGATRRTIEGFHLFENDIPNFPDYIKIKLV
PKT
111111111111000000000000000000000000000000000000000000000000
000000000000000000000000000000000000000000000000000000000000
000
>DM_train2347
MSLKFPDTGLEEKEVAFSIVNHAAKSLGFIHVDQWDYERVMFDYKIVHHEGTFYLRVPAYAV
KGEIPRPSTIVQIMTPILGKYYYPHGVEYEGETFPQAVIDKCNNKLALLAKTIKAEWEGHHH
HHH
100000000000000000000000000000000000000000000000000000000000
000000000000000000000000000000000000000000000000000011111111
111
>DM_train2348
PRTQFSGLRRELPPSVRLLTLARWGPETLLLRLEHQFAVGEDSGRNLSSPVTLDLTNLFSAF
TITNLRETTLAANQLLAYASRLQWTTDTGPTPHPSPSRPVSATITLQPMEIRTFLASVQWEE
DG
000000000000000000000000000000000000000000000000000000000000
000000000000000000000000001111111111111100000000000000000011
11
>DM_train2349
MTDQATPNLPSRDFDSTAAFYERLGFGIVFRDAGWMILQRGDLMLEFFAHPGLDPLASWFSC
CLRLDDLAEFYRQCKSVGIQETSSGYPRIHAPELQEWGGTMAALVDPDGTLLRLIQNELLAG
IS
000000000000000000000000000000000000000000000000000000000000

000000000000000000000000000000000000000000000000000000000001111
11
>DM_train2350
MQTGRTEDDLVPTAPSLGTKEGYLTKQGGLVKTWKTRWFTLHRNELKYFKDQMSPEPIRILD
LTECSAVQFDYSQERVNCFCLVFPFRTFYLCAKTGVEADEWIKILRWKLSQIRKQLNQGEGT
IR
11111111111111000000000000000000000000000000000000000000000000
000000000000000000000000000000000000000000000000001111111111
11
>DM_train2351
SHMPASVQLHTAVEMHHWCIPFSVDGQPAPSLRWLFNGSVLNETSFIFTEFLEPAANETVRH
GCLRLNQPTHVNNGNYTLLAANPFGQASASIMAAFMDNPFEFNPEDPIPDTNSTSGDPVEKK
DE
00000000000000000000000000000000000000000000000000000000000000
00000000000000000000000000000000000000000000001111111111111111
11
>DM_train2352
KAMHVAQPAVVLASSRGIASFVCEYASPGKATEVRVTVLRQADSQVTEVCAATYMMGNELTF
LDDSICTGTSSGNQVNLTIQGLRAMDTGLYICKVELMYPPPYYLGIGNGTQIYVIDPEPCPD
SD
00000000000000000000000000111100000000000001110000000000000000000
000000000000000000000000000000000000000000000000000000000001111
11
>DM_train2354
MRHISPEELIALHDANISRYGGLPGMSDPGRAEAIIGRVQARVAYEEITDLFEVSATYLVAT
ARGYIFNDANKRTALNSALLFLRRNGVQVFDSPELADLTVGAATGEISVSSVADTLRRLYGS
AD
00000000000000000000000001000000000000000000000000000000000000
00000000000000000000000000000000000000000000000000000000000001
11
>DM_train2355
GKPDISAKDLRNIMYDHLPGFGTAFHQLVQVICKLGKDSNSLDIIHAEFQASLAEGDSPQCA
LIQITKRVPIFQDAAPPVIHIRSRGDIPRACQKSLRPVPPSPKIDRGWVCVFQLQDGKTLGL
KI
11100000000000000000000000000000000000000000000000000000000000
00000000000000000000000000000000000000000000000000000000000000
00
>DM_train2356
GMGFLDALMGNASEVDLGKLAAELSPILGDNEELQLAYKMVRDLFVFTSKRLILIDKQGVTG
KKVSYHSIPYKAIVHFQVETAGTFDMDAELKLWISGQHEPLVKELKRGTDVVGIQKTIARYA
LG
11111111111111000000000000000000000000000000000000000000000000
00000000000000000000000000000000000000000000000000000000000000
00

>DM_train2357
RVSVTDYKRLLDSGAFHLLLDVRPQVEVDICRLPHALHIPLKHLERRDAESLKLLKEAIWEE
KQGTQEGAAVPIYVICKLGNDSQKAVKILQSLSAAQELDPLTVRDVVGGLMAWAAKIDGTFP
QY
00000000000000000000000000000000000000000000000000000000000000
00001110000000000000000000000000000000000000000000000000000000
00
>DM_train2358
THSQLDQLLTGLVDRVAEVDHAVVLSEDGLVVSKSTGFLRDDAERLAATASGLMSLSKGVSM
DFRRGPVRQALIEMGKGYLILTAAGPGAHLVVLTRQGADVGVVAYQMNMLVKKIGEHLSAPP
RG
10000000000000000000000000000000000000000000000000000000111111
11111110000000000000000000000000000000000000000000000011111111
11
>DM_train2359
GAPSQGATANAETQTTIPLGDTSQNALDWPGVYEGVLPCASCEGIQTTLTLQADNSFELKSI
YLGKDESIFKVAGKFDWDSNGSKITLSDGSKYLVGENQLLMLDTEGNRITGGLAEHYILKKK
GM
11111111111111111110000000000000000000000000000000000000000000
00000000000000000000000000000000000000000000000000000000000000
00
>DM_train2360
PTMEGPLRRKTLLKEGRKPALSSWTRYWVVLSGATLLYYGAKSLRGTDRKHYKSTPGKKVSI
VGWMVQLPDDPEHPDIFQLNNPDKGNVYKFQTGSRFHAILWHKHLDDACKSSRPQVPANLMS
FE
00000000000000000000000000000000000000000000000000000000000000
00000000000000000000000000000000000000000000000000000111111111
11
>DM_train2361
MPAIFTHEGKVEGVPGNYPLTAENLFRIGLALCTLWILDKEIEEPTLSIPETNFVTLALSVG
FMNAGGSVNVGKGGDIKLFLQKGEIYVLEFQPLSETDIKKLESILFGRAPIPKKTGEDIGSF
KC
11000000000000000000000000000000000000000000000000000000000000
00000000000000000000000000000000000000000000000000000000000000
00
>DM_train2362
MAFSGTWQVYAQENYEEFLRAISLPEEVIKLAKDVKPVTEIQQNGSDFTITSKTPGKTVTNS
FTIGKEAEITTMDGKKLKCIVKLDGGKLVCRTDRFSHIQEIKAGEMVETLTVGGTTMIRKSK
KI
10000000000000000000000000000000000000000000000000000000000000
00000000000000000000000000000000000000000000000000000000000000
00
>DM_train2363
DIEKEILDLAAATERLNLTDALNSNPAGNLYDWRSSNSYPWTQKLNLHLTITATGQKYRILA

SKIVDFNIYSNNFNNLVKLEQSLGDGVKDHYVDISLDAGQYVLVMKANSSYSGNYPYSILFQ
KF
111000000000000000000000000000000000000000000000000000000000
000000000000000000000000000000000000000000000000000000000000
00
>DM_train2364
MPYKNILTLISVNNDNFENYFRKIFLDVRSSGSKKTTINVFTEIQYQELVTLIREALLENID
IGYELFLWKKNEVDIFLKNLEKSEVDGLLVYCDDENKVFMSKIVDNLPTAIKRNLIKDFCRK
LS
110000000000000000000000000000000000000000000000000000000000
000000000000000000000000000000000000000000000000000000000000
00
>DM_train2365
MSDSHHKPVWDRTHHAKMATGIGDPQCFKGMAGKSKFNVGDRVRIKDLPDLFYTRTMTYTRG
ATGTIVRLVYESPAAEDEAFGNEENVEWFYSIVFAQKDLWPEYSDTFANDTLETEIPERYLE
KA
111111100000000000000000000000000000000000000000000000000000
000000000000000000000000000000000000000000000000000000000000
00
>DM_train2366
MTNKQVEISMAEWDVMNIIWDKKSVSANEIVVEIQKYKEVSDKTIRTLITRLYKKEIIKRYK
SENIYFYSSNIKEDDIKMKTAKTFLNKLYGGDMKSLVLNFAKNEELNNKEIEELRDILNDIS
KK
111100000000000000000000000000000000000000000000000000000000
000000000000000000000000000000000000000000000000000000000000
00
>DM_train2367
MKDPLLNSLIYVSRYYGLANSPEALVNGLPLSDGKLTPFLLPRAAERAGLVAKENRAELEKI
SSLILPAILVLKGGDSCVLNSINMETREAEVTTLESGMVPISIPLEDLLEQYTGRYFLVKKQ
FR
000000000000000000000000000000000000000000000000000000000000
000000000000000000000000000000000000000000000000000000000000
11
>DM_train2368
GSRESTGYLEELEKERSLLLADLDKEEKEKDWYYAQLQNLTKRIDSLPLTENFSLQTDMTRR
QLEYEARQIRVAMEEQLGTCQDMEKRAQRRIARIQQIEKDILRIRQLLQSQATEAERSSQNK
H
111100000000000000000000000000000000000000000011111000000000
000000000000000000000000000000000000000000000000001111111111
1
>DM_train2369
TTEENSKSEALLDIPMLEQYLELVGPKLITDGLAVFEKMMPGYVSVLESNLTAQDKKGIVEE
GHKIKGAAGSVGLRHLQQLGQQIQSPDLPAWEDNVGEWIEEMKEEWRHDVEVLKAWVAKATK
K

1111110000000000000000000000000000000000000000000000000000000000
0000000000000000000000000000000000000000000000000000000000000000
0
>DM_train2371
RAKQQAAKSERRVRELTEATNDILWEFTADLSEVLVINSAYEDIWGRSVAKLRENPHDFLNG
IHPEDRELMKDTMQSLMDGESADVECRVNATEEYQRWVWIQGEPITNDAGETVRVAGFARDI
T
1111111110000000000000000000000000000000000000000000000000000000
0000000000000000000000000000000000000000000000000000000000000000
0
>DM_train2372
NLYFQGHMRKIFLACPYSHADAEVVEQRFRACNEVAATIVRAGHVVFSQVSMSHPINLCLAE
LDRAAIGRLWAPVDAFYMDHLEELIVLDLPGWRDSAGIRREMEFFEAGGQRVSLWSEVEHEF
R
1111110000000000000000000000000000000000000000000000000000000000
0000000000000000000000000000000000000000000000000000000000000000
0
>DM_train2373
MGSDKIHHHHHHMFPYKIVDDVVILMPNKELNIENAHLFKKWVFDEFLNKGYNKIFLVLSDV
ESIDSFSLGVIVNILKSISSSGGFFALVSPNEKVERVLSLTNLDRIVKIYDTISEAMEEVRR
K
1111111000000000000000000000000000000000000000000000000000000000
0000000000000000000000000000000000000000000000000000000000000000
1
>DM_train2374
AECSVDIQGNDQMQFNTNAITVDKSCKQFTVNLSHPGNLPKNVMGHNWVLSTAADMQGVVTD
GMASGLDKDYLKPDDSRVIAHTKLIGSGEKDSVTFDVSKLKEGEQYMFFCAAHAAMKGTLTL
K
0000000000000000000000000000000000000000000000000000000000000000
0000000000000000000000000000000000000000011100000000000000000000
0
>DM_train2375
GMGVNMRAETESRIFSVDEYVRPSNGEPIRSVVLETNDSVVVVWHAHPGQEIASHVHPHGQD
TWTVISGEAEYHQGNGIVTHLKAGDIAIAKPGQVHGAMNSGPEPFIFVSVVAPGNAGFALAE
K
1111111111000000000000000000000000000000000000000000000000000000
0000000000000000000000000000000000000000000000000000000000000000
0
>DM_train2376
DVTPRRDAEYPPPEFLEAMKPLREICIKKTGVTEEAIIEFSDGKVHEDENLKCYMNCLFHEA
KVVDDTGHVHLEKLHDALPDSMHDIALHMGKRCLYPEGENLCEKAFWLHKCWKESDPKHYFL
I
1111111110000000000000000000000000000000000000000000000000000000
0000000000000000000000000000000000000000000000000000000000000000

0
>DM_train2377
DLYFNPRFLADDPQAVADLSRFENGQELPPGTYRVDIYLNNGYMATRDVTFNTGDSEQGIVP
CLTRAQLASMGLNTASVAGMNLLADDACVPLTTMVQDATAHLDVGQQRLNLTIPQAFMSNRA
R
00000000011111111100000000000000000000000000000000000000000000
00000000000000000000000000000000000000000000000000000000000000
0
>DM_train2378
MRVEDVMVTDVDTIDITASLEDVLRNYVENAKGSSVVVKEGVRVGIVTTWDVLEAIAEGDDL
AEVKVWEVMERDLVTISPRATIKEAAEKMVKNVVWRLLVEEDDEIIGVISATDILRAKMAKR
Y
10000000000000000000000000000000000000000000000000000000000000
00000000000000000000000000000000000000000000000000000000000111
1
>DM_train2379
GSSGSSGEKITKVYELGNEPERKLWVDRYLTFMEERGSPVSSLPAVGKKPLDLFRLYVCVKE
IGGLAQVNKNKKWRELATNLNVGTSSSAASSLKKQYIQYLFAFECKIERGEEPPPEVFSTGD
T
11111100000000000000000000000000000000000000000000000000000000
00000000000000000000000000000000000000000000000000000000001111
1
>DM_train2380
ASENIEVHMLNKGAEGAMVFEPAYIKANPGDTVTFIPVDKGHNVESIKDMIPCGACKFKSKI
NENYVLTVTQPGAYLVKCTPHYAMGMIALIAVGDSPANLDQIVSAKKPKIVQERLEKVIASA
K
00000000000000000000000000000000000000000000000000000000000000
00000000000000000000000000000000000000000000000000000000000011
1
>DM_train2381
MSLSIGRTCWAIAEGYIPPYGNGPEPQFISHETVCILNAGDEDAHVEITIYYSDKEPVGPYR
LTVPARRTKHVRFNDLNDPAPIPHDTDFASVIQSNVPIVVQHTRLDSRQAENALLSTIAYAN
T
11100000000000000001111111111110000000000000000000000000000000
00000000000000000000000000000000000000000001111111111111111111
1
>DM_train2382
SNAMALGFGMKMELQQFLDALASSPEKIEFETTMAVIEDNYDFTPAAFTNGNTQNDANENNG
SCKIFAFGLLNALDKEATLACFGRFYREDVLLHPENNDHQNIRNFMVTGWEGIQFETSALTA
K
11111000000000000000000000000000000000000000000000000000000000
00000000000000000000000000000000000000000000000000000000000000
0
>DM_train2383

MTGRIVHFEIPFDDGDRARAFYRDAFGWAIAEIPDMDYSMVTTGPVGESGMPDEPGYINGGM
MQRGEVTTPVVTVDVESIESALERIESLGGKTVTGRTPVGNMGFAAYFTDSEGNVVGLWETA
R
00000000000000000000000000000000011000000000000000000000000000
00000000000000000000000000000000000000000000000000000000000000
0
>DM_train2384
GPLGSMPSEKTFKQRRSFEQRVEDVRLIREQHPTKIPVIIERYKGEKQLPVLDKTKFLVPDH
VNMSELIKIIRRRLQLNANQAFFLLVNGHSMVSVSTPISEVYESERDEDGFLYMVYASQETF
G
11111111100000000000000000000000000000000000000000000000000000
00000000000000000000000000000000000000000000000000000000000000
0
>DM_train2385
ESVKAMWGVVTDSQTEIVALAKVRNEDVVPIVVSGYHYTIEMNGVKVADGYENSPVTVKPAS
ATTLKFSLRLNNSFLREWWVTHIANGEKTKIRVAIKPTIEIGGRDVEVPVFLRESEFTTKLL
S
00000000000000000000000000000000000000000000000000000000000000
00000000000000000000000000000000000000000010000000000000000000
1
>DM_train2386
IVVYTDREVYGAVGSQVTLHCSFWSSEWVSDDISFTWRYQPEGGRDAISIFHYAKGQPYIDE
VGTFKERIQWVGDPSWKDGSIVIHNLDYSDNGTFTCDVKNPPDIVGKTSQVTLYVFEKVPTR
00000000000000000000000000000000000000000000000000000000000000
00000000000000000000000000000000000000001111000000000000011111
>DM_train2387
SGPWMCYPGQAFQVPALPACRPLLRLQCNGSQVPEAVLRDCCQQLAHISEWCRCGALYSMLD
SMYKEHGAQEGQAGTGAFPRCRREVVKLTAASITAVCRLPIVVDASGDGAYVCKDVAAYPDA
11100000000000000000000000000000000000000000000000000000000000
00000011111111100000000000000000000000000000000000000000000000
>DM_train2388
TESGTAAQDPEALAAEIGPVKQVSLGEQIDAALAQQGEQLFNTYCTACHRLDERFIGPALRD
VTKRRGPVYIMNVMLNPNGMIQRHPVMKQLVQEYGTMMTDMALSEEQARAILEYLRQVAENQ
11111111000000000000000000000000000000000000000000000000000000
00000000000000000000000000000000000000000000000000000000000000
>DM_train2389
LRPKALTQVLSQANTGGVQSTLLLNNEGSLLAYSGYGDTDARVTAAIASNIWAAADRNGNQA
FNEDSLKFILMDCMEGRVAITRVANLLLCMYAKETVGFGMLKAKAQALVQYLEEPLTQVAAS
00000000000000000000000000000000000000000000000000000000000000
00000000000000000000000000000000000000000000000000000011111111
>DM_train2390
GLPVEVRGSNGAFYKGFVKDVHEDSVTIFFENNWQSERQIPFGDVRLPPPADYNKEITEGDE
VEVYSRANEQEPCGWWLARVRMMKGDFYVIEYAACDATYNEIVTLERLRPVNPNPLATKGSF
00000000000000000000000000000000000000000000000000011110000000

000000011100000000000000000000000011110000000000000000011111
>DM_train2391
MSWIEPIISHFCQDLGVPTSSPLSPLIQLEMAQSGTLQLEQHGATLTLWLARSLAWHRCEDA
MVKALTLTAAQKSGALPLRAGWLGESQLVLFVSLDERSLTLPLLHQAFEQLLRLQQEVLAPP
100000000000000000000000000000000000000000000000000000000000
000000000000000000000000000000000000000000000000000000000011
>DM_train2392
GHMSDLVTKFESLIISKYPVSFTKEQSAQAAQWESVLKSGQIQPHLDQLNLVLRDNTFIVST
LYPTSTDVHVFEVALPLIKDLVASSKDVKSTYTTYRHILRWIDYMQNLLEVSSTDKLEINHD
111000000000000000000011111111000000000000000000000000000000
000000000000000000000000000000000000000000000000000000000011
>DM_train2393
MGGSGTDIVLNDLPFVDGPPAEGQSRISWIKNGEEILGADTQYGSEGSMNRPTVSVLRNVEV
LDKNIGILKTSLETANSDIKTIQEAGYIPEAPRDGQAYVRKDGEWVLLSTFLGSSGSGLVPR
111111000000000000000000000000000000000000000000000000000000
000000000000000000000000000000000000000000000000001111111111
>DM_train2394
ALEGTEFARSEGASALASVNPLKTTVEEALSRGWSVKSGTGTEDATKKEVPLGVAADANKLG
TIALKPDPADGTADITLTFTMGGAGPKNKGKIITLTRTAADGLWKCTSDQDEQFIPKGCSR
111000000000000000000000000000000000000000000000000000000000
00000000000000000000000000000000000000000000000000000000000
>DM_train2395
AITKINDCFELLSMVTYADKLKSLIKKEFSISFEEFAVLTYISENKEKEYYLKDIINHLNYK
QPQVVKAVKILSQEDYFDKKRNEHDERTVLILVNAQQRKKIESLLSRVNKRITEANNEIEL
000000000000000000000000000000000000000000000000000001111111
11100000000000000000000000000000000000000000001111111111111
>DM_train2398
KACTRECGNLGFGICPRSEGSPLNPICINCCSGYKGCNYYNSFGKFICEGESDPKRPNACTF
NCDPNIAYSRCPRSQGKSLIYPTGCTTCCTGYKGCYYFGKDGKFVCEGESDEPKANMYPVM
000000000000000000000000000000000000000000000000000000000000
00000000001111111111110000000000000000000000000000001111111
>DM_train2399
MPHLVIEATANLRLETSPGELLEQANAALFASGQFGEADIKSRFVTLEAYRQGTAAVERAYL
HACLSILDGRDAATRQALGESLCEVLAGAVAGGGEEGVQVSVEVREMERASYAKRVVARQR
000000000000000000000000000000000000000000000000000000000000
00000000000000000000000000000000000000000000000000000000011
>DM_train2400
NALRRDPNLPVHIRGWLHKQDSSGLRLWKRRWFVLSGHCLFYYKDSREESVLGSVLLPSYNI
RPDGPGAPRGRRFTFTAEHPGMRTYVLAADTLEDLRGWLRALGRASRAEGDDYGQPRSPAR
110000000000000000000000000000000000000000000000000000000000
00000000000000000000000000000000000000000000011111111111111
>DM_train2401
GSHMVGQLSEGAIAAIMQKGDTNIKPILQVINIRPITTGNSPPRYRLLMSDGLNTLSSFMLA
TQLNPLVEEEQLSSNCVCQIHRFIVNTLKDGRRVVILMELEVLKSAEAVGVKIGNPVPYNE

111000000000000000000000000000000000111000000000000000000000
00000000000000000000000000000000000000000000000000000000000
>DM_train2402
GHMNSFVGLRVVAKWSSNGYFYSGKITRDVGAGKYKLLFDDGYECDVLGKDILLCDPIPLDT
EVTALSEDEYFSAGVVKGHRKESGELYYSIEKEGQRKWYKRMAVILSLEQGNRLREQYGLG
111100000000000000000000000000000000000000000000000000000000
00000000000000000000000000000000000000000000000000000000000
>DM_train2403
GMGSLAEWYQRIPTPDDLTRVESLFANMQAQFPQLKLEFKWNQPMFTDHGTFIMGFNPSKKH
LAVAIEPQTMTRFIPQIDKAGYDHSQIIRFPWHKPLDEQLIHDLIAYTIDQKKDATTFWQR
110000000000000000000000000000000000000000000000000000000000
00000000000000000000000000000000000000000000000000000000000
>DM_train2404
MSKLDLHQMTTQDLVALFAKVTVEQDDALLGNQISRFNRLFGVMAEIADELKARDGDQRTAL
LSLFEYPNMQVRLQAAKLTLAVAPVKAREQLEAIVSSKWFPQAGDAGMCLDLLDDGTFKPK
110000000000000000000000000000000000000000000000000000000000
00000000000000000000000000000000000000000000000000000000000
>DM_train2405
MRNYIQGIDHVQVAAPVGCEEEARAFYGETIGMEEIPKPEELKKRGGCWFKCGNQEIHIGVE
QNFNPAKRAHPAFYVLKIDEFKQELIKQGIEVIDDHARPDVIRFYVSDPFGNRIEFMENKN
000000000000000000000000000000000000000000000000000000000000
00000000000000000000000000000000000000000000000000000000001
>DM_train2406
GDFAKEYADALYDSLGHSVLICDRDVYIAVSGSSKKDYLNKSISEMLERTMDQRSSVLESDA
KSVQLVNGIDEDMNSYTVGPIVANGDPIGAVVIFSKDQTMGEVEHKAVETAAGFLARQMEQ
100000000000000000000000000000000000000000000000000000000000
00000000000000000000000000000000000000000000000000000011111
>DM_train2407
MGARRMNAFDIISGSPGFNLSGLFGDARKYDRVERFVSAWTAERVVERLEEIVSAENLTVAK
KETWGMKIEGQKGNFAMVVEINQLTDELVMIEVRKRQRAAASGRDLWTDTLRPFFVELVHE
111000000000000000000000000000000000000000000000000000000000
00000000000000000000000000000000000000011100000000000000001
>DM_train2408
GMPQHDKSRLVRIDTGPMINPVAGKPSRPIAGDASFRTVTAFEGGQGKVESGVWESTSGSFQ
SNTTGYIEYCHIIEGEARLVDPDGTVHAVKAGDAFIMPEGYTGRWEVDRHVKKIYFVTHLA
111100000000000000000000000000000000000000000000000000000000
00000000000000000000000000000000000000000000000000000000001
>DM_train2409
PMFIVNTNVPRASVPDGFLSELTQQLAQATGKPPQYIAVHVVPDQLMAFGGSSEPCALCSLH
SIGKIGGAQNRSYSKLLCGLLAERLRISPDRVYINYYDMNAANVGWNNSTFALEHHHHHH
000000000000000000000000000000000000000000000000000000000000
0000000000000000000000000000000000000000000000000011111111
>DM_train2410
MEGSHHHHHHSQAQSKAQVLQSVAGQTLTVRCQYPPTGSLYEKKGWCKEASALVCIRLVTSS

KPRTMAWTSRFTIWDDPDAGFFTVTMTDLREEDSGHYWCRIYRPSDNSVSKSVRFYLVVS
11111111111110000000000000000000000000000000000000000000000000
000000000000000000000000000000000000000000000000000000000000
>DM_train2411
MVRGIRGAITVEEDTPEAIHQATRELLLKMLEANGIQSYEELAAVIFTVTEDLTSAFPAEAA
RQIGMHRVPLLSAREVPVPGSLPRVIRVLALWNTDTPQDRVRHVYLREAVRLRPDLESAQ
00000000000000000000000000000000000000000000000000000000000000
000000000000000000000000000000000000000000000000000001111111
>DM_train2412
EPYSLTAQQSSLIREDKSNAKLWNEVLASLKDRPASGSPFQLFLSKVEETFQCICCQELVFR
PITTVCQHNVCKDCLDRSFRAQVFSCPACRYDLGRSYAMQVNQPLQTVLNQLFPGYGNGR
11100000000000000000000000000111000000000000000000000000000000
000000000000000000000000000000000000000000000000000000000000
>DM_train2413
GADEEESTPCDNVETDQQTFACAAFNKQVAERELQSAYDELIERMRDQFGDEAGLMSRIEAA
EKVWSQLRDADCKVETHAEQPGSNAYQIAWNSCIAQRSDERAEYLRSLGSQNSDEPAPED
11111111111111000000000000000000000000000000000000000000000000
000000000000000000000000000000000000000000000000000111111111
>DM_train2414
GMEPITQAFAQQFSREWIDAWNAHDLDAILSHYADGFEMSSPMIVQIAGEPSGRLRGKEQVG
AYWREALRMIPDLHFEWIATLAGVDSVAIHYRGAKGRLALEVFHFGPDRRVVKALAHYAG
11100000000000000000000000000000000000000000000001000000000000
000000000000000000000000000000000000000000000000000000000000
>DM_train2415
MKKKDTCEIFGYDEEKVNRIQGDLQTVDISGVSQILKAIADENRAKITYALCQDEELCVCDI
ANILGVTIANASHHLRTLYKQGVVNFRKEGKLALYSLGDEHIRQIMMIALAHKKEVKVNV
11111111110000000000000000000000000000000000000000000000000000
000000000000000000011111111111111000000000000000000000001111
>DM_train2416
APVLENRRARHDYEILETYEAGIALKGTEVKSLRAGKVDFTGSFARFEDGELYLENLYIAPY
EKGSYANVDPRRKRKLLLHKHELRRLLGKVEQKGLTLVPLKIYFNERGYAKVLLGLARGK
11000000000000000000000000000000000000000000000000000000000001
111111000000000000000000000000000000000000000000000000000000
>DM_train2417
MEKATKRNDEECEVQLNIKRNSKHSAWTGELFKIECPVKYCVHRPNVTWCKHNGTIWVPLEV
GPQLYTSWEENRSVPVFVLHFKPIHLSDNGSYSCSTNFNSQVINSHSVTIHVRERSQNSS
11111111111000000000000000000000000000000000000000000000000000
000000000000000000000000000000000000000000000000000001111111
>DM_train2418
ARAQLSEAMTLASGLKTKVSDIFSQDGSCPANTAATAGIEKDTDINGKYVAKVTTGGTAAAS
GGCTIVATMKASDVATPLRGKTLTLTLGNADKGSYTWACTSNADNKYLPKTCQTATTTTP
00000000000000000000000000000000000000000000000000000000000000
000000000000000000000000000000000000000000000000000000000001
>DM_train2419

MDIVFIEQLSVITTIGVYDWEQTIEQKLVFDIEMAWDNRKAAKSDDVADCLSYADIAETVVS
HVEGARFALVERVAEEVAELLLARFNSPWVRIKLSKPGAVARAANVGVIIERGNNLKENN
00000000000000000000000000000000000000000000000000000000000000
000000000000000000000000000000000000000000000000000001111111
>DM_train2420
SLRVEETEVFKKYFKNLTDRERAVFEGGITLGALFHQFVGTPVSKYNKESLERAIEEAMKNQ
PCVYDIKVKIRNVGEKYVSLDGKMLDVDLKIKINKTVAHLKLEYIPEIDYPLMYVKKFEE
00000000000000000000000000000000000000000000000000000000000000
000000000000000000000000000000000000000000000000000000000001
>DM_train2421
MSKTIVLSVGEATRTLTEIQSTADRQIFEEKVGPLVGRLRLTASLRQNGAKTAYRVNLKLDQ
ADVVDSGLPKVRYTQVWSHDVTIVANSTEASRKSLYDLTKSLVATSQVEDLVVNLVPLGR
10000000000000000000000000000000000000000000000000000000000000
000011110000000000000000000000000000000000000000000000000000
>DM_train2422
NGGVGVDVELITSINVENDTFIERNFTPQEIEYCSAQPSVQSSFAGTWSAKEAVFKSLGVKS
LGGGAALKDIEIVRVNKNAPAVELHGNAKKAAEEAGVTDVKVSISHDDLQAVAVAVSTKK
10000000000000000000000000000000000000000000000000000000001111
111111000000001111000000000000000000000000000000000000000001
>DM_train2423
GMPTDDIVQLLKGQEEAWNRGDLDAYMQGYWQNEQLMLISNGKFRNGWDETLAAYKKNYPDK
ESLGELKFTIKEIKMLSNYAAMVVGRWDLKRLKDTPTGVFTLLVEKIDDRWVITMDHSSD
10000000000000000000000000000000000000000000000000000000000000
000000000000000000000000000000000000000000000000000000000000
>DM_train2424
GMARKRLIIEMGMGIDQHGQEPTIAASRAVRNAIAHNALPGVWEVAGLSHPNEMIIEVQVAV
PYPEQVREEEVLAVLPFGRKTLTVESGGMIVQGRAIPELNDKNDEMLIAIAAVTVLIENE
00000000000000000000000000000000000000000000000000000000000000
000000000000000000000000000000000000000000000000000000000111
>DM_train2425
QGQFFREIENLKEYFNASSPDVAKGGPLFSEILKNWKDESDKKIIQSQIVSFYFKLFENLKD
NQVIQRSMDIIKQDMFQKFLNGSSEKLEDFKKLIQIPVDDLQIQRKAINELIKVMNDLS
00000000000000001111111000000000000000000000000000000000000000
00000000000000000000000000000000000000000000000000000000000
>DM_train2426
GSHRTSGRVAVEEVDEEGKFVRLRNKSNEDQSMGNWQIKRQNGDDPLLTYRFPPKFTLKAGQ
VVTIWAAGAGATHSPPTDLVWKAQNTWGCGNSLRTALINSTGEEVAMRKLVRSVTVVED
00000000000000000000000000000000000000000000000000000000000000
00000000000000000000000000000000000000000000000000011111111
>DM_train2427
QGVQGSAHAINKAGSLRMQSYRLLAAVPLSEKDKPLIKEMEQTAFSAELTRAAERDGQLAQL
QGLQDYWRNELIPALMRAQNRETVSADVSQFVAGLDQLVSGFDRTTEMRIETEHHHHHH
11110000000000000000000000000000000000000000000000000000000000
00000000000000000000000000000000000000000000000000001111111

>DM_train2428
SVDMMDLPRSRINAGMLAQFIDKPVCFVGRLEKIHPTGKMFILSDGEGKNGTIELMEPLDEE
ISGIVEVVGRVTAKATILCTSYVQFKEDSHPFDLGLYNEAVKIIHDFPQFYPLGIVQHD
10000000000000000000000000000000000000000000000000000000000000
00000000000000000000000000000000000000000000000000000011111
>DM_train2430
DKKAYFYHSSFQILNVEYTEALNSPATHEYRTLSERIEAMITDEFRGSSLKSEFIRTHVVKL
RKEGTGVVADVVMKFRSSKRNNRKVMKTRIQSVLRRLSSSGNLEIAPSNEITSLTDQDT
10000000000000000000000000000000000000000000000000000000000000
00000000000000000111100000000000000000000000000000000000001
>DM_train2431
GMREDMKDNVVKDKSLEFAVRIVNLYKFLVNEQKEFVMSKQILRSGTSIGANIREAEQAQSR
ADFINKLNIALKEANETEYWLELLIRTEYITREQYESINNDSTEINKLLISIIKTTKNN
11111110000000000000000000000000000000000000000000000000000000
00000000000000000000000000000000000000000000000000000111111
>DM_train2432
MRVAIPAEDDRGIKSNVSKHFGRSRYFVFVDIEGEDVKNVEVVEVPFEEHGPGDLPNFIKDH
GAKIVLTYGIGRRAIEYFNSLGISVVTGVYGRISDVIKAFIGGKLKIDYDWKEKIEKEH
00000000000000000000000000000000000000000000000111110000000000
00000000000000000000000000000000000000000000000000000111111
>DM_train2433
MNAPDRFELFLLGEGESKLKIDPDTKAPNAVVITFEKEDHTLGNLIRAELLNDRKVLFAAYK
VEHPFFARFKLRIQTTEGYDPKDALKNACNSIINKLGALKTNFETEWNLQTLAADDAF
00000000000000000000000000000000000000000000000000000000000000
0000000000000000000000000000000000000000000000000000111111
>DM_train2435
NKPKNLDASITSIIHEIGVPAHIKGYLYLREAIAMVYHDIELLGSITKVLYPDIAKKYNTTA
SRVERAIRHAIEVAWSRGNLESISSLFGYTVSVSKAKPTNSEFIAMVADKLRLEHKAS
00000000000000000000000000000000000000000000000000000000000000
0000000000000000000011111111111111000000000000000000001111
>DM_train2436
MEGGKLQSTFVFEEIGRRLKDIGPEVVKKVNAVFEWHITKGGNIGAKWTIDLKSGSGKVYQG
PAKGAADTTIILSDEDFMEVVLGKLDPQKAFFSGRLKARGNIMLSQKLQMILKDYAKL
11111000000000000000000000000000000000000000000000000000000000
0000000000000000000000000000000000000000000000000000000000
>DM_train2437
NPLASLTTDKNDLYINWLKSLSFFQTNSSCAEALVKVIPHYHNKLIDFSQVLQLVFSASEKF
PIQENQPLPEQLMFLSNLEKQTPFAKAVGSSIYKLVTGKNLSLDFASQILKEASILEH
11111111111100000000000000000000000000000000000000000000000000
0000000000000000000000000000000000000000000000000000000001
>DM_train2438
SSPGKPPRLVGGPMDASVEEEGVRRALDFAVGEYNKASNDMYHSRACQVVRARKQIVAGVNY
FLDVELCRTTCTKTQPNLDNCPFHDQPHLKRKAFCSFQIYAVPWQGTMTLSKSTCQDA
11111111111000000000000000000000000000000000000000000000000000

000000000000000110000000000000000000000000000000000000000000
>DM_train2439
MKGMSKMPQFNLRWPGGGPQFNLRWPREVLDLVRKVAEENGRSVNSEIYQRVMESFKKEGRI
GGTGGSGGGREVLDLVRKVAEENGRSVNSEIYQRVMESFKKEGRIGAHHHHHHKNQHE
11111100000000000000000000000000000000000000000000000000000000
011111000000000000000000000000000000000000000011111111111111
>DM_train2440
MVDIVKRRDWEKKEKKKIAIERIDTLFTLAERVARYSPDLAKRYVELALEIQKKAKVKIPRK
WKRRYCKRCHTFLIPGVNARVRLRTKRMPHVVITCLECGYIMRYPYLREVKQKRKKAT
11100000000000000000000000000000000000000000000000000000000000
000000000000000000000000000000000000000000000000001111111111
>DM_train2441
MQAYSNPGYSSFPSPTGLEPSCKSCGAHFANTARKQTCLDCKKNFCMTCSSQVGNGPRLCLL
CQRFRATAFQREELMKMKVKDLRDYLSLHDISTEMCREKEELVLLVLGQQPVISQEDR
11111111111111111110000000000000000000000000000000001111000000
000000000000000000000000000000000000000000000000000000111111
>DM_train2442
MMRKLKTLPPTLRDKNRYIAFEIISDGDFTKDEVKELIWKSSLEVLGETGTAIVKPWLIKFD
PNTKTGIVRSDREYVEYLRFALMLVSEFNGKRLIIRTLGVSGTIKRLKRKFLAKYGWK
11000000000000000000000000000000000000000000000000000000000000
000000000000000000000000000000000000000000000000000000000000
>DM_train2443
SNAMDTQTLNSDLRVFMHHIYEFEKGVRSMVLATLANDDIPYAEERLRSRQIPYFAQPTPNT
ERTNLFFGCKECMEAIRLFVSGRSLNSLTPEEDFIIGAMLGYDICRQCERYCRRKSNS
11111100000000000000000000000000000000000000000000000000000000
000000000000000000000000000000000000000000000000000000000111
>DM_train2444
GHMVRKSIDRSTAAEITRGIGNVFADLGMPDAEERQTKLRLAYALNAVIDRARLSQAAAAAR
LGINQPKVSALRNYKLEGFSVERLMTLLNALDQDVEIVIRKKPRSRAAARISVVAAGS
11111111111111111111111111111000000000000000000000000000000000
000000000000000000000000000000000000000000000000000000000011
>DM_train2445
WVAFDKQVLSFDAYLEEEVLDKSQTNYRIRYYKIYFYPEDDTIQVNEPEVKNSGLLQGTSIR
RHRITLPPPDEDQFYTVYHFNVGTEVVFYGRTFKIYDCDAFTRNFLRKIGVKVNPPVQ
00000000000000000000000000000000000000000000000011111100000000
000000000000000000000000000000000000000000000000000000000001
>DM_train2448
GGPGNEKLKEKENNDSSDKATVIPNFNTTMQGSLLGDDSRDYYSFEVKEEGEVNIELDKKDE
FGVTWTLHPESNINDRITYGQVDGNKVSNKVKLRPGKYYLLVYKYSGSGNYELRVNK
11111111111111111110000000000000000000000000000000000000000000
00000000000111100000000000000000000000000000000000000000000
>DM_train2449
MEEKVGNLKPNMESVNVTVRVLEASEARQIQTKNGVRTISEAIVGDETGRVKLTLWGKHAGS
IKEGQVVKIENAWTTAFKGQVQLNAGSKTKIAEASEDGFPESSQIPENTPTAPQQMR

000000000000000000000000000000000000000000000000000000000000000
00000000000000000000000000000000000000000000000000000011111
>DM_train2450
EDPFGPGCEEAGCPEGSACNIITDRCTCSGVRCRVHCPHGFQRSRYGCEFCKCRLEPMKATC
DISECPEGMMCSRLTNKCDCKIDINCRKTCPNGLKRDKLGCEYCECRPKRKLIPRLS
111111000000000000000000000000000000000000000000000000000000000
000000000000000000000000000000000000000000000000000111111111
>DM_train2452
SKQPDKQRKSQRRAPLHERHKQVRATLSADLREEYGQRNVRVNAGDTVEVLRGDFAGEEGEV
INVDLDKAVIHVEDVTLEKTDGEEVPRPLDTSNVRVTDLDLEDEKREARLESEDDSA
000000000000000000000000000000000000111000000000000000000000000
00000000000000000000000000000000000000000000000000000111111
>DM_train2453
APLAVNDGDMRLADGGATNQGRVEIFYRGQWGTVCDNLWDLTDASVVCRALGFENATQALGR
AAFGQGSGPIMLDEVQCTGTEASLADCKSLGWLKSNCRHERDAGVVCTNETRSTHTL
111000000000000000000000000000000000000000000000000000000000000
00000000000000000000000000000000000000000000000000000111100
>DM_train2454
RDVVSFEQPEFSVSRGDQVARIPVIRRVLDGGKSQVSYRTQDGTAQGNRDYIPVEGELLFQP
GEAWKELQVKLLELQEVDSLLRGRQVRRFHVQLSNPKFGAHLGQPHSTTIIIRDPDE
000000000000000000000000000000000000000000000000000000000000000
00000000000000011111111100000000000000000000000000000000011
>DM_train2455
PMNIKGLFEVAVKVKNLEKSSQFYTEILGFEAGLLDSARRWNFLWVSGRAGMVVLQEEKENW
QQQHFSFRVEKSEIEPLKKALESKGVSVHGPVNQEWMQAVSLYFADPNGHALEFTAL
111000000000000000000000000000000000000000000000000000000000000
00000000000000000000000000000000000000000000000000000000000
>DM_train2456
MTDYDKLSNLATTFEFPDLTVEIKGPDVVGVNKLAEYEVHVKNLGGIGVPSTKVRVYINGTL
YKNWTVSLGPKEEKVLTFNWTPTQEGMYRINATVDEENTVVELNENNNVATFDVSVV
111111111110000000000000000000000000000000000000000000000000000
00000000000000000000000000000000000000000000000000000000000
>DM_train2457
MADRIELRGLTVHGRHGVYDHERVAGQRFVIDVTVWIDLAEAANSDDLADTYDYVRLASRAA
EIVAGPPRKLIETVGAEIADHVMDDQRVHAVEVAVHKPQAPIPQTFDDVAVVIRRSR
100000000000000000000000000000000000000000000000000000000000000
00000000000000000000000000000000000000000000000000000000000
>DM_train2458
LRIPNIATYTGTIQGKGEVCIIGNKEGKTRGGELYAVLHSTNVNADMTLILLRNVGGNGWGE
IKRNDIDKPLKYEDYYTSGLSWIWKIKNNSSETSNYSLDATVHDDKEDSDVLTKCPV
110000000000000000000000000000000000000000000000000000000000000
00000000000000000000000000000000000000000000000000000000000
>DM_train2459
MRRVEKVIIVEGRSDKQKVAAVLNEPVVIVCTNGTISDARLEELADELEGYDVYLLADADEA

GEKLRRQFRRMFPEAEHLYIDRAYREVAAAPIWHLAQVLLRARFDVRIESLMRGRGE
00000000000000000000000000000000000000000000000000000000000000
000000000000000000000000000000000000000000000000000011111
>DM_train2460
QGHMAVRRVVANIATPEPARAQAFYGDILGMPVAMDHGWIVTHASPLEAHAQVSFAREGGSG
TDVPDLSIEVDNFDEVHARILKAGLPIEYGPVTEAWGVQRLFLRDPFGKLINILSHA
11110000000000000000000000000000000000000000000000000000000000
000000000000000000000000000000000000000000000000000000011
>DM_train2461
SMSDVTIVKEGWVQKRGEYIKNWRPRYFLLKTDGSFIGYKEKPQDVDLPYPLNNFSVAKCQL
MKTERPKPNTFIIRCLQWTTVIERTFHVDTPEEREEWTEAIQAVADRLQRQEEERMN
11110000000000000000000000000000000000000000000000000000000000
000000000000000000000000000000000000000000000000000001111
>DM_train2462
MDVFLMIRRHKTTIFTDAKESSTVFELKRIVEGILKRPPDEQRLYKDDQLLDDGKTLGECGF
TSQTARPQAPATVGLAFRADDTFEALCIEPFSSPPELPDVMKPQDSGSSANEQAVQ
00000000000000000000000000000000000000000000000000000000000000
00000000000000000000000000000000000011111111111111111111
>DM_train2463
MSLLNKPKSEMTPEELQKREEEEFNTGPLSVLTQSVKNNTQVLINCRNNKKLLGRVKAFDRH
CNMVLENVKEMWTEVPKSGKGKKKSKPVNKDRYISKMFLRGDSVIVVLRNPLIAGK
11111111111111111111111110000000000000000000000000000000000000
00000000000111111111111111111100000000000000000000000000
>DM_train2464
MRHLKSGRKLNRHSSHRLALYRNQAKSLLTHGRITTTVPKAKELRGFVDHLIHLAKRGDLHA
RRLVLRDLQDVKLVRKLFDEIAPRYRDRQGGYTRVLKLAERRRGDGAPLALVELVE
11111111111110000000000000000000000000000000000000000000000000
00000000000000000000000000000000000000000000000000000000
>DM_train2465
MTKEPRNLSEWIKELKKASREAVILVEGKNDKKALSKFSIKNVIDLSGKRYADVVDMLEGKW
EKVILLFDLDTHGERINQKMKELLSSQGFLVDENFRNFLKKWNIIHIEEINGGKNS
11110000000000000000000000000000000000000000000000000000000000
00000000000000000000000000000000000000000000000000111111
>DM_train2466
MPTRVNDGVDADEVTFVNRFTVHGAPAEFESVFARTAAFFARQPGFVRHTLLRERDKDNSYV
NIAVWTDHDAFRRALAQPGFLPHATALRALSTSEHGLFTARQTLPEGGDTTGSGHR
11111111110000000000000000000000000000000000000000000000000000
00000000000000000000000000000000000000000000011111111111
>DM_train2467
EGSCMSITINPSRPSVNNIGQVDIDSVILGRPGAIGSWELNNFITIGLNRVNADTVRVNIRN
TGRTNRLIITQWDNTVTRGDVYELFGDYALIQGRGSFCLNIRSDTGRENWRMQLEN
00111111111111111000000000000000000000000000000000000000000000
00000000000000000000000000000000000000000000000000000000
>DM_train2468

MEAVVVEREAKGMKEIAIQEKDLTLQWRGNTGKLVKVRLKNTRAMEMWYNKQITEENIQEIT
TLNIIKNGKSLALEVYPEKSIYVKPNLGRINVPVFFIKTPINRGVFEEIFGETLKA
11111111111000000000000000000000000000000000000000000000000000
00000000000000000000000001110000000000000000000000011111
>DM_train2469
MRGSGSSTVDELTAAFTGGAATGEGGLTLTAPEIAENGNTVPIEVKAPGAVAIMLLAAGNPE
PAVATFNFGPAAADQRAATRIRLAQTQDVIALAKMADGSVVKAQTTVKVTIGGCGG
11111100000000000000000000000000000000000000000000000000000000
00000000000000000000000000000000000000000000000000000011
>DM_train2471
MAFQLEMVTRETVVIRLFGELDHHAVEQIRAKISTAIFQGAVTTIIWNFERLSFMDSSGVGL
VLGRMRELEAVAGRTILLNPSPTMRKVFQFSGLGPWMMDATEEEAIDRVRGIVNG
00000000000000000000000000000000000000000000000000000000000000
0000000000000000000000000000001110000000000000000000111
>DM_train2472
LMGDDKVKKEVGRASWKYFHTLLARFPDEPTPEEREKLHTFIGLYAELYPCGECSYHFVKLI
EKYPVQTSSRTAAAMWGCHIHNKVNEYLKKDIYDCATILEDYDCGCSDSDGKRVS
11111000000000000000000000000000000000000000000000000000000000
0000000000000000000000000000000000000000000000111111111
>DM_train2473
MSLSPPGVRLFYDPRGHHAGAINELCWGLEEQGVPCQTITYDGGGDAAALGALAARSSPLRV
GIGLSASGEIALTHAQLPADAPLATGHVTDSDDQLRTLGANAGQLVKVLPLSERN
11110000000000000000000000000000000000000000000000000000000000
0000000000000000000000000000000000000000000000000000000
>DM_train2474
MSAMNARQGILSLALKDKPALYSAYMPFVKGGGIFVPTPKRYMLGDEVFLLLTLPDSSERLP
VAGKVIWTTPAGAQGNRAAGIGVQFPDGPEGEAVRNKIETLLAGLTTSDKPTHTM
11111111000000000000000000000000000000000000000000000000000000
0000000000011111100000000000000000000000000111111111111
>DM_train2475
MNSDPTRERILHGLITLYILKELVKRPMHGYELQKSMFETTGQALPQGSIYILLKTMKERGF
VISESSVNEKGQQLTVYHITDAGKKFLCDHSQALQLARKIIDDLLSTVDIIENRN
11111110000000000000000000000000000000000000001000000000000000
0000000000000000000000000000000000000000000000000111111
>DM_train2476
HCDLPCGVYDPAQARIEAESVKAVQEKMAGNDDPHFQTRATVIKEQRAELAKHHVSVLWSDY
FKPPHFEKYPELHQLVNDTLKAMSAAKGSKDPATGQKALDYIAQIDKIFWETKKA
00000000000000000000000000000000000000000000000000000000000000
0000000000000000000000000000000000000000000000000000011
>DM_train2477
GSSEEESFFVQVHDVSPEQPRTVIKAPRVSTAQDVIQQTLCKAKYSLSILSNPNPSDYVLLE
EVVKDTTNKKTTTPKSSQRVLLDQECVFQAQSKWKGAGKFILKLKEQVQASREDK
11110000000000000000000000000000000000000000000000000000000000
0011111111111111000000000000111111111100000000011111111

>DM_train2478
MDRTHERVLQAMAENLGEGLPRAIPLLAEKAPGLLLEHGRSWTYAMPEKGALDEKTRTLILL
GIALATGSEACVKAMAHRAKRLGLSKEALLETLKIARQAQANAVLGHAAPLLEVL
111110000000000000000000000000000000000000000000000000000000000
0000000000000000000000000000000000000000000000000000000
>DM_train2479
SNAMSANDKLTILWTTDNKDTVFNMLAMYALNSKNRGWWKHINIILWGASVKLVANDTQVQT
EILEMLQSGITIEACQDCCENFGVASIITNLGITVRYMGIPLTEYLKNGEKILSI
100000000000000000000000000000000000000000000000000000000000000
0000000000000000000000000000000000000000000000000000000
>DM_train2480
QGHMVSAEGRNRKAVLCQRCGSRVLQPGTALFSRRQLFLPSMRKKPALSDGSNPDGDLLQEH
WLVEDMFIFENVGFTKDVGNIKFLVCADCEIGPIGWHCLDDKNSFYVALERVSHE
000000000000000000000000000000000000000011111111100000000
0000000000000000000000000000000000000000000000000000000
>DM_train2481
MAVNFYDVAYDLENALRGSEEFTRLKNLYDEVNADESAKRMFENFRDVQLRLQQKQMAGEEI
TQEEVTQAQKTVALVQQHEKISQLMEAEQRMSMLIGELNKIIMKPLEELYGSVEG
11000000000000000000000000000000000000000000000000111111111111
111111000000000000000000000000000000000000000000011111
>DM_train2482
MPKNKGKGGKNRRRGKNDSEGDKRELVFKEEGQEYGQVQRMLGNGRLDAYCFDGQKRLCHIR
GKMRKKVWVNPGDIVLVSLRDFQDSKGDIILKYTPDEARALKSKGEIPETTKINE
111111111111111111111000000000000000000000000000000000000000
0000000000000000000000000000000000000000000000000000011
>DM_train2483
MQNIPPQVQAMLGQLESYQQQLQLVVQQKQKVQLELTEAKKALDEIESLPDDAVVYKTVGTL
IVKTTKDKAVAELKEKIETLEVRLNALERQEKKLNEKLKELTAQIQSALRPPTAG
000000000000000000000000000000000000000000000000000000000000000
0000000000000000000000000000000000000000000000000000111
>DM_train2484
SLTFYPAWLTVSEGANATFTCSLSNWSEDLMLNWNRLSPSNQTEKQAAFSNGLSQPVQDARF
QIIQLPNRHDFHMNILDTRRNDSGIYLCGAISLHPKLKIEESPGAELVVTERILE
000000000000000000000000000000000000000000000000000000000000000
0000000000000000000000000000000000000000000000000011111
>DM_train2485
ARINGPDECGRVIKDTSGSISNTDRQKNLCTWTILMKPDQKVRMAIPYLNLACGKEYVEVFD
GLLSGPSYGKLCAGAAIVFLSTANTMTIKYNRISGNSSSPFLIYFYGSSPGSEY
000000000000000000000000000000000000000000000000000000000000000
000000000000000000000000000000000000000000000000001111
>DM_train2487
MATQRAHRAETPAHPNRLWIWEKHVYLDEFRRSWLPVVIKSNEKFQVILRQEDVTLGEAMSP
SQLVPYELPLMWQLYPKDRYRSADSMYWQILYHIKFRDVEDMLLELIDSESNDE
111111100000000000000000000000000000000000000000000000000000000

0000000000000000000000000000000000000000000000011111111
>DM_train2488
MSEAKELIKKMCDLQNSNEEIQKEMAGWSGVVQYKLDGEEFYVEYKSDGTCEFKEGVHSSPT
FTVVAPPDFWLAVLKGQEDPVSGFMMGKYRIEGNIMEAQRLAGVIKKFQGKFEL
00000000000000000000000000000000000000000000000000000000000000
000000000000000000000000000000000000000000000000111111
>DM_train2489
NNTPVTNKLKAYGDANFNFTNNSIADAEKQVQEAYKGLLNLNEKNASDKLLVEDNTAATVGN
LRKLGWVLSSKNGTRNEKSQQVKHADEVLFEGKGGVQVTSTSENGKHTITFALA
11000000000000000000000000000000000000000000001110000000000000
000000000000000000000000000000000000000000000000000001
>DM_train2490
MAEIPKEMLRAQTNVILLNVLKQGDNYVYGIIKQVKEASNGEMELNEATLYTIFKRLEKDGI
ISSYWGDESQGGRRKYYRLTEIGHENMRLAFESWSRVDKIIENLEANKKSEAIK
11110000000000000000000000000000000000000000000000000000000000
000000011110000000000000000000000000000000000001111111
>DM_train2491
MNDKKIELLTTYLSLYIDHHTVLADMQNATGKYVVLDVRNAPAQVKKDQIKGAIAMPAKDLA
TRIGELDPAKTYVVYDWTGGTTLGKTALLVLLSAGFEAYELAGALEGWKGMQLP
10000000000000000000000000000000000000001111111100000000000000
000000000000000000000000000000000000000000111111111111
>DM_train2492
MGSDKIHHHHHHMFYEIRTYRLKNGAIPAYLKVVEDEGIEIQKSHLGELVGYFFSEIGPINE
IVHIWAFSSLDDRAERRARLMADPRWLSFLPKIRDLIEVAENKIMKPARFSPLM
11111100000000000000000000000000000000000000000000000000000000
000000000000000000000000000000000000000000000000000000
>DM_train2493
GHMLYINSFLDRMGEIIRGEKSVEEADKLLDQKNIFEMFRSDCEEILNLYKSGKAEKEEVQR
NFYLLKTYVVSQLSIHFERLKEFAESKGFKIEKKLDPEVINEIALYIDRVEKEV
00000000000000000000000000000000000000000000000000000000000000
000000000000000000000000000011100000000000000000000000
>DM_train2494
GSPGIRMSLKSDEVFAKIAKRLESIDPANRQVEHVYKFRITQGGKVVKNWVMDLKNVKLVES
DDAAEATLTMEDDIMFAIGTGALPAKEAMAQDKMEVDGQVELIFLLEPFIASLK
11100000000000000000000000000000000000000000000000000000000000
000000000000000000000000000000000000000000000000000000
>DM_train2495
GMQQSEHFSFGEQTEIEDIGGGLKRQMLGFNHELMAVKIWFDKGAEGYVHAHRHSQVSYVVE
GEFHVNVDGVIKVLTAGDSFFVPPHVDHGAVCPTGGILIDTFSPAREDFVEGLS
10000000000000000000000000000000000000000000000000000000000000
000000000000000000000000000000000000000000000000001111
>DM_train2496
MVERLLEIIERSLRKCPWLEKQSIETLLEALASEIEEVAEAVKKNDLANLEEEIGDMIYDAL
LVAAVAQRDYGIDLESAIQKVVEKISHRKPWLFWEEKISLEEAEKIWKERKKKI

00000000000000000000000000000000000000000000000000000000000000
000000000000000000000000000000000000000000000000000011
>DM_train2497
MERQPTTDRMIQEYVPGKQVTLAHLIANPGKDLFKKLGLQDAVSAIGILTITPSEASIIACD
IATKSGAVEIGFLDRFTGAVVLTGDVSAVEYALKQVTRTLGEMMQFTTCSITRT
11111100000000000000000000000000000000000000000000000000000000
000000000000000000000000000000000000000000000000000000
>DM_train2498
MDAENRVIINVGGIRHETYKATLKKIPATRLSRLTEGMLNYDPVLNEYFFDRHPGVFAQIIN
YYRSGKLHYPTDVCGPLFEEELEFWGLDSNQVEPCCWMTYTAHRDTQETLAVL
11100000000000000000000000000000000000000000000000000000000000
00000000000000000000000000000000000000000000011111111
>DM_train2499
QSIYIELKNTGSLNQVFSSTQNSSIVIKFGAVWCKPCNKIKEYFKNQLNYYYVTLVDIDVDI
HPKLNDQHNIKALPTFEFYFNLNNEWVLVHTVEGANQNDIEKAFQKYCLEKAK
00000000000000000001000000000000000000000000000000000000000000
00000000000000000000000000000000000000000000000000011
>DM_train2500
RTLPGEGTQVHPRAPLLQILKVAGAQEEVFTVKEVMHYLGQYIMMKQLYDKQRQHIVHCHDD
PLGELLEVGSFSVKNPSPLYEMLKRNLVILNNSDAAKNLSVGKDSNESPSEDP
11111110000000000000000000000000000000000000000000000000000000
00000000000000000000000000000011111111111111111111111
>DM_train2501
NVCQPTRFISRHNIEGIFTFVDHRCVATVGYQPQELLGKNIVEFCHPEDQQLLRDSFQQVVK
LKGQVLSVMFRFRSKNQEWLWMRTSSFTFQNPYSDEIEYIICTNTNVKNSSQE
10000000000000000000000000000000000000000000000000000000000000
00000000000000000000000000000000000000000000000000111
>DM_train2502
MVTKTITEQRAEVRIFAGNDPAHTATGSSGISSPTPALTPLMLDEATGKLVVWDGQKAGSAV
GILVLPLEGTETALTYYKSGTFATEAIHWPESVDEHKKANAFAGSALSHAALP
11111111111100000000000000000000000000000000000000000000000000
00000000000000000000000000000000000000000000000000000
>DM_train2503
SNAMSYINNIEHAKVLDLTQEVMIEQDQMLSRTLVQRQDLGITVFSLDKGQEIGRHSSPGDA
MVTILSGLAEITIDQETYRVAEGQTIVMPAGIPHALYAVEAFQMLLVVVKPEA
10000000000000000000000000000000000000000000000000000000000000
00000000000000000000000000000000000000000000000000000
>DM_train2505
PKTQRGIYHNLKESEYVASNTDVTFFFSSELYLNKFLDGYQEYRKKFNKKIERVAVTPWNMD
MLADITFYSEVEKRGFHAWLKGDNATWREVHVYALRIMTKPNTLDWSRIQKPR
00000000000000000000000000000000000000000000000000000000000000
00000000000000000000000000000000000000000000000001111
>DM_train2506
MTLIYKILSRAEWDAAKAQGRFEGSAVDLADGFIHLSAGEQAQETAAKWFRGQANLVLLAVE

AEPLGEDLKWEASRGGARFPHLYRPLLVSEVTREADLDLDADGVPQLGDHLAL
100000000000000000000000000000000000000000000000000000000000
00000000000000000000000000000000000000000000000000000
>DM_train2507
MKYKLNVLLAEIALIGTGNHYHEEANCIAEWLHLKGEEEAVQLIRLSSLMNRGDYASALQQG
NKLAYPDLEPWLALCEYRLGLGSALESRLNRLARSQDPRIQTFVNGMREQLKT
110000000000000000000000000000000000000000000000000000000000
00000000000000000000000000000000000000000000000000001
>DM_train2508
ETGSKGQTAGELYQRWERYRRECQETLAAAEPPSGLACNGSFDMYVCWDYAAPNATARASCP
WYLPWHHHVAAGFVLRQCGSDGQWGLWRDHTQCENPEKNEAFLDQRLILERLQ
111110000000000000000000000000000000000000000000000000000000
00000000000000000000000000000000000001111111111111111
>DM_train2509
MATATTIRRKLRTRRKVRTTTAASGRLRLSVYRSSKHIYAQIIDDSRGQTLAAASSAALKSG
NKTDTAAAVGKALAAAAAEKGIKQVVFDRGSYKYHGRVKALADAAREGGLDF
111000000000000000000000000000000000000000000000000000000000
0000000000000000000000000000000000000000000000000000
>DM_train2510
GMKTQITTRDLVLEFIHALNTENFPAAKKRLNENFTFNGPMGHREGSERYMNDMEKMKFKYV
VHKMFEEGNDVCLIYDINMNGKTIAASGLYHLEKGEITSLHVYFDPRPLFEE
100000000000000000000000000000000000000000000000000000000000
0000000000000000000000000000000000000000000000000001
>DM_train2511
MNNKHATSAVHEIIREICRLVDSGHSMTRDQFHELSEQERFIAFLAEKYSSTIKLYYLADSS
PLFEKDTSSFIENAFGRHANTVVMEDFGLKSNALLLAINICLAILREINGEV
000000000000000000000000000000000000000000000000000000000000
0000000000000000000000000000000000000000000000000111
>DM_train2512
HKGTLYVVATPLGNLDDMTFRAVNTLRNAGAIACEDTRRTSILLKHFGIEGKRLVSYHSFNE
ERAVRQVIELLEEGSDVALVTDAGTPAISDPGYTMASAAHAAGLPVVPVPGA
000000000000000000000000000000000000000000000000000000001000
0000000000000000000000111111111000000000000000000011
>DM_train2513
SNAMGYMFIETKTFTVKEGTSNIVVERFTGEGIIEKFEGFIDLSVLVKKVRRGDEEVVVMIR
WESEEAWKNWETSEEHLAGHRAGRGKPKPDHIINVDHAVYYVKSSKAAYQQS
111100000000000000000000000000000000000000000000000000000000
0000000000000000000111111111000000000000000000001111
>DM_train2514
MGSSSSGLVPAGSHMRLTPHEQERLLLSYAAELARRRARGLRLNHPEAIAVIADHILEGAR
DGRTVAELMASGREVLGRDDVMEGVPEMLAEVQVEATFPDGTKLVTVHQPIA
111111111111111000000000000000000000000000000000000000000000
0000000000000000000000000000000000000000000000000000
>DM_train2515

GMIRLLYLLVKPAGMSDETFRAECLRHYEMSHDVPGLHKYEVRLVAEQPTDTHVPFFDIGHV
DAIGECWFKDDAAYATYMASDIRKAWFEHGKTFIGQLKPFRTAPVAGDEPAS
000000000000000000000000000000000000000000000001111111111100
0000000000000000000000000000000000000000000000111111
>DM_train2516
MKNLSFPLLFLFFLVPELLGSSMPLCPIDEAIDKKIKQDFNSLFPNAIKNIGLNCWTVSSRG
KLASCPEGTAVLSCSCGSACGSWDIREEKVCHCQCARIDWTAARCCKLQVAS
111111111111111111111110000000000000000000000000000000000000
0000000000000000000000000000000000000000000000000000
>DM_train2517
NWVNVISDLKKIEDLIQSMHIDATLYTESDVHPSCKVTAMKCFLLELQVISLESGDASIHDT
VENLIILANNSLSSNGNVTESGCKECEELEEKNIKEFLQSFVHIVQMFINTS
000000000000000000000000000000000000000000000000000000000000
0000000000000111110000000000000000000000000000000000
>DM_train2518
GAMDPEFTLSSEKKAKKVRFYRNGDRYFKGIVYAISPDRFRSFEALLADLTRTLSDNVNLPQ
GVRTIYTIDGLKKISSLDQLVEGESYVCGSIEPFKKLEYTKNVNPNWSVNV
111111111111000000000000000000000000000000000000000000000000
000000000000000000000000000000000000000000000000000
>DM_train2519
DAPSALYAPSALVLTVGKGVSATTAAPERAVTLTCAPGPSGTHPAAGSACADLAAVGGDLNA
LTRGEDVMCPMVYDPVLLTVDGVWQGKRVSYERVFSNECEMNAHGSSVFAF
111110000000000000000000000000000000000000000000000000000000
000000000000000000000000000000000000000000000000000
>DM_train2520
TCKMSQLERNIETIINTFHQYSVKLGHPDTLNQGEFKELVRKDLQNFLKKENKNEKVIEHIM
EDLDTNADKQLSFEEFIMLMARLTWASHEKMHEGDEGPGHHHKPGLGEGTP
111000000000000000000000000000000000000000000000000000000000
000000000000000000000001111111111111111111111111111
>DM_train2521
MVDLEQEFALGGRCLAFHGPLMYEAKILKIWDPSSKMYTSIPNDKPGGSSQATKEIKPQKLG
EDESIPEEIINGKCFFIHYQGWKSSWDEWVGYDRIRAYNEENIAMKKRLAN
111111000000000000000000000000000000000011111111111000000000
000000000000000000000000000000000000000000000000000
>DM_train2522
ATAFVPALVASGLPNEKFCFEGFLPQKKGRMTKLKSLVDEHRTMVFYESPHRLLKTLTQFAE
YFGPERQVSVSREISKIHEETVRGTLSELIEHFTATDPRGEIVIVLAGIDD
000000000000000000000000000000000000000000000000000000000000
000000000000000000000000000000000000000000000000011
>DM_train2523
NSPQEEVELKKLKHLEKSVEKIADQLEELNKELTGIQQGFLPKDLQAEALCKLDRRVKATIE
QFMKILEEIDTLILPENFKDSRLKRKGLVKKVQAFLAECDTVEQNICQETE
000000000000000000000000000000000000000000000000000000000000
000000000000000000000000000000000000000000000000111

>DM_train2524
NGLITKIWGTAGWTFNHAVTFGYPLNPTSDDKRRYKNYFISLGDVLPCRLCRESYKKFITTG
KTALTNEVLRNRHTLTKWFYDVHNAVNNKLEVDYGLSYEDVVNKYESFRAK
00000000000000000000000000000000000000000000000000000000000000
000000000000000000000000000000000000000000000000001
>DM_train2525
MSLILVYSTFPNEEKALEIGRKLLEKRLIACFNAFEIRSGYWWKGEIVQDKEWAAIFKTTEE
KEKELYEELRKLHPYETPAIFTLKVENVLTEYMNWLRESVLEGGSHHHHHH
10000000000000000000000000000000000000000000000000000000000000
000000000000000000000000000000000000000011111111111
>DM_train2526
GSHMGQNDLVKTLRMNYLFDFYQSLLTNKQRNYLELFYLEDYSLSEIADTFNVSRQAVYDNI
RRTGDLVEDYEKKLELYQKFEQRREIYDEMKQHLSNPEQIQRYIQQLEDLE
11111110000000000000000000000000000000000000000000000000000000
000000000000000000000000000000000000000000000000000
>DM_train2527
LETPGQYLVYNGDLVEYEADHMAQLQRVHGFLMNDCLLVATWLPQRRGMYRYNALYPLDRLA
VVNVKDNPPMKDMFKLLMFPESRIFQAENAKIKREWLEVLEETKRALSDKR
11110000000000000000000000000000000000000000000000000000000000
000000000000000000000000000000000000000000000000000
>DM_train2528
MNKPLTPSTYIRNLNVGILRKLSDFIDPQEGWKKLAVAIKKPSGDDRYNQFHIRRFEALLQT
GLSPTCELLFDWGTTNCTVGDLVDLLVQIELFAPATLLLPDAVPQTVKSLP
11000000000000000000000000000000000000000000000000000000000000
000000000000000000000000000000000000000000000111111
>DM_train2529
MQYATLELNNAFKVLFSLRQVQAAEMVIAPGDREGGPDNRHRGADQWLFVVDGAGEAIVDGH
TQALQAGSLIAIERGQAHEIRNTGDTPLKTVNFYHPPAYDAQGEPLPAGEG
10000000000000000000000000000000000000000000000000000000000000
000000000000000000000000000000000000000000000000011
>DM_train2530
GMARRDISGAVDFAYLEGFAAGDFAVVDEVLALFREQAALWAPMLDPTHPGWKDAVHTVKGA
ARGVGAFNLGEVCERCEAGQESLEGVRTALDAALLDIAAYAHEQALRSLKG
11111111000000000000000000000000000000000000000000000000000000
000000000000000000000000000000000000000000000000001
>DM_train2531
ATVSVFPGARLLTIGDANGEIQRHAEQQALRLEVRAGPDAAGIALYSHEDVCVFKCSVSRET
ECSRVGRQSFIITLGCNSVLIQFATPHDFCSFYNILKTCRGHTLERSVFSE
00000000000000000000000000000000000000000000000000000000000000
000000000000000000000000000000000000000011111111111
>DM_train2532
EDKYTDKYDNINLDEILANKRLLVAYVNCVMERGKCSPEGKELKEHLQDAIENGCKKCTENQ
EKGAYRVIEHLIKNEIEIWRELTAKYDPTGNWRKKYEDRAKAAGIVIPEE
11111111100000000000000000000000000000000000000000000000000000

00000000000000000000000000000000000000000000000111
>DM_train2533
MDGEEKTYGGCEGPDAMYVKLISSDGHEFIVKREHALTSGTIKAMLSGPGQFAENETNEVNF
REIPSHVLSKVCMYFTYKVRYTNSSTEIPEFPIAPEIALELLMAANFLDC
11111111111111110000000000000000000000000000000000011111111000000
00000000000000000000000000000000000000000000000000
>DM_train2534
MHFSIPETESRSGDSGGSAYVAYNIHVNGVLHCRVRYSQLLGLHEQLRKEYGANVLPAFPPK
KLFSLTPAEVEQRREQLEKYMQAVRQDPLLGSSETFNSFLRRAQQETQQV
10000000000001111110000000000000000000000000000000000000000000000
00000000000000000000000000000000000000000000000011
>DM_train2535
WSATNEEDDLSVEAEIAHQIAESFSKKYKFPSRSSGIFLYNFEQLKMNLDDIVKEAKNVPGV
TRLAHDGSKIPLRCVLGWVALANSKKFQLLVEADKLSKIMQDDLNRYTSC
00000000000000000000000000000000000000000000000000000000000000000
00000000000000000000000000000000000000000000000001
>DM_train2536
LKSESAEKTREVLWQQYYASNPPDHAVLEVLATPVREALLARFGQHQGSVVPAIDLPELRSV
LQQFDSFGKRWEAILLQVLEGIKPNESQVGLPYLSELINKELMILLPSNS
11110000000000000000000000000000000000000000000000000000000000000
00000000000000000000000011111111000000000000000000001111
>DM_train2537
MLQQNNIVYFDLDKYDIRSDFAQMLDAHANFLRSNPSYKVTVEGHADERGTPEYNISLGERR
ANAVKMYLQGKGVSADQISIVSYGKEKPAVLGHDEAAYSKNRRAVLVYLE
11110000000000000000000000000000000000000000000000000000000000000
00000000000000000000000000000000000000000000000001
>DM_train2538
MNKTKSEHIKQQALDLFTRLQFLLQKHDTIEPYQYVLDILETGISKTKHNQQTPERQARVVY
NKIASQALVDKLHFTAEENKVLAAINELAHSQKGWGEFNMLDTTNTWPSQ
11111100000000000000000000000000000000000000000000000000000000000
000000000000000000000000000000000011111111111111111
>DM_train2539
GMDKLVKYQELVKKLLTNYASDDVSDQDVEVQLILDTERNHYQWMNVGWQGLNRIYRCVIHF
DIKDGKIWLQQNLTDRNPAEELVMMGVPREDIVLGLQAPYKRQYTDYGVA
11100000000000000000000000000000000000000000000000000000000000000
00000000000000000000000000000000000000000000000000
>DM_train2540
MDTSLPSAILNNMVTATLILDDGLAIRYANPAAELLFSQSAKRIVEQSLSQLIQHASLDLAL
LTQPLQSGQSITDSDVTFVVDGRPLMLEVTVSPITWQRQLMLLVEMRKID
11000000000000000000000000000000000000000000000000000000000000000
00000000000000000000000000000000000000000000000001
>DM_train2541
MAVRIRLAKFGRKHHPIYRIVVMDAKSPREGKYIDILGTYDPKRKVLINVYPEKVKEWVLKG
VELSHRAKAILWNHGILKEVVPEGYEMKRVGDYYVFEKRESKKSKGGEAA

10000000000000000000000001111110000000000000000000000000000000
00000000000000000000000000000000000000001111111111
>DM_train2542
MGKATTEEQKLIEDVNASFRAAMATTANVPPADKYKTFEAAFTVSSKRNLADAVSKAPQLVP
KLDEVYNAAYNAADHAAPEDKYEAFVLHFSEALRIIAGTPEVHAVKPGA
11100000000000000000000000000000000000000000000000000000000000
0000000000000000000000000000000000000000000001111
>DM_train2543
ATDAASLVGELQALDAEYQNLANQEEARFNEERAQADAARQALAQNEQVYNELSQRAQRLQA
EANTRFYKSQYQEAASKYEDALKKLEAEMEQQKAVISDFEKIQALRAGN
11100000000000000000000000000000000000000000000000000000000000
0000000000000000000000000000000000000000000000011
>DM_train2544
MAHHHHHHVDDDDKMSLLDAHIPQLIASHTAFAAKAGLMRHTIGQAEQQAMSAQAFHQGESA
AAFQGAHARFVAAAAKVNTLLDIAQANLGEAAGTYVAADAAAASSYTGF
11111111111111111111111111111100000000000000000000000000000000
0000000000000000000000000000111111111111111111111
>DM_train2545
MDGASEIELVFRPHPTLMEKDDSAQTRYIKTSGNATVDHLSKYLAVRLALEELRSKGESNQM
NLDTASEKQYTIYIATASGQFTVLNGSFSLELVSEKYWKVNKPMELYYA
11110000000000000000111000000000000000000000000000000111111111
1110000000000000000000000000000000000000000000000
>DM_train2546
MKMQTYYYVLASRRFLLQEEPIEEVLKERTRHYHEQEKEIDFWLVPQPAFLEAPEFADIKAK
CPQPAAAIISTNSQFITWLKLRLEYVVTGEFSAPSETIPNPLASLATAS
00000000000000000000000000000000000000000000000000000000000000
0000000000000000000000000000000000000000000000111
>DM_train2547
MDFREVIEQRYHQLLSRYIAELTKTSLYQAQKFSRKTIEHQIPPEEIISIHRKVLKELYPSL
PEDVFHSLDFLIEVMIGYGMAYQEHQTLRGIQQEIKSEIEIAANVQQTL
11110000000000000000000000000000000000000000000000000000000000
0000000000000000000000011111111111111111111111111
>DM_train2548
SNAMATGGLAIIQSMKHKLPPSERKLADYILAHPHKAIESTVNEISALANSSDAAVIRLCKS
LGLKGFQDLKMRVAGDLAKPTFQGYRDIVPHEPLPSISEKTAGNAIQAI
11110000000000000000000000000000000000000000000000000000000000
0000000000000000000000001111111111111111111111111
>DM_train2549
GQDCPDSSEEVVGVSGKPVQLRPSNIQTKDVSVQWKKTEQGSHRKIEILNWYNDGPSWSNVS
FSDIYGFDYGDFALSIKSAKLQDSGHYLLEITNTGGKVCNKNFQLLILD
11000000000000000000000000000000000000000000000000000000000000
0000000000000000000000000000000000000000000000001
>DM_train2550
SFEKICSLDDIWVGEMETFETSDGTEVLIVNSEEHGVKAYQAMCPHQEILLSEGSYEGGVIT

CRAHLWTFNDGTGHGINPDDAALAEYPVEVKGDDIYVSTKGILPNKAHS
000000000000000000000000000000000000000000000000000000000000
0000000000000000000000000000000000000000000000011
>DM_train2551
GSSGSSGPGSLSGTEQAEMKMAVISEHLGLSWAELARELQFSVEDINRIRVENPNSLLEQSV
ALLNLWVIREGQNANMENLYTALQSIDRGEIVNMLEGSGRQSRNLKPDR
111111110000000000000000000000000000000000000000000000000000
0000000000000000000000000000000000001111111111111
>DM_train2552
MSIAVGMIETLGFPAVVEAADSMVKAARVTLVGYEKIGSGRVTVIVRGDVSEVQASVTAGIE
NIRRVNGGEVLSNHIIARPHENLEYVLPIRYTEAVEQFREIVNPSIIRR
110000000000000000000000000000000000000000000000000000000000
0000000000000000000000000000000111111111111111111
>DM_train2553
MSMEDPFFVVKGEVQKAVNTAQGLFQRWTELLQGPSAATREEIDWTTNELRNNLRSIEWDLE
DLDETISIVEANPRKFNLDATELSIRKAFITSTRQIVRDMKDQMSASS
000000000000000000000000000000000000000000000000000000000000
000000000000111100000000000000000000000000000000
>DM_train2554
MERGEIWLVSLDPTAGHEQQGTRPVLIVTPAAFNRVTRLPVVVPVTSGGNFARTAGFAVSLD
GVGIRTTGVVRCDQPRTIDMKARGGKRLERVPETIMNEVLGRLSTILT
000000000000000000000000000000000000000000000111110000000000
000000000000000000000000000000000000000000000000
>DM_train2555
AEFKHVFVCVQDRPPGHPQGSCAQRGSREVFQAFMEKIQTDPQLFMTTVITPTGCMNACMMG
PVVVVYPDGVWYGQVKPEDVDEIVEKHLKGGEPVERLVISKGKPPGMF
000000000000000000000000000000000000000000000000000000000000
000000000000000000000000000000000000000001111111
>DM_train2556
CILRFIACNGQTRAVQSRGDYQKTLAIALKKFSLEDASKFIVCVSQSSRIKLITEEEFKQIC
FNSSSPERDRLIIVPKEKPCPSFEDLRRSWEIELAQPAALSSQSSLSP
000000000000000000000000000000000000000000000000000000111111
111111100000000000000000000000000111111111111111
>DM_train2557
MSENANDIVTLRAQGGSDAPSDQPLNQVITNSTRQFNIELIRVQPRGEMMQVWIQPLPFSQL
VSWIAYLQERQGVSVDAIDIDRGKVNGVVEVKRLQLKRGGLEHHHHHH
111111111111111111111100000000000000000000000000000000000000
000000000000000000000001111000000000001111111111
>DM_train2558
GGGGGGMLNRVFLEGEIESSCWSVKKTGFLVTIKQMRFFGERLFTDYYVIYANGQLAYELEK
HTKKYKTISIEGILRTYLERKSEIWKTTIEIVKIFNPKNEIVIDYKEI
111111000000000000000000000000000000000000000000000000000000
000000000000000001111110000000000000000001111111
>DM_train2559

GMAEPKFTSFTTADFINDVDMELFIDAVEKTAPVWVKEMKSRGLLKFSMNRVWNKGEVFRVV
MTYEYKDRASFEANIAYLEDTFGKNPVFLQLVTTAKFTTSRCLVVMEV
11100000000000000000000000000000000000000000000000000000000000
000000000000000000000000000000000000000000000000
>DM_train2560
AGRPAVSASSRWLEGIRKWYYNAAGFNKYGLMRDDTIYENDDVKEAIRRLPENLYDDRMFRI
KRALDLNMRQQILPKEQWTKYEEDVPYLEPYLKEVIRERKEREEWDK
11111111100000000000000000000000000000000000000000000000000000
00000000000000000000000000000000000000000000000
>DM_train2561
SHGSHETDEEFDARWVTYFNKPDIDAWELRKGMNTLVGYDLVPEPKIIDAALRACRRLNDFA
SAVRILEVVKDKAGPHKEIYPYVIQELRPTLNELGISTPEELGLDKV
11110000000000000000000000000000000000000000000000000000000000
00000000000000000000000000000000000000000000000
>DM_train2562
GSHMTTKKASDSAFWLEVEGNSMTAPTGSKPSFPDGMLILVDPEQAVEPGDFCIARLGGDEF
TFKKLIRDSGQVFLQPLNPQYPMIPCNESCSVVGKVIASQWPEETFG
11111111000000000000000000000000000000000000000000000000000000
00000000000000000000000000000000000000000000000
>DM_train2563
DLHEILHEAVPLDANEREILELKEDAFAQRRREIETRLRAANGKLADAIAKNPAWSPEVEAA
TQEVERAAGDLQRATLVHVFEMRAGLKPEHRPAYDRVLIDALRRGSQ
00000000000000000000000000000000000000000000000000000000000000
00000000000000000000000000000000000000000000111
>DM_train2564
MKTLMVFDPAMAASTGVCGTDVDQALVDFSTDVQWLKQSGVQIERFNLAQQPMSFVQNEKVK
AFIEASGAEGLPLLLLDGETVMAGRYPKRAELARWFGIPLDKVGLAP
00000000001111111111111000000000000000000000000000000000000000
00000000000000000000000000000000000000000001111
>DM_train2565
MAYRALMVLRMDPADAEHVAAAFAEHDTTELPLEIGVRRRVLFRFHDLYMHLIEADDDIMER
LYQARSHPLFQEVNERVGQYLTPYAQDWEELKDSKAEVFYSWTAPDS
10000000000000000000000000000000000000000000000000000000000000
00000000000000000000000000000000000000000000011
>DM_train2566
PSSPSIDQVEPYSSTAQVQFDEPEATGGVPILKYKAEWRAVGEEVWHSKWYDAKEASMEGIV
TIVGLKPETTYAVRLAALNGKGLGEISAASEFKTQPVREPSAPKLEG
00000000000000000000000111100000000000000000000000000000000000
00000000000000000000000000000000000001111111111
>DM_train2567
MVNFKDKSMPTAIEKALDFIGGMNTSASVPHSMDESTAKGILKYLHDLGVPVSPEVVVARGE
QEGWNPEFTKKVAGWAEKVASGNRILIKNPEYFSTYMQEQLKELVLE
11111111110000000000000000000000000000000000000000000000000000
00000000000000000000000000000000000000000000000

>DM_train2568
GSDYIIKEKTVLLQKKDSEGFGFVLRGAKAQTPIEEFTPTPAFPALQYLESVDEGGVAWRAG
LRMGDFLIEVNGQNVVKVGHRQVVNMIRQGGNTLMVKVVMVTRHPDM
00000000000000000000000000001111100000000000000000000000000000
00000000000000000000000000000000000000000000000
>DM_train2569
KAINNIVASFSSVNDAITQTAEAIHTVTIALNKIQDVVNQQGSALNHLTSQLRHNFQSGGRG
SGRGGNLTYLNLSSELKQLEAKTASLFQTTVELQGLIDQINSTYVDL
00000000000000000000000000000000000000000000000000001111111111
11111110000000000000000000000000000000000001111
>DM_train2570
NNEIMPGKMKVKATKGEGDGGITSEGNALYNNEGGRAFMYAYVTTKPGMKYVKWEHDSGVVT
VELEPPCRFVIDTPTGPQIKYLYFVKNLNNLRRGAVLGYIGATVRLQ
11111110000000000000000000000000111100000000000000000000000000
00000000000000000000000000000000000000000000000
>DM_train2571
MTDLPAGDRLTGIPSHILNSSPSDRQINQLAQRLGPEWEPMVLSLGLSQTDIYRCKANHPHN
VQSQVVEAFIRWRQRFGKQATFQSLHNGLRAVEVDPSLLLHMLEELE
11111111111111100000000000000000000000000000000000000000000000
00000000000000000000000000000000000000000000111
>DM_train2572
MEEAKGPVKHVLLASFKDGVSPEKIEELIKGYANLVNLIEPMKAFHWGKDVSIENLHQGYTH
IFESTFESKEAVAEYIAHPAHVEFATIFLGSLDKVLVIDYKPTSVSL
11111100000000000000000000000000000000000000000000000000000000
00000000000000000000000000000000000000000000000
>DM_train2573
SMFNNELMADVHFVVGPPGATRTVPAHKYVLAVGSSVFYAMFYGDLAEVKSEIHIPDVEPAA
FLILLKYMYSDEIDLEADTVLATLYAAKKYIVPALAKACVNFLETSL
00000000000000000000000000000000000000000001111110000000000000
00000000000000000000000000000000000000000000000
>DM_train2574
MKWEMGLQEEYIELIKAGKKKIEGRLYDEKRRQIKPGDIIIFEGGKLKVKVKGIRVYSSFKE
MLEKEGIENVLPGVKSIEEGVKVYRQFYDEEREKKYGVVAIEIEPIE
00000000000000000000000000000000000000000000000000000000000000
00000000000000000000000000000000000000000000001
>DM_train2575
HMSNPLGELVKALEKLSFKPSDVRIYSLLLERGGMRVSEIARELDLSARFVRDRLKVLLKRG
FVRREIVEKGWVGYIYSAEKPEKVLKEFKSSILGEIERIEKMFTDGS
11000000000000000000000000000000000000000000000000000000000000
00000000000000000000000000000000000000000000111
>DM_train2576
MAEESFYGVTLTAESDSVTWDVDEDYARGQKLVIKQILLGAEAKENEFNVVEVNTPKDSVQI
PIAVLKAGETRAVNPDVEFYESKVTFKLIKGSGPVYIHGHNIKDDV
11100000000000000000011111110000000000000000000000000000000000

00000000000000000000000000000000000000000001111
>DM_train2577
MASMSNGTPDIIVNAQINSEDENVLDFIIEDEYYLKKRGVGAHIIKVASSPQLRLLYKNAYS
TVSCGNYGVLCNLVQNGEYDLNAIMFNCAEIKLNKGQMLFQTKIWR
11111100000000000000000000000000000000000000000000000000000000
0000000000000000000000000000000000000000000000
>DM_train2578
MGPMPKLADRKLCADQECSHPISMAVALQDYMAPDCRFLTIHRGQVVYVFSKLKGRGRLFWG
GSVQGDYYGDLAARLGYFPSSIVREDQTLKPGKVDVKTDKWDFYCQ
11100000000000000000000000000000000000000000000000000000000000
0000000000000000000000000000000000000000000000
>DM_train2579
MEITSDMEEDKDLMLKLLDKNGFVLKKVEIYRSNYLAILEKRTNGIRNFEINNNGNMRIFGY
KMMEHHIQKFTDIGMSCKIAKNGNVYLDIKRSAENIEAVITVASEL
11111000000000000000000000000000000000000000000000000000000000
0000000000000000000000000000000000000000000000
>DM_train2581
DDPIGLFVMRPQDGEVTVGGSITFSARVAGASLLKPPVVKWFKGKWVDLSSKVGQHLQLHDS
YDRASKVYLFELHITDAQPAFTGSYRCEVSTKDKFDCSNFNLTVHE
00000000000000000000000000000001000000000000000000000000000000
0000000000000000000000000000000000000000000000
>DM_train2582
MLRYGDTEICIDPSESVLHLLGKKYTMLIISVLGNGSTRQNFNDIRSSIPGISSTILSRRIK
DLIDSGLVERRSGQITTYALTEKGMNVRNSLMPLLQYISVLDRNGD
00000000000000000000000000000000000000000000000000000000000000
0000000000000000000000000000000000000000000011
>DM_train2584
MSSAQFDHVTVIKKSNVYFGGLCISHTVQFEDGTKKTLGVILPTEQPLTFETHVPERMEIIS
GECRVKIADSTESELFRAGQSFYVPGNSLFKIETDEVLDYVCHLEG
11000000000000000000000000000000000000000000000000000000000000
0000000000000000000000000000000000000000000001
>DM_train2585
MLTVIAEIRTRPGQHHRQAVLDQFAKIVPTVLKEEGCHGYAPMVDCAAGVSFQSMAPDSIVM
IEQWESIAHLEAHLQTPHMKAYSEAVKGDVLEMNIRILQPGISGRV
00000000000000000000000000000000000000000000000000000000000000
0000000000000000000000000000000000000000011111
>DM_train2586
MSTSADYKINQQQIVCVASFLSKEGKTEALIAALASLIPDTRREAGCIRYELNVSRDEPRRV
TFVEKFVDIAAFDEHCAKDAIQHYFHQVMPELVESFHVETYHQVIA
11111000000000000000000000000000000000000000000000000000000000
0000000000000000000000000000000000000000000000
>DM_train2587
MHRTTRIKITELNPHLMCVLCGGYFIDATTIIECLHSFCKTCIVRYLETSKYCPICDVQVHK
TRPLLNIRSDKTLQDIVYKLVPGLFKNEMKRRRDFYAAHPSADAAN

11111000000000000000000000000000000000000000000000000000000000
00000000000000000000000000000000000000000011111
>DM_train2588
MLAKNKGKIPGLKIDQKIRGQMPERGWTEDDIKNTVSNGATGTSFDKRSPKKTPPDYLGRND
PATVYGSPGKYVVVNDRTGEVTQISDKTDPGWVDDSRIQWGNKNDQ
11111111111000000000000000000000000000000000000000000000000000
0000000000000000000000000000000000000000001111
>DM_train2589
RYCKRTIPPGYKVDQVFGPRTKGKEGNFGDDKMNEEGIKDGRVTAMLNLVPSSHACLFGSRV
TPKLQPDGLHLKFEFTTVVPRDDPQFDNYVKICDQCVDGVGTRPKD
00000000000000000000000000000000000000000000000000000000000000
0000000000000000000000000000000000000000000001
>DM_train2590
ADDAKPRVKVPSSAKAGETVTVKALISHKMESGQRKDADGKLIPRSIINRFTCELNGVNVVD
VAIDPAVSTNPYFEFDAKVDAAGEFKFTWYDDDGSVYEDVKPIAVA
11000000000000000000000000000000001111111000000000000000000000
0000000000000000000000000000000000000000000000
>DM_train2591
AQQSPYSAAMAEQRHQEWLRFVDLLKNAYQNDLHPLLNLMLTPDEREALGTRVRIVEELLR
GEMSQRELKNELGAGIATITRGSNSLKAAPVELRQWLEEVLLKSD
11100000000000000000000000000000000000000000000000000000000000
000000000000000000000000000000000000000000000
>DM_train2592
YEVPKAKIDVFYPKGFEVSIPDEEGITLFAFHGKLNEEMEGLEAGTWARDIVKAKNGRWTFR
DRITALKPGDTLYYWTYVIYNGLGYREDDGSFVVNGYSGNNASPH
00000000000000000000000000000000000000000000000000000000000000
000000000000000000000000000000000000000111111
>DM_train2593
GLFDFLKRKEVKEEEKIEILSKKPAGKVVVEEVVNIMGKDVIIGTVESGMIGVGFKVKGPSG
IGGIVRIERNREKVEFAIAGDRIGISIEGKIGKVKKGDVLEIYQT
11111111111111110000000000000000000000000000000000000000000000
000000000000000000000000000000000000000000000
>DM_train2594
MNDMGSSEVNDENKEKEARYSVMTKSELEALAVSAIREHRRLLWADQAVYEEWLRASDDPSI
SGPVLQTLQDEYVARQKRSEAQQEELSDILDALGFVPDVPFDDDN
11111111111111110000000000000000000000000000000000000000000000
000000000000000000000000000000000000000011111
>DM_train2595
PLGSQFWVTVQRTEAAERCGLHGSYVLRVEAERLTLLTVGAQSQILEPLLSWPYTLLRRYGR
DKVMFSFEAGRRCPSGPGTFTFQTAQGNDIFQAVETAIHRQKAQG
00000000000000000000000000000000000000001111000000000000000000
000000000000000000000000000000000000000001111
>DM_train2596
ENLYFMLEMEIKITEVKENKLIGRKEIYFEIYHPGEPTPSRKDVKGKLVAMLDLNPETTVIQ

YIRSYFGSYKSKGYAKYYYDKDRMLYIEPEYILIRDGIIEKKEGE
11111111000000000000000000000000000000000000000000000000000000
000000000000000000000000000000000000000011111
>DM_train2599
MAPTQLDIKVKALKRLTKEEGYYQQELKDQEAHVAKLKEDKSVDPYDLKKQEEVLDDTKRLL
PTLYEKIREFKEDLEQFLKTYQGTEDVSDARSAITSAQELLDSK
11100000000000000000000000000000000000000000000000000000000000
00000000000000000000000000000000000000000000
>DM_train2600
MIPGEYHVKPGQIALNTGRATCRVVVENHGDRPIQVGSHYHFAEVNPALKFDRQQAAGYRLN
IPAGTAVRFEPGQKREVELVAFAGHRAVFGFRGEVMGPLEVNDE
00000000000000000000000000000000000000000000000000000000000000
00000000000000000000000000000000000000011111
>DM_train2601
MGKTNDWLDFDQLAEEKVRDALKPPSMYKVILVNDDYTPMEFVIDVLQKFFSYDVERATQLM
LAVAYQGKAICGVFTAEVAETKVAMVNKYARENEHPLLCTLEKA
11111111111111111111110000000000000000000000000000000000000000
00000000000000000000000000000000000000000000
>DM_train2602
MAQKKPKRNLSALKRHRQSLKRRLRNKAKKSAIKTLSKKAIQLAQEGKAEEALKIMRKAESL
IDKAAKGSTLHKNAAARRKSRLMRKVRQLLEAAGAPLIGGGLSA
11111110000000000000000000000000000000000000000000000000000000
00000000000000000000000000000000000000000000
>DM_train2603
VLLESEQFLTELTRLFQKCRTSGSVYITLKKYDGRTKPIPKKGTVEGFEPADNKCLLRATDG
KKKISTVVSSKEVNKFQMAYSNLLRANMDGLKKRDKKNKTKKTK
00000000000000000000000000000000011111111111111111110000000000
00000000000000000000000000000000111111111111
>DM_train2604
HFGGYIQKQAQTERKSGKRQTEREKKKKILAERRKVLAIDHLNEDQLREKAKELWQTIYNLE
AEKFDLQEKFKQQKYEINVLRNRINDNQKVSKTRGKAKVTGRWK
11111111111111111111000000000000000000000000000000000000000000
00000000000000000000000000011111111111111111
>DM_train2605
GSVSKWVRLNVGGTYFLTTRQTLCRDPKSFLYRLCQADPDLDSDKDETGAYLIDRDPTYFGP
VLNYLRHGKLVINKDLAEEGVLEEAEFYNITSLIKLVKDKIRER
11100000000000000000000000000000000111111110000000000000000000
00000000000000000000000000000000000000000000
>DM_train2606
VETKCPNLDIVTSSGEFHCSGCVEHMPEFSYMYWLAKDMKSDEDTKFIEHLGDGINEDETVR
TTDGGITTLRKVLHVTDTNKFAHYRFTCVLTTLDGVSKKNIWLK
00000000000000000000000000000000000000000000000000000000000000
00000000000000000000000000000000000000000001
>DM_train2607

GMTESDRDQEASMREVVIVKSTPQRGKFNAFAELVGKLVSETRDFPGCLGAYLMLAPERNEQ
VVMHIWETPDALEAYLTWRADRGDFLEINEYLEVEQDFKTYQLA
11111111111100000000000000000000000000000000000000000000000000
00000000000000000000000000000000000000000000
>DM_train2608
TKAIPWKINWQTMAFEYIGPQIEALLGWPQGSWKSVEDWATRMHPEDQEWVVNFCVKQSECG
VDHEADYRALHRDGHYVWIRDVVHVVRDDSGEVEALIGFMFDIS
10000000000000000000000000000000000000000000000000000000000000
00000000000000000000000000000000000000000000
>DM_train2609
PIEPQSDAHVLKSRLEWGEPAFTILDVRDRSTYNDGHIMGAMAMPIEDLVDRASSSLEKSRD
IYVYGAGDEQTSQAVNLLRSAGFEHVSELKGGLAAWKAIGGPTE
11111100000000011100000000000000000000000000000000000000000000
00000000000000000000000000000000000000000000
>DM_train2610
GMNKKTQLLEVIAALPEELVDQALNYVQMLQNPIQITPGVCGGQARIRNTRIPVWTLVAYRQ
QGAPDKELLANYPGLTAEDLSAAWHYYEQNPEQIDREIAQDDLV
10000000000000000000000000000000000000000000000000000000000000
00000000000000000000000000000000000000001111
>DM_train2611
SMALGPFPAMENQVLVIRIKIPNSGAVDWTVHSGPQLLFRDVLDVIGQVLPEATTTAFEYED
EDGDRITVRSDEEMKAMLSYYYSTVMEQQVNGQLIEPLQIFPRA
11111111111110000000000000000000010000000000000000000000000000
00000000000000000000000000000000000000000000
>DM_train2612
MITVDITVNDEGKVTDVIMDGHADHGEYGHDIVCAGASAVLFGSVNAIIGLTSERPDINYDD
NGGHFHIRSVDTNNDEAQLILQTMLVSLQTIEEEYNENIRLNYK
00000000000000000000011111111110000000000000000000000000000000
00000000000000000000000000000000000000000000
>DM_train2613
MTLEKHAFKMQLNPGMEAEYRKRHDEIWPELVDLLHQSGASDYSIHLDRETNTLFGVLTRPK
DHTMASLPDHPVMKKWWAHMADIMATNPDNSPVQSDLVTLFHMP
10000000000000000000000000000000000000000000000000000000000000
00000000000000000000000000000000000000000000
>DM_train2614
VGFAPELYCGLENCYDVLEVNREEFDKQKLAKAYRALARKHHPDRVKNKEEKLLAEERFRVI
ATAYETLKDDEAKTNYDYYLDHPDQRFYNYYQYYRLRAAPKVDL
00000000000000000000000000000000000000000000000000000000000000
00000000000000000000000000000000000001111111
>DM_train2615
MGEETINSTQNKTRTKTTRPKAVYLWTVSDVLKWYRRHCGEYTQYEQLFAQHDITGRALLRI
TDSSLQRMGVTDNRDREAIWREIVKQRLKTDIMEIRDMERLNIY
11111111111111111110000000000000000000000000000000000000000000
00000000000000000000000000000000000000000001

>DM_train2616
EAEASNMSSAHKCIDTNVPENAACYRYLDGTEEWRCLLGFKEVGGKCVPASITCEENNGGCA
PEAECTMDDKKEVECKCTKEGSEPLFEGVFCSSSSGPHHHHHH
111111100000000000000000000000000000000000000000000000000000
0000000000000000000000000000000011111111111
>DM_train2617
GMKNEVFFGEGMKVVAAAYPDLYDIIVKLNDTVFTGKTLDYKTQKLIAIGIVASRCDEVAIE
KQMKSAMKELGITKEEIADVLRVVLLTSGMPAFTKAMKILEKL
111111111000000000000000000000000000000000000000000000000000
0000000000000000000000000000000000000000000
>DM_train2618
MPVVKINAIEVPAGAGPELEKRFAHRAHAVENSPGFLGFQLLRPVKGEERYFVVTHWESDEA
FQAWANGPAIAAHAGHRANPVATGASLLEFEVVLDVGGTGKTA
100000000000000000000000000000000000000000000000000000000000
0000000000000000000000000000000000000111111
>DM_train2619
GPHMATADRDILARLHKAVTSHYHAITQEFENFDTMKTNTISREEFRAICNRRVQILTDEQF
DRLWNEMPVNAKGRLKYPDFLSRFSSETAATPMATGDSAVAQR
111100000000000000000000000000000000000000000000000000000000
0000000000000000000000000111111111111111111
>DM_train2620
MSPFKKTYVLKLYVAGNTPNSVRALKMLKNILEQEFQGVYALKVIDVLKNPQLAEEDKILAT
PTLAKILPPPVRKIIGDLSDREKVLIGLDLLYDEIREREAEDQ
111110000000000000000000000000000000000000000000000000000000
0000000000000000000000000000000000111111111
>DM_train2621
QTLDIQRGATLFNRACAACHDTGGNIIQPGATLFTKDLERNGVDTEEEIYRVTYFGKGRMPG
FGEKCTPRGQCTFGPRLQDEEIKLLAEFVKFQADQGWPTVSTD
110000000000000000000000000000000000000000000000000000000000
0000000000000000000000000000000000000001111
>DM_train2622
GHMTETKPRIAIRYCTQCNWLLRAGWMAQEILQTFASDIGEVSLIPSTGGLFEITVDGTIIW
ERKRDGGFPGPKELKQRIRDLIDPERDLGHVDRTKHEGLDTGS
111100000000000000000000000000000000000000000000000000000000
0000000000000000000000011111111111111111111
>DM_train2623
MDKPILKDSMKLFEALGTIKSRSMFGGFGLFADETMFALVVNNQLHIRADQQTSSDFETQGL
KPYVYKKRGFPVVTKYYAISSELWESSDRLIEVAKKSLENAKL
110000000000000000000000000000000000000000000000000000000000
0000000000000000000000000000000000000000001
>DM_train2624
GAMGKCPTKVVLLRNMVGAGEVDEDLEVETKEECEKYGKVGKCVIFEIPGAPDDEAVRIFLE
FERVESAIKAVVDLNGRYFGGRVVKACFYNLDKFRVLDLAEQV
100000000000000000000011100000000000000000000000000000000000

0000000000000000000000000000000000000000000
>DM_train2625
GMINMKVAISMDVDKISNSFEDCKYFLIVRIDDNEVKSTKVIFNDESGKKSIVKENVNAIIC
KNISEENYKKFSKKIEIYHAEGDDVDKNISLFIEGELSKISNP
10000000000000000000000000000000000000000000000000000000000000
0000000000000000000000000000000000000000000
>DM_train2626
YRALFEHAIDGIFIMDAEGHYLDVNPAICSAIGYTRDEFLALDWGVLSRGVDSGWAAASLAR
IVGGEPLREERTVWTRNGDQLTVELSAHLLPDGKILGIARDVS
00000000000000000000000000000000000000000000000000000000000000
0000000000000000000000000000000000000000011
>DM_train2627
GSMSSDAQWLTAEERDQLIPGLKAAGWSELSERDAIYKEFSFKNFNQAFGFMSRVALQAEKM
NHHPEWFNVYNKVQITLTSHDCGGLTKRDVKLAQFIEKAAASL
11111000000000000000000000000000000000000000000000000000000000
0000000000000000000000000000000000000000011
>DM_train2628
SQTIALLNIYRNPQNSSQSADGLRSAVSDVEMQEHYDEFFEEVFTEMEEKYGEVEEMNVCDN
LGDHLVGNVYVKFRREEDAEKAVIDLNNRWFNGQPIHAELSP
00000000000000111111000000000000000000000000000000000000000000
000000000000000000000000000000000000000000
>DM_train2629
SKGMALRSVGGMVIESPRNETEHWLLETVGRQAQQAGIGMPTVAIYDSADINAFATGAKRDD
SLVAVSTGLLHNMTRDEAEAVLAHEVSHIANGDMVTMTLMQG
11100000000000000000000000000000000000000000000000000000000000
000000000000000000000000000000000000000000
>DM_train2630
MNRDQEKIQIENEMNAMHGTIKEDILKDFEEFKGYLKKQVNRGKKLGLDDGKLVKSAAILGD
YLAKHEEPQNGEEMLLQELWSVADEDEKEHLAQLLVKLVDKQ
11111111111111111111000000000000000000000000000000000000000000
000000000000000000000000000000000000000000
>DM_train2631
MARPNFRYTHYDLKELRAGTTLEISLSSVNNVRLMTGANFQRFTELLDFKYLGGVAKKSPIR
IAVPETMHWHLIIDAEGHSGLAESSVKMLPAQPQATLTRKAS
11100000000000000000000000000000000000000000000000000000000000
000000000000000000000000000000000000000000
>DM_train2632
MSHTILLVQPTKRPEGRTYADYESVNECMEGVCKMYEEHLKRMNPNSPSITYDISQLFDFID
DLADLSCLVYRADTQTYQPYNKDWIKEKIYVLLRRQAQQAGK
10000000000000000000000000000000000000000000000000000000000000
000000000000000000000000000000000000000111
>DM_train2633
GHMTKLALFVRLEAKPGQEAALADFLASALPLANAESGTTAWFALKFGPSTFGVFDAFADEA
GRQAHLNGQIAAALMANAATLLSSPPNIEKVELLAAKLPAGS

11000000000000000000000000000000000000000000000000000000000000
000000000000000000000000000000000000000011
>DM_train2634
MSGRGKGGKGLGKGGAKRHRKVLRDNIQGITKPAIRRLARRGGVKRISGLIYEETRGVLKVF
LENVIRDAVTYTEHAKRKTVTAMDVVYALKRQERTLYGFGG
11111111111111111111111110000000000000000000000000000000000000
00000000000000000000000000000000000000000
>DM_train2635
GAKPGKKTRGRVKIKMEFIDNKLRRYTTFSKRKTGIMKKAYELSTLTGTQVLLLVASETGHV
YTFATRKLQPMITSETGKALIQTCLNSPDSPPRSDPTTDQR
11111000000000000000000000000000000000000000000000000000000000
00000000000000000000000000011111111111111
>DM_train2636
MTARNPLEAQAWALLEAVYDPELGLDVVNLGLIYDLVVEPPRAYVRMTLTTPGCPLHDSLGE
AVRQALSRLPGVEEVEVEVTFEPPWTLARLSEKARRLLGWG
11000000000000000000000000000000000000000000000000000000000000
00000000000000000000000000000000000000011
>DM_train2637
NGYTMQRDNQKTLAVYMFEEINRDVEYLSGRLSEKELKDKYRYYGRGYVRITDKDGQVITYE
DGSVQDKTVFLTNEGANKLGWKLEFLIDEKMFEEEILEKQN
11000000000000000000000000000000000000000000000000000000000000
00000000000000000000000000000000000001111
>DM_train2638
MSTLADQALHNNNVGPIIRAGDLVEPVIETAEIDNPGKEITVEDRRAYVRIAAEGELILTRK
TLEEQLGRPFNMQELEINLASFAGQIQADEDQIRFYFDKTM
10000000000000000000000000000000000000000000000000000000000000
00000000000000000000000000000000000000000
>DM_train2639
GKKQPSLHYTAAQLLEKGVLVEIEDLPASHFRNVIFDITPGDEAGKFEVNAKFLGVDMERFQ
LHYQDLLQLQYEGVAVMKLFNKAKVNVNLLIFLLNKKFLRK
11100000000000000000000000000000000000000000000000000000000000
00000000000000000000000000000000000000001
>DM_train2640
MKSITTDELKNKLLESKPVQIVDVRTDEETAMGYIPNAKLIPMDTIPDNLNSFNKNEIYYIV
CAGGVRSAKVVEYLEANGIDAVNVEGGMHAWGDEGLEIKSI
00000000000000000000000000000000000000000000000000000000000000
00000000000000000000000000000000000000001
>DM_train2641
TNRLVLSGTVCRAPLRKVSPSGIPHCQFVLEHRSVQEEAGFHRQAWCQMPVIVSGHENQAIT
HSITVGSRITVQGFISCHKAKNGLSKMVLHAEQIELIDSGD
00000000000000000000000000000000000000000000000000000000000000
00000000000000000000000000000000000001111
>DM_train2642
GPLGSEQIGKLRSELEMVSGNVRVMSEMLTELVPTQAEPADLELLQELNRTCRAMQQRVLEL

IPQIANEQLTEELLIVNDNLNNVFLRHERFERFRTGQTTKA
0000000000000000000000000000000000000000000000000000000000000
0000000000000000000000000000000000011111
>DM_train2643
SATEPPKIVWNEGKRRFETEDHEAFIEYKMRNNGKVMDLVHTYVPSFKRGLGLASHLCVAAF
EHASSHSISIIPSCSYVSDTFLPRNPSWKPLIHSEVFKSSI
1111000000000000000000000000000000000000000000000000000000000
0000000000000000000000000000000000001111
>DM_train2644
SNAMEFAPRSVVIEEFIDTLEPMMEAYGLDQVGIFEEHGEGNRYYVGYTINKDDEMITIHMP
FVKNERGELALEKQEWTVRKDGREKKGFHSLQEAMEEVIHS
1110000000000000000000000000000000000000000000000000000000000
0000000000000000000000000000000000000000
>DM_train2645
SANVTQAFGRRGPEQTQGNFGDQDLIRQGTDYKHWPQIAQFAPSASAFFGMSRIGMEVTPSG
TWLTYHGAIKLDDKDPQFKDNVILLNKHIDAYKTFPPTEPK
1111110000000000000000000000000000000000000000000000000000000
0000000000000000000000000000000000000001
>DM_train2646
DYKDDDDKVKLYKTNKYGTLYKSESASFTANTDIITRLTGPFRSMPQSGVLRKGLTIKYDEV
MKQDGHVWVGYNTNSGKRVYLPVRTWNESTGELGPLWGTIK
1111000000000000000000000000000000000000000000000000000000000
0000000000000000000000000000000000000000
>DM_train2647
AHHHHHHGSSTRGDLIRILGEIEEKMNELKMDGFNPDIILFGREAYNFLSNLLKKEMEEEGP
FTHVSNIKIEILEELGGDAVVIDSKVLGLVPGAAKRIKIIK
1111111110000000000000000000000000000000000000000000000000000
0000000000000000000000000000000000000000
>DM_train2648
GMTMFLHVVMMEFDDGIDAGFFRTVDEYVARMKRECDGLLLYHFGENVAARSQGYTHATSSA
FVDAAAHDAYQVCPAHVAMKAFMGPRIKRVVVYDGEVPAIG
0000000000000000000000000000000000000000000000000000000000000
0000000000000000000000000000000000000001
>DM_train2649
KILAVTACPTGIAHTFMAADALKEKAKELGVEIKVETNGSSGIKHKLTAQEIEDAPAIIVAA
DKQVEMERFKGKRVLQVPVTAGIRRPQELIEKAMNQDAPIY
0000000000011000000000000000000000000000000000000000000000000
0000000000000000000000000000000000000000
>DM_train2650
QDKKTMMMNEEDDGNGMDELLAVLGYKVRSSEMADVAQKLEQLEVMMSNVQEDDLSQLATET
VHYNPAELYTWLDSMLTDLNPPSSNAEYDLKAIPGDAILNQ
1111111111111110000000000000000000000000000000000111111100000
0000000000000000000111111111111111111111
>DM_train2651

MKRVMIGAVAALTMLSGQALAGDAAAGQAKAAVCAACHGADGNATIPGYPNLKGQNEQYIVS
SIKAYKNKERSGGLAAVMQAQASLLSDDDIANLAAYYSSLK
111111111111111111110000000000000000000000000000000000000000
00000000000000000000000000000000000000111
>DM_train2653
VNFEVKDQTLMMELVPERLRGETATFDIEADGKVYVEKGRRVTARHIRQLEKDGVNFIEVPV
EYIVGKVSAKDYVNEATGELIITANQEISLEALANLSQAG
000000000000000000000000000000000000000000000000000000000000
0000000000000000000000000000000000000011
>DM_train2654
MDSAKLIFINQINDCKDGQKLRFLGCVQSYKNGILRLIDGSSSVTCDVTVVLPDVSIQKHEW
LNIVGRKRQDGIVDVLLIRSAVGINLPRYRQMVSERQKCD
110000000000000000000000000000000000000000000000000000000000
0000000000000000000000000000000000000000
>DM_train2655
MRGRLQGVELIARDWIGLMVEVVESPNHSEVGIKGEVVDETQNTLKIMTEKGLKVVAKRGRT
FRVWYKGKIMRIKGDLINFRPEDRIKRGLMMLKRAKGVWI
111100000000000000000000000000000000000000000000000000000000
0000000000000000000000000000000000000000
>DM_train2656
GAVTGPRKGGSRRNAWGNQSYAELISQAIESAPEKRLTLAQIYEWMVRTVPYFKDKGDSNSS
AGWKNSIRHNLSLHSKFIKVHNEATGKSSWWMLNPEGGKS
111111111110000000000000000000000000000000000000000000000000
0000000000000000000000000000000000111111
>DM_train2657
EAQSEKEEAPKPLHKVVVCVSKKLSKKQSELNGIAASLGADYRRSFDETVTHFIYQGRPNDT
NREYKSVKERGVHIVSEHWLLDCAQECKHLPESLYPHTYN
111111111100000000000000000000000000000000000000000000000000
0000000000000000000000000000000000000000
>DM_train2658
GMTSPVAVIARFMPRPDARSALRALLDAMITPTRAEDGCRSYDLYESADGGELVLFERYRSR
IALDEHRGSPHYLNYRAQVGELLTRPVAVTVLAPLDEASA
111000000000000000000000000000000000000000000000000000000000
0000000000000000000000000000000000000001
>DM_train2659
MKAYLGLYTARLETPARSLKEKRALIKPALERLKARFPVSAARLYGLDAWGYEVVGFTLLGN
DPAWVEETMRAAARFLAEAGGFQVALEEFRLEAFELDGLL
000000000000000000000000000000000000000000000000000000000000
0000000000000000000000000000000000001111
>DM_train2660
MSILHTVNKSPFERNSLESCLKFATEGASVLLFEDGIYAALAGTRVESQVTEALGKLKLYVL
GPDLKARGFSDERVIPGISVVDYAGFVDLTTECDTVQAWL
100000000000000000000000000000000000000000000000000000000000
0000000000000000000000000000000000000000

>DM_train2661
HHMGNLNRCIADIVSLFITVMDKLRLEIRAMDEIQPDLRELMETMNRMSHLPPDFEGREKVS
QWLQKLSSMSASDELDDSQVRQMLFDLESAYNAFNRFLHS
110000000000000000000000000000000000000000000000000000000000
0000000000000000000000000000000000000001
>DM_train2662
MTQEVDQQILLQQLKSDYRQILLSYFTTDKALKEKIDKFINAVFCANIPVPEIIEIHMELID
EFSKQLRLEGRGDETLMDYRLTLIDILAHLCEAYRGAIFK
111000000000000000000000000011000000000000000000000000000000
0000000011100110000000000000000000000001
>DM_train2663
MKPRDINKLIASKIFGYEIKDDNIIKDGRYRLGIPLYSQNIESAWQVVEKLEYDVKVTKTDL
KPKYQVHVFVPGGVKMVFAETAPMAICKGALASVDIELQD
000000000000000000000000000000000000000000000000000000000000
0000000000000000000000000000000000111111
>DM_train2664
MELEDANVTKKVELRPLIGLTRGLPPTDLETITIDAIRTHRRLVEKADELFQALPETYKTGQ
ACGGPQHIRYIEASIEMHAQMSALNTLISILGFIPKVVVN
111111111110000000000000000000000000000000000000000000000000
0000000000000000000000000000000000000111
>DM_train2666
MKKHIIIKTIPKKEEIISRDLCDCIYYYDNSVICKPIGPSKVYVSTSLENLEKCLQLHYFKK
LVKNIEIFDEVHNSKPNCDKCLIVEIGGVYFVRRVNGVPR
000000000000000000000000000000000000000000000000000000000000
0000000000000000000000000000000000011111
>DM_train2667
RPGADCEVCKEFLNRFYKSLIDRGVNFSLDTIEKELISFCLDTKGKENRLCYYLGATKDAAT
KILSEVTRPMSVHMPAMKICEKLKKLDSQICELKYEKTLD
111000000000000000000000000000000000000000000000000000000000
0000000000000000000000000000000000000011
>DM_train2668
ITCPPPMSVEHADIWVKSYSLYSRERYICNSGFKRKAGTSSLTECVLNKATNVAHWTTPSLK
CIRDPALVHQRPAPPSTVTTAGVTPQPESLSPSGKEPAAS
000000000000000000000000000000000000000000000000000000000000
0000000000011111111111111111111111111111
>DM_train2669
MRRYEVNIVLNPNLDQSQLALEKEIIQRALENYGARVEKVAILGLMVLAYPIAKDPQGYFLW
YQVEMPEDRVNDLARELRIRDNVRRVMVVKSQEPFLANA
000000000000000000000000000000000000000000000000000000000000
000000000000000000000000000000011111111
>DM_train2670
QALSYREAVLRAVDRLNEQSSEANLYRLLELDQPPKADEDPGTPKPVSFTVKETVCPRPTRQ
PPELCDFKENGRVKQCVGTVTLDQIKDPLDITCNEVQGV
000000000000000000000000000000011111111100000000000000000000

000000000000000000000001111000000000000
>DM_train2671
MSPTSILDIRQGPKEPFRDYVDRFYKTLRAEQASQEVKNWMTETLLVQNANPDCKTILKALG
PGATLEEMMTACQGVGGPGHKARVLAEAMSQVTNPATIM
111000000000000000000000000000000000000000000000000000000000
000000000000111111111111111111111111111
>DM_train2672
MGRKLELTKAEKHVHNFMMDTQLTKRVKNAAANVLRETWLIYKNTKLVKKIDHAKVRKHQRK
FLQAIHQLRSVKMEQRKLNDQANTLVDLAKTQLEHHHHH
111111111111111111100000000000000000000000000000000000000000
000000000000000000000000000000000000000
>DM_train2673
GTEEYRIGEIFLAATEENKPQVFANAEKIVEQLKQGGSFVAYARQYSEASTAAVGGDLGWIR
LAQLPTELATTAASMGPGQLAGPVEIRGGFSILYLIDKR
100000000000000000000000000000000000000000000000000000000000
000000000000000000000000000000000000000
>DM_train2674
GMSDPRTQPLEIRPLMISRVMEVDWADGHTSRLTFEHLRVECPCAECKGHTPDQAQIVTGKE
HVSVVEVVPVGHYAVQLHFSDGHNTGIFTWEYLRRLDAE
110000000000000000000000000000000000000000111111111100000000
000000000000000000000000000000000000000
>DM_train2675
MGSDKIHHHHHHMEVKIEKPTPEKLKELSVEKWPIWEKEVSEFDWYYDTNETCYILEGKVEV
TTEDGKKYVIEKGDLVTFPKGLRCRWKVLEPVRKHYNLF
111111111111100000000000000000000000000000000000000000000000
000000000000000000000000000000000000000
>DM_train2676
MKKWSDTEVFEMLKRMYARVYGLVQGVGFRKFVQIHAIRLGIKGYAKNLPDGSVEVVAEGYE
EALSKLLERIKQGPPAAEVEKVDYSFSEYKGEFEDFETY
111111111110000000000000000000000000000000000000000000000000
000000000000000000000000000000000000000
>DM_train2677
RVQSGKIDCGDDAGWAKVPSDDPGRDNTRELAKNITFASPYCRPPVVLLSITQLDVEQSQNL
RVIARLYSVSPSGFKASCYTWHNTKVYSMSISWISIENY
000000000000000000000000000000000000000000000000000000000000
000000000000000000000000000000000000011
>DM_train2678
TTPVPVSGSLEVKVNDWGSGAEYDVTLNLDGQYDWTVKVKLAPGATVGSFWSANKQEGNGYV
IFTPVSWNKGPTATFGFIVNGPQGDKVEEITLEINGQVI
111100000000000000000000000000000000000000000000000000000000
000000000000000000000000000000000000000
>DM_train2679
QPVISAQEQETQIVLYGKLVEARQKHANKMDVPPAILATNKILVDMAKMRPTTVENVKRIDG
VSEGKAAMLAPLLEVIKHFCQTNSVQTDLFSSTKPQEEQ

000000000000000000000000000000000000000000000000000000000000000
00000000000000000000000000000001111111
>DM_train2680
MKTPEDCTGLADIREAIDRIDLDIVQALGRRMDYVKAASRFKASEAAIPAPERVAAMLPERA
RWAEENGLDAPFVEGLFAQIIHWYIAEQIKYWRQTRGAA
000000000000000000000000000000000000000111111111100000000000
00000000000000000000000000000001111111
>DM_train2681
GAMLYLIFYDITDDNLRNRVAEFLKKKGLDRIQYSVFMGDLNSSRLKDVEAGLKIIGNRKKL
QEDERFFILIVPITENQFRERIVIGYSGSEREEKSNVVW
100000000000000000000000000000000000000000000000000000000000000
000000000000000000000000011111111111
>DM_train2682
SNAEDITETSPDKWLIDGDTPLDEVERAIGYELPEGDYETISGLLFDHANALLKTGDVIEIP
LDFEPEDYLNNTSPTQRILRITVLEVERNVPVKLALALL
111000000000000000000000000000000000000000000000000000000000000
0000000000000000000000000000000000000
>DM_train2683
MKKRLTESQFQEAIQGLEVGQQTIEIARGVLVDGKPQATFATSLGLTRGAVSQAVHRVWAAF
EDKNLPEGYARVTAVLPEHQAYIVRKWEADAKKKQETKR
100000000000000000000000000000000000000000000000000000000000000
00000000000000000000000000000000111111
>DM_train2684
GAMAQVPTDPGYFSVLLDVKHFSPEEISVKVVGDHVEVHARHEERPDEHGFIAREFHRRYRL
PPGVDPAAVTSALSPEGVLSIQATPASAQASLPSPPAAK
000000000000000000000000000000000000000000000000000000000000000
0000000000000000000000000000000011111
>DM_train2685
GSHMYYVIFAQDIPNTLEKRLAVREQHLARLKQLQAENRLLTAGPNPAIDDENPSEAGFTGS
TVIAQFENLQAAKDWAAQDPYVEAGVYADVIVKPFKKVF
100000000000000000000000000000000000000000000000000000000000000
0000000000000000000000000000000000000
>DM_train2686
GSSSSLPLYNQQVGDCCIIRVSLDVDNGNMYKSILVTSQDKAPTVIRKAMDKHNLDEDEPED
YELLQIISEDHKLKIPENANVFYAMNSAANYDFILKKR
111111111111100000000000000000000000000000000000000000000000000
000000000000000000000000000000000000
>DM_train2687
SNTQAERSIIGMIDMFHKYTRRDDKIDKPSLLTMMKENFPNFLSACDKKGTNYLADVFEKKD
KNEDKKIDFSEFLSLLGDIATDYHKQSHGAAPCSGGSQ
000000000000000000000000000000000000000000000000000000000000000
000000000000000000000000000000001111
>DM_train2688
SSAEQSFLFSREEADTLRLKVEELEGERSRLEEEKRMLEAQLERRALQGDYDQSRTKVLHMS

LNPTSVARQRLREDHSQLQAECERLRGLLRAMERGGTV
111111110000000000000000000000000000000000000000000000000000
00000000000000000000000000000000011111
>DM_train2689
MEKMSSGTPTPSNVVLIGKKPVMNYVLAALTLLNQGVSEIVIKARGRAISKAVDTVEIVRNR
FLPDKIEIKEIRVGSQVVTSQDGRQSRVSTIEIAIRKK
111111111110000000000000000000000000000000000000000000000000
00000000000000000000000000000000000000
>DM_train2690
MRGSHHHHHHGSMAQKTFKVTADSGIHARPATVLVQTASKYDADVNLEYNGKTVNLKSIMGV
MSLGIAKGAEITISASGADENDALNALEETMKSERLGE
111111111111110000000000000000000000000000000000000000000000
00000000000000000000000000000000000000
>DM_train2691
ELIRVAILWHEMWHEGLEEASRLYFGERNVKGMFEVLEPLHAMMERGPQTLKETSFNQAYGR
DLMEAQEWCRKYMKSGNVKDLTQAWDLYYHVFRRISKQ
111111100000000000000000000000000000000000000000000000000000
00000000000000000000000000000000000000
>DM_train2692
MSLTDSKVKNAKSLEKEYKLTDGFGMHLLVHPNGSKYWRLSYRFEKKQRLLALGVYPAVSLA
DARQRRDEAKKLLAAGIDPSAKKQADNKTIQEKRNNTR
100000000000000000000000000000000000000000000000000000000000
00000000000000000000000000000000001111
>DM_train2693
GAHVSRVEKLPKDYQIVYKEIQKYLFKVGPVELNEGIGLLSEILGFFEEGAAAGKGVLDVTG
TDVAAFCDALIGDSKTYADLYQESIQQHVDKAMKNMKD
110000000000000000000000000000000000000000000000000000000000
00000000000000000000000000000011111111
>DM_train2694
SMDNLVLIRMKPDENGRFGFNVKGGYDQKMPVIVSRVAPGTPADLCVPRLNEGDQVVLINGR
DIAEHTHDQVVLFIKASCERHSGELMLLVRPNAVESTV
100000000000010000000000000000000000000000000000000000000000
00000000000000000111111000000000000000
>DM_train2695
GAMGNLRLIGVPESDVENGTKLENTLQDIIQENFPNLARQANVQIQEIQRTPQRYSSRRATP
RHIIVRFTKVEMKEKMLRAAREKGRVTLKGKPIRLTVD
100000000000000000000000000000000000000000000000000000000000
00000000000000000000000000000000000000
>DM_train2696
MAEPGIDKLFGMVDSKYRLTVVVAKRAQQLLRHRFKNTVLEPEERPKMRTLEGLYDDPNAVT
WAMKELLTGRLFFGENLVPEDRLQKEMERLYPTEEEA
000000000000000000000000000000000000000000000000000000000000
0000000000000000000000000000000111111
>DM_train2697

AKVKIKPLEDKILVQAGEAETMTPSGLVIPENAKEKPQEGTVVAVGPGRWDEDGAKRIPVDV
SEGDIVIYSKYGGTEIKYNGEEYLILSARDVLAVVSK
00000000000000001111111111111111110000000000000000000000000000
0000000000000000000000000000000000000
>DM_train2698
MSRSRQPPLVTGISPNEGIPWTKVTIRGENLGTGPTDLIGLTICGHNCLLTAEWMSASKIVC
RVGQAKNDKGDIIVTTKSGGRGTSTVSFKLLKPEKIG
11100000000000000000000000000000000000000000000000000000000000
0000000000000000000000000000000001111
>DM_train2699
PITGPHIAYTEAVSDTQIMLKWTYIPSSNNNTPIQGFYIYYRPTDSDNDSDYKRDVVEGSKQ
WHMIGHLQPETSYDIKMQCFNEGGESEFSNVMICETK
11100000000000000000000111111111000000000000000000000000000000
0000000000000000000000000000000000000
>DM_train2700
AELRLLMFEQPGCLYCARWDAEIAPQYPLTDEGRAAPVQRLQMRDPLPPGLELARPVTFTPT
FVLMAGDVESGRLEGYPGEDFFWPMLARLIGQAEPGQ
10000000000000000000000000000000000000000000000000000000000000
0000000000000000000000000000000000011
>DM_train2701
HRVINHPYYFPFNGKQAEDYLRSKERGDFVIRQSSRGDDHLAITWKLDKDLFQHVDIQEMEK
ENPLAMGKVLVVEGQRYHDLDQIIVEYLQNKIRLLNE
00000000000000000000000000000000000000000000000000000000000011
1111110000000000000000000000000000001
>DM_train2702
STMPLAELGARLYREKACFSCHSIDGSRLVGPSFKGLYGSTRTFEDGTTAVADENYLRESIL
QPGAKVVQGYPNVMPASYASLSEREVAALIEFIKQQQ
11000000000000000000000000000000000000000000000000000000000000
0000000000000000000000000000000000000
>DM_train2703
MNERLKEKLAVLPEQPGCYLMKDKHGTVIYVGKAKSLKERVRSYFTGTHDGKTQRLVEEIAD
FEYIVTSSNAEALILEMNLIKKHDPKYNVMLKDDKSY
11111110000000000000000000000000000000000000000000000000000000
0000000000000000000000000000000001111
>DM_train2704
GHMAKGYWIAQVDVRDSERYKDYVSTAKPAFERFGANFLARGGSVTELEGTARARNVVIEFP
SVQHAIDCYNSPEYQAAAKIRQEVADAEMMIVEGIGS
11000000000000000000000000000000000000000000000000000000000000
0000000000000000000000000000000000001
>DM_train2705
SDLQRLKFIRHARQLGFSLESIRELLSIRIDPEHHTCQESKGIVQERLQEVEARIAELQSMQ
RSLQRLNDACCGTAHSSVYCSILEALEQGASGVKSGC
11000000000000000000000000000000000000000000000000000000000000
0000000000000000000000000000011111111

>DM_train2706
MMLIDCPNCGPRNENEFKYGGEAHVAYPADPHALSDKQWSRYLFYRQNKKGIFAERWVHAAG
CRKWFNALRDTVTYEFKAIYPAGAPRPEIHSAEGGTR
000000000000000000000000000000000000000000000000000000000000000
0000000000000000000000000000011111111
>DM_train2707
ASWSHPQFEKSGGGGGLVPRGSKRADAGTASALAASQLPQATMPGKSMVAIAGSSYQGQNGL
AIGVSRISDNGKVIIRLSGTTNSQGKTGVAAGVGYQW
111111111111111111111000000000000000000000000000000000000000000
0000000000000000000000000000000000000
>DM_train2708
SDFANLSSTNKELSPQYNWVACGILEGGLKAAGVLEEGQYNRELAEAIAAKGEGFWTTQFPQ
IGDWNEDQAAALADRAQTCGLVKADTYLSELSSNFSS
111111111111000000000000000000000000000000000000000000000000000
000000000000000000000000001111111111
>DM_train2709
MTVFVDHKIEYMSLEDDAELLKTMAHPMRLKIVNELYKHKALNVTQIIQILKLPQSTVSQHL
CKMRGKVLKRNRQGLEIYYSINNPKVEGIIKLLNPIQ
111111111000000000000000000000000000000000000000000000000000000
0000000000111111100000000000000000001
>DM_train2710
MFEQRVNSDVLTVSTVNSQDQVTQKPLRDSVKQALKNYFAQLNGQDVNDLYELVLAEVEQPL
LDMVMQYTRGNATRAALMMGINRGTLRKKLKKYGMN
111111111000011111111111000000000000000000011110000000000000000
000000000000000000000000000000000000
>DM_train2711
MPSKKKKYNARFPPARIKKIMQTDEEIGKVAAAVPVIISRALELFLESLLKKACQVTQSRNA
KTMTTSHLKQCIELEGDPAANKARKEAELAAATAEQ
111111111000000000000000000000000000000000000000000000000000011
100000000000011111111111111111111111
>DM_train2712
DPNLWTVKCKIGEERATAISLMRKFIAYQFTDTPLQIKSVVAPEHVKGYIYVEAYKQTHVKQ
AIEGVGNLRLGYWNQQMVPIKEMTDVLKVVKEVANL
000000000000000000000000000000000000000000000000000000000000000
000000000000000000000000000000001111
>DM_train2713
MNTLELSARVLECGAMRHTPAGLPALELLLVHESEVVEAGHPRRVELTISAVALGDLALLLA
DTPLGTEMQVQGFLAPARKDSVKVLKLHLQQARRIAG
000000000000000000000000000000000001111110000000000000000000000
000000000000000000000000000000000011
>DM_train2714
MNSCILQATVVEAPQLRYAQDNQTPVAEMVVQFPGLSSKDAPARLKVVGWGAVAQELQDRCR
LNDEVVLEGRLRINSLLKPDGNREKQTELTVTRVHH
000000000000000000000000000000000001111100000000000000000000000

000000000000000111111110000000000000
>DM_train2715
SPLAVGLAAHLRQIGGTMYGAYWCPHCQDQKELFGAAFDQVPYVECSPNGPGTPQAQECTEA
GITSYPTWIINGRTYTGVRSLEALAVASGYPLEEGR
000000000000000000000000000000000000000000000000000000000000
000000000000000000000000000000001111
>DM_train2716
SWVWNQMFVLEEFSGPEPILVGRLHTDLDPGSKKIKYILSGDGAGTIFQINDITGDIHAIKR
LDREEKAEYTLTAQAVDFETNKPLEPPSEFIIKVQD
000000000000000000000000000011110000000000000000000000000000
000000000000000000000000000000000000
>DM_train2717
MRGSGMASSYSKWFNLEKIHSIEVQSLPEFFTNRIPSKTPEVYMRYRNFMVNSYRLNPNEYF
SVTTARRNVSGDAAALFRLHKFLTKWGLINYQVDSK
111111111100000000000000000000000000000000000000000000000000
000000000000000000000000000000000111
>DM_train2718
MEIIPSESHPHIQLLKSNRELLVTHIRNTQCLVDNLLKNDYFSAEDAEIVCACPTQPDKVRK
ILDLVQSKGEEVSEFFLYLLQQLADAYVDLRPWLLE
111110000000000000000000000000000000000000000000000000000000
000000000000000000000000000000000000
>DM_train2720
MSLKTTLPISLDWSTEEVIDVVHFFQAIEQAYDQGIAREDLLGKYRRFKEIVPSKSEEKQLF
RAYEQENDVSCYQTIKKAREEMEEHIQMEGHHHHHH
111111110000000000000000000000000000000000000000000000000000
000000000000000000000000000011111111
>DM_train2721
ATYKVTLINEAEGLNKTIEVPDDQYILDAAEEAGIDLPYSCRAGACSTCAGKLISGTVDQSD
QSFLDDDQIEAGYVLTCVAYPTSDCVIETHKEEELY
000000000111000000000000000000000000000000000000000000000000
000000000000000000000000000000000000
>DM_train2722
LSIPEDYQARLQPNRVEGSYPLVRMEFTGATVDAPLMSQISRKYNIDVSILSSDLDYAGGVK
FGMMVAELFGNEQDDSAAIEYLRENNVKVEVLGYVL
100000000000000000000000000000000000000000000000000000000000
000000000000000000000000000000000000
>DM_train2723
MSNLKWFSGGDDRRKKAEVIITELLDDLEIDLGNESLRKVLGSYLKKLKNEGTSVPLVLSRM
NIEISNAIKKDGVSLNENQSKKLKELMSISNIRYGY
111111111000000000000000000000000000000000000000000000000000
000000000000000000000000000000111111
>DM_train2724
PLKPEEHEDILNKLLDPELAQSERTEALQQLRVNYGSFVSEYNDLTKSHEKLAAEKDDLIVS
NSKLFRQIGLTEKQEEDHKKADISETITIEDLEAK

000000000000000000000000000000000000000000000000000000000000
00000000000111111111111111111111111
>DM_train2725
MKSGIYQIKNTLNNKVYVGSAKDFEKAWKRHFKDLEKGCHSSIKLQRSFNKHGNVFECSILE
EIPYEKDLIIERENFWIKELNSKINGYNIADATFG
000000000000000000000000000000000000000000000000000000000000
00000000000000000000000000000001111
>DM_train2726
GSHMEEIRNLTVKDFRVQELPLARIKKIMKLDEDVKMISAEAPVLFAKAAQIFITELTLRAW
IHTEDNKRRTLQRNDIAMAITKFDQFDFLIDIVPR
111111111111111111100000000000000000000000000000000000000000
00000000000000000000000000000000000
>DM_train2727
MEPNTPTARTFDWMKVKRNPPKTAKVSEPGLGSPSGLRTNFTTRQLTELEKEFHFNKYLSRA
RRVEIAATLELNETQVKIWFQNRRMKQKKREREGG
111111100000000011111111111111111110000000000000000000000000
00000000000000000000000000000000000
>DM_train2728
MDAKARNCLLQHREALEKDIKTSYIMDHMISDGFLTISEEEKVRNEPTQQQRAAMLIKMILK
KDNDSYVSFYNALLHEGYKDLAALLHDGIPVVSSS
000000000000000000000000000000000000000000000000000000000000
00000000000000000000000000000011111
>DM_train2729
RDSGTVWGALGHGINLNIPNFQMTDDIDEVRWERGSTLVAEFKRKPFLKSGAFEILANGDLK
IKNLTRDDSGTYNVTVYSTNGTRILDKALDLRILE
111000000000000000000000000000000000000000000000000000000000
00000000000000000000000000000000000
>DM_train2730
MDFSQLGGLLDGMKKEFSQLEEKNKDTIHTSKSGGGMVSVSFNGLGELVDLQIDDSLLEDKE
AMQIYLMSALNDGYKAVEENRKNLAFNMLGNFAKL
111111100000000000000000000000000000000000000000000000000000
00000000000000000000000000000111111
>DM_train2731
GMNHNYVEVKYPAIFRDEGTYWDVRFPDVPAAQTFGASVQVAADNAANALAIALFEQSLPPA
SDPQYWRLASTEFVVWITMADVQFGPGADTPEPMN
111111100000000000000000000000000000000000000000000000000000
00000000000000000000000000011111111
>DM_train2732
ADPALADVCRTKLPSQAQDTLALIAKNGPYPYNRDGVVFENRESRLPKKGNGYYHEFTVVTP
GSNDRGTRRVVTGGYGEQYWSPDHYATFQEIDPRC
111000000000000000000000000000000000000000000000000000000000
00000000000000000000000000000000000
>DM_train2733
GMKRPGAIPTVQIDNERVKVTEWRFPPGGETGWHRHSMDYVVVPMTTGPLLLETPEGSVTSQ

LTRGVSYTRPEGVEHNVINPSDTEFVFVEIEIKAA
111000000000000000000000000000000000000000000000000000000000
00000000000000000000000000000000011
>DM_train2734
MGEWEIIDIGPFTQNLGKFAVDEENKIGQYGRLTFNKVIRPCMKKTIYENEGFREIKGYEYQ
LYVYASDKLFRADISEDYKTRGRKLLRFNGPVPPP
100000000000000000000000000000000000000000000000000000000000
00000000000000000000000000000000000
>DM_train2735
MAMIKMSPEEIRAKSQSYGQGSDQIRQILSDLTRAQGEIAANWEGQAFSRFEEQFQQLSPKV
EKFAQLLEEIKQQLNSTADAVQEQDQQLSNNFGLQ
110000000000000000000000000000000000000000000000000000000000
00000000000000000000000111111111111
>DM_train2736
GPSRVVYLGSIPYDQTEEQILDLCSNVGPVINLKMMFDPQTGRSKGYAFIEFRDLESSASAV
RNLNGYQLGSRFLKCGYSSNSDISGVSQQQQQQYN
000000000000000000000000000000000000000000000000000000000000
00000000000000000000000001111111111
>DM_train2737
GKVVIYSEPDVSEKCIEVFSDIQDCSSWSLSPVILIKVVRGCWILYEQPNFEGHSIPLEEGE
LELSGLWGIEDILERHEEAESDKPVVIGSIRHVV
000000000111000000000000000000000000000000000000000000000000
0000000000000000000000000000000000
>DM_train2738
HETRFEAAVKVIQSLPKNGSFQPTNEMMLKFYSFYKQATEGPCKLSRPGFWDPIGRYKWDAW
SSLGDMTKEEAMIAYVEEMKKIIETMPMTEKVEE
000000000000000000000000000000000000000000000000000000000000
0000000000000000000000000111111111
>DM_train2739
SETRKTEVPSDKLELLLDIPLKVTVELGRTRMTLKRVLEMIHGSIIELDKLTGEPVDILVNG
KLIARGEVVVIDENFGVRITEIVSPKERLELLNE
111111111111111000000000000000000000000000000000000000000000
0000000000000000000000000000000000
>DM_train2740
GPMATESPNSVQKIVVHLRATGGAPILKQSKFKVSGSDKFANVIDFLRRQLHSDSLFVYVNS
AFSPNPDESVIDLYNNFGFDGKLVVNYACSMAWG
111111111110000000000000000000000000000000000000000000000000
0000000000000000000000000000000001
>DM_train2741
GHMSLEEWIKADSLEKADEYHKRYNYAVTNPVRRKILRMLDKGRSEEEIMQTLSLSKKQLDY
HLKVLEAGFCIERVGERWVVTDAGKIVDKIRGGS
000000000000000000000000000000000000000000000000000000000000
0000000000000000000000000011111111
>DM_train2742

EVNEKAPAQARPTVFRWTGGGKEVYLSGSFNNWSKLPMTRSQNNFVAILDLPEGEHQYKFFV
DGQWTHDPSEPIVTSQLGTVNNIIQVKKTDFEVF
11111111000000000000000000000000000000000000000000000000000000
0000000000000000000000000001111111
>DM_train2743
MGTEPPENCQDDFNFNYVSDQEIEVYHVDKGWSAGWNYVCLNDYCLPGNKSNGAFRKTFNAV
LGQDYKLTFKVEDRYGQGQQILDRNITFTTQVCN
11111110000000000000000000000000000000000000000000000000000000
0000000000000000000000000000000001
>DM_train2744
GSHMSTLKEVQDNITLHEQRLVTTRQKLKDAERAVELDPDDVNKSTLQSRRAAVSALETKLG
ELKRELADLIAAQKLASKPVDPTGIEPDDHLKEK
11111000000000000000000000000000000000000000000000000000000000
0000000000000011111111111111111111
>DM_train2745
MDVLQEQVFKDLKSRGFKIIEQLDDKIFIAEKKERYLFYVMVEGVEVTIQTLLSVINMGETL
SMPVVLALVSNDGTVTYYYVRKIRLPRNIYAEAV
00000000000000000000000000000000000000000000000000000000000000
0000000000000000000000000000000111
>DM_train2746
GSHMLEQKKANIYKGKITRIEPSLEAAFVDYGAERHGFLPLKEIAREYFPANYSAHGRPNIK
DVLREGQEVIVQIDKEERGNKGAALTTFISLAGS
11111111000000000000000000000000000000000000000000011111100000
0000000000000000000000000000000000
>DM_train2747
AMSDLELRQCLPCGPGGAGRCFGPSICCGDELGCFVGTAEALRCQEENYLPSPCQSGQKPCG
SGGRCAAAGICCNDESCVTEPECREGVGFPRRV
11111100000000000000000000000000000000000000000000000000000000
000000000000000000000001111111111
>DM_train2748
VPDPRGIIINLDEGELCLNSAQCKSNCCQHDTILSLLRCALKARENSECSAFTLYGVYYKCP
CERGLTCEGDKSLVGSITNTNFGICHNVGRSDS
11111000000000000000000000000000000000000000000000000000000000
000000000000000000000000000011111
>DM_train2749
MGSDKIHHHHHHMDLELKELQSKMKEMYFEKDSQRGIYATFTWLVEEVGELAEALLSNNLDS
IQEELADVIAWTVSIANLEGIDIEEALKKKYKL
11111111111100000000000000000000000000000000000000000000000000
000000000000000000000000000000000
>DM_train2750
MAYGKAHLEAQLKRALAEEIQALEDPRLFLLTVEAVRLSKDGSVLSVYVEAFREEEGALRAL
SRAERRLVAALARRVRMRRLPRLEFLPWRASPA
11000000000000000000000000000000000000000000000000000000000000
000000000000000000000000000000111

>DM_train2751
AKKKSKLEIIQAILEACKSGSPKTRIMYGANLSYALTGRYIKMLMDLEIIRQEGKQYMLTKK
GEELLEDIRKFNEMRKNMDQLKEKINSVLSIRQ
11000000000000000000000000000000000000000000000000000000000000
000000000000000000000000000000111
>DM_train2752
MEDERIKLLFKEKALEILMTIYYESLGGNDVYIQYIASKVNSPHSYVWLIIKKFEEAKMVEC
ELEGRTKIIRLTDKGQKIAQQIKSIIDIMENDT
11110000000000000000000000000000000000000000000000000000000000
000000000000000000000000000001111
>DM_train2753
KQKVINVKEVRLSPTIEEHDFNTKLRNARKFLEKGDKVKATIRFKGRAITHKEIGQRVLDRL
SEACADIAVVETAPKMDGRNMFLVLAPKNDNK
11110000000000000000000000000000000000000000000000000000000000
00000000000000000000000000000000
>DM_train2754
KPAKPAAKSIVTLDVKPWDDETNLEEMVANVKAIEMEGLTWGAHQFIPIGFGIKKLQINCVV
EDDKVSLDDLQQSIEEDEDHVQSTDIAAMQKL
11110000000000000000000000000000000000000000000000000000000000
00000000000000000000000000000000
>DM_train2755
GHMSRTVMCRKYHEELPGLDRPPYPGAKGEDIYNNVSRKAWDEWQKHQTMLINERRLNMMNA
EDRKFLQQEMDKFLSGEDYAKADGYVPPSAGS
00000000000000000000000000000000000000000000000000000000000000
00000000000000000001111111111111
>DM_train2756
GSRSLQLDKLVNEMTQHYENSVPEDLTVHVGDIVAAPLPTNGSWYRARVLGTLENGNLDLYF
VDFGDNGDCPLKDLRALRSDFLSLPFQAIECS
11110000000000000000000000000000000000000000000000000000000000
00000000000000000000000000000001
>DM_train2757
EDDEDYGDLGGCPFLVAENKTGYPTIVACKQDCNGTTETAPNGTRCFSIGDEGLRRMTANLP
YDCPLGQCSNGDCIPKETYEVCYRRNWRDKKN
11111111110000000000000000000000000000000000000000000000000000
00000000000000000000000000111111
>DM_train2758
GASIVGSWVEPVPGLEGQVQGIKMEEGGVASSVNMATLVYESWKQEGTKLILTGKSIGNGQT
IEFVDTMDIKRLTADSLVLDNQGMEIRYAKQK
10000000000000000000000000000000000000000000000000000000000000
00000000000000000000000000000001
>DM_train2759
MRKVRPEELPALLEEGVLVVDVRPADRRSTPLPFAAEWVPLEKIQKGEHGLPRRPLLLVCEK
GLLSQVAALYLEAEGYEAMSLEGGLQALTQGK
00000000000000000000000001110000000000000000000000000000000000

00000000000000000000000000011111
>DM_train2760
EDMEIAYPITCGESKAILLWKKFVCPGINVKCVKFNDQLISPKHFVHLAGKSTLKDWKRAIR
LGGIMLRKMMDSGQIDFYQHDKVCSNTCRSTK
10000000000000000000000000000000000000000000000000000000000000
00000000000000000000000000001111
>DM_train2761
QQRKVLTLEKGDNQTFGFEIQTYGLHHREEQRVEMVTFVARVHESSPAQLAGLTPGDTIASV
NGLNVEGIRHREIVDIIKASGNVLRLETLYGT
00000000000000000000000001111111000000000000000000000000000000
00000000000000000000000000000000
>DM_train2762
GGAHKVRAGGPGLERAEAGVPAEFSIWTREAGAGGLAIAVEGPSKAEISFEDRKDGSCGVAY
VVQEPGDYEVSVKFNEEHIPDSPFVVPVASPS
00000000000000000000000000000000000000000000000000000000000000
00000000000000000000000000000001
>DM_train2763
MIKVTVTNSFFEVTGHAPDKTLCASVSLLTQHVANFLKAEKKAKIKKESGYLKVKFEELENC
EVKVLAAMVRSLKELEQKFPSQIRVEVIDNGS
00000000000000000000000000000000000000000000000000000000000000
00000000000000000000000000000111
>DM_train2764
GVDRDYLQSEYGVLKAGQCYKVVRSFRDYRNINYERGDVMRFLGSNFVPYESGLSLFFDKNG
SERQIMLCVRPEFQMEIAHHLDSYFCKLDDN
11100000000000000000000000000000000000000000000000000000000000
0000000000000000000000000000111
>DM_train2765
MQQSLAVKTFEDLFAELGDRARTRPADSTTVAALDGGVHALGKKLLEEAGEVWLAAEHESND
ALAEEISQLLYWTQVLMISRGLSLDDVYRKL
11111100000000000000000000000000000000000000000000000000000000
0000000000000000000000000000000
>DM_train2766
MASTRGSGRPWSAKENKAFERALAVYDKDTPDRWANVARAVEGRTPEEVKKHYEILVEDIKY
IESGKVPFPNYRTTGGNMKTDEKRFRNLKIR
11111110000000000000000000000000000000000000000000000000000000
0000000011111111111111111111111
>DM_train2767
GSRMSAEASARPLRVGSRVEVIGKGHRGTVAYVGATLFATGKWVGVILDEAKGKNDGTVQGR
KYFTCDEGHGIFVRQSQIQVFEDGADTTSPE
11111111111100000000000000000000000000000000000000000000000000
0000000000000000000001111111111
>DM_train2768
GSHMNTEVSSEIYQWVRDELKRAGISQAVFARVAFNRTQGLLSEILRKEEDPKTASQSLLVN
LRAMQNFLQLPEAERDRIYQDERERSLRKRK

11111100000000000000000000000000000000000000000000000000000000
0000000000000000000000000000111
>DM_train2769
AEKCFDHAAGTSYVVGETWEKPYQGWMMVDCTCLGEGSGRITCTSRNRCNDQDTRTSYRIGD
TWSKKDNRGNLLQCICTGNGRGEWKCERHTS
00000000000000000000000000000000000000000000000000000000000000
0000000000000000000000000000111
>DM_train2770
APDTQDCPECTLQENPFFSQPGAPILQCMGCCFSRAYPTPLRSKKTMLVQKNVTSESTCCVA
KSYNRVTVMGGFKVENHTACHCSTCYYHKS
11111000000000000000000000000000000000000000000000000000000000
000000000000000000000000000000
>DM_train2771
ISEFGSEPIDPSKLEFARALYDFVPENPEMEVALKKGDLMAILSKKDPLGRDSDWWKVRTKN
GNIGYIPYNYIEIIKRRKKIEHVDDETRTH
11111111110000000000000000000000000000000000000000000000000000
000000000000000001111111111111
>DM_train2772
WKDKEFQVLFVLTILTLISGTIFYSTVEGLRPIDALYFSVVTLTTVGDGNFSPQTDFGKIFT
ILYIFIGIGLVFGFIHKLAVNVQLPSILSN
11110000000000000000000000000000000000000000000000000000000000
000000000000000000000000000000
>DM_train2773
MSTISRDSCPALRAGVRLQHDRARDQWVLLAPERVVELDDIALVVAQRYDGTQSLAQIAQTL
AAEFDADASEIETDVIELTTTLHQKRLLRL
11000000000000000000000000000000000000000000000000000000000000
000000000000000000000000000000
>DM_train2774
KNTGDEVVAIISQNGKVIREIPLTGHKGNEQFTIKGKGAQYNLMEVDGERIRIKEDNSPDQV
GVKMGWKSKAGDTIVCLPHKVFVEIKSTQK
11100000000000000000000000000000000000000000000000000000000000
000000000000000000000000000011
>DM_train2775
MPLVGFMKEKKRATFYLYKNIDGRKLRYLLHKLENVENVDIDTLRRAIEAEKKYKRSITLTE
EEEVIIQRLGKSANLLLNCELVKLDEGERA
11111000000000000000000000000000000000000000000000000000000000
000000000000000000000000011111
>DM_train2776
SNAMVKETKLYDLLGVSPSANEQELKKGYRKAALKYHPDKPTGDTEKFKEISEAFEILNDPQ
KREIYDQYGLEAARSGGPSFGPGGPGGAGG
11100000000000000000000000000000000000000000000000000000000000
000000000000000000000001111111
>DM_train2777
MSKMLYRVELIITNKEGVRDPEGETIQRYVVSRFSDKIIETRAGKYLVFRVNSSSQQEATEL

VKKLADEMRLYNPIVHKIEIRANRIEDSSN
111000000000000000000000000000000000000000000000000000000000000
00000000000000000000000001111
>DM_train2778
SASDFEERVVTIPLRDARAEPNHKRADKAMILIREHLAKHFSVDEDAVRLDPSINEAAWARG
RANTPSKIRVRAARFEEEGEAIVEAETAE
111111000000000000000000000000000000000000000000000000000000000
0000000000000000000000000111
>DM_train2779
MAKNTSCGVQLRIRGKVQGVGFRPFVWQLAQQLNLHGDVCNDGDGVEVRLREDPETFLVQLY
QHCPPLARIDSVEREPFIWSQLPTEFTIR
111000000000000000000000000000000000000000000000000000000000000
0000000000000000000000000000
>DM_train2780
GSHMDINNKARIHWACRRGMRELDISIMPFFEHEYDSLSDDEKRIFIRLLECDDPDLFNWLM
NHGKPADAELEMMVRLIQTRNRERGPVAI
111000000000000000000000000000000000000000000000000000000000000
0000000000000000000000000001
>DM_train2781
AHHHHHHENLYFQAGSTTLKLLRKEIDKIDNQIISLLKKRLEIAQAIGKIKKELNLPIEDRK
REEEVLRRAGEFREIFEKILEVSKDVQRL
111111111111110000000000000000000000000000000000000000000000000
0000000000000000000000000001
>DM_train2782
SNAMKAIITVVGKDKSGIVAGVSGKIAELGLNIDDISQTVLDEYFTMMAVVSSDEKQDFTYL
RNEFEAFGQTLNVKINIQSAAIFEAMYNI
110000000000000000000000000000000000000000000000000000000000000
0000000000000000000000011111
>DM_train2783
LNSAPTPRDVVANAPAPVQAAVAGAQEYAAQAGLNTEELAVDALYNAIKVRLAGTGLGIPPQ
IEAFYQANRTNFNGFYMANRGAIDFIFSM
000000000000000000000000000000000000000000000000000000011100000
0000000000000000000000000001
>DM_train2784
MEIMDEIKVNLQKEVSLEEAERYAKNIASKYGDGILLSVHDSKTGYRAPEVYCCGEKPWEVY
ACNRGANLKISVNQFEFYFRIEVEGQAKY
110000000000000000000000000000000000000000000000000000000000000
00000000000000000000000001111
>DM_train2785
MNKKELIDRVAKKAGAKKKDVKLILDTILETITEALAKGEKVQIVGFGSFEVRKAAARKGVN
PQTRKPITIPERKVPKFKPGKALKEKVK
000000000000000000000000000000000000000000000000000001111111111
111111111111100000000000000
>DM_train2786

MAVTYEKTFEIEIINELSASVYNRVLNYVLNHELNKNDSQLLEVNLLNQLKLAKRVNLFDYS
LEELQAVHEYWRSMNRYSKQVLNKEKVA
00000000000000000000000000000000000000000000000000000000000000
0000000000000000000000001111
>DM_train2787
MQKEQLSALMDGETLDSELLNELAHNPEMQKTWESYHLIRDSMRGDTPEVLHFDISSRVMAA
IEEEPVRQPATLIPEAQPAPHQWQKMPF
00000000000000000000000000000000000000000000000000000000000000
0011111111111111111111111111
>DM_train2788
VELLQVDADLQAEIVGKYNADLQKAVQIEEKKASEIATEAVKEHVTAEYEERYAEHEEHDRI
MRDVAEILEQMEHAEVRRLITEDKVRPD
11110000000000000000000000000000000000000000000000000000000000
0000000000000000000011111111
>DM_train2789
DVITVYKDCNYTGFSGGLTIGDYNLARLNSLGVLNDDISSLRITQGYQAILYQDDNFGGAST
VINSDNSCLNTTWNDKVSSIRVIANGTT
00000000000000000000000000000000000000000000000000000000000000
0000000000000000000000001111
>DM_train2790
MGKRLYAVAYDIPDDTRRVKLANLLKSYGERVQLSVFECYLDERLLEDLRRRARRLLDLGQD
ALRIYPVAGQVEVLGVGPLPELREVQVL
10000000000000000000000000000000000000000000000000000000000000
0000000000000000000001111111
>DM_train2791
QFSDMRISINQTPGKSLDFGFTIKWDIPGIFVASVEAGSPAEFSQLQVDDEIIAINNTKFSY
NDSKEWEEAMAKAQETGHLVMDVRRYGK
00000000000000000000000000000000000000000000000000000000000000
0000000000000000000000000001
>DM_train2792
SNAEEKVVAEQQADGTWLMDGWISIRKASNLLEHDLVDEAERYSTLGGYLLWQFGYIPAAGE
QITVDGLIFEIVSVNKHNIGKVRVHRTQ
11110000000000000000000000000000000110000000000000000000000000
0000000000000000000000000001
>DM_train2793
DEVENLYTQVADNEYLVQGRMLIDEFNEVFETDLHMSDVDTMAGYLITALGTIPDEGEKPSF
EVGNIKLTAEEMEGTRLLVLRVHFYDEE
11110000000000000000000000000000000000000000000000000000000000
0000000000000000000000000011
>DM_train2794
MNLEMLESQPVPEIDTLREEIDRLDAEILALVKRRAEVSKAIGKARMASGGTRLVHSREMKV
IERYSELGPDGKDLAILLLRLGRGRLGH
11111111111100000000000000000000000000000000000000000000000000
0000000000000000000000000001

>DM_train2795
AELKEFLNKPAATSDVPPAPAGFDFDAAKKLVDVRCNKCHTLDSVADLFRTKYKKTGQVNLI
VKRMQGFPGSGISDDDAKTIGIWLHEKF
111111111111100000000000000000000000000000000000000000000000
0000000000000000000000000000
>DM_train2796
VKVKVKYFARFRQLAGVDEEEIELPEGARVRDLIEEIKKRHEKFKEEVFGEGYDEDADVNIA
VNGRYVSWDEELKDGDVVGVFPPVSGG
000000000000000000000000000000000000000000000000000000000000
000000000000000000000000011
>DM_train2797
GPLGSQVRVKAYYRGDIMITHFEPSISFEGLCNEVRDMCSFDNEQLFTMKWIDEEGDPCTVS
SQLELEEAFRLYELNKDSELLIHVFPC
111110000000000000000000000000000000000000000000000000000000
000000000000000000000000001
>DM_train2799
GSAQLLKSVFVKNVGWATQLTSGAVWVQFNDGSQLVMQAGVSSISYTSPDGQTTRYGENEKL
PEYIKQKLQLLSSILLMFSNPTPVFQ
111111100000000000000000000000000000000000000000000000000000
00000000000000000000111111
>DM_train2801
MSDAFTDVAKMKKIKEEIKAHEGQVVEMTLENGRKRQKNRLGKLIEVYPSLFIVEFGDVEGD
KQVNVYVESFTYSDILTEKNLIHYLD
000000000000000000000000000000011111100000000000000000000000
00000000000000000000000000
>DM_train2802
GAMVLSQRQRDELNRAIADYLRSNGYEEAYSVFKKEAELDMNEELDKKYAGLLEKKWTSVIR
LQKKVMELESKLNEAKEEFTSGGPLG
111000000000000000000000000000000000000000000000000000000000
00000000000000011111111111
>DM_train2803
LSSNSSMNPKSLTDPKLLKNIPMWLKSLRLHKYSDALSGTPWIELIYLDDETLEKKGVLALG
ARRKLLKAFGIVIDYKERDLIDRSAY
111110000000000000000000000000000000000000000000000000000000
00000000000000000000000000
>DM_train2804
NPWWAAFSRVCKDMNLTLEPEIMPAAGDNRYIRAVGVPALGFSPMNRTPVLLHDHDERLHEA
VFLRGVDIYTRLLPALASVPALPSDS
000000000000000000000000000000000000000000000000000000000000
00000000000000000000000111
>DM_train2806
MPICKEEKQKVIQEFARFPGDTGSTEVQVALLTLRINRLSEHLKVHKKDHHSHRGLLMMVGQ
RRRLLRYLQREDPERYRALIEKLGI
111000000000000000000000000000000000000000000000000000000000

0000000000000000000000000
>DM_train2807
QLTQKNKSADGLVFNVVTQDMINKSTKPYRGHRFTKENVRILESWFAKNIENPYLDTKGLEN
LMKNTSLSRIQIKNWVSNRRRKEKT
111111111100000000000000000000000000000000000000000000000000
0000000000000000000000000
>DM_train2808
GTGAESTAERSARFERDALEFLDQMYSAALRMTRNPADAEDLVQETYAKAYASFHQFREGTN
LKAWLYRILTNTFINSYRKKQREPQ
111110000000000000000000000000000000000000000000000000000000
0000000000000000001111111
>DM_train2809
VKSKAKKTVDKHSDEYKIRRERNNIAVRKSRDKAKMRNLETQHKVLELTAENERLQKKVEQL
SRELSTLRNLFKQLPEPLLASSGHC
111111111000000000000000000000000000000000000000000000000000
0000000000000111111111111
>DM_train2810
MPYTVRFTTTARRDLHKLPPRILAAVVEFAFGDLSREPLRVGKPLRRELAGTFSARRGTYRL
LYRIDDEHTTVVILRVDHRADIYRR
100000000000000000000000000000000000000000000000000000000000
0000000000000000000111111
>DM_train2811
GAMEPQVTLNVTFKNEIQSFLVSDPENTTWADIEAMVKVSFDLNTIQIKYLDEENEEVSINS
QGEYEEALKMAVKQGNQLQMQVHEG
100000000000000000000000000000000000000000000000000000000000
0000000000000000000000000
>DM_train2812
SNAMKVTDVRLRKIQTDGRMKALVSITLDEAFVIHDLRVIEGNSGLFVAMPSKRTPDGEFRD
IAHPINSDMRQEIQDAVMKVYDETD
110000000000000000000000000000000000000000000000000000000000
0000000000000000000000000
>DM_train2813
MSLDCRERIEKDLELLEKNLMEMKSIKLSDDEEAVVERALNYRDDSVYYLEKGDHITSFGCI
TYAHGLLDSLRMLHRIIEGHHHHHH
111100000000000000000000000000000000000000000000000000000000
0000000000000000001111111
>DM_train2814
AADAPAQLDPAGEKLYRSACVVCHASGVANAPKLGDKQAWAPFLAQGADALLATVLKGKGAM
PPRGGTAADEATLRAAVAYMMDAAR
111111000000000000000000000000000000000000000000000000000000
0000000000000000000000000
>DM_train2815
ADGKAIFQQKGCGSCHQANVDTVGPSLKKIAQAYAGKEDQLIKFLKGEAPAIVDPAKEAIMK
PQLTMLKGLSDAELKALADFILSHK

00000000000000000000000000000000000000000000000000000000000000
0000000000000000000000001
>DM_train2817
ASCFALISGTANQVKCYRFRVKKNHRHRYENCTTTWFTVADNGAERQGQAQILITFGSPSQR
QDFLKHVPLPPGMNISGFTASLDF
00000000000000000000000000000000000001111111111110000000000000
000000000000000000000000
>DM_train2818
MSRVQMDTEEVREFVGHLERFKELLREEVNSLSNHFHNLESWRDARRDKFSEVLDNLKSTFN
EFDEAAQEQIAWLKERIRVLEEDY
11110000000000000000000000000000000000000000000000000000000000
000000000000000000000000
>DM_train2819
MVRVPLTAEELERGQRLGELLRSARGDMSMVTVAFDAGISVETLRKIETGRIATPAFFTIAA
VARVLDLSLDDVAAVVTFGPVSTS
11110000000000000000000000000000000000000000000000000000000000
000000000000000000000011
>DM_train2820
SNAMRRNEDSWLIDGATPLEDVMRALNIHTFPRDENYETIGGFMMYMLRKIPKKTDFVLYDK
YKFEIIDTENFRIDQLMVSFRKDV
11111111000000000000000000000000000000000000000000000000000000
000000000000000000000001
>DM_train2821
MVRVDQNLFNEVMYLLDELSQDITVPKNVRKVAQDSKAKLSQENESLDLRCATVLSMLDEMA
NDPNVPAHGRTDLYTIISKLEALS
10000000000000000000000000000000000000000000000000000000000000
000000000000000000000000
>DM_train2822
GAGQSSPATGSQNQSGNTGSIINNYYMQQYQNSMDTQLGNDAISGGSNEGSTDTTSTHTTNT
QNNDWFSKLASSAFSGLFGALLA
11111111111111000000000000000000000000011111111111111111111111
11000000000000000000000
>DM_train2823
PQYQTWEEFSRAAEKLYLADPMKARVVLKYRHSDGNLCVKVTDDLVCLVYKTDQAQDVKKIE
KFHSQLMRLMVAKEARNVTMETE
11100000000000000000000000000000000000000000000000000000000000
00000000000011111111111
>DM_train2824
AEDIQAKIKNYQTAPFDSRFPNQNQTRNCWQNYLDFHRCEKAMTAKGGDVSVCEWYRRVYKS
LCPISWVSTWDDRRAEGTFPGKI
11111111110000000000000000000000000000000000000000000000000000
00000000000000000000000
>DM_train2825
SGGSGAGGGELFATHCAGCHPQGGNTVHPEKTLARARREANGIRTVRDVAAYIRNPGPGMPA

FGEAMIPPADALKIGEYVVASFP
1111110000000000000000000000000000000000000000000000000000000000
00000000000000000000000
>DM_train2826
VKTPGTRYVTHKQLDEKLKNFVTKTEFKEFQTVVMESFAVQNQNIDAQGEQIKELQVEQKAQ
GKTLQLILEALQGINKRLDNLES
1111100000000000000000000000000000000000000000000000000000000000
00000000000000000000000
>DM_train2827
ASAAKGDHGGTGARTWRFLTFGLALPSVALCTLNSWLHSGHRERPAFIPYHHLRIRTKPFSW
GDGNHTFFHNPRVNPLPTGYEKP
0000000000000000000000000000000000000000000000000000000000000000
00000000000000000000001
>DM_train2828
MGKLPESPKRAEEIWQQSVIGDYLARFKNDRVKALKAMEMTWNNMEKKEKLMWIKKAAEDQK
RYERELSEMRAPPAATNSSKKLE
1110000000000000000000000000000000000000000000000000000000000000
00000001111111111111111
>DM_train2829
GSHMSTGDFLTKGIELVQKAIDLDTATQYEEAYTAYYNGLDYLMLALKYEKNPKSKDLIRAK
FTEYLNRAEQLKKHLESEEANAA
1000000000000000000000000000000000000000000000000000000000000000
00000000000000000000011
>DM_train2830
ALLITKKCINCDMCEPECPNEAISMGDHIYEINSDKCTECVGHYETPTCQKVCPIPNTIVKD
PAHVETEEQLWDKFVLMHHADKI
0000000000000000000000000000000000000000000000000000000000000000
00000000000000000011111
>DM_train2831
IRVNVLYSTPACYLWELNKANLSWSVKKDDFFPYADGPYMFWTGYFSSRPALKRYERLSYNF
LQVCNQLEALAGPAANVGPYGS
0000000000000000000000000000000000000000000000000000000000000000
0000000000000111111111
>DM_train2832
MTNLSDIIEKETGKQLVIQESILMLPEEVEEVIGNKPESDILVHTAYDESTDENVMLLTSDA
PEYKPWALVIQDSNGENKIKML
1111111111111100000000000000000000000000000000000000000000000000
0000000000000000000000
>DM_train2833
KPEFDPILLRPVDDLELTVRSANCLKAEAIHYIGDLVQRTEVELLKTPNLGKKSLTEIKDVL
ASRGLSLGMRLENWPPASIADE
1111000000000000000000000000000000000000000000000000000000000000
0000000011111111111111
>DM_train2834

SWDVIKHPHVTEKAMNDMDFQNKLQFAVDDRASKGEVADAVEEQYDVTVEQVNTQNTMDGEK
KAVVRLSEDDDAQEVASRIGVF
000000000000000000000000000000000000000000000000000000000000
0000000000000000111111
>DM_train2835
CEVDALKGTNESLERQMREMEENFAVEAANYQDTIGRLQDEIQNMKEEMARHLREYQDLLNV
KMALDIEIATYRKLLEGEESRI
111111111000000000000000000000000000000000000000000000000000
0000000000000000011111
>DM_train2837
MTKTLRTPEHVYLCQRLRQARLDAGLTQADLAERLDKPQSFVAKVETRERRLDVIEFAKWMA
ACEGLDVVSEIVATIAEGRAQA
111100000000000000000000000000000000000000000000000000000000
0000000000000000000011
>DM_train2838
MYKVKVYVSLKESVLDPQGSAVQHALHSMTYNEVQDVRIGKYMELTIEKSDRDLDVLVKEMC
EKLLANTVIEDYRYEVEEVVAQ
000000000000000000000000000000000000000000000000000000000000
0000000000000000001111
>DM_train2839
MSAFPVHAAFEKDFLVQLVVVDLNDSMDQVAEKVAYHCVNRRVAPREGVMRVRKHRSTELFP
RDMTIAESGLNPTEVIDVVFEE
100000000000000000000000000000000000000000000000000000000000
0000000000000000000000
>DM_train2841
MSYEKEFLKDFEDWVKTQIQVNQLAMATSQEVAQEDGDERAKDAFIRYESKLDAYEFLLGKF
DNYKNGKAFHDIPDELFGARHY
000000000000000000000000000000000111100000000000000000000000
0000000000000001111111
>DM_train2842
MKLIWSEESWDDYLYWQETDKRIVKKINELIKDTRRTPFEGKGKPEPLKHNLSGFWSRRITE
EHRLVYAVTDDSLLIAACRYHY
000000000000000000000000000000000000000000000000000000000000
0000000000000000000001
>DM_train2843
GAMATDWLGSIVSINCGDSLGVYQGRVSAVDQVSQTISLTRPFHNGVKCLVPEVTFRAGDIT
ELKILEIPGPGDNQHFGDLHQT
111100000000000000000000000000000000000000000000000000000000
0000000000000011111111
>DM_train2844
MRGSHHHHHHGSDETGKELVLALYDYQEKSPREVTMKKGDILTLLNSTNKDWWKVEVNDRQG
FVPAAYVKKLAAAWSHPQFEK
111111111111111100000000000000000000000000000000000000000000
000000000000000000000

>DM_train2846
MSAEGYQYRALYDYKKEREEDIDLHLGDILTVNKGSLVALGFSDGQEARPEEIGWLNGYNET
TGERGDFPGTYVEYIGRKKIS
1000000000000000000000000000000000000000000000000000000000000000
000000000000000000001
>DM_train2847
PLTASMLASAPPQEQKQMLGERLFPLIQAMHPTLAGKITGMLLEIDNSELLHMLESPESLRS
KVDEAVAVLQAHQAKEAAQKA
1111111110000000000000000000000000000000000000000000000000000000
000000000000000001111
>DM_train2848
PESSENKNLGGVTQESFDLMIKENPSSQYWKEVAEKRRKALYEALKENEKLHKEIEQKDNEI
ARLKKENKELAEVAEHVQYMA
1111111111111111111111000000000000000000000000000000000000000000
000000000000000000000
>DM_train2849
SCELYHYQECVRGTTVILKEPCPSGTYEGNSPFHPLADNKFALTCTSTHFAFACADGTRHTY
QLRARSVSPKLFIRQEEVQQE
1000000000000000000000000000000000000000000000000000000000000000
000000011111111111111
>DM_train2850
MGSDKIHHHHHHMKMRSQLLIVLQEHLRNSGLTQFKAAELLGVTQPRVSDLMRGKIDLFSLE
SLIDMITSIGLKVEINIKDAA
1111100000000000000000000000000000000000000000000000000000000000
000000000000000000011
>DM_train2851
MGSLPCDICKDVVTAAGDMLKDNATEEEILVYLEKTCDWLPKPNMSASCKEIVDSYLPVILD
IIKGEMSRPGEVCSALNLCES
1000000000000000000000000000000000000000000000000000000000000000
000000000000000000000
>DM_train2852
SNAIRSIWENNGFGLMSSKTMTDFDYWISDFEKIGASQKEAEQLIVKAIEIAIDANARNYNY
INAILKDWEQRGFKSVEEREA
0000000000000000000000000000000000000000000000000000000000000000
000000000000000111111
>DM_train2853
MQRFNPLSKLKRALMDAFVKIDSASHMIVLKTMPGNAQAIGALMDNLDWDEMMGTICGDDTI
LIICRTPEDTEGVKNRLLELL
1111000000000000000000000000000000000000000000000000000000000000
000000000000000000000
>DM_train2854
DSTTKELIKKLAEINKCENEISAKYCDHMIHEEIPLKTCTKEKTRNLCCAVSDYCMSYFTYD
SEEYYDCTKREFDDPSYTCFR
1000000000000000000000000000000111000000000000000000000000000000

000000000000000000000
>DM_train2855
MASAIVLINTDAGGEDEVFERLKSMSEVTEVHVVYGVYDIVVKVEADSMDKLKDFVTNTIRK
LPKVRSTLTMIIVEGKSLVKK
100000000000000000000000000000000000000000000000000000000000
000000000000000000001
>DM_train2857
QSRGEKRTAHNAIEKRYRSSINDKIIELKDLVVGTEAKLNKSAVLRKAIDYIRFLQHSNQKL
KQENLSLRTAVHKSKSLKDL
000000000000000000000000000000000000000000000000000000000000
00000000000000111111
>DM_train2858
GRTNLIVNYLPQNMTQDELRSLFSSIGEVESAKLIRDKVAGHSLGYGFVNYVTAKDAERAIN
TLNGLRLQSKTIKVSYARPS
000000000000000000000000000000000001111110000000000000000000
00000000000000000001
>DM_train2859
KTPLVNESLKKFLNTKDGRLVASLVAEFLQFFNLDFTLAVFQPETSTLQGLEGRENLARDLG
IIEAEGTVGGPLLLEVIRRW
111100000000000000000000000000000000000000000111000000000000
00000000000000000000
>DM_train2860
SSSGKPGPTVIKVQNMPFTVSIDEILDFFYGYQVIPGSVCLKYNEKGMPTGEAMVAFESRDE
ATAAVIDLNDRPIGSRKVKL
111111110000000000000000000000000000000000000000000000000000
00000000000000000000
>DM_train2861
STSLPTSNEYQNEKLANELKSLLDELNVNELATGSLNTYYKRTIKISGQKAMYALKSKDFKK
MSEAKYQLQKIYNEIDEALK
111111110000000000000000000000000000000000000000000000000000
00000000000000000000
>DM_train2862
KGMTPPKTVNFKMKGVADAAFSHEFHLGMYKCNECHTKLFAYKAGAKRFTMADMDKGKSCGA
CHNGKDAFSSASDCGKCHPG
000000000000000000000000000000000000000000000000000000000000
00000000000000000001
>DM_train2863
GPLGSPEFGYWITCCPTCDVDINTWVPFYSTELNKPAMIYCSHGDGHWVHAQCMDLEERTLI
HLSEGSNKYYCNEHVQIARA
111111110000000000000000000000000000000000000000000000000000
00000000000000000000
>DM_train2864
MFPAQEEADRTVFVGNLEARVREEILYELFLQAGPLTKVTICKDREGKPKSFGFVCFKHPES
VSYAIALLNGIRLYGRPINV

111100000000000000000000000000000000000000000000000000000000
00000000000000000000
>DM_train2865
REVKEVDLRGLTVAEALLEVDQALEEARALGLSTLRLLHGKGTGALRQAIREALRRDKRVES
FADAPPGEGGHGVTVVALRP
110000000000000000000000000000000000000000000000000000000000
00000000000000000000
>DM_train2866
TGTSEMAPALVAAFGGKENITNLDACITRLRVSVADVSKVDQAGLKKLGAAGVVVAGSGVQA
IFGTKSDNLKTEMDEYIRNH
000000000000000000000000000000000000000000000000000000000000
00000000000000000001
>DM_train2867
ELITVHIVKGPMGFGFTIADSPGGGGQRVKQIVDSPRSRGLKEGDLIVEVNKKNVQALTHNQ
VVDMLVESPKGSEVTLLVQR
000000000000000000000000000000000110000000000000000000000000
00000000000000000000
>DM_train2869
GSHMLSIYQDQEQRILKFLEELGEGKATTAHDLSGKLGTPKKEINRVLYSLAKKGKLQKEAG
TPPLWKIAVSTQAWNQHSG
111111100000000000000000000000000000000000000000000000000000
0000000011111111111
>DM_train2870
MNREGAPGKSPEEMYIQQKVRVLLMLRKMGSNLTASEEEFLRTYAGVVNSQLSQLPPHSIDQ
GAEDVVMAFSRSETEDRRQ
111111110000000000000000000000000000000000000000000011111000
0000000000000000111
>DM_train2871
GRQFGHLTRVRHVITYSLSPFEQRAFPHYFSKGIPNVLRRTRACILRVAPPFVAFYLVYTWG
TQEFEKSKRKNPAAYENDR
000000000000000000000000000000000000000000000000000000000000
0000000000000111111
>DM_train2872
PSKTSNTIRVFLPNKQRTVVNVRNGMSLHDCLMKALKVRGLQPECCAVFRLLHEHKGKKARL
DWNTDAASLIGEELQVDFL
111110000000000000000000000000000000000000000000000000000000
0000000000000000000
>DM_train2873
MRITIYTRNDCVQCHATKRAMENRGFDFEMINVDRVPEAAEALRAQGFRQLPVVIAGDLSWS
GFRPDMINRLHPAPHAASA
000000000000000000000000000000000000000000000000000000000000
0000000000000011111
>DM_train2874
TTPIVHLKGDANTLKCLRYRFKKHCTLYTAVSSTWHWTGHNVKHKSAIVTLTYDSEWQRDQF

LSQVKIPKTITVSTGFMSI
0000000000000000000000000000000000000111111100000000000000000
0000000000000000000
>DM_train2875
MDAIKKKMQMLKLDKENALDRAEQAEADKKAAEERSKQLEDELVALQKKLKGTEDELDKYSE
SLKDAQEKLELADKKATDC
0000000000000000000000000000000000000000000000000000000000000
0000000000000001111
>DM_train2876
GAMAEAEAEVTLRELQEALEEEVLTRQSLSREMEAIRTDNQNFASQLREAEARNRDLEAHVR
QLQERMELLQAEGATAVTG
1111111100000000000000000000000000000000000000000000000000000
0000000001111111111
>DM_train2877
MEKTRVNGTDRPPISSWSVDDVSNFIRELPGCQDYVDDFIQQEIDGQALLLLKEKHLVNAMG
MKLGPARKIVAKVESIKEV
1111111111110000000000000000000000000000000000000000000000000
0000000000000000111
>DM_train2878
GSMDYKDDDDKKTNWLKRIYRVRPCVKCKVAPRDWKVKNKHLRIYNMCKTCFNNSIDIGDDT
YHGHVDWLMYADSKEISNT
1111110000000000000000000000000000000000000000000000000000000
0000000000000111111
>DM_train2879
SRSTHNEMEKNRRAHLRLSLEKLKGLVPLGPDSSRHTTLSLLTKAKLHIKKLEDSDRKAVHQ
IDQLQREQRHLKRQLEKL
0000000000000000000000000000000000000000000000000000000000000
000000000000000111
>DM_train2880
DSQWGSGKNLYDKVCGHCHKPEVGVGPVLEGRGLPEAYIKDIVRNGFRAMPAFPASYVDDES
LTQVAEYLSSLPAPAAQP
0000000000000000000000000000000000000000000000000000000000000
000000000000111111
>DM_train2881
GSYHADPSLVSFLTGLGCPNCIEYFTSQGLQSIYHLQNLTIEDLGALKIPEQYRMTIWRGLQ
DLKQGHDYSTAQQLLRSS
1111111000000000000000000000000000000000000000000000000000000
001111111111111111
>DM_train2882
KERPEISLPSDFEHTIHVGFDAVTGEFTGMPEQWARLLQTSNITKSEQKKNPQAVLDVLEFY
NSKKTSNSQKYMSFTDKS
1111111100000000000000000000000000000000000000000000000000000
000000000000000000
>DM_train2883

MCLSIPSQVVAVDNERQSVTVDTLGVRRDVSSHLMTEPLAIGDYVLIHIGFVMNKIDRNDAL
QSLELYQEIVSKLENETT
0000000000000000000000000000000000000000000000000000000000000
000000000000001111
>DM_train2884
AIDTYEFASDAERERFRNLTQELRCPKCQNQDIADSNAPIAADLRKQIYGQLQQGKSDGEIV
DYMVARYGDFVRYKPPVN
0000000000000000000000000000000000000000000000000000000000000
000000000000000011
>DM_train2885
AVYGIDAMNPSSRDDFTEFGKLLKDKITQYEKSLYYASFLEVLVRDVCISLEIDDLKKITNS
LTVLCSEKQKQEKQSKAK
1110000000000000000000000000000000000000000000000000000000000
000000000011111111
>DM_train2886
MEPVSKWSPSQVVDWMKGLDDCLQQYIKNFEREKISGDQLLRITHQELEDLGVSRIGHQELI
LEAVDLLCALNYGLETEN
1000000000000000000000000000000000000000000000000000000000000
000000000000001111
>DM_train2887
RPAIRKRLLKPKVLDSSPRALVGHRAEVLEDVGATSGQVRLDGSIWSARSMDPTHTFAEGEI
VSVIDIQGTTAIVWKEA
1111111111111111000000000000000000000000000000000000000000000
00000000000000000
>DM_train2888
PTVVDQIRLWQLELDRVITYEGSLYSDFETSQEYNLLSKYAQDIGVLLWKDDKKKKFFISKE
GNSQVLDFAKRKLKKKQ
1111111111110000000000000000000000000000000000000000000000000
00000000000011111
>DM_train2889
MITIDALGQVCPIPVIRAKKALAELGEAGGVVTVLVDNDISRQNLQKMAEGMGYQSEYLEKD
NGVIEVTIVAGEGCAVE
0000000000000000000000000000000000000000000000000000000000000
00000000000011111
>DM_train2890
TDFLAGIRIVGEDKNGMTNQITGVISKFDTNIRTIVLNAKDGIFTCNLMIFVKNTDKLTTLM
DKLRKVQGVFTVERLSN
0000000000000000000000000000000000000000000000000000000000000
00000000000000011
>DM_train2891
IPLEQGIEFLSVNVEEDSPVVGKKLKDLPLPRDSIIAAIVRGGVLVVPRGDTEILSGDKLYV
IVSAEAKETVEETLLGR
0000000000000000000000000000000000000000000000000000000000000
00000000000000011

>DM_train2892
MKIPIEELEDRVFVNCNTSITWVEGTVGTLLSDITRLDLGKRILDPRGIYRCNGTDIYKDKE
STVQVHYRMCQSCVELD
100000000000000000000000000000000000000000000000000111111110
00000000001111111
>DM_train2893
MLQCYNCPNPTADCKTAVNCSSDFDACLITKAGLQVYNKCWKFEHCNFNDVTTRLRENELTY
YCCKKDLCNFNEQLENG
000000000000000000000000000000000000000000000000000000000000
00000000000000001
>DM_train2894
KVTLVKSRKNEEYGLRLASHIFVKEISQDSLAARDGNIQEGDVVLKINGTVTENMSLTDAKT
LIERSKGKLKMVVQRDE
000000000000000000000000000000000000000000000000000000000000
00000000000000011
>DM_train2895
MNAIDIAINKLGSVSALAASLGVRQSAISNWRARGRVPAERCIDIERVTNGAVICRELRPDV
FGASPAGHRPEASNAAA
000000000000000000000000000000000000000000000000000000000000
01111111111111111
>DM_train2896
MTVATDNNPTPEAVADLKKKVRKLNSKAGQMKMDLHDLAEGLPTDYENLVETAEKTYEIFRE
LDQLKKKLNIWEETLK
111111111111110000000000000000000000000000000000000000000000
0000000000000111
>DM_train2897
MHEQLSPRDQELEARLVELETRLSFQEQALTELSEALADARLTGARNAELIRHLLEDLGKVR
STLFADAADEPPPPHY
111111100000000000000000000000000000000000000000000000001111
1111111111111111
>DM_train2898
MEEAKQKVVDFLNSKSGSKSKFYFNDFTDLFPDMKQREVKKILTALVNDEVLEYWSSGSTTM
YGLKGAGKQAAAEHED
000000000000000000000000000000000000000000000000000000000000
0000000000001111
>DM_train2899
GHMLIPHDLLEADTLNNLLEDFVTREGTDNGDETPLDVRVERARHALRRGEAVILFDPESQQ
CQLMLRSEVPAELLRD
000000000000000000000000111111100000000000000000000000000000
0000000000000000
>DM_train2900
GSMWIQVRTMDGRQTHTVDSLSRLTKVEELRRKIQELFHVEPGLQRLFYRGKQMEDGHTLFD
YEVRLNDTIQLLVRQS
000000000000000000000000000000000000000000000000000000000000

0000000000001111
>DM_train2901
MKTDGEFVSEHLMELITQQNLTINRVATLAGLNQSTVNAMFEGRSKRPTITTIRKVCGTLGI
SVHDFFDFPPYNEVEK
00000000000000000000000000000000000000000000000000000000000000
0000000000000011
>DM_train2902
MLMVVPLSEMGPGDKGIVVNILGGHNARQKLVSMGLTPGATIQVLESHPMGPIIISVGGVRF
AIGKGLAGRVMVRKL
10000000000000000000001110000000000000000000000000000000000000
000000000000000
>DM_train2903
MKYCKFCCKAVTGVKLIHVPKCAIKRKLWEQSLGCSLGENSQICDTHFNDSQWKAAPAKGQT
FKRRRLNADAVPSKV
00000000000000000000000000000000000000000000000000000000110000
000000000000000
>DM_train2904
ALGKLQDVVNQNAQALNTLVKQLSSNFGAISSVLNDISGGRGGDISGINASVVNIQKEIDRL
NEVAKNLNESLIDLQ
00000000000000000000000000000000000000000000000000000000000000
000000000000011
>DM_train2905
YTNSDFVVIKALEDGVNVIGLTRGADTRFHHSEKLDKGEVLIAQFTEHTSAIKVRGKAYIQT
RHGVIESEGKKAAAM
11100000000000000000000000000000000000000000000000000000000000
000011111111111
>DM_train2906
QAKEPCVESLVSQYFQTVTDYGKDLMEKVKSPELQAEAKSYFEKSKEQLTPLIKKAGTELVN
FLSYFVELGTQPATQ
11100011000000100000000000000000000000000000000000000000000000
000000000000001
>DM_train2907
AMDPEFMGREVENLILENTQLLETKNALNIVKNDLIAKVDELTCEKDVLQGELEAVKQAKLK
LEEKNRELEEELRKA
11111000000000000000000000000000000000000000000000000000000000
000001111111111
>DM_train2908
MPMISCDMRYGRTDEQKRALSAGLLRVISEATGEPRENIFFVIREGSGINFVQHGEHLPDYV
PGNANDKALIAKLK
10000000000000000000000000000000000000000000000000000000000000
01111111111111
>DM_train2909
RFYVKDHRNKAMINLHIQKDNPKIVHAFDMEDLGDKAVYCRCWRSKKFPFCDGAHTKHNEET
GDNVGPLIIKKKET

000000000000000000000000000000000000000000000000000000000000000
00000000001111
>DM_train2910
MKLYFNGGKKKKIRAVDFVGTIAKIDGVSADDIGIITIMDNASYVEILNGKGPHVLKVMKNT
TVKGKQLKVNKANK
000000000011110000000000000000000000000000000000000000000000000
00111000000001
>DM_train2911
MQPDALNVPSTCCFTFSSKKISLQRLKSYVITTSRCPQKAVIFRTKLGKEICADPKEKWVQN
YMKHLGRKAHTLKT
111000000000000000000000000000000000000000000000000000000000000
00000011111111
>DM_train2912
MEKKALTLAERQRLIVEGLPHVSATLARRLLKHFGSVERVFTASVAELMKVEGIGEKIAKEI
RRVITAPYIEDEE
111100000000000000000000000000000000000000000000000000000000000
0000000000111
>DM_train2913
NECVSKGFGCLPQSDCPQEARLSYGGCSTVCCDLSKLTGCKGKGGECNPLDRQCKELQAESA
SCGKGQKCCVWLH
000000000000000000000000000000000000000000000000000000000000000
0000000000001
>DM_train2914
MRVEALGKVAPLPQAQTPVSLNEASLEELMALPGIGPVLARRIVEGRPYARVEDLLKVKGIG
PATLERLRPYLRP
111111111100000000000000000000000000000000000000000000000000000
0000000000000
>DM_train2915
VDQADDAAIQQTLAKMGIKSSDIQPAPVAGMKTVLTNSGVLYITDDGKHIIQGPMYDVSGTA
PVNVTNKMLLKQL
111100000000000000000000000000000000000000000000000000000000000
0000000000001
>DM_train2916
GKSNGEKKYIVGFKQGFKSCAKKEDVISEKGGKLQKCFKYVDAASATLNEKAVEELKKDPSV
AYVEEDKLYRAL
111110000000000000000000000000000000000000000000000000000000000
000000000000
>DM_train2917
KVFVGRCTEDMTAEELQQFFCQYGEVVDVFIPKPFRAFAFVTFADDKVAQSLCGEDLIIKGI
SVHISNAEPKHN
000000000000000000000000000000000000000000000000000000000000000
000000001111
>DM_train2918
VDMSNVVKTYDLQDGSKVHVFKDGKMGMENKFGKSMNMPEGKVMETRDGTKIIMKGNEIFRL

DEALRKGHSEGG
000000000000000000000000000000000000000000000000000000000000
000011111111
>DM_train2919
YDGTKCKAAGNCWEPKPGFPEKIAGSKYDPKHDPKELNKQADSIKQMEERNKKRVENFKKTG
KFEYDVAKISAN
000000000000000000000000000000000000000000000000000000000000
000000000001
>DM_train2920
MQSHHDHYADLVKFGQRLRELRTAKGLSQETLAFLSGLDRSYVGGVERGQRNVSLVNILKLA
TALDIEPRELFC
111111100000000000000000000000000000000000000000000000000000
000000000001
>DM_train2921
MSELFSVPYFIENLKQHIEMNQSEDKIHAMNSYYRSVVSTLVQDQLTKNAVVLKRIQHLDEA
YNKVKRGESKLE
110000000000000000000000000000000000000000000000000000000000
000000011111
>DM_train2922
AVQQNKELNFKLREKQNEIFELKKIAETLRSKLEKYVDITKKLEDQNLNLQIKISDLEKKLS
DANSTFKEMRFP
110000000000000000000000000000000000000000000000000000000000
001111111111
>DM_train2923
QELRLRVQGKEKHQMLEISLSPDSPLKVLMSHYEEAMGLSGHKLSFFFDGTKLSGKELPADL
GLESGDLIEVWG
110000000000000000000000000000000000000000000000000000000000
000000000000
>DM_train2924
MSDVLIRDIPDDVLASLDAIAARLGLSRTEYIRRRLAQDAQTARVTVTAADLRRLRGAVAGL
GDPELMRQAWR
100000000000000000000000000000000000000000000000000000000000
00000000000
>DM_train2925
MAKKDGAIEVEGRVVEPLPNAMFRIELENGHKVLAHISGKMRQHYIRILPEDRVVVELSPYD
LSRGRIVYRYK
111111000000000000000000000000000000000000000000000000000000
00000000000
>DM_train2926
KNVTLTIRLLMHGKEVGSIIGKKGESVKKMREESGARINISEGNCPERIITLAGPTNAIFKA
FAMIIDKLEED
110000000000000000000000000000000000000000000000000000000000
00000000000
>DM_train2927

GPLGSDLKDAEAVQKFFLEEIQLGEELLAQGDYEKGVDHLTNAIAVCGQPQQLLQVLQQTLP
PPVFQMLLTKL
111110000000000000000000000000000000000000000000000000000000
00000000000
>DM_train2928
GSGEGSEDDPLYDEAVRFVTESRRASISAVQRKLKIGYNRAARMIEAMEMAGVVTPMNTNGS
REVIAPAPVRD
111111110000000000000000000000000000000000000000000000000000
00000001111
>DM_train2929
MSNNQQMMLNRIKVVLAEKQRTNRWLAEQMGKSENTISRWCSNKSQPSLDMLVKVAELLNVD
PRQLINGKIKI
111111000000000000000000000000000000000000000000000000000000
00000011111
>DM_train2930
NKYDWFEFFLNCGVDVSNCQRYTINFDREQLTEDMMPDINNSMLRTLGLREGDIVRVMKHLD
KKFGRENIAS
100000000000000000000000000000000000000000000000000000000000
0000011111
>DM_train2931
AMATRQEEMIDHRLTDREWAEEWKHLDHLLNCIMDMVEKTRRSLTVLRRCQEADREELNYWI
RRYSDAEDLK
111111111100000000000000000000000000000000000000000000000000
0000000111
>DM_train2932
MKKHHHHHHDNATDDPVDLVYAAEKIIQKRVKKGVVEYRVKWKGWNQRYNTWEPEVNILDRR
LIDIYEQTNK
111111111111111110000000000000000000000000000000000000000000
0000001111
>DM_train2933
MVSGEVRTKKVPLDTNHKRFYDAFAQGAGKLDLDRQCVECHHEKPGGIPFPKNHPVKPADGP
MRCLFCHKFK
111110000000000000000000000000000000000000000000000000000000
0000000000
>DM_train2934
MAGYLNNIALNLEIVLKNKADSPEVSETLVTRICENLLLSKEVSFLKADGSVENFKLSDMEY
EITNTEELPE
100000000000000000000000000000000000000000000000000000000000
0000000001
>DM_train2935
QPRRPALRQCCNQLRQVDRPCVCPVLRQAAQQVLQRQIIQGPQQLRRLFDAARNLPNICNIP
NIGACPFRAW
111100000000000000000000000000000000000000000000000000000000
0000000001

>DM_train2936
ARARKGALVQCDPSIKALILQIDAKMSDIVLEELDDTHLLVNPSKVEFVKHELNRLLSKNIY
NPMDEEENQ
000000000000000000000000000000000000000000000000000000000000
011111111
>DM_train2937
LKYMVPGARVTRGLDWKWRDQDGSPQGEGTVTGELHNGWIDVTWDAGGSNSYRMGAEGKFDL
KLAPGYDPD
000000000000000000000000000000000000000000000000000000000000
000000001
>DM_train2938
AVCPTGLFSNPLCCATNVLDLIGVDCKTPTIAVDTGAIFQAHCASKGSKPLCCVAPVADQAL
LCQKAIGTF
000000000000000000000000000000000000000000000000000000000000
000000001
>DM_train2939
MNLTVNGKPSTVDGAESLNVTELLSALKVAQAEYVTVELNGEVLEREAFDATTVKDGDAVEF
LYFMGGGK
000000000000000000000000000000000000000000000000000000000000
00000001
>DM_train2940
PFIECHIATGLSVARKQQLIRDVIDVTNKSIGSDPKIINVLLVEHAEANMSISGRIHGEAAS
TERTPAVS
000000000000000000000000000000000000000000000000000000001111
11111111
>DM_train2941
MSRFDSGLPSVRQVQLLIKDQTPVEIKLLTGDSLFGTIRWQDTDGLGLVDDSERSTIVRLAA
IAYITPRR
111100000000000000000000000000000000000000000000000000000000
00000001
>DM_train2942
AHAAIDTFTKYLDIDEDFATVLVEEGFSTLEELAYVPMKELLEIEGLDEPTVEALRERAKNA
LATIAQAQ
000000000000000000000000000000000000000000000000000000000000
00000011
>DM_train2943
GPETLCGAELVDALQFVCGDRGFYFNKPTGYGSSSRRAPQTGIVDECCFRSCDLRRLEMYCA
PLKPAKSA
100000000000000000000000000000000000000000000000000000000000
01111111
>DM_train2944
EAEEDGDLQCLCVKTTSQVRPRHITSLEVIKAGPHCAVPQLIATLKNGRKICLDLQAPLYKK
IIKKLLES
111111100000000000000000000000000000000000000000000000000000

00000000
>DM_train2945
RTDRLEVCREYQRGNCNRGENDCRFAHPADSTMIDTNDNTVTVCMDYIKGRCSREKCKYFHP
PAHLQAK
11000000000000000000000000000000000000000000000000000000000000
0000001
>DM_train2946
TEDVSDLPVEEQLRRLQEERTCKVCMDKEVSIVFIPCGHLVVCKDCAPSLRKCPICRSTIKG
TVRTFLS
11110000000000000000000000000000000000000000000000000000000000
0000000
>DM_train2947
GDSVEKIVLELSGLSCHHCVARVKKALEEAGAKVEKVDLNEAVVAGNKEDVDKYIKAVEAAG
YQAKLRS
10000000000000000000000000000000000000000000000000000000000000
0000000
>DM_train2948
HLNVLAKALYDNVAESPDELSFRKGDIMTVLEQDTQGLDGWWLCSLHGRQGIVPGNRLKILV
GMYDKKP
10000000000000000000000000000000000000000000000000000000000000
0000000
>DM_train2949
GPLGSAEGWLPLSGGQGQSEDSDPLLQQIHNITSFIRQAKAAGRMDEVRTLQENLRQLQDEY
DQQQTEK
11111100000000000000000000000000000000000000000000000000000000
0000011
>DM_train2950
MEKDLLDKLGQHLVWRMGRAEDEDVLVVRVGLASATPRFRELPRLLNLPEAEMRRLVQEGRV
RVEWVEE
11111111111000000000000000000000000000000000000000000000000000
0000000
>DM_train2951
GSHMPDWRQFCELHAQAAAVDFAHKFCRFLRDNPAYDTPDAGASFSRHFAANFLDVFGEEVR
RVLVAGP
11110000000000000000000000000000000000000000000000000000000000
0000111
>DM_train2952
ASNFDCCLGYTDRILHPKFIVGFTRQLANEGCDINAIIFHTKKKLSVCANPKQTWVKYIVRL
LSKKVKN
11110000000000000000000000000000000000000000000000000000000000
0000000
>DM_train2953
MEQKTLQVEGMSCQHCVKAVETSVGELDGVSAVHVNLEAGKVDVSFDADKVSVKDIADAIED
QGYDVAK

00000000000000000000000000000000000000000000000000000000000000
0000001
>DM_train2954
MSEPEVPFKVVAQFPYKSDYEDDLNFEKDQEIIVTSVEDAEWYFGEYQDSNGDVIEGIFPKS
FVAVQG
11000000000000000000000000000000000000000000000000000000000000
000000
>DM_train2955
MVFKKVLLTGTSEESFTAAADDAIDRAEDTLDNVVWAEVVDQGVEIGAVEERTYQTEVQVAF
ELDGSQ
10000000000000000000000000000000000000000000000000000000000000
000111
>DM_train2957
GSKLQEVEIRNKDLEGQLSEMEQRLEKSQSEQDAFRSNLKTLLEILDGKIFELTELRDNLAK
LLECS
11100000000000000000000000000000000000000000000000000000000000
00111
>DM_train2958
GSHMLSTGDNLEQKILQVLSDDGGPVAIFQLVKKCQVPKKTLNQVLYRLKKEDRVSSPSPKY
WSIGG
11111111000000000000000000000000000000000000000000000000000000
00001
>DM_train2959
SNAMMTKKERIAIQRSMAEEALGKLKAIRQLCGAEDSSDSSDMQEVEIWTNRIKELEDWLWG
ESPIA
11111000000000000000000000000000000111110000000000000000000000
00000
>DM_train2960
MKKIPLSKYLEEHGTQSALAAALGVNQSAISQMVRAGRSIEITLYEDGRVEANEIRPIPARP
KRTAA
00000000000000000000000000000000000000000000000000000000000111
11111
>DM_train2961
ADPGAEPEASIDDLDAEALIRMALGPRNTMTSSNEQLVDALRASLKENEELRKESRRRADRR
QEPM
11111111100000000000000000000000000000000000000000000000000000
0000
>DM_train2962
MVKVKFKYKGEEKEVDTSKIKKVWRVGKFVSFTYDDNGKTGRGAVSEKDAPKELLDMLARAE
REKK
10000000000000000000000000000000000000000000000000000000000000
0011
>DM_train2963
MQILFNDQAMQCAAGQTVHELLEQLDQRQAGAALAINQQIVPREQWAQHIVQDGDQILLFQV

IAGG
1000000000000000000000000000000000000000000000000000000000000
0000
>DM_train2964
GPLGSENDLGITAVALYDYQAAGDDEISFDPDDIITNIEMIDDGWWRGVCKGRYGLFPANYV
ELRQ
1111110000000000000000000000000000000000000000000000000000000
0000
>DM_train2965
SFSSLYKTFFKRNAVFVGTIFAGAFVFQTVFDTAITSWYENHNKGKLWKDVKARIAAGDGDD
DDE
0000000000000000000000000000000000000000000000000000000011111
111
>DM_train2966
GLKWTDSREIGEALYDAYPDLDPKTVRFTDMHQWICDLEDFDDDPQASNEKILEAILLVWLD
EAE
0000000000000000000000000000000000000000000000000000000000000
001
>DM_train2967
MNQPIELSLEQQFSIRSFATQVQNMSHDQAKDFLVKLYEQMVVREATYQELLKHQWGLDSGS
TPA
1111000000000000000000000000000000000000000000000000011111111
111
>DM_train2968
MVRDAYIAQNYNCVYHCARDAYCNELCTKNGAKSGSCPYLGEHKFACYCKDLPDNVPIRVPG
KCH
1000000000000000000000000000000000000000000000000000000000000
000
>DM_train2969
GSHMRRRVRAILPYTKVPDTDEISFLKGDMFIVHNELEDGWMWVTNLRTDEQGLIVEDLVEE
VGR
1111000000000000000000000000000000000000000000000000000000000
001
>DM_train2970
TQLEEQLHNVETVRSITMQLEMALTKLKKDMMRGGDAKQYQVWQRESKALESAIAIIHYVAG
DLK
1111111100000000000000000000000000000000000000000000000000000
111
>DM_train2971
GHSSSDPVYPCGICTNEVNDDQDAILCEASCQKWFHRICTGMTETAYGLLTAEASAVWGCDT
CMA
1111111100000000000000000000000000000000000000000000000000000
000
>DM_train2973

PQFSQQREEDIYRFLKDNGPQRALVIAQALGMRTAKDVNRDLYRMKSRHLLDMDEQSKAWTI
YR
000000000000000000000000000000000000000000000000000000000000
01
>DM_train2974
KSSHTLKTANSYTDVTVSNSTKKAIRESNQYTDHKFHQLENRLDKLEKRLLKLLASSAALNS
LF
000000000000000000000000000000000000000000000000000011111111
11
>DM_train2975
GSHMPVPSFGEAMAYFAMVKRYLTSFPIDDRVQSHILHLEHDLVHVTRKNHARQAGVRGLGH
QS
111100000000000000000000000000000000000000000001111111111111
11
>DM_train2976
MVWLNGEPRPLEGKTLKEVLEEMGVELKGVAVLLNEEAFLGLEVPDRPLRDGDVVEVVALMQ
GG
000000000000000000000000000000000000000000000000000000000000
01
>DM_train2977
GSHMEANENILKLKLYRSLGVILDLENDQVLINRKNDGNIDILPLDNNLSDFYKTKYIWERL
GK
111110000000000000000000000000000000000000000000000000000000
00
>DM_train2978
GSGAKSLFSTAFSESLAAEISSVSRDELMEAIQKQEEINFRLQDYIDRIIVAIMETNPSILE
VK
111111111111111111111000000000000000000000000000000000000000
00
>DM_train2979
DAPANPCCDAATCKLTTGSQCADGLCCDQCKFMKEGTVCRRARGDDLDDYCNGISAGCPRNP
FH
111000000000000000000000000000000000000000000000000000000111
11
>DM_train2980
GSMLILTRRVGETLMIGDEVTVTVLGVKGNQVRIGVNAPKEVSVHREEIYQRIQAEKSQPTS
Y
000000000000000000000000000000000000000000000000000000001111
1
>DM_train2981
GSHMKEPKLSPKQERFIEEYFINDMNATKAAIAAGYSKNSASAIGAENLQKPAIRARIDARL
K
111111100000000000000000000000000000000000000000000000000000
0

>DM_train2982
GTKQEKTALNMARFIRSQTLTLLEKLNELDADEQADICESLHDHADELYRSCLARFGDDGEN
L
10000000000000000000000000000000000000000000000000000000001111
1
>DM_train2983
YTDCTESGQNLCLCEGSNVCGQGNKCILGSDGEKNQCVTGEGTPKPQSHNDGDFEEIPEEYL
Q
00000000000000000000000000000000000000000000000011100000000000
0
>DM_train2984
GPLGSSDGAVTLQEYLELKKALATSEAKVQQLMKVNSSLSDELRKLQREIHKLQAENLQLRQ
P
11111100000000000000000000000000000000000000000000000000000000
1
>DM_train2985
GSQEQKGDAPTCGICHKTKFADGCGHNCSYCQTKFCARCGGRVSLRSNKVMWVCNLCRKQQE
11111111000000000000000000000000000000000000000000000000000000
>DM_train2986
GSEFAIDPNEPTYCLCNQVSYGEMIGCDNEQCPIEWFHFSCVSLTYKPKGKWYCPKCRGDNE
11111111100000000000000000000000000000000000000000000000000001
>DM_train2987
VFPSNATFGMGDRVRKKSGAAWQGQIVGWYCTNLTPEGYAVESEAHPGSVQIYPVAALERIN
11110000000000000000000000000000000000000000000000000000000000
>DM_train2988
EMELEIDRNLDQIQQVSNRLKKMALTTGKELDSQQKRLNNIEESTDDLDINLHMNTNRLAGI
00000000000000000000000000000000000000000000000000000000000001
>DM_train2989
MASLLTETMPFRMTMEGTVNGHHFKCTGKGEGNPFEGTQDMKIEVIEGGPLPFAFDILSTSC
11110000000000000000000000000000000000000000000000000000000000
>DM_train2991
GSHMLCAISGKVPRRPVLSPKSRTIFEKSLLEQYVKDTGNDPITNEPLSIEEIVEIVPSAQ
1110000000000000000000000000000000000000000000000000000000111
>DM_train2992
SGEKTVVCKHWLRGLCKKGDQCEFLHEYDMTKMSECYFYSKFGECSNKECPFLHIDPESKI
0000000000000000000000000000000000000000000000000000000001111
>DM_train2993
MGEPTEFEYLRKVLFEYMMGRETKTMAKVITTVLKFPDDQTQKILEREDARLMSWLRSSS
100000000000000000000000000000000000000000000000000011111111
>DM_train2994
MVEMRMKKCPKCGLYTLKEICPKCGEKTVIPKPPKFSLEDRWGKYRRMLKRALKNKNKAE
110000000000000000000000000000000000000000000000000000000111
>DM_train2995
GPLGSPEFAIAKFDYVGRSARELSFKKGASLLLYHRASEDWWEGRHNGIDGLVPHQYIVV

111110000000000000000000000000000000000000000000000000000000
>DM_train2996
MDAKFLEILVCPLCKGPLVFDKSKDELICKGDRLAFPIKDGIPMMLESEARELAPEEEVK
111000000000000000000000000000000000000000000000000000000000
>DM_train2997
QKGQVGIFSFQNNYADSATTFRILAHLDEQRYPLPNGAAEKNLPSLFEGFKATVSIIQQR
111111111111111100000000000000000001111000000000000000000000
>DM_train2998
GMERGIVQYDFMAESQDELTIKSGDKVYILDDKKSKDWWMCQLVDSGKSGLVPAQFIEPV
110000000000000000000000000000000000000000000000000000000000

The IDRs validation dataset.

```
>DM_validata9
CGVGFIANLRGKPDHTLVEQALKALGCMEHRGGCSADNDSGDGAGVMTAIPRELLAQWFNTR
NLPMPDGDRLGVGMVFLPQEPSAREVARAYVEEVVRLEKLTVLGWREVPVNSDVLGIQAKNN
QPHIEQILVTCPEGCAGDELDRRLYIARSIIGKKLAEDFYVCSFSCRTIVYKGMVRSIILGE
FYLDLKNPGYTSNFAVYHRRFSTNTMPKWPLAQPMRLLGHNGEINTLLGNINWMAAREKELE
VSGWTKAELEALTPIVNQANSDSYNLDSALELLVRTGRSPLEAAMILVPEAYKNQPALKDYP
EISDFHDYYSGLQEPWDGPALLVFSDGKIVGAGLDRNGLRPARYCITKDDYIVLGSEAGVVD
LPEVDIVEKGRLAPGQMIAVDLAEQKILKNYQIKQQAAQKYPYGEWIKIQRQTVASDSFAEK
TLFNDAQTVLQQQAAFGYTAEDVEMVVVPMASQGKEPTFCMGDDTPLAVLSHKPRLLYDYFK
QRFAQVTNPPIDPLRENLVMSLAMFLGKRGNLLEPKAESARTIKLRSPLVNEVELQAIKTGQ
LQVAEVSTLYDLDGVNSLEDALTNLVKTAIATVQAGAEILVLTDRPNGAILTENQSFIPPLL
AVGAVHHHLIRAGLRLKASLIVDTAQCWSTHHFACLVGYGASAICPYLALESVRQWWLDEKT
QKLMENGRLDRIDLPTALKNYRQSVEAGLFKILSKMGISLLASYHGAQIFEAIGLGAELVEY
AFAGTTSRVGGLTIADVAGEVMVFHGMAFPEMAKKLENFGFVNYRPGGEYHMNSPEMSKSLH
KAVAAYKVGGNGNNGEAYDHYELYRQYLKDRPVTALRDLLDFNADQPAISLEEVESVESIVK
RFCTGGMSLGALSREAHETLAIAMNRLGAKSNSGEGGEDVVRYLTLDDVDSEGNSPTLPHLH
GLQNGDTANSAIKQIASGRFGVTPEYLMSGKQLEIKMAQGAKPGEGGQLPGKKVSEYIAMLR
RSKPGVTLISPPPHHDIYSIEDLAQLIYDLHQINPEAQVSVKLVAEIGIGTIAAGVAKANAD
IIQISGHDGGTGASPLSSIKHAGSPWELGVTEVHRVLMENQLRDRVLLRADGGLKTGWDVVM
AALMGAEEYGFGSIAMIAEGCIMARVCHTNNCPVGVATQQERLRQRFKGVPGQVVNFFYFIA
EEVRSLLAHLGYRSLDDIIGRTDLLKVRSDVQLSKTQNLTLDCLLNLPDTKQNRQWLNHEPV
HSNGPVLDDDILADPDIQEAINHQTTATKTYRLVNTDRTVGTRLSGAIAKKYGNNGFEGNIT
LNFQGAAGQSFGAFNLDGMTLHLQGEANDYVGKGMNGGEIVIVPHPQASFAPEDNVIIGNTC
LYGATGGNLYANGRAGERFAVRNSVGKAVIEGAGDHCCEYMTGGVIVVLGPVGRNVGAGMTG
GLAYFLDEVGDLPEKINPEIITLQRITASKGEEQLKSLITAHVEHTGSPKGKAILANWSDYL
GKFWQAVPPSEKDSPEANNDVSLTGEKTLTSV
000000000000000000000000000000000000000000000000000000000000
000000000000000000000000000000000000000000000000000000000000
000000000000000000000000000000000000000000000000000000000000
000000000000000000000000000000000000000000000000000000000000
000000000000000000000000000000000000000000000000000000000000
000000000000000000000000000000000000000000000000000000000000
000000000000000000000000000000000000000000000000111111111111
111100000000000000000000000000000000000000000000000000000000
000000000000000000000000000000000000111110000000000000000000
000000000000000000000000000000000000000000000000000000000000
000000000000000000000000000000000000000000000000000000000000
000000000000000000000000000000000000000000000000000000000000
000000000000000000000000000000000000000000000000000000000000
000000111111111110000000000000000000000000000000000000000000
000000000000000000000000000000000000000000000000000000000000
000000000000000000000000000000000000000000000000000000000000
000000000000000000000000000000000000000000000000000000000000
```

000000000000000000000000000000000000000000000000000000000000
000000000000000000000000000000000000000000000000000000000000
000000000000000000000000000000000000000000000000000000000000
000000000000000000000000000000000000000000000000000000000000
000000000000000000000000000000000000000000000000000000000000
000000000000000000000000000000000000000000000000000000000000
000000000000000000000000000000000000000000000000000000000000
00000000000000000001111111111111
>DM_validata10
MSDLANSEKYYDEDPYGFEDESAPITAEDSWAVISAFFREKGLVSQQLDSFNQFVDYTLQDI
ICEDSTLILEQLAQHTTESDNISRKYEISFGKIYVTKPMVNESDGVTHALYPQEARLRNLTY
SSGLFVDVKKRTYEAIDVPGRELKYELIAEESEDDSESGKVFIGRLPIMLRSKNCYLSEATE
SDLYKLKECPFDMGGYFIINGSEKVLIAQERSAGNIVQVFKKAAPSPISHVAEIRSALEKGS
RFISTLQVKLYGREGSSARTIKATLPYIKQDIPIVIIFRALGIIPDGEILEHICYDVNDWQM
LEMLKPCVEDGFVIQDRETALDFIGRRGTALGIKKEKRIQYAKDILQKEFLPHITQLEGFES
RKAFFLGYMINRLLLCALDRKDQDDRDHFGKKRLDLAGPLLAQLFKTLFKKLTKDIFRYMQR
TVEEAHDFNMKLAINAKTITSGLKYALATGNWGEQKKAMSSRAGVSQVLNRYTYSSTLSHLR
RTNTPIGRDGKLAKPRQLHNTHWGLVCPAETPEGQACGLVKNLSLMSCISVGTDPMPIITFL
SEWGMEPLEDYVPHQSPDATRVFVNGVWHGVHRNPARLMETLRTLRRKGDINPEVSMIRDIR
EKELKIFTDAGRVYRPLFIVEDDESLGHKELKVRKGHIAKLMATEYQDIEGGFEDVEEYTWS
SLLNEGLVEYIDAEEEESILIAMQPEDLEPAEANEENDLDVDPAKRIRVSHHATTFTHCEIH
PSMILGVAASIIPFPDHNQSPRNTYQSAMGKQAMGVFLTNYNVRMDTMANILYYPQKPLGTT
RAMEYLKFRELPAGQNAIVAIACYSGYNQEDSMIMNQSSIDRGLFRSLFFRSYMDQEKKYGM
SITETFEKPQRTNTLRMKHGTYDKLDDDGLIAPGVRVSGEDVIIGKTTPISPDEEELGQRTA
YHSKRDASTPLRSTENGIVDQVLVTTNQDGLKFVKVRVRTTKIPQIGDKFASRHGQKGTIGI
TYRREDMPFTAEGIVPDLIINPHAIPSRMTVAHLIECLLSKVAALSGNEGDASPFTDITVEG
ISKLLREHGYQSRGFEVMYNGHTGKKLMAQIFFGPTYYQRLRHMVDDKIHARARGPMQVLTR
QPVEGRSRDGGLRFGEMERDCMIAHGAASFLKERLMEASDAFRVHICGICGLMTVIAKLNHN
QFECKGCDNKIDIYQIHIPYAAKLLFQELMAMNITPRLYTDRSRDF
111111111111111110000000000000000000000000000000000000000000
000000001111111111111111110000000000000000000000000000000000
000000000000011111111111111111111111111000000000000000000000
000000000000000000000000000000000000000000000000000000000000
000000000000000000000000000000000000000000000000000000000000
000000000000000000000000000000000000000000000000000000000000
000000000000000000000000000000000000000000000000000000001111
111111111110000000000000000000001111111111100000000000000000
000000111111000000000000000000000000000000000000000000000000
000000000000000000000000000000000000000000000000000000000000
000000000000000000000000000000000000000000000011111111100000
000000000000000000000000000000111111111000000000000000000000
000000000000000000000000000000000000000000000000000000000000
000000000000000000000000000000000000000000000000000000000000
000000000000000000000000000000000000000000000001111111111111

110000000000000000000000000000000000000000000000000000000000
000000000000000000000000000000000000000000000000000000000000
000000000000000000000000000000000000000000000000000000111111
111111111100000000000000000000000000000000000000000000000000
0000000000000000000000000000000000000000000000
>DM_validata12
MAPVLSKDVADIESILALNPRTQSHAALHSTLAKKLDKKHWKRNPDKNCFHCEKLENNFDDI
KHTTLGERGALREAMRCLKCADAPCQKSCPTHLDIKSFITSISNKNYYGAAKMIFSDNPLGL
TCGMVCPTSDLCVGGCNLYATEEGSINIGGLQQFASEVFKAMNIPQIRNPCLPSQEKMPEAY
SAKIALLGAGPASISCASFLARLGYSDITIFEKQEYVGGLSTSEIPQFRLPYDVVNFEIELM
KDLGVKIICGKSLSENEITLNTLKEEGYKAAFIGIGLPEPKTDDIFQGLTQDQGFYTSKDFL
PLVAKSSKAGMCACHSPLPSIRGAVIVLGAGDTAFDCATSALRCGARRVFLVFRKGFVNIRA
VPEEVELAKEEKCEFLPFLSPRKVIVKGGRIVAVQFVRTEQDETGKWNEDEDQIVHLKADVV
ISAFGSVLRDPKVKEALSPIKFNRWDLPEVDPETMQTSEPWVFAGGDIVGMANTTVESVNDG
KQASWYIHKYIQAQYGASVSAKPELPLFYTPVDLVDISVEMAGLKFINPFGLASAAPTTSSS
MIRRAFEAGWGFALTKTFSLDKDIVTNVSPRIVRGTTSGPMYGPGQSSFLNIELISEKTAAY
WCQSVTELKADFPDNIVIASIMCSYNKNDWMELSRKAEASGADALELNLSCPHGMGERGMGL
ACGQDPELVRNICRWVRQAVQIPFFAKLTPNVTDIVSIARAAKEGGADGVTATNTVSGLMGL
KADGTPWPAVGAGKRTTYGGVSGTAIRPIALRAVTTIARALPGFPILATGGIDSAESGLQFL
HSGASVLQVCSAVQNQDFTVIQDYCTGLKALLYLKSIEELQGWDGQSPGTESHQKGKPVPRI
AELMGKKLPNFGPYLEQRKKIIAEEKMRLKEQNAAFPPLERKPFIPKKPIPAIKDVIGKALQ
YLGTFGELSNIEQVVAVIDEEMCINCGKCYMTCNDSGYQAIQFDPETHLPTVTDTCTGCTLC
LSVCPIIDCIRMVSRTTPYEPKRGLPLAVNPVC
000000000000000000000000000000000000000000000000000000000000
000000000000000000000000000000000000000000000000000000000000
000000000000000000000000000000000000000000000000000000000000
000000000000000000000000000000000000000000000000000000000000
000000000000000000000000000000000000000000000000000000000000
000000000000000000000000000000000000000000000000000000000000
000000000000000000000000000000000000000000000000000000000000
000000000000000000000000000000000000000000000000000000000000
000000000000000000000000000000000000000000000000000000000000
000000000000000000000000000000000000000000000000000000000000
000000000000000000000000000000000000000000000000000011111000
000000000000000000000000000000000000000000000000000000000000
000000000000000000000000000000000000000000000000000000000000
000000000000000000000000000000000000000000000000000000000000
000000000000000000000000000000001111110000000000000000000000
000000000000000000000000000000000000000000000000000000000000
000000000000000000000000011111111
>DM_validata13
MPIFRFTALAMTLGLLSAPYNAMAATSNPAFDPKNLMQSEIYHFAQNNPLADFSSDKNSILT
LSDKRSIMGNQSLLWKWKGGSSFTLHKKLIVPTDKEASKAWGRSSTPVFSFWLYNEKPIDGY
LTIDFGEKLISTSEAQAGFKVKLDFTGWRAVGVSLNNDLENREMTLNATNTSSDGTQDSIGR

SLGAKVDSIRFKAPSNVSQGEIYIDRIMFSVDDARYQWSDYQVKTRLSEPEIQFHNVKPQLP
VTPENLAAIDLIRQRLINEFVGGEKETNLALEENISKLKSDFDALNIHTLANGGTQGRHLIT
DKQIIIYQPENLNSQDKQLFDNYVILGNYTTLMFNISRAYVLEKDPTQKAQLKQMYLLMTKH
LLDQGFVKGSALVTTHHWGYSSRWWYISTLLMSDALKEANLQTQVYDSLLWYSREFKSSFDM
KVSADSSDLDYFNTLSRQHLALLLLEPDDQKRINLVNTFSHYITGALTQVPPGGKDGLRPDG
TAWRHEGNYPGYSFPAFKNASQLIYLLRDTPFSVGESGWNNLKKAMVSAWIYSNPEVGLPLA
GRHPFNSPSLKSVAQGYYWLAMSAKSSPDKTLASIYLAISDKTQNESTAIFGETITPASLPQ
GFYAFNGGAFGIHRWQDKMVTLKAYNTNVWSSEIYNKDNRYGRYQSHGVAQIVSNGSQLSQG
YQQEGWDWNRMEGATTIHLPLKDLDSPKPHTLMQRGERGFSGTSSLEGQYGMMAFNLIYPAN
LERFDPNFTAKKSVLAADNHLIFIGSNINSSDKNKNVETTLFQHAITPTLNTLWINGQKIEN
MPYQTTLQQGDWLIDSNGNGYLITQAEKVNVSRQHQVSAENKNRQPTEGNFSSAWIDHSTRP
KDASYEYMVFLDATPEKMGEMAQKFRENNGLYQVLRKDKDVHIILDKLSNVTGYAFYQPASI
EDKWIKKVNKPAIVMTHRQKDTLIVSAVTPDLNMTRQKAATPVTINVTINGKWQSADKNSEV
KYQVSGDNTELTFTSYFGIPQEIKLSPLP
11111111111111111111111100000000000000000000000000000000000000
00000000000000000000000000000000000000000000000000000000000000
00000000000000000000000000000000000000000111111111111111111111
11111000000000000000000000000000000000000000000000000000000000
00000000000000000000000000000000000000000000000000000000000000
00000000000000000000000000000000000000000000000000000000000000
00000000000000000000000000000000000000000000000000000000000000
00000000000000000000000000000000000000000000000000000000000000
00000000000000000000000000000000000000000000000000000000000000
00000000000000000000000000000000000000000000000000000000000000
00000000000000000000000000000000000000000000000000000000000000
00000000000000000000000000000000000000000000000000000000000000
00000000000000000000000000000000000000000000000000000000000000
00000000000000000000000000000000000000000000000000000000000000
00000000000000000000000000000000000000000000000000000000000000
00000000000000000000000000000000000000000000000000000000000000
00000000000000000000000000000
>DM_validata14
FNLDVDSPAEYSGPEGSYFGFAVDFFVPSASSRMFLLVGAPKANTTQPGIVEGGQVLKCDWS
STRRCQPIEFDATGNRDYAKDDPLEFKSHQWFGASVRSKQDKILACAPLYHWRTEMKQEREP
VGTCFLQDGTKTVEYAPCRSQDIDADGQGFCQGGFSIDFTKADRVLLGGPGSFYWQGQLISD
QVAEIVSKYDPNVYSIKYNNQLATRTAQAIFDDSYLGYSVAVGDFNGDGIDDFVSGVPRAAR
TLGMVYIYDGKNMSSLYNFTGEQMAAYFGFSVAATDINGDDYADVFIGAPLFMDRGSDGKLQ
EVGQVSVSLQRASGDFQTTKLNGFEVFARFGSAIAPLGDLDQDGFNDIAIAAPYGGEDKKGI
VYIFNGRSTGLNAVPSQILEGQWAARSMPPSFGYSMKGATDIDKNGYPDLIVGAFGVDRAIL
YRARPVITVNAGLEVYPSILNQDNKTCSLPGTALKVSCFNVRFCLKADGKGVLPRKLNFQVE
LLLDKLKQKGAIRRALFLYSRSPSHSKNMTISRGGLMQCEELIAYLRDESEFRDKLTPITIF
MEYRLDYRTAADTTGLQPILNQFTPANISRQAHILLDCGEDNVCKPKLEVSVDSDQKKIYIG
DDNPLTLIVKAQNQGEGAYEAELIVSIPLQADFIGVVRNNEALARLSCAFKTENQTRQVVCD
LGNPMKAGTQLLAGLRFSVHQQSEMDTSVKFDLQIQSSNLFDKVSPVVSHKVDLAVLAAVEI

RGVSSPDHVFLPIPNWEHKENPETEEDVGPVVQHIYELRNNGPSSFSKAMLHLQWPYKYNNN
TLLYILHYDIDGPMNCTSDMEINPLRIKISSLQTTEKNDTVAGQGERDHLITKRDLALSEGD
IHTLGCGVAQCLKIVCQVGRLDRGKSAILYVKSLLWTETFMNKENQNHSYSLKSSASFNVIE
FPYKNLPIEDITNSTLVTTNVTWGIQP
000000000000000000000000000000000000000000000000000000000000
000000000000000000000000000000000000000000000000000000000000
000000000000000000000000000000000000000000000000000000000000
000000000000000000000000000000000000000000000000000000000000
000000000000000000000000000000000000000000000000000000000000
000000000000000000000000000000000000000000000000000000000000
000000000000000000000000000000000000000000000000000000000000
000000000000000000000000000000000000000000000000000000000000
000000000000000000000000000000000000000000000000000000000000
000000000000000000000000000000000000000000000000000000000000
000000000000000000000000000000000000000000000000000000000000
000000000000000000000000000000000000000000000000000000000000
000000000000000000000000000000000000000000000000000000000000
000000000000000000000000000000011111111111111111111111111110
000000000000000000000000000000000000000000000000000000000000
00000000000000000000000000
>DM_validata15
MAGRNWNASWIWGGQEESPRNEWRCFRGSFDAPASVEGPAMLHITADSRYVLFVNGEQVGRG
PVRSWPKEQFYDSYDIGGQLRPGVRNTIAVLVLHFGVSNFYYLRGRGGLIAEIEADGRTLAA
TDAAWRTERLGGQRSNSPRMACQQGFGEVIDARELAEDWALPAFDDGGWAQARSIGPAGTAP
WTSLVPRDIPFLTEEKLYPASIQSLSRVKAPKYAAALDLRNQMVPESVNHANPVSYCGYVAT
ILTLETSGVVTLGFPTGVRGSGVWVDGVLQTEWTGVQPERYYSLNLAAGEHLVLVDITSSDH
GGSSHFAIDSEAAFTLRSPAGDNGVPLATIGTFDQSEYIDHRPGRRMQTDHPDYRALPEAAP
TAAALEAFASWVKPFEPSLYTEENVFGSNVWRTLAERRAVPRSVLNAILPVPEPGVLPVFED
GDCELVIDLGAERSGFIGFELEAPAGTIIDAYGVEYMREGYTQHTYGLDNTFRYICREGRQS
YVSPVRRGFRYLFLTVRGNSAPVKLHEIYIRQSTYPVAEQGSFRCSDALLNATWEISRHTTR
LCMEDTFVDCPSYEQVFWVGDSRNEALVNYYVFGETEIVERCLNLVPGSADETPLYLDQVPS
AWSSVIPNWTFFWILACREYAAHTGNEAFAARIWPAVKHTLTHYLEHIDDSGLLNMAGWNLL
DWAPIDQPNEGIVTHQNLFLVKALRDSRALAAAAGATEEADAFAARADLLAETINAVLWDEE
KRAYIDCIHADGRRSDVYSMQTQVVAYLCGVAQGEREAVIEGYLSSPPPAFVQIGSPFMSFF
YYEALEKAGRQTLMLDDIRRNYGQMLRYDATTCWEMYPNFAENRSNPDMLTRSHCHAWSAAP
GYFLGSSILGVKRGADGWRTVDIAPQPCDLTWAEGVVPLPQGGHIAVSWEFVSAGKLKLRIE
APEDIEVNVTLPEGIEGEVTQVKYMS
110000000000000000000000000000000000000000000000000000000000
000000000000000000000000000000000000000000000000000000000000
000000000000000000000000000000000000000000000000000000000000
000000000000000000000000000000000000000000000000000000000000
000000000000000000000000000000000000000000000000000000000000
000000000000000000000000000000000000000000000000000000000000
000000000000000000000000000000000000000000000000000000000000

000000000000000000000000000000000000000000000000000000000000
000000000000000000000000000000000000000000000000000000000000
000000000000000000000000000000000000000000000000000000000000
000000000000000000000000000000000000000000000000000000000000
000000000000000000000000000000000000000000000000000000000000
000000000000000000000000000000000000000000000000000000000000
000000000000000000000000000000000000000000000000000000000000
000000000000000000000000000000000000000000000000000000000000
00000000000000000000000000
>DM_validata16
MGSSHHHHHHSSGLEVLFQGPAMATKKVHIISHSHWDREWYMAYEQHHMRLINLIDDLLEVF
QTDPDFHSFHLDGQTIILDDYLKVRPEREPEIRQAIASGKLRIGPFYILQDDFLTSSESNVR
NMLIGKEDCDRWGASVPLGYFPDTFGNMGQTPQLMLKAGLQAAAFGRGIRPTGFNNQVDTSE
KYSSQFSEISWQGPDNSRILGLLFANWYSNGNEIPTTEAEARLFWDKKLADAERFASTKHLL
MMNGCDHQPVQLDVTKAIALANQLYPDYEFVHSCFEDYLADLADDLPENLSTVQGEITSQET
DGWYTLANTASARIYLKQANTRVSRQLENITEPLAAMAYEVTSTYPHDQLRYAWKTLMQNHP
HDSICGCSVDSVHREMMTRFEKAYEVGHYLAKEAAKQIADAIDTRDFPMDSQPFVLFNTSGH
SKTSVAELSLTWKKYHFGQRFPKEVYQEAQEYLARLSQSFQIIDTSGQVRPEAEILGTSIAF
DYDLPKRSFREPYFAIKVRLRLPITLPAMSWKTLALKLGNETTPSETVSLYDDSNQCLENGF
LKVMIQTDGRLTITDKQSGLIYQDLLRFEDCGDIGNEYISRQPNHDQPFYADQGTIKLNIIS
NTAQVAELEIQQTFAIPISADKLLQAEMEAVIDITERQARRSQEKAELTLTTLIRMEKNNPR
LQFTTRFDNQMTNHRLRVLFPTHLKTDHHLADSIFETVKRPNHPDATFWKNPSNPQHQECFV
SLFDGENGVTIGNYGLNEYEILPDTNTIAITLLRSVGEMGDWGYFPTPEAQCLGKHSLSYSF
ESITKQTQFASYWRAQEGQVPVITTQTNQHEGTLAAEYSYLTGTNDQVALTAFKRRLADNAL
ITRSYNLSNDKTCDFSLSLPNYNAKVTNLLEKDSKQSTPSQLGKAEILTLAWKKQ
111111111111111111111111000000000000000000000000000000000000
000000000000000000000000000000000000000000000000000000000000
000000000000000000000000000000000000000000000000000000000000
000000000000000000000000000000000000000000000000000000000000
000000000000000000000000000000000000000000000000000000000000
000000000000000000000000000000000000000000000000000000000000
000000000000000000000000000000000000000000000000000000000000
000000000000000000000000000000000000000000000000000000000000
000000000000000000000000000000000000011111111000000000000000
000000000000000000000000000000000000000000000000000000000000
000000000000000000000000000000000000000000000000000000000000
000000000000000000000000000000000000000000000000000000000000
000000000000000000000000000000000000000000000000000000000000
000000000000000000000000000000000000000000000000000000000000
0000000000000000000000000000000000000000000000000000000
>DM_validata17
MIAAQLLAYYFTELKDDQVKKIDKYLYAMRLSDETLIDIMTRFRKEMKNGLSRDFNPTATVK
MLPTFVRSIPDGSEKGDFIALDLGGSSFRILRVQVNHEKNQNVHMESEVYDTPENIVHGSGS
QLFDHVAECLGDFMEKRKIKDKKLPVGFTFSFPCQQSKIDEAILITWTKRFKASGVEGADVV

KLLNKAIKKRGDYDANIVAVVNDTVGTMMTCGYDDQHCEVGLIIGTGTNACYMEELRHIDLV
EGDEGRMCINTEWGAFGDDGSLEDIRTEFDRAIDAYSLNPGKQLFEKMVSGMYLGELVRLIL
VKMAKEGLLFEGRITPELLTRGKFNTSDVSAIEKNKEGLHNAKEILTRLGVEPSDDDCVSVQ
HVCTIVSFRSANLVAATLGAILNRLRDNKGTPRLRTTVGVDGSLYKTHPQYSRRFHKTLRRL
VPDSDVRFLLSESGSGKGAAMVTAVAYRLAEQHRQIEETLAHFHLTKDMLLEVKKRMRAEME
LGLRKQTHNNAVVKMLPSFVRRTPDGTENGDFLALDLGGANFRVLLVKIRSGKKRTVEMHNK
IYAIPIEIMQGTGEELFDHIVSCISDFLDYMGIKGPRMPLGFTFSFPCQQTSLDAGILITWT
KGFKATDCVGHDVVTLLRDAIKRREEFDLDVVAVVNDTVGTMMTCAYEEPTCEVGLIVGTGS
NACYMEEMKNVEMVEGDQGQMCINMEWGAFGDNGCLDDIRTHYDRLVDEYSLNAGKQRYEKM
ISGMYLGEIVRNILIDFTKKGFLFRGQISETLKTRGIFETKFLSQIESDRLALLQVRAILQQ
LGLNSTCDDSILVKTVCGVVSRRAAQLCGAGMAAVVDKIRENRGLDRLNVTVGVDGTLYKLH
PHFSRIMHQTVKELSPKCNVSFLLSEDGSGKGAALITAVGVRLRTEASS
11111111111111100000000000000000000000000000000000000000000000
00000000000000000000000000000000000000000000000000000000000000
00000000000000000000000000000000000000000000000000000000000000
00000000000000000000000000000000000000000000000000000000000000
00000000000000000000000000000000000000000000000000000000000000
00000000000000000000000000000000000000000000000000000000000000
00000000000000000000000000000000000000000000000000000000000000
00000000000000000000000000000000000000000000000000000000000000
00000000000000000000000000000000000000000000000000000000000000
00000000000000000000000000000000000000000000000000000000000000
00000000000000000000000000000000000000000000000000000000000000
00000000000000000000000000000000000000000000000000000000000000
00000000000000000000000000000000000000000000000000000000000000
00000000000000000000000000000000000000000000000000000000000000
0000000000000000000000000000000000000000000001111
>DM_validata22
MLEEYRKHVAERAAEGIAPKPLDANQMAALVELLKNPPAGEEEFLLDLLTNRVPPGVDEAAY
VKAGFLAAIAKGEAKSPLLTPEKAIELLGTMQGGYNIHPLIDALDDAKLAPIAAKALSHTLL
MFDNFYDVEEKAKAGNEYAKQVMQSWADAEWFLNRPALAEKLTVTVFKVTGETNTDDLSPAP
DAWSRPDIPLHALAMLKNAREGIEPDQPGVVGPIKQIEALQQKGFPLAYVGDVVGTGSSRKS
ATNSVLWFMGDDIPHVPNKRGGGLCLGGKIAPIFFNTMEDAGALPIEVDVSNLNMGDVIDVY
PYKGEVRNHETGELLATFELKTDVLIDEVRAGGRIPLIIGRGLTTKAREALGLPHSDVFRQA
KDVAESDRGFSLAQKMVGRACGVKGIRPGAYCEPKMTSVGSQDTTGPMTRDELKDLACLGFS
ADLVMQSFCHTAAYPKPVDVNTHHTLPDFIMNRGGVSLRPGDGVIHSWLNRMLLPDTVGTGG
DSHTRFPIGISFPAGSGLVAFAAATGVMPLDMPESVLVRFKGKMQPGITLRDLVHAIPLYAI
KQGLLTVEKKGKKNIFSGRILEIEGLPDLKVEQAFELTDASAERSAAGCTIKLNKEPIIEYL
NSNIVLLKWMIAEGYGDRRTLERRIQGMEKWLANPELLEADADAEYAAVIDIDLADIKEPIL
CAPNDPDDARPLSAVQGEKIDEVFIGSCMTNIGHFRAAGKLLDAHKGQLPTRLWVAPPTRMD
AAQLTEEGYYSVFGKSGARIEIPGCSLCMGNQARVADGATVVSTSTRNFPNRLGTGANVFLA
SAELAAVAALIGKLPTPEEYQTYVAQVDKTAVDTYRYLNFNQLSQYTEKADGVIFQTAV
00000000000000000000000000000000000000000000000000000000000000
00000000000000000000000000000000000000000000000000000000000000

00000000000000000000000000000000000000000000000000000000000000
00000000000000000000000000000000000000000000000000000000000000
00000000000000000000000000000000000000000000000000000000000000
00000000000000000000000000000000000000000000000000000000000000
00000000000000000000000000000000000000000000000000000000000000
00000000000000000000000000000000000000000000000000000000000000
00000000000000000000000000000000000000000000000000000000000000
00000000000000000000000000000000000000000000000000000000000000
00000000000000000000000000000000000000000000000000000000000000
00000000000000000000000000000000000000000000000000000000000000
00000000000000000000000000000000000000000000000000000000000000
00000000000000000000000000000000000000000000000000000000111
>DM_validata23
MSAIENFDAHTPMMQQYLRLKAQHPEILLFYRMGDFYELFYDDAKRASQLLDISLTKRGASA
GEPIPMAGIPYHAVENYLAKLVNQGESVAICEQIGDPATSKGPVERKVVRIVTPGTISDEAL
LQERQDNLLAAIWQDSKGFGYATLDISSGRFRLSEPADRETMAAELQRTNPAELLYAEDFAE
MSLIEGRRGLRRRPLWEFEIDTARQQLNLQFGTRDLVGFGVENAPRGLCAAGCLLQYAKDTQ
RTTLPHIRSITMEREQDSIIMDAATRRNLEITQNLAGGAENTLASVLDCTVTPMGSRMLKRW
LHMPVRDTRVLLERQQTIGALQDFTAGLQPVLRQVGDLERILARLALRTARPRDLARMRHAF
QQLPELRAQLETVDSAPVQALREKMGEFAELRDLLERAIIDTPPVLVRDGGVIASGYNEELD
EWRALADGATDYLERLEVRERERTGLDTLKVGFNAVHGYYIQISRGQSHLAPINYMRRQTLK
NAERYIIPELKEYEDKVLTSKGKALALEKQLYEELFDLLLPHLEALQQSASALAELDVLVNL
AERAYTLNYTCPTFIDKPGIRITEGRHPVVEQVLNEPFIANPLNLSPQRRMLIITGPNMGGK
STYMRQTALIALMAYIGSYVPAQKVEIGPIDRIFTRVGAADDLASGRSTFMVEMTETANILH
NATEYSLVLMDEIGAGTSTYDGLSLAWACAENLANKIKALTLFATHYFELTQLPEKMEGVAN
VHLDALEHGDTIAFMHSVQDGAASKSYGLAVAALAGVPKEVIKRARQKLRELESIS
11111111000000000000000000000000000000000000000000000000111111
11000000000000000000000000000000011111111100000000000000000000
00000000000000000000000000000000000000000000000000000000000000
00000000000000000000000000000000000000000000000000000000000000
00000000000000000000000000000000000000000000000000000000000000
00000000000000000000000000000000000000000000000000000000000000
00000000000000000000000000000000000000000000000000000000000000
00000000000000000000000000000000000000000000000000000000000000
00000000000000000000000000000000000000000000000000000000000000
00000000000000000000000000000000000000000000000000000000000000
00000000000000000000000000000000000000011111110000000000000000
00000000000000000000000000000000000000000000000000000000000000
00000000000000000000000000000000000000000000000000000000
>DM_validata25
MSRRRHSDENDGGQPHKRRKTSDANETEDHLESLICKVGEKSACSLESNLEGLAGVLEADLP
NYKSKILRLLCTVARLLPEKLTIYTTLVGLLNARNYNFGGEFVEAMIRQLKESLKANNYNEA
VYLVRFLSDLVNCHVIAAPSMVAMFENFVSVTQEEDVPQVRRDWYVYAFLSSLPWVGKELYE
KKDAEMDRIFANTESYLKRRQKTHVPMLQVWTADKPHPQEEYLDCLWAQIQKLKKDRWQERH

ILRPYLAFDSILCEALQHNLPPFTPPPHTEDSVYPMPRVIFRMFDYTDDPEGPVMPGSHSVE
RFVIEENLHCIIKSHWKERKTCAAQLVSYPGKNKIPLNYHIVEVIFAELFQLPAPPHIDVMY
TTLLIELCKLQPGSLPQVLAQATEMLYMRLDTMNTTCVDRFINWFSHHLSNFQFRWSWEDWS
DCLSQDPESPKPKFVREVLEKCMRLSYHQRILDIVPPTFSALCPANPTCIYKYGDESSNSLP
GHSVALCLAVAFKSKATNDEIFSILKDVPNPNQDDDDDEGFSFNPLKIEVFVQTLLHLAAKS
FSHSFSALAKFHEVFKTLAESDEGKLHVLRVMFEVWRNHPQMIAVLVDKMIRTQIVDCAAVA
NWIFSSELSRDFTRLFVWEILHSTIRKMNKHVLKIQKELEEAKEKLARQHKRRSDDDDRSSD
RKDGVLEEQIERLQEKVESAQSEQKNLFLVIFQRFIMILTEHLVRCETDGTSVLTPWYKNCI
ERLQQIFLQHHQIIQQYMVTLENLLFTAELDPHILAVFQQFCALQA
11111111111111111111111110000000000000000000000000000000000000
00000000000000000000000000000000000000000000000000000000000000
00000000000000000000000000000000000000000000000000000000000000
00000000000000000000000000000000000000000000000000000000000000
00000000000000000000000000000000000000000000000000000000000000
00000000000000000000000000000000000000000000000000000000000000
00000000000000000000000000000000000000000000000000000000000000
00000000000000000000000000000000000000000000000000000000000000
00000000000000000000000000000011111111111100000000000000000000
00000000000000000000000000000000000000000000000000000000000000
00000000000000000000000000000000000000000000011111111111111111
11110000000000000000000000000000000000000000000000000000000000
0000000000000000000000000000000000000000000000
>DM_validata28
ASMTGGQQMGRGSGSEFMENETHYAEAVIDNGAFGTRTIRFETGRLARQAAGSAVAYLDDDT
MVLSATTASKNPKDQLDFFPLTVDVEERMYAAGKIPGSFFRREGRPSEDAILTCRLIDRPLR
PSFKKGLRNEIQVVATIMALNPDHLYDVVAINAASASTQLAGLPFSGPIGGVRVALIRGQWV
AFPTHTELEDAVFDMVVAGRVLEDGDVAIMMVEAEATEKTIQLVKDGAEAPTEEVVAAGLDA
AKPFIKVLCKAQADLAAKAAKPTGEFPVFLDYQDDVLEALSAAVRPELSAALTIAGKQDREA
ELDRVKALAAEKLLPEFEGREKEISAAYRALTKSLVRERVIAEKKRIDGRGVTDIRTLAAEV
EAIPRVHGSALFERGETQILGVTTLNMLRMEQQLDTLSPVTRKRYMHNYNFPPYSVGETGRV
GSPKRREIGHGALAERAIVPVLPTREEFPYAIRQVSEALGSNGSTSMGSVCASTMSLLNAGV
PLKAPVAGIAMGLISQEINGETHYVALTDILGAEDAFGDMDFKVAGTKEFVTALQLDTKLDG
IPASVLAAALKQARDARLHILDVMMEAIDTPDEMSPNAPRIITVKIPVDKIGEVIGPKRQMI
NQIQEDTGAEITIEDDGTIYIGAADGPAAEAARATINGIANPTSPEVGERILGSVVKTTTFG
AFVSLLPGKDGLLHISQIRKLAGGKRVENVEDVLGVGQKVQVEIAEIDSRGKLSLIPVIEGE
EAASDEKKDDAEQ
11111111111111111110000000000000000000000000000000000000000000
00000000000000000000000000000000000000000000000000000000000000
00000000000000000000000000000000000000000000000000000000000000
00000000000000000000000000000000000000000000000000000000000000
00000000000000000000000000000000000000000000000000000000000000
00000000000000000000000000000000000000000000000000000000000000
00000000000000000000000000000000000000000000000000000000000000
00000000000000000000000000000000000000000000000000000000000000

00000000000000000000000000000000000000000000000000000000000000
00000000000000000000000000000000000000000000111111111111111111
00000000000111111110000000000001111111111111111111110000000000
00000011100000111111111111111111100000000000000000001111111111
1111111111111
>DM_validata29
MTISPPEREPKVRVVVDNDPVPTSFEKWAKPGHFDRTLARGPQTTTWIWNLHALAHDFDTHT
SDLEDISRKIFSAHFGHLAVVFIWLSGMYFHGAKFSNYEAWLADPTGIKPSAQVVWPIVGQG
ILNGDVGGGFHGIQITSGLFQLWRASGITNEFQLYCTAIGGLVMAGLMLFAGWFHYHKRAPK
LEWFQNVESMLNHHLAGLLGLGSLAWAGHQIHVSLPINKLLDAGVAAKDIPLPHEFILNPSL
MAELYPKVDWGFFSGVIPFFTFNWAAYSDFLTFNGGLNPVTGGLWLSDTAHHHLAIAVLFII
AGHMYRTNWGIGHSLKEILEAHKGPFTGAGHKGLYEVLTTSWHAQLAINLAMMGSLSIIVAQ
HMYAMPPYPYLATDYPTQLSLFTHHMWIGGFLVVGGAAHGAIFMVRDYDPAMNQNNVLDRVL
RHRDAIISHLNWVCIFLGFHSFGLYVHNDTMRAFGRPQDMFSDTGIQLQPVFAQWVQNLHTL
APGGTAPNAAATASVAFGGDVVAVGGKVAMMPIVLGTADFMVHHIHAFTIHVTVLILLKGVL
FARSSRLIPDKANLGFRFPCDGPGRGGTCQVSGWDHVFLGLFWMYNCISVVIFHFSWKMQSD
VWGTVAPDGTVSHITGGNFAQSAITINGWLRDFLWAQASQVIGSYGSALSAYGLLFLGAHFI
WAFSLMFLFSGRGYWQELIESIVWAHNKLKVAPAIQPRALSIIQGRAVGVAHYLLGGIATTW
AFFLARIISVG
11111111111100000000000000000000000000000000000000000000000000
00000000000000000000000000000000000000000000000000000000000000
00000000000000000000000000000000000000000000000000000000000000
00000000000000000000000000000000000000000000000000000000000000
00000000000000111000000000000000000000000000000000000000000000
00000000000000000000000000000000000000000000000000000000000000
00000000000000000000000000000000000000000000000000000000000000
00000000000000000000000000000000000000000000000000000000000000
00000000000000000000000000000000000000000000000000000000000000
00000000000000000000000000000000000000000000000000000000000000
00000000000000000000000000000000000000000000000000000000000000
00000000000000000000000000000000000000000000000000000000000000
00000000000
>DM_validata30
MTSTGRFTLPSEENFAEKTKELAELWGADAIRNSDGTHLDEAVLALGKKIYNAYFPTRAHNE
WITLHMDETPQVYLLTDRILAESDTVDIPLMESFFAEQLKPNRDADPHKYWEVVDRTTGEVV
DSANWTLDADEDTVHVSGVAAWHEYTVSFLAYIIWDPVEMYNHLTNDWGDKEHEIPFDIYHP
ATRKFVFDTFEQWLKDSPQTDVVRFTTFFYQFTLLFDEKRREKVVDWFGCACTVSPRALDDF
EAKYGYRLRPEDFVDGGAYNSAWRVPRKAQRDWIDFLSGFVRENVKQLADMSHAAGKEAMMF
LGDQWIGTEPYKDGFDELGLDAVVGSIGDGTTTRMIADIPGVKYTEGRFLPYFFPDTFYEGN
DPSIEGLDNWRKARRAILRSPISRMGYGGYLSLAAKFPKFVDTVTHIANEFRDIHDRTGGVA
AEGELNVAILNSWGKMRSWMAFTVAHALPNKQTYSYYGILESLSGMRVNVRFISFDDVLAHG
IDSDIDVIINGGPVDTAFTGGDVWTNPKLVETVRAWVRGGGAFVGVGEPSSAPRFQTGRFFQ
LADVIGVDEERYQTLSVDKYFPPVVPDHFITADVPVDPAAREAWEQAGYRIPLSGCGGGQSI
KPLGGIDFGEPVLNTYPVNENVTLLRADGGQVQLATNDYGKGRGVYISGLPYSAANARLLER

VLFYASHNEDKYAAWSSSNPECEVAHFPEQGLYCVINNTDQPQKTTVTLADGTTEDFDLPDS
GIAWREA
110000000000000000000000000000001111111100000000000000000000
000000000000000000000000000000000000000000000000000000000000
000000000000000000000000000000000000000000000000000000000000
000000000000000000000000000000000000000000000000000000000000
000000000000000000000000000000000000000000000000000000000000
000000000000000000000000000000000000000000000000000000000000
000000000000000000000000000000000000000000000000000000000000
000000000000000000000000000000000000000000000000000000000000
000000000000000000000000000000000000000000000000000000000000
000000000000000000000000000000000000000000000000000000000000
000000000000000000000000000000000000000000000000000000000000
000000000000000000000000000000000000000000000000000000000000
0000001
>DM_validata31
MSLTKVSSLGYPRLGENREWKKLIEAYWAGKVSKNDLFAGAKELRLDFLKKQLNAGLDLIPV
GDFSLYDHILDLSVQFNIIPKRFAKEPIDIDLYFAIARGNKENVASSMKKWFNTNYHYIVPE
WSKQRPKLNNNRLLDLYLEAREVVGDKAKPVITGPITYVALSTGVEDFTAAVKSLLPLYKQV
FTELVKAGASYIQVDEPIFVTDEGKDYLQAAKAVYAYFAKEVPDAKFIFQTYFEGLIDSQVL
SQLPVDAFGLDFVYGLEENLEAIKTGAFKGKEIFAGVIDGRNIWSSDFVKTSALLETIEEQS
AALTIQPSCSLLHVPVTTKNETDLDPVLRNGLAFADEKLTEVKRLAEHLDGREDPAYDLHIA
HFDALQAADFRNVKLEDLSRVATKRPSDFAKRRDIQQEKLHLPLLPTTTIGSFPQSREIRRT
RLAWKRGDISDAEYKQFIQAEIERWIRIQEDLDLDVLVHGEFERVDMVEFFGQKLAGFTTTK
FGWVQSYGSRAVKPPIIYGDVQHLEPITVEETVYAQSLTDRPVKGMLTGPITITNWSFERTD
IPRDQLFNQIGLAIKDEIKLLENAGIAIIQVDEAALREGLPLRKSKQKAYLDDAVHAFHIAT
SSVKDETQIHTHMCYSKFDEIIDAIRALDADVISIETSRSHGDIIESFETAVYPLGIGLGVY
DIHSPRVPTKEEVVANIERPLRQLSPTQFWVNPDCGLKTRQEPETIAALKVLVAATKEVRQK
LGNEG
110000000000000000000000000000000000000000000000000000000000
000000000000000000000000000000000000000000000000000000000000
000000000000000000000000000000000000000000000000000000000000
000000000000000000000000000000000000000000000000000000000000
000000000000000000000000000000000000000000000000000000000000
000000000000000000000000000000000000000000000000000000000000
000000000000000000000000000000000000000000000000000001111111
111111111100000000000000000000000000000000000000000000000000
000000000000000000000000000000000000000000000000000000000000
000000000000000000000000000000000000000000000000000000000000
000000000000000000000000000000000111111111111111111100000000
000000000000000000000000000000000000000000000000000000000000
11111
>DM_validata32
MAKIIWTRTDEAPLLATYSLKPVVEAFAATAGIEVETRDISLAGRILAQFPERLTEDQKVGN

ALAELGELAKTPEANIIKLPNISASVPQLKAAIKELQDQGYDIPELPDNATTDEEKDILARY
NAVKGSAVNPVLREGNSDRRAPIAVKNFVKKFPHRMGEWSADSKTNVATMDANDFRHNEKSI
ILDAADEVQIKHIAADGTETILKDSLKLLEGEVLDGTVLSAKALDAFLLEQVARAKAEGILF
SAHLKATMMKVSDPIIFGHVVRAYFADVFAQYGEQLLAAGLNGENGLAAILSGLESLDNGEE
IKAAFEKGLEDGPDLAMVNSARGITNLHVPSDVIVDASMPAMIRTSGHMWNKDDQEQDTLAI
IPDSSYAGVYQTVIEDCRKNGAFDPTTMGTVPNVGLMAQKAEEYGSHDKTFRIEADGVVQVV
SSNGDVLIEHDVEANDIWRACQVKDAPIQDWVKLAVTRSRLSGMPAVFWLDPERAHDRNLAS
LVEKYLADHDTEGLDIQILSPVEATQLSIDRIRRGEDTISVTGNVLRDYNTDLFPILELGTS
AKMLSVVPLMAGGGLFETGAGGSAPKHVQQVQEENHLRWDSLGEFLALAESFRHELNNNGNT
KAGVLADALDKATEKLLNEEKSPSRKVGEIDNRGSHFWLTKFWADELAAQTEDADLAATFAP
VAEALNTGAADIDAALLAVQGGATDLGGYYSPNEEKLTNIMRPVAQFNEIVDALKK
10000000000000000000000000000000000000000000000000000000000000
00000000000000000000000000000000000000000000000000000000000000
00000000000000000000000000000000000000000000000000000000000000
00000000000000000000000000000000000000000000000000000000000000
00000000000000000000000000000000000000000000000000000000000000
00000000000000000000000000000000000000000000000000000000000000
00000000000000000000000000000000000000000000000000000000000000
00000000000000000000000000000000000000000000000000000000000000
00000000000000000000000000000000000000000000000000000000000000
00000000000000000000000000000000000000000000000000000000000000
00000000000000000000000000000000000000000000000000000000000000
00000000000000000000000000000000000000000000000000000011
>DM_validata33
RSPGTLPRKAGVFSDLSNQELKAVHSFLWSKKELRLQPSSTTTMAKNTVFLIEMLLPKKYHV
LRFLDKGERHPVREARAVIFFGDQEHPNVTEFAVGPLPGPCYMRALSPRPGYQSSWASRPIS
TAEYALLYHTLQEATKPLHQFFLNTTGFSFQDCHDRCLAFTDVAPRGVASGQRRSWLIIQRY
VEGYFLHPTGLELLVDHGSTDAGHWAVEQVWYNGKFYGSPEELARKYADGEVDVVVLEDPLP
GGKGHDSTEEPPLFSSHKPRGDFPSPIHVSGPRLVQPHGPRFRLEGNAVLYGGWSFAFRLRS
SSGLQVLNVHFGGERIAYEVSVQEAVALYGGHTPAGMQTKYLDVGWGLGSVTHELAPGIDCP
ETATFLDTFHYYDADDPVHYPRALCLFEMPTGVPLRRHFNSNFKGGFNFYAGLKGQVLVLRT
TSTVYNYDYIWDFIFYPNGVMEAKMHATGYVHATFYTPEGLRHGTRLHTHLIGNIHTHLVHY
RVDLDVAGTKNSFQTLQMKLENITNPWSPRHRVVQPTLEQTQYSWERQAAFRFKRKLPKYLL
FTSPQENPWGHKRSYRLQIHSMADQVLPPGWQEEQAITWARYPLAVTKYRESELCSSSIYHQ
NDPWDPPVVFEQFLHNNENIENEDLVAWVTVGFLHIPHSEDIPNTATPGNSVGFLLRPFNFF
PEDPSLASRDTVIVWPRDNGPNYVQRWIPEDRDCSMPPPFSYNGTYRPV
11111110000000000000000000000000000000000000000000000000000000
00000000000000000000000000000000000000000000000000000000000000
00000000000000000000000000000000000000000000000000000000000000
00000000000000000000000000000000000000000000000000000000011111
11111111100000000000000000000000000000000000000000000000000000
00000000000000000000000000000000000000000000000000000000000000
00000000000000000000000000000000000000000000000000000000000000
00000000000000000000000000000000000000000000000000000000000000

000000000000000000000000000000000000000000000000000000000000
000000000000000000000000000000000000000000000000000000000000
000000000000000000000000000000000000000000000000000000000000
000000000000000000000000000000000000000000000000
>DM_validata34
MADTQYILPNDIGVSSLDCREAFRLLSPTERLYAYHLSRAAWYGGLAVLLQTSPEAPYIYAL
LSRLFRAQDPDQLHQHALAEGLTEEEYQAFLVYAAGVYSNMGNYKSFGDTKFVPNLPKEKLE
RVILGSEAAQQHPEEVRGLWQTCGELMFSLEPRLRHLGLGKEGITTYFSGNCTMEDAKLAQD
FLDSQNLSAYNTRLFKEVDGEGKPYYEVRLASVLGSEPSLDSEVTSKLKSYEFRGSPFQVTR
GDYAPILQKVVEQLEKAKAYAANSHQGQMLAQYIESFTQGSIEAHKRGSRFWIQDKGPIVES
YIGFIESYRDPFGSRGEFEGFVAVVNKAMSAKFERLVASAEQLLKELPWPPTFEKDKFLTPD
FTSLDVLTFAGSGIPAGINIPNYDDLRQTEGFKNVSLGNVLAVAYATQREKLTFLEEDDKDL
YILWKGPSFDVQVGLHELLGHGSGKLFVQDEKGAFNFDQETVINPETGEQIQSWYRSGETWD
SKFSTIASSYEECRAESVGLYLCLHPQVLEIFGFEGADAEDVIYVNWLNMVRAGLLALEFYT
PEAFNWRQAHMQARFVILRVLLEAGEGLVTITPTTGSDGRPDARVRLDRSKIRSVGKPALER
FLRRLQVLKSTGDVAGGRALYEGYATVTDAPPECFLTLRDTVLLRKESRKLIVQPNTRLEGS
DVQLLEYEASAAGLIRSFSERFPEDGPELEEILTQLATADARFW
111100000000000000000000000000000000000000000000000000000000
000000000000000000000000000000000000000000000000000000000000
000000000000000000000000000000000000000000000000000000000000
000000000000000000000000000000000000111100000000000000000000
000000000000000000000000000000000000000000000000000000000000
000000000000000000000000000000000000000000000000000000000000
000000000000000000000000000000000000000000000000000000000000
000000000000000000000000000000000000000000000000000000000000
000000000000000000000000000000000000000000000000000000000000
000000000000000000000000000000000000000000000000000000000000
000000000000000000000000000000000000000000000000000000000000
00000000000000000000000000000000000000000000
>DM_validata35
MRVDELRVDERIKSTLKERGIESFYPPQAEALKSGILEGKNALISIPTASGKTLIAEIAMVH
RILTQGGKAVYIVPLKALAEEKFQEFQDWEKIGLRVAMATGDYDSKDEWLGKYDIIIATAEK
FDSLLRHGSSWIKDVKILVADEIHLIGSRDRGATLEVILAHMLGKAQIIGLSATIGNPEELA
EWLNAELIVSDWRPVKLRRGVFYQGFVTWEDGSIDRFSSWEELVYDAIRKKKGALIFVNMRR
KAERVALELSKKVKSLLTKPEIRALNELADSLEENPTNEKLAKAIRGGVAFHHAGLGRDERV
LVEENFRKGIIKAVVATPTLSAGINTPAFRVIIRDIWRYSDFGMERIPIIEVHQMLGRAGRP
KYDEVGEGIIVSTSDDPREVMNHYIFGKPEKLFSQLSNESNLRSQVLALIATFGYSTVEEIL
KFISNTFYAYQRKDTYSLEEKIRNILYFLLENEFIEISLEDKIRPLSLGIRTAKLYIDPYTA
KMFKDKMEEVVKDPNPIGIFHLISLTPDITPFNYSKREFERLEEEYYEFKDRLYFDDPYISG
YDPYLERKFFRAFKTALVLLAWINEVPEGEIVEKYSVEPGDIYRIVETAEWLVYSLKEIAKV
LGAYEIVDYLETLRVRVKYGIREELIPLMQLPLVGRRRARALYNSGFRSIEDISQARPEELL
KIEGIGVKTVEAIFKFLGKNVKISEKPRKSTLDYFLKS
000000000000000000000000000000000000000000000000000000000000
000000000000000000000000000000000000000000000000000000000000

00000000000000000000000000000000000000000000000000000000000000
00000000000000000000000000000000000000000000000000000000000000
00000000000000000000000000000000000000000000000000000000000000
00000000000000000000000000000000000000000000000000000000000000
00000000000000000000000000000000000000000000000000000000000000
00000000000000000000000000000000000000000000000000000000000000
00000000000000000000000000000000000000000000000000000000000000
00000000000000000000000000000000000000000000000000000000000000
00000000000000000000000000000000000000000000000000000000000000
00000000000000000011111111111111111111
>DM_validata36
VSNAKLLGGSPYSPFRIGLEGVWTPEVLKARASVIGKPIGESYKRILAKLQRIHNSNILDER
QGLMHELMELIDLYEESQPSSERLNAFRELRTQLEKALYLPEMEALKKQILQIPNKGSGAAR
FLLRTAMNEMAGKTSESTADLIRFALQDTVISAPFRGYAGAIPEAIDFPVKYVIEDISVFDK
IQTNYWELPAYESWNEGSNSALLPGLLRESQSKGMLSKCRIIENSLYIGHSYEEMFYSISPY
SNQVGGPYELYPFTFFSMLQEVQGDLGFEQAFATRNFFNTLVSDRLSLMENTMLLTESFDYT
PWDAIYGDINYDEQFAAMSINERIEKCMNTYRGVAFQNSSKSIDFFLNNLTTFIDNGLTEIA
ISDLPYDIVQQEISQFLQGSNEWKTLDAMLFNLDKGDINGAFRKLLQSAKDNNIKFRAIGHS
DNSVPPFNNPYKSLYYKGNIIAEAIEKLDREGQKFVVFADSSLLNSTPGTGRPMPGLVQYLK
IPATVVDSDGAWQFLPDVASSRVPIEVTELENWQVLTPPQGKILGLKQFKLTAGFPTEQSRL
PLLENSVSEDLREELMQKIDAIKNDVKMNSLVCMEAGSCDSVSPKVAARLKDMGLEAGMGAS
ITWWRREGGMEFSHQMHTTASFKFAGKEFAVDASHLQFVHDQLDTTILILPVDDWALEIAQR
NRAINPFVEYVSKTGNMLALFMPPLFTKPRLTRAL
11111100000000000000000000000000000000000000000000000000000000
00000000000000000000000000000000000000000000000000000000000000
00000000000000000000000000000000000000000000000000000000000000
00000000000000000000000000000000000000000000000000000000000000
00000000000000000000000000000000000000000000000000000000000000
00000000000000000000000000000000000000000000000000000000000000
00000000000000000000000000000000000000000000000000000000000000
00000000000000000000000000000000000000000000000000000000000000
00000000000000000000000000000000000000000000000000000000000000
00000000000000000000000000000000000000000000000000000000000000
00000000000000000000000000000000000000000000000000000000000000
00000000000000000000000000000000000
>DM_validata37
QQKLTSPDNNLVMTFQVDSKGAPTYELTYKNKVVIKPSTLGLELKKEDNTRTDFDWVDRRDL
TKLDSKTNLYDGFEVKDTQTATFDETWQPVWGEEKEIRNHYNELAVTLYQPMNDRSIVIRFR
LFNDGLGFRYEFPQQKSLNYFVIKEEHSQFGMNGDHIAFWIPGDYDTQEYDYTISRLSEIRG
LMKEAITPNSSQTPFSQTGVQTALMMKTDDGLYINLHEAALVDYSCMHLNLDDKNMVFESWL
TPDAKGDKGYMQTPCNTPWRTIIVSDDARNILASRITLNLNEPCKIADAASWVKPVKYIGVW
WDMITGKGSWAYTDELTSVKLGETDYSKTKPNGKHSANTANVKRYIDFAAAHGFDAVLVEGW
NEGWEDWFGNSKDYVFDFVTPYPDFDVKEIHRYAARKGIKMMMHHETSASVRNYERHMDKAY
QFMADNGYNSVKSGYVGNIIPRGEHHYGQWMNNHYLYAVKKAADYKIMVNAHEATRPTGICR

TYPNLIGNESARGTEYESFGGNKVYHTTILPFTRLVGGPMDYTPGIFETHCNKMNPANNSQV
RSTIARQLALYVTMYSPLQMAADIPENYERFMDAFQFIKDVALDWDETNYLEAEPGEYITIA
RKAKDTDDWYVGCTAGENGHTSKLVFDFLTPGKQYIATVYADAKDADWKENPQAYTIKKGIL
TNKSKLNLHAANGGGYAISIKEVKDKSEAKGLKRL
0000000000000000000000000000000000000000000000001111111111111111
0000000000000000000000000000000000000000000000000000000000000000
0000000000000000000000000000000000000000000000000000000000000000
0000000000000000000000000000000000000000000000000000000000000000
0000000000000000000000000000000000000000000000000000000000000000
0000000000000000000000000000000000000000000000000000000000000000
0000000000000000000000000000000000000000000000000000000000000000
0000000000000000000000000000000000000000000000000000000000000000
0000000000000000000000000000000000000000000000000000000000000000
0000000000000000000000000000000000000000000000000000000000000000
0000000000000000000000000000000000000000000000000000000000000000
00000000000000000000000000000000000
>DM_validata38
AVEPKEDTITVTAAPAPQESAWGPAATIAARQSATGTKTDTPIQKVPQSISVVTAEEMALHQ
PKSVKEALSYTPGVSVGTRGASNTYDHLIIRGFAAEGQSQNNYLNGLKLQGNFYNDAVIDPY
MLERAEIMRGPVSVLYGKSSPGGLLNMVSKRPTTEPLKEVQFKAGTDSLFQTGFDFSDSLDD
DGVYSYRLTGLARSANAQQKGSEEQRYAIAPAFTWRPDDKTNFTFLSYFQNEPETGYYGWLP
KEGTVEPLPNGKRLPTDFNEGAKNNTYSRNEKMVGYSFDHEFNDTFTVRQNLRFAENKTSQN
SVYGYGVCSDPANAYSKQCAALAPADKGHYLARKYVVDDEKLQNFSVDTQLQSKFATGDIDH
TLLTGVDFMRMRNDINAWFGYDDSVPLLNLYNPVNTDFDFNAKDPANSGPYRILNKQKQTGV
YVQDQAQWDKVLVTLGGRYDWADQESLNRVAGTTDKRDDKQFTWRGGVNYLFDNGVTPYFSY
SESFEPSSQVGKDGNIFAPSKGKQYEVGVKYVPEDRPIVVTGAVYNLTKTNNLMADPEGSFF
SVEGGEIRARGVEIEAKRPLSASVNVVGSYTYTDAEYTTDTTYKGNTPAQVPKHMASLWADY
TFFDGPLSGLTLGTGGRYTGSSYGDPANSFKVGSYTVVDALVRYDLARVGMAGSNVALHVNN
LFDREYVASCFNTYGCFWGAERQVVATATFRF
1111111111111111111000000000000000000000000000000000000000000000
0000000000000000000000000000000000000000000000000000000000000000
0000000000000000000000000000000000000000000000000000000000000000
0000000000000000000000000000000000000000000000000000000000000000
0000000000000000000000000000000000000000000000000000000000000000
0000000000000000000000000000000000000000000000000000000000000000
0000000000000000000000000000000000000000000000000000000000000000
0000000000000000000000000000000000000000000000000000000000000000
0000000000000000000000000000000000000000000000000000000000000000
0000000000000000000000000000000000000000000000000000000000000000
0000000000000000000000000000000000000000000000000000000000000000
00000000000000000000000000000000
>DM_validata39
DGNNLNNPVLLLDTFDFGPREDNWFFYPGGNIGLYCPYSSKGAPEEDSAMVFVSNEVGEHSI
TTRDLSVNENTIIQFEINVGCSTDSSSADPVRLEFSRDFGATWHLLLPLCYHSSSLVSSLCS

TEHHPSSTYYAGTTQGWRREVVHFGKLHLCGSVRFRWYQGFYPAGSQPVTWAIDNVYIGPQC
EEMCYGHGSCINGTKCICDPGYSGPTCKISTKNPDFLKDDFEGQLESDRFLLMSGGKPSRKC
GILSSGNNLFFNEDGLRMLVTRDLDLSHARFVQFFMRLGCGKGVPDPRSQPVLLQYSLNGGL
SWSLLQEFLFSNSSNVGRYIALEMPLKARSGSTRLRWWQPSENGHFYSPWVIDQILIGGNIS
GNTVLEDDFSTLDSRKWLLHPGGTKMPVCGSTGDALVFIEKASTRYVVTTDIAVNEDSFLQI
DFAASCSVTDSCYAIELEYSVDLGLSWHPLVRDCLPTNVECSRYHLQRILVSDTFNKWTRIT
LPLPSYTRSQATRFRWHQPAPFDKQQTWAIDNVYIGDGCLDMCSGHGRCVQGSCVCDEQWGG
LYCDEPETSLPTQLKDNFNRAPSNQNWLTVSGGKLSTVCGAVASGLALHFSGGCSRLLVTVD
LNLTNAEFIQFYFMYGCLITPSNRNQGVLLEYSVNGGITWNLLMEIFYDQYSKPGFVNILLP
PDAKEIATRFRWWQPRHDGLDQNDWAIDNVLI
11111111000000000000000000000000000000000000000000000000000000
00000000000000000000000000000000000000000000000000000000000000
00000000000000000000000000000000000000000000000000000000000000
00000000000000000000000000000000000000000000000000000000000000
00000000000000000000000000000000000000000000000000000000000000
00000000000000000000000000000000000000000000000000000000000000
00000000000000000000000000000000000000000000000000000000000000
00000000000000000000000000000000000000000011100000000000000000
00000000000000000000000000000000000000000000000000000000000000
00000000000000000000000000000000000000000000000000000000000000
00000000000000000000000000000000000000000000000000000000000000
00000000000000000000000000000000
>DM_validata40
MKNLTTIKQTNKNVKQERRKKYADLAIQGTNNSSIASKRSVELLYLPKLSSANNFQMDKNNK
LLEYFKFFVPKKIKRSPCINRGYWLRLFAIRSRLNSIIEQTPQDKKIVVVNLGCGYDPLPFQ
LLDTNNIQSQQYHDRVSFIDIDYSDLLKIKIELIKTIPELSKIIGLSEDKDYVDDSNVDFLT
TPKYLARPCDLNDSKMFSTLLNECQLYDPNVVKVFVAEVSLAYMKPERSDSIIEATSKMENS
HFIILEQLIPKGPFEPFSKQMLAHFKRNDSPLQSVLKYNTIESQVQRFNKLGFAYVNVGDMF
QLWESADEATKKELLKVEPFDELEEFHLFCHHYVLCHATNYKEFAFTQGFLFDRSISEINLT
VDEDYQLLECECPINRKFGDVDVAGNDVFYMGGSNPYRVNEILQLSIHYDKIDMKNIEVSSS
EVPVARMCHTFTTISRNNQLLLIGGRKAPHQGLSDNWIFDMKTREWSMIKSLSHTRFRHSAC
SLPDGNVLILGGVTEGPAMLLYNVTEEIFKDVTPKDEFFQNSLVSAGLEFDPVSKQGIILGG
GFMDQTTVSDKAIIFKYDAENATEPITVIKKLQHPLFQRYGSQIKYITPRKLLIVGGTSPSG
LFDRTNSIISLDPLSETLTSIPISRRIWEDHSLMLAGFSLVSTSMGTIHIIGGGATCYGFGS
VTNVGLKLIAIAK
11111111111111100000000000000000000000000000000000000000000000
00000000000000000000000000000000000000000000000000000000000000
00000000000000000000000000000000000000000000000000000000000000
00000000000000000000000000000000000000000000000000000000000000
00000000000000000000000000000000000000000000000000000000000000
00000000000000000000000000000000000000000000000000000011110000
00000000000000000000000000000000000000000000000000000000000000
00000000000000000000000000000000000000000000000000000000000000
00000000000000000000000000000000000000000000000000000000000000

00000000000000000000000000000000000000000000000000000000000000
00000000000000000000000000000000000000000001110000000000000000
0000000000001
>DM_validata44
MASKDKEINNTIDAIEDKNFKQVYKDSSYISKSDNGEVEMTERPIKIYNSLGVKDINIQDRK
IKKVSKNKKRVDAQYKIKTNYGNIDRNVQFNFVKEDGMWKLDWDHSVIIPGMQKDQSIHIEN
LKSERGKILDRNNVELANTGTAYEIGIVPKNVSKKDYKAIAKELSISEDYIKQQMDQNWVQD
DTFVPLKTVKKMDEYLSDFAKKFHLTTNETESRNYPLEKATSHLLGYVGPINSEELKQKEYK
GYKDDAVIGKKGLEKLYDKKLQHEDGYRVTIVDDNSNTIAHTLIEKKKKDGKDIQLTIDAKV
QKSIYNNMKNDYGSGTAIHPQTGELLALVSTPSYDVYPFMYGMSNEEYNKLTEDKKEPLLNK
FQITTSPGSTQKILTAMIGLNNKTLDDKTSYKIDGKGWQKDKSWGGYNVTRYEVVNGNIDLK
QAIESSDNIFFARVALELGSKKFEKGMKKLGVGEDIPSDYPFYNAQISNKNLDNEILLADSG
YGQGEILINPVQILSIYSALENNGNINAPHLLKDTKNKVWKKNIISKENINLLTDGMQQVVN
KTHKEDIYRSYANLIGKSGTAELKMKQGETGRQIGWFISYDKDNPNMMMAINVKDVQDKGMA
SYNAKISGKVYDELYENGNKKYDIDE
11110000000000000000000000000000000000000000000000000000000000
00011111100000000000000000000000000000000000000000000000000000
00000000000000000000000000000000000000000000000000000000000000
00000000000000000000000000000000000000000000000000000000000000
00000000000000000000000000000000001000000000000000000000000000
00000000000000000000000000000000000000000000000000000000000000
00000000000000000000000000000000000000000000000000000000000000
00000000000000000000000000000000000000000000000111100000000000
00000000000000000000000000000000000000000000000000000000000000
00000000000000000000000011111100000000000000000000000000000000
00000000000000000000000000
>DM_validata45
PSTIQTASPFRLASAGEISEVQGILRTAGLLGPEKRIAYLGVLDPARGAGSEAEDRRFRVFI
HDVSGARPQEVTVSVTNGTVISAVELDTAATGELPVLEEEFEVVEQLLATDERWLKALAARN
LDVSKVRVAPLSAGVFEYAEERGRRILRGLAFVQDFPEDSAWAHPVDGLVAYVDVVSKEVTR
VIDTGVFPVPAEHGNYTDPELTGPLRTTQKPISITQPEGPSFTVTGGNHIEWEKWSLDVGFD
VREGVVLHNIAFRDGDRLRPIINRASIAEMVVPYGDPSPIRSWQNYFDTGEYLVGQYANSLE
LGCDCLGDITYLSPVISDAFGNPREIRNGICMHEEDWGILAKHSDLWSGINYTRRNRRMVIS
FFTTIGNYDYGFYWYLYLDGTIEFEAKATGVVFTSAFPEGGSDNISQLAPGLGAPFHQHIFS
ARLDMAIDGFTNRVEEEDVVRQTMGPGNERGNAFSRKRTVLTRESEAVREADARTGRTWIIS
NPESKNRLNEPVGYKLHAHNQPTLLADPGSSIARRAAFATKDLWVTRYADDERYPTGDFVNQ
HSGGAGLPSYIAQDRDIDGQDIVVWHTFGLTHFPRVEDWPIMPVDTVGFKLRPEGFFDRSPV
LDVPANPSQSGSHCHGSNWSHPQFEK
11111100000000000000000000000000000000000000000000000000000000
00000000000000000000000000000000000000000000000000000000000000
00000000000000000000000000000000000000000000000000000000000000
00000000000000000000000000000000000000000000000000000000000000
00000000000000000000000000000000000000000000000000000000000000
00000000000000000000000000000000000000000000000000000000000000

000000000000000000000000000000000000000000000000000000000000
000000000000000000000000000000000000000000000000000000000000
000000000000000000000000000000000000000000000000000000000000
000000000000000000000000000000000000000000000000000000000000
0000001111111111111111111
>DM_validata46
DDLPYRFGLGKADITGEAAEVGMMGYSSLEQKTAGIHMRQWARAFVIEEAASGRRLVYVNTD
LGMIFQAVHLKVLARLKAKYPGVYDENNVMLAATHTHSGPGGFSHYAMYNLSVLGFQEKTFN
AIVDGIVRSIERAQARLQPGRLFYGSGELRNASRNRSLLSHLKNPDIAGYEDGIDPQMSVLS
FVDANGELAGAISWFPVHSTSMTNANHLISPDNKGYASYHWEHDVSRKSGFVAAFAQTNAGN
LSPNLNLKPGSGPFDNEFDNTREIGLRQFAKAYEIAGQAQEEVLGELDSRFRFVDFTRLPIR
PEFTDGQPRQLCTAAIGTSLAAGSTEDGPGPLGLEEGNNPFLSALGGLLTGVPPQELVQCQA
EKTILADTGNKKPYPWTPTVLPIQMFRIGQLELLGAPAEFTVMAGVRIRRAVQAASEAAGIR
HVVFNGYANAYASYVTTREEYAAQEYEGGSTLYGPWTQAAYQQLFVDMAVALRERLPVETSA
IAPDLSCCQMNFQTGVVADDPYIGKSFGDVLQQPRESYRIGDKVTVAFVTGHPKNDLRTEKT
FLEVVNIGKDGKQTPVTVATDNDWDTQYRWERVGISASKATISWSIPPGTEPGHYYIRHYGN
AKNFWTQKISEIGGSTRSFEVLGTTP
000000000000000000000000000000000000000000000000000000000000
000000000000000000000000000000000000000000000000000000000000
000000000000000000000000000000000000000000000000000000000000
000000000000000000000000000000000000000000000000000000000000
000000000000000000000000000000000000000000000000000000000000
000000000000000001111111111111110000000000000000000000000000
000000000000000000000000000000000000000000000000000000000000
000000000000000000000000000000000000000000000000000000000000
000000000000000000000000000000000000000000000000000000000000
000000000000000000000000000000000000000000000000000000000000
0000000000000000000000001
>DM_validata48
MKLSDLISRWIDVEPSKNAQIILRDRYFMKDLDGNYLETKWEDVARRVARVVATAELLNPSY
KKNEKLDRIKEWEDIFFRVLKARLFIPNSPTLFNAGLGVKHDLLWKPIDQMTLEDYEEIYRS
RNHLHMLSACFVVPVGDSIEEIFEAVKEYALITKVGGGVGSNFSELRPKGSFVAGTHGKASG
PVSFMHVFNSAISVVKQGSRRRGALMGILNINHPDIEEFIDAKKENTGEAVLNFFNLSVGFP
MDKKEILKLYEEDGELELSHPRSTIRKKVKIRELFRKIATNAWKSGDPGLAFLGEMNKYYPL
YPHRKINSTNPCGEIGLSDYEACNLGSIDVAKFYNNGFVDLEALQELVQIAVRFLDNVIDVN
VFPIDKITKAVKESRRLGLGIMGFADLLYKLEIPYNSQEARDFAANLMAFIALHAHRTSYEL
GKEKGNFPLLEISRYRTEDNFVPFAMGMSNYDDEIREVMKMTKEFRRNVALLTIAPTGSISN
IADTSSGLEPNFLLAYTRFVTKEDGTKEPLLYVNQVLREKLNPEILKRIEKELIEKGSLKDI
PDVPEKIKKVFVVALDIDPMDHLLMQDAFQRYVDNNISKTINMPQSATVDDVLNVYLEALRT
NVRGITVYRDGSLQTQVLTKALKT
000000000000000000000000000000000000000000000000000000000000
000000000000000000000000000000000000000000000000000000000000
000000000000000000000000000000000000000000000000000000000000
000000000000000000000000000000000000000000111111000000000000

000000000000000000000000000000000000000000000000000000000000
000000000000000000000000000000000000000000000000000000000000
000000000000000000000000000000000000000000000000000000000000
000000000000000000000000000000000000000000000000000000000000
000000000000000000011111111100000000000000000000000000000000
000000000000000000000000000000000000000000000000000000000000
000000000000011111111111
>DM_validata49
MVYYPKKYELYKADEVPTEVVETDILIIGGGFSGCGAAYEAAYWAKLGGLKVTLVEKAAVER
SGAVAQGLSAINTYIDLTGRSERQNTLEDYVRYVTLDMMGLAREDLVADYARHVDGTVHLFE
KWGLPIWKTPDGKYVREGQWQIMIHGESYKPIIAEAAKMAVGEENIYERVFIFELLKDKNDP
NAVAGAVGFSVREPKFYVFKAKAVILATGGATLLFRPRSTGEAAGRTWYAIFDTGSGYYMGL
KAGAMLTQFEHRFIPFRFKDGYGPVGAWFLFFKCKAKNAYGEEYIKTRAAELEKYKPYGAAQ
PIPTPLRNHQVMLEIMDGNQPIYMHTEEALAELAGGDKKKLKHIYEEAFEDFLDMTVSQALL
WACQNIDPQEQPSEAAPAEPYIMGSHSGEAGFWVCGPEDLMPEEYAKLFPLKYNRMTTVKGL
FAIGDCAGANPHKFSSGSFTEGRIAAKAAVRFILEQKPNPEIDDAVVEELKKKAYAPMERFM
QYKDLSTADDVNPEYILPWQGLVRLQKIMDEYAAGIATIYKTNEKMLQRALELLAFLKEDLE
KLAARDLHELMRAWELVHRVWTAEAHVRHMLFRKETRWPGYYYRTDYPELNDEEWKCFVCSK
YDAEKDEWTFEKVPYVQVIEWSF
100000000000000000000000000000000000000000000000000000000000
000000000000000000000000000000000000000000000000000000000000
000000000000000000000000000000000000000000000000000000000000
000000000000000000000000000000000000000000000000000000000000
000000000000000000000000000000000000000000000000000000000000
000000000000000000000000000000000000000000000000000000000000
000000000000000000000000000000000000000000000000000000000000
000000000000000000000000000000000000000000000000000000000000
000000000000000000000000000000000000000000000000000000000000
000000000000000000000000000000000000000000000000000000000000
00000000000000000000000
>DM_validata50
MRPVFPLILSAVLFLSCFFGARQTEASASKRAIDANQIVNRMSLDEKLGQMLMPDFRNWQKE
GESSPQALTKMNDEVASLVKKYQFGGIILFAENVKTTKQTVQLTDDYQKASPKIPLMLSIDQ
EGGIVTRLGEGTNFPGNMALGAARSRINAYQTGSIIGKELSALGINTDFSPVVDINNNPDNP
VIGVRSFSSNRELTSRLGLYTMKGLQRQDIASALKHFPGHGDTDVDSHYGLPLVSHGQERLR
EVELYPFQKAIDAGADMVMTAHVQFPAFDDTTYKSKLDGSDILVPATLSKKVMTGLLRQEMG
FNGVIVTDALNMKAIADHFGQEEAVVMAVKAGVDIALMPASVTSLKEEQKFARVIQALKEAV
KNGDIPEQQINNSVERIISLKIKRGMYPARNSDSTKEKIAKAKKIVGSKQHLKAEKKLAEKA
VTVLKNEQHTLPFKPKKGSRILIVAPYEEQTASIEQTIHDLIKRKKIKPVSLSKMNFASQVF
KTEHEKQVKEADYIITGSYVVKNDPVVNDGVIDDTISDSSKWATVFPRAVMKAALQHNKPFV
LMSLRNPYDAANFEEAKALIAVYGFKGYANGRYLQPNIPAGVMAIFGQAKPKGTLPVDIPSV
TKPGNTLYPLGYGLNIKTGRPL
111111111111111111111111000000000000000000000000000000000000
000000000000000000000000000000000000000000000000000000000000

00000000000000000000000000000000000000000000000000000000000000
00000000000000000000000000000000000000000000000000000000000000
00000000000000000000000000000000000000000000000000000000000000
00000000000000000000000000000000000000000000000000000000000000
00000000000000000000000000000000000000000000000000000000000000
00000000000000000000000000000000000000000000000000000000000000
00000000000000000000000000000000000000000000000000000000000000
00000000000000000000000000000000000000000000000000000000000000
0000000000000000000000
>DM_validata51
MQFHLNGFRPGNPLIAPASPLAPAHTEAVPSQVDVLIVGCGPAGLTLAAQLAAFPDIRTCIV
EQKEGPMELGQADGIACRTMEMFEAFEFADSILKEACWINDVTFWKPDPGQPGRIARHGRVQ
DTEDGLSEFPHVILNQARVHDHYLERMRNSPSRLEPHYARRVLDVKVDHGAADYPVTVTLER
CDAAHAGQIETVQARYVVGCDGARSNVRRAIGRQLVGDSANQAWGVMDVLAVTDFPDVRYKV
AIQSEQGNVLIIPREGGHLVRFYVEMDKLDADERVASRNITVEQLIATAQRVLHPYKLEVKN
VPWWSVYEIGQRICAKYDDVVDAVATPDSPLPRVFIAGDACHTHSPKAGQGMNFSMQDSFNL
GWKLAAVLRKQCAPELLHTYSSERQVVAQQLIDFDREWAKMFSDPAKEGGQGGVDPKEFQKY
FEQHGRFTAGVGTHYAPSLLTGQAKHQALASGFTVGMRFHSAPVVRVCDAKPVQLGHCGKAD
GRWRLYAFAAQNDLAQPESGLLALCRFLEGDAASPLRRFTPAGQDIDSIFDLRAVFPQAYTE
VALETLPALLLPPKGQLGMIDYEKVFSPDLKNAGQDIFELRGIDRQQGALVVVRPDQYVAQV
LPLGDHAALSAYFESFMRA
00000000000000000000000000000000000000000000000000000000000000
00000000000000000000000000000000000000000000000000000000000000
00000000000000000000000000000000000000000000000000000000000000
00000000000000000000000000000000000000000000000000000000000000
00000000000000000000000000011111111111000000000000000000000000
00000000000000000000000000000000000000000000000000000000000000
00000000000000000000000000000000000000001111111111111100000000
00000000000000000000000000000000000000000000000000000000000000
00000000000000000000000000000000000000000000000000000000000000
00000000000000000000000000000000000000000000000000000000000000
0000000000000000000
>DM_validata52
SSTDQGTNPADTDDLTPTTLSLAGDFPKATEEQWEREVEKVLNRGRPPEKQLTFAECLKRLT
VHTVDGIDIVPMYRPKDAPKKLGYPGVAPFTRGTTVRNGDMDAWDVRALHEDPDEKFTRKAI
LEGLERGVTSLLLRVDPDAIAPEHLDEVLSDVLLEMTKVEVFSRYDQGAAAEALVSVYERSD
KPAKDLALNLGLDPIGFAALQGTEPDLTVLGDWVRRLAKFSPDSRAVTIDANIYHNAGAGDV
AELAWALATGAEYVRALVEQGFTATEAFDTINFRVTATHDQFLTIARLRALREAWARIGEVF
GVDEDKRGARQNAITSWRELTREDPYVNILRGSIATFSASVGGAESITTLPFTQALGLPEDD
FPLRIARNTGIVLAEEVNIGRVNDPAGGSYYVESLTRSLADAAWKEFQEVEKLGGMSKAVMT
EHVTKVLDACNAERAKRLANRKQPITAVSEFPMIGARSIETKPFPAAPARKGLAWHRDSEVF
EQLMDRSTSVSERPKVFLACLGTRRDFGGREGFSSPVWHIAGIDTPQVEGGTTAEIVEAFKK
SGAQVADLCSSAKVYAQQGLEVAKALKAAGAKALYLSGAFKEFGDDAAEAEKLIDGRLFMGM
DVVDTLSSTLDILGVAK

11111111111111111000000000000000000000000000000000000000000
000000000000000000000000000000000000000000000000000000000000
000000000000000000000000000000000000000000000000000000000000
000000000000000000000000000000000000000000000000000000000000
000000000000000000000000000000000000000000000000000000000000
000000000000000000000000000000000000000000000000000000000000
000000000000000000000000000000000000000000000000000000000000
000000000000000000000000000000000000000000000000000000000000
000000000000000000000000000000000000000000000000000000000000
000000000000000000000000000000000000000000000000000000000000
00000000000000000
>DM_validata53
MATFLGLSSKQEKALVRLDKYLNLGEIAVSLVTDSATSIKVEGRQGYYQVSYKQPHQLYRAL
ALLSAALRSGQDEVQIEEEAAYEDLAYMADCSRNAVLNLSSAKKMIEVLALMGYSTFELYME
DTYEIENQPYFGYFRGRYTVAELQEIEDYAADFDMSFVPCIQTLAHLSAFVKWGIKEVQELR
DVEDILLIGEEKVYDLIEGMFQTMAHLHTRKINIGMDEAHLVGLGRYLIKHGFQNRSLLMCQ
HLERVLDIADKYGFNCQMWSDMFFKLMSADGQYDRDVEIPEETRVYLDRLKERVTLVYWDYY
QDSEEKYNRNFQNHHKISQDIAFAGGAWKWIGFTPHNHFSRLVAIEANKACRKNQVKEVIVT
GWGDNGGETSQFSVLPALQIWAELAYRNDLKKVSEHFLVSTGLDFDDFMKIDLANLLPDLPD
NLSGINPNRYVLYQDVLCPLLEQHIRPEKDKQHFASSAQQLGEISKRAGEYAYIFETQAQLN
ALLALKISITSGIQKAYRNGDKEHLSALAEKDFPQLYQMVEDFSDQFSRQWQQENKIFGLDT
IDIRFGGLLKRIKRAQERLEQFISGQIDCVEELEQEILPFNDFYKDQGLTATTANQWHLIAT
ASTIYTT
100000000000000000000000000000000000000000000000000000000000
000000000000000000000000000000000000000000000000000000000000
000000000000000000000000000000000000000000000000000000000000
000000000000000000000000000000000000000000000000000000000000
000000000000000000000000001111111111100000000000000000000000
000000000000000000000000000000000000000000000000000000000000
000000000000000000000000000000000000000000000000000000000000
000000000000000000000000000000000000000000000000000000000000
000000000000000000000000000000000000000000000000000000000000
000000000000000000000000000000000000000000000000000000000000
0000000
>DM_validata54
SELNEKLATAWEGFTKGDWQNEVNVRDFIQKNYTPYEGDESFLAGATEATTTLWDKVMEGVK
LENRTHAPVDFDTAVASTITSHDAGYINKQLEKIVGLQTEAPLKRALIPFGGIKMIEGSCKA
YNRELDPMIKKIFTEYRKTHNQGVFDVYTPDILRCRKSGVLTGLPDAYGRGRIIGDYRRVAL
YGIDYLMKDKLAQFTSLQADLENGVNLEQTIRLREEIAEQHRALGQMKEMAAKYGYDISGPA
TNAQEAIQWTYFGYLAAVKSQNGAAMSFGRTSTFLDVYIERDLKAGKITEQEAQEMVDHLVM
KLRMVRFLRTPEYDELFSGDPIWATESIGGMGLDGRTLVTKNSFRFLNTLYTMGPSPEPNMT
ILWSEKLPLNFKKFAAKVSIDTSSLQYENDDLMRPDFNNDDYAIACCVSPMIVGKQMQFFGA
RANLAKTMLYAINGGVDEKLKMQVGPKSEPIKGDVLNYDEVMERMDHFMDWLAKQYITALNI
IHYMHDKYSYEASLMALHDRDVIRTMACGIAGLSVAADSLSAIKYAKVKPIRDEDGLAIDFE

IEGEYPQFGNNDPRVDDLAVDLVERFMKKIQKLHTYRDAIPTQSVLTITSNVVYGKKTGNTP
DGRR
111000000000000000000000000000000000000000000000000000000000
000000000000000000000000000000000000000000000000000000000000
000000000000000000000000000000000000000000000000000000000000
000000000000000000000000000000000000000000000000000000000000
000000000000000000000000000000000000000000000000000000000000
000000000000000000000000000000000000000000000000000000000000
000000000000000000000000000000000000000000000000000000000000
000000000000000000000000000000000000000000000000000000000000
000000000000000000000000000000000000000000000000000000000000
000000000000000000000000000000000000000000000000000000111111
1111
>DM_validata55
LEEDKIKGMFNPKIWDKTFQDGLKKEIEDSQPYNWGTIHELVNDDLLRAVRKEIETEIHFTK
KETDIYRVNQSGDLANLSGLDWDDLSRLPNLFKLRQILYSKQYRDFFGYVTKAGKLSGSKTD
MSINTYTKGCHLLTHDDVIGSRRISFILYLPDPDRKWKSHYGGGLRLFPSILPNVPHSDPSA
KLVPQFNQIAFFKVLPGFSFHDVEEVKVDKHRLSIQGWYHIPQVGEEGYIPGEEEAWVRNNT
STLAQIESNVLEDFEFPKDERNILSFHEVKHFEKMLKGDAGAKTDNTPKESMTSVISDSVKL
SEAEFTYLSQYISPEHLSSKGIEKLQKQFVENSSLQIESFLNDDKSELLKKVIKQKELEQEC
PYHSKDVKAPWKTAIPPHKARYLYIDGKEYRNFQTEADILEALNNNDLPNFQFTKDAIKIIS
DASGNSRENNFDAELALIDLAVFHKSTIFKKYLALLTSLCPVSEQILIRRFRPGMDFTLATK
CRFNELLKSNPDIIDAVLEGTLCLTPSAGWESGELGGYELYMMDDDEDNKQYLKEDVEDASV
YRADDSGDSVLINDPPAWNTFNLVLRDESVLEFVKYVSWSAKSSRWDVKMKWDVKSCDEDGQ
EDEA
111000000000000000000000000000000000000000000000000000000000
000000000000000000000000000000000000000000000000000000000000
000000000000000000000000000000000000000000000000000000000000
000000000000000000000000000000000000000000000000000000000000
011111110000000000000000000000000000111111111111111111111000
000000000000000000000000000000000000000000000000000000000000
000000000000000000000000000000000000000000000000000000000000
000000000000000000000000000000000000000000000000000000000000
000000000000000000000000000000000000000000011111111111111111
111111100000000000000000000000000000000000000000000000011111
1111
>DM_validata57
MSSISLKEIIPPQPSTQRNFTTHLSYDPTTNAIAYPCGKSAFVRCLDDGDSKVPPVVQFTGH
GSSVVTTVKFSPIKGSQYLCSGDESGKVIVWGWTFDKESNSVEVNVKSEFQVLAGPISDISW
DFEGRRLCVVGEGRDNFGVFISWDSGNSLGEVSGHSQRINACHLKQSRPMRSMTVGDDGSVV
FYQGPPFKFSASDRTHHKQGSFVRDVEFSPDSGEFVITVGSDRKISCFDGKSGEFLKYIEDD
QEPVQGGIFALSWLDSQKFATVGADATIRVWDVTTSKCVQKWTLDKQQLGNQQVGVVATGNG
RIISLSLDGTLNFYELGHDEVLKTISGHNKGITALTVNPLISGSYDGRIMEWSSSSMHQDHS
NLIVSLDNSKAQEYSSISWDDTLKVNGITKHEFGSQPKVASANNDGFTAVLTNDDDLLILQS

FTGDIIKSVRLNSPGSAVSLSQNYVAVGLEEGNTIQVFKLSDLEVSFDLKTPLRAKPSYISI
SPSETYIAAGDVMGKILLYDLQSREVKTSRWAFRTSKINAISWKPAEKGANEEEIEEDLVAT
GSLDTNIFIYSVKRPMKIIKALNAHKDGVNNLLWETPSTLVSSGADACIKRWNVVLE
00000000000000000000000000000000000000000000000000000000000000
00000000000000000000000000000000000000000000000000000000000000
00000000000000000000000000000000000000000000000000000000000000
00000000000000000000000000000000000000000000000000000000000000
00000000000000000000000000000000000000000000000000000000000000
00000000000000000000000000000000000000000000000000000000000000
00000000000000000000000000000000000000000000000000000000000000
00000000000000000000000000000000000000000000000000000000000000
00000000000000000000000000000000000000000000000111111000000000
000000000000000000000000000000000000000000000000000000000
>DM_validata59
MASTANMISQLKKLSIAEPAVAKDSHPDVNIVDLMRNYISQELSKISGVDSSLIFPALEWTN
TMERGDLLIPIPRLRIKGANPKDLAVQWAEKFPCGDFLEKVEANGPFIQFFFNPQFLAKLVI
PDILTRKEDYGSCKLVENKKVIIEFSSPNIAKPFHAGHLRSTIIGGFLANLYEKLGWEVIRM
NYLGDWGKQFGLLAVGFERYGNEEALVKDPIHHLFDVYVRINKDIEEEGDSIPLEQSTNGKA
REYFKRMEDGDEEALKIWKRFREFSIEKYIDTYARLNIKYDVYSGESQVSKESMLKAIDLFK
EKGLTHEDKGAVLIDLTKFNKKLGKAIVQKSDGTTLYLTRDVGAAMDRYEKYHFDKMIYVIA
SQQDLHAAQFFEILKQMGFEWAKDLQHVNFGMVQGMSTRKGTVVFLDNILEETKEKMHEVMK
KNENKYAQIEHPEEVADLVGISAVMIQDMQGKRINNYEFKWERMLSFEGDTGPYLQYAHSRL
RSVERNASGITQEKWINADFSLLKEPAAKLLIRLLGQYPDVLRNAIKTHEPTTVVTYLFKLT
HQVSSCYDVLWVAGQTEELATARLALYGAARQVLYNGMRLLGLTPVERM
11110000000000000000000000000000000000000000000000000000000000
00000000000000000000000000000000000000000000000000000000000000
00000000000000000000000000000000000000000000000000000000000000
00000000000000000000000000000000000000000000000000000000000000
00000000000000000000000000000000000000000000000000000000000000
00000000000000000000000000000000000000000000000000000000000000
00000000000000000000000000000000000000000000000000000000000000
00000000000000000000000000000000000000000000000000000000000000
00000000000000000000000000000000000000000000000000000000000000
000000000000000000000000000000000000000000000000
>DM_validata60
KSVAHHEDVYSHNLPPMDEKEMALYKLYRPERVTPKKRSAELLKEPRLNKGMGFSLYERQYL
GLHGLLPPAFMTQEQQAYRVITKLREQPNDLARYIQLDGLQDRNEKLFYRVVCDHVKELMPI
VYTPTVGLACQNFGYIYRKPKGLYITINDNSVSKIYQILSNWHEEDVRAIVVTDGERILGLG
DLGAYGIGIPVGKLALYVALGGVQPKWCLPVLLDVGTNNMDLLNDPFYIGLRHKRVRGKDYD
TLLDNFMKACTKKYGQKTLIQFEDFANPNAFRLLDKYQDKYTMFNDDIQGTASVIVAGLLTC
TRVTKKLVSQEKYLFFGAGAASTGIAEMIVHQMQNEGISKEEACNRIYLMDIDGLVTKNRKE
MNPRHVQFAKDMPETTSILEVIRAARPGALIGASTVRGAFNEEVIRAMAEINERPIIFALSN
PTSKAECTAEEAYTFTNGAALYASGSPFPNFELNGHTYKPGQGNNAYIFPGVALGTILFQIR
HVDNDLFLLAAKKVASCVTEDSLKVGRVYPQLKEIREISIQIAVEMAKYCYKNGTANLYPQP

EDLEKYVRAQVYNTEYEELINATYDWPEQDMRHGFPVPVVRHDSMDG
000000000000000000000000000000000000000000000000000000000000
000000000000000000000000000000000000000000000000000000000000
000000000000000000000000000000000000000000000000000000000000
000000000000000000000000000000000000000000000000000000000000
000000000000000000000000000000000000000000000000000000000000
000000000000000000000000000000000000000000000000000000000000
000000000000000000000000000000000000000000000000000000000000
000000000000000000000000000000000000000000000000000000000000
000000000000000000000000000000000000000000000000000000000000
00000000000000000000000000000000000111111111111
>DM_validata61
DYVLYKDATKPVEDRVADLLGRMTLAEKIGQMTQIERLVATPDVLRDNFIGSLLSGGGSVPR
KGATAKEWQDMVDGFQKACMSTRLGIPMIYGIDAVHGQNNVYGATIFPHNVGLGATRDPYLV
KRIGEATALEVRATGIQYAFAPCIAVCRDPRWGRCYESYSEDRRIVQSMTELIPGLQGDVPK
DFTSGMPFVAGKNKVAACAKHFVGDGGTVDGINENNTIINREGLMNIHMPAYKNAMDKGVST
VMISYSSWNGVKMHANQDLVTGYLKDTLKFKGFVISDWEGIDRITTPAGSDYSYSVKASILA
GLDMIMVPNKYQQFISILTGHVNGGVIPMSRIDDAVTRILRVKFTMGLFENPYADPAMAEQL
GKQEHRDLAREAARKSLVLLKNGKTSTDAPLLPLPKKAPKILVAGSHADNLGYQCGGWTIEW
QGDTGRTTVGTTILEAVKAAVDPSTVVVFAENPDAEFVKSGGFSYAIVAVGEHPYTETKGDN
LNLTIPEPGLSTVQAVCGGVRCATVLISGRPVVVQPLLAASDALVAAWLPGSEGQGVTDALF
GDFGFTGRLPRTWFKSVDQLPMNVGDAHYDPLFRLGYGLTTNATKKY
000000000000000000000000000000000000000000000000000000000000
000000000000000000000000000000000000000000000000000000000000
000000000000000000000000000000000000000000000000000000000000
000000000000000000000000000000000000000000000000000000000000
000000000000000000000000000000000000000000000000000000000000
000000000000000000000000000000000000000000000000000000000000
000000000000000000000000000000000000000000000000000000000000
000000000000000000000000000000000000000000000000000000000000
000000000000000000000000000000000000000000000000000000000000
00000000000000000000000000000000000000000000111
>DM_validata62
MTIEKEIEGLIHKTNKDLLNENANKDSRVFPTQRDLMAGIVSKHIAKNMVPSFIMKAHESGI
IHVHDIDYSPALPFTNCCLVDLKGMLENGFKLGNAQIETPKSIGVATAIMAQITAQVASHQY
GGTTFANVDKVLSPYVKRTYAKHIEDAEKWQIADALNYAQSKTEKDVYDAFQAYEYEVNTLF
SSNGQTPFVTITFGTGTDWTERMIQKAILKNRIKGLGRDGITPIFPKLVMFVEEGVNLYKDD
PNYDIKQLALECASKRMYPDIISAKNNKAITGSSVPVSPMGCRSFLSVWKDSTGNEILDGRN
NLGVVTLNLPRIALDSYIGTQFNEQKFVELFNERMDLCFEALMCRISSLKGVKATVAPILYQ
EGAFGVRLKPDDDIIELFKNGRSSVSLGYIGIHELNILVGRDIGREILTKMNAHLKQWTERT
GFAFSLYSTPAENLCYRFCKLDTEKYGSVKDVTDKGWYTNSFHVSVEENITPFEKISREAPY
HFIATGGHISYVELPDMKNNLKGLEAVWDYAAQHLDYFGVNMPVDKCFTCGSTHEMTPTENG
FVCSICGETDPKKMNTIRRTCAYLGNPNERGFNLGKNKEIMHRVKHQ
111111111111111111111111100000000000000000000000000000000000

00000000000000000000000000000000000000000000000000000000000000
00000000000000000000000000000000000000000000000000000000000000
00000000000000000000000000000000000000000000000000000000000000
00000000000000000000000000000000000000000000000000000000000000
00000000000000000000000000000000000000000000000000000000000000
00000000000000000000000000000000000000000000000000000000000000
00000000000000000000000000000000000000000000000000000000000000
00000000000000000000000000000000000000000000000111111111111111
11111111111110000000000000000111111111111111111
>DM_validata63
DPDQFRAIIESPEGAGHVGYQYRRNTGSTMRMVSDVLDERVSLWDFHCDPSGNVIQPGPNVD
SRQYLQAAIDYVSSNGGGTITIPAGYTWYLGSYGVGGIAGHSGIIQLRSNVNLNIEGRIHLS
PFFDLKPFQVFVGFDNGDPASSGNLENCHIYGHGVVDFGGYEFGASSQLRNGVAFGRSYNCS
VTGITFQNGDVTWAITLGWNGYGSNCYVRKCRFINLVNSSVNADHSTVYVNCPYSGVESCYF
SMSSSFARNIACSVELHQHDTFYRGSTVNGYCRGAYVVMHAAEAAGAGSYAYNMQVENNIAV
IYGQFVILGSDVTATVSGHLNDVIVSGNIVSIGERAAFSAPFGAFIDIGPDNSGASNVQDIQ
RVLVTGNSFYAPANITDSAAITLRANLNGCTFIANNFDCRYMVYNAPGTTSPVVQNLVWDKS
NVIGGTHANQRAGQNLFDMQFASVVNSTIEVQLSCEDLSMFSCILFPASCQLSYSKITVDSA
WTKSMSNTAVFEGNQQAGANVYVSYPATVNLTSYNTQGAVPFFSTDTNYAWVTSAYSLSINE
NLDFSPPATYTNKANGQLVGVGYNEIGGVRSVSVRLMLQRQV
11000000000000000000000000000000000000000000000000000000000000
00000000000000000000000000000000000000000000000000000000000000
00000000000000000000000000000000000000000000000000000000000000
00000000000000000000000000000000000000000000000000000000000000
00000000000000000000000000000000000000000000000000000000000000
00000000000000000000000000000000000000000000000000000000000000
00000000000000000000000000000000000000000000000000000000000000
00000000000000000000000000000000000000000000000000000000000000
00000000000000000000000000000000000000000000000000000000000000
000000000000000000000000000000000000000000
>DM_validata64
GSMGKALVIVESPAKAKTINKYLGSDYVVKSSVGHIRDLPTSGSAAKKSADSTSTKTAKKPK
KDERGALVNRMGVDPWHNWEAHYEVLPGKEKVVSELKQLAEKADHIYLATDLDREGEAIAWH
LREVIGGDDARYSRVVFNEITKNAIRQAFNKPGELNIDRVNAQQARRFMDRVVGYMVSPLLW
KKIARGLSAGRVQSVAVRLVVEREREIKAFVPEEFWEVDASTTTPSGEALALQVTHQNDKPF
RPVNKEQTQAAVSLLEKARYSVLEREDKPTTSKPGAPFITSTLQQAASTRLGFGVKKTMMMA
QRLYEAGYITYMRTDSTNLSQDAVNMVRGYISDNFGKKYLPESPNQYASKENSQEAHEAIRP
SDVNVMAESLKDMEADAQKLYQLIWRQFVACQMTPAKYDSTTLTVGAGDFRLKARGRILRFD
GWTKVMPALRKGDEDRILPAVNKGDALTLVELTPAQHFTKPPARFSEASLVKELEKRGIGRP
STYASIISTIQDRGYVRVENRRFYAEKMGEIVTDRLEENFRELMNYDFTAQMENSLDQVANH
EAEWKAVLDHFFSDFTQQLDKAEKDPEEGGMRPNQMVLTSI
11100000000000000000000000000000000000001111111111111111111111
11111000000000000000000000000000000000000000000000000000000000
00000000000000000000000000000000000000000000000000000000000000

0000000000000000000000000000000000000000000000000000000000000000
0000000000000000000000000000000000000000000000000000000000000000
0000000000000000000000000000000000000000000000000000000000000000
0000000000000000000000000000000000000000000000000000000000000000
0000001111111111000000000000000000000000000000000000000000000000
0000000000000000000000000000000000000000000000000000000000000000
000000000000000011111111111111111111111111
>DM_validata65
SLPPLPGPDKKVPGMDIKYDVVIVGSGPIGCTYARELVGAGYKVAMFDIGEIDSGLKIGAHK
KNTVEYQKNIDKFVNVIQGQLMSVSVPVNTLVVDTLSPTSWQASTFFVRNGSNPEQDPLRNL
SGQAVTRVVGGMSTHWTCATPRFDREQRPLLVKDDADADDAEWDRLYTKAESYFQTGTDQFK
ESIRHNLVLNKLAEEYKGQRDFQQIPLAATRRSPTFVEWSSANTVFDLQNRPNTDAPEERFN
LFPAVACERVVRNALNSEIESLHIHDLISGDRFEIKADVYVLTAGAVHNTQLLVNSGFGQLG
RPNPTNPPELLPSLGSYITEQSLVFCQTVMSTELIDSVKSDMTIRGTPGELTYSVTYTPGAS
TNKHPDWWNEKVKNHMMQHQEDPLPIPFEDPEPQVTTLFQPSHPWHTQIHRDAFSYGAVQQS
IDSRLIVDWRFFGRTEPKEENKLWFSDKITDAYNMPQPTFDFRFPAGRTSKEAEDMMTDMCV
MSAKIGGFLPGSLPQFMEPGLVLHLGGTHRMGFDEKEDNCCVNTDSRVFGFKNLFLGGCGNI
PTAYGANPTLTAMSLAIKSCEYIKQNFTPSPFTSEAQ
1111111111111100000000000000000000000000000000000000000000000000
0000000000000000000000000000000000000000000000000000000000000000
0000000000000000000000000000000000000000000000000000000000000000
0000000000000000000000000000000000000000000000000000000000000000
0000000000000000000000000000000000000000000000000000000000000000
0000000000000000000000000000000000000000000000000000000000000000
0000000000000000000000000000000000000000000000000000000000000000
0000000000000000000000000000000000000000000000000000000000000000
0000000000000000000000000000000000000000000000000000000000000000
00000000000000000000000000000000001111
>DM_validata66
MQKRIALSFPEEVLEHVFSFIQLDKDRNSVSLVCKSWYEIERWCRRKVFIGNCYAVSPATVI
RRFPKVRSVELKGKPHFADFNLVPDGWGGYVYPWIEAMSSSYTWLEEIRLKRMVVTDDCLEL
IAKSFKNFKVLVLSSCEGFSTDGLAAIAATCRNLKELDLRESDVDDVSGHWLSHFPDTYTSL
VSLNISCLASEVSFSALERLVTRCPNLKSLKLNRAVPLEKLATLLQRAPQLEELGTGGYTAE
VRPDVYSGLSVALSGCKELRCLSGFWDAVPAYLPAVYSVCSRLTTLNLSYATVQSYDLVKLL
CQCPKLQRLWVLDYIEDAGLEVLASTCKDLRELRVFPSEPFVMEPNVALTEQGLVSVSMGCP
KLESVLYFCRQMTNAALITIARNRPNMTRFRLCIIEPKAPDYLTLEPLDIGFGAIVEHCKDL
RRLSLSGLLTDKVFEYIGTYAKKMEMLSVAFAGDSDLGMHHVLSGCDSLRKLEIRDCPFGDK
ALLANASKLETMRSLWMSSCSVSFGACKLLGQKMPKLNVEVIDERGAPDSRPESCPVERVFI
YRTVAGPRFDMPGFVWNMDQDSTMRFSRQIITTNGL
1111111110000000000000000000000000000000000000000000000000000000
0000000000000000000000000000000000000000000000000000000000000000
0000000000000000000000000000000000000000000000000000000000000000
0000000000000000000000000000000000000000000000000000000000000000
0000000000000000000000000000000000000000000000000000000000000000

00000000000000000000000000000000000000000000000000000000000000
00000000000000000000000000000000000000000000000000000000000000
00000000000000000000000000000000000000000000000000000000000000
00000000000000000000000000000000000000000000000000000000000000
000000000000000000111111111111111111
>DM_validata67
MNMYSMPGYFQNMPTIGKELVNPNPENEQEIKAVESDIHESIKKALDAGITSEEKLNERGQL
SAMQRINALIDPGTWCPLNSLFNPENNKFGTTNIVNGLGRVDGKWVYIVASDNKKMAGAWVP
GQAENLIRCSDAAKMMHLPLIYLLNCSGVEFPNQDKVYPNRRGGGTPFFRNSELNQLGIPVI
VGIYGTNPAGGGYHSISPTILIAHQDANMAVGGAGILSGMNPKGYIDDEAAEQIIAAQIENS
KLKVPAPGSVPIHYDETGFFREVYQNDLGVIDGIKKYISYLPAYNLEFFRVDTPKAPQLPAE
DLYSIIPMNQKRPYDIYEVIARLFDNSEFSEYKKGYGPEMVTGLAKVNGLLVGVIANVQGLL
MNYPEYKQNSVGIGGKLYRQGLIKMNEFVTLCARDRIPLIWLQDTTGIDVGDEAEKAELLGL
GQSLIYSIENSKLPSLEITIRKASAAAHYVLGGPQGNNTNVFSIGTGACEYYVMPGETAANA
MYSRKLVKAKKAGEDLQPIIGKMNDMIQMYTDKSRPKYCTEKGMVDEIVDMTEVRPYIQAFT
EAAYQNPQSICPMHQMLTPRSTREFETFGK
11000000000000000000000000000000000000000000000000000000000000
00000000000000000000000000000000000000000000000000000000000000
00000000000000000000000000000000000000000000000000000000000000
00000000000000000000000000000000000111111111111110000000000000
00000000000000000000000000000000000000000000000000000000000000
00000000000000000000000000000000000000000000000000000000000000
00000000000000000000000000000000000000000000000000000000000000
00000000000000000000000000000000000000000000000000000000000000
00000000011111100000000000000000000000000000000000000000000000
000000000000000000000000000001
>DM_validata70
AGFKPAPPAGQLGAVIVDPYGNAPLTALVDLDSHVISDVKVTVHGKGEKGVEISYPVGQESL
KTYDGVPIFGLYQKFANKVTVEWKENGKVMKDDYVVHTSAIVNNYMDNRSISDLQQTKVIKV
APGFEDRLYLVNTHTFTAQGSDLHWHGEKDKNAGILDAGPATGALPFDIAPFTFIVDTEGEY
RWWLDQDTFYDGRDRDINKRGYLMGIRETPRGTFTAVQGQHWYEFDMMGQVLEDHKLPRGFA
DATHESIETPNGTVLLRVGKSNYRRDDGVHVTTIRDHILEVDKSGRVVDVWDLTKILDPKRD
ALLGALDAGAVCVNVDLAHAGQQAKLEPDTPFGDALGVGPGRNWAHVNSIAYDAKDDSIILS
SRHQGVVKIGRDKQVKWILAPSKGWEKPLASKLLKPVDANGKPITCNENGLCENSDFDFTYT
QHTAWISSKGTLTIFDNGDGRHLEQPALPTMKYSRFVEYKIDEKKGTVQQVWEYGKERGYDF
YSPITSIIEYQADRNTMFGFGGSIHLFDVGQPTVGKLNEIDYKTKEVKVEIDVLSDKPNQTH
YRALLVRPQQMFK
00000000000000000000000000000000000000000000000000000000000000
00000000000000000000000000000000000000000000000000000000000000
00000000000000000000000000000000000000000000000000000000000000
00000000000000000000000000000000000000000000000000000000000000
00000000000000000000000000000000000000000000000000000000000000
00000000001111111000000000000000000000000000000000000000000000
00000000000000000000000000000000000000000000000000000000000000

000000000000000000000000000000000000000000000000000000000000
000000000000000000000000000000000000000000000000000000000000
0000000000000
>DM_validata71
MSGCRAQNAPGGIPVTPKSSYSGPIVVDPVTRIEGHLRIEVEVENGKVKNAYSSSTLFRGLE
IILKGRDPRDAQHFTQRTCGVCTYTHALASTRCVDNAVGVHIPKNATYIRNLVLGAQYLHDH
IVHFYHLHALDFVDVTAALKADPAKAAKVASSISPRKTTAADLKAVQDKLKTFVETGQLGPF
TNAYFLGGHPAYYLDPETNLIATAHYLEALRLQVKAARAMAVFGAKNPHTQFTVVGGVTCYD
ALTPQRIAEFEALWKETKAFVDEVYIPDLLVVAAAYKDWTQYGGTDNFITFGEFPKDEYDLN
SRFFKPGVVFKRDFKNIKPFDKMQIEEHVRHSWYEGAEARHPWKGQTQPKYTDLHGDDRYSW
MKAPRYMGEPMETGPLAQVLIAYSQGHPKVKAVTDAVLAKLGVGPEALFSTLGRTAARGIET
AVIAEYVGVMLQEYKDNIAKGDNVICAPWEMPKQAEGVGFVNAPRGGLSHWIRIEDGKIGNF
QLVVPSTWTLGPRCDKNNVSPVEASLIGTPVADAKRPVEILRTVHSFDPCIACGVHVIDGHT
NEVHKFRIL
111111111111111111000000000000000000000000000000000000000000
000000000000000000000000000000000000000000000000000000000000
000000000000000000000000000000000000000000000000000000000000
000000000000000000000000000000000000000000000000000000000000
000000000000000000000000000000000000000000000000000000000000
000000000000000000000000000000000000000000000000000000000000
000000000000000000000000000000000000000000000000000000000000
000000000000000000000000000000000000000000000000000000000000
000000000000000000000000000000000000000000000000000000111111
111111111
>DM_validata72
MVRSGNKAAVVLCMDVGFTMSNSIPGIESPFEQAKKVITMFVQRQVFAENKDEIALVLFGTD
GTDNPLSGGDQYQNITVHRHLMLPDFDLLEDIESKIQPGSQQADFLDALIVSMDVIQHETIG
KKFEKRHIEIFTDLSSRFSKSQLDIIIHSLKKCDISLQFFLPFSLGKEDGSGDRGDGPFRLG
GHGPSFPLKGITEQQKEGLEIVKMVMISLEGEDGLDEIYSFSESLRKLCVFKKIERHSIHWP
CRLTIGSNLSIRIAAYKSILQERVKKTWTVVDAKTLKKEDIQKETVYCLNDDDETEVLKEDI
IQGFRYGSDIVPFSKVDEEQMKYKSEGKCFSVLGFCKSSQVQRRFFMGNQVLKVFAARDDEA
AAVALSSLIHALDDLDMVAIVRYAYDKRANPQVGVAFPHIKHNYECLVYVQLPFMEDLRQYM
FSSLKNSKKYAPTEAQLNAVDALIDSMSLAKKDEKTDTLEDLFPTTKIPNPRFQRLFQCLLH
RALHPREPLPPIQQHIWNMLNPPAEVTTKSQIPLSKIKTLFPLIEAKKKDQVTAQEIFQDNH
EDGPTAK
111110000000000000000000000000000000000000000000000000000000
000000000000000000000000000000000000000000000000000000000000
000000000000000000000000000000000000000000011111111111100000
000000000000000000000000000000000000000000000000000000000000
000000000000000000000000000000000000000000000000000000000000
000000000000011100000000000000000000000000000000000000000000
000000000000000000000000000000000000000000000000000000000000
000000000000000000000000000000000000000000000000000000000000
000000000000000000000000000000000000000000001111111111111111

1111111
>DM_validata74
MAVRASEISRVYEAYPEKKATLYFLVLGFLALIVGSLFGPFQALNYGNVDAYPLLKRLLPFV
QSYYQGLTLHGVLNAIVFTQLFAQAIMVYLPARELNMRPNMGLMWLSWWMAFIGLVVAALPL
LANEATVLYTFYPPLKGHWAFYLGASVFVLSTWVSIYIVLDLWRRWKAANPGKVTPLVTYMA
VVFWLMWFLASLGLVLEAVLFLLPWSFGLVEGVDPLVARTLFWWTGHPIVYFWLLPAYAIIY
TILPKQAGGKLVSDPMARLAFLLFLLLSTPVGFHHQFADPGIDPTWKMIHSVLTLFVAVPSL
MTAFTVAASLEFAGRLRGGRGLFGWIRALPWDNPAFVAPVLGLLGFIPGGAGGIVNASFTLD
YVVHNTAWVPGHFHLQVASLVTLTAMGSLYWLLPNLTGKPISDAQRRLGLAVVWLWFLGMMI
MAVGLHWAGLLNVPRRAYIAQVPDAYPHAAVPMVFNVLAGIVLLVALLLFIYGLFSVLLSRE
RKPELAEAPLPFAEVISGPEDRRLVLAMDRIGFWFAVAAILVVLAYGPTLVQLFGHLNPVPG
WRLW
11111111111110000000000000000000000000000000000000000000000000
00000000000000000000000000000000000000000000000000000000000000
00000000000000000000000000000000000000000000000000000000000000
00000000000000000000000000000000000000000000000000000000000000
00000000000000000000000000000000000000000000000000000000000000
00000000000000000000000000000000000000000000000000000000000000
00000000000000000000000000000000000000000000000000000000000000
00000000000000000000000000000000000000000000000000000000000001
11110000000000000000000000000000000000000000000000000000000000
0000
>DM_validata75
MSKTQEFRPLTLPPKLSLSDFNEFIQDIIRIVGSENVEVISSKDQIVDGSYMKPTHTHDPTH
VMDQDYFLASAIVAPRNVADVQSIVGLANKFSFPLWPISIGRNSGYGGAAPRVSGSVVLDMG
KNMNRVLEVNVEGAYCVVEPGVTYHDLHNYLEANNLRDKLWLDVPDLGGGSVLGNAVERGVG
YTPYGDHWMMHSGMEVVLANGELLRTGMGALPDPKRPETMGLKPEDQPWSKIAHLFPYGFGP
YIDGLFSQSNMGIVTKIGIWLMPNPGGYQSYLITLPKDGDLKQAVDIIRPLRLGMALQNVPT
IRHILLDAAVLGDKRSYSSRTEPLSDEELDKIAKQLNLGRWNFYGALYGPEPIRRVLWETIK
DAFSAIPGVKFYFPEDTPENSVLRVRDKTMQGIPTYDELKWIDWLPNGAHLFFSPIAKVSGE
DAMMQYAVTKKRCQEAGLDFIGTFTVGMREMHHIVCIVFNKKDLIQKRKVQWLMRTLIDDCA
ANGWGEYRTHLAFMDQIMETYNWNNSSFLRFNEVLKNAVDPNGIIAPGKSGVWPSQYSHVTW
KL
11111000000000000000000000000000000000000111111111000000000000
00000000000000000000000000000000000000000000000000000000000000
00000000000000000000000000000000000000000000000000000000000000
00000000000000000000000000000000000000000000000000000000000000
00000000000000000000000000000000000000000000000000000000000000
00000000000000000000000000000000000000000000000000000000000000
00000000000000000000000000000000000000000000000000000000000000
00000000000000000000000000000000000000000000000000000000000000
00000000000000000000000000000000000000000000000000000000000000
00
>DM_validata76

MTDNNKYRDVEIRAPRGNKLTAKSWLTEAPLRMLMNNLDPQVAENPKELVVYGGIGRAARNW
ECYDKIVETLTRLEDDETLLVQSGKPVGVFKTHSNAPRVLIANSNLVPHWANWEHFNELDAK
GLAMYGQMTAGSWIYIGSQGIVQGTYETFVEAGRQHYGGSLKGKWVLTAGLGGMGGAQPLAA
TLAGACSLNIESQQSRIDFRLETRYVDEQATDLDDALVRIAKYTAEGKAISIALHGNAAEIL
PELVKRGVRPDMVTDQTSAHDPLNGYLPAGWTWEQYRDRAQTEPAAVVKAAKQSMAVHVQAM
LDFQKQGVPTFDYGNNIRQMAKEEGVADAFDFPGFVPAYIRPLFCRGVGPFRWAALSGEAED
IYKTDAKVKELIPDDAHLHRWLDMARERISFQGLPARICWVGLGLRAKLGLAFNEMVRSGEL
SAPVVIGRDHLDSGSVSSPNAETEAMRDGSDAVSDWPLLNALLNTAGGATWVSLHHGGGVGM
GFSQHSGMVIVCDGTDEAAERIARVLTNDPGTGVMRHADAGYDIAIDCAKEQGLDLPMITG
111100000000000000000000000000000000000000000000000000000000
000000000000000000000000000000000000000000000000000000000000
000000000000000000000000000000000000000000000000000000000000
000000000000000000000000000000000000000000000000000000000000
000000000000000000000000000000000000000000000000000000000000
000000000000000000000000000000000000000000000000000000000000
000000000000000000000000000000000000000000000000000000000000
000000000000000000000000000000000000000000000000000000000000
00000000000000000000000000000000000000000000000000000000000
>DM_validata77
MSEAEARPTNFIRQIIDEDLASGKHTTVHTRFPPEPNGYLHIGHAKSICLNFGIAQDYKGQC
NLRFDDTNPVKEDIEYVESIKNDVEWLGFHWSGNVRYSSDYFDQLHAYAIELINKGLAYVDE
LTPEQIREYRGTLTQPGKNSPYRDRSVEENLALFEKMRAGGFEEGKACLRAKIDMASPFIVM
RDPVLYRIKFAEHHQTGNKWCIYPMYDFTHCISDALEGITHSLCTLEFQDNRRLYDWVLDNI
TIPVHPRQYEFSRLNLEYTVMSKRKLNLLVTDKHVEGWDDPRMPTISGLRRRGYTAASIREF
CKRIGVTKQDNTIEMASLESCIREDLNENAPRAMAVIDPVKLVIENYQGEGEMVTMPNHPNK
PEMGSRQVPFSGEIWIDRADFREEANKQYKRLVLGKEVRLRNAYVIKAERVEKDAEGNITTI
FCTYDADTLSKDPADGRKVKGVIHWVSAAHALPVEIRLYDRLFSVPNPGAADDFLSVINPES
LVIKQGFAEPSLKDAVAGKAFQFEREGYFCLDSRHSTAEKPVFNRTVGLRDTWAKVGE
111111110000000000000000000000000000000000000000000000000000
000000000000000000000000000000000000000000000000000000000000
000000000000000000000000000000000000000000000000000000000000
000000000000000000000000000000000000000000000000000000000000
000000000000000000000000000000000000000000000000000000000000
000000000000000000000000000000000000000000000000000000000000
000000000000000000000000000000000000000000000000000000000000
000000000111111111110000000000000000000000000000000000000000
0000000000000000000000000000000000000000000000000000111111
>DM_validata78
MALRVAAFDLDGVLALPSIAGAFRRSEEALALPRDFLLGAYQTEFPEGPTEQLMKGKITFSQ
WVPLMDESYRKSSKACGANLPENFSISQIFSQAMAARSINRPMLQAAIALKKKGFTTCIVTN
NWLDDGDKRDSLAQMMCELSQHFDFLIESCQVGMIKPEPQIYNFLLDTLKAKPNEVVFLDDF
GSNLKPARDMGMVTILVHNTASALRELEKVTGTQFPEAPLPVPCNPNDVSHGYVTVKPGIRL
HFVEMGSGPALCLCHGFPESWFSWRYQIPALAQAGFRVLAIDMKGYGDSSSPPEIEEYAMEL
LCKEMVTFLDKLGIPQAVFIGHDWAGVMVWNMALFYPERVRAVASLNTPFMPPDPDVSPMKV

IRSIPVFNYQLYFQEPGVAEAELEKNMSRTFKSFFRASDETGFIAVHKATEIGGILVNTPED
PNLSKITTEEEIEFYIQQFKKTGFRGPLNWYRNTERNWKWSCKGLGRKILVPALMVTAEKDI
VLRPEMSKNMEKWIPFLKRGHIEDCGHWTQIEKPTEVNQILIKWLQTEVQNPSVTSKI
11100000000000000001111111111111111111111111111000000000000000
01111111111111111111111111100000000000000000000000000000000000
00000000000000000000000000000000000000000000000000000000000000
00000000000000000000000000000000000000000000000000000000000000
00000000000000000000000000000000000000000000000000000000000000
00000000000000000000000000000000000000000000000000000000000000
00000000000000000000000000000000000000000000000000000000000000
00000000000000000000000000000000000000000000000000000000000000
0000000000000000000000000000000000000000000000001111111111
>DM_validata79
MSSAEEKLFMKALKEKFEESPEEKYTKFYIFGGWKQSERKKEFKEWADKIVEERGVPHYNPD
IGVPLGQRKLMSYQVSGTDVFVEGDDLTFVNNAAMQQMWDDIRRTVIVGMDTAHRVLERRLG
KEVTPETINEYMETLNHALPGGAVVQEHMVEIHPGLTWDCYAKIITGDLELADEIDDKFLID
IEKLFPEEQAEQLIKAIGNRTYQVCRMPTIVGHVCDGATMYRWAAMQIAMSFICAYKIAAGE
AAVSDFAFASKHAEVINMGEMLPARRARGENEPGGVPFGVLADCVQTMRKYPDDPAKVALEV
IAAGAMLYDQIWLGSYMSGGVGFTQYATAVYPDNILDDYVYYGLEYVEDKYGIAEAEPSMDV
VKDVATEVTLYGLEQYERYPAAMETHFGGSQRAAVCAAAAGCSTAFATGHAQAGLNGWYLSQ
ILHKEGQGRLGFYGYALQDQCGAANSLSVRSDEGLPLELRGPNYPNYAMNVGHLGEYAGIVQ
AAHAARGDAFCVHPVIKVAFADENLVFDFTEPRKEFAKGALREFEPAGERDLIVPAE
11111110000000000000000000000000000000000000000000000000000000
00000000000000000000000000000000000000000000000000000000000000
00000000000000000000000000000000000000000000000000000000000000
00000000000000000000000000000000000000000000000000000000000000
00000000000000000000000000000000000000000000000000000000000000
00000000000000000000000000000000000000000000000000000000000000
00000000000000000000000000000000000000000000000000000000000000
00000000000000000000000000000000000000000000000000000000000000
000000000000000000000000000000000000000000000000000000000
>DM_validata82
MASWSHPQFEKGASTSLYKKAGLMDSQRSTLVHWFRKGLRLHDNPALSHIFTAANAAPGRYF
VRPIFILDPGILDWMQVGANRWRFLQQTLEDLDNQLRKLNSRLFVVRGKPAEVFPRIFKSWR
VEMLTFETDIEPYSVTRDAAVQKLAKAEGVRVETHCSHTIYNPELVIAKNLGKAPITYQKFL
GIVEQLKVPKVLGVPEKLKNMPTPPKDEVEQKDSAAYDCPTMKQLVKRPEELGPNKFPGGET
EALRRMEESLKDEIWVARFEKPNTAPNSLEPSTTVLSPYLKFGCLSARLFNQKLKEIIKRQP
KHSQPPVSLIGQLMWREFYYTVAAAEPNFDRMLGNVYCMQIPWQEHPDHLEAWTHGRTGYPF
IDAIMRQLRQEGWIHHLARHAVACFLTRGDLWISWEEGQRVFEQLLLDQDWALNAGNWMWLS
ASAFFHQYFRVYSPVAFGKKTDPQGHYIRKYVPELSKYPAGCIYEPWKASLVDQRAYGCVLG
TDYPHRIVKHEVVHKENIKRMGAAYKVNREVRTGKEEESSFEEKSET
11111111111111111111111100000000000000000000000000000000000000
00000000000000000000000000000000000000000000000000000000000000
00000000000000000000000000000000000000000000000000000000000000

00000000000000000000000000000000000000000000000000000000000000
00000000000000000000000000000000000000000000000000000000000000
00000000000000000000000000000000000000000000000000000000000000
00000000000000000000000000000000000000000000000000000000000000
00000000000000000000000000000000000000000000000000000000000000
00000000000000000000000000000000000011111111111
>DM_validata85
MMMASKDATSSVDGASGAGQLVPEVNASDPLAMDPVAGSSTAVATAGQVNPIDPWIINNFVQ
APQGEFTISPNNTPGDVLFDLSLGPHLNPFLLHLSQMYNGWVGNMRVRIMLAGNAFTAGKII
VSCIPPGFGSHNLTIAQATLFPHVIADVRTLDPIEVPLEDVRNVLFHNNDRNQQTMRLVCML
YTPLRTGGGTGDSFVVAGRVMTCPSPDFNFLFLVPPTVEQKTRPFTLPNLPLSSLSNSRAPL
PISSMGISPDNVQSVQFQNGRCTLDGRLVGTTPVSLSHVAKIRGTSNGTVINLTELDGTPFH
PFEGPAPIGFPDLGGCDWHINMTQFGHSSQTQYDVDTTPDTFVPHLGSIQANGIGSGNYVGV
LSWISPPSHPSGSQVDLWKIPNYGSSITEATHLAPSVYPPGFGEVLVFFMSKMPGPAYNLP
CLLPQEYISHLASEQAPTVGEAALLHYVDPDTGRNLGEFKAYPDGFLTCVPNGASSGPQQLP
INGVFVFVSWVSRFYQLKPVGTASSARGRLGLRR
11111111111111111111111111110000000000000000000000000000000000
00000000000000000000000000000000000000000000000000000000000000
00000000000000000000000000000000000000000000000000000000000000
00000000000000000000000000000000000000000000000000000000000000
00000000000000000000000000000000000000000000000000000000000000
00000000000000000000000000000000000000000000000000000000000000
00000000000000000000000000000000000000000000000000000000000000
00000000000000000000000000000000000000000000000000000000000000
0000000000000000000000001111111111
>DM_validata86
MALSTATKAATDALAANRAPTSVNAQEVHRWLQSFNWDFKNNRTKYATKYKMANETKEQFKL
IAKEYARMEAVKDERQFGSLQDALTRLNAGVRVHPKWNETMKVVSNFLEVGEYNAIAATGML
WDSAQAAEQKNGYLAQVLDEIRHTHQCAYVNYYFAKNGQDPAGHNDARRTRTIGPLWKGMKR
VFSDGFISGDAVECSLNLQLVGEACFTNPLIVAVTEWAAANGDEITPTVFLSIETDELRHMA
NGYQTVVSIANDPASAKYLNTDLNNAFWTQQKYFTPVLGMLFEYGSKFKVEPWVKTWDRWVY
EDWGGIWIGRLGKYGVESPRSLKDAKQDAYWAHHDLYLLAYALWPTGFFRLALPDQEEMEWF
EANYPGWYDHYGKIYEEWRARGCEDPSSGFIPLMWFIENNHPIYIDRVSQVPFCPSLAKGAS
TLRVHEYNGEMHTFSDQWGERMWLAEPERYECQNIFEQYEGRELSEVIAELHGLRSDGKTLI
AQPHVRGDKLWTLDDIKRLNCVFKNPVKAFN
11111111111111111000000000000000000000000000000000000000000000
00000000000000000000000000000000000000000000000000000000000000
00000000000000000000000000000000000000000000000000000000000000
00000000000000000000000000000000000000000000000000000000000000
00000000000000000000000000000000000000000000000000000000000000
00000000000000000000000000000000000000000000000000000000000000
00000000000000000000000000000000000000000000000000000000000000
00000000000000000000000000000000000000000000000000000000000000
0000000000000000000000000000000

>DM_validata87
INDLDISPIDPQTGGSFDQQGVFVKGYAMLGVTGQKGIDGSPDLDGQDEGESGFYRTTFNCN
ELPTDECLWAWQKNQDIPQLTSISWSPSSQRTEWVYVRLGYDITQYNFFLDQTEGMTDAETL
RQRAEIRFLRALHYWYFLDLFGKAPFKEHFSNDLPVEKKGTELYTYIQNELNEIEADMYEPR
QAPFGRADKAANWLLRARLYLNAGVYTGQTDYAKAEEYASKVIGSAYKLCTNYSELFMADND
ENENAMQEIILPIRQDGVKTRNYGGSTYLVCGTRVAGMPRMGTTNGWSCIFARAAMVQKFFS
NLEDVPMLPADVEIPTKGLDTDEQIDAFDAEHGIRTEDMIKAAGDDRALLYSGVGGGRRKIQ
TDAISGFTDGLSIVKWQNYRSDGKPVSHATYPDTDIPLFRLAEAYLTRAEAIFRQGGDATGD
INELRKRANCTRKVQTVTEQELIDEWAREFYLEGRRRSDLVRFGMFTTNKYLWDWKGGAMNG
TSVASYYNKYPIPVSDINNNRNMSQNEGYK
111111111111111100000000000000001111111111000000000000000000
000000000000000000000000000000000000000000000000000000000000
000000000000000000000000000000000000000000000000000000000000
000000000000000000000000000000000000000000000000000000000000
000000000000000000000000000000000000000000000000000000000000
000000000000000000000000000000000000000000000000000000000000
000000000000000000000000000000000000000000000000000000000000
000000000000000000000000000000000000000000000000000000000000
000000000000000000000000000000
>DM_validata88
MKARRSNELSKLRMRFFSALNHTSEIDLHTLFDNLKSNLTLGSIEHLQEGSVTYAIIQELLK
GADAQKKIESFLKGAIKNVIHPGVIKGLTPNEINWNVAKAYPEYYEHEKLPDVTFGGFKVRD
SNEFKFKTNVQTSIWFSIKPELFMPSKQQEALKRRREQYPGCKIRLIYSSSLLNPEANRQMK
AFAKKQNISLIDIDSVKTDSPLYPLIKAELANLGMGGNPAAASDLCRWIPELFNEGFYVDID
LPVDSSKIVEGHQITGGVPIMLNMGSIISEPIAPHHRRQEAVCMATDIIAYANDRETQVMMD
TVALHLKNIYDDPYTALKDTPLAQTAFFNRCEEEGKNIFELRKGLQDAFRSDSLLELYVFLG
PAKFKEVFKLKETQIKYIDDHISEFNEHDLLLHLISDNPSEINQHTLDFGRAKVMYMDIAKE
HYSAFYKPLVEEISGPGAIYNALGGASNFTTTHRRSTGPMLPTTPPRVLQVFCDAHDKGPFV
SDNIARWQTNVRELGVLNREGLSWLPSVG
111110000000000000000000000000000000000000000000000000000000
000000000000000000000000000000000000000000000000000000000000
000000000000000000000000000000000000000000000000000000000000
000000000000000000000000000000000000000000000000000000000000
000000000000000000000000000000000000000000000000000000000000
000000000000000000000000000000000000000000000000000000000000
000000000000000000000000000000000000000000000000000000000000
000000000000000000000000000000000000000000000000000000000000
00000000000000000000000000000
>DM_validata89
MVHWADYIADKIIRERGEKEKYVVESGITPSGYVHVGNFRELFTAYIVGHALRDKGYEVRHI
HMWDDYDRFRKVPRNVPQEWKDYLGMPISEVPDPWGCHESYAEHFMRKFEEEVEKLGIEVDL
LYASELYKRGEYSEEIRLAFEKRDKIMEILNKYREIAKQPPLPENWWPAMVYCPEHRREAEI
IEWDGGWKVKYKCPEGHEGWVDIRSGNVKLRWRVDWPMRWSHFGVDFEPAGKDHLVAGSSYD
TGKEIIKEVYGKEAPLSLMYEFVGIKGQKGKMSGSKGNVILLSDLYEVLEPGLVRFIYARHR

PNKEIKIDLGLGILNLYDEFEKVERIYFGVEGGKGDDEELRRTYELSMPKKPERLVAQAPFR
FLAVLVQLPHLTEEDIINVLIKQGHIPRDLSKEDVERVKLRINLARNWVKKYAPEDVKFSIL
EKPPEVEVSEDVREAMNEVAEWLENHEEFSVEEFNNILFEVAKRRGISSREWFSTLYRLFIG
KERGPRLASFLASLDRSFVIKRLRLEG
000000000000000000000000000000000000000000000000000000000000
000000000000000000000000000000000000000000000000000000000000
000000000000000000000000000000000000000000000000000000000000
000000000000000000000000000000000000000000000000000000000000
000000000000000000000000000111111111000000000000000000000000
000000000000000000000000000000011111000000000000000000000000
000000000000000000000000000000000000000000000000000000000000
000000000000000000000000000000000000000000000000000000000000
000000000000000000000000000
>DM_validata90
MASMEEPPLLPGENIKDMAKDVTYICPFTGAVRGTLTVTNYRLYFKSMERDPPFVLDASLGV
INRVEKIGGASSRGENSYGLETVCKDIRNLRFAHKPEGRTRRSIFENLMKYAFPVSNNLPLF
AFEYKEVFPENGWKLYDPLLEYRRQGIPNESWRITKINERYELCDTYPALLVVPANIPDEEL
KRVASFRSRGRIPVLSWIHPESQATITRCSQPMVGVSGKRSKEDEKYLQAIMDSNAQSHKIF
IFDARPSVNAVANKAKGGGYESEDAYQNAELVFLDIHNIHVMRESLRKLKEIVYPNIEETHW
LSNLESTHWLEHIKLILAGALRIADKVESGKTSVVVHSSDGWDRTAQLTSLAMLMLDGYYRT
IRGFEVLVEKEWLSFGHRFQLRVGHGDKNHADADRSPVFLQFIDCVWQMTRQFPTAFEFNEY
FLITILDHLYSCLFGTFLCNSEQQRGKENLPKRTVSLWSYINSQLEDFTNPLYGSYSNHVLY
PVASMRHLELWVGYYIRWNPRAAALE
111000000000000000000000000000000000000000000000000000000000
000000000000000000000000000000000000000000000000000000000000
000000000000000000000000000000000000000000000000000000000000
000000000000000000000000000000000000000000000000000000000000
000000000000000000000000000000000000000000000000000000000000
000000000000000000000000000000000000000000000000000000000000
000000000000000000000000000000000000000000000000000000000000
000000000000000000000000000000000000000000000000000000000000
00000000000000000000111111
>DM_validata92
MNTVSPAKKKVIIIGAGIAGLKAASTLHQNGIQDCLVLEARDRVGGRLQTVTGYQGRKYDIG
ASWHHDTLTNPLFLEEAQLSLNDGRTRFVFDDDNFIYIDEERGRVDHDKELLLEIVDNEMSK
FAELEFHQHLGVSDCSFFQLVMKYLLQRRQFLTNDQIRYLPQLCRYLELWHGLDWKLLSAKD
TYFGHQGRNAFALNYDSVVQRIAQSFPQNWLKLSCEVKSITREPSKNVTVNCEDGTVYNADY
VIITVPQSVLNLSVQPEKNLRGRIEFQPPLKPVIQDAFDKIHFGALGKVIFEFEECCWSNES
SKIVTLANSTNEFVEIVRNAENLDELDSMLEREDSQKHTSVTCWSQPLFFVNLSKSTGVASF
MMLMQAPLTNHIESIREDKERLFSFFQPVLNKIMKCLDSEDVIDGMRPIENIANANKPVLRN
IIVSNWTRDPYSRGAYSACFPGDDPVDMVVAMSNGQDSRIRFAGEHTIMDGAGCAYGAWESG
RREATRISDLLKLEHHHHHH
111110000000000000000000000000000000000000000000000000000000
000000000000000000000000000000000000000000000000000000000000

00000011111110000000000000000000000000000000000000000000000000
00000000000000000000000000000000000000000000000000000000000000
00000000000000000000000000000000000000000000000000000000000000
00000000000000000000000000000011111111100000000000000000000000
00000000000000000000000000000000000000000000000111111000000000
00000000000000000000011110000000000000000000000000000000000000
00000000000000000111
>DM_validata94
MGAPVLPAAFGFLASARTGGGRAPGPVFATRGSHTDIDTPQGERSLAATLVHAPSVAPDRAV
ARSLTGAPTTAVLAGEIYNRDELLSVLPAGPAPEGDAELVLRLLERYDLHAFRLVNGRFATV
VRTGDRVLLATDHAGSVPLYTCVAPGEVRASTEAKALAAHRDPKGFPLADARRVAGLTGVYQ
VPAGAVMDIDLGSGTAVTHRTWTPGLSRRILPEGEAVAAVRAALEKAVAQRVTPGDTPLVVL
SGGIDSSGVAACAHRAAGELDTVSMGTDTSNEFREARAVVDHLRTRHREITIPTTELLAQLP
YAVWASESVDPDIIEYLLPLTALYRALDGPERRILTGYGADIPLGGMHREDRLPALDTVLAH
DMATFDGLNEMSPVLSTLAGHWTTHPYWDREVLDLLVSLEAGLKRRHGRDKWVLRAAMADAL
PAETVNRPKLGVHEGSGTTSSFSRLLLDHGVAEDRVHEAKRQVVRELFDLTVGGGRHPSEVD
TDDVVRSVADRTARGAA
00000000000000000000111100000000000000000000000000000000000000
00000000000000000000000000000000000000000000000000000000000000
00000000000000000000000000000000000000001111000000000000000000
00000000000000000000000000000000000000000000000000000000000000
00000000000000000000000000000000000000000000000000000000000000
00000000000000000000000000000000000000000000000000000000000000
00000000000000000000000000000000000000000000000000000000000000
00000000011111111110000000000000011111000000000000000000000000
00000000001111111
>DM_validata96
MKMLNKLAGYLLPIMVLLNVAPCLGQVVASNETLYQVVKEVKPGGLVQIADGTYKDVQLIVS
NSGKSGLPITIKALNPGKVFFTGDAKVELRGEHLILEGIWFKDGNRAIQAWKSHGPGLVAIY
GSYNRITACVFDCFDEANSAYITTSLTEDGKVPQHCRIDHCSFTDKITFDQVINLNNTARAI
KDGSVGGPGMYHRVDHCFFSNPQKPGNAGGGIRIGYYRNDIGRCLVDSNLFMRQDSEAEIIT
SKSQENVYYGNTYLNCQGTMNFRHGDHQVAINNFYIGNDQRFGYGGMFVWGSRHVIACNYFE
LSETIKSRGNAALYLNPGAMASEHALAFDMLIANNAFINVNGYAIHFNPLDERRKEYCAANR
LKFETPHQLMLKGNLFFKDKPYVYPFFKDDYFIAGKNSWTGNVALGVEKGIPVNISANRSAY
KPVKIKDIQPIEGIALDLNALISKGITGKPLSWDEVRPYWLKEMPGTYALTARLSADRAAKF
KAVIKRNKEH
11111111111111111111111110000000000000000000000000000000000000
00000000000000000000000000000000000000000000000000000000000000
00000000000000000000000000000000000000000000000000000000000000
00000000000000000000000000000000000000000000000000000000000000
00000000000000000000000000000000000000000000000000000000000000
00000000000000000000000000000000000000000000000000000000000000
00000000000000000000000000000000000000000000000000000000000000
00000000000000000000000000000000000000000000000000000000000000

0000000000
>DM_validata97
ADSATPHLDAVEQTLRQVSPGLEGDVWERTSGNKLDGSAADPSDWLLQTPGCWGDDKCADRV
GTKRLLAKMTENIGNATRTVDISTLAPFPNGAFQDAIVAGLKESAAKGNKLKVRILVGAAPV
YHMNVIPSKYRDELTAKLGKAAENITLNVASMTTSKTAFSWNHSKILVVDGQSALTGGINSW
KDDYLDTTHPVSDVDLALTGPAAGSAGRYLDTLWTWTCQNKSNIASVWFAASGNAGCMPTMH
KDTNPKASPATGNVPVIAVGGLGVGIKDVDPKSTFRPDLPTASDTKCVVGLHDNTNADRDYD
TVNPEESALRALVASAKGHIEISQQDLNATCPPLPRYDIRLYDALAAKMAAGVKVRIVVSDP
ANRGAVGSGGYSQIKSLSEISDTLRNRLANITGGQQAAKTAMCSNLQLATFRSSPNGKWADG
HPYAQHHKLVSVDSSTFYIGSKNLYPSWLQDFGYIVESPEAAKQLDAKLLDPQWKYSQETAT
VDYARGICNA
11000000000000000000000000000000000000000000000000000000000000
00000000000000000000000000000000000000000000000000000000000001
10000000000000000000000000000000000000000000000000000000000000
00000000000000000000000000000000000000000000000000000000000000
00000000000000000000000000000000000000000000000000000000000000
00000000000000000000000000000000000000000000000000000000000000
00011111100000000000000000000000000000000000000000000000000000
00000000000000000000000000000000000000000000000000000000000000
0000000000
>DM_validata98
DNGGYVPAVVIGTGYGAAVSALRLGEAGVQTLMLEMGQLWNQPGPDGNIFCGMLNPDKRSSW
FKNRTEAPLGSFLWLDVVNRNIDPYAGVLDRVNYDQMSVYVGRGVGGGSLVNGGMAVEPKRS
YFEEILPRVDSSEMYDRYFPRANSMLRVNHIDTKWFEDTEWYKFARVSREQAGKAGLGTVFV
PNVYDFGYMQREAAGEVPKSALATEVIYGNNHGKQSLDKTYLAAALGTGKVTIQTLHQVKTI
RQTKDGGYALTVEQKDTDGKLLATKEISCRYLFLGAGSLGSTELLVRARDTGTLPNLNSEVG
AGWGPNGNIMTARANHMWNPTGAHQSSIPALGIDAWDNSDSSVFAEIAPMPAGLETWVSLYL
AITKNPQRGTFVYDAATDRAKLNWTRDQNAPAVNAAKALFDRINKANGTIYRYDLFGTQLKA
FADDFCYHPLGGCVLGKATDDYGRVAGYKNLYVTDGSLIPGSVGVNPFVTITALAERNVERI
IKQDVTAS
11100000000000000000000000000000000000000000000000000000000000
00000000000000000000000000000000000000000000000000000000000000
00000000000000000000000000000000000000000000000000000000000000
00000000000000000000000000000000000000000000000000000000000000
00000000000000000000000000000000000000000000000000000000000000
00000000000000000000000000000000000000000000000000000000000000
00000000000000000000000000000000000000000000000000000000000000
00000000000000000000000000000000000000000000000000000000000000
00000111
>DM_validata99
SDALKVRWPDFNQEAYVGGTMVRSGQDPYARNKFNQVESDKLRMDRAIPDTRHDQCQRKQWR
VDLPATSVVITFHNEARSALLRTVVSVLKKSPPHLIKEIILVDDYSNDPEDGALLGKIEKVR
VLRNDRREGLMRSRVRGADAAQAKVLTFLDSHCECNEHWLEPLLERVAEDRTRVVSPIIDVI
NMDNFQYVGASADLKGGFDWNLVFKWDYMTPEQRRSRQGNPVAPIKTPMIAGGLFVMDKFYF

EELGKYDMMMDVWGGENLEISFRVWQCGGSLEIIPCSRVGHVFRKQHPYTFPGGSGTVFARN
TRRAAEVWMDEYKNFYYAAVPSARNVPYGNIQSRLELRKKLSCKPFKWYLENVYPELRVPDH
QDIAFGALQQGTNCLDTLGHFADGVVGVYECHNAGGNQEWALTKEKSVKHMDLCLTVVDRAP
GSLIKLQGCRENDSRQKWEQIEGNSKLRHVGSNLCLDSRTAKSGGLSVEVCGPALSQQWKFT
LNLQQ
111100000000000000000000000000000000000000000000000000000000
000000000000000000000000000000000000000000000000000000000000
000000000000000000000000000000000000000000000000000000000000
000000000000000000000000000000000000000000000000000000000000
000000000000000000000000000000000000000000000000000000000000
000000000000000000000000000000000000000000000000000000000000
000000000000000000000000000000000000000000000000000000000000
000000000000000000000000000000000000000000000000000000000000
00111
>DM_validata100
MIRFKKTKLIASIAMALCLFSQPVISFSKDITDKNQSIDSGISSLSYNRNEVLASNGDKIES
FVPKEGKKAGNKFIVVERQKRSLTTSPVDISIIDSVNDRTYPGALQLADKALVENRPTILMV
KRKPININIDLPGLKGENSIKVDDPTYGKVSGAIDELVSKWNEKYSSTHTLPARTQYSESMV
YSKSQISSALNVNAKVLENSLGVDFNAVANNEKKVMILAYKQIFYTVSADLPKNPSDLFDDS
VTFNDLKQKGVSNEAPPLMVSNVAYGRTIYVKLETTSSSKDVQAAFKALIKNTDIKNSQQYK
DIYENSSFTAVVLGGDAQEHNKVVTKDFDEIRKVIKDNATFSTKNPAYPISYTSVFLKDNSV
AAVHNKTDYIETTSTEYSKGKINLDHSGAYVAQFEVAWDEVSYDKEGNEVLTHKTWDGNYQD
KTAHYSTVIPLEANARNIRIKARECTGLAWEWWRDVISEYDVPLTNNINVSIWGTTLYPGSS
ITYN
111111111111111111111111111100000000000000000000000000000000
000000000000000000000000000000000000000000000000000000000000
000000000000000000000000000000000000000000000000000000000000
000000000000000000000000000000000000000000000000000000000000
000000000000000000000000000000000000000000000000000000000000
000000000000000000000000000000000000000000000000000000000000
000000000000000000000000000000000000000000000000000000000000
000000000000000000000000000000000000000000000000000000000000
0000
>DM_validata101
MAMHPRKDWYELTRATNWTPSYVTEEQLFPERMSGHMGIPLEKWESYDEPYKTSYPEYVSIQ
REKDAGAYSVKAALERAKIYENSDPGWISTLKSHYGAIAVGEYAAVTGEGRMARFSKAPGNR
NMATFGMMDELRHGQLQLFFPHEYCKKDRQFDWAWRAYHSNEWAAIAAKHFFDDIITGRDAI
SVAIMLTFSFETGFANMQFLGLAADAAEAGDYTFANLISSIQTDESRHAQQGGPALQLLIEN
GKREEAQKKVDMAIWRAWRLFAVLTGPVMDYYTPLEDRSQSFKEFMYEWIIGQFERSLIDLG
LDKPWYWDLFLKDIDELHHSYHMGVWYWRTTAWWNPAAGVTPEERDWLEEKYPGWNKRWGRC
WDVITENVLNDRMDLVSPETLPSVCNMSQIPLVGVPGDDWNIEVFSLEHNGRLYHFGSEVDR
WVFQQDPVQYQNHMNIVDRFLAGQIQPMTLEGALKYMGFQSIEEMGKDAHDFAWADKCKPAM
KKSA
100000000000000000000000000000000000000000000000000000000000

00000000000000000000000000000000000000000000000000000000000000
00000000000000000000000000000000000000000000000000000000000000
00000000000000000000000000000000000000000000000000000000000000
00000000000000000000000000000000000000000000000000000000000000
00000000000000000000000000000000000000000000000000000000000000
00000000000000000000000000000000000000000000000000000000000000
00000000000000000000000000000000000000000000000000000000001111
1111
>DM_validata102
GSEDTMDRINKDHDHTTSVASRFILADVITSTAFSNASGDINTYASSYIEYEVGVDNQLYYA
EVRENEPSSSSTFNNSWNGIYSSLKNARIIIDQCGEGGRDHGNDVTRGMAEVMAAYNCALIA
DFFGDAPCSQAALVDEKGSPVYMTPKMDTQQEIYTQIISYLDDAIANLQKEDLADVTEQDFL
YAGDADKWLKFAYGLKARYTMRLINRSSNKSADYEKVLDYVSKSFTSADDQAAFDIYDSNNI
NPFYGFYNSRAGFGASTSLGTKLLAYNDPRANRAFFTPIVDKKRSQVAANDPSLVPAPNGSP
DQSTSKYGISAFVYAKTAPTLLMSYHELMFLKAEALCRLNRDAEDALKEAVVAGLLNAENSI
SIAIKELGSGLNTNSSEVITETSAGKYFDDVVKAKYAANPLQETMIQKYLAMWGASGEATET
YNDFRRMKGLNENFITLTNPNNSSKFPLRYPYGNSDTAANPEVKAAYGNGDYVYSEPVWWAG
GSR
11111111111111111000000000000000000000000000000000000000000000
00000000000000000000000000000000000000000000000000000000000000
00000000000011111111110000000000000000000000000000000000000000
00000000000000000000000000000000000000000000000000000000000000
00000000000000000000000000000000000000000000000000000000000000
00000000000000000000000000000000000000000000000000000000000000
00000000000000000000000000000000000000000000000000000000000000
00000000000000000000000000000000000000000000000000000000000000
000
>DM_validata103
MFESAEVGHSIDKDTYEKAVIELREALLEAQFELKQQARFPVIILINGIEGAGKGETVKLLN
EWMDPRLIEVQSFLRPSDEELERPPQWRFWRRLPPKGRTGIFFGNWYSQMLYARVEGHIKEA
KLDQAIDAAERFERMLCDEGALLFKFWFHLSKKQLKERLKALEKDPQHSWKLSPLDWKQSEV
YDRFVHYGERVLRRTSRDYAPWYVVEGADERYRALTVGRILLEGLQAALATKERAKRQPHAA
PLVSSLDNRGLLDSLDLGQYLDKDAYKEQLAAEQARLAGLIRDKRFRQHSLVAVFEGNDAAG
KGGAIRRVTDALDPRQYHIVPIAAPTEEERAQPYLWRFWRHIPARRQFTIFDRSWYGRVLVE
RIEGFCAPADWLRAYGEINDFEEQLSEYGIIVVKFWLAIDKQTQMERFKEREKTPYKRYKIT
EEDWRNRDKWDQYVDAVGDMVDRTSTEIAPWTLVEANDKRFARVKVLRTINDAIEAAYKKDK
00000000000000000000000000000000000000000000000000000000000000
00000000000000000000000000000000000000000000000000000000000000
00000000000000000000000000000000000000011111111111111111111110
00000000000000000000000000000000000000000000000000001111111111
11111100000000000000000000000000000000000000000000000000000000
00000000000000000000000000000000000000000000000000000000000000
00000000000000000000000000000000000000000000000000000000000000
00000000000000000000000000000000000000000000000000000000000000

>DM_validata104
DISSISEITYSDGTVIASIESDMLRQDFLPSGTVTGISQDYLYFTTLAEAQERMYDYLAQRD
NVSAKELKNEATQKFYRDLAAKEIENGGYKITTTIDQKIHSAMQSAVADYGYLLDDGTGRVE
VGNVLMDNQTGAILGFVGGRNYQENQNNHAFDTKRSPASTTKPLLAYGIAIDQGLMGSETIL
SNYPTNFANGNPIMYANSKGTGMMTLGEALNYSWNIPAYWTYRMLRENGVDVKGYMEKMGYE
IPEYGIESLPMGGGIEVTVAQHTNGYQTLANNGVYHQKHVISKIEAADGRVVYEYQDKPVQV
YSKATATIMQGLLREVLSSRVTTTFKSNLTSLNPTLANADWIGKTGTTNQDENMWLMLSTPR
LTLGGWIGHDDNHSLSQQAGYSNNSNYMAHLVNAIQQASPSIWGNERFALDPSVVKSEVLKS
TGQKPGKVSVEGKEVEVTGSTVTSYWANKSGAPATSYRFAIGGSDADYQNAWSSIVGSLP
111111100011111111111111111111111111110000000000000000000000
000000000000000000000000000000000000000000000000000000000000
000000000000000000000000000000000000000000000000000000000000
000000000000000000000000000000000000000000000000000000000000
000000000000000000000000000000000000000000000000000000000000
000000000000000000000000000000000000000000000000000000000000
000000000000000000000000000000000000000000000000000000000000
0000000000000000000000000000000000000000000000000000000011
>DM_validata105
MDVNNSKPNDYGTLVKIEQKLFNNANTLKTTTPIKHVVIIFQENNSFDRYFGMYPNAKNPEG
EPKFVAKENTPNVNGLTKQLLENNPNTKNPYRLDRNFQPCSQNHEYHQEISSFNGGLMNKFV
EHGGHDNDTYKQNCDGQVMGYYDGNTVTALWNYAQNFALNDNTFGTTFGPSTPGALNLVAGA
NGPAMSPSGNLENIENNYIIDDPNPYYDDCSYGTSKSGDTNTAVAKITDGYNIGHYLTQKGI
TWGWFQGGFKPTSYSGKTAICDAMSTNKFGVKSRDYIPHHEPFNYWKETSNPHHLAPSDDKY
IGSNDQANHQYDISEFWKALDQNNMPAVSYLKAPGYQDGHGGYSNPLDEQEWLVNTINRIQQ
SKDWDSTAIIIIYDDSDGDYDHVYSPKSQFSDIKGRQGYGPRLPMLVISPYAKANYVDHSLL
NQASVLKFIEYNWGIGSVSKYSNDKYSNNILNMFDFNKEQKTLKLILDPKTGLVMDKLN
111110000001111111000000000000000000000000000000000000000000
000000000000000000000000000000000000000000000000000000000000
000011111111100000000000000000000000000000000000000000000000
000000000000000000000000000000000000000000000000000000000000
000000000000000000000000000000000000000000000000000000000000
000000000000000000000000000000000000000000000000000000000000
000000000000000000000000000000000000000000000000000000000000
000000000000000000000000000000000000000000000000000001111
>DM_validata106
LMVGGLLLAAASNNVQAEALQPDPAWQQGKLDNGFSWQLLATPQRPSDRIELRLIVNTGSLS
ENTQEVGFAHLLPRLALMSSASFTPAQLQSLWQQGIDNERPLPPAITSYDFTLYSLSLPNNR
PDLLKDALAWLSDTAGNLAVSEQTVNAALNTATDPIATFPQNIQEPWWRYRLKGSSLIGHDP
GQPVTQPVDVEKLKQFYQQWYTPDAMTLYVVGNVDSRSIAAQISKAFSELKGKRTAPAAVAT
LAPLPPEPVSLMNEQAAQDTLSLMWDTPWHPIQDSMALSRYWRSDLAREALFWHIKQVLEKN
NQKNLKLGFDCRVQYQRAQCAIHLNTPVENLTANMTFVARELAALRANGLSQAEFDALMTQK
NDQLSKLFATYARTDTDILMSQRLRSQQSGVVDIAPEQYQKLRQAFLSGLTLAELNRELKQQ
LSQDTTLVLMQPKGEPEVNVKALQEIYNGIMAPQTVAEEEVAPAEAVETAPVMPTTAQ
000000011111111000000000000000000000000000000000000000000000

000000000000000000000000000000000000000000000000000000000000
000000000000000000000000000000000000000000000000000000000000
000000000000000000000000000000000000000000000000000000000000
000000000000000000000000000000000000000000000000000000000000
000000000000000000000000000000000000000000000000000000000000
000000000000000000000000000000000000000000000000000000000000
00000000000000000000000000000001111111111111111111111111
>DM_validata107
MAHYPPSKDQLNELIQEVNQWAITNGLSMYPPKFEENPSNASVSPVTIYPTPIPRKCFDEAV
QIQPVFNELYARITQDMAQPDSYLHKTTEALALSDSEFTGKLWSLYLATLKSAQYKKQNFRL
GIFRSDYLIDKKKGTEQIKQVEFNTVSVSFAGLSEKVDRLHSYLNRANKYDPKGPIYNDQNM
VISDSGYLLSKALAKAVESYKSQQSSSTTSDPIVAFIVQRNERNVFDQKVLELNLLEKFGTK
SVRLTFDDVNDKLFIDDKTGKLFIRDTEQEIAVVYYRTGYTTTDYTSEKDWEARLFLEKSFA
IKAPDLLTQLSGSKKIQQLLTDEGVLGKYISDAEKKSSLLKTFVKIYPLDDTKLGREGKRLA
LSEPSKYVLKPQREGGGNNVYKENIPNFLKGIEERHWDAYILMELIEPELNENNIILRDNKS
YNEPIISELGIYGCVLFNDEQVLSNEFSGSLLRSKFNTSNEGGVAAGFGCLDSIILY
111100000000000000000000000000000000000000000000000000000000
000000000000000000000001111111111111000000000000000000000000
000000000000111000000000000000000000000000000000000000000000
000000000000000000000000000000000000000000000000000000000000
000000000000000000000000000000000000000000000000000000000000
000000000000000000000000000000000000000000000000000000000000
000000000000111100000000000000000000000000000000000000000000
00000000000000000000000000000000001111111111110000000000
>DM_validata109
MTFRNCVAVDLGASSGRVMLARYERECRSLTLREIHRFNNGLHSQNGYVTWDVDSLESAIRL
GLNKVCAAGIAIDSIGIDTWGVDFVLLDQQGQRVGLPVAYRDSRTNGLMAQAQQQLGKRDIY
QRSGIQFLPFNTLYQLRALTEQQPELIPHIAHALLMPDYFSYRLTGKMNWEYTNATTTQLVN
INSDDWDESLLAWSGANKAWFGRPTHPGNVIGHWICPQGNEIPVVAVASHDTASAVIASPLN
GSRAAYLSSGTWSLMGFESQTPFTNDTALAANITNEGGAEGRYRVLKNIMGLWLLQRVLQER
QINDLPALIAATQALPACRFIINPNDDRFINPDEMCSEIQAACREMAQPIPESDAELARCIF
DSLALLYADVLHELAQLRGEDFSQLHIVGGGCQNTLLNQLCADACGIRVIAGPVEASTLGNI
GIQLMTLDELNNVDDFRQVVSTTANLTTFTPNPDSEIAHYVALIHSTRQTKELCA
100000000000000000000000000000000000000000000000000000000000
000000000000000000000000000000000000000000000000000000000000
000000000000000000000000000000000000000000000000000000000000
000000000000000000000000000000000000000000000000000000000000
000000000000000000000000000000000000000000000000000000000000
000000000000000000000000000000000000000000000000000000000000
000000000000000000000000000000000000000000000000000000000000
00000000000000000000000000000000000000000000011111111
>DM_validata110
MEATPTPADLFSEDYLVDTLDGLTVDDQQAVLASLSFSKFLKHAKVRDWCAQAKIQPSMPAL
RMAYNYFLFSKVGEFIGSEDVCNFFVDRVFGGVRLLDVASVYAACSQMNAHQRHHICCLVER

ATSSQSLNPVWDALRDGIISSSKFHWAVKQQNTSKKIFSPWPITNNHFVAGPLAFGLRCEEV
VKTLLATLLHPDETNCLDYGFMQSPQNGIFGVSLDFAANVKTDTEGRLQFDPNCKVYEIKCR
FKYTFAKMECDPIYAAYQRLYEAPGKLALKDFFYSISKPAVEYVGLGKLPSESDYLVAYDQE
WEACPRKKRKLTPLHNLIRECILHNSTTESDVYVLTDPQDTRGQISIKARFKANLFVNVRHS
YFYQVLLQSSIVEEYIGLDSGIPRLGSPKYYIATGFFRKRGYQDPVNCTIGGDALDPHVEIP
TLLIVTPVYFPRGAKHRLLHQAANFWSRSAKDTFPYIKWDFSYLSANVPHSP
11111100000000000000000000000000000000000000000000000000000000
00000000000000000000000000000000000000000000000000000000000000
00000000000000000000000000000000000000001111111111111110000000
00000000000000000000000000000000000000000000000000000000000000
00000000000000000000000000000000000000000000000000000000000000
00001111000000000000000000000000000000000000000000000000000000
00000000000000000000000000000000000000000000000000000000000000
0000000000000000000000000000000000000000000000011111
>DM_validata111
SGSETPDLTVATRTGSKDLPIRNIPGNYGLPIVGPIKDRWDYFYDQGAEEFFKSRIRKYNST
VYRVNMPPGAFIAENPQVVALLDGKSFPVLFDVDKVEKKDLLTGTYMPSTELTGGYRILSYL
DPSEPKHEKLKNLLFFLLKSSRNRIFPEFQATYSELFDSLEKELSLKGKADFGGSSDGTAFN
FLARAFYGTNPADTKLKADAPGLITKWVLFNLHPLLSIGLPRVIEEPLIHTFSLPPALVKSD
YQRLYEFFLESAGEILVEADKLGISREEATHNLLFATCFNTWGGMKILFPNMVKRIGRAGHQ
VHNRLAEEIRSVIKSNGGELTMGAIEKMELTKSVVYECLRFEPPVTAQYGRAKKDLVIESHD
AAFKVKAGEMLYGYQPLATRDPKIFDRADEFVPERFVGEEGEKLLRHVLWSNGPETETPTVG
NKQCAGKDFVVLVARLFVIEIFRRYDSFDIEVGTSPLGSSVNFSSLRKASF
11111111111111111100000000000000000000000000000000000000000000
00000000000000000000000000000000000000000000000000000000000000
00000000000000000000000000000000000000000000000000000000000000
00000000000000000000000000000000000000000000000000000000000000
00000000000000000000000000000000000000000000000000000000000000
00000000000000000000000000000000000000000000000000000000000000
00000000000000000000000000000000000000000000000000000000000000
000000000000000000000000000000000000000000000000011
>DM_validata112
MNPAAEAEFNILLATDSYKVTHYKQYPPNTSKVYSYFECREKKTENSKLRKVKYEETVFYGL
QYILNKYLKGKVVTKEKIQEAKDVYKEHFQDDVFNEKGWNYILEKYDGHLPIEIKAVPEGFV
IPRGNVLFTVENTDPECYWLTNWIETILVQSWYPITVATNSREQKKILAKYLLETSGNLDGL
EYKLHDFGYRGVSSQETAGIGASAHLVNFKGTDTVAGLALIKKYYGTKDPVPGYSVPAAEHS
TITAWGKDHEKDAFEHIVTQFSSVPVSVVSDSYDIYNACEKIWGEDLRHLIVSRSTQAPLII
RPDSGNPLDTVLKVLEILGKKFPVTENSKGYKLLPPYLRVIQGDGVDINTLQEIVEGMKQKM
WSIENIAFGSGGGLLQKLTRDLLNCSFKCSYVVTNGLGINVFKDPVADPNKRSKKGRLSLHR
TPAGNFVTLEEGKGDLEEYGQDLLHTVFKNGKVTKSYSFDEIRKNAQLNI
11111111000000000000000000000000000000000111111111110000000000
00000000000000000000000000000000000000000000000000000000000000
00000000000000000000000000000000000000000000000000000000000000
00000000000000000000000000000000000000000000000000000000000000

00000000000000000000000000000000000000000000000000000000000000
00000000000000000000000000000000000000000000000000000000000000
00000000000000000000000000000000000000000000000000000000000000
00000000000000000000000000000000000000000000000000
>DM_validata113
LNPPSAFSGAALACLEKMQASGVEEKCIHIFLIQHALVRKGETGYIPEKSISPVESLPFLQG
IETKGENTALLRQAVVLKLNGGLGTGMGLNGPKSLLQVKNGQTFLDFTALQLEHFRQVRNCN
VPFMLMNSFSTSGETKNFLRKYPTLYEVFDSDIELMQNRVPKIRQDNFFPVTYEADPTCEWV
PPGHGDVYTVLYSSGKLDYLLGKGYRYMFISNGDNLGATLDVRLLDYMHEKQLGFLMEVCRR
TESDKKGGHLAYKDVIDETTGQTRRRFVLRESAQCPKEDEDSFQNIAKHCFFNTNNIWINLM
ELKKMMDEQLGVLRLPVMRNPKTVNPQDSQSTKVYQLEVAMGAAISLFDRSEAVVVPRERFA
PVKTCSDLLALRSDAYQVTEDQRLVLCEERNGKPPAIDLDGEHYKMIDGFEKLVKGGVPSLR
QCTSLTVRGLVEFGADVSVRGNVVIKNLKEEPLIIGSGRVLDNEVVVVE
11111110000000000000000000000000000000000000000000000000000011
11111111000000000000000000000000000000000000000000000000000011
00000000000000000000000000000000000000000000000000000000000000
00000000000000000000000000000000000000000000000000000000000000
00000000000000000111111000000000000000000000000000000000000000
00000000000000000000000000000000000000000000000000000000000000
00000000000000000000000000000000000000000000000000000000000000
0000000000000000000000000000000000000000000000001
>DM_validata114
SIDETRAHLLLKEKMMRLGGRLVLNTKEELANERLMTLKIAEMKEAMRTLIFPPSMHFFQAK
HLIERSQVFNILRMMPKGAALHLHDIGIVTMDWLVRNVTYRPHCHICFTPRGIMQFRFAHPT
PRPSEKCSKWILLEDYRKRVQNVTEFDDSLLRNFTLVTQHPEVIYTNQNVVWSKFETIFFTI
SGLIHYAPVFRDYVFRSMQEFYEDNVLYMEIRARLLPVYELSGEHHDEEWSVKTYQEVAQKF
VETHPEFIGIKIIYSDHRSKDVAVIAESIRMAMGLRIKFPTVVAGFDLVGHEDTGHSLHDYK
EALMIPAKDGVKLPYFFHAGETDWQGTSIDRNILDALMLNTTRIGHGFALSKHPAVRTYSWK
KDIPIEVCPISNQVLKLVSDLRNHPVATLMATGHPMVISSDDPAMFGAKGLSYDFYEVFMGI
GGMKADLRTLKQLAMNSIKYSTLLESEKNTFMEIWKKRWDKFIADVATK
00000000000000000000000000000000000000000000000000000000000000
00000000000000000000000000000000000000000000000000000000000000
00000000000000000000000000000000000000000000000000000000000000
00000000000000000000000000000000000000000000000000000000000000
00000000000000000000000000000000000000000000000000000000000000
00000000000000000000000000000000000000000000000000000000000000
00000000000000000000000000000000000000000000000000000000000000
0000000000000000000000000000000000000000000000001
>DM_validata115
QMRLPSADVYRFAEPDSEENIIFEENMQPKAGIPIIKAGTVIKLIERLTYHMYADPNFVRTF
LTTYRSFCKPQELLSLIIERFEIPEPEPTEADRIAIENGDQPLSAELKRFRKEYIQPVQLRV
LNVCRHWVEHHFYDFERDAYLLQRMEEFIGTVRGKAMKKWVESITKIIQRKKIARDNGPGHN
ITFQSSPPTVEWHISRPGHIETFDLLTLHPIEIARQLTLLESDLYRAVQPSELVGSVWTKED
KEINSPNLLKMIRHTTNLTLWFEKCIVETENLEERVAVVSRIIEILQVFQELNNFNGVLEVV

SAMNSSPVYRLDHTFEQIPSRQKKILEEAHELSEDHYKKYLAKLRSINPPCVPFFGIYLTNI
LKTEEGNPEVLKRHGKELINFSKRRKVAEITGEIQQYQNQPYCLRVESDIKRFFENLNPMGN
SMEKEFTDYLFNKSLEIEPRNPKPLPRFPKKYSYPLKSPGVRPSNPR
00000000000000000000000001111110000000000000000000000000000000
00000000000000000000000000111111111111111111100000000000000000
00000000000000000000000000000000000000000000000000000011111100
00000000000000000000000000000000000000000000000000000000000000
00000000000000000000000000000000000000000000000000000000000000
00000000000000000000000000000000000000000000000000000000000000
00000000000000000000000000000000000000000000000000000000000000
00000000000000000000000000000000000000000000000
>DM_validata116
MGKNLSLRQDDAQHALSANTSSAYNSVYDFLRYHDRGDGLTVNGKTSYSIDQAAAQITRENV
SWNGTNVFGKSANLTFKFLQSVSSIPSGDTGFVKFNAEQIEQAKLSLQSWSDVANLTFTEVT
GNKSANITFGNYTRDASGNLDYGTQAYAYYPGNYQGAGSSWYNYNQSNIRNPGSEEYGRQTF
THEIGHALGLAHPGEYNAGEGDPSYNDAVYAEDSYQFSIMSYWGENETGADYNGHYGGAPMI
DDIAAIQRLYGANMTTRTGDSVYGFNSNTDRDFYTATDSSKALIFSVWDAGGTDTFDFSGYS
NNQRINLNEGSFSDVGGLKGNVSIAHGVTIENAIGGSGNDILVGNSADNILQGGAGNDVLYG
GAGADTLYGGAGRDTFVYGSGQDSTVAAYDWIADFQKGIDKIDLSAFRNEGQLSFVQDQFTG
KGQEVMLQWDAANSITNLWLHEAGHSSVDFLVRIVGQAAQSDIIV
11111111111111111111111100000000000000000000000000000000000000
00000000000000000000000000000000000000000000000000000000000000
00000000000000000000000000000000000000000000000000000000000000
00000000000000000000000000000000000000000000000000000000000000
00000000000000000000000000000000000000000000000000000000000000
00000000000000000000000000000000000000000000000000000000000000
00000000000000000000000000000000000000000000000000000000000000
000000000000000000000000000000000000000000000
>DM_validata117
MVLDIQLFRDETGANIIRESQRRRFADPDIVDAIIEADKKWRRTQFLTEASKKLINICSKAV
GAKKKAKEADGDTSEIPPQVKEAYENGTLKGEQVEQLCVLQLKQLSKDLSDQVAGLAKEAQQ
LEEERDKLMLNVGNILHESVPIAQDEETGNTVVRTFGNTTKRAKLNHVSIMERLGMMDTSKA
VTSMAGGRSYVLKGGLVQLQVALVSYSLDFLVKRGYTPFYPPFFLNRDVMGEVAQLSQFDEE
LYQVSGDGDKKYLIATSEMPIAAYHRGRWFTELKEPLKYAGMSTCFRKEAGAHGRDTLGIFR
VHQFDKIEQFVVCSPRQEESWRHLEDMITTSEEFNKSLGLPYRVVNICSGALNNAAAKKYDL
EAWFPASGAFRELVSCSNCTDYQSQSVNCRYGPNLRGTAAQNVKEYCHMLNGTLCAITRTMC
CICENYQTEEGVVIPDVLRPYMMGIEMIRFENNAQAEGTTPDKGE
00000000000000000000000000000000000000000000000000000000000000
00000000011111111111111111111111111110000000000000000000000000
00000000000000000000000000000000000000000000000000000000000000
00000000000000000000000000000000000000000000000000000000000000
00000000000000000000000000000000000000000000000000000000000000
00000000000000000000000000000000000000000000000000000000000000
00000000000000000000000000000000000011110000000000000000000000

0000000000000000000000000000000000111111111111
>DM_validata118
VVRDVNWGALRIAVSTEDLTDPAYHCARVGQHVKDHAGAIVTCTAEDILTNEKRDILVKHLI
PQAVQLHTERLKVQQVQGKWKVTDMVGDICGDFKVPQAHITEGFSNTDFVMYVASVPSEEGV
LAWATTCQTFSDGHPAVGVINIPAANIASRYDQLVTRVVTHEMAHALGFSGPFFEDARIVAN
VPNVRGKNFDVPVINSSTAVAKAREQYGCDTLEYLEVEDQGGAGSAGSHIKMRNAQDELMAP
AAAAGYYTALTMAIFQDLGFYQADFSKAEVMPWGQNAGCAFLTNKCMEQSVTQWPAMFCNES
EDAIRCPTSRLSLGACGVTRHPGLPPYWQYFTDPSLAGVSAFMDYCPVVVPYSDGSCTQRAS
EAHASLLPFNVFSDAARCIDGAFRPKATDGIVKSYAGLCANVQCDTATRTYSVQVHGSNDYT
NCTPGLRVELSTVSNAFEGGGYITCPPYVEVCQGNVQAAKDGGN
0000000000000000000000000000000000000000000000000000000000000000
0000000000000000000000000000000000000000000000000000000000000000
0000000000000000000000000000000000000000000000000000000000000000
0000000000000000000000000000000000000000000000000000000000000000
0000000000000000000000000000000000000000000000000000000000000011
1100000000000000000000000000000000000000000000000000000000000000
0000000000000000000000000000111111000000000000000000000000000000
000000000000000000000000000000000000000000111
>DM_validata119
GMIDLDFRKLTIEECLKLSEEEREKLPQLSLETIKRLDPHVKAFISVRENVSVEKKGKFWGI
PVAIKDNILTLGMRTTCASRILENYESVFDATVVKKMKEAGFVVVGKANLDEFAMGSSTERS
AFFPTRNPWDLERVPGGSSGGSAAAVSAGMVVAALGSDTGGSVRQPASLCGVVGYKPTYGLV
SRYGLVAFASSLDQIGPITKTVRDAAILMEIISGRDENDATTVNRKVDFLSEIEEGVSGMKF
AVPEEIYEHDIEEGVSERFEEALKLLERLGAKVERVKIPHIKYSVATYYVIAPAEASSNLAR
FDGVKYGLRIKEKGLREMYMKTRNVGFGEEVRRRIMIGTFTLSAAYYEAYFNKAMKVRRKIS
DELNEVLSQYDAILTPTSPVTAFKIGEIKDPLTYYLMDIFTIPANLAGLPAISVPFGFSNNL
PVGVQVIGRRFADGKVFRIARAIEKNSPYNENGMFPLPEVKA
0000000000000000000000000000000000000000000000000000000000000000
0000000000000000000000000000000000000000000000000000000000000000
0000000000000000000000000000000000000000000000000000000000000000
0000000000000000000000000000000000000000000000000000000000000000
0000000000000000000000000000000000000000000000000000000000000111
1111111111111111111110000000000000000000000000000000000000000000
0000000000000000000000000000000000000000000000000000000000000000
0000000000000000000000000000000000000000000
>DM_validata120
MGSSSSGLVPRGSHMSGSGSGSGTALTPSYLKDDDGRSLILRGFNTASSAKSAPDGMPQFTE
ADLAREYADMGTNFVRFLISWRSVEPAPGVYDQQYLDRVEDRVGWYAERGYKVMLDMHQDVY
SGAITPEGNSGNGAGAIGNGAPAWATYMDGLPVEPQPRWELYYIQPGVMRAFDNFWNTTGKH
PELVEHYAKAWRAVADRFADNDAVVAYDLMNEPFGGSLQGPAFEAGPLAAMYQRTTDAIRQV
DQDTWVCVAPQAIGVNQGLPSGLTKIDDPRAGQQRIAYCPHLYPLPLDIGDGHEGLARTLTD
VTIDAWRANTAHTARVLGDVPIILGSFGLDTTLPGARDYIERVYGTAREMGAGVSYWSSDPG
PWGPYLPDGTQTLLVDTLNKPYPRAVAGTPTEWSSTSDRLQLTIEPDAAITAPTEIYLPEAG
FPGDVHVEGADVVGWDRQSRLLTVRTPADSGNVTVTVTPAA

11111111111111111111111111000000000000000000000000000000000000
000000000000000000000000000000000000000000000000000000000000
000000000000000000000000000000000000000000000000000000000000
000000000000000000000000000000000000000000000000000000000000
000000000000000000000000000000000000000000000000000000000000
000000000000000000000000000000000000000000000000000000000000
000000000000000000000000000000000000000000000000000000000000
00000000000000000000000000000000000000000
>DM_validata121
MRGLAVLVLLVFAVQVAAAEDFRPVVFVHGLAGSAGQFESQGMRFAANGYPAEYVKTFEYDT
ISWALVVETDMLFSGLGSEFGLNISQIIDPETLDKILSKSRERLIDETFSRLDRVIDEALAE
SGADKVDLVGHSMGTFFLVRYVNSSPERAAKVAHLILLDGVWGVDAPEGIPTLAVFGNPKAL
PALGLPEEKVVYNATNVYFNNMTHVQLCTSPETFAVMFEFINGYKPATTDIVPQDGDYVKVK
GKFLAFATNGDVSGWLSIYPIDENGKRLTRLPVKFMRVKGDFEVRLRKGQLYEFQFRKDFSP
IIYHYYRAPFVRDDLWARFLVSKPPLDVELLILPERLSPAAKETSGLLLIRYKEMIGEYDEE
IGGVDEVYVNGVNVCTERICPIERAVNGLWVFDRGADGKSDLDREVVRYSIMPFMSAADLVV
PAEGTISIAVKSRTGGEESFTIPAWSADRHSIIVQFSDYI
111111111111111111110000000000000000000000000000000000000000
000000000000000000000000000000000000000000000000000000000000
000000000000000000000000000000000000000000000000000000000000
000000000000000000000000000000000000000000000000000000000000
000000000000000000000000000000000000000000000000000000000000
000000000000000000000000000000000000000000000000000000000000
000000000000000000000000000000000000000000000000000000000000
0000000000000000000000000000000000000000
>DM_validata122
GASAKEKFTSLSPAEFFKRNPELAGFPNPARALYQTVRELIENSLDATDVHGILPNIKITID
LIDDARQIYKVNVVDNGIGIPPQEVPNAFGRVLYSSKYVNRQTRGMYGLGVKAAVLYSQMHQ
DKPIEIETSPVNSKRIYTFKLKIDINKNEPIIVERGSVENTRGFHGTSVAISIPGDWPKAKS
RIYEYIKRTYIITPYAEFIFKDPEGNVTYYPRLTNKIPKPPQEVKPHPYGVDREEIKILINN
LKRDYTIKEFLVNEFQSIGDTTADKILELAGLKPNKKVKNLTEEEITRLVETFKKYEDFRSP
SADSLSVIGEDLIELGLKKIFNPDFAASITRKPKAYQGHPFIVEAGVAFGGSIPVGEEPIVL
RYANKIPLIYDEKSDVIWKVVEELDWKRYGIESDQYQMVVMVHLCSTKIPYKSAGKESIAEV
EDIEKEIKNALMEVARKLKQYLSEKRKEQEAKKKLLA
111111111100000000000000000000000000000000000000000000000000
000000000000000000000000000000000000000000000000000000000000
000000000000000000000000000000000000000000000000000000000000
000000000000000000000000000000000000000000000000000000000000
000000000000000000000000000000000000000000000000000000000000
000000000000000000000000000000000000000000000000000000000000
000000000000000000000000000000000000000000000000000000000000
0000000000000000000000000000000000000
>DM_validata123
MHHHHHHASENLAFQGAMASSIDISKINSWNKEFQSDLTHQLATTVLKNYNADDALLNKTRL

QKQDNRVFNTVVSTDSTPVTNQKSSGRAWLFAATNQLRLNVLSELNLKEFELSQAYLFFYDK
LEKANYFLDQIVSSADQDIDSRLVQYLLAAPTEDGGQYSMFLNLVKKYGLIPKDLYGDLPYS
TTASRKWNSLLTTKLREFAETLRTALKERSADDSIIVTLREQMQREIFRLMSLFMDIPPVQP
NEQFTWEYVDKDKKIHTIKSTPLEFASKYAKLDPSTPVSLINDPRHPYGKLIKIDRLGNVLG
GDAVIYLNVDNETLSKLVVKRLQNNKAVFFGSHTPKFMDKKTGVMDIELWNYPAIGYNLPQQ
KASRIRYHESLMTHAMLITGCHVDETSKLPLRYRVENSWGKDSGKDGLYVMTQKYFEEYCFQ
IVVDINELPKELASKFTSGKEEPIVLPIWDPMGALAK
11111111111100000000000000000000000000000000000000000000000000
00000000000000000000000000000000000000000000000000000000000000
00000000000000000000000000000000000000000000000000000000000000
00000000000000000000000000000000000000000000000000000000000000
00000000000000000000000000000000000000000000000000000000000000
00000000000000000000000000000000000000000000000000000000000000
00000000000000000000000000000000000000000000000000000000000000
0000000000000000000000000000000000000
>DM_validata124
AEPVYPDQLRLFSLGQGVCGDKYRPVNREEAQSVKSNIVGMMGQWQISGLANGWVIMGPGYN
GEIKPGTASNTWCYPTNPVTGEIPTLSALDIPDGDEVDVQWRLVHDSANFIKPTSYLAHYLG
YAWVGGNHSQYVGEDMDVTRDGDGWVIRGNNDGGCDGYRCGDKTAIKVSNFAYNLDPDSFKH
GDVTQSDRQLVKTVVGWAVNDSDTPQSGYDVTLRYDTATNWSKTNTYGLSEKVTTKNKFKWP
LVGETELSIEIAANQSWASQNGGSTTTSLSQSVRPTVPARSKIPVKIELYKADISYPYEFKA
DVSYDLTLSGFLRWGGNAWYTHPDNRPNWNHTFVIGPYKDKASSIRYQWDKRYIPGEVKWWD
WNWTIQQNGLSTMQNNLARVLRPVRAGITGDFSAESQFAGNIEIGAPVPLAADSKVRRARSV
DGAGQGLRLEIPLDAQELSGLGFNNVSLSVTPAANQ
00000000000000000000000000000000000000000000000000000000000000
00000000000000000000000000000000000000000000000000000000000000
00000000000000000000000000000000000000000000000000000000000000
00000000000000000000011100000000000000000000000000000000000000
00000000000000000000000000000000000000000000000000000000000000
00000000000000000000000000000000000000000000000000000000000000
00000000000000000000000000000000000000000000000000111111111111
111110000000000000000000000000000000
>DM_validata126
SPLSLPSTKLNPSPDHYAVWGKAIMAENNRRVGPEHMFRTAIRAQQQLQGLADKWTPDAKVY
CCGSMVTYGQMERGSDLDLACMFDDPYPSHEVQAKRTDKLRTVIKRYVPHYLRNNLLGLTEA
RTPVVKLRFANDEKVARARYTPLSEEEDRKARTALLDVRNQCVGDNDVEYIAEKMGRDNVEG
IRVDRTTYGCRIAIQCTSKEQMIEAIGFFPDGKIMTRGMREDYTRDVLDVRFVPEMFMYRWD
ISFVGYGVKNSYLIRHYLHNGPVAARHTAMAVKAWGKATNVGAGSGAMLTSYAVTVMFIYYL
LVTRQVLWVDPWSLPHPAHLPRYPDFSPLYDCDPTELGRLLHGFFIFYAHHFDYEREVVSLN
RNRRSYRSDIGWNFPQNKKGTFSYNFCIEDPYEDVGTGGLNLVRHLHPAKFQLVKQEFLRAA
QCMERFLPTNAPEKSILGVKRADLRHFERDRDRE
11111111110000000000000000000000000000000000000000000000000000
00000000000000000000000000000000000000000000000000000000000000
00000000000000000000000000000000000000000000000000000000000000

000000000000000000000000000000000000000000000000000000000000
000000000000000000000000000000000000001111100000000000000000
000000000000000000000000000000000000000000000000000000000000
000000000000000000000000000000000000000000000000000000000000
000000000000000000111111111111111
>DM_validata127
MDKKQVTDLRSELLDSRFGAKSISTIAESKRFPLHEMRDDVAFQIINDELYLDGNARQNLAT
FCQTWDDENVHKLMDLSINKNWIDKEEYPQSAAIDLRCVNMVADLWHAPAPKNGQAVGTNTI
GSSEACMLGGMAMKWRWRKRMEAAGKPTDKPNLVCGPVQICWHKFARYWDVELREIPMRPGQ
LFMDPKRMIEACDENTIGVVPTFGVTYTGNYEFPQPLHDALDKFQADTGIDIDMHIDAASGG
FLAPFVAPDIVWDFRLPRVKSISASGHKFGLAPLGCGWVIWRDEEALPQELVFNVDYLGGQI
GTFAINFSRPAGQVIAQYYEFLRLGREGYTKVQNASYQVAAYLADEIAKLGPYEFICTGRPD
EGIPAVCFKLKDGEDPGYTLYDLSERLRLRGWQVPAFTLGGEATDIVVMRIMCRRGFEMDFA
ELLLEDYKASLKYLSDHPKLQGIAQQNSFKHT
111000000000000000000000000000000000000000000000000000000000
000000000000000000000000000000000000000000000000000000000000
000000000000000000000000000000000000000000000000000000000000
000000000000000000000000000000000000000000000000000000000000
000000000000000000000000000000000000000000000000000000000000
000000000000000000000000000000000000000000000000000000000000
000000000000000000000000000000000000000000000000000000000000
0000000000000000001111111111111
>DM_validata128
MEPSSLELPADTVQRIAAELKCHPTDERVALHLDEEDKLRHFRECFYIPKIQDLPPVDLSLV
NKDENAIYFLGNSLGLQPKMVKTYLEEELDKWAKIAAYGHEVGKRPWITGDESIVGLMKDIV
GANEKEIALMNALTVNLHLLMLSFFKPTPKRYKILLEAKAFPSDHYAIESQLQLHGLNIEES
MRMIKPREGEETLRIEDILEVIEKEGDSIAVILFSGVHFYTGQHFNIPAITKAGQAKGCYVG
FDLAHAVGNVELYLHDWGVDFACWCSYKYLNAGAGGIAGAFIHEKHAHTIKPALVGWFGHEL
STRFKMDNKLQLIPGVCGFRISNPPILLVCSLHASLEIFKQATMKALRKKSVLLTGYLEYLI
KHNYGKDKAATKKPVVNIITPSHVEERGCQLTITFSVPNKDVFQELEKRGVVCDKRNPNGIR
VAPVPLYNSFHDVYKFTNLLTSILDSAETKN
111110000000000000000000000000000000000000000000000000000000
000000000000000000000000000000000000000000000000000000000000
000000000000000000000000000000000000000000000000000000000000
000000000000000000000000000000000000000000000000000000000000
000000000000000000000000000000000000000000000000000000000000
000000000000000000000000000000000000000000000000000000000000
000001111111110000000000000000000000000000000000000000000000
0000000000000000000000000011111
>DM_validata132
STQEQTPQICVVGSGPAGFYTAQHLLKHHSRAHVDIYEKQLVPFGLVRFGVAPDHPEVKNVI
NTFTQTARSDRCAFYGNVEVGRDVTVQELQDAYHAVVLSYGAEDHQALDIPGEELPGVFSAR
AFVGWYNGLPENRELAPDLSCDTAVILGQGNVALDVARILLTPPDHLEKTDITEAALGALRQ
SRVKTVWIVGRRGPLQVAFTIKELREMIQLPGTRPMLDPADFLGLQDRIKEAARPRKRLMEL

LLRTATEKPGVEEAARRASASRAWGLRFFRSPQQVLPSPDGRRAAGIRLAVTRLEGIGEATR
AVPTGDVEDLPCGLVLSSIGYKSRPIDPSVPFDPKLGVVPNMEGRVVDVPGLYCSGWVKRGP
TGVITTTMTDSFLTGQILLQDLKAGHLPSGPRPGSAFIKALLDSRGVWPVSFSDWEKLDAEE
VSRGQASGKPREKLLDPQEMLRLLGH
111110000000000000000000000000000000000000000000000000000000
000000000000000000000000000000000000000000000000000000000000
000000000000000000000000000000000000000000000000000000000000
000000000000000000000000000000000000000000000000000000000000
000000000000000000000000000000000000000000000000000000000000
000000000000000000000000000000000000000000000000000000000000
000000000000000000000000000000000000000000000000000000000000
00000000000000000000000000
>DM_validata133
MGSSSSGLVPRGSHMFTTKLAEKVVSAWKAKISQPALKAAQDGVIDTVAAALGGVTEHSVQV
ALKYVAATGGSGDSKLWGVNQRSNMFDAAFVNGMAAHAIDFDDSFPVMRGHPSSSLVPAIFA
VGEHVGANGHNCLKSYVLGIEVVATLGRAVGKGHYLAGWHPTSTLGVFGATTAAALLLGADE
EQLRNAWGIAASNSCGIIKNFGTMTKPMHTGSAARNGVLSAWLSMQSFTGCQTVFDDAEGIL
AMYGAQPGPELFNAMQKFGTPWAIIAPGLYKKSWPSCYANHKPLAGLFAIMKEHGLTGQDIS
HVDVGFLPGVEKPLLYMDPRTTEEAKFSIEANIGAALLDGEVSLASFEIEHLDRPAMRAAMK
KVTRFDMPSETTFSGTTGYTDIVVHTADGKIERRIEATPGSLEDPMDDAHLERKFKDCTAWM
PFGESGLLFDRLRSLTADQGIKTVQP
111111111111100000000000000000000000000000000000000000000000
000000000000000000000000000000000000000000000000000000000000
000000000000000000000000000000000000000000000000000000000000
000000000000000000000000000000000000000000000000000000000000
000000000000000000000000000000000000000000000000000000000000
000000000000000000000000000000000000000000000000000000000000
000000000000000000000000000000000000000000000000000000000000
00000000000000000000000000
>DM_validata134
MAATSSTSSQSFDIELDIIGQQPPLLSIYTQISLVYPVSDPSQYPTIVSTLEEGLKRLSQTF
PWVAGQVKTEGISEGNTGTSKIIPYEETPRLVVKDLRDDSSAPTIEGLRKAGFPLEMFDENV
VAPRKTLAIGPGNGPNDPKPVLLLQLNFIKGGLILTVNGQHGAMDMTGQDAIIRLLSKACRN
ESFTEEEISAMNLDRKTVVPLLENYKVGPELDHQIAKPAPAGDAPPAPAKATWAFFSFTPKA
LSELKDAATKTLDASSKFVSTDDALSAFIWQSTSRVRLARLDASTPTEFCRAVDMRGPMGVS
STYPGLLQNMTYHDSTVAEIANEPLGATASRLRSELNSDRLRRRTQALATYMHGLPDKSSVS
LTADANPSSSIMLSSWAKVGCWEYDFGFGLGKPESVRRPRFEPFESLMYFMPKKPDGEFTAS
ISLRDEDMERLKADEEWTKYAKYIG
111111111110000000000000000000000000000000000000000000000000
000000000001110000000000000000000000000000000000000000000000
000000001111111110000000000000000000000000000000000000000000
000000000000000000000011110000000111111111111111100000000000
000001111111111100000000000000000111111111000000000000000000
000000000000000000000000000000000000000000000000000000000000

000111000000000000000000000000000000000000000000000000000000
0000000000000000000000000
>DM_validata135
MTFRKSFDCYDFYDRAKVGEKCTQDDWDLMKIPMKAMELKQKYGLDFKGEFIPTDKDMMEKL
FKAGFEMLLECGIYCTDTHRIVKYTEDEIWDAINNVQKEFVLGTGRDAVNVRKRSVGDKAKP
IVQGGPTGSPISEDVFMPVHMSYALEKEVDTIVNGVMTSVRGKSPIPKSPYEVLAAKTETRL
IKNACAMAGRPGMGVKGPETSLSAQGNISADCTGGMTCTDSHEVSQLNELKIDLDAISVIAH
YKGNSDIIMDEQMPIFGGYAGGIEETTIVDVATHINAVLMSSASWHLDGPVHIRWGSTNTRE
TLMIAGWACATISEFTDILSGNQYYPCAGPCTEMCLLEASAQSITDTASGREILSGVASAKG
VVTDKTTGMEARMMGEVARATAGVEISEVNVILDKLVSLYEKNYASAPAGKTFQECYDVKTV
TPTEEYMQVYDGARKKLEDLGLVF
100000000000000000000000000000000000000000000000000000000000
000000000000000000000000000000000000000000000000000000000000
000000000000000000000000000000000000000000000000000000000000
000000000000000000000000000000000000000000000000000000000000
000000000000000000000000000000000000000000000000000000000000
000000000000000000000000000000000000000000000000000000000000
000000000000000000000000000000000000000000000000000000000000
000000000000000000000000
>DM_validata136
MITMTNWESLYEKALDKVEASIRKVRGVLLAYNTNIDAIKYLKREDLEKRIEKVGKEEVLRY
SEELPKEIETIPQLLGSILWSIKRGKAAELLVVSREVREYMRKWGWDELRMGGQVGIMANLL
GGVYGIPVIAHVPQLSELQASLFLDGPIYVPTFERGELRLIHPREFRKGEEDCIHYIYEFPR
NFKVLDFEAPRENRFIGAADDYNPILYVREEWIERFEEIAKRSELAIISGLHPLTQENHGKP
IKLVREHLKILNDLGIRAHLEFAFTPDEVVRLEIVKLLKHFYSVGLNEVELASVVSVMGEKE
LAERIISKDPADPIAVIEGLLKLIKETGVKRIHFHTYGYYLALTREKGEHVRDALLFSALAA
ATKAMKGNIEKLSDIREGLAVPIGEQGLEVEKILEKEFSLRDGIGSIEDYQLTFIPTKVVKK
PKSTVGIGDTISSSAFVSEFSLH
111111000000000000000000000000000000000000000000000000000000
000000000000000000000000000000000000000000000000000000000000
000000000000000000000000000000011111100000000111000000000000
000000000000000000000000000000000000000000000000000000000000
000000000000000000000000000000000000000000000000000000000000
000000000000000000000000000000000000000000000000000000000000
000000000000000000000000000000000000000000000000000000001111
11111000000000000000000
>DM_validata137
GSIFKPAPHKARLPAAEIDPTYRRLRWQIFLGIFFGYAAYYLVRKNFALAMPYLVEQGFSRG
DLGFALSGISIAYGFSKFIMGSVSDRSNPRVFLPAGLILAAAVMLFMGFVPWATSSIAVMFV
LLFLCGWFQGMGWPPCGRTMVHWWSQKERGGIVSVWNCAHNVGGGIPPLLFLLGMAWFNDWH
AALYMPAFCAILVALFAFAMMRDTPQSCGLPPIEEYKNDYPDDYNEKAEQELTAKQIFMQYV
LPNKLLWYIAIANVFVYLLRYGILDWSPTYLKEVKHFALDKSSWAYFLYEYAGIPGTLLCGW
MSDKVFRGNRGATGVFFMTLVTIATIVYWMNPAGNPTVDMICMIVIGFLIYGPVMLIGLHAL
ELAPKKAAGTAAGFTGLFGYLGGSVAASAIVGYTVDFFGWDGGFMVMIGGSILAVILLIVVM

IGEKRRHEQLLQELVPR
11100000000000000000000000000000000000000000000000000000000000
00000000000000000000000000000000000000000000000000000000000000
00000000000000000000000000000000000000000000000000000000000000
00000000000000000000000000000000000000011111111111110000000000
00000000000000000000000000000000000000000000000000000000000000
00000000000000000000000000000000000000000000000000000000000000
00000000000000000000000000000000000000000000000000000000000000
00000000000000000
>DM_validata138
RSPVFTDNSSPPAVPQSFQVAHLHAPTGSGKSTKVPAAYAAQGYKVLVLNPSVAATLGFGAY
MSKAHGVDPNIRTGVRTITTGSPITYSTYGKFLADGGCSGGAYDIIICDECHSTDATSILGI
GTVLDQAETAGARLVVLATATPPGSVTVSHPNIEEVALSTTGEIPFYGKAIPLEVIKGGRHL
IFCHSKKKCDELAAKLVALGINAVAYYRGLDVSVIPTNGDVVVVSTDALMTGFTGDFDSVID
CNTCVTQTVDFSLDPTFTIETTTLPQDAVSRTQRRGRTGRGKPGIYRFVAPGERPSGMFDSS
VLCECYDAGCAWYELMPAETTVRLRAYMNTPGLPVCQDHLEFWEGVFTGLTHIDAHFLSQTK
QSGENFPYLVAYQATVCARAQAPPPSWDQMWKCLIRLKPTLHGPTPLLYRLGAVQNEVTLTH
PITKYIMTCMSADLEVV
11111111000000000000000000000000000000000000000000000111111111
11111111111111111111000000000000000000000000000000000000000000
00000000000000000000000000000000000000000000000000000000000000
00000000000000000000000000000000000000000000000000000000000000
00000000000000000000000000000000000000000000000000000000000000
00000000000000000000000000000000000000000000000000000000000000
00000000000000000000000000000000000000000000000000000000000000
00000000000000000
>DM_validata140
EIPLKYGATNEGKRQDPAMQKFRDNRLGAFIHWGLYAIPGGEWNGKVYGGAAEWLKSWAKVP
ADEWLKLMDQWNPTKFDAKKWAKMAKEMGTKYVKITTKHHEGFCLWPSKYTKYTVANTPYKR
DILGELVKAYNDEGIDVHFYFSVMDWSNPDYRYDIKSKEDSIAFSRFLEFTDNQLKELATRY
PTVKDFWFDGTWDASVKKNGWWTAHAEQMLKELVPGVAINSRLRADDKGKRHFDSNGRLMGD
YESGYERRLPDPVKDLKVTQWDWEACMTIPENQWGYHKDWSLSYVKTPIEVIDRIVHAVSMG
GNMVVNFGPQADGDFRPEEKAMATAIGKWMNRYGKAVYACDYAGFEKQDWGYYTRGKNDEVY
MVVFNQPYSERLIVKTPKGITVEKATLLTTGEDITVVETTRNEYNVSVPKKNPGEPYVIQLK
VRAAKGTKSIYRDALT
00000000000000000000000000000000000000000000000000000000000000
00000000000000000000000000000000000000000000000000000000000000
00000000000000000000000000000000000000000000000000000000000000
00000000000000000000000000000000000000000000000000000000000000
00000000000000000000000000000000000000000000000000000000000000
00000000000000000000000000000000000000000000000000000000000000
00000000000000000000000000000000000000000000000000000000000000
0001111111111111
>DM_validata141

KMTWTMKAAEEAEAVANINCSEHGRAFLDGIISEGSPKCECNTCYTGPDCSEKIQGCSADVA
SGDGLFLEEYWKQHKEASAVLVSPWHRMSYFFNPVSNFISFELEKTIKELHEVVGNAAAKDR
YIVFGVGVTQLIHGLVISLSPNMTATPDAPESKVVAHAPFYPVFREQTKYFDKKGYVWAGNA
ANYVNVSNPEQYIEMVTSPNNPEGLLRHAVIKGCKSIYDMVYYWPHYTPIKYKADEDILLFT
MSKFTGHSGSRFGWALIKDESVYNNLLNYMTKNTEGTPRETQLRSLKVLKEVVAMVKTQKGT
MRDLNTFGFKKLRERWVNITALLDQSDRFSYQELPQSEYCNYFRRMRPPSPSYAWVKCEWEE
DKDCYQTFQNGRINTQNGVGFEASSRYVRLSLIKTQDDFDQLMYYLKDMVKAKRKTPLIKQL
FIDQTETASRRPFI
00000000000000000000000000000000000000000000000000000000000000
00000000000000000000000000000000000000000000000000000000000000
00000000000000000000000000000000000000000000000000000000000000
00000000000000000000000000000000000000000000000000000000000000
00000000000000000000000000000000000000000000000000000000000000
00000000000000000000000000000000000000000000000000000000000000
00000000000000000000000000000000000000000000000000000111111111
11111111111111
>DM_validata142
MTEPSLSPAVQTFWKWLQEEGVITAKTPVKASVVTEGLGLVALKDISRNDVILQVPKRLWIN
PDAVAASEIGRVCSELKPWLSVILFLIRERSREDSVWKHYFGILPQETDSTIYWSEEELQEL
QGSQLLKTTVSVKEYVKNECLKLEQEIILPNKRLFPDPVTLDDFFWAFGILRSRAFSRLRNE
NLVVVPMADLINHSAGVTTEDHAYEVKGAAGLFSWDYLFSLKSPLSVKAGEQVYIQYDLNKS
NAELALDYGFIEPNENRHAYTLTLEISESDPFFDDKLDVAESNGFAQTAYFDIFYNRTLPPG
LLPYLRLVALGGTDAFLLESLFRDTIWGHLELSVSRDNEELLCKAVREACKSALAGYHTTIE
QDRELKEGNLDSRLAIAVGIREGEKMVLQQIDGIFEQKELELDQLEYYQERRLKDLGLCGEN
GDILENLYFQ
11111000000000000000000000000000000000000000000000000000000000
00000000000000000000000000000000000000000000000000000000000000
00000000000000000000000000000000000000000000000000000000000111
00000000000000000000000000111111111100000000000000000000000000
00000000000000000000000000000000000000000000000000000000000000
00000000000000000000000000000000000000000000000000000000000000
00000000000000000000000000000000000000000000000000000000000000
0000000000
>DM_validata143
MMNWEHLLSLKRQGDTAKRLRIEQDDTRLGFEVDYDRIIFSAPFRSLQDKTQVIPLSKTDFV
HTRLTHSLEVSVVGRSLGRMVGKKLLEKYPHLEQVYGYKFNDFGAIVAAAALAHDIGNPPFG
HSGEKAIGEFFKNGYGKRYKDSLTAKEYQDLIKFEGNANGFKVLSQSKPGAQGGLRLSYATL
GAFMKYPKESLPHKPSDHIADKKYGFFQSERALFEDVAQELGLLKRSTTDDVSWSRHPLAYL
VEAADDICYTIIDFEDGINLGLIPEEYALEYMVKLVGQTIDRNKYNALQETSDRVSYLRALA
IGTLINESVDTFMKYEEEILAGTFDQSLIDKSNYQAQITDIINLSIERIYNSREVIEKEIAG
YEILSTLLEARCRALDNNDTHYNQLIQQLLAPNDHSEKSLYENLIQICAEVSTMTDGKALRN
YKKIKGLDVK
10000000000000000000000000000000000000000000000000000001110000
00000000000000000000000000000000000000000000000000000000000000

000000000000000000000000000000000000000000000000000000000000
000000000001111111111100000000000000000000001111000000000000
000000000000000000000000011111111111111111000000000000000000
000000000000000000000000000000000000000000000000000000000000
000000000000000000000000000000011111100000000000000000000000
0000000111
>DM_validata144
MAREAKDTVDLYDDRGNCVAEEVPIEVLSPMRNEAIQSIVNDIKRTVAVDLEGIENALQNAT
VGGKGMKIPGREMDVDIVDNAEAIADEIEKMIRVYQDDDTNVEPMYDGKRLLVQLPSERVKV
MADPYSGTLQAGMAVVHAIIDVCEVDMWDANMVKAAVFGRYPQTIDYFGGNVASMLDVPMKQ
EGVGYALRNIMVNHIVAATRKNTMQAVCLAATLQQTAMFEMGDALGPFERLHLLGYAYQGLN
ADNMVYDIVKKHGKEGTVGTVVREVVERALEDGVIEVKEELPSFKVYKANDMDLWNAYAAAG
LVAAVMVNQGAARAAQGVSATILYYNDLLEYETGLPGVDFGRAEGTAVGFSFFSHSIYGGGG
PGIFHGNHIVTRHSKGFAIPPVAAAMALDAGTQMFSPEVTSKLIGDVFGEIDEFREPMKYIT
EAAAEEAKR
111111000000000000000000000000000000000000000000000000000000
000000000000000000000000000000000000000000000000000000000000
000000000000000000000000000000000000000000000000000000000000
000000000000000000000000000000000000000000000000000000000000
000000000000000000000000000000000000000000000000000000000000
000000000000000000000000000000000000000000000000000000000000
000000000000000000000000000000000000000000000000000000000000
000000000
>DM_validata145
MGSDKIHHHHHHMSEPVIKSLLDTDMYKITMHAAVFTNFPDVTVTYKYTNRSSQLTFNKEAI
NWLKEQFSYLGNLRFTEEEIEYLKQEIPYLPSAYIKYISSSNYKLHPEEQISFTSEEIEGKP
THYKLKILVSGSWKDTILYEIPLLSLISEAYFKFVDIDWDYENQLEQAEKKAETLFDNGIRF
SEFGTRRRRSLKAQDLIMQGIMKAVNGNPDRNKSLLLGTSNILFAKKYGVKPIGTVAHEWVM
GVASISEDYLHANKNAMDCWINTFGAKNAGLALTDTFGTDDFLKSFRPPYSDAYVGVRQDSG
DPVEYTKKISHHYHDVLKLPKFSKIICYSDSLNVEKAITYSHAAKENGMLATFGIGTNFTND
FRKKSEPQVKSEPLNIVIKLLEVNGNHAIKISDNLGKNMGDPATVKRVKEELGYTERSWSGD
NEAHRWT
111111111110000000000000000000000000000000000000000000000000
000000000000000000000000000000000000000000000000000000000000
000000000000000000000000000000000000000000000000000000000000
000000000000000000000000000000000000000000000000000000000000
000000000000000000000000000000000000000000000000000000000000
000000000000000000000000000000000000000000000000000000000000
000000000000000000000000000000000000000000000000000001111111
1111111
>DM_validata146
MVAESNSAVAPTANVATSPAHEHFVRVNGGHFELQGKPYVITGVNMWYAAYLGAPNEVGDRD
RLAKELDNLKAIGVNNLRVLAVSEKSEINSAVKPAVTNGFGNYDETLLQGLDYLLVELAKRD
MTVVLYFNNFWQWSGGMTQYMAWIEGEPVQDPNVTNEWEAFMAKSASFYRSEKAQQEYRKTL

EKIITRVNSINGKAYVDDATIMSWQLANEPRPGNSQTTAEEKQIYIDWVHAAAAYIKTLDAH
HLVSSGSEGEMGSVNDMQVFIDAHATPDIDYLTYHMWIRNWSWFDKTKPAETWPSAWEKAQN
YMRAHIDVAKQLNKPLVLEEFGLDRDMGSYAMDSTTEYRDNYFRGVFELMLASLEQGEPSAG
YNIWAWNGYGRTTRANYWWQEGDDFMGDPPQEEQGMYGVFDTDTSTIAIMKEFNARFQPKLE
HHHHHH
11111111111111111111100000000000000000000000000000000000000000
00000000000000000000000000000000000000000000000000000000000000
00000000000000000000000000000000000000000000000000000000000000
00000000000000000000000000000000000000000000000000000000000000
00000000000000000000000000000000000000000000000000000000000000
00000000000000000000000000000000000000000000000000000000000000
00000000000000000000000000000000000000000000000000000000000111
111111
>DM_validata147
MAVTKSSSLLIVGAGTWGTSTALHLARRGYTNVTVLDPYPVPSAISAGNDVNKVISSGQYSN
NKDEIEVNEILAEEAFNGWKNDPLFKPYYHDTGLLMSACSQEGLDRLGVRVRPGEDPNLVEL
TRPEQFRKLAPEGVLQGDFPGWKGYFARSGAGWAHARNALVAAAREAQRMGVKFVTGTPQGR
VVTLIFENNDVKGAVTADGKIWRAERTFLCAGASAGQFLDFKNQLRPTAWTLVHIALKPEER
ALYKNIPVIFNIERGFFFEPDEERGEIKICDEHPGYTNMVQSADGTMMSIPFEKTQIPKEAE
TRVRALLKETMPQLADRPFSFARICWCADTANREFLIDRHPQYHSLVLGCGASGRGFKYLPS
IGNLIVDAMEGKVPQKIHELIKWNPDIAANRNWRDTLGRFGGPNRVMDFHDVKEWTNVQYRD
ISKL
10000000000000000000000000000000000000000000000000000000000000
00000000000000000000000000000000000000000000000000000000000000
00000000000000000000000000000000000000000000000000000000000000
00000000000000000000000000000000000000000000000000000000000000
00000000000000000000000000000000000000000000000000000000000000
00000000000000000000000000000000000000000000000000000000000000
00000000000000000000000000000000000000000000000000000000000000
0001
>DM_validata148
MADDPSAADRNVEIWKIKKLIKSLEAARGNGTSMISLIIPPKDQISRVAKMLADEFGTASNI
KSRVNRLSVLGAITSVQQRLKLYNKVPPNGLVVYCGTIVTEEGKEKKVNIDFEPFKPINTSL
YLCDNKFHTEALTALLSDDSKFGFIVIDGSGALFGTLQGNTREVLHKFTVDLPKKHGRGGQS
ALRFARLRMEKRHNYVRKVAETAVQLFISGDKVNVAGLVLAGSADFKTELSQSDMFDQRLQS
KVLKLVDISYGGENGFNQAIELSTEVLSNVKFIQEKKLIGRYFDEISQDTGKYCFGVEDTLK
ALEMGAVEILIVYENLDIMRYVLHCQGTEEEKILYLTPEQEKDKSHFTDKETGQEHELIESM
PLLEWFANNYKKFGATLEIVTDKSQEGSQFVKGFGGIGGILRYRVDFQGMEYQGGDDEFFDL
DDY
11110000000000000000000000000000000000000000000000000000000000
00000000000000000000000000000000000000000000000000000000000000
00000000000000000000000000000000000000000000000000000000000000
00000000000000000000000000000000000000000000000000000000000000
00000000000000000000000000000000000000000000000000000000000000

00000000000000000000000111111111000000000000000001111111111000
0000000000000000000000000000000000000000000000000111111111111111
111
>DM_validata149
GMNFPIDEKLIREKQNELHIKDLGMASIRDLVALVTNLEKATGTKFCRMEMGVPGLPAPQIG
IETEIQKLREGVASIYPNLDGLPELKQEASRFAKLFVNIDIPARACVPTVGSMQGCFVSFLV
ANRTHKNREYGTLFIDPGFNLNKLQCRILGQKFESFDLFEYRGEKLREKLESYLQTGQFCSI
IYSNPNNPTWQCMTDEELRIIGELATKHDVIVIEDLAYFGMDFRKDYSHPGEPLYQPSVANY
TDNYILALSSSKAFSYAGQRIGVLMISGKLYEREYPDLEESFGRLRFGEALSSSALYALSSG
ATHSAQWGMAAMLKACNDGEYNFRDSVIEYGRKARIMKKMFLDNGFNIVYDKDGNEPLADGF
YFTVGYKGMDSSKLIEKFVRYGMCAITLKTTGSKRNEAMRICTSLLPESQFPDLEKRLQMLN
AEG
11000000000000000000000000000000000000000000000000000000000000
00000000000000000000000000000000000000000000000000000000000000
00000000000000000000000000000000000000000000000000000000000000
00000000000000000000000000000000000000000000000000000000000000
00000000000000000000000000000000000000000000000000000000000000
00000000000000000000000000000000000000000000000000000000000000
00000000000000000000000000000000000000000000000000000000000000
000
>DM_validata151
SPAIMLQRDFITALPARGLDHIAENILSYLDAKSLCAAELVCKEWYRVTSDGMLWKKLIERM
VRTDSLWRGLAERRGWGQYLFKNKPPDGNAPPNSFYRALYPKIIQDIETIESNWRCGRHSLQ
RIHCRSETSKGVYCLQYDDQKIVSGLRDNTIKIWDKNTLECKRILTGHTGSVLCLQYDERVI
ITGSSDSTVRVWDVNTGEMLNTLIHHCEAVLHLRFNNGMMVTCSKDRSIAVWDMASPTDITL
RRVLVGHRAAVNVVDFDDKYIVSASGDRTIKVWNTSTCEFVRTLNGHKRGIACLQYRDRLVV
SGSSDNTIRLWDIECGACLRVLEGHEELVRCIRFDNKRIVSGAYDGKIKVWDLVAALDPRAP
AGTLCLRTLVEHSGRVFRLQFDEFQIVSSSHDDTILIWDFLNDPAAQAEPPRSPSRTYTYIS
R
00000000000000000000000000000000000000000000000000000000000000
00000000000000000000011111111100000000000000000000000000000000
00000000000000000000000000000000000000000000000000000000000000
00000000000000000000000000000000000000000000000000000000000000
00000000000000000000000000000000000000000000000000000000000000
00000000000000000000000000000000000000000000000000000000000000
00000000000000000000000000000000000000011111111111111111111111
1
>DM_validata152
MVVMIKLRDELGTATTDSAQKILLLGSGELGKEIAIEAQRLGVEVVAVDRYANAPAMQVAHR
SYVGNMMDKDFLWSVVEREKPDAIIPEIEAINLDALFEFEKDGYFVVPNARATWIAMHRERL
RETLVKEAKVPTSRYMYATTLDELYEACEKIGYPCHTKAIMSSSGKGSYFVKGPEDIPKAWE
EAKTKARGSAEKIIVEEHIDFDVEVTELAVRHFDENGEIVTTFPKPVGHYQIDGDYHASWQP
AEISEKAEREVYRIAKRITDVLGGLGIFGVEMFVKGDKVWANEVSPRPHDTGMVTLASHPPG
FSEFALHLRAVLGLPIPGEWVDGYRLFPMLIPAATHVIKAKVSGYSPRFRGLVKALSVPNAT

VRLFGKPEAYVGRRLGIALAWDKDVEVAKRKAEMVAHMIELRTRSSDWHDQNYEKRKHLLR
100000000000000000000000000000000000000000000000000000000000
000000000000000000000000000000000000000000000000000000000000
000000000000000000000000000000000000000001111110000000000000
011111111100000000000000000000000000000000000000000000000000
000000000000000000000000000000000000000000000000000000000000
000000000000000000000000000000000000000000000000000000000000
00000000000000000000000000000000000000000000000011111111111
>DM_validata153
FVRKTNKNRDMPLDSDVFRVPPGYNAPQQVHITQGDLVGRAMIISWVTMDEPGSSAVRYWSE
KNGRKRIAKGKMSTYRFFNYSSGFIHHTTIRKLKYNTKYYYEVGLRNTTRRFSFITPPQTGL
DVPYTFGLIGDLGQSFDSNTTLSHYELSPKKGQTVLFVGDLSYADRYPNHDNVRWDTWGRFT
ERSVAYQPWIWTAGNHEIEFAPEINETEPFKPFSYRYHVPYEASQSTSPFWYSIKRASAHII
VLSSYSAYGRGTPQYTWLKKELRKVKRSETPWLIVLMHSPLYNSYNHHFMEGEAMRTKFEAW
FVKYKVDVVFAGHVHAYERSERVSNIAYKITDGLCTPVKDQSAPVYITIGDAGNYGVIDSNM
IQPQPEYSAFREASFGHGMFDIKNRTHAHFSWNRNQDGVAVEADSVWFFNRHWYPVDDST
111111110000000000000000000000000000000000000000000000000000
000000000000000000000000000000000000000000000000000000000000
000000000000000000000000000000000000000000000000000000000000
000000000000000000000000000000000000000000000000000000000000
000000000000000000000000000000000000000000000000000000000000
000000000000000000000000000000000000000000000000000000000000
0000000000000000000000000000000000000000000000000000000000
>DM_validata154
GSHMTIQKVHGREVLDSRGNPTVEVEVTTEKGVFRSAVPSGASTGVYEACELRDGDKKRYVG
KGCLQAVKNVNEVIGPALIGRDELKQEELDTLMLRLDGTPNKGKLGANAILGCSMAISKAAA
AAKGVPLYRYLASLAGTKELRLPVPCFNVINGGKHAGNALPFQEFMIAPVKATSFSEALRMG
SEVYHSLKGIIKKKYGQDAVNVGDEGGFAPPIKDINEPLPILMEAIEEAGHRGKFAICMDCA
ASETYDEKKQQYNLTFKSPEPTWVTAEQLRETYCKWAHDYPIVSIEDPYDQDDFAGFAGITE
ALKGKTQIVGDDLTVTNTERIKMAIEKKACNSLLLKINQIGTISEAIASSKLCMENGWSVMV
SHRSGETEDTYIADLVVALGSGQIKTGAPCRGERTAKLNQLLRIEEELGAHAKFGFPGWS
000000000000000000000000000000000000000000000000000000000000
000000000000000000000000000000000000000000000000000000000000
000000000000000000000000000000000000000000000000000000000000
000000000000000000000000000000000000000000000000000000000000
000000000000001111111000000000000000000000000000000000000000
000000000000000000000000000000000000000000000000000000000000
0000000000000000000000000000000000000000000000000000000000
>DM_validata155
GMMVPHLTTALTGPLLTLEKRLLDNMPRIEHWFRSQWQEYGAPFYASVDLRNAGFKLAPVDT
NLFPGGFNNLNPDFLPLCIQAAMVAVEKICPDARRLLLIPENHTRNTFYLRNVHALTHILRQ
AGLEVRIGSIAPEITAPTFLETHDGHSILLEPVRRKANRLELDNFDSCAILLNNDLSGGIPD
ILQGLEQSLIPPLHAGWATRRKSNHFTAYDRVVEEFAPLIDIDPWLLNPYFDTCGGLDFHAR
LGEEQLAEKVDSLLAKIRRKYAEYGVKQEPFVIVKADAGTYGMGIMTVKSADDVRDLNRKQR

NKMSVVKEGLKVSEVILQEGVYTFEHLKDAVAEPVIYMMDHFVVGGFYRVHTSRGADENLNA
PGMHFEPLTFETPCSTPDCAGAPDAAPNRFYAYGVVARLALLAATIELQETDPDLLDERT
00000000000000000000000000000000000000000000000000000000000000
00000000000000000000000000000000000000000000000000000000000000
00000000000000000000000000000000000000000000000000000000000000
00000000000000000000000000000000000000000000000000000000000000
00000000000000000000000000000000000000000000000000000000000000
00000000000000000000000000000000000000000000000000000000000000
000000000000000000000000000000000000000000000000000011111111
>DM_validata156
MSNVQTSAEREIVDLSHLAFDCGMLGRLKTVSWTPVIAGDSFELDAVGALRLSPLRRGLAID
SKVDFFTFYIPHRHVYGDQWIQFMRDGVNAQPLPSVTCNRYPDHAGYVGTIVPANNRIPKFL
HQSYLNIYNNYFRAPWMPERTEANPSNLNEDDARYGFRCCHLKNIWSAPLPPETKLAEEMGI
ESNSIDIMGLQAAYAQLHTEQERTYFMQRYRDVISSFGGSTSYDADNRPLLVMHTDFWASGY
DVDGTDQSSLGQFSGRVQQTFKHSVPRFFVPEHGVMMTLALIRFPPISPLEHHYLAGKSQLT
YTDLAGDPALIGNLPPREISYRDLFRDGRSGIKIKVAESIWYRTHPDYVNFKYHDLHGFPFL
DDAPGTSTGDNLQEAILVRHQDYDACFQSQQLLQWNKQARYNVSVYRHMPTVRDSIMTS
11111111100000000000000000000000000000000000000000000000000000
00000000000000000000000000000000000000000000000000000000000000
00000000000000000000000000000000000000000000000000000000000000
00000000000000000000000000000000000000000000000000000000000000
00000000000000000000000000000000000000000000000000000000000000
00000000000000000000000000000000000000000000000000000000000000
00000000000000000000000000000000000000000000000000000000000
>DM_validata157
MAKHVVVIGGGVGGIATAYNLRNLMPDLKITLISDRPYFGFTPAFPHLAMGWRKFEDISVPL
APLLPKFNIEFINEKAESIDPDANTVTTQSGKKIEYDYLVIATGPKLVFGAEGQEENSTSIC
TAEHALETQKKLQELYANPGPVVIGAIPGVSCFGPAYEFALMLHYELKKRGIRYKVPMTFIT
SEPYLGHFGVGGIGASKRLVEDLFAERNIDWIANVAVKAIEPDKVIYEDLNGNTHEVPAKFT
MFMPSFQGPEVVASAGDKVANPANKMVIVNRCFQNPTYKNIFGVGVVTAIPPIEKTPIPTGV
PKTGMMIEQMAMAVAHNIVNDIRNNPDKYAPRLSAICIADFGEDAGFFFADPVIPPRERVIT
KMGKWAHYFKTAFEKYFLWKVRNGNIAPSFEEKVLEIFLKVHPIELCKDCEGAPGSRC
10000000000000000000000000000000000000000000000000000000000000
00000000000000000000000000000000000000000000000000000000000000
00000000000000000000000000000000000000000000000000000000000000
00000000000000000000000000000000000000000000000000000000000000
00000000000000000000000000000000000000000000000000000000000000
00000000000000000000000000000000000000000000000000000000000000
0000000000000000000000000000000000000000000000000000000000
>DM_validata158
GSSMANNALQTIINARLPGEEGLWQIHLQDGKISAIDAQSGVMPITENSLDAEQGLVIPPFV
EPHIHLDTTQTAGQPNWNQSGTLFEGIERWAERKALLTHDDVKQRAWQTLKWQIANGIQHVR
THVDVSDATLTALKAMLEVKQEVAPWIDLQIVAFPQEGILSYPNGEALLEEALRLGADVVGA
IPHFEFTREYGVESLHKTFALAQKYDRLIDVHCDEIDDEQSRFVETVAALAHHEGMGARVTA

SHTTAMHSYNGAYTSRLFRLLKMSGINFVANPLVNIHLQGRFDTYPKRRGITRVKEMLESGI
NVCFGHDGVFDPWYPLGTANMLQVLHMGLHVCQLMGYGQINDGLNLITHHSARTLNLQDYGI
AAGNSANLIILPAENGFDALRRQVPVRYSVRGGKVIASTQPAQTTVYLEQPEAIDYKR
111111100000000000000000000000000000000000000000000000000000
000000000000000000000000000000000000000000000000000000000000
000000000000000000000000000000000000000000000000000000000000
000000000000000000000000000000000000000000000000000000000000
000000000000000000000000000000000000000000000000000000000000
000000000000000000000000000000000000000000000000000000000000
0000000000000000000000000000000000000000000000000000000000
>DM_validata159
MESKAKTTVTFHSGILTIGGTVIEVAYKDAHIFFDFGTEFRPELDLPDDHIETLINNRLVPE
LKDLYDPRLGYEYHGAEDKDYQHTAVFLSHAHLDHSRMINYLDPAVPLYTLKETKMILNSLN
RKGDFLIPSPFEEKNFTREMIGLNKNDVIKVGEISVEIVPVDHDAYGASALLIRTPDHFITY
TGDLRLHGHNREETLAFCEKAKHTELLMMEGVSISFPEREPDPAQIAVVSEEDLVQHLVRLE
LENPNRQITFNGYPANVERFAKIIEKSPRTVVLEANMAALLLEVFGIEVRYYYAESGKIPEL
NPALEIPYDTLLKDKTDYLWQVVNQFDNLQEGSLYIHSDAQPLGDFDPQYRVFLDLLAKKDI
TFVRLACSGHAIPEDLDKIIALIEPQVLVPIHTLKPEKLENPYGERILPERGEQIVL
111100000000000000000000000000000000000000000000000000000000
000000000000000000000000000000000000000000000000000000000000
000000000000000000000000000000000000000000000000000000000000
000000000000000000000000000000000000000000000000000000000000
000000000000000000000000000000000000000000000000000000000000
000000000000000000000000000000000000000000000000000000000000
000000000000000000000000000000000000000000000000000000000
>DM_validata160
EGDVESIEKFLSTFKILPPLRDYKEFGPIQEIVRSPNMGNLRGKLIATLMENEPNSITSSAV
SPGETPYLITGSDQGVIKIWNLKEIIVGEVYSSSLTYDCSSTVTQITMIPNFDAFAVSSKDG
QIIVLKVNHYQQESEVKFLNCECIRKINLKNFGKNEYAVRMRAFVNEEKSLLVALTNLSRVI
IFDIRTLERLQIIENSPRHGAVSSICIDEECCVLILGTTRGIIDIWDIRFNVLIRSWSFGDH
APITHVEVCQFYGKNSVIVVGGSSKTFLTIWNFVKGHCQYAFINSDEQPSMEHFLPIEKGLE
ELNFCGIRSLNALSTISVSNDKILLTDEATSSIVMFSLNELSSSKAVISPSRFSDVFIPTQV
TANLTMLLRKMKRTSTHSVDDSLYHHDIINSISTCEVDETPLLVACDNSGLIGIFQ
111100000000001000000000000000000000000000000000000000000000
000000000000000000000000000000000000000000000000000000000000
000000000000000000000000000000000000000000000000000000000000
000000000000000000000000000000000000000000000000000000000000
000000000000000000000000000000000000000000000000000000000000
000000000000000000000000000000000000000000000000110000000000
00000000000011111111111110000000000000000000000000000000
>DM_validata161
MKFTNLTAKEFGAFTDSMPYSHFTQTVGHYELKLAEGYETHLVGIKNNNNEVIAACLLTAVP
VMKVFKYFYSNRGPVIDYENQELVHFFFNELSKYVKKHRCLYLHIDPYLPYQYLNHDGEITG
NAGNDWFFDKMSNLGFEHTGFHKGFDPVLQIRYHSVLDLKDKTADDIIKNMDGLRKRNTKKV

KKNGVKVRFLSEEELPIFRSFMEDTSESKAFADRDDKFYYNRLKYYKDRVLVPLAYINFDEY
IKELNEERDILNKDLNKALKDIEKRPENKKAHNKRDNLQQQLDANEQKIEEGKRLQEEHGNE
LPISAGFFFINPFEVVYYAGGTSNAFRHFAGSYAVQWEMINYALNHGIDRYNFYGVSGKFTE
DAEDAGVVKFKKGYNAEIIEYVGDFIKPINKPVYAAYTALKKVKDRIFHHHHHH
00000000000000000000000000000000000000000000000000000000000000
00000000000000000000000000000000000000000000000000000000000000
00000000000000000000000000000000000000000000000000000000000000
00000000000000000000001111111111100000000000000000000000000000
00000000000000000000000000000000000000000000000000000000000000
00000000000000000000000000000000000000000000000000000000000000
000000000000000000000000000000000000000011111111111111
>DM_validata163
MKTVVIGAGLGGLLSAARLSKAGHEVEVFERLPITGGRFTNLSYKGFQLSSGAFHMLPNGPG
GPLACFLKEVEASVNIVRSEMTTVRVPLKKGNPDYVKGFKDISFNDFPSLLSYKDRMKIALL
IVSTRKNRPSGSSLQAWIKSQVSDEWLIKFADSFCGWALSLKSDEVPVEEVFEIIENMYRFG
GTGIPEGGCKGIIDALETVISANGGKIHTGQEVSKILIENGKAAGIIADDRIHDADLVISNL
GHAATAVLCSEALSKEADAAYFKMVGTLQPSAGIKICLAADEPLVGHTGVLLTPYTRRINGV
NEVTQADPELAPPGKHLTMCHQYVAPENVKNLESEIEMGLEDLKEIFPGKRYEVLLIQSYHD
EWPVNRAASGTDPGNETPFSGLYVVGDGAKGKGGIEVEGVALGVMSVMEKVLG
00000000000000000000000000000000000000000000000000000000000000
00000000000000000000000000000000000000000000000000000000000000
00000000000000000000000000000000000000000000000000000000000000
00000000000000000000000000000000000000000000000000000000000000
00000000000000000000000000000000000000000000000000000000000000
00000000000000000000000000000000000000000000000000000000000000
00000000000000000000000000000000000000000000000000111
>DM_validata165
FRRKEDLDPPLALLPLKGLREAAALLEEALRQGKRIRVHGDYDADGLTGTAILVRGLAALGA
DVHPFIPHRLEEGYGVLMERVPEHLEASDLFLTVDCGITNHAELRELLENGVEVIVTDHHTP
GKTPPPGLVVHPALTPDLKEKPTGAGVAFLLLWALHERLGLPPPLEYADLAAVGTIADVAPL
WGWNRALVKEGLARIPASSWVGLRLLAEAVGYTGKAVEVAFRIAPRINAASRLGEAEKALRL
LLTDDAAEAQALVGELHRLNARRQTLEEAMLRKLLPQADPEAKAIVLLDPEGHPGVMGIVAS
RILEATLRPVFLVAQGKGTVRSLAPISAVEALRSAEDLLLRYGGHKEAAGFAMDEALFPAFK
ARVEAYAARFPDPVREVALLDLLPEPGLLPQVFRELALLEPYGEGNPEPLFL
11111111100000000000000000000000000000000000000000000000000000
00000000000000000000000000000000000000000000000000000000000000
00000000000000000000000000000000000000000000000000000000000000
00000000000000000000000000000000000000000000000000000000000000
00000000000000000000000000000000000000000000000000000000000000
00000000000000000000000000000000000000000000000000000000000000
0000000000000000000000111111111111111111111111111111
>DM_validata166
SKSKPDTSAPTSPKVTYKAPVPSGEVYFADSFDRGTLSGWILSKAKKDDTDDEIAKYDGKWE
VDEMKETKLPGDKGLVLMSRAKHHAISAKLNKPFLFDTKPLIVQYEVNFQNGIECGGAYVKL

LSKTPELNLDQFHDKTPYTIMFGPDKCGEDYKLHFIFRHKNPKTGVYEEKHAKRPDADLKTY
FTDKKTHLYTLILNPDNSFEILVDQSIVNSGNLLNDMTPPVNPSREIEDPEDQKPEDWDERP
KIPDPDAVKPDDWNEDAPAKIPDEEATKPDGWLDDEPEYVPDPDAEKPEDWDEDMDGEWEAP
QIANPKCESAPGCGVWQRPMIDNPNYKGKWKPPMIDNPNYQGIWKPRKIPNPDFFEDLEPFK
MTPFSAIGLELWSMTSDIFFDNFIVCGDRRVVDDWANDGWGLKKAADGAAEP
11111111111111110000000000000000000000000000001111111111100000
00000000000000000000000000000000000000000000000000000000000000
00000000000000000000000000000000000000000000000000000000000000
00000000000000000000000000000000111111100000000000000000000000
00000000000000000000000000000000000000000000000000000000000000
00000000000000000000000000000000000000000000000000000000000000
0000000000000000000000000000000000000000001111111111
>DM_validata167
MGFKLVGFSKFVRKNPKSDKFKVKRFHHIEFWCGDATNVARRFSWGLGMRFSAKSDLSTGNM
VHASYLLTSGDLRFLFTAPYSPSLSAGEIKPTTTASIPSFDHGSCRSFFSSHGLGVRAVAIE
VEDAESAFSISVANGAIPSSPPIVLNEAVTIAEVKLYGDVVLRYVSYKAEDTEKSEFLPGFE
RVEDASSFPLDYGIRRLDHAVGNVPELGPALTYVAGFTGFHQFAEFTADDVGTAESGLNSAV
LASNDEMVLLPINEPVHGTKRKSQIQTYLEHNEGAGLQHLALMSEDIFRTLREMRKRSSIGG
FDFMPSPPPTYYQNLKKRVGDVLSDDQIKECEELGILVDRDDQGTLLQIFTKPLGDRPTIFI
EIIQRVGCMMKDEEGKAYQSGGCGGFGKGNFSELFKSIEEYEKTLEAKQLVG
11111111111111000000000000000000000000000000000000000000000000
00000000000000000000000000000000000000000000000000000000000000
00000000000000000000000000000000000000000000000011111110000000
00000000000000000000000000000000000000000000111111111110000000
00000000000000000011100000000000000000000000000000000000000000
00000000000000000000000000000000000000000000000000000000000000
0000000001111111110000000000000000001111111111111111
>DM_validata169
MKKVMLATALFLGLTPAGANAADLGHQTLGSNDGWGAYSTGTTGGSKASSSNVYTVSNRNQL
VSALGKETNTTPKIIYIKGTIDMNVDDNLKPLGLNDYKDPEYDLDKYLKAYDPSTWGKKEPS
GTQEEARARSQKNQKARVMVDIPANTTIVGSGTNAKVVGGNFQIKSDNVIIRNIEFQDAYDY
FPQWDPTDGSSGNWNSQYDNITINGGTHIWIDHCTFNDGSRPDSTSPKYYGRKYQHHDGQTD
ASNGANYITMSYNYYHDHDKSSIFGSSDSKTSDDGKLKITLHHNRYKNIVQKAPRVRFGQVH
VYNNYYEGSTSSSSYPFSYAWGIGKSSKIYAQNNVIDVPGLSAAKTISVFSGGTALYDSGTL
LNGTQINASAANGLSSSVGWTPSLHGSIDASANVKSNVINQAGAGKLN
11111111111111111111100000000000000000000000000000000000000000
00000000000000000000000000000000000000000000000000000000000000
00000000000000000000000000000000000000000000000000000000000000
00000000000000000000000000000000000000000000000000000000000000
00000000000000000000000000000000000000000000000000000000000000
00000000000000000000000000000000000000000000000000000000000000
000000000000000000000000000000000000000000000000
>DM_validata170
MQEGAYRFIRNPNVSAEAIRKAGAMQTVKLAQEFPELLAIEDTTSLSYRHQVAEELGKLGSI

QDKSRGWWVHSVLLLEATTFRTVGLLHQEWWMRPDDPADADEKESGKWLAAAATSRLRMGSM
MSNVIAVCDREADIHAYLQDKLAHNERFVVRSKHPRKDVESGLYLYDHLKNQPELGGYQISI
PQKGVVDKRGKRKNRPARKASLSLRSGRITLKQGNITLNAVLAEEINPPKGETPLKWLLLTS
EPVESLAQALRVIDIYTHRWRIEEFHKAWKTGAGAERQRMEEPDNLERMVSILSFVAVRLLQ
LRESFTLPQALRAQGLLKEAEHVESQSAETVLTPDECQLLGYLDKGKRKRKEKGSLQWAYMA
IARLGGFMDSKRTGIASWGALWEGWEALQSKLDGFLAAKDLMAQGIKI
111111111111110000000000000000000000000000000001111111111111
111111000000000000000000000000000000000000000000000000000000
000000000000000000000000000000000000000000000000000000000000
000001111111110000000000000000000000000000000000000000000000
000000000000000000000000000000000000000000000000000000000000
000000000000000000000000000000000000000000000000000000000000
000000000000000000000000000000000000000000001111
>DM_validata171
MELADVGAAASSQGVHDQVLPTPNASSRVIVHVDLDCFYAQVEMISNPELKDKPLGVQQKYL
VVTCNYEARKLGVKKLMNVRDAKEKCPQLVLVNGEDLTRYREMSYKVTELLEEFSPVVERLG
FDENFVDLTEMVEKRLQQLQSDELSAVTVSGHVYNNQSINLLDVLHIRLLVGSQIAAEMREA
MYNQLGLTGCAGVASNKLLAKLVSGVFKPNQQTVLLPESCQHLIHSLNHIKEIPGIGYKTAK
CLEALGINSVRDLQTFSPKILEKELGISVAQRIQKLSFGEDNSPVILSGPPQSFSEEDSFKK
CSSEVEAKNKIEELLASLLNRVCQDGRKPHTVRLIIRRYSSEKHYGRESRQCPIPSHVIQKL
GTGNYDVMTPMVDILMKLFRNMVNVKMPFHLTLLSVCFCNLKALNTAK
111111111111111111111111100000000000000000000000000000000000
000000000000000000000000000000000000000000000000000000000000
000000000000000000000000000000000000000000000000000000000000
000000000000000000000000000000000000000000000000000000000000
000000000000000000000000000000000000000000000000000000000000
000000000000000000000001110000000000000111110000000000000001
111110000000000000000000000000000000000000111111
>DM_validata173
EETIPLQTLRCYNDYTSHITCRWADTQDAQRLVNVTLIRRVNEDLLEPVSCDLSDDMPWSAC
PHPRCVPRRCVIPCQSFVVTDVDYFSFQPDRPLGTRLTVTLTQHVQPPEPRDLQISTDQDHF
LLTWSVALGSPQSHWLSPGDLEFEVVYKRLQDSWEDAAILLSNTSQATLGPEHLMPSSTYVA
RVRTRLAPGSRLSGRPSKWSPEVCWDSQPGDEAQPQNLECFFDGAAVLSCSWEVRKEVASSV
SFGLFYKPSPDAGSAVLLREEECSPVLREGLGSLHTRHHCQIPVPDPATHGQYIVSVQPRRA
EKHIKSSVNIQMAPPSLQVTKDGDSYSLRWETMKMRYEHIDHTFEIQYRKDTATWKDSKTET
LQNAHSMALPALEPSTRYWARVRVRTSRTGYNGIWSEWSEARSWDTE
000000000000000000000000000000000000000000000000000000000000
000000000000000000000000000000000000000000000000000000000000
000000000000000000000000000000000000000000000000000000000000
000000000000000000000000000000000000000000000000000000000000
000000000011111111000000000000000000000000000000000000000000
000000000000000000000000000000000000000000000000000000000000
00000000000000000000000000000000000000000000111
>DM_validata175

MDTAGDTKTTDDGGVVNITYMHRLPDSEGMTLVNDIVAKWNKQHPDIQVKATKFDGKASDMI
KKLETDVKSGEAPDLAQVGYAELPEVFTKGLLQDVTQYAEQYKNDFASGPYSLVQVGGKAYG
LPQDTGPLVYFYNKAEFEKLGITEIPQTADEFIAAAKTAAAAGKYIMSYQPDEAGNMISGLA
GASGGWYKVKGDSWVVNTETDGSKATADFYQQLLDAKAATTNPRWDPSFDASIKDGSLIGTV
AAAWEAPLFMTSSGGTGSGEWQVAQLGDWFGNAGKTGPDGGSAVAVLKNSKHPKEAMEFLDW
FNTQVPDLVSQGLVPAATTEDAETPSEWSTFFGGQDIMKEFKTANNNMGDFTYMPGFSAVAA
KMNETAAKATDGSGKVADIFSDAQTTSVDTLKNFGLSVSE
1111111110000000000000000000000000000000000000000000000000000000
0000000000000000000000000000000000000000000000000000000000000000
0000000000000000000000000000000000000000000000000000000000000000
0000000000000000000000000000000000000000000000000000000000000000
0000000000000000000000000000000000000000000000000000000000000000
0000000000000000000000000000000000000000000000000000000000000000
0000000000111000000000000000000000000001
>DM_validata176
MEFTTGLMSLDTALNEMLSRVTPLTAQETLPLVQCFGRILASDVVSPLDVPGFDNSAMDGYA
VRLADIASGQPLPVAGKSFAGQPYHGEWPAGTCIRIMTGAPVPEGCEAVVMQEQTEQMDNGV
RFTAEVRSGQNIRRRGEDISAGAVVFPAGTRLTTAELPVIASLGIAEVPVIRKVRVALFSTG
DELQLPGQPLGDGQIYDTNRLAVHLMLEQLGCEVINLGIIRDDPHALRAAFIEADSQADVVI
SSGGVSVGEADYTKTILEELGEIAFWKLAIKPGKPFAFGKLSNSWFCGLPGNPVSATLTFYQ
LVQPLLAKLSGNTASGLPARQRVRTASRLKKTPGRLDFQRGVLQRNADGELEVTTTGHQGSH
IFSSFSLGNCFIVLERDRGNVEVGEWVEVEPFNALFGGL
1111110000000000000000000000000000000000000000000000000000000000
0000000000000000000000000000000000000000000000000000000000000000
0000000000000000000000000000000000000000000000000000000000000000
0000000000000000000000000000000000000000000000000000000000000000
0000000000000000000000000000000000000000000000000000000000000000
0000000000000000000000000000000000000000000000000000000000000000
000000000000000000000000000000000000000
>DM_validata177
QSEPELKLESVVIVSRHGVRAPTKATQLMQDVTPDAWPTWPVKLGWLTPRGGELIAYLGHYQ
RQRLVADGLLAKKGCPQSGQVAIIADVDERTRKTGEAFAAGLAPDCAITVHTQADTSSPDPL
FNPLKTGVCQLDNANVTDAILSRAGGSIADFTGHRQTAFRELERVLNFPQSNLCLKREKQDE
SCSLTQALPSELKVSADNVSLTGAVSLASMLTEIFLLQQAQGMPEPGWGRITDSHQWNTLLS
LHNAQFYLLQRTPEVARSRATPLLDLIKTALTPHPPQKQAYGVTLPTSVLFIAGHDTNLANL
GGALELNWTLPGQPDNTPPGGELVFERWRRLSDNSQWIQVSLVFQTLQQMRDKTPLSLNTPP
GEVKLTLAGCEERNAQGMCSLAGFTQIVNEARIPACSL
1110000000000000000000000000000000000000000000000000000000000000
0000000000000000000000000000000000000000000000000000000000000000
0000000000000000000000000000000000000000000000000000000001111100
0000000000000000000000000000000000000000000000000000000000000000
0000000000000000000000000000000000000000000000000000000000000000
0000000000000000000000000000000000000000000000000000000000000000
00000000000000000000000000000000000000

>DM_validata178
LNSAAQAQSSDPVVIGCPAPLTGIVAADGIEFQRGIQMAADEINAVGGILGRPIELVFADTQ
SKGVDVVIQSAQRLIDRDNASALIAGYNLENGTALHDVAADAGVIAMHANTVAVHDEMVKSD
PDRYWGTFQYDPPETLYGGGFLKFLKDIEDNGEFSRPNNKIAIITGPGIYSVNIANAIRDGA
GEYGYDVSLFETVAIPVSDWGPTLAKLRADPPAVIVVTHFYPQDQALFMNQFMTDPTNSLVY
LQYGASLAAFRDIAGDNSVGVTYATVLGTLQDEMGDAFAKAYKERYGDLSSTASGCQTYSAL
YAYSIAAALAGGPGAPYDDVQNKAVADRLRSLIFRGPVGTMRFHADTQSAWSYPTETNDPSL
GMPHIFSQIFDKAEDGVLIAPAPYKKAGFKMPPWMKG
111111111100000000000000000000000000000000000000000000000000
000000000000000000000000000000000000000000000000000000000000
000000000000000000000000000000000000000000000000000000000000
000000000000000000000000000000000000000000000000000000000000
000000000000000000000000000000000000000000000000000000000000
000000000000000000000000000000000000000000000000000000000000
0000000000000000000000000000000000011
>DM_validata179
MKVLVINAGSSSLKYQLIDMTNESALAVGLCERIGIDNSIITQKKFDGKKLEKLTDLPTHKD
ALEEVVKALTDDEFGVIKDMGEINAVGHRVVHGGEKFTTSALYDEGVEKAIKDCFELAPLHN
PPNMMGISACAEIMPGTPMVIVFDTAFHQTMPPYAYMYALPYDLYEKHGVRKYGFHGTSHKY
VAERAALMLGKPAEETKIITCHLGNGSSITAVEGGKSVETSMGFTPLEGLAMGTRCGSIDPA
IVPFLMEKEGLTTREIDTLMNKKSGVLGVSGLSNDFRDLDEAASKGNRKAELALEIFAYKVK
KFIGEYSAVLNGADAVVFTAGIGENSASIRKRILTGLDGIGIKIDDEKNKIRGQEIDISTPD
AKVRVFVIPTNEELAIARETKEIVETEVKLRSSIPV
000000000000000000000000000000000000000000000000000000000000
000000000000000000000000000000000000000000000000000000000000
000000000000000000000000000000000000000000000000000000000000
000000000000000000000000000000000000000000000000000000000000
000000000000000000000000000000000000000000000000000000000000
000000000000000000000000000000000000000000000000000000000000
000000000000000000000000001111111111
>DM_validata181
GGVDRWPEYYPETGRDIWIDSVMRQEYLWYRDMPSPAAPDYFQKPEAFLKKAVASMDNGFSK
IDSLLDEPIPSYGFDYTLYKVLDNDTAYNALISYVVPGSPAEEAGLQRGHWIMMMNGDYITK
KVESELLQGSTRQLQIGVYKEVVGEDGEVTGGVVPIGETTMPASRSLVDKPVHRFEIIPWNG
KKVGYLMYNEFKAGPTTDSQAYNDDLRRAFRDFQTGGVNEFVLDLRYNTGGSLDCAQLLCTM
LAPADKMNQLLALLRYSDKRVEANQDLTFNPELIQSGANLNLSTVYVLTTNATRGAAEMVIN
CLNPYMKVVLIGTKTAGEYVATKPFVHPTDRFILNLVVCNVYNAEEKSDYATGFKPTYEYNE
DSYLSTYLPFGNTNETLLNAALKIMSGITDK
111100000001111000000000000000000000000000000000000000000000
000000000000000000000000000000000000000000000000000000000000
000000000000000000000001111100000000000000000000000000000000
000000000000000000000000000000000000000000000000000000000000
000000000000000000000000000000000000000000000000000000000000
000000000000000000000000000000000000000000000000000000000000

0000000000000000000000000000001
>DM_validata183
MSYYHHHHHHDYDPTTENLYFQGPAVGIDLGTTYSCVGVFQHGKVEIIANDQGNRTTPSYVA
FTDTERLIGDAAKNQVAMNPTNTVFDAKRLIGRRFDDAVVQSDMKHWPFMVVNDAGRPKVQV
EYKGETKSFYPEEVSSMVLTKMKEIAEAYLGKTVTNAVVTVPAYFNDSQRQATKDAGTIAGL
NVLRIINEPTAAAIAYGLDKKVGAERNVLIFDLGGGTFDVSILTIEDGIFEVKSTAGDTHLG
GEDFDNRMVNHFIAEFKRKHKKDISENKRAVRRLRTACERAKRTLSSSTQASIEIDSLYEGI
DFYTSITRARFEELNADLFRGTLDPVEKALRDAKLDKSQIHDIVLVGGSTRIPKIQKLLQDF
FNGKELNKSINPDEAVAYGAAVQAAILS
11111111111111111111111000000000000000000000000000000000000000
000000000000000000000000000000000000000000000000000000000000
000000000000000000000000000000000000000000000000000000000000
000000000000000000000000000000000000000000000000000000000000
000000000000000000000000000000000000000000000000000000000000
000000000000000000000000000000000000000000000000000000000000
0000000000000000000000000000
>DM_validata184
MNGPVDGLCDHSLSEEGAFMFTSESVGEGHPDKICDQISDAVLDAHLKQDPNAKVACETVCK
TGMVLLCGEITSMAMIDYQRVVRDTIKHIGYDDSAKGFDFKTCNVLVALEQQSPDIAQCVHL
DRNEEDVGAGDQGLMFGYATDETEECMPLTIVLAHKLNTRMADLRRSGVLPWLRPDSKTQVT
VQYVQDNGAVIPVRVHTIVISVQHNEDITLEAMREALKEQVIKAVVPAKYLDEDTIYHLQPS
GRFVIGGPQGDAGVTGRKIIVDTYGGWGAHGGGAFSGKDYTKVDRSAAYAARWVAKSLVKAG
LCRRVLVQVSYAIGVAEPLSISIFTYGTSKKTERELLEVVNKNFDLRPGVIVRDLDLKKPIY
QKTACYGHFGRSEFPWEVPKKLVF
111111111111111100000000000000000000000000000000000000000000
000000000000000000000000000000000000000000000000000011111111
111100000000000000000000000000000000000000000000000000000000
000000000000000000000000000000000000000000000000000000000000
000000000000000000000000000000000000000000000000000000000000
000000000000000000000000000000000000000000000000000000000000
000000000000000000000000
>DM_validata185
IGLDQIWDDLRAGIQQVYTRQSMAKSRYMELYTHVYNYCTSVHQSNQARGAGVPPSKSKKGQ
TPGGAQFVGLELYKRLKEFLKNYLTNLLKDGEDLMDESVLKFYTQQWEDYRFSSKVLNGICA
YLNRHWVRRECDEGRKGIYEIYSLALVTWRDCLFRPLNKQVTNAVLKLIEKERNGETINTRL
ISGVVQSYVELGLNEDDAFAKGPTLTVYKESFESQFLADTERFYTRESTEFLQQNPVTEYMK
KAEARLLEEQRRVQVYLHESTQDELARKCEQVLIEKHLEIFHTEFQNLLDADKNEDLGRMYN
LVSRIQDGLGELKKLLETHIHNQGLAAIEKCGEAALNDPKMYVQTVLDVHKKYNALVMSAFN
NDAGFVAALDKACGRFINNNAVTK
000000000000000000000000000000000000000111111111111111111111
111110000000000000000000000000000000000000000000000000000000
000000000001111000000000000000000000000000000000000000000000
000000000000000011111111000000000000000000000000000000000000
000000000000000000000000000000000000000000000000000000000000

000000000000000000000000000000000000000000000000000000000000
000000000000000000000000
>DM_validata186
GMVEPMTPRFKYTKASQENIQQLGNILEQCFVMSFGDSEIYVKGIGLENFRVIYREQKVAGG
LAILPMGQWWGGQRVPMAGIAAVGIAPEYRGDGAAIALIQHTLQEISEQDIPISVLYPATQR
LYRKAGYEQAGSSCVWEIPTDSIQIQHASLPLEPVVLKNNPIFHELYQQQAQLTHGYLDRHP
AIWQGLNRTLDTETLYSYLIGDKDKPQGYIIFTQERTRDGSILRIRDWVTLSNPAVQSFWTF
IANHRSQIDKVTWKSSVIDALTLLLPEQSATIRSQDRWMLRIVNVCKALEARGYPLGVEAEL
HLEVQDDLLATNQGKFILSVANGKSEVTKGGKGELQLDIKGLASLYTSLFTPRQLQLTGKLQ
ATETALLKATQIFAGESPWMIDFF
111111110000000000000000000000000000000000000000000000000000
000000000000000000000000000000000000000000000000000000000000
000000000000000000000000000000000000000000000000000000000000
000000000000000000000000000000000000000000000000000000000000
000000000000000000000000000000000000000000000000000000000000
000000000000000000000000000000000000000000000000000000000000
000000000000000000000000
>DM_validata187
MLEVLFQGPSSTVTIEYFNQKKEMTKTLEEITRDFEKENPKIKVKVVNVPNAGEVLKTRVLA
GDVPDVVNIYPQSIELQEWAKAGVFEDLSNKDYLKRVKNGYAEKYAVNEKVYNVPFTANAYG
IYYNKDKFEELGLKVPETWDEFEQLVKDIVAKGQTPFGIAGADAWTLNGYNQLAFATATGGG
KEANQYLRYSQPNAIKLSDPIMKDDIKVMDILRINGSKQKNWEGAGYTDVIGAFARGDVLMT
PNGSWAITAINEQKPNFKIGTFMIPGKEKGQSLTVGAGDLAWSISATTKHPKEANAFVEYMT
RPEVMQKYYDVDGSPTAIEGVKQAGEDSPLAGMTEYAFTDRHLVWLQQYWTSEADFHTLTMN
YVLTGDKQGMVNDLNAFFNPMKM
111110000000000000000000000000000000000000000000000000000000
000000000000000000000000000000000000000000000000000000000000
000000000000000000000000000000000000000000000000000000000000
000000000000000000000000000000000000000000000000000000000000
000000000000000000000000000000000000000000000000000000000000
000000000000000000000000000000000000000000000000000000000000
00000000000000000000000
>DM_validata188
AIIRKNVNSLTPSDIKELRDAMAKVQADTSDNGYQKIASYHGIPLSCHYENGTAYACCQHGM
VTFPNWHRLLTKQMEDALVAKGSHVGIPYWDWTTTFANLPVLVTEEKDNSFHHAHIDVANTD
TTRSPRAQLFDDPDKGDKSFFYRQIALALEQTDFCDFEIQFEIGHNAIHSWVGGSSPYGMST
LHYTSYDPLFYLHHSNTDRIWSVWQALQKYRGLPYNTANCEINKLVKPLKPFNLDTNPNAVT
KAHSTGATSFDYHKLGYDYDNLNFHGMTIPELEEHLKEIQHEDRVFAGFLLRTIGQSADVNF
DVCTKDGECTFGGTFCILGGEHEMFWAFDRLFKYDITTSLKHLRLDAHDDFDIKVTIKGIDG
HVLSNKYLSPPTVFLAPAKTTH
000000000000000000000000000000000000000000000000000000000000
000000000000000000000000000000000000000000000000000000000000
000000000011111111000000000000000000000000000000000000000000
000000000000000000000000000000000000000000000000000000000000

000000000000000000000000000000000000000000000000000000000000
000000000000000000000000000000000000000000000000000000000000
000000000000000001111
>DM_validata189
MSTRESFNPESYELDKSFRLTRFTELKGTGCKVPQDVLQKLLESLQENHFQEDEQFLGAVMP
RLGIGMDTCVIPLRHGGLSLVQTTDYIYPIVDDPYMMGRIACANVLSDLYAMGVTECDNMLM
LLGVSNKMTDRERDKVMPLIIQGFKDAAEEAGTSVTGGQTVLNPWIVLGGVATTVCQPNEFI
MPDNAVPGDVLVLTKPLGTQVAVAVHQWLDIPEKWNKIKLVVTQEDVELAYQEAMMNMARLN
RTAAGLMHTFNAHAATDITGFGILGHAQNLAKQQRNEVSFVIHNLPVLAKMAAVSKACGNMF
GLMHGTCPETSGGLLICLPREQAARFCAEIKSPKYGEGHQAWIIGIVEKGNRTARIIDKPRI
IEVAPQVATQNVNPTPGATS
111111110000000000000000000000000000000000011111111111111100
000000000000000000000000000000000000000000000000000000000000
000000000000000000000000000000000000000000000000000000000000
000000000000000000000000000111111111111000000000000000000000
000000000000000000000000000000000000000000000000000000000000
000000000000000000000000000000001110000000000000000000000000
00000111111111111111
>DM_validata190
MTAAQGNSNETLFSSYKMGRFDLSHRVVLAPMTRCRALNGVPNAALAEYYAQRTTPGGFLIS
EGTMVSPGSAGFPHVPGIYSDEQVEAWKQVVEAVHAKGGFIFCQLWHVGRASHAVYQPNGGS
PISSTNKPISENRWRVLLPDGSHVKYPKPRALEASEIPRVVEDYCLSALNAIRAGFDGIEIH
GAHGYLIDQFLKDGINDRTDQYGGSIANRCRFLKQVVEGVVSAIGASKVGVRVSPAIDHLDA
TDSDPLSLGLAVVGMLNKLQGVNGSKLAYLHVTQPRYHAYGQTESGRQGSDEEEAKLMKSLR
MAYNGTFMSSGGFNKELGMQAVQQGDADLVSYGRLFIANPDLVSRFKIDGELNKYNRKTFYT
QDPVVGYTDYPFLAPFSRL
111111100000000000000000000000000000000000000000000000000000
000000000000000000000000000000000000000000000000000000000000
000000000000000000000000000000000000000000000000000000000000
000000000000000000000000000000000000000000000000000000000000
000000000000000000000000000000000011111111111111100000000000
000000000000000000000000000000000000000000000000000000000000
0000000000000001111
>DM_validata192
MAIVVDDSVFSPSYVPKRLPHREQQLQQLDILLGNWLRNPGHHYPRATLLGRPGTGKTVTLR
KLWELYKDKTTARFVYINGFIYRNFTAIIGEIARSLNIPFPRRGLSRDEFLALLVEHLRERD
LYMFLVLDDAFNLAPDILSTFIRLGQEADKLGAFRIALVIVGHNDAVLNNLDPSTRGIMGKY
VIRFSPYTKDQIFDILLDRAKAGLAEGSYSEDILQMIADITGAQTPLDTNRGDARLAIDILY
RSAYAAQQNGRKHIAPEDVRKSSKEVLFGISEEVLIGLPLHEKLFLLAIVRSLKISHTPYIT
FGDAEESYKIVCEEYGERPRVHSQLWSYLNDLREKGIVETRQNKRGEGVRGRTTLISIGTEP
LDTLEAVITKLIKEELR
000000000000000000000000000000000000000000000000000000000000
000000000000000000000000000000000000000000000000000000000000
000000000000000000000000000000000000000000000000000000000000

000000000000000000000000000000000000000000000000000000000000
000000000000000000000000000000000000000000000000000000000000
000000000000000000000000000000000000000000111111110000000000
00000000000000000
>DM_validata193
MVEATAQETDRPRFSFSIAAREGKARTGTIEMKRGVIRTPAFMPVGTAATVKALKPETVRAT
GADIILGNTYHLMLRPGAERIAKLGGLHSFMGWDRPILTDSGGFQVMSLSSLTKQSEEGVTF
KSHLDGSRHMLSPERSIEIQHLLGSDIVMAFDECTPYPATPSRAASSMERSMRWAKRSRDAF
DSRKEQAENAALFGIQQGSVFENLRQQSADALAEIGFDGYAVGGLAGGEGQDEMFRVLDFSV
PMLPDDKPHYLMGVGKPDDIVGAVERGIDMFDCVLPTRSGRNGQAFTWDGPINIRNARFSED
LKPLDSECHCAVCQKWSRAYIHHLIRAGEILGAMLMTEHNIAFYQQLMQKIRDSISEGRFSQ
FAQDFRARYFARNS
111111111100000000000000000000000000000000000000000000000000
000000000000000000000000000000000000000000000011111100000000
001111111100000000000000000000000000000000000000000000000000
000000000000000000000000000000000000000000000000000000000000
000000000000000000000000000000011111111100000000000000000000
000000000000000000000000000000000000000000000000000000000000
00000000001111
>DM_validata194
MGSHQERPLIGLLFSETGVTADIERSQRYGALLAVEQLNREGGVGGRPIETLSQDPGGDPDR
YRLCAEDFIRNRGVRFLVGCYMSHTRKAVMPVVERADALLCYPNPYEGFEYSPNIVYGGPAP
NQNSAPLAAYLIRHYGERVVFIGSDYIYPRESNHVMRHLYRQHGGTVLEEIYIPLYPSDDDL
QRAVERIYQARADVVFSTVVGTGTAELYRAIARRYGDGRRPPIASLTTSEAEVAKMESDVAE
GQVVVAPYFSSIDTPASRAFVQACHGFFPENATITAWAEAAYWQTLLLGRAAQAAGNWRVED
VQRHLYDIDIDAPQGPVRVERQNNHSRLSSRIAEIDARGVFQVRWQSPEPIRPDPYVVVHNL
DDWSASMGGGPLP
111111000000000000000000000000000000000000000000000000000000
000000000000000000000000000000000000000000000000000000000000
000000000000000000000000000000000000000000000000000000000000
000000000000000000000000000000000000000000000000000000000000
000000000000000000000000000000000000000000000000000000000000
000000000000000000000000000000000000000000000000000000000000
0001111111111
>DM_validata196
MTRRPDRKDVATVDELHASATKLVGLDDFGTDDDNYREALGVLLDAYQGEAGLTVLGSKMNR
FFLRGALVARLLSQSAWKQYPEHVDVAIKRPIFVTGLVRTGTTALHRLLGADPAHQGLHMWL
AEYPQPRPPRETWESNPLYRQLDADFTQHHAENPGYTGLHFMAAYELEECWQLLRQSLHSVS
YEALAHVPSYADWLSRQDWTPSYCRHRRNLQLIGLNDAEKRWVLKNPSHLFALDALMATYPD
ALVVQTHRPVETIMASMCSLAQHTTEGWSTKFVGAQIGADAMDTWSRGLERFNAARAKYDSA
QFYDVDYHDLIADPLGTVADIYRHFGLTLSDEARQAMTTVHAESQSGARAPKHSYSLADYGL
TVEMVKERFAGL
111111000000000000000000000000000000000000000000000000000000
000000000000000000000000000000000000000000000000000000000000

00000000000000000000000000000000000000000000000000000000000000
00000000000000000000000000000000000000000000000000000000000000
00000000000000000000000000000000000000000000000000000000000000
000000000000000000000000000000000000000111111111111100000000000
000000000000
>DM_validata197
LTLGLAPALSTHSSGVSTQSVDLSQIKRGDEIQAHCLTPAETEVTECAGILKDVLSKNLHEL
QGLCNVKNKMGVPWVSVEELGQEIITGRLPFPSVGGTPVNDLVRVLVVAESNTPEETPEEEF
YAYVELQTELYTFGLSDDNVVFTSDYMTVWMIDIPKSYVDVGMLTRATFLEQWPGAKVTVMI
PYSSTFTWCGELGAISEESAPQPSLSARSPVCKNSARYSTSKFCEVDGCTAETGMEKMSLLT
PFGGPPQQAKMNTCPCYYKYSVSPLPAMDHLILADLAGLDSLTSPVYVMAAYFDSTHENPVR
PSSKLYHCALQMTSHDGVWTSTSSEQCPIRLVEGQSQNVLQVRVAPTSMPNLVGVSLMLEGQ
QYRLEYFGDH
11111111111000000000000000000000000000000000000000000000000000
00000000000000000000000000000000000000000000000000000000000000
00000000000000000000000000000000000000000000000000000000000000
00000000000000000000000000000000000000000000000000000000000000
00000000000000000000000000000000000000000000000000000000000000
00000000000000000000000000000000000000000000000000000000000000
0000000000
>DM_validata198
DFDVKRKYFRQELERLDEGLRKEDMAVHVRRDHVFEDSYRELHRKSPEEMKNRLYIVFEGEE
GQDAGGLLREWYMIISREMFNPMYALFRTSPGDRVTYTINPSSHANPNHLSYFKFVGRIVAK
AVYDNRLLECYFTRSFYKHILGKSVRYTDMESEDYHFYQGLVYLLENDVSTLGYDLTFSTEV
QEFGVAEVRDLKPNGANILVTEENKKEYVHLVCQMRMTGAIRKQLAAFLEGFYEIIPKRLIS
IFTEQELELLISGLPTIDIDDLKSNTEYHKYQSNSIQIQWFWRALRSFDQADRAKFLQFVTG
TSKVPLQGFAALEGMNGIQKFQIHRDDRSTDRLPSAHTCFNQLDLPAYESFEKLRHMLLLAI
QEASEGFGLA
00000000000000000000000000000000000000000000000000000000000000
00000000000000000000000000000000000000000011111000000000000000
00000000000000000000000000000000000000000000000000000000000000
00000000000000000000000000000000000000000000000000000000000000
00000000000000000000000000000000000000000000000000000000000000
00000000000000000000000000000000000000000000000000000000000000
0011111111
>DM_validata200
NSSPSPVPGSQNVPAPAVKKISQYACQRRTTLNNYNQLFTDALDILAENDELRENEGSCLAF
MRASSVLKSLPFPITSMKDTEGIPCLGDKVKSIIEGIIEDGESSEAKAVLNDERYKSFKLFT
SVFGVGLKTAEKWFRMGFRTLSKIQSDKSLRFTQMQKAGFLYYEDLVSCVNRPEAEAVSMLV
KEAVVTFLPDALVTMTGGFRRGKMTGHDVDFLITSPEATEDEEQQLLHKVTDFWKQQGLLLY
CDILESTFEKFKQPSRKVDALDHFQKCFLILKLDHGRVHSEKSGQQEGKGWKAIRVDLVMCP
YDRRAFALLGWTGSRQFERDLRRYATHERKMMLDNHALYDRTKRVFLEAESEEEIFAHLGLD
YIEPWERNA
11111111111111111100000000000000000000000000000000000000000000

0000000000000000000000000000000000000000000000000000000000000
0000000000000000000000000000000000000000000000000000000000000
0000000000000000000000000000000000000000000000000000000000000
0000000000000000000000000000000000000000001110000000000000000
0000000000000000000000000000000000000000000000000000000000000
000000000
>DM_validata201
LPVLAPVTKDPATSLYTIPFHDGASLVLDVAGPLVWSTCDGGQPPAEIPCSSPTCLLANAYP
APGCPAPSCGSDKHDKPCTAYPYNPVSGACAAGSLSHTRFVANTTDGSKPVSKVNVGVLAAC
APSKLLASLPRGSTGVAGLANSGLALPAQVASAQKVANRFLLCLPTGGPGVAIFGGGPVPWP
QFTQSMPYTPLVTKGGSPAHYISARSIVVGDTRVPVPEGALATGGVMLSTRLPYVLLRPDVY
RPLMDAFTKALAAQHANGAPVARAVEAVAPFGVCYDTKTLGNNLGGYAVPNVQLGLDGGSDW
TMTGKNSMVDVKQGTACVAFVEMKGVAAGDGRAPAVILGGAQMEDFVLDFDMEKKRLGFSRL
PHFTGCGGL
0000000000000000000000000000000000000000000000000000000000000
0000000001111110000000000000000000000000000000000000000000000
0000000000000000000000000000000000000000000000000000000000000
0000000000000000000000000000000000000000000000000000000000000
0000000000000011111110000000000000000000000000000000000000000
0000000000000000000000000001111110000000000000000000000000000
000000000
>DM_validata202
AHSDASSDITLKVAIYPYVPDPARFQAAVLDQWQRQEPGVKLEFTDWDSYSADPPDDLDVFV
LDSIFLSHFVDAGYLLPFGSQDIDQAEDVLPFALQGAKRNGEVYGLPQILCTNLLFYRKGDL
KIGQVDNIYELYKKIGTSHSEQIPPPQNKGLLINMAGGTTKASMYLEALIDVTGQYTEYDLL
PPLDPLNDKVIRGLRLLINMAGEKPSQYVPEDGDAYVRASWFAQGSGRAFIGYSESMMRMGD
YAEQVRFKPISSSAGQDIPLFYSDVVSVNSKTAHPELAKKLANVMASADTVEQALRPQADGQ
YPQYLLPARHQVYEALMQDYPIYSELAQIVNKPSNRVFRLGPEVRTWLKDAKQVLPEALGLT
DVSSLAS
1111111100000000000000000000000000000000000000000000000000000
0000000000000000000000000000000000000000000000000000000000000
0000000000000000000000000000000000000000000000000000000000000
0000000000000000000000000000000000000000000000000000000000000
0000000000000000000000000000000000000000000000000000000000000
0000000000000000000000000000000000000000000000000000000000011
1111111
>DM_validata203
GMHSALQLRSRIKSSGELELSLDSIDTPHPGPDEVLIRIEASPLNPSDLGLLFGAADMSTAK
ASGTAERPIVTARVPEGAMRSMAGRLDASMPVGNEGAGVVVEAGSSPAAQALMGKTVAAIGG
AMYSQYRCIPADQCLVLPEGATPADGASSFVNPLTALGMVETMRLEGHSALVHTAAASNLGQ
MLNQICLKDGIKLVNIVRKQEQADLLKAQGAVHVCNAASPTFMQDLTEALVSTGATIAFDAT
GGGKLGGQILTCMEAALNKSAREYSRYGSTTHKQVYLYGGLDTSPTEFNRNFGMAWGMGGWL
LFPFLQKIGRERANALKQRVVAELKTTFASHYSKEISLAEVLDLDMIAVYNKRATGEKYLIN
PNKGLAG

110000000000000000000000000000000000000000000000000000000000
000000000000000000000000000000000000000000000000000000000000
000000000000000000000000000000000000000000000000000000000000
000000000000000000000000000000000000000000000000000000000000
000000000000000000000000000000000000000000000000000000000000
000000000000000000000000000000000000000000000000000000000000
0000011
>DM_validata204
ATPVTPYYGPGHITFDWCGFGDSRSDCTNPQSPMSLDIPQQLCPKFSSKSSSSMFLSLHWNN
HSSFVSYDYFNCGVEKVFYEGVNFSPRKQYSCWDEGVDGWIELKTRFYTKLYQMATTSRCIK
LIQLQAPSSLPTLQAGVCRTNKQLPDNPRLALLSDTVPTSVQFVLPGSSGTTICTKHLVPFC
YLNHGCFTTGGSCLPFGVSYVSDSFYYGYYDATPQIGSTESHDYVCDYLFMEPGTYNASTVG
KFLVYPTKSYCMDTMNITVPVQAVQSIWSEQYASDDAIGQACKAPYCIFYNKTTPYTVTNGS
DANHGDDEVRMMMQGLLRNSSCISPQGSTPLALYSTEMIYEPNYGSCPQFYKLFDTSGNENI
DVISSS
100000000000000000000000000000000000000000000000000000000000
000000000000000000000000000000000000000000000000000000000000
000000000000000000000000000000000000000000000000000000000000
000000000000000000000000000000000111110000000000000000000000
000000000000000000000000000000000000000000000000000000000000
000000000000000000000000000000000000000000000000000011111111
111111
>DM_validata205
MGSDKIHHHHHHMKVTVTTLELKDKITIASKALAKKSVKPILAGFLFEVKDGNFYICATDLE
TGVKATVNAAEISGEARFVVPGDVIQKMVKVLPDEITELSLEGDALVISSGSTVFRITTMPA
DEFPEITPAESGITFEVDTSLLEEMVEKVIFAAAKDEFMRNLNGVFWELHKNLLRLVASDGF
RLALAEEQIENEEEASFLLSLKSMKEVQNVLDNTTEPTITVRYDGRRVSLSTNDVETVMRVV
DAEFPDYKRVIPETFKTKVVVSRKELRESLKRVMVIASKGSESVKFEIEENVMRLVSKSPDY
GEVVDEVEVQKEGEDLVIAFNPKFIEDVLKHIETEEIEMNFVDSTSPCQINPLDISGYLYIV
MPIRLA
111111111110000000000000000000000000000000000000000000000000
000000000000000000000000000000000000000000000000000000000000
000000000000000000000000000000000000000000000000000000000000
000000000000000000000000000000000000000000000000000000000000
000000000000000000000000000000000000000000000000000000000000
000000000000000000000000000000000000000000000000000000000000
000000
>DM_validata206
MEGGMKRVVLAFGTRPEATKMAPVYLALRGIPGLKPLVLLTGQHREQLRQALSLFGIQEDRN
LDVMQERQALPDLAARILPQAARALKEMGADYVLVHGDTLTTFAVAWAAFLEGIPVGHVEAG
LRSGNLKEPFPEEANRRLTDVLTDLDFAPTPLAKANLLKEGKREEGILVTGQTGVDAVLLAA
KLGRLPEGLPEGPYVTVTMHRRENWPLLSDLAQALKRVAEAFPHLTFVYPVHLNPVVREAVF
PVLKGVRNFVLLDPLEYGSMAALMRASLLLVTDSGGLQEEGAALGVPVVVLRNVTERPEGLK
AGILKLAGTDPEGVYRVVKGLLENPEELSRMRKAKNPYGDGKAGLMVARGVAWRLGLGPRPE

DWLP
111100000000000000000000000000000000000000000000000000000000
000000000000000000000000000000000000000000000000000000000000
000000000000000000000000000000000000000000000000000000000000
000000000000000000000000000000000000000000000000000000000000
000000000000000000000000000000000000000000000000000000000000
000000000000000000000000000000000000000000000000000000000000
0000
>DM_validata208
TAQTKNTQTLMPLTERVNVQADSARINQIIDGCWVAVGTNKPHAIQRDFTNLFDGKPSYRFE
LKTEDNTLEGYAKGETKGRAEFSYCYATSDDFRGLPADVYQKAQITKTVYHHGKGACPQGSS
RDYEFSVYIPSSLDSNVSTIFAQWHGMPDRTLVQTPQGEVKKLTVDEFVELEKTTFFKKNVG
HEKVARLDKQGNPVKDKNGKPVYKAGKPNGWLVEQGGYPPLAFGFSGGLFYIKANSDRKWLT
DKDDRCNANPGKTPVMKPLTSEYKASTIAYKLPFADFPKDCWITFRVHIDWTVYGKEAETIV
KPGMLDVRMDYQEQGKKVSKHIVDNEKILIGRNDEDGYYFKFGIYRVGDSTVPVCYNLAGYS
ER
000000000000000000000000000000000000000000000000000000000000
000000000000000000000000000000000000000000000000000000000000
000000000000000000000000000000000000000000000000000000000000
000000000000000000000000000000000000000000000000000000000000
000000000000000000000000000000000000000000000000000000000000
000000000000010000000000000000000000000000000000000000000000
00
>DM_validata209
VCEKGFEAGDNKLGGALNAKHVEKYGDNFKNGMHKPEFHEDGLHKPMEVGGKKFESGFHYLL
ECHELGGKNASGGYGGPLCEDPYGSEVQAMTEKLLKEADSDRTLCFNNFQDPCPQLTKEQVA
MCKGFDYGDKTLKLPCGPLPWPAGLPEPGYVPKTNPLHGRWITVSGGQAAFIKEAIKSGMLG
AAEANKIVADTDHHQTGGMYLRINQFGDVCTVDASVAKFARAKRTWKSGHYFYEPLVSGGNL
LGVWVLPEEYRKIGFFWEMESGRCFRIERRAFPVGPYTFMRQATEVGGKISFVFYVKVSNDP
ESDPIPLQSRDYTALAGRDNAPTNLGKPYPTLAKDLDYPKKRDGWLEKNEKEMLRQRNIVSS
TF
110000000000000000000000000000000000000000000000000000000000
000000000000000000000000000000000000000000000000000000000000
000000000000000000000000000000000000000000000000000000000000
000000000000000000000000000000000000000000000000000000000000
000000000000000000000000000000000000000000000000000000000000
000000000000000000000000000000000000000000011111111111111111
11
>DM_validata210
MLSDYEEFLRLEKARKIILEILNEKGRDALYDLSGLSGGFLIDEKDKALLNTYIGSSYFAEK
VNEYGLKHLGGDENDKCVGFNRTSSAILATILALKPKKVIHYLPELPGHPSIERSCKIVNAK
YFESDKVGEILNKIDKDTLVIITGSTMDLKVIELENFKKVINTAKNKEAIVFVDDASGARVR
LLFNQPPALKLGADLVVTSTDKLMEGPRGGLLAGKKELVDKIYIEGTKFGLEAQPPLLAGIY
RALKNFNLERIRKAFERAKNFDLSKIEKLNKELKAIDDNINIVYERTPTGFVIKRVYKDDTI

NIKKLIEIGFNLLKNYGIITITVAGMPGASKSLRIDLTSRDAERIDDNYIIKAIVESIKMAF
KS
11111111000000000000000000000000000000000000000000000000000000
00000000000000000000000000000000000000000000000000000000000000
00000000000000000000000000000000000000000000000000000000000000
00000000000000000000000000000000000000000000000000000000000000
00000000000000000000000000000000000000000000000000000000000000
00000000000000000000000000000000000000000000000000000000000000
00
>DM_validata211
GMEWQAEQAYNHLPPLPLDSKLAELAETLPILKACIPARAALAELKQAGELLPNQGLLINLL
PLLEAQGSSEIENIVTTTDKLFQYAQEDSQADPMTKEALRYRTALYQCFTQLSNRPLCVTTA
LEICSTIKSVQMDVRKVPGTSLTNQATGEVIYTPPAGESVIRDLLSNWEAFLHNQDDVDPLI
KMAMAHYQFEAIHPFIDGNGRTGRVLNILYLIDQQLLSAPILYLSRYIVAHKQDYYRLLLNV
TTQQEWQPWIIFILNAVEQTAKWTTHKIAAARELIAHTTEYVRQQLPKIYSHELVQVIFEQP
YCRIQNLVESGLAKRQTASVYLKQLCDIGVLEEVQSGKEKLFVHPKFVTLMTKDSNQFSRYA
L
10000111111000000000000000000000000000000000000000000000000000
00000000000000000000000000000000000000000000000000000000000000
00000000000000000000000000000000000000000000000000000000000000
00000000000000000000000000000000000000000000000000000000000000
00000000000000000000000000000000000000000000000000000000000000
00000000000000000000000000000000000000000000000000000000000001
1
>DM_validata212
SGLQGMDVVHGTATMQVDGNKTIIRNSVDAIINWKQFNIDQNEMVQFLQENNNSAVFNRVTS
NQISQLKGILDSNGQVFLINPNGITIGKDAIINTNGFTASTLDISNENIKARNFTFEQTKDK
ALAEIVNHGLITVGKDGSVNLIGGKVKNEGVISVNGGSISLLAGQKITISDIINPTITYSIA
APENEAVNLGDIFAKGGNINVRAATIRNQGKLSADSVSKDKSGNIVLSAKEGEAEIGGVISA
QNQQAKGGKLMITGDKVTLKTGAVIDLSGKEGGETYLGGDERGEGKNGIQLAKKTSLEKGST
INVSGKEKGGRAIVWGDIALIDGNINAQGSGDIAKTGGFVETSGHDLFIKDNAIVDAKEWLL
D
10000000000000000000000000000000000000000000000000000000000000
00000000000000000000000000000000000000000000000000000000000000
00000000000000000000000000000000000000000000000000000000000000
00000000000000000000000000000000000000000000000000000000000000
00000000000000000000000000000000000000000000000000000000000000
00000000000000000000000000000000000000000000000000000000000000
0
>DM_validata213
MKNVGFIGWRGMVGSVLMDRMSQENDFENLNPVFFTTSQAGQKAPVFGGKDAGDLKSAFDIE
ELKKLDIIVTCQGGDYTNEVYPKLKATGWDGYWVDAASALRMKDDAIIVLDPVNQHVISEGL
KKGIKTFVGGNCTVSLMLMAIGGLFEKDLVEWISVATYQAASGAGAKNMRELLSQMGLLEQA
VSSELKDPASSILDIERKVTAKMRADNFPTDNFGAALGGSLIPWIDKLLPETGQTKEEWKGY

AETNKILGLSDNPIPVDGLCVRIGALRCNSQAFTIKLKKDLPLEEIEQIIASHNEWVKVIPN
DKEITLRELTPAKVTGTLSVPVGRLRKLAMGPEYLAAFTVGDQLLWGAAEPVRRILKQLVA
000000000000000000000000000000000000011111111111111100000000
000000000000000000000000000000000000000000000000000000000000
000000000000000000000000000000000000000000000000000000000000
000000000000000000000000000000000000000000000000000000000000
000000000000000000000000000000000000000000000000000000000000
00000000000000000000000000000000000000000000000000000000000
>DM_validata214
MIKEFSDPLYGFVRVGEAGLRLIDSFPFQRLRYVKQLGLAYLVFPSAQHTRFEHSLGVYHIT
ERICESLKVKEKELVKLAGLLHDLGHPPFSHTTEVLLPRERSHEDFTERVIKETEIYEILKQ
DYSHEDIERLVRITLGKPEDEEEKLLSEIITGEFGSDRMDYLRRDAYFCGVSYGFFDYDRLI
STLRVYENKVVVDESGLRALENFLISRYFMYVQVYFHKVVRILSIHLVEFLKKLISQEDFTD
INNFLRLNDAFVISELFKRKAFREDFERIFQRKHFKTLLSTENYEKFSETKERLLEKFPQEK
VRFDEVEKEVYGGNIYVLSSEGLKKAHELSPLIASLKPIKLYRIYVDRQLWEKARSELKLS
000000000000000000000000000000000000000000000000000000000000
000000000000000000000000000000000000000000000000000000000000
000000000000000000000000000000000000000000000000000000000000
000000000000000000000000000000000000000000000000000000000000
000000000000000000000000000000000000000000000000000000000000
00000000000000000000000000000000000000000000000000000000011
>DM_validata215
MSESPMFAANGMPKVNQGAEEDVRILGYDPLASPALLQVQIPATPTSLETAKRGRREAIDII
TGKDDRVLVIVGPCSIHDLEAAQEYALRLKKLSDELKGDLSIIMRAYLEKPRTTVGWKGLIN
DPDVNNTFNINKGLQSARQLFVNLTNIGLPIGSEMLDTISPQYLADLVSFGAIGARTTESQL
HRELASGLSFPVGFKNGTDGTLNVAVDACQAAAHSHHFMGVTKHGVAAITTTKGNEHCFVIL
RGGKKGTNYDAKSVAEAKAQLPAGSNGLMIDYSHGNSNKDFRNQPKVNDVVCEQIANGENAI
TGVMIESNINEGNQGIPAEGKAGLKYGVSITDACIGWETTEDVLRKLAAAVRQRREVNKK
111111111111111111111000000000000000000000000000000000000000
000000000000000000000000000000000000000000000000000000000000
000000000000000000000000000000000000000000000000000000000000
000000000000000000000000000000000000000000000000000000000000
000000000000000000000000000000000000000000000000000000000000
0000000000000111111111110000000000000000000000000000000000
>DM_validata216
AHCTEYMNAPKKLPADVAEELATTAQKLVQAGKGILAADESTQTIKKRFDNIKLENTIENRA
SYRDLLFGTKGLGKFISGAILFEETLFQKNEAGVPMVNLLHNENIIPGIKVDKGLVNIPCTD
EEKSTQGLDGLAERCKEYYKAGARFAKWRTVLVIDTAKGKPTDLSIHETAWGLARYASICQQ
NRLVPIVEPEILADGPHSIEVCAVVTQKVLSCVFKALQENGVLLEGALLKPNMVTAGYECTA
KTTTQDVGFLTVRTLRRTVPPALPGVVFLSGGQSEEEASVNLNSINALGPHPWALTFSYGRA
LQASVLNTWQGKKENVAKAREVLLQRAEANSLATYGKYKGGAGGENAGASLYEKKYVY
111111111111000000000000000000000000000000000000000000000000
000000000000000000000000000000000000000000000000000000000000
000000000000000000000000000000000000000000000000000000000000

00000000000000000000000000000000000000000000000000000000000000
00000000000000000000000000000000000000000000000000000000000000
0000000000000000000000000000000000000000000011111111111111
>DM_validata217
AMKPAQKGKTFSNVEIFDPPTNYRDPQVLYARPLELSDGTLLGTWENYSPEPPNVWFPIVKS
KDGGKTWKEISKVKDTQNNWGLRYQPQLYELPRAFGKYPKGTVLCSGSSIPSDLSETLIEVY
ASRDKGYTWEFVSHVALGGEALPNPGLTPVWEPFLMTYKEKLILYYSDQRDNATHSQKLVHQ
TTTDLKKWSKVVDDTKYADYYARPGMPTVAKLPNNEYIYVYEYGGGPNPPAGSDYWFPVYYR
LSKDPQKFLNKAHHQIVSNDGTTPAGSPYVVWTPYGGKNGTIVVSCGTRSEIFTNQALGDAS
AWKKWDVPQPTAYTRSLLTFQKDPDLLMIMGAGILPPAGGKNTVSASVVRLSEVMKS
00000110000000000000000000000000000000000000000000000000000000
00000000000000000000000000000000000000000000000000000000000000
00000000000000000000000000000000000000000000000000000000000000
00000000000000000000000000000000000000000000000000000000000000
00000000000000000000000000000000000000000000000000000000000000
000000000000000000000000000000000000000000000000000000000
>DM_validata218
ALFGTKDTTTAHSDYEIILEGGSSSWGQVKGRAKVNVPAAIPLLPTDCNIRIDAKPLDAQKG
VVRFTTKIESVVDSVKNTLNVEVDIANETKDRRIAVGEGSLSVGDFSHSFSFEGSVVNMYYY
RSDAVRRNIPNPIYMQGRQFHDILMKVPLDNNDLVDTWEGFQQSISGGGANFGDWIREFWFI
GPAFAAINEGGQRISPIVVNSSNVEGGEKGPVGVTRWKFSHAGSGVVDSISRWTELFPVEQL
NKPASIEGGFRSDSQGIEVKVDGNLPGVSRDAGGGLRRILNHPLIPLVHHGMVGKFNDFTVD
TQLKIVLPKGYKIRYAAPQFRSQNLEEYRWSGGAYARWVEHVCKGGTGQFEVLYAQ
11111100000000000000000000000000000000000000000000000000001110
00000000000000000000000000000000000000000000000000000000000000
00000000000000000000000000000000000000000001110000000000000000
00000000000000000000000001110000000000000000000000000000000000
00000000000000000000000000000000000000000000000000000000000000
00000000000000000000000000000000000000000000000000000000
>DM_validata220
MSLADISVIQSLPLSGSQAVTGRALNAGARLYFDWLNLNGGINGETIRLVARDDEQKIEQTV
RNVRDMARVDNPVALLTVVGTANVEALMREGVLAEARLPLVGPATGASSMTTDPLVFPIKAS
YQQEIDKMITALVTIGVTRIGVLYQEDALGKEAITGVERTLKAHALAITAMASYPRNTANVG
PAVDKLLAADVQAIFLGATAEPAAQFVRQYRARGGEAQLLGLSSIDPGILQKVAGLDAVRGY
SLALVMPNPGKSVNPVIREFNRARAAVGAKDVDLSFRAVEGFVAAKVLAEAIRRAGPKPTRE
QVRHALTELRDYDVGGGFTVDFTDRSRPGSHYIELGVVGPNGLVIQEGHHHHHH
11110000000000000000000000000000000000000000000000000000000000
00000000000000000000000000000000000000000000000000000000000000
00000000000000000000000000000000000000000000000000000000000000
00000000000000000000000000000000000000000000000000000000000000
00000000000000000000000000000000000000000000000000000000000000
000000000000000000000000000000000000000000000001111111
>DM_validata221
NAQKRSPTGIVLMNMGGPSKVEETYDFLYQLFADNDLIPISAKYQKTIAKYIAKFRTPKIEK

QYREIGGGSPIRKWSEYQATEVCKILDKTCPETAPHKPYVAFRYAKPLTAETYKQMLKDGVK
KAVAFSQYPHFSYSTTGSSINELWRQIKALDSERSISWSVIDRWPTNEGLIKAFSENITKKL
QEFPQPVRDKVVLLFSAHSLPMDVVNTGDAYPAEVAATVYNIMQKLKFKNPYRLVWQSQVGP
KPWLGAQTAEIAEFLGPKVDGLMFIPIAFTSDHIETLHEIDLGVIGESEYKDKFKRCESLNG
NQTFIEGMADLVKSHLQSNQLYSNQLPLDFALGKSNDPVKDLSLVFGNHEST
11111100000000000000000000000000000000000000000000000000000000
00000000000000000000000000000000000000000000000000000000000000
00000000000000000000000000000000000000000000000000000000000000
00000000000000000000000000000000000000000000000000000000000000
00000000000000000000000000000000000000000000000000000000000000
0000000000000000000000000000000000000000000000000000
>DM_validata222
KTTISVIKADIGSLAGHHIVHPDTMAAANKVLASAKEQGIILDYYITHVGDDLQLIMTHTRG
ELDTKVHETAWNAFKEAAKVAKDLGLYAAGQDLLSDSFSGNVRGLGPGVAEMEIEERASEPI
AIFMADKTEPGAYNLPLYKMFADPFNTPGLVIDPTMHGGFKFEVLDVYQGEAVMLSAPQEIY
DLLALIGTPARYVIRRVYRNEDNLLAAVVSIERLNLIAGKYVGKDDPVMIVRLQHGLPALGE
ALEAFAFPHLVPGWMRGSHYGPLMPVSQRDAKATRFDGPPRLLGLGFNVKNGRLVGPTDLFD
DPAFDETRRLANIVADYMRRHGPFMPHRLEPTEMEYTTLPLILEKLKDRFKK
00000000000000000000000000000000000000000000000000000000000000
00000000000000000000000000000000000000000000000000000000000000
00000000000000000000000000000000000000000000000000000000000000
00000000000000000000000000000000000001110000000000000000000000
00000000000000000000000000000000000000000000000000000000000000
0000000000000000000000000000000000000000000000000000
>DM_validata223
MERTLDRVGVFAATHAAVAASDPLQARALVLQLPGLNRNKDVPGIVGLLREFLPVRGLPSGW
GFVEAAAAMRDIGFFLGSLKRHGHEPAEVVPGLEPVLLDLARATNLPPRETLLHVTVWNPTA
ADAQRSYTGLPDEAHLLESVRISMAALEAAIALTVELFDVSLRSPEFAQRSDELEAYLQKMV
ESIVYAYRFISPQVFYDELRPFYEPIRVGGQSYLGPGAVEMPLFVLEHVLWGSQSDDQTYRE
FKETYLPYVLPAYRAVYARFSGEPALIDRALDEARAVGTRDEHVRAGLTALERVFKVLLRFR
APHLKLAERAYEVGQSGPEIGSGGYAPSMLGELLTLTYAARSRVRAALDES
11100000000000000000000000000000000000000000000000000000000000
00000000000000000000000000000000000000000000000000000000000000
00000000000000000000000000000000000000000000000000000000000000
00000000000000000000000000000000000000000000000000000000000000
00000000000000000000000000000000000000000000000000000000000000
000000000000011111111111100000000000000000000000011
>DM_validata224
GMHLPGGIVENVRKRPGMYCGDVGEYGLHHLVYFLLDVAYEEARRGECRDVVLEVGGDGSIA
LFCTSRTVTAENLVRVATGAGFLGRPPGDGWGWDSMLVVSLALSSRYQVDIWADGRQWRVMG
EHGHPQGEGAAVTPMEPMPVSAERGVRVHFVPDATIFEVLAFDRARLSRRCNELAALAPGLR
VSFADLQRGERTLWHLPGGVAQWAHVLTEARPQLHPEPVVFDFTWDGLRVQCALQWCEDEDS
TLLSFANAVRTVRHGAHVKGVTQALRGALAKLSGETRGAFPWARVAQGLTAIVAVSGPRRQM
AFAGPTKELLAIPGLEEAIRKQLQPLFIELLREHPVTPALLARRTSGS

1111111000000000000000000000000000000000000000000000000000000
000001111111111111111111111111111000000000000000001000000000
000000011111110000000000000000000000000000000000000000000000
000000000000000000000000000000000000000000000000000000000000
000000000000000000000000000000000000000000000000000000000000
00000000000000000000000000000000000000000000111
>DM_validata225
GMLFDTSPKDNRKDFFDREKEIEKLKGLRAPITLVLGLRRTGKSSIIKIGINELNLPYIYLD
LRKFEERNYISYKDFLLELQKEINKLVKRLPSLLKALKNIQGIVIMGNEIKFNWNRKDRLSF
ANLLESFEQASKDNVIIVLDEAQELVKLRGVNLLPALAYAYDNLKRIKFIMSGSEMGLLYDY
LRVEDPESPLFGRAFSTVELKPFSREEAIEFLRRGFQEADIDFKDYEVVYEKIGGIPGWLTY
FGFIYLDNKNLDFAINQTLEYAKKLILKEFENFLHGREIARKRYLNIMRTLSKCGKWSDVKR
ALELEEGIEISDSEIYNYLTQLTKHSWIIKEGEKYCPSEPLISLAFS
000000000000000000000000000000000000000000000000000000000000
000000000000000000000000000000000000000000000000000111110000
000000000000000000000000000000000000000000000000000000000000
000000000000000000000000000000000000000000000000000000000000
000000000000000000000000000000000000000000000000000000000000
00000000000000000000000000000000000000000000000
>DM_validata227
MASAEVVSKLEAAFAKLQNASDCHSLLKKYLTKEVFDQLKGKQTKMGATLMDVIQSGVENLD
SGIGVYAPDAESYTLFAALLDPIIEDYHKGFKPSDKQPPKDSGDLNTFIDVDPDKKYVISTR
VRCGRSLEGYPFNPCLKKQQYEEMESRVKGQLESMSGELRGKYYPLTGMTKETQKQLIDDHF
LFKEGDRFLQAAHACKFWPTGRGIYHNDAKTFLVWVNEEDHLRIISMQKGGNLKEVFGRLVT
AVGVIEEKVKFSRDDRLGFLTFCPTNLGTTIRASVHIKLPKLGADRKKLEEVAAKYNLQVRG
TAGEHSDSPDGVYDISNKRRLGLSEYEAVKEMQDGILELIKAEESAR
000000000000000000000000000000000000000000000000000000000000
000000000000000000000000000000000000000000000000000000000000
000000000000000000000000000000000000000000000000000000000000
000000000000000000000000000000000000000000000000000000000000
000000000000000000000000000000000000001111100000000000000001
11111111110000000000000000000000000000000000011
>DM_validata228
SVLPETPVPFKSGTGAIDNDTVYIGLGSAGTAWYKLDTQAKDKKWTALAAFPGGPRDQATSA
FIDGNLYVFGGIGKNSEGLTQVFNDVHKYNPKTNSWVKLMSHAPMGMAGHVTFVHNGKAYVT
GGVNQNIFNGYFEDLNEAGKDSTAIDKINAHYFDKKAEDYFFNKFLLSFDPSTQQWSYAGES
PWYGTAGAAVVNKGDKTWLINGEAKPGLRTDAVFELDFTGNNLKWNKLAPVSSPDGVAGGFA
GISNDSLIFAGGAGFKGSRENYQNGKNYAHEGLKKSYSTDIHLWHNGKWDKSGELSQGRAYG
VSLPWNNSLLIIGGETAGGKAVTDSVLITVKDNKVTVQNLEHHHHHH
000000000000000000000000000000000000000000000000000000000000
000000000000000000000000000000000000000000000000000000000000
000000000000000000000000000000000000000000000000000000000000
000000000000000000000000000000000000011100000000000000000000
000000000000000000000000000000000000000000011100000000000000

000000000000000000000000000000100000000011111111
>DM_validata229
MVAVERVESLAKSGIISIPKEYIRPKEELESINDVFLEEKKEDGPQVPTIDLKNIESDDEKI
RENCIEELKKASLDWGVMHLINHGIPADLMERVKKAGEEFFSLSVEEKEKYANDQATGKIQG
YGSKLANNASGQLEWEDYFFHLAYPEEKRDLSIWPKTPSDYIEATSEYAKCLRLLATKVFKA
LSVGLGLEPDRLEKEVGGLEELLLQMKINYYPKCPQPELALGVEAHTDVSALTFILHNMVPG
LQLFYEGKWVTAKCVPDSIVMHIGDTLEILSNGKYKSILHRGLVNKEKVRISWAVFCEPPKD
KIVLKPLPEMVSVESPAKFPPRTFAQHIEHKLFGKEQEELVSEKND
000000000000000000000000000000000000000000000000000000000000
000000000000000000000000000000000000000000000000000000000000
000000000000000000000000000000000000000000000000000000000000
000000000000000000000000000000000000000000000000000000000000
000000000000000000000000000000000000000000000000000000000000
0000000000000000000000000000000000000111111111
>DM_validata230
GPHMSVEQTETAELPASDSINPGNLQLVSELKNPSGSCSVDVSAMFWERAGCKEPCIITACE
DVVSLWKALDAWQWEKLYTWHFAEVPVLQIVPVPDVYNLVCVALGNLEIREIRALFCSSDDE
SEKQVLLKSGNIKAVLGLTKRRLVSSSGTLSDQQVEVMTFAEDGGGKENQFLMPPEETILTF
AEVQGMQEALLGTTIMNNIVIWNLKTGQLLKKMHIDDSYQASVCHKAYSEMGLLFIVLSHPC
AKESESLRSPVFQLIVINPKTTLSVGVMLYCLPPGQAGRFLEGDVKDHCAAAILTSGTIAIW
DLLLGQCTALLPPVSDQHWSFVKWSGTDSHLLAGQKDGNIFVYHYS
111111111111111111111100000000000000000000000011110000000000
000000000000000000000000000000000000000000000000000000011111
100000000000000000000000000000000000000000000000000000000000
000000000000000000000000000000000000111000000000000000000111
111111111000000000000000000000000000000000000000000000000000
0000000000000011000000000000000000000000000000
>DM_validata231
MKFGNFLLTYQPPELSQTEVMKRLVNLGKASEGCGFDTVWLLEHHFTEFGLLGNPYVAAAHL
LGATETLNVGTAAIVLPTAHPVRQAEDVNLLDQMSKGRFRFGICRGLYDKDFRVFGTDMDNS
RALMDCWYDLMKEGFNEGYIAADNEHIKFPKIQLNPSAYTQGGAPVYVVAESASTTEWAAER
GLPMILSWIINTHEKKAQLDLYNEVATEHGYDVTKIDHCLSYITSVDHDSNRAKDICRNFLG
HWYDSYVNATKIFDDSDQTKGYDFNKGQWRDFVLKGHKDTNRRIDYSYEINPVGTPEECIAI
IQQDIDATGIDNICCGFEANGSEEEIIASMKLFQSDVMPYLKEKQ
000000000000000000000000000000000000000000000000000000000000
000000000000000000000000000000000000000000000000000000000000
000000000000000000000000000000000000000000000000000000000000
000000000000000000000000000000000000000000000000000000000000
000000000000011111111111111111111111111111000000000000000000
000000000000000000000000000000000000000000000
>DM_validata232
MLIGEIMDLNLKNFLEDREEIIRDAKRKDEKSFKDFKKIVEEIKERENKDKIVCDFTEYNPL
HKGHKYALEKGKEHGIFISVLPGPLERSGRGIPYFLNRYIRAEMAIRAGADIVVEGPPMGIM
GSGQYMRCLIKMFYSLGAEIIPRGYIPEKTMEKVIDCINKGYHIQVKPYKIICIETGEILGE

KLNIDNYVIASMSQMIYKLNREGLKFNPKFVFVKRLEGISGTKIREAIFSGKFEDIKNMLPK
TTLSILKELYDNGKLNELILKRFEDRILETANEYDLYEYLPSNVAEILEKKRPFNNIEEIKN
SLPYGFSRHFRERILSKLEARIPNETLSKYINNYPAKIKILAVKL
00000000000000000000000000000000000000000000000000000000000000
00000000000000000000000000000000000000000000000000000000000000
00000000000000000000000000000000000000000000000000000000000000
00000000000000000000000000000000000000000000000000000000000000
00000000000000000000000000000000000000000000000000000000000000
000000000000000000000000000000000000000000001
>DM_validata233
MSKVNEITRESWILSTFPEWGTWLNEEIEQTVVEPNTFSMWWLGCTGIWLKSAGNTNLSIDF
WCGTGKKTQKNRLMNTQHQMMRMGGVEALQPNLRTSIFPLDPFAIKEIDAVLASHDHADHID
VNVAAAVLQNCGEHVKFIGPQACVDLWLGWGVPQERCIVAKVGDVLEIGDVKIRVLDSFDRT
ALVTLPKGVSSYDKAILDGMDERAVNYLIETSGGSVYHSGDSHYSNYYAKHGNDYQIDVALL
SYGENPRGVTDKMTSSDVLRAAESLDCQVVVPFHHDIWANFQNDPREIEVLWNMKKDRLQYQ
FAPFFWQVGGKYTYPTDKGRMHYQHFRGFQDIFKNEPELPYKAFL
11000000000000000000000000000000000000000000000000000000000000
00000000000000000000000000000000000000000000000000000000000000
00000000000000000000000000000000000000000000000000000000000000
00000000000000000000000000000000000000000000000000000000000000
00000000000000000000000000000000000000000000000000000000000000
000000000000000000000000000000000000000000000
>DM_validata234
MEAAHFFEGTEKLLEVWFSRQQPDANQGSGDLRTIPRSEWDILLKDVQCSIISVTKTDKQEA
YVLSEASMFVSKRRFILKTCGTTLLLKALVPLLKLARDYSGFDSIQSFFYSRKNFMKPSHQG
YPHRNFQEEIEFLNAIFPNGAAYCMGRMNSDCWYLYTLDFPESRVISQPDQTLEILMSELDP
AVMDQFYMKDGVTAKDVTRESGIRDLIPGSVIDATMFNPCGYSMNGMKSDGTYWTIHITPEP
EFSYVSFETNLSQTSYDDLIRKVVEVFKPGKFVTTLFVNQSSKCRTVLASPQKIEGFKRLDC
QSAMFNDYNFVFTSFAKKQQQQQSHICRSQMVTSQQTSSVVRQT
11100000000000000000011111100000000000000000000000000000000000
00000000000000000000000000000000000000000000000000000000000000
00000000000000000000000000000000000000000000000000000000000000
00000000000000000000000000000000000000000000000000000000000000
00000000000000000000000000000000000000000000000000000000000000
00000000000000000011111111111111111111111111
>DM_validata235
MMESAIGEHLQCPRTLTRRVPDTYTPPFPMWVGRADDALQQVVMGYLGVQFRDEDQRPAALQ
AMRDIVAGFDLPDGPAHHDLTHHIDNQGYENLIVVGYWKDVSSQHRWSTSTPIASWWESEDR
LSDGLGFFREIVAPRAEQFETLYAFQEDLPGVGAVMDGISGEINEHGYWGSMRERFPISQTD
WMQASGELRVIAGDPAVGGRVVVRGHDNIALIRSGQDWADAEADERSLYLDEILPTLQSGMD
FLRDNGPAVGCYSNRFVRNIDIDGNFLDLSYNIGHWASLDQLERWSESHPTHLRIFTTFFRV
AAGLSKLRLYHEVSVFDAADQLYEYINCHPGTGMLRDAVTIAEH
10000000000000000000000000000000000000000000000000000000000000
00000000000000000000000000000000000000000000000000000000000000

000000000000000000000000000000000000000000000000000000000000
000000000000000000000000000000000000000000000000000000000000
000000000000000000000000000000000000000000000000000000000000
0000000000000000000000000000000000000000001
>DM_validata236
MGSSHHHHHHSSGLVPRGSNLKIKLQKRRDEVNTCLCIGLDPDEADIKSFMQSEKQNGYQSV
KKNLSNSGSSSSSSNSSSGKGELFAPQMGGQMLLAETPPKEAQEKDEFFYFFNHFCFYIINE
TKEYALAYKMNFAFYLPYGSLGVDVLKNVFDYLHHLNVPTILDIKMNDIGNTVKHYRKFIFD
YLRSDSCTANIYMGTQMLRDICLDEECKRYYSTFVLVKTTNADSHIFQNRLSLDGKEAYVVI
AEEVQKMAKQLHLEENGEFVGFVVGANCYDEIKKIRELFPDCYILAPGVGAQKGDLRKMLCN
GYSKNYEKVLINVGRAITKSGSPQQAAREYHQQIKEVLAELQE
111111111111111111100000000000000000000000000000000000000000
000000001111111111111000000000000001000000000000000000000000
000000000000000000000000000000000000000000000000000000000000
000000000000000000000000000000000000000000000000000000000000
000000000000000000000000000000000000000000000000000000000000
0000000000000000000000000000000000000000011
>DM_validata237
MVDTHKLADDVLQLLDNRIEDNYRVCVILVGSPGSGKSTIAEELCQIINEKYHTFLSEHPNV
IEVNDRLKPMVNLVDSLKTLQPNKVAEMIENQGLFKDHVEDVNFQPVKYSALTSNNEECTAV
VARGGTANAIRIAAVDNPVNVNKLAQDSINIAQIVPMDGFHLSRRCLDLFKDPQTAHKRRGS
PSTFDSNNFLQLCKILAKTSLCKVSSHHKFYSTSSVFEKLSKTFSQTIPDIFVPGFNHALKD
PTPDQYCISKFTRIVILEGLYLLYDQENWKKIYKTLADTGALLVYKIDIDYEATEERVAKRH
LQSGLVTTIAEGREKFRSNDLLNGRDIDNHLIKVDNIVHIRND
100000000000000000000000000000000000000000000000000000000000
000000000000000000000000000000000000000000000000011111000000
000000000000001111111111110000000000000000000000000000000000
000000000000000000000000111111100000000000000000000000000000
000000000000000000000000000000000000000000000000000000000000
0000000000000000000000000000000000000000000
>DM_validata238
GSHMPIQVLPPQLANQIAAGEVVERPASVVKELVENSLDAGATRIDIDIERGGAKLIRIRDN
GCGIKKDELALALARHATSKIASLDDLEAIISLGFRGEALASISSVSRLTLTSRTAEQQEAW
QAYAEGRDMNVTVKPAAHPVGTTLEVLDLFYNTPARRKFLRTEKTEFNHIDEIIRRIALARF
DVTINLSHNGKIVRQYRAVPEGGQKERRLGAICGTAFLEQALAIEWQHGDLTLRGWVADPNH
TTPALAEIQYCYVNGRMMRDRLINHAIRQACEDKLGADQQPAFVLYLEIDPHQVDVNVHPAK
HEVRFHQSRLVHDFIYQGVLSVLQQQLETPLPLDDEPQPAPR
111111111111111111111100000000000000000000000000000000000000
000000000000001111111111111000000000000000000000000000000000
000111110000000000000000000001111111111000000000000000000000
000000000000000000000000000000000000000000000000000000000000
000000000000000000000000000000000000000000000000000011111111
111110000000000000000000111111111111111111
>DM_validata239

MIGKRFFQTTSKKIAFAFDIDGVLFRGKKPIAGASDALKLLNRNKIPYILLTNGGGFSERAR
TEFISSKLDVDVSPLQIIQSHTPYKSLVNKYSRILAVGTPSVRGVAEGYGFQDVVHQTDIVR
YNRDIAPFSGLSDEQVMEYSRDIPDLTTKKFDAVLVFNDPHDWAADIQIISDAINSENGMLN
TLRNEKSGKPSIPIYFSNQDLLWANPYKLNRFGQGAFRLLVRRLYLELNGEPLQDYTLGKPT
KLTYDFAHHVLIDWEKRLSGKIGQSVKQKLPLLGTKPSTSPFHAVFMVGDNPASDIIGAQNY
GWNSCLVKTGVYNEGDDLKECKPTLIVNDVFDAVTKTLEKYA
11111111111110000000000000000000000000000000000000000000000000
00000000000000000000000000000000000000000000000000000000000000
00000000000000000000000000000000000000000000000000000000000000
00000000000000000000000000000000000000000000000000000000000000
00000000000000000001111111111111111111110000000000000000000000
000000000000000000000000000000000000000000
>DM_validata240
MRGSHHHHHHGSENLYFQSGAMEIREQLNLGGIVNAQNAQLSNCSDGAAQLESCGTAPDLKG
ITGWLNTPGNKPIDLKSLRGKVVLIDFWAYSCINCQRAIPHVVGWYQAYKDSGLAVIGVHTP
EYAFEKVPGNVAKGAANLGISYPIALDNNYATWTNYRNRYWPAEYLIDATGTVRHIKFGEGD
YNVTETLVRQLLNDAKPGVKLPQPSSTTTPDLTPRAALTPETYFGVGKVVNYGGGGAYDEGS
AVFDYPPSLAANSFALRGRWALDYQGATSDGNDAAIKLNYHAKDVYIVVGGTGTLTVVRDGK
PATLPISGPPTTHQVVAGYRLASETLEVRPSKGLQVFSFTYG
11111111111111111110000000000000000000000000000000000000000000
00000000000000000000000000000000000000000000000000000000000000
00000000000000000000000000000000000000000000000000000000000000
00000000000000000000000000000000000000000000000000000000000000
00000000000000000000000000000000000000000000000000000000001111
100000000000000000000000000000000000000000
>DM_validata241
MNGIDDLLNINDRIKQVQNERNELASKLQNLKQSLASNDTGGGSGGGSDSSDLLQREAILAN
ELNILDNLKTFLNLIKEVKTNLNILELENCYYSLQSLRKKMRNNAAYLKQSFNFQQSISTYV
DTLHLELVSTLYKILTNGFWKITENSIQFTPTVEWGKDKVHIEYDTFMDFVAQQYFPKGSLD
NQAWFILDMTSADSQEQVRAKLNTIMKEYMNLSRIVSMIKNSIFISGKEISYENEKNILVFS
KSSSHGQHCVSTVLTSFEAVCDFMLDGLAFRDRKTLSYELGPLFNTEFTKFVKNNASIILES
LDSPLKNLVSVINNKLTRLVAKSEVTNWTHSGKEIQDLLMN
11111111000000000000000000000000111111111111111111111000000000
00000000000000000000000000000000000000000000000000000000000000
00000000000000000000000000000000000000000000000000000000000000
00000000000000000000000000000000000000000000000000000000000000
00111111000000000000000000000000000000000000000000000000000000
00000000000000000000000000000000000000001
>DM_validata242
MTTAVARLQPSASGAKTRPVFPFSAIVGQEDMKLALLLTAVDPGIGGVLVFGDRGTGKSTAV
RALAALLPEIEAVEGCPVSSPNVEMIPDWATVLSTNVIRKPTPVVDLPLGVSEDRVVGALDI
ERAISKGEKAFEPGLLARANRGYLYIDECNLLEDHIVDLLLDVAQSGENVVERDGLSIRHPA
RFVLVGSGNPEEGDLRPQLLDRFGLSVEVLSPRDVETRVEVIRRRDTYDADPKAFLEEWRPK
DMDIRNQILEARERLPKVEAPNTALYDCAALCIALGSDGLRGELTLLRSARALAALEGATAV

GRDHLKRVATMALSHRLRRDPLDEAGSTARVARTVEETLP
11111111111111110000000000000000000000000000000000000000000000
00000000000000000000000000000000000000000000000000000000000000
00000000000000000000000000000000000000000000000000000000000000
00000000000000000000000000000000000000000000000000000000000000
00000000000000000000000000000000000000000000000000000000000000
0000000000000000001111111111100000000000
>DM_validata243
TPDNNKTVREFNVYWNVPTFMCHKYGLRFEEVSEKYGILQNWMDKFRGEEIAILYDPGMFPA
LLKDPNGNVVARNGGVPQLGNLTKHLQVFRDHLINQIPDKSFPGVGVIDFESWRPIFRQNWA
SLQPYKKLSVEVVRREHPFWDDQRVEQEAKRRFEKYGQLFMEETLKAAKRMRPAANWGYYAY
PYCYNLTPNQPSAQCEATTMQENDKMSWLFESEDVLLPSVYLRWNLTSGERVGLVGGRVKEA
LRIARQMTTSRKKVLPYYWYKYQDRRDTDLSRADLEATLRKITDLGADGFIIWGSSDDINTK
AKCLQFREYLNNELGPAVKRIALNNNANDRLTVDVSVDQV
11111111100000000000000000000000000000000000000000000000000000
00111111100000000000000000000000000000000000000000000000000000
00000000000000000000000000000000000000000000000000000000000000
00000000000000000000000000000000000000000000000000000000000000
00000000000000000000000000000000000000000000000000000000000000
0000000000000000000011111111111111111111
>DM_validata245
MKSDYDIPTTENLYFQGSAANAAGPASRYDVTLDQSDAELVEEIAWKLATQATGRPDDAEWV
EAARNAWHAWPATLRRDLAGFRRDSGPDGAIVLRGLPVDSMGLPPTPRVNGSVQREASLGAA
VLLMTACGLGDPGAFLPEKNGALVQDVVPVPGMEEFQGNAGSTLLTFHNENAFHEHRPDFVM
LLCLRADPTGRAGLRTACVRRVLPLLSDSTVDALWAPEFRTAPPPSFQLSGPEEAPAPVLLG
DRSDPDLRVDLAATEPVTERAAEALRELQAHFDATAVTHRLLPGELAIVDNRVTVHGRTEFT
PRYDGTDRWLQRTFVLTDLRRSRAMRPADGYVLGAAPQPA
11111111111111111111111111000000000000000000000000000000000000
00000000000000000000000000000000000000000000000000000000000000
00000000000000000000000000000000000000000000000000000000000000
00000000000000000000000000000000000000000000000011111100000000
00000000000000000000000000000000000000000000000000000000000000
0000000000000000000000000000000000000111
>DM_validata247
ADTGPALKAGHEYMIVTNYPNNLHVVDVASDTVYKSCVMPDKFGPGTAMMAPDNRTAYVLNN
HYGDIYGIDLDTCKNTFHANLSSVPGEVGRSMYSFAISPDGKEVYATVNPTQRLNDHYVVKP
PRLEVFSTADGLEAKPVRTFPMPRQVYLMRAADDGSLYVAGPDIYKMDVKTGKYTVALPLRN
WNRKGYSAPDVLYFWPHQSPRHEFSMLYTIARFKDDKQDPATADLLYGYLSVDLKTGKTHTQ
EFADLTELYFTGLRSPKDPNQIYGVLNRLAKYDLKQRKLIKAANLDHTYYCVAFDKKGDKLY
LGGTFNDLAVFNPDTLEKVKNIKLPGGDMSTTTPQVFIR
11100000000000000000000000000000000000000000000000000000000000
00000000000000000000000000000000000000000000000000000000000000
00000000000000000000000000000000000000000000000000000000000000
00000000000000000000000000000000011111110000000000000000000000

00000000000000000000000000000000000000000000000000000000000000
000000000000000000000000000000000000000
>DM_validata248
GRSRLLEDFRNNRYPNLQLREIAGHIMEFSQDQHGSRFIQLKLERATPAERQLVFNEILQAA
YQLMVDVFGNYVIQKFFEFGSLEQKLALAERIRGHVLSLALQMYGCRVIQKALEFIPSDQQN
EMVRELDGHVLKCVKDQNGNHVVQKCIECVQPQSLQFIIDAFKGQVFALSTHPYGCRVIQRI
LEHCLPDQTLPILEELHQHTEQLVQDQYGNYVIQHVLEHGRPEDKSKIVAEIRGNVLVLSQH
KFASNVVEKCVTHASRTERAVLIDEVCTMNDGPHSALYTMMKDQYANYVVQKMIDVAEPGQR
KIVMHKIRPHIATLRKYTYGKHILAKLEKYYMKNGVDLG
00000000000000000000000000000000000000000000000000000000000000
00000000000000000000000000000000000000000000000000000000000000
00000000000000000000000000000000000000000000000000000000000000
00000000000000000000000000000000000000000000000000000000000000
00000000000000000000000000000000000000000000000000000000000000
000000000000111111111111111111111111111
>DM_validata249
MIEFDNLTYLHGKPQGTGLLKANPEDFVVVEDLGFEPDGEGEHILVRILKNGCNTRFVADAL
AKFLKIHAREVSFAGQKDKHAVTEQWLCARVPGKEMPDLSAFQLEGCQVLEYARHKRKLRLG
ALKGNAFTLVLREVSNRDDVEQRLIDICVKGVPNYFGAQRFGIGGSNLQGAQRWAQTNTPVR
DRNKRSFWLSAARSALFNQIVAERLKKADVNQVVDGDALQLAGRGSWFVATTEELAELQRRV
NDKELMITAALPGSGEWGTQREALAFEQAAVAAETELQALLVREKVEAARRAMLLYPQQLSW
NWWDDVTVEIRFWLPAGSFATSVVRELINTTGDYAHIAE
00000000000000000000000000000000000000000000000000000000000000
00000000000000000000000000000000000000000000000000000000000000
00000000000000000000000000000000000000000000000000000011111111
11000000000000000000000000000000000000000000000000000000000000
00000000000000000000000000000000000000000000000000000000000000
000000000000000000000000000000111111111
>DM_validata250
MELRHTPARDLDKFIEDHLLPNTCFRTQVKEAIDIVCRFLKERCFQGTADPVRVSKVVKGGS
SGKGTTLRGRSDADLVVFLTKLTSFEDQLRRRGEFIQEIRRQLEACQREQKFKVTFEVQSPR
RENPRALSFVLSSPQLQQEVEFDVLPAFDALGQWTPGYKPNPEIYVQLIKECKSRGKEGEFS
TCFTELQRDFLRNRPTKLKSLIRLVKHWYQTCKKTHGNKLPPQYALELLTVYAWEQGSRKTD
FSTAQGFQTVLELVLKHQKLCIFWEAYYDFTNPVVGRCMLQQLKKPRPVILDPADPTGNVGG
GDTHSWQRLAQEARVWLGYPCCKNLDGSLVGAWTMLQKI
00000000000000000000000000000000000000000000000111000000000000
01111111100000000000000000000000000000000000000000000000000001
11111000000000000000000000000000000000000000000000000000000000
00000000000000000000000000000000000000000000000000000000000000
00000000000000000000000000000000000000000000000000000000000000
000000000000000000000000000000000000111
>DM_validata251
PRRRDPQDDVYLDITDRLCFAILYSRPKSASNVHYFSIDNELEYENFYADFGPLNLAMVYRY
CCKINKKLKSITMLRKKIVHFTGSDQRKQANAAFLVGCYMVIYLGRTPEEAYRILIFGETSY

IPFRDAAYGSCNFYITLLDCFHAVKKAMQYGFLNFNSFNLDEYEHYEKAENGDLNWIIPDRF
IAFCGPHSRARLESGYHQHSPETYIQYFKNHNVTTIIRLNKRMYDAKRFTDAGFDHHDLFFA
DGSTPTDAIVKEFLDICENAEGAIAVHCKAGLGRTGTLIACYIMKHYRMTAAETIAWVRICR
PGSVIGPQQQFLVMKQTNLWLEGDYFRQKLKGQENGQH
111000000000000000000000000000000000000000000000000000000000
000000000000000000000000000000000000000000000000000000000000
000000000000000000000000000000000000000000000000000000000000
000000000000000000000000000000000000000000000000000000000000
000000000000000000000000000000000000000000000000000000000000
00000000000000000000000000000001111111
>DM_validata253
MGSSGSMNHYTRWLELKEQNPGKYARDIAGLMNIREAELAFARVTHDAWRMHGDIREILAAL
ESVGETKCICRNEYAVHEQVGTFTNQHLNGHAGLILNPRALDLRLFLNQWASVFHIKENTAR
GERQSIQFFDHQGDALLKVYATDNTDMAAWSELLARFITDENTPLELKAVDAPVVQTRADAT
VVEQEWRAMTDVHQFFTLLKRHNLTRQQAFNLVADDLACKVSNSALAQILESAQQDGNEIMV
FVGNRGCVQIFTGVVEKVVPMKGWLNIFNPTFTLHLLEESIAEAWVTRKPTSDGYVTSLELF
AHDGTQIAQLYGQRTEGEQEQAQWRKQIASLIPEGVAA
111110000000000000000000000000000000000000000000000000000000
000000000000000000000000000000000000000000000000000000000000
000000000000000000000000000000000000000000000000111111100000
000000000000000000000000000000000000000000000000000000000000
000000000000000000000000000000000000000000000000000000000000
00000000000000000000000000000000011111
>DM_validata254
YDPIHEYVNHELRKRENEFSEHKNVKIFVASYNLNGCSATTKLENWLFPENTPLADIYVVGF
QEIVQLTPQQVISADPAKRREWESCVKRLLNGKCTSGPGYVQLRSGQLVGTALMIFCKESCL
PSIKNVEGTVKKTGLGGVSGNKGAVAIRFDYEDTGLCFITSHLAAGYTNYDERDHDYRTIAS
GLRFRRGRSIFNHDYVVWFGDFNYRISLTYEEVVPCIAQGKLSYLFEYDQLNKQMLTGKVFP
FFSELPITFPPTYKFDIGTDIYDTSDKHRVPAWTDRILYRGELVPHSYQSVPLYYSDHRPIY
ATYEANIVKVDREKKKILFEELYNQRKQEVRDASQTS
000000000000000000000000000000000000000000000000000000000000
000000011111000000000000000000000000000000000000000000000000
000000000000000111100000000000000000000000000000000000000000
000000000000000000000000000000000000000000000000000000000000
000000000000000000000000000000000000000000000000000000000000
0000000000000000000000000000000000000
>DM_validata255
TAPSQVLKIRRPDDWHLHLRDGDMLKTVVPYTSEIYGRAIVMPNLAPPVTTVEAAVAYRQRI
LDAVPAGHDFTPLMTCYLTDSLDPNELERGFNEGVFTAAKLYPANATANSSHGVTSVDAIMP
VLERMEKIGMPLLVHGEVTHADIDIFDREARFIESVMEPLRQRLTALKVVFEHITTKDAADY
VRDGNERLAATITPQHLMFNRNHMLVGGVRPHLYCLPILKRNIHQQALRELVASGFNRVFLG
TDSAPHARHRKESSCGCAGCFNAPTALGSYATVFEEMNALQHFEAFCSVNGPQFYGLPVNDT
FIELVREEQQVAESIALTDDTLVPFLAGETVRWSVKQ
111000000000000000000000000000000000000000000000000000000000

00000000000000000000000000000000000000000000111111100000000000
00000000000000000000000000000000000000000000000000000000000000
00000000000000000000000000000000000000000000000000000000000000
00000000000000000000000000000000000000000000000000000000000000
0000000000000000000000000000000000001
>DM_validata256
MSLKMETGKLLMELSNLDGPSGYETNVVSYIKSVIEPFVDEAKTTRHGSLIGYKKGKGIGKL
AFFAHVDEIGFVVSKVEGQFARLEPVGGVDPKVVYASKVRIYTKNGIERGVIGMLAPHLQDS
ESRKKVPTYDEIFVDLSLCERGVRVGDIAVIDQTAFETNGKVVGKALDNRASCGVLVKVLEF
LKRYDHPWDVYVVFSVQEETGCLGALTGAYEINPDAAIVMDVTFASEPPFSDHIELGKGPVI
GLGPVVDRNLVQKIIEIAKKHNVSLQEEAVGGRSGTETDFVQLVRNGVRTSLISIPLKYMHT
PVEMVDPRDVEELARLLSLVAVELEVEGGSHHHHHH
11110000000000000000000000000000000000000000000000000000000000
00000000000000000000000000111111110000000000000000000000000000
00000000000000000000000000000000000000000000000000000000000000
00000000000000000000000000000000000000000000000000000000000000
00000000000000000000000000000000011110000000000000000000000000
000000000000000000000000011111111111
>DM_validata257
APIQAPEISKCVVPPADLPPGAVVDNCCPPVASNIVDYKLPAVTTMKVRPAAHTMDKDAIAK
FAKAVELMKALPADDPRNFYQQALVHCAYCNGGYDQVNFPDQEIQVHNSWLFFPFHRWYLYF
YERILGKLIGDPSFGLPFWNWDNPGGMVLPDFLNDSTSSLYDSNRNQSHLPPVVVDLGYNGA
DTDVTDQQRITDNLALMYKQMVTNAGTAELFLGKAYRAGDAPSPGAGSIETSPHIPIHRWVG
DPRNTNNEDMGNFYSAGRDIAFYCHHSNVDRMWTIWQQLAGKPRKRDYTDSDWLNATFLFYD
ENGQAVKVRIGDSLDNQKMGYKYAKTPLPWLDSKP
00000000000000000000000000000000000000000000000000000000000000
00000000000000000000000000000000000000000000000000000000000000
00000000000000000000000000000000000000000000000000000000000000
00000000000000000000000000000000000000000000000000000000000000
00000000000000000000000000000000000000001111100000000000000000
00000000000000000000000000000001111
>DM_validata258
MKQLSVSLAILCAVATSEAFAADELQQRAQGLFKPVPAKAPTLKGNPASPVKVELGKMLYFD
PRLSASHLISCNTCHNVGLGGGDLQATSTGHGWQKGPRNAPTVLNSVFNTAQFWDGRAKDLA
EQAKGPVQAPVEMNNTPDQVVKTLNSIPDYVALFKKAFPGEKDPVTFDNMAKAIEVFEATLI
TPDSPFDQYLKGKKKALDGKQTAGLKLFLDKGCVACHGGLNLGGTGYFPFGVVEKPAENILP
LGDKGRFAVTNTAKDEYVFRAPSLRNVAITYPYFHSGVVWSLKEAVAVMGSAQFGIKLSDDE
SEAIAAFLGSLTGKQPKVVYPIMPASTDATPRPRL
11111111111111111111000000000000000000000000000000000000000000
00000000000000000000000000000000000000000000000000000000000001
11111110000000000000000000000000000000000000000000000000000000
00000000000000000000000000000000000000000000000000000001111111
11110000000000000000000000000000000000000000000000000000000000
00000000000000000000000000000000000

>DM_validata260
MIRLYPEQLRAQLNEGLRAAYLLLGNDPLLLQESQDAVRQVAAAQGFEEHHTFSIDPNTDWN
AIFSLCQAMSLFASRQTLLLLLPENGPNAAINEQLLTLTGLLHDDLLLIVRGNKLSKAQENA
AWFTALANRSVQVTCQTPEQAQLPRWVAARAKQLNLELDDAANQVLCYCYEGNLLALAQALE
RLSLLWPDGKLTLPRVEQAVNDAAHFTPFHWVDALLMGKSKRALHILQQLRLEGSEPVILLR
TLQRELLLLVNLKRQSAHTPLRALFDKHRVWQNRRGMMGEALNRLSQTQLRQAVQLLTRTEL
TLKQDYGQSVWAELEGLSLLLCHKPLADVFIDG
000000000000000000000000000000000000000000000000000000000000
000000000000000000000000000000000000000000000000000000000000
000000000000000000000000000000000000000000000000000000000000
000000000000000000000001110000000000000000000000000000000000
000000000000000000000000000000000000000000000000000000000000
000000001110000000000001111111111
>DM_validata261
MEVSCGQAESSEKPNAEDMTSKDYYFDSYAHFGIHEEMLKDEVRTLTYRNSMFHNRHLFKDK
VVLDVGSGTGILCMFAAKAGARKVIGIECSSISDYAVKIVKANKLDHVVTIIKGKVEEVELP
VEKVDIIISEWMGYCLFYESMLNTVLHARDKWLAPDGLIFPDRATLYVTAIEDRQYKDYKIH
WWENVYGFDMSCIKDVAIKEPLVDVVDPKQLVTNACLIKEVDIYTVKVEDLTFTSPFCLQVK
RNDYVHALVAYFNIEFTRCHKRTGFSTSPESPYTHWKQTVFYMEDYLTVKTGEEIFGTIGMR
PNAKNNRDLDFTIDLDFKGQLCELSCSTDYRMR
111111111111111111111111111000000000000000000000000000000000
000000000000000000000000000000000000000000000000000000000000
000000000000000000000000000000000000000000000000000000000000
000000000000000000000000000000000000000000000000000000000000
000000000000000000000000000000000000000000000000000000000000
000000000000000000000000000000000
>DM_validata262
AGALWEIEKELFTKLPAPSSAINSHLQPAKPKVPQKKPSKWDPPAEFKVDLSTAVSYNDIGD
INWKNLQQFKGIERSEKGTEGLFFVETESGVFIVKRSTNIESETFCSLLCMRLGLHAPKVRV
VSSNSEEGTNMLECLAAIDKSFRVITTLANQANILLMELVRGITLNKLTTTSAPEVLTKSTM
QQLGSLMALDVIVNNSDRLPIAWTNEGNLDNIMLSERGATVVPIDSKIIPLDASHPHGERVR
ELLRTLIAHPGHESSQFHSIRDIITLYTGYDVGTEGSISMQEGFLATVRECASFDLDAFERE
LLSWQESLQKCHNLSISPQAIPFILRMLRIFH
000000000000000000000000000000011111111111111100000000000000
000000000000000000000000000000000000000000000000000000000000
000000000000000000000000000000000000000000000000000000000000
000000000000000000000000000000000000000000000000000000000000
000000000000000000000000000000000000000000000000000000000000
00000000000000000000000000000000
>DM_validata263
NIPEIENANLKPALKDSVLPDGFYSTTNHPTHVKVNDEWIEVANPKMDAVIVVYPEEKRAET
KVIRKVKKGDFVLIGHNGIRVMPPEKSREAGQLFEFMNSEVSSEKPKEAIIKRIAKEMHEIR
EEYKKTGTGGIAIVGGPAIIHTGGGPALAKMVELGYIQAILAGNALATHDIESALYGTSLGV
NIKTAKPVTGGHKHHIYAINAINDAGNIKNAVESGVLKEGIMYQCIKNNIPYVLAGSIRDDG

PIPDVITDSMVAQDKMRTTVMDKKMVIMLSTLLHSVATGNLMPSYIKTVCVDIQPSTVTKLM
DRGTSQAIGVVTDVGVFLVLLLKELERLELQE
000000000000000000000000000000000000000000000000000000000000
000000000000000000000000011111111111110000000000000000000000
000000000000000000000000000000000000000000000000000000000000
000000000000000000000000000000000000000000000000000000000000
000000000000000000000000000000000000000000000000000000000000
00000000000000000000000000000011
>DM_validata264
MATFVRNAWYVAALPEELSEKPLGRTILDTPLALYRQPDGVVAALLDICPHRFAPLSDGILV
NGHLQCPYHGLEFDGGGQCVHNPHGNGARPASLNVRSFPVVERDALIWIWPGDPALADPGAI
PDFGCRVDPAYRTVGGYGHVDCNYKLLVDNLMDLGHAQYVHRANAQTDAFDRLEREVIVGDG
EIQALMKIPGGTPSVLMAKFLRGANTPVDAWNDIRWNKVSAMLNFIAVAPEGTPKEQSIHSR
GTHILTPETEASCHYFFGSSRNFGIDDPEMDGVLRSWQAQALVKEDKVVVEAIERRAYVEA
NGIRPAMLSCDEAAVRVSREIEKLEQLEAAR
100000000000000000000000000000000000000000000000000000000000
000000000000000000000000000000000000000000000000000000000000
000000000000000000000000000000011111111111111111111000000000
000000000000000000000111110000000000000000000000000000000000
000000000000000000000000000000000000000000000000000000000000
0000000000000000000000000000001
>DM_validata265
MPFPFGKSHKSPADIVKNLKESMAVLEKQDISDKKAEKATEEVSKNLVAMKEILYGTNEKEP
QTEAVAQLAQELYNSGLLSTLVADLQLIDFEGKKDVAQIFNNILRRQIGTRTPTVEYICTQQ
NILFMLLKGYESPEIALNCGIMLRECIRHEPLAKIILWSEQFYDFFRYVEMSTFDIASDAFA
TFKDLLTRHKLLSAEFLEQHYDRFFSEYEKLLHSENYVTKRQSLKLLGELLLDRHNFTIMTK
YISKPENLKLMMNLLRDKSRNIQFEAFHVFKVFVANPNKTQPILDILLKNQAKLIEFLSKFQ
NDRTEDEQFNDEKTYLVKQIRDLKRPAQQEA
111111110000000000000000000111000000000000000000000001111111
000000000000000000000000000000000000000000000000000000000000
000000000000000000000000000000000000000000000000000000000000
000000000000000000000000000000000000000000000000000000000000
000000000000000000000000000000000000000000000000000000000000
0001000000000000000000000000011
>DM_validata266
MSLINEATLAESIRRLRQGERATLAQAMTLVESRHPRHQALSTQLLDAIMPYCGNTLRLGVT
GTPGAGKSTFLEAFGMLLIREGLKVAVIAVDPSSPVTGGSILGDKTRMNDLARAEAAFIRPV
PSSGHLGGASQRARELMLLCEAAGYDVVIVETVGVGQSETEVARMVDCFISLQIAGGGDDLQ
GIKKGLMEVADLIVINKDDGDNHTNVAIARHMYESALHILRRKYDEWQPRVLTCSALEKRGI
DEIWHAIIDFKTALTASGRLQQVRQQQSVEWLRKQTEEEVLNHLFANEDFDRYYRQTLLAVK
NNTLSPRTGLRQLSEFIQTQYFDEGHHHHHH
100000000000000000000000000000000000000000000000000000000000
000000000000000000000000000000001111111111111110000000000000
001111100000000000000000000000000000000000000000000001111110

000000000000000000000000000000000000000000000000000000000000
000000000000000000000000000000000000000000000000000000000000
000000000000000000011111111111
>DM_validata268
MKLTGKQTWEFENPLFVNSSGTAVGPKEKEGPLGHLFDKSYDEMHCNQKNWEMAERKLMEDA
VQSALSKQNLKKEDIDIFLAGDLLNQNVTANYVARHLKIPFLCLFGACSTSMESIAISSALI
DGGFAKRALAATSSHNATAERQFRYPTEYGGQKPGTATSTVTGSGAVVLSQQPGGIKITSAT
VGRVIDLGITDSQDMGSAMAPAAADTIKQHLEDLGRTPDDYDLILTGDLSGVGSPILKDLLK
EEGINVGTKHNDCGLMIYTPDQQVFAGGSGCACSAVVTFAHIFKEIEAGRLNRVLVVATGAL
LSPTIIQQKESIPCIAHGVVFERAERGNA
000000000000000000000000000000000000000000000000000000000000
000000000000000000000000000000000000000000000000000000000000
000000000000000000000001111111111111100000000000000000000000
000000000000000000000000000000000000000000000000000000000000
000000000000000000001111111110000000000000000000000000000000
00000000000000000000000011111
>DM_validata270
GHMRRELAIEFSRVTESAALAGYKWLGRGDKNTADGAAVNAMRIMLNQVNIDGTIVIGEGEI
DEAPMLYIGEKVGTGRGDAVDIAVDPIEGTRMTAMGQANALAVLAVGDKGCFLNAPDMYMEK
LIVGPGAKGTIDLNLPLADNLRNVAAALGKPLSELTVTILAKPRHDAVIAEMQQLGVRVFAI
PDGDVAASILTCMPDSEVDVLYGIGGAPEGVVSAAVIRALDGDMNGRLLARHDVKGDNEENR
RIGEQELARCKAMGIEAGKVLRLGDMARSDNVIFSATGITKGDLLEGISRKGNIATTETLLI
RGKSRTIRRIQSIHYLDRKDPEMQVHIL
000000000000000000000000000000000000000000000000000000000001
110000000000000000000000000011111100000000000000000000000000
000000000000000000000000000000000000000000000000000000000000
000000000000000000000000000000000000000000000000111111111111
111111111111100000000000000000000000000000000000000000000000
0000000000000000011111111111
>DM_validata271
MRGSHHHHHHGSGSMHQRQTLYQLMDGLNTFYQQSLQQPVATSARQYLEKRGLSHEVIARFA
IGFAPPGWDNVLKRFGGNPENRQSLIDAGMLVTNDQGRSYDRFRERVMFPIRDKRGRVIGFG
GRVLGNDTPKYLNSPETDIFHKGRQLYGLYEAQQDNAEPNRLLVVEGYMDVVALAQYGINYA
VASLGTSTTADHIQLLFRATNNVICCYDGDRAGRDAAWRALETALPYMTDGRQLRFMFLPDG
EDPDTLVRKEGKEAFEARMEQAMPLSAFLFNSLMPQVDLSTPDGRARLSTLALPLISQVPGE
TLRIYLRQELGNKLGILDDSQLERLMPK
111111111111111111100000000000000000000000000000000000000000
000000000000000000000000000000000111000000000000000000000000
000000000000000000000000000000000000000000000000000000000000
000000000000000000000000000000000000000000000000000000000000
000000000000000000000000000000000000000000000000000000000000
0000000000000000000000111111
>DM_validata272
MNLQRFPRYPLTFGPTPIQPLARLSKHLGGKVHLYAKREDCNSGLAFGGNKTRKLEYLIPEA

LAQGCDTLVSIGGIQSNQTRQVAAVAAHLGMKCVLVQENWVNYSDAVYDRVGNIQMSRILGA
DVRLVPDGFDIGFRRSWEDALESVRAAGGKPYAIPAGCSDHPLGGLGFVGFAEEVRAQEAEL
GFKFDYVVVCSVTGSTQAGMVVGFAADGRADRVIGVDASAKPAQTREQITRIARQTAEKVGL
ERDIMRADVVLDERFAGPEYGLPNEGTLEAIRLCARTEGMLTDPVYEGKSMHGMIEMVRNGE
FPEGSRVLYAHLGGVPALNGYSFIFRDG
000000000000000000000000000000000000000000000000000000000000
000000000000000000000000000000000000000000000000000000000000
000001111111111000000000000000000000000000000000000000000000
000000000000000000000000000000000000000000000000000000000000
000000000000000000000000000000000000000000000000000000000000
0000000000000000000000000000
>DM_validata273
DYFQGAMGSKPAYSFHVTADGQMQPVPFPPDALIGPGIPRHARQINTLNHGEVVCAVTISNP
TRHVYTGGKGCVKVWDISHPGNKSPVSQLDCLNRDNYIRSCKLLPDGCTLIVGGEASTLSIW
DLAAPTPRIKAELTSSAPACYALAISPDSKVCFSCCSDGNIAVWDLHNQTLVRQFQGHTDGA
SCIDISNDGTKLWTGGLDNTVRSWDLREGRQLQQHDFTSQIFSLGYCPTGEWLAVGMESSNV
EVLHVNKPDKYQLHLHESCVLSLKFAYCGKWFVSTGKDNLLNAWRTPYGASIFQSKESSSVL
SCDISVDDKYIVTGSGDKKATVYEVIY
000000000000000001111100000000000000000000000000000000000000
000000000000000000011100000000000000000000000000000000000000
000111000000000000000000000000000000000000000000000000000000
000000000000000000000000000000000000000000000000000000000000
000000000000000000000000000000000000000000000000000000000000
000000000000000000000000000
>DM_validata274
VASGYGGIRVGGPYFDDLSKGQVFDWAPGVTLSLGLAAAHQSIVGNRLRLALDSDLCAAVTG
MPGPLAHPGLVCDVAIGQSTLATQRVKANLFYRGLRFHRFPAVGDTLYTRTEVVGLRANSPK
PGRAPTGLAGLRMTTIDRTDRLVLDFYRCAMLPASPDWKPGAVPGDDLSRIGADAPAPAADP
TAHWDGAVFRKRVPGPHFDAGIAGAVLHSTADLVSGAPELARLTLNIAATHHDWRVSGRRLV
YGGHTIGLALAQATRLLPNLATVLDWESCDHTAPVHEGDTLYSELHIESAQAHADGGVLGLR
SLVYAVSDSASEPDRQVLDWRFSALQF
111111100000000000000000000000000000000000000000000000000000
000000000000000000000000000000000000000000000000000000000000
000000000000000000000000000000000000000000000000000000000000
000000000000000000000000000000000000000000000000000000000000
000000000000000000000000000000000000000000000000000000000000
000000000000000000000000000
>DM_validata275
MREEQPHLATTWAARGWVEEEGIGSATLGRLVRAWPRRAAVVNKADILDEWADYDTLVPDYP
LEIVPFAEHPLFLAAEPHQRQRVLTGMWIGYNERVIATEQLIAEPAFDLVMHGVFPGSDDPL
IRKSVQQAIVDESFHTYMHMLAIDRTRELRKISERPPQPELVTYRRLRRVLADMPEQWERDI
AVLVWGAVAETCINALLALLARDATIQPMHSLITTLHLRDETAHGSIVVEVVRELYARMNEQ
QRRALVRCLPIALEAFAEQDLSALLLELNAAGIRGAEEIVGDLRSTAGGTRLVRDFSGARKM
VEQLGLDDAVDFDFPERPDWSPHTPR

111111111111111111111100000000000000000000000000000000000000
000000000000000000000000000000000000000000000000000000000000
000000000000000000000000000000000000000000000000000000000000
000000000000000000000000000000000000000000000000000000000000
000000000000000000000000000000000000000001111111000000000000
0000000000000000000111111
>DM_validata276
MIAVKTCGKLYWAGEYAILEPGQLALIKDIPIYMRAEIAFSDSYRIYSDMFDFAVDLRPNPD
YSLIQETIALMGDFLAVRGQNLRPFSLAIYGKMEREGKKFGLGSSGSVVVLVVKALLALYNL
SVDQNLLFKLTSAVLLKRGDNGSMGDLACIAAEDLVLYQSFDRQKVAAWLEEENLATVLERD
WGFSISQVKPTLECDFLVGWTKEVAVSSHMVQQIKQNINQNFLTSSKETVVSLVEALEQGKS
EKIIEQVEVASKLLEGLSTDIYTPLLRQLKEASQDLQAVAKSSGAGGGDCGIALSFDAQSTK
TLKNRWADLGIELLYQERIGHDDKS
000000000000000000000000000000000000000000000000000000000000
000000000000000000000000000000000000000000000000000000000000
000000000000000000000000000000000000000000000000000000000000
000000000000000000000000000000000000000000000000000000000000
000000000000000000000000000000000000000000000000000000000000
000000000000000000111111
>DM_validata277
GMTKLPATSDIYISFFMFTTNLQPDNLDYRRIVVAHIKKLQRFGYSGFEFPIAPGLPENYAQ
DLENYTNLRHYLDSEGLENVKISTNVGATRTFDPSSNYPEQRQEALEYLKSRVDITAALGGE
IMMGPIVIPYGVFPTTDFNEPIWSDELQEHLKVRYANAQPILDKLGEYAEIKKVKLAIEPIT
HWETPGPNKLSQLIEFLKGVKSKQVGVVIDSAHEILDGEGPEIFKTQVEYLAQQGRLHYVQV
SPPDRGALHTSWLPWKSFLTPIVKVYDGPIAVEIFNAIPAFTNSLRLTRRKFWIPDEDPPNQ
YPNAYDIADEAIKVTRKELKKIGSK
111111110000000000000000000000000000000000000000000000000000
000000000000000000000000000000000000000000000000000000000000
000000000000000000000000000000000000000000000000000000000000
000000000000000000000000000000000000000000000000000000000000
000000000000000000000000000000000000000000000000000000000000
000000000000000000000011
>DM_validata280
MSSGVDLGTENLYFQSNAMYEIKGHHHISMVTKNANENNHFYKNVLGLRRVKMTVNQDDPSM
YHLFYGDKTGSPGTELSFFEIPLVGRTYRGTNAITRIGLLVPSEDSLHYWKERFEKFDVKHS
EMTTYANRPALQFEDAEGLRLVLLVSNGEKVEHWETWEKSEVPAKHQIQGMGSVELTVRRLD
KMASTLTEIFGYTEVSRNDQEAIFQSIKGEAFGEIVVKYLDGPTEKPGRGSIHHLAIRVKND
AELAYWEEQVKQRGFHSSGIIDRFYFKSLYFRESNGILFEIATDGPGFTVDGDVEHLGEKLD
LPPFLEDQRAEIEANLAPIEEK
111100000000000000000000000000000000000000000000000000000000
000000000000000000000000000000000000000000000000000000000000
000000000000000000000000000000000000000000000000000000000000
000000000000000000000000000000000000000000000000000000000000
000000000000000000000000000000000000000000000000000000000000

0000000000000000000000
>DM_validata282
THNAHNWRLGQAPANWYNDTYPLSPPQRTPAGIRYRIAVIADLDTESRAQEENTWFSYLKKG
YLTLSDSGDKVAVEWDKDHGVLESHLAEKGRGMELSDLIVFNGKLYSVDDRTGVVYQIEGSK
AVPWVILSDGDGTVEKGFKAEWLAVKDERLYVGGLGKEWTTTTGDVVNENPEWVKVVGYKGS
VDHENWVSNYNALRAAAGIQPPGYLIHESACWSDTLQRWFFLPRRASQERYSEKDDERKGAN
LLLSASPDFGDIAVSHVGAVVPTHGFSSFKFIPNTDDQIIVALKSEEDSGRVASYIMAFTLD
GRFLLPETKIGSVKYEGIEFI
111111111111111000000000000000000000000000000000000000000000
000000000000000000000000000000000000000000000000000000000000
000000000000000000000000000000000000000000000000000000000000
000000000000000000000000000000000000000000000000000000000000
000000000000000000000000000000000000000000000000000000000000
000000000000000000000
>DM_validata283
MNTRNFSLPQLQNLPIEEARIVADALAVHATSRQIDSAASKLAALAEAGLKGDRQAYAAYQQ
LLYVLSLSDDVATAQTRRWLARAIYRVEERFMPAADLSRALSEEDFQKRLEQEIAAQSRERH
PMSQYVFSGSASRAQLQVFLRHQWFRTFRLYRDAADLLVNLTDVDEAAALARYLYGELGEED
EKGSHPRLLAKLLEAIGLEADFQAVSTMPEEIAYLNNRARAFRHAEVGWGLAVFYITELVVP
GNHEKLYRALLQAGLSEDQAEYYKVHISLVPPRAKREWQLIARRIPDVQFQNAFLTSLSQHF
RVERAYYDAIWEEMQSVK
100000000000000000000000000000000000000000000000000000000000
000000000000000000000000000000000000000000000000000000011000
000000000000000000000000000000000000000000000000000000000000
000000000000000000000000000000000000000000000000000000000000
000000000000000000000000000000111000000000000000000000000000
000000000000000001
>DM_validata284
MSFESKKPMRTWSHLAEMRKKPSEYDIVSRKLHYSTNNPDSPWELSPDSPMNLWYKQYRNAS
PLKHDNWDAFTDPDQLVYRTYNLMQDGQESYVQSLFDQFNEREHDQMVREGWEHTMARCYSP
LRYLFHCLQMSSAYVQQMAPASTISNCCILQTADSLRWLTHTAYRTHELSLTYPDAGLGEHE
RELWEKEPGWQGLRELMEKQLTAFDWGEAFVSLNLVVKPMIVESIFKPLQQQAWENNDTLLP
LLIDSQLKDAERHSRWSKALVKHALENPDNHAVIEGWIEKWRPLADRAAEAYLSMLSSDILH
AQYLERSTSLRASILTV
110000000000000000000000000000000000000000000000000000000000
000000000000000000000000000000000000000000000000000000000000
000000000000000000000000000000000000000000000000000000000000
000000000000000000000000000000000000000000000000000000000000
000000000000000000000000000000000000000000000000000000001111
11111111111111111
>DM_validata285
DNQPEPGNTSAGEGEVLTVFHAGSLSVPFEELEAEFEAQHPGVDVQREAAGSAQSVRKITEL
GKKADVLASADYALIPSLMVPEYADWYAAFARNQMILAYTNESKYGDEINTDNWYEILRRPD
VRYGFSNPNDDPAGYRSQMVTQLAESYYNDDMIYDDLMLANTGMTLTTEENGTALIHVPASE

EISPNTSKIMLRSMEVELSSALETGEIDYLYIYRSVAEQHGFEYVALPPAIDLSSLEYADNY
SKVQVEMVNGEVVTGSPIVYGVTIPNNAENSELATEFVALLLGETGQQIFIENGQPPIVPAI
AEGKDSMPEELQALVV
11111111111111000000000000000000000000000000000000000000000000
00000000000000000000000000000000000000000000000000000000000000
00000000000000000000000000000000000000000000000000000000000000
00000000000000000000000000000000000000000000000000000000000000
00000000000000000000000000000000000000000000000000000000000000
0000000000000000
>DM_validata287
GPLGSMTTKQTVSLFIWLPESKQKTLFISTKNHTQFELNNIIFDVTLSTELPDKEPNAIITK
RTHPVGKMADEMRKYEKDHPKVLFLESSAIHDMMSSREEINALLIKNNIPIPNSFSVKSKEE
VIQLLQSKQLILPFIVKPENAQGTFNAHQMKIVLEQEGIDDIHFPCLCQHYINHNNKIVKVF
CIGNTLKWQTRTSLPNVHRCGIKSVDFNNQHLEDILSWPEGVIDKQDIIENSANRFGSKILE
DPILLNLTSEAEMRDLAYKVRCALGVQLCGIDFIKENEQGNPLVVDVNVFPSYGGKVDFDWF
VEKVALCYTEVAKI
11111111100000000000000000000000000000000000000000000000000000
00000000000000000000000000000000000000000000000000000000000000
00000000000000000000000000000000000000000000000000000000000000
00000000000000000000000000000000000000000000000000000000000000
00000000000000000000000000000000000000000000000000000000000000
00000000001111
>DM_validata289
MNTQKPFENHLKSVDDLKTTYEEYRAGFIAFALEKNKRSTPYIERARALKVAASVAKTPKDL
LYLEDIQDALLYASGISDKAKKFLTEDDKKESINNLIENFLEPAGEEFIDELIFRYLLFQGD
SLGGTMRNIAGALAQQKLTRAIISALDIANIPYKWLDSRDKKYTNWMDKPEDDYELETFAKG
ISWTINGKHRTLMYNITVSLVKKNVDICLFNCEPEIYTPQKVHQQPEKYLLLGELKGGIDPA
GADEHWKTANTALTRIRNKFSEKGLSPKTIFIGAAIEHSMAEEIWDQLQSGSLTNSANLTKT
EQVGSLCRWIINI
11110000000000000000000000000000000000000000000000000000000000
00000000000000000000000000000000000000000000000000000000000000
00000000000000000000000000000000000000000000000000000000000000
00000000000000000000000000000000001111111110000000000000000000
00000000000000000000000000000000000000000000000000000000000000
0000000000000
>DM_validata292
HGDNNSKPKPGGNSGNRGNNGDGASAKVGEITITPDNSKPGRYISSNPEYSLLAKLIDAESI
KGTEVYTFHTRKGQYVKVTVPDSNIDKMRVDYVNWKGPKYNNKLVKRFVSQFLLFRKEEKEK
NEKEALLKASELVSGMGDKLGEYLGVKYKNVAKEVANDIKNFHGRNIRSYNEAMASLNKVLA
NPKMKVNKSDKDAIVNAWKQVNAKDMANKIGNLGKAFKVADLAIKVEKIREKSIEGYNTGNW
GPLLLEVESWIIGGVVAGVAISLFGAVLSFLPISGLAVTALGVIGIMTISYLSSFIDANRVS
NINNIISSVIR
11111111111111111111111100000000000000000000000000000000000000
00000000000000000000000000000000000000000000000000000000000000

00000000000000000000000000000000000000000000000000000000000000
00000000000000000000000000000000000000000000000000000000000000
00000000000000000000000000000000000000000000000000000000000000
00000000000
>DM_validata293
EPQHNVMQMGGDFANNPNAQQFIDKMVNKHGFDRQQLQEILSQAKRLDSVLRLMDNQAPTTS
VKPPSGPNGAWLRYRKKFITPDNVQNGVVFWNQYEDALNRAWQVYGVPPEIIVGIIGVETRW
GRVMGKTRILDALATLSFNYPRRAEYFSGELETFLLMARDEQDDPLNLKGSFAGAMGYGQFM
PSSYKQYAVDFSGDGHINLWDPVDAIGSVANYFKAHGWVKGDQVAVMANGQAPGLPNGFKTK
YSISQLAAAGLTPQQPLGNHQQASLLRLDVGTGYQYWYGLPNFYTITRYNHSTHYAMAVWQL
GQAVALARVQ
00000000000000000000000000000000000000000000000000000000111111
11111000000000000000000000000000000000000000000000000000000000
00000000000000000000000000000000000000000000000000000000000000
00000000000000000000000000000000000000000000000000000000000000
00000000000000000000000000000000000000000000000000000000000000
0000000000
>DM_validata295
MGRIESAFDLGFIRGMTFGFVGQHGTWGTDEARASMRALAEQPFNWVTLAFAGLMEHPGDPA
IAYGPPVTVSDDEIASMAELAHALGLKVCLKPTVNCRDGTWRGEIRFEKEHGPDLESWEAWF
GSYSDMMAHYAHVAKRTGCEMFCVGCEMTTAEPHEAMWRETIARVRTEYDGLVTYNCNHGRE
EHVRFWDAVDLISSSAYYPIDRWRDRVPVLREVAEAHEKPLFFMEVGCPSRSGSGACPWDYR
HPGAVCLDEQARFYEAMFAAMPDEPWFKGYMLWEWPWKLYPREAASEDGSYCIYGKPAEDVV
ARAFSAIANR
11110000000000000000000000000000000000000000000000000000000000
00000000000000000000000000000000000000000000000000000000000000
00000000000000000000000000000000000000000000000000000000000000
00000000000000000000000000000000000000000000000000000000000000
00000000000000000000000000000000000000000000000000000000000000
0000000000
>DM_validata299
GSHMTALDWRSALTADEQRSVRALVTATTAVDGVAPVGEQVLRELGQQRTEHLLVAGSRPGG
PIIGYLNLSPPRGAGGAMAELVVHPQSRRRGIGTAMARAALAKTAGRNQFWAHGTLDPARAT
ASALGLVGVRELIQMRRPLRDIPEPTIPDGVVIRTYAGTSDDAELLRVNNAAFAGHPEQGGW
TAVQLAERRGEAWFDPDGLILAFGDSPRERPGRLLGFHWTKVHPDHPGLGEVYVLGVDPAAQ
RRGLGQMLTSIGIVSLARRLGGRKTLDPAVEPAVLLYVESDNVAAVRTYQSLGFTTYSVDTA
YALAGTDN
11111000000000000000000000000000000000000000000000000000000000
00000000000111000000000000000000000000000000000000000000000000
00000000000000000000000000000000000000000000000000000000000000
00000000000000000000000001111110000000000000000000000000000000
00000000000000000000111111111000000000000000000000000000000000
00011111
>DM_validata300

MTELPLQFAEPDDDLERVRATLYSLDPDGDRTAGVLRDTLDQLYDGQRTGRWNFDQLHKTEK
THMGTLVEINLHREFQFGDGFETDYEIAGVQVDCKFSMSQGAWMLPPESIGHICLVIWASDQ
QCAWTAGLVKVIPQFLGTANRDLKRRLTPEGRAQVVKLWPDHGKLQENLLLHIPGDVRDQIF
SAKSSRGNQHGQARVNELFRRVHGRLIGRAVIATVAQQDDFMKRVRGSGGARSILRPEGIII
LGHQDNDPKVANDLGLPVPRKGQVVAARVVPADEGDQRQTAEIQGRRWAVAVPGDPIVEAPV
VPRKSAE
111111111000000000000000000000000000000000000000000000000000
000000000000000000000000000000000000000000000000000000000000
000000000000000000000000000000000000000000000000000000000000
000011110000000000000000000000000000000000000000000000000000
000001111100000000000000000000000000000000000000000000000000
0111111
>DM_validata301
EVVLVNVTENFNMWKNDMVEQMHEDIISLWDQSLKPCVKLTPLCVGAGSCNTSVITQACPKV
SFEPIPIHYCAPAGFAILKCNNKTFNGTGPCTNVSTVQCTHGIRPVVSTQLLLNGSLAEEEV
VIRSVNFTDNAKTIIVQLNTSVEINCTGAGHCNIARAKWNNTLKQIASKLREQFGNNKTIIF
KQSSGGDPEIVTHSFNCGGEFFYCNSTQLFNSTWFNSTWSTEGSNNTEGSDTITLPCRIKQI
INMWQKVGKAMYAPPISGQIRCSSNITGLLLTRDGGNSNNESEIFRPGGGDMRDNWRSELYK
YKVVKIE
000000000000000000000000000000000000000000000000000000000000
000000000000000000000000000000000000000000000000000000000000
000000000000000000000000000000000000000000000000000000000000
000000000000000000000000000000000011111111111100000000000000
000000000000000000000000000000000000000000000000000000000000
0000001
>DM_validata303
AAIVAGPKQNCEPDLMPYARPFAVGKRTCSGIVGLPSGSDPAFSQPKSVLDAGLTCQGASPS
SVSKPILLVPGTGTTGPQSFDSNWIPLSAQLGYTPCWISPPPFMLNDTQVNTEYMVNAITTL
YAGSGNNKLPVLTWSQGGLVAQWGLTFFPSIRSKVDRLMAFAPDYKGTVLAGPLDALAVSAP
SVWQQTTGSALTTALRNAGGLTQIVPTTNLYSATDEIVQPQVSNSPLDSSYLFNGKNVQAQA
VCGPLFVIDHAGSLTSQFSYVVGRSALRSTTGQARSADYGITDCNPLPANDLTPEQKVAAAA
LLAPAA
111110000000000000000000000000000000000000000000000000000000
000000000000000000000000000000000000000000000000000000000000
000000000000000000000000000000000000000000000000011111100000
000000000000000000000000000000000000000000000000000000000000
000000000000000000000000000000000000000000000000011111111111
111111
>DM_validata304
VNYAAGLSPYADKGKCGLPEIFDPPEELERKVWELARLVWQSSSVVFHTGAGISTASGIPDF
RGPHGVWTMEERGLAPKFDTTFESARPTQTHMALVQLERVGLLRFLVSQNVDGLHVRSGFPR
DKLAELHGNMFVEECAKCKTQYVRDTVVGTMGLKATGRLCTVAKARGLRACRGELRDTILDW
EDSLPDRDLALADEASRNADLSITLGTSLQIRPSGNLPLATKRRGGRLVIVNLQPTKHDRHA
DLRIHGYVDEVMTRLMKHLGLEIPAWDGPRVLERALPPLPRPPTPKLEPKEESPTRINGSIP

AGPKQE
111111111111100000000000000000000000000000000000000000000000
000000000000000000000000000000000000000000000000000000000000
000000000000000000000000000000000000000011111111000000000000
000000000000000000000000000000000000000000000000000000000000
000000000000000000000000000000000000000000001111111111111111
111111
>DM_validata305
ANMPPGSVEQVAAKVVPSVVMLETDLGRQSEEGSGIILSAEGLILTNNHVIAAAAKPPLGSP
PPKTTVTFSDGRTAPFTVVGADPTSDIAVVRVQGVSGLTPISLGSSSDLRVGQPVLAIGSPL
GLEGTVTTGIVSALNRPVSTTGEAGNQNTVLDAIQTDAAINPGNSGGALVNMNAQLVGVNSA
IATLGADSADAQSGSIGLGFAIPVDQAKRIADELISTGKASHASLGVQVTNDKDTLGAKIVE
VVAGGAAANAGVPKGVVVTKVDDRPINSADALVAAVRSKAPGATVALTFQDPSGGSRTVQVT
LGKAEQ
111110000000000000000000011110000000000000000000000000000010
000000000000000000000000000000000000000000000000000000000000
000000000000000000001111110000000000000000000000000000000000
000011111111000000000000000000000000000000000000011110000000
011100011110000000000010000000000000000000111111111111111111
000011
>DM_validata306
RDFSPVPWSQYFESMEDVEVENETGKDTFRVYKSGSEGPVLLLLHGGGHSALSWAVFTAAII
SRVQCRIVALDLRSHGETKVKNPEDLSAETMAKDVGNVVEAMYGDLPPPIMLIGHSMGGAIA
VHTASSNLVPSLLGLCMIDVVEGTAMDALNSMQNFLRGRPKTFKSLENAIEWSVKSGQIRNL
ESARVSMVGQVKQCEGITSPEGSKKDHPYTWRIELAKTEKYWDGWFRGLSNLFLSCPIPKLL
LLAGVDRLDKDLTIGQMQGKFQMQVLPQCGHAVHEDAPDKVAEAVATFLIRHRFAEPIGGFQ
CVFPGC
000000000000000000000000000000000000000000000000000000000000
000000000000000000000000000000000000000000000000000000000000
000000000000000000000000000000000000000000000000000000000000
000000000000000111111111111000000000000000000000000000000000
000000000000000000000000000000000000000000000000000000001111
111111
>DM_validata308
DQLIPVINKLQDVFNTLGSDPLDLPQIVVVGSQSSGKSSVLENIVGRDFLPRGSGIVTRRPL
ILQLTHLPIADDGSQTQEWGEFLHKPNDMFYDFSEIREEIIRDTDRMTGKNKGISAQPINLK
IYSPHVVNLTLVDLPGITKVPVGDQPTDIEQQIRRMVMAYIKKQNAIIVAVTPANTDLANSD
ALQLAKEVDPEGKRTIGVITKLDLMDKGTDAMEVLTGRVIPLTLGFIGVINRSQEDIIAKKS
IRESLKSEILYFKNHPIYKSIANRSGTAYLSKTLNKLLMFHIRDTLPDLKVKVSKMLSDVQG
ELSTY
000000000000000000000000000000000000000000000000011111111000
000000000000000000000000000000000000000000011111111000000000
000000000000000111111111000000000000000000000000000000000000
000000000000000000000000000000000000000000000000000000000000

00000000000000000000000000000000000000000000000000000000001111
11111
>DM_validata309
MEVKTIKVFTTVDNTNLHTQLVDMSMTYGQQFGPTYLDGADVTKIKPHVNHEGKTFFVLPSD
DTLRSEAFEYYHTLDESFLGRYMSALNHTKKWKFPQVGGLTSIKWADNNCYLSSVLLALQQL
EVKFNAPALQEAYYRARAGDAANFCALILAYSNKTVGELGDVRETMTHLLQHANLESAKRVL
NVVCKHCGQKTTTLTGVEAVMYMGTLSYDNLKTGVSIPCVCGRDATQYLVQQESSFVMMSAP
PAEYKLQQGTFLCANEYTGNYQCGHYTHITAKETLYRIDGAHLTKMSEYKGPVTDVFYKETS
YTTTI
000000000000000000000000000000000000000000000000000000000000
000000000000000000000000000000000000000000000000000000000000
000000000000000000000000000000000000000000000000000000000000
000000000000000000000000000000000000000000000000000000000000
000000000000000000000000000000000000000000000000000000000000
00011
>DM_validata311
GMAGAIASRMSFSSLKRKQPKTFTVRIVTMDAEMEFNCEMKWKGKDLFDLVCRTLGLRETWF
FGLQYTIKDTVAWLKMDKKVLDHDVSKEEPVTFHFLAKFYPENAEEELVQEITQHLFFLQVK
KQILDEKIYCPPEASVLLASYAVQAKYGDYDPSVHKRGFLAQEELLPKRVINLYQMTPEMWE
ERITAWYAEHRGRARDEAEMEYLKIAQDLEMYGVNYFAIRNKKGTELLLGVDALGLHIYDPE
NRLTPKISFPWNEIRNISYSDKEFTIKPLDKKIDVFKFNSSKLRVNKLILQLCIGNHDLFMR
RRKA
111111111111111111110000000000000000000000000000000000000000
000000000000000000000000000000000000000000000000000000000000
000000000000000000000000000000000000000000000000000000000000
000000000000000000000000000000000000000000000000000000000000
000000000000000000000000000000000000000000000000000000000000
0000
>DM_validata312
MRKPILAALTLAGLGLAQEFITIGSGSTTGVYFPVATGIAKLVNDANVGIRANARSTGGSVA
NINAINAGEFEMALAQNDIAYYAYQGCCIPAFEGKPVKTIRALAALYPEVVHVVARKDAGIR
TVADLKGKRVVVGDVGSGTEQNARQILEAYGLTFDDLGQAIRVSASQGIQLMQDKRADALFY
TVGLGASAIQQLALTTPIALVAVDLNRIQAIAKKYPFYVGFNIPGGTYKGVDVTTPTVAVQA
MLIASERLSEETVYKFMKAVFGNLEAFKKIHPNLERFFGLEKAVKGLPIPLHPGAERFYKEA
GVLK
111111111111111100000000000000000000000000000000000000000000
000000000000000000000000000000000000000000000000000000000000
000000000000000000000000000000000000000000000000000000000000
000000000000000000000000000000000000000000000000000000000000
000000000000000000000000000000000000000000000000000000000000
0000
>DM_validata313
PTITPNKLTLKIGRAEGRPGDTVEIPVNLYGVPQKGIASGDFVVSYDPNVLEIIEIEPGELI
VDPNPTKSFDTAVYPDRKMIVFLFAEDSGTGAYAITEDGVFATIVAKVKEGAPEGFSAIEIS

EFGAFADNDLVEVETDLINGGVLVTNKPVIEGYKVSGYILPDFSFDATVAPLVKAGFKVEIV
GTELYAVTDANGYFEITGVPANASGYTLKISRATYLDRVIANVVVTGDTSVSTSQAPIMMWV
GDIVKDNSINLLDVAEVIRCFNATKGSANYVEELDINRNGAINMQDIMIVHKHFGATSSDYD
AQ
111111000000000000000000000000000000000000000000000000000000
000000000000000000000000000000000000000000000000000000000000
000000000000000000000000000000000000000000000000000000000000
000000000000000000000000000000000000000000000000000000000000
000000000000000000000000000000000000000000000000000000000001
11
>DM_validata314
MVPAESNAVQAKLAKTLQRFENKIKAGDYYEAHQTLRTIANRYVRSKSYEHAIELISQGALS
FLKAKQGGSGTDLIFYLLEVYDLAEVKVDDISVARLVRLIAELDPSEPNLKDVITGMNNWSI
KFSEYKFGDPYLHNTIGSKLLEGDFVYEAERYFMLGTHDSMIKYVDLLWDWLCQVDDIEDST
VAEFFSRLVFNYLFISNISFAHESKDIFLERFIEKFHPKYEKIDKNGYEIVFFEDYSDLNFL
QLLLITCQTKDKSYFLNLKNHYLDFSQAYKSELEFLGQEYFNIVAPKQTNFLQDMMSGFLGG
SK
111111111111000000000000000000000000000000000000000000000000
000000000000000000000000000000000000000000000000000000000000
000000000000000000000000000000000000000000000000000000000000
000000000000000000000000000000000000000000000000000000000000
000000000000000000000000000000000000000000111111111111111111
11
>DM_validata315
TSCVAEEPIKKIAIFGGTHGNELTGVFLVTHWLKNGAEVHRAGLEVKPFITNPRAVEKCTRY
IDCDLNRVFDLENLSKEMSEDLPYEVRRAQEINHLFGPKNSDDAYDVVFDLHNTTSNMGCTL
ILGDSGNDFLIQMFHYIKTCMAPLPCSVYLIEHPSLKYATTRSIAKYPVGIEVGPQPHGVLR
ADILDQMRRMLKHALDFIQRFNEGKEFPPCAIDVYKIMEKVDYPRNESGDVAAVIHPNLQDQ
DWKPLHPGDPVFVSLDGKVIPLGGDCTVYPVFVNEAAYYEKKEAFAKTTKLTLNAKSIRSTL
H
110000000000000000000000000000000000000000000000000000000000
000000000000000000000000000000000000000000000000000000000000
000000000000000000000000000000000000000000000000000000000000
000000000000000000000000000000000000000000000000000000000000
000000000000000000000000000000000000000000000000000000000001
1
>DM_validata316
MGSKNESTASKASGTASEKKKIEYLDKTYEVTVPTDKIAITGSVESMEDAKLLDVHPQGAIS
FSGKFPDMFKDITDKAEPTGEKMEPNIEKILEMKPDVILASTKFPEKTLQKISTAGTTIPVS
HISSNWKENMMLLAQLTGKEKKAKKIIADYEQDLKETKTKINDKAKDSKALVIRIRQGNIYI
YPEQVYFNSTLYGDLGLKAPNEVKAAKAQELISLEKLSEMNPDHIFVQFSDDENADKPDALK
DLEKNPIWKSLKAVKEDHVYVNSVDPLAQGGTAWSKVRFLKAAAEKLTQNKLAAALEHHHHH
H
111111111111111100000000000000000000000000000000000000000000

00000000000000000000000000000000000000000000000000000000000000
00000000000000000000000000000000000000000000000000000000000000
00000000000000000000000000000000000000000000000000000000000000
00000000000000000000000000000000000000000000000000011111111111
1
>DM_validata317
MARKRSNTYRSINEIQRQKRNRWFIHYLNYLQSLAYQLFEWENLPPTINPSFLEKSIHQFGY
VGFYKDPVISYIACNGALSGQRDVYNQATVFRAASPVYQKEFKLYNYRDMKEEDMGVVIYNN
DMAFPTTPTLELFAAELAELKEIISVNQNAQKTPVLIRANDNNQLSLKQVYNQYEGNAPVIF
AHEALDSDSIEVFKTDAPYVVDKLNAQKNAVWNEMMTFLGIKNANLEKKERMVTDEVSSNDE
QIESSGTVFLKSREEACEKINELYGLNVKVKFRYDIVEQMRRELQQIENVSRGTSDGETNE
11111111111111110000000000000000000000000000000000000000000000
00000000000000000000000000000000000000000000000000000000000000
00000000000000000000000000000000000000000111000000000000000000
00000000000000000000000000000000000000000001111111111111110000
0000000000000000000000000000000000001111111111111111111111111
>DM_validata318
MDWGRFVEEKVREIRETVGDSKAIIALSGGVDSSTAAVLAHKAIGDRLHAVFVNTGFLRKGE
PEFVVKTFRDEFGMNLHYVDAQDRFFSALKGVTDPEEKRKIIGRVFIEVFEEVAKKIGAEYL
IQGTIAPDWIESQGKIKSHHNVGGLPEKLNLKLIEPLRDLYKDEVRELAKFLGLPEKIYNRM
PFPGPGLAVRVIGEVTPEKIRIVREANAIVEEEVERAGLRPWQAFAVLLGVKTVGVQGDIRA
YKETIAVRIVESIDGMTANAMNVPWEVLQRIAFRITSEIPEVGRVLYDITNKPPATIEFE
11111000000000000000000000000000000000000000000000000000000000
00000000000000000000000000000000000000000000000000000000000000
00000111111111111111111111111100000000000000000000000000000000
00000000000000000000000000000000000000000000000000000000000000
000000000000000000000000000000000000000000000000000000000000
>DM_validata319
MSEAADVERVYAAMEEAAGLLGVACARDKIYPLLSTFQDTLVEGGSVVVFSMASGRHSTELD
FSISVPTSHGDPYATVVEKGLFPATGHPVDDLLADTQKHLPVSMFAIDGEVTGGFKKTYAFF
PTDNMPGVAELSAIPSMPPAVAENAELFARYGLDKVQMTSMDYKKRQVNLYFSELSAQTLEA
ESVLALVRELGLHVPNELGLKFCKRSFSVYPTLNWETGKIDRLCFAVISNDPTLVPSSDEGD
IEKFHNYATKAPYAYVGEKRTLVYGLTLSPKEEYYKLGAYYHITDVQRGLLKAFDSLED
11000000000000000000000000000000000000000000000000000000000000
00000000000000000000000000000000000000000000000000000000000000
00000000000000000000000000000000000000000000000000000000000000
00000000000000000000000000000000000000000000000000000000000000
00000000000000000000000000000000000000000000000000000000001111
>DM_validata320
GTDDEEKDNNGQGSVSVETNYLPIADPYVMFYNNKYYAYGTGGTTAGEGFACFSSDDLKNWK
REGQALSATDSYGTWGFWAPEVYYVESKKKFYLFYSAEEHICVATSTTPEGPFRQEVKQPIW
SEKSIDTSLFIDDDGTPYLYFVRFTDGNVIWVAQMTDDLMSIKTETLNQCIKAEVSWELLQG
KVAEGPSLLKKNGVYYLIYSANHYENKGYGVGYATSDTPMGPWVKYSKNPLLQGDAATGLVG
TGHGAPFQCKDGSWKYIFHAHWSAAEIQPRTSYIKDFAISDQGVVTISGTVIKPRVLK

111111111111111000000000000000000000000000000000000000000000
000000000000000000000000000000000000000000000000000000000000
000000000000000000000000000000000000000000000000000000000000
000000000000000000000000000000000000000000000000000000000000
0000000000000000000000000000000000000000000000000000000
>DM_validata321
EVSTTQNDTLKVMTHNVYMLSTNLYPNWGQTERADLIGAADYIKNQDVVILNEVFDNSASDR
LLGNLKKEYPNQTAVLGRSSGSEWDKTLGNYSSSTPEDGGVAIVSKWPIAEKIQYVFAKGCG
PDNLSNKGFVYTKIKKNDRFVHVIGTHLQAEDSMCGKTSPASVRTNQLKEIQDFIKNKNIPN
NEYVLIGGDMNVNKINAENNNDSEYASMFKTLNASVPSYTGHTATWDATTNSIAKYNFPDSP
AEYLDYIIASKDHANPSYIENKVLQPKSPQWTVTSWFQKYTYNDYSDHYPVEATISMK
111111000000000000000000000000000000000000000000000000000000
000000000000000000000000000000000000000000000000000000000001
111000000000000000000000000000011111110000000000000000000000
000000000000000000000000000000000000000000000000000000000000
0000000000000000000000000000000000000000000000000000001
>DM_validata323
MDSHCRVRPAGPAVPADCDPPRITHAALAARLGDARLLTLYDQATWSEGPAWWEAQRTLVWS
DLVGRRVLGWREDGTVDVLLDATAFTNGNAVDAQQRLVHCEHGRRAITRSDADGQAHLLVGR
YAGKRLNSPNDLIVARDGAIWFTDPPFGLRKPSQGCPADPELAHHSVYRLPPDGSPLQRMAD
LDHPNGLAFSPDEQTLYVSQTPEQGHGSVEITAFAWRDGALHDRRHFASVPDGLPDGFCVDR
GGWLWSSSGTGVCVFDSDGQLLGHIPTPGTASNCTFDQAQQRLFITGGPCLWMLPLP
111100000000000000000000000000000000000000000000000000000000
000000000000000000000000000000000000000000000000000000000000
000000000000000000000000000000000000000000000000000000000000
000000000000000000000111100000000000000000000000000000000000
0000000000000000000000000000000000000000000000000000000
>DM_validata324
MIPFKDITLADRDTITAFTMKSDRRNCDLSFSNLCSWRFLYDTQFAVIDDFLVFKFWAGEQL
AYMMPVGNGDLKAVLRKLIEDADKEKHNFCMLGVCSNMRADLEAILPERFIFTEDRAYADYI
YLRSDLATLKGKKFQAKRNHINRFRNTYPDYEYTPITPDRIQECLDLEAEWCKVNNCDQQEG
TGNERRALIYALHNFEALGLTGGILHVNGKIVAFTFGMPINHETFGVHVEKADTSIDGAYAM
INYEFANRIPEQYIYINREEDLGIEGLRKAKLSYQPVTILEKYMACLKDHPMDMVRW
000000000000000000000000000000000000000000000000000000000000
000000000000000000000000000000000000000000000000000000000000
000000000000000000000000000000000000000000000000000000000000
000000000000000000000000000000000000000000000000000000000000
0000000000000000000000000000000000000000000000111111111
>DM_validata326
HSGRPVLGSSHHHHHHSSGMMLRVAVPNKGALSEPATEILAEAGYRRRTDSKDLTVIDPVNN
VEFFFLRPKDIAIYVGSGELDFGITGRDLVCDSGAQVRERLALGFGSSSFRYAAPAGRNWTT
ADLAGMRIATAYPNLVRKDLATKGIEATVIRLDGAVEISVQLGVADAIADVVGSGRTLSQHD
LVAFGEPLCDSEAVLIERAGTDGQDQTEARDQLVARVQGVVFGQQYLMLDYDCPRSALKKAT
AITPGLESPTIAPLADPDWVAIRALVPRRDVNGIMDELAAIGAKAILASDIRFCRF

1111111111111111111000000000000000000000000000000000000000000
000000000000000000000000000000000000000000000000000000000000
000000000000000000000000000000000000000000000000000000000000
000000000000000001111111110000000000000000000000000000000000
000000000000000000000000000000000000000000000000000000

>DM_validata327

GALGVVDEVHAGKEIGLQIYSLSQELYKGDVAANLRKVKDMGYSKLELAGYGKGAIGGVPMM
DFKKMAEDAGLKIISSHVNPVDTSISDPFKAMIFKYSKEVTPKIMEYWKATAADHAKLGCKY
LIQPMMPTITTHDEAKLVCDIFNQASDVIKAEGIATGFGYHNHNMEFNRVATKEQQEKVKGN
PFAAFMKVGDQIYDLMLKDTDPSKVYFEMDVYWTVMGQNDPVEYMQKHPDRIKVLHIKDRAV
FGQSGMMNFEMIFKQMYANGIKDYFVELEQMPDGRTQFAGVKDCADYLIKAPFVK

111111111110000000000000000000000000000000000000000000000000
000000000000000000000000000000000000000000000000000000000000
000000000000000000000000000000000000000000000000000000111111
111100000000000000000000000000000000000000000000000000000000
00000000000000000000000000000000000000000000000000000

>DM_validata328

MSETPRLLFVHAHPDDESLSNGATIAHYTSRGAQVHVVTCTLGEEGEVIGDRWAQLTADHAD
QLGGYRIGELTAALRALGVSAPIYLGGAGRWRDSGMAGTDQRSQRRFVDADPRQTVGALVAI
IRELRPHVVVTYDPNGGYGHPDHVHTHTVTTAAVAAAGVGSGTADHPGDPWTVPKFYWTVLG
LSALISGARALVPDDLRPEWVLPRADEIAFGYSDDGIDAVVEADEQARAAKVAALAAHATQV
VVGPTGRAAALSNNLALPILADEHYVLAGGSAGARDERGWETDLLAGLGFTASGT

100000000000000000000000000000000000000000000000000000000000
000000000000000000000000000000000111111100000000000000000000
000000000000000000000000000000000000001111000000000000000000
000000000000000000011111111000000000000000000000000000000000
00000000000000000000000000000000000000000000000011111

>DM_validata329

SSSSTAAASAKKILVKHVTVIGGGLMGAGIAQVAAATGHTVVLVDQTEDILAKSKKGIEESL
RKVAKKKFAENPKAGDEFVEKTLSSISTSTDAASVVHSTDLVVEAIVENLKVKSELFKRLDK
FAAEHTIFASNTSSLQITSLANATTRQDRFAGLHFFNPVPLMKLVEVVKTPMTSQKTLESLV
DFSKTLGKHPVSCKDTPGFIVNRLLVPYLIEAVRLYERGDASKEDIDTAMKLGAGYPMGPFE
LLDYVGLDTTKFIIDGWHEMDSQNPLFQPSPAMNKLVAENKFGKKTGEGFYKYK

111111111110000000000000000000000000000000000000000000000000
000000000000000000000000000000000000000000000000000000000000
000000000000000000000000000000000000000000000000000000000000
000000000000000000000000000000000000111110000000000011110000
0000000000000000001111111110000000000000000001111111

>DM_validata330

MARVTTGITSSHIPALGAAIQTGTSDNDYWGPVFKGYQPIRDWIKQPGNMPDVVILVYNDHA
SAFDMNIIPTFAIGCAETFKPADEGWGPRPVPDVKGHPDLAWHIAQSLILDEFDMTIMNQMD
VDHGCTVPLSMIFGEPEEWPCKVIPFPVNVVTYPPPSGKRCFALGDSIRAAVESFPEDLNVH
VWGTGGMSHQLQGPRAGLINKEFDLNFIDKLISDPEELSKMPHIQYLRESGSEGVELVMWLI
MRGALPEKVRDLYTFYHIPASNTALGAMILQPEETAGTPLEPRKVMSGHSLAQA

00000000000000000000000000000000000000000000000000000000000000
00000000000000000000000000000000000000000000000000000000000000
00000000000000000000000000000000000000000000000000000000000000
00000000000000000000000000000000000000000000000000000000000000
000000000000000000000000000000000000000000000000000111
>DM_validata331
MTTYTFDQVEEAIEQLYPDFTINTIEISGEGNDCIAYEINGNFIFKFPKHSRASINLLNEVT
VLKTIHNELSLPIPEVVFTGMPSEMCQMSFAGFTKIKGVPLTPLLLKNLPKQSQDQAAKDLA
RFLSELHSINISGFKSNLVLDFREKINEDNKKIKKLLSRELKGHQMKKVDDFYRDILDNEIY
FKYYPCLIHNDFSSDHILFDTEKNTICGIIDFGDAAISDPDNDFISLMEDDEEYGMEFVSKI
LNHYKHKDIPTVLEKYMMKEKYWSFEKIIYGKEYGYMDWYEEGLNEIRSIKIK
10000000000000000000000000000000000000000000000000000000000000
00000000000000000000000000000000000000000000000000000000000000
00000000000000000000000000000000000000000000000000000000000000
00000000000000000000000000000000000000000000000000000000000000
00000000000000000000000000000000000000000000000011111
>DM_validata332
MSNAALATAPHALELDVHPVAGRIGAEIRGVKLSPDLDAATVEAIQAALVRHKVIFFRGQTH
LDDQSQEGFAKLLGEPVAHPTVPVVDGTRYLLQLDGAQGQRANSWHTDVTFVEAYPKASILR
SVVAPASGGDTVWANTAAAYQELPEPLRELADKLWAVHSNEYDYASLKPDIDPAKLERHRKV
FTSTVYETEHPVVRVHPISGERALQLGHFVKRIKGYSLADSQHLFAVLQGHVTRLENTVRWR
WEAGDVAIWDNRATQHYAVDDYGTQPRIVRRVTLAGEVPVGVDGQLSRTTRKG
11111111111100000000000000000000000000000000000000000000000000
00000000000000000111111111111100001111110000000000000000000000
00000000000000000000000000000000000000000111111111111111111111
11111000000000000000000000000000000000000000000000000000000000
00000000000000000000000000000000000000000000000000011
>DM_validata333
MGSASEQRVTLTNADKVLYPATGTTKSDIFDYYAGVAEVMLGHIAGRPATRKRWPNGVDQPA
FFEKQLALSAPPWLSRATVAHRSGTTTYPIIDSATGLAWIAQQAALEVHVPQWRFVAEPGSG
ELNPGPATRLVFDLDPGEGVMMAQLAEVARAVRDLLADIGLVTFPVTSGSKGLHLYTPLDEP
VSSRGATVLAKRVAQRLEQAMPALVTSTMTKSLRAGKVFVDWSQNSGSKTTIAPYSLRGRTH
PTVAAPRTWAELDDPALRQLSYDEVLTRIARDGDLLERLDADAPVADRLTRY
11111100000000000000000000000000000000000000000000000000000000
00000000000000000000000000000000000000000000000000000000000000
00000000000000000000000000000000000000000000000000000000000000
00000000000000000000000000000000000000000000000000000000000000
0000000000000000000000000000000000000000111111111111
>DM_validata334
DISSTEIAVYWGQREDGLLRDTCKTNNYKIVFISFLDKFGCEIRKPELELEGVCGPSVGNPC
SFLESQIKECQRMGVKVFLALGGPKGTYSACSADYAKDLAEYLHTYFLSERREGPLGKVALD
GIHFDIQKPVDELNWDNLLEELYQIKDVYQSTFLLSAAPGCLSPDEYLDNAIQTRHFDYIFV
RFYNDRSCQYSTGNIQRIRNAWLSWTKSVYPRDKNLFLELPASQATAPGGGYIPPSALIGQV
LPYLPDLQTRYAGIALWNRQADKETGYSTNIIRYLNATAMPFTSNLLKYPS

00000000000000000000000000000000000000000000000000000000000000
00000000000000000000000000000000000000000000000000000000000000
00000000000000000000000000000000000000000000000000000000000000
00000000000000000000000000000000000000000000000000000000000000
000000000000000000000000000000000001111111111111111
>DM_validata335
MTMAGQTINDRLLAARHSLAGQGLAKSVCKATTEECIGPKKKHLDYLVHCANEPNVSIPHLA
NLLIERSQNANWVVVYKSLITTHHLMAYGNERFMQYLASSNSTFNLSSFLDKGTVQDGGMGV
PGGRMGYDMSPFIRRYAKYLNEKSLSYRAMAFDFCKVKRGKEEGSLRSMNAEKLLKTLPVLQ
AQLDALLEFDCQSNDLSNGVINMSFMLLFRDLIRLFACYNDGIINLLEKYFDMNKKHARDAL
DLYKKFLVRMDRVGEFLKVAENVGIDKGDIPDLTKAPSSLLDALEQHLATL
11111111111111111111100000000000000000000000000000000000000000
00000000000000000000000000000000000000000000000000000011111000
00000000000000000000000000000000000000111100000000000000000000
00000000000000000000000000000000000000000000000000000000000000
000000000000000000000000000000000000000000000000000
>DM_validata336
MSLVPATNYIYTPLNQLKGGTIVNVYGVVKFFKPPYLSKGTDYCSVVTIVDQTNVKLTCLLF
SGNYEALPIIYKNGDIVRFHRLKIQVYKKETQGITSSGFASLTFEGTLGAPIIPRTSSKYFN
FTTEDHKMVEALRVWASTHMSPSWTLLKLCDVQPMQYFDLTCQLLGKAEVDGASFLLKVWDG
TRTPFPSWRVLIQDLVLEGDLSHIHRLQNLTIDILVYDNHVHVARSLKVGSFLRIYSLHTKL
QSMNSENQTMLSLEFHLHGGTSYGRGIRVLPESNSDVDQLKKDLESANLTA
11111000000000000000000000000000000000000000000000000000000000
00000000000000000000000000000000000000000000000000000000000000
00000000000000000000000100000000000000000000000000000000000000
00000000000001100000000000000000000000000000000000000000000000
000000000000000000000000000000000000000000000000000
>DM_validata337
MMAIATKRRGVAAVMSLGVATMTAVPALAQDVLGDLPVIGKPVNGGMNFQPASSPLAHDQQW
LDHFVLYIITAVTIFVCLLLLICIVRFNRRANPVPARFTHNTPIEVIWTLVPVLILVAIGAF
SLPILFRSQEMPNDPDLVIKAIGHQWYWSYEYPNDGVAFDALMLEKEALADAGYSEDEYLLA
TDNPVVVPVGKKVLVQVTATDVIHAWTIPAFAVKQDAVPGRIAQLWFSVDQEGVYFGQCSEL
CGINHAYMPIVVKAVSQEKYEAWLAGAKEEFAADASDYLPASPVKLASAE
11111111111111111111111111111000000000000000000000000000000000
00000000000000000000000000000000000000000000000000000000000000
00000000000000000000000000000000000000000000000000000000000000
00000000000000000000000000000000000000000000000000000000000000
00000000000000000000000000000000011111111111111111
>DM_validata338
MPVDKNLRDLEPGIHTDLEGRLTYGGYLRLDQLLSAQQPLSEPAHHDEMLFIIQHQTSELWL
KLLAHELRAAIVHLQRDEVWQCRKVLARSKQVLRQLTEQWSVLETLTPSEYMGFRDVLGPSS
GFQSLQYRYIEFLLGNKNPQMLQVFAYDPAGQARLREVLEAPSLYEEFLRYLARFGHAIPQQ
YQARDWTAAHVADDTLRPVFERIYENTDRYWREYSLCEDLVDVETQFQLWRFRHMRTVMRVI
GFKRGTGGSSGVGFLQQALALTFFPELFDVRTSVGVDNRPPQGSADAGKR

11111111111111111111000000000000000000000000000000000000000000
00000000000000000000000000000000000000000000000000000000000000
00000000000000000000000000000000000000000000000000000000000000
00000000000000000000000000000000000000000000000000000000000000
00111111000000000000000000000000000000000011111111
>DM_validata339
GMRESYANENQFGFKTINSDIHKIVIVGGYGKLGGLFARYLRASGYPISILDREDWAVAESI
LANADVVIVSVPINLTLETIERLKPYLTENMLLADLTSVKREPLAKMLEVHTGAVLGLHPMF
GADIASMAKQVVVRCDGRFPERYEWLLEQIQIWGAKIYQTNATEHDHNMTYIQALRHFSTFA
NGLHLSKQPINLANLLALSSPIYRLELAMIGRLFAQDAELYADIIMDKSENLAVIETLKQTY
DEALTFFENNDRQGFIDAFHKVRDWFGDYSEQFLKESRQLLQQANDLKQG
11111111111100000000000000000000000000000000000000000000000000
00000000000000000000000000000000000000000000000000000000000000
00000000000000000000000000000000000000000000000000000000000000
00000000000000000000000000000000000000000000001111000000000000
00000000000000000000000000000000000000000000111111
>DM_validata340
MASDKFPVAENPSSSFKYESAVQYRPAPDSYLNPCPQAGRIVKETYTGINGTKSLNVYLPYG
YDPNKKYNIFYLMHGGGENENTIFSNDVKLQNILDHAIMNGELEPLIVVTPTFNGGNCTAQN
FYQEFRQNVIPFVESKYSTYAESTTPQGIAASRMHRGFGGFAMGGLTTWYVMVNCLDYVAYF
MPLSGDYWYGNSPQDKANSIAEAINRSGLSKREYFVFAATGSEDIAYANMNPQIEAMKALPH
FDYTSDFSKGNFYFLVAPGATHWWGYVRHYIYDALPYFFHELEHHHHHH
11111111111111000000000000000000000000000000000000000000000000
00000000000000000000000000000000000000000000000000000000000000
00000000000000000000000000000000000000000000000000000000000000
00000000000000000000000000000000000000000000000000000000000000
0000000000000000000000000000000000000000000000000
>DM_validata341
STETLSFTPDNINADISLGTLSGKTKERVYLAEEGGRKVSQLDWKFNNAAIIKGAINWDLMP
QISIGAAGWTTLGSRGGNMVDQDWMDSSNPGTWTDEARHPDTQLNYANEFDLNIKGWLLNEP
NYRLGLMAGYQESRYSFTARGGSYIYSSEEGFRDDIGSFPNGERAIGYKQRFKMPYIGLTGS
YRYEDFELGGTFKYSGWVESSDNDEHYDPKGRITYRSKVKDQNYYSVAVNAGYYVTPNAKVY
VEGAWNRVTNKKGNTSLYDHNNNTSDYSKNGAGIENYNFITTAGLKYTF
11111111110000000000000000000000000000000000000000000000000000
00000000000000000000000000000000000000000000000000000000000000
00000000000000000000000000000000000000000000000000000000000000
00000000000000000000000000000000000000000000000000000000000000
0000000000000000000000000000000000000000000000000
>DM_validata342
KTTYNRWWKKKVWFGRKDMSYKVIVDSCGEFTPEMKADGGFEHVALGIQIEDTQWTDDDSLK
QEELLLKIAESTSCAKTSCPSPERYMESYHCDAERIYVVTLSAELSGSYNSAVLGKNLYEEE
YGEKQIHVFNSRSASVGETLIALKVQQCEKAGMTFEEVVESVECYIEEQHTYFVLENLDTLR
KNGRLTGIKSLVAGALNIKPIMGSTPQGTICQKEKARGMKKALVKMADCVAADVVNAGDKIL
AIAHCNCEERAKEVQRLLKERFAVKSSFIVDTSGISTVYANDGGIIVVV

11111111111111110000000000000000000000000000000000000000000000
00000000000000000000000000000000000000000000000000000000000000
00000000000000000000000000000000000000000000000000000000000000
00000000001100000000000000000000000000000000000000000000000000
0000000000000000000000000000000000000000000000000
>DM_validata343
MGSDKIHHHHHHVIIESRIEKGKPVVGMETTVFVHGLPRKEAIELFRRAKEISREKGFQLAV
IGILKGKIVAGMSEEELEAMMREGADKVGTREIPIVVAEGKNAATTVSATIFLSRRIGIEVV
VTGGTGGVHPGRVDVSQDLTEMSSSRAVLVSSGIKSILDVEATFEMLETLEIPLVGFRTNEF
PLFFSRKSGRRVPRIENVEEVLKIYESMKEMELEKTLMVLNPVPEEYEIPHDEIERLLEKIE
LEVEGKEVTPFLLKKLVEMTNGRTLKANLALLEENVKLAGEIAVKLKRS
11110000000000000000000000000000000000000000000000000000000000
00000000000000000000000000000000000000000000000000000000000000
00000000000000000000000000000000000000000000000000000000000000
00000000000000000000000000000000000000000000000000000000000000
0000000000000000000000000000000000000000000000001
>DM_validata344
GAHMNDIKQLLWNGELNVLVSIDPSFLMKGSPREIAVLRIRVPRETYLVNYMPLIWNKIKSF
LSFDPLTDSEKYFWFEHNKTPIPWNYPVGVLFDCLAGKSATFTTSFENQVKDVLTFLRIHLV
MGDSLPPTIIPIASSKTQAEKFWFHQWKQVCFILNGSSKAIMSLSVNEARKFWGSVITRNFQ
DFIEISNKISSSRPRHIPLIIQTSRTSGTFRISQPTISMTGVNPTLKDIEGDILDVKEGING
NDVMVICQGIEIPWHMLLYDLYSKLRSFDGFLYITLVPIKGGDKASSEL
11000000000000000000000000000000000000000000000000000000000000
00000111000000000000000000000000000000001111111110000000000000
00000000000000000000000000000000000000000000000000000000000000
00000000000000000000000000000000000000000000000000000000000111
1000000000000000000000000000000000000000111111111
>DM_validata345
MGSDKIHHHHHHMTNANGNLKKCPITISSYTLGTEVSFPKRVKVAAENGFDGIGLRAENYVD
ALAAGLTDEDMLRILDEHNMKVTEVEYITQWGTAEDRTAEQQKKEQTTFHMARLFGVKHINC
GLLEKIPEEQIIVALGELCDRAEELIIGLEFMPYSGVADLQAAWRVAEACGRDNAQLICDTW
HWARANQTAESIKNVPADRIVSIQLCDVHETPYKELREESLHDRLAPGEGYGDTVGFAKILK
EHGVNPRVMGVEVISDSMVATGLEYAALKVYNATKKVLDEAWPEISPR
11111111111111111111100000000000000000000000000000000000000000
00000000000000000000000000000000000000000000000000000000000000
00000000000000000000000000000000000000000000000000000000000000
00000000000000000000000000000000000000000000000000000000000000
000000000000000000000000000000000000000000000000
>DM_validata346
MRNYCTKTFGSAFSVTVVPTLKDNFSYLINDHTTHTLAAVDVNADYKPILTYIEEHLKQQGN
ADVTYTFSTILSTHKHWDHSGGNAKLKAELEAMNSTVPVVVVGGANDSIPAVTKPVREGDRV
QVGDLSVEVIDAPCHTRGHVLYKVQHPQHPNDGVALFTGDTMFIAGIGAFFEGDEKDMCRAM
EKVYHIHKGNDYALDKVTFIFPGHEYTSGFMTFSEKTFPDRASDDLAFIQAQRAKYAAAVKT
GDPSVPSSLAEEKRQNLFLRVADPAFVAKMNQGNAHALMMYLYNACD

00000000000000000000000000000000000000000000000000000000011111
11100000000000000000000000000000001100000000000000000000000000
00000000000000000000000000000000000000000000000000000000000000
00000000000000000000000000000000000000000000000000000000000000
00000000000000000000000000000000000000000000011
>DM_validata347
MASQLDSGTPAREASSALRDGYRQAGVSGRVRSYLDLLAGLSDFREHFDGSDGFSLDLVDMA
DGPGEVTVICCAGTAAISGPHEFTRLAGALRGIAPVRAVPQPGYEEGEPLPSSMAAVAAVQA
DAVIRTQGDKPFVVAGHSAGALMAYALATELLDRGHPPRGVVLIDVYPPGHQDAMNAWLEEL
TATLFDRETVRMDDTRLTALGAYDRLTGQWRPRETGLPTLLVSAGEPMGPWPDDSWKPTWPF
EHDTVAVPGDHFTMVQEHADAIARHIDAWLGGGNSSSVDKLAAALE
11111111111111111111111110000000000000000000000000000000000000
00000000000000000000000000000000000000000000000000000000000000
00000000000000000000000000000000000000000000000000000000000000
00000111100000000000000000000000000000000000000000000000000000
0000000000000000000000000000000011111111111111
>DM_validata348
MHAFPLTLDNRLAEALPLWRNLARTDRAPRRNIDLADWKADWRELIAALDRFSRSHGYRQPF
AAQGHAALENAWAWGQAAENASTLLLKAIDRGLAGAELRSIYLETAALWLDYSRLLGAARDS
LREQGEVDFETAPALAPRTGQYPFALQLLAMGVLLDAQELIPALVEEVLQFDTDRLLDYLGA
AALGLTSASEETFHPRPFGQLRAFFEEADGSDAQALAPYLQSQYREFFQLSPKAQKKTRRLT
GPYAWGWWAMEVSALGVLYGWDDGVLRASPHYLGDLVDYARARGDA
11111000000000000000000000000000000000000000000000000000000000
00000000000000000000000000000000000000000000000000000000000000
00000111110000000000000000000000000000000000000000000000000000
00000000000000000000000000011000000000000000000000000000000000
0000000000000000000000000000000000000000000001
>DM_validata349
MSLRRGIYHIENAGVPSAIDLKDGSSSDGTPIVGWQFTPDTINWHQLWLAEPIPNVADTFTL
CNLFSGTYMDLYNGSSEAGTAVNGWQGTAFTTNPHQLWTIKKSSDGTSYKIQNYGSKTFVDL
VNGDSSDGAKIAGWTGTWDEGNPHQKWYFNRMSVSSAEAQAAIARNPHIHGTYRGYILDGEY
LVLPNATFTQIWKDSGLPGSKWREQIYDCDDFAIMKAAVGKWGADSWKANGFAIFCGVMLG
VNKAGDAAHAYNFTLTKDHADIVFFEPQNGGYLNDIGYDSYMAFY
10000000000000000000000000000000000000000000000000000000000000
00000000000000000000000000000000000000000000000000000000000000
00000000000000000000000000000000000000000000000000000000000000
00000000000000000000000000000000000000000000000000000000000000
000000000000000000000000000000000000000000000
>DM_validata350
MVVSIGVFDGVHIGHQKVLRTMKEIAFFRKDDSLIYTISYPPEYFLPDFPGLLMTVESRVEM
LSRYARTVVLDFFRIKDLTPEGFVERYLSGVSAVVVGRDFRFGKNASGNASFLRKKGVEVYE
IEDVVVQGKRVSSSLIRNLVQEGRVEEIPAYLGRYFEIEGIVHKDREFGRKLGFPTANIDRG
NEKLVDLKRGVYLVRVHLPDGKKKFGVMNVGFRPTVGDARNVKYEVYILDFEGDLYGQRLKL
EVLKFMRDEKKFDSIEELKAAIDQDVKSARNMIDDIINSKFEKEG

100000000000000000000000000000000000000000000000000000000000
000000000000000000000000000000000000000000000000000000000000
000000000000000000000000000000000000000011111111111000000000
000000000000000000000000000000001111110000000000000000000000
00000000000111100000000000000000000000011111
>DM_validata351
SIERLGYLGFAVKDVPAWDHFLTKSVGLMAAGSAGDAALYRADQRAWRIAVQPGELDDLAYA
GLEVDDAAALERMADKLRQAGVAFTRGDEALMQQRKVMGLLCLQDPFGLPLEIYYGPAEIFH
EPFLPSAPVSGFVTGDQGIGHFVRCVPDTAKAMAFYTEVLGFVLSDIIDIQMGPETSVPAHF
LHCNGRHHTIALAAFPIPKRIHHFMLQANTIDDVGYAFDRLDAAGRITSLLGRHTNDQTLSF
YADTPSPMIEVEFGWGPRTVDSSWTVARHSRTAMWGHKSVRGQR
000000000000000000000000000000000000000000000000000000000000
000000000000000000000000000000000000000000000000000000000000
000000000000000000000000000000000000000000000000111111100000
000000000000000000000000000000000000000000000000000000000000
00000000000000000000111000000000000000001111
>DM_validata352
GKQQNFVRQTTKYWVHPDNITELKLIILKHLPVLVFNTNKEFEREDSAITSIYFDNENLDLY
YGRLRKDEGAEAHRLRWYGGMSTDTIFVERKTHREDWTGEKSVKARFALKERHVNDFLKGKY
TVDQVFAKMRKEGKKPMNEIENLEALASEIQYVMLKKKLRPVVRSFYNRTAFQLPGDARVRI
SLDTELTMVREDNFDGVDRTHKNWRRTDIGVDWPFKQLDDKDICRFPYAVLEVKLQTQLGQE
PPEWVRELVGSHLVEPVPKFSKFIHGVATLLNDKVDSIPFWLPQ
111110000000000000000000000000000111111100000000000000000000
000000000000000000000000000000000000000000000000000000000000
000000000000000000000000000000000000000000000000000000000000
000000000000000000000000000000000000000000000000000000000000
0000000000000000000000000000000000000011111
>DM_validata353
KIGVLQFVSHPSLDLIYKGIQDGLAEEGYKDDQVKIDFMNSEGDQSKVATMSKQLVANGNDL
VVGIATPAAQGLASATKDLPVIMAAITDPIGANLVKDLKKPGGNVTGVSDHNPAQQQVELIK
ALTPNVKTIGALYSSSEDNSKTQVEEFKAYAEKAGLTVETFAVPSTNEIASTVTVMTSKVDA
IWVPIDNTIASGFPTVVSSNQSSKKPIYPSATAMVEVGGLASVVIDQHDLGVATGKMIVQVL
KGAKPADTPVNVFSTGKSVINKKIAQELGITIPESVLKEAGQVI
000000000000000000000000000011110000000000000000000000000000
000000000000000000000000000000000000000000000000000000000000
000000000000000000000000000000000000000000000000000000000000
000000000000000000000000000000000000000000000000000000000000
0000000000000000000000000000000000000000000
>DM_validata354
MSKKHPIISVTGSSGAGTSTVKHTFDQIFRREGVKAVSIEGDAFHRFNRADMKAELDRRYAA
GDATFSHFSYEANELKELERVFREYGETGQGRTRTYVHDDAEAARTGVAPGNFTDWRDFDSD
SHLLFYEGLHGAVVNSEVNIAGLADLKIGVVPVINLEWIQKIHRDRATRGYTTEAVTDVILR
RMHAYVHCIVPQFSQTDINFQRVPVVDTSNPFIARWIPTADESVVVIRFNRPRGIDFPYLTS
MIHGSWMSRANSIVVPGNKLDLAMQLILTPLIDRVVRESKVA

```
00000000000000111000000000000000000000000000000000000000000000
00000000000000000000000000000000000001111110000000000000000000
00000000000000000000000000000000000000000000000000000000000000
00000000000000000000000000000000000000000000000000000000000000
000000000000000000000000000000000000000000
>DM_validata356
MSLIRMALAAVCALLFSITTMTPFVQASEITISGSTSVARIMDVLAEKYNQQHPETYVAVQG
VGSTAGISLLKKGVADIAMTSRYLTESEAQNTLHTFTLAFDGLAIVVNQANPVTNLTREQLY
GIYKGQITNWKQVGGNDQKIAVVTREASSGTRYSFESLMGLTKTVKDREVSDVAPTALVVNS
NSMMKTLVNHNTQAVGFISIGSVDKSVKAIQFEKADPTSDNIAKHTYQLSRPFLILHYSDNA
DEQTKEFIAFLKSESAKKLIVEYGYIMPSDVEEGGSHHHHHH
11111111111111111111111111110000000000000000000000000000000000
00000000000000000000000000000000000000000000000000000000000000
00000000000000000000000000000000000000000001110000000000000000
00000000000000000000000000000000000000000000000000000000000000
000000000000000000000000000011111111111111
>DM_validata357
MRDLDREETYLVDRTGLALELRDLVGTGPVPGEAYPGPHAALGYGEGQFAALLSGLPDWGEE
GTLFLLEGGYDLGEAAGMALLAETGRARVVRVGFRPGVEVHIPPSPLAPYRYLRFLLLATGR
EEVLRSVDEALLEERRRLGPEVPVEENPAKFLAYTLLERLPLFYSPLFRPLEGAVQTLFARV
AKSLSLTPPPSALEFFLVGLEARHEQGDPLAAVLLGPGEEAALAKEILESRVDALAEVPATG
ANRLAQVMALWYRMAWTAYYLALLYGVDPGDHGLLERLREVT
00000000000000000000000000000000000000000000000000000000000000
00000000000000000001100000000000000000000000000000000000000000
00000000000000000000000000000000000000000000000000000000000000
00000000000000000000000000000000000000000000000000000000000000
000000000000000000000000000000000000111111
>DM_validata358
GMNQAMDTKQIAPFTLALPEGEALPLVCDSPHSGTFYPADFGAVVAPERLRGGEDTHVDALW
EAVPRVGGTLLAATFPRVYIDPNRMLDDIDPAQLEGPWPTPLAPGEKTRLGYGLIWSNVDAA
TPIYDRKLTVAEVQRRINRYYRPYHAALTEAVEGAYQRFGAVWHLNLHSMPNNAYERLKIQS
PRPLADFVLGDRDGTTCEPGLVDLVERELREKGYTVARNDPYKGVQLIAQIGRPAERRNSLQ
IEIRRPLYMEEGTRERNEGFATLQRDLTLLTLRIAEYVRRGV
11111111110000000000000000000000000000000000000000000000000000
00000000000000000000000000000000000000000000011011110000000001
10000000000000000000000000000000000000000000000000000000000000
00000000000000000000000000000000000000000000100000000000000000
000000000000000000000000000000000000000000
>DM_validata360
MAMRQCAIYGKGGIGKSTTTQNLVAALAEMGKKVMIVGCDPKADSTRLILHSKAQNTIMEMA
AEAGTVEDLELEDVLKAGYGGVKCVESGGPEPGVGCAGRGVITAINFLEEEGAYEDDLDFVF
YDVGDVVCGGFAMPIRENKAQEIYIVCSGEMMAMYAANNISKGIVKYANSGSVRLGGLICNS
RNTDREDELIIALANKLGTQMIHFVPRDNVVQRAEIRRMTVIEYDPKAKQADEYRALARKVV
DNKLLVIPNPITMDELEELLMEFGIMEVEDESIVGKTAEEV
```

00000000000000000000000000000000000000000000000001111100000000
00000000000000000000000000000000000000000000000000000011100000
00000000000000000000000000000000000000000000000000000000000000
11110000000000000000000000000000000000000000000000000000000000
00000000000000000000000111111111111111111
>DM_validata361
MVDIIIAEHAGFCFGVKRAVKLAEESLKESQGKVYTLGPIIHNPQEVNRLKNLGVFPSQGEE
FKEGDTVIIRSHGIPPEKEEALRKKGLKVIDATCPYVKAVHEAVCQLTREGYFVVLVGEKNH
PEVIGTLGYLRACNGKGIVVETLEDIGEALKHERVGIVAQTTQNEEFFKEVVGEIALWVKEV
KVINTICNATSLRQESVKKLAPEVDVMIIIGGKNSGNTRRLYYISKELNPNTYHIETAEELQ
PEWFRGVKRVGISAGASTPDWIIEQVKSRIQEICEGQLVSS
00000000000000000000000000000000000000000000000000000000000000
00000000000000000000000000000000000000000000000000000000000000
00000000000000000000000000000000000000000000000000000000000000
00000000000000000000000000000000000000000000000000000000000000
00000000000000000000000000000000011111111
>DM_validata363
KEFTLDFSTAKTYVDSLNVIRSAIGTPLQTISSGGTSLLMIDSGTGDNLFAVDVRGIDPEEG
RFNNLRLIVERNNLYVTGFVNRTNNVFYRFADFSHVTFPGTTAVTLSGDSSYTTLQRVAGIS
RTGMQINRHSLTTSYLDLMSHSGTSLTQSVARAMLRFVTVTAEALRFRQIQRGFRTTLDDLS
GRSYVMTAEDVDLTLNWGRLSSVLPDYHGQDSVRVGRISFGSINAILGSVALILNCHHHASR
VARMASDEFPSMCPADGRVRGITHNKILWDSSTLGAILM
00000000000000000000000000000000000000000111100000000000000000
00000000000000000000000000000000000000000000000000000000000000
00000000000000000000000000000000000000000000000000000000001111
11000000000000000000000000000000000000000000000000000000111111
111111110000000000000000000000000000000
>DM_validata364
MANIVNFTDKQFENRLNDNLEELIQGKKAVESPTAFLLGGQPGSGKTSLRSAIFEETQGNVI
VIDNDTFKQQHPNFDELVKLYEKDVVKHVTPYSNRMTEAIISRLSDQGYNLVIEGTGRTTDV
PIQTATMLQAKGYETKMYVMAVPKINSYLGTIERYETMYADDPMTARATPKQAHDIVVKNLP
TNLETLHKTGLFSDIRLYNREGVKLYSSLETPSISPKETLEKELNRKVSGKEIQPTLERIEQ
KMVLNKHQETPEFKAIQQKLESLQPPTPPIPKTPKLPGI
00000000000000000000000000000000000000000000000000000000000000
00000000000000000000000000000000000000000000000000000000000000
00000000000000000000000000000000000000000000000000000000000000
00000000000000000000000000000000000000000000000000000000000000
000000000000000000000001111111111111111
>DM_validata365
ASMTGGQQMGRGSASLPACPEESPLLVGPMLIEFNMPVDLELVAKQNPNVKMGGRYAPRDCV
SPHKVAIIIPFRNRQEHLKYWLYYLHPVLQRQQLDYGIYVINQAGDTIFNRAKLLNVGFQEA
LKDYDYTCFVFSDVDLIPMNDHNAYRCFSQPRHISVAMDKFGFSLPYVQYFGGVSALSKQQF
LTINGFPNNYWGWGGEDDDIFNRLVFRGMSISRPNAVVGTTRHIRHSRDKKNEPNPQRFDRI
AHTKETMLSDGLNSLTYQVLDVQRYPLYTQITVDIGTPS

111111111100000000000000000000000000000000000000000000000000
000000000000000000000000000000000000000000000000000000000000
000000000000000000000000000000000000000000000000000000000000
000000000000000000000000000000000000000000011111111100000000
00000000000000000000000000000000000000
>DM_validata366
MTMMDMNFKYCHKIMKKHSKSFSYAFDLLPEDQRKAVWAIYAVCRKIDDSIDVYGDIQFLNQ
IKEDIQSIEKYPYEYHHFQSDRRIMMALQHVAQHKNIAFQSFYNLIDTVYKDQHFTMFETDA
ELFGYCYGVAGTVGEVLTPILSDHETHQTYDVARRLGESLQLINILRDVGEDFENERIYFSK
QRLKQYEVDIAEVYQNGVNNHYIDLWEYYAAIAEKDFRDVMDQIKVFSIEAQPIIELAARIY
IEILDEVRQANYTLHERVFVEKRKKAKLFHEINSKYHRI
000000000000000000000000000000000000000000000000000000000000
000000000000000000000000000000000000000000000000000000000000
000000000000000000000000000000000000000000000000000000000000
000000000000000000000000000000000000000000000000000000000000
00000000000000000000000000000000000111
>DM_validata367
MNRLKNMSKYSDAKELASLTLGKKTEYANQYDPSLLQPVPRSLNRNDLHLSATLPFQGCDIW
TLYELSWLNQKGLPQVAIGEVSIPATSANLIESKSFKLYLNSYNQTRFASWDEVQTRLVHDL
SACAGETVTVNVKSLNEYTAEPIVTMQGECIDDQDIEIANYEFDDALLQGAAQGEEVSEVLH
SHLLKSNCLITNQPDWGSVEIAYHGAKMNREALLRYLVSFREHNEFHEQCVERIFTDIMRYC
QPQSLTVYARYTRRGGLDINPFRSSHQSAPNHNQRMARQ
111111111111111111111111111000000000000000000000000000000000
000000000000000000000000000000000000000000000000000000000000
000000000000000000000000000000000000000000000000000000000000
000000011111000000000000000000000000000000000000000000000000
00000000000000000000000000000000000000
>DM_validata368
MIAITGATGQLGHYVIESLMKTVPASQIVAIVRNPAKAQALAAQGITVRQADYGDEAALTSA
LQGVEKLLLISSSEVGQRAPQHRNVINAAKAAGVKFIAYTSLLHADTSPLGLADEHIETEKM
LADSGIVYTLLRNGWYSENYLASAPAALEHGVFIGAAGDGKIASATRADYAAAAARVISEAG
HEGKVYELAGDSAWTLTQLAAELTKQSGKQVTYQNLSEADFAAALKSVGLPDGLADMLADSD
VGASKGGLFDDSKTLSKLIGHPTTTLAESVSHLFNVNN
000000000000000000000000000000000000000000000000000000000000
000000000001111111100000000000000000000000000000000000000000
000000000000000000000000000000000000000000000000000000000000
000000000000000000000000000000000000000000000000000000000000
0000000000000000000000000000000000111
>DM_validata369
GMVKDKNDVGPKTVAILGAGGKMGARITRKIHDSAHHLAAIEIAPEGRDRLQGMGIPLTDGD
GWIDEADVVVLALPDNIIEKVAEDIVPRVRPGTIVLILDAAAPYAGVMPERADITYFIGHPC
HPPLFNDETDPAARTDYHGGIAKQAIVCALMQGPEEHYAIGADICETMWSPVTRTHRVTTEQ
LAILEPGLSEMVAMPFVETMVHAVDECADRYGIDRQAALDFMIGHLNVEIAMWFGYSPKVPS
DAALRLMEFAKDIVVKEDWREALNPAKVKQAAELIAGK

111111110000000000000000000000000000000000000000000000000000
000000000000000000000000000000000000000000000000000000000000
000000000000000000000000000000000000000000000000000000000000
000000000000000000000000000000000000000000000000000000000011
10000000000000000000000000000000000001
>DM_validata370
MKYIGAHVSAAGGLANAAIRAAEIDATAFALFTKNQRQWRAAPLTTQTIDEFKAACEKYHYT
SAQILPHDSYLINLGHPVTEALEKSRDAFIDEMQRCEQLGLSLLNFHPGSHLMQISEEDCLA
RIAESINIALDKTQGVTAVIENTAGQGSNLGFKFEHLAAIIDGVEDKSRVGVCIDTCHAFAA
GYDLRTPAECEKTFADFARTVGFKYLRGMHLNDAKSTFGSRVDRHHSLGEGNIGHDAFRWIM
QDDRFDGIPLILETINPDIWAEEIAWLKAQQTEKAVA
000000000000000000000000000000000000000000000000000000000000
000000000000000000000000000000000000000000000000000000000000
000000000000000000000000000000000000000000000000000000000000
000000000000000000000000000000000000000000000000000000000000
0000000000000000000000000000000111111
>DM_validata371
DNNIENIGDGAEVVKRTEDTSSDKWGVTQNIQFDFVKDKKYNKDALILKMQGFINSKTTYYN
YKNTDHIKAMRWPFQYNIGLKTNDPNVDLINYLPKNKIDSVNVSQTLGYNIGGNFNSGPSTG
GNGSFNYSKTISYNQQNYISEVEHQNSKSVQWGIKANSFITSLGKMSGHDPNLFVGYKPYSQ
NPRDYFVPDNELPPLVHSGFNPSFIATVSHEKGSGDTSEFEITYGRNMDVTHATRRTTHYGN
SYLEGSRIHNAFVNRNYTVKYEVNWKTHEIKVKGHN
110000011000000000000000000000000000000000000000000000000000
000000000000000000000000000000000000000000000000000000011111
111000000000000000000000000000000000000111000000000000000000
000000000000000000000000000000000000000000000000000000011110
000000000000000000000000000000000000
>DM_validata373
MTVAKAIFIKCGNLGTSMMMDMLLDERADREDVEFRVVGTSVKMDPECVEAAVEMALDIAED
FEPDFIVYGGPNPAAPGPSKAREMLADSEYPAVIIGDAPGLKVKDEMEEQGLGYILVKPDAM
LGARREFLDPVEMAIYNADLMKVLAATGVFRVVQEAFDELIEKAKEDEISENDLPKLVIDRN
TLLEREEFENPYAMVKAMAALEIAENVADVSVEGCFVEQDKERYVPIVASAHEMMRKAAELA
DEARELEKSNDAVLRTPHAPDGKVLSKRKFMEDPE
100000000000000000000000000000000000000000000000000000000000
000000000000000000000000000000000000000000000000000000000000
000000000000000000000000000000000000000000000000000000000000
000000000000000000000000000000000000000000000000000000000000
00000000000000000000000000000000000
>DM_validata374
SLLSYQHGYHAGNFADVIKHITLTRLLAYLTHKDKPLFYLETHSGRGIYDLKDKQSLKTEEY
KEGINPVWLDRENLPSLFLEYISVIKQINLNSTLSYYPGSPYFAINQLRSQDRLYLCELHPT
EYNFLLKLPHFNKKVYVNHTDGVSKLNALLPPPEKRGLIFIDPSYERKEEYKEIPYAIKNAY
SKFSTGLYCVWYPVVNKAWTEQFLRKMREISSKSVRIELHLNPLINEGMTGCGLWIINPPYT
FPSEIKLVLETLTTYFNPGSSSYMIESGSKLCHEL

11111111100000000000000000000000000000000000000000000011110000
00000000000000000000000000000000000000000000000000000000000000
00000000000000000000000000000000000000000000000000000000000000
00000000000000000000000000000000000000000000000000000000000000
00000000000000000000000000000000111
>DM_validata375
MASNYNSGLKIGAWVGTQPSESAIKSFQELQGRKLDIVHQFINWSTDFSWVRPYADAVYNNG
SILMITWEPWEYNTVDIKNGKADAYITRMAQDMKAYGKEIWLRPLHEANGDWYPWAIGYSSR
VNTNETYIAAFRHIVDIFRANGATNVKWVFNVNCDNVGNGTSYLGHYPGDNYVDYTSIDGYN
WGTTQSWGSQWQSFDQVFSRAYQALASINKPIIIAEFASAEIGGNKARWITEAYNSIRTSYN
KVIAAVWFHENKETDWRINSSPEALAAYREAIGAG
11111110000000000000000000000000000000000000000000000000000000
00000000000000000000000000000000000000000000000000000000000000
00000000000000000000000000000000000000000000000000000000000000
00000000000000000000000000000000000000000000000000000000000000
00000000000000000000000000000000111
>DM_validata376
MLNEKAIQDVDLYFQNIHGPEGRLASNETFDIVPGLSKDGAVQYQTYQFNEAPKHLQKQVKA
GRILMERFVAVASAAVNKKAPSNKEKYHYDIWKEVSNQLIPAFFTDPIKGEQNLNTTVKGVE
VAKSVIQFAGNVIAGNVTGFATFLQNFGNGLSAEMNKTQANYNYLYAYSTHDLFQDTSGNVF
YKPRFLIYGTHFKQEQKKIATSCASYQEVNLEFGVDTVGGTFRIEEYFSNETFKKKVDNFLD
KYEGKAIDDADSYFDDIFNGVKPNKNYVYHHHHHH
11111111100000000001111111000000000000000000000000000000000000
00000000000000000000000000000000000000000000000000000000000000
00000000000000000000000000000000000000000000000000000000000000
00000000000000000000000000000000000000000000000000000000000000
00000000000000000000000000000111111
>DM_validata377
MGSDKIHHHHHHMAVNRFRLENGKEELALYQIQLLKDQSHNENEEDKVSSSSFRQRILGNLL
QPPNERPELPSGLYVLGLTGISGSGKSSVAQRLKNLGAYIIDSDHLGHRAYAPGGPAYQPVV
EAFGTDILHKDGTINRKVLGSRVFGNKKQMKILTDIVWPVIAKLAREEMDVAVAKGKTLCVI
DAAMLLEAGWQSMVHEVWTVVIPETEAVRRIVERDGLSEAAAQSRLQSQMSGQQLVEQSNVV
LSTLWESHVTQSQVEKAWNLLQKRLPKAYQTRN
11111111111111111111111111100000000001111111111111111100000000
00000000000000000000000000000000000000000000000000000000000000
00000000000000000000000000000000000000000000000000000000000000
00000000000000000000000000000000000000000000000000000000000000
000000000000000000000000001111111
>DM_validata378
MQNVLIVGVGFMGGSFAKSLRRSGFKGKIYGYDINPESISKAVDLGIIDEGTTSIAKVEDFS
PDFVMLSSPVRTFREIAKKLSYILSEDATVTDQGSVKGKLVYDLENILGKRFVGGHPIAGTE
KSGVEYSLDNLYEGKKVILTPTKKTDKKRLKLVKRVWEDVGGVVEYMSPELHDYVFGVVSHL
PHAVAFALVDTLIHMSTPEVDLFKYPGGGFKDFTRIAKSDPIMWRDIFLENKENVMKAIEGF
EKSLNHLKELIVREAEEELVEYLKEVKIKRMEI

00000000000000000000000000000000000000000000000000000000000000
00000000000000000000000000000000000000000000000000000000000000
00000000000000000000000000000000000000000000000000000000000000
00000000000000000000000000000000011100000000000000000000000000
000000000000000000000000000000000
>DM_validata379
MSATGPFSIGERVQLTDAKGRRYTMSLTPGAEFHTHRGSIAHDAVIGLEQGSVVKSSNGALF
LVLRPLLVDYVMSMPRGPQVIYPKDAAQIVHEGDIFPGARVLEAGAGSGALTLSLLRAVGPA
GQVISYEQRADHAEHARRNVSGCYGQPPDNWRLVVSDLADSELPDGSVDRAVLDMLAPWEVL
DAVSRLLVAGGVLMVYVATVTQLSRIVEALRAKQCWTEPRAWETLQRGWNVVGLAVRPQHSM
RGHTAFLVATRRLAPGAVAPAPLGRKREGRDG
11100000000000000000000000000000000000000000000000000000000000
00000000000000000000000000000000000000000000000000000000000000
00000000000000000000000000000000000000000000000000000000000000
00000000000000000000000000000000000000000000000000000000000000
00000000000000000001111111111111
>DM_validata380
MNIIGFSKALFSTWIYYSPERILFDAGEGVSTTLGSKVYAFKYVFLTHGHVDHIAGLWGVVN
IRNNGMGDREKPLDVFYPEGNRAVEEYTEFIKRANPDLRFSFNVHPLKEGERVFLRNAGGFK
RYVQPFRTKHVSSEVSFGYHIFEVRRKLKKEFQGLDSKEISRLVKEKGRDFVTEEYHKKVLT
ISGDSLALDPEEIRGTELLIHECTFLDARDRRYKNHAAIDEVMESVKAAGVKKVILYHISTR
YIRQLKSVIKKYREEMPDVEILYMDPRKVFEM
00000000000000000000000000000000000000000000000000000000000000
00001111100000000000000000000000000000000000000000000000000000
00000000111111000000000000000000000000000000000000000000000000
00000000000000000000000000000000000000000000000000000000000000
00000000000000000000000000000000
>DM_validata381
GAMDRLRAAEHPRPDYVLLHISDTHLIGGDRRLYGAVDADDRLGELLEQLNQSGLRPDAIVF
TGDLADKGEPAAYRKLRGLVEPFAAQLGAELVWVMGNHDDRAELRKFLLDEAPSMAPLDRVC
MIDGLRIIVLDTSVPGHHHGEIRASQLGWLAEELATPAPDGTILALHHPPIPSVLDMAVTVE
LRDQAALGRVLRGTDVRAILAGHLHYSTNATFVGIPVSVASATCYTQDLTVAAGGTRGRDGA
QGCNLVHVYPDTVVHSVIPLGGGETVGTFVSP
11111111111000000000000000011111111110000000000000000000000000
00000000000000000000000000000000000000000000000000000000000000
00000000000000000000000000000000000000000000000000000000000000
00000000000000000000000000000000000000000000111111111111111111
10000000000000000001111111111111
>DM_validata382
GETSVPPGSALVGPSCVMDDFRDPQRWKECAKQGKMPCYFDLIEENVYLTERKKNKSHRDIK
RMQCECTPLSKDERAQGEIACGEDCLNRLLMIECSSRCPNGDYCSNRRFQRKQHADVEVILT
EKKGWGLRAAKDLPSNTFVLEYCGEVLDHKEFKARVKEYARNKNIHYYFMALKNDEIIDATQ
KGNCSRFMNHSCEPNCETQKWTVNGQLRVGFFTTKLVPSGSELTFDYQFQRYGKEAQKCFCG
SANCRGYLGGENRVSIRAAGGKMKKERSRK

1111111111111000000000000000000000000000000000000011111111111
110000000000000000000000000000000000000000000000000000000000
000000000000000000000000000000000000000000000000000000000000
000000000000000000000000000000000000000000000000000000000000
000000000011111111111111111111
>DM_validata383
MEACPHIRYAFQNDKLLLQQASVGRLTLVNKTTILLRPMKTTTVDLGLYARPPEGHGLMLWG
STSRPVTSHVGIIDPGYTGELRLILQNQRRYNSTLRPSELKIHLAAFRYATPQMEEDKGPIN
HPQYPGDVGLDVSLPKDLALFPHQTVSVTLTVPPPSIPHHRPTIFGRSGLAMQGILVKPCRW
RRGGVDVSLTNFSDQTVFLNKYRRFCQLVYLHKHHLTSFYSPHSDAGVLGPRSLFRWASCTF
EEVPSLAMGDSGLSEALEGRQGRGFGSSGQ
111000000000000000000000000000000000000000000000000000000000
000000000000000000000000000000000000000000000000000011110000
000000000000000000000000000000000000000000000000000000000000
000000000000000000000000000000000000000000000000000000000000
000000001111111111111111111111
>DM_validata384
PNIADKGSVFYHFSATSFDSVDGTRHYRVWTAVPNTTAPASGYPILYMLDGNAVMDRLDDEL
LKQLSEKTPPVIVAVGYQTNLPFDLNSRAYDYTPAAESRKTDLHSGRFSRKSGGSNNFRQLL
ETRIAPKVEQGLNIDRQRRGLWGHSYGGLFVLDSWLSSSYFRSYYSASPSLGRGYDALLSRV
TAVEPLQFCTKHLAIMEGSATQGDNRETHAVGVLSKIHTTLTILKDKGVNAVFWDFPNLGHG
PMFNASFRQALLDISGENANYTAGCHELSH
000000000000000000000000000000000000000000000000000000000000
000000000000000000000000000000000000000000011111000000000000
000000000000000000000000000000000000000000000000000000000000
000000000000000000001111111111100000000000000000000000000000
000000000000000001111111111111
>DM_validata385
GSTKVKYVDKIHIGNYEIDAWYFSPFPEDYGKQPKLWLCEYCLKYMKYEKSYRFHLGQCQWR
QPPGKEIYRKSNISVHEVDGKDHKIYCQNLCLLAKLFLDHRTLYFDVEPFVFYILTEVDRQG
AHIVGYFSKEKESPDGNNVACILTLPPYQRRGYGKFLIAFSYELSKLESTVGSPEKPLSDLG
KLSYRSYWSWVLLENLRDFRGTLSIKDLSQMTSITQNDIISTLQSLNMVKYWKGQHVICVTP
KLVEEHLKSAQYKKPPITVDSVCLKWAPPK
111110000000000000000000000000000000000000000000000000000000
000000000000000000000000000000000000000000000000000000000000
000000000000000000000000000000000000000000000000000000000000
000000000000000001111110000000000000000000000000011110000000
000000000000000000000000000011
>DM_validata386
MIGKKSFQTNVYRMSKFDTYIFNNLYINDYKMFWIDSGIAKLIDKNCLVSYEINSSSIILLK
KNSIQRFSLTSLSDENINVSVITISDSFIRSLKSYILGDLMIRNLYSENKDLLLWNCEHNDI
AVLSEVVNGFREINYSDEFLKVFFSGFFSKVEKKYNSIFITDDLDAMEKISCLVKSDITRNW
RWADICGELRTNRMILKKELESRGVKFRELINSIRISYSISLMKTGEFKIKQIAYQSGFASV
SYFSTVFKSTMNVAPSEYLFMLTGVAEK

11100000000000000000000000000000000000000000000000000000000000
00000000000000000000000000000000000000111111111100000000000000
00000000000000000000000000000000000000000000000000000000000000
00000000000000000000000000000000000000000000000000000000000000
0000000000000000000000000111
>DM_validata387
KGAELGILIKNADALEVAEKVTAVIFDKTGTLTKGKPEVTDLVPLNGDERELLRLAAIAERR
SEHPIAEAIVKKALEHGIELGEPEKVEVIAGEGVVADGILVGNKRLMEDFGVAVSNEVELAL
EKLEREAKTAVIVARNGRVEGIIAVSDTLKESAKPAVQELKRMGIKVGMITGDNWRSAEAIS
RELNLDLVIAEVLPHQKSEEVKKLQAKEVVAFVGDGINDAPALAQADLGIAVGSGSDVAVES
GDIVLIRDDLRDVVAAIQLSRKTMSKIK
11000000000000000000000000000000000000000000000000000000000000
00000000000000000000000000000000000000000000000000000000000000
00000000000000000000000000000000000000000000000000000000000000
00000000000000000000000000000000000000000000000000000000000000
0000000000000000000000111111
>DM_validata388
ARIRAPQEATVKEVHDAPAVRGSIIANMLQEHDNPFTLYPYDTNYLIYTQTSDLNKEAIASY
DWAENARKDEVKFQLSLAFPLWRGILGPNSVLGASYTQKSWWQLSNSEESSPFRETNYEPQL
FLGFATDYRFAGWTLRDVEMGYNHDSNGRSDPTSRSWNRLYTRLMAENGNWLVEVKPWYVVG
NTDDNPDITKYMGYYQLKIGYHLGDAVLSAKGQYNWNTGYGGAELGLSYPITKHVRLYTQVY
SGYGESLIDYNFNQTRVGVGVMLNDLF
11111111111111111100000000000000000000000000000000000000000000
00000000000000000000000000000000000000000000000000000000000000
00000000000000000000000000000000000000000000000000000000000000
00000000000000000000000000000000000000000000000000000000000000
000000000000000000000000000
>DM_validata389
MSLITVNTLQKMKAAGEKIAMLTAYESSFAALMDDAGVEMLLVGDSLGMAVQGRKSTLPVSL
RDMCYHTECVARGAKNAMIVSDLPFGAYQQSKEQAFAAAAELMAAGAHMVKLEGGVWMAETT
EFLQMRGIPVCAHIGLTPQSVFAFGGYKVQGRGGKAQALLNDAKAHDDAGAAVVLMECVLAE
LAKKVTETVSCPTIGIGAGADCDGQVLVMHDMLGIFPGKTAKFVKNFMQGHDSVQAAVRAYV
AEVKAKTFPAAEHIFADEGGSHHHHHH
10000000000000000000000000000000000000000000000000000000000000
00000000000000100000000000000000000000000000000000000000000000
00000000000000011111111111111111111000000000000000000000000000
00000000000000000000000000000000000000000000000000000000000000
000000000001111111111111111
>DM_validata390
SNTSWRKSEVLAVPLQPTLQQEVILARMEQILASRALTDDERAQLLYERGVLYDSLGLRALA
RNDFSQALAIRPDMPEVFNYLGIYLTQAGNFDAAYEAFDSVLELDPTYNYAHLNRGIALYYG
GRDKLAQDDLLAFYQDDPNDPFRSLWLYLAEQKLDEKQAKEVLKQHFEKSDKEQWGWNIVEF
YLGNISEQTLMERLKADATDNTSLAEHLSETNFYLGKYYLSLGDLDSATALFKLAVANNVHN
FVEHRYALLELSLLGQDQDDLAESDQQ

111111000000000000000000000000000000000000000000000000000000
000000000000000000000000000000000000000000000000000000000000
000000000000000000000000000000000000000000000000000000000000
000000000000000000000000000000000000000000000000000000000000
000000000000000001111111111
>DM_validata391
MAQKPVADALTLELEPVVEANMTRHLDTEDIWFAHDYVPFDQGENFAFLGGRDWDPSQSTLP
RTITDACEILLILKDNLAGHHRELVEHFILEDWWGRWLGRWTAEEHLHAIALREYLVVTREV
DPVANEDVRVQHVMKGYRAEKYTQVETLVYMAFYERCGAVFCRNLAAQIEEPILAGLIDRIA
RDEVRHEEFFANLVTHCLDYTRDETIAAIAARAADLDVLGADIEAYRDKLQNVADAGIFGKP
QLRQLISDRITAWGLAGEPSLKQFVTG
111111100000000000000000000000000000000000000000000000000000
000000000000000011111111111111100000000000000000000000000000
000000000011111111110000000000000000000000000000000000000000
000000000000000000000000000000000000000000000000000000000000
000000000000000000000000001
>DM_validata392
GMTSRSPFESFVWQSEIFNCQSNDIDAFYAQLAEEVNRLGLKKNTLGSVDSFAINLYQSASQ
RSDLPSLLISSGFHGEEAAGPWGMLHFLRGLQPALFERVNLSLLPLVNPTGFKAGHRFNRFG
ENPNRGFTLENGKPTPNEHTSLEGKLLLEHAQLLCAASRDGILTCHEDVLMNETYVYSFEPT
QTPGRFSLGLRDALGQYFKLAKDGFIDECPVTDGVIFNHFDTSFEAFLVRSGAKLAACSETP
GQEDFDRRVQANSAAMGQFIAHCAPIL
100000000000000000000000000000000000000000000000000000000000
000000000000000000000000000000000000000000000000000000000000
000000000111111111000000000000000000000000000000000000000000
000000000000000000000011000000000000000000000000000000000000
000000000000000000000000001
>DM_validata393
GMTRRIAICAPSTPFTREDSARVIALAAAEFPDLSLSFHEQCFASEGHFAGSDALRLSAFLE
CANDDAFEAVWFVRGGYGANRIAEDALARLGRAASAKQYLGYSDAGTLLAALYAHRIGRSVH
APMPVDIRRPEGESAVRRTLGWLAGAREGLEPTLGAGAPAVAFNLMTLAMLCGTRLLPDLSG
HVVMIEEVAEHHYAVDRLLFHVTSCLADAGIAGLRLGRVSDVPENDRPFGCSVEEMARHWCH
RAGIAFLGTADIGHDVDNRIVPFGLA
000000000000000000000000000000000000000000000000000000000000
000000000000000000000000000000000000000000000000000000000000
000000000000000000000000000000000001100000000000000000000000
000000000000000000000000000000000000000000000000000000000000
00000000000000000000000011
>DM_validata395
MSEIVKFNPVMASGFGAYIDHRDFLEAKTETIKNLLMRQGFVVVKNLDIDSDTFRDIYSAYG
TIVEYADEKIGVGFGYRDTLKLEGEKGKIVTGRGQLPFHADGGLLLSQVDQVFLYAAEIKNV
KFRGATTVCDHALACQEMPAHLLRVLEEETFEVRVLERGYYVDVSPDGWFKVPVFTDLGWVR
KMLIYFPFDEGQPASWEPRIVGFTDHETQAFFQELGAFLKQPRYYYKHFWEDGDLLIMDNRR
VIHEREEFNDDDIVRRLYRGQTADI

000000000000000000000000000000000000000000000000000000000000
000011111111111100000000000000000000000000000000000000000000
000000000000000000000000000000000000111111111110000000000000
000000000000000000000000000000000000000000000000000000000000
0000000000000000000000000
>DM_validata398
ERLTLDTRLRQALERNELVLHYQPIVELASGRIVGGEALVRWEDPERGLVMPSAFIPAAEDT
GLIVALSDWVLEACCTQLRAWQQQGRAADDLTLSVNISTRQFEGEHLTRAVDRALARSGLRP
DCLELEITENVMLVMTDEVRTCLDALRARGVRLALDDFGTGYSSLSYLSQLPFHGLKIDQSF
VRKIPAHPSETQIVTTILALARGLGMEVVAEGIETAQQYAFLRDRGCEFGQGNLMSTPQAAD
AFASLLDRQKASGQRPVHGHETAP
110000000000000000000000000000000000000000001111111111111100
000000000000000000000000000000000000000000000000000000000000
000000000000000000000000000000000000000000000000000000000000
000000000000000000000000000000000000000000000000000000000000
000000000000111111111111
>DM_validata399
GTQPTFGFTVNWKFSESTTVFTGQCFIDRNGKEVLKTMWLLRSSVNDIGDDWKATRVGINIF
TRLRTQKEGGSGGSARKCSLTGKWTNDLGSNMTIGAVNSRGEFTGTYITAVTATSNEIKESP
LHGTQNTINKSGGSTTVFTGQCFIDRNGKEVLKTMWLLRSSVNDIGDDWKATRVGINIFTRL
RTQKEGGSGGSARKCSLTGKWTNDLGSNMTIGAVNSRGEFTGTYITAVTATSNEIKESPLHG
TQNTINKRTQPTFGFTVNWKFSE
100000000000000000000000000000000000000000000000000000000000
000111111111111100000000000000000000000000000000000000000000
000000001111000000000000000000000000000000000000000000000000
111111111100000000000000000000000000000000000000000000000000
00000000000000000000001
>DM_validata400
QLQQQEDDRILGLPGQPNGVAFGMYGGYVTIDDNNGRALYYWFQEADTADPAAAPLVLWLNG
GPGCSSIGLGAMQELGAFRVHTNGESLLLNEYAWNKAANILFAESPAGVGFSYSNTSSDLSM
GDDKMAQDTYTFLVKWFERFPHYNYREFYIAGESGHFIPQLSQVVYRNRNNSPFINFQGLLV
SSGLTNDHEDMIGMFESWWHHGLISDETRDSGLKVCPGTSFMHPTPECTEVWNKALAEQGNI
NPYTIYTPTCDREPSPYQRRFW
111000000000000000000000000000000000000000000000000000000000
000000000000000000000000000000000000000000000000000000000000
000000000000000000000000000000000000000000000000000000000000
000000000000000000000000000000000000000000000000000000000000
0000000000000000000000
>DM_validata401
VLLKEYRVILPVSVDEYQVGQLYSVAEASKNETGGGEGVEVLVNEPYEKDDGEKGQYTHKIY
HLQSKVPTFVRMLAPEGALNIHEKAWNAYPYCRTVITNEYMKEDFLIKIETWHKPDLGTQEN
VHKLEPEAWKHVEAIYIDIADRSQVLSKDYKAEEDPAKFKSVKTGRGPLGPNWKQELVNQKD
CPYMCAYKLVTVKFKWWGLQNKVENFIHKQEKRLFTNFHRQLFCWLDKWVDLTMDDIRRMEE
ETKRQLDEMRQKDPVKGMTADD

000000000000000000000000000000000000000000000000000000000000
000000000000000000000000000000000000000000000000000000000000
000000000000000000000000000000000000000000000000000000000000
000000000000000000000000000000000000000000000000000000000000
0000000011111111111111
>DM_validata403
MKEYLTRSNISYFSIGSGTPIIFLHGLSLDKQSTCLFFEPLSNVGQYQRIYLDLPGMGNSDP
ISPSTSDNVLETLIEAIEEIIGARRFILYGHSYGGYLAQAIAFHLKDQTLGVFLTCPVITAD
HSKRLTGKHINILEEDINPVENKEYFADFLSMNVIINNQAWHDYQNLIIPGLQKEDKTFIDQ
LQNNYSFTFEEKLKNINYQFPFKIMVGRNDQVVGYQEQLKLINHNENGEIVLLNRTGHNLMI
DQREAVGFHFDLFLDELNSNN
110000000000000000000000000000000000000000000000000000000000
000000000000000000000000000000000000000000000000000000000000
000000000000000000000000000000000000000000000000000000000000
000000000000000000000000000000000000000000000000000000000000
000000000000000000111
>DM_validata404
MGHHHHHHGGHMSLLTFVGLGLWDVKDISVKGLEAVREADEVYVEYYTSKLLSSIEEMEEFF
GKRVVELERSDLEENSFRLIERAKSKSVVLLVPGDPMVATTHSAIKLEAERKGVKTRIIHGA
SISTAVCGLTGLHNYRFGKSATVSWHRSQTPVNVIKANRSIDAHTLLFLDLHPEPMTIGHAV
ENLIAEDAQMKDLYAVGIARAGSGEEVVKCDRLENLKKIDFGKPLHVMVVLAKTLHFMEFEC
LREFADAPAELERLVAEGGS
111111111111000000000000000000000000000000000000000000000000
000000000000000000000000000000000000000000000000000000000000
000000000000000000000000000000000000000000000000000000000000
000000000000000000000000000000000000000000000000000000000000
00000000000000011111
>DM_validata406
GSHMAVLQKSSPLQENKDFYILDTHTQKKISFEDMILELLKADVILLGEKHDEVKHKISQVM
IFNALEGNLSSQNINFDVALEMLASTEQNHLDKAFKNKKTIKANELTNALNWDKVWKWKDYE
QFVNVVFYSKSKILGANLSRSEITSIYNGAQPLKGYVSTTNEVKKQLFDIISLSHKLNPEEN
KELLDKLVEIQQFKDRRMADVLVHHVNKVLLLAGSYHTSKKIGIPLHIQDFKSSKKIVVVNL
SYGEIDLKDSDYVLIYKGEE
111111111110000000000000000000000000000000000000000000000000
000000000000000000000000000000000000000000000000000000000000
000000000000000000000000000000000000000000000000000000000000
000000000000000000000000000000000000000000000000000000000000
00000000000000000011
>DM_validata407
WKANLLVPVEDPRELMGTFDFLRDITYPKGSVKLLGLAGNTDKENLLSQLPSISEGFQEEGV
FSSWTIIDTAEFEENLVVGMEALTGSFFRPSILFLRLPENRDRDEEIREIIRKASMYRMGVL
LFSKHPQAGLGRQNLINLWIENRGLDWDISMELGNMDLALLIAYKLKSNWKASLSFMTFAPT
AIQAQAAENFLQSLAELARIPNVKMQVLRENPIKSSKLPFASLHIFSLDPNPDLDLARHLME
KAGSSCIFALDSGEENALA

00000000000000000000000000000000000000111000000000000000000000
00000001111100000000000000000000000000000000000000000000000000
00000000000000000000000111000000000000000000000000000000000000
00000000000000000000000000000000000000000000000000000000000000
0000000000000000000
>DM_validata408
MLKQKELIANVKNLTESDERITACMMYGSFTKGEGDQYSDIEFYIFLKHSITSNFDSSNWLF
DVAPYLMLYKNEYGTEVVIFDNLIRGEFHFLSEKDMNIIPSFKDSGYIPDTKAMLIYDETGQ
LENYLSEISGARPNRLTEENANFLLCNFSNLWLMGINVLKRGEYARSLELLSQLQKNTLQLI
RMAEKNADNWLNMSKNLEKEISLENYKKFAKTTARLDKVELFEAYKNSLLLVMDLQSHLIEQ
YNLKVTHDILERLLNYISE
10000000000000000000000000000000000000000000000000000000000000
00000000000000000000000000000000000000000000000000000000000000
00000000000000000000000000000000000000000000000000000000000000
00000000000000000000000000000000000000000000000000000000000000
0000000000000000000
>DM_validata409
GMMFKWPWKADDESGNAEMPWEQALAIPVLAHLSSTEQHKLTQMAARFLQQKRLVALQGLEL
TPLHQARIAMLFCLPVLELGIEWLDGFHEVLIYPAPFIVDDEWEDDIGLVHNQRVVQSGQSW
QQGPVVLNWLDIQDSFDASGFNLVVHEVAHKLDTRNGDRASGVPLIPLREVAGWEHDLHAAM
NNIQDEIDLVGESAASIDAYAATDPAECFAVLSEYFFSAPELFAPRFPALWQRFCHFYRQDP
LARRRENGLQDEGDRRIVH
11111111111111110000000000000000000000000000000000000000000000
00000000000000000000000000000000000001111111111000000000011111
00000000000000000000000000000000000000000000000000000000000000
00000000000000000000000000000000000000000000000000000000000000
0000001111111111111
>DM_validata410
MALSMPLNGLKEEDKEPLIELFVKAGSDGESIGNCPFSQRLFMILWLKGVVFSVTTVDLKRK
PADLQNLAPGTHPPFITFNSEVKTDVNKIEEFLEEVLCPPKYLKLSPKHPESNTAGMDIFAK
FSAYIKNSRPEANEALERGLLKTLQKLDEYLNSPLPDEIDENSMEDIKFSTRKFLDGNEMTL
ADCNLLPKLHIVKVVAKKYRNFDIPKEMTGIWRYLTNAYSRDEFTNTCPSDKEVEIAYSDVA
KRLPSKVPKGEFQHTGGRY
11111111111111100000000000000000000000000000000000000000000000
00000000000000000000000000000000000000000000000000000000000000
00000000000000000000000000000000000111111111111110000000000000
00000000000000000000000000000000000000000000000000000000000000
0000000001111111111
>DM_validata411
MFDKHTHTLIAQRLDQAEKQREQIRAISLDYPEITIEDAYAVQREWVRLKIAEGRTLKGHKI
GLTSKAMQASSQISEPDYGALLDDMFFHDGSDIPTDRFIVPRIEVELAFVLAKPLRGPNCTL
FDVYNATDYVIPALELIDARCHNIDPETQRPRKVFDTISDNAANAGVILGGRPIKPDELDLR
WISALMYRNGVIEETGVAAGVLNHPANGVAWLANKLAPYDVQLEAGQIILGGSFTRPVPARK
GDTFHVDYGNMGSISCRFV

00000000000000000000000000000000000000000000000000000000000000
00000000000000000000000000000000000000000000000000000000000000
0000000000000000000000001111111000000000000000000000000000000000
00000000000000000000000000000000000000000000000000000000000000
0000000000000000000
>DM_validata412
TRQEIFQEQLAAVPEFRGLGPLFKSSPEPVALTESETEYVIRCTKHTFTNHMVFQFDCTNTL
NDQTLENVTVQMEPTEAYEVLCYVPARSLPYNQPGTCYTLVALPKEDPTAVACTFSCMMKFT
VKDCDPTTGETDDEGYEDEYVLEDLEVTVADHIQKVMKLNFEAAWDEVGDEFEKEETFTLST
IKTLEEAVGNIVKFLGMHPCERSDKVPDNKNTHTLLLAGVFRGGHDILVRSRLLLLDTVTMQ
VTARSLEELPVDIILASVG
00000000000000000000000000000000000000000000000000000000000000
00000000000000000000000000000000000000000000000000000000000000
00000000000000000000000000000000000000000000000000000000000000
00000000000000000000000000000000000000000000000000000000000000
0000000000000000001
>DM_validata413
MSCYIYWDKIKRIASRLEGMNYHFDEMDTSGVMPLLDEIEEIAHDSTIDFESAKHILDDAEM
NHALSLIRKFYVNLGMKLEMEKAQEVIESDSPWETLRSFYFYPRYLELLKNEAALGRFRRGE
RAVFIGGGPLPLTGILLSHVYGMRVNVVEIEPDIAELSRKVIEGLGVDGVNVITGDETVIDG
LEFDVLMVAALAEPKRRVFRNIHRYVDTETRIIYRTYTGMRAILYAPVSDDDITGFRRAGVV
LPSGKVNNTSVLVFKCPD
10000000000000000000000000000000000000000000000000000000000000
00000000000000000000000000000000000000000000000000000000000000
00000000000000000000000000000000000000000000000000000000000000
00000000000000000000000000000000000000000000000000000000000000
000000000000000001
>DM_validata414
CSVHAECRDYATGFCCRCVANYTGNGRQCVAEGSPQRVNGKVKGRIFVGSSQVPVVFENTDL
HSYVVMNHGRSYTAISTIPETVGYSLLPLAPIGGIIGWMFAVEQDGFKNGFSITGGEFTRQA
EVTFLGHPGKLVLKQQFSGIDEHGHLTISTELEGRVPQIPYGASVHIEPYTELYHYSSSVIT
SSSTREYTVMEPDQDGAAPSHTHIYQWRQTITFQECAHDDARPALPSTQQLSVDSVFVLYNK
EERILRYALSNSIGPVR
00000001111111100000000000000000000000000000000000000000000000
00000000000000000000000000000000000000000000000000000000000000
00000000000000000000000000000000000000000000000000000000000000
00000000000011111100000000000000000011111100000000000000000000
00000000000000000
>DM_validata416
MAKMRISPELKKLIEKYRCVKDTEGMSPAKVYKLVGENENLYLKMTDSRYKGTTYDVEREKD
MMLWLEGKLPVPKVLHFERHDGWSNLLMSEADGVLCSEEYEDEQSPEKIIELYAECIRLFHS
IDISDCPYTNSLDSRLAELDYLLNNDLADVDCENWEEDTPFKDPRELYDFLKTEKPEEELVF
SHGDLGDSNIFVKDGKVSGFIDLGRSGRADKWYDIAFCVRSIREDIGEEQYVELFFDLLGIK
PDWEKIKYYILLDELF

111100000000000000000000000000000000000000000000000000000000
000000000000000000000000000000000000000000000000000000000000
000000000000000000000000000000000000000000000000000000000000
000000000000000000000000000000000000000000000000000000000000
0000000000000000
>DM_validata417
LRVGFIGFGEVAQTLASRLRSRGVEVVTSLEGRSPSTIERARTVGVTETSEEDVYSCPVVIS
AVTPGVALGAARRAGRHVRGIYVDINNISPETVRMASSLIEKGGFVDAAIMGSVRRKGADIR
IIASGRDAEEFMKLNRYGLNIEVRGREPGDASAIKMLRSSYTKGVSALLWETLTAAHRLGLE
EDVLEMLEYTEGNDFRESAISRLKSSCIHARRRYEEMKEVQDMLAEVIDPVMPTCIIRIFDK
LKDVKVSADARLQGCA
000000000000000000000000000000000000000000000000000000000000
000000000000000000000000000000000000000000000000000000000000
000000000000000000000000000000000000000000000000000000000000
000000000000000000000000000000000000000000000000000000000000
0011111100000000
>DM_validata419
GLAESLELPKAIQDQLPRLQQLLKTLEEGLEGLEDAPPVELQLLHECSPSQMDLLCAQLQLP
QLSDLGLLRLCTWLLALSPDLSLSNATVLTRSLFLGRILSLTSSASRLLTTALTSFAAKYTY
PVCSALLDPVLQAPGTGPAQTELLCCLVKMESLEPDAQVLMLGQILELPWKEETFLVLQSLL
ERQVEMTPEKFSVLMEKLCKKGLAATTSMAYAKLMLTVMTKYQANITETQRLGLAMALEPNT
TFLRKSLKAALKHLGP
110000000000000000000000000111111100000000000000000000000000
000000000000000000000000000000000000000000000000000000000000
000000000000000000000000000000000000000000000000000000000000
000000000000000000011111000000000000000000000000000000000000
0000000000000001
>DM_validata420
MPQSAVMNVMVQAAMKAGRSLVRDYGEVQNLQVSLKGPADYVSQADRKAEKIIFNELSKARP
KFGFLMEESEEIIGEDSQHRFIVDPLDGTTNFLHGIPFFAVSIALESQGKIVAGVIYNPIND
ELFTAERGSGAFFNDRRCRVSARRRLEDCVIATGMPHLGRPGHGTYLIELRNVMAEVSGIRR
FGTAALDLAYVAAGRTDGFWEDNLQIWDMAAGILMVREAGGFVTDKEGGNDIFRKKNIIAGN
EHIRIKLERALKKGI
111000000000000000000000001111000000000000000000000000000000
000000000000000000000000000000000000000000000000000000000000
000000000000000000000000000000000000011000000000000000000000
000000000000000000000000000000000000000000000000000000000000
000000000000000
>DM_validata421
GMTQTTPQPDPSRLRDELVRLHGKASPEWDSLVRLDPRFVDAYLKFAGVPQRRNHLDDKTRA
FIALAADACATQLYAPGVARHIERALSFGATREELIEVLELVSTIGIHTSNVGVPVLLEVLE
EEGLRKGAPPLDERRQKLKAEFETNRGYWHPTWEGLLELDPDLFEAYVEFSSVPWRTGVLSP
KIKEFMYCAFDASATHLYVPGLKLHIRNALRYGATAEELMELLEIVSVTGIHGAELGAPLLE
AALKRSGAAAGEPHA

111111000000000000000000000000000000000000000000000000000000
000000000000000000000000000000000000000000000000000000000000
000000000000000000000000000000000000000000000000000000000000
000000000000000000000000000000000000000000000000000000000000
000000011111111
>DM_validata422
MSSALLRAAYAYDRLRAHPPEVAGQIATAMASAVHPKGEEPVFLELGVGTGRIALPLIARGY
RYIALDADAAMLEVFRQKIAGVDRKVQVVQADARAIPLPDESVHGVIVVHLWHLVPDWPKVL
AEAIRVLKPGGALLEGWDQAEASPEWTLQERWRAFAAEEGFPVERGLHAKRLKEVEEALRRL
GLKPRTREVARWREERTPREALEALSERLYSFTQGLPEPVHARVMERLWAWAEAELGDLDRP
FPVEKRFLLRVSRLG
100000000000000000000000000000000000000000000000000000000000
000000000000000000000000000000000000000000000000000000000000
000000000000000000000000000000000000000000000000000000000000
000000000000000000000000000000000000000000000000000000000000
000000000000001
>DM_validata423
KNSINTIKLIDDIIALHNDPKGNKLLWNDNWQDKIINRDLANIFEKIDESVSELGGLEMYQE
MVGVNPYDPTEPVCGLSAQNIFKLMTEGEHAVDPVEMAQTGKIDGNEFAESVDQLSSAKNYV
ALVNDRRLGHMFLIDIPSNDQETVGYIYQSDLGQGALPPLKIADWLNSRGKDAVSLNKLKKL
LSREFNLLSDDEKRALISETLDIHKDVSNVELDRIKRDRGVDIYLTEYDVNNFYENIETLKS
KLSNYDKKLSKPK
110000000000000000000000000000000000000000000000000000000000
000000000000000000000000000000000000000000000000000000000000
000000000000000000000000000000000000000000000000000000000000
000000000000000000000000000000000000000000000000000000000000
0011111111111
>DM_validata424
MRRLEEDAVRVAYAGLRRKEAFKALAEKLGFTPLLFPVQATEKVPVPEYRDQVRALAQGVDL
FLATTGVGVRDLLEAGKALGLDLEGPLAKAFRLARGAKAARALKEAGLPPHAVGDGTSKSLL
PLLPQGRGVAALQLYGKPLPLLENALAERGYRVLPLMPYRHLPDPEGILRLEEALLRGEVDA
LAFVAAIQVEFLFEGAKDPKALREALNTRVKALAVGRVTADALREWGVKPFYVDETERLGSL
LQGFKRALQKEVA
111111100000000000000000000000000000000000000000000000000000
000000000000000000000000000000000000000000000000000000000000
000000000000000000000000000000000000000000000000000000000000
000000000000000000000000000000000000000000000000000000000000
0000000000000
>DM_validata425
GAMGYKDNIRHGVCWIYYPDGGSLVGEVNEDGEMTGEKIAYVYPDERTALYGKFIDGEMIEG
KLATLMSTEEGRPHFELMPGNSVYHFDKSTSSCISTNALLPDPYESERVYVAESLISSAGEG
LFSKVAVGPNTVMSFYNGVRITHQEVDSRDWALNGNTLSLDEETVIDVPEPYNHVSKYCASL
GHKANHSFTPNCIYDMFVHPRFGPIKCIRTLRAVEADEELTVAYGYDHSPPGKSGPEAPEWY
QVELKAFQATQQK

1111111111000000000000000000000000000000000000000000000000000
000000000000000000000000000000000000000000000000000000000000
000000000000000000000000000000000000000000000000000000000000
000000000000000000000000000000000000000000000000111110000000
0000000000011
>DM_validata428
SQFVGSDGMGGDAAAPGAAGTQAELPHRRFEYKYSFKGPHLVQSDGTVPFWAHAGNAIPSAD
QIRIAPSLKSQRGSVWTKTKAAFENWEVEVTFRVTGRGRIGADGLAIWYTENQGLDGPVFGS
ADMWNGVGIFFDSFDNDGKKNNPAIVVVGNNGQINYDHQNDGATQALASCQRDFRNKPYPVR
AKITYYQKTLTVMINNGFTPDKNDYEFCAKVENMVIPTQGHFGISAATGGLADDHDVLSFLT
FQLTEPGKEPPT
111111111111111111111111100000000000000000000000000000000000
000000000000000000000000000000000000000000000000000000000000
000000000000000011111000000000000000000000000000000000000000
000000000000000000000000000000000000000000000000000000000000
000001111111
>DM_validata429
TIYVVTPTYARLVQKAELVRLSQTLSLVPRLHWLLVEDAEGPTPLVSGLLAASGLLFTHLVV
LTPKAQRLREGEPGWVHPRGVEQRNKALDWLRGRGGAVGGEKDPPPPGTQGVVYFADDDNTY
SRELFEEMRWTRGVSVWPVGLVGGLRFEGPQVQDGRVVGFHTAWEPSRPFPVDMAGFAVALP
LLLDKPNAQFDSTAPRGHLESSLLSHLVDPKDLEPRAANCTRVLVWHTRTEKPKMKQEEQLQ
RQGRGSDPAIEV
000000000000000000000000000000000000000000000000000000000000
000111111111111110000000000000000000000000000000000000000000
000000000000000000000000000000000000000000000000000000000000
000000000000000000000000000000000000000000000000000000000000
000000000000
>DM_validata430
MSKSVFVGELTWKEYEARVAAGDCVLMLPVGALEQHGHHMCMNVDVLLPTAVCKRVAERIGA
LVMPGLQYGYKSQQKSGGGNHFPGTTSLDGATLTGTVQDIIRELARHGARRLVLMNGHYENS
MFIVEGIDLALRELRYAGIQDFKVVVLSYWDFVKDPAVIQQLYPEGFLGWDIEHGGVFETSL
MLALYPDLVDLDRVVDHPPATFPPYDVFPVDPARTPAPGTLSSAKTASREKGELILEVCVQG
IADAIREEFPPT
110000000000000000000000000000000000000000000000000000000000
000000000000000000000000000000000000000000000000000000000000
000000000000000000000000000000000000000000000000000000000000
000000000000000000000000000000000000000000000000000000000000
000000000001
>DM_validata431
MEGGSITPGEDKPEIQSPIPPNQIFILSGQSNMAGRGGVFKDHHNNRWVWDKILPPECAPNS
SILRLSADLRWEEAHEPLHVDIDTGKVCGVGPGMAFANAVKNRLETDSAVIGLVPCASGGTA
IKEWERGSHLYERMVKRTEESRKCGGEIKAVLWYQGESDVLDIHDAESYGNNMDRLIKNLRH
DLNLPSLPIIQVAIASGGGYIDKVREAQLGLKLSNVVCVDAKGLPLKSDNLHLTTEAQVQLG
LSLAQAYLSNFC

1111111111111111111000000000000000000000000000000000000000000
000000000000000000000000000000000000000000011100000000000000
000000000000000000000000000000000000000000000000000000000000
000000000000000000000000000000000000000000000000000000000000
000000000000
>DM_validata432
VTAVPTDSAEKPAVADAGVRSVTRVIDLLELFDAAHPTRSLKELVEGTKLPKTTVVRLVATM
CARSVLTSRADGSYSLGPEMLRWVRLAGRTWAPPEEVVDIMRQLSADTGETVNLYIRQGLSR
VVVAQCESTATVRSVIPLGVPYPLWAGAAGKILLLAAPELIDDVAADSPHGPEFADQLREKV
EDGRERGYQLVHGERELGSSGLSFPLVDSHGTVVAALTLGGPTGRFTEDRTPHYIECTRAAA
EEISAIGLPGLD
111111111111111110000000000000000000000000000000000000000000
000000000000000000000000000000000000000000000000000000000000
000000000000000000000000000000000000000000000000000000000000
000000000000000000000000000000000000000000000000000000000000
000000000001
>DM_validata433
GMTEAFPALVRQDDARYAITVGPDLAVGPPGHAYLFGGASMALALDVAAETVGRPVVQGSLQ
FVSFTPLGSVLDLTVEVLQSGRTLAQARVAGTVDGRLVFHSGISLGMREGFSARQWALAPPV
PQPDNCPPCTTLPAQDDNARYLEGIEVREAGGPEVPSGRTRLWLRRKDGAPLDAASLAMFAD
FLPIALGRATGCSGGGNSLDNSLRITGAAAPGWCLCDMIIPSSASGFAQGQVTLWDQSGRLL
ATGAQSLLLKG
111100000000000000000000000000000000000000000000000000000000
000000000000000000000000000000000000000000000000000000000000
000000000000000000000000000000000000000000000000000000000000
000000000000011000000000000000000000000000000000000000000000
00000000000
>DM_validata434
MQQSVCRARPIWWGTQRRGSETMAGAAVKYLSQEEAQAVDQELFNEYQFSVDQLMELAGLSC
ATAIAKAYPPTSMSKSPPTVLVICGPGNNGGDGLVCARHLKLFGYQPTIYYPKRPNKPLFTG
LVTQCQKMDIPFLGEMPPEPMMVDELYELVVDAIFGFSFKGDVREPFHSILSVLSGLTVPIA
SIDIPSGWDVEKGNPSGIQPDLLISLTAPKKSATHFTGRYHYLGGRFVPPALEKKYQLNLPS
YPDTECVYRLQ
111111111111111111111111100000000000000000000000000000000000
000000000000000000000000000000000000000000000000000000000000
000000000000000000000000000000000000000000000000000000000000
000000000000000000000000000000000000000000000000000000000000
00000000000
>DM_validata435
MKDLKKIESYLDKLRIKEKDGEERKIYAEVLDGRTLKTLYKLSAKGYITAMGGVISTGKEAN
VFYADGVFDGKPVAMAVKIYRIETSEFDKMDEYLYGDERFDMRRISPKEKVFIWTEKEFRNL
ERAKEAGVSVPQPYTYMKNVLLMEFIGEDELPAPTLVELGRELKELDVEGIFNDVVENVKRL
YQEAELVHADLSEYNIMYIDKVYFIDMGQAVTLRHPMAESYLERDVRNIIRFFSKYGVKADF
EEMLKEVKGE

000000000000000000000000000000000000000000000000000000000000
000000000000000000000111111100000000111111111000000000000000
000000000000000000000000000000000000000000000000000000000000
000000000000000000000000000000000000000000000000000000000000
0000000000
>DM_validata436
MKEKSIEDIVFEKFQPYINWSIDKLCEHFSINKGEKGLNYRIASAILNLKGKTTKSKPFPEV
EEFEKSSIVVKTVHFNKKNVNKESMSFGAFKFEELANEEWEDSEGYPSAQWRNFLLETRFLF
FVVKEDEDGVDIFKGIKFFSMPEEDINGPVKRMWDDTVKKLKEGVTLEAVPDKSTKDGWRIK
NNFVDKSDDLICHVRPHTNNRDYRGGSNADKLPKKINWINRPDSDDYSDEWMTKQSFWINND
YIKKQVEDLL
111100000000000000000000000000000000000000000011111111111111
111100000000000000000000000000000000000000000000000000000000
000000000000000000000000000000000000000000000000000000000000
000000000000000000000000000000000000000000000000000000000000
0000000000
>DM_validata438
SFIKPIYQDINSILIGQKVKRPKSGTLSGHAAGEPFEKLVYKFLKENLSDLTFKQYEYLNDL
FMKNPAIIGHEARYKLFNSPTLLFLLSRGKAATENWSIENLFEEKQNDTADILLVKDQFYEL
LDVKTRNISKSAQAPNIISAYKLAQTCAKMIDNKEFDLFDINYLEVDWELNGEDLVCVSTSF
AELFKSEPSELYINWAAAMQIQFHVRDLDQGFNGTREEWAKSYLKHFVTQAEQRAISMIDKF
VKPFKKYIL
000000000000000000011111111111000000000000000000000000000000
011111110000000000000000000000000000011100000000000000000000
000000000000000000000000000000000000000000000000000000000000
000000000000000000000000000000000000000000000000000000000000
000000000
>DM_validata439
MLTFALTIVRHGETQYNRDKLLQGQGIDTPLSDTGHQQAAAAGRYLKDLHFTNVFVSNLQRA
IQTAEIILGNNLHSSATEMILDPLLRERGFGVAEGRPKEHLKNMANAAGQSCRDYTPPGGET
LEQVKTRFKMFLKSLFQRMFEEHGSALSSVPSEADQPVIAGLADDGAQNVPVHALMVSHGAF
IRISVRHLVEDLQCCLPAGLKMNQVFSPCPNTGISRFIFTIHREESVLRATRIQGVFINRKD
HLEEVKNSD
100000000000000111111111111100000000000000000000000000000000
000000000000000000000000000001111111111111111111111111000000
000000000000000000000001111111111100000000000000000000000000
000000000000000000000000000000000000000000000000000000000000
001111111
>DM_validata440
GALPDPVKPKAPKAVNPFHLGMAGYTFVNFDLDTTLKTLERLDIHYLCIKDFHLPLNSTDEQ
IRAFHDKCAAHKVTGYAVGPIYMKSEEEIDRAFDYAKRVGVKLIVGVPNYELLPYVDKKVKE
YDFHYAIHLHGPDIKTYPDATDVWVHTKDLDPRIGMCLDVGHDLRNGCDPVADLKKYHTRVF
DMHIKDVTDSSKAGVGIEIGRGKIDFPALIRMMREVNYTGMCSLEYEKDMKDPFLGIAESIG
YFKAVSDLT

111111000000000000000000000000000000000000000000000000000000
000000000000000000000000000000000000000000000000000000000000
000000000000000000000000000000000000000000000000000000000000
000000000000000000000000000000000000000000000000000000000000
000000000
>DM_validata441
MSLRIPRIYHPISLENQTQCYLSEDAANHVARVLRMTEGEQLELFDGSNHIYPAKIIESNKK
SVKVEILGRELADKESHLKIHLGQVISRGERMEFTIQKSVELGVNVITPLWSERCGVKLDAE
RMDKKIQQWQKIAIAACEQCGRNIVPEIRPLMKLQDWCAENDGALKLNLHPRAHYSIKTLPT
IPAGGVRLLIGSEGGLSAQEIAQTEQQGFTEILLGKRVLRTETASLAAISALQICFGDLGEE
GGSHHHHHH
111100000000000000000000000000000000000000000000000000011111
100000000000000000000000000110000000000000000000000000000000
000000000000000000000000000000000000000000000000000000000000
000000000000000000000000000000000000000000000000000000000000
011111111
>DM_validata442
EMETYVNKLHEGSTYTAAVQYNVLEKDDDPASLTIWVPMFQSSMPADLLIKELANVNILVKQ
ISTPKGPSLRVMINSRSAVLAQMPSKFTICANVSLDDRSKLAYDVTTPCEIKACSLTCLKSK
NMLTTVKDLTMKTLNPTHDIIALCEFENIVTSKKVIIPTYLRSISVRNKDLNTLENITTTEF
KNAITNAKIIPYSGLLLVITVTDNKGAFKYIKPQSQFIVDLGAYLEKESIYYVTTNWKHTAT
RFAIKPRED
000000000000000000000000000000000000000000000000000000000000
000000000000000000000000000000000000100000000000000000000000
000000000000000000000000000000000000000000000000000000000000
000000000000000000000000000000000000000000000000000000000000
000000001
>DM_validata443
FWRQTWSGPGTTKRFPETVLARCVKYTEIHPEMRHVDCQSVWDAFKGAFISKHPCDITEEDY
QPLMKLGTQTVPCNKILLWSRIKDLAHQFTQVQRDMFTLEDTLLGYLADDLTWCGEFDTSKI
NYQSCPDWRKDCSNNPVSVFWKTVSRRFAEAACDVVHVMLDGSRSKIFDKDSTFGSVEVHNL
QPEKVQTLEAWVIHGGREDSRDLCQDPTIKELESIISKRNIQFSCKNIYRPDKFLQCVKNPE
DSSCTSEI
000000000000000000000000000000000000000000000000000000000000
000000000000000000000000000000000000000000000000000000000000
000000000000000000000000000000000000000000000000000000000000
000000000000000000000000000000000000000000000001111111111111
11111111
>DM_validata444
MDNSHDSDSVFLYHIPCDNCGSSDGNSLFSDGHTFCYVCEKWTAGNEDTKERASKRKPSGGK
PGTYNVWNFGESNGRYSALTARGISKETCQKAGYWIAKVDGVMYQVADYRDQNGNIVSQKVR
DKDKNFKTTGSHKSDALFGKHLWNGGKKIVVTEGEIDMLTVMELQDCKYPVVSLGHGASAAK
KTCAANYEYFDQFEQIILMFDMDEAGRKAVEEAAQVLPAGKVRVAVLPCKDANECHLNGHDR
EIMEQVW

111111111000000000000000000000000000000111111100000000000000
000000000000000000000000000000000000000000000000000000000000
000000000000000000000000000000000000000000000000000000000000
000000000000000000000000000000000000000000000000000000000000
0000000
>DM_validata445
SKAVTGMRTIDGKKYYFNTNTAEAATGWQTIDGKKYYFNTNTSIASTGYTIINDKHFYFNTD
GIMQIGVFKGPDGFEYFAPANTDANNIEGQAIRYQNRFLYLHDNIYYFGNNSKAATGWVTID
GRRYYFEPNTAIGANGYKIIDNKNFYFRNGLPQIGVFKGPNGFEYFAPANTDANNIDGQAIR
YQNRFLHLLGNIYYFGNNSKAVTGWQTINGNMYYFMPDTAMAAAGGLFEIDGVIYFFGVDGV
KAPGIYG
111110000000000000000000000000000000000000000000000000000000
000000000000000000000000000000000000000000000000000000000000
000000000000000000000000000000000000000000000000000000000000
000000000000000000000000000000000000000000000000000000000000
0011111
>DM_validata446
QGHMRVEVFPVEGLPLIKEGDDLAELISSRVRFEDGDVLVVCSTVISKAEGRIRRLEEFNPS
ERAKEIAARIGKPAEFVQAVLEESEEVLLDFPFLLVKAKFGNVCVNAGIDASNVEEGSLLLP
PLDPDGSAEKLRRRILELTGKRVGVIITDTNGRCFRRGVVGFAIGISGVKAMKDWIGRKDLY
GRELEVTVECVADEIAAFANLLMGEGGDGIPAVVVRGLNVAGEGSMEEIYRSEEEDVIRRCL
KRCLGS
111100000000000000000000000000000000000000000000000000000000
000000000000000000000000000000000000000000000000000000000000
000000000000000000000000000000000000000000000000000000000000
000000000000000000000000000000000000000000000000000000000000
000011
>DM_validata447
DKGKSVKSTEKSVGMPPKTPKTDNNAHVDNEFLILQVNDAVFPIGSYTHSFGLETYIQQKKV
TNKESALEYLKANLSSQFLYTEMLSLKLTYESALQQDLKKILGVEEVIMLSTSPMELRLANQ
KLGNRFIKTLQAMNELDMGEFFNAYAQKTKDPTHATSYGVFAASLGIELKKALRHYLYAQTS
NMVINCVKSVPLSQNDGQKILLSLQSPFNQLIEKTLELDESHLCTASVQNDIKAMQHESLYS
RLYMS
111111111111111111111100000000000000000000001111100000000000
000000000000000000000000000000000000000000000000000000000000
000000000000000000000000000000000000000000000000000000000000
000000000000000000000000000000000000000000001111111111111111
11111
>DM_validata448
WMTGHHHHHHRQKWEWKVGTGLNGFGSVLNDLTNGGTKLTITVTGNKPILLGRTKEAFATPV
TSGVDGIPHIAFTDYEGASVELRNPDGETEKGLAYFVLPMKNAEGTKVGSVKVNASYAGALG
RGGVTSADGELMSLFAEGSHAIFYGGLPTNVKNSELKGGSAAAARTELFGSLSKNDILGQIQ
RVNANITSLVNVPGSFNENMAYTDGSVVSVAYALGIANGQTIEATFNQAVTTSTQWSAPLNV
AITYY

1111111111000000000000000001000000000000000000000000000000000001
1111100000000000000000000000000000000000000000000000000000000000
0000000000000000111111000000000000000000000000000000000000000000
0000000000001111111111111000000000000000000000000001100000000000
00000
>DM_validata449
AGHMEAVIEKECSALGGLFQTIISDMKGSYPVWEDFINKAGKLQSQLRTTVVAAAAFLDAFQ
KVADMATNTRGGTREIGSALTRMCMRHRSIEAKLRQFSSALIDCLINPLQEQMEEWKKVANQ
LDKDHAKEYKKARQEIKKKSSDTLKLQKKAKKVDAQGRGDIQPQLDSALQDVNDKYLLLEET
EKQAVRKALIEERGRFCTFISMLRPVIEEEISMLGEITHLQTISEDLKSLTMDPHKLPSSSE
QVILD
0000000000000000000000000000000000000000000000000000000000000000
0000000000000000000000000000000000000000000000000000000000000000
0000000000000000000000000000000000011111111111111000000000000000
0000000000000000000000000000000000000000000000000000000000000111
11111
>DM_validata450
EPREFVLRPAPQGRTVRCRLTRDKKGMDRGMYPSYFLHLDTEKKVFLLAGRKRKRSKTANYL
ISIDPTNLSRGGENFIGKLRSNLLGNRFTVFDNGQNPQRGYSTNVASLRQELAAVIYETNVL
GFRGPRRMTVIIPGMSAENERVPIRPRNASDGLLVRWQNKTLESLIELHNKPPVWNDDSGSY
TLNFQGRVTQASVKNFQIVHADDPDYIVLQFGRVAEDAFTLDYRYPLCALQAFAIALSSFDG
KLACE
1000000000000000000000001111111110000000000000000000000000000000
0000000111111000000000000000000000000000000000000000000000001111
1111100000000000000000000000000000000000000000000000000000000000
0000000000000000000000000000000000000000000000000000000000000001
11111
>DM_validata451
AQELQTANEFTVHTDLSSISSTRAFLKEKHKAAKHIGVRADIPFDANQGIRLEAGFGRSKKN
IINLETDENKLGKTKNVKLPTGVPENRIDLYTGYTYTQTLSDSLNFRVGAGLGFESSKDSIK
TTKHTLHSSRQSWLAKVHADLLSQLGNGWYINPWSEVKFDLNSRYKLNTGVTNLKKDINQKT
NGWGFGLGANIGKKLGESASIEAGPFYKQRTYKESGEFSVTTKSGDVSLTIPKTSIREYGLR
VGIKF
1111110000000000000000000000000000000000000000000000000000000000
0000000000000000000000000000000000000000000000000000000000000001
1111100000000000000000000000000000000000000000000001111111000000
0000000000000000110000000000000000000000011111110000000000000000
00000
>DM_validata452
MSLTLTLTGTGGAQGVPAWGCECAACARARRSPQYRRQPCSGVVKFNDAITLIDAGLHDLAD
RWSPGSFQQFLLTHYHMDHVQGLFPLRWGVGDPIPVYGPPDEQGCDDLFKHPGLLDFSHTVE
PFVVFDLQGLQVTPLPLNHSKLTFGYLLETAHSRVAWLSDTAGLPEKTLKFLRNNQPQVMVM
DCSHPPRADAPRNHCDLNTVLALNQVIRSPRVILTHISHQFDAWLMENALPSGFEVGFDGME
IGVA

1000000000000000000000000000000000000000000000000000000000000000
0000000000000000000000000000000000000000000000000000000000000000
0000000000000000000000000000000000000000000000000000000000000000
0000000000000000000000000000000000000000000000000000000000000000
0011
>DM_validata454
AQTAAAATSRIKKFSIYRWDPDKPGDKPRMQTYEVDLNKCGPMVLDALIKIKNELDSTLTFR
RSCREGICGSCAMNIAGGNTLACTKKIDPDLSKTTKIYPLPHMYVVKDLVPDLSNFYAQYKS
IEPYLKKKDESKQGKEQYLQSIEDRQKLDGLYECILCACCSTSCPSYWWNGDKYLGPAVLMQ
AYRWMIDSRDDYTEERLAQLQDPFSLYRCHTIMNCTRTCPKGLNPGKAIAEIKKMMATYKEK
AAAA
1111111000000000000000000000000000000000000000000000000000000000
0000000000000000000000000000000000000000000000000000000000000000
0000000000000000000000000000000000000000000000000000000000000000
0000000000000000000000000000000000000000000000000000000000000011
1111
>DM_validata456
AMRTIYVIGIGTGSPEFLTLQAISGLRHAQAIVALDKGEQKSDLLALRQKIVDTHAPGTPIY
AVTDPERDRNPDNYEEEVRRWHAERAHLLASTIRERTPDDGAVAFLVWGDPSLYDSTLRIIE
HMRNLEDLHADVKVIPGITAVQVLTAEHGILINRIGEAIHITTGRNLPETSAKDRRNCVVML
DGKTAWQDVATEHTYMWWGAFLGTEQQVLRKGYVHEIGAQVAELKQQLRTEHGWIMDTYLLR
ELD
0000000000000000000000000000000000000111000000000000000000000000
0000111111111100000000000000000000000000000000000000000000000000
0000000000000000000000000000000000000000000000000000000000000000
0000000000000000000000000000000000000000000000000000000000000000
000
>DM_validata457
GMTISRNYDQEIKDTAGHKYAYNFDFDVMHPFMVRAFTPFFRPGNLLELGSFKGDFTSRLQE
HFNDITCVEASEEAISHAQGRLKDGITYIHSRFEDAQLPRRYDNIVLTHVLEHIDDPVALLK
RINDDWLAEGGRLFLVCPNANAVSRQIAVKMGIISHNSAVTEAEFAHGHRCTYALDTLERDA
SRAGLQVTYRSGIFFKALANFQWDQILQTDILSKEYLDGCYQLGQQYPDLCASIFLLCEKGI
NQ
1111111111111111111111100000000000000000000000000000000000000000
0000000000000000000000000000000000000000000000000000000000000000
0000000000000000000000000000000000000000000000000000000000000000
0000000000000000000000000000000000000000000000000000000000000001
11
>DM_validata458
LNKTLFLGYAFLLTMATTSDKLASRSKLPDGPTGSSEEEEEFLEIPPFNKQYTESQLRAGAG
YILEDFNEAQCNTAYQCLLIADQHCRTRKYFLCLASGIPCVSHVWVHDSCHANQLQNYRNYL
LPAGYSLEEQRILDWQPRENPFQNLKVLLVSDQQQNFLELWSEILMTGGAASVKQHHSSAHN
KDIALGVFDVVVTDPSCPASVLKCAEALQLPVVSQEWVIQCLIVGERIGFKQHPKYKHDYVS
H

00000000000000000111111111111111111111111111110000000000000000
00000000111000000000000000000000000000000000000000000000000000
00000000000000000000000000000000000000000000000000000000000000
00000000000000000000000000000000000000000000000000000000000011
1
>DM_validata459
MGFMIEHWDFSTPMATQETTTAEHIQPNHWYHCERLHPDIRGWLEDNHVPRATVDHLLADES
RPSFHPLDDDNFMLILRGINMNENASPEDMLSIRILYFQGALISTRKIPSRAIMEIRQALAE
HKGPKSLASLLNQIIEGLNGKIDLYLDTIEETLNEFDVNDESTYNHIAAQKALISIKRFIRP
QQYAIRDLIESESELVTSRPHQYRFAHNNITRINETIEFYLGEVALFQDEIKHNRDEKTNKN
S
10000000000000000000000000000000000000000000000000000000000000
00000000000000000000000000000000000000000000000000000000000000
00000000000000000000000000000000000000000000000000000000000000
00000000000000000000000000000000000000000000000000000000011111
1
>DM_validata460
HGNQLTAYTLRLGDNCLVLSQRLGEWCGHAPELEIDLALANIGLDLLGQARNFLSYAAELAG
EGDEDTLAFTRDERQFSNLLLVEQPNGNFADTIARQYFIDAWHVALFTRLMESRDPQLAAIS
AKAIKEARYHLRFSRGWLERLGNGTDVSGQKMQQAINKLWRFTAELFDADEIDIALSEEGIA
VDPRTLRAAWEAEVFAGINEATLNVPQEQAYRTGGKKGLHTEHLGPMLAEMQYLQRVLPGQQ
W
00000000000000000000000000000000000000000000000000000000000000
00000000000000000000000000000000000000000000000000000000000000
00000000000000000000000000000000000000000000000000000000000000
00000000000000000000000000000000000000000000000000000000001111
1
>DM_validata461
MKSCAILAALGCLAGSVLGHGQVQNFTINGQYNQGFILDYYYQKQNTGHFPNVAGWYAEDLD
LGFISPDQYTTPDIVCHKNAAPGAISATAAAGSNIVFQWGPGVWPHPYGPIVTYVVECSGSC
TTVNKNNLRWVKIQEAGINYNTQVWAQQDLINQGNKWTVKIPSSLRPGNYVFRHELLAAHGA
SSANGMQNYPQCVNIAVTGSGTKALPAGTPATQLYKPTDPGILFNPYTTITSYTIPGPALWQ
G
11111111111111111110000000000000000000000000000000000000000000
00000000000000000000000000000000000000000000000000000000000000
00000000000000000000000000000000000000000000000000000000000000
00000000000000000000000000000000000000000000000000000000000001
1
>DM_validata462
MFKKYSSLENHYNSKFIEKLYSLGLTGGEWVAREKIHGTNFSLIIERDKVTCAKRTGPILPA
EDFFGYEIILKNYADSIKAVQDIMETSAVVSYQVFGEFAGPGIQKNVDYCDKDFYVFDIIVT
TESGDVTYVDDYMMESFCNTFKFKMAPLLGRGKFEELIKLPNDLDSVVQDYNFTVDHAGLVD
ANKCVWNAEAKGEVFTAEGYVLKPCYPSWLRNGNRVAIKCKNSKFSEKKKSDKPIKAKVELS
E

000000000000000000000000000000000000000000000000000000000000
000000000000000000000000000000000000000000000000000000000000
000000000000000000000000000000000000000000000000000000000000
000000000000000000000000000000000000000000000111111111111111
1
>DM_validata464
MALLEICCYSMECALTAQQNGADRVELCAAPKEGGLTPSLGVLKSVRQRVTIPVHPIIRPRG
GDFCYSDGEFAAILEDVRTVRELGFPGLVTGVLDVDGNVDMPRMEKIMAAAGPLAVTFHRAF
DMCANPLYTLNNLAELGIARVLTSGQKSDALQGLSKIMELIAHRDAPIIMAGAGVRAENLHH
FLDAGVLEVHSSAGAWQASPMRYRNQGLSMSSDEHADEYSRYIVDGAAVAEMKGIIERHQAK
100000000000000000000000000000000000000000000000000000000000
000000000000000000000000000000000000000000000000000000000000
000000000000000000000000000000000000000000000000000000000000
000000000000000000000000111111111111100000000000000000000000
>DM_validata465
DLPDKIETNVYCNLTPEQAAMYKAEVENLFNNIDSVTGIKRKGMILSTLLKLKQIVDHPALL
KGGEQSVRRSGKMIRTMEIIEEALDEGDKIAIFTQFVDMGKIIRNIIEKELNTEVPFLYGEL
SKKERDDIISKFQNNPSVKFIVLSVKAGGFGINLTSANRVIHFDRWWNPAVEDQATDRVYRI
GQTRNVIVHKLISVGTLEEKIDQLLAFKRSLFKDIISSGDSWITELSTEELRKVIELSVGGY
111000000000000000000000000000000000000000000000000000000000
000000000000000000000000000000000000000000000000000000000000
000000000000000000000000000000000000000000000000001111111111
111100000000000000000000000000000000000000000000000000000111
>DM_validata466
MRGSHHHHHHGSVTCTGCVDLDELSFEKTVERFPYSVVKFDIASPYGEKHEAFTAFSKSAHK
ATKDLLIATVGVKDYGELENKALGDRYKVDDKNFPSIFLFKGNADEYVQLPSHVDVTLDNLK
AFVSANTPLYIGRDGCIKEFNEVLKNYANIPDAEQLKLIEKLQAKQEQLTDPEQQQNARAYL
IYMRKIHEVGYDFLEEETKRLLRLKAGKVTEAKKEELLRKLNILEVFRVHKVTKTAPEKEEL
111111111111100000000000000000000000000000000000000000000000
000000000000000000000000000000000000000000000000000000000000
000000000000000000000000000000000000000000000000000000000000
000000000000000000000000000000000000000000000001111111111111
>DM_validata467
MSLETKTPNDAKNQVQTHERMKTSSEHVTPLDFNYPIHIVQAPQNHHVVGILTPRIQVSDNL
KPYIDKFQDALINQIQTIFEKRGYQVLRFQDEKALNAQDKRKIFSVLDLKGWVGILEDLKMN
LKDPNNPNLDTLVDQSSGSVWFNFYEPESNRVVHDFAVEVGTFQAMTYTYKHNNSGGLNSSN
SIIHEYLEKNKEDAIHKILNRMYAVVMKKAVTELTKENIDKYREAIDRMKGFKSSMPQKKEG
111111111111111110000000000000000000000000000000000000000000
000000000000000000000000000000000000000000000000000000000000
000000000000000000000000000000000000000000000000000000000000
000000000000000000000000000000000000000000000000000111111111
>DM_validata468
MLIIILTGLPGVGKSTFSKNLAKILSKNNIDVIVLGSDLIRESFPVWKEKYEEFIKKSTYRLI
DSALKNYWVIVDDTNYYNSMRRDLINIAKKYNKNYAIIYLKASLDVLIRRNIERGEKIPNEV

IKKMYEKFDEPGKKYKWDEPFLIIDTTKDIDFNEIAKKLIEKSKEIPKFYVLEENKNKNNNI
SDKIDKETRKIVSEYIKSKKLDKDKIKEVVELRKEFLKKIKKMEEVDADRVLKEFKDLLNSY
00000000000000000000000000000000000000000000000000000000000000
00000000000000000000000000000000000000000000000000000000000000
00000000000000000000000000000000000000000000000001111111111100
00000000000000000000000000000000000000000011100000000000000000
>DM_validata470
TESPADSFLKVELELNLKLSNLVFQDPVQYVYNPLVYAWAPHENYVQTYCKSKKEVLFLGMN
PGPFGMAQTGVPFGEVNHVRDWLQIEGPVSKPEVEHPKRRIRGFECPQSEVSGARFWSLFKS
LCGQPETFFKHCFVHNHCPLIFMNHSGKNLTPTDLPKAQRDTLLEICDEALCQAVRVLGVKL
VIGVGRFSEQRARKALMAEGIDVTVKGIMHPSPRNPQANKGWEGIVRGQLLELGVLSLLTG
11100000000000000000000000000000000000000000000000000000000000
00000000000000000000000000000000000000000000000000000000000000
00000000000000000000000000000000000000000000000000000000000000
0000000000000000000000000000000000000000000000000000000000000
>DM_validata471
MMKIVPPKPFFFEAGERAVLLLHGFTGNSADVRMLGRFLESKGYTCHAPIYKGHGVPPEELV
HTGPDDWWQDVMNGYEFLKNKGYEKIAVAGLSLGGVFSLKLGYTVPIEGIVTMCAPMYIKSE
ETMYEGVLEYAREYKKREGKSEEQIEQEMEKFKQTPMKTLKALQELIADVRDHLDLIYAPTF
VVQARHDEMINPDSANIIYNEIESPVKQIKWYEQSGHVITLDQEKDQLHEDIYAFLESLDW
11111000000000000000000000000000000000000000000000000000000000
00000000000000000000000000000000000000000000000000000000000000
00000000000000000000000000000000000000000000000000000000000000
0000000000000000000000000000000000000000000000000000000000000
>DM_validata473
APVTSPAWADDPPATVYRYDSRPPEDVFQNGFTAWGNNDNVLEHLTGRSCQVGSSNSAFVST
SSSRRYTEVYLEHRMQEAVEAERAGRGTGHFIGYIYEVRADNNFYGAASSYFEYVDTYGDNA
GRILAGALATYQSEYLAHRRIPPENIRRVTRVYHNGITGETTTTEYSNARYVSQQTRANPNP
YTSRRSVASIVGTLVRMAPVVGACMARQAESSEAMAAWSERAGEAMVLVYYESIAYSF
11111111110000000000000000000000000000000000000000000000000000
00000000000000000000000000000000000000000000000000000000000000
00000000000000000000000000000000000000000000000000000000000000
0000000000000000000000000000000001111111111000000000000000
>DM_validata474
VTAGAAPRILKHQVVRAELDRMLDGMRIGDPFPAEREIAEQFEVARETVRQALRELLIDGRV
ERRGRTTVVARPKIRQPLGMGSYTEAAKAQGLSAGRILVAWSDLTADEVLAGVLGVDVGAPV
LQLERVLTTDGVRVGLETTKLPAQRYPGLRETFDHEASLYAEIRSRGIAFTRTVDTIDTALP
DAREAALLGADARTPMFLLNRVSYDQDDVAIEQRRSLYRGDRMTFTAVMHAKNSAIVS
11111111111110000000000000000000000000000000000000000000000000
00000000000000000000000000000000000000000000000000000000000000
00000000000000000000000000000000000000000000000000000000000000
0000000000000000000000000000000000000000000000000000111111
>DM_validata475
MSKLELLPAVDVRDGQAVRLVHGESGTETSYGSPLEAALAWQRSGAEWLHLVDLDAAFGTGD

NRALIAEVAQAMDIKVELSGGIRDDDTLAAALATGCTRVNLGTAALETPEWVAKVIAEHGDK
IAVGLDVRGTTLRGRGWTRDGGDLYETLDRLNKEGCARYVVTDIAKDGTLQGPNLELLKNVC
AATDRPVVASGGVSSLDDLRAIAGLVPAGVEGAIVGKALYAKAFTLEEALEATSLEPG
10000000000000000001111111100000000000000000000000000000000000
00000000000000000000000000000000000000000000000000000000000000
00000000000000000000000000000000000000000000111111100000000000
0000000000000000000000000000000000000000000000000000001111
>DM_validata476
AQVPASGKLLVDPRYSTNDQEESILQDIITRFPNVVLMKQTAQLRAMMTIIRDKETPKEEFV
FYADRLIRLLIEEALNELPFQKKEVTTPLDVSYHGVSFYSKICGVSIVRAGESMESGLRAVC
RGVRIGKILIQRDETTAEPKLIYEKLPADIRERWVMLLDPMCATAGSVCKAIEVLLRLGVKE
ERIIFVNILAAPQGIERVFKEYPKVRMVTAAVDICLNSRYYIVPGIGDFGDRYFGTM
11111111111111111110000000000000000000000000000000000000000000
00000000000000000000000000000000000000000000000000000000000000
00000000000001111000000000000000000000000000000000000000000000
000000000000000000000000000000000000000000000000000000000
>DM_validata477
IDFHVHLDLYPDPVAVARACEERQLTVLSVTTTPAAWRGTLALAAGRPHVWTALGFHPEVVS
ERAADLPWFDRYLPETRFVGEVGLDGSPSLRGTWTQQFAVFQHILRRCEDHGGRILSIHSRR
AESEVLNCLEANPRSGTPILHWYSGSVTELRRAISLGCWFSVGPTMVRTQKGAALIRSMPRD
RVLTETDGPFLELDGQAALPWDVKSVVEGLSKIWQIPASEVERIVKENVSRLLGTVR
00000000000000000000000000000000000000000000000000000000000000
00000000000000000000000000000000000000000000000000000000000000
00000000000000000000000000000000000000000000000000000000000000
000000000000000000000000000000000000000000000000000000011
>DM_validata478
MTVQIPKLLFLHGFLQNGKVFSEKSSGIRKLLKKANVQCDYIDAPVLLEKKDLPFEMDDEKW
QATLDADVNRAWFYHSEISHELDISEGLKSVVDHIKANGPYDGIVGLSQGAALSSIITNKIS
ELVPDHPQFKVSVVISGYSFTEPDPEHPGELRITEKFRDSFAVKPDMKTKMIFIYGASDQAV
PSVRSKYLYDIYLKAQNGNKEKVLAYEHPGGHMVPNKKDIIRPIVEQITSSLQEASE
11000000000000000000000000000000000000000000000000000000000000
00000000000000000000000000000000000000000000000000000000000000
00000000000000000000000000000000000000000000000000000000000000
000000000000000000000000000000000000000000000000000001111
>DM_validata479
GMAQNIYDQPDFFAGYSQLGRSIEGLDGAAEWPALRAMLPEVGGLRIVDLGCGFGWFCRWAH
EHGASYVLGLDLSEKMLARARAAGPDTGITYERADLDKLHLPQDSFDLAYSSLALHYVEDVA
RLFRTVHQALSPGGHFVFSTEHPIYMAPARPGWAIDAEGRRTWPIDRYLVEGPRKTDWLAKG
VVKHHRTVGTTLNALIRSGFAIEHVEEFCPTDAQITARPELAEELDRPMFLLVSARR
11111111111111111111111100000000000000000000000000000000000000
00000000000000000000000000000000000000000000000000000000000000
00000000000000000000000000000000000111000000000000000000000000
000000000000000000000000000000000000000000000000000000000
>DM_validata480

MNKIFIYAGVRNHNSKTLEYTKRLSSIISSRNNVDISFRTPFNSELEISNSDSEELFKKGID
RQSNADDGGVIKKELLESDIIIISSPVYLQNVSVDTKNFIERIGGWSHLFRLAGKFVVTLDV
AESNGSDNVSEYLRDIFSYMGGQILHQVSITNSLKDIAEAQLMEATYKIEDVLEGKIKYKTT
DYQERAYQTLKLILENYDSEHFEKMYWEKKRLFEANSLEEWYYVENIKLEHHHHHH
000000000000000000000000000000000000000000000000000000000000
000000000000000000000000000000000000000000000000000000000000
000000000000000000000000000000000000000000000000000000000000
00000000000000000000000000000000000000000000001111111111
>DM_validata481
HPHAMPDMGPDMNKVPWMGDEQIAMLVYPGMTVMDLVGPHCMFGSLMGAKIYIVAKSLDPVT
SDAGLAIVPTATFGTCPRDLTVLFAPGGTDGTLAAASDAETLAFMADRGARAKYITSVCSGS
LILGAAGLLKGYKATSHWSCRDALAGFGAIPTEARVVRDRNRITGAGVTAGLDFGLSMVAEL
RDQTYAECAQLMSEYDPDPPFNAGSMKTAPAHVRTAMIELVAEFTKKADALAGFPK
000000000000000000000000000000000000000000000000000000000000
000000000000000000000000000000000000000000000000000000000000
000000000000000000000000000000000000000000000000000000000000
00000000000000000000000000000000000000000000000000000011
>DM_validata482
MSGLHILAFGAHADDVEIGMAGTIAKYTKQGYEVGICDLTEADLSSNGTIELRKEEAKVAAR
IMGVKTRLNLAMPDRGLYMKEEYIREIVKVIRTYKPKLVFAPYYEDRHPDHANCAKLVEEAI
FSAGIRKYMPELSPHRVESFYNYMINGFHKPNFCIDISEYLSIKVEALEAYESQFSTGSDGV
KTPLTEGYVETVIAREKMFGKEVGVLYAEGFMSKKPVLLHADLLGGCKLGHHHHHH
100000000000000000000000000000000000000000000000000000000000
000000000000000000000000000000000000000000000000000000000000
000000000000000000000000000000000000000000000000000000000000
00000000000000000000000000000000000000000000001111111111
>DM_validata483
MPTKQQLLYEGKAKKIYATDEPDVLWVEYKDSATAFNGEKKATIAGKGRLNNEISSLLFLKL
REAGIANHFIEKLSPTEQLVRRVTIIPLEVVVRNVVAGSLAKRIGLEEGTPLEAPLVEFYYK
NDDLGDPLLLEDHIFILKLASREEVAALKQAALAVNDVLRLHFAERNVRLIDFKLEFGRTAD
GAILLADEISPDTCRLWDAKTNEKLDKDVFRRDLGSLTDAYEVILQRLGGESACTK
000000000000000000000000000000000000000000000000000000000000
000000000000000000000000000000000000000000000000000000000000
000000000000000000000000000000000000000000000000000000000000
00000000000000000000000000000000000000000000000000011111
>DM_validata484
GMRSNRVTQHPPLQLSLPVHLPDDETFTSYYPAAGNDELIGALKSAASGDGVQAIYLWGPVK
SGRTHLIHAACARANELERRSFYIPLGIHASISTALLEGLEQFDLICIDDVDAVAGHPLWEE
AIFDLYNRVAEQKRGSLIVSASASPMEAGFVLPDLVSRMHWGLTYQLQPMMDDEKLAALQRR
AAMRGLQLPEDVGRFLLNRMARDLRTLFDVLDRLDKASMVHQRKLTIPFVKEMLRL
111111111111000000000000000000000000000000000000000000000000
000000000000000000000000000000000000000000000000000000000000
000000000000000000000000000000000000000000000000000000000000
00000000000000000000000000000000000000000000000000000000

>DM_validata485
GMSESCILFFVKYPEPGKVKTRLGEVVGNDKAAMLYRHFVQDMLQGLARLHADLHICYVPGD
ADLPEKFKAWLGPQHMFAAQQGLDLGERMKHAMQKAFDDGYDRVVLMGSDIPDYPCELVQKA
LNDLQHYDAAIGPAFDGGYYLIGFRKDSFCPDVFDGIRWGEADVYQPTVEKMRRARLEVLQL
PDWNDVDTVWDLNVLYRTNKNSSFRRSSTYALLRENDALIRQYDIDLPGMAPVEKE
100000000000000000000000000000000000000000000000000000000000
000000000000000000000000000000000000000000000000000000000000
000000000000000000000000000000000000000000000000000000000000
00000000000000000000000000000000000000000000001111111111
>DM_validata486
MDVAGLTDCGLIRKSNQDAFYIDEKHQRFFIVADGMGGHAGGEEASRLAVDHIRQYLETHLE
DLQHDPVTLLRQAFLAANHAIVEQQRQNSARADMGTTAVVILLDEKGDRAWCAHVGDSRIYR
WRKDQLQQITSDHTWIAQAVQLGSLTIEQARQHPWRHVLSQCLGREDLSQIDIQPIDLEPGD
RLLLCSDGLTEELTDDVISIYLSEPNVQKAAAALVDAAKTHGGRDNVTVVVISV
000000000000000000000000000000000000111000000000000000000000
000000000000000000000000000000000000000000000000000000000000
000000000000000000011111111111111110000000000000000000000000
000000000000000000000000000000000000000000000000000000
>DM_validata487
ESQPDPKPDELHKSSKFTGLMENMKVLYDDNHVSAINVKSIDQFLYFDLIYSIKDTKLGNYD
NVRVEFKNKDLADKYKDKYVDVFGANYYYQCYFSKKTNDINSHQTDKRKTCMYGGVTEHNGN
QLDKYRSITVRVFEDGKNLLSFDVQTNKKKVTAQELDYLTRHYLVKNKKLYEFNNSPYETGY
IKFIENENSFWYDMMPAPGDKFDQSKYLMMYNDNKMVDSKDVKIEVYLTTKKK
000000000000000000000000000000000000000000000000000001111100
000000000000000000000000000000000011111111111000000000000000
000000000000000000000000000000000000000000000000000000000000
00000000000000000000000000000000000000000000000000111
>DM_validata488
ASKGEELFTGVVPILVELDGDVNGHKFSVSGEGEGDATYGKLTLKFICTTGKLPVPWPTLVT
TFSYGVQCFSRYPDHMKRHDFFKSAMPEGYVQERTIFFKDDGNYKTRAEVKFEGDTLVNRIE
LKGIDFKEDGNILGHKLEYNYNSHNVYIMADKQKNGIKVNFKIRHNIEDGSVQLADHYQQNT
PIGDGPVLLPDNHYLSTQSALSKDPNEKRDHMVLLEFVTAAGITHGMDELYK
000000000000000000000000000000000000000000000000000000000000
000000000000000000000000000000000000000000000000000000000000
000000000000000000000000000000000000000000000000000000000000
0000000000000000000000000000000000000000000011111111
>DM_validata489
MTGSNSHITILTLNINGLNSAIKRHRLASWIKSQDPSVCCIQETHLTCRDTHRLKIKGWRKI
YQANGKQKKAGVAILVSDKTDFKPTKIKRDKEGHYIMVKGSIQQEELTILNIYAPNTGAPRF
IKQVLSDLQRDLDSHTLIMGDFNTPLSTLDRSTRQKVNKDTQELNSALHQADLIDIYRTLHP
KSTEYTFFSAPHHTYSKIDHIVGSKALLSKCKRTEIITNYLSDHSAIKLELR
111110000000000000000000000000000000000000000000000000000000
000000000000000000000000000000000000000000000000000000000000
000000000000000000000000000000000000000000000000000000000000

0000000000000000000000000000000000000000000000000000
>DM_validata490
PEATSCETEGSSISFTVEKAGHVVRFNCPSTLEEIKPAYEAGDSTKVCTTADCSNEAALKDV
LKSASLAQAEGSGPSGGNDFTLTVDALPEAETSVFFLCQRTGASRSARRLGTAVPSDKCGVH
ILVKAAPQAPVCSAQDHTLELQITAANSDTSFVCGGTFNVIKPANAAKVLQGDSCETEVDLV
SLVPHASRSALEQSGLIKLSVTDLPQQQQKLCYRCEDSSQKACKVLVTVSAS
111000000000000000000000000000000000000000000000000000000000
000000000001110000000000000000000000000000111111110000000000
000000000000000000000000000000000000000000000000000000000000
0000000000000000000000000000000000000000000000000000
>DM_validata491
MVDNRMRFTIDQNMQFPLVEIDLEHGGSVYLQQGSMVYHTENVTLNTKLNGKGSGLGKLVGA
IGRSMVSGESMFITQAMSNGDGKLALAPNTPGQIVALELGEKQYRLNDGAFLALDGSAQYKM
ERQNIGKALFGGQGGLFVMTTEGLGTLLANSFGSIKKITLDGGTMTIDNAHVVAWSRELDYD
IHLENGFMQSIGTGEGVVNTFRGHGEIYIQSLNLEQFAGTLKRYLPTSSN
111100000000000000000000000000000000000000000001111000000000
000000000000000000000000000000000000000000000000000000000000
000000111111000000000000000000000000000000000000000000000000
00000000000000000000000000000000000000000000011111
>DM_validata492
KECTIVPSNHYGPIPGIPVGTMWRFRVQVSESGVHRPHVAGIHGRSNDGAYSLVLAGGYEDD
VDHGNFFTYTGSGGRDLSGNKRTAEQSCDQKLTNTNRALALNCFAPINDQEGAEAKDWRSGK
PVRVVRNVKGGKNSKYAPAEGNRYDGIYKVVKYWPEKGKSGFLVWRYLLRRDDDEPGPWTKE
GKDRIKKLGLTMQYPEGYLEALANREREKENSKREEEEQQEGGFASPRTG
100000000000000000000000000000000000000000000000000000000000
000000000000001111111111000000000000000000000000000000000000
000000000000000000000000000000000000000000000000000000000000
00000000000000000000001111111111111111111111111111
>DM_validata493
AQPDIKFASKEYGVTIGESRIIYPLDAAGVMVSVKNTQDYPVLIQSRIYDENKEKESEDPFV
VTPPLFRLDAKQQNSLRIAQAGGVFPRDKESLKWLCVKGIPPKDEDIWVDDATNKQKFNPDK
DVGVFVQFAINNCIKLLVRPNELKGTPIQFAENLSWKVDGGKLIAENPSPFYMNIGELTFGG
KSIPSHYIPPKSTWAFDLPKGLAGARNVSWRIINDQGGLDRLYSKNVTL
111111000000000000000000000000000000000000000001111111110000
000000000000000000000000000000000000000111111111111111111110
000000000000000000000000000000000000000000000000000000000000
0000000000000000000111111100000000000000000000000
>DM_validata494
MKTKEDMQELYFPTPKLIEWENGVRQYSTVRGDTEVLMSYVPPHTNVEPHQHKEVQIGMVVS
GELMMTVGDVTRKMTALESAYIAPPHVPHGARNDTDQEVIAIDIKRLKADETYTSPEDYFLD
IFKTRDLLPGMEVTFFVEDWVEIMLAKIPGNGGEMPFHKHRNEQIGICIGGGYDMTVEGCTV
EMKFGTAYFCEPREDHGAINRSEKESKSINIFFPPRYNRAKAKKMKADE
100000000000000000000000000000000000000000000000000000000000
000000000000000000000000000000000000000000000000000000000000

000000000000000000000000000000000000000000000000000000000000
0000000000000000000000000000000000000001111111111
>DM_validata495
GHMAQSETVERILDAAEQLFAEKGFAETSLRLITSKAGVNLAAVNYHFGSKKALIQAVFSRF
LGPFCASLEKELDRRQAKPEAQHATLEDLLHLLVSQAMAVKPRSGNDLSIFMRLLGLAFSQS
QGHLRKYLEEVYGKVFRRYMLLVNEAAPKLPPIELFWRVHFMLGAAAFSMSGIKALRAMAET
DFGVNTSTEQVMHLMVPFFAAGMRAESGIDDPLLAGAQLRPRNKTPAKA
111000000000000000000000000000000000000000000000000000000000
000000000000000000000000000000000000000000000000000000000000
000000000000000000000000000000000000000000000000000000000000
0000000000000000000000000000001111111111111111111
>DM_validata496
DHTLRYPVKPEEMDWSELYPEFFAPLTQNQSHDDPKDKKEKRAQAQVEFADIGCGYGGLLVE
LSPLFPDTLILGLEIRVKVSDYVQDRIRALRAAPAGGFQNIACLRSNAMKHLPNFFYKGQLT
KMFFLFPDPHFKRTKHKWRIISPTLLAEYAYVLRVGGLVYTITDVLELHDWMCTHFEEHPLF
ERVPLEDLSEDPVVGHLGTSTEEGKKVLRNGGKNFPAIFRRIQDPVLQ
111110000000000000000011111111111111111111000000000000000000
000000000000000000000000000000000000000000000000000000000000
000000001111111111100000000000000000000000000000000000000000
000000000000000000000000000000000000000000000000
>DM_validata497
MPDKPTKEKLMEQLKGGIIVSCQALPGEPLYSETGGIMPLMAKAAQEAGAVGIRANSVRDIK
EIQAITDLPIIGIIKKDYPPQEPFITATMTEVDQLAALNIAVIAMDCTKRDRHDGLDIASFI
RQVKEKYPNQLLMADISTFDEGLVAHQAGIDFVGTTLSGYTPYSRQEAGPDVALIEALCKAG
IAVIAEGKIHSPEEAKKINDLGVAGIVVGGAITRPKEIAERFIEALKS
111000000000000000000000000000000000000000000000000000000000
000000000000000000000000000000000000000000000000000000000000
000000000000000000000000000000000000000000000000000000000000
000000000000000000000000000000000000000000000001
>DM_validata498
DYKDDDDKQSFSSSSSCTEEENKHHMGIDVIIKVTKQDQTPTNDKICQSVTEVTESEDESEE
VVKGDPTTYYTVVGGGLTMDFGFTKCPKISSISEYSDGNTVNARLSSVSPGQGKDSPAITRE
EALSMIKDCEMSINIKCSEEEKDSNIKTHPVLGSNISHKKVSYEDIIGSTIVDTKCVKNLEI
SVRIGDMCKESSELEVKDGFKYVDGSASEDAADDTSLINSAKLIACV
111111111000000000000000000000000000000000000000000000000000
000000000000000000000000000000000000000000000000000000000000
000000000000000000000000000000000000000000000000000000000000
00000000000000000000000000000000000000000000000
>DM_validata499
MPSTDSIPKSNFDAIPDVIQAFKNGEFVVVLDDPSRENEADLIIAAESVTTEQMAFMVRHSS
GLICAPLTPERTTALDLPQMVTHNADPRGTAYTVSVDAEHPSTTTGISAHDRALACRMLAAP
DAQPSHFRRPGHVFPLRAVAGGVRARRGHTEAGVELCRLAGKRPVAVISEIVDDGQEVEGRA
VRAAPGMLRGDECVAFARRWGLKVCTIEDMIAHVEKTEGKLETNGSG
111111111100000000000000000000000000000000000000000000000000

000000000000000000000111111100000000000000000000000000000000000
00000000000000000000000000000000000000000000000000000000000000
00000000000000000000000000000000000000011111111
>DM_validata500
GKNVMVEPHRHEGVFICRGKEDALVTKNLVPGESVYGEKRVSISEGDDKIEYRAWNPFRSKL
AAAILGGVDQIHIKPGAKVLYLGAASGTTVSHVSDIVGPDGLVYAVEFSHRSGRDLINLAKK
RTNIIPVIEDARHPHKYRMLIAMVDVIFADVAQPDQTRIVALNAHTFLRNGGHFVISIKANC
IDSTASAEAVFASEVKKMQQENMKPQEQLTLEPYERDHAVVVGVYRP
11110000000000000011111000000000000000000000111000000000000000
00000000000000000000000000000000000000000000000000000000000000
00000000000000000000000000000000000000000000000000000000000000
00000000000000000000000000000000000000000000000
>DM_validata501
MTKNPRNNKPKKILDSSYKSKTIWQNYIDALFETFPQLEISEVWAKWDGGNVTKDGGDAKLT
ANIRTGEHFLKAREAHIVDPNSDIYNTILYPKTGADLPCFGMDLMKFSDKKVIIVFDFQHPR
EKYLFSVDGLPEDDGKYRFFEMGNHFSKNIFVRYCKPDEVDQYLDTFKLYLTKYKEMIDNNK
PVGEDTTVYSDFDTYMTELDPVRGYMKNKFGEGRSEAFVNDFLFSYK
11111111111111111111100000000000000000000000000000011111000000
00000000000000000000000000000000000000000000000000000000000000
00000000000000000000000000000000000000000000000000000000000000
00000000000000000001100000000000000000000000000
>DM_validata502
MADPSLNNPVVIQATRLDASILPRNVFSKSYLLYVIAQGTDVGAIAGKANEAGQGAYDAQVK
NDEQDVELADHEARIKQLRIDVDDHESRITANTKAITALNVRVTTAEGEIASLQTNVSALDG
RVTTAENNISALQADYVSKTATTSQSLASPLNVTTSYSVGGKKVVGARQTGWTAATGTANKG
VFDADLTFAVSDTYTQSEIQAIANALITERRRTKAMEDALRAHGLID
00000000000000000000000000000000000000000000000000000000000000
00000000000000000000000000000000000000000000000000000000000000
00000000000000000000000000000000000000000000000000000000000000
00000000111111111111000000000000000000000000000
>DM_validata503
TTLSCKVTSVEAITDTVYRVRIVPDAAFSFRAGQYLMVVMDERDKRPFSMASTPDEKGFIEL
HIGASEINLYAKAVMDRILKDHQIVVDIPHGEAWLRDDEERPMILIAGGTGFSYARSILLTA
LARNPNRDITIYWGGREEQHLYDLCELEALSLKHPGLQVVPVVEQPEAGWRGRTGTVLTAVL
QDHGTLAEHDIYIAGRFEMAKIARDLFCSERNAREDRLFGDAFAFI
00000000000000000000000000000000000000000000000000000000000000
00011111100000000000000000000000000000000000000000000000000000
00000000000000000000000000000000000000000000000000000000000000
0000000000000000000000000000000000000000000000
>DM_validata504
QVKPFSFVRDTDPVQLVNFLQSEHPQTIAVVLSYLDPPVAAQILGALPEELQTEVLKRIALL
ERTSPEVVKEIERNLEKKISGFVSRTFSKVGGIDTAAEIMNNLDRTTEKKIMDKLVQENPEL
ADEIRRRMFVFEDILKLDDRSIQLVLREVDTRDLALALKGASDELKEKIFKNMSKRAAALLK
DELEYMGPVRLKDVEEAQQKIINIIRRLEEAGEIVIARGGGEELIM

11111111111000000000000000000000000000000000000000000000000000
00000000000000000000000000000000000000000000000000000000000000
00000000000000000000000000000000000000000000000000000000000000
0000000000000000000000000000000000000011111111
>DM_validata505
MGSDKIHHHHHHMEKTVIYLLEDGYVDFVVEKIRTKMEKLLEEKDKIFVVLAGGRTPLPVYE
KLAEQKFPWNRIHFFLSDERYVPLDSDQSNFRNINEVLFSRAKIPSGNVHYVDTSLPIEKAC
EKYEREIRSATDQFDLAILGMGPDGHVASIFDLETGNKDNLVTFTDPSGDPKVPRVTLTFRA
LNTSLYVLFLIRGKEKINRLTEILKDTPLPAYFVRGKEKTVWFVGK
11111111111111000000000000000000000000000000000000000000000000
00000000000000000000000000000000000000000000000000000000000000
00000000000000000000000000000000000000000000000000000000000000
0000000000000000000000000000000000000000000000
>DM_validata506
GAMAGSIRSKLSAIDVRQLGTVDYRTAWQLQRELADARVAGGADTLLLLEHPAVYTAGRRTE
THERPIDGTPVVDTDRGGKITWHGPGQLVGYPIIGLAEPLDVVNYVRRLEESLIQVCADLGL
HAGRVDGRSGVWLPGRPARKVAAIGVRVSRATTLHGFALNCDCDLAAFTAIVPCGISDAAVT
SLSAELGRTVTVDEVRATVAAAVCAALDGVLPVGDRVPSHAVPSPL
00000000000000000000000000000000000000000000000000000000000000
00000000000000000000000000000000000000000000000000000000000000
00000000000000000000000000000000000000000000000000000000000000
0000000000000000000000000000000011111111111111
>DM_validata507
SNAMKILVTSGGTSEAIDSVRSITNHSTGHLGKIITETLLSAGYEVCLITTKRALKPEPHPN
LSIREITNTKDLLIEMQERVQDYQVLIHSMAVSDYTPVYMTGLEEVQASSNLKEFLSKQNHQ
AKISSTDEVQVLFLKKTPKIISLVKEWNPTIHLIGFKLLVDVTEDHLVDIARKSLIKNQADL
IIANDLTQISADQHRAIFVEKNQLQTVQTKEEIAELLLEKIQAYHS
10000000000000000000000000000000000000000000000000000000000000
00000000000000000000000000000000000000000000000000000000011111
11111100000000000010000000000000000000000000000000000000000000
0000000000000000000000000000000000000000000000
>DM_validata508
LALVATGNDTTTKPDLYYLKNSEAINSLALLPPPPAVGSIAFLNDQAMYEQGRLLRNTERGK
LAAEDANLSSGDVANAFSGAFGSPITEKDAPALHKLLTNMIEDAGDLATRSAKDHYMRIRPF
AFYGVSTCNTTEQDKLSKNGSYPSGHTSTGWATALVLAEINPQRQNEILKRGYELGQSRVIC
GYHWQSDVDAARVVGSAVVATLHTNPAFQQQLQKAKAEFAQHQKK
11111100000000000000000000000000000000000000000000000000000000
00000000000000000000000000000000000000000000000000000000000000
00000000011111110000000000000000000000000000000000000000000000
000000000000000000000000000000000000000001111
>DM_validata509
MDVNLYGPGGPHTALKDIANKYSEKTGVKVNVNFGPQATWFEKAKKDADILFGASDQSALAI
ASDFGKDFNVSKIKPLYFREAIILTQKGNPLKIKGLKDLANKKVRIVVPEGAGKSNTSGTGV
WEDMIGRTQDIKTIQNFRNNIVAFVPNSGSARKLFAQDQADAWITWIDWSKSNPDIGTAVAI

EKDLVVYRTFNVIAKEGASKETQDFIAYLSSKEAKEIFKKYGWRE
10000000000000000000000000000000000000000000000000000000000000
00000000000000000000000000000000000000000000000000000000000000
00000000000000000000000000000001100000000000000000000000000000
000000000000000000000000000000000000000000000
>DM_validata510
GMNSPKNSLYEEKSGHYYNAVNPNLLKHIKKEWKEVLDIGCSSGALGAAIKENGTRVSGIEA
FPEAAEQAKEKLDHVVLGDIETMDMPYEEEQFDCVIFGDVLEHLFDPWAVIEKVKPYIKQNG
VILASIPNVSHISVLAPLLAGNWTYTEYGLLDKTHIRFFTFNEMLRMFLKAGYSISKVDRVY
VDHKMYEPLIEELYGICKKYRLGSGFMAETVVFQYIIEAEKSQL
11111111111111111110000000000000000000000000000000000000000000
00000000000000000000000000000000000000000000000000000000000000
00000000000000000000000000000000000000000000000000000000000000
00000000000000000000000000000000000000000000
>DM_validata512
MTNIEILENMLKLQQKLNDETNGLNWENGYTKEGKLISWRRCIYMECAELIDSFTWKHWKNI
SSLTNWENVRIEIVDIWHFILSLLLEEYRDKNNKDFKAIATEVNAVSVFQDFCKEEEYPNEG
DIYGILNDIELIIHKCSGFGFNLGELLSTYFTLAIKCGLNLEILYKTYIGKNVLNIFRQNNG
YKDGSYKKTWNGKEDNEVLAQILEQELDFDTIYKKLEECYKKA
00000000000000000000000000000000000000000000000000000000000000
00000000000000000000000000111111110000000000000000000011111110
00000000000000000000000000000000000000000000000000000000000000
0000000000000000000000000111110000000000000
>DM_validata513
MPAAAPGKNFDLSHWKLQLPDANTTEISSANLGLGYTSQYFYTDTDGAMTFWAPTTGGTTAN
SSYPRSELREMLDPSNSKVNWGWQGTHTMKLSGKTVQLPSSGKIIVAQIHGIMDDGTNAPPL
VKAVFQDGQLDMQVKQNSDGTGSDVHNYFTGIKLGDLYNMEIRVTDGVAYVTMNGDTRSVDF
VGKDAGWKNLKYYFKAGNYVQDNTSTGGSAIAKLYSLSVSHSN
11100000000000000000000000000000000000000000000000000000000000
00000000000000000000000000000000000000000000000000000000000000
00000000000000000000000000000000000000000000000000000000000000
0000000000000000000000000000000000000000000
>DM_validata514
MGSSGSPVRASIEPLTWENAFFGVNSAIVRITSEAPLLTPDALAPWSRVQAKIAASNTGELD
ALQQLGFSLVEGEVDLALPVNNVSDSGAVVAQETDIPALRQLASAAFAQSRFRAPWYAPDAS
GRFYAQWIENAVRGTFDHQCLILRAASGDIRGYVSLRELNATDARIGLLAGRGAGAELMQTA
LNWAYARGKTTLRVATQMGNTAALKRYIQSGANVESTAYWLYR
11111110000000000000000000000000000000000000000000000000000000
00000000000000000000000000000000000000000000000000000000000000
00000000000000000000000000000000000000000000000000000000000000
0000000000000000000000000000000000000000000
>DM_validata515
GSSGSSGMCGRFAQSQTREDYLALLAEDIERDIPYDPEPIGRYNVAPGTKVLLLSERDEHLH
LDPVFWGYAPGWWDKPPLINARVETAATSRMFKPLWQHGRAICFADGWFEWKKEGDKKQPFF

IYRADGQPIFMAAIGSTPFERGDEAEGFLIVTAAADQGLVDIHDRRPLVLSPEAAREWMRQE
ISGKEASEIAASGCVPANQFSWHPVSRAVGNVKNQGAELIQPV
11111111100000000000000000000000000000000000000000000000000000
00000000000111100000000000000000000000000000000000001111110000
00000000000000000000000000000000000000000000000000000000000000
0000000000000000000000000000000000000000000
>DM_validata516
RQNWEEADSMKSSVESLQNRVTELESVDKSAGQVARNTGLLESQLSRHDQMLSVHDIRLADM
DLRFQVLETASYNGVLIWKIRDYKRRKQEAVMGKTLSLYSQPFYTGYFGYKMCARVYLNGDG
MGKGTHLSLFFVIMRGEYDALLPWPFKQKVTLMLMDQGSSRRHLGDAFKPDPNSSSFKKPTG
EMNIASGCPVFVAQTVLENGTYIKDDTIFIKVIVDTSDLPDP
11111111111111111111111000000000000000000000000000000000000000
00000000000000000000000000000000000000000000000000000000000000
00000000000000000000000000000000000000000000000000000000000000
000000000000000000000000000000000000000000
>DM_validata518
DYKDDDDKLHSQANLMRLKSDLFNRSPMYPGPTKDDPLTVTLGFTLQDIVKADSSTNEVDLV
YYEQQRWKLNSLMWDPNEYGNITDFRTSAADIWTPDITAYSSTRPVQVLSPQIAVVTHDGSV
MFIPAQRLSFMCDPTGVDSEEGATCAVKFGSWVYSGFEIDLKTDTDQVDLSSYYASSKYEIL
SATQTRQVQHYSCCPEPYIDVNLVVKFRERRAGNGFFRNLFD
11111100000000000000000000110000000000000000000000000000000000
00000000000000000000000000000000000000000000000000000000000000
00000000000000000000000000000000000000000000000000000000000000
000000000011110000000000000000011111111111
>DM_validata519
MIERGKFRSLTLINWNGFFARTFDLDELVTTLSGGNGAGKSTTMAAFVTALIPDLTLLHFRN
TTEAGATSGSRDKGLHGKLKAGVCYSMLDTINSRHQRVVVGVRLQQVAGRDRKVDIKPFAIQ
GLPMSVQPTQLVTETLNERQARVLPLNELKDKLEAMEGVQFKQFNSITDYHSLMFDLGIIAR
RLRSASDRSKFYRLIEASLYGGISSAITRSLRDYLLPENSG
11100000000000000000000000000000000000000000000000000000001111
11111111111111000000000000000000000000000000000000000000000000
00000000000000000000000000000000000000000000000000000000000000
00000000000000000000000000000000000000111
>DM_validata520
MCSHYQALKDQERMRKYFAAHPSAEVPADMWPRYMGAFIRRPLEWDSGDEAVPEREAATGRW
GMIPPGTRPEKLAEASKKNTSNARSETAHQLWTFRNAWAKAQHCIIPADAIYEPDWRSGKAV
PTRFTRADGAPLGIAGLWDRYRNAAGEWIDSYTMLTINADDDPLFRDYHQAGKEKRMVVILP
DGAYGDWLTAPATDTRDFLLPYPADRLVAAAVKLEHHHHHH
10000000000000000000000000000000000000000001111111100000000000
00000000000000000000000000000000000000000000000000000000000000
00000000000000000000000000000000000000000000000000000000000000
00000000000000000000000000000000001111111
>DM_validata521
MTAIAPVITIDGPSGAGKGTLCKAMAEALQWHLLDSGAIYRVLALAALHHHVDVASEDALVP

LASHLDVRFVSTNGNLEVILEGEDVSGEIRTQEVANAASQVAAFPRVREALLRRQRAFRELP
GLIADGRDMGTVVFPDAPVKIFLDASSEERAHRRMLQLQEKGFSVNFERLLAEIKERDDRDR
NMAVAPLVPAADALVLDSTTLSIEQVIEKALQYARQKLALA
110000000000000000000000000000000000000000000000000000000000000
000000000000000000000000000000000000000000000000000000000000000
000000000000000000000000000000000000000000000000000000001111111
11111100000000000000000000000000000000000
>DM_validata522
GMPVNSILGVFAKSPIKPLQEHMDKVYDCASLLVPFFEATITGNWDDAVQIRKQISLAEKQG
DSLKREIRLTLPSGLFMPVERTDLLELLTQQDKIANKAKDISGRVIGRQLLIPQALQVPFIA
YLQRCIDAVGLAQQVINELDDLLEAGFRGREVDFVAKMINELDIIEEDTDDLQIQLRRQLFA
LESELNPVDVMFLYKTIEWVGGLADLAERVGSRLELMLARV
111111111111110000000000000000000000000000000000000000000000000
000000000000000000000000000000000000000000000000000000000000000
000000000000000000000000001100000000000000000000000000000000000
00000000000000000000000000000000000000000
>DM_validata523
GSHMLTKQQIRYCLDEMAKMFPDAHCELVHRNPFELLIAVVLSAQCTDALVNKVTKRLFEKY
RTPHDYIAVPLEELEQDIRSIGLYRNKARNIQKLCAMLIDKYNGEVPRDRDELMKLPGVGRK
TANVVVSVAFGVPAIAVDTHVERVSKRLGFCRWDDSVLEVEKTLMKIIPKEEWSITHHRMIF
FGRYHCKAQSPQCPSCPLLHLCREGKKRMRKREEKAANQK
111000000000000000000000000000000000000000000000000000000000000
000000000000000000000000000000000000000000000000000000000000000
000000000000000000000000000000000000000000000000000000000000000
0000000000000000000000000001111111111
>DM_validata524
QQKILENIRGIGTNTMTIFNGNGFGDRRSRHIQNLKISDANTLSKQSYIQSVTPNTSSSGIL
VVGNKSFTSANLYGIGEQYFDVEGLKLKQGRLLTEDDVDQSNQVVVLDESAKKAIFANENPL
GKTVIFNKRPFRVIGVVSDQQLGGFPGNSLNLYSPYSTVLNKITGGSRIGSITVKISDDVNS
TVAEKSLTELLKSLHGKKDFFIMNSDTIKQTIENTTGTMK
100000000000000000000000000000000000000000000000000000000000000
000000000000000000000000000000000000000000000000000000000000000
000000000000000000001111111100000000000000000000000000000000000
0000000000000000000000000000000000000111
>DM_validata525
MQLLSGIVQQQNNLLRAIEAQQHLLQLTVWGIKQLQARILAGGSGGHTTWMEWDREINNYTS
LIHSLIEESQNQQEKNEQELLEGSSGGQLLSGIVQQQNNLLRAIEAQQHLLQLTVWGIKQLQ
ARILAGGSGGHTTWMEWDREINNYTSLIHSLIEESQNQQEKNEQELLEGSSGGQLLSGIVQQ
QNNLLRAIEAQQHLLQLTVWGIKQLQARILAGGHHHHHHG
000000000000000000000000000000000000000001111110000000000000000
000000000000000000000011110000000000000000000000000000000000000
000011111111110000000000000000000000000000000001100000000000000
0000000000000000000000000001111111111
>DM_validata527

MSLTFSILAHDPETGAIGGAAATGSLCVGGWVLRGDLNAGMSASQGAAPSTFWGEEVLQHLR
DGSHPEDAVNHVTSQDSGRAYRQLAAMDLLGNAAAFTGSENQDIKGSVTFASGIASGNMLGD
NSVLGAMTEAFVASDLTFERRLLAALIAAEGAGSDFRGLLSAAMLVLHPDRPPVTLRIDYHP
DNPIGALEQLYQKATTGDYADWARQVPVLSDKERILDEG
10000000000000000000000000000000000000000000000000000000000000
00000000000000000000000000000000000000000000000000000000000000
00000000000000000000000000000000111110000000000000000000000000
00000000000000000000000000000000000011
>DM_validata529
MTEIYTSVLSYRLLEGKAYSDADTRSLDRMMRSIDEFFSANPGYINFHIYRSYRTDSDVIFW
YSSRNPDLMILAKERVQASMRPIAVSSFSSISIYDESPYNAMNKKLEDSLRLPPLRYFVAYP
MSKTPDWYLLDFDTRKEIMHEHIKMALNHPDEKGIRSYTTYSFGIGDQEFVVLYEIPDIAAW
SRVTEKLREARARKWIIKETPILLGRLVDAGDIAGFLL
10000000000000000000000000000000000000000000000000000000000000
00000000000000000000000000000000000000000000000000000000000000
00000000000000000000000001111111100000000000000000000000000000
00000000000000000000000000000000000000
>DM_validata530
MSKEKRMPFFKKTEHTPPYDVVPSMRPVVLVGPSLKGYEVTDMMQKALFDFLKHRFEGRISI
TRVTADISLAKRSVLNNPSKHAIIERSNTRSSLAEVQSEIERIFELARTLQLVVLDADTINH
PAQLSKTSLAPIIVYVKISSPKVLQRLIKSRGKSQAKHLNVQMVAADKLAQCPPQESFDVIL
DENQLEDACEHLADYLEAYWKATHPPSSNLPNPLLSRT
11111000011111100000000000000000000000000000000000000000000000
00000000001111111111100000000000000000000000000000000000000000
00000000000000000000000000000001111100000000000000000000000000
00000000000000000000000000111111111100
>DM_validata531
GSSCGNDESDPDSTVTIAMATVEKQPQYDAPYLVLDNGEKLWVVQHIVPYRDLKAGERIFGN
YSFLEAGESGFAYNIRLNDYTLVPVQKIIGLNPDNMDSIGNMKVQIKDMWPSDDYLNVRFML
NFPSPQKPILNLVVNEMIPWTKDGYAHLELRYNNNGSQGRLVPGMVSFKLDDYSPENSELKG
IKVLVNPVDGEEKTYIFSYPLTGEDVPGFNPLDLAELK
11111111111110000000000000000000000000000000000000000000000000
00000000000000000000000000000000000000000000000000000000000000
00000000000000000000000000000000000000000000000000000000000000
00000000000000000000000000000000000000
>DM_validata532
MSLSYTTQQIIEKLRELKIVPVIALDNADDILPLADTLAKNGLSVAEITFRSEAAADAIRLL
RANRPDFLIAAGTVLTAEQVVLAKSSGADFVVTPGLNPKIVKLCQDLNFPITPGVNNPMAIE
IALEMGISAVKFFPAEASGGVKMIKALLGPYAQLQIMPTGGIGLHNIRDYLAIPNIVACGGS
WFVEKKLIQSNNWDEIGRLVREVIDIIKEGGSHHHHHH
11100000000000000000000000000000000000000000000000000000000000
00000000000000000000000000000000000000000000000000000000000000
00000000000000000000000000000000000000000000000000000000000000
00000000000000000000000000000111111111

>DM_validata534
GFGELKEEWRGEVVHLSWSPRAFLLKNFLSDEECDYIVEKARPKMVKSSVVDNESGKSVDSE
IRTSTGTWFAKGEDSVISKIEKRVAQVTMIPLENHEGLQVLHYHDGQKYEPHYDYFHDPVNA
GPEHGGQRVVTMLMYLTTVEEGGETVLPNAEQKVTGDGWSECAKRGLAVKPIKGDALMFYSL
KPDGSNDPASLHGSCPTLKGDKWSATKWIHVAPIGGRH
111111000000000000000000000000000000000000011111111111111111
110000000000000000000000000000000000000000110000000000000000
000000000000000000000000000000000000000000000000000000000000
00000000000000000000000000000000000011
>DM_validata535
ATETSFIIDAFNKTNLILQGDATVSSNGNLQLSYNSYDSMSRAFYSAPIQIRDSTTGNVASF
DTNFTMNIRTHRQANSAVGLDFVLVPVQPESKGDTVTVEFDTFLSRISIDVNNNDIKSVPWD
VHDYDGQNAEVRITYNSSTKVFSVSLSNPSTGKSNNVSTTVELEKEVYDWVSVGFSATSGAY
QWSYETHDVLSWSFSSKFINLKDQKSERSNIVLNKIL
000000000000000000000000000000000000000000000000000000000000
000000000000111000000000000111111000000000000000000000000000
000000000000000000000000000000000000000000000000000000000000
0000000000000000001111111111111111111
>DM_validata536
GHMLRGLRIIAENKIGVLRDLTTIIAEEGGNITFAQTFLIKHGEHEGKALIYFEIEGGDFEK
ILERVKTFDYIIEIEEEESFERVFGKRVIILGGGALVSQVAIGAISEADRHNLRGERISVDT
MPVVGEEEIAEAVKAVSRLHRAEVLVLAGGIMGGKITEEVKKLRKSGIRVISLSMFGSVPDV
ADVVISDPVMAGTLAVMHISEKAKFDLDRVKGRRIGK
111111000000000000000000000111000000000000000000000011111111
000000000000000000000000000000000000000000000000000000000000
000000000000000000000000000000000000000000000000000000000000
0000000000000000000000000000000000000
>DM_validata537
GMIGDGIENSPLLTDLAFPYRLLGAGKESRECLFLLHGSGVDETTLVPLARRIAPTATLVAA
RGRIPQEDGFRWFERIDPTRFEQKSILAETAAFAAFTNEAAKRHGLNLDHATFLGYSNGANL
VSSLMLLHPGIVRLAALLRPMPVLDHVPATDLAGIRTLIIAGAADETYGPFVPALVTLLSRH
GAEVDARIIPSGHDIGDPDAAIVRQWLAGPIAIAQAD
111111100000000000000000000000000000000000000000000000000000
000000000000000000000000000000000000000000000000000000000000
000000000000000000000000000000000000000000000000000000000000
0000000000000000000000000000001111111
>DM_validata540
TNAKASKRVYITFTGYDKKPSIDNLKKLDMSITSNPSKCTHLIAPRILRTSKFLCSIPYGPC
VVTMDWINSCLKTHEIVDEEPYLLNDPEKELELGCTLESALKRARAQGPSLLEDYVVYLTSK
TVAPENVPAVISIVKSNGGVCSTLNVYNKRLARHLEDGNVVLITCNEDSHIWTNFLDNASQN
KTIFLQNYDWLIKTVLRQEIDVNDRIADEFARAV
111111000000000000000000000000000000000000000000000000000000
000000000000000000000000000000000000000000000000000000000000
000000000000000000000000000000000000000000000000000000000000

0000000000000000000000000000000000
>DM_validata541
MGSDKIHHHHHHMKITWFGHACFALEMEGKTIVTDPFDESVGYPIPNVTADVVTESHQHFDH
NAHHLVKGNFRVIDRPGAYTVNGVKIKGVETFHDPSHGRERGKNIVFVFEGEGIKVCHLGDL
GHVLTPAQVEEIGEIDVLLVPVGGTYTIGPKEAKEVADLLNAKVIIPMHYKTKYLKFNLLPV
DDFLKLFDSYERVGNILELFEKPKERKVVVMEVQ
11111111111000000000000000000000000001111100000000000000001111
00000000000000000000000000000000111111111100000000000000000000
00000000000000000000000000000000000000000000000000000000000000
0000000000000000000000000000000001
>DM_validata542
MEKKITGYTTVDISQWHRKEHFEAFQSVAQCTYNQTVQLDITAFLKTVKKNKHKFYPAFIHI
LARLMNAHPEFRMAMKDGELVIWDSVHPCYTVFHEQTETFSSLWSEYHDDFRQFLHIYSQDV
ACYGENLAYFPKGFIENMFFVSANPWVSFTSFDLNVANMDNFFAPVFTMGKYYTQGDKVLMP
LAIQVHHAVCDGFHVGRMLNELQQYCDEWQGGA
11110000000000000000000000000000000000000000000000000000000000
00000000000000000000000000000000000000000000000000000000000000
00000000000000000000000000000000000000000000000000000000000000
000000000000000000000000000000000
>DM_validata543
SGTTNTVAAYNLTWKSTNFKTILEWEPKPVNQVYTVQISTKSGDWKSKCFYTTDTECDLTDE
IVKDVKQTYLARVFSYPAGNVESTGSAGEPLYENSPEFTPYLETNLGQPTIQSFEQVGTKVN
VTVEDERTLVRRNNTFLSLRDVFGKDLIYTLYYWKSSSSGKKTAKTNTNEFLIDVDKGENYC
FSVQAVIPSRTVNRKSTDSPVECMGQEKGEFRE
11110000000000000000000000000000000000000000000000000000000000
00000000000000000000011111110000000000000000000000000111110000
00000000001110000000000000000000001111110000000000000000000111 0
000000000000000000000000111111111
>DM_validata544
SSSVPSQKTYQGSYGFRLGFLHSGTAKSVTCTYSPALNKMFCQLAKTCPVQLWVDSTPPPGT
RVRAMAIYKQSQHMTEVVRRCPHHERCSDSDGLAPPQHLIRVEGNLRVEYLDDRNTFRHSVV
VPYEPPEVGSDCTTIHYNYMCNSSCMGGMNRRPILTIITLEDSSGNLLGRNSFEVRVCACPG
RDRRTEEENLRKKGEPHHELPPGSTKRALPNNT
00000000000000000000000000000000000000000000000000000000000000
00000000000000000000000000000000000000000000000000000000000000
00000000000000000000000000000000000000000000000000000000000000
000000000011111111111111111111111
>DM_validata545
MSLLNNKAGVISRLADFLGFRPKTGDIDVMNRQSVGSVTISQLAKGFYEPNIESAINDVHNF
SIKDVGTIITNKTGVSPEGVSQTDYWAFSGTVTDDSLPPGSPITVLVFGLPVSATTGMTAIE
FVAKVRVALQEAIASFTAINSYKDHPTDGSKLEVTYLDNQKHVLSTYSTYGITISQEIISES
KPGYGTWNLLGAQTVTLDNQQTPTVFYHFERTA
11111111111000000000000000000000000000000000000000000000000000
00000000000000000000000000000000000000000000000000000000000000

000000000000000000000000000000000000000000000000000000000000
00000000000000000000000000000000
>DM_validata546
LVPELNEKDDDQVQKALASRENTQLMNRDNIEITVRDFKTLAPRRWLNDTIIEFFMKYIEKS
TPNTVAFNSFFYTNLSERGYQGVRRWMKRKKTQIDKLDKIFTPINLNQSHWALGIIDLKKKT
IGYVDSLSNGPNAMSFAILTDLQKYVMEESKHTIGEDFDLIHLDCPQQPNGYDCGIYVCMNT
LYGSADAPLDFDYKDAIRMRRFIAHLILTDALK
000000000000000000000000000110000000000000000000000000000000
000000000000000000000000000000000000000000000000000000000000
000000000000000000000000000000000000000000000000000000000000
00000000000000000000000000000000
>DM_validata547
MAGRRRWSTEQILDAAAELLLAGDAETFSVRKLAASLGTDSSSLYRHFRNKTELLRAVADRI
LLSAMDGYRPEGDWKQRLTAVALRLRESFGQQPQLAAVWGRHGSGGTGSRLMMEEVLQALRA
SGLPDDEIPARYHRLVILISSLITAEGGFGAVGAQEHEQGMEQFRVAVLGADPERFPALSHF
AREIRPLGADRGAAFEEILAAHLAHLEAAAPGS
111100000000000000000000000000000000000000000000000000000000
000000000000000000000000000000000000000000000000000000000000
000000000000000000000000000111111111111110000000000000000000
00000000000000000000000000000011
>DM_validata548
VLDSFEILKALKSLDLLKNAPAWWWPNALKFEALLGAVLTQNTKFEAVLKSLENLKNAFILE
NDDEINLKKIAYIEFSKLAECVRPSGFYNQKAKRLIDLSGNILKDFQSFENFKQEVTREWLL
DQKGIGKESADAILCYACAKEVMVVDKYSYLFLKKLGIEIEDYDELQHFFEKGVQENLNSAL
ALYENTISLAQLYARFHGKIVEFSKQKLELKL
100000000000000000000000000000000000000000000000000000000000
000000000000000000000000000000000000000000000000000000000000
000000000000000000000000000000000000000000000000000000000000
0000000000000000000000000000000
>DM_validata549
MSRHWPLFDLRITTPRLQLQLPTEELCDQLIDTILEGVHDPDRMPFSVPWTRASREDLPFNT
LSHLWQQLAGFKRDDWSLPLAVLVDGRAVGVQALSSKDFPITRQVDSGSWLGLRYQGHGYGT
EMRAAVLYFAFAELEAQVATSRSFVDNPASIAVSRRNGYRDNGLDRVAREGAMAEALLFRLT
RDDWQRHRTVEVRVDGFDRCRPLFGPLEPPRY
100000000000000000000000000000000011111111111111111111000000
000000000000000000000000000000000000000000000000000000000000
000000000000000000000000000000000000000000000000000000000000
0000000000000000000000001111111
>DM_validata550
MATVEPETTPTPNPPTTEEEKTESNQEVANPEHYIKHPLQNRWALWFFKNDKSKTWQANLRL
ISKFDTVEDFWALYNHIQLSSNLMPGCDYSLFKDGIEPMWEDEKNKRGGRWLITLNKQQRRS
DLDRFWLETLLCLIGESFDDYSDDVCGAVVNVRAKGDKIAIWTTECENREAVTHIGRVYKER
LGLPPKIVIGYQSHADTATKSGSTTKNRFVV
111111111111111111111111100000000000000000000000000000000000

00000000000000000000000000000000000000000000000000000000000000
00000000000000000000000000000000000000000000000000000000000000
0000000000000000000111110000000
>DM_validata551
MADAGEGEDEIQFLRTDDEVVLQCTATIHKEQQKLCLAAEGFGNRLCFLESTSNSKNVPPDL
SICTFVLEQSLSVRALQEMLANTVEKSEGQVDVEKWKFMMKTAQGGGHRTLLYGHAILLRHS
YSGMYLCCLSTSRSSTDKLAFDVGLQEDTTGEACWWTIHPASKQRSEGEKVRVGDDLILVSM
SSERYLHLSYGNSSWHVDAAFQQTLWSVAPI
11111111111000000000000000000000000000000000000000000111100000
00000000000000000000000001111111111011111111111000000000000000
00000000000011111100000000000000000000000000000000000000000000
0000000000000000000000000000000
>DM_validata552
LNTSGSGTILIDLSPDDKEFQSVEEEMQSTVREHRDGGHAGGIFNRYNILKIQKVCNKKLWE
RYTHRRKEVSEENHNHANERMLFHGSPFVNAIIHKGFDERHAYIGGMFGAGIYFAENSSKSN
QYVYGIGGTGCPVHKDRSCYICHRQLLFCRVTLGKSFLQFSAMKMAHSPPGHHSVTGRPSV
NGLALAEYVIYRGEQAYPEYLITYQIMRPEG
11111100000000000000000000000000000000000000000000000000000000
00000000000000000000000000000000000000000000000000000000000000
00000000000000000000000000000000000000000000100000000000000000
0000000000000000000000000000001
>DM_validata553
MTTSAASQASLPRGRRTARPSGDDRELAILATAENLLEDRPLADISVDDLAKGAGISRPTFY
FYFPSKEAVLLTLLDRVVNQADMALQTLAENPADTDRENMWRTGINVFFETFGSHKAVTRAG
QAARATSVEVAELWSTFMQKWIAYTAAVIDAERDRGAAPRTLPAHELATALNLMNERTLFAS
FAGEQPSVPEARVLDTLVHIWVTSIYGENR
11111111111111111111100000000000000000000000000000000000000000
00000000000000000000000000000000000000000000000000000000000000
00000000000000000000000000000000000000000000000000000000000000
000000000000000000000000000001
>DM_validata554
LKIYGVYRSRASRPLWLLAELDLPFEHVPVIQANRVAHPHGPEAPLNTASAAYLAVNPLGQI
PCLEEEGLILTESLAITLHIARTQGGQLGPRSEPEDALMVSWSLFAATAVEPPALEIQLIQR
SGGGTSPEGQAAIAIAAERLRRPLARLERHFAAEDYLVGGRFTVADLNLAETLRYGQAHPAL
LEPFPAVAAWLDRCQSRPAFRLMMERRAAE
00000000000000000000000000000000000000001100000000000000000000
00000000000000000000000000000000000000000000000000000000000000
00000000000000000000000000000000000000000000000000000000000000
000000000000000000000000000000
>DM_validata555
MANKQQNFNTGIKDFDFWLSEVEALLASEDYGKDLASVNNLLKKHQLLEADISAHEDRLKDL
NSQADSLMTSSAFDTSQVKDKRETINGRFQRIKSMAAARRAKLNESHRLHQFFRDMDDEESW
IKEKKLLVSSEDYGRDLTGVQNLRKKHKRLEAELAAHEPAIQGVLDTGKKLSDDNTIGKEEI
QQRLAQFVDHWKELKQLAAARGQRLEESLE

100000000000000000000000000000000000000000000000000000000000
000000000000000000000000000000000000000000000000000000000000
000000000000000000000000000000000000000000000000000000000000
000000000000000000000000001111
>DM_validata556
DDYDGIPNLTAVQISPNADPALGLEYPVRPPVPGAGGLNASAKGIHSIAIGATAEAAKGAAV
AVGAGSIATGVNSVAIGPLSKALGDSAVTYGAASTAQKDGVAIGARASTSDTGVAVGFNSKA
DAKNSVAIGHSSHVAANHGYSIAIGDRSKTDRENSVSIGHESLNRQLTHLAAGTKDTDAVNV
AQLKKEIEKTQENTNKRSAELLANANAYAD
111111000000000000000000011111111110000000000000000000000000
000000000000000000000000000000000000000000000000000000000000
000000000000000000000000000000000000000000000000000000000000
000000000111111111111111111111
>DM_validata557
MERNKLARQIIDTCLEMTRLGLNQGTAGNVSVRYQDGMLITPTGIPYEKLTESHIVFIDGNG
KHEEGKLPSSEWRFHMAAYQSRPDANAVVHNHAVHCTAVSILNRSIPAIHYMIAAAGGNSIP
CAPYATFGTRELSEHVALALKNRKATLLQHHGLIACEVNLEKALWLAHEVEVLAQLYLTTLA
ITDPVPVLSDEEIAVVLEKFKTYGLRIEE
000000000000000000000000000000000000000000000000000000000000
000000000000000000000000000000000000000000000000000000000000
000000000000000000000000000000000000000000000000000000000000
00000000000000000000111111111
>DM_validata558
GMKDNYDAPESMLTGRVMYNGEALQLRGNEAVQLQLYQHGYAKHDPINVYVNQDGMYSANLF
DGEYQMITKSGNGPWTSEGRDTINVTVAGNTVQDVEVTPYYLVRDAQMTLEGNKVNASFKVE
KVAGGGIDRVFFMLSTTQFVNDAEHNVDRYDETDNLDAYDETGKLYTFATRDYTDNSMFQTA
LKRGTLFGRICIWPKGSDQGIYSKVIRLK
111111111000000000000000000000000000000000000000000000000000
000000000000000000000000000000000000000000000000000000000000
000000000000000000000000000000000000000000000000000000000000
00000000000000000000000000000
>DM_validata559
AIRLYKLAVALGVFIVSAPAFSHGHHSHGKPLTEVEQKAANGVFDDANVQNRTLSDWDGVWQ
SVYPLLQSGKLDPVFQKKADADKTKTFAEIKDYYHKGYATDIEMIGIEDGIVEFHRNNETTS
CKYDYDGYKILTYKSGKKGVRYLFECKDPESKAPKYIQFSDHIIAPRKSSHFHIFMGNDSQQ
SLLNEMENWPTYYPYQLSSEEVVEEMMSH
111111111111111111111111110000000000000000000000000000000000
000000000000000000000000000000000000000000000000000000000000
000000000000000000000000000000000000000000000000000000000000
00000000000000000000000000000
>DM_validata560
MASKTIKIMPVGDSCTEGMGGGEMGSYRTELYRLLTQAGLSIDFVGSQRSGPSSLPDKDHEG
HSGWTIPQIASNINNWLNTHNPDVVFLWIGGNDLLLNGNLNATGLSNLIDQIFTVKPNVTLF
VADYYPWPEAIKQYNAVIPGIVQQKANAGKKVYFVKLSEIQFDRNTDISWDGLHLSEIGYKK

IANIWYKYTIDILRALAGETQLEHHHHHH
111000000000000000000000000000000000000000000000000000000000
000000000000000000000000000000000000000000000000000000000000
000000000000000000000000000000000000000000000000000000000000
00000000000000000011111111111
>DM_validata561
MAQRLDGARFRYLNEQLYSGPSSAAQRLFQEDPEAFLLYHRGFQSQVKKWPLQPVDRIARDL
RQRPASLVVADFGCGDCRLASSIRNPVHCFDLASLDPRVTVCDMAQVPLEDESVDVAVFCLS
LMGTNIRDFLEEANRVLKPGGLLKVAEVSSRFEDVRTFLRAVTKLGFKIVSKDLTNSHFFLF
DFQKTGPPLVGPKAQLSGLQLQPCLYKRR
100000000000000000000000000000000000000000000000000000000000
000000000000000000000000000000000000000000000000000000000000
000000000000000000000000000000000000000000000000000000000000
00000000000000000000000000111
>DM_validata562
FSDTLEESSPIAAIFDTENLEKISITEGIERGIVDSITGQRLLEAQACTGGIIHPTTGQKLS
LQDAVSQGVIDQDMATRLKPAQKAFIGFEGVKGKKKMSAAEAVKEKWLPYEAGQRFLEFQYL
TGGLVDPEVHGRISTEEAIRKGFIDGRAAQRLQDTSSYAKILTCPKTKLKISYKDAINRSMV
EDITGLRLLEAASVSSKGLPSPYNMSSA
111111100000000000000000000000000000000000000000000000000000
000000000000000000000000000011111110000000000000000000000000
000000000000000000000000000000000000000000000000000000000000
0000000000000000011111111111
>DM_validata563
MELAAGSFSEEQFWEACAELQQPALAGADWQLLVETSGISIYRLLDKKTGLYEYKVFGVLED
CSPTLLADIYMDSDYRKQWDQYVKELYEQECNGETVVYWEVKYPFPMSNRDYVYLRQRRDLD
MEGRKIHVILARSTSMPQLGERSGVIRVKQYKQSLAIESDGKKGSKVFMYYFDNPGGQIPSW
LINWAAKNGVPNFLKDMARACQNYLKKT
111111100000000000000000000000000000000000000000000000000000
000000000000000000000000000000000000000000000000000000000000
000000000000000000000000000000000000000000000000000000000000
0000000000000000000000001111
>DM_validata564
NTGIVSSFFTYTGPAHGTQWDEIDIEFLGKDTTKVQFNYYTNGVGGHEKVISLGFDASKGFH
TYAFDWQPGYIKWYVDGVLKHTATANIPSTPGKIMMNLWNGTGVDDWLGSYNGANPLYAEYD
WVKYTSNQTGGSFFEPFNSYNSGTWEKADGYSNGGVFNCTWRANNVNFTNDGKLKLGLTSSA
YNKFDCAEYRSTNIYGYGLYEVSMKPAK
000000000000000000000000000000000000000000000000000000000000
000000000000000000000000000000000000000011111111111110000000
000000000000000000000000000000000000000000000000000000000000
0000000000000000000000000000
>DM_validata565
FLGDGGDVSFSTRGTQNWTVERLLQAHRQLEERGYVFVGYHGTFLEAAQSIVFGGVRARSQD
LDAIWRGFYIAGDPALAYGYAQDQEPDARGRIRNGALLRVYVPRSSLPGFYRTSLTLAAPEA

AGEVERLIGHPLPLRLDAITGPEEEGGRLETILGWPLAERTVVIPSAIPTDPRNVGGDLDPS
SIPDKEQAISALPDYASQPGKPPREDLK
000000000000000000000000000000000000000000000000000000000000
000000000000000000000111111111000000000000000000000000000000
000000000000000000000000000000000000000000000000000000000000
0000000000000011111111111111
>DM_validata567
LPTSNPAQELEARQLGRTTRDDLINGNSASCADVIFIYARGSTETGNLGTLGPSIASNLESA
FGKDGVWIQGVGGAYRATLGDNALPRGTSSAAIREMLGLFQQANTKCPDATLIAGGYSQGAA
LAAASIEDLDSAIRDKIAGTVLFGYTKNLQNRGRIPNYPADRTKVFCKTGDLVCTGSLIVAA
PHLAYGPDARGPAPEFLIEKVRAVRGSA
111111111111111100000000000000000000000000000000000000000000
000000000000000000000000000000000000000000000000000000000000
000000000000000000000000000000000000000000000000000000000000
0000000000000000000000000111
>DM_validata568
LPLYFGWFLTKKSSETLRKAGQVFLEELGNHKAFKKELRQFVPGDEPREKMDLVTYFGKRPP
GVLHCTTKFCDYGKAPGAEEYAQQDVLKKSYSKAFTLTISALFVTPKTTGARVELSEQQLQL
WPSDVDKLSPTDNLPRGSRAHITLGCAADVEAVQTGLDLLEILRQEKGGSRGEEVGELSRGK
LYSLGNGRWMLTLAKNMEVRAIFTGYYG
000000000000000000000000000000000000000001111100000000000000
000000000000000000000000000000000000000000000000000000000000
000000000000000000000000000000000000000000000000000000000000
0000000000000000000000000000
>DM_validata569
GHMSDKRNPRISSRKQPQQARSSELVASILEAAVQVLASEGAQRFTTARVAERAGVSIGSLY
QYFPNKAAILFRLQSDEWRRTTRLLGEILEDTTRPPLERLRRLVLAFVRSECEEAAIRVALS
DAAPLYRDADEAREVKAEGARVFQAFLREALPEVAEAERSLAGDLLTTTLGAVGKQFSEQPR
SEAEIERYAEALADMLCAYLAALGERGS
111111111111111111111100000000000011111100000000000000000000
000000000000000000000000000000000000000000000000000000000000
000000111100000000000000000000000000000000000000000000000000
0000000000000000000000000111
>DM_validata570
ALDVRGGYRSGSHAYETRLKVSEGWQNGWWASMESNTWNTIHDNKKENAALNDVQVEVNYAI
KLDDQWTVRPGMLTHFSSNGTRYGPYVKLSWDATKDLNFGIRYRYDWKAYRQQDLSGDMSRD
NVHRWDGYVTYHINSDFTFAWQTTLYSKQNDYRYANHKKWATENAFVLQYHMTPDITPYIEY
DYLDRQGVYNGRDNLSENSYRIGVSFKL
000000000000000000000000000000000000000111111111100000000000
000000000000000000000000000000000000000000000000000000000000
000000000000000000000000000000000000000000000000000000000000
0000000000000000000000000000
>DM_validata571
MSLVEAISLWNEGVLAADKKDWKGALDAFSAVQDPHSRICFNIGCMYTILKNMTEAEKAFTR

SINRDKHLAVAYFQRGMLYYQTEKYDLAIKDLKEALIQLRGNQLIDYKILGLQFKLFACEVL
YNIAFMYAKKEEWKKAEEQLALATSMKSEPRHSKIDKAMECVWKQKLYEPVVIPVGRLFRPN
ERQVAQLAKKDYLGKATVVASVVDQDS
000000000000000000000000000000000000000000000000000000000000
000000000000000000000000000000000000000000000000000000000000
000000000000000000000000000000000000000000000000000000000000
000000011111111111111111111
>DM_validata572
AVINHDAVPVWPQPEPADATQALAVRFKPQLDVVNGCQPYPAVDPQGNTSGGLKPSGSQAAA
CRDMSKAQVYSRSGTYNGYYAIMYSWYMPKDSPSTGIGHRHDWENVVVWLDNAASANIVALS
ASAHSGYKKSFPADKSYLDGITAKISYKSTWPLDHELGFTTSAGKQQPLIQWEQMTQAARDA
LESTDFGNANVPFKSNFQDKLVKAFFQ
000000000000000000000000000000000000000000000000000000110000
000000000000000000000000000000000000000000000000000000000000
000000000000000000000000000000000000000000000000000000000000
000000000000000000000000000
>DM_validata573
MTQGAVKTTGKRSRAVSAKKKAILSAALDTFSQFGFHGTRLEQIAELAGVSKTNLLYYFPSK
EALYIAVLRQILDIWLAPLKAFREDFAPLAAIKEYIRLKLEVSRDYPQASRLFCMEMLAGAP
LLMDELTGDLKALIDEKSALIAGWVKSGKLAPIDPQHLIFMIWASTQHYADFAPQVEAVTGA
TLRDEVFFNQTVENVQRIIIEGIRPR
111111111111100000000000000000000000000000000000000000000000
000000000000000000000000000000000000000000000000000000000000
000000000000000000000000000000000000000000000000000000000000
00000000000000000000000000
>DM_validata575
VLRSKSLCHDEIENLLDSDHRELIGDYSKAFLLQTVDGKHQDLKYISPETMVALLTGKFSNI
VDKFVIVDCRYPYEYEGGHIKTAVNLPLERDAESFLLKSPIAPCSLDKRVILIFHCEFSSER
GPRMCRFIRERDRAVNDYPSLYYPEMYILKGGYKEFFPQHPNFCEPQDYRPMNHEAFKDELK
TFRLKTRSWAGERSRRELCSRLQDQ
111111111111111111000000000000000000000000000000000000000000
000000000000000000000000000000000000000000000000000000000000
000000000000000000000000000000000000000000000000000000000000
0000000001111111111111111
>DM_validata576
ADAPFEGRKKCSSCHKAQAQSWKDTAHAKAMESLKPNVKKEAKQKAKLDPAKDYTQDKDCVG
CHVDGFGQKGGYTIESPKPMLTGVGCESCHGPGRNFRGDHRKSGQAFEKSGKKTPRKDLAKK
GQDFHFEERCSACHLNYEGSPWKGAKAPYTPFTPEVDAKYTFKFDEMVKEVKAMHEHYKLEG
VFEGEPKFKFHDEFQASAKPAKKGK
000000000000000000000000000000000000000000000000000000000000
000000000000000000000000000000000000000000000000000000000000
000000000000000000000000000000000000000000000000000000000000
0000000000000000000000111
>DM_validata577

QNSQFQSLEQVKRRPAHLMALLQHVALQFEPGPLLCCLHADMLGSLGPKEAKKAFLDFYHSF
LEKTAVLRVPVPPNVAFELDRTRADLISEDVQRRFVQEVVQSQQVAVGRQLEDFRSKRLMGM
TPWEQELAQLEAWVGRDRASYEARERHVAERLLMHLEEMQHTISTDEEKSAAVVNAIGLYMR
HLGVRTKSGDKKSGRNFFRKKVMGN
00000000000000000000000000000000000000000000000000000000000000
00000000000000000000000000000000000000000000000000000000000000
00000000000000000000000000000000000000000000000000000000000000
0000001111111111111111111
>DM_validata578
MQEPLAAGAVILRRFAFNAAEQLIRDINDVASQSPFRQMVTPGGYTMSVAMTNCGHLGWTTH
RQGYLYSPIDPQTNKPWPAMPQSFHNLCQRAATAAGYPDFQPDACLINRYAPGAKLSLHQDK
DEPDLRAPIVSVSLGLPAIFQFGGLKRNDPLKRLLLEHGDVVVWGGESRLFYHGIQPLKAGF
HPLTIDCRYNLTFRQAGKKENLYFQ
11110000000000000000000000000000000000000000000000000000000000
00000000000000000000000000000000000000000000000000000000000000
00000000000000000000000000000000000000000000000000000000000000
0000000000000000011111111
>DM_validata579
MWLTKLVLNPASRAARRDLANPYEMHRTLSKAVSRALEEGRERLLWRLEPARGLEPPVVLVQ
TLTEPDWSVLDEGYAQVFPPKPFHPALKPGQRLRFRLRANPAKRLAATGKRVALKTPAEKVA
WLERRLEEGGFRLLEGERGPWVQILQDTFLEVRRKKDGEEAGKLLQVQAVLFEGRLEVVDPE
RALATLRRGVGPGKALGLGLLSVAP
00000000000000000000000000000000000000000000000000000000000000
00000000000000000000000000000000000000000000011111111100000000
00000000000000000000000000000001111111111111000000000000000000
0000000000000000000000000
>DM_validata580
ILQTTVDGNSTAISNLKSDISSNGLAITDLQDRVKSLESTASHGLSFSPPLSVADGVVSLDM
DPYFCSQRVSLTSYSAEAQLMQFRWMARGTNGSSDTIDMTVNAHCHGRRTDYMMSSTGNLTV
TSNVVLLTFDLSDITHIPSDLARLVPSAGFQAASFPVDVSFTRDSATHAYQAYGVYSSSRVF
TITFPTGGDGTANIRSLTVRTGIDT
11111100000000000000000000000000000000000000000000000000000000
00000000000000000000000000000000000000000000000000000000000000
00000000000000000000000000000000000000000000000000000000000000
0000000000000000000000000
>DM_validata581
PSTLELLKDVHLGLPVPCHDPARLALLSGHYLYYHYGCDGLDDRGWGCGYRTLQTLCSWPGG
QSSGVPGLPALQGALEAMGDKPPGFRGSRNWIGCVEASLCLEHFGGPQGRLCHLPRGVGLRG
EEERLYSHFTTGGGPVMVGGDADAQSKALLGICEGPGSEVYVLILDPHYWGTPKNRCELQAA
GWVGWQKVKSVFDSNSFYNLCFTR
10000000000000000000000000000000000000000000000000000000000000
00000000000000000000000000000000000000000000000000000000000000
00000000000000000000000000000000000000000000000000000000000000
000000000000000000000000

>DM_validata582
MNEADYLRLLTRQAEQANDFLSNARKWDRERWVCQRFLEALNVPYRQEDFAAPGEQPPDVLF
KGAGFEVFFVLDEGRRLNEEWREELTRRRQAVSLRQLIRREERPQRIAAAELQARLAPTLRK
KAHNYSERGIDHGELDLLAFVNLKRAVPDFNTPFPPPTEYLRQGWRSLSMVGPTFARVLFAH
SGAPEFLRANLGRSILFDAGVGL
000000000000000000000000000000000000000000000000000000000000
000000000000011111111111111111111111111100000000000000000000
000000000000000000000000000000000000000000000000000000000000
00000000000000000000000
>DM_validata583
GHMTKLFSKVKESIEGIKMPSTLTINGKAPIVAYAELIAARIVNALAPNSIAIKLVDDKKAP
AAKLDDATEDVFNKITSKFAAIFDNGDKEQVAKWVNLAQKELVIKNFAKLSQSLETLDSQLN
LRTFILGGLKYSAADVACWGALRSNGMCGSIIKNKVDVNVSRWYTLLEMDPIFGEAHDFLSK
SLLELKKSANVGKKKETHKANFE
111111111111110000000000000000000000000000000000000000000000
000000000000000000000000000000000000000000000000000000000000
000000000000000000000000000000000000000000000000000000000000
00000001111111111111111
>DM_validata584
RVTLNIATNADSLGTWFLDAVSKFTGGSDYLVNIAVDDQDHTVEWLRGGRVLAAVTAHDKPV
QGCRVTPLGVLRYHATASPDFMARHFADGVTPAALARAPGLTFNQKDRLQASWIRTALGEDV
SYPTHWLPSTDGFVKASLAGMGWGLNPVQLVAEHLAAGRLVELMPGTPLDIPLYWQVNRLAA
ERLAGLTANMVGTARVVLMPVG
000000000000000000000000000000000000000000000000000000000000
000000000000000000000000000000000000000000000000000000000000
000000000000000000000000000000000000000000000000000000000000
0000000000000000000001
>DM_validata585
MSTPSVHCLKPSPLHLPSGIPGSPGRQRRHTLPANEFRCLTPEDAAGVFEIEREAFISVSGN
CPLNLDEVQHFLTLCPELSLGWFVEGRLVAFIIGSLWDEERLTQESLALHRPRGHSAHLHAL
AVHRSFRQQGKGSVLLWRYLHHVGAQPAVRRAVLMCEDALVPFYQRFGFHPAGPCAIVVGSL
TFTEMHCSLRGHAALRRNSDR
111111111111111111111111111100000000000000000000000000000000
000000000000000000000000000000000000000000000000000000000000
000000000000000000000000000000000000000000000000000000000000
000000000111111111111
>DM_validata586
MAEHDFETISSETLHTGAIFALRRDQVRMPGGGIVTREVVEHFGAVAIVAMDDNGNIPMVYQ
YRHTYGRRLWELPAGLLDVAGEPPHLTAARELREEVGLQASTWQVLVDLDTAPGFSDESVRV
YLATGLREVGRPEAHHEEADMTMGWYPIAEAARRVLRGEIVNSIAIAGVLAVHAVTTGFAQP
RPLDTEWIDRPTAFAARRAER
111110000000000000000000000011111000000000000000000000000000
000000000000000000000000000000000000000000000000000000000000
000000000001111111111000000000000000000000000000000000000000

000000000000000000000
>DM_validata587
MDKDCEMKRTTLDSPLGKLELSGCEQGLHEIKLLGKGTSAADAVEVPAPAAVLGGPEPLMQC
TAWLNAYFHQPEAIEEFPVPALHHPVFQQESFTRQVLWKLLKVVKFGEVISYQQLAALAGNP
KAARAVGGAMRGNPVPILIPCHRVVCSSGAVGNYSGGLAVKEWLLAHEGHRLGKPGLGGSSG
LAGAWLKGAGATSGSPPAGRN
111100000000000000000000000000000011111111000000000000000000
000000000000000000000000000000000000000000000000000000000000
000000000000000000000000000000000000000000000000000000011111
111111111111111111111
>DM_validata588
GPGSMAEEIRTMIIGTSSAFRANVLREHFGDRFRNFVLLPPDIDEKAYRAADPFELTESIAR
AKMKAVLEKARQHSPPISGPAIALTFDQVVVKGDEVREKPLSTEQCRSFIASYSGGGVRTVA
TYALCVVGTENVLVAHNETETFFSKFGDDIVERTLERGACMNSAGGLVVEDEDMSRHVVRIV
GTSYGVRGMEPAVVEKLLSQL
111111000000000000000000000000000000000000000000000000000000
000000000000011111100000000000000000000000000000000000000000
000000000000000000000000000000000000000000000000000000000000
000000000000000000000
>DM_validata589
MDNEKGLLIVLSGPSGVGKGTVRKRIFEDPSTSYKYSISMTTRQMREGEVDGVDYFFKTRDA
FEALIKDDQFIEYAEYVGNYYGTPVQYVKDTMDEGHDVFLEIEVEGAKQVRKKFPDALFIFL
APPSLEHLRERLVGRGTESDEKIQSRINEARKEVEMMNLYDYVVVNDEVELAKNRIQCIVEA
EHLKRERVEAKYRKMILEAKK
111000000000000000000000000000000000000000000000000000000000
000000000000000000000000000000000000000000000000000000000000
000011111111111111111111111111000000000000000000000000000000
000000000001111111111
>DM_validata590
SNASQEIVAGELERCFLAMPESVLPIVTMEERNDLCRRAGHLSGFTHTASLESSLGGTVTFL
LNRNFIRIQTSTVGEVFMRILPFSDSSSVICVVTTVLHPVADSRIDFYTTEWKPLKTDRFWQ
QPRIEDFFLPHTDRQSYAYQAIYASLTPSYMQVSLSEESDTLSIRQTVTETLAEEEKPLAAI
FLSPEPLVYRWQSGRFVRQVR
111111110000000000000000000000000000000000000000000000000000
000000000000000000000000000000000000000000000000000000000000
000000000000000000000000000000000000000000000000000000000000
000000000000000000011
>DM_validata591
MAHHHHHHDDKSVCDSDNKEYMGIEVYVEATLDEPLRQTTCESKIHKYGASVSNGGLNISVD
LLNCFLNFHTVGVYTNRDTVYAKFASLDPWTTEPINSMTHDDLVKLTEECIVDIYLKCEVDK
TKDFMKTNGNRLKPRDFKTVPPSNVGSMIELQSDYCVNDVTTYVKIYDECGNIKQHSIPTLR
DYFTTKNGQPRKILKKKFDNC
111111111111100000000000000000000000000000000000000000000000
000000000000000000000000000000000000000000000000000000000000

000000000000000000000000000000000000000000000000000000000000
000000000000000000000
>DM_validata592
SDAMTTFLQRDEFAVTARVLGALFYYSPESHETAPLVQALLNDDWQAQWPLDAEALAPVAAM
FKTHSEESLPQAWQRLFIGPYALPSPPWGSVWLDRESVLFGDSTLALRQWMRENGIQFEMQQ
NEPEDHFGSLLLLAAWLAENDRHHECEQLLAWHLFPWSSRFLDVFIDHAGHPFYQALGQLAR
LTLAQWQAQLIIPVAVKPLFR
000000000000000000000000000000000000000000000000000000000000
000000000000000000000000000000000000000000000000000000011111
100000000000000000000000000000000000000000000000000000000000
000000000000000000000
>DM_validata594
MVFKIKDEWGEFLVRLARRAIEEYLKTGKEIEPPKDTPPELWEKMGVFVTLNRYNVPPQTAL
RGCIGFPTPIYPLVEATIKAAIYSAVDDPRFPPVKLEEMDNLVVEVSVLTPPELIEGPPEER
PRKIKVGRDGLIVEKGIYSGLLLPQVPVEWGWDEEEFLAETCWKAGLPPDCWLDEDTKVYKF
TAEIFEEEYPRGPIKRKPLV
110000000000000000000000000000000000000000000000000000000000
000000000000000000000000000000000000000000000000000000000000
000000000000000000000000000000000000000000000000000000000000
00000000000000000001
>DM_validata595
TVESNWGGAILIGSDFDTVSATANVPSASGGSSAAGTAWVGIDGDTCQTAILQTGFDWYGDG
TYDAWYEWYPEVSDDFSGITISEGDSIQMSVTATSDTSGSATLENLTTGQKVSKSFSNESSG
SLCRTNAEFIIEDFEECNSNGSDCEFVPFASFSPAVEFTDCSVTSDGESVSLDDAQITQVII
NNQDVTDCSVSGTTVSCSYV
000000000000000000000000000000000000000000000000000000000000
000000000000000011000000000000000000000000000000000000000000
000000000000000000000000000000000000000000000000000000000000
00000000000000000000
>DM_validata596
GAMGNICNSGGVSRTYSPPTSPVYGSGVSSPSRFVGQYTLTSIHQLSSEERENFLDAHDPMR
VYDLNSETSVYRTTQREYVRNGYATGNPNSGAIIALHEELQESPYAQHIGARPDQADAYRPR
TAHVSSLNTPSLNVMAGQGALSALRGYAGSDHVTTEMRLGDFLDQGGKVYSDTSAMSAGGDS
VEALIVTLPKGRKVPVNILD
111111111111111111111111111111000000000000000000000000000000
000000000000000000000000000000000000000000000000000000000000
000000000000000000000000111111100000000000000000000011110000
00000000000000000000
>DM_validata597
LRPVLPTQSAHDPPAVHLSNGPGQEPIAVMTFDLTKITKTSSSFEVRTWDPEGVIFYGDTNP
KDDWFMLGLRDGRPEIQLHNHWAQLTVGAGPRLDDGRWHQVEVKMEGDSVLLEVDGEEVLRL
RQVSGPLTSKRHPIMRIALGGLLFPASNLRLPLVPALDGCLRRDSWLDKQAEISASAPTSLR
SCDVESNPGIFLPPGTQAE
111111111111000000000000000000000000000000000000000000000000

000000000000000000000000000000000000000000000000000000000000
000000011100000000000000000000000000000000000000000000000000
0011111111111111111
>DM_validata598
MTLELLQAQAQNCTACRLMEGRTRVVFGEGNPDAKLMIVGEGPGEEEDKTGRPFVGKAGQLL
NRILEAAGIPREEVYITNIVKCRPPQNRAPLPDEAKICTDKWLLKQIELIAPQIIVPLGAVA
AEFFLGEKVSITKVRGKWYEWHGIKVFPMFHPAYLLRNPSRAPGSPKHLTWLDIQEVKRALD
ALPPKERRPVKAVSQEPLF
100000000000000000000000000000000000000000000000000000000000
000000000000000000000000000000000000000000000000000000000000
000000000000000000000000000000000000000000000000000000000000
0000000111111111111
>DM_validata599
TMAGRSMQAARCPTDELSLSNCAVVSEKDYQSGQHVIVRTSPNHKYIFTLRTHPSVVPGSVA
FSLPQRKWAGLSIGQEIEVALYSFDKAKQCIGTMTIEIDFLQKKNIDSNPYDTDKMAAEFIQ
QFNNQAFSVGQQLVFSFNDKLFGLLVKDIEAMDPSILKGEPASGKRQKIEVGLVVGNSQVAF
EKAENSSLNLIGKAKTKE
000000000000000000000000000000000000000000000000000000000000
000000000000000000000000000000000000000000000000000000000000
000000000000000000000000000000000011111111111100000000000000
000000000000000000
>DM_validata600
MGSSEQELKAIVKDLGCGPYFLGTYDKRFPGFVSPHKLACAIVNTAGRETGGVHWMAFAWNP
RSKTCYLFEPFGFSDQRLKQVYQFEYESLLRRSAIASSPDRCITLEKSTQSVQGPNSAACGL
FCCMFLHAFANWPQTPMDHNPTMNLITGVPNSMLNSPQVQPTLRRNQEQLYSFLERHSPYFR
SHSAQIRSATSFCHLKNM
100000000000000000000000000000000000000000000000000000000000
000000000000000000000000000000000000000000000000000000000000
000000000000000000000000000000000000000000000000000000000000
000000000000000000
>DM_validata601
MANNTLIETPLRHSPAEHLDTVMDAASVAGRVELREIAFTTQISLRCAPGTQAHAALAAATG
AGLPAKVGEVAGEAQGTAVLWLAPDEFLATSAENTELGGVLSAALGDAPGQVVDLSANRSVL
ELTGPDAPLVLRKSCPADLHPRAFAVNQAIVTSVANIPVLLWRTGEQAWRIMPRASFTEHTV
HWLVDAMSEFASEAVILE
111111110000000000000000000000000000000000000000000000000000
000000000000000000000000000000000000000000000000000000000000
000000000000000000000000000000000000000000000000000000000000
000000000000111111
>DM_validata602
GMSPTVLPATQADFPKIVDVLVEAFANDPTFLRWIPQPDPGSAKLRALFELQIEKQYAVAGN
IDVARDSEGEIVGVALWDRPDGNHSAKDQAAMLPRLVSIFGIKAAQVAWTDLSSARFHPKFP
HWYLYTVATSSSARGTGVGSALLNHGIARAGDEAIYLEATSTRAAQLYNRLGFVPLGYIPSD
DDGTPELAMWKPPAMPTV

1000000000000000000000000000000000000000000000000000000000000000
0000000000000000000001111111111110000000001000111111111100000000
0000000000000000000000000000000000000000000000000000000000000000
000000000000000000
>DM_validata604
AGEARLEEAVNRWVLKFYFHEALRAFRGSRYGDFRQIRDIMQALLVRPLGKEHTVSRLLRVM
QCLSRIEEGENLDCSFDMEAELTPLESAINVLEMIKTEFTLTEAVVESSRKLVKEAAVIICI
KNKEFEKASKILKKHMSKDPTTQKLRNDLLNIIREKNLAHPVIQNFSYETFQQKMLRFLESH
LDDAEPYLLTMAKKALK
0000000000000000000000000000000000000000000000000001111000000000
0000000000000000000000000000000000000000000000000000000000000000
0000000000000000111111111000000000000000000000000000000000000000
00000000000000000
>DM_validata605
MNRSIFNVILGGFGSEGGVAAAGGAAGDRSVKAGSAEDAAFIMKNASKVIIVPGYGMAVAQA
QHALREMADVLKKEGVEVSYAIHPVAGRMPGHMNVLLAEANVPYDEVFELEEINSSFQTADV
AFVIGANDVTNPAAKTDPSSPIYGMPILDVEKAGTVLFIKRSMASGYAGVENELFFRNNTMM
LFGDAKKMTEQIVQAMN
1111111111111111111111111110000000000000000000000000000000000000
0000000000000000000000000000000000000000000000000000000000000000
0000000000000000000000000000000000000000000000000000000000000000
00000000000000000
>DM_validata606
PQVTLWQRPLVTIKIGGQLKEALLDTGADDTVLEEMSLPGRWKPKMIGGIGGFIKVRQYDQI
LIEICGHKAIGTVLVGPTPVNIIGRDLLTQIGMTLNFGGSSGPQVTLWQRPLVTIKIGGQLK
EALLDTGADDTVLEEMSLPGRWKPKMIGGIGGFIKVRQYDQILIEICGHKAIGTVLVGPTPV
NIIGRDLLTQIGMTLNF
0000000000000000000000000000000000000000000000000000000000000000
0000000000000000000000000000000000000111110000000000000000000000
0000000000000000000000000000000000000000000000000000000000000000
00000000000000000
>DM_validata607
MPDPAAPGTLRPGGRTARIREAVLLAAGDALAADGFDALDLGEIARRAGVGKTTVYRRWGTP
GGLAADLLADMAEQSLPRADTGALEEDLRANARLVVRTLDDPRQGRLFRALIAASLCNEQAA
EALHRFYAVRVDEWAGCVRDAVARGEVPDGTDPHGVVAAVSAPLYYALLNTGRSLTEADADR
AARAASTAARAGVWVTG
1111111111111000000000000000000000000000000000000000000000000000
0000000000000000000000000000000000000000000000000000000000000000
0000000000000000000000000000000000000000000000000000000000000000
00000000000000000
>DM_validata608
MRIVEVYSHLNGLEYIQVHLPHIWEEIQEIIVSIDAEACRTKESKEKTKQGQILYSPVALNE
AFKEKLEAKGWKESRTNYYVTADPKLIRETLSLEPEEQKKVIEAAGKEALKSYNQTDFVKDR
VAIEVQFGKYSFVAYDLFVKHMAFYVSDKIDVGVEILPMKELSKEMSSGISYYEGELYNVIR

QGRGVPAVPLVLIGIAP
000000000000000000000000000000000000000011111111110000000000
000000000000000000000000000000000000000000000000000000000000
000000000000000000000000000000000000000000000000000000000000
00000000000000000
>DM_validata609
MGKSELSGRLNWQALAGLKASGAEQNLYNVFNAVFEGTKYVLYEKPKHLKNLYAQVVLPDDV
IKEIFNPLIDLSTTQWGVSPAFAIENTETHKILFGEIKRQDGWVEGKDPSAGRGNAHERSCK
LFTPGLLKAYRTIGGINDEEILPFWVVFEGDITRDPKRVREITFWYDHYQDNYFMWRPNESG
EKLVQHFNEKLKKYLD
111110000000000000000000000000000000000000000000000000000000
000000000000000000000000000000000000000000000000000000000000
000000000000000000000000000000000000000000000000000000000000
0000000000000000
>DM_validata610
MKELSKSDRTRQFIIESTAPVFNVKGLAGTSLTDLTEATNLTKGSIYGNFENKEAVAIAAFD
YNWGHVKSVLTAKVQACNTYKEMLLVYSSMYNDADGSLFPVGGCPLLNTTIEADDTHDALRK
KAGEAILSWKKNLVTIIKKGIQAKEFRPDTDVTKIAFSMIALVEGAILIHRATKNRAYSDYV
FESLEDLIAGIEVKKK
100000000000000000000000000000000000000000000000000000000000
000000000000000000000000000000000000000000000000000000000000
000000000000000000000000000000000000000000000000000000000000
0000000000000001
>DM_validata611
GSHMSNAKTKLGITKYSIVTNSNDSVTLRLMTEHDLAMLYEWLNRSHIVEWWGGEEARPTLA
DVQEQYLPSVLAQESVTPYIAMLNGEPIGYAQSYVALGSGDGWWEEETDPGVRGIDQLLANA
SQLGKGLGTKLVRALVELLFNDPEVTKIQTDPSPSNLRAIRCYEKAGFERQGTVTTPDGPAV
YMVQTRQAFERTRSDA
111111111111111111111100000000000000000000000000000000000000
000000000000000000000000000000000000000000000000000000000000
000000000000000000000000000000000000000000000000000000000000
0000000000000000
>DM_validata612
MNLMTTITGVVLAGGKARRMGGVDKGLLELNGKPLWQHVADALMTQLSHVVVNANRHQEIYQ
ASGLKVIEDSLADYPGPLAGMLSVMQQEAGEWFLFCPCDTPYIPPDLAARLNHQRKDAPVVW
VHDGERDHPTIALVNRAIEPLLLEYLQAGERRVMVFMRLAGGHAVDFSDHKDAFVNVNTPEE
LARWQEKRSHHHHHH
111000000000000000000000000000000000000000000000000000000000
000000000000000000000000000000000000000000000000000000000000
000000000000000000000000000000000000000000000000000000000000
000001111111111
>DM_validata614
MGSDKIHHHHHHMEKKIVYFNKPGRENTEETLRLAVERAKELGIKHLVVASSYGDTAMKALE
MAEGLEVVVVTYHTGFVREGENTMPPEVEEELRKRGAKIVRQSHILSGLERSISRKLGGVSR

TEAIAEALRSLFGHGLKVCVEITIMAADSGAIPIEEVVAVGGRSRGADTAVVIRPAHMNNFF
DAEIKEIICMPRNKR
111111111110000000000000000000000000000000000000000000000000
000000000000000000000000000000000000000000000000000000000000
000000000000000000000000000000000000000000000000000000000000
000000000000000
>DM_validata615
NYRSRAAYKLIELDNKYLFLKKNKIILDIGCYPGSWCQVILERTKNYKNKIIGIDKKIMDPI
PNVYFIQGEIGKDNMNNIKNINYIDNMNNNSVDYKLKEILQDKKIDIILSDAAVPCIGNKID
DHLNSCELTLSITHFMEQYINIGGTYIVKMYLGSQTNNLKTYLKGMFQLVHTTKPKASRNES
REIYLVCKNFLGRKK
100000000000000000000000000000000000000000000000000000000000
000000000000000000111111111110000000000000000000000000000000
000000000000000000000000000000000000000000000000000000111110
000000000000011
>DM_validata616
LFTVTAPKEVYTVDVGSSVSLECDFDRRECTELEGIRASLQKVENDTSLQSERATLLEEQLP
LGKALFHIPSVQVRDSGQYRCLVICGAAWDYKYLTVKVKASYMRIDTRILEVPGTGEVQLTC
QARGYPLAEVSWQNVSVPANTSHIRTPEGLYQVTSVLRLKPQPSRNFSCMFWNAHMKELTSA
IIDPLSRMEPKVPRT
000000000000000000000000000000111000000000000011000000000000000
000000000000000000000000000000000000000000000000000000000000
000000000000000000000000000000000000000000000000000000000000
000011111111111
>DM_validata617
MERTELLKPRTLADLIRILHELFAGDEVNVEEVQAVLEAYESNPAEWALYAKFDQYRYTRNL
VDQGNGKFNLMILCWGEGHGSSIHDHTDSHCFLKLLQGNLKETLFDWPDKKSNEMIKKSERT
LRENQCAYINDSIGLHRVENVSHTEPAVSLHLYSPPFDTCHAFDQRTGHKNKVTMTFHSKFG
IRTPFTTSGSLENN
111000000000000000000000000000000000000000000000000000000000
000000000000000000000000000000000000000000000000000000000000
000000000000000000000000000000000000000000000000000000000000
00001111111111
>DM_validata618
GHMSELRILRAVDYPRMPWKNGAGSTEEIARDGGDGLDGFGWRLSIADVGESGGFSGFAGYQ
RIISVLEGGGMRLRVDGAESAPLRARQAFAFSGDSEVHCTLLDGAIRDFNLIYAPRRHRARL
QWLRVEGELDWHGTASTLLLFAQQDGVAISLQGQPRGQLAAHDCLCAEGLQGLQHWRLTAHE
PAWVCAVELDSLGS
111100000000000000000000000000111111000000000000000000000000
000000000000000000000000000000000000000000000000000000000000
000000000000000000000000000000000000000000000000000000000000
00000000000001
>DM_validata619
MTNSNNRQSEYPVDPLFLDRWSPRAFDGSPMPKEHLLTILDAAHWAPSASNHQPWRFVYAHK

DSEDWPLFVELLMEGNQKWAKNASVLLFVISRDHTISHEGEKKPSATHSFDAGAAWFSLAMQ
AHLLGYHAHGMGGIFKDRIVEKLDIPDGFKVEAGVAIGTLTDKSILPDDLAEREVPSKRVPL
ADVAFEGRFTGKAD
000000000000000000000000000000000000000000000000000000000000
000000000000000000000000000000000000000000000000000000000000
000000000000000000000000000000000000000000000000000000000000
00000000000011
>DM_validata620
MAHHHHHHMGGPPTIEELKREKIIPHVFPDENVDLTVDMYISFKSGKEVNHGNILDLAGTGS
VPRNIKFSEEPPEDYCYILFMIDPDFPSRRRPDGRDYVHWAVSGIKSKELVKGTDKNCITLL
PYVGPSIKKGTGLHRISFILSLVKEENKGNVTGVPLYRGEHYITRVKFNNCQSAYNVIQMND
MKIVGFNWCQMRRK
111111111000000000000000000000000000000000000000000000000000
000000000000000000000000000000000000000000000000000000000000
000000000000000000000000000000000000000000000000000000000000
00000000000000
>DM_validata621
MVSLRLPLILLSLLAISFSCSAAPPPVYDTEGHELSADGSYYVLPASPGHGGGLTMAPRVLP
CPLLVAQETDERRKGFPVRFTPWGGAAAPEDRTIRVSTDVRIRFNAATICVQSTEWHVGDEP
LTGARRVVTGPLIGPSPSGRENAFRVEKYGGGYKLVSCRDSCQDLGVSRDGARAWLGASQPP
HVVVFKKARPSPPE
111111111111111111111100000000000000000000000000000000000000
000000000000000000000000000000000000000000000000000000000111
111100000000111111000000000000000000000000000000000000000000
00000000111111
>DM_validata622
ASITGDALVALPEGESVRIADIVPGARPNSDNAIDLKVLDRHGNPVLADRLFHSGEHPVYAV
RTVEGLRVTGTANHPLLCLVDVAGVPTLLWKLIDEIKPGDYAVIQRSAFSVDCAGFARGKPE
FAPTTYTVGVPGLVRFLEAHHRDPDAKAIADELTDGRFYYAKVASVTDAGVQPVYSLRVDTA
DHAFITNGFVSHN
000000000000000000000000000000000000000000000000000000000000
000000000000000000000000000000000000000000000000111111111111
111111000000000000000000000000000000000000000000000000000000
0000000000000
>DM_validata623
ASDDNVFQPVDQLPEDLIPSSIQVLKFSGKYLKLEQDKAYFDWPGFKTAIDNYTGEDLSFDK
YDQSTINQQSQEVGAMVDKIAKFLHDAFAAVVDLSKLAAIILNTFTNLEEESSSGFLQFNTN
NVKKNSSWEYRVLFSVPFGDNAPSYFYSLVTTILITADIEEKTGWWGLTSSTKKNFAVQIDA
LELVVKKGFKAPN
111100000000000000000000000000000000000000000000000000000000
000000000000000000000000000000000000000000000000000000000000
000000000000000000111000000000000000000000000000000000000000
0000000000001
>DM_validata624

MRLFYAVFLPEEVRAALVEAQTKVRPFRGWKPVPPHQLHLTLLFLGERPEEELPDYLALGHR
LARLEAPFRARLRGTGYFPNEGTPRVWFAKAEAEGFLRLAEGLRAGVEELLGEEAVRIPGWD
KPFKPHITLARRKAPAPRVPPVLFGLEWPVEGFALVRSELKPKGPVYTVLEKFSLRGEHGRE
QAQGPGERPEGD
00000000000000000000000000000000000000000000000000000000000000
00000000000000000000000000000000000000000000000000000000000000
00000000000000000000000000000000000000000000000000000000000111
111111111111
>DM_validata625
TNKKNLRVVALAPTGRYFASIISSLEILETAAEFAEFQGFMTHVVTPNNRPLIGRGGISVQP
TAQWQSFDFTNILIIGSIGDPLESLDKIDPALFDWIRELHLKGSKIVAIDTGIFVVAKAGLL
QQNKAVMHSYFAHLFGELFPEIMLMTEQKALIDGNVYLSSGPYSHSSVMLEIVEEYFGKHTR
NLGNQFLSTIES
11110000000000000000000000000000000111100000000000000000000000
00000000000000000000000000000000000000000000000000000000000000
00000000000000000000000000000000000000000000000000000000001111
111100000000
>DM_validata626
MSNTKTASTGFAELLKDRREQVKMDHAALASLLGETPETVAAWENGEGGELTLTQLGRIAHV
LGTSIGALTPPAGNDLDDGVIIQMPDERPILKGVRDNVDYYVYNCLVRTKRAPSLVPLVVDV
LTDNPDDAKFNSGHAGNEFLFVLEGEIHMKWGDKENPKEALLPTGASMFVEEHVPHAFTAAK
GTGSAKLIAVNF
11110000000000000000000000000000000000000000000000000000000000
00000000000000000000000000000000000000000000000000000000000000
00000000000000000000000000000000000000000000000000000000000000
000000000000
>DM_validata627
AEPRYKADIGGGSLKLPESRIIAGLLLEGVTEDQWRHAIEVENVLQRRSPGTAKRQSSLMRN
RLETMGPELWQMVRDGSTQVAIQAVFAAAIKHSTLLGDFLDLVVRDQFRMFRPDLPRKMWDQ
YLEQCRNRDPLMPVWQDSTANKLADCVYRILVEVGYITDSKTYRLKSVRISGEVMSYLRENN
EQYVIRCIQVSI
11000000000000000000000000000000000000000000000111110000000000
00000000000000000000000000000000000000000000000000000000000000
00000000000011110000000000000000000001111000000000000000000000
000000000001
>DM_validata629
MRGSHHHHHHGSPPAAPITLWTGPGPSINGFINDTPVIRCFICLTRDSNLVTVNASFVGEGG
YRIVSPTQSQFSLIMEFDQFGQLMSTGNINSTTTWGEKPWGNNTVQPRPSHTWKLCMPNREV
YSTPAATISRCGLDSIAVDGAPSRSIDCMLIINKPKGVATYTLTFRFLNFNRLSGGTLFKTD
VLTFTYVGENQ
11111111111111100000000000000000000000000000000000000000000000
00000000000000000000000000000000000000000000000000000000000000
00000000000000000000000000000000000000000000000000000000000000
00000000000

>DM_validata630
SNAVRGFLIVGNKAFTQPFSLNDLPGAGRMDVLCRCTSQALFISHGIRRDVEVYLLLLGPPS
PPKSILIKGDEVRRMSPDERNVAGHIKKALAVECGKSWKKVHSGVYVSRKGLEELIEELSEK
YSIIYLKEDGVDISNAQLPPNPLFVIGDHEGLTEEQEKVVERYAALKLSLSPLSLLAEQCVV
IAHHHLDRLQF
11000000000000000000000000000000000000000000000000000000000000
00000000000000000000000000000000000000000000000000000000000000
00000000000000000000000000000000000000000000000000000000000000
00000000000
>DM_validata631
MSANSLLGSLRELQVLVLNPPGEVSDALVLQLIRIGCSVRQCWPPPEAFDVPVDVVFTSIFQ
NRHHDEIAALLAAGTPRTTLVALVEYESPAVLSQIIELECHGVITQPLDAHRVLPVLVSARR
ISEEMAKLKQKTEQLQDRIAGQARINQAKVLLMQRHGWDEREAHQHLSREAMKRREPILKIA
QELLGNEPSA
00000000000000000000000000000000000000000000000000000000000000
00000000000000000000000000000000000000000000000000000000000000
00000000000000000000000000000000000000000000000000000000000000
0000111111
>DM_validata632
MSKSESPKEPEQLRKLFIGGLSFETTDESLRSHFEQWGTLTDCVVMRDPNTKRSRGFGFVTY
ATVEEVDAAMNARPHKVDGRVVEPKRAVSREDSQRPGAHLTVKKIFVGGIKEDTEEHHLRDY
FEQYGKIEVIEIMTDRGSGKKRGFAFVTFDDHDSVDKIVIQKYHTVNGHNCEVRKALSKQEM
ASASSSQRGR
11111100000000000000000000000000000000000000000000000000000000
00000000000000000000000000000111111111100000000000000000000000
00000000000000000000000000000000000000000000000000000000011111
1111111111
>DM_validata633
MSDERYQQRQQKVKDRVDARVAQAQEERGIIIVFTGNGKGKTTAAFGTAARAVGHGKNVGVV
QFIKGTWPNGERNLLEPHGVEFQVMATGFTWETQNREADTAACMAVWQHGKRMLADPLLDMV
VLDELTYMVAYDYLPLEEVISALNARPGHQTVIITGRGCHRDILDLADTVSELRPVKHAFDA
GVKAQMGIDY
11111111111111111111111110000000000000000000000000000000000000
00000000000000000000000000000000000000000000000000000000000000
00000000000000000000000000000000000000000000000000000000000111
1111111111
>DM_validata634
NLIPTVIEQTNRGERAYDIYSRLLKDRIIMLGSAIDDNVANSIVSQLLFLAAEDPEKEISLY
INSPGGSITAGMAIYDTMQFIKPKVSTICIGMAASMGAFLLAAGEKGKRYALPNSEVMIHQP
LGGAQGQATEIEIAAKRILLLRDKLNKVLAERTGQPLEVIERDTDRDNFKSAEEALEYGLID
KILTHTEDKK
11111111111111111000000000000000000000000000000000000000000000
00000000000000000000000000000000000000000000000000000000000000
00000000000000000000000000000000000000000000000000000000000000

0000011111
>DM_validata635
MQLKRVAEAKLPTPWGDFLMVGFEELATGHDHVALVYGDISGHTPVLARVHSECLTGDALFS
LRCDCGFQLEAALTQIAEEGRGILLYHRQEGRNIGLLNKIRAYALQDQGYDTVEANHQLGFA
ADERDFTLCADMFKLLGVNEVRLLTNNPKKVEILTEAGINIVERVPLIVGRNPNNEHYLDTK
AEKMGHLLNK
000000000000000000000000000000000000000000000000000000000000
000000000000000000000000000000000000000000000000000000000000
000000000000000000000000000000000000000000000000111111111111
1111111111
>DM_validata637
MAGARRLELGEALALGSGWRHVCHALLYAPDPGMLFGRIPLRYAILMQMRFDGRLGFPGGFV
DTQDRSLEDGLNRELREELGEAAAAFRVERTDYRSSHVGSGPRVVAHFYAKRLTLEELLAVE
AGATRAKDHGLEVLGLVRVPLYTLRDGVGGLPTFLENSFIGSAREQLLEALQDLGLLQSGSI
SGLKIPAHH
110000000000000000000000000000000000000000000000000000000000
000000000000000000000000000000000000000000000000000000000000
000000000000000000000000000000000000000000000000000000001111
111111111
>DM_validata638
LAQYPVIGIDDDEFATAKKLITKQEVRAVTLSKLRLQDDLVMWDIGAGSASVSIEASNLMPN
GRIFALERNPQYLGFIRDNLKKFVARNVTLVEAFAPEGLDDLPDPDRVFIGGSGGMLEEIID
AVDRRLKSEGVIVLNAVTLDTLTKAVEFLEDHGYMVEVACVNVAKTKGLTEYKMFESHNPVY
IITAWKSDE
110000000000000000000000000000000000000000000000000000000000
000000000000000000000000000000000000000000000000000000000000
000000000000000000000000000000000000000000000011110000000000
000000011
>DM_validata639
ATDDVSKAYSSPTFDAEALLGTVISAEDPDRVLIEPWATGVDGVILDVGSGTGRWTGHLASL
GHQIEGLEPATRLVELARQTHPSVTFHHGTITDLSDSPKRWAGLLAWYSLIHMGPGELPDAL
VALRMAVEDGGGLLMSFFSGPSLEPMYHPVATAYRWPLPELAQALETAGFQVTSSHWDPRFP
HAYLTAEAS
100000000000000000000000000000000000000000000000000000000000
000000000000000000000000000000000000000000000000000000000000
000000000000000000000000000000000000000000000000000000000000
000000000
>DM_validata640
GLVLPHTEFQPAPKQDSSPHLTSQRPVDMVQLLKKYPIVWQGLLALKNDTAAVQLHFVSGNN
VLAHRSLPLSEGGPPLRIAQRMRLEATQLEGVARRMTVETDYCLLLALPCGRDQEDVVSQTE
SLKAAFITYLQAKQAAGIINVPNPGSNQPAYVLQIFPPCEFSESHLSRLAPDLLASISNISP
HLMIVIASV
111111111111111111111111000000000000000000000000000000000000
000000000011100000000000000000000000000000000000000000000000

000000000000000000000000000000000000000000000000000000000000
000000000
>DM_validata641
MANAIRLPPEAYPMSIAAQKNDDDRQARALAALAEKPDGIVEAIAAKAEVAPAEILAILPQG
AAVSAPADRFDAIWNEMRGWGEILMIVQTGDIVLEVPGHLPEGTESHGWFNIHGDSPIGGHI
KKDNCAAITFVDRGFHGRRSCSVWFMNAAGGAMFKIFVRRDENKELLAGQLAKFEELRDGFR
GLEHHHHHH
111111111100000000000000000000000000011110000000000000000000
000000000000000000000000000000000000000000000000000000000000
000000000000000000000000000000000000000000000000000000000000
111111111
>DM_validata642
ILKELENLSPEEAAHQKAVVETLLQEDPWRVAKMVKSYLQQHNIPQREVVDTTGLNQSHLSQ
HLNKGTPMKTQKRAALYTWYVRKQREVAQQFTHAGQGGLIEEPTGDELPTKKGRRNRFKWGP
ASQQILFQAYERQKNPSKEERETLVEECNRAECIQRGVSPSQAQGLGSNLVTEVRVYNWFAN
RRKEEAFR
000000000000000000000000000000000000000000000000000000000000
000000000000000000000000000000000111111111111111111110000000
000000000000000000000000000000000000000000000000000000000000
00000000
>DM_validata643
GMTLTRDSLLTLEAYAKVRRQEHARVIAHKKRRAVSIGNHLRLLFEDETTIRYQIHEMLHIE
KIFDEDGIQAELDAYLPLVPDGSNLKATLQIEYENETQRRAALARLVGIEDRVFLRVDDEAP
VYAIADEDLERDTAEKTSAVHFLRFELGDAMKAKLKAGAPLSIGCDHPHYPIQAARIDPDVA
ASLAGDLD
111000000000000000000000000000000000000000000000000000000000
000000000000000000000000000000000000000000000000000000000000
000001111111111111100000000000000000000000000000000000000000
00000000
>DM_validata644
MPRPKLKSDDEVLEAATVVLKRCGPIEFTLSGVAKEVGLSRAALIQRFTNRDTLLVRMMERG
VEQVRHYLNAIPIGAGPQGLWEFLQVLVRSMNTRNDFSVNYLISWYELQVPELRTLAIQRNR
AVVEGIRKRLPPGAPAAAELLLHSVIAGATMQWAVDPDGELADHVLAQIAAILCLMFPEHDD
FQLLQAHA
111110000000000000000000000000000000000000000000000000000000
000000000000000000000000000000000000000000000000000000000000
000000000000000000000000000000000000000000000000000000000000
00011111
>DM_validata645
MGGSHHHHHHGDDDDKMTKTFKTLDDFLGTHFIYTYDNGWEYEWYAKNDHTVDYRIHGGMVA
GRWVTDQKADIVMLTEGIYKISWTEPTGTDVALDFMPNEKKLHGTIFFPKWVEEHPEITVTY
QNEHIDLMEQSREKYATYPKLVVPEFANITYMGDAGQNNEDVISEAPYKEMPNDIRNGKYFD
QNYHRLNK
111111111111111110000000000000000000000000000000000000000000

00000000000000000000000000000000000000000000000000000000000000
00000000000000000000000000000000000000000000000000000000000000
00000011
>DM_validata646
MWAFPERFEGRHVRLEPLALAHLPAFLRHYDPEVYRFLSRAPVAPTEEALRAHLEGLLGEPG
RVNWAILFGKEVAGRISVIAPEPEHAKLELGTMLFKPFWGSPANKEAKYLLLRHAFEVLRAE
RVQFKVDLRNERSQRALEALGAVREGVLRKNRRLPDGAFRDDVVYSVLKEEWPGVKARLEAR
LYGASGNP
00000000000000000000000000000000000000000000000000000000000000
00000000000000000000000000000000000000000000000000000000000000
00000000000000000000000000000000000000000000000000000000000000
00111111
>DM_validata647
ILESLVTTLDEQGRINLAPLGPIVLPPQSPGGLPQFLLRPYEGSTTCDNLLASGNAVIHVID
DALLIAKTAIGKVDASDLVVPIPGLEDTHVRLKRCHRWFAVRVTQRAGTPPRHELTARCLAS
GLVDPFFGFNRAKHAVIEAAVAATRLHLLPPEEIEEELERARIAIEKTGGEPEREALQLIRR
HVRESSIS
00000000000000000000000000000000000000000000000000000000000000
00000000000000000000000000000000000000000000000000000000000000
00000000000000000000000000000000000000000000000000000000000000
00001111
>DM_validata648
MAPSMIPFTIKLKTCLKMCIQRLRYAQEKQQAIAKQSRRQVAQLLLTNKEQKAHYRVETLIH
DDIHIELLEILELYCELLLARVQVINDISTEEQLVKEHMDDGINEAIRSLIYAILFVDEVKE
LSQLKDLMAWKINVEFVNGVIADHIDVPEKIIKKCSPSVPKEELVDLYLKEIAKTYDVPYSK
LENSLSS
11111000000000000000000000000000000000000000000000000000000000
00000000000000000000000000000000000000000000000000000000000000
00000000000000000000000000000000000000000000000000000000000000
0000011
>DM_validata649
MGSSSSGRENLYFQGHMVARLLEEHGFETKTNVIVQGNCVEQEIDVVAERDGERYMIECKFH
NIPVYTGLKEAMYTYARFLDVEKHGFTQPWIFTNTKFSEEAKKYAGCVGIKLTGWSYPEKEG
IEVLLESKGLYPITILRIDKEVLDELVRAGLVFCRDVVSAGEEKLREIGLSAKKAREVIAEA
KKVIGGS
11111111100000000000000000000000000000000000000000000000000000
00000000000000000000000000000000000000000000000000000000000000
00000000000000000000000000000000000000000000000000000000000000
0000000
>DM_validata650
MNHLPVVGEDYVEIPDGRPFAPLAGKIEVVEIFGYTCPHCAHFDSKLQAWGARQAKDVRFTL
VPAVFGGVWDPFARAYLAADVLGVAKRSHTAMFEAIHEKGSVPIQNVGPDELAVFYAGYGVQ
PDRFVATFNGPEVEKRFQAARAYALKVRPVGTPTIVVNGRYMVTGHDFEDTLRITDYLVSRE
RAASHGR

1100000000000000000000000000000000000000000000000000000000000
000000000000000000000000000000000000000000000000000000000000
000000000000000000000000000000000000000000000000000000000000
0001111
>DM_validata651
GSNGDNEKMAALEAKICHQIEYYFGDFNLPRDKFLKEQIKLDEGWVPLEIMIKFNRLNRLTT
DFNVIVEALSKSKAELMEISEDKTKIRRSPSKPLPEVTDEYKNDVKNRSVYIKGFPTDATLD
DIKEWLEDKGQVLNIQMRRTLHKAFKGSIFVVFDSIESAKKFVETPGQKYKETDLLILFKDD
YFAKKNE
111111100000000000000000000000000000000000000000000000000000
000000000000000000000000000000000000000000000000000000000000
000000000000000000000000000000000000000000000000000000000011
1111111
>DM_validata652
PVQLNLLYVQARDDILNGSHPVSFDKACEFAGYQCQIQFGPHNEQKHKPGFLELKDFLPKEY
IKQKGERKIFMAHKNCGNMSEIEAKVRYVKLARSLKTYGVSFFLVKEKMKGKNKLVPRLLGI
TKECVMRVDEKTKEVIQEWSLTNIKRWAASPKSFTLDFGDYQDGYYSVQTTEGEQIAQLIAG
YIDIIL
000000000000000000000000000000000000000000000000000000000000
000000000000000000000000000000000000000000000011111000000000
000000000000000000000000000000000000000000000000000000000000
000000
>DM_validata653
AMKPRIYVKVKPERLGAVIGPRGEVKAEIMRRTGTVITVDTENSMVIVEPEAEGIPPVNLMK
AAEVVKAISLGFPPEKAFRLLEEDQILVVVDLKQVVGDSQNHLKRIKGRIIGEGGRARRTIE
EMTDTYINVGEYEVAIIGDYERAMAAKQAIEMLAEGRMHSTVYRHLERIMREIKRRERLKMW
AREEL
000000000000000000000000000000000000000000000000000000000000
000000000000000000000000000000000000000000000000000000000000
000000000000000000000000000000000000000000000000000000000000
00011
>DM_validata654
SNAAQISDGKQYITLDKPVAGEPQVLEFFSFYCPHCYQFEEVLHVSDNVKKKLPEGTKMTKY
HVEFLGPLGKELTQAWAVAMALGVEDKVTVPLFEAVQKTQTVQSAADIRKVFVDAGVKGEDY
DAAWNSFVVKSLVAQQEKAAADLQLQGVPAMFVNGKYQINPQGMDTSSMDVFVQQYADTVKY
LVDKK
111110000000000000000000000000000000000000000000000000000000
000000000000000000000000000000000000000000000000000000000000
000000000000000000000000000000000000000000000000000000000000
00001
>DM_validata655
GTEVEIPVVHDPKGEAVLPSVFEDGTRQGWDWAGESGVKTALTIEEANGSNALSWEFGYPEV
KPSDNWATAPRLDFWKSDLVRGENDYVTFDFYLDPVRATEGAMNINLVFQPPTNGYWVQAPK
TYTINFDELEEANQVNGLYHYEVKINVRDITNIQDDTLLRNMMIIFADVESDFAGRVFVDNV

RFEGA
111110000000000000000000000000000000000000000000000000000001
111110000000000000000000000000000000000000000000000000000000
000000000000000000000000000000000000000000000000000000000000
00001
>DM_validata656
MAAHQLSDFQRNKILRVFNTFYDCNHDGVIEWDDFELAIKKICNLHSWPTDGKKHNEARATL
KLIWDGLRKYADENEDEQVTKEEWLKMWAECVKSVEKGESLPEWLTKYMNFMFDVNDTSGDN
IIDKHEYSTVYMSYGIPKSDCDAAFDTLSDGGKTMVTREIFARLWTEYFVSNDRGAKGNHLF
GTLKL
100000000000000000000000000000000000000000000000000000000000
000000000000000000000000000000000000000000000000000000000000
000000000000000000000000000000000000000000000000000000000000
00000
>DM_validata657
MAEPLLVVGLGNPGANYARTRHNLGFVVADLLAARLGAKFKAHKRSGAEVATGRSAGRSLVL
AKPRCYMNESGRQIGPLAKFYSVAPANIIVIHDDLDLEFGRIRLKIGGGEGGHNGLRSVVAA
LGTKDFQRVRIGIGRPPGRKDPAAFVLENFTPAERAEVPTICEQAADATELLIEQGMEPAQN
RVHAW
000000000000000000000000000000000000000000000000000000000000
000000000000000000000000000000000000000000000000000000000000
000000000000000000000000000000000000000000000000000001111111
11111
>DM_validata658
MNPSAPSYPMASLYVGDLHPDVTEAMLYEKFSPAGPILSIRVCRDMITRRSLGYAYVNFQQP
ADAERALDTMNFDVIKGKPVRIMWSQRDPSLRKSGVGNIFIKNLDKSIDNKALYDTFSAFGN
ILSCKVVCDENGSKGYGFVHFETQEAAERAIEKMNGMLLNDRKVFVGRFKSRKEREAELGAR
AKEF
111111111100000000000000000000000000000000000000000000000000
000000000000000000000000000000000000011111111110000000000000
000000001111000000000000011111111111111110000001111111111111
1111
>DM_validata659
MLKNDDFVIAKNQLGNIVPNSVGVIRAVNGKSAMVLFIGLNELKRVDFSELEAIDIYRTGKG
YDKKICNICHILKNTDGFEINQTDAKGRKTTRPSCRECRKNIDGVKLSSTEKKKMDEIAPPK
GSVFTCPICEKRSIVGVTANLVHDHNHDTGWGREWICDSCNTGLGRFKDNPKFLEKVIEYLK
KYEK
100000000000000000000000000000000000000000000000000000000000
000000000000000000000000000000000000000000000000000000000000
000000000000000000000000000000000000000000000000000000000000
0001
>DM_validata660
MKGMKYHDYIWDLGGTLLDNYETSTAAFVETLALYGITQDHDSVYQALKVSTPFAIETFAPN
LENFLEKYKENEARELEHPILFEGVSDLLEDISNQGGRHFLVSHRNDQVLEILEKTSIAAYF

TEVVTSSSGFKRKPNPESMLYLREKYQISSGLVIGDRPIDIEAGQAAGLDTHLFTSIVNLRQ
VLDI
111000000000000000000000000000000000000000000000000000000000
000000000000000000000000000000000000000000000000000000000000
000000000000000000000000000000000000000000000000000000000000
0000
>DM_validata661
MEEVKKDVYSVWALPDEESEPRFKKLMEALRSEFTGPRFVPHVTVAVSAYLTADEAKKMFES
ACDGLKAYTATVDRVSTGTFFFQCVFLLLQTTPEVMEAGEHCKNHFNCSTTTPYMPHLSLLY
AELTEEEKKNAQEKAYTLDSSLDGLSFRLNRLALCKTDTEDKTLETWETVAVCNLNPGSHHH
HHH
000000000000000000000000000000000000000000000000000000000000
000000000000000000000000000000000000000111111111100000000000
000000000000000000000000000000000000000000000000000000011111
111
>DM_validata662
MSRGTQACINAAHTVSGIIGDLDTTIMFATAGTLHSDGDGSFADHREHILQTAKALVEDTKV
LVTGAAGTQDQLANAAQNAVSTITQLAEAVKRGACSLGSTQPDSQVMVINAVKDVASALGDL
INCTKLASGKSINDPSMQDLKESARVMVLNVSSLLKTVKAVEDEHTRGTRAMEATVEAISQE
IRA
111000000000000000000000000000000000111000000000000000000000
000000000000000000000000000000000000000000000000000000000000
000000001111111000000000000000000000000000000000000000000000
000
>DM_validata663
GMSYQNANYSAFYVSEPFSESNLGANSTHDFVYYNMLRMWKGEDNSFPFNDAHDKTYNVRDG
SDWEKTLKPRLHTRLDNSKNIILFLSSITANSRALREEMNYGIGTKGLPVIVIYPDYDKKSD
IVDSNGNFKKQIKDLWDKLPAFRDNMSSVATLHIPCTKSVIISALNNEDFMVNTMADAEKYY
YKP
111000000000000000000000000000000000000000000000000000000000
000000000000000000000000000000000000000000000000000000000000
000000000000000000000000000000000000000000000000000000000000
000
>DM_validata664
FDVIGGNAYTIGGRSRCSIGFAVNGGFITAGHCGRTGATTANPTGTFAGSSFPGNDYAFVRT
GAGVNLLAQVNNYSGGRVQVAGHTAAPVGSAVCRSGSTTGWHCGTITALNSSVTYPEGTVRG
LIRTTVCAEPGDSGGSLLAGNQAQGVTSGGSGNCRTGGTTFFQPVNPILQAYGLRMITTDSG
SSP
000000000000000000000000000000000000000000000000000000000000
000000000000000000000000000000000000000000000000000000000000
000000000000000000000000000000000000000000000000000000000111
111
>DM_validata666
MDSSEVVKVKQASIPAPGSILSQPNTEQSPAIVLPFQFEATTFGTAETAAQVSLQTADPITK

LTAPYRHAQIVECKAILTPTDLAVSNPLTVYLAWVPANSPATPTQILRVYGGQSFVLGGAIS
AAKTIEVPLNLDSVNRMLKDSVTYTDTPKLLAYSRAPTNPSKIPTASIQISGRIRLSKPMLI
AN
11111111111111111111111111110000000000000000000000000000000000
00000000000000000000000000000000000000000000000000000000000000
00000000000000000000000000000000000000000000000000000000000000
00
>DM_validata667
MRPLTEEETRVMFEKIAKYIGENLQLLVDRPDGTYCFRLHNDRVYYVSEKIMKLAANISGDK
LVSLGTCFGKFTKTHKFRLHVTALDYLAPYAKYKVWIKPGAEQSFLYGNHVLKSGLGRITEN
TSQYQGVVVYSMADIPLGFGVAAKSTQDCRKVDPMAIVVFHQADIGEYVRHEETLTLEHHHH
HH
00000000000000000000000000000000000000000000000000000000000000
00000000000000000000000000000000000000000000000000000000000000
00000000000000000000000000000000000000000000000000001111111111
11
>DM_validata669
MKGTIVGTWIKTLRDLYGNDVVDESLKSVGWEPDRVITPLEDIDDDEVRRIFAKVSEKTGKN
VNEIWREVGRQNIKTFSEWFPSYFAGRRLVNFLMMMDEVHLQLTKMIKGATPPRLIAKPVAK
DAIEMEYVSKRKMYDYFLGLIEGSSKFFKEEISVEEVERGEKDGFSRLKVRIKFKNPVFEYK
KN
00000000000000000000000000000000000000000000000000000000000000
00000000000000000000000000000000000000000000000000000000000000
00000000000000000000000000000000000000000000000000000000000000
01
>DM_validata670
MRGSHHHHHHRSPSKVQELSVYEINELDRHSPKILKNAFSLMFGLGDLVPFTNKLYTGDLKK
RVGITAGLCVVIEHVPEKKGERFEATYSFYFGDYGHLSVQGPYLTYEDSFLAITGGAGIFEG
AYGQVKLQQLVYPTKLFYTFYLKGLANDLPLELTGTPVPPSKDIEPAPEAKALEPSGVISNY
TN
11111111111111000000000000000000000000000000000000000000000000
00000000000000000000000000000000000000000000000000000000000000
00000000000000000000000000000000000000000000000000000000000000
00
>DM_validata671
MGSIEEVLLEERLIGYLDPGAEKVLARINRPSKIVSTSSCTGRITLIEGEAHWLRNGARVAY
KTHHPISRSEVERVLRRGFTNLWLKVTGPILHLRVEGWQCAKSLLEAARRNGFKHSGVISIA
EDSRLVIEIMSSQSMSVPLVMEGARIVGDDALDMLIEKANTILVESRIGLDTFSREVEELVE
CF
11111111111111111000000000000000000000000000000000000011110000
00000000000000000000000000000000000000000000000000000000000000
00000000000000000000000000000000000000000000000000000000000000
01
>DM_validata672

GSLSWKRCAGCGGKIADRFLLYAMDSYWHSRCLKCSSCQAQLGDIGTSSYTKSGMILCRNDY
IRLFGNSGACSACGQSIPASELVMRAQGNVYHLKCFTCSTCRNRLVPGDRFHYINGSLFCEH
DRPTALINGHLNSGGSGGSGGSGGDVMVVGEPTLMGGEFGDEDERLITRLENTQFDAANGID
DE
111000000000000000000000000000000000000000000000000000000000
000000000000000000000000000000000000000000000000000000000000
000000011111111111111100000000000000000000000000000001111111111
11
>DM_validata674
DESDNAFIRASRALTDKVTDLLGGLFSKTEMSEVLTEILRVDPAFDKDRFLKQCENDIIPNV
LEAMISGELDILKDWCYEATYSQLAHPIQQAKALGLQFHSRILDIDNVDLAMGKMVEQGPVL
IITFQAQLVMVVRNPKGEVVEGDPDKVLRMLYVWALCRDQDELNPYAAWRLLDISASSTEQI
L
111100000000000000000000000000000000000000000000000000000000
000000000000000000000000000000000000000000000000000000000000
000000000000000000000000000000000000000000000000000000000000
1
>DM_validata676
GPGGEDVIDSLQLNELLNAGEYKIGELTFQSIRSSQELQKKNTIVNLFGIVKDFTPSRQSLH
GTKDWVTTVYLWDPTCDTSSIGLQIHLFSKQGNDLPVIKQVGQPLLLHQITLRSYRDRTQGL
SKDQFRYALWPDFSSNSKDTLCPQPMPRLMKTGDKEEQFALLLNKIWDEQTNKHKNGELLST
S
111111000000000000000000000000000000000000000000000000000000
000000000000000000000000000000000000000000000000000000000000
000000000000000000000000000000000000000000000000001111111111
1
>DM_validata677
GMPELAVQKVVVHPLVLLSVVDHFNRIGKVGNQKRVVGVLLGSWQKKVLDVSNSFAVPFDED
DKDDSVWFLDHDYLENMYGMFKKVNARERIVGWYHTGPKLHKNDIAINELMKRYCPNSVLVI
IDVKPKDLGLPTEAYISVEEVHDDGTPTSKTFEHVTSEIGAEEAEEVGVEHLLRDIKDTTVG
T
100000000000000000000000000000000000000000000000000000000000
000000000000000000000000000000000000000000000000000000000000
000000010000000000000011110000000000000000000000000000001111
1
>DM_validata678
MTKQYLTHRCLIAPPEMADDFFANTVIYLARHDEEGAQGIIINRPAGIQIKELLNDLDIDAD
NVNPHEVLQGGPLRPEAGFVLHTGQPTWHSSIAVGENVCITTSKDILDAIAHNEGVGRYQIA
LGYASWGKNQLEDEIARGDWLICDADMDLIFNLPYDDRWDAAYKKIGVDRTWLASEIGHALE
111100000000000000000000000000000000000000000000000000000000
000000000000000000000000000000000000000000000000000000000000
000000000000000000000000000000000000000000000000000001111111
>DM_validata679
MPEITESERAYHLRKMKTRMQRVDVTGDGFISREDYELIAVRIAKIAKLSAEKAEETRQEFL

RVADQLGLAPGVRISVEEAAVNATDSLLKMKGEEKAMAVIQSLIMYDCIDTDKDGYVSLPEF
KAFLQAVGPDLTDDKAITCFNTLDFNKNGQISRDEFLVTVNDFLFGLEETALANAFYGDLVD
11000000000000000000000000000000000000000000000000000000000000
00000000000000000000000000000000000000000000000000000000000000
00000000000000000000000000000000000000000000000000000000000000
>DM_validata680
MAGVAEFNDKGELLLPKNYREWVMVGTQVTPNELNDGKAPFTEIRTVYVDPESYAHWKKTGE
FRDGTVTVKELVSVGDRKGPGSGNGYFMGDYIGLEASVKDSQRFANEPGNWAFYIFYVPDTP
LVAAAKNLPTAECAACHKENAKTDMVFTQFYPVLRAAKATGESGVVAPKKLAAALEHHHHHH
10000000000000000000000000000001111111111000000000000000000000
00000000000000000000111000000000000000000000000000000000000000
00000000000000000000000000000000000000000000000111111111111111
>DM_validata681
ATSWLLDGHLRAYTDDLARRLRGEPNAHLLHFADSQVVTMLSSADPDQQARAQRLLAGDDIP
PIVFLPINQPNAHWSLLVVDRRNKDAVAAYHYDSMAQKDPQQRYLADMAAYHLGLDYQQTHE
MPIAIQSDGYSAGDHVLTGIEVLAHRVLDGTFDYAGGRDLTDIEPDRGLIRDRLAQAEQAPA
10000000000000000000000000000000000000000000000000000000000000
00000000000000000000000000000000000000000000000000000000000000
00000001100000000000000000000000000000000000000000000000011111
>DM_validata682
NAMPPIIKRRVMRKIIIASQNPAKVNAVRSAFSTVFPDQEWEFIGVSVPSEVADQPMSDEET
KQGALNRVRNAKQRHPGAEYYVGLEAGIEENKTFAWMIVESDQQRGESRSACLMLPPLVLER
LRQAKELGDVMDEVFGTENIKQKGGAIGLLTRHHLTRSTVYHQALILALIPFINPEHYPSA
11111111110000000000000000000000000000000000000000000000000000
00000000000000000000000000000000000000000000000000000000000000
0000100000000000111111100000000000000000000000000000000000011
>DM_validata684
MDVSFEFEHYQVRLIKSSDAVTIANYFMRNRHHLAPWEPKRSHAFFTPEGWKQRLLQLVELH
KHNLAFYFVVVDKNEHKIIGTVSYSNITRFPFHAGHVGYSLDSEYQGKGIMRRAVNVTIDWM
FKAQNLHRIMAAYIPRNEKSAKVLAALGFVKEGEAKKYLYINGAWEDHILTSKINDDWKP
10000000000000000000000000000000000000000000000000000000000000
00000000000000000000000000000000000000000000000000000000000000
000000000000000000000000000000000000000000000000000000000000
>DM_validata685
GAAASIQTTVNTLSERISSKLEQEANASAQTKCDIEIGNFYIRQNHGCNLTVKNMCSADADA
QLDAVLSAATETYSGLTPEQKAYVPAMFTAALNIQTSVNTVVRDFENYVKQTCNSSAVVDNK
LKIQNVIIDECYGAPGSPTNLEFINTGSSKGNCAIKALMQLTTKATTQIAPKQVAGTGVQ
11100000000000000000000000000000000000000000000000000000000000
00000000000000000000000000000000000000000000000000000000000000
000000000000000000000000000000000000000000000001111111111111
>DM_validata686
MAEPQPASSGLTDETAFSCCSDPDPSTKDFLLQQTMLRIKDPKKSLDFYTRVLGLTLLQKLD
FPAMKFSLYFLAYEDKNDIPKDKSEKTAWTFSRKATLELTHNWGTEDDETQSYHNGNSDPRG
FGHIGIAVPDVYSACKRFEELGVKFVKKPDDGKMKGLAFIQDPDGYWIEIILNPNKIATII

11111111000000000000000000000000000000000000000000000000000000
00000000000000000000000000000000000000000000000000000000000000
000000000000000000000000000000000000000000000000000000000000
>DM_validata687
EIPLCANLVPVPITNATLDQITGKWFYIASAFRNEEYNKSVQEIQATFFYFTPNKTEDTIFL
REYQTRQDQCIYNTTYLNVQRENGTISRYVGGQEHFAHLLILRDTKTYMLAFDVNDEKNWGL
SVYADKPETTKEQLGEFYEALDCLRIPKSDVVYTDWKKDKCEPLEKQHEKERKQEEGES
00000000000000000000000000000000000000000000000000000000000000
00000000000000000000000000000000000000000000000000000000000000
00000000000000000000000000000000000000000000000000011111111
>DM_validata688
SDSTAGTDAVAVGGTFQFHSPDGKMEIFYDEADRQQLPDIGGDSLMEEGTQINLSDFENQVV
ILNAWGQWCAPCRSESDDLQIIHEELQAAGNGDTPGGTVLGINVRDYSRDIAQDFVTDNGLD
YPSIYDPPFMTAASLGGVPASVIPTTIVLDKQHRPAAVFLREVTSKDVLDVALPLVDEA
11111111111100000000000000000000000000000000000000000000000000
00000000000000000000000000000011100000000000000000000000000000
00000000000000000000000000000000000000000000000000000000001
>DM_validata689
MYEVELKGYANDEIFEKVRETFEFMRKEIHEDIYYQHPCRDFSKTDEALRIRIKRFNGHNEV
FLTYKGPKIDEKSKTRLEIEVEIQEDVDKYFELLDRLGFKEVLKVVKTREKYYVEKGVTITL
DEVEGLGKFIEIETLVKEKDEIPEAVEKLEKILRELGVEKFERRSYLELLLEKRTELNI
11000000000000000000000000000000000000000000000000000000000000
00000001111111110000000000000000000000000000000000000000000000
00000000000000001111111100000000000000000000000000000000001
>DM_validata690
PSKFSKEQLRTFQMIHENFGRALSTYLSGRLRTFVDVEISIDQLTYEEFIRSVMIPSFIVIF
TGDVFEGSAIFEMRLDLFYTMLDIIMGGPGENPPNRPPTEIETSIMRKEVTNMLTLLAQAWS
DFQYFIPSIENVETNPQFVQIVPPNEIVLLVTASVSWGEFTSFINVCWPFSLLEPLLEK
00000000000000000000000000000000000000000000000000000000000000
00000000000000000000000000001111000000000000000000000000000000
00000000000000000000000000000000000000000000000000000000000
>DM_validata692
MSLSDDIKVLGISGSLRSGSYNSAALQEAIGLVPPGMSIELADISGIPLYNEDVYALGFPPA
VERFREQIRAADALLFATPEYNYSMAGVLKNAIDWASRPPEQPFSGKPAAILGASAGRFGTA
RAQYHLRQTLVFLDVHPLNKPEVMISSAQNAFDAQGRLLDDKARELIQQQLQALQLWVR
11111100000000000000000000000000000000000000000000000000000000
00000000000000000000000000000000000000000000000000000000000000
00000000000000000000000000000000000000000000000000000000111
>DM_validata694
APPSNLMQLPWRQGYSWQPNGAHSNTGSGYPYSSFDASYDWPRWGSATYSVVAAHAGTVRVL
SRCQVRVTHPSGWATNYYHMDQIQVSNGQQVSADTKLGVYAGNINTALCEGGSSTGPHLHFS
LLYNGAFVSLQGASFGPYRINVGTSNYDNDCRRYYFYNQSAGTTHCAFRPLYNPGLAL
10000000000000000000000000000000000000000000000000000000000000
00000000000000000000000000000000000000000000000000000000000000

00000000000000000000000000000000000000000000000000000001
>DM_validata695
MSETAPLPSASSALEDKAASAPVVGIIMGSQSDWETMRHADALLTELEIPHETLIVSAHRTP
DRLADYARTAAERGLNVIIAGAGGAAHLPGMCAAWTRLPVLGVPVESRALKGMDSLLSIVQM
PGGVPVGTLAIGASGAKNAALLAASILALYNPALAARLETWRALQTASVPNSPITEDK
111111111111111111100000000000000000000000000000000000000000
000000000000000000000000000000000000000000000000000000000000
00000000000000000000000000000000000000000000000000001111
>DM_validata696
MPYIEKLELKGFKSYGNKKVVIPFSKGFTAIVGANGSGKSNIGDAILFVLGGLSAKAMRASR
ISDLIFAGSKNEPPAKYAEVAIYFNNEDRGFPIDEDEVVIRRRVYPDGRSSYWLNGRRATRS
EILDILTAAMISPDGYNIVLQGDITKFIKMSPLERRLLIDDISGIAEYDSKKEKALEE
000000000000000000000000000000000000000000000000000000111111
111111111111110000000000000000000000000000000000000000000000
00000000000000000000000000000000000000000111111111111111
>DM_validata697
MKKIILTIGCPGSGKSTWAREFIAKNPGFYNINRDDYRQSIMAHEERDEYKYTKKKEGIVTG
MQFDTAKSILYGGDSVKGVIISDTNLNPERRLAWETFAKEYGWKVEHKVFDVPWTELVKRNS
KRGTKAVPIDVLRSMYKSMREYLGLPVYNGTPGKPKAVIFDVDGTLAKMNGRGPYDL
000000000000000000000000000000000000000000000000000000000000
000000000000000000000000000000000000000000000000000000000000
0000000000000000000000000001111111111111111111111111111
>DM_validata698
MMTMRGLLVGRMQPFHRGALQVIKSILEEVDELIICIGSAQLSHSIRDPFTAGERVMMLTKA
LSENGIPASRYYIIPVQDIECNALWVGHIKMLTPPFDRVYSGNPLVQRLFSEDGYEVTAPPL
FYRDRYSGTEVRRRMLDDGDWRSLLPESVVEVIDEINGVERIKHLAKKEVSELGGIS
111000000000000000000000000000000000000000000000000000000000
000000000000000000000000000000000000000000000000000000000001
1111100000000000000000000000000000000000000001111111111
>DM_validata699
AEHNPVVMVHGIGGASFNFAGIKSYLVSQGWSRDKLYAVDFWDKTGTNYNNGPVLSRFVQKV
LDETGAKKVDIVAHSMGGANTLYYIKNLDGGNKVANVVTLGGANRLTTGKALPGTDPNQKIL
YTSIYSSADMIVMNYLSRLDGARNVQIHGVGHIGLLYSSQVNSLIKEGLNGGGQNTN
110000000000000000000000000000000000000000000000000000000000
000000000000000000000000000000000000000000000000000000000000
0000000000000000000000000000000000000000000000000000000
>DM_validata700
MGSSTGFHHADHVNYSSNLNKEEILEQLLLSYEGLSDGQVNWVCNLSNASSLIWHAYKSLAV
DINWAGFYVTQASEENTLILGPFQGKVACQMIQFGKGVCGTAASTKETQIVPDVNKYPGHIA
CDGETKSEIVVPIISNDGKTLGVIDIDCLDYEGFDHVDKEFLEKLAKLINKSCVFK
111000000000000000000000000000000000000000000000000000000000
000000000000000000000000000000000000000000000000000000000000
000000000000000000000000000000000000000000000000000000
>DM_validata702

MILVTGGARSGKSRHAEALIGDAPQVLYIATSQILDDEMAARIQHHKDGRPAHWRTAECWRH
LDTLITADLAPDDAILLECITTMVTNLLFALGGENDPEQWDYAAMERAIDDEIQILIAACQR
CPAKVVLVTNEVGMGIVPENRLARHFRDIAGRVNQRLAAAADEVWLVVSGIGVKIK
000000000000000000000000000000000111111111000000000000011100
000000000000000000000000000000111000000000000000000000000000
000000000000000000000000000000000000000000000000000000
>DM_validata703
MTEMELKLIKIDTSHYFEKKPGLGERVDYAGRCFYNKFQRVNAMLTSSLIQKHLKREIEIAH
NLILRNDKVENIVFDYNGRNPERFYHKAQLLLREEGFMNFTAYNTKTPGHLHLYVHKGHTEL
GEGERLVKTLSMKLAQGLPKEWKVFPSNEWPKEFNILALPYEVFAKERGSSWAKHL
110000000000000000000000000000000000000000000000000000000000
000000000000000000000000000000000000000000000000000000000000
000000000000000000000000000000000000000000000000000011
>DM_validata704
SANENNLIWIDLEMTGLDPERDRIIEIATLVTDANLNILAEGPTIAVHQSDEQLALMDDWNV
RTHTASGLVERVKASTMGDREAELATLEFLKQWVPAGKSPICGNSIGQDRRFLFKYMPELEA
YFHYRYLDVSTLKELARRWKPEILDGFTKQGTHQAMDDIRESVAELAYYREHFIKL
100000000000000000000000000000000000000000000000000000000000
000000000000000000000000000000000000000000000000000000000000
000000000000000000000000000000000000000000000000000000
>DM_validata705
GMTRQRIAIDMDEVLADTLGAVVKAVNERADLNIKMESLNGKKLKHMIPEHEGLVMDILKEP
GFFRNLDVMPHAQEVVKQLNEHYDIYIATAAMDVPTSFHDKYEWLLEYFPFLDPQHFVFCGR
KNIILADYLIDDNPKQLEIFEGKSIMFTASHNVYEHRFERVSGWRDVKNYFNSIEK
110000000000000000000000000000000000000000111111110000000000
000000000000000000000000000000110000000000000000000000000000
000000000000000000000000000000000000000000000000000001
>DM_validata706
MAGNREDRKAKVIEVLNKARAMELHAIHQYMNQHYSLDDMDYGELAANMKLIAIDEMRHAEN
FAERIKELGGEPTTQKEGKVVTGQAVPVIYESDADQEDATIEAYSQFLKVCKEQGDIVTARL
FERIIEEEQAHLTYYENIGSHIKNLGDTYLAKIAGTPSSTGTASKGFVTATPAAE
111000000000000000000000000000000000000000000000000000000000
000000000000000000000000000000000000000000000000000000000000
00000000000000000000000000000000000000000000001111111
>DM_validata707
GSPEFPGRENLYFQGREPVLDGPYQPTTFNPPVDYWMLLAPTAAGVVVEGTNNTDRWLATIL
VEPNVTSETRSYTLFGTQEQITIANASQTQWKFIDVVKTTQNGSYSQYGPLQSTPKLYAVMK
HNGKIYTYNGETPNVTTKYYSTTNYDSVNMTAFCDFYIIPREEESTCTEYINNGL
111111111111111111100000000000000000000000000000000000000000
000000000000000000000000000000000000000000000000000000000000
00000000000000000000000000000000000000000000000000000
>DM_validata708
GEDKKGGTKPSNEAALTKTENLDFRLSFNKIKVTTDQNHFSGGTSIEQLKQWFGDPNKSEQR
NAGNITLDSYTWVKDGAVINAQLYKNSTVARSISNFSFSREAKIGKEDYDELKIGESYKKIV

EKLGEPDVLSQSMSSDKEEMQTVWSSGIKTKSSSATIELYFENGLLKNKTQKDLE
11111111111111111000000000000000000000000000000000000000000000
00000000000000000000000000000000000000000000000000000000000000
0000000000000000000000000000000000000000000000000000000
>DM_validata709
MHAGEAVQQLKKAFETVASFDFRDALSKASTPVTVVATNGPFGLAGLTCSAVCSVCDRPPTV
LLCINRKSYAAGIIKSNGVLSVNWLAAGQAVISQTFAGVGSVPMEERFADKGWQTIATGAPY
RMDAAVSFDCTIANIVDVGSHSVIFAEVVARNHAEECTPLIYHRRQYATTRSLAE
11111111111111000000000000000000000000000000000000000000000000
00000000000000000000000000000000000000000000000000000000000000
0000000000000000000000000000000000000000000000000000011
>DM_validata710
GMNQNKITAGGLEFLVRFAAPTDRLKINDLMIDTARWLKESGSTQWSDILHGFDVHNIEQRI
ELGEVALFETEAGALAGAMIIRKTPSDWDTDLWEDLAIDKAYYLHRIMVSRAFSGISLSKQM
IYFAEKLGIEMSVPFIRLDCIESNETLNQMYVRYGFQFSGKKNGFYLYQKELSQK
11110000000000000000000000000000000000000000000000000000000000
00000000000000000000000000000000000000000000000000000000000000
0000000000000000000000000000000000000000000000000000111
>DM_validata711
REATVLSYDGSMFMKIQLPVVMHTEAEDVSLRFRSQRAYGILMATTSRDSADTLRLELDAGR
VKLTVNLKGPETLFAGYNLNDNEWHTVRVVRRGKSLKLTVDDQQAMTGQMAGDHTRLEFHNI
ETGIITERRYLSSVPSNFIGHLQSLTFNGMAYIDLCKNGDIDYCELNARFGFRNI
11000000000000000000000000000000000000000000000000000000000000
00000000000000000000000000000000000000000000000000000000000000
0000000000000000000000000000000000000000000000000000111
>DM_validata712
QYRIIDFKRDKDGIPGRVATIEYDPNRSANIALINYADGEKRYIIAPKNLKVGMEIMSGPDA
DIKIGNALPLENIPVGTLVHNIELKPGRGGQLVRAAGTSAQVLGKEGKYVIVRLASGEVRMI
LGKCRATVGEVGNGGRTDKPFVKAGNKHHKMKARGTKWPNVRGVAMNAVDHPFG
00000000000000000000000000000000000000000000000000000000000000
00000000000000000000000000000000000000000000000000000000000000
000000000000000000000000000000000111110000000000000000
>DM_validata713
MKIIITGEPGVGKTTLVKKIVERLGKRAIGFWTEEVRDPETKKRTGFRIITTEGKKKIFSSK
FFTSKKLVGSYGVNVQYFEELAIPILERAYREAKKDRRKVIIIDEIGKMELFSKKFRDLVRQ
IMHDPNVNVVATIPIRDVHPLVKEIRRLPGAVLIELTPENRDVILEDILSLLER
00000000000000000000000000000000000001111110000000000000000000
00000000000000000000000000000000000000000000000000000000000000
000000000000000000000000000000000000000000000000000000
>DM_validata714
MSSGVDLGTENLYFQSNAMKDTFRLENQTIYFGTERAISASPQTIWRYLTETDKLKQWFPEL
EIGELGVNGFWRFILPDFEETMPFTDYAEEKYLGVTWDTGIIYFDLKEQAPHQTLLVFSESL
PENFTTPRHKDIAGWSIVLNRLKQVVETPDAAPEKIDFPQIENHYLEKLTNLEN
11111110000000000000000000000000000000000000000000000000000000

00000000000000000000000000000000000000000000000000000000000000
000000000000000000000000000000000000000000000000000000
>DM_validata715
EKQPNIDELKKRMEQSRLNKLRGDLDQLIESDPKLRALRPHLKIDLVQEGLRIQIIDSQNRP
MFKTGSAEVEPYMRDILRAIAPVLNGIPNRISLAGHTDDFPYANGEKGYSNWELSADRANAS
RRELVAGGLDNGKVLRVVGMAATMRLSDRGPDDAINRRISLLVLNKQAEQAILH
11100000000000000000000000000000000000000000000000000000000000
00000000000000000000000000000000000000000000000000000000000000
000000000000000000000000000000000000000000000000000000
>DM_validata716
AEAQNDPLLPGYSFNAHLVAGLTPIEANGYLDFFIDRPLGMKGYILNLTIRGQGVVKNQGRE
FVCRPGDILLFPPGEIHHYGRHPEAREWYHQWVYFRPRAYWHEWLNWPSIFANTGFFRPDEA
HQPHFSDLFGQIINAGQGEGRYSELLAINLLEQLLLRRMEAINESLHPPMDNR
11110000000000000000000000000000000000000000000000000000000000
00000000000000000000000000000000000000000000000000000000000000
00000000000000000000000000000000000000000001111111111
>DM_validata717
MDLGEFPAPDPSVLVQNFNISDFNGKWYITSGLNPTFDAFDCQLHEFHTEGDNKLVGNISWR
IKTLDSGFFTRSAVQKFVQDPNQPGVLYNHDNEYLHYQDDWYILSSKIENKPEDYIFVYYRG
RNDAWDGYGGAVVYTRSSVLPNSIIPELEKAAKSIGRDFSTFIRTDNTCGPEP
11111100000000000000000000000000000000000000000000000000000000
00000000000000000000000000000011111111000000000000000000000000
00000000000000000000000000000000000000000000000000011
>DM_validata718
GPVDKAGTTDNDNSQTDKTGPFSGLKFKQNSFLSTVPSVTNMHSMHFDARETFLGVIRKALE
PDTSTPFPVRRAFDGLRAEILPNDTIKSAALKAQCSDIDKHPELKAKMETLKEVITHHPQKE
KLAEIALQFAREAGLTRLKGETDYVLSNVLDGLIGDGSWRAGPAYESYLNKPG
11111111111111111111000000000000000000000000000000000000000000
00000000000000000000000000000000000000000000000000000000000000
00000000000000000000000000000000000000000111111111111
>DM_validata719
SSESESGGDFHEMREPRIEKVVVHMGIGHGGRDLANAEDILGEITGQMPVRTKAKRTVGEFD
IREGDPIGAKVTLRDEMAEEFLQTALPLAELATSQFDDTGNFSFGVEEHTEFPSQEYDPSIG
IYGLDVTVNLVRPGYRVAKRDKASRSIPTKHRLNPADAVAFIESTYDVEVSE
11111111100000000000000000000111110000000000000000000000000000
00000000000000000000000000000000000000000000011111111111111111
11100000000000000000000000000000011111111111111111111
>DM_validata720
RTEDLPKAVVFLEPQWYSVLEKDSVTLKCQGAYSPEDNSTQWFHNESLISSQASSYFIDAAT
VNDSGEYRCQTNLSTLSDPVQLEVHIGWLLLQAPRWVFKEEDPIHLRCHSWKNTALHKVTYL
QNGKDRKYFHHNSDFHIPKATLKDSGSYFCRGLVGSKNVSSETVNITITQGL
11110000000000000000000000000000000000000000000000000000000000
00000000000000000000000000000000000000000000000000000000000000
0000000000000000000000000000000000000000000000011111

>DM_validata721
GHMAGASHDIHPPKVEYFDLRDHTNTDPKGFVRHVDHYRVEPWGLYMARTSDHPQFHYLESW
LLPDLGLRASIFHYHPYHQRDQDHYVDIGTFTRGDDVWKSEDHYLDLVVRTGRDTELLDVDE
LMEAHTTGLLDTATAEQAILTATTAIDGIAAHGHDLGRWLASIGMPIDWRGS
111111111100000000000000000000000000000000000000000000000000
000000000000000000000000000000000000000000000000000000000000
0000000000000000000000000000000000000000000000000001
>DM_validata722
GMSRGHCILAHGFESGPDALKVTALAEVAERLGWTHERPDFTDLDARRDLGQLGDVRGRLQR
LLEIARAATEKGPVVLAGSSLGSYIAAQVSLQVPTRALFLMVPPTKMGPLPALDAAAVPISI
VHAWHDELIPAADVIAWAQARSARLLLVDDGHRLGAHVQAASRAFAELLQSL
100000000000000000000000000000000000000000000000000000000000
000000000011100000000000000000000000000000000000000000000000
0000000000000000000000000000000000000000000000000000
>DM_validata723
MTPLGPASSLPQSFLLKCLEQVRKIQGDGAALQEKLCATYKLCHPEELVLLGHSLGIPWAPL
SSCPSQALQLAGCLSQLHSGLFLYQGLLQALEGISPELGPTLDTLQLDVADFATTIWQQMEE
LGMAPALQPTQGAMPAFASAFQRRAGGVLVASHLQSFLEVSYRVLRHLAQP
111111000000000000000000000000000000000000000000000000000000
000000000000000000000000000000000000000000000000000000000000
000011111111000000000000000000000000000000000000000
>DM_validata724
MSLTLRLAEHRDLEAVVAIYNSTIASRMVTADTEPVTPEDRMEWFSGHTESRPLYVAEDENG
NVAAWISFETFYGRPAYNKTAEVSIYIDEACRGKGVGSYLLQEALRIAPNLGIRSLMAFIFG
HNKPSLKLFEKHGFAEWGLFPGIAEMDGKRYDLKILGRELSEGGSHHHHHH
100000000000000000000000001111111000000000000000000000000000
000000000000000000000000000000000000000000000000000000000000
000000000000000000000000000000000000000000111111111
>DM_validata725
MELDDFDPEDKEILSWDINDVKLPQNVKTTDWFQEWPDSYVKHIYSSDDRNAQRHLSSWAMR
NTNNHNSRILKKSCLGVVVCSRDCSTEEGRKIYLRPAICDKARQKQQRKSCPNCNGPLKLIP
CRGHGGFPVTNFWRHDGRFIFFQSKGEHDHPRPETKLEAEARRAMKKVHM
111111111111100000000000000000000000000000000000000000000000
000000000000000000000000000000000000000000000000000000000000
00000000000000000000000000000000000000000000001111
>DM_validata726
GMREMQIMSQPEQSRLDLFIDRMVSQRACLEHAIAQTAGLSGPVYELGLGNGRTYHHLRQHV
QGREIYVFERAVASHPDSTPPEAQLILGDIRETLPATLERFGATASLVHADLGGHNREKNDR
FARLISPLIEPHLAQGGLMVSSDRMYFEGLEELPLPPGAVVGRCFIYRRG
111111111110000000000000000000000000000000000000000000000000
000000000000000000000000000000000000000000000000000100000000
00000000000000000000000000000000000000000000000001
>DM_validata727
RADKASSIELKFDRNKGEVGDILIGTVRINNIKNFAGFQVNIVYDPKVLMAVDPETGKEFTS

STFPPGRTVLKNNAYGPIQIADNDPEKGILNFALAYSYIAGYKETGVAEESGIIAKIGFKIL
QKKSTAVKFQDTLSMPGAISGTQLFDWDGEVITGYEVIQPDVLSLGDEPY
11111000000000000000000000000000000000000000000000000000000000
00000000000000000000000000000000000000000000000000000000000000
00000000000000000000000000000000000000000000111111
>DM_validata728
PTNIHKIHEVQKKLQEEVSIVLIDIADIIVNPKKENGYSRDLYTLNSLIDSSISETYDNINN
TLLSDTRFFLEHMDIIKSQRDILENLYSYVSQLNSTPPQAHILSAFIHKIGYTEFEAETGNL
LLEELKRLMISMKNQPLPVDRTEFENRAILFLCLTELKQFLVNRKHAQML
11111000000000000000000000000000000000000000000000000000000000
00000000000000000000000000000000000000000000000000000000100000
00000000000000000000000000000000000000000000000000
>DM_validata729
NSSKSLEAELVRDRQELIDARKKELKAYMMMGVTAIKPLYDSDVNGSNKQAAKEILKAMRFE
SDGYFFAYDSQGINTLHAIKPSLEGKNLYDLKDENGVAVIAGLIDASQKGDGFLYFSWHKPT
INAQAPKLGYAEYLQKWDWVLGTGIYIDDIDQQVAMQRELRTQELNQHTI
11111110000000000000000000000000000000000000000000000000000000
00000000000000000000000000000000000000000000000000000000000000
00000000000000000000000000000001111111111111111111
>DM_validata730
MAAAAELSLLEKSLGLSKGNKYSAQGERQIPVLQTNNGPSLMGLTTIAAHLVKQANKEYLLG
STAEEKAMVQQWLEYRVTQVDGHSSKNDIHTLLMDLNSYLEDKVYLTGYNFTLADILLYYGL
HRFIVDLTVQEKEKYLNVSRWFCHIQHYPGIRQHLSSVVFIKNRLYTNSH
00000000000000000000000000000000000000000000000000000000000000
00000000000000000000001111000000000000000000000000000000000000
00000000000000000000000000000000000000000000001111
>DM_validata731
MEEAVLTGVATDKSEAKVTVLGISDKPGEAAKVFRALADAEINIDMVLQNVSSVEDGTTDIT
FTCPRSDGRRAMEILKKLQVQGNWTNVLYDDQVGKVSLVGAGMKSHPGVTAEFMEALRDVNV
NIELISTSEIRISVLIREDDLDAAARALHEQFQLGGEDEAVVYAGTGRH
10000000000000000000000000000000000000000000000000000000000000
00000000000000000000000000000000000000000000000000000000000000
0000000000000000000000000000000000111100000011111
>DM_validata732
RGSHHHHHHGSAPIKVGDAIPAVEVFEGEPGNKVNLAELFKGKKGVLFGVPGAFTPGSSKTH
LPGFVEQAEALKAKGVQVVACLSVNDAFVTGEWGRAHKAEGKVRLLADPTGAFGKETDLLLD
DSLVSIFGNRRLKRFSMVVQDGIVKALNVEPDGTGLTCSLAPNIISQL
11111111110000000000000000000000000000000000000000000000000000
00000000000000000000000000000000000000000000000000000000000000
000000000000000000000000000000000000000000000000
>DM_validata733
GHMIQESTMEMTNAQRLILSNQYKMMTMLDPANAERYRRLQTIIERGYGLQMRELDREFGEL
KEETCRTIIDIMEMYHALHVSWSNLQDQQSIDERRVTFLGFDAATEARYLGYVRFMVNVEGR
YTHFDAGTHGFNAQTPMWEKYQRMLNVWHACPRQYHLSANEINQIINA

111110000000000000000000000000000000000000000000000000000000
000000000000000000000000000000000000000000000000000000000000
000000000000000000000000000000000000000000000000
>DM_validata734
MNTQTRVSDTTVREITEWLYMEAELLDAGKYREWLALVTEDLSYVVPIRVTREREAVTDVVE
GMTHMDDDADSMEMRVLRLETEYAWAEDPPSRSRHFVTNVRVATGDSEDEFKVTSNLLLYRT
RGDVATYDVLSGERTDVLRRAGDSFLMAKRVVLLDQTTIMTHNLALIM
111110000000000000000000000000000000000000000000000000000000
000000000000000000000000000000000000000000000000000000000000
000000000000000000000000000000000000000000000000
>DM_validata735
MDYFTLFGLPARYQLDTQALSLRFQDLQRQYHPDKFASGSQAEQLAAVQQSATINQAWQTLR
HPLMRAEYLLSLHGFDLASEQHTVRDTAFLMEQLELREELDEIEQAKDEARLESFIKRVKKM
FDTRHQLMVEQLDNETWDAAADTCRKLRFLDKLRSSAEQLEEKLLDF
000000000000000000000000000001111111111111000000000000000000
000000000000000000000000000000000000000000000000000000000000
00000000000000000000000000000000000000000000000
>DM_validata736
GSHMGIQETDPLTQLSLPPGFRFYPTDEELMVQYLCRKAAGYDFSLQLIAEIDLYKFDPWVL
PNKALFGEKEWYFFSPRDRKYPNGSRPNRVAGSGYWKATGTDKIISTEGQRVGIKKALVFYI
GKAPKGTKTNWIMHEYRLIEPSRRNGSTKLDDWVLCRIYKKQSSAQK
000000011110000000000000000000000000000000000000000000000000
000000000000000000011111110000000000000000000000000000000000
00000000000000000000001111111100000000000011111
>DM_validata737
MQEKYNFGKVSSQHKNYSKIETMLRPKGFDKLDHYFRTELDIDLTDETIELLLNSVKAAFGK
LFYGAEQRARWNGRDFIALADLNITKALEEHIKNFQKIEQDMGVDELLEYIAFIPPVEMNVG
EDLKSEYRNIMGGLLLMHADVIKKATGERKPSREAMEFVAQIVDKVF
111111111111111111100000000000000000000000000000000000000000
000000000000000000000000000000000000000000000000000000000000
00000000000000000000000000000000000000000000000
>DM_validata738
RAVPGGSSPAWTQCQQLSQKLCTLAWSAHPLVGHMDLREEGDEETTNDVPHIQCGDGCDPQG
LRDNSQFCLQRIHQGLIFYEKLLGSDIFTGEPSLLPDSPVGQLHASLLGLSQLLQPEGHHWE
TQQIPSLSPSQPWQRLLLRFKILRSLQAFVAVAARVFAHGAATLSP
111111110000000000000000000000011111111111111100000000000000
000000000000000000000000000000000000000000000000000000111111
1111110000000000000000000000000000000000000011
>DM_validata739
MREVLVPYAGVSPSVDSTAFIAGNARIIGDVCIGKNASIWYGTVLRGDVDKIEVGEGTNIQD
NTVVHTDSMHGDTVIGKFVTIGHSCILHACTLGNNAFVGMGSIVMDRAVMEEGSMLAAGSLL
TRGKIVKSGELWAGRPAKFLRMMTEEEILYLQKSAENYIALSRGYL
000000000000000000000000000000000000000000000000000000000000
000000111100000000000000000000000000000000000000000000000000

0000000000000000000000000000000000000000000000
>DM_validata740
GHMNEYFFPKLTAVEALAPYRLRTTWSTGEVLEVDVGDILRKIPDLAPILDPEAFARVHIAE
WEGSVEWFDTEFGRDNVYAWAKEQAGEVSHEMFGDWMHRNNLSLTTAAEALGISRRMVSYYR
TAHKIIPRTIWLACLGWEATRPETKTLPRTLPAAYAKGVSASLSGS
11111000000000000000000000000000000000000000000000000000000000
00000000000000000000000000000000000000000000000000000000000000
0000000000000000000000000000000011111111111111
>DM_validata741
GPLGSGGLFFNALKNCKENFTVLQTIRQQQSTLNGSWVALLQTRNTLNRAGIRYMMDQNNIG
SGSTVAELMESASISLKQAEKNWADYEALPRDPRQSTAAAAEIKRNYDIYHNALAELIQLLG
AGKINEFFDQPTQGYQDGFEKQYVAYMEQNDRLHDIAVSDNNASYS
11111111110000000000000000000000000000000000000000000000000000
00000000000000000000000000000000000000000000000000000000000000
0000000000000000000000000000000000000000011111
>DM_validata742
GSKGSRIELGDVTPHNIKQLKRLNQVIFPVSYNDKFYKDVLEVGELAKLAYFNDIAVGAVCC
RVDHSQNQKRLYIMTLGCLAPYRRLGIGTKMLNHVLNICEKDGTFDNIYLHVQISNESAIDF
YRKFGFEIIETKKNYYKRIEPADAHVLQKNLKVPSGQNADVQKTDN
11110000000000000000000000000000000000000000000000000000000000
00000000000000000000000000000000000000000000000000000000000000
0000000000000000000000000000000011111111111111
>DM_validata743
MSEQKQDVAATEEQQPVLQIQRIYVKDVSFEAPNLPHIFQQEWKPKLGFDLSTETTQVGDDL
YEVVLNISVETTLEDSGDVAFICEVKQAGVFTISGLEDVQMAHCLTSQCPNMLFPYARELVS
NLVNRGTFPALNLSPVNFDALFVEYMNRQQAENAEEKSEEEQTKH
11111111111111110000000000000000000000000000000000000000000000
00000000000000000000000000000000000000000000000000000000000000
000000000000000000000000000111111111111111111
>DM_validata744
MAHHHHHHMSANGKISVPEAVVNWLFKVIQPIYNDGRTTFHDSLALLDNFHSLRPRTRVFTH
SDGTPQLLLSIYGTISTGEDGSSPHSIPVIMWVPSMYPVKPPFISINLENFDMNTISSSLPI
QEYIDSNGWIALPILHCWDPAAMNLIMVVQELMSLLHEPPQDQAP
11111111111111100000000000000000000000000000000000000000000000
00000000000000000011111100000000000000000000000000000001111000
000000000000000000000000000000000000000000011
>DM_validata745
MVRLEDYGTWDEALKRLEASRKALLALLREADPAWLSAPLREGAWTPLMVAEHVALVEDSTA
RVLRRLRRLAAGENLPPVPVKPGEFKDGKPQAPEGVRPKGGLSLEEVLALLDRARAFLLEEV
AKADPQNPATFPHPFFGELNPLGWLRAAAYHEAHHLKALQASLPR
11111111000000000000000000000000000000011100000000000000000000
00000000000111111111111111111111111111111000000000000000000000
000000000000000000000000000000000000000000011
>DM_validata746

MSPGERFLDWLKRLQGQKAWTAARAAFRRSLAFPPGAYPRAMPYVEPFLAKGDWRQEEREAH
YLVAALYALKDGDHQVGRTLARALWEKAQGSASVEKRFLALLEADRDQIAFRLRQAVALVEG
GIDFARLLDDLLRWFSPERHVQARWAREYYGAGASEEEKKKEVEA
00000000000000000000000000000000000000000000000000000000000000
00000000000000000000000000000000000000000000000000000000000000
000000000000000000000000000000001111111111111
>DM_validata747
GSPEFMGEQPIFSTRAHVFQIDPNTKKNWVPTSKHAVTVSYFYDSTRNVYRIISLDGSKAII
NSTITPNMTFTKTSQKFGQWADSRANTVYGLGFSSEHHLSKFAEKFQEFKEAARLAKEKSQE
KMELTSTPSQESAGGDLQSPLTPESINGTDDERTPDVTQNSEPR
11100000000000000000000000000000000000000000000000000000000000
00000000000000000000000000000000000000000000000000000000000000
00000000011111000000000011111111111111111111
>DM_validata748
MSTIPSEIINWTILNEIISMDDDDSDFSKGLIIQFIDQAQTTFAQMQRQLDGEKNLTELDNL
GHFLKGSSAALGLQRIAWVCERIQNLGRKMEHFFPNKTELVNTLSDKSIINGINIDEDDEEI
KIQVDDKDENSIYLILIAKALNQSRLEFKLARIELSKYYNTNLH
00000000000000000000000000000000000000000000000000000000000000
00000000000000000000000000000000000000000000000000000000000000
00111100000000000000000000000000000000000000
>DM_validata749
MEPPNLYPVKLYVYDLSKGLARRLSPIMLGKQLEGIWHTSIVVHKDEFFFGSGGISSCPPGG
TLLGPPDSVVDVGSTEVTEEIFLEYLSSLGESLFRGEAYNLFEHNCNTFSNEVAQFLTGRKI
PSYITDLPSEVLSTPFGQALRPLLDSIQIQPPGGSSVGRPNGQS
11110000000000000000000000000000000000000000000000000000000000
00000000000000000000000000000000000000000000000000000000000000
00000000000000000000000011111111111111111111
>DM_validata750
PKVNLYATFRDLTGKSQLELPGATVGEVLENLVRAYPALKEELFEGEGLAERVSVFLEGRDV
RYLQGLSTPLSPGATLDLFPPVAGGGFERTFGAFPPWLLERYLEEWGGTREGEGVYRLPGAV
VRFREVEPLKVGSLSIPQLRVEVEGEEAERWFERIAFAASRGGG
00000000000000000000000000000000000000000000000000000000000000
00000000000000000000011110000000000000000000000000000000000000
00000000000000000000000000000000000000001111
>DM_validata751
MKLQVLPLSQEAFSAYGDVIETQQRDFFHINNGLVERYHDLALVEILEQDCTLISINRAQPA
NLPLTIHELERHPLGTQAFIPMKGEVFVVVVALGDDKPDLSTLRAFITNGEQGVNYHRNVWH
HPLFAWQRVTDFLTIDRGGSDNCDVESIPEQELCFALEHHHHHH
00000000000000000000000000000011110000000000000000000000000000
00000000000000000000000000000000000000000000000000000000000000
00000000000000000011000000000000000011111111
>DM_validata752
CLAEGTRIFDPVTGTTHRIEDVVDGRKPIHVVAAAKDGTLHARPVVSWFDQGTRDVIGLRIA
GGAILWATPDHKVLTEYGWRAAGELRKGDRVAQPRRFDGFGDSAPIPARVQALADALDDKFL

HDMLAEELRYSVIREVLPTRRARTFDLEVEELHTLVAEGVVVHN
000000000000000000000000000000000000000000000000000000000000
000000000000000000000000000000000000000011111111111111111111
11111000000000000000000000000000000000000000
>DM_validata753
TYNGCSSSEQSALAAAASAAQSYVAESLSYLQTHTAATPRYTTWFGSYISSRHSTVLQHYTD
MNSNDFSSYSFDCTCTAAGTFAYVYPNRFGTVYLCGAFWKAPTTGTDSQAGTLVHESSHFTR
NGGTKDYAYGQAAAKSLATMDPDKAVMNADNHEYFSENNPAQS
111100000000000000000000000000000000000000000000000000000000
000000000000000000000000000000000000000000000000000000000000
0000000000000000000000000000000000000000000
>DM_validata754
MPLLDSFKVDHTKMNAPAVRIAKTMLTPKGDNITVFDLRFCIPNKEILSPKGIHTLEHLFAG
FMRDHLNGDSIEIIDISPMGCRTGFYMSLIGTPNEQKVSEAWLASMQDVLGVQDQASIPELN
IYQCGSYTEHSLEDAHEIAKNVIARGIGVNKNEDLSLDNSLLK
111111110000000000000000000000000000000000000000000000000000
000000000000000000000000000000000000000000000000000000000000
0000000000000000000000000000000111111111111
>DM_validata755
GSPGISGGGGGILSLLCVGVKKAKFDGAQEKFNTYVTLKVQNVESTTIAVRGSQPSWEQDFM
FEINRLDLGLTVEVWNKGLIWDTMVGTVWIPLRTIRQSNEEGPGEWLTLDSQAIMADSEICG
TKDPTFHRILLDAHFELPLDIPEEEARYWAKKLEQLNAKLNSS
111111111110000000000000000000000000000000000000000000000000
000000000000000000000000000000000000000000000000000000000000
0000000000000000000000000000000000000000000
>DM_validata756
MSDKFNQFINRVLSHEGGYANHPKDPGGETNWGITKRTAQANGYNGSMRAMTREQAISIYRK
AFWERYRADQMPEAVAFQFFDACVNHGYGNAARMLQRAAGVPDDGVIGAVSLKAINSLPEND
LLLRFNAERLVFYTKLGTFTSFGKGWVRRVAQNLIHASADNTD
110000000000000000000000000000000000000000000000000000000000
000000000000000000000000000000000000000000000000000000000000
0000000000000000111111100000000000000001111
>DM_validata757
GMGYTLAEFLAHAIALETEAAERYVELADMMEAHNNLDTATVFRDMARFSTLHGDEIKQRSR
ALELPKLMSWQYRWKTPPEVGDENDIHYLMTPYHALRYARDNEIRGMEYYKEAAANSADPEV
KRLGADFAAEEAEHVVALDKWIEKTPRPSITWSEDADPAQCVD
111000000000000000000000000000000000000000000000000000000000
000000000000000000000001110000000000000000000000000000000000
0000000000000000000000000000000111111111111
>DM_validata758
GMALTPDQMPDRAEVVFTCNGKAVGKMRNELDVAMVKPFEERFALATDEGAFHGGDASAPPP
LALFIAGLTGCVMTQIRAFAKRLKVTVTDLDVECRVVWDWAKAGPVYETGPKSFEIDIILHS
PDPIEAQQALIEAAKKGCFLEQTLGQANTIRHRLKVGDTFIDA
111111111110000000000000000000000000000000000000001111100000

000000000000000000000000000000000000000000000000000000000000000
000000000000000000000000000000000000000000
>DM_validata759
ADQVDPQLKFAMQRDLGIFPTQLPQYLQTEKLARTQAAAIEREFGAQFAGSWIERNEDGSFK
LVAATSGARKSSTLGGVEVRNVRYSLKQLQSAMEQLDAGANARVKGVSKPLDGVQSWYVDPR
SNAVVVKVDDGATDAGVDFVALSGADSAQVRIESSPGKLQTT
111111111111111100000000000000000000000000000000000000000000000
000000000000000000000000000000000000000111111111111000000000000
000000000000000000000000000000000011111111
>DM_validata760
APPRLICDSRVLERYLLEAKEAEKITTGCAEHCSLNEKITVPDTKVNFYAWKRMEVGQQAVE
VWQGLALLSEAVLRGQALLVKSSQPWEPLQLHVDKAVSGLRSLTTLLRALGAQKEAISPPDA
ASAAPLRTITADTFRKLFRVYSNFLRGKLKLYTGEACRTGDR
000000000000000000000000000000000000000000000000000000000000000
000000000000000000000000000000000000000000000000000000000000001
111111000000000000000000000000000000001111
>DM_validata761
ESNPDPTTYPDVELSPPPRISLRSLLTAQPVKNDHYDSHNYLSTHWELIDYKGKEYEKLRDG
GTLVQFKVVGAAKCFAFLGKGTTDCKDTDHTVFNLIPTNTGAFLIKDALLGFCITSHDFDDL
KLEPCGGSVSGRTFSLAYQWGILPPFGPSKILIPPVRRNQGS
111100000000000000000000000000000000000000000000000000000000000
000000000000000000000000000000000000000000000000000000000000000
000000000000000000000000000000000011111111
>DM_validata762
MVVAVYPGTFDPLTRGHEDLVRRASSIFDTLVVGVADSRAKKPFFSLEERLKIANEVLGHYP
NVKVMGFTGLLKDFVRANDARVIVRGLRAVSDFEYEFQMAGMNRYLLPDVETMFMTPSDQYQ
FISGTIVREIAQLGGDVSKFVFPSVEKWLTEKVAAMAQGPSA
000000000000000000000000000000000000000000000000000000000000000
000000000000000000000000000000000000000000000000000000000000000
000000000000000000000000000000000000011111
>DM_validata763
GSSGSSGMTPAVKLLEKNKISFQIHTYEHDPAETNFGDEVVKKLGLNPDQVYKTLLVAVNGD
MKHLAVAVTPVAGQLDLKKVAKALGAKKVEMADPMVAQRSTGYLVGGISPLGQKKRLPTIID
APAQEFATIYVSGGKRGLDIELAAGDLAKILDAKFADIARRD
111111100000000000000000000000000000000000000000000000000000000
000000000000000000000000000000000000000000000000000000000000000
000000000000000000000000000000000000000001
>DM_validata764
DGTSQLAELVDAAAERLEVADPVAAFKWRAQLPIEDSGRVEQQLAKLGEDARSQHIDPDYVT
RVFDDQIRATEAIEYSRFSDWKLNPASAPPEPPDLSASRSAIDSLNNRMLSQIWSHWSLLSA
PSCAAQLDRAKRDIVRSRHLDSLYQRALTTATQSYCQALPPA
110000000000000000000000000000000000000000000000000000000000000
000000000000000000000000000000000000000000000000000000000000000
000000000000000000000000000000000000000000

>DM_validata765
MLNMWKVRELVDKATNVVMNYSEIESKVREATNDDPWGPSGQLMGEIAKATFMYEQFPELMN
MLWSRMLKDNKKNWRRVYKSLLLLAYLIRNGSERVVTSAREHIYDLRSLENYHFVDEHGKDQ
GINIRQKVKELVEFAQDDDRLREERKKAKKNKDKYVGVSSD
11111111111111110000000000000000000000000000000000000000000000
00000000000000000000000000000000000000000000000000000000000000
00000000000000000000000000001111111111111
>DM_validata766
SNAMAVDMFIKIGDVKGESKDKTHAEEIDVLAWSWGMSQSGSMHMGGGGGAGKVNVQDLSFT
KYIDKSTPNLMMACSSGKHYPQAKLTIRKAGGENQVEYLIITLKEVLVSSVSTGGSGGEDRL
TENVTLNFAQVQVDYQPQKADGAKDGGPVKYGWNIRQNVQA
11110000000000000000000000000000000000000001111100000000000000
00000000000000000000000000000000000000000000000000000000000000
00000000000000000000000000000000000000000
>DM_validata768
SNAMQSDHKEKIAILVDVQNVYYTCREAYRSNFDYNQFWYVATQEKEVVSAKAYAIASNDPK
QRQFHHILRGVGFEVMLKPYIQRRDGSAKGDWDVGITLDAIEIAPDVDRVILVSGDGDFSLL
VERIQQRYNKKVTVYGVPRLTSQTLIDCADNFVAIDDDFLL
11100000000000000000000000000000000000000000000000000000000000
00000000000000000000000000000000000000000000000000000000000000
00000000000000000000000000000000000000000
>DM_validata771
GMGMEHKTYEADLFVWCQQQADGLRALSRSRRDLPDDLDLEHIAEEIEDMGRSELREATSLV
RQICVRVIMAMSAPEAPDRARWRSEVVSWHNLLLDTITPGMIDRIDIGVIWRRAVSEAKAAL
IEINVAPQAGLSFQAPLPADHFLDEDFDYDATVARLGPTA
11110000000000000000000000000000000000000000000000000000000000
00000000000000000000000000000000000000000000000000000000000000
0000000000000000000000000000000000000011
>DM_validata772
SCVRDNSLVRDISQMPQSSYGIEGLSHITVAGALNHGMKEVEVWLQTISPGQRTPIHRHSCE
EVFTVLKGKGTLLMGSSSLKYPGQPQEIPFFQNTTFSIPVNDPHQVWNSDEHEDLQVLVIIS
RPPAKIFLYDDWSMPHTAAVLKFPFVWDEDCFEAAKEQL
00000000000000000000000000000000000000000000000000000000000000
00000000000000000000000000000000000000000000000000000000000000
000000000000000000000000000000000001111
>DM_validata775
MGSDKIHHHHHHMKVIETKYSGKLEVAEDRLIAFDQGIPAFEDEKEFVLLPFAAGTPYYTLQ
STKTVDLAFIIVNPFSFFPEYRVKLPEATIAQLNITNENDVAIFSLLTVKEPFSETTVNLQA
PIVINANKQMGKQLVLGDTAYNRKQPLFQKELVLAKEAK
11111111100000000000000000000000000000000000000000000000000000
00000000000000000000000000000000000000000000000000000000000000
000000000000000000000000000000000011111
>DM_validata776
MVEPSLVLYGAPYERAVEVLEETLRETGARYALLIDRKGFVLAHKEALWAPKPPPLDTLATL

VAGNAAATQALAKLLGEARFQEEVHQGERMGLYVDEAGEHALLVLVFDETAPLGKVKLHGKR
ASEALARIAEEALANPPRLALDTEYREGAEALLDDLLRN
11111000000000000000000000000000000000000000000000000000000000
00000000000000000000000000000000000000000000000000000000000000
000000000000001111111111111111111111111
>DM_validata777
VRYRFLRLAPDEEGEGGRAESRILECRRLRAPAEIARALELRAGETVVTIRRQLSMNHMPTV
IDDLWLPGTHFRGLTLELLTASKAPLYGLFESEFGVSMVRADEKLRAVAASPEIAPLLGVEP
GRPLLQVDRISYTYGDRPMEVRRGLYLTDHYHYRNSLN
00000000000011111100000000000000000000000000000000000000000000
00000000000000000000000000000000000000000000000000000000000000
00000000000000000000000000000000000000
>DM_validata778
MSDLDTPTPSPHPVLLNLEQFLPYRLSVLSNRISGNIAKVYGDRYGMAIPEWRVITILALYP
GSSASEVSDRTAMDKVAVSRAVARLLERGFIRRETHGDDRRRSMLALSPAGRQVYETVAPLV
NEMEQRLMSVFSAEEQQTLERLIDRLAKDGLPRMASKD
11111111111000000000000000000000000000000000000000000000000000
00000000000000000000000000000000011111111100000000000000000000
00000000000000000000000000000000000111
>DM_validata779
GTKQRADIVMSEAEIADFVNSSRTGTLATIGPDGQPHLTAMWYAVIDGEIWLETKAKSQKAV
NLRRDPRVSFLLEDGDTYDTLRGVSFEGVAEIVEEPEALHRVGVSVWERYTGPYTDECKPMV
DQMMNKRVGVRIVARRTRSWDHRKLGLPHMSVGGSTAP
10000000000000000000000000000000000000000000000000000000000000
00000000000000000000000000000000000000000000000000000000000000
00000000000000000000000000000000000001
>DM_validata780
MSGMRVYLGADHAGYELKQRIIEHLKQTGHEPIDCGALRYDADDDYPAFCIAAATRTVADPG
SLGIVLGGSGNGEQIAANKVPGARCALAWSVQTAALAREHNNAQLIGIGGRMHTVAEALAIV
DAFVTTPWSKAQRHQRRIDILAEYERTHEAPPVPGAPA
11000000000000000000000000000000000000000000000000000000000000
00000000000000000000000000000000000000000000000000000000000000
00000000000000000000000000000000000111
>DM_validata781
MLTMQEIKAHYRFTDEDAELLGSLFPLAETNKERLADQFYDYLLGIPETAEFLKEDLVLQKL
KQTHQDWFVSLFAGSYDNRYIHNLQKIGHAHVRVGLNAHYVNVAMNVVRQFTLSIIQDNFPD
PEERRQRREAVEKILDINLDIMSASYREEEMRKFFVSH
00000000000000000000000000000000000000000000000000000000000000
00000000000000000000000000000000000000000000000000000000000000
00000000000000000000000000001111111111
>DM_validata782
MAQAFVDATWPQAAKAAQSLGVPAHFLVAQAALETGWGKSQIRNKDGTPSYNLFNIKAGSNW
TGKVVEARTVEYENGQRKVRVERFRAYDSYEQAFQDYADLVGNSPRYAKVAGKTDGHAFARA
LQEGGYATDPSYADKLARVINGNALRQRLMASAASARG

10000000000000000000000000000000000000000000000000000000000000
00000000001111110000000000000000000000000000000000000000000000
00000000000000000000000000000000000000
>DM_validata783
GSHMESLTQYIPDEFSMLRFGKKFAEILLKLHTEKAIMVYLNGDLGAGKTTLTRGMLQGIGH
QGNVKSPTYTLVEEYNIAGKMIYHFDLYRLADPEELEFMGIRDYFNTDSICLIEWSEKGQGI
LPEADILVNIDYYDDARNIELIAQTNLGKNIISAFSN
11100000000000000000000000000000000000000000000000000000000000
00000000000000000000000000000000000000000000000000000000000000
0000000000000000000000000000000000000
>DM_validata785
RSVAHLFIDETSSEVLDELYRVSKEYTHSRPQAQRVIKDLIKVAIKVAVLHRNGSFGPSELA
LATRFRQKLRQGAMTALSFGEVDFTFEAAVLAGLLTECRDVLLELVEHHLTPKSHGRIRHVF
DHFSDPGLLTALYGPDFTQHLGKICDGLRKLLDEGKL
11111100000000000000000000000000000000000000000000000000000000
00000000000000000000000000000000000000000000000000000000000000
0000000000000000000000000000000000000
>DM_validata786
MQVFIMRHGDAALDAASDSVRPLTTNGCDESRLMANWLKGQKVEIERVLVSPFLRAEQTLEE
VGDCLNLPSSAEVLPELTPCGDVGLVSAYLQALTNEGVASVLVISHLPLVGYLVAELCPGET
PPMFTTSAIASVTLDESGNGTFNWQMSPCNLKMAKAI
00000000000000000000000000000000000000000000000000000000000000
00000000000000000000000000000000000000000000000000000000000000
0000000000000000000000000000000011111
>DM_validata787
MGSDKIHHHHHHMAHIFVYGTLKRGQPNHKVMLDHSHGLAAFRGRGCTVESFPLVIAGEHNI
PWLLYLPGKGHCVTGEIYEVDEQMLRFLDDFEDCPSMYQRTALQVQVLEWEGDGDPGDSVQC
FVYTTATYAPEWLFLPYHESYDSEGPHGLRYNPRENR
11111111110000000000000000000000000000000000000000000000000000
00000000000000000000000000000000000000000000000000011100000000
0000000000000000000000000000000000000
>DM_validata788
GMSDIKKLGSSWIINWFFGFNQIPTNEDSSIYMKSVLTCAKADGVISPEEKDWALGFCASWG
VADWVIEDLKTYEADEALEEVIARSPQVSMAQRDILLSAIWVSAADGELHEKEKAKIRKMAT
ILGIKEEIVDQLEQLYYYEAALRQKRLNLLYPQKSPY
10000000000000000000000000000000000000000000000000000000000000
00000000000000000000000000000000000000000000000000000000000000
0000000000000000000000000000000000000
>DM_validata789
MLDIVLYEPEIPQNTGNIIRLCANTGFRLHLIEPLGFTWDDKRLRRSGLDYHEFAEIKRHKT
FEAFLESEKPKRLFALTTKGCPAHSQVKFKLGDYLMFGPETRGIPMSILNEMPMEQKIRIPM
TANSRSMNLSNSVAVTVYEAWRQLGYKGAVNLPEVK
00000000000000000000000000000000000000000000000000000000000000
00000000000000000000000000000000000000000000000000000000000000

00000000000000000000000000000001111
>DM_validata790
SPGQGTQSENSCTHFPGNLPNMLRDLRDAFSRVKTFFQMKDQLDNLLLKESLLEDFKGYLGC
QALSEMIQFYLEEVMPQAENQDPDIKAHVNSLGENLKTLRLRLRRCHRFLPCENKSKAVEQV
KNAFNKLQEKGIYKAMSEFDIFINYIEAYMTMKIRN
11111111111111110000000000000000000001111111000000000000000000
00000000000000000000000000000000000000000000011100000000000000
000000000000000000000000000000000000
>DM_validata791
MNDQIKIRKATKEDWEKIYQLYNSLSDEDLYLRFFHLYRITEEDAKKIASNEDHVTFLAEVD
GKVVGEASLHKDGEFSLVVHRNYRTLGIGTLLVKTLIEEAKKSGLSTVKFYTLPENTPMIKI
GRKLGFKMRFYEDEVYGEMRLTERELNVNLATFSAP
11110000000000000000000000000000000000000011111111110000000000
00000000000000000000000000000000000000000000000000000000000000
000000000000000000000011111111111111
>DM_validata792
MTDAVLELPAATFDLPLSLSGGTQTTLRAHAGHWLVIYFYPKDSTPGSTTEGLDFNALLPEF
DKAGAKILGVSRDSVKSHDNFSAKQGFAFPLVSDGDEALCRAFDVIKEKNMYGKQVLGIERS
TFLLSPEGQVVQAWRKVKVAGHADAVLAALKAHAKQ
11000000000000000000000000000000000000000000000000000000000000
00000000000000000000000000000000000000000000000000000000000000
000000000000000000000000000000000001
>DM_validata793
TKVKREATKVQRDISAFKKVMQNISLAVNKFNVDIERYVGGDASHLLADGNVLIKATLDGVQ
SLQNEPPLSSMEALALVGPVQDLSNQIMLAIQNLIDKKEPLVQAGFGGKVENNLRQQEEAAQ
KLSELVSTKVPHELADISRQLSDGIAAGIKKGIDAF
11111000000000000000000000000000000000000000000000000000000000
00000000000000000000000000000000000000000000000000000000000000
000000000000000000000000000000000000
>DM_validata794
GSMADEALFLLLHNEMVSGVYKSAEQGEVENGRCITKLENMGFRVGQGLIERFTKDTARFKD
ELDIMKFICKDFWTTVFKKQIDNLRTNHQGIYVLQDNKFRLLTQMSAGKQYLEHASKYLAFT
CGLIRGGLSNLGIKSIVTAEVSSMPACKFQVMIQKL
00000000000000000000000000000000000000000000000000000111111111
00000000000000000000000000000000000000000000011111110000000000
000000000000000000000000000000000001
>DM_validata795
GHMNAQARFSQNLLDQGSHPTSEKLLSVLRPASGHVADALGITEGENVIHLRTLRRVNGVAL
CLIDHYFADLTLWPTLQRFDSGSLHDFLREQTGIALRRSQTRISARRAQAKECQRLEIPNMS
PLLCVRTLNHRDGESSPAEYSVSLTRADMIEFTMEH
10000000000000000000000000000000000000000000000000000000000000
00000000000000000000000000000000000000000000000000000000000000
000000000000000000000000000000000000
>DM_validata796

MEIRKKLVVPSKYGTKCPYTMKPKYITVHNTYNDAPAENEVNYMITNNNEVSFHVAVDDKQA
IQGIPWERNAWACGDGNGPGNRESISVEICYSKSGGDRYYKAENNAVDVVRQLMSMYNIPIE
NVRTHQSWSGKYCPHRMLAEGRWGAFIQKVKSGNV
000000000000000000000000000000000000000000000000000000000000
000000000000000000000000000000000000000000000000000000000000
00000000000000000000000000000000011
>DM_validata798
NSFLQDVPYWMLQNRSEYITQGVDSSHIVDGKKTEEIEKIATKRATIRVAQNIVHKLKEAYL
SKTNRIKQKITNEMFIQMTQPIYDSLMNVDRLGIYINPNNEEVFALVRARGFDKDALSEGLH
KMSLDNQAVSILVAKVEEIFKDSVNYGDVKVPIAM
100000000000000000000000000000000000000000000000000000000000
000000000000000000000000000000000000000000000000000000000000
00000000000000000000000011111111111
>DM_validata799
MDFECQFVCELKELAPVPALLIRTQTTMSELGSLFEAGYHDILQLLAGQGKSPSGPPFARYF
GMSAGTFEVEFGFPVEGGVEGSGRVVTGLTPSGKAASSLYIGPYGEIEAVYDALMKWVDDNG
FDLSGEAYEIYLDNPAETAPDQLRTRVSLMLHES
111110000000000000000000000000000000000000000000000000000000
000000000000000000000000000000000000000000000000000000000000
0000000000000000000000000000000000
>DM_validata800
TGFLVSFMVDARGGSMRGSRHNGLRVVIPPRTCAAPTRITCRLVKPQKLSTPPPLAEEEGLA
SRIIALGPTGAQFLSPVIVEIPHFASHGRGDRELVVLRSENGSVWKEHRSRYGESYLDQILN
GMDEELGSLEELEKKRVCRIITTDFPLYFVIMSR
100000000000000000000000000000000000000000000000000000000000
000000000000000000000000111111110000000000000000000000000000
0000000000000000000000000000000000
>DM_validata801
MDITNIKEMNYEEVFSITITVDKPILIGQDDIVGRRQLIPIISGKVSGNNFNGKVLPGGIDS
QIVRPDGKCELSARYAIRLDDGAAIYIENNGIRTVPDEYIEAVKSGEFVDPNAYYFRTIPTF
ETYSPKYKWMMNHIFVCCASRLPENVLLKFYKIS
111111000000000000000000000000000000000000000000000000000000
000000000000000000000000000000000011111111111111000000000000
0000000000000000000000110000000000
>DM_validata802
LENENQLTHQFPTLSRWNPMFISDVHKISFHPHLQRYIGFWMGFPIRWIQIVGYIAAIDIYE
GKHVLTVDDCSGMVLRVVFIIQDDFSMSKRAISMSPGNVVCVFGKINSFRSEVELIAQSFEE
LRDPNDEWKAWQKRMRYKKNLTKISKNHHSIIRT
111111111111110000000000000000001110000000000000000000000000
000000000000000000000000000000000000000000000000000000000000
0000000000000000000000000000011111
>DM_validata803
GMEYQLQQLASLTLVGIKETYENGRQAQQHIAGFWQRCYQEGVIADLQLKNNGDLAGILGLC
IPELDGKMSYMIAVTGDNSADIAKYDVITLASSKYMVFEAQGAVPKAVQQKMEEVHHYIHQY

QANTVKSAPFFELYQDGDTTSEKYITEIWMPVKG
00000000000000000000000000000000000000000000000000000000000000
00000000000000001111100000000000000000000000000000000000000000
0000000000000000000000000000000000
>DM_validata804
EQAPQYVDETISLSAKTLFGFDKDSLRAEAQDNLKVLAQRLSRTNIQSVRVEGHTDFMGSDK
YNQALSERRAYVVANNLVSNGVPVSRISAVGLGESQAQMTQVCEAEVAKLGAKVSKAKKREA
LIACIEPDRRVDVKIRSIVTRQVVPAHNHHQHLE
11100000000000000000000000000000000000000000000000000000000000
00000000000000000000000000000000000000000000000000000000000000
0000000000000000000111111111111111
>DM_validata805
MGSDKIHHHHHHMSQLEKAQAEYAGFIQEFQSAIISTISEQGIPNGSYAPFVIDDAKNIYIY
VSGLAVHTKNIEANPLVNVLFVDDEAKTNQIFARRRLSFDCTATLIERESQKWNQVVDQFQE
RFGQIIEVLRGLADFRIFQLTPKEGRFVIGFGA
11111111111111111111110000000000000000000000000000000000000000
00000000000000000000000000000000000000000000000000000000000000
000000000000000000000000000000000
>DM_validata806
VRSSSRTPSDMPVAHVVANPQAEGQLQWLNRRANALLANGVELRDNQLVVPSEGLYLIYSQV
LFSGQGCPSTHVLLTHTISRISTTHNQPVNLLSAIRSPCQRETPEGAEANPWYEPIYLGGVF
QLEPGDRLSAEINRPDYLDFAESGQVYFGIIAL
11111100000000000000000000000000000000000000000000000000000000
00000000000000000000000000000000000000001111111000000000000000
000000000000000000000000000000000
>DM_validata807
GMPDSPTLLDLFAEDIGHANQLLQLVDEEFQALERRELPVLQQLLGAKQPLMQQLERNGRAR
AEILREAGVSLDREGLARYARERADGAELLARGDELGELLERCQQANLRNGRIIRANQASTG
SLLNILRGQDAPSLYDSRGGTASSSRQRPLSQA
10000000000000000000000000000000000000000000000000000000000000
00000000000000000000000000000000000000000000000000000110000000
000000011111111111111111111111111
>DM_validata808
GFEERSGVVPCGTPWGQWYQTLEEVFIEVQVPPGTRAQDIQCGLQSRHVALAVGGREILKGK
LFDSTIADEGTWTLEDRKMVRIVLTKTKRDAANCWTSLLESEYAADPWVQDQMQRKLTLERF
QKENPGFDFSGAEISGNYTKGGPDFSNLGNDGT
11000000000000000000000000000000000000000000000000000000000000
00000000000000000000000000000000000000000000000000000000000000
000000000111111111111111111111111
>DM_validata809
AASSAQCDFGGPFQAYKSVNGPGNGGYYLRKTTKPGTPECAYVLVPQNTLSEGQSTSFTYGK
LQNGQMIQLTATVTVNGDKIEVTGAGQDLSGTTTVLFSDYRSCDVMRGPDGNYELWVHSSAI
NLQSYGCCDTKFAQVAGGRPIHHTWQTYCPPLP
11111000000000000000000000000000001000000000000000000000000000

00000000000000000000000001110000000000000000000000000000000000
000000000000000000000000000000000
>DM_validata810
YRQSQSRAARLRLLVDTGQELIQLPPEAMRKCVLQRACAFVAMDHGLLLEWGADNGVQTTAR
HGSKERLSTLETTADPLAIGPQWLERPGTHLPCVLLLPLRGADEGSFGTLVLANSVAISAPD
GEDIESLQLLATLLAAHLENNRLLEALVARDRT
00000000000000000000000000000000000000000000000000000100000000
00000000001110000000000000000000000000000000000000000000000000
000000000000000000000000000011111
>DM_validata812
AIRSDFSNEDIYDNIDPDTISFPPKIATTDLFLPLFFHFGSTRQFMDKLHEVISGDYEPSQA
EKLVQDLCDETGIRKNFSTSILTCLSGDLMVFPRYFLNMFKDNVNPPPNVPGIWTHDDDESL
KSNDQEQIRKLVKKHGTGRMEMRKRFFEKDLL
11000000000000000000000000000000000000000000000000000000000000
00000000000000000000000000000000000000000000000000000000000000
00000000000000000000000000000111
>DM_validata813
MSDVIEGRLKELGFTLPVAAAPAANYVPFTISGNLLYVSGQLPMESGKIAVTGLVGRDVDVA
SAQRAAELCAVNILAQVKAALNGDLSKIRRVIKLNGFVASVPEFVEQHLVINGASNLIATVL
GEPGRHARAAVGMASLPFNASVEIDAIVEIDV
00000000000000000111111100000000000000000000000000000000000000
00000000000000000000000000000000000000000000000000000000000000
00000000000000000000000000000000
>DM_validata814
GMTAPTLSRAAMEKVIRTYYDGCNEADEAKMIACFVPEAVHYFPAGMYGGAFRGAAQIAHRW
RTAVETLGSYWTIDALVIDAETAEAAIEWTHFKTNQDKVLRGAECVEFDRASGLIREIRAFY
ASPQAEGIARLELGDFDYAGRGYRVTSPRKPA
11110000000000000000000000000000000000000000000000000000000000
00000000000000000000000000000000000000000000000000000000000000
00000000000000000000000000000000
>DM_validata815
GMSSSGIEIAKPFVTATTNVLSTMAGIQPIPGQPYVKKNNVAKGDVSAVVGITGHKNGSISV
TFTKQCAIAVVKAMLGDDIQDIIQDTKDAVGEVTNMISGQARAALSEMGMTFQGATPSVIMG
DGHTISHVTKSPVIAIPFKTNHGEFTVEFCLE
11111100000000000000000000000000000000000000000000000000000000
00000000000000000000000000000000000000000000000000000000000000
00000000000000000000000000000000
>DM_validata816
SSGEHEASRVLRERDYRWEGTEEEAYKAEGTHFSGARRQTLVGRPAGQEAPAFETRYFEVEP
GGYTTLERHEHTHVVMVVRGHAEVVLDDRVEPLTPLDCVYIAPHAWHQIHATGANEPLGFLC
IVDSDRDRPQRPDADDLARMCADPAVARRIRT
11111000000000000000000001111111111100000000000000000000000000
00000000000000000000000000000000000000000000000000000000000000
00000000000000000000000000000000

>DM_validata817
SEMSVVEYEVVSKNLTSKMSHELLFSVKKRWFVKPFRHDRQLGKLHYKLLPGNYIKFGLYVL
KNQDYARFEIAWVHVDKDGKIEERTVYSIETYWHIFIDIENDLNCPYVLAKFIEMRPEFHKT
AWVEESNYSIAEDDIQMVESIKRYLERKIASD
111000000000000000000000000000000000000000000000000000000000
000000000000000000000000000000000000000000000000000000000000
00000000000000000000000000000111
>DM_validata818
MEEINTKEVAQRITTELKRYSIPQAIFAQRVLCRSQGTLSDLLRNPKPWSKLKSGRETFRRM
WKWLQEPEFQRMSALRLAACKRKEQEHGKDRGNTPKKPRLVFTDVQRRTLHAIFKENKRPSK
ELQITISQQLGLELSTVSNFFMNARRRSLDKW
000000000000000000000000000000000000000000000000000000000000
000000000000000001111111111111111111000000000000000000000000
00000000000000000000000000000001
>DM_validata819
MAREVKLTKAGYERLMQQLERERERLQEATKILQELMESSDDYDDSGLEAAKQEKARIEARI
DSLEDILSRAVILEEGSGEVIGLGSVVELEDPLSGERLSVQVVSPAEANVLDTPMKISDASP
MGKALLGHRVGDVLSLDTPKGKREFRVVAIHG
110000000000000000000000000000000000000000000000000000000000
000000000000000000000000000000000000000000000000000000000000
00000000000000000000000000000000
>DM_validata820
GMPSEKQPQSKGNKMAILKLDEHLYISPQLTKADAEQIAQLGIKTIICNRPDREEESQPDFA
QIKQWLEQAGVTGFHHQPVTARDIQKHDVETFRQLIGQAEYPVLAYCRTGTRCSLLWGFRRA
AEGMPVDEIIRRAQAAGVNLENFRERLDNARV
111111111111100000000000000000000000000000000000000000000000
000000000000000000000000000000000000000000000000000000000000
00000000000000000000000000000000
>DM_validata821
MPFSISGRKGSEMMAKLDELKQKLTAKQIQAAYLLVENELMESNNEEKRTQDEMANELGINR
TTLWEWRTKNQDFIAFKSEVADSFLAEKREQVYSKLMQLILGPQPSVKAMQLYMQRFGLLTD
KKVIEGDLGNATRTNAEIEEQLQKLKKLTGE
111111111111111111111100000000000000001111111000000000000000
000000011111000000000000000000000000000000000000000000000000
0000000001111111111111111111111
>DM_validata822
MSRSQLTILTNICLIEDLETQRVVMQYRAPENNRWSGYAFPGGHVENDEAFAESVIREIYEE
TGLTIQNPQLVGIKNWPLDTGGRYIVICYKATEFSGTLQSSEEGEVSWVQKDQIPNLNLAYD
MLPLMEMMEAPDKSEFFYPRRTEDDWEKKIF
000000000000000000000000000011111000000000000000000000000000
000000000000000000000000000000000000000000000000000000000000
0000000000000000000000000000000
>DM_validata823
MVKEFNTQTELSVRLEALWAVLSKDFITVVPKVLPHIVKDVQLIEGDGGVGTILIFNFLPEV

SPSYQREEITEFDESSHEIGLQVIEGGYLSQGLSYYKTTFKLSEIEEDKTLVNVKISYDHDS
DIEEKVTPTKTSQSTLMYLRRLERYLSNGSA
000000000000000000000000000000000000000000000000000000000000
000000000000000000000000000000000000000000000000000000000011
1111100000000000000000000001111
>DM_validata824
GMTTPPARTAKQRIQDTLNRLELDVDAWVSTAGADGGAPYLVPLSYLWDGETFLVATPAASP
TGRNLSETGRVRLGIGPTRDLVLVEGTALPLEPAGLPDGVGDTFAEKTGFDPRRLTTSYLYF
RISPRRVQAWREANELSGRELMRDGEWLVTD
100000000000000000000000000000000000000000000000000000000000
000000000000000000000000000000000000000000000000000000000000
0000000000000000000000000000111
>DM_validata825
RLHAGVWGLKVRYEGSFEVSKTPEEVFEFLTDPKRFSRAFPGFKSVEVEDGSFTIELRLSLG
PLRGDARVRASFEDLEKPSKATVKGSGRGAGSTLDFTLRFAVEPSGGGSRVSWVFEGNVGGL
AASMGGRVLDSLARRMINDVISGVKRELGEA
111000000000000000000000000000000000000000000000000000000000
000000000000000000000000000000000000000000000000000000000000
0000000000000000000000000000000
>DM_validata826
GMDDLTNLAARLRLLEDREEIRELIARYGPLADSGDAEALSELWVEDGEYAVVGFATAKGRA
AIAALIDGQTHRALMADGCAHFLGPATVTVEGDTATARCHSVVFRCVSGTFGSHRVSANRWT
FRRTPAGWRAVRRENALLDGSAAARALLQFR
000000000000000000000000000000000000000000000000000000000000
000000000000000000000000000000000000000000000000000000000000
0000000000000000000000000000001
>DM_validata827
GISYSVEADPDTTAKAMLRERQMSFKHSKAIAREIKGKTAGEAVDYLEAVIEGDQPVPFKQH
NSGVGHKSKVDGWDAGRYPEKASKAFLDLLENAVGNADHQGFDGEAMTIKHVAAHKVGEQQG
RKPRAMGRASAWNSPQVDVELILEEPEVED
000000000000000000000000000000000000000000000000000000000000
000000000000000000000000000000000000000000000000000000000000
0000000000000000000000001111
>DM_validata828
AEELVKPYNGDPFVGHLSTPISDSGLVKTFIGNLPAYRQGLSPILRGLEVGMAHGYFLIGPW
VKLGPLRDSDVANLGGLISGIALILVATACLAAYGLVSFQKGGSSSDPLKTSEGWSQFTAGF
FVGAMGSAFVAFFLLENFLVVDGIMTGLFN
111000000000000000000000000000000000000000000000000000000000
000000000000000000000000000000000000000000000000000000000000
000000000000000000000000000000
>DM_validata829
KHVEPQDAISPDNYMDMLGLEARDRTMYELVIYRKNDKDKGPWKRYDLNGRSCYLVGRELGH
SLDTDLDDRTEIVVADIGIPEETSSKQHCVIQFRNVRGILKCYVMDLDSSNGTCLNNVVIPG
ARYIELRSGDVLTLSEFEEDNDYELIFMNV

000000000000000000000000000000000000000000000000000000000000
111111111000000000000000000000000000000000000000000000000000
000000000000000000000000000000
>DM_validata830
SPQVRTAHIGDVPVLVRLMSEFYQEAGFALPHDAAIRAFKALLGKPDLGRIWLIAEGTESVG
YIVLTLGFSMEYGGLRGFVDDFFVRPNARGKGLGAAALQTVKQGCCDLGVRALLVETGPEDH
PARGVYSRAGFEESGRMLLGQALAPPIHEA
000000000000000000000000000000000000000000000000000000000000
000000000000000000000000000000000000000000000000000000011111
111000000000000000000000000001
>DM_validata832
MSLMIKVEDMMTRHPHTLLRTHTLNDAKHLMEALDIRHVPIVDANKKLLGIVSQRDLLAAQE
SSLQRSAQGDSLAFETPLFEVMHTDVTSVAPQAGLKESAIYMQKHKIGCLPVVAKDVLVGII
TDSDFVTIAINLLELQEESEPDELDEEQEG
100000000000000000000000000000000000000000000000000000000000
000011111111100000000000000000000000000000000000000000000000
000000000000000000111111111111
>DM_validata833
LSSKYSRNTELRRVEDNDIYRLAKILDENSCWRKLMSIIPKGMDVQACSGAGCLNFPAEIKK
GFKYTAQDVFQIDEAANRLPPDQSKSQMMIDEWKTSGKLNERPTVGVLLQLLVQAELFSAAD
FVALDFLNESTPARPVDGPGALISLELLE
000000000000000000000000000000000000000000000000000000000000
000000000000000000000000000000000000000000000000000000000000
00000000000000000000000000111
>DM_validata834
GAMDSSEMSTICDKTLNPSFCLKFLNTKFASANLQALAKTTLDSTQARATQTLKKLQSIIDG
GVDPRSKLAYRSCVDEYESAIGNLEEAFEHLASGDGMGMNMKVSAALDGADTCLDDVKRLRS
VDSSVVNNSKTIKNLCGIALVISNMLPRN
111100000000000000000000000000000000000000000000000000000000
000000000000000000000000000000000000000000000000000000000000
00000000000000000000000000000
>DM_validata835
MNIHVTIREATEGDLEQMVHMLADDVLGRKRERYEKPLPVSYVRAFKEIKKDKNNELIVACN
GEEIVGMLQVTFTPYLTYQGSWRATIEGVRTHSAARGQGIGSQLVCWAIERAKERGCHLIQL
TTDKQRPDALRFYEQLGFKASHEGLKMHF
111000000000000000000000000000000000000000000000000000000000
000000000000000000000000000000000000000000000000000000000000
00000000000000000000000000000
>DM_validata836
GHMTWKPNVTVAAVIEQDDKYLLVEEIPRGTAIKLNQPAGHLEPGESIIQACSREVLEETGH
SFLPEVLTGIYHWTCASNGTTYLRFTFSGQVVSFDPDRKLDTGIVRAAWFSIDEIRAKQAMH
RTPLVMQCIEDYHAGKRYPLDILQYYDGS
111110000000000000000000000000000000000000000000000000000000
000000000000000000000000000000000000000000000000000000000000

00000000000000000000000000000
>DM_validata837
GMCVEKTPWELVIDFHGHTCPDIALGYRIAQLAQREMGIRPAPDSECLVKAYTQSCALDAIQ
VLNKATIGRHALIIEETHRYMYQFHFTGTQDIHQFTVSPAVLDHLETLRHPDLSPRERQNKV
LEGVQYVLTLEESAFCHYDKIPGQLSKIV
111100000000000000000000000000000000000000000000000000000000
000000000000000000000000000000000000000000000000000000000000
00000000000000000000000000001
>DM_validata838
YFQGMLKNIDPALNADVLHALRAMGHGDTLVISDTNFPSDSVARQTTVGKVLHIDNVSAARA
MKAILSVLPLDTPLQPSVGRMEVMGAPDQLEPVQVEVQQEIDAAEGKSAPMYGIERFAFYEK
AKQAYCVITTGETRFYGCFLLTKGVIPPK
000000000000000000000000000000000000000000000000000000000000
000000000000000000000000000000000000000000000000000000000000
00000000000000000000000000011
>DM_validata839
MDTTTATTARTDTLLQLDNQLCFALYSANLAMHKLYRGLLKALDLTYPQYLVMLVLWETDER
SVSEIGERLYLDSATLTPLLKRLQAAGLVTRTRAASDERQVIIALTETGRALRSKAGAVPEQ
VFCASASSLDELRQLKQELEKLRSSLGAG
111111111111111100000000000000000000000000000000000000000000
000000000000000000000000000000001111110000000000000000000000
00000000000000000000001111111
>DM_validata840
MVIPAEANIIVGYSHFIKTVEDLNEIIRTHVPGSKYGIGFSEASGDRLIRYDGNDDDLVKAC
IENIRRISAGHTFVILIRNAYPINILNAVKMCQEVGSIFAATANPLQIIVYKGERGNGVLGV
IDGYSPVGVESDADIEKRRQFLRRIGYKE
000000000000000000000000000000000000000000000000000000000000
000000000000000000000000000000000000000000000000000000000000
00000000000111111111111111111
>DM_validata841
DRLGQPPSGYGVLLSVHEDKTVDVFTSGRKMRLTCSPNIDTDTLALGQTVRLNEALTIVEAG
TYEQVGEISTLREVLDDGLRALVVGHADEERIVWLAAPLAAVFADPEADIIAYDADSPTRKL
RPGDSLLVDTKAGYAFERIPKAEVEDLVL
111111000000000000000000000000000000000000000000000000000000
000000000000000000000000000000000000000000000111111111111111
11000000000000000000000111111
>DM_validata842
MDYCNKKHTAEEYVKISDNNYVPFPEAFSDGGITWQLLHSSPETSSWTAIFNCPAGSSFASH
IHAGPGEYFLTKGKMEVRGGEQEGGSTAYAPSYGFESSGALHGKTFFPVESQFYMTFLGPLN
FIDDNGKVIASIGWAEAQGAWLATKNEAA
000000000000000000000000000000000000000000000000000000000000
000000000000000000000000000000000000000000000000000000000000
00000000000000000000000001111
>DM_validata843

MDIKRYCPVTDSELPADHVYFKFRSEIEAAEAYLGLAISEGIKVRETREILDIIDTVYNSLS
DPESKLNDFQEKRLNFTEEDWYDIKEKANNGNRWSLYMFLARSAVDSAVYWSYRMKETEEFK
EIVKEEMISKLLKAGYVILRESLGEVRL
000000000000000000000000000000000000000000000000000000000000
000000000000000000000000000000000000000000000000000000000000
0000000000000000000000001111
>DM_validata844
PHMSGLKKFFPYSTNVLKGAAADIALPSLAGKTVFFYFSASWCPPCRAFTPQLIDFYKAHAE
KKNFEVMLISWDESAEDFKDYYAKMPWLALPFEDRKGMEFLTTGFDVKSIPTLVGVEADSGN
IITTQARTMVVKDPEAKDFPWPNVEAKK
111000000000000000000000000000000000000000000000000000000000
000000000000000000000000000000000000000000000000000000000000
0000000000000000000000011111
>DM_validata845
MSEKYIVTWDMLQIHARKLASRLMPSEQWKGIIAVSRGGLVPGALLARELGIRHVDTVAISS
YDHDNQRELKVLKRAEGDGEGFIVIDDLVDTGGTAVAIREMYPKAHFVTIFAKPAGRPLVDD
YVVDIPQDTWIEQPWDMGVVFVPPISGR
000000000000000000000000000000000000000000000000000000000011
111111111111111100000000000001110000000000000000000000000000
0000000000000000000000000000
>DM_validata846
MSQVTEQSVRFQTALASIKLIQASAVLDLTEDDFDFLTSNKVWIATDRSRARRCVEACVYGT
LDFVGYPRFPAPVEFIAAVIAYYVHPVNIQTACLIMEGAEFTENIINGVERPVKAAELFAFT
LRVRAGNTDVLTDAEENVRQKLRAEGVM
111110000000000000000000000000000000000000000000000000000000
000000000000000000000000000000000000000000000000000000000000
0000000000000011111111111111
>DM_validata847
MGSSSSGRENLYFQGHMADRQLLHTAHIPVRWGDMDSYGHVNNTLYFQYLEEARVAWFETLG
IDLEGAAEGPVVLQSLHTYLKPVVHPATVVVELYAGRLGTSSLVLEHRLHTLEDPQGTYGEG
HCKLVWVRHAENRSTPVPDSIRAAIAGS
111111111111111111110000000000000000000000000000000000000000
000000000000000000000000000000000000000000000000000000000000
0000000000000000000000000011
>DM_validata848
GSMETCENVDCGPGKKCRMNKKNKPRCVCAPDCSNITWKGPVCGLDGKTYRNECALLKARCK
EQPELEVQYQGKCKKTCRDVFCPGSSTCVVDQTNNAYCVTCNRICPEPSSSEQSLCGNDGVT
YSSACHLRKATCLLGRSIGLAYEGKCIK
111100000000000000000000000000000000000000000000000000000000
000000000000000000000000000000000000000000000000000000000000
0000000000000000000000000000
>DM_validata849
GQQPVNHLVKEIDMLLKEYLLSGDISEAEHCLKELEVPHFHHELVYEAIVMVLESTGESAFK
MILDLLKSLWKSSTITIDQMKRGYERIYNEIPDINLDVPHSYSVLERFVEECFQAGIISKQL

RDLCPSRSRGRKRFVSEGDGGRLKPESY
110000000000000000000000000000000000000000000000000000000000
000000000000000000000000000000000000000000000000000000000000
0000000111111111111111111111
>DM_validata850
MVDEVAKKKSKKEKDKDSKLEKALKAQNDLIWNIKDELKKVCSTNDLKELLIFNKQQVPSGE
SAILDRVADGMVFGALLPCEECSGQLVFKSDAYYCTGDVTAWTKCMVKTQTPNRKEWVTPKE
FREISYLKKLKVKKQDRIFPPETSASVA
111111111100000000000000000000000000000000000000000000000000
000000000000000000000000000000000000000000000000000000000000
0000000000000000000001111111
>DM_validata853
MMSKNILDMRNTVQIGIVVRDIEESLQNYAEFFGVEKPQWFWTDDYSKAHTKFNGRPTKARA
KLAFFELGPLQLELIEPDENPSTWREFLDKNGEGIHHIAFVVKDMDRKVEELYRKGMKVIQK
GDFEGGRYAYIDTLRALKVMIELLENY
100000000000000000000000000000000000000000000000000000000000
000000000000000000000000000000000000000000000000000000000000
000000000000000000000000000
>DM_validata855
MGSDKIHHHHHHMKSYRKELWFHTKRRREFINITPLLEECVRESGIKEGLLLCNAMHITASV
FINDDEPGLHHDFEVWLEKLAPEKPYSQYKHNDTGEDNADAHLKRTIMGREVVIAITDRKMD
LGPWEQVFYGEFDGMRPKRVLVKIIGE
111111111111000000000000000000000000000000000000000000000000
000000000000000000000000000000000000000000000000000000000000
000000000000000000000000000
>DM_validata856
GPLGSNIEQYIHDLDSNSFELDLQFSEDEKRLLLEKQAGGNPWHQFVENNLILKMGPVDKRK
GLFARRRQLLLTEGPHLYYVDPVNKVLKGEIPWSQELRPEAKNFKTFFVHTPNRTYYLMDPS
GNAHKWCRKIQEVWRQRYQSHPDAAVQ
111111000000000000000000000000000000000000000000000000000000
000000000000000000000000000000000000000000000000000000000000
000000000000000000011111111
>DM_validata857
GSHMASTPDKFNDGNLNIAYAKPTTQSSVDYNGDPNRAVDGNRNGNFNSGSVTHTRADNPSW
WEVDLKKMDKVGLVKIYNRTDAETQRLSNFDVILYDNNRNEVAKKHVNNLSGESVSLDFKEK
GARYIKVKLLTSGVPLSLAEVEVFRES
111111111111110000000000000000000000000000000000000000000000
000000000000000000000000000000000000000000000000000000000000
000000000000000000000000000
>DM_validata858
GMANKVEHRLSEQQMKALTDLPLVFLITHDQSKSWPITHAISWVYAKDETTIRFAIEADSLL
VKTLADHPVFTLIFFADQSTYSLTCTDVAAWETTARLPLKVALYEGQIKEVRDILFYGAAVS
DRPRVYKTYDEAAAMQLDQQIQDILKG
111100000000000000000000000000000000000000000000000000000000

00000000000000000000000000000000000000000000000000000000000000
000000000000000000000000000
>DM_validata859
EKVAVETSVYLSELEWKSASTGYGEIQKDASCDGNTITLKGENGEKVSYDKGIGTHAHSEIV
YSLEGLDYYDYFETFVGVDQEMAGTVASISFEVYLDNEKVFDSGLMTGDTTQKHVKVPIAGK
NTLKLVVKDGGDSIGSDHGSFGDAKLT
11110000000000000000000000000000000000000000000000000000000000
00000000000000000000000000000000000000000000000000000000000000
000000000000000000000000000
>DM_validata860
VQTVTTEDGETVKVFEDLQGFETFIANETEDDDFDHLHCKLNYYPPFVLHESHEDPEKISDA
ANSHSKKFVRHLHQHIEKHLLKDIKQAVRKPELKFHEKSKEETFDKITWHYGEETEYHGRPF
KIDVQVVCTHEDAMVFVDYKTHPVGAN
11111000000000000000000000000000000000000000000000000000000000
00000000000000000000000000000000000000000000000000000000000000
000000000000000000000000000
>DM_validata861
GMTDTNLVEMRAIERMMFDYSYHLDMNHPEELAALFVEDCEVSYAPNFGATGRDAYKKTLEG
IGTFFRGTSHHNSNICIDFVSETEANVRSVVLAIHRYTKERPDGILYGQYFDTVVKVDGQWK
FKRRELRTTMTTDYHVRAANPIGRAE
10000000000000000000000000000000000000000000000000000000000000
00000000000000000000000000000000000000000000000000000000000000
00000000000000000000000000
>DM_validata862
MWRRHYKLVPQIDTRDCGPAVLASVAKHYGSNYSIAYLRELSKTNKQGTTALGIVEAAKKLG
FETRSIKADMTLFDYNDLTYPFIVHVIKGKRLQHYYVVYGSQNNQLIIGDPDPSVKVTRMSK
ERFQSEWTGLAIFLAPQPNYKPHKGE
11110000000000000000000000000000000000000000000000000000000000
00000000000000000000000000000000000000000000000000000000000000
00000000000000000111111111
>DM_validata863
DSESDCTGSEPVDAFQAFSEGKEAYVLVRSTDPKARDCLKGEPAGEKQDNTLPVMMTFKQGT
DWASTDWTFTLDGAKVTATLGQLTQNREVVYDSQSHHCHVDKVEKEVPDYEMWMLDAGGLEV
EVECCRQKLEELASGRNQMYPHLKDC
11111000000000000000000000000000000000000000000000000000000000
00000000000000000000000000000000000000000000000000000000000000
00000000000000000000000000
>DM_validata864
MSSDAISLRAAGPGDLPGLLELYQVLNPSDPELTTQEAGAVFAAMLAQPGLTIFVATENGKP
VATATLLIVPNLTRAARPYAFIENVVTLEARRGRGYGRTVVRHAIETAFGANCYKVMLLTGR
HDPAVHAFYESCGFVQNKTGFQIRQD
11111000000000000000000000000000000000000000000000000000000000
00000000000000000000000000000000000000000000000000000000000000
00000000000000000000000001

>DM_validata865
MPSFVNPEKCDGCKALERTACEYICPNDLMTLDKEKMKAYNREPDMCWECYSCVKMCPQGAI
DVRGYVDYSPLGGACVPMRGTSDIMWTVKYRNGKVLRFKFAIRTTPWGSIQPFEGFPEPTEE
ALKSELLAGEPEIIGTSEFPQVKKKA
10000000000000000000000000000000000000000000000000000000000000
00000000000000000000000000000000000000000000000000000000000000
00000000000000000000000000
>DM_validata866
GMELKDIMHILEDMKVGVFATLDEYGNPHARHAHITAANEEGIFFMTSPETHFYDQLMGDQR
VAMTAISEEGYLIQVVRVEGTARPVENDYLKTVFADNPYYQHIYKDESSDTMQVFQIYAGHG
FYHSLTQGHKYIFSIGQGEHSEVRAL
10000000000000000000000000000000000000000000000000000000000000
00000000000000000000000000000000000000000000011111000000000000
00000000000000001111100000
>DM_validata867
MATVYDVPGDLLVERVAQRLKEIPEIKPPEWAPFVKTGRHKERLPEQEDWWYYRVASILRRV
YLDGPVGIERLRTYYGGRKNRGHAPERFYKAGGSIIRKALQQLEAAGFVEKVPGKGRVITPK
GRSFLDKIATELKKELEEIIPELKKY
10000000000000000000000000000000000000111111000000000000000000
00000000000000000111100000000000000000000000000000000000000000
00000000000000000000000000
>DM_validata868
ATWTCINQQLNPKTNKWEDKRLLYSQAKAESNSHHAPLSDGKTGSSYPHWFTNGYDGNGKLI
KGRTPIKFGKADCDRPPKHSQNGMGKDDHYLLEFPTFPDGHDYKFDSKKPKENPGPARVIYT
YPNKVFCGIVAHQRGNQGDLRLCSH
00000000001111111000000000000000000000000000000000000000000000
00000000000000000000000000000000000000000000000000000000000000
0000000000000000000000000
>DM_validata870
MDSYSITNVKYLDPTELHRWMQEGHTTTLREPFQVVDVRGSDYMGGHIKDGWHYAYSRLKQD
PEYLRELKHRLLEKQADGRGALNVIFHCMLSQQRGPSAAMLLLRSLDTAELSRCRLWVLRGG
FSRWQSVYGDDESVTAGYLPDLWR
11110000000000000000000000000000000000000000000000000000000000
00000000000000000000000000000000000000000000000000000000000000
000000000000000000000000
>DM_validata871
GSLHEEADHRVTDHGPRPFVVNIEDETKRNRAFRRALWTGDHLQVTLMSIQVGEDIGLEIHP
HLDQFLRVEEGRGLVQMGHRQDNLHFQEEVFDDYAILIPAGTWHNVRNTGNRPLKLYSIYAP
PQHPHGTVHETKAIAMAAEEHHHL
11111111110000000000000000000000000000000000000000000000000000
00000000000000000000000000000000000000000000000000000000000000
000000000000000000111111
>DM_validata872
AAAALRDQLTALLSSMFSQGLVDEQFQQLQMLQDEGGTPGFVSEVVTLFCDDADRIINEIAT

LLEQPVVNFDKVDAYVHQLKGSSASVGAQKVKFTCMQFRQFCQDKSRDGCLMALAVVRNDFY
DLRNKFQTMLQLEQQIQAYDPKQQ
1000000000000000000000000000000001111000000000000000000000000
0000000000000000000000000000000000000000000000000000000000000
000000000000000001111111
>DM_validata873
MDTPNKDDSIIRFSVSLQQNLLDELDNRIIKNGYSSRSELVRDMIREKLVEDNWAEDNPNDE
SKIAVLVVIYDGGQRELNQRMIDIQHASGTHVLCTTHIHMDEHNCLETIILQGNSFEIQRLQ
LEIGGLRGVKFAKLTKASSFEYNE
1111111100000000000000000000000000000000000000000000000000000
0000000000000000000000000000000000000000000000000000000000000
000000000000000000111111
>DM_validata874
MLVKTISRTIESESGFLQPTLDVIATLPADDRSKKIPISLVVGFKQEASLNSSSSLSCYYYA
IPLMRDRHINLKSGGSNVVGIPLLDTKDDRIRDMARHMATIISERFNRPCYVTWSSLPSEDP
SMLVANHLYILKKCLDLLKTELGE
0000000000001111100000000000000000000000000000001111100000000000
0000011111111110000000000000000000000000000000000000000000000
000000000000000000000001
>DM_validata875
MVITKIVGHIDDLSHQIKKVDWLEVEWEDLNKRILRKETENGTDIAIKLENSGTLRYGDVLY
ESDDTLIAIRTKLEKVYVIKPQTMQEMGKMAFEIGNRHTMCIIEDDEILVRYDKTLEKLIDE
VGVSYEQSERRFKEPFKYRGHQH
0000000000000000000000000000000000000000000000000000000000000
0000000000000000000000000000000000000000000000000000000000000
00000000000000000011111
>DM_validata876
MISIPKDEFQQIFVKDLMISSEKVAHVQIGNGLEHALLVLVKSGYSAIPVLDPMYKLHGLIS
TAMILDGILGLERIEFERLEEMKVEQVMKQDIPVLKLEDSFAKALEMTIDHPFICAVNEDGY
FEGILTRRAILKLLNKKVRQHNR
1111000000000000000000000000000000000000000000000000000000000
0000000000000000000000000000000000000000000000000000000000000
00000000000000000011111
>DM_validata877
MGSDKIHHHHHHMVDDIFERGSKGSSDFFTGNVWVKMLVTDENGVFNTQVYDVVFEPGARTH
WHSHPGGQILIVTRGKGFYQERGKPARILKKGDVVEIPPNVVHWHGAAPDEELVHIGISTQV
HLGPAEWLGSVTEEEYRKATEGK
1111111111111110000000000000000000000000000000000000000000000
0000000000000000000000000000000000000000000000000000000000000
00000000000000000000000
>DM_validata878
MSTEDFYLRYYVGHKGKFGHEFLEFEFRPDGKLRYANNSNYKNDTMIRKEAFVHQSVMEELK
RIIIDSEIMQEDDLPWPPPDRVGRQELEIVIGDEHISFTTSKTGSLVDVNRSKDPEGLRCFY
YLVQDLKCLVFSLIGLHFKIKPI

111000000000011111100000000000000001111111100000000000000000
000000000000000000000000000000000000000000000000000000000000
00000000000000000000111
>DM_validata879
GMSISSKAKEILTQFTREVWSEGNIEASDKYIAPKYTVLHDPGDPWEGRELDVAGYKERVKT
LRAAFPDQCFDIQGLFADGDAVVMTWLWTATHKEDIPGFPSTGKQIKMSGATVYYFDGNRLT
GHWQITDRLGVYQQLRQAAAGA
100000000000000000000000000000000000000000000000000000000000
000000000000000000000000000000000000000000000000000000000000
0000000000000000000111
>DM_validata880
EARGFTAEDFALSHPGGALGRKLLLRVNDIMHTGDEIPHVGLQATLRDALLEITRKNLGMTA
ICDDDMNIIGIFTDGDLRRVFDTGVDMRDASIADVMTRGGIRIRPGTLAVDALNLMQSRHIT
CVLVADGDHLLGVVHMHDLLRA
111111111111111100000000000000000000000000000000000000000000
000000000000000000000000000000000000000000000000000000000000
0000000000000000000001
>DM_validata881
MARSEQIAAVRRMVEAYNTGKTDDVADYIHPEYMNPGTLEFTSLRGPELFAINVAWVKKTFS
EEARLEEVGIEERADWVRARLVLYGRHVGEMVGMAPTGRLFSGEQIHLLHFVDGKIHHHRDW
PDYQGTYRQLGEPWPETEHRRP
111000000000000000000000000000000000000000000000000000000000
000000000000000000000000000000000000000000000000000000000000
0000000000000000000111
>DM_validata882
AEAAAAQAPAAGPGQMSFTFASPTQVFFNSANVRQVDVPTQTGAFGILAAHVPTLQVLRPGL
VVVHAEDGTTSKYFVSSGSVTVNADSSVQLLAEEAVTLDMLDLGAAKANLEKAQSELLGAAD
EATRAEIQIRIEANEALVKALE
111111111111111100000000000001111000000011111111111111100000
000111111111100000011111111100000000000000000000000000000000
0000000000000000000001
>DM_validata883
MGVTVQDICFAFLQNYYERMRTDPSKLAYFYASTAELTHTNYQSKSTNEKDDVLPTVKVTGR
ENINKFFSRNDAKVRSLKLKLDTIDFQYTGHLHKSILIMATGEMFWTGTPVYKFCQTFILLP
SSNGSTFDITNDIIRFISNSFK
111000000000000000000000000000000000000000111111000000000000
000000000000000000000000000000000000000000000000000000000000
0111000000000000000001
>DM_validata884
MSVAEFDADVIVDARDCIMGRVASQVAEQALDGETVAVVNAERAVITGREEQIVEKYEKRVD
IGNDNGYFYPKRPDGIFKRTIRGMLPHKKQRGREAFESVRVYLGNPYDEDGEVLDGTSLDRL
SNIKFVTLGEISETLGANKTW
111000000000000000000000000000000000000000000000000000000000
000000000000000000000000000000000000000000000011111111111111

11111100000000000001111
>DM_validata885
GMSWEKSGKEGSLLRWYDVMEAERYEYTVGPAGEQFFNGLKQNKIIGSKCSKCGRIFVPARS
YCEHCFVKIENYVEINKDEAYVDSYTIIYNDDEGNKLAQPVYIALIRFPNIEGGLLCYAEGN
VKVGAKAKILSFQWPLRVKVD
11111111110000000000000000000010000000000000000000000000000000000
00000000000000000000000000000000000000000000000000000000000000000
000000000000000000000
>DM_validata886
GHMSYSITLRVYQTNRDRGYFSIVEKTVWHFANGGTWSEANGAHTLTMGGSGTSGMLRFMST
KGERITVAVGVHNYKRWCDVVTGLKPDETALVINPQYYNNGGRDYVREKQLAEYSVTSAIGT
KVEVVYTVAEGNNLEANVIFS
11000000000000000000000000000000000000000000000000000000000000000
00000000000000000000000000000000000000000000000000000000000000000
000000000000000000000
>DM_validata887
MAHHHHHHMEYWHYVETTSSGQPLLREGEKDIFIDQSVGLYHGKSKILQRQRGRIFLTSQRI
IYIDDAKPTQNSLGLELDDLAYVNYSSGFLTRSPRLILFFKDPSSSTEFVQLSFRKSDGVLF
SQATERALENILTEKNKHIFN
11111100000000000000000000000000000000000000000000000000000000000
00000000000000000000000000000000000000000000000000000000000000000
000000000000011111111
>DM_validata888
MKSASDPFDLKRFVYAQAPVYRSVVEELRAGRKRGHWMWFVFPQLRGLGSSPLAVRYGISSL
EEAQAYLQHDLLGPRLHECTGLVNQVQGRSIEEIFGPPDDLKLCSSMTLFARATDANQDFVA
LLAKYYGGGEDRRTVALLAVT
11111000000000000000000000000000000000000000000000000000000000000
00000000000000000000000000000000000000000000000000000000000000000
000000000000000000000
>DM_validata889
MSGDYRITLEELRYISVFHSITGVTAYRCIVDEENNRLIFLVSEGEAGRAIGRGGRLIKLLR
EALGKNIEVVEYSSDLERIVKNLFPGVKIESINVRERNGVKQVVIKVSEDDKGAAIGKGGKN
VKRARLVLSKLFGVEKVVIR
11111100000000000000000000000000000000000000000000000000000000000
00000000000000000000000000000000000000000000000000000000000000000
00000000000000000000
>DM_validata890
MKDLKGTKTAENLKQGFIGESMANRRYLYFAKRADEEGYPEIAGLLRSIAEGETAHAFGHLD
FIRQGGLTDPATDKPIGTLEQMIESAIAGETYEWTQMYPGFAKVAREEGFPEVAEWFETLAR
AEKSHAEKFQNVLKQLKGGT
11000000000000000000000000000000000000000000000000000000000000000
00000000000000000000000000000000000000000000000000000000000000000
00000000000000011111
>DM_validata891

MAYWLCITNEDNWKVIKEKKIWGVAERYKNTINKVKVGDKLIIYEIQRSGKDYKPPYIRGVY
EVVSEVYKDSSKIFKPTPRNPNEKFPYRVKLKEIKVFEPPINFKELIPKLKFITNKKRWSGH
LMGKAMREIPEEDYKLIVGN
000000000000000000000000000000000000000000000000000000000000
000000000000000000000000000000000000000000000000000000000001
10000000000000000000
>DM_validata892
VSNLIVNGTAENGMDGWPDWGYPVSAVPEAAYGGTKGFKLSGGKQAGMGQKVALKPNTTYIL
GAWGKFTAKPGTYCDVIVQYHLKDANNTYVQNILRFTETDWTYKQVVFTTPDAFGSDPEFVL
WKDDASNADFYADNITLVEV
110000000000000000000000000000000000000000000000000000000000
000000000000000000000000000000000000000000000000000000000000
00000000000000000001
>DM_validata893
MKSNEHDDCQVTNPSTGHLFDLSSLSGRAGFTAAYSEKGLVYMSICGENENCPPGVGACFGQ
TRISVGKANKRLRYVDQVLQLVYKDGSPCPSKSGLSYKSVISFVCRPEAGPTNRPMLISLDK
QTCTLFFSWHTPLACEQAT
111111000000000000000000000000000000000000000000000000000000
000000000000000000000000000000000000000000000111110000000000
0000000000000000111
>DM_validata894
DSRGCARWCPQDSSCVNATACRCNPGFSSFSEIITTPMETCDDINECATLSKVSCGKFSDCW
NTEGSYDCVCSPGYEPVSGAKTFKNESENTCQDVDECSSGQHQCDSSTVCFNTVGSYSCRCR
PGWKPRHGIPNNQKDTVCE
110000000000000000000000000000000000000000000001111000000000
000000000000000000000000000000000000000000000000000000000000
0000000000000000000
>DM_validata895
MVANPPKPFEVHESGAYLHGTKAELKVGDRLVPGRESNFEAGRIMNHIYITQTLDAAVWGAE
LAAGEGRGRIFIVEPEGAIEDDPNVTDKKLPGNPTRSYRTREPVWIVGELTDWVGHPPEQLA
AMRQGLEELRRKGLAVIYD
111110000000000000000000000000000000000000000000000000000000
000000000000000000000000000000000000000000000000000000000000
0000000000000000000
>DM_validata896
SNAQLGGDMKTIAIADRTGEYEQLFKENDEFRFVHAEKTAEEYRKMGADKSGIDAVLEIRQD
LLEDPNAVAIYGYKQLPASVSNHISRILSDYLSDKKIASYNIPDIKQILADSKIELSVHTYK
WSEDGTNERTSGELASGIS
111111100000000000000000000000000000000000000000000000000000
000000000000000000000000000000000000000000000000000000000000
0000011111111111111
>DM_validata897
MHHHHHHMKEIATEYSFIKYTELELDDNGSIKQLSIPNKYNVIYAIAINDELVYIGKTKNLR
KRINYYRTAINRKDKTSDSTKSALIHSALKEGSKVEFYARQCFNLSMTNELGTMTIATIDLE

APLFIKLFNPPWNIQHKKK
111100000000000000000000000000000000000000000000000000000000
000000000000111110000000000000000000000000000000000000000000
0000000000000011111
>DM_validata898
GMKNVTKASIDDLDSIVHIDIDVIGNDSRRNYIKHSIDEGRCVIVKEDNSISGFLTYDTNFF
DCTFLSLIIVSPTKRRRGYASSLLSYMLSHSPTQKIFSSTNESNESMQKVFNANGFIRSGIV
ENLDEGDPEIIFYTKKLRA
000000000000000000000000000000000000000000000000000000000000
000000000000000000000000000000000000000000000000000000000000
0000000000000000001
>DM_validata899
NFNRFTQRAKKAIDLAFESAKSLGHNIVGSEHILLGLLREEEGIAAKVLSKVGFTEAYLEGK
IVDMEGKGEEISEDIVLSPRSKQILELSGMFANKLKTNYIGTEHILLAIIQEGEGIANKILN
YAGVNDRTLAQLTIDMMG
000000000000000000000000000000000000000000000000000000000000
000000000001100000000000000000000000000000000000000000000000
000000000000000000
>DM_validata901
MGSDKIHHHHHHMKSLGRHLVAEFYECDREVLDNVQLIEQEMKQAAYESGATIVTSTFHRFL
PYGVSGVVVISESHLTIHTWPEYGYAAIDLFTCGEDVDPWKAFEHLKKALKAKRVHVVEHER
GRYDEIGIPEDSPHKAAV
111111111111000000000000000000000000000000000000000000000000
000000000000000000000000000000000000000000000000000000000000
000000111111111111
>DM_validata902
GSHMAKEEIIWESLSVDVGSQGNPGIVEYKGVDTKTGEVLFEREPIPIGTNNMGEFLAIVHG
LRYLKERNSRKPIYSNSQTAIKWVKDKKAKSTLVRNEETALIWKLVDEAEEWLNTHTYETPI
LKWQTDKWGEIKADYGRK
111111000000000000000000000000000000000000000000000000000000
000000000000000000000000000000000000000000000000000000000000
000000000000000111
>DM_validata903
MGSSSSGLVPRGSPYTLSKRINGKWSGQSAGGCGNFQETHKNNPIYQFHIEKTGPLLIELRG
PRQYSVGFEVVTVSTLGDPGPHGFLRKSSGDYRCGFCYLELENIPSGIFNIIPSTFLPKQEG
PFFLDFNSIIPIKITQLQ
111100000000000000000000000000000000000000000000000000000000
000000000000000000000000000000000000000000000000000000000000
000000000000000000
>DM_validata904
MYKHILVAVDLSEESPILLKKAVGIAKRHDAKLSIIHVDVNFSDLYTGLIDVNMSSMQDRIS
TETQKALLDLAESVDYPISEKLSGSGDLGQVLSDAIEQYDVDLLVTGHHQDFWSKLMSSTRQ
VMNTIKIDMLVVPLRDE
000000000000000000000000000000000000000000000000000001111111

11000000000000000000000000000000000000000000000000000000000000
00000000000001111
>DM_validata905
ATKIDKEACRAAYNLVRDDGSAVIWVTFKYDGSTIVPGEQGAEYQHFIQQCTDDVRLFAFVR
FTTGDAMSKRSKFALITWIGENVSGLQRAKTGTDKTLVKEVVQNFAKEFVISDRKELEEDFI
KSELKKAGGANYDAQTE
00000000000000000000000000000000000000000000000000000000000000
00000000000000000000000000000000000000000000000000000000000000
00000001111111111
>DM_validata906
MRFRPFTEEDLDRLNRLAGKRPVSLGALRFFARTGHSFLAEEGEEPMGFALAQAVWQGEATT
VLVTRIEGRSVEALRGLLRAVVKSAYDAGVYEVALHLDPERKELEEALKAEGFALGPLVLAV
RVLGSRGARGETRGVLE
00000000000000000000000000000000000000000000000000000000000000
00000000000000000000000000000000000000000000000000000000000000
00000011111111111
>DM_validata907
SNAMSTKTMKEKAVELLQKCEVVTLASVNKEGYPRPVPMSKIAAEGISTIWMSTGADSLKTI
DFLSNPKAGLCFQEKGDSVALMGEVEVVTDEKLKQELWQDWFIEHFPGGPTDPGYVLLKFTA
NHATYWIEGTFIHKKLD
11111100000000000000000000000000000000000000000000000000000000
00000000000000000000000000000000000000000000000000000000000000
00000000000000001
>DM_validata908
MGDTFIRHIALLGFEKRFVPSQHYVYMFLVKWQDLSEKVVYRRFTEIYEFHKTLKEMFPIEA
GAINPENRIIPHLPAPKWFDGQRAAENRQGTLTEYCSTLMSLPTKISRCPHLLDFFKVRPDD
LKLPTDNQTKKPETYLM
10000000000000000000000000000000000000000000000000000000000000
00000000000000000000000000000000000000000000000000000000000000
00000000000000000
>DM_validata909
GHMSIKPKIMISSLDAERLEILLETLSQNAFPGRDDLEAELARAEVVDPEEIPPTVVTMNST
VRFRVESSAEEFCLTLVYPKDVDTSGEKISILAPVGSALLGLAQGDEIEWPKPGGGVLRVRI
VEVTYQPERSGEYYRGS
11111000000000000000000000011100000000000000000000000000000000
00000000000000000000000000000000000000000000000000000000000000
00000111111111111
>DM_validata910
MSNITIYHNPASGTSRNTLEMIRNSGTEPTIILYLENPPSRDELVKLIADMGISVRALLRKN
VEPYEQLGLAEDKFTDDQLIDFMLQHPILINRPIVVTPLGTRLCRPSEVVLDILQDAQKGAF
TKEDGEKVVDEAGKRLK
11000000000000000000000000000000000000000000000000000000000000
00000000000000000000000000000000000000000000000000000000000000
00000000000000001

>DM_validata911
GGGGGMNLSLSDLHRQVSRLVQQESGDCTGKLRGNVAANKETTFQGLTIASGARESEKVFAQ
TVLSHVANVVLTQEDTAKLLQSTVKHNLNNYDLRSVGNGNSVLVSLRSDQMTLQDAKVLLEA
ALRQESGARGHHHHHH
111111000000000000000000000000000000000000000000000000000000
000000000000000000000000000000000000000000000000000000000000
0000001111111111
>DM_validata912
MTSMTQSLREVIKAMTKARNFERVLGKITLVSAAPGKVICEMKVEEEHTNAIGTLHGGLTAT
LVDNISTMALLCTERGAPGVSVDMNITYMSPAKLGEDIVITAHVLKQGKTLAFTSVDLTNKA
TGKLIAQGRHTKHLGN
111000000000000000000000000000000000000000000000000000000000
000000000000000000000000000000000000000000000000000000000000
0000000000000001
>DM_validata913
ILVPIDISDKEFTERIISHVESEARIDDAEVHFLTVIPSLPYYASLGMAYTAELPGMDELRE
GSETQLKEIAKKFSIPEDRMHFHVAEGSPKDKILALAKSLPADLVIIASHRPDITTYLLGSN
AAAVVRHAECSVLVVR
000000000000000000000000000000000000011111111111111110000000
000000000000000000000000000000000000000000000000000000000000
0000000000000000
>DM_validata914
MGSSSQDPAKSNSIIVSPRQRGNPVLKFVRNVPWEFGDVIPDYVLGQSTCALFLSLRYHNLH
PDYIHGRLQSLGKNFALRVLLVQVDVKDPQQALKELAKMCILADCTLILAWSPEEAGRYLET
YKAYEQKPADLLMEKL
111111111110000000000000000000000000000000000000000000000000
000000000000000000000000000000000000000000000000000000000000
0000000000000000
>DM_validata915
SQISDMTRDGLANKALAVARTLADSPEIRQGLQKKPQESGIQAIAEAVRKRNDLLFIVVTDM
QSLRYSHPEAQRIGQPFKGDDILKALNGEENVAINRGFLAQALRVFTPIYDENHKQIGVVAI
GLELSRVTQQINDSRW
111100000000000000000000000000000000000000000000000000000000
000000000000000000000000000000000000000000000000000000000000
0000000000000111
>DM_validata916
GVLRYTLRARGHPNVTAGHRTTFEVTVDPEIGETADCIIGVSSSDSISTLPDEMKRAIARES
SLVRVILRTENGYDEIRGYGHPELTLDHPTDIVCRKSDYICSRTLMIRADKAAFDLDENLVR
DLRKGRELKVEIIVEY
000000000000000000000000000000000000000000000000000000000001
000000000000000000000000000000000000000000000000000000000000
0000000000000000
>DM_validata917
MENKIIYFLCTGNSCRSQMAEGWAKQYLGDEWKVYSAGIEAHGLNPNAVKAMKEVGIDISNQ

TSDIIDSDILNNADLVVTLCGDAADKCPMTPPHVKREHWGFDDPARAQGTEEEKWAFFQRVR
DEIGNRLKEFAETGK
111000000000000000000000000000000000000000000000000000000000000
000000000000000000011111111111100000000000000000000000000000000
000000000000000
>DM_validata918
MWKAVMNAWNGTESQSKNVSNIQSYSFEDMKRIVGKHDPNVVLVDVREPSEYSIVHIPASIN
VPYRSHPDAFALDPLEFEKQIGIPKPDSAKELIFYCASGKRGGEAQKVASSHGYSNTSLYPG
SMNDWVSHGGDKLDL
111111111111111111000000000000000000000000000000000000000000000
000000000000000000000000000000000000000000000000000000000000000
000000000000000
>DM_validata919
MTQTTQSPALIASQSSWRCVQAHDREGWLALMADDVVIEDPIGKSVTNPDGSGIKGKEAVGA
FFDTHIAANRLTVTCEETFPSSSPDEIAHILVLHSEFDGGFTSEVRGVFTYRVNKAGLITNM
RGYWNLDMMTFGNQE
111100000000000000000000000000000000000000000000000000000000000
000000000000000000000000000000000000000000000000000000000000000
000000000000011
>DM_validata920
MIEGLSHMTFIVRDLERMTRILEGVFDAREVYASDTEQFSLSREKFFLIGDIWVAIMQGEKL
AERSYNHIAFKIDDADFDRYAERVGKLGLDMRPPRPRVEGEGRSIYFYDDDNHMFELHTGTL
TERLARKAKGLEAAQ
000000000000000000000000000000001111111000000000000000000000000
000000000000000000000000000000000000111000000000000000000000000
000001111111111
>DM_validata921
GTMGKSKGTRFERDLLVELWKAGFAAIRVAGSGVSPFPCPDIVAGNGRTYLAIEVKMRKELP
LYLSADEVEQLVTFARGFGAEAYVALKLPRKKWRFFPVQMLERTEKNFKIDESVYPLGLEIA
EVAGKFFQERFGEKV
111111000000000000000000000000000000000000000000000000000000000
000000000000000000000000000000000000000000000000000000000000000
000111111111111
>DM_validata922
AEVAPGDVAIDGQGHVARPLTDAPGDPVEGRRLMTDRSVGNCIACHEVTEMADAQFPGTVGP
SLDGVAARYPEAMIRGILVNSKNVFPETVMPAYYRVEGFNRPGIAFTSKPIEGEIRPLMTAG
QIEDVVAYLMTLTQ
000000000000000000000000000000000000000000000000000011100000000
000000000000000000000000000000000000000000000000000000000000000
00000000000000
>DM_validata923
AMTYHLDVVSAEQQMFSGLVEKIQVTGSEGELGIYPGHAPLLTAIKPGMIRIVKQHGHEEFI
YLSGGILEVQPGNVTVLADTAIRGQDLDEARAMEAKRKAEEHISSSHGDVDYAQASAELAKA
IAQLRVIELTKKAM

000000000000000000000000000000000000000000000000000000000000
000000000000000000000000000000000000000011111100000000000000
00000000001111
>DM_validata924
MSGTLMAFDFGTKSIGVAVGQRITGTARPLPAIKAQDGTPDWNIIERLLKEWQPDEIIVGLP
LNMDGTEQPLTARARKFANRIHGRFGVEVKLHDERLSTVEARSGLFEQGGYRALNKGKVDSA
SAVIILESYMEQGY
100000000000000000000000000000000000000000000000000000000000
000000000000000000000000000000000000000111111111111111110000
00000000000000
>DM_validata925
MIKSKGDVNILLNFRHNINGEDLIIAVAQDHETGEVLMVAYMNREALRRTLETGTAHYWSTS
RGKLWLKGESSGHVQRVKDVLVDCDGDAVVLKVEQEGGACHTGYRSCFYRSIDGDELKVRED
AVKVFDPEEIYGDG
111111100000000000000000000000000000000000000000000000000000
000000000000000000000000000000000000000000000000000000000000
00000001111111
>DM_validata926
GPLGSSRVSVLSMDFEDIYRFFQDPPPHYLSKELAVCYVLAVLRHEDSYGTELIQHLETHWP
NYRLSDTVLYTALKFLEDEQIISGYWKKVEGRGRPRRMYQLAQANDDRSRDLAQLWERYLSS
SAATDRQLIPVEAR
111111111111000000000000000000000000000000000000000000000000
000000000000000000000000000000000000000000000000000000000011
11111111111111
>DM_validata927
AELTPEVLTVPLNSEGKTITLTEKQYLEGKRLFQYACASCHVGGITKTNPSLDLRTETLALA
TPPRDNIEGLVDYMKNPTTYDGEQEIAEVHPSLRSADIFPKMRNLTEKDLVAIAGHILVEPK
ILGDKWGGGKVYY
000000000000111111111000000000000000000000000000000000000000
000000000000000000000000000000000000000000000000000000000000
0111111111111
>DM_validata928
GSHMRKIPKVGHTFFQKPESCPPVPGGSMKLDIGIINENQRVSMSRNIESRSTSPWNYTVTW
DPNRYPSEVVQAQCRNLGCINAQGKEDISMNSVPIQQETLVVRRKHQGCSVSFQLEKVLVTV
GCTCVTPVIHHVQ
111111111111000000000111000000000000000000000000000000000000
000000000000000000000000000000000000000000000000000000000000
0000000011111
>DM_validata929
MFKTILLAYDGSEHARRAAEVAKAEAEAHGARLIVVHAYEPVPDYLGEPFFEEALRRRLERA
EGVLEEARALTGVPKEDALLLEGVPAEAILQAARAEKADLIVMGTRGLGALGSLFLGSQSQR
VVAEAPCPVLLVR
000000000000000000000000000000000000000111111111111110000000
000000000000000000000000000000000000000000000000000000000000

0000000000000
>DM_validata931
MEEIKCLLCRYLKERQEKFISDWKKKVIIERERDPYKEEIIKNGEHLLSAFIMYLKEEISLQE
IEITSKKIARERIDAKVNIAEFIHNTNVAKIEIMNILTLLNPDLQQYQALVKKINQFFDHLI
YYTVHSYYEQKA
111100000000000000000000000000000000000000000000000000000000
000000000000000000000000000000000000000000000000000000000000
000000000000
>DM_validata932
MMSNQTVYQFIAENQNELLQLWTDTLKELSEQESYQLTDQVYENISKEYIDILLLSVKDENA
AESQISELALRAVQIGLSMKFLATALAEFWKRLYTKMNDKRLPDQESTELIWQIDRFFSPIN
TEIFNQYSISWE
110000000000000000000000000000000000000000000000000000000000
000000000000000000000000000000000000001111110000000000000000
000000000000
>DM_validata933
MMKVEKEIKTNQDIDVVMTIFSDPAFTIPQIFPGIASIKCIEPEIFEAEGKFLAFSYKVKGR
VYKGVDEVRIIYDSDRGNGILYIRKKDNNTLQIILEHDNKLTAFLGKPYVSSNLDRLAENID
EIIRLERIKRKI
110000000000000000000000000000000000000000000000000000000000
000000000000000000000000000000000000000000000000000000000000
000000000000
>DM_validata934
MSEVVKKFMYLNRKAPYGTIYAWEALEVVLIGAAFDQDVCVLFLDDGVYQLTRGQDTKGIGM
KNFSPTYRTLGDYEVRRIYVDRDSLEARGLTQDDLVEIAFEDMETEEEFDNIVEVIDSARVS
ELMNESDAVFSF
111100000000000000000000000000000000000000000000000000000000
000000000000000000000000000000000000000000000000000000000000
000000000000
>DM_validata935
SSIIRKVISTSKAPAAIGAYSQAVLVDRTIYVSGQIGMDPSSGQLVPGGVAEEAKQALKNLG
EILKAAGCDFTNVVKTTVLLADINDFGTVNEIYKTYFQGNLPARAAYQVAALPKGSRIEIEA
IAVQGPFTTAGL
000000000000000000000000000000000000000000000000000000000000
000000000000000000000000000000000000000000000000000000000000
000000001111
>DM_validata936
SNAMRKLNNHDVHKRYQDRLEEDVEFTINYELPLSCLWSTIKDFSSDFEEKTEAFFILFKEL
LRRGHLKLQRDGQIIGHTPEEWEQIFREVWPEYEIEPNPLPGYAPFDIGMWLTVEAPAYAVW
IDPEDGSEYWAG
111111000000000000000000000000000000000000000000000000000000
000000000000000000000000000000000000000000000000000000000000
111111111111
>DM_validata937

TIAIVIGTHGWAAEQLLKTAEMLLGEQENVGWIDFVPGENAETLIEKYNAQLAKLDTTKGVL
FLVDTWGGSPFNAASRIVVDKEHYEVIAGVNIPMLVETLMARDDDPSFDELVALAVETGREG
VKALKAKPFAG
000000000000000000000000000000000000000000000000000000000000
000000000000000000000000000000000000000000000000000000000000
00000111111
>DM_validata938
GMSTQENVQIVKDFFAAMGRGDKKGLLAVSAEDIEWIIPGEWPLAGTHRGHAALAALLQKAS
EMVEISYPEPPEFVAQGERVLVVGFATGRVKSTNRTFEDDWVFAITVRKSKVTSIREYIDTL
ALARATNFNAT
000000000000000000000000000000000000000000000000000000000000
000000000000000000000000000000000000000000000000000000000000
00000001111
>DM_validata939
GMSHSLKDFLEACETLGTLRLIVTSSAAVLEARGKIEKLFYAELAKGKYANMHTEGFEFHLN
MEKITQVKFETGEAKRGNFTTYAIRFLDEKQESALSLFLQWGKPGEYEPGQVEAWHTLKEKY
GEVWEPLPVQL
100000000000000000000000000000000000000000000000000000000000
000000000000001100000000000000000000000000000000000000000000
00000000000
>DM_validata940
SDRAEVRNIPFKLGMYLTVGGVVNSNATRFSINVGESTDSIAMHMDHRFSYGADQNVLVLNS
LVHNVGWQQEERSKKFPFTKGDHFQITITFDTHTFYIQLSNGETVEFPNRNKDAAFNLIYLA
GDARLTFVRLE
100000000000000000000000000000000000000000000000000000000000
000000000000000000000000000000000000000000000000000000000000
00000000000
>DM_validata941
LEEVIEQLREANEPVPVPLELPDEDQLVEIEEQLFINIPFVFKEFLLTVSDVVYGSLEPVTV
TDPQSHTYLPEVCATAWDLGVPRELIPICQDGEDYYCVEEDGTVLLWSAEEELVTEESWESV
WHWARDVWLES
000000000000000000000000000000000000000000000000000000000000
000000000000000000000000000000000000000000000001110000000000
00000000000
>DM_validata942
MKKQQFIDMQEQGTSTIPNLLLTHYKQLGLNETELILLLKIKMHLEKGSYFPTPNQLQEGMS
ISVEECTNRLRMFIQKGFLFIEECEDQNGIKFEKYSLQPLWGKLYEYIQLAQNQTQERKAEG
EQKLEHHHHHH
000000000000000000000000000000000000000000000000000000000000
000000000000000000000000000000000000000000000000000111111111
11111111111
>DM_validata943
MTAPEQTFRNKKQRKQQVKLRKPGFAVAKYVRMSPRKVRLVVDVIRGKSVQDAEDLLRFIPR
SASEPVAKVLNSAKANALHNDEMLEDRLFVKEAYVDAGPTLKRLIPRARGSANIIKKRTSHI

TIIVAEKGNK
111100000000000000000000000000000000000000000000000000000000
000000000000000000000000000000000000000000000000000000000000
0000000000
>DM_validata944
APKNELVQKFQVYYLGNVPVAKPVGVDVINGALESVLSSSSREQWTPSHVSVAPATLTILHQ
QTEAVLGECRVRFLSFLAVGRDVHTFAFIMAAGPASFCCHMFWCEPNAASLSEAVQAACMLR
YQKCLDARSQ
111111111000000000000000000000000000000000000000000000000000
000000000000000000000000000000000000000000000000000000000000
0000000001
>DM_validata945
GQQKNMENKTLNENIPEMIISLEKEALASTDPMAFVELSDTDVIYFDPSLETKIEGLEQLRT
YYKGMQLPPADHFDMIRPVVQVAQNIAVLTFNLDSYLSDKVIKWNCTEVYRRNPDNQWKIIQ
THWSYVKPLD
110000000000000000000000000000000000000000000000000000000000
000000000000000000000000000000000000000000000000000000000000
0000000000
>DM_validata946
SDAFDEMDEIWALYADDGAQALDAMEASLLALQAGEDAAAHVGPLFRAVHTFKGNSRVLGLS
VVESRAHLCEDLIGLVRDAGVPMDGEIVEILLFASDTLRAMLEETAASRADVEGTGSEALMD
QLRSKIARCS
111111000000000000000000000000000000000000000000000000000000
000000000000000000000000000000000000000000000000000000000000
0000000000
>DM_validata947
GHMNAADRVMQSYGRCCASTGFFDDFYRHFLASSPQIRAKFATTDMTAQKHLLRAGIMNLVM
YARGMSDSKLRALGASHSRAALDIRPELYDLWLDALLMAVAEHDRDCDAETRDAWRDVMGRG
IAVIKSYYGS
111000000000000000000000000000000000000000000000000000000000
000000000000000000000000000000000000000000000000000000000000
0000000000
>DM_validata948
MEFKKVAKETAITLQSYLTYQAVRLISQQLSETNPGQAIWLGEFSKRHPIQESDLYLEAMML
ENKELVLRILTVRENLAEGVLEFLPEMVLSQIKQSNGNHRRSLLERLTQVDSSSTDQTEPNP
GESDTSEDSE
000000000000000000000000000000000000000000000000000000000000
000000000000000000000000000000000000000000000011111111111111
1111111111
>DM_validata949
MGSDKIHHHHHHMKEGTGMVVRSSEITPERISNMRGGKGEVEMAHLLSKEAMHNKARLFARM
KLPPGSSVGLHKHEGEFEIYYILLGEGVFHDNGKDVPIKAGDVCFTDSGESHSIENTGNTDL
EFLAVIILL
111111111111111110000000000000000000000000000000000000000000

00000000000000000000000000000000000000000000000000000000000000
000000000
>DM_validata950
APTSSSTKKTQLQLEHLLLDLQMILNGINNCKNPKLTRMLTFKFYMPKKATELKHLQCLEEE
LKPLEEVLNLAQSKNFHLRPRDLISNINVIVLELKGSETTFMCEYADETATIVEFLNRWITF
CQSIISTLT
11111100000000000000000000000000000000000000000000000000000000
00000000000111111000000000000000001111110000000000000000000000
000000000
>DM_validata951
MSWQTYVDEHLMCDIDGQGEELAASAIVGHDGSVWAQSSSFPQFKPQEITGIMKDFEEPGHL
APTGLHLGGIKYMVIQGEAGAVIRGKKGSGGITIKKTGQALVFGIYEEPVTPGQCNMVVERL
GDYLIDQGL
00000000000011111111100000000000000000000000000000000000000000
00000000000000000000000000000000000000000000000000000000000000
000000000
>DM_validata952
GSHMKVEDPKDFPSELLSFLSHAVFSNRTLACFAIYTTKEKAALLYKKIMEKYSVTFISRHN
SYNHNILFFLTPHRHRVSAINNYAQKLCTFSFLICKGVNKEYLMYSALTRDPFSVIEESLPG
GLKEHDFNP
11111111000000000000000000000000000000000000000000000000000000
00000000000000000000000000000000000000000000000000000000000001
111111111
>DM_validata953
SLSEDKMDLYLQQGMYGPLETKPDERHLFLGSLRERVVLALTKGQVLRSKPYKEAEHELKNS
HNVTLLINGELQYQSYSSYIQMASRYGVPFKIVSDLQFHTPLGIVIAADIAVNRELIYIQDD
IYNRSVLKS
11100000000000000000000000000000000000000000000000000000000000
00000000000000000000000000000000000000000000000000000000000000
000000011
>DM_validata954
MSGSLARAAARPNAPTLVDEATVDDFIAHSGKIVVLFFRGDAVRFPEAADLAVVLPELINAF
PGRLVAAEVAAEAERGLMARFGVAVCPSLAVVQPERTLGVIAKIQDWSSYLAQIGAMLAEVD
QPGEAELQS
11111111111111000000000000000000000000000000000000000000000000
00000000000000000000000000000000000000000000000000000000000000
001111111
>DM_validata955
MVDNKVTQSNKLIESSHTLTLNEKRLVLCAASLIDSRKPLPKDGYLTIRADTFAEVFGIDVK
HAYAALDDAATKLFNRDIRRYVKGKVVERMRWVFHVKYREGQGCVELGFSPTIIPHLTMLHK
EFTSYQLK
11111110000000000000000000000000000000000000000000000000000000
00000000000000000000000000000000000000000000000000000000000000
00000000

>DM_validata956
FENATSDAINQDMMLYIERIAKIIQKLPKRVHINVRGFTDDTPLVKTRFKSHYELAANRAYR
VMKVLIQYGVNPNQLSFSSYGSTNPIAPNDSLENRMKNNRVEIFFSTDANDLSKIHSILDNE
FNPHKQQE
000000000000000000000000000000000000000000111100000000000000
000000000000000000000000000000000000000000000000000000000000
00011111
>DM_validata957
GHMGSADSYTSRPSDSDVSLEEDREAVRREAERQAQAQLEKAKTKPVAFAVRTNVRYSAAQE
DDVPVPGMAISFEAKDFLHVKEKFNNDWWIGRLVKEGCEIGFIPSPVKLENMRLQHEQRAKQ
GKFYSSKS
111111111111111111111111110000000000000000000000000000000000
000000000000000000000000000000000000000000000000000000000001
11111111
>DM_validata958
MHGLQDDPDLQALLKGSQLLKVKSSSWRRERFYKLQEDCKTIWQESRKVMRSPESQLFSIED
IQEVRMGHRTEGLEKFARDIPEDRCFSIVFKDQRNTLDLIAPSPADAQHWVQGLRKIIHHSG
SMDQRQK
000000000000000000000000000000000000000000000000000000000000
000000000000000000000000000000000000000000000000000000000111
1111111
>DM_validata959
CPERELQRREEEANVVLTGTVEEIMNVDPVHHTYSCKVRVWRYLKGKDIVTHEILLDGGNKV
VIGGFGDPLICDNQVSTGDTRIFFVNPAPQYMWPAHRNELMLNSSLMRITLRNLEEVEHCVE
EHRKLLA
111100000000000000000000000000000000000000000000000000000000
000000000000000000000000000000000000000000000000000000000000
0000000
>DM_validata960
MNLPTAQEVQGLMARYIELVDVGDIEAIVQMYADDATVEDPFGQPPIHGREQIAAFSRQGLG
GGKVRACLTGPVRASHNGCGAMPFRVEMVWNGQPCALDVIDVMRFDEHGRIQTMQAYWSEVN
LSVREPQ
000000000000000000000000000000000000000000000000000000000011
110000000000000000000000000000000000000000000000000000000000
0011111
>DM_validata961
GMMNTTTLDRYFDLFDASRTDEKAFDDLISLFSDEITFVLNGQEQHGIDAWKQFVRMVFTAN
QDIKHMYAGWVPSETGDTMETRWAVCGKSADGSVFTQDGTDIARLNADGKIVYLANVPDDTA
MFNQYND
111110000000000000000000000000000000000000000000000000000000
000000000000000000000000000000000000000000000000000000000001
1111111
>DM_validata962
GMSSKEEILSILEAFASTERMGSFFLDNATADFLFIRPSGNPLDAKGFENMWSSGDLVLESA

EITKVHKFELLGSNAAICVFTLGSKFTYKGTQNDDLPTVTSIFKKIDEKWKVAWMQRSSGQS
DMTLWNE
100000000000000000000000000000000000000000000000000000000000
000000000000000000000000000000000000000000000000000000000000
0000011
>DM_validata963
MVRVGMRAAPRVSLEALKAALGGLKLSEAKVYLITDWQDKRDQARYALLLHTGKKDLLVPDA
FGPAFPGGEEALSELVGLLLAQGARRFYEAVVSPGEMTALLDLPPEELLKRVMAIANPTDPG
IYLKRAA
000000000000000000000000000000000000000000000000000000000000
000000000000000000000000000000000000000000000000000000000000
0001111
>DM_validata964
SGRHERDAFDTLFDHAPDKLNVVKKTLITFVNKHLNKLNLEVTELETQFADGVYLVLLMGLL
EGYFVPLHSFFLTPDSFEQKVLNVSFAFELMQDGGLEKPKPRPEDIVNCDLKSTLRVLYNLF
TKYRNVE
111110000000000000000000000000000000000000000000000000000000
000000000000000000000000000000000000000000000000000000000000
0000000
>DM_validata966
SNAMRETFEIGEIVTGIYKTGKYIGEVTNSRPGSYVVKVLAVLKHPVQGDLHNVKQANVPFF
HERRALAFREQTNIPEQMVKKYEGEIPDYTESLKLALETQMNSFSEDDSPFAERSLETLQQL
KKDYKL
111111000000000000000000000000000000000001111111111111
100000000000000000000000000000000000000000000000000000000000
000000
>DM_validata967
MTMSRRNTDAITIHSILDWIEDNLESPLSLEKVSERSGYSKWHLQRMFKKETGHSLGQYIRS
RKMTEIAQKLKESNEPILYLAERYGFESQQTLTRTFKNYFDVPPHKYRMTNMQGESRFLHPL
NHYNS
111111110000000000000000000000000000000000000000000000000000
000000000000000000000000000000000000000000000000000000000000
11111
>DM_validata968
GQSAKEAIEAANADFVKAYNSKDAAGVASKYMDDAAAFPPDMARVDGRQNIQKLWQGAMDMG
ISELKLTTLDVQESGDFAFESGSFSLKAPGKDSKLVDAAGKYVVVWRKGQDGGWKLYRDIWN
SDPAK
111000000000000000000000000000000000000000000000000000000000
000000000000000000000000000000000000000000000000000000000000
00000
>DM_validata969
SCSSPQVEIPTTYSPEKIAQLQVYVNPIAVARDGMEKRLQGLIADQNWVDTQTYIHGPLGQL
RRDMLGLASSLLPKDQDKAKTLAKEVFGHLERLDAAAKDRNGSQAKIQYQEALADFDSFLNL
LPQAS

111111111110000000000000000000000000000000000000000000000000
000000000000000000000000000000000000000000000000000000000000
00111
>DM_validata970
TSTMGNLQTAINDKSGTLASQNFLDADEQKRNAYNQAVSAAETILNKQTGPNTAKTAVEQAL
NNVNNAKHALNGTQNLNNAKQAAITAINGASDLNQKQKDALKAQANGAQRVSNAQDVQHNAT
ELNTA
110000000000000000000000000000000000000000011111110000000000
000000000000000000000000000000000000000000000000000000000000
00001
>DM_validata971
SNAMGWKMQATVTEIGKHAIDDSEKMIILFGETATDTLKQHAVIQSFPEKDQVTLAEGDHLK
IGDTNYTITKVGSFANSNLQSIAHSTLIFADAPTDEDDVIRNGVYLTPHQLPKITIGTTIDY
LVNGA
111111100000000000000000000000000000000000000000000000000000
000000000000000000000000000000000000000000000000000000000000
00111
>DM_validata972
KQEVQAWDGEVRQVSKHAFSLKQLDNPARIPPCGWKCSKCDMRENLWLNLTDGSILCGRRYF
DGSGGNNHAVEHYRETGYPLAVKLGTITPDGADVYSYDEDDMVLDPSLAEHLSHFGIDMLKM
QKTDK
111111111100000000000000000000000000000000000000000000000000
000000000000000000000000000000000000000000000000000000000000
00011
>DM_validata974
GSSEDKITVHFINRDGETLTTKGKIGDSLLDVVVQNNLDIDGFGACEGTLACSTCHLIFEQH
IFEKLEAITDEENDMLDLAYGLTDRSRLGCQICLTKAMDNMTVRVPDAVSDARESIDMGMNS
SKIE
111100000000000000000000000000000000000000000000000000000000
000000000000000000000000000000000000000000000011111111111111
1111
>DM_validata975
MAREFKRSDRVAQEIQKEIAVILQREVKDPRIGMVTVSDVEVSSDLSYAKIFVTFLFDHDEM
AIEQGMKGLEKASPYIRSLLGKAMRLRIVPEIRFIYDQSLVEGMRMSNLVTNVVREDEKKHV
EESN
111111000000000000000000000000000000000000000000000000000000
000000000000000000000000000000000000000000111111111111111111
1111
>DM_validata976
MSHQIRSKKTLWELVGRNKDALRDFLKEHRGTILLRDIASEHKVVYKPIFKRYNGDPDLIED
NSNDVEHWYDYHLERYWNTPELKKEFYKKFGPVDLNQPIILAKPLRQHNRGDLVHLLPQFVV
PVYN
111111100000000000000000000000000000000000000000000000000000
000000000000000000000000000000000000000000000000000000000000

0000
>DM_validata977
GIRLKDELINIKQILEAADIMFIYEEEKWPENISLLNENILSMCLKEAVTNVVKHSQAKTCR
VDIQQLWKEVVITVSDDGTFKGEENSFSKGHGLLGMRERLEFANGSLHIDTENGTKLTMAIP
NNSK
000000000000000000000000000000000000000000000000000000000000
000000000000000000000000000000000000000000000000000000000000
0111
>DM_validata978
MGRELSFFLQKESAGFFLGMDAPAGSSVACGSEVLRAVPVGTVDAAKEKHIPVVEVHGHEVK
VKVGSVAHPMTPEHYIAWVCLKTRKGIQLKELPVDGAPEVTFALTADDQVLEAYEFCNLHGV
WSGK
110000000011111000000000000000000000000000000000000000000000
000000000000000000000000000000000000000000000000000000000000
0000
>DM_validata979
ATHTAQTQTHLNFTQIKTVDELNQALVEAKGKPVMLDLYADWCVACKEFEKYTFSDPQVQKA
LADTVLLQANVTANDAQDVALLKHLNVLGLPTILFFDGQGQEHPQARVTGFMDAETFSAHLR
DRQP
111111111000000000000000000000000000000000000000000000000000
000000000000000000000000000000000000000000000000000000000000
0011
>DM_validata980
MADFARSLSITTPEEMIEKAKGETAYLPCKFTLSPEDQGPLDIEWLISPADNQKVDQVIILY
SGDKIYDDYYPDLKGRVHFTSNDLKSGDASINVTNLQLSDIGTYQCKVKKAPGVANKKIHLV
VLVK
111111000000000000000000000000000000000000000000000000000000
000000000000000000000000000000000000000000000000000000000000
0011
>DM_validata981
MPKSFYDAVGGAKTFDAIVSRFYAQVAEDEVLRRVYPEDDLAGAEERLRMFLEQYWGGPRTY
SEQRGHPRLRMRHAPFRISLIERDAFLRCMHTAVASIDSETLDDEHRRELLDYLEMAAHSLV
NSPF
110000000000000000000000000000000000000000000000000000000000
000000000000000000000000000000000000000000000000000000000000
0000
>DM_validata982
GQKSNRKDYSLTMQSSVTVQEGMCVHVRCSFSYPVDSQTDSDPVHGYWFRAGNDISWKAPVA
TNNPAWAVQEETRDRFHLLGDPQTKNCTLSIRDARMSDAGRYFFRMEKGNIKWNYKYDQLSV
NVT
111111000000000000000000000000000011100000000000011111100000
000000000000000000000000000000000000000000000000000000000000
000
>DM_validata983

GPTGTGESKCPLMVKVLDAVRGSPAINVAVHVFRKAADDTWEPFASGKTSESSDSHGLTTEE
EFVEGIYKVEIDTKSYWKALGISPFHEHAEVVFTANDSGPRRYTIAALLSPYSYSTTAVVTN
PKE
111111111100000000000000000000000000000000000000000000000000
000000000000000000000000000000000000000000000000000000000000
111
>DM_validata984
GSPQDLTVSLIPVSGLKAGKNAPSAKIAKLVVNSTTLKEFGVRGISNNVVDSTGTAWRVAGK
NTGKEIGVGLSSDSLRRSDSTEKWNGVNWMTFNSNDTLDIVLTGPAQNVTADTYPITLDVVG
YQP
110000000000011111000000000000000000000000000000000000000001
111110000000000000000000000000000000000000001000000000000000
000
>DM_validata985
GMDMKKHGLSIGINRIESVFFVTLKAIGTLTHEDYLVITPMLEGALSQVDQPKVSLFLDATE
LDGWDLRAAWDDLKLGLKHKSEFERVAILGNKDWQEWAAKIGSWFIAGEIKYFEDEDDALKW
LRY
111000000000000000000000000000000000000000000000000000000000
000000000000000001100000000000000000000000000000000000000000
000
>DM_validata986
MIRTMLQGKLHRVKVTHADLHYEGSCAIDQDFLDAAGILENEAIDIWNVTNGKRFSTYAIAA
ERGSRIISVNGAAAHCASVGDIVIIASFVTMPDEEARTWRPNVAYFEGDNEMKRTAKAIPVQ
VA
000000000000000000000000000000000000000000000000000000000000
000000000000000000000000000000000000000000000000000111111111
11
>DM_validata987
MADEIAKAQVARPGGDTIFGKIIRKEIPAKIIFEDDQCLAFHDISPQAPTHFLVIPKKHISQ
ISAAEDADESLLGHLMIVGKKCAADLGLKKGYRMVVNEGSDGGQSVYHVHLHVLGGRQMNWP
PG
111111111111000000000000000000000000000000000000000000000000
000000000000000000000000000000000000000000000000000000000000
00
>DM_validata988
PTNDQLTDLQEAHFVVFESEENSESVMDGFVEHPFYTATLNGQKYVVMKTKDDSYWKDLIVE
GKRVTTVSKDPKNNSRTLIFPYIPDKAVYNAIVKVVVANIGYEGQYHVRIINQDINTKDDDT
SQ
111110000000000000000000000000000000000000000000000000000000
000000000000000000000000000000000000000000000000000001111111
11
>DM_validata990
MQPASAKWYDRRDYVFIEFCVEDSKDVNVNFEKSKLTFSCLGGSDNFKHLNEIDLFHCIDPN
DSKHKRTDRSILCCLRKGESGQSWPRLTKERAKLNWLSVDFNNWKDWEDDSDEDMSNFDRFS

E
000000000000000000000000000000000000000000000000000000000000
000000000000000000000000000000000000000000000011111111111111
1
>DM_validata991
MAPMLSGLLARLVKLLLGRHGSALHWRAAGAATVLLVIVLLAGSYLAVLAERGAPGAQLITY
PRALWWSVETATTVGYGDLYPVTLWGRCVAVVVMVAGITSFGLVTAALATWFVGREQERRGH
F
111111111111111111111100000000000000000000000000000000000000
000000000000000000000000000000000000000000000000000000001111
1
>DM_validata992
MNGNHSAPAIAIAVIDGCDGLWREVLLGIEEEGIPFRLQHHPAGEVVDSAWQAARSSPLLVG
IACDRHMLVVHYKNLPASAPLFTLMHHQDSQAHRNTGNNAARLVKGIPFRDLNSEATGEQQD
E
111100000000000000000000000000000000000000000000000000000000
000000000000000000000000000000000000000000000000111111111111
1
>DM_validata993
MASTVSNTSKLEKPVSLIWGCELNEQNKTFEFKVEDDEEKCEHQLALRTVCLGDKAKDEFHI
VEIVTQEEGAEKSVPIATLKPSILPMATMVGIELTPPVTFRLKAGSGPLYISGQHVAMEEDY
111111111111110000000000000000011111111000000000000000000000
000001111110000000000000000000000000000000000000000000111111
>DM_validata994
ASNVANDGERPSRVVELTDETFDSIVMDPEKDVFVLYYVPWSRHSVAAMRLWDDLSMSQSQK
RNHLTFVAARIDGEKYPDVIERMRVSGFPTMRYYTRIDKQEPFEYSGQRYLSLVDSFVFQNT
111111111000000000000000000000000000000000000000000000000000
000000000000000000000000000000000000000000000000000000000000
>DM_validata995
GSSMKISRGLLKTILEAAKSAHPDEFIALLSGSKDVMDELIFLPFVSGSVSAVIHLDMLPIG
MKVFGTVHSHPSPSCRPSEEDLSLFTRFGKYHIIVCYPYDENSWKCYNRKGEEVELEVVEKD
000000000000000000000000000000000000000000111111111111111110
000000000000000000000000000000000000000000000000000000000011
>DM_validata996
MAKLTSAVPVLTARDVAEAVEFWTDRLGFSRVFVEDDFAGVVRDDVTLFISAVQDQVVPDNT
QAWVWVRGLDELYAEWSEVVSTNFRDASGPAMTEIVEQPWGREFALRDPAGNCVHFVAEEQD
100000000000000000000000000000000000000000000000000000000000
000000000000000000000000000000000000000000000000000000000111
>DM_validata997
ADDLKRFLYKKLPSVEGLHAIVVSDRDGVPVIKVANDNAPEHALRPGFLSTFALATDQGSKL
GLSKNKSIICYYNTYQVVQFNRLPLVVSFIASSSANTGLIVSLEKELAPLFEELRQVVEVS
100000000000000000000000000000000000000000000000000000000000
00000000000000000000000000000000000000000000000000000000111
>DM_validata998

SDIHQITPFLHIPDMQEALTLFCDTLGFELKYRHSNYAYLELSGCGLRLLEEPARKIIPDGI
ARVAICIDVSDIDSLHTKLSPALENLPADQVEPLKNMPYGQREFQVRMPDGDWLNFTAPLA
11000000000000000000000000000000000000000000000000000111111111
0000000000000000000000000000000000000000000000000000000000000
>DM_validata999
MARYEEVSVSGFEEFHRAVEQHNGKTIFAYFTGSKDAGGKSWCPDCVQAEPVVREGLKHISE
GCVFIYCQVGEKPYWKDPNNDFRKNLKVTAVPTLLKYGTPQKLVESECLQANLVEMLFSED
11100000000000000000000000000000000000000000000000000000000000
0000000000000000000000000000000000000000000000000000000000001
>DM_validata1000
ERKRREKRLEETSSRLEALFENSPDMIDVLDADGTICEVNQRFCAELGYDESEVLGRSIWEF
DLMFDAEDVQTQLSGFSVDERRKFEGLYERRDGSTMSVEVHLLRFNLEGEDRFLAISRDIT
11111111000000000000000000000000000000000000000000000000000000
0000000000000000000000000000000000000000000000000000000000001
>DM_validata1001
VTDNNGAVKFPQLCKFCDVRFSTCDNQKSCMSNCSITSICEKPQEVCVAVWRKNDENITLET
VCHDPKLPYHDFILEDAASPKCIMKEKKKPGETFFMCSCSSDECNDNIIFSEEYNTSNPD
11111111110000000000000000000000000000000000000000000000000000
000000000000000000000000000000000000000000000000000000111111
>DM_validata1002
KKTSGNQDEILVIRKGWLTINNIGIMKGGSKEYWFVLTAENLSWYKDDEEKEKKYMLSVDNL
KLRDVEKGFMSSKHIFALFNTEQRNVYKDYRQLELACETQEEVDSWKASFLRAGVYPERV
11111111100000000000000000000000000000000000000000000000000000
000000000000000000000000000000000000000000000000000000000000
>DM_validata1003
MKEEKRSSTGFLVKQRAFLKLYMITMTEQERLYGLKLLEVLRSEFKEIGFKPNHTEVYRSLH
ELLDDGILKQIKVKKEGAKLQEVVLYQFKDYEAAKLYKKQLKVELDRSKKLIEKALSDNF
11111110000000000000000000000000000000000000000000000000000000
000000000000000000000000000000000000000000000000000000000000
>DM_validata1004
ENEIERPEDMRVRTLANKSKMKVSIVQQIDRKVALDDIAVSHGLDFPELLSEVETIVYSGTR
INIDYFINEVMDEDHLEDIFEYFKESTTDSLEEAMQELGKDYSEEEIRLVRIKFLSEMAN
11111111110000000000000000000000000000000000000000000000000000
000000000000000000000000000000000000000000000000000000000011
>DM_validata1005
WGLSVERSTYFLAPADRHYLADYARQAEDAWRREGAAGAERFRKELSAKEDTWVALVGPHLE
SLGSTPLSAEESSHLTFMRKLDWPMSRRLQDELPYVSIEFPGHPEQGRLVIQLPERLLPG
11111111100000000000000000000000000000000000000000000000000000
000000000000000000000000000000000000000000000000000000000001
>DM_validata1006
GPKLAALDPIASQFSQLRTISKADAESEELGFKDAADHHTDDVTHCLFGGELSLSNPDQQVI
GLAGNPTDTSQPYSQEGNKDLAFMDMKKLAQFLAGKPEHPMTRETLNAENIAKYAFRIVP
11000000000000000000000111111000000011110000000000000000000000
000000000000001111100000000000000000000000000000000000000000

>DM_validata1007
GMKEDTIEIAGFHAHVYFDAASRDVAARVREGLGARFEVQLGRWFDKPIGPHPKGMYQVAFL
PNQFDKVVPWLMLNREGLDILVHPETGDAVSDHAVYSLWLGAALALNIEFLRQLSSTSSN
11000000000000000000000000000000000000000000000000000000000000
000000000000000000000000000000000000000000000000000000011111
>DM_validata1009
GMTTSEHIAALTALVETYVMAMTRGDRPALERIFFGKASEVGHYEGELLWNSRDAFIAMCED
AADAETDPFWAISSVSVQGDIAMLHVENDWAGMRFDDFLTVLLHEGSWRIVSKVYRIR
10000000000000000000000000000000000000000000000000000000000000
0000000000000000000000000000000000000000000000000000000000
>DM_validata1010
ASLVVPNLMGNKDKADRQKVMSDLVALESTLDMYRLDNNRYPTTEQGLRALVSKPTVQPEPR
NYRQDGYIRRLPQDPWGGDYQLLNPGQYSDIDIFSPGPDGVPNTEDDIGNWTLGNAQP
10000000000000000000000000000000000000000000000000000000000000
0000000000000000000000000000000000000000000000000000011111
>DM_validata1011
MDIRYFGTTPRYSEAVGANGLIFLSGMVPENGETAAEQTADVLAQIDRWLAECGSDKAHVLD
AVIYLRDMGDYAEMNGVWDAWVAAGRTPARACVEARLARPEWRVEIKITAVKRDAATA
00000000000000000000000000000000000000000000000000000000000000
0000000000000000000000000000000000000000000000000000011111
>DM_validata1012
IGIPFPDHSSDILSGLNEQRTQGLLCDVVILVEGREFPTHRSVLAACSQYFKKLFTSGAVVD
QQNVYEIDFVSAEALTALMDFAYTATLTVSTANVGDILSAARLLEIPAVSHVCADLLD
00000000000000000000000000000000000000000000000000000000011111
0000000000000000000000000000000000000000000000000000000000
>DM_validata1013
GSHMKDPRPLNDKAFIQQCIRQLCEFLTENGYAHNVSMKSLQAPSVKDFLKIFTFLYGFLCP
SYELPDTKFEEEVPRIFKDLGYPFALSKSSMYTVGAPHTWPHIVAALVWLIDCIKIHT
11111000000000000000000000000000000000000000000000000000000000
0000000000000000000000000000000000000000000000000000000001
>DM_validata1014
MALKTDQILDKLNEKLAQVDRSKRSFTVILFVHLRQEGKVVRSVVLDFNDLKISEIELAVTS
TADYPAERIDASITIDDNDFYLVATKETSFAALIEQGKVDITGNKQAFLTLDEKFRNK
10000000000000000000000000000000000000000000000000000000000000
0000000000000000000000000000000000000000000000000000000000
>DM_validata1015
TWGVSSPKNVQGLSGSCLLIPCIFSYPADVPVSNGITAIWYYDYSGKRQVVIHSGDPKLVDK
RFRGRAELMGNMDHKVCNLLLKDLKPEDSGTYNFRFEISDSNRWLDVKGTTVTVTTD
00000000000000000000000000000001110000000000000000000000000000
000000000000000000000000000000000000000000000000000000000
>DM_validata1016
YAEHKSHRGEYSVCDSESLWVTDKSSAIDIRGHQVTVLGEIKTGNSPVKQYFYETRCKEARP
VKNGCRGIDDKHWNSQCKTSQTYVRALTSENNKLVGWRWIRIDTSCVCALSRKIGRT
11111111110000000000000000000000000000000111111000000000000111

```
111000000000000000000000000000000000000000000000001111111
>DM_validata1017
GMSPQSMLTSPQHPRRTTMVISHGTLSASAEHAAHLRQLLVHIAQATRQEDGCLLYLVSEDL
SQPGHFLITEHWDNLGAMHTHLALPGVTQAIDALKHLNVTDLKITAYEAGEAINIMG
111111111111111111000000000000000000000000000000000000000000
000000000000000000000000000000000000000000000000000000000
>DM_validata1018
SNAMFAPQGLAQFIKVNVTLENGEPVFIYTDANGQVCQGDITVTQAGTITYLLNDQTLKGLK
FVGVGFVTPFDGIIDAVTISSDGMLVQLVDLDKTPGTTKFQFVLSNTANTLLVLSPD
111111110000000000000000000000000000000000000000000000000000
000000000000000000000000000000000000000000000000000000000
>DM_validata1019
GSAPRVQEELCLVCGDRASGYHYNALTCEGCKGFFRRSVTKSAVYCCKFGRACEMDMYMRRK
CQECRLKKCLAVGMRPECVVPENQCAMKRREKKAQKEKDKMTTSPSSQHGSPGIHRD
111000000000000000000000000000000000000000000000000000000000
000000000000000000000000000001111111111111111111111111111
>DM_validata1020
MAQYPEQLNGIFQALADPTRRAVLGRLSRGPATVSELAKPFDMALPSFMKHIHFLEDSGWIR
THKQGRVRTCAIEKEPFTAVEAWLAEQQELWESRTDRLEQFVTEHTVTAHAKATPQ
111110000000000000000000000000000000000000000000000000000000
00000000000000000000000000000000001111111111111111111111
>DM_validata1021
GSDDSYARVRAVVMTRDDSSGGWLQLGGGGLSSVTVSKTLQPGDSGGTEFLVHGERLRDKTV
VLECVLRRDLVYNKVTPTFHHWRIGDKKFGLTFQSPADARAFDRGIRRAIEDLSQG
111100000000000000000000000000000000011111110000000000000000
00000000000000000000000000000000000000000000000000000000
>DM_validata1022
PSQVAFEIRGTLLPGEVFAICGSCDALGNWNPQNAVALLPENDTGESMLWKATIVLSRGVSV
QYRYFKGYFLEPKTIGGPCQVIVHKWETHLQPRSITPLESEIIIDDGQFGIHNGVE
000000000000000000000000000000000000000111110000000000000000
00000000000000000000000000000110000000000000000000111111
>DM_validata1023
SAQFASVTIRNAQTGRLLDSNYNGNVYTLPANGGNYQRWTGPGDGTVRNAQTGRCLDSNYDG
AVYTLPCNGGSYQKWLFYSNGYIQNVETGRVLDSNYNGNVYTLPANGGNYQKWYTG
110000000000000000000000000000000000000000000000000000000000
00000000000000000000000000000000000000000000000000000000
>DM_validata1025
TSSDTQNNKTLAAMKNFAEQYAKRTDTYFCSDLSVTAVVIEGLARHKEELGSPLCPCRHYED
KEAEVKNTFWNCPCVPMRERKECHCMLFLTPDNDFAGDAQDIPMETLEEVKASMA
111111000000000000000000000000000000000000000000000000000000
0000000000000000000000000000000000000000000000000000000
>DM_validata1026
GMIEKVYEFKRDAKTKVVEKLVNTEHVQINHIVLPRGEQMPKHYSNSYVHLIIIKGEMTLTL
EDQEPHNYKEGNIVYVPFNVKMLIQNINSDILEFFVVKAPHPKKLNAPEDPIKCE
```

000000000000000000000000000000000000000000000000000000000000
0000000000000000000000000000000000000000000000011111111
>DM_validata1027
MAMSNMTYNNVFDHAYEMLKENIRYDDIRDTDDLHDAIHMAADNAVPHYYADIFSVMASEGI
DLEFEDSGLMPDTKDVIRILQARIYEQLTIDLWEDAEDLLNEYLEEVEEYEEDEE
111110000000000000000000000000000000000000000000000000000000
0000000000000000000000000000000000000000000000000111111
>DM_validata1030
MQPWGRLLRLGAEEGEPHVLLRKREWTIGRRRGCDLSFPSNKLVSGDHCRIVVDEKSGQVTL
EDTSTSGTVINKLKVVKKQTCPLQTGDVIYLVYRKNEPEHNVAYLYESLSEKQG
000000000000000000000000000000000000000000000000000000000000
000000000000000000000000000000000000000000000000000111
>DM_validata1031
FTPVTKAAVINTFDGVADYLIRYKRLPNDYITKSQASALGWVASKGDLAEVAPGKSIGGDVF
SNREGRLPSAGSRTWREADINYVSGFRNADRLVYSSDWLIYKTTDHYATFTRIR
111111110000000000000000000000000000000000000000000000000000
000000000000000000000000000000000000000000000000000000
>DM_validata1032
KALYSKPGSKNGRWNPETHKFCKCGVRIQTSAYTCSKCRNRSGENNSFFNHKHSDITKSKIS
EKMKGKKPSNIKKISCDGVIFDCAADAARHFKISSGLVTYRVKSDKWNWFYINA
111111111111111111100000000000000000000000000000000000000000
000000000000000000000000000000000000000000000000000000
>DM_validata1033
KENGKRTEVKIGAKTSVMKEKDGKLFTGKANKETNKVDGANATEDADEGKGLVTAKDVIDAV
NKTGWRIKTTDANGQNGDFATVASGTNVTFASGNGTTATVTNGTDGITVKYDAK
111111000000000000000000000000000000000000000000000000000000
000000000000000000000000000000000000000000000000000000
>DM_validata1034
MSYKEKSVRIAVYGTLRKGKPLHWYLKGAKFLGEDWIEGYQLYFEYLPYAVKGKGKLKVEVY
EVDKETFERINEIEIGTGYRLVEVSTKFGKAFLWEWGSKPRGKRIKSGDFDEIR
111111000000000000000000000000000000000000000000000000000000
000000000000000000000000000000000000000000000000000000
>DM_validata1035
GMTGAVETESMKTVRIREKIKKFLGDRPRNTAEILEHINSTMRHGTTSQQLGNVLSKDKDIV
KVGYIKRSGILSGGYDICEWATRNWVAEHCPEWTEGQPIILNEEGDFTLGPLPE
111111000000000000000000000000000000000000000000000000000000
000000000000000000000000000000000001111111111111111111
>DM_validata1036
MGFAFKRGISTPDLALITRQLATLVQSGMPLEECLRAVAEQSEKPRIRTMLVAVRAKVTEGY
TLSDSLGDYPHVFDELFRSMVAAGEKSGHLDSVLERLADYAENRQKMRSKLQQA
111111100000000000000000000000000000000000000000000000000000
000000000000000000000000000000000000000000000000000000
>DM_validata1037
MTLKQVIVVRDDLKLSRGKLAVQVAHAAIIGYLKSDSSLRRKWLDEGQKKVVLKVKSLEELL

GIKHKAESLGLVTGLVQDAGLTEVPPGTITAVVIGPDEERKIDKVTGNLPLLK
10000000000000000000000000000000000000000000000000000000000000
00000000000000000000000000000000000000000000001100001
>DM_validata1038
MNRHSAGQSTPVRVATLDQLKPGVPTAFDVDGDEVMVVRDGDSVYAISNLCSHAEAYLDMGV
FHAESLEIECPLHVGRFDVRTGAPTALPCVLPVRAYDVVVDGTEILVAPKEAD
11111111000000000000000000000000000000000000000000000000000000
00000000000000000000000000000000000000000000000000111
>DM_validata1039
HHHHHHVGVERTMAEDIHHWSECEAIIERLYPELERRLAKVKPDLLIARQGVKLKFDDFQQT
TQEHVWPRLNKADLIATARKTWDERRGGRGVRLVGLHVTLLDPQMERQLVLGL
11110000000000000000000000000000000000000000000000000000000000
00000000000000000000000000000000000000000000000000000
>DM_validata1040
GHMQRERTAESSLRVISKEKNSITVEMINYDNTLLRTLVEEILKDDQVDEARYYIKHPVIDN
PQIYVRVKSGKPQSAIKRAVRKLSKLYEDLGTQFQKEFQRYESDHMIKAVEGS
11111111100000000000000000000000000000000000000000000000000000
00000000000000000000000000000000000000000000011111111
>DM_validata1041
SSSNSEKEWHIVPVSKDYFSIPNDLLWSFNTTNKSINVYSKCISGKAVYSFNAGKFMGNFNV
KEVDGCFMDAQKIAIDKLFSMLKDGVVLKGNKINDTILIEKDGEVKLKLIRGI
11111000000000000000000000000000000000000000000000000000000000
00000000000000000000000000000000000000000000000000000
>DM_validata1042
GSTYPPTPPNVTRLSDESVMLRWMVPRNDGLPIVIFKVQYRMVGKRKNWQTTNDNIPYGKPK
WNSELGKSFTASVTDLKPQHTYRFRILAVYSNNDNKESNTSAKFYLQPGAALD
00000000000000111000000000000000000000000000111000000000000000
00000000000000000000000000000000000000000000000011111
>DM_validata1043
MESHTAVQGLAGHPVTLPCIYSTHLGGIVPMCWGLGECRHSYCIRSLIWTNGYTVTHQRNSR
YQLKGNISEGNVSLTIENTVVGDGGPYCCVVEIPGAFHFVDYMLEVKPELVPR
11111000000000000000000000000000000000000000000000000000000000
00000000000000000000000000000000000000000000000000111
>DM_validata1044
MLSPELLAKAFPFHFAFSRNREIVQTGEVLERISPEPLVGKLIEQHFQINRPKILIDFDAIS
KQPRALFILEFLHNGMQLKGQMMYQPEEEVIFFLGSPWITDTTSLAPLGIKLK
00000000000000000000000000000000000000000000000000000000000000
00000000000000000000000000000000000000000000011111111
>DM_validata1045
MPELEVKGKKLRLDEDGFLQDWEEWDEEVAEALAKDTRFSPQPIELTEEHWKIIRYLRDYFI
KYGVAPPVRMLVKHCKKEVRPDCNLQYIYKLFPQGPAKDACRIAGLPKPTGCV
10000000000000000000000000000000000000000000000000000000000000
00000000000000000000000000000000000000000000000000000
>DM_validata1047

GMSLPHLSSGEVASVLPLGKQLTQTPSAALFKEHRLEVMRMVLPAGKQVGSHSVAGPSTIQC
LEGEVEIGVDGAQRRLHQGDLLYLGAGAAHDVNAITNTSLLVTVVLVDRGGS
11111110000000000000000000000000000000000000000000000000000000
0000000000000000000000000000000000000000000000011111
>DM_validata1048
GMAYGSSNPEDLAKNFTKDLYSGDTKSVMSYIDLSEAKSDEEKTFVSDKITQVVAENAAKAK
RMGGVKDIQIEEKTINKDSAKIRVLVLFNNDNNQSSNVFLAKKDRKWLVLLK
11111100000000000000000000000000000000000000000000000000000000
0000000000000000000000000000000000000000000000000000
>DM_validata1049
MTSFAQLDIKSEELAIVKTILQQLVPDYTVWAFGSRVKGKAKKYSDLDLAIISEEPLDFLAR
DRLKEAFSESDLPWRVDLLDWATTSEDFREIIRKVYVVIQEKEKTVEKPTAL
11111000000000000000000000000000000000000000000000000000000000
0000000000000000000000000000000000000000000011111111
>DM_validata1050
QGMTVGTYAELASVFAALSDETRWEILTELGRADQSASSLATRLPVSRQAIAKHLNALQACG
LVESVKVGREIRYRALGAELNKTARTLERIGAEWDRRLAAIKQIAESMEEGS
11110000000000000000000000000000000000000000000000000000000000
0000000000000000000000000000000000000000000000001111
>DM_validata1051
GMARLKNLPQERPLPLASLIEARENQVLSMALAQSDRVQISLFSFADGESVSEEEYFGDTLY
LILQGEAVITFDDQKIDLVPEDVLMVPAHKIHAIAGKGRFKMLQITLIDEER
11000000000000000000000000000000000000000000000000000000000000
0000000000000000000000000000000000000000000000000111
>DM_validata1052
TVPGATTTSSAATETMENVKKCKNFLSTLIKLASSGKQSTETAANVKELVQNLLDGKIEAED
FTSRLYRELNSSPQPYLVPFLKRSLPALRQLTPDSAAFIQQSQQQPPPPTSQ
11111111000000000000000000000000000000000000000000000000000000
0000000000000000000000000000000000000000000011111111
>DM_validata1053
MPITWMRMRPWLEMQINSNQIPGLIWINKEEMIFQIPWKHAAKHGWDINKDACLFRSWAIHT
GRYKAGEKEPDPKTWKANFRCAMNSLPDIEEVKDQSRNKGSSAVRVYRMLP
11111110000000000000000000000000000000000000000000000000000000
000000000000000000000000000000000000000000000000000
>DM_validata1054
AMGIKHLNLTVADVVAAREFLEKYFGLTCSGTRGNAFAVMRDNDGFILTLMKGKEVQYPKTF
HVGFPQESEEQVDKINQRLKEDGFLVEPPKHAHAYTFYVEAPGGFTIEVMC
10000000000000000000000000000000000000000000000000000000000000
000000000000000000000000000000001000000000000000000
>DM_validata1055
MTDPALRAATLLQLHFAFNGPFGDAMAEQLKPLAESINQEPGFLWKVWTESEKNHEAGGIYL
FTDEKSALAYLEKHTARLKNLGVEEVVAKVFDVNEPLSQINQAKLAGLCGR
11111111000000000000000000000000000000000000000000000000000000
000000000000000000000000000000000000000000011111111

>DM_validata1056
AFACSFPPSEIPGSKECLAEALQKHQGFKKKSYALICAYLNYKEDAENYERAAEDFDSAVKC
TGCKEGVDLHEGNPELIEEGFEKFLASLKIDRKALGSLCTLFQKLYAIPHN
000000000000000000000000000000000000000000000000000000000000
000000000000000000000000000000000000000000000000001
>DM_validata1057
MKKLLEELETKKTTIEEERNEITQRIGELKNKIGDLKTAIEELKKAKGKCPVCGRELTDEHR
EELLSKYHLDLNNSKNTLAKLIDRKSELERELRRIDMEIKRLTPLLTVAE
000000000000000000000000000000000000000000000000000000000000
00000000000000000000000000000000000000001111111111
>DM_validata1058
IRMYFEPGHYTVMENCGEFEVRVVRRGDISTYASVEYETQDGTASAGTDFVGRKGLLSFPPG
VDEQRFRIEVIDDDVFEEDECFYIRLFNPSEGVKLAVPMIATVMILDDDH
000000000000000000000000000000000000000000000000000000000000
00000000000011111000000000000000000000000000001111
>DM_validata1059
MIGQRIKQYRKEKGYSLSELAEKAGVAKSYLSSIERNLQTNPSIQFLEKVSAVLDVSVHTLL
DEKHETEYDGQLDSEWEKLVRDAMTSGVSKKQFREFLDYQKWRKSQKEE
000000000000000000000000000000000000000000000000000000000000
0000001111100000000000000000000000000000000000111
>DM_validata1060
MSEVQQLPIRAVGEYVILVSEPAQAGDEEVTESGLIIGKRVQGEVPELCVVHSVGPDVPEGF
CEVGDLTSLPVGQIRNVPHPFVALGLKQPKEIKQKFVTCHYKAIPCLYK
111100000000000000000000000000000000000000000000000000000000
0000000000000000000000000000000000000000000000000
>DM_validata1061
VAQFPTPFGGSLDTWAITVEERAKHDQQFHSLKPISGFITGDQARNFFFQSGLPQPVLAQIW
ALADMNNDGRMDQVEFSIAMKLIKLKLQGYQLPSALPPVMKQQPVAISS
111110000000000000000000000000000000000000000000000000000000
0000000000000000000000000000000000000000011111111
>DM_validata1062
MDDSREESQFIVDDVSKTIKEAIETTIGGNAYQHDKVNNWTGQVVENCLTVLTKEQKPYKYI
VTAMIMQKNGAGLHTASSCYWNNDTDGSCTVRWENKTMYCIVSVFGLAV
111111100000000000000000000000000000000000000000000000000000
0000000000000000000000000000000000000000000000000
>DM_validata1063
GEFPLQTKTDIHAVLASNGRIIYISANSKLHLGYLQGEMIGSFLKTFLHEEDQFLVESYFYN
EHHLMPCTFRFIKKDHTIVWVEAAVEIVTTRAERTEREIILKMKVLEEE
111111110000000000000000000000000000000000000000000000000000
0000000000000000000000000001111110000000000000111
>DM_validata1064
MHSLQNVIPQQQAHIAELQVYNNKLERDLQNKIGSLTSSIEWYLRSMELDPEIKADIEQQIN
SIDAINPLHAFDDLESVIRNLISDYDKLFLMFKGLIQRSNYQYSFGSE
000000000000000000000000000000000000000000000000000000000000

000000000000000000000000000000000000000000000111
>DM_validata1065
VSRYVPDMGDLIWVDFDPTKGSEQAGHRPAVVLSPFMYNNKTGMCLCVPCTTQSKGYPFEVV
LSGQERDGVALADQVKSIAWRARGATKKGTVAPEELQLIKAKINVLIG
000000000000000011111111110000000000000000000000000000000000
000000000000000000000000000000000000000000000000
>DM_validata1066
MRRRENMDVNKAIRTAVDTGKVILGSKRTIKFVKHGEGKLVVLAGNIPKDLEEDVKYYAKLS
NIPVYQHKITSLELGAVCGKPFPVAALLVLDEGLSNIMELVEKKEGGE
111111000000000000000000000000000000000000000000000000000000
000000000000000000000000000000000000000000011111
>DM_validata1067
MGEVEAPGRLWLESPPGEAPPIFLPSDGQALVLGRGPLTQVTDRKCSRTQVELVADPETRTV
AVKQLGVNPSTTGTQELKPGLEGSLGVGDTLYLVNGLHPLTLRWEETR
111111100000000000000000000000000000000000000000000000000000
000000000000010000000000000000000000000000000011
>DM_validata1068
GMEAHMKSHNLLEAVRFDDQRFVMELVHESENFKIVSFTFKAGQELPVHSHNIEGELNIVVL
EGEGEFVGDGDAVIPAPRGAVLVAPISTPHGVRAVTDMKVLVTIAPPI
111111000000000000000000000000000000000000000000000000000000
000000000000000000000000000000000000000000000000
>DM_validata1069
GMITSIARQSIILKCLRQKSVLVSNYELYYTAGLAKKCFGIAVDADMEPKQLLEELQKHIDK
VSPADEQEKYLIHLLGNYEPDDTHDEQTVELFHMGETEEHIWQVSIT
000000000000000000000000000000000000000000000000000000000000
00000000000000000000000000000000000000000000011
>DM_validata1070
EQGGFSGPSATQSQAGGFQGPNGSVTTVESAKSLRDDTWVTLRGNIVERISDDLYVFKDASG
TINVDIDHKRWNGVTVTPKDTVEIQGEVDKDWNSVEIDVKQIRKVNP
100000000111111111111100000000000000000000000000000000000000
00000000000000000000000000000000000000000000011
>DM_validata1071
GAEFEEPRVIDLWDLAQSANLTDKELEAFREELKHFEAKIEKHNHYQKQLEIAHEKLRHAES
VGDGERVSRSREKHALLEGRTKELGYTVKKHLQDLSGRISRARHNEL
000000000000000000000000000000000000000000000000000000000000
00000000000000000000000000000000000000000000111
>DM_validata1072
MTNKLFEHTVLYDSGDAFFELKGNASMKLSPKAAIEVCNEAAKKGLWILGIDGGHWLNPGFR
IDSSASWTYDMPEEYKSKIPENNRLAIENIKDDIENGYTAFIITLKM
111000000000000000000000000000000000000000000000000000000000
00000000000000000000000000000000000000000000000
>DM_validata1073
MSLTADPPACTVPAAGVSSTHKLVNGGAEKIVFKIKSSNNNEYRIAPVFGFVDPSGSKDVVI
TRTAGAPKEDKLVVHFASAPADATDAQAAFVAVAPAGTVTIPMSATA

11000000000000000000000000000000000000000000000000000000000000
00000000000000000000000000000000000000000000000
>DM_validata1074
GANTTETPVKDVELDGRWDDPIRSAATNCPITVFTDGYLLTLKNASPDRDMTIRITDMAKGG
VVYENDIPEVQSAYITISIANFPAEEYKLEITGTPSGHLTGYFTKE
11111110000000000001111111100000000000000000000000000000000000
0000000000000000000000000000000000000000000000
>DM_validata1075
MRMDEFYTKVYDAVCEIPYGKVSTYGEIARYVGMPSYARQVGQAMKHLHPETHVPWHRVINS
RGTISKRDISAGEQRQKDRLEEEGVEIYQTSLGEYKLNLPEYMWKP
10000000000000000000000000000000000000000000000000000000000000
0000000000000000000000000000000000000000000000
>DM_validata1076
GMKGENMMVKLIALYEQPEDKQAFDEHYFNTHAPLTRKIPGLRDMKVTRIVGSPMGESKFYL
MCEMYYDDHESLQQAMRTDEGKASGKDAMKFAGKLLTLMIGEEMDE
11111100000000000000000000000000000000000000000000000000000000
0000000000000000000000000000000000000000000011
>DM_validata1077
MVTYTTLLDKPISESAPRKAPEPLLREALGAALRSFRADKGVTLRELAEASRVSPGYLSELE
RGRKEVSSELLASVCHALGASVADVLIEAAGSMALQAAQEDLARV
11111111111111111111100000000000000000000000000000000000000000
000000000000000000000000000000000000111111111
>DM_validata1078
DDMIILKGVNIFPIQIETILLQFKELGSDYLITLETAESNDEMTVEVELSQLFTDDYGRLQA
LTREITRQLKDEILVTPRVKLVPKGALPKSEGKAVRVKDLRKTF
00000000000000000000000000000000000000000000000000000000000000
00000000000000000000000000000011100000000000
>DM_validata1079
MGSDKIHHHHHHMPKVTVSIKVVPAVEDGRLHEVIDRAIEKISSWGMKYEVGPSNTTVEGEF
EEIMDRVKELARYLEQFAKRFVLQLDIDYKAGGITIEEKVSKYR
11111111111110000000000000000000000000000000000000000000000000
00000000000000000000000000000000000000000000
>DM_validata1080
ANETTKQESPVVDTDINAVTNYIVGMCQKFLQKGEKVTPSSKLEELRTREDRLWDCLDTVEF
VLDVEEIFDVTVPDEVADNFQTLQEIADFVVSERAKAGKFMKDQ
11111111110000000000000000000000000000000000000000000000000000
00000000000000000000000000000000000000111111
>DM_validata1082
MNREEMTLLGFEIVAYAGDARSKLLEALKAAENGDFAKADSLVVEAGSCIAEAHSSQTGMLA
REASGEELPYSVTMMHGQLHLMTTILLKDVIHHLIELYKRGAK
00000000000000000000000000000000000000000000000000000011111000
0000000000000000000000000000000000000000000
>DM_validata1083
MPKIRIKLRGFDHKTLDASAQKIVEAARRSGAQVSGPIPLPTRVRRFTVIRGPFKHKDSREH

FELRTHNRLVDIINPNRKTIEQLMTLDLPTGVEIEIKTVGGGR
00000000000000000000000000000000000000000000000000000000000000
0000000000000000000000000000000000000011111
>DM_validata1084
MSQFTLYKNKDKSSAKTYPYFVDVQSDLLDNLNTRLVIPLTPIELLDKKAPSHLCPTIHIDE
GDFIMLTQQMTSVPVKILSEPVNELSTFRNEIIAAIDFLITGI
1000000000000000000000000000000000000000000011111110000000000
0000000000000000000000000000000000000000000
>DM_validata1085
MDGNEEMGGITQTPYKVSISGTTVILTCPQYPGSEILWQHNDKNIGGDEDDKNIGSDEDHLS
LKEFSELEQSGYYVCYPRGSKPEDANFYLYLRARVCENCMEMD
11111111111000000000000000000000000000000000000000000000000000
0000000000000000000000000000000000001111111
>DM_validata1086
MKSHKMMGGGISMHYITACLKIISDKDLNEIMKEFKKLEEETNKEEGCITFHAYPLEPSERK
IMLWEIWENEEAVKIHFTKKHTIDVQKQELTEVEWLMKSNVND
11111111111000000000000000000000000000000000000000000000000000
0000000000000000000000000000000000000000001
>DM_validata1087
TYVIAEPCVDVKDKACIEECPVDCIYEGARMLYIHPDECVDCGACEPVCPVEAIYYEDDVPD
QWSSYAQANADFFAELGSPGGASKVGQTDNDPQAIKDLPPQGE
00000000000000000000000000000000000000000000000000000000000000
0000000000000000000000000000000000000000011
>DM_validata1088
QQLPQPSLSLHPSQGVSLGDTVTLRCHLPRMAAWVQLWLNGTLRFDKEKDKEQDAAEFSFAV
TNLEDAGTYQCRYQVSEPLWTSNQSDPVELVLTIEGRHHHHHH
11000000000000000000000000000000000000000000000000000000000000
0000000000000000000000000000000001111111111
>DM_validata1089
MDLSTKQKQHLKGLAHPLKPVVLLGSNGLTEGVLAEIEQALEHHELIKVKIATEDRETKTLI
VEAIVRETGACNVQVIGKTLVLYRPTKERKISLPLEHHHHHH
00000000000000000000000000000000000000000000000000000000000000
000000000000000000000000000000000000111111
>DM_validata1090
AHPELKSSVPQADSAVAAPEKIQLNFSENLTVKFSGAKLTMTGMKGMSSHSPMPVAAKVAPG
ADPKSMVIIPREPLPAGTYRVDWRAVSSDTHPITGNYTFTVK
00000000000000000000000000000000000000000000111000000000000000
000000000000000000000000000000000000000000
>DM_validata1091
MSQEQYTENLKVIVAEKLAGIPNFNEDIKYVAEYIVLLIVNGGTVESVVDELASLFDSVSRD
TLANVVQTAFFALEALQQGESAENIVSKIRMMNAQSLGQSDI
11111000000000000000000000000000000000000000000000000000000000
000000000000000000000000000000000111111111
>DM_validata1092

GMDNWQDITDDRLVRPIHPGEVIADILDDLDINTANFAEILGVSNQTIQEVINGQRSITVDI
AIRLGKALGNGPRLWLNLQQKVDLWYALQSHKEEYEQVMTLV
11111111111110000000000000000000000000000000000000000000000000
000000000000000000000000000000000000000000
>DM_validata1093
MADIDQASKTEMEAAAFRHLLRHLDEHKDVQNIDLMIQADFCRNCLAKWLMEAATEQGVELD
YDGAREYVYGMPFAEWKTLYQKPASEAQLAAFEAKQAARKR
00000000000000000000000000000000000000000000000000000000000000
00000000000000000000000001111111111111111
>DM_validata1095
GSSKAASLHWTSERAVSALLLGLLPAAYLYPGPAVDYSLAAALTLHGHWGLGQVITDYVHGD
TPIKVANTGLYVLSAITFTGLCYFNYYDVGICKAVAMLWSI
11000000000000000000000000000000000000000000000000000000000000
00000000000000000000000000000000000000000
>DM_validata1096
MTLHSWLDRYEKILASRGIKQKTLINYMSKIKAIRRGLPDAPLEDITTKEIAAMLNGYIDEG
KAASAKLIRSTLSDAFREAIAEGHITTNHVAATRAAKSEVR
00000000000000000000000000000000000000000000000000000000000000
00000000000000000000000000000001111111111
>DM_validata1097
GSRRASVGSMARPAFVNKLWSMVNDKSNEKFIHWSTSGESIVVPNRERFVQEVLPKYFKHSN
FASFVRQLNMYGWHKVQDVKSGSMLSNNDSRWEFENERHA
11111111100000000000000000000000000000000000000000000000000000
0000000000000000011111111100000000000000
>DM_validata1098
MAGSGVAKQIFALDFEIFGRVQGVFFRKHTSHEAKRLGVRGWCMNTRDGTVKGQLEAPMMNL
MEMKHWLENNRIPNAKVSKAEFSQIQEIEDYTFTSFDIKH
11111000000000000000000000000000000000000000000000000000000000
0000000000000000000000000000000000000000
>DM_validata1099
GSHMASGDATDITIYYKTGWTHPHIHYSLNQGAWTTLPGVPLTKSEYEGYVKVTIEAEEGSQ
LRAAFNNGSGQWDNNQGRDYDFSSGVHTLADGRILSGTPK
11111110000000000000000000000000000000000000000000000000000000
0000000000000000000000000000000000000001
>DM_validata1100
SDTGRPFVEMYSEIPEIIHMTEGRELVIPCRVTSPNITVTLKKFPLDTLIPDGKRIIWDSRK
GFIISNATYKEIGLLTCEATVNGHLYKTNYLTHRQTNTI
11100000000000000000000000000000000000000000000000000000000000
000000000000000000000000000000000001111
>DM_validata1101
MAAPILKDVVAYVEVWSSNGTENYSKTFTTQLVDMGAKVSKTFNKQVTHVIFKDGYQSTWDK
AQKRGVKLVSVLWVEKCRTAGAHIDESLFPAANMNEHLS
11100000000000000000000000000000000000000000000000000000000000
000000000000000000000000000000000111111

>DM_validata1102
MRGSHHHHGSRLQSGKMTGIVKWFNADKGFGFITPDDGSKDVFVHFSAGSSGAAVRGNPQQG
DRVEGKIKSITDFGIFIGLDGGIDGLVHLSDISWAQAEA
111111111111111000000000000000000000000000000000000000000000
000000000000000000000000000000000000001
>DM_validata1103
MPKYAPHVYTEQAQIATLEHWVKLLDGQERVRIELDDGSMIAGTVAVRPTIQTYRDEQEREG
SNGQLRIDHLDASQEPQWIWMDRIVAVHPMPLGAPQVMP
000000000000000000000000000000000000000000000000000000000000
000000000000000000000000000000011111111
>DM_validata1104
MRLTEEEVQALLEKADGWKLADERWIVKKYRFQDYLQGIEFVRRIAAISENANHHPFISIDY
KLITVKLSSWRAKGLTKLDFDLAKQYDEVYNQMKQGEGE
000000000000000000000000000000000000000000000000000000000000
000000000000000000000000000000000011111
>DM_validata1105
AMGQTVTTPLSLTLGHWKDVERIAHNQSVDVKKRRWVTFCSAEWPTFNVGWPRDGTFNRDLI
TQVKIKVFSPGPHGHPDQVPYIVTWEALAFDPPPWVKP
111111000000000000000000000000000000000000000000000000000000
00000000000000000000000000000000000000
>DM_validata1106
GSHMKTRKIPLRKSVVSNEVIDKRDLLRIVKNKEGQVFIDPTGKANGRGAYIKLDNAEALEA
KKKKVFNRSFSMEVEESFYDELIAYVDHKVKRRELGLE
111111000000000000000000000000000000000000000000000000000000
00000000000000000000000000000000000000
>DM_validata1107
MGRFVVWPSELDSRLSRKYGRIVPRSIAVESPRVEEIVRAAEELKFKVIRVEEDKLNPRLSG
IDEELRTFGMIVLESPYGKSKSLKLIAQKIREFRRRSA
100000000000000000000000000000000000000000000000000000111111
11110000000000000000000000000000000011
>DM_validata1108
MQTNVPKFTAVAEQVSAVLSQYGITGPNRAIYQGFGLKVARALNRLGGGPALVNMINGLKAY
YISAFNANPTVLDAVTDIITGSPTGYVSGGSGHHHHHH
111111100000000000000000000000000000000000000000000000000000
00000000000000000000000000001111111111
>DM_validata1109
SEAEAKLWQHLRAGRLNGYKFRRQQPMGNYIVDFMCVTPKLIVEADGGQHAEQAVYDHARTV
YLNSLGFTVLRFWNHEILQQTNDVLAEILRVLQELEKQ
000000000000000000000000000000000000000000000111111100000000
00000000000000000000000000000000000001
>DM_validata1110
MTKVTREEVEHIANLARLQISPEETEEMANTLESILDFAKQNDSADTEGVEPTYHVLDLQNV
LREDKAIKGIPQELALKNAKETEDGQFKVPTIMNEEDA
110000000000000000000000000000000000000000000000000000000000

0000000000000000000000000000000111111
>DM_validata1111
MGSDKIHHHHHHMERKKLIAKFVEIASEKMGKDLETVDEENTFKELGFDSIDVIDLVMFFED
EFALRIEDEEISKIRKVKDLIDIVIKKLEEIDDEVSEG
11111111100000000000000000000000000000000000000000000000000000
0000000000000000000000000000000111111
>DM_validata1112
MGSSSQDPLVPRGSEVKPEVKPETHINLKVSDGSSEIFFKIKKTTPLRRLMEAFAKRQGKEM
DSLRFLYDGIRIQADQTPEDLDMEDNDIIEAHREQIGG
11111100000000000110000000000000000000000000000000000000000000
00000000000000000000000000000000001111
>DM_validata1113
ALCAFKDPYQQDLGIGESRISHENGTILCSKGSTCYGLWEKSKGDINLVKQGCWSHIGDPQE
CHYEECVVTTTPPSIQNGTYRFCCCSTDLCNVNFTENF
00000000011111111111111000000000000000001111111000000000000000000
000000000111111111100000000000000000011
>DM_validata1115
MSLVEATLEVIGGKWKCVILCHLTHGKKRTSELKRLMPNITQKMLTQQLRELEADGVINRIV
YNQVPPKVEYELSEYGRSLEGILDMLCAWGANHINRV
00000000000000000000000000000000000000000000000000000000000000
0001110000000000000000000000000000001
>DM_validata1116
MSKGRDILTKTIILALREVAPGLEAVLEAHLRATLNSGIELAYDDPQKFKEAVSKLFGEYSA
RLLEMVIISKLKGRLGEDIEANSLEELVSEIRKIYGE
11000000000000000000000000000000000000000000000000000000000000
000000000000000011000000000000000000
>DM_validata1118
GMSAPVTLINPFKVPADKLEAAIEYWEAHRDFMAQQPGYLSTQLHQSIDEGATYQLINVAIW
QSEADFYQAAQKMRQALGHVQVEGLCGNPALYRVIRT
00000000000000000000000000000000000000000000000000000000000000
000000000000000000111100000000000000000
>DM_validata1119
APLAAPSKPIMVTVEEQRSQSVRPGADVTFICTAKSKSPAYTLVWTRLHNGKLPSRAMDFNG
ILTIRNVQPSDAGTYVCTGSNMFAMDQGTATLHVQV
11111111000000000000000000000000000000000000000000000000000000
000000000000000000000000000000000000
>DM_validata1120
MEKRTPHTRLSQVKKLVNAGQVRTTRSALLNADELGLDFDGMCNVIIGLSESDFYKSMTTYS
DHTIWQDVYRPRLVTGQVYLKITVIHDVLIVSFKEK
10000000000000000000000000000000000000000000000000000000000011
100000000000000000000000000000000111
>DM_validata1121
MTGPKQQPLPPDVEGREDAIEVLRAFVLDGGLSIAFMRAFEDPEMWGLLLVDIARHAARSYA
RESEYTEDEALERIVEMFEAELSRPTDTGATTERTQ

0000000000000000000000000000000000000010000000000000000000000
000000000000000000000000111111111111
>DM_validata1122
MNLKDMEQNSAKAVVLLKAMANERRLQILCMLHNQELSVGELCAKLQLSQSALSQHLAWLRR
DGLVTTRKEAQTVYYTLKSEEVKAMIKLLHSLYCEE
1111000000000000000000000000000000000000000000000000000000000
000000000000000000000000000000000011
>DM_validata1123
GIPPKIEALPSDISIDEGKVLTVACAFTGEPTPEVTWSCGGRKIHSQEQGRFHIENTDDLTT
LIIMDVQKQDGGLYTLSLGNEFGSDSATVNIHIRSI
0000000000000000000000000000000000000000000000000000000000000
000000000000000000000000000000000001
>DM_validata1124
GVYHREARSGKYKLTYAEAKAVCEFEGGHLATYKQLEAARKIGFHVCAAGWMAKGRVGYPIV
KPGPNCGFGKTGIIDYGIRLNRSERWDAYCYNPHAK
0000000000000000000000000000000000000000000000000000000000001
111111111111000000000000000000000001
>DM_validata1126
GLGFMALDEDLRIIYVNSGCLRHVRRSRDELLGRVVTEVLPETQGSYFDALCRKVLATGREQ
QTRVDSLYSPGMTIEVTAAADSGALVVHFRDVTAE
0000000000000000000000000000000000000000000000000000000000000
00000000000000000000000000000000011
>DM_validata1127
KNIEETELVVEEMLEQYPNGKIVRLLFHGEQAKLPIISHIVQEYQVEVSIIQGNIQQTKQGA
VGSLYIQLLGEEQNILAAIEGLRKLRVETEVIGNE
1111111000000000000000000000000000000000000000000000000000000
00000000000000000000000000000000000
>DM_validata1129
MSQIMYNYPAMMAHAGDMAGYAGTLQSLGADIASEQAVLSSAWQGDTGITYQGWQTQWNQAL
EDLVRAYQSMSGTHESNTMAMLARDGAEAAKWGG
1111111111111111111000000000000000000000000000000000000000000
0000000000001111111111111111111111
>DM_validata1130
MANIKIRQETPTAFYIKVHDTDNVAIIVNDNGLKAGTRFPDGLELIEHIPQGHKVALLDIPA
NGEIIRYGEVIGYAVRAIPRGSWIDESMVVLPEA
0000000000000000000000000000000000000000000000000000000000000
0000000000000000000000000000000111
>DM_validata1131
GAHRGRVTRRNIIWHELIGLRVRIVGSTHPAFVGIEGYVIDETRNMLVIAGDRIWKVPKDVS
IFEFEADDGTKIKIPGERLVGRPEMRLKKRWKKW
1111100000000000000000000000000000000000000000000000000000000
0000000000000000000000000000000000
>DM_validata1132
GMSDPGNEQNGDGIDPAIVEVLLVLREAGIENGATPWSLPKIAKRAQLPMSVLRRVLTQLQA

AGLADVSVEADGRGHASLTQEGAALAAQLFPDPF
111111111111000000000000000000000000000000000000000000000000
0000000000000000000000000000000111
>DM_validata1133
RSKKGDKNGKGLRHFSMKVCEKVQRKGTTSYNEVADELVSEFTNSNNHLAADSAYDQKNIRR
RVYDALNVLMAMNIISKEKKEIKWIGLPTNSAQ
111111110000000000000000000000000000000000000000000000000000
000000000000000000000000000011111
>DM_validata1134
MAVPETRPNHTIYINNLNEKIKKDELKKSLYAIFSQFGQILDILVSRSLKMRGQAFVIFKEV
SSATNALRSMQGFPFYDKPMRIQYAKTDSDIIA
111110000000000000000000000000000000000000000000000000000000
000000000000000000000000000001111
>DM_validata1135
NYKISELMPNLSGTINAEVVTAYPKKEFSRKDGTKGQLKSLFLKDDTGSIRGTLWNELADFE
VKKGDIAEVSGYVKQGYSGLEISVDNIGIIEKS
000000000000000000000000000111100000000000000000000000000000
000000000000000110000000000000000
>DM_validata1136
VRPVVGEIAANSIAAEAQIAPGTELKAVDGIETPDWDAVRLQMVDKIGDESTTITVAPFGSD
QRRDVKLDLRHWAFEPDKEDPVSSLGIRPRGPQ
000000000000000000000000000000000000000000000000000000001111
100000000000000000000000000000111
>DM_validata1137
GSRTGSESSRPNTALLDPSSPEFSAVVSVGDWLQAIKMDRYKDNFTAAGYTTLEAVVHMSQD
DLARIGITAITHQNKILSSVQAMRTQMQQMHG
111111111111111111111100000000000000000000000000000000000000
00000000000000000000000000000000
>DM_validata1138
MELKNSISDYTEAEFVQLLKEIEKENVAATDDVLDVLLEHFVKITEHPDGTDLIYYPSDNRD
DSPEGIVKEIKEWRAANGKPGFKQGKPGFKQG
000000000000000000000000000000000000000000000000000000000000
00000000000000000000000001111111
>DM_validata1139
VPPFQDCILSFLGFSDEEKTNMEEMTEMQGGKYLPLGDERCTHLVVEENIVKDLPFEPSKKL
YVVKQEWFWGSIQMDARAGETMYLYEKANTPE
000000000000000000000000000000000000000000000000000000000000
00000000000000000000000000011111
>DM_validata1140
MGSDKIHHHHHHLPLFKFAIDVQYRSNVRDPRGETIERVLREEKGLPVKKLRLGKSIHLEVE
AENKEKAYEIVKKACEELLVNPVVEEYEVREL
111111111000000000000000000000000000000000000000000000000000
00000000000000000000000000000000
>DM_validata1141

SNAENTDEDALMVTREDGSFLIDGTLPIEELREVLGAELPDGEENNYHTLAGMCISYFGRIP
HVGEYFDWAGWRIEIVDLDGARIDKLLLQRLN
11111100000000000000000000000000000011111100000000000000000000
00000000000000000000000000000000
>DM_validata1142
MRDYSELEIFEGNPLDKWNDIIFHASKKLSKKELERLLELLALLETFIEKEDLEEKFESFAK
ALRIDEELQQKIESRKTDIVIQSMANILSGNE
11111111111100000000000000000000000000000000000000000000000000
00000000000000000000000000000011
>DM_validata1143
MKLSELQSHIKEFDYAPEQSEHYFFKLIEEVGELSESIRKGKSGQPTLDELKGSVAEELYDV
LYYVCALANIHGVNLEKTHELKEVLNKVKYNR
11111111111100000000000000000000000000000000000000000000000000
00000000000000000000000000000111
>DM_validata1144
GLMDEGLPPEEVERIRAFLQERIRGRALEVHDLKTRRAGPRSFLEFHLVVRGDTPVEEAHRL
CDELERALAQAFPGLQATIHVEPEGERKRTNP
11110000000000000000000000000000000000000000000000000000000000
00000000000000000000000001111111
>DM_validata1145
MHHHHHHMSESSDISAMQPVNPKPFLKGLVNHRVGVKLKFNSTEYRGTLVSTDNYFNLQLNE
AEEFVAGVSHGTLGEIFIRCNNVLYIRELPN
11111111111111111111111110000000000000000000000000000000000000
0000000000000000000000000000000
>DM_validata1146
MNVTVSIPTILRPHTGGQKSVSASGDTLGAVISDLEANYSGISERLMDPSSPGKLHRFVNIY
VNDEDVRFSGGLATAIADGDSVTILPAVAGG
10000000000000000000000000000000000000000000000000000000000000
0000000000000000000000000000000
>DM_validata1147
MRILPISTIKGKLNEFVDAVSSTQDQITITKNGAPAAVLVGADEWESLQETLYWLAQPGIRE
SIAEADADIASGRTYGEDEIRAEFGVPRRPH
00000000000000000000000000000000000000000000000000000000000000
0000000000000000000000000000011
>DM_validata1149
MVKVTYDGVYVMSVKDDVPAADVLHAGDLITEIDGNAFKSSQEFIDYIHSKKVGDTVKINYK
HGDKNEQADIKLTAIDKKGTPGIGITLVDDH
10000000000000000000000000000000000000000000000000000000000000
0000000000000000000000000000000
>DM_validata1150
GAMALIEVEKPLYGVEVFVGETAHFEIELSEPDVHGQWKLKGQPLAASPDCEIIEDGKKHIL
ILHNCQLGMTGEVSFQAANTKSAANLKVKEL
11110000000000000000000000000000000000000000000000000000000000
0000000000000000000000000000000

>DM_validata1151
SNAEEELPFKVLGDGSYLFEGKTSLSDVRHYLDLPENAFGELGDEVDTLSGLFLEIKQELPH
VGDTAVYEPFRFQVTQMDKRRIIEIKIFPFE
111111000000000000000000000000000000000000000000000000000000
0000000000000000000000000000000
>DM_validata1152
MGHHHHHHSGQRKRGRQSKDEQLASDNELPVSAFQISEMSLSELQQVLKNESLSEYQRQLIR
KIRRRGKNKVAARTCRQRRTDRHDKMSHYI
111111111111110000000000000000000000000000000000000000000000
000000000000000000000000001111
>DM_validata1153
GKDYPFARFNKTVQVPVYSEQEYQLYLHDDAWTKAETDHLFDLSRRFDLRFVVIHDRYDHQQ
FKKRSVEDLKERYYHICAKLANVRAVPGTD
111111111111100000000000000000000000000000000000000000000000
000000000000000000000000011111
>DM_validata1154
MTVGKSSKMLQHIDYRMRCILQDGRIFIGTFKAFDKHMNLILCDCDEFRKIKPKNSKQAERE
EKRVLGLVLLRGENLVSMTVEGPPPKDTG
111111000000000000000000000000000000000000000000000000000000
00000000000000000000000001111
>DM_validata1155
LLGTHGDSGAAGTVFTTVEDLGSKILLTCSLDDSATEVTGHRWLKGGVVLKEDALPGQKTEF
KVDSDDQWGEYSCVFLPEPMGTANIQLHG
111111111110000000000000000000000100000000000000000000000000
00000000000000000000000000000
>DM_validata1156
SNADEDDSADNIHAVSSERWRIHAATEIEDINTFFGTEYSSEEADTIGGLVIQELGHLPVRG
EKVLIGGLQFTVARADNRRLHTLMATRVK
111111111000000000000000000000000000000000000000000000000000
00000000000000000000000000001
>DM_validata1157
AEEYGYIVTDQKPLSLAAGVKLLEILAEHVHMSSGSFINISVVGPALTFRIRHNEQNLSLAD
VTQQAGLVKSELEAQTGLQILQTGVGQRE
100000000000000000000000000000000000000000000000000000000000
00000000000000000000000000001
>DM_validata1158
MKKAVINGEQIRSISDLHQTLKKELALPEYYGENLDALWDALTGWVEYPLVLEWRQFEQSKQ
LTENGAESVLQVFREAKAEGADITIILS
000000000000000000000000000000000000000000000000000000011111
1111000000000000000000000000
>DM_validata1159
FVNKDQIAKDVKQFYDQALQQAVVDDDANNAKAVVKTFHETLDCCGSSTLTALTTSVLKNNL
CPSGSNIISNLFKEDCHQKIDDLFSGKH
000000000000000000000000111100000000000000000000000000000000

0000000000000000000000000000
>DM_validata1160
ASGPRALSRNQPQYPARAQALRIEGQVKVKFDVTPDGRVDNVQILSAKPANMFEREVKNAMR
RWRYEPGKPGSGIVVNILFKINGTTEIQ
11000000000000000000000000000000000000000000000000000000000000
0000000000000000000000000000
>DM_validata1161
SNAMDGQQLNRLLLEWIGAWDPFGLGKDAYDVEAASVLQAVYETEDARTLAARIQSIYEFAF
DEPIPFPHCLKLARRLLELKQAASCPLP
11111000000000000000000000000000000000000000000000000000000000
0000000000000000000000001111
>DM_validata1162
SSVQTAATSWGTVPSIRVYTANNGKITERCWDGKGWYTGAFNEPGDNVSVTSWLVGSAIHIR
VYASTGTTTTEWCWDGNGWTKGAYTSTN
00000000000000000000000000000000000000000000000000000000000000
0000000000000000000000000011
>DM_validata1163
MTEKLNEIVVRKTKNVEDHVLDVIVLFNQGIDEVILKGTGREISKAVDVYNSLKDRLGDGVQ
LVNVQTGSEVRDRRRISYILLRLKRVY
11100000000000000000000000000000000000000000000000000000000000
000000000000000000000000000
>DM_validata1164
ADAQKAADNKKPVNSWTCEDFLAVDESFQPTAVGFAEALNNKDKPEDAVLDVQGIATVTPAI
VQACTQDKQANFKDKVKGEWDKIKKDM
11111111100000000000000000000000000000000000000000000000000000
000000000000000000000001111
>DM_validata1165
MHHHHHHSSGRENLYFQGSARLRALRQRQLDRAAAAVEPDVVVKRQEALAAARLKMQEELNA
QVEKHKEKLKQLEEEKRRQKIEMWDSM
11111111111111111000000000000000000000000000000000000000000000
000000000000000000000000000
>DM_validata1166
KLQICVEPTSQKLMPGSTLVLQCVAVGSPIPHYQWFKNELPLTHETKKLYMVPYVDLEHQGT
YWCHVYNDRDSQDSKKVEIIIDELNNL
00000000000000000000000000000000000000000000000000000000000000
000000000000000000000011111
>DM_validata1167
SMSNYSVSLVGPAPWGFRLQGGKDFNMPLTISSLKDGGKAAQANVRIGDVVLSIDGINAQGM
THLEAQNKIKGCTGSLNMTLQRESDL
11000000000000000000000000000000000000000000000000000000000000
00000000000000000000000000
>DM_validata1168
IEPVLENVQPNSAASKAGLQAGDRIVKVDGQPLTQWVTFVMLRDNPGKSLALEIERQGSPL
SLTLIPESKPGNGKAIGFVGIEPKVI

110000000000000000000000000000000000000000000000000000000000
00000000000000000000000000
>DM_validata1169
MIDVSSQRVNVQRPLDALGNSLNSPVIIKLKGDREFRGVLKSFDLHMNLVLNDAEELEDGEV
TRRLGTVLIRGDNIVYISRGKLAAA
111111111000000000000000000000000000000000000000000000000000
0000000000000000000111111
>DM_validata1170
ARRKRRNFNKQATEILNEYFYSHLSNPYPSEEAKEELAKKCGITVSQVSNWFGNKRIRYKKN
IGKFQEEANIYAAKTAVTATNVSAH
000000000000000000000000000000000000000000000000000000000000
0000000000000111111111111
>DM_validata1171
AERSLLTGEEGWRTYKATGPRLSLPRLVALLKGQGLEVGKVAEAEGGFYVDLRPEARPEVAG
LRLEPARRVEGLLEIPSRTRRPARA
000000000000000000000000000000000000000000000000000000000000
0000000000000111111111111
>DM_validata1172
MDGMGTLTRYLEEAMARARYELIADEEPYYGEIPDLPGVWATGKSLKECEANLQAALEDWLL
FLLSRGETPPPLGEVRIELPHGEAA
111000000000000000000000000000000000000000000000000000000000
0000000000000000011111111
>DM_validata1174
GPHMSIVEVKSKFDAEFRRFALPRASVSGFQEFSRLLRAVHQIPGLDVLLGYTDAHGDLLPL
TNDDSLHRALASGPPPLRLLVQKR
111100000000000000000000000000000000000000000000000000000000
000000000000000000000000
>DM_validata1175
GHMEKRFYILTIVVEDREKAYRQVNELLHNFSEDILLRVGYPVREENMAIIFLVLKTDNDTI
GALSGKLGQISGVRVKTVPLKRGS
111000000000000000000000000000000000000000000000000000000000
000000000000000000000111
>DM_validata1177
PRRGAVAHANSIVQQLVSEGADISHTRNMLRNAMNGDAVAFSRVEQNIFRQHFPNMPMHGIS
RDSELAIELRGALRRAVHQQAAS
111000000000000000000000000000000000000000000000000000000000
00000000000000000001111
>DM_validata1178
GTQGFAVLSYVYEHEKRDLASRIVSTQHHHHDLSVATLHVHINHDDCLEIAVLKGDMGDVQH
FADDVIAQRGVRHGHLQCLPKED
100000000000000111000000000000000000000000000000000000000000
00000000000000000000001
>DM_validata1179
MVLATLPDTTWTPSVGLVVILCNLFAIALGRYAIQSRGKGPGLPIALPALFEGFGLPELLAT

TSFGHLLAAGVVSGLQYAGAL
111111111111111111000000000000000000000000111111111100000000
000000000000000111111
>DM_validata1180
GMEQQHPTIHTLKIETEFFKAVKERRKTFEIRKNDRNFQVGDILILEEYMNGMYLDDECEAE
VIYITDYAQREGYVVLGIELH
111100000000000000000000000000000000000000000000000000000000
000000000000000000000
>DM_validata1181
GHMFEKIRKILADIEDSQNEIEMLLKLANLSLGDFIEIKRGSMDMPKGVNEAFFTQLSEEVE
RLKELINALNKIKKGLLVFGS
110000000000000000000000000000000000000000000000000000000000
000000000000000000000
>DM_validata1182
GSHMANKRNEALRIESALLNKIAMLGTEKTAEAVGVDKSQISRWKRDWIPKFSMLLAVLEWG
VVDDDMARLARQVAAILTNKK
111111100000000000000000000000000000000000000000000000000000
000000000000000000111
>DM_validata1183
SGHHEDSTDEASESSKPCCDRCECTKSIPPQCRCSDVRLNSCHSACKSCACTFSIPAQCFCG
DINDFCYKPCKSSHSDDDDWN
111111111111111100000000000000000000000000000000000000000000
000000000011111111111
>DM_validata1185
MSKGHSLQDPYLNTLRKERVPVSIYLVNGIKLQGQIESFDQFVILLKNTVSQMVYKHAISTV
VPSRPVRLPSGDQPAEPGNA
111100000000000000000000000000000000000000000000000000000000
00000001111111111111
>DM_validata1186
MEKTRANSHLRSQPIDWTIEEVIQYIESNDNSLAVHGDLFRKHEIDGKALLRLNSERMMKYM
GLKLGPALKICNLVNKVNGR
111111111111100000000000000000000000000000000000000000000000
00000000000000000011
>DM_validata1187
EDPEVLAKNKGCVACHAIDTKMVGPAYKDVAAKFAGQAGAEAELAQRIKNGSQGVWGPIPMP
PNAVSDDEAQTLAKWVLSQK
100000000000000000000000000000000000000000000000000000000000
00000000000000000000
>DM_validata1188
EDISDSESEPGIALKRKQRRSRTTFSASQLDELERAFERTQYPDIYTREELAQRTNLTEARI
QVWFQNRRARLRKQHTSVS
111111111111111110000000000000000000000000000000000000000000
0000000000000111111
>DM_validata1189

RKRNRIPLSCTICRKRKVKCDKLRPHCQQCTKTGVAHLCHYMEQTWAEEAEKELLKDNELKK
LRERVKSLEKTLSKVHSSP
111100000000000000000000000000000000000000000000000000000000
0000000000001111111
>DM_validata1190
SELDAKLNKLGVDRIAISPYKQWTRGYMEPGNIGNGYVTGLKVDAGVRDKSDNNVLDGIVSY
DRAETKNAYIGQINMTTAS
000000000000000000000000000000000000000000000111111111111111
1000000000000000000
>DM_validata1191
PHLIKVTVKTPKDKEDFSVTDTCTIQQLKEEISQRFKAHPDQLVLIFAGKILKDPDSLAQCG
VRDGLTVHLVIKRQHRAMG
000000000000000000000000000000000000000000000000000000000000
0000000000000000001
>DM_validata1192
LDLHGLHVDEALEHLMRVLEKKTEEFKQNGGKPYLSVITGRGNHSQGGVARIKPAVIKYLIS
HSFRFSEIKPGCLKVMLK
000000000000000000000000000000000000000111111111000000000000
000000000000000001
>DM_validata1193
SQRGVLEDEQMYQRCCNLFEKFFPSSSYRRPVGISSMVEAMVSRARIDARIDFESGRIKKEE
FTEIMKICSTIEELRRQK
111111100000000000000000000000000000000000000000000000000000
000000000000000000
>DM_validata1194
GSSALLDTDEVGSELDDSDDDYLISEGEEDGPDENLMLCLYDKVTRTKARWKCSLKDGVVTI
NRNDYTFQKAQVEAEWV
111111111111111111110000111111110000000000000000000000000000
00000000000000000
>DM_validata1195
MGGYVNIKTFTHPAGEGKEVKGMEVSVPFEIYSNEHRIADAHYQTFPSEKAAYTTVVTDAAD
WRTKNAAMFTPTPVSG
100000000000000000000000000000000000000000000000000000000000
0000000000000011
>DM_validata1196
QAVGLPHGFCIQCNRKTWSNCSIGHRCLPYHMTCYTLYKPDENGEMKWAVKGCARMCPTAKS
GERVKCCTGASCNSD
111000000000000000000000000000000000000000000000000000000000
000000000000000
>DM_validata1197
MKCPVCHQGEMVSGIKDIPYTFRGRKTVLKGIHGLYCVHCEESIMNKEESDAFMAQVKAFRA
SVNAETVAPEFIVK
000000000000000000001111110000000000000000000000000000000000
00011111111111

>DM_validata1198
MAHHHHHHMASADVVSTWVCPICMVSNETQGEFTKDTLPTPICINCGVPADYELTKSSINCS
NAIDPNANPRNQFG
11111111111111000000000000000000000000000000000000000000000001
11111111111111
>DM_validata1199
MQIFVKTLTGKTITLEVEPSDTIENVKAKIQDKEGIPPDQQRLIFAGKQLEDGRTLSDYNIQ
KESTLHLVLRLRGG
00000000000000000000000000000000000000000000000000000000000000
00000000001111
>DM_validata1200
KLITLLLRSSKSEDLRLSIPVDFTVKDLIKRYCTEVKISFHERIRLEFEGEWLDPNDQVQST
ELEDEDQVSVVLD
00000000000000000000000000000000000000000000000000000000000000
0000000000001
>DM_validata1201
MDLLSCTVNDAEIFSLVKKEVLSLNTNDYTTAISLSNRLKINKKKINQQLYKLQKEDTVKMV
PSNPPKWFKNYNC
11111000000000000000000000000000000000000000000000000000000000
0000000000000
>DM_validata1202
GSKEESEKPRGFARGLEPERIIGATDSSGELMFLMKWKNSDEADLVPAKEANVKCPQVVISF
YEERLTWHSYPS
11111111000000000000000000000000000000000000000000000000000000
000000000000
>DM_validata1203
GHMTSVFDRDDIQFQVVVNHEEQYSIWPEYKEIPQGWRAAGKSGLKKDCLAYIEEVWTDMRP
LSLRQHMDKAAG
11111111100000000000000000000000000000000000000000000000000000
000000001111
>DM_validata1204
QDPLSGLTDQERTLLGLLSEGLTNKQIADRMFLAEKTVKNYVSRLLAKLGMERRTQAAVFAT
ELKRSRPPGDGP
11111110000000000000000000000000000000000000000000000000111111
111111111111
>DM_validata1206
HDKVVVLEAKNGNVTFDHKKHAGVKGECKACHETEAGGKIAGMGKDWAHKTCTGCHKEMGKG
PTKCGECHKK
11100000000000000000000000000000000000000000000000000000000000
0000000001
>DM_validata1207
FNCNKREGPCSQRSLCECDPNLQLGRHSDQLWHYNLRTNRCERGGYRDNCNSHSSSGACVMA
CERIHHHHHH
00000000000000000000000000000000000000000000000000000000000000

0000111111
>DM_validata1208
MDPNTVSSFQVDCFLWHVRKRLADQELGDAPFLDRLRRDQKSLRGRGNTLGLDIETATRAGK
QIVERILEEE
100000000000000000000000000000000000000000000000000000000000000
0000000000
>DM_validata1209
GSTSDVANLANEKEELNNKLKEAQEQLSRLKDEEISAAAIKAQFEKQLLTERTLKTQAVNKL
AEIMNRKEP
000000000000000000000000000000000000000000000000000000000000000
000000111
>DM_validata1210
MKPEELVRHFGDVEKAAVGVGVTPGAVYQWLQAGEIPPLRQSDIEVRTAYKLKSDFTSQRMG
KEGHNSGTK
000000000000000000000000000000000000000000000000000000000000000
001111111
>DM_validata1211
ASVNAETVAPEFIVKVRKKLSLTQKEASEIFGGGVNAFSRYEKGNAQPHPSTIKLLRVLDKH
PELLNEIR
111100000000000000000000000000000000000000000000000000000000000
00000000
>DM_validata1212
ESAFNQTEFNKLLLECVVKTQSSVAKILGIESLSPHVSGNSKFEYANMVEDIREKVSSEMER
FFPKNDDE
110000000000000000000000000000000000000000000000000000000000000
00000000
>DM_validata1213
MGKVYKKVELVGTSEEGLEAAIQAALARARKTLRHLDWFEVKEIRGTIGEAGVKEYQVVLEV
GFALEET
100000000000000000000000000000000000000000000000000000000000000
0000011
>DM_validata1214
MPKHEFSVDMTCGGCAEAVSRVLNKLGGVKYDIDLPNKKVCIESEHSMDTLLATLKKTGKTV
SYLGLE
100000000000000000000000000000000000000000000000000000000000000
000001
>DM_validata1215
MNTQLMGERIRARRKKLKIRQAALGKMVGVSNVAISQWERSETEPNGENLLALSKALQCSPD
YLLKGD
110000000000000000000000000000000000000000000000000000000000000
000000
>DM_validata1217
GTTPAGPVWTAVFDYEAAGDEELTLRRGDRVQVLSQDCAVSGDEGWWTGQLPSGRVGVFPSN
YVAP

111100000000000000000000000000000000111111000000000000000000
0000
>DM_validata1218
GSPLGQDVFLDYCQKLLEKFRYPWELMPLMYVILKDADANIEEASRRIEEGQYVVNEYSRQH
NLN
111110000000000000000000000000000000000000000000000000000000
001
>DM_validata1220
GSSGSSGGYFDAHALAMDYRSLGFRECLAEVARYLSIIEGLDASDPLRVRLVSHLNNYASQR
EA
111110000000000000000000000000000000000000000000000000000000
11
>DM_validata1221
NLLEEESAVLGQAVTNLMLSGDNVNNKNIILSLIHSLETTSDILKADVIRKTLEIVLRYTAD
DM
000000000000000000000000000000000000000000000000000000000000
11
>DM_validata1222
MDNRQFLSLTGVSKVQSFDPKEILLETIQGVLSIKGEKLGIKHLDLKAGQVEVEGLIDALVY
P
110000000000000000000000000000000000000000000000000000000000
0
>DM_validata1223
GSHGGIEGVKGAASGVVGELARARLALDERGQKLSDLEERTAAMMSSADSFSKHAHEMMLKY
K
111100000000000000000000000000000000000000000000000000000000
1
>DM_validata1224
MAEPGDLRHDLNQQERATLSSNVQRFFMIGHGSLTADAGGLTYTVSWVPTKQIQRKVAPSAG
P
111000000000000000000000000000000000000000000000000000001111
1
>DM_validata1225
MVDYIVEYDYDAVHDDELTIRVGEIIRNVKKLQEEGWLEGELNGRRGMFPDNFVKEIKRETE
100000000000000000000000000000000000000000000000000000000111
>DM_validata1227
IKKEQKLIQAQNLVREFEKTHTVSAHRKAQKAVNLVSFEYKVKKMVLQERIDNVLKQGLVR
10000000000000000000000000000000000000000000000000000000000
>DM_validata1228
EQTHRAIFRFVPRHEDELELEVDDPLLVELQAEDYWYEAYNMRTGARGVFPAYYAIEVTK
0000000000000000000000000000000000000000000000000000000011
>DM_validata1229
SEKACRHCHYITSEDRCPVCGSRDLSEEWFDLVIIVDVENSEIAKKIGAKVPGKYAIRVR
1000000000000000000000000000000000000000000000000000000000
