## Supplementary Information S2 for "TransDFL: Identification of Disordered Flexible Linkers in Proteins by Transfer Learning"

**Supplementary Information S2**. The TE64 independent test set.
Line 1: >protein ID
Line 2: protein sequence
Line 3: annotations of disordered flexible linker (DFL) regions, where '1' denotes a DFL, '0' denotes a disordered residue annotated with functions other than DFL, 'x' denotes other residues.

```
>DP02359
MDAGFFRGTSAEQDNRFSNKQKKLLKQLKFAECLEKKVDMSKVNLEVIKPWITKRVTEILGF
EDDVVIEFIFNQLEVKNPDSKMMQINLTGFLNGKNAREFMGELWPLLLSAQENIAGIPSAFL
ELKKEEIKQRQIEQEKLASMKKQDEDKDKRDKEEKESSREKRERSRSPRRRKSRSPSPRRRS
SPVRRERKRSHSRSPRHRTKSRSPSPAPEKKEKTPELPEPSVKVKEPSVQEATSTSDILKVP
KPEPIPEPKEPSPEKNSKKEKEKEKTRPRSRSRSKSRSRTRSRSPSHTRPRRRHRSRSRSYS
PRRRPSPRRRPSPRRRTPPRRMPPPPRHRRSRSPVRRRRRSSASLSGSSSSSSSSRSRSPPK
KPPKRTSSPPRKTRRLSPSASPPRRRHRPSPPATPPPKTRHSPTPQQSNRTRKSRVSVSPGR
TSGKVTKHKGTEKRESPSPAPKPRKVELSESEEDKGGKMAAADSVQQRRQYRRQNQQSSSDS
GSSSSSEDERPKRSHVKNGEVGRRRRHSPSRSASPSPRKRQKETSPRGRRRRSPSPPPTRRR
RSPSPAPPPRRRRTPTPPPRRRTPSPPPRRRSPSPRRYSPPIQRRYSPSPPPKRRTASPPPP
PKRRASPSPPPKRRVSHSPPPKQRSSPVTKRRSPSLSSKHRKGSSPSRSTREARSPQPNKRH
SPSPRPRAPQTSSSPPPVRRGASSSPQRRQSPSPSTRPIRRVSRTPEPKKIKKAASPSPQSV
RRVSSSRSVSGSPEPAAKKPPAPPSPVQSQSPSTNWSPAVPVKKAKSPTPSPSPPRNSDQEG
GGKKKKKKKDKKHKKDKKHKKHKKHKKEKAVAAAAAAAVTPAAIAAATTTLAQEEPVAAPEP
KKETESEAEDNLDDLEKHLREKALRSMRKAQVSPQS
000000000000000000000000000000000000000000000000000000000000
000000000000000000000000000000000000000000000000000000000000
000000000000000000000000000000000000000000000000000000000000
000000000000000000000000000000000000000000000000000000000000
000000000000000000000000000000000000000000000000000000000000
000000000000000000000000000000000000000000000000000000000000
000000000000000000000000000000000000000000000000000000000000
000000000000000000000000000000000000000000000000000000000000
000000000000000000000000000000000000000000000000000000000000
000000000000000000000000000000000000000000000000000000000000
000000000000000000000000000000000000000000000000000000000000
000000000000000000000000000000000000000000000000000000000000
000000000000000000000000000000000000000000000000000000000000
000000000000000000000000000000000000000000000000000000000000
000000000000000000000000000000000000
>DP02371
MADKMDMSLDDIIKLNRSQRGGRGGGRGRGRAGSQGGRGGAVQAAARVNRGGGPMRNRPAIA
RGAAGGGRNRPAPYSRPKQLPDKWQHDLFDSGFGGGAGVETGGKLLVSNLDFGVSDADIQEL
FAEFGTLKKAAVHYDRSGRSLGTADVHFERKADALKAMKQYNGVPLDGRPMNIQLVTSQIDT
QRRPAQSINRGGMTRNRGSGGFGGGGTRRGTRGGSRGRGRGTGRNSKQQLSAEELDAQLDAY
NARMDTS
```

xxxxxxxxxxxxxxxxxxxxxxxxxxxxxxxxxxxxxxxxxxxxxxxxxxxxxxxxxxxx
xxxxxxxxxxxxxx0000000000000000000000000000000000000000000000
00000000000000000000000000000000000000000000000000000000xxxx
xxxxxxxxxxxxxxxxxxxxxxxxxxxxxxxxxxxxxxxxxxxxxxxxxxxxxxxxxxxx
xxxxxxx
>DP02373
MPCRREEEEEAGEEAEGEEEEEDSFLLLQQSVALGSSGEVDRLVAQIGETLQLDAAQHSPAS
PCGPPGAPLRAPGPLAAAVPADKARSPAVPLLLPPALAETVGPAPPGVLRCALGDRGRVRGR
AAPYCVAELATGPSALSPLPPQADLDGPPGAGKQGIPQPLSGPCRRGWLRGAAASRRLQQRR
GSQPETRTGDDDPHRLLQQLVLSGNLIKEAVRRLHSRRLQLRAKLPQRPLLGPLSAPVHEPP
SPRSPRAACSDPGASGRAQLRTGDGVLVPGS
000000000000000000000000000000000000000000000000000000000000
000000000000000000000000000000000000000000000000000000000000
000000000000000000000000000000000000000000000000000000000000
000000000000000000000000000000000000000000000000000000000000
0000000000000000000000000000000
>DP02414
MDYNRMNSFLEYPLCNRGPSAYSAHSAPTSFPPSSAQAVDSYASEGRYGGGLSSPAFQQNSG
YPAQQPPSTLGVPFPSSAPSGYAPAACSPSYGPSQYYPLGQSEGDGGYFHPSSYGAQLGGLS
DGYGAGGAGPGPYPPQHPPYGNEQTASFAPAYADLLSEDKETPCPSEPNTPTARTFDWMKVK
RNPPKTAKVSEPGLGSPSGLRTNFTTRQLTELEKEFHFNKYLSRARRVEIAATLELNETQVK
IWFQNRRMKQKKREREEGRVPPAPPGCPKEAAGDASDQSTCTSPEASPSSVTS
xxxxxxxxxxxxxxxxxxxxxxxxxxxxxxxxxxxxxxxxxxxxxxxxxxxxxxxxxxxx
xxxxxxxxxxxxxxxxxxxxxxxxxxxxxxxxxxxxxxxxxxxxxxxxxxxxxxxxxxxx
xxxxxxxxxxxxxxxxxxxxxxxxxxxxxxxxxxxxxxxxxxxxxxxxxxxxxxxxxxxx
11111111111111111111xxxxxxxxxxxxxxxxxxxxxxxxxxxxxxxxxxxxxxxx
xxxxxxxxxxxxxxxxxxxxxxxxxxxxxxxxxxxxxxxxxxxxxxxxxxxxx
>DP02511
MVELSKKVTGKLDKTTPGIQIWRIENMEMVPVPTKSYGNFYEGDCYVLLSTRKTGSGFSYNI
HYWLGKNSSQDEQGAAAIYTTQMDEYLGSVAVQHREVQGHESETFRAYFKQGLIYKQGGVAS
GMKHVETNTYNVQRLLHVKGKKNVVAAEVEMSWKSFNLGDVFLLDLGQLIIQWNGPESNRAE
RLRAMTLAKDIRDRERAGRAKVGVVEGENEAASPELMQALTHVLGEKKNIKAATPDEQVHQA
LNSALKLYHVSDASGNLVIQEVAIRPLTQDMLQHEDCYILDQAGLKIFVWKGKNANKEEKQQ
AMSRALGFIKAKNYLASTSVETENDGSESAVFRQLFQKWTVPNQTSGLGKTHTVGKVAKVEQ
VKFDATTMHVKPEVAAQQKMVDDGSGEAEVWRVENQELVPVEKRWLGHFYGGDCYLVLYTYY
VGPKVNRIIYIWQGRHASTDELAASAYQAVFLDQKYNNEPVQVRVTMGKEPAHLMAIFKGKM
VVYENGSSRAGGTEPASSTRLFHVHGTNEYNTKAFEVPVRAASLNSNDVFVLKTPSSCYLWY
GKGCSGDEREMGKMVADIISKTEKPVVAEGQEPPEFWVALGGKTSYANSKRLQEENPSVPPR
LFECSNKTGRFLATEIVDFTQDDLDENDVYLLDTWDQIFFWIGKGANESEKEAAAETAQEYL
RSHPGSRDLDTPIIVVKQGFEPPTFTGWFMAWDPLCWSDRKSYDELKAELGDNASIGQLVSG
LTSKNEVFTATTTLVPTKLETFPLDVLVNTAAEDLPRGVDPSRKENHLSDEDFKAVFGMTRS
AFANLPLWKQQNLKKEKGLF
xxxxxxxxxxxxxxxxxxxxxxxxxxxxxxxxxxxxxxxxxxxxxxxxxxxxxxxxxxxx
xxxxxxxxxxxxxxxxxxxxxxxxxxxxxxxxxxxxxxxxxxxxxxxxxxxxxxxxxxxx

xxxxxxxxxxxxxxxxxxxxxxxxxxxxxxxxxxxxxxxxxxxxxxxxxxxxxxxxxxxx
xxxxxxxxxxxxxxxxxxxxxxxxxxxxxxxxxxxxxxxxxxxxxxxxxxxxxxxxxxxx
xxxxxxxxxxxxxxxxxxxxxxxxxxxxxxxxxxxxxxxxxxxxxxxxxxxxxxxxxxxx
xxxxxxxxxxxxxxxxxxxxxxxxxxxxxxxxxxxxxxxxxxxxxxxxxxxxxxxxxxxx
xxxxxxxxxxxxxxxxxxxxxxxxxxxxxxxxxxxxxxxxxxxxxxxxxxxxxxxxxxxx
xxxxxxxxxxxxxxxxxxxxxxxxxxxxxxxxxxxxxxxxxxxxxxxxxxxxxxxxxxxx
xxxxxxxxxxxxxxxxxxxxxxxxxxxxxxxxxxxxxxxxxxxxxxxxxxxxxxxxxxxx
xxxxxxxxxxxxxxxxxxxxxxxxxxxxxxxxxxxxxxxxxxxxxxxxxxxxxxxxxxxx
xxxxxxxxxxxxxxxxxxxxxxxxxxxxxxxxxxxxxxxxxxxxxxxxxxxxxxxxxxxx
xxxxxxxxxxxxxxxxxxxxxxxxxxxxxxxxxxxxxx1111111111111111111111
1111111111111111111xxxxxxxxxxxxxxxxxxxxxxxxxxxxxxxxxxxxxxxxx
xxxxxxxxxxxxxxxxxxxx
>DP02521
MDYQEYQQFLARINTARDACVAKDIDVDLLMARHDYFGRELCKSLNIEYRNDVPFIDIILDI
RPEVDPLTIDAPHITPDNYLYINNVLYIIDYKVSVSNESSVITYDKYYELTRDISDRLSIPI
EIVIIRIDPVSRDLHINSDRFKELYPTIVVDINFNQFFDLKQLLYEKFGDDEEFLLKVAHGD
FTLTAPWCKTGCPEFWKHPIYKEFKMSMPVPERRLFEESVKFNAYESERWNTNLVKIREYTK
KDYSEHISKSAKNIFLASGFYKQPNKNEISEGWTLMVERVQDQREISKSLHDQKPSIHFIWG
AHNPGNSNNATFKLILLSKSLQSIKGISTYTEAFKSLGKMMDIGDKAIEYEEFCMSLKSKAR
SSWKQIMNKKLEPKQINNALVLWEQQFMINNDLIDKSEKLKLFKNFCGIGKHKQFKNKMLED
LEVSKPKILDFDDANMYLASLTMMEQSKKILSKSNGLKPDNFILNEFGSRIKDANKETYDNM
HKIFETGYWQCISDFSTLMKNILSVSQYNRHNTFRIAMCANNNVFAIVFPSADIKTKKATVV
YSIIVLHKEEENIFNPGCLHGTFKCMNGYISISRAIRLDKERCQRIVSSPGLFLTTCLLFKH
DNPTLVMSDIMNFSIYTSLSITKSVLSLTEPARYMIMNSLAISSNVKDYIAEKFSPYTKTLF
SVYMTRLIKNACFDAYDQRQRVQLRDIYLSDYDITQKGIKDNRELTSIWFPGSVTLKEYLTQ
IYLPFYFNAKGLHEKHHVMVDLAKTILEIECEQRENIKEIWSTNCTKQTVNLKILIHSLCKN
LLADTSRHNHLRNRIENRNNFRRSITTISTFTSSKSCLKIGDFRKEKELQSVKQKKILEVQS
RKMRLANPMFVTDEQVCLEVGHCNYEMLRNAMPNYTDYISTKVFDRLYELLDKKVLTDKPVI
EQIMDMMIDHKKFYFTFFNKGQKTSKDREIFVGEYEAKMCMYAVERIAKERCKLNPDEMISE
PGDGKLKVLEQKSEQEIRFLVETTRQKNREIDEAIEALATEGYESNLGKIEKLSLGKAKGLK
MEINADMSKWSAQDVFYKYFWLIALDPILYPQEKERILYFMCNYMDKELILPDELLFNLLDQ
KVAYQNDIIATMTNQLNSNTVLIKRNWLQGNFNYTSSYVHSCAMSVYKEILKEAITLLDGSI
LVNSLVHSDDNQTSITIVQDKMENDKIIDFAMKEFERACLTFGCQANMKKTYVTNCIKEFVS
LFNLYGEPFSIYGRFLLTSVGDCAYIGPYEDLASRISSAQTAIKHGCPPSLAWVSIAISHWM
TSLTYNMLPGQSNDPIDYFPAENRKDIPIELNGVLDAPLSMISTVGLESGNLYFLIKLLSKY
TPVMQKRESVVNQIAEVKNWKVEDLTDNEIFRLKILRYLVLDAEMDPSDIMGETSDMRGRSI
LTPRKFTTAGSLRKLYSFSKYQDRLSSPGGMVELFTYLLEKPELLVTKGEDMKDYMESVIFR
YNSKRFKESLSIQNPAQLFIEQILFSHKPVIDFSGIRDKYINLHDSRALEKEPDILGKVTFT
EAYRLLMRDLSSLELTNDDIQVIYSYIILNDPMMITIANTHILSIYGSPQRRMGMSCSTMPE
FRNLKLIHHSPALVLRAYSKNNPDIQGADPTEMARDLVHLKEFVENTNLEEKMKVRIAMNEA
EKGQRDIVFELKEMTRFYQVCYEYVKSTEHKIKVFILPAKSYTTTDFCSLMQGNLIKDKEWY
TVHYLKQILSGGHKAIMQHNATSEQNIAFECFKLITHFADSFIDSLSRSAFLQLIIDEFSYK
DVKVSKLYDIIKNGYNRTDFIPLLFRTGDLRQADLDKYDAMKSHERVTWNDWQTSRHLDMGS
INLTITGYNRSITIIGEDNKLTYAELCLTRKTPENITISGRKLLGSRHGLKFENMSKIQTYP

GNYYITYRKKDRHQFVYQIHSHESITRRNEEHMAIRTRIYNEITPVCVVNVAEVDGDQRILI
RSLDYLNNDIFSLSRIKVGLDEFATIKKAHFSKMVSFEGPPIKTGLLDLTELMKSQDLLNLN
YDNIRNSNLISFSKLICCEGSDNINDGLEFLSDDPMNFTEGEAIHSTPIFNIYYSKRGERHM
TYRNAIKLLIERETKIFEEAFTFSENGFISPENLGCLEAVVSLIKLLKTNEWSTVIDKCIHI
CLIKNGMDHMYHSFDVPKCFMGNPITRDINWVMFREFINSLPGTDIPPWNVMTENFKKKCIA
LINSKFETQRDFSEFTKLMKKEGGRSNIEFD
xxxxxxxxxxxxxxxxxxxxxxxxxxxxxxxxxxxxxxxxxxxxxxxxxxxxxxxxxxxxxx
xxxxxxxxxxxxxxxxxxxxxxxxxxxxxxxxxxxxxxxxxxxxxxxxxxxxxxxxxxxxxx
xxxxxxxxxxxxxxxxxxxxxxxxxxxxxxxxxxxxxxxxxxxxxxxxxxxxxxxxxxxx11
11111111111111111111111111111111111111111111111111111111111111
1111111111111111111111xxxxxxxxxxxxxxxxxxxxxxxxxxxxxxxxxxxxxxxx
xxxxxxxxxxxxxxxxxxxxxxxxxxxxxxxxxxxxxxxxxxxxxxxxxxxxxxxxxxxxxx
xxxxxxxxxxxxxxxxxxxxxxxxxxxxxxxxxxxxxxxxxxxxxxxxxxxxxxxxxxxxxx
xxxxxxxxxxxxxxxxxxxxxxxxxxxxxxxxxxxxxxxxxxxxxxxxxxxxxxxxxxxxxx
xxxxxxxxxxxxxxxxxxxxxxxxxxxxxxxxxxxxxxxxxxxxxxxxxxxxxxxxxxxxxx
xxxxxxxxxxxxxxxxxxxxxxxxxxxxxxxxxxxxxxxxxxxxxxxxxxxxxxxxxxxxxx
xxxxxxxxxxxxxxxxxxxxxxxxxxxxxxxxxxxxxxxxxxxxxxxxxxxxxxxxxxxxxx
xxxxxxxxxxxxxxxxxxxxxxxxx0000000000xxxxxxxxxxxxxxxxxxxxxxxxxxx
xxxxxxxxxxxxxxxxxxxxxxxxxxxxxxxxxxxxxxxxxxxxxxxxxxxxxxxxxxxxxx
xxxxxxxxxxxxxxxxxxxxxxxxxxxxxxxxxxxxxxxxxxxxxxxxxxxxxxxxxxxxxx
xxxxxxx0000000000000000xxxxxxxxxxxxxxxxxxxxxxxxxxxxxxxxxxxxxxx
xxxxxxxxxxxxxxxxxxx0000000000xxxxxxxxxxxxxxxxxxxxxxxxxxxxxxxxx
xxxxxxxxxxxxxxxxxxxxxxxxxxxxxxxxxxxxxxxxxxxxxxxxxxxxxxxxxxxxxx
xxxxxxxxxxxxxxxxxxxxxxxxxxxxxxxxxxxxxxxxxxxxxxxxxxxxxxxxxxxxxx
xxxxxxxxxxxxxxxxxxxxxxxxxxxxxxxxxxxxxxxxxxxxxxxxxxxxxxxxxxxxxx
xxxxxxxxxxxxxxxxxxxxxxxxxxxxxxxxxxxxxxxxxxxxxxxxxxxxxxxxxxxxxx
xxxxxxxxxxxxxxxxxxxxxxxxxxxxxxxxxxxxxxxxxxxxxxxxxxxxxxxxxxxxxx
xxxxxxxxxxxxxxxxxxxxxxxxxxxxxxxxxxxxxxxxxxxxxxxxxxxxxxxxxxxxxx
xxxxxxxxxxxxxxxxxxxxxxxxxxxxxxxxxxxxxxxxxxx00000000000000000xx
xxxxxxxxxxxxxxxxxxxxxxxxxxxxxxxxxxxxxxxxxxxxxxxxxxxxxxxxxxxxxx
xxxxxxxxxxxxxxxxxxxxxxxxxxxxxxxxxxxxxxxxxxx000000000000xxxxxxx
xxxxxxxxxxxxxxxxxxxxxxxxxxxxxxxxxxxxxxxxxxxxxxxxxxxxxxxxxxxxxx
xxxxxxxxxxxxxxxxxxxxxxxxxxxxxxxxxxxxxxxxxxxxxxxxxxxxxxxxxxxxxx
xxxxxxxxxxxxxxxxxxxxxxxxxxxxxxxxxxxxxxxxxxxxxxxxxxxxxxxxxxxxxx
xxxxxxxxxxxxxx000000000000000000000000000000000000000000000000
00000000000000000000000000000000000000000000000000000000000000
00000000000000000000000000000000000000000000000000000000000000
00000000000000000000000000000000000000000000000000000000000000
00000000000000000000000000000000000000000000000000000000000000
00000000000000000000000000000000000000000000000000000000000000
00000000000000000000000000000000000000000000000000000000000000
00000000000000000000000000000000000000000000000000000000000000
0000000000000000000000000000000
>DP02523

MPAMVEKGPEVSGKRRGRNNAAASASAAAASAAASAACASPAATAASGAAASSASAAAASAA
AAPNNGQNKSLAAAAPNGNSSSNSWEEGSSGSSSDEEHGGGGMRVGPQYQAVVPDFDPAKLA
RRSQERDNLGMLVWSPNQNLSEAKLDEYIAIAKEKHGYNMEQALGMLFWHKHNIEKSLADLP
NFTPFPDEWTVEDKVLFEQAFSFHGKTFHRIQQMLPDKSIASLVKFYYSWKKTRTKTSVMDR
HARKQKREREESEDELEEANGNNPIDIEVDQNKESKKEVPPTETVPQVKKEKHSTQAKNRAK
RKPPKGMFLSQEDVEAVSANATAATTVLRQLDMELVSVKRQIQNIKQTNSALKEKLDGGIEP
YRLPEVIQKCNARWTTEEQLLAVQAIRKYGRDFQAISDVIGNKSVVQVKNFFVNYRRRFNID
EVLQEWEAEHGKEETNGPSNQKPVKSPDNSIKMPEEEDEAPVLDVRYASAS
00000000000000000000000000000000000000000000000000000000000000
00000000000000000000000000000000000000000000000000000000000000
00000000000000000000000000000000000000000000000000000000000000
00000000000000000000000000000000000000000000000000000000000000
00000000000000000000000000000000000000000000000000000000000000
00000000000000000000000000000000000000000000000000000000000000
00000000000000000000000000000000000000000000000000000000000000
000000000000000000000000000000000000000000000000000
>DP02529
MPLVKRNIDPRHLCHTALPRGIKNELECVTNISLANIIRQLSSLSKYAEDIFGELFNEAHSF
SFRVNSLQERVDRLSVSVTQLDPKEEELSLQDITMRKAFRSSTIQDQQLFDRKTLPIPLQET
YDVCEQPPPLNILTPYRDDGKEGLKFYTNPSYFFDLWKEKMLQDTEDKRKEKRKQKQKNLDR
PHEPEKVPRAPHDRRREWQKLAQGPELAEDDANLLHKHIEVANGPASHFETRPQTYVDHMDG
SYSLSALPFSQMSELLTRAEERVLVRPHEPPPPPPMHGAGDAKPIPTCISSATGLIENRPQS
PATGRTPVFVSPTPPPPPPPLPSALSTSSLRASMTSTPPPPVPPPPPPPATALQAPAVPPPP
APLQIAPGVLHPAPPPIAPPLVQPSPPVARAAPVCETVPVHPLPQGEVQGLPPPPPPPPLPP
PGIRPSSPVTVTALAHPPSGLHPTPSTAPGPHVPLMPPSPPSQVIPASEPKRHPSTLPVISD
ARSVLLEAIRKGIQLRKVEEQREQEAKHERIENDVATILSRRIAVEYSDSEDDSEFDEVDWL
E
xxxxxxxxxxxxxxxxxxxxxxxxxxxxxxxxxxxxxxxxxxxxxxxxxxxxxxxxxxxxxx
xxxxxxxxxxxxxxxxxxxxxxxxxxxxxxxxxxxxxxxxxxxxxxxxxxxxxxxxxxxxxx
xxxxxxxxxxxxxxxxxxxxxxxxxxxxxxxxxxxxxxxxxxxxxxxxxxxxxxxxxxxx00
00000000000000000000000000000000000000000000000000000000000000
00000000000000000000000000000000000000000000000000000000000000
00000000000000000000000000000000000000000000000000000000000000
00000000000000000000000000000000000000000000000000000000000000
000000000000000000000000000000000000000000000000000000000000xx
xxxxxxxxxxxxxxxxxxxxxxxxxxxxxxxxxxxxxxxxxxxxxxxx00000000000000
0
>DP02530
MGIELLCLFFLFLGRNDHVQGGCALGGAETCEDCLLIGPQCAWCAQENFTHPSGVGERCDTP
ANLLAKGCQLNFIENPVSQVEILKNKPLSVGRQKNSSDIVQIAPQSLILKLRPGGAQTLQVH
VRQTEDYPVDLYYLMDLSASMDDDLNTIKELGSRLSKEMSKLTSNFRLGFGSFVEKPVSPFV
KTTPEEIANPCSSIPYFCLPTFGFKHILPLTNDAERFNEIVKNQKISANIDTPEGGFDAIMQ
AAVCKEKIGWRNDSLHLLVFVSDADSHFGMDSKLAGIVIPNDGLCHLDSKNEYSMSTVLEYP
TIGQLIDKLVQNNVLLIFAVTQEQVHLYENYAKLIPGATVGLLQKDSGNILQLIISAYEELR

SEVELEVLGDTEGLNLSFTAICNNGTLFQHQKKCSHMKVGDTASFSVTVNIPHCERRSRHII
IKPVGLGDALELLVSPECNCDCQKEVEVNSSKCHHGNGSFQCGVCACHPGHMGPRCECGEDM
LSTDSCKEAPDHPSCSGRGDCYCGQCICHLSPYGNIYGPYCQCDNFSCVRHKGLLCGGNGDC
DCGECVCRSGWTGEYCNCTTSTDSCVSEDGVLCSGRGDCVCGKCVCTNPGASGPTCERCPTC
GDPCNSKRSCIECHLSAAGQAREECVDKCKLAGATISEEEDFSKDGSVSCSLQGENECLITF
LITTDNEGKTIIHSINEKDCPKPPNIPMIMLGVSLAILLIGVVLLCIWKLLVSFHDRKEVAK
FEAERSKAKWQTGTNPLYRGSTSTFKNVTYKHREKQKVDLSTDC
00000000000000000000000000000000000000000000000000000000000000
00000000000000000000000000000000000000000000000000000000000000
00000000000000000000000000000000000000000000000000000000000000
00000000000000000000000000000000000000000000000000000000000000
00000000000000000000000000000000000000000000000000000000000000
00000000000000000000000000000000000000000000000000000000000000
00000000000000000000000000000000000000000000000000000000000000
00000000000000000000000000000000000000000000000000000000000000
00000000000000000000000000000000000000000000000000000000000000
00000000000000000000000000000000000000000000000000000000000000
00000000000000000000000000000000000000000000000000000000000000
00000000000000000000000000000000000000000000000000000000000000
00000000000000000000000000000000000000000000
>DP02548
MKKVLGVILGGLLLLPVVSNAADAQKAADNKKPVNSWTCEDFLAVDESFQPTAVGFAEALNN
KDKPEDAVLDVQGIATVTPAIVQACTQDKQANFKDKVKGEWDKIKKDM
xxxxxxxxxxxxxxxxxxxxx00000000000000000000000000000000000000000
000000000000000000000000000000000000000000000000
>DP02562
MSEGITDIEESQIQTNYDKVVYKFDDMELDENLLRGVFGYGFEEPSAIQQRAIMPIIEGHDV
LAQAQSGTGKTGTFSIAALQRIDTSVKAPQALMLAPTRELALQIQKVVMALAFHMDIKVHAC
IGGTSFVEDAEGLRDAQIVVGTPGRVFDNIQRRRFRTDKIKMFILDEADEMLSSGFKEQIYQ
IFTLLPPTTQVVLLSATMPNDVLEVTTKFMRNPVRILVKKDELTLEGIKQFYVNVEEEEYKY
ECLTDLYDSISVTQAVIFCNTRRKVEELTTKLRNDKFTVSAIYSDLPQQERDTIMKEFRSGS
SRILISTDLLARGIDVQQVSLVINYDLPANKENYIHRIGRGGRFGRKGVAINFVTNEDVGAM
RELEKFYSTQIEELPSDIATLLN
xxxxxxxxxxxxxxxxxxxxxxxxxxxxxxxxxxxxxxxxxxxxxxxxxxxxxxxxxxxxxx
xxxxxxxxxxxxxxxxxxxxxxxxxxxxxxxxxxxxxxxxxxxxxxxxxxxxxxxxxxxxxx
xxxxxxxxxxxxxxxxxxxxxxxxxxxxxxxxxxxxxxxxxxxxxxxxxxxxxxxxxxxxxx
xxxxxxxxxxxxxxxxxxxxxxxxxxxxxxxxxxxx11111111111xxxxxxxxxxxxxxx
xxxxxxxxxxxxxxxxxxxxxxxxxxxxxxxxxxxxxxxxxxxxxxxxxxxxxxxxxxxxxx
xxxxxxxxxxxxxxxxxxxxxxxxxxxxxxxxxxxxxxxxxxxxxxxxxxxxxxxxxxxxxx
xxxxxxxxxxxxxxxxxxxxxxx
>DP02570
MSFLFGSRSSKTFKPKKNIPEGSHQYELLKHAEATLGSGNLRMAVMLPEGEDLNEWVAVNTV
DFFNQINMLYGTITDFCTEESCPVMSAGPKYEYHWADGTNIKKPIKCSAPKYIDYLMTWVQD
QLDDETLFPSKIGVPFPKNFMSVAKTILKRLFRVYAHIYHQHFDPVIQLQEEAHLNTSFKHF

IFFVQEFNLIDRRELAPLQELIEKLTSKDR
xxxxxxxxxx1111111111111xxxxxxxxxxxxxxxxxxxxxxxxxxxxxxxxxxxxx
xxxxxxxxxxxxxxxxxxxxxxxxxxxxxxxxxxxxxxxxxxxxxxxxxxxxxxxxxxxx
xxxxxxxxxxxxxxxxxxxxxxxxxxxxxxxxxxxxxxxxxxxxxxxxxxxxxxxxxxxx
xxxxxxxxxxxxxxxxxxxxxxxxxxxxxx
>DP02591
MSVNSEKSSSSERPEPQQKAPLVPPPPPPPPPPPPPLPDPTPPEPEEEILGSDDEEQEDPAD
YCKGGYHPVKIGDLFNGRYHVIRKLGWGHFSTVWLCWDMQGKRFVAMKVVKSAQHYTETALD
EIKLLKCVRESDPSDPNKDMVVQLIDDFKISGMNGIHVCMVFEVLGHHLLKWIIKSNYQGLP
VRCVKSIIRQVLQGLDYLHSKCKIIHTDIKPENILMCVDDAYVRRMAAEATEWQKAGAPPPS
GSAVSTAPQQKPIGKISKNKKKKLKKKQKRQAELLEKRLQEIEELEREAERKIIEENITSAA
PSNDQDGEYCPEVKLKTTGLEEAAEAETAKDNGEAEDQEEKEDAEKENIEKDEDDVDQELAN
IDPTWIESPKTNGHIENGPFSLEQQLDDEDDDEEDCPNPEEYNLDEPNAESDYTYSSSYEQF
NGELPNGRHKIPESQFPEFSTSLFSGSLEPVACGSVLSEGSPLTEQEESSPSHDRSRTVSAS
STGDLPKAKTRAADLLVNPLDPRNADKIRVKIADLGNACWVHKHFTEDIQTRQYRSIEVLIG
AGYSTPADIWSTACMAFELATGDYLFEPHSGEDYSRDEDHIAHIIELLGSIPRHFALSGKYS
REFFNRRGELRHITKLKPWSLFDVLVEKYGWPHEDAAQFTDFLIPMLEMVPEKRASAGECLR
HPWLNS
000000000000000000000000000000000000000000000000000000000000
000000000000000000xxxxxxxxxxxxxxxxxxxxxxxxxxxxxxxxxxxxxxxxxx
xxxxxxxxxxxxxxxxxxxxxxxxxxxxxxxxxxxxxxxxxxxxxxxxxxxxxxxxxxxx
xxxxxxxxxxxxxxxxxxxxxxxxxxxxxxxxxxxxxxxxxx000000000000000000
000000000000000000000000000000000000000000000000000000000000
000000000000000000000000000000000000000000000000000000000000
000000000000000000000000000000000000000000000000000000000000
000000000000000000000000000000000000000000000000000000000000
000000000000000xxxxxxxxxxxxxxxxxxxxxxxxxxxxxxxxxxxxxxxxxxxxx
xxxxxxxxxxxxxxxxxxxxxxxxxxxxxxxxxxxxxxxxxxxxxxxxxxxxxxxxxxxx
xxxxxxxxxxxxxxxxxxxxxxxxxxxxxxxxxxxxxxxxxxxxxxxxxxxxxxxxxxxx
xxxxxx
>DP02592
MGSSQSVEIPGGGTEGYHVLRVQENSPGHRAGLEPFFDFIVSINGSRLNKDNDTLKDLLKAN
VEKPVKMLIYSSKTLELRETSVTPSNLWGGQGLLGVSIRFCSFDGANENVWHVLEVESNSPA
ALAGLRPHSDYIIGADTVMNESEDLFSLIETHEAKPLKLYVYNTDTDNCREVIITPNSAWGG
EGSLGCGIGYGYLHRIPTRPFEEGKKISLPGQMAGTPITPLKDGFTEVQLSSVNPPSLSPPG
TTGIEQSLTGLSISSTPPAVSSVLSTGVPTVPLLPPQVNQSLTSVPPMNPATTLPGLMPLPA
GLPNLPNLNLNLPAPHIMPGVGLPELVNPGLPPLPSMPPRNLPGIAPLPLPSEFLPSFPLVP
ESSSAASSGELLSSLPPTSNAPSDPATTTAKADAASSLTVDVTPPTAKAPTTVEDRVGDSTP
VSEKPVSAAVDANASESP
xxxxxxxxxxxxxxxxxxxxxxxxxxxxxxxxxxxxxxxxxxxxxxxxxxxxxxxxxxxx
xxxxxxxxxxxxxxxxxxxxxxxxxxxxxxxxxxxxxxxxxxxxxxxxxxxxxxxxxxxx
xxxxxxxxxxxxxxxxxxxxxxxxxxxxxxxxxxxxxxxxxxxxxxxxxxxxxxxxxxxx
xxxxxxxxxxxxxxxxxxxxxx00000000000000000000000000000000000000
000000000000000000000000000000000000000000000000000000000000

000000000000000000000000000000000000000000000000000000000000
000000000000000000000000000000000000000000000000000000000000
000000000000000000
>DP02593
MMAAALGPPEVIAQLENAAKVLMAPPSMVSNEQRQHAEHIFLSFRKSKSPFAVCRHILETSK
VDYVLFQAATAIMEAVVREWVLLEKGSIESLRTFLLTYVLQRPNLQKYVREQILLAVAVIVK
RGSLDKSIDCKSIFHEVSQLISSGNPTVQTLACSILTALLSEFSSSSKTSNIGLSMEFHGNC
KRVFQEEDLRQIFMLTVGVLQEFSRRENLSAQMSSVFQRYLALANQVLSWNFLPPKLGRHYI
AMFESSQNVLLKPTESWREALLDSRVMELFFTVHRKIREDSDMAQDSLQCLAQLASLHGPIF
PDEGSQVDYLAHFIEGLLNTINGIEIEDSEAVGISSIISNLITVFPRNVLTAIPSELFSSFV
NCLTHLTCSFGRSAALEEVLDKDDMVYMEAYDKLLESWLTLVRDDKHFHKGFFTQHAVQVFN
SYIQCHLAAPDGTRNLTANGVASREEEEISELQEDDRDQFSDQLASVGMLGRIAAEHCMPLL
TSLLEERVTRLHGQLQRHQQFLASPGSSTIDNKMLDDLYEDIHWLILVTGYLLADDTQGET
PLIPPEIMEYSIKHSSEVDINTTLQILGSPGEKASSIPGYSRTDSVIRLLSAVLRVSEVESR
AIRADLTHLLSPQMGKDIVWFLKRWAKTYLLVDEKLYDQISLPLSTAFGADTEGSQWIIGYL
LQKVISNLSVWSSEQDLANDTVQLLVTLVERRERANLVIQCENWWNLAKQFASRSPPLNFLS
SPVQRTLMKALVLGGFAHMDTETKQQYWTEVLQPLQQRFLRVINQENFQQMCQQEEVKQEIT
ATLEALCGIAEATQIDNVAILFNFLMDFLNNCIGLMEVYKNTPETVNLIIEVFVEVAHKQIC
YLGESKAMHLYEACLTLLQVYSKNNLGRQRIDVTAEEEQYQDLLLIMELLTNLLSKEFIDFS
DTDEVFRGHEPGQAAGRSVSAADVVLYGVNLILPLMSQDLLKFPTLCNQYYKLITFICEIFP
EKIPQLPEDLFKSLMYSLELGMTSMSSEVCQLCLEALTPLAEQCAKAQETDSPLFLATRHFL
KLVFDMLVLQKHNTEMTTAAGEAFYTLVCLHQAEYSELVETLLSSQQDPVIYQRLADAFNKL
TASSTPPALDRKQKMAFLKSLEEFMANVGGLLCVK
xxxxxxxxxxxxxxxxxxxxxxxxxxxxxxxxxxxxxxxxxxxxxxxxxxxxxxxxxxxx
xxxxxxxxxxxxxxxxxxxxxxxxxxxxxxxxxxxxxxxxxxxxxxxxxxxxxxxxxxxx
xxxxxxxxxxxxxxxxxxxxxxxxxxxxxxxxxxxxxxxxxxx111111111111xxxxx
xxxxxxxxxxxxxxxxxxxxxxxxxxxxxxxxxxxxxxxxxxxxxxxxxxxx00000000
000000000000xxxxxxxxxxxxxxxxxxxxxxxxxxxxxxxxxxxxxxxxxxxxxxxx
xxxxxxxxxxxxxxxxxxxxxxxxxxxxxxxxxxxxxxxxxxxxxxxxxxxxxxxxxxxx
xxxxxxxxxxxxxxxxxxxxxxxxxxxxxxxxxxxxxxxxxxxxxxxxxxxxxxxxxxxx
xxxxxxxxxxxxxxxxxxxxxxxxxxxxxxxxxxxxxxxxxxxxxxxxxxxxxxxxxxxx
xxxxxxxxxx111111111111111111111111xxxxxxxxxxxxxxxxxxxxxxxxxx
xxxxxxxxxxxxxxxxxxxxxxxxxxxxxxxxxxxxxxxxxxxxxxxxxxxxxxxxxxxx
xxxxxxxxxxxxxxxxxxxxxxxxxxxxxxxxxxxxxxxxxxxxxxxxxxxxxxxxxxxx
xxxxxxxxxxxxxxxxxxxxxxxxxxxxxxxxxxxxxxxxxxxxxxxxxxxxxxxxxxxx
xxxxxxxxxxxxxxxx11111111111xxxxxxxxxxxxxxxxxxxxxxxxxxxxxxxxx
xxxxxxxxxxxxxxxxxxxxxxxxxxxxxxxxxxxxxxxxxxxxxxxxxxxxxxxxxxxx
xxxxxxxxxxxxxxxxxxxxxxxxxxxxxxxxxxxxxxxxxxxxxxxxxxxxxxxxxxxx
000000000000000000xxxxxxxxxxxxxxxxxxxxxxxxxxxxxxxxxxxxxxxxxx
xxxxxxxxxxxxxxxxxxxxxxxxxxxxxxxxxxxxxxx1111111111111xxxxxxxx
xxxxxxxxxxxxxxxxxxxxxxxxxxxxxxxxxxxxxxxxxxxxxxxxxxxxxxxxxxxx
xxxxxxxxxxxxxxxxxxxxxxxxxxxxxxxxxxx
>DP02599
MDRFVWTSGLLEINETLVIQQRGVRIYDGEEKIKFDAGTLLLSTHRLIWRDQKNHECCMAIL

LSQIVFIEEQAAGIGKSAKIVVHLHPAPPNKEPGPFQSSKNSYIKLSFKEHGQIEFYRRLSE
EMTQRRWENMPVSQSLQTNRGPQPGRIRAVGIVGIERKLEEKRKETDKNISEAFEDLSKLMI
KAKEMVELSKSIANKIKDKQGDITEDETIRFKSYLLSMGIANPVTRETYGSGTQYHMQLAKQ
LAGILQVPLEERGGIMSLTEVYCLVNRARGMELLSPEDLVNACKMLEALKLPLRLRVFDSGV
MVIELQSHKEEEMVASALETVSEKGSLTSEEFAKLVGMSVLLAKERLLLAEKMGHLCRDDSV
EGLRFYPNLFMTQS
xxxxxxxxxxxxxxxxxxxxxxxxxxxxxxxxxxxxxxxxxxxxxxxxxxxxxxxxxxxx
xxxxxxxxxxxxxxxxxxxxxxxxxxxxxxxxxxxxxxxxxxxxxxxxxxxxxxxxxxxx
xxxxxxxxxxxxxxxxxxxxxxx11111111111111111111xxxxxxxxxxxxxxxxx
xxxxxxxxxxxxxxxxxxxxxxxxxxxxxxxxxxxxxxxxxxxxxxxxxxxxxxxxxxxx
xxxxxxxxxxxxxxxxxxxxxxxxxxxxxxxxxxxxxxxxxxxxxxxxxxxxxxxxxxxx
xxxxxxxxxxxxxxxxxxxxxxxxxxxxxxxxxxxxxxxxxxxxxxxxxxxxxxxxxxxx
xxxxxxxxxxxxxx
>DP02605
MTTCSRQFTSSSSMKGSCGIGGGIGGGSSRISSVLAGGSCRAPSTYGGGLSVSSSRFSSGGA
CGLGGGYGGGFSSSSSSFGSGFGGGYGGGLGAGLGGGFGGGFAGGDGLLVGSEKVTMQNLND
RLASYLDKVRALEEANADLEVKIRDWYQRQRPAEIKDYSPYFKTIEDLRNKILTATVDNANV
LLQIDNARLAADDFRTKYETELNLRMSVEADINGLRRVLDELTLARADLEMQIESLKEELAY
LKKNHEEEMNALRGQVGGDVNVEMDAAPGVDLSRILNEMRDQYEKMAEKNRKDAEEWFFTKT
EELNREVATNSELVQSGKSEISELRRTMQNLEIELQSQLSMKASLENSLEETKGRYCMQLAQ
IQEMIGSVEEQLAQLRCEMEQQNQEYKILLDVKTRLEQEIATYRRLLEGEDAHLSSSQFSSG
SQSSRDVTSSSRQIRTKVMDVHDGKVVSTHEQVLRTKN
xxxxxxxxxxxxxxxxxxxxxxxxxxxxxxxxxxxxxxxxxxxxxxxxxxxxxxxxxxxx
xxxxxxxxxxxxxxxxxxxxxxxxxxxxxxxxxxxxxxxxxxxxxxxxxxxxxxxxxxxx
xxxxxxxxxxxxxxxxxxxxxxxxxxxxxxxxxxxxxxxxxxxxxxxxxxxxxxxxxxxx
xxxxxxxxxxxxxxxxxxxxxxxxxxxxxxxxxxxxxxxxxxxxxxxxxxxxxxxxxxxx
xxxxxxxxxxxxxxxxxxxxxxxxxxxxxxxxxxxxxxxxxxxxx111111111111111
11111111111111111111xxxxxxxxxxxxxxxxxxxxxxxxxxxxxxxxxxxxxxxx
xxxxxxxxxxxxxxxxxxxxxxxxxxxxxxxxxxxxxxxxxxxxxxxxxxxxxxxxxxxx
xxxxxxxxxxxxxxxxxxxxxxxxxxxxxxxxxxxxxx
>DP02622
MLPAATASLLGPLLTACALLPFAQGQTPNYTRPVFLCGGDVKGESGYVASEGFPNLYPPNKE
CIWTITVPEGQTVSLSFRVFDLELHPACRYDALEVFAGSGTSGQRLGRFCGTFRPAPLVAPG
NQVTLRMTTDEGTGGRGFLLWYSGRATSGTEHQFCGGRLEKAQGTLTTPNWPESDYPPGISC
SWHIIAPPDQVIALTFEKFDLEPDTYCRYDSVSVFNGAVSDDSRRLGKFCGDAVPGSISSEG
NELLVQFVSDLSVTADGFSASYKTLPRGTAKEGQGPGPKRGTEPKVKLPPKSQPPEKTEESP
SAPDAPTCPKQCRRTGTLQSNFCASSLVVTATVKSMVREPGEGLAVTVSLIGAYKTGGLDLP
SPPTGASLKFYVPCKQCPPMKKGVSYLLMGQVEENRGPVLPPESFVVLHRPNQDQILTNLSK
RKCPSQPVRAAASQD
xxxxxxxxxxxxxxxxxxxxxxxxxxxxxxxxxxxxxxxxxxxxxxxxxxxxxxxxxxxx
xxxxxxxxxxxxxxxxxxxxxxxxxxxxxxxxxxxxxxxxxxxxxxxxxxxxxxxxxxxx
xxxxxxxxxxxxxxxxxxxxxxxxx1111111111111110000xxxxxxxxxxxxxxxx
xx00000000000xxxxxxxxxxxxxxxxxxx0000000000000xxxxxxx00000000
0000xxxxxxxxxxxxxxx0000000000xxxxxxxxxxxxxxxxxxxxxxxxxxxxxxx

xxxxxxxxxxxxxxxxxxxxxxxxxxxxxxxxxxxxxxxxxxxxxxxxxxxxxxxxxxxx
xxxxxxxxxxxxxxxxxxxxxxxxxxxxxxxxxxxxxxxxxxxxxxxxxxxxxxxxxxxx
xxxxxxxxxxxxxxx
>DP02635
MQRDCIMDYKESCPSVSIPSSDEHREKKKRFTVYKVLVSVGRSEWFVFRRYAEFDKLYNSLK
KQFPAMALKIPAKRIFGDNFDPDFIKQRRAGLNEFIQNLVRYPELYNHPDVRAFLQMDSPRH
QSDPSEDEDERSTSKPHSTSRNINLGPTGNPHAKPTDFDFLKVIGKGSFGKVLLAKRKLDGK
FYAVKVLQKKIVLNRKEQKHIMAERNVLLKNVKHPFLVGLHYSFQTTEKLYFVLDFVNGGEL
FFHLQRERSFPEPRARFYAAEIASALGYLHSIKIVYRDLKPENILLDSMGHVVLTDFGLCKE
GIAISDTTTTFCGTPEYLAPEVIRKQPYDNTVDWWCLGAVLYEMLYGLPPFYCRDVAEMYDN
ILHKPLNLRPGVSLTAWSILEELLEKNRQNRLGAKEDFLEIQNHPFFESLSWTDLVQKKIPP
PFNPNVAGPDDIRNFDAVFTEETVPYSVCVSSDYSIVNASVLEADDAFVGFSYAPPSEDLFL
xxxxxxxxxxxxxxxxxxxxxxxxxxxxxxxxxxxxxxxxxxxxxxxxxxxxxxxxxxxx
xxxxxxxxxxxxxxxxxxxxxxxxxxxxxxxxxxxxxxxxxxxxxxxxxxxxxxxxxxxx
x1111111111111111111111111111111111xxxxxxxxxxxxxxxxxxxxxxxxx
xxxxxxxxxxxxxxxxxxxxxxxxxxxxxxxxxxxxxxxxxxxxxxxxxxxxxxxxxxxx
xxxxxxxxxxxxxxxxxxxxxxxxxxxxxxxxxxxxxxxxxxxxxxxxxxxxxxxxxxxx
xxxxxxxxxxxxxxxxxxxxxxxxxxxxxxxxxxxxxxxxxxxxxxxxxxxxxxxxxxxx
xxxxxxxxxxxxxxxxxxxxxxxxxxxxxxxxxxxxxxxxxxxxxxxxxxxxxxxxxxxx
xxxxxxxxxxxxxxxxxxxxxxxxxxxxxxxxxxxxxxxxxxxxxxxxxxxxxxxxxxxx
>DP02636
MAVTSHHMVPVFVLMSACLATAGPEPSTRCELSPISASHPVQALMESFTVLSGCASRGTTGL
PREVHILNLRSTDQGLGQPQREVTLHLNPIASVHTHHKPVVFLLNSPQPLVWHVKTERLAAG
VPRLFLVSEGSVVQFSSGNFSLTAETEERSFPQENEHLLHWAQKEYGAVTSFTELKIARNIY
IKVGEDQVFPPTCNIGKNFLSLNYLAEYLQPKAAEGCVLASQPHEKEVHIIELISPNSNPYS
TFQVDIIIDIRPAREDPEVVKNLVLILKCKKSVNWVIKSFDVKGNLKVIAPDSIGFGKESER
SMTVTKLVRNDYPSTQENLMKWALDNGYSPVTSYTIAPVANRFHLRLENNEEMRDEEVHTIP
PELRILLGPDHLPALDSPPFQGEIPNGGFPFPFPDIPRRGWKEGEDRIPRPKEPIIPRVQLL
PDHREPEEVQGGVNIALSVKCDNEKMVVAVDKDSFQTNGYSGMELTLLDPSCKAKMNGTHFV
LESPLNGCGTRHRRSAPDGVVYYNSIVVQAPSPGDSSGWPDGYEDLESGDNGFPGDTDEGET
APLSRAGVVVFNCSLRQLRSPSGFQDQLDGNATFNMELYNTDLFLVPSPGVFSVAENEHVYV
EVSVTKADQDLGFAIQTCFISPYSNPDRMSDYTIIENICPKDDSVKFYSSKRVHFPIPHAEV
DKKRFSFVFKSVFNTSLLFLHCELTLCSRNKGSQKLPKCVTPDDACTSLDATMIWTMMQNKK
TFTKPLAVVLQVDYKENVPNMKESSPVPPPPQIFHGLDTLTVMGIAFAAFVIGALLTGALWY
IYSHTGETARRQQVPTSPPASENSSAAHSIGSTQSTPCSSSSTA
xxxxxxxxxxxxxxxxxxxxxxxxxxxxxxxxxxxxxxxxxxxxxxxxxxxxxxxxxxxx
xxxxxxxxxxxxxxxxxxxxxxxxxxxxxxxxxxxxxxxxxxxxxxxxxxxxxxxxxxxx
xxxxxxxxxxxxxxxxxxxxxxxxxxxxxxxxxxxxxxxxxxxxxxxxxxxxxxxxxxxx
xxxxxxxxxxxxxxxxxxxxxxxxxxxxxxxxxxxxxxxxxxxxxxxxxxxxxxxxxxxx
xxxxxxxxxxxxxxxxxxxxxxxxxxxxxxxxxxxxxxxxxxxxxxxxxxxxxxxxxxxx
xxxxxxxxxxxxxxxxxxxxxxxxxxxxxxxxxxxxxxxxxxxxxxxxxxxxxxxxxxxx
xxxxxxxxxxxxxxxxxxxxxxxxxxxxxxxxxxxxxxxxxxxxxxxxxxxxxxxxxxxx
xxxxxxxxxxxxxxxxxxxxxxxxxxxxxxxxxxxxxxxxxxxxxxxxxxxxxxxxxxxx
xxxxxxxxxxxxxxxxxxxxxxxxxxxxxxxxxxxxxxxxxxxxxxxxxxxxxxxxxxxx

xxxxxxxxxxxxxxxxxxxxxxxxxxxxxxxxxxxxxxxxxxxxxxxxxxxxxxxxxxxx
xxxxxxxxxxxxxxxxxxxxxxxxxxxxxxxxxxxxxxxxxxxxxxxxxxxxxxxxxxxx
xxxxxxxxxxxxxxxxxxxxxxxxxxxxxxxxxxxxxxxxxxxxx111111111111111
xxxxxxxxxxxxxxxxxxxxxxxxxxxxxxxxxxxxxxxxxxxxxxxxxxxxxxxxxxxx
xxxxxxxxxxxxxxxxxxxxxxxxxxxxxxxxxxxxxxxxxxxx

>DP02640
MESSSSLKGSALGKLVVTSGLLHSSWSKILEIHNPPYSNHDPGLQVSKKKKDSGLEFQIHRE
EKFTLVVFSAPPICRSSSSDSTLLHVKDKENPFPFLCSENNPSFSLHTPAFNLFTSASTSLT
YLKSELLQTLKSEKPVIITGAALGGSVASLYTLWLLETIEPTLKRPLCITFGSPLIGDASLQ
QILENSVRNSCFLHVVSAQTRIKMDFFKPFGTFLICFDSGCVCIEDHVAVTELLNGVHDSGL
VDYSQVLNRLDQSMLSLADSRLIPEDVIKGIEKRAEMKNLRFDMMFKKLNDMKISMAYIEWY
KKKCKEVKIGYYDRFKTQLAFPSKEFDINIKNHHKSELNRFWKSVVEEVERRPQSDASILKR
RFLFSGNNYRRMIEPLDIAEYYLEGRKEYRTTGRSHHYVMLEKWFGMESILIEKERCKKRDL
SDLLTFDSCFWAEVEDSLIVINQLNTTVGMRDDVREVLTRKLVEFEGYVWEIITKREVSPEI
FLEESSFMKWWKEYKKIKGFNSSYLTEFMNTRKYESYGKSQ
xxxxxxxxxxxxxxxxxxxxxxxxxxxxxxxxx11111111111111111xxxxxxxxxx
xxxxxxxxxxxxxxxxxxxxxxxxxxxxxxxxxxxxxxxxxxxxxxxxxxxxxxxxxxxx
xxxxxxxxxxxxxxxxxxxxxxxxxxxxxxxxxxxxxxxxxxxxxxxxxxxxxxxxxxxx
xxxxxxxxxxxxxxxxxxxxxxxxxxxxxxxxxxxxxxxxxxxxxxxxxxxxxxxxxxxx
xxxxxxxxxxxxxxxxxxxxxxxxxxxxxxxxxxxxxxxxxxxxxxxxxxxxxxxxxxxx
xxxxxxxxxxxxxxxxxxxxxxxxxxxxxxxxxxxxxxxxxxxxxxxxxxxxxxxxxxxx
xxxxxxxxxxxxxxxxxxxxxxxxxxxxxxxxxxxxxxxxxxxxxxxxxxxxxxxxxxxx
xxxxxxxxxxxxxxxxxxxxxxxxxxxxxxxxxxxxxxxxxxxxxxxxxxxxxxxxxxxx
xxxxxxxxxxxxxxxxxxxxxxxxxxxxxxxxxxxxxxxxx

>DP02644
MSSSSSSSTSMIDLMAAIIKGEPVIVSDPANASAYESVAAELSSMLIENRQFAMIVTTSIAV
LIGCIVMLVWRRSGSGNSKRVEPLKPLVIKPREEEIDDGRKKVTIFFGTQTGTAEGFAKALG
EEAKARYEKTRFKIVDLDDYAADDDEYEEKLKKEDVAFFFLATYGDGEPTDNAARFYKWFTE
GNDRGEWLKNLKYGVFGLGNRQYEHFNKVAKVVDDILVEQGAQRLVQVGLGDDDQCIEDDFT
AWREALWPELDTILREEGDTAVATPYTAAVLEYRVSIHDSEDAKFNDINMANGNGYTVFDAQ
HPYKANVAVKRELHTPESDRSCIHLEFDIAGSGLTYETGDHVGVLCDNLSETVDEALRLLDM
SPDTYFSLHAEKEDGTPISSSLPPPFPPCNLRTALTRYACLLSSPKKSALVALAAHASDPTE
AERLKHLASPAGKDEYSKWVVESQRSLLEVMAEFPSAKPPLGVFFAGVAPRLQPRFYSISSS
PKIAETRIHVTCALVYEKMPTGRIHKGVCSTWMKNAVPYEKSENCSSAPIFVRQSNFKLPSD
SKVPIIMIGPGTGLAPFRGFLQERLALVESGVELGPSVLFFGCRNRRMDFIYEEELQRFVES
GALAELSVAFSREGPTKEYVQHKMMDKASDIWNMISQGAYLYVCGDAKGMARDVHRSLHTIA
QEQGSMDSTKAEGFVKNLQTSGRYLRDVW
xxxxxxxxxxxxxxxxxxxxxxxxxxxxxxxxxxxxxxxxxxxxxxxxxxxxxxxxxxxx
xxxxxxxxxx00000000000000000000000000000000000000000000000000
000000000000000000000000000000000000000000000000000000000000
000000000000000000000000000000000000000000000000000000000000
00000000000000111111111111111111111111111111111111111xxxxxxx
xxxxxxxxxxxxxxxxxxxxxxxxxxxxxxxxxxxxxxxxxxxxxxxxxxxxxxxxxxxx
xxxxxxxxxxxxxxxxxxxxxxxxxxxxxxxxxxxxxxxxxxxxxxxxxxxxxxxxxxxx

xxxxxxxxxxxxxxxxxxxxxxxxxxxxxxxxxxxxxxxxxxxxxxxxxxxxxxxxxxxx
xxxxxxxxxxxxxxxxxxxxxxxxxxxxxxxxxxxxxxxxxxxxxxxxxxxxxxxxxxxx
xxxxxxxxxxxxxxxxxxxxxxxxxxxxxxxxxxxxxxxxxxxxxxxxxxxxxxxxxxxx
xxxxxxxxxxxxxxxxxxxxxxxxxxxxxxxxxxxxxxxxxxxxxxxxxxxxxxxxxxxx
xxxxxxxxxxxxxxxxxxxxxxxxxxxx
>DP02661
MGEPGQSPSPRSSHGSPPTLSTLTLLLLLCGHAHSQCKILRCNAEYVSSTLSLRGGGSSGAL
RGGGGGGRGGGVGSGGLCRALRSYALCTRRTARTCRGDLAFHSAVHGIEDLMIQHNCSRQGP
TAPPPPRGPALPGAGSGLPAPDPCDYEGRFSRLHGRPPGFLHCASFGDPHVRSFHHHFHTCR
VQGAWPLLDNDFLFVQATSSPMALGANATATRKLTIIFKNMQECIDQKVYQAEVDNLPVAFE
DGSINGGDRPGGSSLSIQTANPGNHVEIQAAYIGTTIIIRQTAGQLSFSIKVAEDVAMAFSA
EQDLQLCVGGCPPSQRLSRSERNRRGAITIDTARRLCKEGLPVEDAYFHSCVFDVLISGDPN
FTVAAQAALEDARAFLPDLEKLHLFPSDAGVPLSSATLLAPLLSGLFVLWLCIQ
xxxxxxxxxxxxxxxxxxxxxxxxxxxxxxxxxxxxxxxxxxxxxxxxxxxxxxxxxxxx
xxxxxxxxxxxxxxxxxxxxxxxxxxxxxxxxxxxxxxxxxxxxxxxxxxxxxxxxxxxx
xxxxx1111111111111111xxxxxxxxxxxxxxxxxxxxxxxxxxxxxxxxxxxxxxx
xxxxxxxxxxxxxxxxxxxxxxxxxxxxxxxxxxxxxxxxxxxxxxxxxxxxxxxxxxxx
xxxxxxxxxxxxxxxxxxxxxxxxxxxxxxxxxxxxxxxxxxxxxxxxxxxxxxxxxxxx
xxxxxxxxxxxxxxxxxxxxxxxxxxxxxxxxxxxxxxxxxxxxxxxxxxxxxxxxxxxx
xxxxxxxxxxxxxxxxxxxxxxxxxxxxxxxxxxxxxxxxxxxxxxxxxxxx
>DP02664
MKLLEQIEKWAAETPDQTAFVWRDAKITYKQLKEDSDALAHWISSEYPDDRSPIMVYGHMQP
EMIINFLGCVKAGHAYIPVDLSIPADRVQRIAENSGAKLLLSATAVTVTDLPVRIVSEDNLK
DIFFTHKGNTPNPEHAVKGDENFYIIYTSGSTGNPKGVQITYNCLVSFTKWAVEDFNLQTGQ
VFLNQAPFSFDLSVMDIYPSLVTGGTLWAIDKDMIARPKDLFASLEQSDIQVWTSTPSFAEM
CLMEASFSESMLPNMKTFLFCGEVLPNEVARKLIERFPKATIMNTYGPTEATVAVTGIHVTE
EVLDQYKSLPVGYCKSDCRLLIMKEDGTIAPDGEKGEIVIVGPSVSVGYLGSPELTEKAFTM
IDGERAYKTGDAGYVENGLLFYNGRLDFQIKLHGYRMELEEIEHHLRACSYVEGAVIVPIKK
GEKYDYLLAVVVPGEHSFEKEFKLTSAIKKELNERLPNYMIPRKFMYQSSIPMTPNGKVDRK
KLLSEVTA
xxxxxxxxxxxxxxxxxxxxxxxxxxxxxxxxxxxxxxxxxxxxxxxxxxxxxxxxxxxx
xxxxxxxxxxxxxxxxxxxxxxxxxxxxxxxxxxxxxxxxxxxxxxxxxxxxxxxxxxxx
xxxxxxxxxxxxxxxxxxxxxxxxxxxxxxxxxxxxxxxxxxxxxxxxxxxxxxxxxxxx
xxxxxxxxxxxxxxxxxxxxxxxxxxxxxxxxxxxxxxxxxxxxxxxxxxxxxxxxxxxx
xxxxxxxxxxxxxxxxxxxxxxxxxxxxxxxxxxxxxxxxxxxxxxxxxxxxxxxxxxxx
xxxxxxxxxxxxxxxxxxxxxxxxxxxxxxxxxxxxxxxxxxxxxxxxxxxxxxxxxxxx
xxxxxxxxxxxxxxxxxxxxxxx11111111111111111xxxxxxxxxxxxxxxxxxxx
xxxxxxxxxxxxxxxxxxxxxxxxxxxxxxxxxxxxxxxxxxxxxxxxxxxxxxxxxxxx
xxxxxxxx
>DP02693
MAFLDNPTIILAHIRQSHVTSDDTGMCEMVLIDHDVDLEKIHPPSMPGDSGSEIQGSNGETQ
GYVYAQSVDITSSWDFGIRRRSNTAQRLERLRKERQNQIKCKNIQWKERNSKQSAQELKSLF
EKKSLKEKPPISGKQSILSVRLEQCPLQLNNPFNEYSKFDGKGHVGTTATKKIDVYLPLHSS
QDRLLPMTVVTMASARVQDLIGLICWQYTSEGREPKLNDNVSAYCLHIAEDDGEVDTDFPPL

DSNEPIHKFGFSTLALVEKYSSPGLTSKESLFVRINAAHGFSLIQVDNTKVTMKEILLKAVK
RRKGSQKVSGPQYRLEKQSEPNVAVDLDSTLESQSAWEFCLVRENSSRADGVFEEDSQIDIA
TVQDMLSSHHYKSFKVSMIHRLRFTTDVQLGISGDKVEIDPVTNQKASTKFWIKQKPISIDS
DLLCACDLAEEKSPSHAIFKLTYLSNHDYKHLYFESDAATVNEIVLKVNYILESRASTARAD
YFAQKQRKLNRRTSFSFQKEKKSGQQ
000000000000000000000000000000000000000000000000000000000000
000000000000000000000000000000000000000000000000000000000000
000000000000000000000000000000000000000000000000000000000000
000000000000000000000000000000000000000000000000000000000000
000000000000000000000000000000000000000000000000000000000000
000000000000000000000000000000000000000000000000000000000000
000000000000000000000000000000000000000000000000000000000000
000000000000000000000000000000000000000000000000000000000000
00000000000000000000000000
>DP02704
MFAKAFRVKSNTAIKGSDRRKLRADVTTAFPTLGTDQVSELVPGKEELNIVKLYAHKGDAVT
VYVSGGNPILFELEKNLYPTVYTLWSYPDLLPTFTTWPLVLEKLVGGADLMLPGLVMPPAGL
PQVQKGDLCAISLVGNRAPVAIGVAAMSTAEMLTSGLKGRGFSVLHTYQDHLWRSGNKSSPP
SIAPLALDSADLSEEKGSVQMDSTLQGDMRHMTLEGEEENGEVHQAREDKSLSEAPEDTSTR
GLNQDSTDSKTLQEQMDELLQQCFLHALKCRVKKADLPLLTSTFLGSHMFSCCPEGRQLDIK
KSSYKKLSKFLQQMQQEQIIQVKELSKGVESIVAVDWKHPRITSFVIPEPSPTSQTIQEGSR
EQPYHPPDIKPLYCVPASMTLLFQESGHKKGSFLEGSEVRTIVINYAKKNDLVDADNKNLVR
LDPILCDCILEKNEQHTVMKLPWDSLLTRCLEKLQPAYQVTLPGQEPIVKKGRICPIDITLA
QRASNKKVTVVRNLEAYGLDPYSVAAILQQRCQASTTVNPAPGAKDSLQVQIQGNQVHHLGW
LLLEEYQLPRKHIQGLEKALKPGKKK
xxxxxxxxxxxxxxxxxxxxxxxxxxxxxxxxxxxxxxxxxxxxxxxxxxxxxxxxxxxx
xxxxxxxxxxxxxxxxxxxxxxxxxxxxxxxxxxxxxxxxxxxxxxxxxxxxxxxxxxxx
xxxxxxxxxxxxxxxxxxxxxxxxxxxxxxxxxxxxxxxxxxxxxxxxxxxxxxxxx111
111111111111111111111111111111111111111111111111111111111111
1111111111xxxxxxxxxxxxxxxxxxxxxxxxxxxxxxxxxxxxxxxxxxxxxxxxxx
xxxxxxxxxxxxxxxxxxxxxxxxxxxxxxxxxxxx111111111111111111111111
1xxxxxxxxxxxxxxxxxxxxxxxxxxxxxxxxxxxxxxxxxxxxxxxxxxxxxxxxxxx
xxxxxxxxxxxxxxxxxxxxxxxxxxxxxxxxxxxxxxxxxxxxxxxxxxxxxxxxxxxx
xxxxxxxxxxxxxxxxxxxxxxxxxxxxxxxxxxxxxxxxxxxxxxxxxxxxxxxxxxxx
xxxxxxxxxxxxxxxxxxxxxxxxxx
>DP02712
MASMRESDTGLWLHNKLGATDELWAPPSIASLLTAAVIDNIRLCFHGLSSAVKLKLLLGTLH
LPRRTVDEMKGALMEIIQLASLDSDPWVLMVADILKSFPDTGSLNLELEEQNPNVQDILGEL
REKVGECEASAMLPLECQYLNKNALTTLAGPLTPPVKHFQLKRKPKSATLRAELLQKSTETA
QQLKRSAGVPFHAKGRGLLRKMDTTTPLKGIPKQAPFRSPTAPSVFSPTGNRTPIPPSRTLL
RKERGVKLLDISELDMVGAGREAKRRRKTLDAEVVEKPAKEETVVENATPDYAAGLVSTQKL
GSLNNEPALPSTSYLPSTPSVVPASSYIPSSETPPAPSSREASRPPEEPSAPSPTLPAQFKQ
RAPMYNSGLSPATPTPAAPTSPLTPTTPPAVAPTTQTPPVAMVAPQTQAPAQQQPKKNLSLT
REQMFAAQEMFKTANKVTRPEKALILGFMAGSRENPCQEQGDVIQIKLSEHTEDLPKADGQG

STTMLVDTVFEMNYATGQWTRFKKYKPMTNVS
xxxxxxxxxxxxxxxxxxxxxxxxxxxxxxxxxxxxxxxxxxxxxxxxxxxxxxxxxxxx
xxxxxxxxxxxxxxxxxxxxxxxxxxxxxxxxxxxxxxxxxxxxxxxxxxxxxxxxxxxx
xxxxxxxxxxxxxxxxxxxxxxxxxxxxxxxxxxxxxxxxxxxxxxxxxxxxxxxxxxxx
xx0000000000000000000000000000000000000000000000000000000000
000000000000000000000000000000000000000000000000000000000000
000000000000000000000000000000000000000000000000000000000000
000000000000000000000000000000000000000000000000000000000000
000000000000000000000000000000000000000000000000000000000000
00000000000000000000000000000000
>DP02713
MLVIPPGLSEEEEALQKKFNKLKKKKALLALKKQSSSSTTSQGGVKRSLSEQPVMDTATAT
EQAKQLVKSGAISAIKAETKNSGFKRSRTLEGKLKDPEKGPVPTFQPFQRSISADDDLQESS
RRPQRKSLYESFVSSSDRLRELGPDGEEAEGPGAGDGPPRSFDWGYEERSGAHSSASPPRSR
SRDRSHERNRDRDRDRERDRDRDRDRDRERDRDRDRDRDRERDRDRERDRDRDREGPFRR
SDSFPERRAPRKGNTLYVYGEDMTPTLLRGAFSPFGNIIDLSMDPPRNCAFVTYEKMESADQ
AVAELNGTQVESVQLKVNIARKQPMLDAATGKSVWGSLAVQNSPKGCHRDKRTQIVYSDDVY
KENLVDGF
xxxxxxxxxxxxxxxxxxxxxxxxxxxxxxxxxxxxxxxxxxxxxxxxxxxxxxxxxxxx
xxxxxxxxxxxxxxxxxxxxxxxxxxxxxxxxxxxxxxxxxxxxxxxxxxxxxxxxxxxx
xxxxxxxxxxxxxx0000000000000000000000000000000000000000000000
000000000000000000000000000000000000000000000000000000000000
000000000000000000000000000000000000000000000000000000000000
000000000000000000000000000000000000000000000000000xxxxxxxxx
xxxxxxxx
>DP02716
MWIGIISLFPEMFRAITDYGVTGRAVKNGLLSIQSWSPRDFTHDRHRTVDDRPYGGGPGMLM
MVQPLRDAIHAAKAAAGEGAKVIYLSPQGRKLDQAGVSELATNQKLILVCGRYEGIDERVIQ
TEIDEEWSIGDYVLSGGELPAMTLIDSVSRFIPGVLGHEASATEDSFAEGLLDCPHYTRPEV
LEGMEVPPVLLSGNHAEIRRWRLKQSLGRTWLRRPELLENLALTEEQARLLAEFKTEHAQQQ
HKHDGMA
xxxxxxxxxxxxxxxxxxxxxxxxxxxxxxxxxxxxxxxxxxxxxxxxxxxxxxxxxxxx
xxxxxxxxxxxxxxxxxxxxxxxxxxxxxxxxxxxxxxxxxxxxxxxxxxxxxxxxxxxx
xxxxxxxxxxxxxxxxxxxxxxxxxxxxxxxxxxx11111111111xxxxxxxxxxxxxx
xxxxxxxxxxxxxxxxxxxxxxxxxxxxxxxxxxxxxxxxxxxxxxxxxxxxxxxxxxxx
xxxxxxx
>DP02719
MFGCLVAGRLVQTAAQQVAEDKFVFDLPDYESINHVVVFMLGTIPFPEGMGGSVYFSYPDSN
GMPVWQLLGFVTNGKPSAIFKISGLKSGEGSQHPFGAMNIVRTPSVAQIGISVELLDSMAQQ
TPVGNAAVSSVDSFTQFTQKMLDNFYNFASSFAVSQAQMTPSPSEMFIPANVVLKWYENFQR
RLAQNPLFWKT
xxxxxxxxxxxxxxxxxxxxxxxxxxxxxxxxxxxxxxxxxxxxxxxxxxxxxxxxxxxx
xxxxxxxxxxxxxxxxxxxxxxxx000000000000000000xxxxxxxxxxxxxxxxx1
1111111111xxxxxxxxxxxxxxxxxxxxxxxxxxxxxxxxxxxxxxxxxxxxxxxxxx

xxxxxxxxxxx
>DP02721
MNTVPFTSAPIEVTIGIDQYSFNVKENQPFHGIKDIPIGHVHVIHFQHADNSSMRYGYWFDC
RMGNFYIQYDPKDGLYKMMEERDGAKFENIVHNFKERQMMVSYPKIDEDDTWYNLTEFVQMD
KIRKIVRKDENQFSYVDSSMTTVQENELLKSSLQKAGSKMEAKNEDDPAHSLNYTVINFKSR
EAIRPGHEMEDFLDKSYYLNTVMLQGIFKNSSNYFGELQFAFLNAMFFGNYGSSLQWHAMIE
LICSSATVPKHMLDKLDEILYYQIKTLPEQYSDILLNERVWNICLYSSFQKNSLHNTEKIME
NKYPELLGKDNEDDALIYGISDEERDDEDDEHNPTIVGGLYYQRP
xxxxxxxxxxxxxxxxxxxxxxxxxxxxxxxxxxxxxxxxxxxxxxxxxxxxxxxxxxxxxx
xxxxxxxxxxxxxxxxxxxxxxxxxxxxxxxxxxxxxxxxxxxxxxxxxxxxxxxxxxxxxx
xxxxxxxxxxxxxxxxxxxxxxxxxxxx1111111111111111111xxxxxxxxxxxxxxx
xxxxxxxxxxxxxxxxxxxxxxxxxxxxxxxxxxxxxxxxxxxxxxxxxxxxxxxxxxxxxx
xxxxxxxxxxxxxxxxxxxxxxxxxxxxxxxxxxxxxxxxxxxxxxxxxxxxxxxxxxxxxx
xxxxxxxxxxxxxxxxxxxxxxxxxxxxxxxxxxxxxxxxxxxxx
>DP02722
MSGLPPPPPGFEEDSDLALPPPPPPPPGYEIEELDNPMVPSSVNEDTFLPPPPPPPSNFEIN
AEEIVDFTLPPPPPPPGLDELETKAEKKVELHGKRKLDIGKDTFVTRKSRKRAKKMTKKAKR
SNLYTPKAEMPPEHLRKIINTHSDMASKMYNTDKKAFLGALKYLPHAILKLLENMPHPWEQA
KEVKVLYHTSGAITFVNETPRVIEPVYTAQWSATWIAMRREKRDRTHFKRMRFPPFDDDEPP
LSYEQHIENIEPLDPINLPLDSQDDEYVKDWLYDSRPLEEDSKKVNGTSYKKWSFDLPEMSN
LYRLSTPLRDEVTDKNYYYLFDKKSFFNGKALNNAIPGGPKFEPLYPREEEEDYNEFNSIDR
VIFRVPIRSEYKVAFPHLYNSRPRSVRIPWYNNPVSCIIQNDEEYDTPALFFDPSLNPIPHF
IDNNSSLNVSNTKENGDFTLPEDFAPLLAEEEELILPNTKDAMSLYHSPFPFNRTKGKMVRA
QDVALAKKWFLQHPDEEYPVKVKVSYQKLLKNYVLNELHPTLPTNHNKTKLLKSLKNTKYFQ
QTTIDWVEAGLQLCRQGHNMLNLLIHRKGLTYLHLDYNFNLKPTKTLTTKERKKSRLGNSFH
LMRELLKMMKLIVDTHVQFRLGNVDAFQLADGIHYILNHIGQLTGIYRYKYKVMHQIRACKD
LKHIIYYKFNKNLGKGPGCGFWQPAWRVWLNFLRGTIPLLERYIGNLITRQFEGRSNEIVKT
TTKQRLDAYYDLELRNSVMDDILEMMPESIRQKKARTILQHLSEAWRCWKANIPWDVPGMPA
PIKKIIERYIKSKADAWVSAAHYNRERIKRGAHVEKTMVKKNLGRLTRLWIKNEQERQRQIQ
KNGPEITPEEATTIFSVMVEWLESRSFSPIPFPPLTYKNDTKILVLALEDLKDVYASKVRLN
ASEREELALIEEAYDNPHDTLNRIKKYLLTQRVFKPVDITMMENYQNISPVYSVDPLEKITD
AYLDQYLWYEADQRKLFPNWIKPSDSEIPPLLVYKWTQGINNLSEIWDVSRGQSAVLLETTL
GEMAEKIDFTLLNRLLRLIVDPNIADYITAKNNVVINFKDMSHVNKYGLIRGLKFASFIFQY
YGLVIDLLLLGQERATDLAGPANNPNEFMQFKSKEVEKAHPIRLYTRYLDRIYMLFHFEEDE
GEELTDEYLAENPDPNFENSIGYNNRKCWPKDSRMRLIRQDVNLGRAVFWEIQSRVPTSLTS
IKWENAFVSVYSKNNPNLLFSMCGFEVRILPRQRMEEVVSNDEGVWDLVDERTKQRTAKAYL
KVSEEEIKKFDSRIRGILMASGSTTFTKVAAKWNTSLISLFTYFREAIVATEPLLDILVKGE
TRIQNRVKLGLNSKMPTRFPPAVFYTPKELGGLGMISASHILIPASDLSWSKQTDTGITHFR
AGMTHEDEKLIPTIFRYITTWENEFLDSQRVWAEYATKRQEAIQQNRRLAFEELEGSWDRGI
PRISTLFQRDRHTLAYDRGHRIRREFKQYSLERNSPFWWTNSHHDGKLWNLNAYRTDVIQAL
GGIETILEHTLFKGTGFNSWEGLFWEKASGFEDSMQFKKLTHAQRTGLSQIPNRRFTLWWSP
TINRANVYVGFLVQLDLTGIFLHGKIPTLKISLIQIFRAHLWQKIHESIVFDICQILDGELD
VLQIESVTKETVHPRKSYKMNSSAADITMESVHEWEVSKPSLLHETNDSFKGLITNKMWFDV
QLRYGDYDSHDISRYVRAKFLDYTTDNVSMYPSPTGVMIGIDLAYNMYDAYGNWFNGLKPLI

QNSMRTIMKANPALYVLRERIRKGLQIYQSSVQEPFLNSSNYAELFNNDIKLFVDDTNVYRV
TVHKTFEGNVATKAINGCIFTLNPKTGHLFLKIIHTSVWAGQKRLSQLAKWKTAEEVSALVR
SLPKEEQPKQIIVTRKAMLDPLEVHMLDFPNIAIRPTELRLPFSAAMSIDKLSDVVMKATEP
QMVLFNIYDDWLDRISSYTAFSRLTLLLRALKTNEESAKMILLSDPTITIKSYHLWPSFTDE
QWITIESQMRDLILTEYGRKYNVNISALTQTEIKDIILGQNIKAPSVKRQKMAELEAARSEK
QNDEEAAGASTVMKTKTINAQGEEIVVVASADYESQTFSSKNEWRKSAIANTLLYLRLKNIY
VSADDFVEEQNVYVLPKNLLKKFIEISDVKIQVAAFIYGMSAKDHPKVKEIKTVVLVPQLGH
VGSVQISNIPDIGDLPDTEGLELLGWIHTQTEELKFMAASEVATHSKLFADKKRDCIDISIF
STPGSVSLSAYNLTDEGYQWGEENKDIMNVLSEGFEPTFSTHAQLLLSDRITGNFIIPSGNV
WNYTFMGTAFNQEGDYNFKYGIPLEFYNEMHRPVHFLQFSELAGDEELEAEQIDVFS
xxxxxxxxxxxxxxxxxxxxxxxxxxxxxxxxxxxxxxxxxxxxxxxxxxxxxxxxxxxx
xxxxxxxxxxxxxxxxxxxxxxxxxxxxxxxxxxxxxxxxxxxxxxxxxxxxxxxxxxxx
xxxxxxxxxxxxxxxxxxxxxxxxxxxxxxxxxxxxxxxxxxxxxxxxxxxxxxxxxxxx
xxxxxxxxxxxxxxxxxxxxxxxxxxxxxxxxxxxxxxxxxxxxxxxxxxxxxxxxxxxx
xxxxxxxxxxxxxxxxxxxxxxxxxxxxxxxxxxxxxxxxxxxxxxxxxxxxxxxxxxxx
xxxxxxxxxxxxxxxxxxxxxxxxxxxxxxxxxxxxxxxxxxxxxxxxxxxxxxxxxxxx
xxxxxxxxxxxxxxxxxxxxxxxxxxxxxxxxxxxxxxxxxxxxxxxxxxxxxxxxxxxx
xxxxxxxxxxxxxxxxxxxxxxxxxxxxxxxxxxxxxxxxxxxxxxxxxxxxxxxxxxxx
xxxxxxxxxxxxxxxxxxxxxxxxxxxxxxxxxxxxxxxxxxxxxxxxxxxxxxxxxxxx
xxxxxxxxxxxxxxxxxxxxxxxxxxxxxxxxxxxxxxxxxxxxxxxxxxxxxxxxxxxx
xxxxxxxxxxxxxxxxxxxxxxxxxxxxxxxxxxxxxxxxxxxxxxxxxxxxxxxxxxxx
xxxxxxxxxxxxxxxxxxxxxxxxxxxxxxxxxxxxxxxxxxxxxxxxxxxxxxxxxxxx
xxxxxxxxxxxxxxxxxxxxxxxxxxxxxxxxxxxxxxxxxxxxxxxxxxxxxxxxxxxx
xxxxxxxxxxxxxxxxxxxxxxxxxxxxxxxxxxxxxxxxxxxxxxxxxxxxxxxxxxxx
xxxxxxxxxxxxxxxxxxxxxxxxxxxxxxxxxxxxxxxxxxxxxxxxxxxxxxxxxxxx
xxxxxxxxxxxxxxxxxxxxxxxxxxxxxxxxxxxxxxxxxxxxxxxxxxxxxxxxxxxx
xxxxxxxxxxxxxxxxxxxxxxxxxxxxxxxxxxxxxxxxxxxxxxxxxxxxxxxxxxxx
xxxxxxxxxxxxxxxxxxxxxxxxxxxxxxxxxxxxxxxxxxxxxxxxxxxxxxxxxxxx
xxxxxxxxxxxxxxxxxxxxxxxxxxxxxxxxxxxxxxxxxxxxxxxxxxxxxxxxxxxx
xxxxxxxxxxxxxxxxxxxxxxxxxxxxxxxxxxxxxxxxxxxxxxxxxxxxxxxxxxxx
xxxxxxxxxxxxxxxxxxxxxxxxxxxxxxxxxxxxxxxxxxxxxxxxxxxxxxxxxxxx
xxxxxxxxxxxxxxxxxxxxxxxxxxxxxxxxxxxxxxxxxxxxxxxxxxxxxxxxxxxx
xxxxxxxxxxxxxxxxxxxxxxxxxxxxxxxxxxxxxxxxxxxxxxxxxxxxxxxxxxxx
xxxxxxxxxxxxxxxxxxxxxxxxxxxxxxxxxxxxxxxxxxxxxxxxxxxxxxxxxxxx
xxxxxxxxxxxxxxxxxxxxxxxxxxxxxxxxxxxxxxxxxxxxxxxxxxxxxxxxxxxx
xxxxxxxxxxxxxxxxxxxxxxxxxxxxxxxxxxxxxxxxxxxxxxxxxxxxxxxxxxxx
xxxxxxxxxxxxxxxxxxxxxxxxxxxxxxxxxxxxxxxxxxxxxxxxxxxxxxxxxxxx
xxxxxxxxxxxxxxxxxxxxxxxxxxxxxxxxxxxxxxxxxxxxxxxxxxxxxxxxxxxx
xxxxxxxxxxxxxxxxxxxxxxxxxxxxxxxxxxxxxxxxxxxxxxxxxxxxxxxxxxxx
xxxxxxxxxxxxxxxxxxxxxxxxxxxxxxxxxxxxxxxxxxxxxxxxxxxxxxxxxxxx
xxxxxxxxxxxxxxxxxxxxxxxxxxxxxxxxxxxxxxxxxxxxxxxxxxxxxxxxxxxx
xxxxxxxxxxxxxxxxxxxxxxxxxxxxxxxxxxxxxxxxxxxxxxxxxxxxxxxxxxxx
xxxxxxxxxxxxxxxxxxxxxxxxxxxxxxxxxxxxxxxxxxxxxxxxxxxxxxxxxxxx
xxxxxxxxxxxxxxxxxxxxxxxxxxxxxxxxxxxxxxx111111111111111111111

11111111111111111111111111111111111111xxxxxxxxxxxxxxxxxxxxxx
xxxxxxxxxxxxxxxxxxxxxxxxxxxxxxxxxxxxxxxxxxxxxxxxxxxxxxxxxxxx
xxxxxxxxxxxxxxxxxxxxxxxxxxxxxxxxxxxxxxxxxxxxxxxxxxxxxxxxxxxx
xxxxxxxxxxxxxxxxxxxxxxxxxxxxxxxxxxxxxxxxxxxxxxxxxxxxxxxxxxxx
xxxxxxxxxxxxxxxxxxxxxxxxxxxxxxxxxxxxxxxxxxxxxxxxxxxxxxx
>DP02726
MVLSHAVSESDVSVHSTFASRYVRTSLPRFKMPENSIPKEAAYQIINDELMLDGNPRLNLAS
FVTTWMEPECDKLIMSSINKNYVDMDEYPVTTELQNRCVNMIAHLFNAPLEEAETAVGVGTV
GSSEAIMLAGLAFKRKWQNKRKAEGKPVDKPNIVTGANVQVCWEKFARYFEVELKEVKLSEG
YYVMDPQQAVDMVDENTICVAAILGSTLNGEFEDVKLLNDLLVEKNKETGWDTPIHVDAASG
GFIAPFLYPELEWDFRLPLVKSINVSGHKYGLVYAGIGWVIWRNKEDLPEELIFHINYLGAD
QPTFTLNFSKGSSQVIAQYYQLIRLGHEGYRNVMENCRENMIVLREGLEKTERFNIVSKDEG
VPLVAFSLKDSSCHTEFEISDMLRRYGWIVPAYTMPPNAQHITVLRVVIREDFSRTLAERLV
IDIEKVMRELDELPSRVIHKISLGQEKSESNSDNLMVTVKKSDIDKQRDIITGWKKFVADRK
KTSGIC
0000000000000000000000000000000000000000000000000000000xxxxx
xxxxxxxxxxxxxxxxxxxxxxxxxxxxxxxxxxxxxxxxxxxxxxxxxxxxxxxxxxxx
xxxxxxxxxxxxxxxxxxxxxxxxxxxxxxxxxxxxxxxxxxxxxxxxxxxxxxxxxxxx
xxxxxxxxxxxxxxxxxxxxxxxxxxxxxxxxxxxxxxxxxxxxxxxxxxxxxxxxxxxx
xxxxxxxxxxxxxxxxxxxxxxxxxxxxxxxxxxxxxxxxxxxxxxxxxxxxxxxxxxxx
xxxxxxxxxxxxxxxxxxxxxxxxxxxxxxxxxxxxxxxxxxxxxxxxxxxxxxxxxxxx
xxxxxxxxxxxxxxxxxxxxxxxxxxxxxxxxxxxxxxxxxxxxxxxxxxxxxxxxxxxx
xxxxxxxxxxxxx11111111111111111111000000000000000000000000000
000000
>DP02730
MRRFLRPGHDPVRERLKRDLFQFNKTVEHGFPHQPSALGYSPSLRILAIGTRSGAIKLYGAP
GVEFMGLHQENNAVTQIHLLPGQCQLVTLLDDNSLHLWSLKVKGGASELQEDESFTLRGPPG
AAPSATQITVVLPHSSCELLYLGTESGNVFVVQLPAFRALEDRTISSDAVLQRLPEEARHRR
VFEMVEALQEHPRDPNQILIGYSRGLVVIWDLQGSRVLYHFLSSQQLENIWWQRDGRLLVSC
HSDGSYCQWPVSSEAQQPEPLRSLVPYGPFPCKAITRILWLTTRQGLPFTIFQGGMPRASYG
DRHCISVIHDGQQTAFDFTSRVIGFTVLTEADPAATFDDPYALVVLAEEELVVIDLQTAGWP
PVQLPYLASLHCSAITCSHHVSNIPLKLWERIIAAGSRQNAHFSTMEWPIDGGTSLTPAPPQ
RDLLLTGHEDGTVRFWDASGVCLRLLYKLSTVRVFLTDTDPNENFSAQGEDEWPPLRKVGSF
DPYSDDPRLGIQKIFLCKYSGYLAVAGTAGQVLVLELNDEAAEQAVEQVEADLLQDQEGYRW
KGHERLAARSGPVRFEPGFQPFVLVQCQPPAVVTSLALHSEWRLVAFGTSHGFGLFDHQQRR
QVFVKCTLHPSDQLALEGPLSRVKSLKKSLRQSFRRMRRSRVSSRKRHPAGPPGEAQEGSAK
AERPGLQNMELAPVQRKIEARSAEDSFTGFVRTLYFADTYLKDSSRHCPSLWAGTNGGTIYA
FSLRVPPAERRMDEPVRAEQAKEIQLMHRAPVVGILVLDGHSVPLPEPLEVAHDLSKSPDMQ
GSHQLLVVSEEQFKVFTLPKVSAKLKLKLTALEGSRVRRVSVAHFGSRRAEDYGEHHLAVLT
NLGDIQVVSLPLLKPQVRYSCIRREDVSGIASCVFTKYGQGFYLISPSEFERFSLSTKWLVE
PRCLVDSAETKNHRPGNGAGPKKAPSRARNSGTQSDGEEKQPGLVMERALLSDERVLKEIQS
TLEGDRGSGNWRSHRAAVGCSLSNGGAE
xxxxxxxxxxxxxxxxxxxxxxxxxxxxxxxxxxxxxxxxxxxxxxxxxxxxxxxxxxxx
xxxxxxxxxxxxxxxxxxxxxxxxxxxxxxxxxxxxxxxxxxxxxxxxxxxxxxxxxxxx

xxxxxxxxxxxxxxxxxxxxxxxxxxxxxxxxxxxxxxxxxxxxxxxxxxxxxxxxxxxxxx
xxxxxxxxxxxxxxxxxxxxxxxxxxxxxxxxxxxxxxxxxxxxxxxxxxxxxxxxxxxxxx
xxxxxxxxxxxxxxxxxxxxxxxxxxxxxxxxxxxxxxxxxxxxxxxxxxxxxxxxxxxxxx
xxxxxxxxxxxxxxxxxxxxxxxxxxxxxxxxxxxxxxxxxxxxxxxxxxxxxxxxxxxxxx
xxxxxxxxxxxxxxxxxxxxxxxxxxxxxxxxxxxxxxxxxxxxxxxxxxxxxxxxxxxxxx
xxxxxxxxxxxxxxxxxxxxxxxxxxxxxxxxxxxx11111111111111xxxxxxxxxxxx
xxxxxxxxxxxxxxxxxxxxxxxxxxxxxxxxxxxxxxxxxxxxxxxxxxxxxxxxxxxxxx
xxxxxxxxxxxxxxxxxxxxxxxxxxxxxxxxxxxxxxxxxxxxxxxxxxxxxxxxxxxxxx
xxxxxxxxxx1111111111111111111111111111111111111111111111111111
11111111111111111111111111xxxxxxxxxxxxxxxxxxxxxxxxxxxxxxxxxxxx
xxxxxxxxxxxxxxxxxxxxxxxxxxxxxxxxxxxxxxxxxxxxxxxxxxxxxxxxxxxxxx
xxxxxxxxxxxxxxxxxxxxxxxxxxxxxxxxxxxxxxxxxxxxxxxxxxxxxxxxxxxxxx
xxxxxxxxxxxxxxxxxxxxxxxxxxxxxxxxxxxxxxxxxxxxxxxxxxxxxxxxxxxxxx
xxxxxxxxxxxxxxxxxxxxxxxxxxxxxxxxxxxxxxxxxxxxxxxxxxxxxxxxxxxxxx
xxxxxxxxxxxxxxxxxxxxxxxxxxxx
>DP02761
MNRQSGVNAGVQNNPPSRVVYLGSIPYDQTEEQILDLCSNVGPVINLKMMFDPQTGRSKGYA
FIEFRDLESSASAVRNLNGYQLGSRFLKCGYSSNSDISGVSQQQQQQYNNINGNNNNNGNNN
NNSNGPDFQNSGNANFLSQKFPELPSGIDVNINMTTPAMMISSELAKKPKEVQLKFLQKFQE
WTRAHPEDAVSLLELCPQLSFVTAELLLTNGICKVDDLIPLASRPQEEASATNNNSVNEVVD
PAVLNKQKELLKQVLQLNDSQISILPDDERMAIWDLKQKALRGEFGAF
xxxxxxxxxxxxxxxxxxxxxxxxxxxxxxxxxxxxxxxxxxxxxxxxxxxxxxxxxxxxxx
xxxxxxxxxxxxxxxxxxxxxxxxxxxxxxxxxxxxxxxxxxxxxxxxxxxxxxxxxxxxxx
xx000000000000000000000000000000000000000000000000000000000000
0000000000000000000000000000000000000000000000xxxxxxxxxxxxxxxx
xxxxxxxxxxxxxxxxxxxxxxxxxxxxxxxxxxxxxxxxxxxxxxxx
>DP02764
MEALPLLAATTPDHGRHRRLLLLPLLLFLLPAGAVQGWETEERPRTREEECHFYAGGQVYPG
EASRVSVADHSLHLSKAKISKPAPYWEGTAVIDGEFKELKLTDYRGKYLVFFFYPLDFTFVC
PTEIIAFGDRLEEFRSINTEVVACSVDSQFTHLAWINTPRRQGGLGPIRIPLLSDLTHQISK
DYGVYLEDSGHTLRGLFIIDDKGILRQITLNDLPVGRSVDETLRLVQAFQYTDKHGEVCPAG
WKPGSETIIPDPAGKLKYFDKLN
00000000000000000000000000000000000000000000000000000000000000
00000000000000000000000000000000000000000000000000000000000000
00000000000000000000000000000000000000000000000000000000000000
00000000000000000000000000000000000000000000000000000000000000
00000000000000000000000
>DP02782
MESLLQHLDRFSELLAVSSTTYVSTWDPATVRRALQWARYLRHIHRRFGRHGPIRTALERRL
HNQWRQEGGFGRGPVPGLANFQALGHCDVLLSLRLLENRALGDAARYHLVQQLFPGPGVRDA
DEETLQESLARLARRRSAVHMLRFNGYRENPNLQEDSLMKTQAELLLERLQEVGKAEAERPA
RFLSSLWERLPQNNFLKVIAVALLQPPLSRRPQEELEPGIHKSPGEGSQVLVHWLLGNSEVF
AAFCRALPAGLLTLVTSRHPALSPVYLGLLTDWGQRLHYDLQKGIWVGTESQDVPWEELHNR
FQSLCQAPPPLKDKVLTALETCKAQDGDFEVPGLSIWTDLLLALRSGAFRKRQVLGLSAGLS

SV

xxxxxxxxxxxxxxxxxxxxxxxxxxxxxxxxxxxxxxxxxxxxxxxxxxxxxxxxxxxx
xxxxxxxxxxxxxxxxxxxxxxxxxxxxxxxxxxxxxxxxxxxxxxxxxxxxxxxxxxxx
xxxxxxxxxxxxxxxxxxxxxxxxxxxxxx000000000000000000000000000000
000000000000000000000000000000000000000000000000000000000000
000000000000000000000000000000000000000000000000000000000000
000000000000000000000000000000000000000000000xxxxxxxxxxxxxxx
xx

>DP02787

MKSLFKVTLLATTMAVALHAPITFAAEAAKPATAADSKAAFKNDDQKSAYALGASLGRYMEN
SLKEQEKLGIKLDKDQLIAGVQDAFADKSKLSDQEIEQTLQAFEARVKSSAQAKMEKDAADN
EAKGKEYREKFAKEKGVKTSSTGLVYQVVEAGKGEAPKDSDTVVVNYKGTLIDGKEFDNSYT
RGEPLSFRLDGVIPGWTEGLKNIKKGGKIKLVIPPELAYGKAGVPGIPPNSTLVFDVELLDV
KPAPKADAKPEADAKAADSAKK

xxxxxxxxxxxxxx0000000000000000000000000000000000000000000000
00000000000000000000000000000000000000000000000000xxxxxxxxxx
xxxxxxxxxxxxxxxxxxxxxxxxxxxxxxxxxxxxxxxxxxxxxxxxxxxxxxxxxxxx
xxxxxxxxxxxxxxxxxxxxxxxxxxxxxxxxxxxxxxxxxxxxxxxxxxxxxxxxxxxx
xxxx000000000000000000

>DP02808

MVKSTSKTSTKETVTKQPTEEKPIQEKEELALETSSSSSDEEDEKDEDEIEGLAASDDEQSG
THKIKRLNPKKQANEKKSKDKKTLEEYSGIIYVSRLPHGFHEKELSKYFAQFGDLKEVRLAR
NKKTGNSRHYGFLEFVNKEDAMIAQESMNNYLLMGHLLQVRVLPKGAKIEKLYKYKKRVLVE
KGITKPVKQLKDNMKQKHEERIKKLAKSGIEFKW

000000000000000000000000000000000000000000000000000000000000
0000000000000000000000000xxxxxxxxxxxxxxxxxxxxxxxxxxxxxxxxxxx
xxxxxxxxxxxxxxxxxxxxxxxxxxxxxxxxxxxxxxxxxxxxxxxxxxxx00000000
0000000000000000000000000000000000

>DP02820

MQVLTKRYPKNCLLTVMDRYSAVVRNMEQVVMIPSLLRDVQLSGPGGSVQDGAPDLYTYFTM
LKSICVEVDHGLLPREEWQAKVAGNETSEAENDAAETEEAEEDRISEELDLEAQFHLHFCSL
HHILTHLTRKAQEVTRKYQEMTGQVL

000000000000000000000000000000000000000000000000000000000000
000000000000000000000000000000000000000000000000000000000000
00000000000000000000000000

>DP02826

MIRSTLSSWREYLTPITHKSTFLTTGQITPEEFVQAGDYLCHMFPTWKWNEESSDISYRDFL
PKNKQFLIIRKVPCDKRAEQCVEVEGPDVIMKGFAEDGDEDDVLEYIGSETEHVQSTPAGGT
KDSSIDDIDELIQDMEIKEEDENDDTEEFNAKGGLAKDMAQERYYDLYIAYSTSYRVPKMYI
VGFNSNGSPLSPEQMFEDISADYRTKTATIEKLPFYKNSVLSVSIHPCKHANVMKILLDKVR
VVRQRRRKELQEEQELDGVGDWEDLQDDIDDSLRVDQYLIVFLKFITSVTPSIQHDYTMEGW

00000000000000000000xxxxxxxxxxxxxxxxxxxxxxxxxxxxxxxxxxxxxxxx
xxxxxxxxxxxxxxxxxxxxx000000000000000000000000000000000000000
000000000000000000000000000000000000xxxxxxxxxxxxxxxxxxxxxxxx

xxxxxxxxxxxxxxxxxxxxxxxxxxxxxxxxxxxxxxxxxxxxxxxxxxxxxxxxxxxx00
00000000000000000000000000000000xxxxxxxxxxxxxxxxxxxxxxxxxxxxxx
>DP02839
MWIIEAEGDILKGKSRILFPGTYIVGRNVSDDSSHIQVISKSISKRHARFTILTPSEKDYFT
GGPCEFEVKDLDTKFGTKVNEKVVGQNGDSYKEKDLKIQLGKCPFTINAYWRSMCIQFDNPE
MLSQWASNLNLLGIPTGLRDSDATTHFVMNRQAGSSITVGTMYAFLKKTVIIDDSYLQYLST
VKESVIEDASLMPDALECFKNIIKNNDQFPSSPEDCINSLEGFSCAMLNTSSESHHLLELLG
LRISTFMSLGDIDKELISKTDFVVLNNAVYDSEKISFPEGIFCLTIEQLWKIIIERNSRELI
SKEIERLKYATASNSTPQKIIQPQRHIQKNIVDDLFSVKKPLPCSPKSKRVKTLENLSIMDF
VQPKQMFGKEPEGYLSNQSNNGSAQNKKSGDNSEKTKNSLKSSSKKSANTGSGQGKTKVEYV
SYNSVDKGNSSPFKPLELNVVGEKKANAEVDSLPSENVQESEDDKAFEENRRLRNLGSVEYI
RIMSSEKSNANSRHTSKYYSGRKNFKKFQKKASQKAPLQAFLSLSEHKKTEVFDQDDTDLEP
VPRLMSKVESIPAGASSDKSGKSSISKKSSNSFKELSPKTNNDEDDEFNDLKFHF
xxxxxxxxxxxxxxxxxxxxxxxxxxxxxxxxxxxxxxxxxxxxxxxxxxxxxxxxxxxxxx
xxxxxxxxxxxxxxxxxxxxxxxxxxxxxxxxxxxxxxxxxxxxxxxxxxxxxxxxxxxxxx
xxxxxxxxxxxxxxxxxxxxxxxxxxxxxxxxxxxxxxxxxxxxxxxxxxxxxxxxxxxxxx
xxxxxxxxxxxxxxxxxxxxxxxxxxxxxxxxxxxxxxxxxxxxxxxxxxxxxxxxxxxxxx
xxxxxxxxxxxxxxxxxxxxxxxxxx000000000000xxxxxxxxxxxxxxxxxxxxxxxx
xxxxxxxxxxxxxxxxxxx0000000000000000000000000000000000000000000
00000000000000000000000000000000000000000000000000000000000000
00000000000000000000000000000000000000000000000000000000000000
000000000000000000000000000000000000000000000000000000000xxxxx
xxxxxxxxxxxxxxxxxxxxxxxxxxxxxxxxxxxxxxxxxxxxxxxxxxxxxxx
>DP02847
MDQKASYFINEKLFTEVKPVLFTDLIHHLKIGPSMAKKLMFDYYKQTTNAKYNCVVICCYKD
QTIKIIHDLSNIPQQDSIIDCFIYAFNPMDSFIPYYDIIDQKDCLTIKNSYELKVSESSKII
ERTKTLEEKSKPLVRPTARSKTTPEETTGRKSKSKDMGLRSTALLAKMKKDRDDKETSRQNE
LRKRKEENLQKINKQNPEREAQMKELNNLFVEDDLDTEEVNGGSKPNSPKETDSNDKDKNND
DLEDLLETTAEDSLMDVPKIQQTKPSETEHSKEPKSEEEPSSFIDEDGYIVTKRPATSTPPR
KPSPVVKRALSSSKKQETPSSNKRLKKQGTLESFFKRKAK
xxxxxxxxxxxxxxxxxxxxxxxxxxxxxxxxxxxxxxxxxxxxxxxxxxxxxxxxxxxxxx
xxxxxxxxxxxxxxxxxxxxxxxxxxxxxxxxxxxxxxxxxxxxxxxxxxxxxx00000000
00000000000000000000000000000000000000000000000000000000000000
00000000000000000000000000000000000000000000000000000000000000
00000000000000000000000000000000000000000000000000000000000000
0000000000000000000000000000000000000000
>DP02853
MQPSLAPCLICLLVHAAFCAVEGQGWQAFRNDATEVIPGLGEYPEPPPENNQTMNRAENGGR
PPHHPYDAKDVSEYSCRELHYTRFLTDGPCRSAKPVTELVCSGQCGPARLLPNAIGRVKWWR
PNGPDFRCIPDRYRAQRVQLLCPGGAAPRSRKVRLVASCKCKRLTRFHNQSELKDFGPETAR
PQKGRKPRPGARGAKANQAELENAY
000000000000000000000000000000000000000000000000000xxxxxxxxxxx
xxxxxxxxxxxxxxxxxxxxxxxxxxxxxxxxxxxxxxxxxxxxxxxxxxxxxxxxxxxxxx
xxxxxxxxxxxxxxxxxx00000000000000000000000000000000000000000000

0000000000000000000000000
>DP02869
MADGVDHIDIYADVGEEFNQEAEYGGHDQIDLYDDVISPSANNGDAPEDRDYMDTLPPTVGD
DVGKGAAPNVVYTYTGKRIALYIGNLTWWTTDEDLTEAVHSLGVNDILEIKFFENRANGQSK
GFALVGVGSEASSKKLMDLLPKRELHGQNPVVTPCNKQFLSQFEMQSRKTTQSGQMSGEGKA
GPPGGSSRAAFPQGGRGRGRFPGAVPGGDRFPGPAGPGGPPPPFPAGQTPPRPPLGPPGPPG
PPGPPPPGQVLPPPLAGPPNRGDRPPPPVLFPGQPFGQPPLGPLPPGPPPPVPGYGPPPGPP
PPQQGPPPPPGPFPPRPPGPLGPPLTLAPPPHLPGPPPGAPPPAPHVNPAFFPPPTNSGMPT
SDSRGPPPTDPYGRPPPYDRGDYGPPGREMDTARTPLSEAEFEEIMNRNRAISSSAISRAVS
DASAGDYGSAIETLVTAISLIKQSKVSADDRCKVLISSLQDCLHGIESKSYGSGSRRERSRE
RDHSRSREKSRRHKSRSRDRHDDYYRERSRERERHRDRDRDRDRERDREREYRHR
xxxxxxxxxxxx00000000000000000000000000000000000000000000000000
000000000000000000xxxxxxxxxxxxxxxxxxxxxxxxxxxxxxxxxxxxxxxxxxxx
xxxxxxxxxxxxxxxxxxxxxxxxxxxxxxxxxxxxxxxxxxxxxxxx00000000000000
0000000000000000000000000000000000000000000000000xxxxxxxxxxxxx
xxxxxxxxxxxxxxxxxxxxxxxxxxxxxxxxxxxxxxxxxxxxxxxxxxxxxxxxxxxxxx
xxxxxxxxxxxxxxxxxxxxxxxxxxxxxxxxxxxxxxxxxxxxxxxxxxxxxxxxxxxxxx
xxxxxxxxxxxxxxxxxxxxxxxxxxxxxxxxxxxxxxxxxxxxxxxxxxxxxxxxxxxxxx
xxxxxxxxxxxxxxxxxxxxxxxxxxxxxxxxxxxxxxxxxxxxxx0000000000000000
0000000000000000000000000000000000000000000000000000000
>DP02885
MASTSYDYCRVPMEDGDKRCKLLLGIGILVLLIIVILGVPLIIFTIKANSEACRDGLRAVME
CRNVTHLLQQELTEAQKGFQDVEAQAATCNHTVMALMASLDAEKAQGQKKVEELEGEITTLN
HKLQDASAEVERLRRENQVLSVRIADKKYYPSSQDSSSAAAPQLLIVLLGLSALLQ
xxxxxxxxxxxxxxxxxxxxxxxxxxxxxxxxxxxxxxxxxxxxxxxxxxxxxxxxxxxxxx
xxxxxxxxxxxxxxxx11111111111xxxxxxxxxxxxxxxxxxxxxxxxxxxxxxxxxxx
xxxxxxxxxxxxxxxxxxxxxxxxxxxxxxxxxxxxxxxxxxxxxxxxxxxxxxxx
>DP02900
MEKGTVKWFNNAKGFGFICPEGGGEDIFAHYSTIQMDGYRTLKAGQSVQFDVHQGPKGNHAS
VIVPVEVEAAVA
00000000000000000000000000000000000000000000000000000000000000
000000000000
>DP02901
MSAVKKQRIDLRLTDDDKSMIEEAAAISNQSVSQFMLNSASQRAAEVIEQHRRVILNEESWT
RVMDALSNPPSPGEKLKRAAKRLQGM
xxxxxxxxxxxxxxxxxxxxxxxxxxxxxxxxxxxxxxxxxxx0000000000000000000
00000000000000000000000000
>DP02903
MKEQEFSYREAKDVSLDSKGLENSFLSSPNREKTPLFFEGNSNETSGYDQTKNFTHGDGDMS
LGNLSELNVATDLLESLDLRSMYMHGYGHLDSSFSSQHSPDNRKRMSSTSVFKRINSEEEGR
IPSLTYSAGTMNSTSSSTASLKGADIVADYETFNPDQNLAELSFDRSKSSRKRAVEVAEFSR
AKTMSPLEYTVQHPYQSHNELSTNPARARAGSVPNLARIPSDVKPVPPAHLSASSTVGPRIL
PSLPKDTTEDNPALERVETTASLDMDYKPLEPLAPIQEAPVEDTSEPFSSVPEATLDDSDIS
TESLRKKVLAKMEAKRISSGSSYASTLRKVYDFSELSLPTNGKDYDELYLQSSRNSEPEIST

IINDSLQQENMDEDISATSIPKSQAAYGHGSVTYHEVPRYNLTSASVGYSISSQRGRIKSSS
TIDNLSAILSSEDLRHPSMQPVPGTKRTYSNYCENEPNKSSQSLVSSESHNVEGWNYSETGT
VGFYDPSAEISASIDELRQSTPVARDSELLSRAHSFDLNRLDLPSQDKSTSYEVPNGTENQS
PRPVTSLGFVNETFFEEKPKAPLPLGRFYIHLNSILNISISEVHSPIKIIVNTPTQNMQLPW
QAVNGNNRLDHDFAFHVDDNFKVSFMFLDIPIEDKSNGSKGVSATKDVSNGKPAETKSKARK
FFDKLFNRRKKRKLNKAAAVENSKAKKSVVIKKVSGTATLNLGNVKDSCFGKAFNVEIPIIS
RGFLEAIPVKINSIGKRTLGNLTLTCLYIPELSVPEQELPFTLEQATMDLRHVRSNYLYNEG
YLYRLEDSSIRRRFVVLRSKQLNFYAEKGGQYLDTFQLSKTVVSIPMVNFSEAVSNLGLVAG
ILATSVDRRHVQLFADSKKVCQKWLQVMNSRSFALDRGTEKLWLQEYVNFMA
00000000000000000000000000000000000000000000000000000000000000
00000000000000000000000000000000000000000000000000000000000000
00000000000000000000000000000000000000000000000000000000000000
00000000000000000000000000000000000000000000000000000000000000
00000000000000000000000000000000000000000000000000000000000000
00000000000000000000000000000000000000000000000000000000000000
00000000000000000000000000000000000000000000000000000000000000
000000000000000000xxxxxxxxxxxxxxxxxxxxxxxxxxxxxxxxxxxxxxxxxxxx
xxxxxxxxxxxxxxxxxxxxxxxxxxxxxxxxxxxxxxxxxxxxxxxxxxxxxxxxxxxxxx
xxxxxxxxxxxxxxxxxxxxxxxxxxxxxxxxxxxxxxxxxxxxxxxxxxxxxxxxxxxxxx
xxxxxxxxxxxxxxxxxxxxxxxxxxxxxxxxxxxxxxxxxxxxxxxxxxxxxxxxxxxxxx
xxxxxxxxxxxxxxxxxxxxxxxxxxxxxxxxxxxxxxxxxxxxxxxxxxxxxxxxxxxxxx
xxxxxxxxxxxxxxxxxxxxxxxxxxxxxxxxxxxxxxxxxxxxxxxxxxxxxxxxxxxxxx
xxxxxxxxxxxxxxxxxxxxxxxxxxxxxxxxxxxxxxxxxxxxxxxxxxxxxxxxxxxxxx
xxxxxxxxxxxxxxxxxxxxxxxxxxxxxxxxxxxxxxxxxxxxxxxxxxxx
>DP02917
MSKFSLKLGSKTLKKNISKKTKKKNSLQKANLFDWDDAETASLSHKPQSKIKIQSIDKFDLD
EESSSKKKLVIKLSENADTKKNDAPLVEYVTEKEYNEVPVEEFGDALLRGMGWESDSEQDSK
GDKTQSRNKDVSNVSQIHPDGLGIGAKLNKAINVEEASFMPVVKIDKITGTKVDDDKKNKS
xxxxxxxxx00000000000000000000000000000000000000000000000000000
00000000000000000000000000000000000000000000000000000000000000
0000000000000000000000000000000000000000000000000000000000000
>DP02920
MQSLSTPHTISLLLPRTSPSRLSPSLHSLAFPTRLRSLSYSSQTSILPDAGDDFIVGDCLVY
EDGVFEDPYLDKEVTQVAKQERKKNRRGGAKRLDESEIEPENLVPEEWRDIQAEVNLTKKDK
RKIAQEMEFGVRVEKKRQGLIPLRKVDLNDFLTYKEAKLAQLRPVILDKPGNFSDDSGASSD
GETAVSSPSERVAPKNPRWAVYGKGFDHVAKFFNSDKYDPSDKKSDGPRKLLSKEEKFMLNS
RNPDLAVATSKKWLPLHTLAACGEFYLVDSLLKHNLDINATDVGGLTVLHRAIIGKKQAITN
YLLRESANPFVLDDEGATLMHYAVQTASAPTIKLLLLYNADINAQDRDGWTPLHVAVQARRS
DIVKLLLIKGADIEVKNKDGLTPLGLCLYLGREIRTYEVMKLLKEFPLSRHKKRLVTTDEDI
E
xxxxxxxxxxxxxxxxxxxxxxxxxxxxxxxxxxxx00000000000000000000000000
00000000000000000000000000000000000000000000000000000000000000
00000000000000000000000000000000000000000000000000000000000000
000000000000000000000000000000000000000000000xxxxxxxxxxxxxxxxx

xxxxxxxxxxxxxxxxxxxxxxxxxxxxxxxxxxxxxxxxxxxxxxxxxxxxxxxxxxxx
xxxxxxxxxxxxxxxxxxxxxxxxxxxxxxxxxxxxxxxxxxxxxxxxxxxxxxxxxxxx
xxxxxxxxxxxxxxxxxxxxxxxxxxxxxxxxxxxxxxxxxxxxxxxxxxxxxxxxxxxx
x
>DP02923
MFFNTKHTTALCFVTCMAFSSSSIADIVISGTRVIYKSDQKSVNVRLENKGNNPLLVQSWLD
TGDDNAEPGSITVPFTATPPVSRIDAKRGQTIKLMYTASTSLPKDRESVFWFNVLEVPPKPD
AEKVANQSLLQLAFRTRIKLFYRPDGLKGNPSEAPLALKWFWSGSEGKASLRVTNPTPYYVS
FSSGDLEASGKRYPIDVKMIAPFSDEVMKVNGLNGKANSAKVHFYAINDFGGAIEGNARL
xxxxxxxxxxxxxxxxxxxxxxxxxxxxxxxxxxxxxxxxxxxxxxxxxxxxxxxxxxxx
xxxxxxxxxxxxxxxxxxxxxxxxxxxxxxxxxxxxxxxxxxxxxxxxxxxxxxxxx111
11111111xxxxxxxxxxxxxxxxxxxxxxxxxxxxxxxxxxxxxxxxxxxxxxxxxxxx
xxxxxxxxxxxxxxxxxxxxxxxxxxxxxxxxxxxxxxxxxxxxxxxxxxxxxxxxxx
>DP02925
MESLVPGFNEKTHVQLSLPVLQVRDVLVRGFGDSVEEVLSEARQHLKDGTCGLVEVEKGVLP
QLEQPYVFIKRSDARTAPHGHVMVELVAELEGIQYGRSGETLGVLVPHVGEIPVAYRKVLLR
KNGNKGAGGHSYGADLKSFDLGDELGTDPYEDFQENWNTKHSSGVTRELMRELNGGAYTRYV
DNNFCGPDGYPLECIKDLLARAGKASCTLSEQLDFIDTKRGVYCCREHEHEIAWYTERSEKS
YELQTPFEIKLAKKFDTFNGECPNFVFPLNSIIKTIQPRVEKKKLDGFMGRIRSVYPVASPN
ECNQMCLSTLMKCDHCGETSWQTGDFVKATCEFCGTENLTKEGATTCGYLPQNAVVKIYCPA
CHNSEVGPEHSLAEYHNESGLKTILRKGGRTIAFGGCVFSYVGCHNKCAYWVPRASANIGCN
HTGVVGEGSEGLNDNLLEILQKEKVNINIVGDFKLNEEIAIILASFSASTSAFVETVKGLDY
KAFKQIVESCGNFKVTKGKAKKGAWNIGEQKSILSPLYAFASEAARVVRSIFSRTLETAQNS
VRVLQKAAITILDGISQYSLRLIDAMMFTSDLATNNLVVMAYITGGVVQLTSQWLTNIFGTV
YEKLKPVLDWLEEKFKEGVEFLRDGWEIVKFISTCACEIVGGQIVTCAKEIKESVQTFFKLV
NKFLALCADSIIIGGAKLKALNLGETFVTHSKGLYRKCVKSREETGLLMPLKAPKEIIFLEG
ETLPTEVLTEEVVLKTGDLQPLEQPTSEAVEAPLVGTPVCINGLMLLEIKDTEKYCALAPNM
MVTNNTFTLKGGAPTKVTFGDDTVIEVQGYKSVNITFELDERIDKVLNEKCSAYTVELGTEV
NEFACVVADAVIKTLQPVSELLTPLGIDLDEWSMATYYLFDESGEFKLASHMYCSFYPPDED
EEEGDCEEEEFEPSTQYEYGTEDDYQGKPLEFGATSAALQPEEEQEEDWLDDDSQQTVGQQD
GSEDNQTTTIQTIVEVQPQLEMELTPVVQTIEVNSFSGYLKLTDNVYIKNADIVEEAKKVKP
TVVVNAANVYLKHGGGVAGALNKATNNAMQVESDDYIATNGPLKVGGSCVLSGHNLAKHCLH
VVGPNVNKGEDIQLLKSAYENFNQHEVLLAPLLSAGIFGADPIHSLRVCVDTVRTNVYLAVF
DKNLYDKLVSSFLEMKSEKQVEQKIAEIPKEEVKPFITESKPSVEQRKQDDKKIKACVEEVT
TTLEETKFLTENLLLYIDINGNLHPDSATLVSDIDITFLKKDAPYIVGDVVQEGVLTAVVIP
TKKAGGTTEMLAKALRKVPTDNYITTYPGQGLNGYTVEEAKTVLKKCKSAFYILPSIISNEK
QEILGTVSWNLREMLAHAEETRKLMPVCVETKAIVSTIQRKYKGIKIQEGVVDYGARFYFYT
SKTTVASLINTLNDLNETLVTMPLGYVTHGLNLEEAARYMRSLKVPATVSVSSPDAVTAYNG
YLTSSSKTPEEHFIETISLAGSYKDWSYSGQSTQLGIEFLKRGDKSVYYTSNPTTFHLDGEV
ITFDNLKTLLSLREVRTIKVFTTVDNINLHTQVVDMSMTYGQQFGPTYLDGADVTKIKPHNS
HEGKTFYVLPNDDTLRVEAFEYYHTTDPSFLGRYMSALNHTKKWKYPQVNGLTSIKWADNNC
YLATALLTLQQIELKFNPPALQDAYYRARAGEAANFCALILAYCNKTVGELGDVRETMSYLF
QHANLDSCKRVLNVVCKTCGQQQTTLKGVEAVMYMGTLSYEQFKKGVQIPCTCGKQATKYLV
QQESPFVMMSAPPAQYELKHGTFTCASEYTGNYQCGHYKHITSKETLYCIDGALLTKSSEYK

GPITDVFYKENSYTTTIKPVTYKLDGVVCTEIDPKLDNYYKKDNSYFTEQPIDLVPNQPYPN
ASFDNFKFVCDNIKFADDLNQLTGYKKPASRELKVTFFPDLNGDVVAIDYKHYTPSFKKGAK
LLHKPIVWHVNNATNKATYKPNTWCIRCLWSTKPVETSNSFDVLKSEDAQGMDNLACEDLKP
VSEEVVENPTIQKDVLECNVKTTEVVGDIILKPANNSLKITEEVGHTDLMAAYVDNSSLTIK
KPNELSRVLGLKTLATHGLAAVNSVPWDTIANYAKPFLNKVVSTTTNIVTRCLNRVCTNYMP
YFFTLLLQLCTFTRSTNSRIKASMPTTIAKNTVKSVGKFCLEASFNYLKSPNFSKLINIIIW
FLLLSVCLGSLIYSTAALGVLMSNLGMPSYCTGYREGYLNSTNVTIATYCTGSIPCSVCLSG
LDSLDTYPSLETIQITISSFKWDLTAFGLVAEWFLAYILFTRFFYVLGLAAIMQLFFSYFAV
HFISNSWLMWLIINLVQMAPISAMVRMYIFFASFYYVWKSYVHVVDGCNSSTCMMCYKRNRA
TRVECTTIVNGVRRSFYVYANGGKGFCKLHNWNCVNCDTFCAGSTFISDEVARDLSLQFKRP
INPTDQSSYIVDSVTVKNGSIHLYFDKAGQKTYERHSLSHFVNLDNLRANNTKGSLPINVIV
FDGKSKCEESSAKSASVYYSQLMCQPILLLDQALVSDVGDSAEVAVKMFDAYVNTFSSTFNV
PMEKLKTLVATAEAELAKNVSLDNVLSTFISAARQGFVDSDVETKDVVECLKLSHQSDIEVT
GDSCNNYMLTYNKVENMTPRDLGACIDCSARHINAQVAKSHNIALIWNVKDFMSLSEQLRKQ
IRSAAKKNNLPFKLTCATTRQVVNVVTTKIALKGGKIVNNWLKQLIKVTLVFLFVAAIFYLI
TPVHVMSKHTDFSSEIIGYKAIDGGVTRDIASTDTCFANKHADFDTWFSQRGGSYTNDKACP
LIAAVITREVGFVVPGLPGTILRTTNGDFLHFLPRVFSAVGNICYTPSKLIEYTDFATSACV
LAAECTIFKDASGKPVPYCYDTNVLEGSVAYESLRPDTRYVLMDGSIIQFPNTYLEGSVRVV
TTFDSEYCRHGTCERSEAGVCVSTSGRWVLNNDYYRSLPGVFCGVDAVNLLTNMFTPLIQPI
GALDISASIVAGGIVAIVVTCLAYYFMRFRRAFGEYSHVVAFNTLLFLMSFTVLCLTPVYSF
LPGVYSVIYLYLTFYLTNDVSFLAHIQWMVMFTPLVPFWITIAYIICISTKHFYWFFSNYLK
RRVVFNGVSFSTFEEAALCTFLLNKEMYLKLRSDVLLPLTQYNRYLALYNKYKYFSGAMDTT
SYREAACCHLAKALNDFSNSGSDVLYQPPQTSITSAVLQSGFRKMAFPSGKVEGCMVQVTCG
TTTLNGLWLDDVVYCPRHVICTSEDMLNPNYEDLLIRKSNHNFLVQAGNVQLRVIGHSMQNC
VLKLKVDTANPKTPKYKFVRIQPGQTFSVLACYNGSPSGVYQCAMRPNFTIKGSFLNGSCGS
VGFNIDYDCVSFCYMHHMELPTGVHAGTDLEGNFYGPFVDRQTAQAAGTDTTITVNVLAWLY
AAVINGDRWFLNRFTTTLNDFNLVAMKYNYEPLTQDHVDILGPLSAQTGIAVLDMCASLKEL
LQNGMNGRTILGSALLEDEFTPFDVVRQCSGVTFQSAVKRTIKGTHHWLLLTILTSLLVLVQ
STQWSLFFFLYENAFLPFAMGIIAMSAFAMMFVKHKHAFLCLFLLPSLATVAYFNMVYMPAS
WVMRIMTWLDMVDTSLSGFKLKDCVMYASAVVLLILMTARTVYDDGARRVWTLMNVLTLVYK
VYYGNALDQAISMWALIISVTSNYSGVVTTVMFLARGIVFMCVEYCPIFFITGNTLQCIMLV
YCFLGYFCTCYFGLFCLLNRYFRLTLGVYDYLVSTQEFRYMNSQGLLPPKNSIDAFKLNIKL
LGVGGKPCIKVATVQSKMSDVKCTSVVLLSVLQQLRVESSSKLWAQCVQLHNDILLAKDTTE
AFEKMVSLLSVLLSMQGAVDINKLCEEMLDNRATLQAIASEFSSLPSYAAFATAQEAYEQAV
ANGDSEVVLKKLKKSLNVAKSEFDRDAAMQRKLEKMADQAMTQMYKQARSEDKRAKVTSAMQ
TMLFTMLRKLDNDALNNIINNARDGCVPLNIIPLTTAAKLMVVIPDYNTYKNTCDGTTFTYA
SALWEIQQVVDADSKIVQLSEISMDNSPNLAWPLIVTALRANSAVKLQNNELSPVALRQMSC
AAGTTQTACTDDNALAYYNTTKGGRFVLALLSDLQDLKWARFPKSDGTGTIYTELEPPCRFV
TDTPKGPKVKYLYFIKGLNNLNRGMVLGSLAATVRLQAGNATEVPANSTVLSFCAFAVDAAK
AYKDYLASGGQPITNCVKMLCTHTGTGQAITVTPEANMDQESFGGASCCLYCRCHIDHPNPK
GFCDLKGKYVQIPTTCANDPVGFTLKNTVCTVCGMWKGYGCSCDQLREPMLQSADAQSFLNR
VCGVSAARLTPCGTGTSTDVVYRAFDIYNDKVAGFAKFLKTNCCRFQEKDEDDNLIDSYFVV
KRHTFSNYQHEETIYNLLKDCPAVAKHDFFKFRIDGDMVPHISRQRLTKYTMADLVYALRHF
DEGNCDTLKEILVTYNCCDDDYFNKKDWYDFVENPDILRVYANLGERVRQALLKTVQFCDAM

RNAGIVGVLTLDNQDLNGNWYDFGDFIQTTPGSGVPVVDSYYSLLMPILTLTRALTAESHVD
TDLTKPYIKWDLLKYDFTEERLKLFDRYFKYWDQTYHPNCVNCLDDRCILHCANFNVLFSTV
FPPTSFGPLVRKIFVDGVPFVVSTGYHFRELGVVHNQDVNLHSSRLSFKELLVYAADPAMHA
ASGNLLLDKRTTCFSVAALTNNVAFQTVKPGNFNKDFYDFAVSKGFFKEGSSVELKHFFFAQ
DGNAAISDYDYYRYNLPTMCDIRQLLFVVEVVDKYFDCYDGGCINANQVIVNNLDKSAGFPF
NKWGKARLYYDSMSYEDQDALFAYTKRNVIPTITQMNLKYAISAKNRARTVAGVSICSTMTN
RQFHQKLLKSIAATRGATVVIGTSKFYGGWHNMLKTVYSDVENPHLMGWDYPKCDRAMPNML
RIMASLVLARKHTTCCSLSHRFYRLANECAQVLSEMVMCGGSLYVKPGGTSSGDATTAYANS
VFNICQAVTANVNALLSTDGNKIADKYVRNLQHRLYECLYRNRDVDTDFVNEFYAYLRKHFS
MMILSDDAVVCFNSTYASQGLVASIKNFKSVLYYQNNVFMSEAKCWTETDLTKGPHEFCSQH
TMLVKQGDDYVYLPYPDPSRILGAGCFVDDIVKTDGTLMIERFVSLAIDAYPLTKHPNQEYA
DVFHLYLQYIRKLHDELTGHMLDMYSVMLTNDNTSRYWEPEFYEAMYTPHTVLQAVGACVLC
NSQTSLRCGACIRRPFLCCKCCYDHVISTSHKLVLSVNPYVCNAPGCDVTDVTQLYLGGMSY
YCKSHKPPISFPLCANGQVFGLYKNTCVGSDNVTDFNAIATCDWTNAGDYILANTCTERLKL
FAAETLKATEETFKLSYGIATVREVLSDRELHLSWEVGKPRPPLNRNYVFTGYRVTKNSKVQ
IGEYTFEKGDYGDAVVYRGTTTYKLNVGDYFVLTSHTVMPLSAPTLVPQEHYVRITGLYPTL
NISDEFSSNVANYQKVGMQKYSTLQGPPGTGKSHFAIGLALYYPSARIVYTACSHAAVDALC
EKALKYLPIDKCSRIIPARARVECFDKFKVNSTLEQYVFCTVNALPETTADIVVFDEISMAT
NYDLSVVNARLRAKHYVYIGDPAQLPAPRTLLTKGTLEPEYFNSVCRLMKTIGPDMFLGTCR
RCPAEIVDTVSALVYDNKLKAHKDKSAQCFKMFYKGVITHDVSSAINRPQIGVVREFLTRNP
AWRKAVFISPYNSQNAVASKILGLPTQTVDSSQGSEYDYVIFTQTTETAHSCNVNRFNVAIT
RAKVGILCIMSDRDLYDKLQFTSLEIPRRNVATLQAENVTGLFKDCSKVITGLHPTQAPTHL
SVDTKFKTEGLCVDIPGIPKDMTYRRLISMMGFKMNYQVNGYPNMFITREEAIRHVRAWIGF
DVEGCHATREAVGTNLPLQLGFSTGVNLVAVPTGYVDTPNNTDFSRVSAKPPPGDQFKHLIP
LMYKGLPWNVVRIKIVQMLSDTLKNLSDRVVFVLWAHGFELTSMKYFVKIGPERTCCLCDRR
ATCFSTASDTYACWHHSIGFDYVVYNPFMIDVQQWGFTGNLQSNHDLYCQVHGNAHVASCDAI
MTRCLAVHECFVKRVDWTIEYPIIGDELKINAACRKVQHMVVKAALLADKFPVLHDIGNPKA
IKCVPQADVEWKFYDAQPCSDKAYKIEELFYSYATHSDKFTDGVCLFWNCNVDRYPANSIVC
RFDTRVLSNLNLPGCDGGSLYVNKHAFHTPAFDKSAFVNLKQLPFFYYSDSPCESHGKQVVS
DIDYVPLKSATCITRCNLGGAVCRHHANEYRLYLDAYNMMISAGFSLWVYKQFDTYNLWNTF
TRLQSLENVAFNVVNKGHFDGQQGEVPVSIINNTVYTKVDGVDVELFENKTTLPVNVAFELW
AKRNIKPVPEVKILNNLGVDIAANTVIWDYKRDAPAHISTIGVCSMTDIAKKPTETICAPLT
VFFDGRVDGQVDLFRNARNGVLITEGSVKGLQPSVGPKQASLNGVTLIGEAVKTQFNYYKKV
DGVVQQLPETYFTQSRNLQEFKPRSQMEIDFLELAMDEFIERYKLEGYAFEHIVYGDFSHSQ
LGGLHLLIGLAKRFKESPFELEDFIPMDSTVKNYFITDAQTGSSKCVCSVIDLLLDDFVEII
KSQDLSVVSKVVKVTIDYTEISFMLWCKDGHVETFYPKLQSSQAWQPGVAMPNLYKMQRMLL
EKCDLQNYGDSATLPKGIMMNVAKYTQLCQYLNTLTLAVPYNMRVIHFGAGSDKGVAPGTAV
LRQWLPTGTLLVDSDLNDFVSDADSTLIGDCATVHTANKWDLIISDMYDPKTKNVTKENDSK
EGFFTYICGFIQQKLALGGSVAIKITEHSWNADLYKLMGHFAWWTAFVTNVNASSSEAFLIG
CNYLGKPREQIDGYVMHANYIFWRNTNPIQLSSYSLFDMSKFPLKLRGTAVMSLKEGQINDM
ILSLLSKGRLIIRENNRVVISSDVLVNN
000000000000000000000000000000000000000000000000000000000000
000000000000000000000000000000000000000000000000000000000000
00001111111111111111111xxxxxxxxxxxxxxxxxxxxxxxxxxxxxxxxxxxxx

xxxxxxxxxxxxxxxxxxxxxxxxxxxxxxxxxxxxxxxxxxxxxxxxxxxxxxxxxxxxxx
xxxxxxxxxxxxxxxxxxxxxxxxxxxxxxxxxxxxxxxxxxxxxxxxxxxxxxxxxxxxxx
xxxxxxxxxxxxxxxxxxxxxxxxxxxxxxxxxxxxxxxxxxxxxxxxxxxxxxxxxxxxxx
xxxxxxxxxxxxxxxxxxxxxxxxxxxxxxxxxxxxxxxxxxxxxxxxxxxxxxxxxxxxxx
xxxxxxxxxxxxxxxxxxxxxxxxxxxxxxxxxxxxxxxxxxxxxxxxxxxxxxxxxxxxxx
xxxxxxxxxxxxxxxxxxxxxxxxxxxxxxxxxxxxxxxxxxxxxxxxxxxxxxxxxxxxxx
xxxxxxxxxxxxxxxxxxxxxxxxxxxxxxxxxxxxxxxxxxxxxxxxxxxxxxxxxxxxxx
xxxxxxxxxxxxxxxxxxxxxxxxxxxxxxxxxxxxxxxxxxxxxxxxxxxxxxxxxxxxxx
xxxxxxxxxxxxxxxxxxxxxxxxxxxxxxxxxxxxxxxxxxxxxxxxxxxxxxxxxxxxxx
xxxxxxxxxxxxxxxxxxxxxxxxxxxxxxxxxxxxxxxxxxxxxxxxxxxxxxxxxxxxxx
xxxxxxxxxxxxxxxxxxxxxxxxxxxxxxxxxxxxxxxxxxxxxxxxxxxxxxxxxxxxxx
xxxxxxxxxxxxxxxxxxxxxxxxxxxxxxxxxxxxxxxxxxxxxxxxxxxxxxxxxxxxxx
xxxxxxxxxxxxxxxxxxxxxxxxxxxxxxxxxxxxxxxxxxxxxxxxxxxxxxxxxxxxxx
xxxxxxxxxxxxxxxxxxxxxxxxxxxxxxxxxxxxxxxxxxxxxxxxxxxxxxxxxxxxxx
xxxxxxxxxxxxxxxxxxxxxxxxxxxxxxxxxxxxxxxxxxxxxxxxxxxxxxxxxxxxxx
xxxxxxxxxxxxxxxxxxxxxxxxxxxxxxxxxxxxxxxxxxxxxxxxxxxxxxxxxxxxxx
xxxxxxxxxxxxxxxxxxxxxxxxxxxxxxxxxxxxxxxxxxxxxxxxxxxxxxxxxxxxxx
xxxxxxxxxxxxxxxxxxxxxxxxxxxxxxxxxxxxxxxxxxxxxxxxxxxxxxxxxxxxxx
xxxxxxxxxxxxxxxxxxxxxxxxxxxxxxxxxxxxxxxxxxxxxxxxxxxxxxxxxxxxxx
xxxxxxxxxxxxxxxxxxxxxxxxxxxxxxxxxxxxxxxxxxxxxxxxxxxxxxxxxxxxxx
xxxxxxxxxxxxxxxxxxxxxxxxxxxxxxxxxxxxxxxxxxxxxxxxxxxxxxxxxxxxxx
xxxxxxxxxxxxxxxxxxxxxxxxxxxxxxxxxxxxxxxxxxxxxxxxxxxxxxxxxxxxxx
xxxxxxxxxxxxxxxxxxxxxxxxxxxxxxxxxxxxxxxxxxxxxxxxxxxxxxxxxxxxxx
xxxxxxxxxxxxxxxxxxxxxxxxxxxxxxxxxxxxxxxxxxxxxxxxxxxxxxxxxxxxxx
xxxxxxxxxxxxxxxxxxxxxxxxxxxxxxxxxxxxxxxxxxxxxxxxxxxxxxxxxxxxxx
xxxxxxxxxxxxxxxxxxxxxxxxxxxxxxxxxxxxxxxxxxxxx000000000000000xx
xxxxxxxxxxxxxxxxxxxxxxxxxxxxxxxxxxxxxxxxxxxxxxxxxxxxxxxxxxxxxx
xxxxxxxxxxxxxxxxxxxxxxxxxxxxxxxxxxxxxxxxxxxxxxxxxxxxxxxxxxxxxx
xxxxxxxxxxxxxxxxxxxxxxxxxxxxxxxxxxxxxxxxxxxxxxxxxxxxxxxxxxxxxx
xxxxxxxxxxxxxxxxxxxxxxxxxxxxxxxxxxxxxxxxxxxxxxxxxxxxxxxxxxxxxx
xxxxxxxxxxxxxxxxxxxxxxxxxxxxxxxxxxxxxxxxxxxxxxxxxxxxxxxxxxxxxx
xxxxxxxxxxxxxxxxxxxxxxxxxxxxxxxxxxxxxxxxxxxxxxxxxxxxxxxxxxxxxx
xxxxxxxxxxxxxxxxxxxxxxxxxxxxxxxxxxxxxxxxxxxxxxxxxxxxxxxxxxxxxx
xxxxxxxxxxxxxxxxxxxxxxxxxxxxxxxxxxxxxxxxxxxxxxxxxxxxxxxxxxxxxx
xxxxxxxxxxxxxxxxxxxxxxxxxxxxxxxxxxxxxxxxxxxxxxxxxxxxxxxxxxxxxx
xxxxxxxxxxxxxxxxxxxxxxxxxxxxxxxxxxxxxxxxxxxxxxxxxxxxxxxxxxxxxx
xxxxxxxxxxxxxxxxxxxxxxxxxxxxxxxxxxxxxxxxxxxxxxxxxxxxxxxxxxxxxx
xxxxxxxxxxxxxxxxxxxxxxxxxxxxxxxxxxxxxxxxxxxxxxxxxxxxxxxxxxxxxx
xxxxxxxxxxxxxxxxxxxxxxxxxxxxxxxxxxxxxxxxxxxxxxxxxxxxxxxxxxxxxx
xxxxxxxxxxxxxxxxxxxxxxxxxxxxxxxxxxxxxxxxxxxxxxxxxxxxxxxxxxxxxx
xxxxxxxxxxxxxxxxxxxxxxxxxxxxxxxxxxxxxxxxxxxxxxxxxxxxxxxxxxxxxx
xxxxxxxxxxxxxxxxxxxxxxxxxxxxxxxxxxxxxxxxxxxxxxxxxxxxxxxxxxxxxx
xxxxxxxxxxxxxxxxxxxxxxxxxxxxxxxxxxxxxxxxxxxxxxxxxxxxxxxxxxxxxx
xxxxxxxxxxxxxxxxxxxxxxxxxxxxxxxxxxxxxxxxxxxxxxxxxxxxxxxxxxxxxx

xxxxxxxxxxxxxxxxxxxxxxxxxxxxxxxxxxxxxxxxxxxxxxxxxxxxxxxxxxxxxx
xxxxxxxxxxxxxxxxxxxxxxxxxxxxxxxxxxxxxxxxxxxxxxxxxxxxxxxxxxxxxx
xxxxxxxxxxxxxxxxxxxxxxxxxxxxxxxxxxxxxxxxxxxxxxxxxxxxxxxxxxxxxx
xxxxxxxxxxxxxxxxxxxxxxxxxxxxxxxxxxxxxxxxxxxxxxxxxxxxxxxxxxxxxx
xxxxxxxxxxxxxxxxxxxxxxxxxxxxxxxxxxxxxxxxxxxxxxxxxxxxxxxxxxxxxx
xxxxxxxxxxxxxxxxxxxxxxxxxxxxxxxxxxxxxxxxxxxxxxxxxxxxxxxxxxxxxx
xxxxxxxxxxxxxxxxxxxxxxxxxxxxxxxxxxxxxxxxxxxxxxxxxxxxxxxxxxxxxx
xxxxxxxxxxxxxxxxxxxxxxxxxxxxxxxxxxxxxxxxxxxxxxxxxxxxxxxxxxxxxx
xxxxxxxxxxxxxxxxxxxxxxxxxxxxxxxxxxxxxxxxxxxxxxxxxxxxxxxxxxxxxx
xxxxxxxxxxxxxxxxxxxxxxxxxxxxxxxxxxxxxxxxxxxxxxxxxxxxxxxxxxxxxx
xxxxxxxxxxxxxxxxxxxxxxxxxxxxxxxxxxxxxxxxxxxxxxxxxxxxxxxxxxxxxx
xxxxxxxxxxxxxxxxxxxxxxxxxxxxxxxxxxxxxxxxxxxxxxxxxxxxxxxxxxxxxx
xxxxxxxxxxxxxxxxxxxxxxxxxxxxxxxxxxxxxxxxxxxxxxxxxxxxxxxxxxxxxx
xxxxxxxxxxxxxxxxxxxxxxxxxxxxxxxxxxxxxxxxxxxxxxxxxxxxxxxxxxxxxx
xxxxxxxxxxxxxxxxxxxxxxxxxxxxxxxxxxxxxxxxxxxxxxxxxxxxxxxxxxxxxx
xxxxxxxxxxxxxxxxxxxxxxxxxxxxxxxxxxxxxxxxxxxxxxxxxxxxxxxxxxxxxx
xxxxxxxxxxxxxxxxxxxxxxxx00000000000000000000000000000000000000
0000000000000000000000000000000000000000000000000000xxxxxxxxxx
xxxxxxxxxxxxxxxxxxxxxxxxxxxxxxxxxxxxxxxxxxxxxxxxxxxxxxxxxxxxxx
xxxxxxxxxxxxxxxxxxxxxxxxxxxxxxxxxxxxxxxxxxxxxxxxxxxxxxxxxxxxxx
xxxxxxxxxxxxxxxxxxxxxxxxxxxxxxxxxxxxxxxxxxxxxxxxxxxxxxxxxxxxxx
xxxxxxxxxxxxxxxxxxxxxxxxxxxxxxxxxxxxx000000000000000000xxxxxxx
xxxxxxxxxxxxxxxxxxxxxxxxxxxxxxxxxxxxxxxxxxxxxxxxxxxxxxxxxxxxxx
xxxxxxxxxxxxxxxxxxxxxxxxxxxxxxxxxxxxxxxxxxxxxxxxxxxxxxxxxxxxxx
xxxxxxxxxxxxxxxxxxxxxxxxxxxxxxxxxxxxxxxxxxxxxxxxxxxxxxxxxxxxxx
xxxxxxxxxxxxxxxxxxxxxxxxxxxxxxxxxxxxxxxxxxxxxxxxxxxxxxxxxxxxxx
xxxxxxxxxxxxxxxxxxxxxxxxxxxxxxxxxxxxxxxxxxxxxxxxxxxxxxxxxxxxxx
xxxxxxxxxxxxxxxxxxxxxxxxxxxxxxxxxxxxxxxxxxxxxxxxxxxxxxxxxxxxxx
xxxxxxxxxxxxxxxxxxxxxxxxxxxxxxxxxxxxxxxxxxxxxxxxxxxxxxxxxxxxxx
xxxxxxxxxxxxxxxxxxxxxxxxxxxxxxxxxxxxxxxxxxxxxxxxxxxxxxxxxxxxxx
xxxxxxxxxxxxxxxxxxxxxxxxxxxxxxxxxxxxxxxxxxxxxxxxxxxxxxxxxxxxxx
xxxxxxxxxxxxxxxxxxxxxxxxxxxxxxxxxxxxxxxxxxxxxxxxxxxxxxxxxxxxxx
xxxxxxxxxxxxxxxxxxxxxxxxxxxxxxxxxxxxxxxxxxxxxxxxxxxxxxxxxxxxxx
xxxxxxxxxxxxxxxxxxxxxxxxxxxxxxxxxxxxxxxxxxxxxxxxxxxxxxxxxxxxxx
xxxxxxxxxxxxxxxxxxxxxxxxxxxxxxxxxxxxxxxxxxxxxxxxxxxxxxxxxxxxxx
xxxxxxxxxxxxxxxxxxxxxxxxxxxxxxxxxxxxxxxxxxxxxxxxxxxxxxxxxxxxxx
xxxxxxxxxxxxxxxxxxxxxxxxxxxxxxxxxxxxxxxxxxxxxxxxxxxxxxxxxxxxxx
xxxxxxxxxxxxxxxxxxxxxxxxxxxxxxxxxxxxxxxxxxxxxxxxxxxxxxxxxxxxxx
xxxxxxxxxxxxxxxxxxxxxxxxxxxxxxxxxxxxxxxxxxxxxxxxxxxxxxxxxxxxxx
xxxxxxxxxxxxxxxxxxxxxxxxxxxxxxxxxxxxxxxxxxxxxxxxxxxxxxxxxxxxxx
xxxxxxxxxxxxxxxxxxxxxxxxxxxxxxxxxxxxxxxxxxxxxxxxxxxxxxxxxxxxxx
xxxxxxxxxxxxxxxxxxxxxxxxxxxxxxxxxxxxxxxxxxxxxxxxxxxxxxxxxxxxxx
xxxxxxxxxxxxxxxxxxxxxxxxxxxxxxxxxxxxxxxxxxxxxxxxxxxxxxxxxxxxxx
xxxxxxxxxxxxxxxxxxxxxxxxxxxxxxxxxxxxxxxxxxxxxxxxxxxxxxxxxxxxxx

xxxxxxxxxxxxxxxxxxxxxxxxxxxxxxxxxxxxxxxxxxxxxxxxxxxxxxxxxxxx
xxxxxxxxxxxxxxxxxxxxxxxxxxxxxxxxxxxxxxxxxxxxxxxxxxxxxxxxxxxx
xxxxxxxxxxxxxxxxxxxxxxxxxxxxxxxxxxxxxxxxxxxxxxxxxxxxxxxxxxxx
xxxxxxxxxxxxxxxxxxxxxxxxxxxxxxxxxxxxxxxxxxxxxxxxxxxxxxxxxxxx
xxxxxxxxxxxxxxxxxxxxxxxxxxxxxxxxxxxxxxxxxxxxxxxxxxxxxxxxxxxx
xxxxxxxxxxxxxxxxxxxxxxxxxxxxxxxxxxxxxxxxxxxxxxxxxxxxxxxxxxxx
xxxxxxxxxxxxxxxxxxxxxxxxxxxxxxxxxxxxxxxxxxxxxxxxxxxxxxxxxxxx
xxxxxxxxxxxxxxxxxxxxxxxxxxxxxxxxxxxxxxxxxxxxxxxxxxxxxxxxxxxx
xxxxxxxxxxxxxxxxxxxxxxxxxxxxxxxxxxxxxxxxxxxxxxxxxxxxxxxxxxxx
xxxxxxxxxxxxxxxxxxxxxxxxxxxxxxxxxxxxxxxxxxxxxxxxxxxxxxxxxxxx
xxxxxxxxxxxxxxxxxxxxxxxxxxxxxxxxxxxxxxxxxxxxxxxxxxxxxxxxxxxx
xxxxxxxxxxxxxxxxxxxxxxxxxxxxxxxxxxxxxxxxxxxxxxxxxxxxxxxxxxxx
xxxxxxxxxxxxxxxxxxxxxxxxxxxxxxxxxxxxxxxxxxxxxxxxxxxxxxxxxxxx
xxxxxxxxxxxxxxxxxxxxxxxxxxxxxxxxxxxxxxxxxxxxxxxxxxxxxxxxxxxx
xxxxxxxxxxxxxxxxxxxxxxxxxxxxxxxxxxxxxxxxxxxxxxxxxxxxxxxxxxxx
xxxxxxxxxxxxxxxxxxxxxxxxxxxxxxxxxxxxxxxxxxxxxxxxxxxxxxxxxxxx
xxxxxxxxxxxxxxxxxxxxxxxxxxxxxxxxxxxxxxxxxxxxxxxxxxxxxxxxxxxx
xxxxxxxxxxxxxxxxxxxxxxxxxxxxxxxxxxxxxxxxxxxxxxxxxxxxxxxxxxxx
xxxxxxxxxxxxxxxxxxxxxxxxxxxxxxxxxxxxxxxxxxxxxxxxxxxxxxxxxxxx
xxxxxxxxxxxxxxxxxxxxxxxxxxxxxxxxxxxxxxxxxxxxxxxxxxxxxxxxxxxx
xxxxxxxxxxxxxxxxxxxxxxxxxxxxxxxxxxxxxxxxxxxxxxxxxxxxxxxxxxxx
xxxxxxxxxxxxxxxxxxxxxxxxxxxxxxxxxxxxxxxxxxxxxxxxxxxxxxxxxxxx
xxxxxxxxxxxxxxxxxxxxxxxxxxxxxxxxxxxxxxxxxxxxxxxxxxxxxxxxxxxx
xxxxxxxxxxxxxxxxxxxxxxxxxxxx
>DP02946
MTAIFSLAINSNFVLANNDKPDASDDKYADYVVRLGSEHPLNHTQIIELSSAVSRAVLLSYP
NIIDRYTAAATEYTVIDALFHSPTFRHIVSFGLHNQQENLGHIRYTNEYEINNNREDEFSLV
SEVSYDDIKSSNAQQVPLVAFYEAREDRATGTPIVNMGVAPSLFSGRYSWWQEALIHEIVHH
VTGSSDTHEENKQGPTEILAQMVAAELHWAIPTFKGYSDPARVEAIQERDFHSLLNMFQRHG
SELGFLFTRLATIAKGKKASPDFGTLTSFCSEGISSFPKYPDHDDDFNGGGAFFLPSASADS
SVECTFDVLNRIEPVDDSIKFEGGNLLIKNDFKNLNLRVAQLSFLNAKKGSGFYRKNWDSWK
SWYQASSWKNGLNSGLYGYGHDESEGNLIYSPYGITFNDGSFSIGFSSRKHINDNTKDDNFV
KLNNANWSSFYYAGQMFFDKNKRPVALVITEPLNAAFGAGWSYIYKDGKWHYEAQDDWDQRL
FKDSTLSLDPHAPQFIN
xxxxxxxxxxxxxxxxxxxxxxxxxxxxxxxxxxxxxxxxxxxxxxxxxxxxxxxxxxxx
xxxxxxxxxxxxxxxxxxxxxxxxxxxxxxxxxxxxxxxxxxxxxxxxxxxxxxxxxxxx
xxxxxxxxxxxxxxxxxxxxxxxxxxxxxxxxxxxxxxxxxxxxxxxxxxxxxxxxxxxx
xxxxxxxxxxxxxxxxxxxxxxxxxxxxxxxxxxxxxxxxxxxxxxxxxxxxxxxxxxxx
xxxxxxxxxxxxxxxxxxxxxxxxxxxxx1111111111111111111111111111111
1111xxxxxxxxxxxxxxxxxxxxxxxxxxxxxxxxxxxxxxxxxxxxxxxxxxxxxxxx
xxxxxxxxxxxxxxxxxxxxxxxxxxxxxxxxxxxxxxxxxxxxxxxxxxxxxxxxxxxx
xxxxxxxxxxxxxxxxxxxxxxxxxxxxxxxxxxxxxxxxxxxxxxxxxxxxxxxxxxxx
xxxxxxxxxxxxxxxxx
>DP02950

MGSINLRIDDELKARSYAALEKMGVTPSEALRLMLEYIADNERLPFKQTLLSDEDAELVEIV
KERLRNPKPVRVTLDEL
xxxxxxxxxxxxxxxxxxxxxxxxxxxxxxxxxxxxxxxxxx00000000000000000000
00000000000000000
>DP02964
MIMSDPAMLPPGFRFHPTDEELILHYLRNRAAQSPCPVSIIADVDIYKFDPWALPSKASYGD
REWYFFTPRDRKYPNGVRPNRAAGSGYWKATGTDKPIRCSATGESVGVKKALVFYKGRPPKG
IKTNWIMHEYRLAAADAHAANTYRPMKFRNASMRLDDWVLCRIYKKTSQVSPMAVPPLSDHE
LDEPSGAGAYPMSSAGMTMQGGAGGYTLQAAVPGTQRMPKIPSISELLNDYSLAQLFDDSGH
ALMARHDQHAALFGHPIMSQFHVNSSGNNMSQLGQMDSPASTSVARDGAAGKRKRLSEEDGE
HNGSTSQPAAAVTNKKPNSSCFGATTFQVGNNTLQGSLGQPLLHF
xxxxxxxxxxxxxxxxxxxxxxxxxxxxxxxxxxxxxxxxxxxxxxxxxxxxxxxxxxxxxx
xxxxxxxxxxxxxxxxxxxxxxxxxxxxxxxxxxxxxxxxxxxxxxxxxxxxxxxxxxxxxx
xxxxxxxxxxxxxxxxxxxxxxxxxxxxxxxxxxxxxxxxxxxxxxx000000000000000
00000000000000000000000000000000000000000000000000000000000000
00000000000000000000000000000000000000000000000000000000000000
000000000000000000000000000000000000000000000
>DP02965
MSEVSVINQATEVEDAAAGLDLPPGFRFHPTDEEIISHYLTPKALDHRFCSGVIGEVDLNKC
EPWHLPGKAKMGEKEWYFFCHKDRKYPTGTRTNRATMSGYWKATGKDKEIFRGRGILVGMKK
TLVFYLGRAPRGEKTGWVMHEFRLEGRLPHPLPRSAKDEWAVSKVFNKELTATNGAMATAMA
AAPDAGIERVSSFGFISDHFLDAGELPPLMDPPLGGDVDQVIDFNSNSAYATGGRSGSGLEV
KMEQHMPPHMMYSSPYFSLPAANSGDLSPAIRRYCKAEQVSGQTSALSPSRDTGLSTDPNAA
GCAEISSAPTPSSHNQDFLDHLDEYPALNLADIWKY
xxxxxxxxxxxxxxxxxxxxxxxxxxxxxxxxxxxxxxxxxxxxxxxxxxxxxxxxxxxxxx
xxxxxxxxxxxxxxxxxxxxxxxxxxxxxxxxxxxxxxxxxxxxxxxxxxxxxxxxxxxxxx
xxxxxxxxxxxxxxxxxxxxxxxxxxxxxxxxxxxxxxxxxxxxxxxxxxxx0000000000
00000000000000000000000000000000000000000000000000000000000000
00000000000000000000000000000000000000000000000000000000000000
000000000000000000000000000000000000
>DP03027
MKKAKEIAKCVAVASVIMSGSFGLQATSAFADSKGTVENLQNGGKVYNSFKTTYDMKQNIKN
SIKVSFIEDPYADKKIAIVTTDGSNIDAKYTINSGYYNAGLKWPSAYHTEAEITSGDSAQFH
KAAPVNTMTSAKVTSEVGYTLGGSVKVGVNDKGPNADASITGSFAWKESVSYDQVDYKTVLE
THTDKKLNWKVGFQSFNFPEWGIYNRDSFNTFYGNQLFMKSRSYNEGTNNFVSKDTVPALTG
YGFSPNVVAVITADKTETTSDLKITNRRISDQYNIEWVSSKWWGTNNKDTYNEFFTNHYKLD
WKNHQVTLDNQKALEEQMNSINSVNDKLNKGKGKLSLSMNGNQLKATSSNAGYGISYEDKNW
GIFVNGEKVYTFNEKSTVGNISNDINKLNIKGPYIEIKQI
xxxxxxxxxxxxxxxxxxxxxxxxxxxxxxxxxxxxxxxxxxxxxxxxxxxxxxxxxxxxxx
xxxxxxxxxxxxxxxxxxxxxxxxxxxxxxxxxxxxxxxxxxxxxxxxxxxxxxxxxxxxxx
xxxxxxxxxxxxxxxxxxxxxxxxxxxxxxxxxxxxxxxxxxxxxxxxxxxxxxxxxxxxxx
xxxxxxxxxxxxxxxxxxxxxxxxxxxxxxxxxxxxxxxxxxxxxxxxxxxxxxxxxxxxxx
xxxxxxxxxxxxxxxxxxxxxxxxxxxxxxxxxxxxxxxxxxxxxxxxxxxxxxxxxxxxxx
xxxxxxxx11111111111xxxxxxxxxxxxxxxxxxxxxxxxxxxxxxxxxxxxxxxxxxx

xxxxxxxxxxxxxxxxxxxxxxxxxxxxxxxxxxxxxxx

>DP03034

MEIQQTHRKINRPLVSLALVGALVSITPQQSHAAFFTTVIIPAIVGGIATGTAVGTVSGLLG
WGLKQAEEANKTPDKPDKVWRIQAGKGFNEFPNKEYDLYKSLLSSKIDGGWDWGNAATHYWI
KGGQWNKLEVDMKDAVGTYKLSGLRNFTGGDLDVNMQKATLRLGQFNGNSFTSYKDSADRTT
RVDFNAKNILIDNFLEINNRVGSGAGRKASSTVLTLQASEGITSSKNAEISLYDGATLNLAS
NSVKLNGNVWMGRLQYVGAYLAPSYSTINTSKVTGEVNFNHLTVGDHNAAQAGIIASNKTHI
GTLDLWQSAGLNIIAPPEGGYKDKPNNTPSQSGAKNDKQESSQNNSNTQVINPPNSTQKTEV
QPTQVIDGPFAGGKDTVVNIDRINTKADGTIKVGGFKASLTTNAAHLNIGKGGVNLSNQASG
RTLLVENLTGNITVDGPLRVNNQVGGYALAGSSANFEFKAGVDTKNGTATFNNDISLGRFVN
LKVDAHTANFKGIDTGNGGFNTLDFSGVTNKVNINKLITASTNVAVKNFNINELIVKTNGVS
VGEYTHFSEDIGSQSRINTVRLETGTRSIFSGGVKFKSGEKLVIDEFYYSPWNYFDARNIKN
VEITRKFASSTPENPWGTSKLMFNNLTLGQNAVMDYSQFSNLTIQGDFINNQGTINYLVRGG
KVATLNVGNAAAMMFNNDIDSATGFYKPLIKINSAQDLIKNTEHVLLKAKIIGYGNVSTGTN
GISNVNLEEQFKERLALYNNNNRMDTCVVRNTDDIKACGMAIGNQSMVNNPDNYKYLIGKAW
KNIGISKTANGSKISVYYLGNSTPTENGGNTTNLPTNTTNNARFASYALIKNAPFAHSATPN
LVAINQHDFGTIESVFELANRSKDIDTLYANSGAQGRDLLQTLLIDSHDAGYARTMIDATSA
NEITKQLNTATTTLNNIASLEHKTSSLQTLSLSNAMILNSRLVNLSRRHTNNIDSFAKRLQA
LKDQRFASLESAAEVLYQFAPKYEKPTNVWANAIGGASLNNGGNASLYGTSAGVDAYLNGQV
EAIVGGFGSYGYSSFNNQANSLNSGANNTNFGVYSRIFANQHEFDFEAQGALGSDQSSLNFK
SALLRDLNQSYNYLAYSAATRASYGYDFAFFRNALVLKPSVGVSYNHLGSTNFKSNSTNKVA
LSNGSSSQHLFNASANVEARYYYGDTSYFYMNAGVLQEFANFGSSNAVSLNTFKVNATRNPL
NTHARVMMGGELKLAKEVFLNLGVVYLHNLISNIGHFASNLGMRYSF

xxxxxxxxxxxxxxxxxxxxxxxxxxxxxxxxxxxxxxxxxxxxxxxxxxxxxxxxxxxx
xxxxxxxxxxxxxxxxxxxxxxxxxxxxxxxxxxxxxxxxxxxxxxxxxxxxxxxxxxxx
xxxxxxxxxxxxxxxxxxxxxxxxxxxxxxxxxxxxxxxxxxxxxxxxxxxxxxxxxxxx
xxxxxxxxxxxxxxxxxxxxxxxxxxxxxxxxxxxxxxxxxxxxxxxxxxxxxxxxxxxx
xxxxxxxxxxxxxxxxxxxxxxxxxxxxxxxxxxxxxxxxxxxxxxxxxxxxxxxxxxxx
xxxxxxxxxxxxxxxxxxxx00000011111111111100000000000000000xxxxx
xxxxxxxxxxxxxxxxxxxxxxxxxxxxxxxxxxxxxxxxxxxxxxxxxxxxxxxxxxxx
xxxxxxxxxxxxxxxxxxxxxxxxxxxxxxxxxxxxxxxxxxxxxxxxxxxxxxxxxxxx
xxxxxxxxxxxxxxxxxxxxxxxxxxxxxxxxxxxxxxxxxxxxxxxxxxxxxxxxxxxx
xxxxxxxxxxxxxxxxxxxxxxxxxxxxxxxxxxxxxxxxxxxxxxxxxxxxxxxxxxxx
xxxxxxxxxxxxxxxxxxxxxxxxxxxxxxxxxxxxxxxxxxxxxxxxxxxxxxxxxxxx
xxxxxxxxxxxxxxxxxxxxxxxxxxxxxxxxxxxxxxxxxxxxxxxxxxxxxxxxxxxx
xxxxxxxxxxxxxxxxxxxxxxxxxxxxxxxxxxxxxxxxxxxxxxxxxxxxxxxxxxxx
xxxxxxxxxxxxxxxxxxxxxxxxxxxxxxxxxxxxxxxxxxxxxxxxxxxxxxxxxxxx
xxxxxxxxxxxxxxxxxxxxxxxxxxxxxxxxxxxxxxxxxxxxxxxxxxxxxxxxxxxx
xxxxxxxxxxxxxxxxxxxxxxxxxxxxxxxxxxxxxxxxxxxxxxxxxxxxxxxxxxxx
xxxxxxxxxxxxxxxxxxxxxxxxxxxxxxxxxxxxxxxxxxxxxxxxxxxxxxxxxxxx
xxxxxxxxxxxxxxxxxxxxxxxxxxxxxxxxxxxxxxxxxxxxxxxxxxxxxxxxxxxx
xxxxxxxxxxxxxxxxxxxxxxxxxxxxxxxxxxxxxxxxxxxxxxxxxxxxxxxxxxxx
xxxxxxxxxxxxxxxxxxxxxxxxxxxxxxxxxxxxxxxxxxxxxxxxxxxxxxxxxxxx
xxxxxxxxxxxxxxxxxxxxxxxxxxxxxxxxxxxxxxxxxxxxx

>DP03121
MDVGSKEVLMESPPDYSAAPRGRFGIPCCPVHLKRLLIVVVVVVLIVVVIVGALLMGLHMSQ
KHTEMVLEMSIGAPEAQQRLALSEHLVTTATFSIGSTGLVVYDYQQLLIAYKPAPGTCCYIM
KIAPESIPSLEALNRKVHNFQMECSLQAKPAVPTSKLGQAEGRDAGSAPSGGDPAFLGMAVN
TLCGEVPLYYI
xxxxxxxxxxxxxxxxxxxxxxxxxxxxxxxxxxxxxxxxxxxxxxxxxxxxxxxx1111
11111111111111111111111111xxxxxxxxxxxxxxxxxxxxxxxxxxxxxxxxxx
xxxxxxxxxxxxxxxxxxxxxxxxx0000000000000000000000000000000xxxx
xxxxxxxxxxx
>DP03226
MVGSGSHTPYDISNSPSDVNVQPATQLNSTLVEDDDVDNQLFEEAQVTETGFRSPSASDNSC
AYCGIDSAKCVIKCNSCKKWFCNTKNGTSSSHIVNHLVLSHHNVVSLHPDSDLGDTVLECYN
CGRKNVFLLGFVSAKSEAVVVLLCRIPCAQTKNANWDTDQWQPLIEDRQLLSWVAEQPTEEE
KLKARLITPSQISKLEAKWRSNKDATINDIDAPEEQEAIPPLLLRYQDAYEYQRSYGPLIKL
EADYDKQLKESQALEHISVSWSLALNNRHLASFTLSTFESNELKVAIGDEMILWYSGMQHPD
WEGRGYIVRLPNSFQDTFTLELKPSKTPPPTHLTTGFTAEFIWKGTSYDRMQDALKKFAIDK
KSISGYLYYKILGHQVVDISFDVPLPKEFSIPNFAQLNSSQSNAVSHVLQRPLSLIQGPPGT
GKTVTSATIVYHLSKIHKDRILVCAPSNVAVDHLAAKLRDLGLKVVRLTAKSREDVESSVSN
LALHNLVGRGAKGELKNLLKLKDEVGELSASDTKRFVKLVRKTEAEILNKADVVCCTCVGAG
DKRLDTKFRTVLIDESTQASEPECLIPIVKGAKQVILVGDHQQLGPVILERKAADAGLKQSL
FERLISLGHVPIRLEVQYRMNPYLSEFPSNMFYEGSLQNGVTIEQRTVPNSKFPWPIRGIPM
MFWANYGREEISANGTSFLNRIEAMNCERIITKLFRDGVKPEQIGVITPYEGQRAYILQYMQ
MNGSLDKDLYIKVEVASVDAFQGREKDYIILSCVRANEQQAIGFLRDPRRLNVGLTRAKYGL
VILGNPRSLARNTLWNHLLIHFREKGCLVEGTLDNLQLCTVQLVRPQPRKTERPMNAQFNVE
SEMGDFPKFQDFDAQSMVSFSGQIGDFGNAFVDNTELSSYINNEYWNFENFKSAFSQKQNRN
EIDDRNLYQEEASHLNSNFARELQREEQKHELSKDFSNLGI
xxxxxxxxxxxxxxxxxxxxxxxxxxxxxxxxxxxxxxxxxxxxxxxxxxxxxxxxxxxx
xxxxxxxxxxxxxxxxxxxxxxxxxxxxxxxxxxxxxxxxxxxxxxxxxxxxxxxxxxxx
xxxxxxxxxxxxxxxxxxxxxxxxxxxxxxxxxxxxxxxxxxxxxxxxxxxxxxxxxxxx
xxxxxxxxxxxxxxxxxxxxxxxxx111111111111xxxxxxxxxxxxxxxxxxxxxxx
xxxxxxxxxxxxxxxxxxxxxxxxxxxxxxxxxxxxxxxxxxxxxxxxxxxxxxxxxxxx
xxxxxxxxxxxxxxxxxxxxxxxxxxxxxxxxxxxxxxxxxxxxxxxxxxxxxxxxxxxx
xxxxxxxxxxxxxxxxxxxxxxxxxxxxxxxxxxxxxxxxxxxxxxxxxxxxxxxxxxxx
xxxxxxxxxxxxxxxxxxxxxxxxxxxxxxxxxxxxxxxxxxxxxxxxxxxxxxxxxxxx
xxxxxxxxxxxxxxxxxxxxxxxxxxxxxxxxxxxxxxxxxxxxxxxxxxxxxxxxxxxx
xxxxxxxxxxxxxxxxxxxxxxxxxxxxxxxxxxxxxxxxxxxxxxxxxxxxxxxxxxxx
xxxxxxxxxxxxxxxxxxxxxxxxxxxxxxxxxxxxxxxxxxxxxxxxxxxxxxxxxxxx
xxxxxxxxxxxxxxxxxxxxxxxxxxxxxxxxxxxxxxxxxxxxxxxxxxxxxxxxxxxx
xxxxxxxxxxxxxxxxxxxxxxxxxxxxxxxxxxxxxxxxxxxxxxxxxxxxxxxxxxxx
xxxxxxxxxxxxxxxxxxxxxxxxxxxxxxxxxxxxxxxxxxxxxxxxxxxxxxxxxxxx
xxxxxxxxxxxxxxxxxxxxxxxxxxxxxxxxxxxxxxxxxxxxxxxxxxxxxxxxxxxx
xxxxxxxxxxxxxxxxxxxxxxxxxxxxxxxxxxxxxxxxx
>DP03309
MEVSRRKAPPRPPRPAAPLPLLAYLLALAAPGRGADEPVWRSEQAIGAIAASQEDGVFVASG

SCLDQLDYSLEHSLSRLYRDQAGNCTEPVSLAPPARPRPGSSFSKLLLPYREGAAGLGGLLL
TGWTFDRGACEVRPLGNLSRNSLRNGTEVVSCHPQGSTAGVVYRAGRNNRWYLAVAATYVLP
EPETASRCNPAASDHDTAIALKDTEGRSLATQELGRLKLCEGAGSLHFVDAFLWNGSIYFPY
YPYNYTSGAATGWPSMARIAQSTEVLFQGQASLDCGHGHPDGRRLLLSSSLVEALDVWAGVF
SAAAGEGQERRSPTTTALCLFRMSEIQARAKRVSWDFKTAESHCKEGDQPERVQPIASSTLI
HSDLTSVYGTVVMNRTVLFLGTGDGQLLKVILGENLTSNCPEVIYEIKEETPVFYKLVPDPV
KNIYIYLTAGKEVRRIRVANCNKHKSCSECLTATDPHCGWCHSLQRCTFQGDCVHSENLENW
LDISSGAKKCPKIQIIRSSKEKTTVTMVGSFSPRHSKCMVKNVDSSRELCQNKSQPNRTCTC
SIPTRATYKDVSVVNVMFSFGSWNLSDRFNFTNCSSLKECPACVETGCAWCKSARRCIHPFT
ACDPSDYERNQEQCPVAVEKTSGGGRPKENKGNRTNQALQVFYIKSIEPQKVSTLGKSNVIV
TGANFTRASNITMILKGTSTCDKDVIQVSHVLNDTHMKFSLPSSRKEMKDVCIQFDGGNCSS
VGSLSYIALPHCSLIFPATTWISGGQNITMMGRNFDVIDNLIISHELKGNINVSEYCVATYC
GFLAPSLKSSKVRTNVTVKLRVQDTYLDCGTLQYREDPRFTGYRVESEVDTELEVKIQKEND
NFNISKKDIEITLFHGENGQLNCSFENITRNQDLTTILCKIKGIKTASTIANSSKKVRVKLG
NLELYVEQESVPSTWYFLIVLPVLLVIVIFAAVGVTRHKSKELSRKQSQQLELLESELRKEI
RDGFAELQMDKLDVVDSFGTVPFLDYKHFALRTFFPESGGFTHIFTEDMHNRDANDKNESLT
ALDALICNKSFLVTVIHTLEKQKNFSVKDRCLFASFLTIALQTKLVYLTSILEVLTRDLMEQ
CSNMQPKLMLRRTESVVEKLLTNWMSVCLSGFLRETVGEPFYLLVTTLNQKINKGPVDVITC
KALYTLNEDWLLWQVPEFSTVALNVVFEKIPENESADVCRNISVNVLDCDTIGQAKEKIFQA
FLSKNGSPYGLQLNEIGLELQMGTRQKELLDIDSSSVILEDGITKLNTIGHYEISNGSTIKV
FKKIANFTSDVEYSDDHCHLILPDSEAFQDVQGKRHRGKHKFKVKEMYLTKLLSTKVAIHSV
LEKLFRSIWSLPNSRAPFAIKYFFDFLDAQAENKKITDPDVVHIWKTNSLPLRFWVNILKNP
QFVFDIKKTPHIDGCLSVIAQAFMDAFSLTEQQLGKEAPTNKLLYAKDIPTYKEEVKSYYKA
IRDLPPLSSSEMEEFLTQESKKHENEFNEEVALTEIYKYIVKYFDEILNKLERERGLEEAQK
QLLHVKVLFDEKKKCKWM
xxxxxxxxxxxxxxxxxxxxxxxxxxxxxxxxxxxxxxxxxxxxxxxxxxxxxxxxxxxx
xxxxxxxxxxxxxxxxxxxxxxxxxxxxxxxxxxxxxxxxxxxxxxxxxxxxxxxxxxxx
xxxxxxxxxxxxxxxxxxxxxxxxxxxxxxxxxxxxxxxxxxxxxxxxxxxxxxxxxxxx
xxxxxxxxxxxxxxxxxxxxxxxxxxxxxxxxxxxxxxxxxxxxxxxxxxxxxxxxxxxx
xxxxxxxxxxxxxxxxxxxxxxxxxxxxxxxxxxxxxxxxxxxxxxxxxxxxxxxxxxxx
xxxxxxxxxxxxxxxxxxxxxxxxxxxxxxxxxxxxxxxxxxxxxxxxxxxxxxxxxxxx
xxxxxxxxxxxxxxxxxxxxxxxxxxxxxxxxxxxxxxxxxxxxxxxxxxxxxxxxxxxx
xxxxxxxxxxxxxxxxxxxxxxxxxxxxxxxxxxxxxxxxxxxxxxxxxxxxxxxxxxxx
xxxxxxxxxxxxxxxxxxxxxxxxxxxxxxxxxxxxxxxxxxxxxxxxxxxxxxxxxxxx
xxxxxxxxxxxxxxxxxxxxxxxxxxxxxxxxxxxxxxxxxxxxxxxxxxxxxxxxxxxx
xxxxxxxxxxxxxxxxxxxx11111111111111111xxxxxxxxxxxxxxxxxxxxxx
xxxxxxxxxxxxxxxxxxxxxxxxxxxxxxxxxxxxxxxxxxxxxxxxxxxxxxxxxxxx
xxxxxxxxxxxxxxxxxxxxxxxxxxxxxxxxxxxxxxxxxxxxxxxxxxxxxxxxxxxx
xxxxxxxxxxxxxxxxxxxxxxxxxxxxxxxxxxxxxxxxxxxxxxxxxxxxxxxxxxxx
xxxxxxxxxxxxxxxxxxxxxxxxxxxxxxxxxxxxxxxxxxxxxxxxxxxxxxxxxxxx
xxxxxxxxxxxxxxxxxxxxxxxxxxxxxxxxxxxxxxxxxxxxxxxxxxxxxxxxxxxx
xxxxxxxxxxxxxxxxxxxxxxxxxxxxxxxxxxxxxxxxxxxxxxxxxxxxxxxxxxxx
xxxxxxxxxxxxxxxxxxxxxxxxxxxxxxxxxxxxxxxxxxxxxxxxxxxxxxxxxxxx
xxxxxxxxxxxxxxxxxxxxxxxxxxxxxxxxxxxxxxxxxxxxxxxxxxxxxxxxxxxx

xxxxxxxxxxxxxxxxxxxxxxxxxxxxxxxxxxxxxxxxxxxxxxxxxxxxxxxxxxxx
xxxxxxxxxxxxxxxxxxxxxxxxxxxxxxxxxxxxxxxxxxxxxxxxxxxxxxxxxxxx
xxxxxxxxxxxxxxxxxxxxxxxxxxxxxxxxxxxxxxxxxxxxxxxxxxxxxxxxxxxx
xxxxxxxxxxxxxxxxxxxxxxxxxxxxxxxxxxxxxxxxxxxxxxxxxxxxxxxxxxxx
xxxxxxxxxxxxxxxxxxxxxxxxxxxxxxxxxxxxxxxxxxxxxxxxxxxxxxxxxxxx
xxxxxxxxxxxxxxxxxxxxxxxxxxxxxxxxxxxxxxxxxxxxxxxxxxxxxxxxxxxx
xxxxxxxxxxxxxxxxxx
>DP03338
MSDQSQEPTMEEILASIRRIISEDDAPAEPAAEAAPPPPPEPEPEPVSFDDEVLELTDPIAP
EPELPPLETVGDIDVYSPPEPESEPAYTPPPAAPVFDRDEVAEQLVGVSAASAAASAFGSLS
SALLMPKDGRTLEDVVRELLRPLLKEWLDQNLPRIVETKVEEEVQRISRGRGA
000000000000000000000000000000000000000000000000000000000000
000000000000000000000000000000000000000000000000000000000000
000000000xxxxxxxxxxxxxxxxxxxxxxxxxxxxxxxxxxxxxxxxxxxx
>DP03345
MMLSLNNLQNIIYNPVIPFVGTIPDQLDPGTLIVIRGHVPSDADRFQVDLQNGSSMKPRADV
AFHFNPRFKRAGCIVCNTLINEKWGREEITYDTPFKREKSFEIVIMVLKDKFQVAVNGKHTL
LYGHRIGPEKIDTLGIYGKVNIHSIGFSFSSDLQSTQASSLELTEISRENVPKSGTPQLRLP
FAARLNTPMGPGRTVVVKGEVNANAKSFNVDLLAGKSKDIALHLNPRLNIKAFVRNSFLQES
WGEEERNITSFPFSPGMYFEMIIYCDVREFKVAVNGVHSLEYKHRFKELSSIDTLEINGDIH
LLEVRSW
xxxxxxxxxxxxxxxxxxxxxxxxxxxxxxxxxxxxxxxxxxxxxxxxxxxxxxxxxxxx
xxxxxxxxxxxxxxxxxxxxxxxxxxxxxxxxxxxxxxxxxxxxxxxxxxxxxxxxxxxx
xxxxxxxxxxxxxxxxxxxxxxxxxxxx111111111111111111111xxxxxxxxxxx
xxxxxxxxxxxxxxxxxxxxxxxxxxxxxxxxxxxxxxxxxxxxxxxxxxxxxxxxxxxx
xxxxxxxxxxxxxxxxxxxxxxxxxxxxxxxxxxxxxxxxxxxxxxxxxxxxxxxxxxxx
xxxxxxx
