## Supplementary Information S3 for "TransDFL: Identification of Disordered Flexible Linkers in Proteins by Transfer Learning"

**Supplementary Information S3.** The length distribution of DFLs on TR166 dataset (**Figure S1**). The precision-recall curves of TransDFL and the other predictors on TE82 and TE64 datasets in situation-I (**Figure S2**). The precision-recall curves of TransDFL and the other predictors on TE82 and TE64 datasets in situation-II (**Figure S3**). The performance of TransDFL based on different feature combinations on DFL validation dataset (**Table S1**). The performance of TransDFL based on different loss function weight coefficients on DFL validation dataset (**Table S2**). The hyper-parameters of RFPR-IDP (pre-trained) (**Table S3**). The hyper-parameters of TransDFL (fine-tuned) (**Table S4**). The performance of 6 state-of-the-art IDR predictors for predicting DFLs on TE82 dataset (situation-I) (**Table S5**). The performance of 6 state-of-the-art IDR predictors for predicting DFLs on TE82 dataset (situation-II) (**Table S6**).

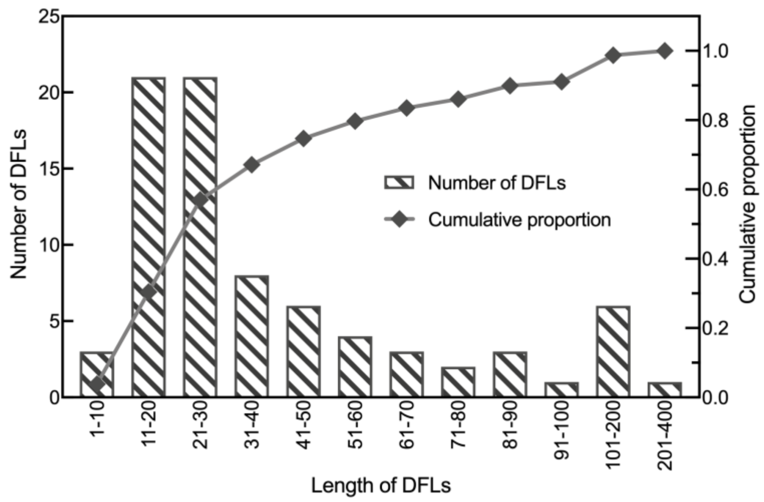


**Figure S1.** The length distribution of DFLs in TR166 dataset (the average length is 47).

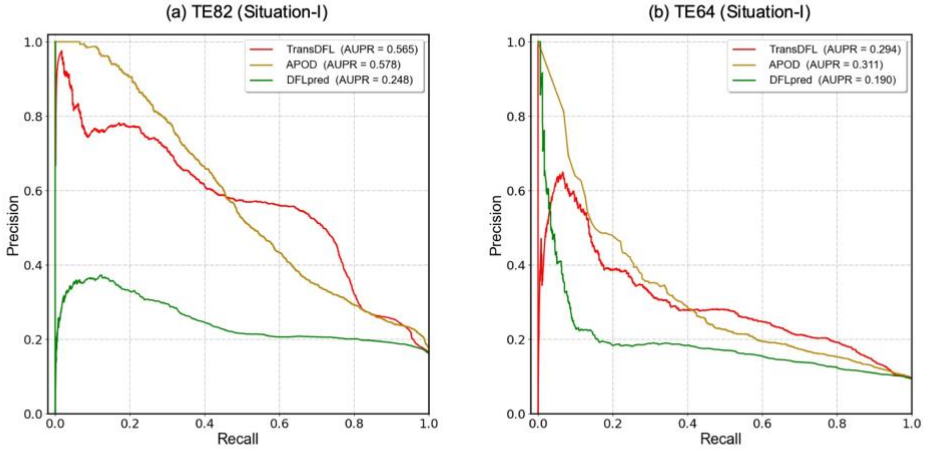


**Figure S2.** The precision-recall curves of TransDFL and the other predictors on TE82 and TE64 datasets in situation-I.

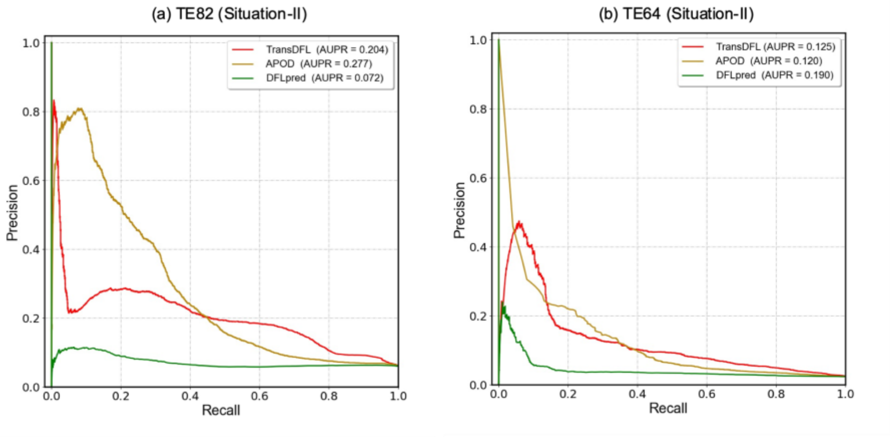


**Figure S3.** The precision-recall curves of TransDFL and the other predictors on TE82 and TE64 datasets in situation-II.

**Table S1.** The performance of TransDFL based on different feature combinations on DFL validation dataset.

| **Features (dimension)** | **AUC** | |
|---|---|---|
| | **Situation-I** | **Situation-II** |
| PSSM (40) | 0.680 | 0.655 |
| Seven (7), SS (4), SA (1) | 0.778 | 0.689 |
| PSSM (40), Seven (7) | 0.752 | 0.702 |
| PSSM (40), Seven (7), SS (4) | 0.814 | 0.744 |
| PSSM (40), Seven (7), SS (4), SA (1) | 0.861 | 0.782 |

**Table S2.** The performance of TransDFL based on different loss function weight coefficients on DFL validation dataset.

| **Weight** | | **AUC** | |
|---|---|---|---|
| **Positive** | **Negative** | **Situation-I** | **Situation-II** |
| 0.5 | 0.5 | 0.833 | 0.761 |
| 0.6 | 0.4 | 0.861 | 0.782 |
| 0.7 | 0.3 | 0.826 | 0.755 |
| 0.8 | 0.2 | 0.805 | 0.718 |

**Table S3.** The hyper-parameters of RFPR-IDP (pre-trained).

<table>
<tr><td colspan="2">Hyper-parameter</td><td>Details</td></tr>
<tr><td colspan="2">Feature sliding window</td><td>9</td></tr>
<tr><td colspan="2">Batch_size</td><td>25</td></tr>
<tr><td colspan="2">Learning_rate</td><td>0.005</td></tr>
<tr><td colspan="2">Input feature dimension</td><td>52</td></tr>
<tr><td rowspan="4">Bi-LSTM layer</td><td rowspan="2">Forward LSTM layer</td><td>RNN_cell: LSTM</td></tr>
<tr><td>Hidden size of the RNN cell: 148</td></tr>
<tr><td rowspan="2">Backward LSTM layer</td><td>RNN_cell: LSTM</td></tr>
<tr><td>Hidden size of the RNN cell: 148</td></tr>
<tr><td>CNN layer</td><td>Filter_size</td><td>[9,52,1,592]</td></tr>
<tr><td rowspan="2">Full connected layer</td><td rowspan="2">Hidden layer 1</td><td>Units: 2</td></tr>
<tr><td>Activation: Tanh</td></tr>
<tr><td colspan="2">Output layer</td><td>Activation: Softmax</td></tr>
</table>

**Table S4.** The hyper-parameters of TransDFL (fine-tuned).

<table>
<tr><th colspan="2">Hyper-parameter</th><th>Details</th></tr>
<tr><td colspan="2">Feature sliding window</td><td>9</td></tr>
<tr><td colspan="2">Batch_size</td><td>25</td></tr>
<tr><td colspan="2">Learning_rate</td><td>0.0008</td></tr>
<tr><td colspan="2">Input feature dimension</td><td>52</td></tr>
<tr><td rowspan="4">Bi-LSTM layer</td><td rowspan="2">Forward LSTM layer</td><td>RNN_cell: LSTM</td></tr>
<tr><td>Hidden size of the RNN cell: 148</td></tr>
<tr><td rowspan="2">Backward LSTM layer</td><td>RNN_cell: LSTM</td></tr>
<tr><td>Hidden size of the RNN cell: 148</td></tr>
<tr><td>CNN layer</td><td>Filter_size</td><td>[9,52,1, 592]</td></tr>
<tr><td rowspan="2">Full connected layer</td><td rowspan="2">Hidden layer 1</td><td>Units: 2</td></tr>
<tr><td>Activation: Tanh</td></tr>
<tr><td colspan="2">Output layer</td><td>Activation: Softmax</td></tr>
<tr><td rowspan="2">Weighted loss</td><td>Positive weight</td><td>0.6</td></tr>
<tr><td>Negative weight</td><td>0.4</td></tr>
</table>

**Table S5.** The performance of 6 state-of-the-art IDR predictors for predicting DFLs on TE82 dataset (situation-I).

| **Predictor*** | **Pre** | **Rec** | **F1** |
|---|---|---|---|
| TransDFL[a] | 0.586 | 0.452 | 0.510 |
| SPINE-D | 0.132 | 0.550 | 0.213 |
| IDP-Seq2seq | 0.114 | 0.468 | 0.183 |
| AUCpreD | 0.087 | 0.245 | 0.128 |
| SPOT-Disoreder | 0.098 | 0.349 | 0.153 |
| DISOPRED3 | 0.089 | 0.286 | 0.136 |
| SPOT-Disoreder2 | 0.092 | 0.345 | 0.145 |

* Ranked by F1 value in descending order.

[a] Proposed DFL predictor.

**Table S6.** The performance of 6 state-of-the-art IDR predictors for predicting DFLs on TE82 dataset (situation-II).

| **Predictor*** | **Pre** | **Rec** | **F1** |
|---|---|---|---|
| TransDFL[a] | 0.149 | 0.727 | 0.247 |
| SPINE-D | 0.098 | 0.550 | 0.166 |
| IDP-Seq2seq | 0.087 | 0.468 | 0.147 |
| SPOT-Disoreder | 0.079 | 0.349 | 0.129 |
| DISOPRED3 | 0.078 | 0.286 | 0.123 |
| SPOT-Disoreder2 | 0.075 | 0.345 | 0.123 |
| AUCpreD | 0.075 | 0.245 | 0.115 |

* Ranked by F1 value in descending order.

[a] Proposed DFL predictor.
